# The interplay between electron transport chain function and iron regulatory factors influences melanin formation in *Cryptococcus neoformans*

**DOI:** 10.1101/2024.02.15.580540

**Authors:** Peng Xue, Eddy Sánchez-León, Guanggan Hu, Christopher WJ Lee, Braydon Black, Anna Brisland, Haohua Li, Won Hee Jung, James W. Kronstad

## Abstract

Mitochondrial functions are critical for the ability of the fungal pathogen *Cryptococcus neoformans* to cause disease. However, mechanistic connections between key functions such as the mitochondrial electron transport chain (ETC) and virulence factor elaboration have yet to be thoroughly characterized. Here, we observed that inhibition of ETC complex III suppressed melanin formation, a major virulence factor. This inhibition was partially blocked upon loss of Cir1 or HapX, two transcription factors that regulate iron acquisition and use. In this regard, loss of Cir1 derepresses the expression of laccase genes as a potential mechanism to restore melanin, while HapX may condition melanin formation by controlling oxidative stress. We hypothesize that ETC dysfunction alters redox homeostasis to influence melanin formation. Consistent with this idea, inhibition of growth by hydrogen peroxide was exacerbated in the presence of the melanin substrate L-DOPA. Additionally, loss of the mitochondrial chaperone Mrj1, which influences the activity of ETC complex III and reduces ROS accumulation, also partially blocked antimycin A inhibition of melanin. The phenotypic impact of mitochondrial dysfunction was consistent with RNA-Seq analyses of WT cells treated with antimycin A or L-DOPA, or cells lacking Cir1 that revealed influences on transcripts encoding mitochondrial functions (e.g., ETC components and proteins for Fe-S cluster assembly). Overall, these findings reveal mitochondria-nuclear communication via ROS and iron regulators to control virulence factor production in *C. neoformans*.

**IMPORTANCE:** There is a growing appreciation of the importance of mitochondrial functions and iron homeostasis in the ability of fungal pathogens to sense the vertebrate host environment and cause disease. Many mitochondrial functions such as heme and iron-sulfur cluster biosynthesis, and the electron transport chain (ETC), are dependent on iron. Connections between factors that regulate iron homeostasis and mitochondrial activities are known in model yeasts and are emerging for fungal pathogens. In this study, we identified connections between iron regulatory transcription factors (e.g., Cir1 and HapX) and the activity of complex III of the ETC that influence the formation of melanin, a key virulence factor in the pathogenic fungus *Cryptococcus neoformans*. This fungus causes meningoencephalitis in immunocompromised people and is a major threat to the HIV/AIDS population. Thus, understanding how mitochondrial functions influence virulence may support new therapeutic approaches to combat diseases caused by *C. neoformans* and other fungi.

## INTRODUCTION

Mitochondria play a central role in respiratory metabolism for fungal pathogens of plants and animals, and support the ability of fungi to sense environmental and host conditions (1–5). The importance of mitochondrial functions in pathogenesis has been demonstrated for a number of fungal pathogens of humans including *Cryptococcus neoformans* and the related species *C. gattii* (1, 6–8). These basidiomycete yeasts have a global impact on human health because of their propensity to cause life-threatening meningoencephalitis in immunocompromised hosts, including the HIV/AIDS population, organ transplant recipients, and patients undergoing chemotherapy (9–11). Key features of these species connected to mitochondrial activities and contributing to disease include the formation of a polysaccharide capsule, titan cells, and the cell-wall pigment melanin (12–14). These connections are demonstrated, for example, by the reduction in capsule formation upon inhibition of the electron transport chain (ETC) in *C. neoformans* and a fascinating “division of labour” role for mitochondria in *C. gattii* (8, 15). In the latter case, a subset of fungal cells displays a tubular mitochondrial morphology in response to oxidative stress provoked by the host. These cells support the proliferation of other *C. gattii* cells with non-tubular organelle morphology within host phagocytic cells. These results, combined with the role of mitochondria in generating reactive oxygen species (ROS) (16–18), indicate a central role for the organelle in fungal pathogenesis.

A number of additional studies demonstrate the role of mitochondria the virulence of *C. neoformans*. For example, a mutation in the promoter of the gene encoding mitochondrial complex I protein NADH dehydrogenase increases capsule and melanin formation (19). Mutations in other regulatory factors also influence mitochondrial functions and virulence. These include mutants lacking components of the kinase module of the mediator complex (Cdk8 and Ssn801) that display altered mitochondrial morphology, reduced growth on acetate, and impaired responses to oxidative and cell wall stressors (6, 7). They also include mutants lacking the transcription factors Mig1 or Gsb1 that demonstrate sensitivity to ETC inhibitors, oxidative stress, and agents that challenge cell wall integrity (20, 21). In this context, recent work identified the heat shock transcription factor Hsf3 as an important regulator of ROS homeostasis in mitochondria, although mutants lacking the factor displayed only minor attenuation of virulence in mice (22). This result is reminiscent of the finding that HapX, another regulator of mitochondrial functions also plays a minor role in virulence (23). Other factors that influence mitochondria in *C. neoformans* include the temperature-responsive J domain protein Mrj1 that supports ETC function, influences shedding of capsule polysaccharide, and impacts cell wall structure (24). A detailed investigation revealed that Mrj1 plays a positive role in maintaining ETC function at the complex III step.

Other mutants in *C. neoformans* display phenotypes that reinforce the known connections between mitochondrial and iron homeostasis. These include mutants lacking Vps45, a Sec1/Munc18 (SM) protein that participates in vesicle fusion; these mutants have increased sensitivity to the ETC inhibitors SHAM (alternative oxidase), antimycin A (complex III), and potassium cyanide (complex IV) (25). Additionally, iron acquisition, cell wall integrity, mitochondrial function and virulence are dependent on Vps45. Similarly, loss of the mitochondrial ABC transporter Atm1 which is required for exporting an iron-sulfur (Fe-S) cluster precursor to the cytoplasm results in impaired growth on non-fermentable carbon sources, enhanced sensitivity to oxidative stress, reduced activity of the cytoplasmic Fe-S protein Leu1, and attenuated virulence in mice (26). Atm1 also plays an important role in the response to copper toxicity that targets Fe-S containing proteins (27).

During infection, pathogenic fungi use transcription factors to sense iron availability and regulate iron transport and homeostasis, ensuring successful competition for iron to support proliferation *in vivo* (28–30). We previously identified Cir1 as an iron-responsive transcription factor in *C. neoformans* (28). Genetic studies revealed that Cir1 is not only a master regulator of iron homeostasis but also regulates key virulence factors including melanin and capsule. In particular, the *cir1Δ* mutant exhibited markedly reduced infectivity, defective capsule production, and impaired growth at 30°C or 37°C. By contrast, the mutant showed enhanced melanin production, and this regulation is due in part to direct binding of Cir1 to the promoter of the laccase gene *LAC1* and repression of transcription (28, 31). Melanin is deposited in the cell wall and is a major virulence factor for *C. neoformans* that contributes to protection against phagocytosis and oxidative killing by phagocytic cells (12, 13, 32–34). *C. neoformans* synthesizes melanin from exogenous catecholamine substrates such as L- 3,4-dihydroxyphenylalanine (L-DOPA) through both laccase-catalyzed reactions and spontaneous oxidation (33).

Cir1 is also a binding partner for the monothiol glutaredoxin Grx4 that participates in iron homeostasis in *C. neoformans* (35). Additionally, mutation of the glutaredoxin (GRX) domain of Grx4 results in defective melanin synthesis. Thus, Grx4 and Cir1 provide a key connection between iron homeostasis and melanin formation, but the underlying signaling mechanisms are poorly understood. In other fungi, Grx4 also interacts with the HapX/Php4 component of the CCAAT binding complex (29, 36). This observation is consistent with the participation of HapX in iron homeostasis in *C. neoformans* including repression of iron-dependent functions (e.g., mitochondrial gene expression) upon iron limitation (23, 31). Loss of HapX also results in reduced susceptibility to agents such as H_2_O_2_ and menadione that cause oxidative stress (20). Our previous RNA-Seq and ChIP-Seq experiments revealed that *HAPX* is also a direct target of Cir1 repression under iron replete conditions (23, 31). Thus, Cir1 and HapX are key iron regulators that control iron homeostasis and mitochondrial functions in *C. neoformans* (37). Furthermore, the pH responsive transcription factor Rim101 also contributes to iron regulation in this pathogen (38, 39).

In this study, we examined the impact of ETC inhibition on melanin formation and found a balancing influence of ROS and the activities of Cir1 and HapX. We also established the influence of the melanin substrate L-DOPA, inhibition of ETC complex III, or loss of Cir1 on the transcript levels of mitochondrial functions. Thus, mitochondrial functions and the iron regulatory factors appear to both influence the proper balance between oxidation and reduction to establish the conditions necessary for melanin formation. In general, the regulation of melanization in fungi is complex (13, 40–42), and our investigation adds new insights into connections between mitochondrial function and virulence factor deployment.

## RESULTS

### Inhibition of ETC complexes I and III impairs melanin formation

Given that melanin formation is influenced by a mutation in the gene for mitochondrial NADH dehydrogenase, a component of complex I of the ETC (19), we hypothesized that mitochondrial functions influence laccase expression, trafficking, and/or activity. In particular, inhibition of ETC complexes I and III generates ROS including superoxide anion radical and hydrogen peroxide, and the subsequent influence on intracellular redox conditions may impact melanin formation (16–18, 22, 43). Additionally, mitochondria play important roles in metal ion homeostasis, heme biosynthesis, and Fe-S cluster biogenesis that may influence the activity of regulatory factors such as Cir1 which control laccase expression (31, 44). We investigated mitochondrial contributions to melanin formation by examining pigment formation on medium with L-DOPA and inhibitors of each ETC complex (Fig. 1A). We found that melanin formation in the wild-type (WT) strain and the *cir1Δ* mutant was inhibited by the complex I inhibitor rotenone, and that growth of the *cir1Δ* mutant was impaired on rotenone compared with the other strains. A more striking reduction in melanin accumulation was observed for the WT strain in the presence of the complex III inhibitors antimycin A or myxothiazol (Fig. 1A). Interestingly, the impact of complex III inhibitors on melanization was partially blocked in the *cir1Δ* and *hapXΔ* deletion mutants (Fig. 1A). Additionally, the growth of the *cir1Δ* mutant was not markedly impaired by antimycin A or myxothiazol compared to treatment with rotenone. A mutant lacking the pH-responsive transcription factor Rim101 was included as a control and did not influence melanin formation in the presence of ETC inhibitors. Overall, we conclude that inhibition of ETC complex III activity impairs melanin formation, Cir1 and HapX influence the phenotypic impact of inhibition, and that inhibition of complex I also has impact on melanin formation.

**FIG 1.**
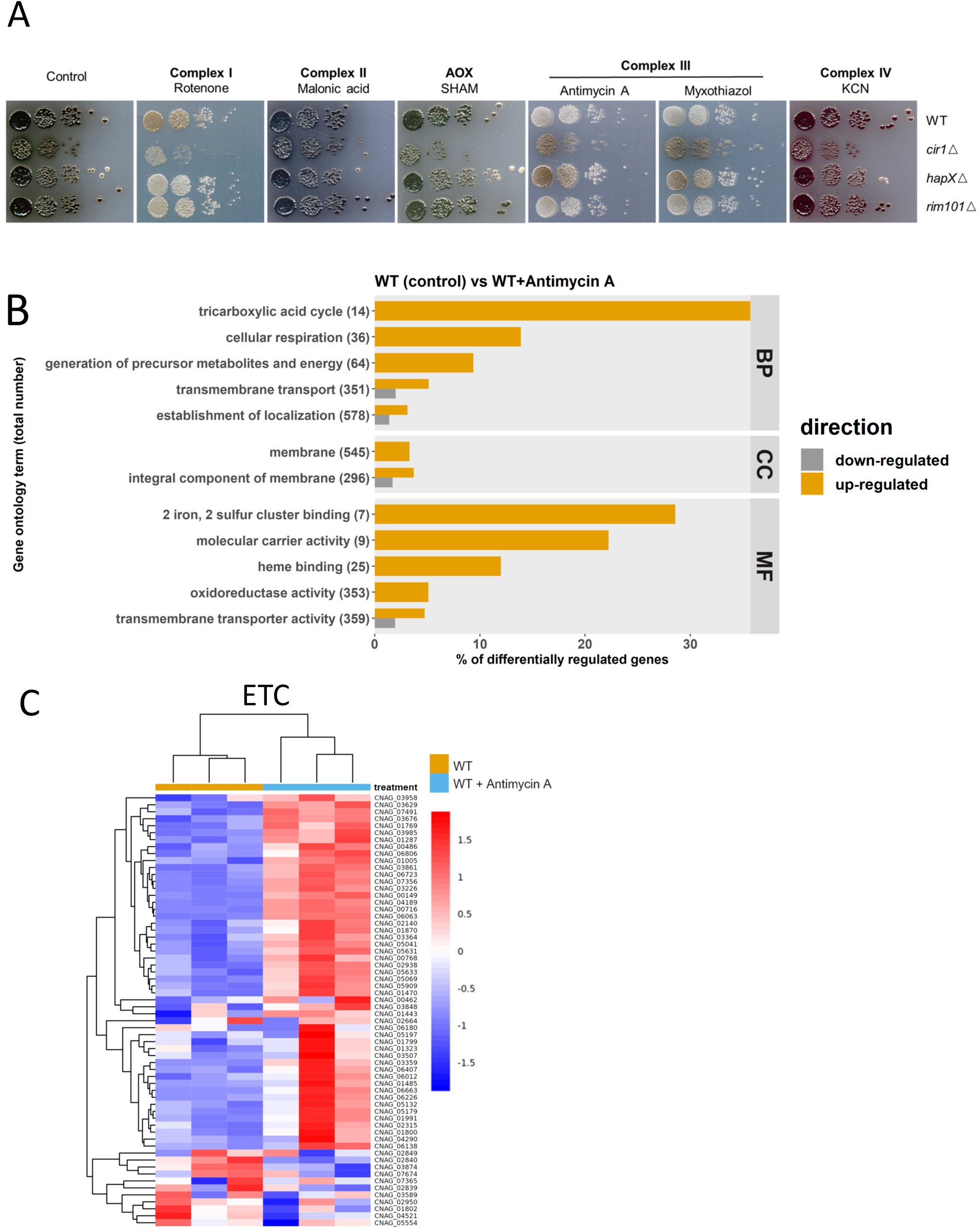
ETC inhibition influences melanin formation and the transcription of genes for mitochondrial functions. (**A**) Spot assays of WT and mutant strains on L-DOPA agar plates with or without inhibitors of the ETC including 1.2 μg/ml rotenone, 100 μM malonic acid, 0.2 mM salicylhydroxamic acid (SHAM), 0.5 μg/ml antimycin A, 0.5 μM myxothiazol, and 100 μM KCN. Plates were incubated for 72 h at 30°C in the dark. (**B**) Gene ontology (GO) categories of the differentially expressed genes identified by RNA-seq analysis of WT cells with and without antimycin A (0.5 μg/ml) treatment. The total number of genes in each functional category is displayed in parentheses, and the percent of all differentially expressed genes is indicated. BP: biological process; CC: cell component; MF: molecular function. (**C**) Heatmap of the expression of genes encoding components of ETC in the WT versus WT with antimycin A treatment. Samples are clustered according to expression similarity and the differences in transcript levels were significant (*P* < 0.05). The corresponding genes are listed in Table S1.

### Inhibition of ETC complex III impacts transcripts for mitochondrial functions, but not laccase genes

To investigate the connection between the ETC and melanin, we focused our subsequent experiments on antimycin A and complex III inhibition by analyzing the transcriptome of the WT strain treated with this inhibitor. In particular, we hypothesized that an influence on the expression of *LAC1* (encoding the major laccase) or regulatory factors for *LAC1* (e.g., Cir1) might account for the melanin defect. Our analysis identified differential transcript levels for 210 genes, with 169 upregulated and 41 downregulated (Fig. S1). Moreover, subsequent analysis of gene ontology (GO) terms revealed up regulation of molecular function categories associated with Fe-S cluster and heme binding, and molecular carrier activity (Fig. 1B). These GO terms prompted an analysis of transcript levels for components of the ETC as well as mitochondrial (ISC) and cytoplasmic (CIA) Fe-S cluster assembly machinery to examine the impact of ETC complex III inhibition. As shown in the heatmaps in Fig. 1C and Fig. S2A, we found that inhibition resulted in elevated transcript levels for ETC components as well as for functions for ISC and CIA machinery in the WT strain. Furthermore, the ETC and mitochondrial ISC assembly pathways were found to be significantly enriched through Gene Set Enrichment Analysis (GSEA). Table S1 lists the specific genes represented by the heatmaps and Fig. S3 documents the output from the GSEA. The RNA-Seq analysis revealed minimal impact of complex III inhibition on the transcript levels of *LAC1* or *CIR1,* thus refuting our hypothesis.

We employed quantitative reverse transcription-PCR (qRT-PCR) to validate the RNA-Seq data. This approach confirmed the transcript levels of the genes CNAG_01881 and CNAG_05199 (ISC pathway genes), *ATM1* (mitochondrial ISC transporter), and CNAG_05840 (CIA pathway gene) in agreement with the RNA-Seq data (Fig. S2B). The transcript levels for the *LAC1* and *CIR1* genes were also confirmed by qRT-PCR (Fig. S2B). A specific examination of the transcript levels for *LAC1, CIR1, LAC2* (a second gene for laccase) and *HAPX* revealed that inhibition of ETC complex III caused only modest reductions in mRNA levels for the *LAC1*, *LAC2,* and *HAPX* genes, and a slight increase in mRNA levels for *CIR1* (Fig. S4). As mentioned, our previous results showed that Cir1 directly and negatively regulates *LAC1* and *HAPX* gene transcription (23, 31). Overall, we conclude that inhibition of complex III has a major influence on transcript levels for mitochondrial functions involved in the ETC and Fe-S biogenesis, but only a minor impact on transcript levels for the laccase genes that is likely insufficient to explain the impaired melanin formation.

### Oxidative stress influences melanin formation

Given the minor influence of antimycin A on *LAC1* and *LAC2* gene transcription and the fact that the inhibitor triggers ROS in *C. neoformans* and other organisms (16–18, 22, 43), we hypothesized that antimycin A influenced melanin by establishing redox conditions that interfere with laccase activity. To support this idea, we examined the transcript levels for genes responsive to oxidative stress and found that the transcript levels for the *CAT1* and *CAT3* genes encoding catalases were elevated ∼2 fold upon antimycin A treatment (Table 1). The transcript levels for these genes are known to be responsive to H_2_O_2_, among a larger group of oxidative stress genes (45). These observations prompted a closer examination of the impact of ROS during melanin formation and we found that the combination of H_2_O_2_ or menadione with the melanin substrate L-DOPA impaired growth in liquid medium (Fig. 2A).

**FIG 2.**
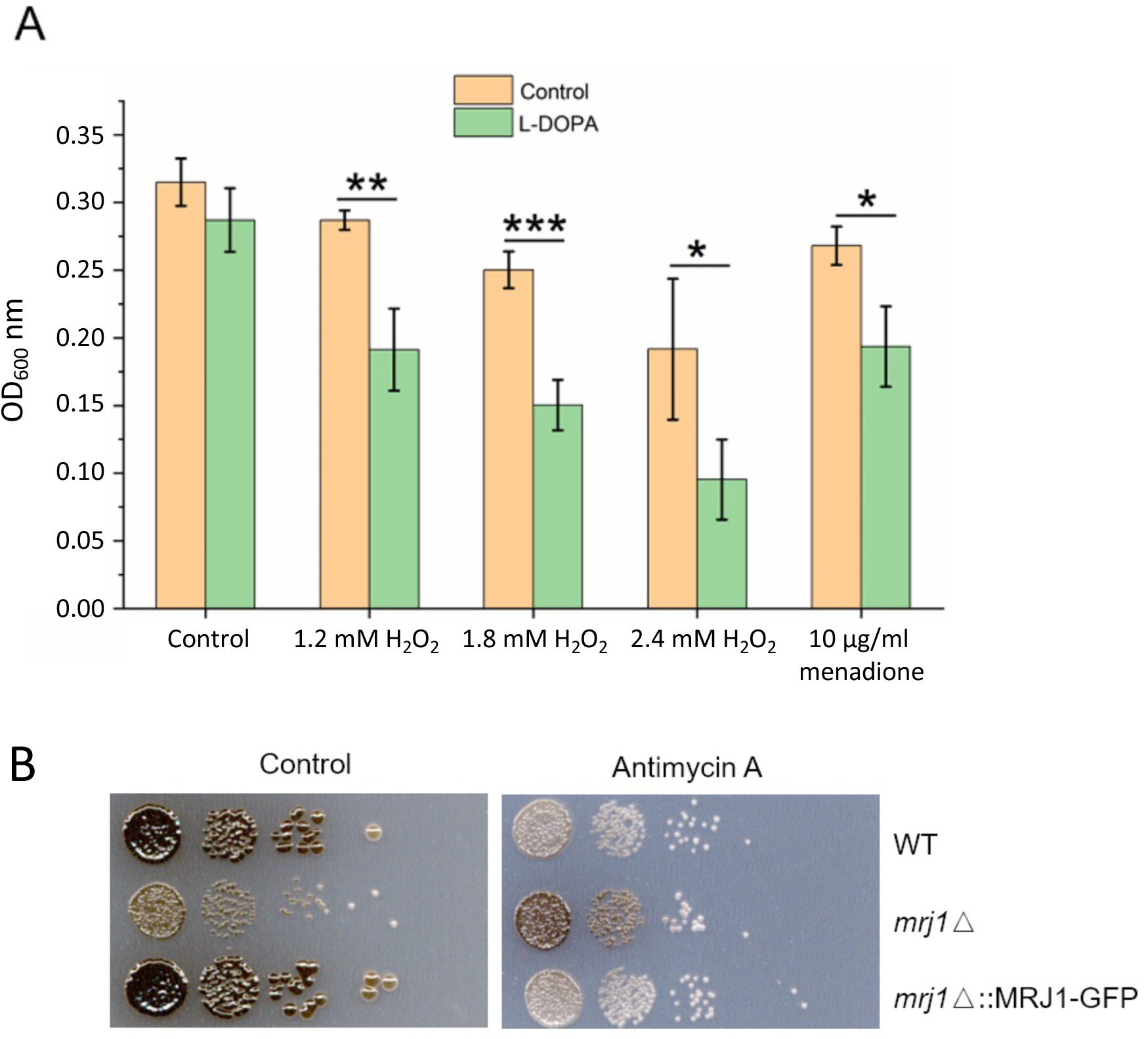
Growth and melanin formation are influenced by L-DOPA, oxidative stress, and the mitochondrial regulator Mrj1. (**A**) The growth of WT cells was tested in liquid medium with and without 0.7 mM L-DOPA in the presence of hydrogen peroxide (H_2_O_2_) or menadione at the indicated concentrations for 18 h at 30°C and 180 rpm in the dark. The assays were performed using 96-well microplates with each well containing a final volume of 200 μl. The initial cell density was set at 1×10^5^ cells/ml. Mean values of three biological replicates are shown ± standard deviation (SD). Significant differences were determined by t tests and are indicated by * (*P* < 0.05), ** (*P* < 0.01), or *** (*P* < 0.005). (**B**) Spot assays of WT, *mrj1Λ* and *mrj1Λ*::*MRJ1-GFP* strains showing that the absence of *MRJ1* rescues melanin formation on L-DOPA plates in the presence of antimycin A (0.5 μg/ml). Plates were incubated for 72 h at 30°C in the dark.

The connection with oxidative stress and redox homeostasis was examined further by using a mutant known to influence the ETC at complex III and to have reduced ROS levels. Specifically, we examined a mutant lacking the *MRJ1* gene encoding mitochondrial respiration J-domain protein 1 that influences the activity of ETC complex III by directly binding to Qcr2, a subunit of ubiquinol cytochrome c reductase (24). We found that the effect of complex III inhibition on melanin formation was partially blocked in the *mrj1*11 mutant, a result consistent with the influence of Mrj1 on ROS accumulation (Fig. 2B). These results further support the idea that melanin formation is conditioned at least in part through a balance in redox homeostasis.

### L-DOPA negatively impacts the transcription of mitochondrial functions and genes encoding catalases

Given that the combination of L-DOPA and H_2_O_2_ impaired growth, we hypothesized that L-DOPA treatment might also influence oxidative stress and mitochondrial functions. We therefore examined the impact of L-DOPA by performing RNA-Seq to compare the transcript profiles for WT cells in the presence and absence of L-DOPA. We found that L-DOPA treatment resulted in differential transcript levels for 1,262 genes, with 836 upregulated and 426 downregulated (Fig. S5). Analysis of the GO categories for the differentially transcribed genes indicated a negative impact of L-DOPA treatment on functions associated with mitochondria (Fig. 3A). This observation was further supported by GSEA using pathways from the KEGG database and homologous gene IDs based on the genome of the *C. neoformans* strain JEC21. This analysis revealed that 15 KEGG terms for mitochondrial functions, including ETC components, were negatively enriched upon growth with L-DOPA (Fig. 3B). The differential regulation of representative genes encoding ETC components (e.g., genes CNAG_01287, CNAG_05041 and CNAG_07491) was confirmed by qRT-PCR (Fig. S6). We also noted the transcript levels of genes encoding Cir1, HapX or Rim101 were not differentially expressed in the RNA-Seq analysis of the response to L-DOPA, and only a minor influence was observed by qRT-PCR (Fig. S6). A summary of the differential transcript levels for mitochondrial functions is presented in Table S2. A specific examination of the genes for oxidative stress (Table 1) also identified the *CAT1, CAT2* and *CAT3* transcripts as being significantly elevated upon treatment with L-DOPA. Together, these results revealed that conditions supporting melanin formation (i.e., the presence of L-DOPA) have an impact on the expression of mitochondrial functions as well as oxidative stress. These results reinforce the idea that mitochondria are responsive to environmental signals, including L-DOPA, and control ROS generation to establish conducive redox conditions that influence virulence factor (melanin) production.

**FIG 3.**
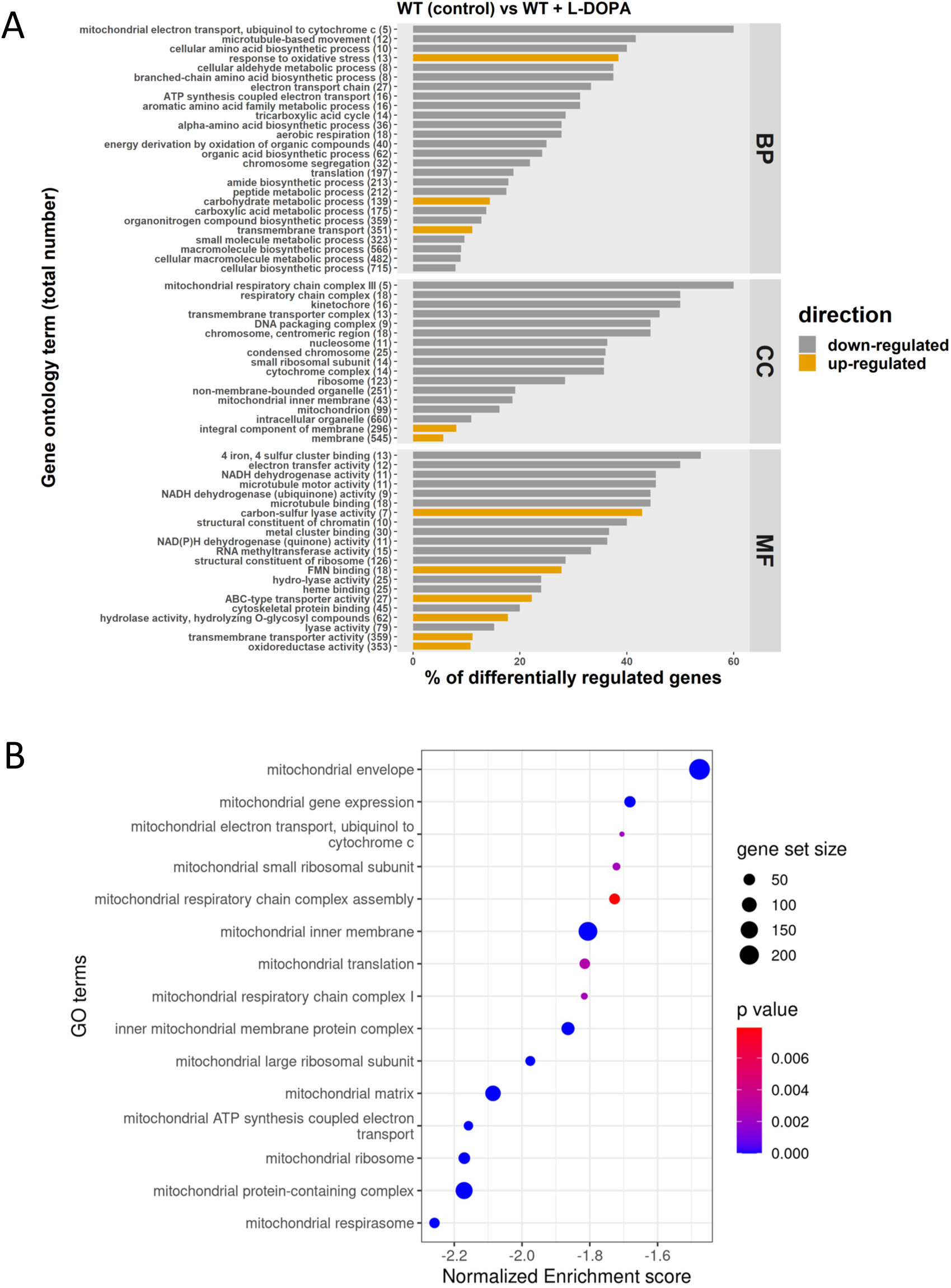
L-DOPA provokes down-regulation of mitochondrial functions. (**A**) Gene ontology (GO) terms for genes with differential transcriptions upon treatment with L-DOPA (0.7 mM). The total number of genes in each functional category is displayed in parentheses, and the percent of all differentially expressed genes is indicated on the x axis. BP: biological process; CC: cell component; MF: molecular function. (**B**) Gene Set Enrichment Analysis demonstrating negative enrichment for pathways directly involved in mitochondrial function. Significantly regulated pathways were determined using a 0.05 cut-off p-value and a false discovery rate of 0.25. The list of regulated genes for components of the ETC and iron-sulfur cluster biogenesis is presented in Table S2.

### Cir1 influences melanin formation through de-repression of *LAC1* transcription and an influence of mitochondrial gene expression

As shown in Fig. 1A, complex III inhibitors impaired melanin formation in the WT strain, and the *cir1Δ* and *hapXΔ* mutants partially restored melanin. Our previous microarray, and RNA-Seq experiments with *cir1Δ* mutant demonstrated that Cir1 represses *LAC1* transcription (23, 31). Furthermore, ChiP-seq analysis revealed that Cir1 directly binds the promoter of the *LAC1* gene to repress transcription (31). These results suggest that de-repression upon loss of Cir1 during antimycin A treatment may contribute to restored melanin formation. We therefore examined the influence of Cir1 on *LAC1* transcription, ETC function and oxidative stress genes by performing an RNA-Seq experiment with WT and the *cir1Δ* mutant under the minimal medium conditions used for the analysis of the impact of antimycin A on the transcriptome (Materials & Methods). We identified 853 differentially expressed genes (710 upregulated and 143 downregulated) and an overview of the analysis is shown in Fig. S7. As with previous transcriptome studies with Cir1 (23, 28, 31), an examination of the GO terms for molecular functions that were upregulated in the *cir1Δ* mutant revealed categories that included oxidoreductase activity (acting on peroxide as acceptor, and acting on single donors with incorporation of molecular oxygen), Fe-S cluster, copper ion binding, heme binding, antioxidant activity, transition metal iron transmembrane transporter activity, and iron ion binding (Fig. 4). For example, the transcript levels of mitochondrial ISC assembly machinery components were significantly up-regulated in the *cir1Δ* mutant compared to the WT, and we note that there are many Fe-S cluster binding proteins in complexes I, II and III of the ETC. Additionally, genes encoding oxidative stress functions including the catalase genes were up-regulated in the *cir1Δ* mutant (Table 1). Heatmaps of the regulation of transcripts for ETC components and mitochondrial ISC assembly pathway components in the *cir1Δ* mutant as compared to WT strain are shown in Fig. 4B and Fig.S8A; the corresponding genes are listed in Table S3 and Fig. S9 presents the GSEA analysis of the ETC and mitochondrial ISC pathways. Overall, these data confirm the influence of Cir1 on mitochondrial functions - an affect that may impact susceptibility to complex III inhibitors such as antimycin A through enhanced expression of specific organelle transcripts.

**FIG 4.**
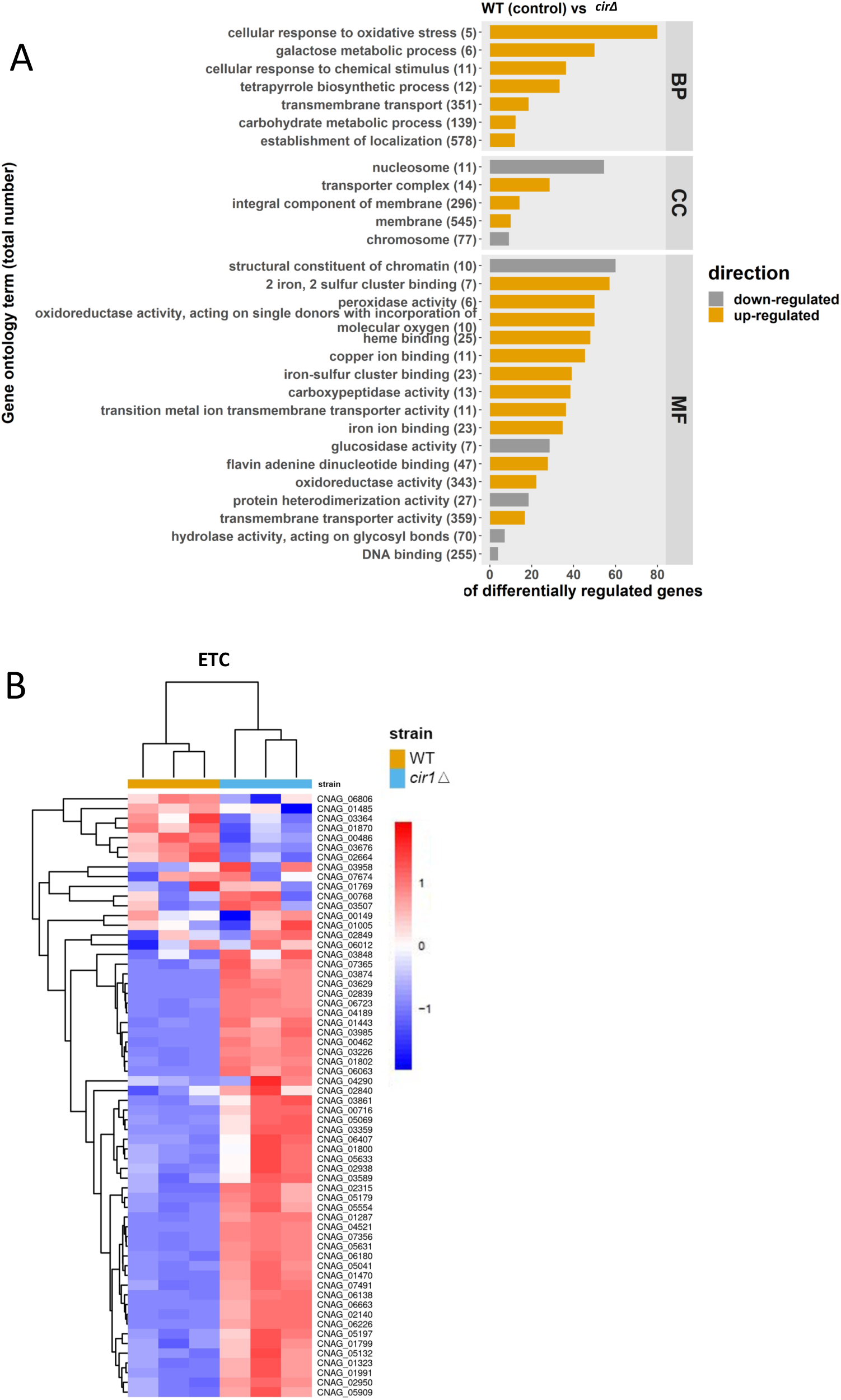
Loss of Cir1 results in up-regulation of mitochondrial functions. **(A)** Gene ontology (GO) terms for RNA-Seq analysis of the WT strain vs the *cir1Δ* mutant. The total number of genes in each functional category is displayed in parentheses, and the percent of all differentially expressed genes is indicated. BP: biological process; CC: cell component; MF: molecular function. (**B**) Heatmap of the expression of genes encoding components of electron transport chain (ETC) in the WT and *cir1Δ* mutant strains. The differences in transcript levels were significant (*P* < 0.05), and the corresponding genes are listed in Table S3.

To confirm the RNA-Seq data, the transcript levels of the ISC pathway gene CNAG_03395, the ETC genes CNAG_06063, CNAG_07491 and CNAG_02839, and *LAC1* were examined by qRT-PCR. We found comparable patterns of regulation between the RNA-Seq and qRT-PCR results (Fig. S8B). Consistent with previous analyses of the regulatory influence of Cir1 (23, 28, 31), we verified the differences between the WT strain and the *cir1Δ* mutant for the transcript levels of the laccase genes *LAC1* and *LAC2*, and confirmed the regulatory influence of Cir1 on the transcription of the genes using qRT-PCR (Fig. S8C). Overall, these results suggest that loss of Cir1 leads to de-repression of *LAC1* to influence melanin in the presence of complex III inhibitors. As mentioned above, an additional influence on the expression of mitochondrial and oxidative stress functions may also condition susceptibility to ETC inhibition.

### Cir1 and HapX respond differently to oxidative stress

In contrast to the regulation of *LAC1* expression by Cir1, previous experiments revealed that HapX does not regulate laccase expression (23, 31). This observation suggests a different mechanism for melanin recovery upon antimycin A treatment, and we hypothesized that an influence of HapX on oxidative stress and the redox environment impacts melanin formation. We therefore examined oxidative stress in more detail by measuring ROS accumulation in the WT and *cir1Δ* and *hapXΔ* mutant strains in response to antimycin A and myxothiazol treatment. Both mutants showed higher levels of DCFDA staining compared to the WT strain indicating an accumulation of ROS (Fig. 5A). The actions of multiple types of ROS (e.g., hydroxyl radicals, H_2_O_2_) can oxidize DCFDA to generate a fluorescent signal (46). In contrast, staining with dihydroethidium (DHE), which is responsive to superoxide accumulation revealed greater levels in the *cir1Δ* mutant and lower endogenous levels/accumulation in the *hapXΔ* mutant. The differences between the mutants prompted an examination of the sensitivities of each strain to oxidative stress caused by H_2_O_2_, menadione, plumbagin and paraquat (Fig. 6A). This experiment revealed greater sensitivity for the *cir1Δ* mutant compared to the WT strain and the *hapXΔ* mutant. Consistent with this differential sensitivity, we found that treatment with H_2_O_2_ provoked enhanced DCFDA straining in the *cir1Δ* mutant but not in the WT strain or the *hapXΔ* mutant (Fig. 6B). These observations are consistent with our previous finding that the *hapXΔ* mutant behaves mainly like WT with regard to sensitivity to oxidative stress (20). HapX is known to control the expression of functions for the response to oxidative stress, including the *CAT1* and *CAT3* catalase genes influenced by antimycin A and L-DOPA (Table 1) (23, 31). We confirmed this regulation by qRT-PCR for a set of the oxidative stress genes from Table 1 (Fig. S10). Overall, these results suggest that antimycin A-induced inhibition of melanin results from changes to the redox environment that are unfavourable for melanin synthesis. Cir1 and particularly HapX appear to modulate this influence by regulating the expression of antioxidant functions, along with the de-repression of laccase expression upon loss of Cir1.

**FIG 5.**
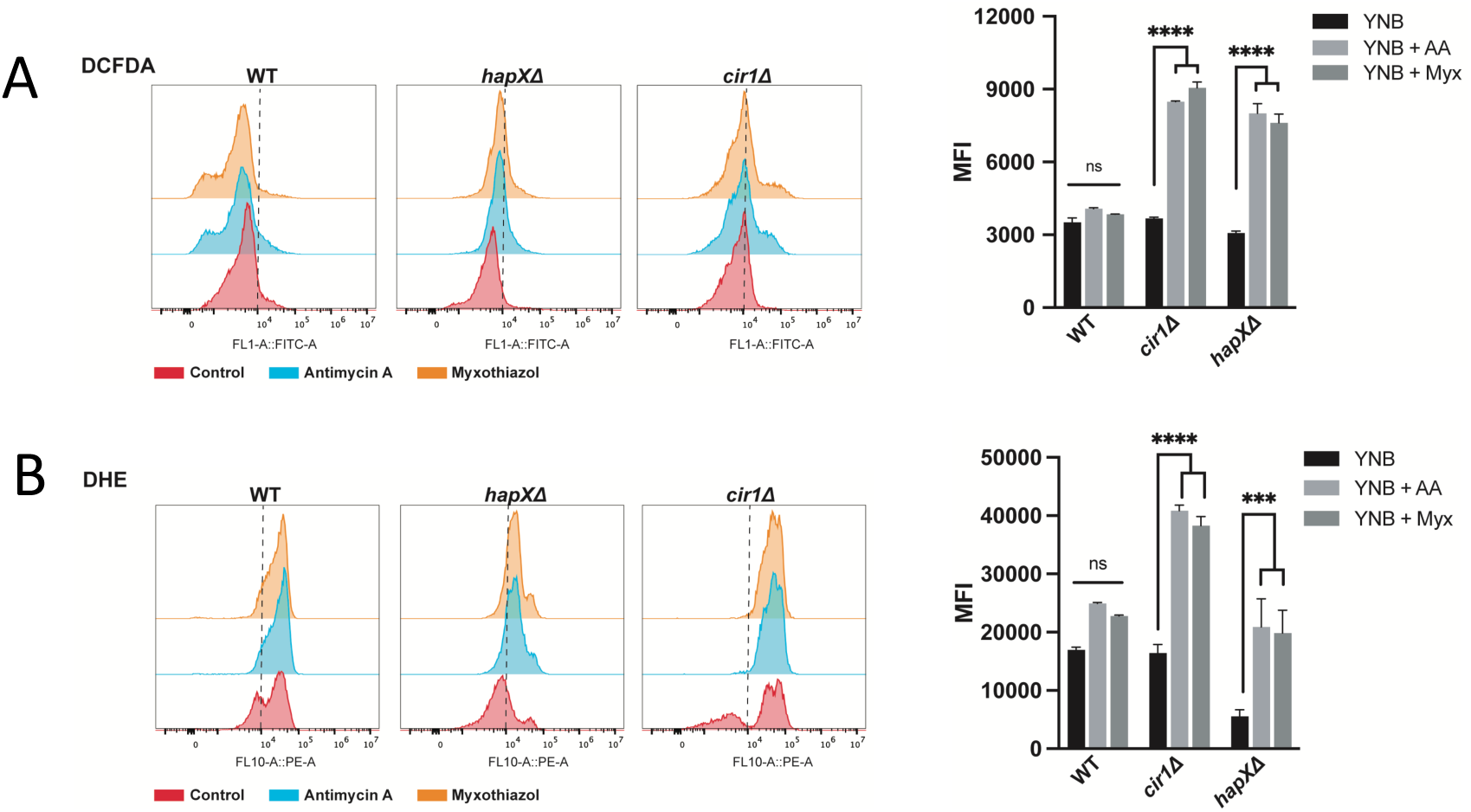
Loss of Cir1 or HapX results in accumulation of ROS in response to complex III inhibition. (**A**) Flow cytometry analysis of WT and mutant cells stained for 1 hour with 2ʹ,7ʹ-Dichlorofluorescein Diacetate (DCFDA, 16 μM) to detect ROS accumulation in response to exposure to antimycin A (AA, 50 μM) or myxothiazol (Myx, 7 μM) for 24 h at 30°C. (**B**) Flow cytometry analysis of WT and mutant cells stained with Dihydroethidium (DHE, 2.5 μg/ml) to detect ROS accumulation in response to ETC-III inhibitors as in (**A**). The data represent the mean fluorescent intensity (MFI, geometric means) from three biological replicates ± standard errors of the means. The statistical comparisons employed a two-way ANOVA test, followed by *post hoc* Šídák’s or Tukey’s multiple comparison tests (*, *P* < 0.05, **, *P* < 0.01 ***, *P* < 0.001; ****, *P* < 0.0001). ns: not significant. The gating strategy for the left panels is shown in Fig. S11.

**FIG 6.**
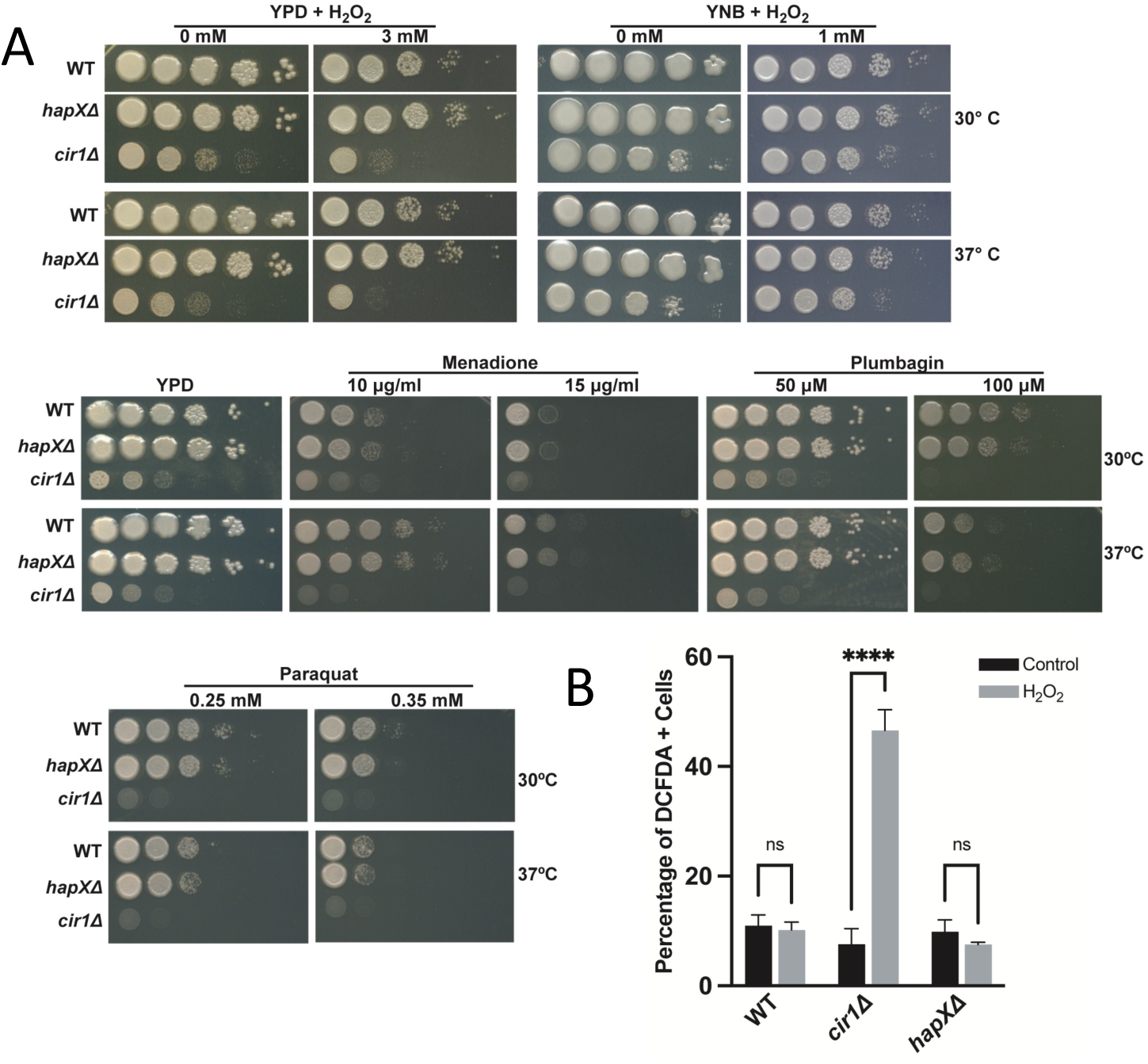
Loss of Cir1 but not HapX causes sensitivity to oxidative stress. (A) Spot assays of the WT strain and the *hapXΛ* or *cir1Λ* mutants on the indicated media were performed with 10-fold serial dilutions from an initial concentration of 2 x 10^7^ cells per ml. Five microliters were spotted into solid YPD or YNB plates supplemented with different compounds and incubated at 30°C and 37°C for 2-3 days before being scanned. The media were supplemented at the indicated concentrations with the following compounds: hydrogen peroxide (H_2_O_2_), menadione, plumbagin, or paraquat. (**B**) Graph of the percentage of DCFDA positive cells upon exposure to H_2_O_2_ [5 mM] for 1 hour at 30°C for the WT strain and the *hapXΛ* and *cir1Λ* mutants as determined by flow cytometry using the gating strategy in Fig. S11. The data represent the average from at least three biological replicates ± standard errors of the means. Statistical comparisons employed a two-way ANOVA test, followed by *post hoc* Šídák’s multiple comparison test (****, *P* < 0.0001). ns: not significant.

## DISCUSSION

Melanin deposition in the cell wall of *C. neoformans* confers protection against various stresses (e.g., oxidative killing by phagocytic cells) and reduces susceptibility to antifungal drugs. These properties are relevant to disease because mutants with defects in melanin formation show reduced virulence in mouse models, and melanin influences dissemination to the brain and aspects of the immune response including phagocytosis (13, 41). In this study, we examined the influence of impaired ETC function on melanin formation with an emphasis on the response of WT cells to antimycin A treatment, a condition known to increase ROS production (22). Given the central role of mitochondria in iron homeostasis, we focused on the influence of iron regulators on melanin formation upon mitochondrial impairment. We found that *cir1Δ* and *hapXΔ* mutants lacking key iron regulators partially restored melanin in the presence of ETC complex III inhibitors. These findings are consistent with our previous discovery of negative regulation of laccase expression by Cir1, as well as the regulation of genes for ETC components and oxidative stress functions by HapX (23, 31). Thus, our analysis suggests two underlying mechanisms for the recovery of melanin in the mutants upon antimycin A treatment: de-repression of *LAC1* by Cir1 and modulation of the redox environment by Cir1 and HapX (Fig. 7). The proposed influence of HapX is consistent with our previous finding of a reciprocal regulatory connection between HapX and the candidate carbon catabolite repressor Mig1 in *C. neoformans* (20). Notably, Mig1 regulates mitochondria functions, and the *mig1Δ* mutant was sensitive to ETC inhibitors and inducers of ROS.

**FIG 7.**
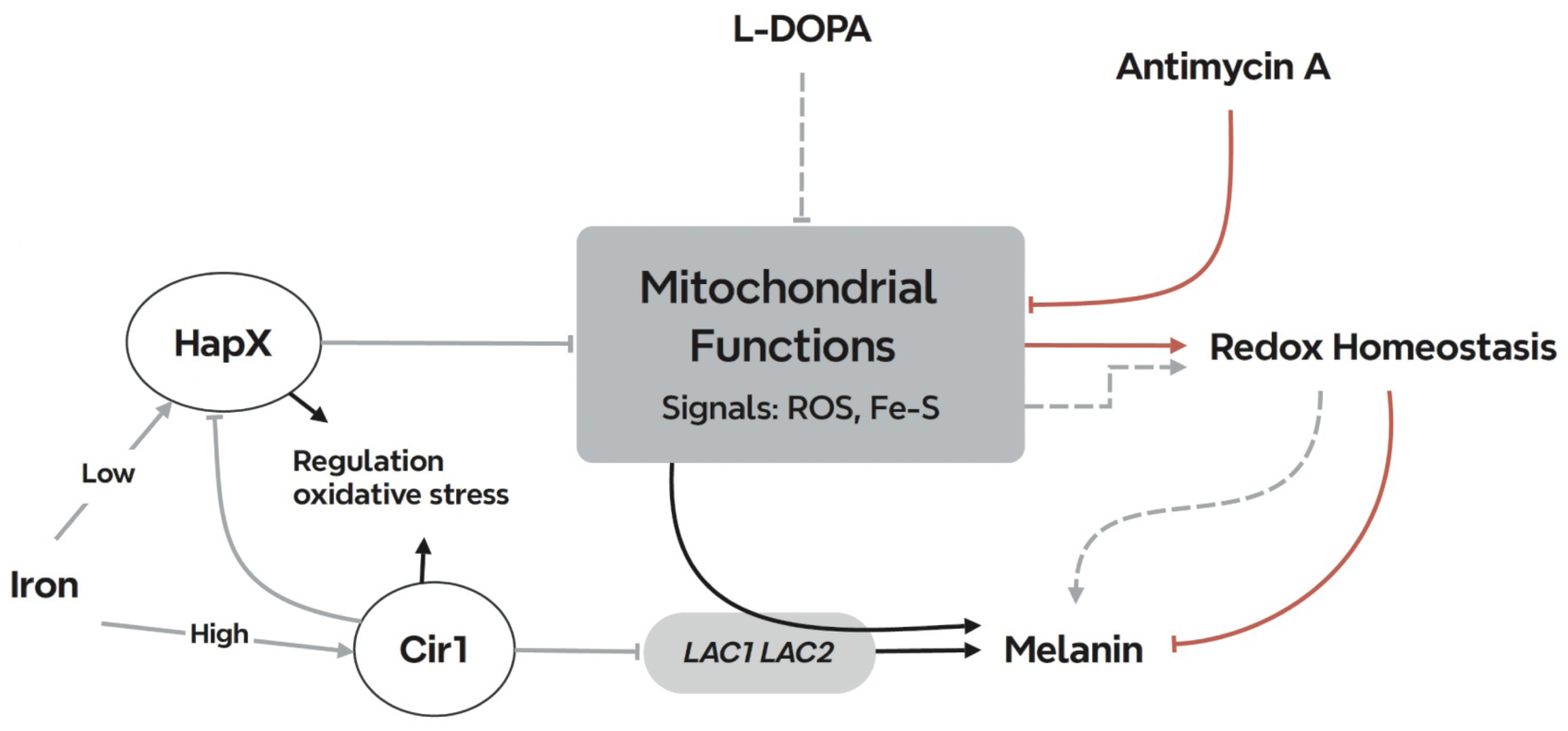
Summary model of the interplay between mitochondrial ETC function and the iron regulators that influence melanin formation in *C. neoformans*. Inhibitors of ETC complex III provoke ROS accumulation which inhibits melanin formation through an influence on laccase activity and/or localization. Cir1 directly represses transcription of the *LAC1* and *LAC2* genes encoding laccases, and loss of Cir1 may derepress the genes to a level sufficient partially restore melanin formation. Cir1 also represses the transcription of the *HAPX* gene (23, 31). HapX is a key regulator of iron-requiring functions in mitochondria, and loss of HapX derepresses genes for the response to oxidative stress. This de-repression may be sufficient to overcome ROS accumulation to partially restore melanin. Other signals including iron-sulfur clusters may be generated by ETC inhibition to include the activities of the iron regulators and the laccases.

In addition to an influence on melanin, there are additional connections between ETC activity and virulence factor expression in C*. neoformans*. For example, regulation of polysaccharide capsule elaboration by ETC complexes I and III and alternative oxidase (AOX) has been reported (15). Along with melanin, the polysaccharide capsule is a major virulence factor for *C. neoformans* (47). Interestingly, ROS also influences titan cell formation further highlighting the connections with virulence (14). These observations reinforce the emerging view that mitochondrial activity is important for fungi to resist host defenses and to cause disease, and are consistent with an impact of iron levels on the size of the polysaccharide capsule and the expression of laccase (48–50).

We focused on Cir1 in our analysis of the impact of ETC inhibition because of the known repression of *LAC1* by the protein (23, 28, 31). Several studies have documented melanin formation by *C. neoformans* in response to a variety of signals including glucose levels, temperature, cell density, and metal ions (calcium, copper, iron) (49, 51). In this context, melanin formation is known to be regulated by a complex network of transcription factors and kinases (42, 51). This network includes the transcription factors Bzp4, Usv101, Mbs1, and Hob1 that activate *LAC1* transcription, and we previously found that Cir1 directly regulates the transcription of *BZP4* (31). Thus, these other factors may counter-balance the repression of *LAC1* by Cir1 to influence melanin formation in response to different environmental and host conditions. Similarly, *HAPX* transcription is also regulated by Cir1, and the influence of HapX on the oxidative stress response was shown to affect melanin formation in the context of ETC inhibition. These observations are consistent with HapX as a key regulator of oxidative stress functions and iron-utilizing mitochondrial functions (23). Our analysis of the transcriptome in the *cir1*11 mutant also revealed regulation of mitochondrial functions and part of this regulation may occur via an influence on HapX expression given that previous RNA-Seq and ChIP-Seq did not identify the same pattern of regulation (31). The transcriptional up-regulation of genes encoding mitochondrial components of the ETC and iron-sulfur cluster synthesis observed for the *cir1*11 mutant was similar to the response to antimycin A treatment.

The observed regulation of oxidative stress functions by HapX and its known role in regulating mitochondrial functions that use iron support the premise that mitochondrial homeostasis orchestrates the necessary cellular environment for melanin formation. The role of HapX in responding to and regulating redox conditions is supported by orthologous studies in *Aspergillus* sp. and *S. cerevisiae* (52, 53). In *Aspergillus nidulans*, for example, the CCAAT binding complex, of which HapX is a regulatory subunit, is responsive to ROS via oxidation of key cysteine residues in a core subunit. HapX in *C. neoformans* interacts with the core subunits of the CCAAT binding complex (Hap2, 3 and 5) as demonstrated by DNA binding experiments (31). We note that Cir1 and HapX also have cysteine-rich regions and ROS may potentially influence the activities of these proteins via oxidation of cysteine residues. Additional experiments are needed to investigate this possibility.

A key finding in our study was the connection between melanin, L-DOPA and oxidative stress as demonstrated by the influence of H_2_O_2_ on growth in combination with L-DOPA, and by the influence on melanin formation in an *mrj1*11 mutant previously demonstrated to have reduced ROS accumulation (24). Previously, the transcriptional response of *C. neoformans* to H_2_O_2_ was analyzed and treatment resulted in the decreased expression of the *LAC1* gene (45). Additionally, a set of genes was identified that were responsive to H_2_O_2_ and that encode functions for oxidative stress resistance. Together, these results support the idea that a balance of ROS is needed to support melanin formation. However, the impact of ETC inhibition and ROS generation is likely complex due to the multitude of potential targets of oxidative damage. For example, we previously reported that Cir1 and HapX are both iron-binding proteins (31), and iron or Fe-S cluster binding to these transcription factors might be affected by inhibition of ETC complex III. Among the oxidative stress genes identified for *C. neoformans* (45), the catalase genes appear to be regulated by antimycin A, L-DOPA and HapX. However, the importance of catalase activity in this context is unclear because a mutant lacking all four identified *CAT* genes is not defective for virulence in mice (54). Oxidative stress in the context of fungal pathogenesis has recently been reviewed (55).

Our analysis of cells exposed to L-DOPA revealed a substantial impact on the transcriptome such that a large number of genes showed differential expression, including the down-regulation of mitochondrial functions. This impact is interesting because laccase activity and melanin may contribute to the neurotropism of *C. neoformans* (56). The possibility that L-DOPA is a signal as well as a substrate is intriguing because our observations of an influence on mitochondria draw parallels with studies on the role of dopamine in neurons and in neurodegenerative diseases (57, 58). For example, examination of the potential toxicity of L-DOPA in the treatment of Parkinson’s disease revealed an impact on mitochondrial and lysosomal activities (59). A previous study examined the transcriptional response of *C. neoformans* to L-DOPA using microarray analysis and, in contrast to our observations, found differential expression of only eight genes (60). Some of these genes encoded functions predicted to be involved in the response to stress and, importantly, this result is consistent with our observed categories of GO terms and our demonstration that cells are more sensitive to H_2_O_2_ in the presence of L-DOPA. The differences in the number of regulated genes could be due to several factors including the use of microarray versus RNA-Seq approaches (and associated differences in evaluating differential expression), the use of different strains (JEC21 versus H99), different preculture conditions, and the timing and level of L-DOPA exposure. However, four of the JEC21 genes had regulated orthologs in the RNA-Seq data for H99. Interestingly, regulation of five of the genes was not observed in a laccase mutant suggesting that formation of melanin may have a regulatory influence (60, 61).

In summary, we have established that impaired ETC complex III activity influences the elaboration of melanin in *C. neoformans* and that iron regulatory factors buffer this influence. Other proteins in the iron regulatory network influence melanin formation in addition to Cir1 and HapX. For example, the monothiol glutaredoxin Grx4 interacts with Cir1 and HapX and has a positive influence on melanin formation (along many other phenotypic contributions such as cell wall integrity) (62). Thus, additional work is needed to investigate the influence of Grx4 on melanin, potentially in partnership with Cir1, HapX and other regulatory proteins. Finally, we also found that L-DOPA treatment impacts the expression of mitochondrial functions thus warranting an investigation into a potential signaling role for this important substrate.

## MATERIALS AND METHODS

### Strains, growth conditions, and spot assays with inhibitors

The following strains were used in this study: WT strain *Cryptococcus neoformans* var. *grubii* H99, deletion mutants *cir1*11 (28), *hapX*Δ (23), *rim101*11 (63, 64), *atm1*11 (26), and *mrj1*11 (16) (24), and complemented mutants *cir1*11::*CIR1* (28), *hapX*Δ::*HAPX* (23), *rim101*11::*RIM101* (63, 64), *atm1*11::*ATM1*-*GFP* (26), and *mrj1*11::*MRJ1-GFP* (24). Routine growth was in YPD or YNB liquid or solid media.

Melanin production was examined in medium with L-DOPA (0.1% glucose, 0.1% L-asparagine, 0.3% KH_2_PO_4_, 0.7 mM L-DOPA, 0.025%MgSO4 · 7H_2_O, pH 5.6). Agar was added at 2% for solid medium. For spot assays, cells from overnight cultures at 30°C were washed twice with liquid YPD medium, density was adjusted to 1×10^5^ cells ml^-1^ and 10-fold serial dilutions were prepared. Next, 5 μl was spotted on the 2% L-DOPA agar plates with or without the ETC inhibitors at the indicated concentrations. Plates were incubated in the dark for 72 h at 30°C, and photographed to document melanin formation.

To examine *C. neoformans* WT (H99) or mutant strains, cells were grown in YPD or YNB for ∼16 hours at 30°C. Precultures were washed, resuspended in H_2_O and 10-fold serial dilutions were performed from an initial concentration of 2 x 10^7^ cells per milliliter. Five μl were spotted into solid YPD or YNB plates supplemented with different compounds and incubated at 30°C and 37°C for 2-3 days before being scanned. For the reactive oxygen stress response, rich (YPD) and minimal (YNB) media were supplemented with the following compounds: hydrogen peroxide solution (H_2_O_2_, 1-5 mM), menadione (10, 15 μg/ml), plumbagin (50, 100 μM), paraquat (0.25, 0.35 mM).

### RNA extraction, RNA sequencing and analysis, and quantitative reverse transcription-PCR

To examine the impact of complex III inhibition on the transcriptome, the WT strain and the *cir1*11 and *hapX* 11 mutants were grown in 50 ml of YPD at 30°C for 16 h. The cells were washed twice with sterile water and then incubated in 50 ml of YNB with 0.06% glucose for 16 h at 30°C. The cells were then diluted to 4.0 x 10^7^ cells in 50 ml of YNB medium (2% glucose) with or without 0.5 μg/ml antimycin A, followed by a 6 h incubation at 30°C. For the analysis of the influence of L-DOPA on the transcriptome, three replicate cultures of WT cells were grown in 50 ml of YPD (2% glucose) at 30°C for 16 h. Cells were then washed twice with sterile water, and diluted to 4.0 x 10^7^ cells in 50 ml of melanin induction medium with or without 0.7 mM L-DOPA. After incubation for 6 h at 30°C, the cells were harvested and flash frozen in liquid nitrogen. The total RNA from frozen cells was extracted by using RNeasy-Mini kit (QIAGEN), followed by treatment with DNase I (TURBO DNA-free™ Kit). RNA-Seq was performed by Genewiz (Azenta Life Sciences, South Plainfield, NJ, USA).

Differentially expressed genes were analyzed in R (version 4.3.2) using the raw gene counts provided from Genewiz and the DEseq2 package. All genes were assessed for significance using DEseq2’s inherent Wald test (pvalue cut-off = 0.05) and normalized between samples using the median of ratios method (65). The identification of enriched pathways was performed by two independent approaches. First, to identify significantly enriched functional groups, the RNA-Seq data were analyzed with respect to KEGG database annotations. For compatibility with the KEGG database, FungiDB was used to convert gene ID’s from strain H99 to strain JEC21. Based on both a ranked genelist (Ranked score = -(log2FoldChange) x log10 (p value)) derived from the DESeq2 output and the KEGG/PATHWAY database, the identification of enriched pathways was performed by Gene Set Enrichment Analysis (GSEA) with 10,000 permutations (66). The output was visualized using Enrichment Map with a Benjamini Hochberg FDR value of 0.25, and a *P*-value cut-off of 0.05 for all comparisons. Gene sets between 10 and 400 were included (67). Gene sets were visualized in heatmaps of the normalized expression generated using R (68) and the DittoSeq package (Bunis et al., 2021). Clustering was performed to organize samples and genes of similar expression (69). Second, we also analyzed sets of over-represented pathways for enrichment in protein functions. *De novo* Gene Ontology (GO) term assignments of predicted proteins were performed by InterProScan 5.26-65 (70). We kept the GO terms with a total term size in the genome of at least five. To test for enrichment, we performed hypergeometric tests. GO terms were only considered significant if the FDR *P*-values were less than <0.001. The R packages GSEABase (71) and GOstats (72) were used to analyze all enrichments, and the R package ggplot2 (73) was used to visualize the outcomes of enrichment tests. Gene expression in the samples for RNA-Seq was verified by qRT-PCR following published procedures (74). The primers for qPCR validation are listed in Table S4.

### RNA extraction and qPCR to analyze oxidative stress

Overnight grown wild type (H99) and *hapXΔ* cells (2.5 mL) in YPD media were washed thrice with low iron water and incubated in 50 mL of YNB media supplemented with BPS (150 μM) for 24 hours at 30°C and 140-150 rpm. Subsequently, cultures were collected, washed twice with YNB-BPS and 500 μL of cells resuspensions were treated in YNB-BPS for 1 hours at 30 C ∼220 rpm. Cells were harvested, washed with ice cold low iron water and pellets were frozen in liquid nitrogen and kept at -80°C. Total RNA was extracted from lysed cells using the RNeasy ® Mini kit (Qiagen) and DNA was removed using Turbo DNA-*free™* Kit (Invitrogen). Synthesis of cDNA was obtained using the High-Capacity cDNA Reverse Transcription Kit (Applied Biosystems ™) using oligo (dT). Real-time PCR (qPCR) was performed using the primers listed in Supplementary Table S5, cDNA and Green-2-Go qPCR Mastermix-low Rox (BioBasic) as described by the manufacturer. Samples were run on an Applied Biosystems 7500 Fast real-time PCR System. Relative gene expression was quantified using the ΔCT method normalized to *ACT1* gene. Statistical analysis was performed using two-way ANOVA test followed by a *post hoc* Šídák’s multiple comparison test.

### Flow cytometry analysis

Flow cytometric measurements were performed using CytoFLEX S (Beckman Coulter) Flow Cytometer equipped with four laser lines (405 nm, 488 nm, 561 nm and 633 nm) fitted with filters FITC (525/40) and PE (585/42). The number of cells measured per experiment was set to 30,000-40,000 unless otherwise stated. For the study of mitochondrial related ROS under the influence of ETC III inhibition, cells were grown overnight at ∼200 rpm and 30°C in YPD media and washed twice with PBS. Subsequently, 0.3 OD cells were grown on minimal media (YNB) for 24 hours and then treated in YNB media with and without the ETC III inhibitors antimycin A (AA, 50 μM) or myxothiazol (Myx, 7 μM) for 24 h at 200 rpm and 30°C. After treatment cells were stained with the intracellular reactive oxygen species (ROS) detector 2ʹ,7ʹ- Dichlorofluorescein Diacetate (DCFDA, 16 μM; Sigma-Aldrich) or Dihydroethidium (DHE, 2.5 μg/ml; EMD Millipore Corp.) for 1 h at 30°C. For the analysis upon ROS stressors, cells were grown on YPD media overnight as mentioned above. Next, cells were grown in YNB media and collected on log phase to subsequently treat them with or without hydrogen peroxide solution (H_2_O_2_, 5 mM; Sigma-Aldrich) for 1 hour at 30°C. Afterwards, cells were washed with PBS and stained with DCFDA as mentioned above. Data analysis and evaluation was conducted using FlowJo software version (10.8.2; 2006-2022). The gating strategy for cryptococcus cells is depicted in Fig. S11. Statistical analysis involved conducting a two-way ANOVA test, followed by *post hoc* Šídák’s or Tukey’s multiple comparison tests. GraphPad Prism software was utilized for the statistical analysis.

### Data availability

The RNA-Seq data have been deposited in Gene Expression Omnibus record GSE222564. The full gene lists for the RNA-Seq data for each condition are given in Supplemental Tables S6-S8

## ACKNOWLEDGMENTS

Research reported in this publication was supported by the National Institute of Allergy and Infectious Diseases of the National Institutes of Health under Award Number R01AI053721 (to J.W.K.). The content is solely the responsibility of the authors and does not necessarily represent the official views of the National Institutes of Health. J.W.K. is a Burroughs Wellcome Fund Scholar in Molecular Pathogenic Mycology and the Power Corporation fellow of the CIFAR program: Fungal Kingdom, Threats & Opportunities.

P.X., E.S.-L., W.J., and J.W.K. designed the study and P.X., E.S.-L., G.H. and B.B. performed the microbiological and transcriptome experiments. H.L. performed qRT-PCR experiments. P.X., E.S.-L., C.W.J.L. B.B., A.B. and J.W.K. performed data analysis. P.X., E.S.-L., W.J. and J.W.K. wrote the manuscript with input from all authors.

## REFERENCES

1. Ma H, Hagen F, Stekel DJ, Johnston SA, Sionov E, Falk R, Polacheck I, Boekhout T, May RC. 2009. The fatal fungal outbreak on Vancouver Island is characterized by enhanced intracellular parasitism driven by mitochondrial regulation. Proc Natl Acad Sci USA 106:12980–12985.

2. Calderone R, Li D, Traven A. 2015. System-level impact of mitochondria on fungal virulence: to metabolism and beyond. FEMS Yeast Res. 15(4):fov027.

3. Verma S, Shakya VPS, Idnurm A. 2018. Exploring and exploiting the connection between mitochondria and the virulence of human pathogenic fungi. Virulence. 9:426–446.

4. Black B, Lee C, Horianopoulos LC, Jung WH, Kronstad JW. 2021. Respiring to infect: Emerging links between mitochondria, the electron transport chain, and fungal pathogenesis. PLoS Pathog. 17:e1009661.

5. Kretschmer M, Damoo D, Sun S, Lee CW, Croll D, Brumer H, Kronstad J. 2022. Organic acids and glucose prime late-stage fungal biotrophy in maize. Science 376:1187–1191.

6. Chang AL, Doering TL. 2018. Maintenance of mitochondrial morphology in *Cryptococcus neoformans* is critical for stress resistance and virulence. MBio 9:e01375–18.

7. Chang AL, Kang Y, Doering TL. 2019. Cdk8 and Ssn801 regulate oxidative stress resistance and virulence in *Cryptococcus neoformans*. MBio 10:e02818–18.

8. Voelz K, Johnston SA, Smith LM, Hall RA, Idnurm A, May RC. 2014. ‘Division of labour’in response to host oxidative burst drives a fatal *Cryptococcus gattii* outbreak. Nat Commun 5:1–12.

9. Rajasingham R, Smith RM, Park BJ, Jarvis JN, Govender NP, Chiller TM, Denning DW, Loyse A, Boulware DR. 2017. Global burden of disease of HIV-associated cryptococcal meningitis: an updated analysis. Lancet Infect Dis 17:873–881.

10. Okurut S, Boulware DR, Olobo J, Meya DB. 2020. Landmark clinical observations and immunopathogenesis pathways linked to HIV and Cryptococcus fatal central nervous system co-infection. Mycoses 63:840–853.

11. Tugume L, Ssebambulidde K, Kasibante J, Ellis J, Wake RM, Gakuru J, Lawrence DS, Abassi M, Rajasingham R, Meya DB, Boulware DR. 2023. Cryptococcal meningitis. Nat Rev Dis Primers. 9:62.

12. Zaragoza O. 2019. Basic principles of the virulence of Cryptococcus. Virulence 10:490–501.

13. Liu S, Youngchim S, Zamith-Miranda D, Nosanchuk JD. 2021. Fungal melanin and the mammalian immune system. J Fungi 7:264.

14. García-Barbazán I, Torres-Cano A, García-Rodas R, Sachse M, Luque D, Megías D, Zaragoza O. 2024. Accumulation of endogenous free radicals is required to induce titan-like cell formation in *Cryptococcus neoformans*. mBio. 15:e0254923.

15. Trevijano-Contador N, Rossi SA, Alves E, Landín-Ferreiroa S, Zaragoza O. 2017. Capsule enlargement in *Cryptococcus neoformans* is dependent on mitochondrial activity. Front Microbiol 8:1423.

16. Pham NA, Robinson BH, Hedley DW. 2000. Simultaneous detection of mitochondrial respiratory chain activity and reactive oxygen in digitonin-permeabilized cells using flow cytometry. Cytometry A: 41:245–251.

17. Lenaz G. 2001. The mitochondrial production of reactive oxygen species: mechanisms and implications in human pathology. IUBMB Life. 52:159–164.

18. Brand MD, Nicholls DG. 2011. Assessing mitochondrial dysfunction in cells. Biochem J. 435:297–312.

19. Merryman M, Crigler J, Seipelt-Thiemann R, McClelland E. 2020. A mutation in *C. neoformans* mitochondrial NADH dehydrogenase results in increased virulence in mice Virulence 11:1366–1378.

20. Caza M, Hu G, Price M, Perfect JR, Kronstad JW. 2016. The zinc finger protein Mig1 regulates mitochondrial function and azole drug susceptibility in the pathogenic fungus *Cryptococcus neoformans*. mSphere. 1:e00080–15.

21. Cheon SA, Thak EJ, Bahn Y-S, Kang HA. 2017. A novel bZIP protein, Gsb1, is required for oxidative stress response, mating, and virulence in the human pathogen *Cryptococcus neoformans*. Sci Rep 7:1–15.

22. Gao X, Fu Y, Sun S, Gu T, Li Y, Sun T, Li H, Du W, Suo C, Li C. 2022. Cryptococcal Hsf3 controls intramitochondrial ROS homeostasis by regulating the respiratory process. Nat Commun 13:1–16.

23. Jung WH, Saikia S, Hu GG, Wang J, Fung CK-Y, D’Souza C, White R, Kronstad JW. 2010. HapX positively and negatively regulates the transcriptional response to iron deprivation in *Cryptococcus neoformans*. PloS Pathog 6:e1001209.

24. Horianopoulos LC, Hu G, Caza M, Schmitt K, Overby P, Johnson JD, Valerius O, Braus GH, Kronstad JW. 2020. The novel J-domain protein Mrj1 is required for mitochondrial respiration and virulence in *Cryptococcus neoformans*. MBio 11:e01127–20.

25. Caza M, Hu G, Nielson ED, Cho M, Jung WH, Kronstad JW. 2018. The Sec1/Munc18 (SM) protein Vps45 is involved in iron uptake, mitochondrial function and virulence in the pathogenic fungus *Cryptococcus neoformans*. PLoS Pathog 14:e1007220.

26. Do E, Park S, Li MH, Wang JM, Ding C, Kronstad JW, Jung WH. 2018. The mitochondrial ABC transporter Atm1 plays a role in iron metabolism and virulence in the human fungal pathogen *Cryptococcus neoformans*. Med Mycol 56:458–468.

27. Garcia-Santamarina S, Uzarska MA, Festa RA, Lill R, Thiele DJ. 2017. *Cryptococcus neoformans* iron-sulfur protein biogenesis machinery is a novel layer of protection against Cu stress. mBio 8:e01742–17.

28. Jung WH, Sham A, White R, Kronstad JW. 2006. Iron regulation of the major virulence factors in the AIDS-associated pathogen *Cryptococcus neoformans*. PLoS Biol 4:e410.

29. Gupta M, Outten CE. 2020. Iron–sulfur cluster signaling: The common thread in fungal iron regulation. Curr Opin Chem Biol 55:189–201.

30. Misslinger M, Hortschansky P, Brakhage AA, Haas H. 2021. Fungal iron homeostasis with a focus on *Aspergillus fumigatus*. Biochim Biophys Acta Mol Cell Res 1868:118885.

31. Do E, Cho YJ, Kim D, Kronstad JW, Jung WH. 2020. A transcriptional regulatory map of iron homeostasis reveals a new control circuit for capsule formation in *Cryptococcus neoformans*. Genetics 215:1171–1189.

32. Wang Y, Aisen P, Casadevall A. 1995. *Cryptococcus neoformans* melanin and virulence: mechanism of action. Infect Immun 63:3131–3136.

33. Kwon-Chung K, Tom W, Costa J. 1983. Utilization of indole compounds by *Cryptococcus neoformans* to produce a melanin-like pigment. J Clin Microbiol 18:1419–1421.

34. Panepinto JC, Williamson PR. 2006. Intersection of fungal fitness and virulence in *Cryptococcus neoformans*. FEMS Yeast Res 6:489–498.

35. Attarian R, Hu GG, Sánchez-León E, Caza M, Croll D, Do E, Bach H, Missall T, Lodge J, Jung WH. 2018. The monothiol glutaredoxin Grx4 regulates iron homeostasis and virulence in *Cryptococcus neoformans*. MBio 9:e02377–18.

36. Dlouhy AC, Beaudoin J, Labbé S, Outten CE. 2017. *Schizosaccharomyces pombe* Grx4 regulates the transcriptional repressor Php4 via [2Fe–2S] cluster binding. Metallomics 9:1096–1105.

37. Jung WH, Sánchez-León E, Kronstad JW. 2021. Coordinated regulation of iron metabolism in *Cryptococcus neoformans* by GATA and CCAAT transcription factors: connections with virulence. Curr Genet 67:583–593.

38. O’Meara TR, Norton D, Price MS, Hay C, Clements MF, Nichols CB, Alspaugh JA. 2010. Interaction of *Cryptococcus neoformans* Rim101 and protein kinase A regulates capsule. PLoS Pathog 6:e1000776.

39. O’Meara TR, Xu W, Selvig KM, O’Meara MJ, Mitchell AP, Alspaugh JA. 2014. The *Cryptococcus neoformans* Rim101 transcription factor directly regulates genes required for adaptation to the host. Mol Cell Biol 34:673–684.

40. Idnurm A. 2011. New surprises from within the black box of fungal melanization. Virulence. 2:261–263.

41. Smith DFQ, Casadevall A. 2019. The role of melanin in fungal pathogenesis for animal hosts. Curr Top Microbiol Immunol. 422:1–30.

42. Lee D, Jang E-H, Lee M, Kim S-W, Lee Y, Lee K-T, Bahn Y-S. 2019. Unraveling melanin biosynthesis and signaling networks in *Cryptococcus neoformans*. mBio 10:e02267–19.

43. Bolter CJ, Chefurka W. 1990. Extramitochondrial release of hydrogen peroxide from insect and mouse liver mitochondria using the respiratory inhibitors phosphine, myxothiazol, and antimycin and spectral analysis of inhibited cytochromes. Arch Biochem Biophys. 278:65–72.

44. Monzel AS, Enríquez JA, Picard M. 2023. Multifaceted mitochondria: moving mitochondrial science beyond function and dysfunction. Nat Metab. 5:546–562.

45. Upadhya R, Campbell LT, Donlin MJ, Aurora R, Lodge JK. 2013. Global transcriptome profile of *Cryptococcus neoformans* during exposure to hydrogen peroxide induced oxidative stress. PloS One 8:e55110.

46. Shehat MG, Tigno-Aranjuez J. 2019. Flow cytometric measurement of ROS production in Macrophages in response to FcγR cross-linking. J Vis Exp. 7:10.3791/59167.

47. Casadevall A, Coelho C, Cordero RJB, Dragotakes Q, Jung E, Vij R, Wear MP. 2019. The capsule of *Cryptococcus neoformans*. Virulence. 10:822–831.

48. Jacobson E, Compton G. 1996. Discordant regulation of phenoloxidase and capsular polysaccharide in *Cryptococcus neoformans*. J Med Vet Mycol 34:289–291.

49. Zhu X, Williamson PR. 2004. Role of laccase in the biology and virulence of Cryptococcus neoformans. FEMS Yeast Res 5:1–10.

50. Vartivarian SE, Anaissie EJ, Cowart RE, Sprigg HA, Tingler MJ, Jacobson ES. 1993. Regulation of cryptococcal capsular polysaccharide by iron. J Infect Dis 167:186–190.

51. Cordero RJ, Camacho E, Casadevall A. 2020. Melanization in *Cryptococcus neoformans* requires complex regulation. mBio 11:e03313–19.

52. Hortschansky P, Haas H, Huber EM, Groll M, Brakhage AA. 2017. The CCAAT-binding complex (CBC) in *Aspergillus* species. Biochim Biophys Acta Gene Regul Mech. 1860:560–570.

53. Chevtzoff C, Yoboue ED, Galinier A, Casteilla L, Daignan-Fornier B, Rigoulet M, Devin A. 2010. Reactive oxygen species-mediated regulation of mitochondrial biogenesis in the yeast *Saccharomyces cerevisiae*. J Biol Chem. 285:1733–1742.

54. Giles SS, Stajich JE, Nichols C, Gerrald QD, Alspaugh JA, Dietrich F, Perfect JR. 2006. The *Cryptococcus neoformans* catalase gene family and its role in antioxidant defense. Eukaryot Cell. 5:1447–1459.

55. Warris A, Ballou ER. 2019. Oxidative responses and fungal infection biology. Semin Cell Dev Biol. 89:34–46.

56. Berg SZ, Berg J. 2023. Melanin: a unifying theory of disease as exemplified by Parkinson’s, Alzheimer’s, and Lewy body dementia. Front Immunol. 14:1228530.

57. Billings JL, Gordon SL, Rawling T, Doble PA, Bush AI, Adlard PA, Finkelstein DI, Hare DJ. 2019. l-3, 4-dihydroxyphenylalanine (l-DOPA) modulates brain iron, dopaminergic neurodegeneration and motor dysfunction in iron overload and mutant alpha-synuclein mouse models of Parkinson’s disease. J Neurochem 150:88–106.

58. Burbulla LF, Song P, Mazzulli JR, Zampese E, Wong YC, Jeon S, Santos DP, Blanz J, Obermaier CD, Strojny C. 2017. Dopamine oxidation mediates mitochondrial and lysosomal dysfunction in Parkinson’s disease. Science 357:1255–1261.

59. Graves SM, Xie Z, Stout KA, Zampese E, Burbulla LF, Shih JC, Kondapalli J, Patriarchi T, Tian L, Brichta L. 2020. Dopamine metabolism by a monoamine oxidase mitochondrial shuttle activates the electron transport chain. Nature Neurosci 23:15–20.

60. Eisenman HC, Chow S-K, Tsé KK, McClelland E, Casadevall A. 2011. The effect of L-DOPA on *Cryptococcus neoformans* growth and gene expression. Virulence 2:329–336.

61. Idnurm A. 2011. New surprises from within the black box of fungal melanization. Virulence. 2:261–263.

62. Hu G, Horianopoulos L, Sánchez-León E, Caza M, Jung W, Kronstad JW. 2021. The monothiol glutaredoxin Grx4 influences thermotolerance, cell wall integrity, and Mpk1 signaling in *Cryptococcus neoformans*. G3 (Bethesda) 11:jkab322.

63. Hu G, Caza M, Cadieux B, Chan V, Liu V, Kronstad J. 2013. *Cryptococcus neoformans* requires the ESCRT protein Vps23 for iron acquisition from heme, for capsule formation, and for virulence. Infect Immun. 81:292–302.

64. Hu G, Caza M, Cadieux B, Bakkeren E, Do E, Jung WH, Kronstad JW. 2015. The endosomal sorting complex required for transport machinery influences haem uptake and capsule elaboration in *Cryptococcus neoformans*. Mol Microbiol 96:973–992.

65. Love MI, Huber W, Anders S. 2014. Moderated estimation of fold change and dispersion for RNA-seq data with DESeq2. Genome Biol 15:1–21.

66. Subramanian A, Tamayo P, Mootha VK, Mukherjee S, Ebert BL, Gillette MA, Paulovich A, Pomeroy SL, Golub TR, Lander ES. 2005. Gene set enrichment analysis: a knowledge-based approach for interpreting genome-wide expression profiles. Proc Natl Acad Sci USA 102:15545–15550.

67. Merico D, Isserlin R, Stueker O, Emili A, Bader GD. 2010. Enrichment map: a network-based method for gene-set enrichment visualization and interpretation. PloS One 5:e13984.

68. Team RC. 2022. R: A Language and Environment for Statistical Computing. R Foundation for Statistical Computing, Vienna, Austria. https://www.R-project.org/.

69. Bunis DG, Andrews J, Fragiadakis GK, Burt TD, Sirota M. 2021. dittoSeq: universal user-friendly single-cell and bulk RNA sequencing visualization toolkit. Bioinform 36:5535–5536.

70. Jones P, Binns D, Chang H-Y, Fraser M, Li W, McAnulla C, McWilliam H, Maslen J, Mitchell A, Nuka G. 2014. InterProScan 5: genome-scale protein function classification. Bioinform 30:1236–1240.

71. Martin M, Seth F, Robert G. 2021. GSEABase: Gene set enrichment data structures and methods. R package version 1.56.0.

72. Falcon S, Gentleman R. 2007. Using GOstats to test gene lists for GO term association. Bioinform 23:257–258.

73. Wickham H. 2009. ggplot2; Elegant graphics for data analysis. P 1-213. In Applied Spatial Data Analysis R. Springer New York, NY.

74. Horianopoulos LC, Lee CW, Schmitt K, Valerius O, Hu G, Caza M, Braus GH, Kronstad JW. 2021. A J domain protein functions as a histone chaperone to maintain genome integrity and the response to DNA damage in a human fungal pathogen. mbio 12:e03273–21.

