## Supplemental Figures for "The interplay between electron transport chain function and iron regulatory factors influences melanin formation in *Cryptococcus neoformans*"

### Figure S1

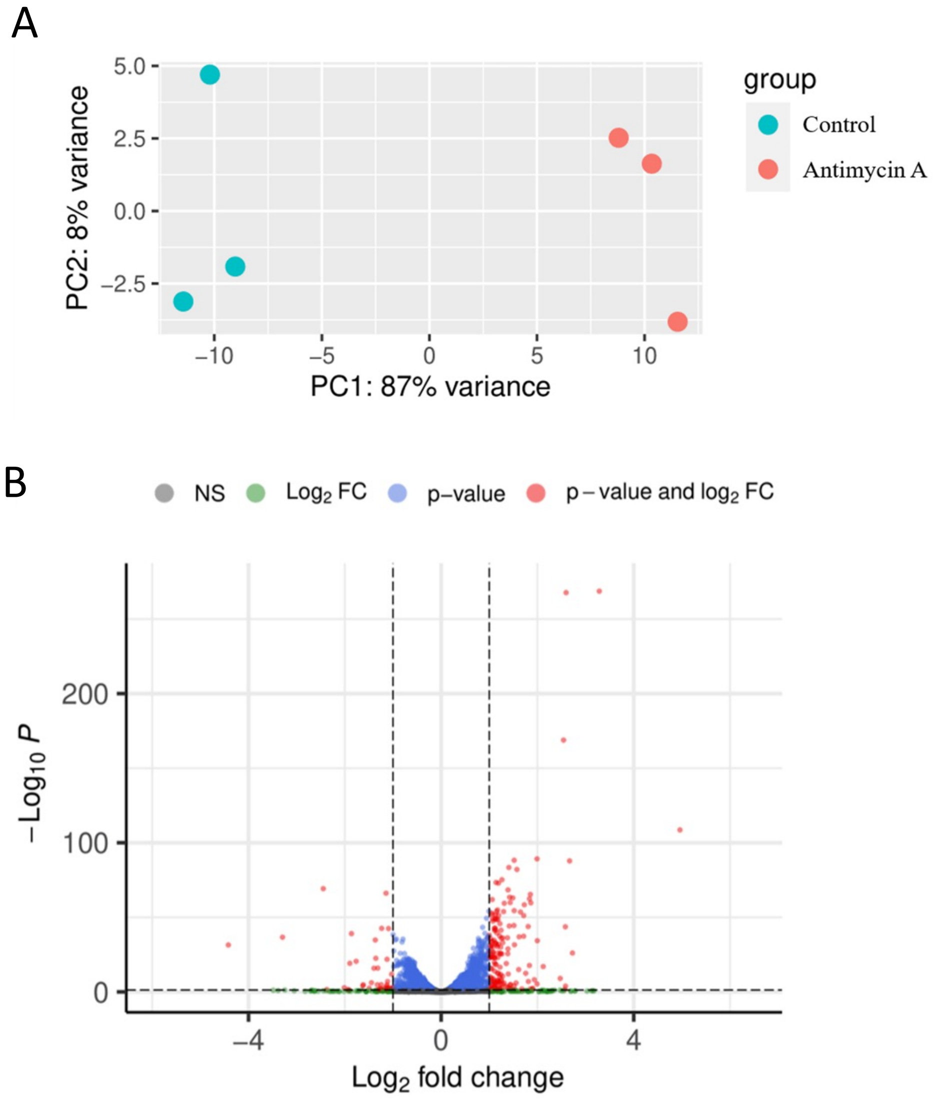


**Figure S1.** The transcriptional response of WT cells to the ETC complex III inhibitor antimycin A. (**A**) Principal Component Analysis (PCA) of the comparison of WT (control) versus antimycin A treated cells showing distinct clustering between the two groups. (**B**) Volcano plot of the differentially expressed genes from (A) displaying a log2 fold change cut off of 1 and revealing upregulation of genes in H99 upon treatment with Antimycin A.

### Figure S2

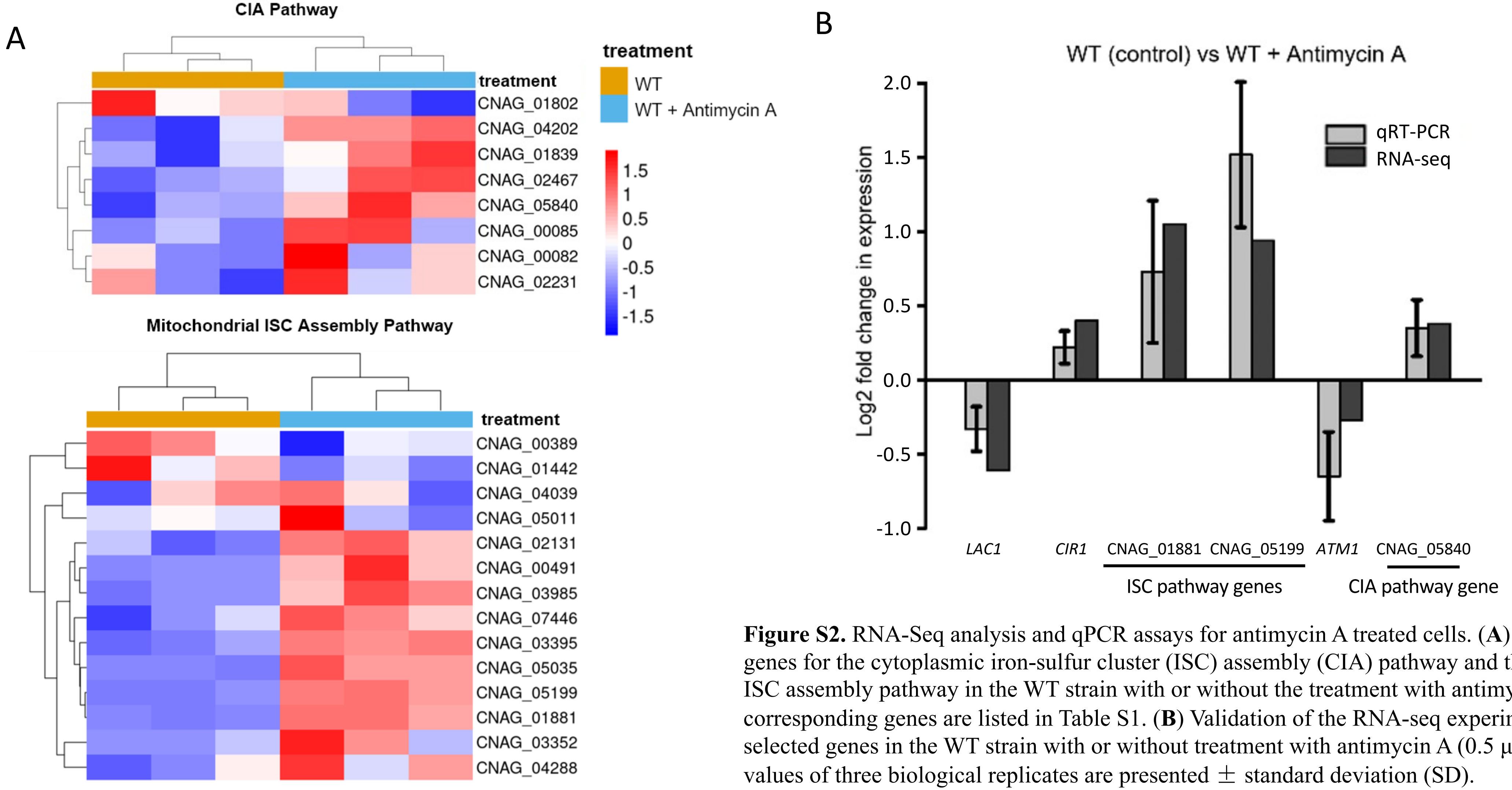


**Figure S2.** RNA-Seq analysis and qPCR assays for antimycin A treated cells. (**A**) Heatmaps of genes for the cytoplasmic iron-sulfur cluster (ISC) assembly (CIA) pathway and the mitochondrial ISC assembly pathway in the WT strain with or without the treatment with antimycin A. The corresponding genes are listed in Table S1. (**B**) Validation of the RNA-seq experiment by qPCR for selected genes in the WT strain with or without treatment with antimycin A (0.5 µg/ml). Mean values of three biological replicates are presented ± standard deviation (SD).

### Figure S3

#### WT vs WT + Antimycin A

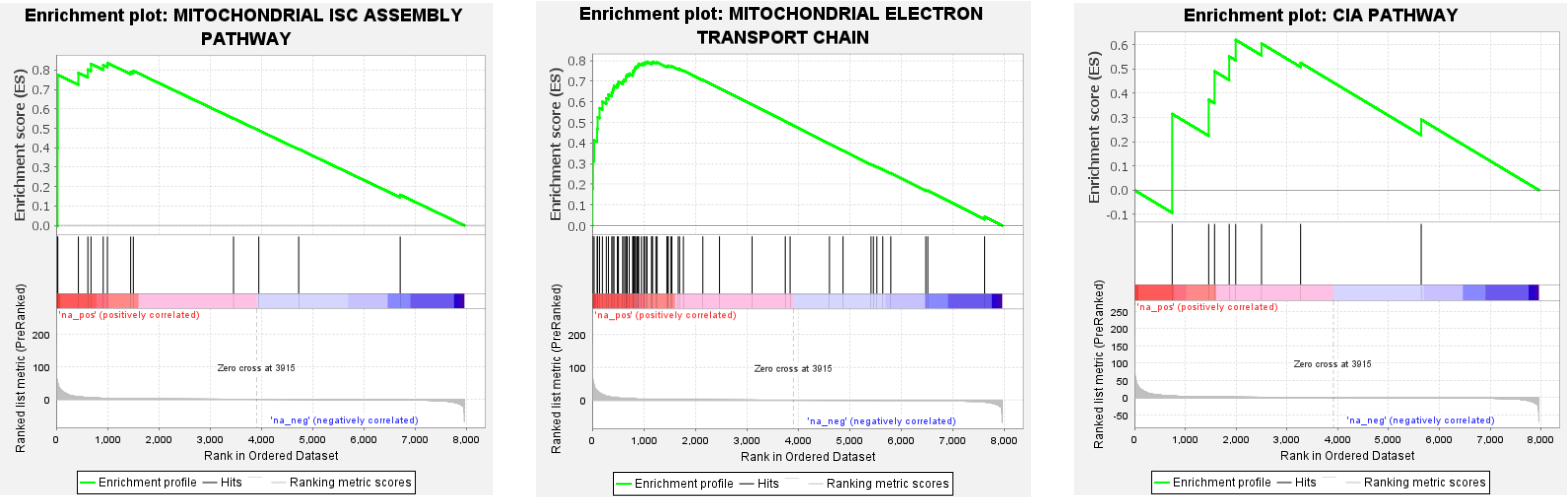


| | |
|---|---|
| Enrichment Score (ES) | 0.8361732 |
| Normalized Enrichment Score (NES) | 1.4800994 |
| Nominal p-value | 0.029363785 |
| FDR q-value | 0.036047716 |
| FWER p-Value | 0.049 |

| | |
|---|---|
| Enrichment Score (ES) | 0.7942508 |
| Normalized Enrichment Score (NES) | 1.6800591 |
| Nominal p-value | 0.0012771392 |
| FDR q-value | 0.0010373443 |
| FWER p-Value | 0.001 |

| | |
|---|---|
| Enrichment Score (ES) | 0.62017757 |
| Normalized Enrichment Score (NES) | 1.0088847 |
| Nominal p-value | 0.5062389 |
| FDR q-value | 0.48392132 |
| FWER p-Value | 0.695 |

**Figure S3.** GSEA enrichment curves of the mitochondrial ISC assembly, ETC, and CIA KEGG pathways showing positive enrichment upon antimycin A treatment. A pre-ranked GSEA was run using 1000 permutations and gene sets larger than 5.

### Figure S4

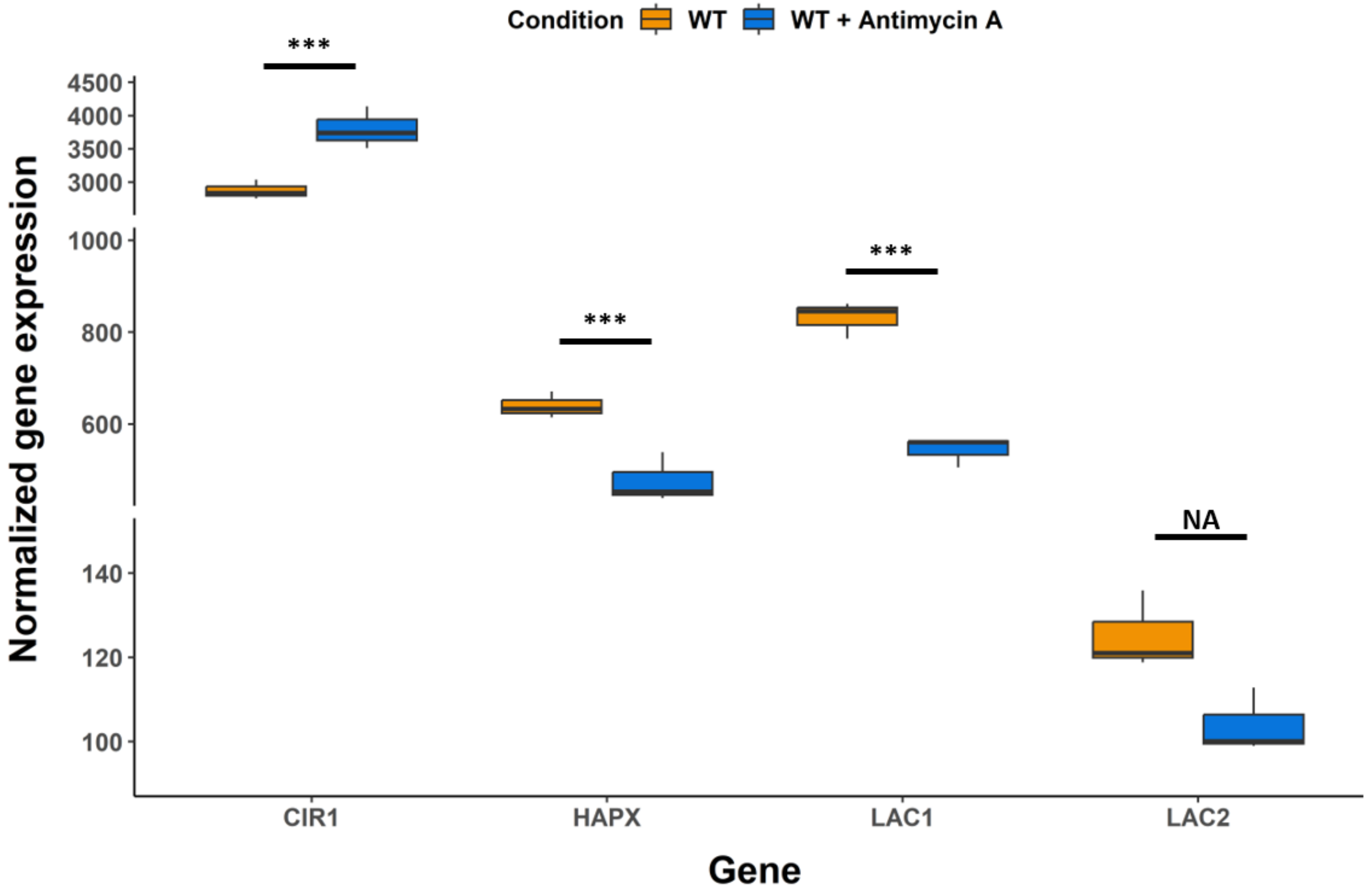


**Figure S4.** RNA-Seq measurements of transcripts for the laccase and transcription factor genes. The normalized gene expression values are shown for the *LAC1*, *LAC2*, *CIR1* and *HAPX* genes in the WT strain with or without treatment with antimycin A (0.5 µg/ml). The differences in transcript levels were significant as determined by the Walds test (***, $P < 0.001$). n=3. Boxes extend from the 25th to the 75th percentile of each group's distribution with the lower and upper whiskers representing the 5th and 95th percentiles respectively.

### Figure S5

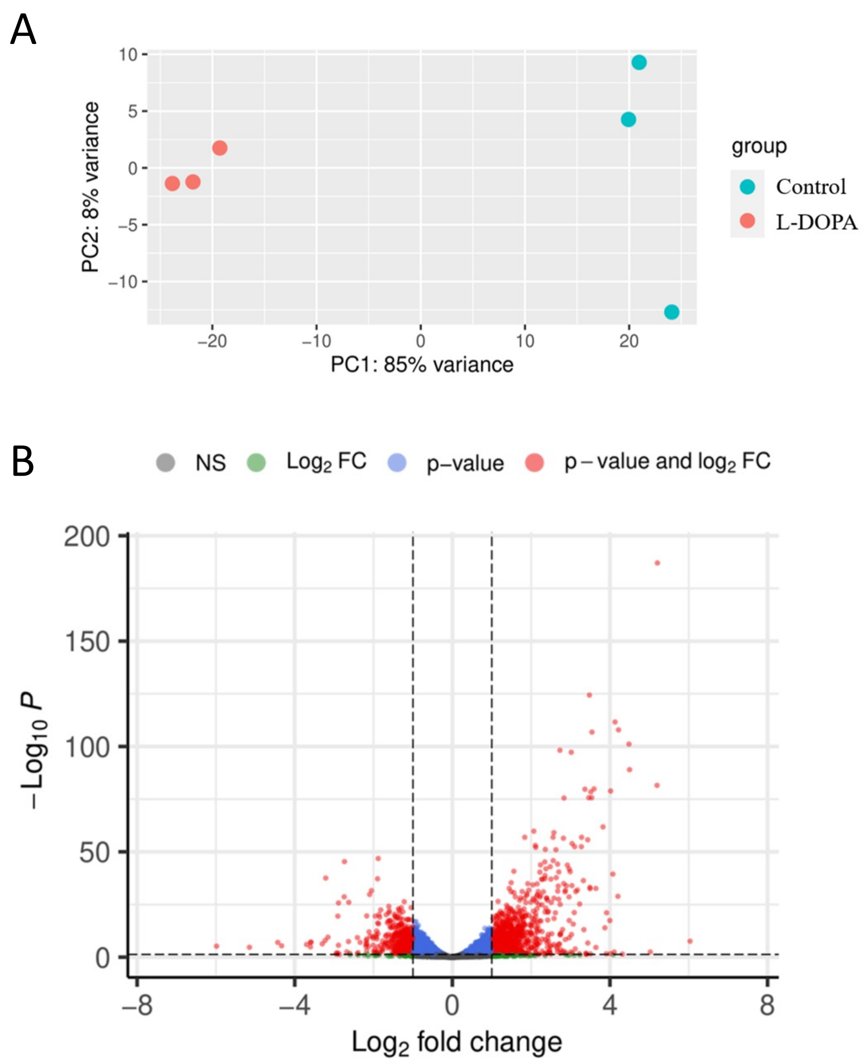


**Figure S5.** The transcriptional response of WT cells to the melanin substrate L-DOPA. (**A**) Principal Component Analysis (PCA) of the comparison of WT (control) versus L-DOPA-treated cells revealing distinct clustering between the two groups. (**B**) Volcano plot of the differentially expressed genes from (A) displaying a log2 fold change cut off of 1 and revealing an upregulated transcriptional response when wildtype H99 is treated with L-DOPA.

### Figure S6

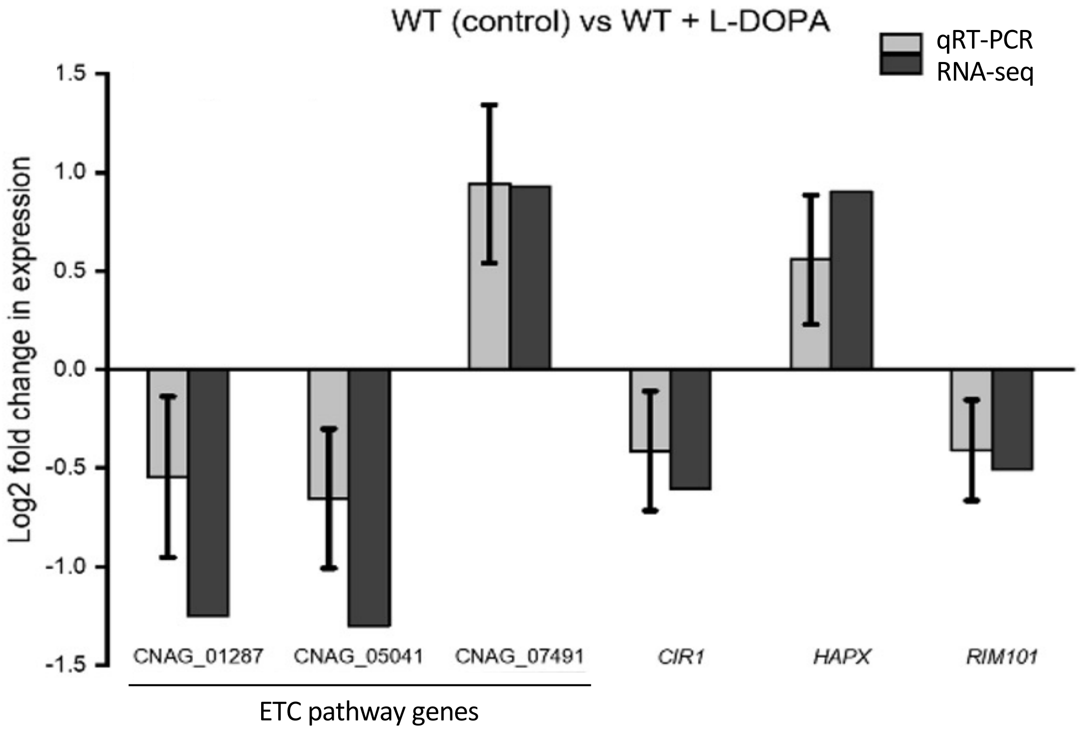


**Figure S6.** Validation of the RNA-seq analysis of the response to L-DOPA by qPCR. The analyzed genes included components of the ETC, as listed in Table S2, and the transcription factors *CIR1*, *HAPX* and *RIM101* employed in the analysis in Fig. 1. Mean values of three biological replicates are indicated ± standard deviation (SD).

### Figure S7

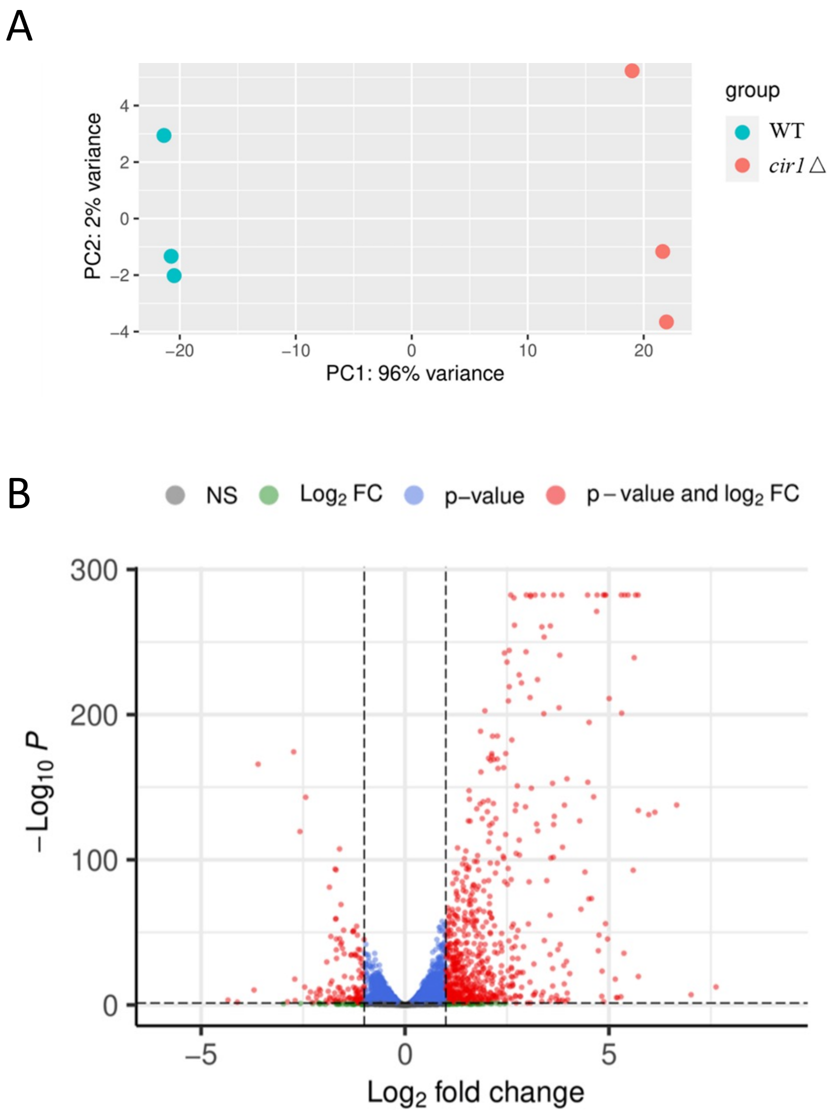


**Figure S7.** Impact of loss of Cir1 on transcript levels for the laccases, components of mitochondrial ISC assembly and the ETC, and the iron regulators HapX and Rim101. (**A**) Principal Component Analysis (PCA) of the comparison of WT (control) versus *cir1D* mutant cells revealed unique clustering between strains and therefore distinct transcriptional profiles. (**B**) Volcano plot of the differentially expressed genes from (A) displaying a log2 fold change cut off of 1 and indicating that loss of Cir1 results in an upregulated transcriptional response.

Figure S8

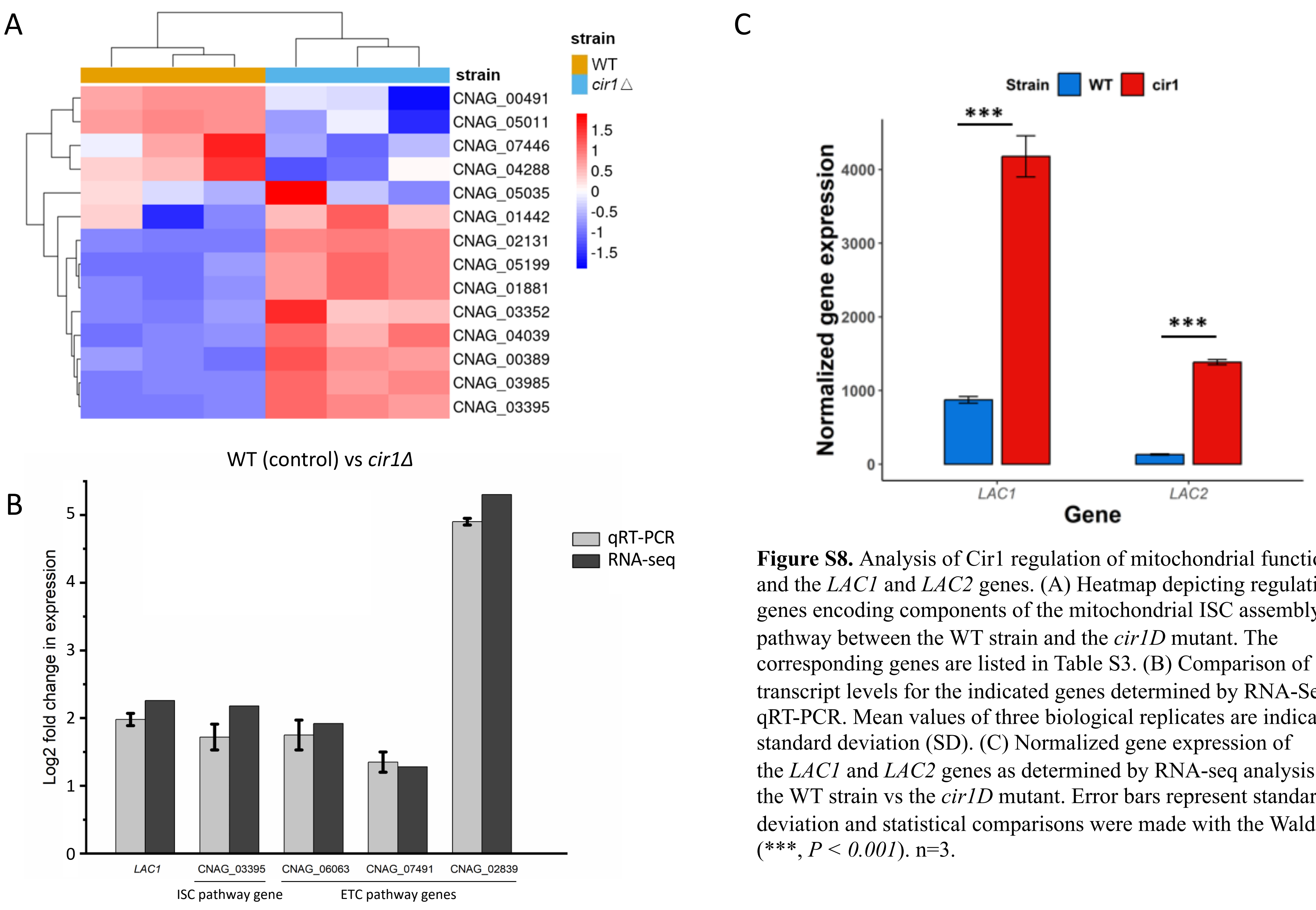


**Figure S8.** Analysis of Cir1 regulation of mitochondrial functions and the *LAC1* and *LAC2* genes. (A) Heatmap depicting regulation of genes encoding components of the mitochondrial ISC assembly pathway between the WT strain and the *cir1D* mutant. The corresponding genes are listed in Table S3. (B) Comparison of transcript levels for the indicated genes determined by RNA-Seq and qRT-PCR. Mean values of three biological replicates are indicated ± standard deviation (SD). (C) Normalized gene expression of the *LAC1* and *LAC2* genes as determined by RNA-seq analysis for the WT strain vs the *cir1D* mutant. Error bars represent standard deviation and statistical comparisons were made with the Walds test (***, $P < 0.001$). n=3.

Figure S9

**WT vs *cir1Δ***

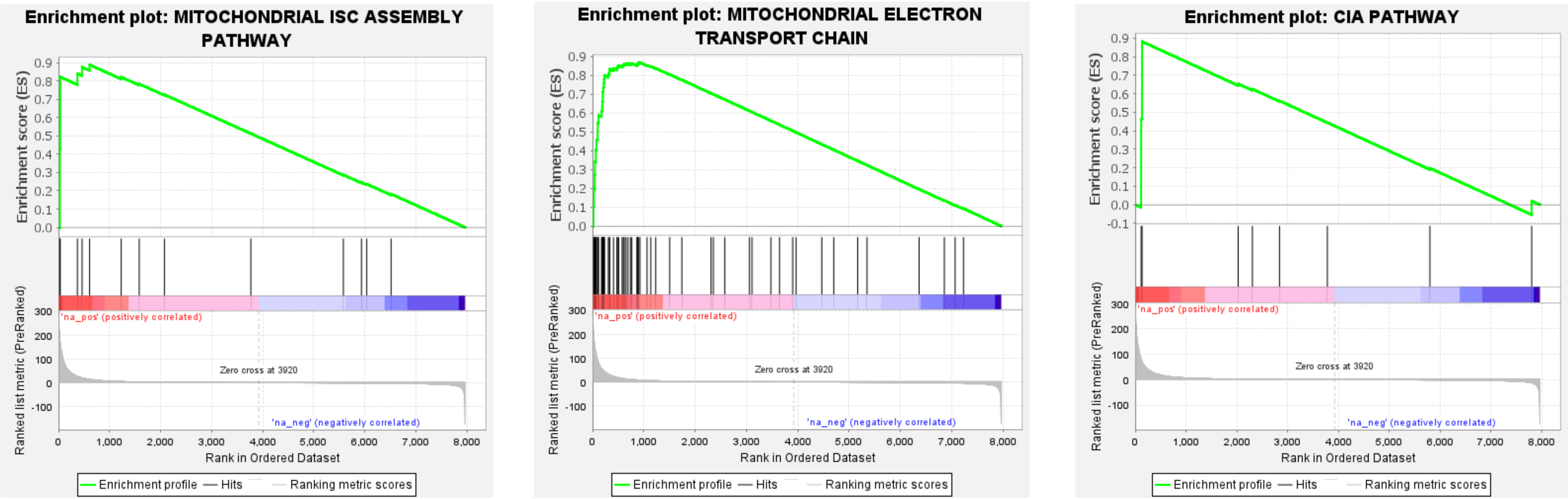


| | |
|---|---|
| Enrichment Score (ES) | 0.8879621 |
| Normalized Enrichment Score (NES) | 1.4434943 |
| Nominal p-value | 0.044604316 |
| FDR q-value | 0.040070567 |
| FWER p-Value | 0.064 |

| | |
|---|---|
| Enrichment Score (ES) | 0.86723256 |
| Normalized Enrichment Score (NES) | 1.5415058 |
| Nominal p-value | 0.0 |
| FDR q-value | 0.0010080645 |
| FWER p-Value | 0.001 |

| | |
|---|---|
| Enrichment Score (ES) | 0.8820039 |
| Normalized Enrichment Score (NES) | 1.3756471 |
| Nominal p-value | 0.10622155 |
| FDR q-value | 0.06804432 |
| FWER p-Value | 0.157 |

**Figure S9.** GSEA enrichment curves of the mitochondrial ISC assembly, ETC, and CIA KEGG pathways showing positive enrichment in the *cir1* mutant. A pre-ranked GSEA was run using 1000 permutations and gene sets larger than 5.

### Figure S10

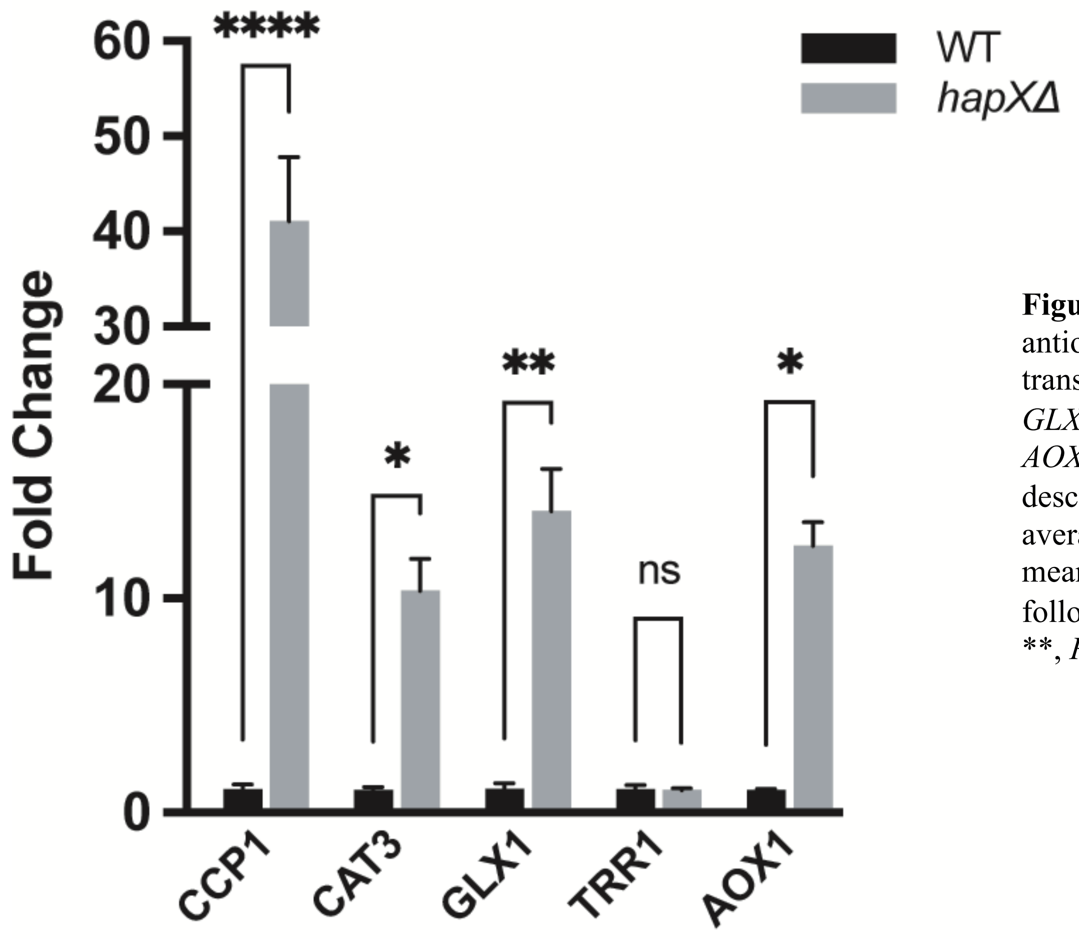


**Figure S10.** Loss of HapX derepresses the transcription of genes for antioxidant functions. qRT-PCR was employed to compare the transcript levels for *CCP1* (cytochrome C peroxidase), *CAT3* (catalase), *GLX1* (lactoylglutathione lyase), *TRR1* (thioredoxin reductase) and *AOX1* (alternative oxidase) in the WT strain and the *hapXD* as described in the Materials and Methods. Fold change represent the averages from three biological replicates ± standard errors of the means. Statistical analysis was performed using two-way ANOVA test followed by a *post hoc* Šídák's multiple comparison test (*, $P < 0.05$; **, $P < 0.01$ ****; $P < 0.0001$). ns: not significant.

### Figure S11

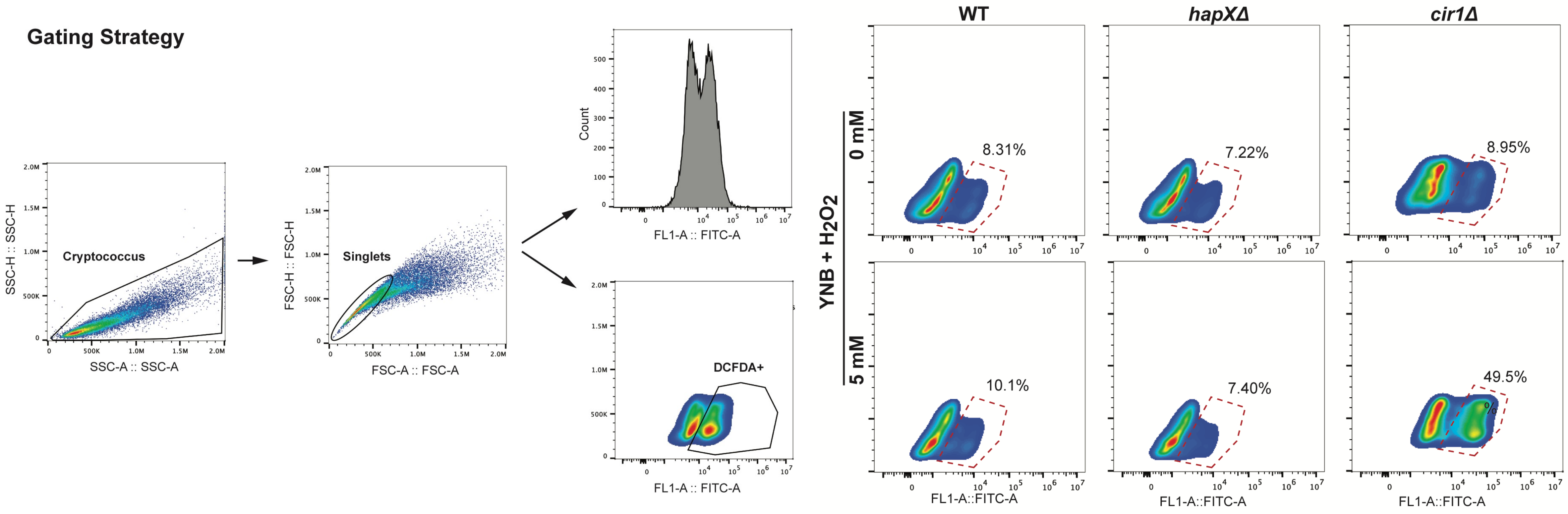


**Figure S11.** Gating strategy for the analysis the response to hydrogen peroxide. The cells were grown on YPD media overnight at 30°C and ~200 rpm. The cells were exposed to hydrogen peroxide ($H_2O_2$, 5 mM; Sigma-Aldrich) in YNB medium for 1 hour at the same temperature, washed with phosphate-buffered saline and stained with DCFDA for flow cytometry analysis as described in the Materials and Methods. Singlets were selected from the cell population and smooth dot plots were used to determine percentage of DCFDA positive cells. Data analysis was conducted using FlowJo software version (10.8.2; 2006-2022).
