## Supplemental Tables for "The interplay between electron transport chain function and iron regulatory factors influences melanin formation in *Cryptococcus neoformans*"

**TABLE S1** The effect of antimycin A on the gene expression of components of the mitochondrial ISC assembly pathway, the CIA pathway and the ETC (KEGG) for the WT strain. The log2 fold change (experimental group/control group) in bold was significantly different (*P* < 0.05).

| Gene ID | Gene name | Function | WT (control) vs WT + antimycin A |
|---|---|---|---|
| **Mitochondrial ISC assembly pathway** | | | |
| CNAG_00389 | *IBA57* | Mitochondrial protein | 0.04 |
| CNAG_01442 | *NFS1* | Cysteine desulferase | **-0.183** |
| CNAG_04039 | *FRR3* | Scaffold protein | -0.001 |
| CNAG_05011 | *FRR4* | Frataxin | 0.031 |
| CNAG_02131 | *ISA1* | Iron sulfur assembly protein 1 | **0.328** |
| CNAG_00491 | *ISA2* | Iron sulfur assembly protein 2 | **0.304** |
| CNAG_03985 | *GRX5* | Monothiol glutaredoxin-5 | **0.297** |
| CNAG_07446 | *ISD11* | Mitochondrial protein | **0.249** |
| CNAG_03395 | *NFU1* | NifU-like protein C | **0.329** |
| CNAG_05035 | *ARH1* | Ferredoxin-NADP reductase | **0.472** |
| CNAG_05199 | *SSQ1* | HSP70-type chaperone | **0.943** |
| CNAG_01881 | *MGE1* | Nucleotide exchange factor co-chaperone | **1.048** |
| CNAG_03352 | *IND1* | Hypothetical protein | **0.2** |
| CNAG_04288 | *JAC1* | J-domain co-chaperone | **0.218** |
| **CIA pathway** | | | |
| CNAG_01802 | *DRE2* | Cytoplasmic protein | -0.102 |
| CNAG_04202 | *NAR1* | Iron hydrogenase | 0.134 |
| CNAG_01839 | *TAH18* | Peptidyl-prolyl cis-trans isomerase NIMA- interacting 1 | **0.154** |
| CNAG_02467 | *MMS19* | DNA repair/transcription protein | **0.142** |
| CNAG_05840 | *CIA2* | Hypothetical protein | **0.379** |
| CNAG_00085 | *CIA1* | Histone chaperone | 0.133 |
| CNAG_00082 | *CFD1* | Cytosolic Fe-S cluster assembly factor | 0.086 |
| CNAG_02231 | *NBP35* | Cytosolic Fe-S cluster assembly factor NBP35 | 0.036 |
| **Mitochondrial ETC** | | | |
| CNAG_03958 | | Hypothetical protein | **0.251** |
| CNAG_03629 | *YHB1* | NADH dehydrogenase (quinone), G subunit | **0.159** |
| CNAG_07491 | | Glutaredoxin | **0.769** |
| CNAG_03676 | | D-lactate dehydrogenase | **0.226** |
| CNAG_01769 | | Solute carrier family 25 (mitochondrial aspartate/glutamate transporter), member 12/13 | **0.396** |

| | | | |
|---|---|---|---|
| CNAG_03985 | *GRX5* | Grx4 family monothiol glutaredoxin | **0.297** |
| CNAG_01287 | | NADH-ubiquinone oxidoreductase 51 kDa subunit, mitochondrial | **0.216** |
| CNAG_00486 | | Succinate dehydrogenase assembly factor 2, mitochondrial | **0.430** |
| CNAG_06806 | *ETF1* | Electron transfer flavoprotein alpha subunit | **0.204** |
| CNAG_01005 | | Glutaredoxin | **0.256** |
| CNAG_03861 | | Cytochrome c oxidase subunit 5a | **0.391** |
| CNAG_06723 | | Succinate dehydrogenase (ubiquinone) membrane anchor subunit | **0.647** |
| CNAG_07356 | | Succinate dehydrogenase, cytochrome b556 subunit | **0.679** |
| CNAG_03226 | | Succinate dehydrogenase [ubiquinone] iron-sulfur subunit, mitochondrial | **1.076** |
| CNAG_00149 | | NADH dehydrogenase (ubiquinone) 1 alpha subcomplex 4 | **0.783** |
| CNAG_04189 | *SDH1* | Succinate dehydrogenase [ubiquinone] flavoprotein subunit, mitochondrial | **1.060** |
| CNAG_00716 | *CYC7* | Cytochrome c | **0.670** |
| CNAG_06063 | | Cytochrome c oxidase assembly protein subunit 15 | **1.407** |
| CNAG_02140 | | NADH dehydrogenase (ubiquinone) 1 alpha subcomplex 12 | **0.133** |
| CNAG_01870 | | Electron transfer flavoprotein beta subunit | **0.333** |
| CNAG_03364 | | Hypothetical protein | 0.162 |
| CNAG_05041 | | NADH-ubiquinone oxidoreductase subunit 8 | **0.302** |
| CNAG_05631 | | NADH-ubiquinone oxidoreductase 49 kDa subunit, mitochondrial | **0.187** |
| CNAG_00768 | | Hypothetical protein | **0.296** |
| CNAG_02938 | | Cytochrome c oxidase subunit 6a | **0.290** |
| CNAG_05633 | | Ubiquinol-cytochrome c reductase subunit 9 | **0.341** |
| CNAG_05069 | | Ubiquinol-cytochrome c reductase subunit 10 | **0.510** |
| CNAG_05909 | *CYT1* | Cytochrome c1, heme protein | **0.440** |
| CNAG_01470 | | NADH dehydrogenase (ubiquinone) flavoprotein 2 | **0.255** |
| CNAG_00462 | | Electron-transferring-flavoprotein dehydrogenase | **0.169** |
| CNAG_03848 | | Glutathione transferase | 0.253 |
| CNAG_01443 | | Hypothetical protein | 0.138 |
| CNAG_02664 | | D-lactate dehydrogenase | 0.003 |
| CNAG_06180 | | NADH dehydrogenase (ubiquinone) Fe-S protein 4 | 0.050 |
| CNAG_05197 | | NADH dehydrogenase (ubiquinone) 1 alpha subcomplex 8 | 0.078 |
| CNAG_01799 | | Ubiquinol-cytochrome c reductase subunit 8 | **0.162** |

| | | | |
|---|---|---|---|
| CNAG_01323 | *QCR7* | Ubiquinol-cytochrome c reductase subunit 7 | **0.169** |
| CNAG_03507 | | Mitochondrial-processing peptidase subunit beta | **0.234** |
| CNAG_03359 | | Cytochrome c oxidase subunit 7 | **0.526** |
| CNAG_06407 | | Cytochrome c oxidase subunit 7c | **0.343** |
| CNAG_06012 | | NADH-cytochrome b5 reductase 1 | **0.217** |
| CNAG_01485 | | D-lactate dehydrogenase (cytochrome) | **0.159** |
| CNAG_06663 | | Ubiquinol-cytochrome c reductase subunit 6 | **0.283** |
| CNAG_06226 | | NADH dehydrogenase (ubiquinone) 1 alpha subcomplex 5 | **0.251** |
| CNAG_05132 | | Cytochrome c oxidase subunit 5b | **0.314** |
| CNAG_05179 | *QCR2* | ubiquinol-cytochrome c reductase core subunit 2 | **0.215** |
| CNAG_01991 | | Cytochrome c oxidase subunit 4 | **0.269** |
| CNAG_02315 | | Ubiquinol-cytochrome c reductase, iron-sulfur subunit | **0.284** |
| CNAG_01800 | | Hypothetical protein | **0.306** |
| CNAG_04290 | | Hypothetical protein | 0.132 |
| CNAG_06138 | | NADH dehydrogenase (ubiquinone) Fe-S protein 6 | **0.206** |
| CNAG_02849 | | Glutathione transferase | -0.283 |
| CNAG_02840 | | Hypothetical protein | **-0.564** |
| CNAG_03874 | | Oxidoreductase | **-0.190** |
| CNAG_07674 | | Hypothetical protein | -0.168 |
| CNAG_07365 | | Oxidoreductase | 0.011 |
| CNAG_02839 | | Hypothetical protein | -0.300 |
| CNAG_03589 | | Adrenodoxin-type ferredoxin | -0.035 |
| CNAG_02950 | *GRX3* | Grx4 family monothiol glutaredoxin | -0.120 |
| CNAG_01802 | | Fe-S cluster assembly protein | -0.102 |
| CNAG_04521 | | Oxidoreductase | **-0.504** |
| CNAG_05554 | | Cytochrome b5 | -0.088 |

**TABLE S2** The impact of L-DOPA on gene expression for components of the mitochondrial ISC assembly pathway, the CIA pathway and the ETC (KEGG) in the WT strain. The log2 fold change (experimental group/control group) in bold was significantly different ($P < 0.05$).

| Gene ID | Gene name | Function | WT (control) vs WT + L-DOPA |
|---|---|---|---|
| **Mitochondrial ISC assembly pathway** | | | |
| CNAG_00389 | *IBA57* | Mitochondrial protein | **-0.468** |
| CNAG_01442 | *NFS1* | Cysteine desulferase | 0.202 |
| CNAG_04039 | *FRR3* | Scaffold protein | **0.459** |

| | | | |
|---|---|---|---|
| CNAG_05011 | *FRR4* | Frataxin | **-0.527** |
| CNAG_02131 | *ISA1* | Iron sulfur assembly protein 1 | **-0.655** |
| CNAG_00491 | *ISA2* | Iron sulfur assembly protein 2 | -0.166 |
| CNAG_03985 | *GRX5* | Monothiol glutaredoxin-5 | -0.310 |
| CNAG_07446 | *ISD11* | Mitochondrial protein | 0.290 |
| CNAG_03395 | *NFU1* | NifU-like protein C | **-0.830** |
| CNAG_05035 | *ARH1* | Ferredoxin-NADP reductase | **-0.564** |
| CNAG_05199 | *SSQ1* | HSP70-type chaperone | **0.624** |
| CNAG_01881 | *MGE1* | Nucleotide exchange factor co-chaperone | 0.257 |
| CNAG_03352 | *IND1* | Hypothetical protein | -0.150 |
| CNAG_04288 | *JAC1* | J-domain co-chaperone | **1.069** |
| **CIA pathway** | | | |
| CNAG_01802 | *DRE2* | Cytoplasmic protein | -0.171 |
| CNAG_04202 | *NAR1* | Iron hydrogenase | -0.080 |
| CNAG_01839 | *TAH18* | Peptidyl-prolyl cis-trans isomerase NIMA- interacting 1 | -0.013 |
| CNAG_02467 | *MMS19* | DNA repair/transcription protein | 0.006 |
| CNAG_05840 | *CIA2* | Hypothetical protein | **-0.326** |
| CNAG_00085 | *CIA1* | Histone chaperone | **-0.433** |
| CNAG_00082 | *CFD1* | Cytosolic Fe-S cluster assembly factor | **0.327** |
| CNAG_02231 | *NBP35* | Cytosolic Fe-S cluster assembly factor NBP35 | 0.148 |

| | | | |
|---|---|---|---|
| **Mitochondrial ETC** | | | |
| CNAG_03958 | | Hypothetical protein | **1.346** |
| CNAG_03629 | *YHB1* | NADH dehydrogenase (quinone), G subunit | **-1.33** |
| CNAG_07491 | | Glutaredoxin | **0.940** |
| CNAG_03676 | | D-lactate dehydrogenase | **-0.363** |
| CNAG_01769 | | Solute carrier family 25 (mitochondrial aspartate/glutamate transporter), member 12/13 | -0.168 |
| CNAG_03985 | *GRX5* | Grx4 family monothiol glutaredoxin | -0.310 |
| CNAG_01287 | | NADH-ubiquinone oxidoreductase 51 kDa subunit, mitochondrial | **-1.255** |
| CNAG_00486 | | Succinate dehydrogenase assembly factor 2, mitochondrial | **-0.631** |
| CNAG_06806 | *ETF1* | Electron transfer flavoprotein alpha subunit | 0.209 |
| CNAG_01005 | | Glutaredoxin | 0.236 |
| CNAG_03861 | | Cytochrome c oxidase subunit 5a | **-0.816** |
| CNAG_06723 | | Succinate dehydrogenase (ubiquinone) membrane anchor subunit | **-1.582** |

| | | | |
|---|---|---|---|
| CNAG_07356 | | Succinate dehydrogenase, cytochrome b556 subunit | **-1.458** |
| CNAG_03226 | | Succinate dehydrogenase [ubiquinone] iron-sulfur subunit, mitochondrial | **-1.694** |
| CNAG_00149 | | NADH dehydrogenase (ubiquinone) 1 alpha subcomplex 4 | **-1.171** |
| CNAG_04189 | *SDH1* | Succinate dehydrogenase [ubiquinone] flavoprotein subunit, mitochondrial | **-1.467** |
| CNAG_00716 | *CYC7* | Cytochrome c | **-1.678** |
| CNAG_06063 | | Cytochrome c oxidase assembly protein subunit 15 | **-0.649** |
| CNAG_02140 | | NADH dehydrogenase (ubiquinone) 1 alpha subcomplex 12 | **-1.258** |
| CNAG_01870 | | Electron transfer flavoprotein beta subunit | -0.488 |
| CNAG_03364 | | Hypothetical protein | **-0.454** |
| CNAG_05041 | | NADH-ubiquinone oxidoreductase subunit 8 | **-1.309** |
| CNAG_05631 | | NADH-ubiquinone oxidoreductase 49 kDa subunit, mitochondrial | **-1.105** |
| CNAG_00768 | | Hypothetical protein | 0.161 |
| CNAG_02938 | | Cytochrome c oxidase subunit 6a | **-0.666** |
| CNAG_05633 | | Ubiquinol-cytochrome c reductase subunit 9 | **-0.929** |
| CNAG_05069 | | Ubiquinol-cytochrome c reductase subunit 10 | **-1.142** |
| CNAG_05909 | *CYT1* | Cytochrome c1, heme protein | **-1.476** |
| CNAG_01470 | | NADH dehydrogenase (ubiquinone) flavoprotein 2 | **-1.564** |
| CNAG_00462 | | Electron-transferring-flavoprotein dehydrogenase | **-1.866** |
| CNAG_03848 | | Glutathione transferase | **-1.548** |
| CNAG_01443 | | Hypothetical protein | 0.197 |
| CNAG_02664 | | D-lactate dehydrogenase | **-0.709** |
| CNAG_06180 | | NADH dehydrogenase (ubiquinone) Fe-S protein 4 | **-1.356** |
| CNAG_05197 | | NADH dehydrogenase (ubiquinone) 1 alpha subcomplex 8 | **-0.969** |
| CNAG_01799 | | Ubiquinol-cytochrome c reductase subunit 8 | **-0.747** |
| CNAG_01323 | *QCR7* | Ubiquinol-cytochrome c reductase subunit 7 | **-1.119** |
| CNAG_03507 | | Mitochondrial-processing peptidase subunit beta | **-0.974** |
| CNAG_03359 | | Cytochrome c oxidase subunit 7 | **-0.900** |
| CNAG_06407 | | Cytochrome c oxidase subunit 7c | **-0.851** |
| CNAG_06012 | | NADH-cytochrome b5 reductase 1 | **-0.591** |
| CNAG_01485 | | D-lactate dehydrogenase (cytochrome) | -0.062 |
| CNAG_06663 | | Ubiquinol-cytochrome c reductase subunit 6 | **-1.391** |
| CNAG_06226 | | NADH dehydrogenase (ubiquinone) 1 alpha subcomplex 5 | **-2.017** |

| | | | |
|---|---|---|---|
| CNAG_05132 | | Cytochrome c oxidase subunit 5b | **-1.190** |
| CNAG_05179 | *QCR2* | ubiquinol-cytochrome c reductase core subunit 2 | **-1.073** |
| CNAG_01991 | | Cytochrome c oxidase subunit 4 | **-0.865** |
| CNAG_02315 | | Ubiquinol-cytochrome c reductase, iron-sulfur subunit | **-1.132** |
| CNAG_01800 | | Hypothetical protein | **-1.634** |
| CNAG_04290 | | Hypothetical protein | **-0.612** |
| CNAG_06138 | | NADH dehydrogenase (ubiquinone) Fe-S protein 6 | **-1.248** |
| CNAG_02849 | | Glutathione transferase | **-3.585** |
| CNAG_02840 | | Hypothetical protein | **1.781** |
| CNAG_03874 | | Oxidoreductase | 0.255 |
| CNAG_07674 | | Hypothetical protein | 0.345 |
| CNAG_07365 | | Oxidoreductase | **0.284** |
| CNAG_02839 | | Hypothetical protein | **4.074** |
| CNAG_03589 | | Adrenodoxin-type ferredoxin | **-0.548** |
| CNAG_02950 | *GRX3* | Grx4 family monothiol glutaredoxin | -0.402 |
| CNAG_01802 | | Fe-S cluster assembly protein | -0.171 |
| CNAG_04521 | | Oxidoreductase | **1.373** |
| CNAG_05554 | | Cytochrome b5 | **-0.962** |

**TABLE S3** The loss of *CIR1* gene affects the gene transcript levels of components of the mitochondrial ISC assembly pathway and the ETC (KEGG). The log2 fold change (experimental group/control group) in bold was significantly different ($P < 0.05$).

| **Gene ID** | **Gene name** | **Function** | **WT (control) vs *cir1*Δ** |
|---|---|---|---|
| **Mitochondrial ISC assembly pathway** | | | |
| CNAG_00491 | *ISA2* | Iron sulfur assembly protein 2 | -0.157 |
| CNAG_05011 | *FRR4* | Frataxin | **-0.207** |
| CNAG_07446 | *ISD11* | Mitochondrial protein | **-0.168** |
| CNAG_04288 | *JAC1* | J-domain co-chaperone | **-0.323** |
| CNAG_05035 | *ARH1* | Ferredoxin-NADP reductase | 0.013 |
| CNAG_01442 | *NFS1* | Cysteine desulferase | **0.169** |
| CNAG_02131 | *ISA1* | Iron sulfur assembly protein 1 | **1.504** |
| CNAG_05199 | *SSQ1* | HSP70-type chaperone | **0.569** |
| CNAG_01881 | *MGE1* | Nucleotide exchange factor co-chaperone | **0.477** |
| CNAG_03352 | *IND1* | Hypothetical protein | **0.391** |
| CNAG_04039 | *FRR3* | Scaffold protein | **0.804** |
| CNAG_00389 | *IBA57* | Mitochondrial protein | **0.236** |
| CNAG_03985 | *GRX5* | Monothiol glutaredoxin-5 | **2.146** |

| | | | |
|---|---|---|---|
| CNAG_03395 | *NFU1* | NifU-like protein C | **2.176** |
| **Mitochondrial ETC** | | | |
| CNAG_06806 | *ETF1* | Electron transfer flavoprotein alpha subunit | -0.09 |
| CNAG_01485 | | D-lactate dehydrogenase (cytochrome) | -0.047 |
| CNAG_03364 | | Hypothetical protein | -0.11 |
| CNAG_01870 | | Electron transfer flavoprotein beta subunit | **-0.237** |
| CNAG_00486 | | Succinate dehydrogenase assembly factor 2, mitochondrial | **-0.375** |
| CNAG_03676 | | D-lactate dehydrogenase | **-0.231** |
| CNAG_02664 | | D-lactate dehydrogenase | **-0.275** |
| CNAG_03958 | | Hypothetical protein | 0.24 |
| CNAG_07674 | | Hypothetical protein | -0.008 |
| CNAG_01769 | | Solute carrier family 25 (mitochondrial aspartate/glutamate transporter), member 12/13 | 0.002 |
| CNAG_00768 | | Hypothetical protein | 0.083 |
| CNAG_03507 | | Mitochondrial-processing peptidase subunit beta | 0.104 |
| CNAG_00149 | | NADH dehydrogenase (ubiquinone) 1 alpha subcomplex 4 | -0.042 |
| CNAG_01005 | | Glutaredoxin | 0.023 |
| CNAG_02849 | | Glutathione transferase | 0.352 |
| CNAG_06012 | | NADH-cytochrome b5 reductase 1 | 0.026 |
| CNAG_03848 | | Glutathione transferase | **0.423** |
| CNAG_07365 | | Oxidoreductase | **0.615** |
| CNAG_03874 | | Oxidoreductase | **2.534** |
| CNAG_03629 | *YHB1* | NADH dehydrogenase (quinone), G subunit | **0.852** |
| CNAG_02839 | | Hypothetical protein | **5.304** |
| CNAG_06723 | | Succinate dehydrogenase (ubiquinone) membrane anchor subunit | **1.283** |
| CNAG_04189 | *SDH1* | Succinate dehydrogenase [ubiquinone] flavoprotein subunit, mitochondrial | **1.287** |
| CNAG_01443 | | Hypothetical protein | **1.197** |
| CNAG_03985 | *GRX5* | Grx4 family monothiol glutaredoxin | **2.146** |
| CNAG_00462 | | Electron-transferring-flavoprotein dehydrogenase | **1.597** |
| CNAG_03226 | | Succinate dehydrogenase [ubiquinone] iron-sulfur subunit, mitochondrial | **1.57** |
| CNAG_01802 | | Fe-S cluster assembly protein | **1.429** |
| CNAG_06063 | | Cytochrome c oxidase assembly protein subunit 15 | **1.916** |
| CNAG_04290 | | Hypothetical protein | 0.131 |
| CNAG_02840 | | Hypothetical protein | **0.39** |
| CNAG_03861 | | Cytochrome c oxidase subunit 5a | **0.355** |
| CNAG_00716 | *CYC7* | Cytochrome c | **1.1** |

| | | | |
|---|---|---|---|
| CNAG_05069 | | Ubiquinol-cytochrome c reductase subunit 10 | **0.835** |
| CNAG_03359 | | Cytochrome c oxidase subunit 7 | **0.716** |
| CNAG_06407 | | Cytochrome c oxidase subunit 7c | **0.53** |
| CNAG_01800 | | Hypothetical protein | **0.635** |
| CNAG_05633 | | Ubiquinol-cytochrome c reductase subunit 9 | **0.6** |
| CNAG_02938 | | Cytochrome c oxidase subunit 6a | **0.354** |
| CNAG_03589 | | Adrenodoxin-type ferredoxin | **0.365** |
| CNAG_02315 | | Ubiquinol-cytochrome c reductase, iron-sulfur subunit | **0.572** |
| CNAG_05197 | | NADH dehydrogenase (ubiquinone) 1 alpha subcomplex 8 | **0.404** |
| CNAG_05554 | | Cytochrome b5 | **0.669** |
| CNAG_01287 | | NADH-ubiquinone oxidoreductase 51 kDa subunit, mitochondrial | **0.781** |
| CNAG_04521 | | Oxidoreductase | **5.652** |
| CNAG_07356 | | Succinate dehydrogenase, cytochrome b556 subunit | **1.207** |
| CNAG_05631 | | NADH-ubiquinone oxidoreductase 49 kDa subunit, mitochondrial | **0.863** |
| CNAG_06180 | | NADH dehydrogenase (ubiquinone) Fe-S protein 4 | **1.064** |
| CNAG_05041 | | NADH-ubiquinone oxidoreductase subunit 8 | **0.983** |
| CNAG_01470 | | NADH dehydrogenase (ubiquinone) flavoprotein 2 | **1.077** |
| CNAG_07491 | | Glutaredoxin | **1.279** |
| CNAG_06138 | | NADH dehydrogenase (ubiquinone) Fe-S protein 6 | **0.937** |
| CNAG_06663 | | Ubiquinol-cytochrome c reductase subunit 6 | **0.63** |
| CNAG_02140 | | NADH dehydrogenase (ubiquinone) 1 alpha subcomplex 12 | **0.975** |
| CNAG_06226 | | NADH dehydrogenase (ubiquinone) 1 alpha subcomplex 5 | **1.053** |
| CNAG_05179 | *QCR2* | ubiquinol-cytochrome c reductase core subunit 2 | **0.464** |
| CNAG_01799 | | Ubiquinol-cytochrome c reductase subunit 8 | **0.415** |
| CNAG_05132 | | Cytochrome c oxidase subunit 5b | **0.463** |
| CNAG_01323 | *QCR7* | Ubiquinol-cytochrome c reductase subunit 7 | **0.682** |
| CNAG_01991 | | Cytochrome c oxidase subunit 4 | **0.468** |
| CNAG_02950 | *GRX3* | Grx4 family monothiol glutaredoxin | **0.399** |
| CNAG_05909 | *CYT1* | Cytochrome c1, heme protein, | **0.461** |

**Table S4. Primers for qPCR validation of RNA-seq data**

| **Name** | **Sequence 5'-3'** | **Reference** |
|---|---|---|
| Cir1_F | ctcagcaaggcgaaaagtct | This Study |

| | | |
|---|---|---|
| Cir1_R | tcctaacgtcaaaactccacaa | This Study |
| HapX_F | cccctacgccaattcctatc | This Study |
| HapX _R | cgttgcctcgtccaataact | This Study |
| Rim101_F | gctgatcaagagttggcttgt | This Study |
| Rim101_R | ccacttcaatcctctccctct | This Study |
| Lac1_F | agggaaagttggctgtggtt | This Study |
| Lac1_R | tcgcgcagatactggaaagt | This Study |
| CNAG_01881_F | aaaacttgaagggcgaacag | This Study |
| CNAG_01881_R | ggcctcaagaccctagaacc | This Study |
| CNAG_05840_F | gagggctttacctccacgat | This Study |
| CNAG_05840_R | aacagccttccaccacactc | This Study |
| CNAG_03395_F | tttggatgaggaagaacttatcg | This Study |
| CNAG_03395_R | cctttccagacttccctgct | This Study |
| CNAG_01287_F | caacgacttgctcaattcca | This Study |
| CNAG_01287_R | aattcgcctaatggcatctc | This Study |
| CNAG_05041_F | caaaaccaggaattctctaccg | This Study |
| CNAG_05041_R | tgaaggttagcagcgatttc | This Study |
| CNAG_07491_F | tgaagaaatgggtgctttcc | This Study |
| CNAG_07491_R | tgatctccgaaccttttcg | This Study |
| CNAG_01769_F | gttgtccacaggccagttct | This Study |
| CNAG_01769_R | taagcgatttggggataagc | This Study |
| CNAG_03848_F | gagataatggcaatgcacga | This Study |
| CNAG_03848_R | tcggcttgttcgtcaccta | This Study |
| CNAG_02849_F | attgagcttgagacgaaaggag | This Study |
| CNAG_02849_R | ggcgtcttcaatccgttg | This Study |
| CNAG_06063_F | gaagattacccttccctgctg | This Study |
| CNAG_06063_R | tcctactgggcttcctcaaa | This Study |
| CNAG_02839_F | atgctcaccttgccttggt | This Study |
| CNAG_02839_R | ttggacaaatacacatcgtcaa | This Study |
| CNAG_04521_F | ctcgcgaggaacattgtagc | This Study |
| CNAG_04521_R | tgtaggaaggctcagtgtcg | This Study |
| CNAG_05199_F | gctcagcttgacgccact | This Study |
| CNAG_05199_R | tggaaaagaccgagagaagc | This Study |

**Table S5. Primers for qPCR analysis of genes involved in the response to oxidative stress.**

| Function | Name | Sequence 5'-3' | Reference |
|---|---|---|---|
| cytochrome c peroxidase | CCP1_F | CCGTCGGAGCTGCATATTAC | This Study |
| cytochrome c peroxidase | CCP1_R | TCGTAACCCTCCTTGTCGAG | This Study |
| catalase 3 | CAT3_F | CAAGGGTTTGGCGTTCATAC | This Study |
| catalase 3 | CAT3_R | GGTCTCTTCGGTGGAAATCA | This Study |
| lactoylglutathione lyase (Glyoxalate I) | GLX1_F | GATGCCAATTTCAAGGGCTA | This Study |
| lactoylglutathione lyase (Glyoxalate I) | GLX1_R | ACTCTCATCCTGCCCTCCTC | This Study |
| thioredoxin-disulfide reductase | TRR1_F | ATGTCTCCCATCGCCAAC | This Study |
| thioredoxin-disulfide reductase | TRR1_R | GTAGAGGACAGGCTCGAGGT | This Study |
| alternative oxidase 1 | AOX1_F | CTCTGCCGTTTCAAGCACTC | This Study |
| alternative oxidase 1 | AOX1_R | CCCCTGCCTTCTGATCCTAC | This Study |
| actin | ACTIN_F | CACCATTGGTAACGAGCGATTC | Horianopolous, 2020 |
| actin | ACTIN_R | TGGTAGTACCACCAGACATGAC | Horianopolous, 2020 |

**Table S6. Data sheets of the RNA-seq data for antimycin A treatment.**

| | baseMean | log2FoldCh | lfcSE | stat | pvalue | padj |
|---|---|---|---|---|---|---|
| CNAG_07304 | 612.5685 | -0.29494 | 0.112575 | -2.61992 | 0.008795 | 0.030559 |
| CNAG_00003 | 534.756 | -0.19602 | 0.099228 | -1.97543 | 0.04822 | 0.121038 |
| CNAG_00005 | 102.6655 | -1.11235 | 0.208394 | -5.33774 | 9.41E-08 | 9.58E-07 |
| CNAG_12001 | 26.87368 | -0.90325 | 0.391048 | -2.30983 | 0.020897 | 0.061774 |
| CNAG_00006 | 4305.682 | 0.190281 | 0.072586 | 2.621461 | 0.008755 | 0.030449 |
| CNAG_00007 | 4644.062 | 0.177629 | 0.060768 | 2.92306 | 0.003466 | 0.013907 |
| CNAG_00010 | 2963.125 | 0.189944 | 0.077087 | 2.464028 | 0.013739 | 0.043941 |
| CNAG_00011 | 604.333 | 0.303674 | 0.112453 | 2.700462 | 0.006924 | 0.024856 |
| CNAG_00012 | 1651.897 | -0.31942 | 0.088189 | -3.62203 | 0.000292 | 0.001613 |
| CNAG_00013 | 507.8788 | 0.530465 | 0.10127 | 5.238128 | 1.62E-07 | 1.59E-06 |
| CNAG_00015 | 226.4114 | 0.373218 | 0.140901 | 2.648795 | 0.008078 | 0.028525 |
| CNAG_00019 | 2990.429 | 0.148963 | 0.071209 | 2.091933 | 0.036445 | 0.096608 |
| CNAG_00020 | 1074.6 | -0.21745 | 0.108093 | -2.01171 | 0.04425 | 0.113297 |
| CNAG_00021 | 938.083 | -0.52979 | 0.090653 | -5.84415 | 5.09E-09 | 6.22E-08 |
| CNAG_00025 | 1209.621 | 0.263638 | 0.078601 | 3.354135 | 0.000796 | 0.003888 |
| CNAG_00030 | 167.1032 | -0.35569 | 0.161896 | -2.19702 | 0.028019 | 0.078967 |
| CNAG_00032 | 108.6737 | -0.40832 | 0.203916 | -2.00241 | 0.045241 | 0.115228 |
| CNAG_00034 | 45017.86 | -0.24964 | 0.071719 | -3.48077 | 0.0005 | 0.002608 |
| CNAG_00035 | 159.1964 | -0.40602 | 0.168876 | -2.40425 | 0.016206 | 0.050167 |
| CNAG_00038 | 785.118 | -0.30876 | 0.09572 | -3.22564 | 0.001257 | 0.005777 |
| CNAG_00039 | 1157.736 | -0.38318 | 0.102317 | -3.74504 | 0.00018 | 0.00104 |
| CNAG_00040 | 4557.666 | -0.76178 | 0.071666 | -10.6295 | 2.17E-26 | 1.30E-24 |
| CNAG_07308 | 142.4692 | 1.393547 | 0.195485 | 7.12867 | 1.01E-12 | 1.91E-11 |
| CNAG_12006 | 223.733 | 1.04596 | 0.170028 | 6.151682 | 7.67E-10 | 1.04E-08 |
| CNAG_00044 | 2577.006 | 0.169627 | 0.067553 | 2.511031 | 0.012038 | 0.039439 |
| CNAG_00046 | 4115.64 | -0.20565 | 0.073753 | -2.7884 | 0.005297 | 0.019741 |
| CNAG_00047 | 578.3429 | 1.175948 | 0.11686 | 10.06287 | 8.06E-24 | 4.08E-22 |
| CNAG_00052 | 668.8482 | 1.387881 | 0.122917 | 11.29119 | 1.45E-29 | 1.01E-27 |
| CNAG_00053 | 532.5399 | 0.348061 | 0.099251 | 3.506883 | 0.000453 | 0.002397 |
| CNAG_00054 | 126.3972 | 0.961799 | 0.185727 | 5.178567 | 2.24E-07 | 2.16E-06 |
| CNAG_00055 | 1084.783 | -0.17307 | 0.086204 | -2.00772 | 0.044674 | 0.114081 |
| CNAG_00057 | 5954.624 | 0.68475 | 0.075798 | 9.033892 | 1.66E-19 | 5.72E-18 |
| CNAG_00059 | 315.0006 | -0.46235 | 0.133703 | -3.45806 | 0.000544 | 0.00281 |
| CNAG_00061 | 26638.51 | 0.908546 | 0.068598 | 13.2445 | 4.86E-40 | 6.65E-38 |
| CNAG_00062 | 3405.8 | -0.22812 | 0.065848 | -3.46435 | 0.000532 | 0.002756 |
| CNAG_00067 | 3079.263 | -0.13164 | 0.066223 | -1.9879 | 0.046823 | 0.118101 |
| CNAG_00068 | 1109.52 | -0.37845 | 0.104762 | -3.61247 | 0.000303 | 0.001663 |
| CNAG_00070 | 1099.423 | 0.188667 | 0.08321 | 2.267369 | 0.023368 | 0.067635 |
| CNAG_00072 | 2526.276 | 0.247378 | 0.085082 | 2.90754 | 0.003643 | 0.014487 |
| CNAG_00073 | 3947.203 | 0.13903 | 0.065553 | 2.120876 | 0.033932 | 0.09125 |
| CNAG_00074 | 4738.152 | 0.174589 | 0.061994 | 2.816242 | 0.004859 | 0.018354 |
| CNAG_00075 | 1959.588 | 0.22914 | 0.093655 | 2.446643 | 0.014419 | 0.045688 |
| CNAG_00076 | 1510.456 | 0.200363 | 0.084751 | 2.364129 | 0.018073 | 0.054793 |
| CNAG_00078 | 2451.457 | 0.477263 | 0.067671 | 7.052691 | 1.75E-12 | 3.22E-11 |
| CNAG_00079 | 679.7749 | 0.809134 | 0.10366 | 7.805653 | 5.92E-15 | 1.35E-13 |
| CNAG_00081 | 2846.796 | -0.19661 | 0.065485 | -3.00229 | 0.00268 | 0.011123 |
| CNAG_00083 | 1491.087 | -0.33131 | 0.078706 | -4.20942 | 2.56E-05 | 0.000178 |

| | | | | | | |
|---|---|---|---|---|---|---|
| CNAG_00084 | 3643.087 | -0.33026 | 0.07206 | -4.58319 | 4.58E-06 | 3.64E-05 |
| CNAG_00086 | 2145.345 | -0.34185 | 0.068481 | -4.99181 | 5.98E-07 | 5.40E-06 |
| CNAG_00087 | 1302.887 | 0.268695 | 0.080521 | 3.33695 | 0.000847 | 0.00411 |
| CNAG_00088 | 3710.145 | 0.197523 | 0.0678 | 2.913335 | 0.003576 | 0.014309 |
| CNAG_00089 | 1447.737 | -0.20811 | 0.081954 | -2.53931 | 0.011107 | 0.036948 |
| CNAG_00091 | 6500.386 | 1.218663 | 0.186992 | 6.517183 | 7.16E-11 | 1.10E-09 |
| CNAG_00093 | 3471.411 | 1.120857 | 0.080934 | 13.84905 | 1.29E-43 | 2.05E-41 |
| CNAG_00094 | 238.4408 | -0.36972 | 0.141182 | -2.61876 | 0.008825 | 0.030609 |
| CNAG_00096 | 627.9364 | 0.269681 | 0.101835 | 2.648208 | 0.008092 | 0.028562 |
| CNAG_00098 | 466.4089 | 0.375887 | 0.107911 | 3.483302 | 0.000495 | 0.002591 |
| CNAG_00100 | 7128.253 | -0.13309 | 0.063947 | -2.08133 | 0.037404 | 0.098561 |
| CNAG_07309 | 2149.173 | 0.271326 | 0.079746 | 3.402373 | 0.000668 | 0.003357 |
| CNAG_07310 | 1971.141 | 0.149273 | 0.071469 | 2.088639 | 0.03674 | 0.09716 |
| CNAG_00104 | 7200.725 | -0.18522 | 0.063465 | -2.91848 | 0.003517 | 0.014098 |
| CNAG_00107 | 2663.085 | 0.930289 | 0.089767 | 10.3634 | 3.64E-25 | 2.01E-23 |
| CNAG_00108 | 4355.37 | -0.13485 | 0.066779 | -2.01939 | 0.043447 | 0.111635 |
| CNAG_00116 | 40784.8 | -0.22366 | 0.081314 | -2.7506 | 0.005949 | 0.021764 |
| CNAG_00117 | 4085.103 | -0.18918 | 0.069073 | -2.73881 | 0.006166 | 0.022467 |
| CNAG_00118 | 343.3595 | -0.24609 | 0.124979 | -1.96906 | 0.048946 | 0.122416 |
| CNAG_00120 | 889.5944 | -0.23844 | 0.104361 | -2.28477 | 0.022326 | 0.065111 |
| CNAG_00121 | 6866.365 | 0.595662 | 0.060555 | 9.836673 | 7.83E-23 | 3.61E-21 |
| CNAG_00122 | 752.1669 | 0.276631 | 0.11081 | 2.496449 | 0.012544 | 0.040927 |
| CNAG_00123 | 514.0227 | 0.843377 | 0.111255 | 7.580548 | 3.44E-14 | 7.42E-13 |
| CNAG_00124 | 1179.171 | 0.354346 | 0.082911 | 4.273802 | 1.92E-05 | 0.000137 |
| CNAG_00126 | 1963.757 | -0.62194 | 0.091651 | -6.78593 | 1.15E-11 | 1.95E-10 |
| CNAG_00130 | 7654.827 | 0.795493 | 0.081378 | 9.775261 | 1.44E-22 | 6.49E-21 |
| CNAG_00134 | 830.2532 | 0.295645 | 0.096073 | 3.077291 | 0.002089 | 0.008923 |
| CNAG_00137 | 1722.345 | 0.232793 | 0.07502 | 3.103099 | 0.001915 | 0.008307 |
| CNAG_00139 | 2816.956 | 0.250353 | 0.076925 | 3.254514 | 0.001136 | 0.005302 |
| CNAG_00142 | 1705.801 | -0.68108 | 0.08859 | -7.68806 | 1.49E-14 | 3.31E-13 |
| CNAG_12024 | 508.1234 | -0.30319 | 0.141595 | -2.14127 | 0.032252 | 0.087822 |
| CNAG_00143 | 4192.941 | -0.18787 | 0.062767 | -2.99309 | 0.002762 | 0.011422 |
| CNAG_00148 | 2884.437 | -0.17552 | 0.080684 | -2.17542 | 0.029599 | 0.082354 |
| CNAG_00149 | 9503.464 | 0.783376 | 0.072579 | 10.79346 | 3.70E-27 | 2.38E-25 |
| CNAG_00150 | 3061.492 | 0.312527 | 0.081104 | 3.853386 | 0.000116 | 0.000706 |
| CNAG_00152 | 839.036 | -0.31734 | 0.090898 | -3.49113 | 0.000481 | 0.002521 |
| CNAG_00156 | 1325.854 | 0.217107 | 0.09473 | 2.291842 | 0.021915 | 0.064222 |
| CNAG_00157 | 1284.159 | 0.236278 | 0.08247 | 2.865022 | 0.00417 | 0.016222 |
| CNAG_00161 | 265.0703 | -0.43068 | 0.141348 | -3.04691 | 0.002312 | 0.009743 |
| CNAG_00162 | 22065.14 | 2.595444 | 0.074148 | 35.00338 | 2.00E-268 | 7.80E-265 |
| CNAG_00164 | 852.8536 | -0.73035 | 0.103953 | -7.02573 | 2.13E-12 | 3.86E-11 |
| CNAG_00165 | 2513.732 | -0.22396 | 0.073254 | -3.05727 | 0.002234 | 0.009453 |
| CNAG_00166 | 1151.103 | -0.5723 | 0.07972 | -7.17891 | 7.03E-13 | 1.34E-11 |
| CNAG_07314 | 498.1941 | 0.365118 | 0.106563 | 3.426305 | 0.000612 | 0.003119 |
| CNAG_00172 | 1096.888 | -0.22997 | 0.115522 | -1.99069 | 0.046515 | 0.117477 |
| CNAG_00173 | 589.2888 | 0.435593 | 0.107228 | 4.062323 | 4.86E-05 | 0.000321 |
| CNAG_00180 | 1004.457 | 0.199758 | 0.093983 | 2.125484 | 0.033546 | 0.090445 |
| CNAG_00183 | 928.6153 | 0.52378 | 0.121474 | 4.311887 | 1.62E-05 | 0.000118 |

| | | | | | | |
|---|---|---|---|---|---|---|
| CNAG_00184 | 2559.452 | -0.19691 | 0.076619 | -2.57002 | 0.010169 | 0.034296 |
| CNAG_00185 | 1430.245 | 0.181852 | 0.079151 | 2.297519 | 0.021589 | 0.063434 |
| CNAG_00187 | 1534.297 | -0.14527 | 0.074044 | -1.96192 | 0.049772 | 0.124176 |
| CNAG_00190 | 1546.54 | 0.430034 | 0.078442 | 5.482161 | 4.20E-08 | 4.50E-07 |
| CNAG_00192 | 2384.082 | -0.66538 | 0.092063 | -7.22748 | 4.92E-13 | 9.62E-12 |
| CNAG_00193 | 4456.988 | -0.16506 | 0.074072 | -2.22835 | 0.025857 | 0.073781 |
| CNAG_00195 | 885.2141 | -0.41049 | 0.089658 | -4.57841 | 4.69E-06 | 3.71E-05 |
| CNAG_07317 | 1758.987 | 0.341201 | 0.087215 | 3.912209 | 9.15E-05 | 0.000567 |
| CNAG_07319 | 2425.49 | 0.609632 | 0.08731 | 6.982382 | 2.90E-12 | 5.15E-11 |
| CNAG_07320 | 1297.58 | 0.293439 | 0.081389 | 3.605408 | 0.000312 | 0.001703 |
| CNAG_07322 | 3408.414 | -0.16154 | 0.067756 | -2.38412 | 0.01712 | 0.052459 |
| CNAG_07326 | 3923.709 | 0.308122 | 0.066725 | 4.617806 | 3.88E-06 | 3.13E-05 |
| CNAG_07333 | 2292.936 | -0.24592 | 0.083986 | -2.92806 | 0.003411 | 0.013699 |
| CNAG_07334 | 639.9327 | -0.20389 | 0.094109 | -2.16658 | 0.030267 | 0.083702 |
| CNAG_07028 | 2694.725 | -0.19425 | 0.070434 | -2.75787 | 0.005818 | 0.021356 |
| CNAG_07338 | 753.2874 | 0.360327 | 0.093361 | 3.859492 | 0.000114 | 0.000691 |
| CNAG_07339 | 1547.973 | 0.439683 | 0.077433 | 5.678232 | 1.36E-08 | 1.57E-07 |
| CNAG_07340 | 396.7786 | 0.295938 | 0.11007 | 2.688637 | 0.007174 | 0.025589 |
| CNAG_07342 | 1684.305 | 0.964963 | 0.07583 | 12.72533 | 4.28E-37 | 5.06E-35 |
| CNAG_00232 | 32143.55 | -0.18428 | 0.073296 | -2.51421 | 0.01193 | 0.0392 |
| CNAG_00233 | 1659.769 | 0.358204 | 0.078809 | 4.545208 | 5.49E-06 | 4.30E-05 |
| CNAG_00234 | 645.4123 | -0.40336 | 0.093407 | -4.31833 | 1.57E-05 | 0.000114 |
| CNAG_00235 | 3472.864 | 0.229618 | 0.066152 | 3.471085 | 0.000518 | 0.002697 |
| CNAG_00236 | 740.1669 | 0.509973 | 0.089849 | 5.675918 | 1.38E-08 | 1.59E-07 |
| CNAG_00237 | 10935.84 | -0.58614 | 0.062854 | -9.32532 | 1.11E-20 | 4.25E-19 |
| CNAG_00238 | 18091.05 | -0.15884 | 0.066482 | -2.38918 | 0.016886 | 0.051902 |
| CNAG_00241 | 1733.516 | -0.24152 | 0.080757 | -2.99066 | 0.002784 | 0.011494 |
| CNAG_00248 | 506.4665 | 0.345364 | 0.106381 | 3.246491 | 0.001168 | 0.005421 |
| CNAG_00249 | 729.7675 | 0.284002 | 0.101151 | 2.807693 | 0.00499 | 0.018784 |
| CNAG_00250 | 1932.82 | 0.275109 | 0.076988 | 3.573389 | 0.000352 | 0.001906 |
| CNAG_00254 | 1225.319 | 0.655775 | 0.093269 | 7.031001 | 2.05E-12 | 3.74E-11 |
| CNAG_00256 | 4157.907 | -0.21348 | 0.088599 | -2.40955 | 0.015972 | 0.04964 |
| CNAG_00257 | 2874.265 | -0.51019 | 0.096154 | -5.30597 | 1.12E-07 | 1.12E-06 |
| CNAG_00259 | 2018.895 | -0.51914 | 0.074272 | -6.98966 | 2.76E-12 | 4.90E-11 |
| CNAG_00260 | 3007.873 | -0.20286 | 0.068439 | -2.96416 | 0.003035 | 0.012362 |
| CNAG_00261 | 9593.965 | 0.539915 | 0.081374 | 6.635005 | 3.24E-11 | 5.15E-10 |
| CNAG_00264 | 1945.682 | 0.312288 | 0.090958 | 3.433306 | 0.000596 | 0.00305 |
| CNAG_00265 | 1540.903 | -0.28047 | 0.082645 | -3.39363 | 0.00069 | 0.003453 |
| CNAG_00268 | 7224.141 | -0.36042 | 0.060252 | -5.98191 | 2.21E-09 | 2.82E-08 |
| CNAG_00271 | 742.249 | -0.22803 | 0.094061 | -2.42427 | 0.015339 | 0.048075 |
| CNAG_00272 | 1532.992 | -0.23822 | 0.096928 | -2.45766 | 0.013985 | 0.044562 |
| CNAG_00275 | 442.0507 | 0.918975 | 0.131282 | 6.999981 | 2.56E-12 | 4.58E-11 |
| CNAG_00280 | 3737.381 | 0.195597 | 0.071559 | 2.733355 | 0.006269 | 0.022788 |
| CNAG_00283 | 352.0719 | -0.84711 | 0.117508 | -7.20894 | 5.64E-13 | 1.09E-11 |
| CNAG_00290 | 1382.506 | 0.285991 | 0.076852 | 3.721331 | 0.000198 | 0.001134 |
| CNAG_00292 | 4623.515 | -0.52873 | 0.073595 | -7.18426 | 6.76E-13 | 1.30E-11 |
| CNAG_00299 | 1284.349 | -0.35778 | 0.099558 | -3.59374 | 0.000326 | 0.001774 |
| CNAG_00301 | 975.2737 | 0.887984 | 0.093119 | 9.536034 | 1.48E-21 | 6.16E-20 |

| | | | | | | |
|---|---|---|---|---|---|---|
| CNAG_00304 | 654.0667 | -0.22983 | 0.101062 | -2.27412 | 0.022959 | 0.066681 |
| CNAG_00305 | 8055.46 | -0.1663 | 0.069457 | -2.39426 | 0.016654 | 0.051208 |
| CNAG_12043 | 31.57865 | -0.6879 | 0.344128 | -1.99898 | 0.045611 | 0.115903 |
| CNAG_00306 | 8806.955 | -0.61466 | 0.187011 | -3.28675 | 0.001014 | 0.004811 |
| CNAG_00307 | 1548.027 | 0.399382 | 0.073427 | 5.439164 | 5.35E-08 | 5.62E-07 |
| CNAG_00309 | 1216.865 | -0.20835 | 0.081493 | -2.55673 | 0.010566 | 0.035435 |
| CNAG_00314 | 716.0909 | 0.379305 | 0.097037 | 3.908873 | 9.27E-05 | 0.000574 |
| CNAG_12046 | 472.7305 | 1.713788 | 0.132903 | 12.89505 | 4.80E-38 | 6.04E-36 |
| CNAG_00317 | 933.9949 | -0.22678 | 0.087129 | -2.60278 | 0.009247 | 0.031817 |
| CNAG_00328 | 977.4552 | 0.198737 | 0.084643 | 2.347937 | 0.018878 | 0.056706 |
| CNAG_00331 | 80.57773 | -0.92553 | 0.226837 | -4.08016 | 4.50E-05 | 0.000298 |
| CNAG_00337 | 1496.629 | 0.233003 | 0.077249 | 3.016265 | 0.002559 | 0.010663 |
| CNAG_00342 | 906.4604 | -0.1833 | 0.087486 | -2.09522 | 0.036151 | 0.096025 |
| CNAG_00346 | 505.6638 | 0.373277 | 0.107707 | 3.465679 | 0.000529 | 0.002744 |
| CNAG_00347 | 1159.904 | -0.25834 | 0.097897 | -2.63894 | 0.008317 | 0.029135 |
| CNAG_07356 | 4529.25 | 0.679481 | 0.068589 | 9.906546 | 3.90E-23 | 1.87E-21 |
| CNAG_00351 | 472.5929 | -0.27446 | 0.107217 | -2.55987 | 0.010471 | 0.035147 |
| CNAG_00358 | 1046.005 | -0.30488 | 0.084489 | -3.60859 | 0.000308 | 0.001684 |
| CNAG_00360 | 221.164 | 0.4254 | 0.173681 | 2.449319 | 0.014313 | 0.045386 |
| CNAG_00363 | 4451.252 | 0.168388 | 0.082392 | 2.043755 | 0.040978 | 0.106526 |
| CNAG_00369 | 771.6722 | -0.18517 | 0.094429 | -1.96094 | 0.049885 | 0.124299 |
| CNAG_00370 | 35031.63 | -0.15093 | 0.073894 | -2.04254 | 0.041098 | 0.106766 |
| CNAG_00371 | 237.6394 | -0.34371 | 0.143296 | -2.39857 | 0.016459 | 0.050809 |
| CNAG_00373 | 785.7733 | 0.357517 | 0.089799 | 3.981287 | 6.85E-05 | 0.000441 |
| CNAG_00375 | 2140.087 | -0.23139 | 0.089707 | -2.57942 | 0.009897 | 0.033575 |
| CNAG_00377 | 5619.067 | -0.15004 | 0.07592 | -1.97626 | 0.048125 | 0.12099 |
| CNAG_00378 | 231.9003 | -0.65013 | 0.13945 | -4.66209 | 3.13E-06 | 2.54E-05 |
| CNAG_00385 | 2059.542 | -0.27524 | 0.070485 | -3.90491 | 9.43E-05 | 0.000582 |
| CNAG_00386 | 7230.443 | -0.15623 | 0.060675 | -2.57492 | 0.010026 | 0.033916 |
| CNAG_00387 | 475.8273 | -0.21478 | 0.106452 | -2.01763 | 0.043629 | 0.111928 |
| CNAG_00388 | 1009.156 | -0.19899 | 0.083934 | -2.37076 | 0.017752 | 0.054037 |
| CNAG_07951 | 1039.823 | -0.48115 | 0.083607 | -5.75491 | 8.67E-09 | 1.02E-07 |
| CNAG_00391 | 283.0611 | -0.45031 | 0.133133 | -3.3824 | 0.000719 | 0.003574 |
| CNAG_00394 | 642.9983 | -0.28869 | 0.097731 | -2.95388 | 0.003138 | 0.012728 |
| CNAG_00400 | 3001.344 | 0.34487 | 0.074452 | 4.632127 | 3.62E-06 | 2.92E-05 |
| CNAG_00404 | 169.9867 | 0.343097 | 0.161665 | 2.122266 | 0.033815 | 0.090998 |
| CNAG_00407 | 7836.201 | 0.338023 | 0.066294 | 5.098864 | 3.42E-07 | 3.22E-06 |
| CNAG_00409 | 1596.008 | 0.567575 | 0.07825 | 7.253355 | 4.07E-13 | 8.01E-12 |
| CNAG_00411 | 1467.664 | -0.19746 | 0.097178 | -2.03194 | 0.04216 | 0.109018 |
| CNAG_00413 | 3851.946 | 0.24036 | 0.072625 | 3.309611 | 0.000934 | 0.004478 |
| CNAG_00414 | 362.2834 | 0.346945 | 0.116499 | 2.978091 | 0.002901 | 0.011907 |
| CNAG_00418 | 19488.02 | -0.41692 | 0.066973 | -6.2253 | 4.81E-10 | 6.65E-09 |
| CNAG_00421 | 757.4651 | 0.292685 | 0.099006 | 2.956244 | 0.003114 | 0.012651 |
| CNAG_00423 | 599.0324 | 0.288689 | 0.097305 | 2.966853 | 0.003009 | 0.012267 |
| CNAG_00429 | 435.1978 | -0.59395 | 0.112616 | -5.27413 | 1.33E-07 | 1.32E-06 |
| CNAG_07359 | 2353.146 | 0.254707 | 0.114801 | 2.218679 | 0.026509 | 0.075254 |
| CNAG_00433 | 252.7549 | -0.77569 | 0.146338 | -5.30069 | 1.15E-07 | 1.15E-06 |
| CNAG_00439 | 2102.633 | -0.22165 | 0.096579 | -2.29503 | 0.021731 | 0.06378 |

| | | | | | | |
|---|---|---|---|---|---|---|
| CNAG_00441 | 13181.91 | -0.55977 | 0.060253 | -9.29041 | 1.54E-20 | 5.82E-19 |
| CNAG_00442 | 2195.8 | 0.344578 | 0.070897 | 4.860278 | 1.17E-06 | 1.01E-05 |
| CNAG_00443 | 264.8006 | -0.30773 | 0.139765 | -2.20178 | 0.027681 | 0.078184 |
| CNAG_12058 | 49.59243 | 0.605379 | 0.280973 | 2.154583 | 0.031194 | 0.085591 |
| CNAG_00447 | 7757.861 | -0.18269 | 0.061152 | -2.98743 | 0.002813 | 0.011603 |
| CNAG_00449 | 1713.257 | -0.55521 | 0.071454 | -7.7702 | 7.84E-15 | 1.78E-13 |
| CNAG_00451 | 439.2307 | -0.56628 | 0.111795 | -5.06531 | 4.08E-07 | 3.78E-06 |
| CNAG_00453 | 3375.061 | 0.91773 | 0.073666 | 12.45807 | 1.26E-35 | 1.32E-33 |
| CNAG_00455 | 2021.609 | -0.47565 | 0.076063 | -6.25334 | 4.02E-10 | 5.63E-09 |
| CNAG_00456 | 3305.442 | -0.19767 | 0.078369 | -2.52225 | 0.011661 | 0.038527 |
| CNAG_00457 | 23872.13 | 0.259532 | 0.077625 | 3.343418 | 0.000828 | 0.004024 |
| CNAG_00461 | 3307.501 | -0.20125 | 0.070036 | -2.87346 | 0.00406 | 0.015874 |
| CNAG_00462 | 6061.184 | 0.169238 | 0.080778 | 2.095089 | 0.036163 | 0.096025 |
| CNAG_12060 | 194.537 | -0.68935 | 0.155256 | -4.44007 | 8.99E-06 | 6.83E-05 |
| CNAG_00465 | 1870.958 | 0.603871 | 0.101681 | 5.938892 | 2.87E-09 | 3.61E-08 |
| CNAG_00467 | 323.3853 | 0.326864 | 0.128916 | 2.535489 | 0.011229 | 0.037242 |
| CNAG_00468 | 1891.303 | 0.222921 | 0.071192 | 3.131273 | 0.001741 | 0.007652 |
| CNAG_00471 | 1754.085 | 0.841611 | 0.083609 | 10.06599 | 7.81E-24 | 3.98E-22 |
| CNAG_00474 | 214.193 | 0.506579 | 0.169162 | 2.994643 | 0.002748 | 0.011375 |
| CNAG_07362 | 9966.675 | -0.41621 | 0.080972 | -5.14013 | 2.75E-07 | 2.63E-06 |
| CNAG_07363 | 11222.87 | 1.12379 | 0.075682 | 14.84874 | 7.09E-50 | 1.50E-47 |
| CNAG_00480 | 1485.43 | 0.810909 | 0.093749 | 8.649831 | 5.16E-18 | 1.50E-16 |
| CNAG_00483 | 17895.43 | -0.13196 | 0.062602 | -2.108 | 0.035031 | 0.093691 |
| CNAG_00485 | 1293.975 | 1.141769 | 0.12783 | 8.931964 | 4.19E-19 | 1.38E-17 |
| CNAG_00486 | 1953.556 | 0.430115 | 0.082976 | 5.183586 | 2.18E-07 | 2.10E-06 |
| CNAG_00491 | 822.6642 | 0.303663 | 0.090725 | 3.347059 | 0.000817 | 0.003979 |
| CNAG_07364 | 670.5411 | -0.2181 | 0.0926 | -2.35528 | 0.018509 | 0.055856 |
| CNAG_00497 | 2584.491 | 0.757063 | 0.076035 | 9.956818 | 2.35E-23 | 1.15E-21 |
| CNAG_00498 | 636.582 | 0.249318 | 0.096121 | 2.593784 | 0.009493 | 0.032477 |
| CNAG_00500 | 2300.725 | 0.171479 | 0.074516 | 2.301222 | 0.021379 | 0.062919 |
| CNAG_00503 | 966.7956 | 0.17297 | 0.082749 | 2.090295 | 0.036591 | 0.096832 |
| CNAG_00505 | 349.4323 | -0.48189 | 0.125645 | -3.8353 | 0.000125 | 0.000756 |
| CNAG_00513 | 2992.838 | -0.5183 | 0.068539 | -7.56201 | 3.97E-14 | 8.49E-13 |
| CNAG_00519 | 2665.757 | -0.56765 | 0.073357 | -7.73823 | 1.01E-14 | 2.27E-13 |
| CNAG_00520 | 4963.452 | -0.17281 | 0.085985 | -2.00981 | 0.044452 | 0.113668 |
| CNAG_00521 | 1297.149 | 0.50388 | 0.080117 | 6.289331 | 3.19E-10 | 4.54E-09 |
| CNAG_00522 | 3246.738 | 0.654199 | 0.068037 | 9.615336 | 6.89E-22 | 2.91E-20 |
| CNAG_00527 | 3783.329 | -0.22197 | 0.069067 | -3.21379 | 0.00131 | 0.005992 |
| CNAG_00528 | 674.6286 | 0.312526 | 0.098636 | 3.168471 | 0.001532 | 0.006849 |
| CNAG_00529 | 1056.068 | -0.54534 | 0.091699 | -5.94708 | 2.73E-09 | 3.44E-08 |
| CNAG_00531 | 3166.48 | 0.566303 | 0.065203 | 8.685169 | 3.78E-18 | 1.13E-16 |
| CNAG_12064 | 38.18052 | 1.133265 | 0.334218 | 3.390799 | 0.000697 | 0.003486 |
| CNAG_00532 | 1118.518 | -0.52198 | 0.081229 | -6.4261 | 1.31E-10 | 1.94E-09 |
| CNAG_00533 | 5147.509 | 0.671883 | 0.06415 | 10.4737 | 1.14E-25 | 6.55E-24 |
| CNAG_00534 | 4457.752 | 0.200872 | 0.071418 | 2.812635 | 0.004914 | 0.018543 |
| CNAG_00537 | 428.7743 | 0.268993 | 0.126718 | 2.122768 | 0.033773 | 0.090916 |
| CNAG_00539 | 547.6385 | 0.919586 | 0.117442 | 7.830145 | 4.87E-15 | 1.13E-13 |
| CNAG_00543 | 1093.06 | -0.31167 | 0.085291 | -3.65423 | 0.000258 | 0.001448 |

| | | | | | | |
|---|---|---|---|---|---|---|
| CNAG_00546 | 1325.496 | -0.21496 | 0.100884 | -2.13076 | 0.033109 | 0.08956 |
| CNAG_07955 | 381.9846 | 0.276029 | 0.11463 | 2.408006 | 0.01604 | 0.049791 |
| CNAG_00550 | 506.3894 | 0.369591 | 0.109544 | 3.373894 | 0.000741 | 0.003668 |
| CNAG_00551 | 5444.647 | -0.23323 | 0.070148 | -3.32481 | 0.000885 | 0.004273 |
| CNAG_00554 | 967.2645 | 0.233493 | 0.085276 | 2.7381 | 0.00618 | 0.022504 |
| CNAG_00557 | 1661.027 | -0.18886 | 0.071771 | -2.63138 | 0.008504 | 0.02964 |
| CNAG_00563 | 1126.387 | 0.424014 | 0.09603 | 4.415426 | 1.01E-05 | 7.62E-05 |
| CNAG_00564 | 820.5287 | 0.205167 | 0.090683 | 2.262461 | 0.023669 | 0.068285 |
| CNAG_00565 | 5806.928 | -0.22285 | 0.063666 | -3.50029 | 0.000465 | 0.002449 |
| CNAG_00566 | 1352.818 | 0.35866 | 0.082727 | 4.335449 | 1.45E-05 | 0.000107 |
| CNAG_00573 | 2364.372 | 0.221272 | 0.083601 | 2.646764 | 0.008127 | 0.028671 |
| CNAG_00575 | 2050.531 | 1.177389 | 0.2127 | 5.535444 | 3.10E-08 | 3.38E-07 |
| CNAG_00577 | 796.546 | 0.234181 | 0.088833 | 2.636207 | 0.008384 | 0.029274 |
| CNAG_00580 | 3576.142 | -0.17545 | 0.084261 | -2.08218 | 0.037326 | 0.098424 |
| CNAG_00581 | 7037.32 | 0.335918 | 0.067263 | 4.994099 | 5.91E-07 | 5.35E-06 |
| CNAG_00582 | 2266.199 | 0.399263 | 0.078253 | 5.102181 | 3.36E-07 | 3.17E-06 |
| CNAG_00583 | 3497.438 | -0.18178 | 0.091986 | -1.97612 | 0.048141 | 0.12099 |
| CNAG_00585 | 1463.196 | -0.70509 | 0.081778 | -8.62202 | 6.58E-18 | 1.90E-16 |
| CNAG_00588 | 292.8255 | 0.789089 | 0.126231 | 6.251136 | 4.07E-10 | 5.70E-09 |
| CNAG_00590 | 2703.997 | -0.25477 | 0.075438 | -3.37714 | 0.000732 | 0.003631 |
| CNAG_00591 | 134.2618 | 0.482809 | 0.203105 | 2.377143 | 0.017447 | 0.053333 |
| CNAG_12079 | 291.1179 | 0.523769 | 0.134516 | 3.893738 | 9.87E-05 | 0.000608 |
| CNAG_00592 | 452.7693 | -0.28727 | 0.116254 | -2.47105 | 0.013472 | 0.043229 |
| CNAG_00595 | 507.2928 | 0.736808 | 0.110246 | 6.683323 | 2.34E-11 | 3.83E-10 |
| CNAG_00596 | 167.5394 | 0.498656 | 0.156825 | 3.179695 | 0.001474 | 0.006639 |
| CNAG_00597 | 280.5505 | -0.56604 | 0.149323 | -3.79071 | 0.00015 | 0.000887 |
| CNAG_00598 | 43.70662 | 0.77968 | 0.338428 | 2.30383 | 0.021232 | 0.062645 |
| CNAG_00602 | 5018.921 | -0.23056 | 0.074598 | -3.09074 | 0.001997 | 0.008604 |
| CNAG_00603 | 1405.753 | -0.48424 | 0.083165 | -5.8226 | 5.79E-09 | 7.00E-08 |
| CNAG_00607 | 1650.294 | -0.40338 | 0.075464 | -5.34533 | 9.03E-08 | 9.21E-07 |
| CNAG_07957 | 1831.576 | -0.40635 | 0.075332 | -5.39407 | 6.89E-08 | 7.12E-07 |
| CNAG_00619 | 683.1582 | -0.21592 | 0.09849 | -2.19234 | 0.028355 | 0.079655 |
| CNAG_07374 | 969.8705 | -0.48408 | 0.091939 | -5.26526 | 1.40E-07 | 1.38E-06 |
| CNAG_00625 | 1886.941 | 0.174455 | 0.088455 | 1.972243 | 0.048582 | 0.121751 |
| CNAG_00626 | 9321.154 | 0.394367 | 0.066618 | 5.919862 | 3.22E-09 | 4.03E-08 |
| CNAG_00627 | 153.0265 | 0.705947 | 0.179771 | 3.92692 | 8.60E-05 | 0.00054 |
| CNAG_07376 | 2418.857 | -0.1776 | 0.070473 | -2.52014 | 0.011731 | 0.038676 |
| CNAG_00633 | 2572.42 | -0.41439 | 0.083984 | -4.93408 | 8.05E-07 | 7.09E-06 |
| CNAG_00637 | 164.6526 | 0.815944 | 0.187966 | 4.340926 | 1.42E-05 | 0.000104 |
| CNAG_00638 | 2734.528 | 0.787371 | 0.087064 | 9.043642 | 1.52E-19 | 5.30E-18 |
| CNAG_00640 | 87776.28 | -0.24537 | 0.072139 | -3.40138 | 0.00067 | 0.003365 |
| CNAG_00641 | 4401.029 | -0.21226 | 0.083453 | -2.54344 | 0.010977 | 0.036561 |
| CNAG_00643 | 1053.121 | -0.29807 | 0.082382 | -3.61815 | 0.000297 | 0.001631 |
| CNAG_00647 | 803.5786 | -0.2379 | 0.106239 | -2.23928 | 0.025138 | 0.072018 |
| CNAG_00652 | 564.0581 | 0.211867 | 0.097038 | 2.183337 | 0.029011 | 0.081087 |
| CNAG_00656 | 58586.95 | -0.24208 | 0.070669 | -3.42551 | 0.000614 | 0.003126 |
| CNAG_00659 | 377.5149 | -0.24284 | 0.115219 | -2.10764 | 0.035062 | 0.093691 |
| CNAG_00662 | 1880.432 | 0.213355 | 0.079122 | 2.696512 | 0.007007 | 0.025095 |

| | | | | | | |
|---|---|---|---|---|---|---|
| CNAG_00663 | 954.551 | 0.3663 | 0.092139 | 3.975516 | 7.02E-05 | 0.000451 |
| CNAG_00666 | 3115.148 | -0.25221 | 0.095624 | -2.63752 | 0.008351 | 0.029213 |
| CNAG_00669 | 791.5782 | -0.57364 | 0.095279 | -6.02066 | 1.74E-09 | 2.24E-08 |
| CNAG_00672 | 46279.72 | -0.30365 | 0.077173 | -3.93467 | 8.33E-05 | 0.000526 |
| CNAG_00673 | 2653.11 | 0.255633 | 0.06627 | 3.85747 | 0.000115 | 0.000695 |
| CNAG_00674 | 1701.648 | 0.384805 | 0.108685 | 3.540561 | 0.000399 | 0.002131 |
| CNAG_00675 | 914.4221 | -0.38955 | 0.09076 | -4.29213 | 1.77E-05 | 0.000128 |
| CNAG_00680 | 1549.456 | -0.19104 | 0.076634 | -2.4929 | 0.01267 | 0.041234 |
| CNAG_00686 | 16716.75 | 0.784895 | 0.081323 | 9.651574 | 4.84E-22 | 2.09E-20 |
| CNAG_00688 | 476.0841 | -0.36744 | 0.104251 | -3.5246 | 0.000424 | 0.002249 |
| CNAG_00696 | 727.6468 | 0.414075 | 0.097906 | 4.229324 | 2.34E-05 | 0.000163 |
| CNAG_00697 | 5290.483 | 0.139692 | 0.06333 | 2.205793 | 0.027399 | 0.077498 |
| CNAG_00698 | 2114.03 | 0.330379 | 0.076775 | 4.303233 | 1.68E-05 | 0.000122 |
| CNAG_12092 | 158.9682 | -0.48365 | 0.167204 | -2.89256 | 0.003821 | 0.015084 |
| CNAG_00700 | 11655.95 | -0.2116 | 0.060259 | -3.51152 | 0.000446 | 0.002357 |
| CNAG_00701 | 2651.525 | 0.133724 | 0.065943 | 2.027878 | 0.042573 | 0.109831 |
| CNAG_00702 | 3577.116 | -0.31984 | 0.080921 | -3.95247 | 7.73E-05 | 0.000492 |
| CNAG_00703 | 41251.74 | -0.16763 | 0.072209 | -2.32151 | 0.02026 | 0.060231 |
| CNAG_00706 | 1619.114 | -0.49625 | 0.075918 | -6.53667 | 6.29E-11 | 9.70E-10 |
| CNAG_00707 | 2168.993 | -0.644 | 0.078093 | -8.24658 | 1.63E-16 | 4.21E-15 |
| CNAG_00708 | 2177.472 | -0.18849 | 0.078725 | -2.39426 | 0.016654 | 0.051208 |
| CNAG_00710 | 2020.283 | 0.669618 | 0.107256 | 6.243151 | 4.29E-10 | 5.99E-09 |
| CNAG_00715 | 1739.219 | 0.276874 | 0.076694 | 3.610123 | 0.000306 | 0.001676 |
| CNAG_00716 | 27174.34 | 0.670161 | 0.058908 | 11.37634 | 5.49E-30 | 3.89E-28 |
| CNAG_00717 | 868.95 | 0.376683 | 0.089229 | 4.221508 | 2.43E-05 | 0.000169 |
| CNAG_00718 | 951.3492 | -0.24184 | 0.08921 | -2.71092 | 0.00671 | 0.02423 |
| CNAG_00719 | 2475.124 | -0.31605 | 0.088406 | -3.57503 | 0.00035 | 0.001897 |
| CNAG_00720 | 1330.905 | -0.23239 | 0.088709 | -2.61969 | 0.008801 | 0.030566 |
| CNAG_07381 | 1452.06 | -0.53468 | 0.092524 | -5.77884 | 7.52E-09 | 8.95E-08 |
| CNAG_12095 | 39.37405 | -0.77632 | 0.338531 | -2.2932 | 0.021836 | 0.064032 |
| CNAG_00727 | 1714.436 | 0.392633 | 0.078102 | 5.0272 | 4.98E-07 | 4.58E-06 |
| CNAG_00728 | 200.4943 | -0.81357 | 0.154043 | -5.28144 | 1.28E-07 | 1.27E-06 |
| CNAG_00730 | 3907.278 | -0.53147 | 0.088445 | -6.00911 | 1.87E-09 | 2.40E-08 |
| CNAG_00732 | 3163.547 | 0.783679 | 0.091612 | 8.554311 | 1.19E-17 | 3.32E-16 |
| CNAG_00733 | 1694.399 | -0.5267 | 0.074388 | -7.08046 | 1.44E-12 | 2.68E-11 |
| CNAG_00738 | 93.14626 | -0.62781 | 0.212019 | -2.9611 | 0.003065 | 0.012466 |
| CNAG_00739 | 1918.053 | -0.17142 | 0.075675 | -2.26516 | 0.023503 | 0.067881 |
| CNAG_00741 | 12329.03 | -0.22346 | 0.0665 | -3.36036 | 0.000778 | 0.003818 |
| CNAG_00743 | 3442.525 | -0.23317 | 0.065307 | -3.57035 | 0.000356 | 0.001923 |
| CNAG_12103 | 46.93698 | 0.611588 | 0.285293 | 2.143718 | 0.032055 | 0.087438 |
| CNAG_00745 | 2578.961 | 0.520137 | 0.088511 | 5.876549 | 4.19E-09 | 5.18E-08 |
| CNAG_00747 | 11315.05 | 0.659842 | 0.064991 | 10.15278 | 3.22E-24 | 1.72E-22 |
| CNAG_00750 | 1143.947 | -0.28169 | 0.091276 | -3.08617 | 0.002028 | 0.008703 |
| CNAG_00752 | 421.9066 | -0.46585 | 0.134905 | -3.4532 | 0.000554 | 0.002855 |
| CNAG_00755 | 689.5308 | 0.305855 | 0.099101 | 3.086291 | 0.002027 | 0.008703 |
| CNAG_12110 | 136.2164 | 0.499738 | 0.189259 | 2.640502 | 0.008278 | 0.029039 |
| CNAG_00760 | 1876.518 | -0.22096 | 0.0738 | -2.99409 | 0.002753 | 0.01139 |
| CNAG_00762 | 687.4904 | -0.68542 | 0.092159 | -7.43735 | 1.03E-13 | 2.14E-12 |

| | | | | | | |
|---|---|---|---|---|---|---|
| CNAG_07960 | 367.2812 | -0.257 | 0.114654 | -2.24154 | 0.024991 | 0.071623 |
| CNAG_00764 | 2707.502 | -0.63216 | 0.070431 | -8.97557 | 2.82E-19 | 9.48E-18 |
| CNAG_00765 | 673.2026 | 0.397478 | 0.098539 | 4.033694 | 5.49E-05 | 0.000359 |
| CNAG_00766 | 606.8175 | 0.205772 | 0.10192 | 2.018945 | 0.043493 | 0.111651 |
| CNAG_00768 | 1027.272 | 0.296037 | 0.086605 | 3.418248 | 0.00063 | 0.003192 |
| CNAG_00771 | 16463.81 | -0.22942 | 0.072143 | -3.18 | 0.001473 | 0.006636 |
| CNAG_00772 | 2633.486 | -0.17685 | 0.085523 | -2.06785 | 0.038654 | 0.101432 |
| CNAG_00773 | 619.876 | 0.443169 | 0.100319 | 4.417579 | 9.98E-06 | 7.56E-05 |
| CNAG_00774 | 6271.495 | -0.39134 | 0.074367 | -5.26225 | 1.42E-07 | 1.41E-06 |
| CNAG_00775 | 1935.152 | -0.59989 | 0.075684 | -7.92625 | 2.26E-15 | 5.37E-14 |
| CNAG_00776 | 15678.65 | 0.328552 | 0.071428 | 4.599747 | 4.23E-06 | 3.39E-05 |
| CNAG_00777 | 2272.424 | -0.34109 | 0.080569 | -4.23353 | 2.30E-05 | 0.000161 |
| CNAG_00779 | 40203.54 | -0.18771 | 0.071218 | -2.63568 | 0.008397 | 0.029306 |
| CNAG_00785 | 32625.59 | -0.24324 | 0.071395 | -3.40697 | 0.000657 | 0.003305 |
| CNAG_00786 | 638.7581 | 0.595714 | 0.099094 | 6.011608 | 1.84E-09 | 2.37E-08 |
| CNAG_00788 | 13613.58 | -0.26998 | 0.070681 | -3.81967 | 0.000134 | 0.000797 |
| CNAG_00789 | 4.154357 | 2.56963 | 1.096795 | 2.342854 | 0.019137 | NA |
| CNAG_00790 | 847.5436 | 0.211892 | 0.089023 | 2.380189 | 0.017304 | 0.052935 |
| CNAG_00791 | 1869.798 | 0.763899 | 0.080454 | 9.494874 | 2.20E-21 | 9.01E-20 |
| CNAG_00793 | 1873.215 | -0.32834 | 0.075963 | -4.32233 | 1.54E-05 | 0.000112 |
| CNAG_00794 | 1282.257 | -0.19165 | 0.076811 | -2.49514 | 0.012591 | 0.041026 |
| CNAG_00796 | 1485.204 | 0.328532 | 0.106946 | 3.07193 | 0.002127 | 0.009055 |
| CNAG_00800 | 320.4779 | 0.469522 | 0.159256 | 2.948226 | 0.003196 | 0.012943 |
| CNAG_00802 | 451.065 | 0.528489 | 0.109167 | 4.841087 | 1.29E-06 | 1.11E-05 |
| CNAG_00803 | 1267.1 | 0.339537 | 0.080264 | 4.230226 | 2.33E-05 | 0.000163 |
| CNAG_00807 | 917.2597 | 0.42145 | 0.093802 | 4.492988 | 7.02E-06 | 5.43E-05 |
| CNAG_00809 | 4190.756 | -0.5999 | 0.067839 | -8.84292 | 9.33E-19 | 2.95E-17 |
| CNAG_00810 | 1328.721 | -0.18331 | 0.082587 | -2.21956 | 0.026449 | 0.075111 |
| CNAG_00815 | 1221.923 | -0.35499 | 0.083522 | -4.25021 | 2.14E-05 | 0.00015 |
| CNAG_00819 | 12415.82 | -0.22419 | 0.068333 | -3.28083 | 0.001035 | 0.004898 |
| CNAG_00821 | 25106.73 | -0.21648 | 0.080202 | -2.6992 | 0.006951 | 0.024928 |
| CNAG_00822 | 4666.532 | -0.35134 | 0.06862 | -5.12008 | 3.05E-07 | 2.90E-06 |
| CNAG_00834 | 1055.267 | 1.077333 | 0.099089 | 10.87238 | 1.56E-27 | 1.04E-25 |
| CNAG_00836 | 504.3689 | -0.30025 | 0.108282 | -2.77287 | 0.005556 | 0.020541 |
| CNAG_00838 | 258.1273 | 0.539385 | 0.139537 | 3.865548 | 0.000111 | 0.000675 |
| CNAG_00841 | 380.5485 | -0.30996 | 0.124754 | -2.48454 | 0.012972 | 0.04204 |
| CNAG_00843 | 254.9845 | 0.40244 | 0.140503 | 2.864289 | 0.004179 | 0.016251 |
| CNAG_00845 | 1948.438 | -0.52362 | 0.074027 | -7.0734 | 1.51E-12 | 2.80E-11 |
| CNAG_00846 | 799.1004 | -0.52182 | 0.091097 | -5.7282 | 1.02E-08 | 1.19E-07 |
| CNAG_00848 | 3790.923 | 1.610045 | 0.200843 | 8.016449 | 1.09E-15 | 2.65E-14 |
| CNAG_00851 | 171.343 | 0.834074 | 0.181028 | 4.607441 | 4.08E-06 | 3.27E-05 |
| CNAG_00854 | 2887.576 | -0.7953 | 0.087952 | -9.0425 | 1.53E-19 | 5.31E-18 |
| CNAG_00855 | 794.9392 | 0.300863 | 0.094326 | 3.189615 | 0.001425 | 0.006453 |
| CNAG_07389 | 552.2446 | -0.46916 | 0.106328 | -4.41241 | 1.02E-05 | 7.70E-05 |
| CNAG_07392 | 403.7308 | 0.350784 | 0.113193 | 3.098997 | 0.001942 | 0.008405 |
| CNAG_07393 | 198.8998 | 0.391989 | 0.155841 | 2.515307 | 0.011893 | 0.039095 |
| CNAG_06800 | 1362.278 | 1.186491 | 0.099666 | 11.9047 | 1.12E-32 | 9.49E-31 |
| CNAG_06798 | 1952.386 | -0.39069 | 0.073855 | -5.28997 | 1.22E-07 | 1.22E-06 |

| | | | | | | |
|---|---|---|---|---|---|---|
| CNAG_06794 | 2128.189 | 0.536676 | 0.077791 | 6.89894 | 5.24E-12 | 9.13E-11 |
| CNAG_06793 | 1335.432 | 0.343373 | 0.08042 | 4.269737 | 1.96E-05 | 0.000139 |
| CNAG_06792 | 1130.182 | 0.184149 | 0.081121 | 2.270041 | 0.023205 | 0.067246 |
| CNAG_06791 | 1764.578 | 1.280478 | 0.100935 | 12.68617 | 7.06E-37 | 8.10E-35 |
| CNAG_06785 | 367.8878 | -0.42888 | 0.126602 | -3.3876 | 0.000705 | 0.003514 |
| CNAG_06784 | 97.09756 | -0.54893 | 0.207851 | -2.64098 | 0.008267 | 0.029039 |
| CNAG_06782 | 1726.689 | 0.253358 | 0.08599 | 2.946356 | 0.003215 | 0.013002 |
| CNAG_06781 | 1308.625 | 0.758784 | 0.083194 | 9.120678 | 7.47E-20 | 2.67E-18 |
| CNAG_06774 | 1213.706 | -0.4784 | 0.078008 | -6.13263 | 8.64E-10 | 1.17E-08 |
| CNAG_06771 | 577.584 | 0.695258 | 0.112082 | 6.203124 | 5.54E-10 | 7.55E-09 |
| CNAG_06770 | 32082.81 | 0.358193 | 0.069326 | 5.166781 | 2.38E-07 | 2.29E-06 |
| CNAG_06769 | 1004.67 | -0.34055 | 0.088114 | -3.8649 | 0.000111 | 0.000677 |
| CNAG_06766 | 2592.628 | -0.6082 | 0.071355 | -8.5236 | 1.55E-17 | 4.27E-16 |
| CNAG_07962 | 110.0085 | -0.7674 | 0.193782 | -3.96013 | 7.49E-05 | 0.000478 |
| CNAG_06765 | 1566.48 | -0.34039 | 0.101108 | -3.36665 | 0.000761 | 0.003751 |
| CNAG_06764 | 2400.663 | -0.19149 | 0.069249 | -2.76523 | 0.005688 | 0.020929 |
| CNAG_06763 | 1200.83 | 0.174076 | 0.080574 | 2.160447 | 0.030738 | 0.084644 |
| CNAG_06762 | 2361.643 | -0.59921 | 0.070295 | -8.52411 | 1.54E-17 | 4.26E-16 |
| CNAG_06761 | 193.1903 | -0.70856 | 0.15235 | -4.65088 | 3.31E-06 | 2.68E-05 |
| CNAG_06760 | 1014.211 | -0.67776 | 0.086651 | -7.82174 | 5.21E-15 | 1.20E-13 |
| CNAG_06759 | 829.0469 | 0.661085 | 0.095106 | 6.951044 | 3.63E-12 | 6.37E-11 |
| CNAG_06758 | 614.7194 | -1.12348 | 0.114969 | -9.772 | 1.48E-22 | 6.66E-21 |
| CNAG_06756 | 803.102 | -0.44055 | 0.092702 | -4.75236 | 2.01E-06 | 1.68E-05 |
| CNAG_06753 | 5342.603 | -0.13216 | 0.061831 | -2.13736 | 0.032568 | 0.088374 |
| CNAG_06751 | 3099.824 | 0.545243 | 0.07267 | 7.502995 | 6.24E-14 | 1.31E-12 |
| CNAG_06749 | 731.5475 | -0.27797 | 0.117257 | -2.37057 | 0.017761 | 0.054037 |
| CNAG_06748 | 2434.781 | -0.55168 | 0.069975 | -7.88406 | 3.17E-15 | 7.43E-14 |
| CNAG_06747 | 33228.71 | -0.20021 | 0.072067 | -2.77805 | 0.005469 | 0.020264 |
| CNAG_06746 | 33053.56 | -0.2203 | 0.062922 | -3.50114 | 0.000463 | 0.002444 |
| CNAG_06745 | 29322.55 | -0.32185 | 0.073616 | -4.372 | 1.23E-05 | 9.17E-05 |
| CNAG_06742 | 2443.88 | -0.5829 | 0.073525 | -7.92794 | 2.23E-15 | 5.32E-14 |
| CNAG_06741 | 2380.036 | -0.73248 | 0.088983 | -8.23177 | 1.84E-16 | 4.75E-15 |
| CNAG_06739 | 3620.945 | 0.760242 | 0.066751 | 11.38925 | 4.73E-30 | 3.39E-28 |
| CNAG_06737 | 1663.72 | 0.333043 | 0.081243 | 4.099349 | 4.14E-05 | 0.000276 |
| CNAG_06732 | 1641.157 | 0.378103 | 0.079203 | 4.773825 | 1.81E-06 | 1.52E-05 |
| CNAG_06729 | 857.7877 | 0.803995 | 0.093705 | 8.58006 | 9.48E-18 | 2.69E-16 |
| CNAG_06723 | 5976.41 | 0.647409 | 0.074706 | 8.66604 | 4.47E-18 | 1.32E-16 |
| CNAG_07465 | 797.5654 | -0.24309 | 0.102864 | -2.36321 | 0.018117 | 0.054899 |
| CNAG_06713 | 3324.841 | -0.18878 | 0.066463 | -2.84045 | 0.004505 | 0.017268 |
| CNAG_06711 | 838.4334 | 0.269618 | 0.088588 | 3.043496 | 0.002338 | 0.009833 |
| CNAG_06708 | 650.5178 | -0.21794 | 0.096023 | -2.26961 | 0.023231 | 0.067296 |
| CNAG_06707 | 162.5951 | -0.41487 | 0.166898 | -2.48576 | 0.012927 | 0.041913 |
| CNAG_06706 | 500.6528 | 0.70518 | 0.124923 | 5.644923 | 1.65E-08 | 1.88E-07 |
| CNAG_06703 | 719.2016 | -0.3001 | 0.096262 | -3.11751 | 0.001824 | 0.007961 |
| CNAG_03595 | 3022.792 | 0.331757 | 0.095017 | 3.491565 | 0.00048 | 0.00252 |
| CNAG_03596 | 9622.514 | 0.787163 | 0.063648 | 12.3675 | 3.92E-35 | 3.87E-33 |
| CNAG_12146 | 16.61192 | -1.49499 | 0.50465 | -2.96244 | 0.003052 | 0.012419 |
| CNAG_03597 | 150.4832 | -0.45708 | 0.182679 | -2.50211 | 0.012345 | 0.040294 |

| | | | | | | |
|---|---|---|---|---|---|---|
| CNAG_03599 | 328.3147 | -0.67456 | 0.128684 | -5.24199 | 1.59E-07 | 1.56E-06 |
| CNAG_03602 | 10830.03 | -0.14842 | 0.064807 | -2.29016 | 0.022012 | 0.064386 |
| CNAG_03603 | 1963.293 | -0.71351 | 0.075248 | -9.48212 | 2.49E-21 | 1.01E-19 |
| CNAG_03606 | 9268.047 | -0.15562 | 0.068441 | -2.27374 | 0.022982 | 0.066704 |
| CNAG_03609 | 748.5521 | 0.441803 | 0.09453 | 4.673661 | 2.96E-06 | 2.42E-05 |
| CNAG_03618 | 290.1191 | -1.35723 | 0.136563 | -9.93848 | 2.83E-23 | 1.37E-21 |
| CNAG_03621 | 12584.74 | -0.17882 | 0.067725 | -2.64038 | 0.008281 | 0.029039 |
| CNAG_03624 | 5735.493 | -0.58752 | 0.065111 | -9.02332 | 1.82E-19 | 6.25E-18 |
| CNAG_03627 | 9724.413 | -0.20169 | 0.070039 | -2.87972 | 0.00398 | 0.015625 |
| CNAG_03629 | 11434.26 | 0.159381 | 0.057699 | 2.762281 | 0.00574 | 0.021083 |
| CNAG_03631 | 1236.268 | 0.492864 | 0.098797 | 4.98863 | 6.08E-07 | 5.48E-06 |
| CNAG_03632 | 1309.275 | 0.478169 | 0.087208 | 5.483058 | 4.18E-08 | 4.48E-07 |
| CNAG_03634 | 1105.446 | -0.59253 | 0.089148 | -6.64661 | 3.00E-11 | 4.78E-10 |
| CNAG_03636 | 1243.738 | -0.4053 | 0.082796 | -4.89511 | 9.83E-07 | 8.54E-06 |
| CNAG_03637 | 1132.807 | 0.308333 | 0.085561 | 3.603659 | 0.000314 | 0.001714 |
| CNAG_03638 | 2047.506 | 0.260431 | 0.081415 | 3.198823 | 0.00138 | 0.006261 |
| CNAG_03639 | 175.446 | -0.59934 | 0.154904 | -3.86908 | 0.000109 | 0.000667 |
| CNAG_03645 | 9165.792 | -0.31231 | 0.071824 | -4.34833 | 1.37E-05 | 0.000101 |
| CNAG_03646 | 1230.406 | -0.18518 | 0.081032 | -2.28524 | 0.022299 | 0.065103 |
| CNAG_03647 | 950.3065 | 0.272456 | 0.085661 | 3.180636 | 0.00147 | 0.006625 |
| CNAG_03654 | 976.2806 | -0.18526 | 0.092241 | -2.00844 | 0.044596 | 0.113922 |
| CNAG_03655 | 3392.419 | -0.36087 | 0.108572 | -3.32373 | 0.000888 | 0.004281 |
| CNAG_03661 | 551.7273 | 0.260743 | 0.099054 | 2.632317 | 0.00848 | 0.029572 |
| CNAG_03663 | 1413.154 | 0.414405 | 0.075321 | 5.501876 | 3.76E-08 | 4.06E-07 |
| CNAG_03664 | 2064.58 | -0.16901 | 0.073915 | -2.28649 | 0.022226 | 0.064914 |
| CNAG_03666 | 363.6211 | 0.336129 | 0.127235 | 2.641784 | 0.008247 | 0.029008 |
| CNAG_03667 | 3018.225 | 0.820822 | 0.076909 | 10.67268 | 1.37E-26 | 8.27E-25 |
| CNAG_03671 | 349.1286 | 0.355573 | 0.133408 | 2.665307 | 0.007692 | 0.027248 |
| CNAG_03673 | 1053.676 | 0.286075 | 0.098199 | 2.91322 | 0.003577 | 0.014309 |
| CNAG_03674 | 5602.077 | 0.773577 | 0.061252 | 12.62951 | 1.45E-36 | 1.60E-34 |
| CNAG_03675 | 3994.448 | -0.66539 | 0.068652 | -9.69218 | 3.26E-22 | 1.41E-20 |
| CNAG_03676 | 930.4278 | 0.225923 | 0.083405 | 2.708744 | 0.006754 | 0.024356 |
| CNAG_03679 | 379.6632 | 0.539081 | 0.117388 | 4.592309 | 4.38E-06 | 3.50E-05 |
| CNAG_03681 | 2957.734 | -0.17743 | 0.06715 | -2.6423 | 0.008235 | 0.028986 |
| CNAG_03683 | 477.2899 | -0.74174 | 0.112937 | -6.5677 | 5.11E-11 | 7.93E-10 |
| CNAG_03684 | 4489.359 | 0.133542 | 0.066305 | 2.014043 | 0.044005 | 0.112817 |
| CNAG_03688 | 2361.523 | 0.655 | 0.080177 | 8.169441 | 3.10E-16 | 7.84E-15 |
| CNAG_03689 | 1850.58 | 0.678223 | 0.082008 | 8.270243 | 1.34E-16 | 3.49E-15 |
| CNAG_07473 | 6013.493 | 0.123536 | 0.061584 | 2.005969 | 0.04486 | 0.114409 |
| CNAG_03692 | 522.706 | 0.700947 | 0.129232 | 5.423949 | 5.83E-08 | 6.10E-07 |
| CNAG_03697 | 316.899 | 0.388532 | 0.129137 | 3.008675 | 0.002624 | 0.010904 |
| CNAG_03699 | 3274.756 | -0.26829 | 0.070824 | -3.7882 | 0.000152 | 0.000892 |
| CNAG_03705 | 3030.614 | 0.408589 | 0.07836 | 5.214285 | 1.85E-07 | 1.80E-06 |
| CNAG_03710 | 1593.907 | -0.38558 | 0.091652 | -4.20703 | 2.59E-05 | 0.000179 |
| CNAG_03716 | 3656.425 | -0.19773 | 0.069062 | -2.86303 | 0.004196 | 0.016299 |
| CNAG_03719 | 1335.766 | -0.41199 | 0.105528 | -3.90405 | 9.46E-05 | 0.000584 |
| CNAG_03724 | 6181.539 | -0.6464 | 0.070054 | -9.22717 | 2.78E-20 | 1.03E-18 |
| CNAG_03725 | 7463.028 | 0.311292 | 0.06402 | 4.862392 | 1.16E-06 | 1.00E-05 |

| | | | | | | |
|---|---|---|---|---|---|---|
| CNAG_03726 | 3376.766 | -0.16112 | 0.076053 | -2.11859 | 0.034125 | 0.091674 |
| CNAG_12161 | 155.8249 | 0.526597 | 0.176201 | 2.988618 | 0.002802 | 0.011565 |
| CNAG_03727 | 1676.123 | 0.26753 | 0.090837 | 2.945162 | 0.003228 | 0.013038 |
| CNAG_03728 | 1059.908 | 0.403179 | 0.107386 | 3.7545 | 0.000174 | 0.001008 |
| CNAG_03732 | 711.6112 | 0.195365 | 0.096494 | 2.024645 | 0.042904 | 0.110466 |
| CNAG_03735 | 1178.029 | -0.31639 | 0.081717 | -3.87172 | 0.000108 | 0.00066 |
| CNAG_03738 | 1574.718 | 0.573028 | 0.081889 | 6.997635 | 2.60E-12 | 4.64E-11 |
| CNAG_03739 | 40818.5 | -0.16876 | 0.069724 | -2.42037 | 0.015505 | 0.048477 |
| CNAG_03747 | 37968.51 | -0.32467 | 0.078127 | -4.15575 | 3.24E-05 | 0.00022 |
| CNAG_03748 | 4431.341 | -0.14169 | 0.065544 | -2.16179 | 0.030634 | 0.084507 |
| CNAG_03753 | 756.8126 | -0.33953 | 0.088731 | -3.82657 | 0.00013 | 0.000779 |
| CNAG_03756 | 1339.068 | 0.167317 | 0.08374 | 1.998046 | 0.045712 | 0.116049 |
| CNAG_03759 | 22.77619 | 1.991348 | 0.434318 | 4.585 | 4.54E-06 | 3.62E-05 |
| CNAG_03762 | 4037.07 | -0.60603 | 0.06692 | -9.05603 | 1.35E-19 | 4.80E-18 |
| CNAG_03763 | 3034.831 | -0.31456 | 0.107894 | -2.91545 | 0.003552 | 0.014221 |
| CNAG_03765 | 7995.571 | 0.399589 | 0.060805 | 6.571674 | 4.98E-11 | 7.74E-10 |
| CNAG_03766 | 993.9255 | -0.37893 | 0.089478 | -4.23494 | 2.29E-05 | 0.00016 |
| CNAG_03769 | 10933.94 | 0.293718 | 0.063035 | 4.659575 | 3.17E-06 | 2.57E-05 |
| CNAG_03770 | 151.1992 | 0.451938 | 0.175174 | 2.579945 | 0.009882 | 0.033543 |
| CNAG_03771 | 17591.8 | 0.85273 | 0.075633 | 11.27453 | 1.75E-29 | 1.21E-27 |
| CNAG_03772 | 10342.69 | -0.44615 | 0.09375 | -4.75897 | 1.95E-06 | 1.63E-05 |
| CNAG_07965 | 15600.03 | -0.54944 | 0.059812 | -9.18608 | 4.07E-20 | 1.50E-18 |
| CNAG_03776 | 769.4586 | -0.28547 | 0.092667 | -3.08058 | 0.002066 | 0.008844 |
| CNAG_03777 | 370.284 | -0.29614 | 0.123443 | -2.399 | 0.01644 | 0.05077 |
| CNAG_07481 | 674.5656 | 0.211585 | 0.106152 | 1.993231 | 0.046236 | 0.117038 |
| CNAG_03780 | 45593.19 | -0.19803 | 0.086831 | -2.28059 | 0.022573 | 0.065681 |
| CNAG_03781 | 1567.282 | -0.18217 | 0.081438 | -2.23695 | 0.02529 | 0.072426 |
| CNAG_03782 | 24.86599 | 1.012364 | 0.453884 | 2.230446 | 0.025718 | 0.07341 |
| CNAG_03785 | 2662.209 | 0.345125 | 0.066133 | 5.218665 | 1.80E-07 | 1.76E-06 |
| CNAG_03786 | 779.7361 | 0.319038 | 0.091927 | 3.470577 | 0.000519 | 0.0027 |
| CNAG_03788 | 687.7813 | -0.24606 | 0.097463 | -2.52464 | 0.011582 | 0.038298 |
| CNAG_03789 | 2432.117 | -0.63008 | 0.071222 | -8.84663 | 9.02E-19 | 2.86E-17 |
| CNAG_03794 | 1089.516 | 0.73261 | 0.095198 | 7.69568 | 1.41E-14 | 3.13E-13 |
| CNAG_03805 | 2623.038 | 0.300821 | 0.070041 | 4.29492 | 1.75E-05 | 0.000126 |
| CNAG_03806 | 930.6822 | 0.225441 | 0.084446 | 2.669642 | 0.007593 | 0.026911 |
| CNAG_03811 | 1358.567 | -0.17059 | 0.083001 | -2.05528 | 0.039852 | 0.104086 |
| CNAG_03812 | 4068.274 | -0.13243 | 0.06574 | -2.0144 | 0.043968 | 0.112759 |
| CNAG_03816 | 2478.317 | -0.13709 | 0.068858 | -1.99084 | 0.046498 | 0.117472 |
| CNAG_03817 | 2348.592 | -0.48999 | 0.079335 | -6.17622 | 6.57E-10 | 8.93E-09 |
| CNAG_03819 | 8314.676 | -0.99748 | 0.07627 | -13.0784 | 4.38E-39 | 5.79E-37 |
| CNAG_03820 | 4262.519 | -0.45817 | 0.089033 | -5.14611 | 2.66E-07 | 2.55E-06 |
| CNAG_03822 | 1138.594 | -0.1667 | 0.079086 | -2.10777 | 0.035051 | 0.093691 |
| CNAG_03824 | 3990.487 | 0.864979 | 0.073451 | 11.77625 | 5.17E-32 | 4.12E-30 |
| CNAG_03832 | 2213.725 | 0.308546 | 0.072041 | 4.282898 | 1.84E-05 | 0.000132 |
| CNAG_03837 | 292.1742 | 0.314903 | 0.137457 | 2.290912 | 0.021969 | 0.064295 |
| CNAG_03839 | 2086.017 | -0.26454 | 0.089875 | -2.94346 | 0.003246 | 0.013104 |
| CNAG_03840 | 1335.133 | -0.17368 | 0.077775 | -2.23314 | 0.02554 | 0.073008 |
| CNAG_03844 | 620.2855 | 0.361203 | 0.099583 | 3.627171 | 0.000287 | 0.00159 |

| | | | | | | |
|---|---|---|---|---|---|---|
| CNAG_03847 | 2223.398 | 0.377316 | 0.074122 | 5.090492 | 3.57E-07 | 3.34E-06 |
| CNAG_03849 | 1210.465 | -0.27317 | 0.125908 | -2.16964 | 0.030034 | 0.083145 |
| CNAG_03856 | 7412.917 | 0.155485 | 0.06185 | 2.513915 | 0.01194 | 0.039216 |
| CNAG_03857 | 3044.118 | 0.657157 | 0.06685 | 9.830326 | 8.33E-23 | 3.83E-21 |
| CNAG_03858 | 2917.452 | 0.223348 | 0.065654 | 3.401915 | 0.000669 | 0.00336 |
| CNAG_03861 | 12358.27 | 0.391206 | 0.064411 | 6.073603 | 1.25E-09 | 1.64E-08 |
| CNAG_03864 | 2099.176 | 0.200032 | 0.075502 | 2.649365 | 0.008064 | 0.02849 |
| CNAG_03865 | 2106.21 | 0.525276 | 0.078828 | 6.663558 | 2.67E-11 | 4.30E-10 |
| CNAG_07966 | 234.5567 | -0.28623 | 0.145656 | -1.96514 | 0.049398 | 0.1234 |
| CNAG_03872 | 1426.128 | 0.225958 | 0.075896 | 2.977186 | 0.002909 | 0.011936 |
| CNAG_03873 | 4409.615 | 0.664223 | 0.097357 | 6.822547 | 8.94E-12 | 1.53E-10 |
| CNAG_03874 | 732.5458 | -0.18982 | 0.092362 | -2.05513 | 0.039867 | 0.104088 |
| CNAG_03875 | 1776.339 | 0.15213 | 0.073257 | 2.076653 | 0.037834 | 0.099579 |
| CNAG_03876 | 3186.111 | 0.147583 | 0.070176 | 2.103039 | 0.035462 | 0.094543 |
| CNAG_03878 | 1535.734 | -0.38775 | 0.076486 | -5.0696 | 3.99E-07 | 3.71E-06 |
| CNAG_07482 | 1013.667 | 0.237412 | 0.09863 | 2.407093 | 0.01608 | 0.049837 |
| CNAG_07483 | 1253.032 | 0.25992 | 0.096133 | 2.70376 | 0.006856 | 0.024668 |
| CNAG_03881 | 1310.787 | 1.028894 | 0.089005 | 11.55991 | 6.58E-31 | 4.98E-29 |
| CNAG_03882 | 905.2851 | 0.259041 | 0.084934 | 3.049918 | 0.002289 | 0.009656 |
| CNAG_03890 | 1809.047 | -0.18731 | 0.074313 | -2.52057 | 0.011717 | 0.038646 |
| CNAG_03891 | 23498.27 | 0.864233 | 0.075245 | 11.48551 | 1.56E-30 | 1.14E-28 |
| CNAG_03892 | 13132.3 | 0.991397 | 0.063603 | 15.58723 | 8.89E-55 | 2.57E-52 |
| CNAG_03893 | 726.5878 | 0.423286 | 0.092129 | 4.594504 | 4.34E-06 | 3.47E-05 |
| CNAG_03896 | 286.2776 | -0.31816 | 0.127732 | -2.49082 | 0.012745 | 0.041407 |
| CNAG_12183 | 75.13418 | 0.671566 | 0.233912 | 2.871015 | 0.004092 | 0.015981 |
| CNAG_03897 | 1334.79 | -0.49801 | 0.087613 | -5.68421 | 1.31E-08 | 1.52E-07 |
| CNAG_03899 | 3477.703 | -0.44658 | 0.063712 | -7.00934 | 2.39E-12 | 4.32E-11 |
| CNAG_03900 | 1299.777 | 0.1803 | 0.086015 | 2.096148 | 0.036069 | 0.095873 |
| CNAG_03901 | 749.5176 | 0.420775 | 0.094503 | 4.452518 | 8.49E-06 | 6.48E-05 |
| CNAG_03902 | 3214.907 | 0.170186 | 0.06445 | 2.640583 | 0.008276 | 0.029039 |
| CNAG_03904 | 2765.176 | -0.2769 | 0.071957 | -3.84815 | 0.000119 | 0.000719 |
| CNAG_03909 | 470.5902 | 0.480626 | 0.113339 | 4.240617 | 2.23E-05 | 0.000156 |
| CNAG_03910 | 2050.421 | 1.128724 | 0.243556 | 4.634351 | 3.58E-06 | 2.90E-05 |
| CNAG_03911 | 144.0035 | -0.57732 | 0.178421 | -3.23572 | 0.001213 | 0.005603 |
| CNAG_03912 | 318.7043 | -0.40525 | 0.123499 | -3.28138 | 0.001033 | 0.004892 |
| CNAG_03915 | 377.9713 | -0.73851 | 0.116136 | -6.359 | 2.03E-10 | 2.96E-09 |
| CNAG_03916 | 14646.56 | 0.579307 | 0.065374 | 8.861448 | 7.90E-19 | 2.54E-17 |
| CNAG_03917 | 1801.355 | -0.18748 | 0.075943 | -2.46873 | 0.013559 | 0.043439 |
| CNAG_03920 | 7099.025 | -0.2809 | 0.061932 | -4.53558 | 5.74E-06 | 4.49E-05 |
| CNAG_03921 | 714.8555 | -0.72656 | 0.105514 | -6.88593 | 5.74E-12 | 9.96E-11 |
| CNAG_03926 | 3776.781 | -0.29599 | 0.067577 | -4.38011 | 1.19E-05 | 8.86E-05 |
| CNAG_03929 | 769.7246 | 0.381856 | 0.088106 | 4.334024 | 1.46E-05 | 0.000107 |
| CNAG_03935 | 2927.916 | 0.413123 | 0.065605 | 6.297126 | 3.03E-10 | 4.34E-09 |
| CNAG_03937 | 1100.464 | 0.38356 | 0.101203 | 3.789988 | 0.000151 | 0.000887 |
| CNAG_03940 | 3536.671 | -0.21615 | 0.068454 | -3.15759 | 0.001591 | 0.007074 |
| CNAG_03941 | 3787.776 | -0.17157 | 0.070403 | -2.43693 | 0.014813 | 0.046744 |
| CNAG_03944 | 8080.587 | -0.35794 | 0.062633 | -5.71485 | 1.10E-08 | 1.28E-07 |
| CNAG_03946 | 1110.741 | 0.365304 | 0.095543 | 3.82346 | 0.000132 | 0.000786 |

| | | | | | | |
|---|---|---|---|---|---|---|
| CNAG_07486 | 313.9964 | -0.27267 | 0.135627 | -2.0104 | 0.044388 | 0.113613 |
| CNAG_03948 | 4663.777 | -0.18006 | 0.077893 | -2.31168 | 0.020795 | 0.061542 |
| CNAG_03949 | 1212.055 | -0.21306 | 0.077456 | -2.75071 | 0.005947 | 0.021764 |
| CNAG_03950 | 5031.526 | -0.20797 | 0.086761 | -2.39704 | 0.016528 | 0.050962 |
| CNAG_03951 | 2001.498 | -0.21758 | 0.071019 | -3.06361 | 0.002187 | 0.00927 |
| CNAG_03952 | 2028.987 | -0.1688 | 0.080088 | -2.10768 | 0.035059 | 0.093691 |
| CNAG_03954 | 861.0056 | -0.62881 | 0.099124 | -6.3436 | 2.24E-10 | 3.26E-09 |
| CNAG_03957 | 1928.837 | -0.2343 | 0.082497 | -2.8401 | 0.00451 | 0.017278 |
| CNAG_03958 | 1204.863 | 0.251464 | 0.092947 | 2.705465 | 0.006821 | 0.024553 |
| CNAG_03959 | 3808.838 | 0.172415 | 0.068514 | 2.516485 | 0.011853 | 0.038998 |
| CNAG_03964 | 1293.078 | -0.78294 | 0.079748 | -9.81761 | 9.46E-23 | 4.32E-21 |
| CNAG_03967 | 2719.355 | 0.239615 | 0.067945 | 3.52659 | 0.000421 | 0.002236 |
| CNAG_03968 | 3705.262 | -0.18282 | 0.063983 | -2.85738 | 0.004272 | 0.016511 |
| CNAG_07487 | 17159.41 | 0.166272 | 0.076506 | 2.173311 | 0.029757 | 0.082642 |
| CNAG_03974 | 1226.37 | 0.541067 | 0.087004 | 6.218897 | 5.01E-10 | 6.90E-09 |
| CNAG_03975 | 3228.631 | -0.57435 | 0.082075 | -6.99789 | 2.60E-12 | 4.64E-11 |
| CNAG_03976 | 1174.509 | -0.2538 | 0.105412 | -2.40771 | 0.016053 | 0.049793 |
| CNAG_03978 | 891.7131 | 0.245575 | 0.084817 | 2.895355 | 0.003787 | 0.014988 |
| CNAG_03985 | 2602.199 | 0.297393 | 0.069379 | 4.286513 | 1.81E-05 | 0.000131 |
| CNAG_03987 | 4781.514 | 0.26988 | 0.06427 | 4.199177 | 2.68E-05 | 0.000185 |
| CNAG_12196 | 62.48028 | 0.663213 | 0.278399 | 2.382241 | 0.017208 | 0.052703 |
| CNAG_03991 | 1189.065 | 0.449317 | 0.081123 | 5.538693 | 3.05E-08 | 3.33E-07 |
| CNAG_03992 | 585.8678 | 0.242237 | 0.098077 | 2.469856 | 0.013517 | 0.043356 |
| CNAG_03994 | 919.8753 | 0.381361 | 0.104544 | 3.647851 | 0.000264 | 0.001481 |
| CNAG_03997 | 100.2087 | 0.426888 | 0.201801 | 2.115384 | 0.034397 | 0.092246 |
| CNAG_04001 | 3267.217 | -0.14825 | 0.075575 | -1.96162 | 0.049807 | 0.124224 |
| CNAG_04004 | 97320.54 | -0.24133 | 0.070683 | -3.41424 | 0.00064 | 0.003231 |
| CNAG_04005 | 3047.628 | -0.22061 | 0.071179 | -3.09934 | 0.00194 | 0.008399 |
| CNAG_04010 | 3889.719 | -0.14412 | 0.062206 | -2.31684 | 0.020513 | 0.060864 |
| CNAG_04014 | 4185.401 | -0.14613 | 0.067051 | -2.17946 | 0.029297 | 0.081655 |
| CNAG_04018 | 1331.148 | 0.260015 | 0.094883 | 2.740375 | 0.006137 | 0.022411 |
| CNAG_07489 | 1033.5 | 0.248538 | 0.091335 | 2.721155 | 0.006505 | 0.023558 |
| CNAG_04021 | 28426.89 | -0.27104 | 0.072212 | -3.75345 | 0.000174 | 0.00101 |
| CNAG_04023 | 1806.917 | -0.34537 | 0.083185 | -4.1518 | 3.30E-05 | 0.000223 |
| CNAG_04027 | 915.3423 | 2.668334 | 0.133734 | 19.95259 | 1.42E-88 | 1.59E-85 |
| CNAG_04030 | 753.844 | 0.259721 | 0.103404 | 2.511719 | 0.012014 | 0.039395 |
| CNAG_04032 | 6588.442 | 0.367678 | 0.058926 | 6.23964 | 4.39E-10 | 6.11E-09 |
| CNAG_04033 | 2392.557 | 0.179906 | 0.067892 | 2.649878 | 0.008052 | 0.028459 |
| CNAG_04035 | 1040.089 | 0.246028 | 0.087065 | 2.825788 | 0.004716 | 0.017955 |
| CNAG_04037 | 1687.088 | -0.31236 | 0.077325 | -4.03956 | 5.36E-05 | 0.00035 |
| CNAG_04038 | 309.81 | 0.573937 | 0.131632 | 4.360166 | 1.30E-05 | 9.64E-05 |
| CNAG_04043 | 365.9154 | 0.756038 | 0.115997 | 6.517717 | 7.14E-11 | 1.10E-09 |
| CNAG_04044 | 1706.885 | -0.34511 | 0.08073 | -4.2749 | 1.91E-05 | 0.000137 |
| CNAG_04049 | 1639.928 | -0.30213 | 0.076392 | -3.95493 | 7.66E-05 | 0.000488 |
| CNAG_04051 | 2312.351 | -0.30155 | 0.100101 | -3.01243 | 0.002592 | 0.010775 |
| CNAG_04053 | 779.1944 | -0.35753 | 0.090287 | -3.95993 | 7.50E-05 | 0.000478 |
| CNAG_07491 | 809.9762 | 0.769215 | 0.107359 | 7.164878 | 7.79E-13 | 1.49E-11 |
| CNAG_04062 | 2345.018 | -0.5626 | 0.069276 | -8.1211 | 4.62E-16 | 1.15E-14 |

| | | | | | | |
|---|---|---|---|---|---|---|
| CNAG_04064 | 6695.034 | 0.397971 | 0.069606 | 5.717513 | 1.08E-08 | 1.26E-07 |
| CNAG_04066 | 4044.871 | 0.181425 | 0.069942 | 2.593948 | 0.009488 | 0.032476 |
| CNAG_04067 | 534.5013 | 0.677951 | 0.124086 | 5.46357 | 4.67E-08 | 4.96E-07 |
| CNAG_04068 | 16110.71 | -0.28912 | 0.092524 | -3.1248 | 0.001779 | 0.007788 |
| CNAG_04070 | 175.55 | 0.394393 | 0.186734 | 2.112055 | 0.034682 | 0.092881 |
| CNAG_04071 | 2281.342 | -0.17505 | 0.083698 | -2.09146 | 0.036487 | 0.096653 |
| CNAG_04072 | 1935.743 | -0.5854 | 0.072795 | -8.04173 | 8.86E-16 | 2.17E-14 |
| CNAG_07494 | 2767.6 | -0.43099 | 0.070259 | -6.13425 | 8.56E-10 | 1.16E-08 |
| CNAG_04076 | 946.4165 | 0.755389 | 0.095342 | 7.922981 | 2.32E-15 | 5.50E-14 |
| CNAG_04077 | 597.4135 | -0.25198 | 0.110363 | -2.28324 | 0.022417 | 0.065276 |
| CNAG_04082 | 7351.283 | -0.14558 | 0.066838 | -2.17809 | 0.029399 | 0.081904 |
| CNAG_12208 | 123.9626 | 0.739389 | 0.184209 | 4.013857 | 5.97E-05 | 0.000388 |
| CNAG_04090 | 3230.467 | 0.380047 | 0.066516 | 5.713597 | 1.11E-08 | 1.29E-07 |
| CNAG_04094 | 289.1225 | -0.87833 | 0.131739 | -6.66715 | 2.61E-11 | 4.22E-10 |
| CNAG_04096 | 41.59658 | -1.13563 | 0.309782 | -3.66591 | 0.000246 | 0.001388 |
| CNAG_04098 | 844.9568 | 1.236488 | 0.107398 | 11.51311 | 1.13E-30 | 8.34E-29 |
| CNAG_04099 | 431.0719 | 0.401212 | 0.108929 | 3.683239 | 0.00023 | 0.001304 |
| CNAG_07499 | 1911.161 | -0.15348 | 0.077444 | -1.98184 | 0.047498 | 0.119572 |
| CNAG_06963 | 11360.45 | 0.547769 | 0.07235 | 7.57113 | 3.70E-14 | 7.93E-13 |
| CNAG_12214 | 86.44692 | 0.96741 | 0.238388 | 4.058126 | 4.95E-05 | 0.000326 |
| CNAG_03083 | 254.0751 | 0.953703 | 0.13729 | 6.946613 | 3.74E-12 | 6.56E-11 |
| CNAG_03082 | 110.0644 | 1.104808 | 0.241155 | 4.581322 | 4.62E-06 | 3.67E-05 |
| CNAG_03081 | 3109.727 | 0.330667 | 0.067023 | 4.933651 | 8.07E-07 | 7.09E-06 |
| CNAG_12216 | 60.24921 | -0.82628 | 0.254768 | -3.24326 | 0.001182 | 0.005479 |
| CNAG_03079 | 889.5137 | 0.480338 | 0.089346 | 5.376168 | 7.61E-08 | 7.81E-07 |
| CNAG_03069 | 856.3378 | -0.17463 | 0.089033 | -1.96138 | 0.049835 | 0.12425 |
| CNAG_03068 | 343.8929 | -2.44858 | 0.13848 | -17.6818 | 5.79E-70 | 3.47E-67 |
| CNAG_07502 | 7344.71 | -0.33842 | 0.060091 | -5.63187 | 1.78E-08 | 2.00E-07 |
| CNAG_03064 | 1818.841 | -0.53609 | 0.074335 | -7.21183 | 5.52E-13 | 1.07E-11 |
| CNAG_03060 | 1297.152 | 0.633781 | 0.11743 | 5.397092 | 6.77E-08 | 7.04E-07 |
| CNAG_03059 | 888.7763 | 0.523066 | 0.087463 | 5.980423 | 2.23E-09 | 2.84E-08 |
| CNAG_03058 | 19398.56 | 0.899914 | 0.0851 | 10.57476 | 3.90E-26 | 2.27E-24 |
| CNAG_03057 | 301.2376 | -0.43693 | 0.148882 | -2.93475 | 0.003338 | 0.013456 |
| CNAG_03055 | 1460.934 | -0.36646 | 0.079365 | -4.61747 | 3.88E-06 | 3.13E-05 |
| CNAG_03053 | 30529.76 | -0.24513 | 0.086886 | -2.82133 | 0.004783 | 0.018148 |
| CNAG_03051 | 742.2131 | -0.23839 | 0.092676 | -2.57233 | 0.010102 | 0.034142 |
| CNAG_03048 | 1920.995 | 0.180463 | 0.073554 | 2.453478 | 0.014148 | 0.044993 |
| CNAG_12222 | 71.70462 | 0.861097 | 0.242003 | 3.558206 | 0.000373 | 0.002001 |
| CNAG_03046 | 823.6451 | 0.397051 | 0.095634 | 4.151785 | 3.30E-05 | 0.000223 |
| CNAG_03045 | 958.8969 | 0.297925 | 0.097065 | 3.069336 | 0.002145 | 0.009114 |
| CNAG_03044 | 385.5059 | 0.270674 | 0.122684 | 2.206275 | 0.027365 | 0.077447 |
| CNAG_03041 | 102.8582 | -0.45877 | 0.199236 | -2.30267 | 0.021298 | 0.062814 |
| CNAG_03040 | 3222.885 | 0.721749 | 0.072736 | 9.922851 | 3.31E-23 | 1.60E-21 |
| CNAG_03034 | 393.3073 | -0.34392 | 0.118814 | -2.89463 | 0.003796 | 0.015007 |
| CNAG_03024 | 1361.834 | 0.220438 | 0.110222 | 1.999942 | 0.045507 | 0.115792 |
| CNAG_03021 | 34.35458 | -1.10818 | 0.394593 | -2.80842 | 0.004979 | 0.018751 |
| CNAG_03020 | 746.6725 | -0.23652 | 0.088545 | -2.67117 | 0.007559 | 0.026813 |
| CNAG_03017 | 1434.653 | -0.17687 | 0.083255 | -2.1245 | 0.033629 | 0.090559 |

| | | | | | | |
|---|---|---|---|---|---|---|
| CNAG_03012 | 53787.08 | 0.169298 | 0.078283 | 2.162634 | 0.030569 | 0.084387 |
| CNAG_12229 | 44.35708 | 0.933307 | 0.301726 | 3.093224 | 0.00198 | 0.008546 |
| CNAG_03011 | 100.6007 | -0.60838 | 0.204895 | -2.96924 | 0.002985 | 0.012204 |
| CNAG_03009 | 674.3976 | -0.22917 | 0.093182 | -2.45935 | 0.013919 | 0.044372 |
| CNAG_03007 | 2351.133 | -0.80596 | 0.086408 | -9.32733 | 1.09E-20 | 4.19E-19 |
| CNAG_03006 | 123.2279 | -0.44017 | 0.209907 | -2.09696 | 0.035997 | 0.095746 |
| CNAG_03005 | 140.2646 | -0.59855 | 0.191433 | -3.12668 | 0.001768 | 0.007756 |
| CNAG_03004 | 1550.451 | 0.172176 | 0.080889 | 2.128549 | 0.033292 | 0.089961 |
| CNAG_03002 | 908.3608 | 0.521199 | 0.093437 | 5.578084 | 2.43E-08 | 2.68E-07 |
| CNAG_03001 | 2932.842 | -0.37101 | 0.06648 | -5.58077 | 2.39E-08 | 2.64E-07 |
| CNAG_03000 | 53342.13 | -0.28734 | 0.076674 | -3.74751 | 0.000179 | 0.001032 |
| CNAG_02994 | 10977.62 | -0.29354 | 0.068859 | -4.26285 | 2.02E-05 | 0.000142 |
| CNAG_02993 | 1198.604 | 0.211299 | 0.080528 | 2.623923 | 0.008692 | 0.03027 |
| CNAG_02989 | 1627.282 | 0.668088 | 0.074167 | 9.007839 | 2.10E-19 | 7.10E-18 |
| CNAG_02986 | 1538.365 | 0.694969 | 0.09512 | 7.306226 | 2.75E-13 | 5.54E-12 |
| CNAG_02983 | 1848.518 | -0.38379 | 0.073525 | -5.2199 | 1.79E-07 | 1.75E-06 |
| CNAG_02982 | 2209.84 | -0.67316 | 0.075688 | -8.89388 | 5.90E-19 | 1.91E-17 |
| CNAG_02978 | 229.1467 | 0.786054 | 0.1477 | 5.321964 | 1.03E-07 | 1.03E-06 |
| CNAG_02977 | 2373.228 | -0.19927 | 0.075466 | -2.64058 | 0.008277 | 0.029039 |
| CNAG_02975 | 1378.165 | 0.230427 | 0.079428 | 2.901098 | 0.003719 | 0.014761 |
| CNAG_02973 | 976.6564 | -0.21088 | 0.087852 | -2.40041 | 0.016377 | 0.050596 |
| CNAG_07506 | 1808.011 | -0.27265 | 0.083929 | -3.24864 | 0.00116 | 0.005391 |
| CNAG_07507 | 868.9187 | -0.24055 | 0.09986 | -2.40884 | 0.016003 | 0.049717 |
| CNAG_02969 | 679.8529 | 0.597766 | 0.103231 | 5.790587 | 7.01E-09 | 8.37E-08 |
| CNAG_02968 | 420.3352 | 0.734289 | 0.122454 | 5.99646 | 2.02E-09 | 2.58E-08 |
| CNAG_02967 | 1753.718 | 0.74947 | 0.090421 | 8.288695 | 1.14E-16 | 3.03E-15 |
| CNAG_02961 | 2178.288 | 0.249783 | 0.088673 | 2.816893 | 0.004849 | 0.018325 |
| CNAG_02960 | 404.0688 | -0.72005 | 0.150438 | -4.78633 | 1.70E-06 | 1.44E-05 |
| CNAG_02959 | 1039.974 | -0.47545 | 0.111342 | -4.27018 | 1.95E-05 | 0.000139 |
| CNAG_02958 | 623.3195 | -0.30444 | 0.112617 | -2.70334 | 0.006865 | 0.024687 |
| CNAG_07509 | 2137.696 | 0.179646 | 0.083936 | 2.140275 | 0.032333 | 0.087979 |
| CNAG_07510 | 4040.496 | -0.20968 | 0.070478 | -2.97509 | 0.002929 | 0.012007 |
| CNAG_02955 | 659.7225 | 0.503867 | 0.112079 | 4.495662 | 6.94E-06 | 5.37E-05 |
| CNAG_02953 | 2143.694 | 0.49588 | 0.100807 | 4.919121 | 8.69E-07 | 7.61E-06 |
| CNAG_02952 | 1852.938 | 0.456471 | 0.075633 | 6.035327 | 1.59E-09 | 2.06E-08 |
| CNAG_02951 | 793.6218 | -0.22032 | 0.094074 | -2.34198 | 0.019182 | 0.05753 |
| CNAG_02948 | 9538.479 | -0.29217 | 0.077469 | -3.77149 | 0.000162 | 0.000946 |
| CNAG_07511 | 5248.76 | -0.25659 | 0.062493 | -4.10584 | 4.03E-05 | 0.000269 |
| CNAG_07512 | 610.1153 | 0.95452 | 0.115248 | 8.282301 | 1.21E-16 | 3.17E-15 |
| CNAG_02944 | 1309.696 | 1.727391 | 0.106522 | 16.21633 | 3.87E-59 | 1.26E-56 |
| CNAG_02943 | 19957.68 | 0.777394 | 0.065918 | 11.79332 | 4.23E-32 | 3.40E-30 |
| CNAG_02942 | 2097.371 | 0.878572 | 0.105312 | 8.342596 | 7.27E-17 | 1.94E-15 |
| CNAG_02938 | 8978.003 | 0.29043 | 0.066132 | 4.391673 | 1.12E-05 | 8.44E-05 |
| CNAG_02937 | 3743.041 | -0.14094 | 0.062678 | -2.24858 | 0.024539 | 0.070484 |
| CNAG_02935 | 1261.718 | 0.226015 | 0.081575 | 2.770641 | 0.005595 | 0.020643 |
| CNAG_02933 | 700.1752 | -0.53254 | 0.128097 | -4.1573 | 3.22E-05 | 0.000219 |
| CNAG_02928 | 53254.89 | -0.27125 | 0.074905 | -3.62121 | 0.000293 | 0.001617 |
| CNAG_02927 | 1131.412 | -0.19991 | 0.08096 | -2.46928 | 0.013538 | 0.043402 |

| | | | | | | |
|---|---|---|---|---|---|---|
| CNAG_02925 | 5170.05 | -0.34389 | 0.100503 | -3.4217 | 0.000622 | 0.003162 |
| CNAG_12238 | 121.4433 | 0.528872 | 0.190989 | 2.769125 | 0.005621 | 0.02072 |
| CNAG_02918 | 10056.93 | -0.68543 | 0.090102 | -7.60726 | 2.80E-14 | 6.07E-13 |
| CNAG_02914 | 1153.026 | -0.22063 | 0.087578 | -2.51927 | 0.01176 | 0.038747 |
| CNAG_07513 | 1317.866 | -0.3071 | 0.098069 | -3.13147 | 0.001739 | 0.007651 |
| CNAG_02907 | 1067.163 | -0.26282 | 0.084231 | -3.12018 | 0.001807 | 0.007893 |
| CNAG_02906 | 1285.618 | -0.17605 | 0.080639 | -2.18321 | 0.029021 | 0.081087 |
| CNAG_02905 | 2204.39 | 0.867928 | 0.071259 | 12.17999 | 3.97E-34 | 3.61E-32 |
| CNAG_02903 | 2107.816 | 1.041823 | 0.085019 | 12.25403 | 1.60E-34 | 1.50E-32 |
| CNAG_02901 | 2611.215 | -0.16886 | 0.070824 | -2.38429 | 0.017112 | 0.052459 |
| CNAG_02896 | 4051.794 | -1.09357 | 0.079321 | -13.7868 | 3.06E-43 | 4.60E-41 |
| CNAG_02895 | 1711.411 | 0.193412 | 0.074319 | 2.602457 | 0.009256 | 0.031817 |
| CNAG_02894 | 628.9958 | -0.30929 | 0.097404 | -3.17531 | 0.001497 | 0.006717 |
| CNAG_02892 | 491.3511 | 0.477304 | 0.108577 | 4.395982 | 1.10E-05 | 8.28E-05 |
| CNAG_02891 | 2734.09 | 0.272945 | 0.073325 | 3.722391 | 0.000197 | 0.00113 |
| CNAG_02890 | 1988.077 | -0.22259 | 0.086084 | -2.58568 | 0.009719 | 0.033091 |
| CNAG_02889 | 90.63 | 0.513544 | 0.217507 | 2.361044 | 0.018224 | 0.055123 |
| CNAG_07514 | 2744.256 | -0.30125 | 0.081371 | -3.70216 | 0.000214 | 0.001215 |
| CNAG_02883 | 863.9885 | 0.261996 | 0.09456 | 2.770684 | 0.005594 | 0.020643 |
| CNAG_02877 | 1595.014 | 0.856439 | 0.107863 | 7.940058 | 2.02E-15 | 4.85E-14 |
| CNAG_02875 | 1227.912 | -0.52332 | 0.082948 | -6.30899 | 2.81E-10 | 4.04E-09 |
| CNAG_02873 | 639.4204 | 0.285054 | 0.099315 | 2.870198 | 0.004102 | 0.015993 |
| CNAG_02870 | 2595.684 | -0.42865 | 0.073836 | -5.80543 | 6.42E-09 | 7.70E-08 |
| CNAG_12248 | 140.4698 | -0.38445 | 0.183621 | -2.09374 | 0.036283 | 0.096246 |
| CNAG_02867 | 1497.7 | -0.34009 | 0.079586 | -4.27323 | 1.93E-05 | 0.000137 |
| CNAG_12249 | 29.10372 | 0.91138 | 0.378507 | 2.407831 | 0.016048 | 0.049793 |
| CNAG_02865 | 166.4979 | 0.497521 | 0.167665 | 2.967357 | 0.003004 | 0.012254 |
| CNAG_02864 | 665.8394 | 1.184091 | 0.110266 | 10.73847 | 6.71E-27 | 4.23E-25 |
| CNAG_12250 | 1324.977 | 0.308833 | 0.083602 | 3.694098 | 0.000221 | 0.001253 |
| CNAG_02855 | 1669.761 | -0.31404 | 0.087518 | -3.58828 | 0.000333 | 0.00181 |
| CNAG_07515 | 154.5172 | 0.456855 | 0.218023 | 2.09544 | 0.036132 | 0.096007 |
| CNAG_02853 | 5659.685 | -0.40567 | 0.068147 | -5.95285 | 2.64E-09 | 3.34E-08 |
| CNAG_02852 | 781.7599 | 0.55658 | 0.088781 | 6.269136 | 3.63E-10 | 5.11E-09 |
| CNAG_12257 | 83.42431 | -0.52539 | 0.223524 | -2.35048 | 0.018749 | 0.056385 |
| CNAG_02848 | 275.0223 | -0.38728 | 0.132997 | -2.91191 | 0.003592 | 0.014354 |
| CNAG_02846 | 391.7429 | 0.387826 | 0.119295 | 3.250969 | 0.00115 | 0.005353 |
| CNAG_02845 | 1804.597 | -0.31674 | 0.07868 | -4.02565 | 5.68E-05 | 0.00037 |
| CNAG_12259 | 66.61398 | 1.082382 | 0.262735 | 4.119676 | 3.79E-05 | 0.000254 |
| CNAG_02843 | 6126.195 | 0.481071 | 0.07069 | 6.805364 | 1.01E-11 | 1.71E-10 |
| CNAG_02841 | 1056.314 | -0.21081 | 0.080248 | -2.62704 | 0.008613 | 0.030007 |
| CNAG_02840 | 508.7701 | -0.56364 | 0.116878 | -4.82241 | 1.42E-06 | 1.21E-05 |
| CNAG_02838 | 1378.483 | -0.26257 | 0.083343 | -3.15047 | 0.00163 | 0.007232 |
| CNAG_07517 | 901.7415 | 0.32607 | 0.106432 | 3.063649 | 0.002187 | 0.00927 |
| CNAG_07518 | 590.618 | -0.28348 | 0.095499 | -2.96842 | 0.002993 | 0.012224 |
| CNAG_07519 | 681.5623 | -0.73072 | 0.101668 | -7.18732 | 6.61E-13 | 1.28E-11 |
| CNAG_12261 | 35.95173 | 0.77385 | 0.326032 | 2.373543 | 0.017618 | 0.053729 |
| CNAG_02834 | 793.3931 | 0.586138 | 0.091423 | 6.41125 | 1.44E-10 | 2.14E-09 |
| CNAG_02833 | 2537.199 | 0.236889 | 0.07383 | 3.208577 | 0.001334 | 0.006088 |

| | | | | | | |
|---|---|---|---|---|---|---|
| CNAG_02830 | 5048.459 | -0.84837 | 0.07948 | -10.674 | 1.35E-26 | 8.21E-25 |
| CNAG_02829 | 1300.954 | -0.19209 | 0.080102 | -2.39804 | 0.016483 | 0.050843 |
| CNAG_02826 | 644.6544 | -0.23571 | 0.095734 | -2.46214 | 0.013811 | 0.044121 |
| CNAG_02825 | 5341.608 | -0.17289 | 0.064394 | -2.68488 | 0.007256 | 0.025867 |
| CNAG_07521 | 312.0581 | 0.632645 | 0.123354 | 5.128701 | 2.92E-07 | 2.78E-06 |
| CNAG_02822 | 4830.221 | -0.34276 | 0.083387 | -4.1105 | 3.95E-05 | 0.000264 |
| CNAG_02821 | 390.2564 | -0.25819 | 0.113881 | -2.2672 | 0.023378 | 0.067635 |
| CNAG_02819 | 763.6513 | 0.366166 | 0.09005 | 4.066231 | 4.78E-05 | 0.000316 |
| CNAG_02818 | 1612.381 | 0.279995 | 0.090917 | 3.079662 | 0.002072 | 0.008867 |
| CNAG_02815 | 4826.023 | 1.178814 | 0.07989 | 14.75544 | 2.84E-49 | 5.40E-47 |
| CNAG_02812 | 4180.661 | 0.209544 | 0.074931 | 2.796481 | 0.005166 | 0.019309 |
| CNAG_02810 | 9721.295 | -0.28542 | 0.07503 | -3.80407 | 0.000142 | 0.000843 |
| CNAG_02809 | 1715.239 | -0.51965 | 0.079194 | -6.56175 | 5.32E-11 | 8.22E-10 |
| CNAG_07522 | 2229.698 | 0.507439 | 0.075505 | 6.720626 | 1.81E-11 | 3.00E-10 |
| CNAG_02804 | 210.1616 | -0.40956 | 0.144931 | -2.82592 | 0.004715 | 0.017955 |
| CNAG_02802 | 209.4344 | 0.311851 | 0.143019 | 2.180491 | 0.029221 | 0.08156 |
| CNAG_02801 | 8599.758 | 0.188695 | 0.074106 | 2.546281 | 0.010888 | 0.036331 |
| CNAG_02795 | 485.925 | -0.48062 | 0.142833 | -3.36488 | 0.000766 | 0.00377 |
| CNAG_02792 | 1144.949 | -0.27893 | 0.087764 | -3.17817 | 0.001482 | 0.006666 |
| CNAG_02791 | 687.8071 | 0.378806 | 0.104425 | 3.627539 | 0.000286 | 0.001589 |
| CNAG_02790 | 1855.07 | 0.237734 | 0.081704 | 2.909686 | 0.003618 | 0.014427 |
| CNAG_02783 | 1165.656 | 0.736411 | 0.081535 | 9.03184 | 1.69E-19 | 5.80E-18 |
| CNAG_02782 | 1348.789 | -0.18005 | 0.078517 | -2.29311 | 0.021842 | 0.064032 |
| CNAG_02777 | 12358.24 | 2.579655 | 0.184575 | 13.97623 | 2.18E-44 | 3.62E-42 |
| CNAG_12269 | 192.7133 | 0.361174 | 0.153595 | 2.35147 | 0.018699 | 0.056279 |
| CNAG_12271 | 24.17472 | 1.140298 | 0.447836 | 2.546241 | 0.010889 | 0.036331 |
| CNAG_02775 | 3048.512 | 0.84803 | 0.078331 | 10.8262 | 2.59E-27 | 1.68E-25 |
| CNAG_02773 | 2939.181 | -0.49722 | 0.073326 | -6.78088 | 1.19E-11 | 2.02E-10 |
| CNAG_02771 | 991.2768 | -0.22646 | 0.111122 | -2.03795 | 0.041555 | 0.107837 |
| CNAG_02768 | 163.9349 | 1.083247 | 0.177763 | 6.09377 | 1.10E-09 | 1.46E-08 |
| CNAG_02765 | 32.49517 | -0.79733 | 0.351548 | -2.26806 | 0.023325 | 0.067544 |
| CNAG_02763 | 4205.357 | -0.14139 | 0.064071 | -2.20681 | 0.027327 | 0.077381 |
| CNAG_02758 | 216.0153 | -0.86339 | 0.154549 | -5.58648 | 2.32E-08 | 2.57E-07 |
| CNAG_02757 | 696.4869 | 0.527825 | 0.096885 | 5.447968 | 5.09E-08 | 5.38E-07 |
| CNAG_02753 | 4926.643 | 0.354819 | 0.075699 | 4.687256 | 2.77E-06 | 2.28E-05 |
| CNAG_02752 | 1982.097 | -0.2745 | 0.083205 | -3.2991 | 0.00097 | 0.004621 |
| CNAG_02751 | 2358.949 | 1.24405 | 0.091525 | 13.59245 | 4.44E-42 | 6.42E-40 |
| CNAG_02744 | 699.989 | -0.24488 | 0.10036 | -2.44001 | 0.014687 | 0.046457 |
| CNAG_02740 | 502.3593 | -0.38184 | 0.109068 | -3.50093 | 0.000464 | 0.002445 |
| CNAG_02736 | 6492.628 | -0.18074 | 0.062743 | -2.8806 | 0.003969 | 0.015597 |
| CNAG_02733 | 83.42826 | 0.701884 | 0.214435 | 3.273182 | 0.001063 | 0.005013 |
| CNAG_07524 | 1170.987 | -0.34971 | 0.082955 | -4.2156 | 2.49E-05 | 0.000173 |
| CNAG_02728 | 540.639 | 0.4717 | 0.103299 | 4.566341 | 4.96E-06 | 3.91E-05 |
| CNAG_02726 | 4336.638 | -0.2035 | 0.068401 | -2.97505 | 0.002929 | 0.012007 |
| CNAG_02725 | 3149.911 | -0.25427 | 0.064756 | -3.92661 | 8.62E-05 | 0.00054 |
| CNAG_02721 | 2143.955 | -0.20965 | 0.103338 | -2.02883 | 0.042476 | 0.109617 |
| CNAG_02720 | 2705.102 | -0.49583 | 0.067747 | -7.31877 | 2.50E-13 | 5.10E-12 |
| CNAG_12278 | 29.67872 | -0.92538 | 0.395796 | -2.33801 | 0.019387 | 0.058078 |

| | | | | | | |
|---|---|---|---|---|---|---|
| CNAG_02718 | 1278.595 | -0.34819 | 0.083359 | -4.17701 | 2.95E-05 | 0.000203 |
| CNAG_02715 | 956.8664 | 0.185622 | 0.087235 | 2.12785 | 0.03335 | 0.090033 |
| CNAG_02712 | 2429.706 | -0.27396 | 0.075571 | -3.62525 | 0.000289 | 0.001598 |
| CNAG_02710 | 8362.756 | -0.16097 | 0.060476 | -2.66163 | 0.007776 | 0.027515 |
| CNAG_02708 | 636.4037 | 0.238607 | 0.093785 | 2.54419 | 0.010953 | 0.036498 |
| CNAG_02705 | 179.9337 | -0.6436 | 0.181786 | -3.54044 | 0.000399 | 0.002131 |
| CNAG_02703 | 2006.917 | -0.28183 | 0.086264 | -3.26707 | 0.001087 | 0.005099 |
| CNAG_02699 | 1463.542 | -0.35672 | 0.09073 | -3.93163 | 8.44E-05 | 0.000532 |
| CNAG_02698 | 1417.241 | -0.20637 | 0.080111 | -2.57604 | 0.009994 | 0.033821 |
| CNAG_02697 | 848.0925 | 0.28575 | 0.094474 | 3.024633 | 0.002489 | 0.010417 |
| CNAG_02693 | 797.7102 | -0.58027 | 0.086965 | -6.6725 | 2.51E-11 | 4.10E-10 |
| CNAG_02692 | 3972.489 | 0.601417 | 0.084157 | 7.146407 | 8.91E-13 | 1.69E-11 |
| CNAG_02691 | 604.6408 | -0.42075 | 0.125009 | -3.36577 | 0.000763 | 0.003761 |
| CNAG_02690 | 1062.004 | 0.635777 | 0.084118 | 7.558121 | 4.09E-14 | 8.72E-13 |
| CNAG_02687 | 2610.357 | 0.210328 | 0.068615 | 3.065339 | 0.002174 | 0.009227 |
| CNAG_02686 | 3728.89 | -0.15233 | 0.065248 | -2.33464 | 0.019562 | 0.058471 |
| CNAG_02685 | 1089.73 | 1.02615 | 0.123795 | 8.289131 | 1.14E-16 | 3.03E-15 |
| CNAG_02682 | 1632.489 | 0.211906 | 0.101571 | 2.086271 | 0.036954 | 0.097627 |
| CNAG_02679 | 763.5832 | -0.19376 | 0.090628 | -2.13793 | 0.032522 | 0.088322 |
| CNAG_02678 | 568.0491 | 0.387637 | 0.101238 | 3.828977 | 0.000129 | 0.000773 |
| CNAG_02677 | 3952.004 | -0.14203 | 0.065719 | -2.1611 | 0.030687 | 0.084563 |
| CNAG_02675 | 3860.221 | -0.2783 | 0.095036 | -2.9284 | 0.003407 | 0.013692 |
| CNAG_02673 | 6197.522 | 0.656687 | 0.073247 | 8.965344 | 3.09E-19 | 1.03E-17 |
| CNAG_02672 | 3085.922 | -0.5239 | 0.069463 | -7.54215 | 4.62E-14 | 9.83E-13 |
| CNAG_02667 | 453.824 | 0.515393 | 0.10684 | 4.823974 | 1.41E-06 | 1.20E-05 |
| CNAG_02663 | 1195.626 | 0.706495 | 0.093895 | 7.524324 | 5.30E-14 | 1.12E-12 |
| CNAG_02662 | 1113.138 | 0.35742 | 0.084298 | 4.239951 | 2.24E-05 | 0.000157 |
| CNAG_12295 | 71.41533 | 1.195637 | 0.238266 | 5.018074 | 5.22E-07 | 4.79E-06 |
| CNAG_12296 | 34.80168 | 0.951973 | 0.340768 | 2.793609 | 0.005212 | 0.019453 |
| CNAG_02657 | 7702.972 | -0.29933 | 0.072407 | -4.13397 | 3.57E-05 | 0.00024 |
| CNAG_02655 | 2185.557 | 0.457 | 0.089355 | 5.114461 | 3.15E-07 | 2.98E-06 |
| CNAG_07536 | 971.1652 | -0.64122 | 0.098994 | -6.47741 | 9.33E-11 | 1.41E-09 |
| CNAG_07537 | 687.4279 | -0.22587 | 0.101152 | -2.23299 | 0.02555 | 0.073009 |
| CNAG_07538 | 1049.983 | 1.213079 | 0.093741 | 12.94072 | 2.65E-38 | 3.39E-36 |
| CNAG_07540 | 532.2419 | 0.971177 | 0.117624 | 8.256624 | 1.50E-16 | 3.89E-15 |
| CNAG_07541 | 1175.308 | -0.19868 | 0.084542 | -2.35002 | 0.018772 | 0.056432 |
| CNAG_07542 | 175.5068 | 0.448396 | 0.15717 | 2.852938 | 0.004332 | 0.016702 |
| CNAG_07544 | 54.50682 | -0.94786 | 0.288343 | -3.28726 | 0.001012 | 0.004805 |
| CNAG_07545 | 891.41 | -0.22961 | 0.094699 | -2.4246 | 0.015325 | 0.048075 |
| CNAG_07546 | 1326.666 | 0.18749 | 0.07854 | 2.387201 | 0.016977 | 0.052141 |
| CNAG_07547 | 930.9774 | 0.199077 | 0.098119 | 2.028939 | 0.042465 | 0.109617 |
| CNAG_07548 | 680.0482 | -0.26956 | 0.093274 | -2.88999 | 0.003853 | 0.0152 |
| CNAG_07549 | 342.6896 | 0.446336 | 0.136364 | 3.273125 | 0.001064 | 0.005013 |
| CNAG_07551 | 2476.24 | 0.47475 | 0.079589 | 5.965002 | 2.45E-09 | 3.10E-08 |
| CNAG_07557 | 1092.645 | 0.203931 | 0.102165 | 1.996098 | 0.045923 | 0.116479 |
| CNAG_07559 | 2932.935 | 0.879209 | 0.084507 | 10.40398 | 2.38E-25 | 1.34E-23 |
| CNAG_07561 | 54988.21 | 0.384186 | 0.069419 | 5.534325 | 3.12E-08 | 3.40E-07 |
| CNAG_07562 | 1461.472 | -0.28548 | 0.087085 | -3.27816 | 0.001045 | 0.004939 |

| | | | | | | |
|---|---|---|---|---|---|---|
| CNAG_07567 | 1287.427 | -0.48943 | 0.086634 | -5.64942 | 1.61E-08 | 1.84E-07 |
| CNAG_07572 | 1177.495 | -0.38373 | 0.083795 | -4.57931 | 4.67E-06 | 3.70E-05 |
| CNAG_02609 | 1522.678 | 0.159429 | 0.078489 | 2.03122 | 0.042233 | 0.10917 |
| CNAG_07573 | 905.2858 | 0.167471 | 0.084371 | 1.984946 | 0.04715 | 0.118812 |
| CNAG_02605 | 692.3926 | -0.1947 | 0.094941 | -2.05076 | 0.040291 | 0.104949 |
| CNAG_02595 | 375.4471 | -0.48823 | 0.129198 | -3.77894 | 0.000157 | 0.000922 |
| CNAG_02593 | 895.1216 | 0.4698 | 0.090236 | 5.206361 | 1.93E-07 | 1.87E-06 |
| CNAG_02591 | 213.7167 | 0.767592 | 0.150772 | 5.091094 | 3.56E-07 | 3.33E-06 |
| CNAG_02590 | 221.3735 | -0.33921 | 0.139316 | -2.43481 | 0.0149 | 0.047 |
| CNAG_02588 | 483.3827 | 0.288086 | 0.110358 | 2.61048 | 0.009042 | 0.03118 |
| CNAG_02587 | 801.4698 | -0.88676 | 0.09678 | -9.16263 | 5.06E-20 | 1.85E-18 |
| CNAG_02585 | 2794.314 | 0.433447 | 0.075494 | 5.741464 | 9.39E-09 | 1.11E-07 |
| CNAG_02583 | 558.5392 | -0.51627 | 0.100417 | -5.14123 | 2.73E-07 | 2.61E-06 |
| CNAG_02581 | 1943.746 | 0.198378 | 0.080068 | 2.477631 | 0.013226 | 0.04265 |
| CNAG_02580 | 1688.932 | -0.19652 | 0.07302 | -2.69127 | 0.007118 | 0.025423 |
| CNAG_02578 | 992.0442 | -0.19104 | 0.084838 | -2.25183 | 0.024333 | 0.069969 |
| CNAG_02577 | 3306.424 | 0.226025 | 0.074907 | 3.017418 | 0.002549 | 0.010628 |
| CNAG_02576 | 1086.862 | 0.370075 | 0.087762 | 4.216826 | 2.48E-05 | 0.000172 |
| CNAG_02573 | 481.5636 | -0.34291 | 0.10664 | -3.21561 | 0.001302 | 0.005958 |
| CNAG_02568 | 3728.768 | 0.186761 | 0.068278 | 2.735288 | 0.006233 | 0.022686 |
| CNAG_02564 | 979.5262 | -0.31823 | 0.083161 | -3.82673 | 0.00013 | 0.000779 |
| CNAG_12315 | 63.92682 | 0.966102 | 0.266424 | 3.626189 | 0.000288 | 0.001594 |
| CNAG_06880 | 1010.998 | 0.243049 | 0.090722 | 2.679044 | 0.007383 | 0.026238 |
| CNAG_06881 | 1757.738 | 0.216571 | 0.078798 | 2.748428 | 0.005988 | 0.021899 |
| CNAG_06882 | 899.716 | -0.21338 | 0.088116 | -2.42157 | 0.015454 | 0.048376 |
| CNAG_07576 | 574.9583 | 0.576505 | 0.09924 | 5.809187 | 6.28E-09 | 7.54E-08 |
| CNAG_06884 | 899.1461 | 0.406591 | 0.1192 | 3.411008 | 0.000647 | 0.003265 |
| CNAG_07580 | 1235.278 | -0.23211 | 0.082239 | -2.82238 | 0.004767 | 0.018111 |
| CNAG_06889 | 1647.547 | -0.17414 | 0.080727 | -2.15712 | 0.030996 | 0.085235 |
| CNAG_06892 | 1364.027 | 0.175023 | 0.085896 | 2.037612 | 0.041589 | 0.107837 |
| CNAG_06893 | 1485.213 | 0.421914 | 0.081149 | 5.19923 | 2.00E-07 | 1.94E-06 |
| CNAG_06897 | 1146.162 | -0.23013 | 0.082889 | -2.77642 | 0.005496 | 0.020357 |
| CNAG_06899 | 5235.737 | -0.38454 | 0.072994 | -5.2681 | 1.38E-07 | 1.37E-06 |
| CNAG_06901 | 307.7398 | -0.40429 | 0.129009 | -3.13379 | 0.001726 | 0.007604 |
| CNAG_06902 | 1114.303 | -0.27005 | 0.088382 | -3.05544 | 0.002247 | 0.009506 |
| CNAG_06905 | 1769.345 | 0.21001 | 0.07942 | 2.644297 | 0.008186 | 0.028842 |
| CNAG_06906 | 10758.95 | -0.27705 | 0.066066 | -4.1936 | 2.75E-05 | 0.000189 |
| CNAG_06912 | 230.0502 | -0.28567 | 0.143877 | -1.98552 | 0.047086 | 0.118728 |
| CNAG_06916 | 914.6969 | -0.85542 | 0.091441 | -9.35486 | 8.37E-21 | 3.27E-19 |
| CNAG_06918 | 83.49238 | 0.592922 | 0.229153 | 2.587452 | 0.009669 | 0.032977 |
| CNAG_06919 | 8685.02 | -0.40314 | 0.071767 | -5.6174 | 1.94E-08 | 2.16E-07 |
| CNAG_06920 | 3953.243 | -0.22756 | 0.071383 | -3.18795 | 0.001433 | 0.006482 |
| CNAG_06921 | 668.0039 | -0.29569 | 0.113467 | -2.60592 | 0.009163 | 0.03157 |
| CNAG_06923 | 6632.694 | 0.627027 | 0.078173 | 8.021034 | 1.05E-15 | 2.56E-14 |
| CNAG_06924 | 1545.042 | 0.720561 | 0.09307 | 7.742101 | 9.78E-15 | 2.21E-13 |
| CNAG_06929 | 1749.845 | -0.28082 | 0.083526 | -3.36205 | 0.000774 | 0.003804 |
| CNAG_06931 | 148.3578 | -0.96055 | 0.168315 | -5.70683 | 1.15E-08 | 1.34E-07 |
| CNAG_07583 | 678.2891 | 0.223261 | 0.095577 | 2.335928 | 0.019495 | 0.058358 |

| | | | | | | |
|---|---|---|---|---|---|---|
| CNAG_06935 | 211.1937 | 0.50574 | 0.149225 | 3.389117 | 0.000701 | 0.003499 |
| CNAG_06936 | 360.3186 | 1.103107 | 0.118647 | 9.297396 | 1.44E-20 | 5.48E-19 |
| CNAG_04933 | 132.5097 | -0.3955 | 0.18681 | -2.1171 | 0.034251 | 0.091981 |
| CNAG_04938 | 1779.796 | 0.804734 | 0.090351 | 8.90671 | 5.26E-19 | 1.72E-17 |
| CNAG_07794 | 1710.201 | 0.208952 | 0.08797 | 2.375277 | 0.017536 | 0.053561 |
| CNAG_04943 | 1399.81 | 0.500331 | 0.077664 | 6.442261 | 1.18E-10 | 1.76E-09 |
| CNAG_04944 | 932.6575 | 0.427995 | 0.08399 | 5.095772 | 3.47E-07 | 3.27E-06 |
| CNAG_04949 | 1461.06 | 0.185074 | 0.085587 | 2.162417 | 0.030586 | 0.084404 |
| CNAG_04950 | 1895.153 | -0.26128 | 0.077399 | -3.37577 | 0.000736 | 0.003645 |
| CNAG_04953 | 10708.26 | 0.242249 | 0.061308 | 3.951316 | 7.77E-05 | 0.000493 |
| CNAG_04955 | 740.7077 | -0.23939 | 0.109609 | -2.184 | 0.028962 | 0.080983 |
| CNAG_04956 | 298.4401 | 0.332725 | 0.135085 | 2.463081 | 0.013775 | 0.044039 |
| CNAG_04958 | 2082.171 | 0.284582 | 0.091978 | 3.09401 | 0.001975 | 0.008528 |
| CNAG_04959 | 1131.308 | 0.277764 | 0.083308 | 3.334176 | 0.000856 | 0.004147 |
| CNAG_04960 | 161.015 | 0.358955 | 0.177182 | 2.025919 | 0.042773 | 0.110165 |
| CNAG_04961 | 2640.223 | -0.22829 | 0.07361 | -3.10129 | 0.001927 | 0.008349 |
| CNAG_04962 | 3992.073 | -0.2412 | 0.066563 | -3.62357 | 0.000291 | 0.001606 |
| CNAG_04963 | 4424.22 | 0.155398 | 0.078384 | 1.982525 | 0.047421 | 0.119416 |
| CNAG_07797 | 1333.802 | -0.19263 | 0.08482 | -2.27105 | 0.023144 | 0.067093 |
| CNAG_04968 | 511.0898 | 0.219045 | 0.102419 | 2.138704 | 0.03246 | 0.088263 |
| CNAG_04969 | 12240.71 | -0.26371 | 0.062673 | -4.20772 | 2.58E-05 | 0.000179 |
| CNAG_04970 | 1582.158 | 0.782122 | 0.090343 | 8.657269 | 4.83E-18 | 1.42E-16 |
| CNAG_12342 | 24.71968 | 0.906704 | 0.413563 | 2.192421 | 0.028349 | 0.079655 |
| CNAG_04976 | 10771.84 | -0.15914 | 0.07709 | -2.06436 | 0.038984 | 0.102022 |
| CNAG_04979 | 386.6205 | -0.37906 | 0.117201 | -3.23429 | 0.001219 | 0.005628 |
| CNAG_04980 | 1125.67 | -0.20522 | 0.082911 | -2.47521 | 0.013316 | 0.04287 |
| CNAG_04981 | 4314.743 | 1.10499 | 0.080112 | 13.79302 | 2.81E-43 | 4.30E-41 |
| CNAG_04983 | 1030.255 | -0.17944 | 0.090836 | -1.97544 | 0.048218 | 0.121038 |
| CNAG_04984 | 1961.876 | -0.23321 | 0.115047 | -2.02705 | 0.042657 | 0.109976 |
| CNAG_04985 | 14805.23 | -0.21693 | 0.075237 | -2.88323 | 0.003936 | 0.015475 |
| CNAG_04986 | 1160.86 | 0.198698 | 0.093956 | 2.114787 | 0.034448 | 0.092351 |
| CNAG_04987 | 506.0312 | 0.364563 | 0.11626 | 3.135752 | 0.001714 | 0.007558 |
| CNAG_04990 | 3116.71 | -0.16197 | 0.075312 | -2.15062 | 0.031507 | 0.086303 |
| CNAG_07801 | 2567.609 | 0.266241 | 0.073481 | 3.62329 | 0.000291 | 0.001607 |
| CNAG_04997 | 954.7278 | -0.32651 | 0.087851 | -3.71668 | 0.000202 | 0.001152 |
| CNAG_04999 | 345.2777 | 0.331038 | 0.120596 | 2.745029 | 0.006051 | 0.022116 |
| CNAG_05003 | 602.3496 | 0.304283 | 0.095056 | 3.201095 | 0.001369 | 0.006232 |
| CNAG_05005 | 601.5458 | 0.754883 | 0.138987 | 5.431333 | 5.59E-08 | 5.87E-07 |
| CNAG_05006 | 779.1432 | -0.30264 | 0.095625 | -3.16483 | 0.001552 | 0.006928 |
| CNAG_05008 | 262.2668 | 0.395781 | 0.141353 | 2.799947 | 0.005111 | 0.01914 |
| CNAG_05009 | 510.5186 | -0.31075 | 0.134714 | -2.30672 | 0.02107 | 0.062248 |
| CNAG_05022 | 36.78081 | 1.006033 | 0.335633 | 2.997418 | 0.002723 | 0.01129 |
| CNAG_05023 | 1130.735 | 0.179099 | 0.089654 | 1.997673 | 0.045752 | 0.116114 |
| CNAG_05026 | 406.6229 | -0.26275 | 0.111525 | -2.35599 | 0.018473 | 0.05577 |
| CNAG_05028 | 4300.713 | -0.14665 | 0.069792 | -2.10128 | 0.035616 | 0.094799 |
| CNAG_05029 | 888.7235 | 0.35149 | 0.086998 | 4.040229 | 5.34E-05 | 0.00035 |
| CNAG_05031 | 1955.264 | 1.119459 | 0.075742 | 14.78 | 1.97E-49 | 3.95E-47 |
| CNAG_05035 | 999.952 | 0.47161 | 0.083401 | 5.654733 | 1.56E-08 | 1.79E-07 |

| | | | | | | |
|---|---|---|---|---|---|---|
| CNAG_05038 | 387.7382 | 0.34283 | 0.118881 | 2.883814 | 0.003929 | 0.015454 |
| CNAG_05041 | 7307.114 | 0.302493 | 0.066094 | 4.57668 | 4.72E-06 | 3.73E-05 |
| CNAG_05042 | 1044.369 | 0.480203 | 0.090539 | 5.303803 | 1.13E-07 | 1.13E-06 |
| CNAG_05045 | 686.8902 | -0.20894 | 0.092774 | -2.25211 | 0.024315 | 0.069944 |
| CNAG_07804 | 3129.051 | 0.205293 | 0.071308 | 2.878953 | 0.00399 | 0.015655 |
| CNAG_07805 | 4404.804 | 0.291235 | 0.073099 | 3.984126 | 6.77E-05 | 0.000436 |
| CNAG_05049 | 538.5547 | -0.299 | 0.105373 | -2.83753 | 0.004546 | 0.017409 |
| CNAG_05050 | 3190.65 | 0.228671 | 0.07028 | 3.253701 | 0.001139 | 0.00531 |
| CNAG_07807 | 31560.86 | -0.17662 | 0.065535 | -2.69502 | 0.007038 | 0.025185 |
| CNAG_05055 | 709.963 | -0.1912 | 0.097085 | -1.96946 | 0.0489 | 0.122416 |
| CNAG_05057 | 905.9663 | -0.49695 | 0.090672 | -5.48074 | 4.24E-08 | 4.53E-07 |
| CNAG_05058 | 1411.315 | 0.179016 | 0.075754 | 2.363125 | 0.018122 | 0.054899 |
| CNAG_05059 | 16533.38 | 0.576143 | 0.084864 | 6.789011 | 1.13E-11 | 1.92E-10 |
| CNAG_05067 | 45.722 | 0.738967 | 0.29488 | 2.505995 | 0.012211 | 0.039938 |
| CNAG_05069 | 4628.986 | 0.510283 | 0.076012 | 6.71321 | 1.90E-11 | 3.15E-10 |
| CNAG_05070 | 9645.389 | -0.32041 | 0.065341 | -4.90369 | 9.41E-07 | 8.19E-06 |
| CNAG_07808 | 2328.736 | 0.183596 | 0.081318 | 2.257742 | 0.023962 | 0.069028 |
| CNAG_05076 | 972.9452 | -0.28951 | 0.083795 | -3.45495 | 0.00055 | 0.002841 |
| CNAG_05077 | 2165.163 | 0.479163 | 0.068214 | 7.024408 | 2.15E-12 | 3.89E-11 |
| CNAG_05079 | 90.41938 | -0.74796 | 0.249935 | -2.99263 | 0.002766 | 0.011432 |
| CNAG_05080 | 4038.661 | -0.16283 | 0.069219 | -2.35247 | 0.018649 | 0.056215 |
| CNAG_05085 | 1253.515 | -0.5667 | 0.0879 | -6.44715 | 1.14E-10 | 1.71E-09 |
| CNAG_05086 | 679.9745 | -0.40234 | 0.107949 | -3.72711 | 0.000194 | 0.00111 |
| CNAG_05087 | 38.00667 | -1.08644 | 0.335665 | -3.23668 | 0.001209 | 0.005594 |
| CNAG_05090 | 2079.57 | 0.216985 | 0.071143 | 3.049989 | 0.002288 | 0.009656 |
| CNAG_05095 | 2106.006 | 0.557242 | 0.090158 | 6.180745 | 6.38E-10 | 8.69E-09 |
| CNAG_05096 | 1651.993 | 0.599581 | 0.075555 | 7.935706 | 2.09E-15 | 5.01E-14 |
| CNAG_05097 | 1596.785 | 0.582263 | 0.079962 | 7.281793 | 3.29E-13 | 6.54E-12 |
| CNAG_05098 | 1158.271 | 0.189519 | 0.08675 | 2.18465 | 0.028915 | 0.080907 |
| CNAG_05101 | 3765.752 | -0.43944 | 0.06651 | -6.60721 | 3.92E-11 | 6.14E-10 |
| CNAG_05105 | 8984.687 | -0.1757 | 0.060593 | -2.89974 | 0.003735 | 0.014802 |
| CNAG_05106 | 337.8493 | 0.451889 | 0.124592 | 3.626952 | 0.000287 | 0.001591 |
| CNAG_05109 | 3455.164 | -0.23071 | 0.067536 | -3.41612 | 0.000635 | 0.003213 |
| CNAG_05111 | 3016.011 | -0.2345 | 0.06823 | -3.43682 | 0.000589 | 0.003022 |
| CNAG_12368 | 104.6363 | 1.010448 | 0.197084 | 5.126994 | 2.94E-07 | 2.80E-06 |
| CNAG_05118 | 1039.759 | -0.2194 | 0.083097 | -2.64031 | 0.008283 | 0.029039 |
| CNAG_05119 | 425.0046 | 0.71783 | 0.114035 | 6.294831 | 3.08E-10 | 4.39E-09 |
| CNAG_05120 | 2150.165 | -0.18339 | 0.079377 | -2.31032 | 0.02087 | 0.061718 |
| CNAG_05122 | 1916.819 | -0.21798 | 0.080776 | -2.69861 | 0.006963 | 0.02496 |
| CNAG_05125 | 2652.066 | -0.3915 | 0.082 | -4.77441 | 1.80E-06 | 1.52E-05 |
| CNAG_07813 | 2001.37 | -0.298 | 0.075152 | -3.96533 | 7.33E-05 | 0.000468 |
| CNAG_05132 | 10439.88 | 0.313847 | 0.08263 | 3.798233 | 0.000146 | 0.000861 |
| CNAG_07815 | 665.1246 | 0.215523 | 0.107132 | 2.011754 | 0.044246 | 0.113297 |
| CNAG_05134 | 1320.155 | 0.190163 | 0.076304 | 2.492172 | 0.012696 | 0.041285 |
| CNAG_05138 | 2610.326 | 0.822543 | 0.073821 | 11.14238 | 7.80E-29 | 5.34E-27 |
| CNAG_05139 | 1333.814 | 0.218846 | 0.095363 | 2.294872 | 0.02174 | 0.063783 |
| CNAG_05140 | 7996.965 | -0.49761 | 0.069558 | -7.15392 | 8.43E-13 | 1.61E-11 |
| CNAG_05145 | 1116.123 | -0.25812 | 0.085998 | -3.00145 | 0.002687 | 0.011148 |

| | | | | | | |
|---|---|---|---|---|---|---|
| CNAG_05147 | 132.2967 | -0.41471 | 0.192826 | -2.1507 | 0.0315 | 0.086303 |
| CNAG_05150 | 2780.9 | 0.30656 | 0.06748 | 4.542965 | 5.55E-06 | 4.35E-05 |
| CNAG_05151 | 1421.463 | -0.39196 | 0.083647 | -4.68586 | 2.79E-06 | 2.29E-05 |
| CNAG_05152 | 2216.838 | 0.276439 | 0.070355 | 3.929218 | 8.52E-05 | 0.000535 |
| CNAG_05153 | 1358.111 | 0.583145 | 0.087042 | 6.699598 | 2.09E-11 | 3.45E-10 |
| CNAG_05155 | 2626.422 | 0.537101 | 0.091394 | 5.876752 | 4.18E-09 | 5.18E-08 |
| CNAG_05156 | 1554.698 | 0.410592 | 0.076147 | 5.392115 | 6.96E-08 | 7.19E-07 |
| CNAG_05157 | 129.6411 | -0.69987 | 0.183549 | -3.81297 | 0.000137 | 0.000816 |
| CNAG_05158 | 555.116 | -0.2667 | 0.107834 | -2.4732 | 0.013391 | 0.043014 |
| CNAG_12377 | 137.5893 | 0.482398 | 0.19475 | 2.477018 | 0.013249 | 0.042706 |
| CNAG_05166 | 4477.525 | -0.34849 | 0.082898 | -4.20382 | 2.62E-05 | 0.000182 |
| CNAG_05167 | 4308.248 | 1.195562 | 0.078739 | 15.18396 | 4.52E-52 | 1.04E-49 |
| CNAG_05169 | 917.3416 | 0.405942 | 0.088455 | 4.589255 | 4.45E-06 | 3.55E-05 |
| CNAG_12381 | 33.90345 | 0.736731 | 0.334591 | 2.201888 | 0.027673 | 0.078184 |
| CNAG_05170 | 701.1259 | -0.34382 | 0.097645 | -3.52109 | 0.00043 | 0.002277 |
| CNAG_05173 | 967.5685 | 0.390184 | 0.087157 | 4.476778 | 7.58E-06 | 5.83E-05 |
| CNAG_07817 | 1768.445 | -0.31097 | 0.085934 | -3.61876 | 0.000296 | 0.001629 |
| CNAG_05176 | 1605.701 | 0.214595 | 0.081988 | 2.617397 | 0.00886 | 0.030718 |
| CNAG_05177 | 1103.534 | -0.46094 | 0.09519 | -4.8423 | 1.28E-06 | 1.10E-05 |
| CNAG_05178 | 171.9958 | 0.527684 | 0.15309 | 3.446884 | 0.000567 | 0.002919 |
| CNAG_05179 | 15571.52 | 0.215029 | 0.069061 | 3.113628 | 0.001848 | 0.008052 |
| CNAG_07974 | 440.3412 | 0.41916 | 0.108622 | 3.858894 | 0.000114 | 0.000692 |
| CNAG_05180 | 445.3923 | 0.279097 | 0.107666 | 2.592247 | 0.009535 | 0.032552 |
| CNAG_05183 | 705.4336 | 0.552924 | 0.095415 | 5.794934 | 6.83E-09 | 8.17E-08 |
| CNAG_05187 | 1232.865 | -0.2463 | 0.083141 | -2.96251 | 0.003051 | 0.012419 |
| CNAG_05194 | 2611.688 | 0.443034 | 0.07122 | 6.22067 | 4.95E-10 | 6.84E-09 |
| CNAG_05196 | 1457.837 | -0.18187 | 0.077619 | -2.34315 | 0.019122 | 0.057373 |
| CNAG_05199 | 32174.55 | 0.942953 | 0.063371 | 14.87996 | 4.45E-50 | 9.64E-48 |
| CNAG_05202 | 3252.115 | -0.53674 | 0.073445 | -7.30803 | 2.71E-13 | 5.48E-12 |
| CNAG_05215 | 1343.298 | 0.198734 | 0.078888 | 2.519198 | 0.011762 | 0.038747 |
| CNAG_05222 | 1382.961 | -0.24682 | 0.084159 | -2.93273 | 0.00336 | 0.013537 |
| CNAG_05227 | 1093.793 | -0.17473 | 0.087072 | -2.00671 | 0.044781 | 0.114299 |
| CNAG_05228 | 1450.917 | -0.76461 | 0.090737 | -8.42669 | 3.56E-17 | 9.64E-16 |
| CNAG_05229 | 1002.915 | 0.566358 | 0.090151 | 6.282287 | 3.34E-10 | 4.72E-09 |
| CNAG_05232 | 83113.67 | -0.21754 | 0.07636 | -2.84883 | 0.004388 | 0.016861 |
| CNAG_05234 | 2703.301 | -0.22927 | 0.081388 | -2.81702 | 0.004847 | 0.018325 |
| CNAG_05236 | 2184.484 | 0.370264 | 0.074399 | 4.976763 | 6.47E-07 | 5.80E-06 |
| CNAG_05239 | 528.453 | -0.22757 | 0.103877 | -2.19073 | 0.028472 | 0.079954 |
| CNAG_05242 | 3854.686 | -0.13748 | 0.068909 | -1.99511 | 0.046031 | 0.116557 |
| CNAG_07975 | 241.2252 | 0.858883 | 0.139151 | 6.172307 | 6.73E-10 | 9.13E-09 |
| CNAG_07976 | 57.89136 | 0.567793 | 0.273443 | 2.076455 | 0.037852 | 0.099594 |
| CNAG_07822 | 765.0994 | 0.198484 | 0.094945 | 2.090515 | 0.036572 | 0.096813 |
| CNAG_05251 | 793.7828 | 1.714329 | 0.113054 | 15.16379 | 6.14E-52 | 1.37E-49 |
| CNAG_05252 | 1324.931 | -0.52079 | 0.078328 | -6.64881 | 2.95E-11 | 4.72E-10 |
| CNAG_05253 | 578.8798 | 0.465322 | 0.117661 | 3.954749 | 7.66E-05 | 0.000488 |
| CNAG_05254 | 1287.639 | -0.52606 | 0.090196 | -5.8324 | 5.46E-09 | 6.63E-08 |
| CNAG_05258 | 42.82346 | -0.76241 | 0.317877 | -2.39843 | 0.016465 | 0.050809 |
| CNAG_07823 | 170.1142 | -0.63996 | 0.162189 | -3.94577 | 7.95E-05 | 0.000504 |

| | | | | | | |
|---|---|---|---|---|---|---|
| CNAG_05260 | 3458.246 | 0.344833 | 0.069484 | 4.962784 | 6.95E-07 | 6.21E-06 |
| CNAG_05262 | 2985.466 | -0.21163 | 0.075104 | -2.81789 | 0.004834 | 0.018287 |
| CNAG_05263 | 85.43951 | 0.661248 | 0.227064 | 2.912163 | 0.003589 | 0.01435 |
| CNAG_05264 | 208.7717 | -0.58553 | 0.152236 | -3.8462 | 0.00012 | 0.000724 |
| CNAG_05267 | 2793.939 | 0.262249 | 0.07333 | 3.576283 | 0.000349 | 0.00189 |
| CNAG_05268 | 66.95014 | -0.59696 | 0.252316 | -2.36594 | 0.017984 | 0.05461 |
| CNAG_05269 | 4351.715 | -0.13795 | 0.064161 | -2.15009 | 0.031548 | 0.086326 |
| CNAG_05270 | 2349.649 | -0.39659 | 0.074748 | -5.30566 | 1.12E-07 | 1.12E-06 |
| CNAG_05273 | 1996.68 | 0.250466 | 0.090666 | 2.762504 | 0.005736 | 0.021083 |
| CNAG_05274 | 903.4998 | -0.36689 | 0.097129 | -3.7774 | 0.000158 | 0.000927 |
| CNAG_05285 | 3684.935 | 0.171999 | 0.073038 | 2.354925 | 0.018526 | 0.055887 |
| CNAG_05286 | 440.0067 | 0.281441 | 0.107741 | 2.612191 | 0.008996 | 0.031038 |
| CNAG_05287 | 658.2214 | 0.283389 | 0.097723 | 2.899938 | 0.003732 | 0.0148 |
| CNAG_05290 | 948.066 | 1.377274 | 0.108365 | 12.70956 | 5.23E-37 | 6.09E-35 |
| CNAG_05291 | 1420.816 | -0.54233 | 0.109299 | -4.96193 | 6.98E-07 | 6.23E-06 |
| CNAG_05292 | 5265.282 | 0.372219 | 0.066895 | 5.564226 | 2.63E-08 | 2.89E-07 |
| CNAG_05294 | 940.6769 | 0.218433 | 0.093684 | 2.331585 | 0.019723 | 0.058903 |
| CNAG_05297 | 476.1889 | 0.310614 | 0.125181 | 2.481307 | 0.01309 | 0.042283 |
| CNAG_05298 | 318.0931 | -0.27647 | 0.125775 | -2.19812 | 0.027941 | 0.078803 |
| CNAG_05299 | 90.14263 | 0.628823 | 0.253559 | 2.479987 | 0.013139 | 0.042422 |
| CNAG_05303 | 126.176 | 0.367653 | 0.183003 | 2.009006 | 0.044536 | 0.113806 |
| CNAG_12402 | 23.95934 | 1.380445 | 0.405055 | 3.408045 | 0.000654 | 0.003294 |
| CNAG_05304 | 166.5381 | 0.414854 | 0.180245 | 2.301604 | 0.021358 | 0.062901 |
| CNAG_05309 | 3810.435 | 0.536049 | 0.073655 | 7.277805 | 3.39E-13 | 6.72E-12 |
| CNAG_05310 | 56.92236 | -1.44712 | 0.291776 | -4.95969 | 7.06E-07 | 6.30E-06 |
| CNAG_05312 | 3033.231 | -0.66176 | 0.083825 | -7.89457 | 2.91E-15 | 6.85E-14 |
| CNAG_05313 | 932.0297 | -0.53524 | 0.112027 | -4.77778 | 1.77E-06 | 1.50E-05 |
| CNAG_05315 | 276.4301 | 0.88012 | 0.149371 | 5.89219 | 3.81E-09 | 4.74E-08 |
| CNAG_05317 | 521.6301 | -0.60925 | 0.102431 | -5.94791 | 2.72E-09 | 3.43E-08 |
| CNAG_05318 | 594.3719 | -0.30178 | 0.108599 | -2.77882 | 0.005456 | 0.020226 |
| CNAG_05321 | 170.7561 | -0.38788 | 0.155426 | -2.4956 | 0.012575 | 0.040991 |
| CNAG_07830 | 714.148 | 0.215423 | 0.091183 | 2.362542 | 0.01815 | 0.054943 |
| CNAG_05324 | 39.15003 | -1.00705 | 0.330442 | -3.04758 | 0.002307 | 0.009726 |
| CNAG_05325 | 182.443 | -0.53678 | 0.152256 | -3.52554 | 0.000423 | 0.002242 |
| CNAG_06876 | 461.1527 | 0.293192 | 0.104126 | 2.815731 | 0.004867 | 0.018374 |
| CNAG_06875 | 645.1746 | 0.593469 | 0.097811 | 6.067484 | 1.30E-09 | 1.70E-08 |
| CNAG_06874 | 698.1442 | 0.43823 | 0.097088 | 4.513739 | 6.37E-06 | 4.95E-05 |
| CNAG_06873 | 87.74281 | 1.114458 | 0.224993 | 4.953298 | 7.30E-07 | 6.48E-06 |
| CNAG_06872 | 48.83584 | 0.789081 | 0.281852 | 2.799629 | 0.005116 | 0.019149 |
| CNAG_06871 | 1300.032 | -0.27366 | 0.082608 | -3.31278 | 0.000924 | 0.004433 |
| CNAG_12410 | 146.3223 | -0.41754 | 0.181535 | -2.30007 | 0.021444 | 0.06308 |
| CNAG_06868 | 1899.638 | 0.834133 | 0.077345 | 10.78463 | 4.07E-27 | 2.60E-25 |
| CNAG_06867 | 668.9084 | 0.304458 | 0.109854 | 2.771468 | 0.00558 | 0.02061 |
| CNAG_06866 | 2108.361 | 0.34367 | 0.075681 | 4.54103 | 5.60E-06 | 4.38E-05 |
| CNAG_06865 | 971.9362 | -0.46505 | 0.08489 | -5.4782 | 4.30E-08 | 4.59E-07 |
| CNAG_06864 | 2379.576 | -0.2594 | 0.071726 | -3.61654 | 0.000299 | 0.00164 |
| CNAG_06860 | 2454.22 | -0.18314 | 0.083494 | -2.19349 | 0.028272 | 0.079566 |
| CNAG_07979 | 1145.398 | -0.18524 | 0.083191 | -2.22674 | 0.025965 | 0.073979 |

| | | | | | | |
|---|---|---|---|---|---|---|
| CNAG_06853 | 218.799 | 0.299338 | 0.144412 | 2.072805 | 0.03819 | 0.100383 |
| CNAG_07401 | 217.5578 | 0.323623 | 0.148446 | 2.180064 | 0.029253 | 0.081618 |
| CNAG_06846 | 448.4575 | 0.270946 | 0.114163 | 2.373328 | 0.017629 | 0.053736 |
| CNAG_12415 | 230.7369 | -0.29922 | 0.145722 | -2.0534 | 0.040034 | 0.104385 |
| CNAG_06838 | 1260.051 | -0.30658 | 0.082075 | -3.73543 | 0.000187 | 0.001077 |
| CNAG_06835 | 606.3847 | 0.751151 | 0.103992 | 7.223193 | 5.08E-13 | 9.91E-12 |
| CNAG_06832 | 919.6752 | -0.75861 | 0.084116 | -9.01859 | 1.91E-19 | 6.49E-18 |
| CNAG_06831 | 159.6815 | -0.51632 | 0.168208 | -3.06955 | 0.002144 | 0.009112 |
| CNAG_06830 | 4672.78 | -0.26563 | 0.062398 | -4.25709 | 2.07E-05 | 0.000146 |
| CNAG_06828 | 1534.709 | 0.385337 | 0.089986 | 4.282178 | 1.85E-05 | 0.000133 |
| CNAG_06824 | 1325.068 | -0.36618 | 0.096971 | -3.77614 | 0.000159 | 0.000931 |
| CNAG_06821 | 1175.33 | -0.5673 | 0.077982 | -7.27481 | 3.47E-13 | 6.85E-12 |
| CNAG_06820 | 965.4402 | -0.50577 | 0.089535 | -5.64882 | 1.62E-08 | 1.84E-07 |
| CNAG_07405 | 198.233 | 0.508251 | 0.149977 | 3.388853 | 0.000702 | 0.0035 |
| CNAG_06818 | 443.0468 | 0.389004 | 0.10614 | 3.665007 | 0.000247 | 0.001392 |
| CNAG_06817 | 546.4015 | 1.807398 | 0.129078 | 14.00241 | 1.51E-44 | 2.56E-42 |
| CNAG_12420 | 105.0874 | 0.989222 | 0.197205 | 5.0162 | 5.27E-07 | 4.83E-06 |
| CNAG_06811 | 21513.71 | -0.20119 | 0.080384 | -2.50287 | 0.012319 | 0.040225 |
| CNAG_06807 | 289.627 | -0.35184 | 0.134529 | -2.61532 | 0.008914 | 0.030824 |
| CNAG_06806 | 4596.488 | 0.203891 | 0.067561 | 3.017896 | 0.002545 | 0.010617 |
| CNAG_06971 | 3264.594 | -0.3404 | 0.090266 | -3.7711 | 0.000163 | 0.000947 |
| CNAG_06980 | 2562.227 | -0.30437 | 0.069991 | -4.34868 | 1.37E-05 | 0.000101 |
| CNAG_07409 | 2796.087 | -0.38015 | 0.078264 | -4.85723 | 1.19E-06 | 1.02E-05 |
| CNAG_07005 | 696.7689 | 0.271701 | 0.090722 | 2.994865 | 0.002746 | 0.011373 |
| CNAG_07004 | 27309.86 | 0.861306 | 0.061898 | 13.91489 | 5.14E-44 | 8.36E-42 |
| CNAG_07015 | 1317.905 | -0.28916 | 0.092502 | -3.12602 | 0.001772 | 0.007767 |
| CNAG_07411 | 3388.1 | -0.19513 | 0.093707 | -2.08235 | 0.037311 | 0.098424 |
| CNAG_01454 | 3590.459 | -0.31048 | 0.090359 | -3.43606 | 0.00059 | 0.003028 |
| CNAG_01451 | 1164.839 | 0.31423 | 0.11599 | 2.709121 | 0.006746 | 0.02434 |
| CNAG_07413 | 7999.756 | -0.60708 | 0.062926 | -9.64746 | 5.04E-22 | 2.16E-20 |
| CNAG_01446 | 5879.527 | 0.969181 | 0.084784 | 11.43118 | 2.92E-30 | 2.11E-28 |
| CNAG_01442 | 2350.075 | -0.18259 | 0.078411 | -2.32862 | 0.019879 | 0.059304 |
| CNAG_01440 | 454.1115 | 0.286396 | 0.108412 | 2.641739 | 0.008248 | 0.029008 |
| CNAG_01437 | 2663.192 | -0.33679 | 0.068117 | -4.9443 | 7.64E-07 | 6.77E-06 |
| CNAG_01432 | 2614.233 | -0.41613 | 0.070603 | -5.89401 | 3.77E-09 | 4.69E-08 |
| CNAG_01428 | 38911.13 | -0.18774 | 0.07258 | -2.58668 | 0.009691 | 0.033024 |
| CNAG_12433 | 29.73471 | 1.211151 | 0.381193 | 3.177261 | 0.001487 | 0.006683 |
| CNAG_01422 | 2086.58 | 0.632073 | 0.073785 | 8.566403 | 1.07E-17 | 3.01E-16 |
| CNAG_01419 | 1328.89 | 0.488479 | 0.08365 | 5.839544 | 5.23E-09 | 6.36E-08 |
| CNAG_01417 | 785.9455 | 0.452109 | 0.109453 | 4.130602 | 3.62E-05 | 0.000243 |
| CNAG_01415 | 1449.127 | -0.3351 | 0.109122 | -3.07089 | 0.002134 | 0.009081 |
| CNAG_01413 | 7738.643 | -0.17888 | 0.061935 | -2.88816 | 0.003875 | 0.015281 |
| CNAG_01412 | 2156.698 | 0.420988 | 0.075786 | 5.554942 | 2.78E-08 | 3.04E-07 |
| CNAG_01411 | 191.735 | -0.67058 | 0.198548 | -3.37743 | 0.000732 | 0.00363 |
| CNAG_07417 | 118.0202 | -0.66074 | 0.206274 | -3.20322 | 0.001359 | 0.006199 |
| CNAG_07418 | 819.9973 | -0.20794 | 0.095619 | -2.17466 | 0.029656 | 0.082407 |
| CNAG_01408 | 1311.702 | -0.38983 | 0.09882 | -3.94481 | 7.99E-05 | 0.000506 |
| CNAG_01404 | 9495.195 | -0.16448 | 0.078239 | -2.10227 | 0.03553 | 0.094601 |

| | | | | | | |
|---|---|---|---|---|---|---|
| CNAG_01403 | 1511.366 | 0.298866 | 0.074565 | 4.008131 | 6.12E-05 | 0.000397 |
| CNAG_01399 | 986.8728 | 0.258445 | 0.100635 | 2.568131 | 0.010225 | 0.034447 |
| CNAG_01398 | 3684.181 | -0.42077 | 0.084221 | -4.99601 | 5.85E-07 | 5.31E-06 |
| CNAG_01397 | 4707.667 | 0.398275 | 0.074435 | 5.350617 | 8.77E-08 | 8.95E-07 |
| CNAG_01387 | 2425.255 | 0.53798 | 0.092379 | 5.823617 | 5.76E-09 | 6.97E-08 |
| CNAG_12434 | 198.4384 | -0.48401 | 0.166474 | -2.90744 | 0.003644 | 0.014487 |
| CNAG_01380 | 893.7104 | 0.270456 | 0.08554 | 3.161761 | 0.001568 | 0.006985 |
| CNAG_01379 | 976.4956 | -0.29369 | 0.089012 | -3.29948 | 0.000969 | 0.004618 |
| CNAG_01377 | 2606.867 | -0.16286 | 0.080491 | -2.02338 | 0.043034 | 0.110765 |
| CNAG_01376 | 6893.488 | -0.19839 | 0.081833 | -2.42429 | 0.015338 | 0.048075 |
| CNAG_01375 | 1296.458 | 0.16784 | 0.083947 | 1.999354 | 0.04557 | 0.115903 |
| CNAG_01365 | 936.4963 | -0.2998 | 0.08718 | -3.43885 | 0.000584 | 0.003001 |
| CNAG_01361 | 6849.202 | -0.20313 | 0.066205 | -3.06816 | 0.002154 | 0.009145 |
| CNAG_01356 | 663.9807 | -0.25624 | 0.103203 | -2.48287 | 0.013033 | 0.042203 |
| CNAG_01354 | 4796.911 | -1.14426 | 0.066213 | -17.2814 | 6.50E-67 | 3.38E-64 |
| CNAG_01353 | 479.1572 | -0.7556 | 0.111984 | -6.74745 | 1.50E-11 | 2.51E-10 |
| CNAG_07423 | 696.2413 | 0.201605 | 0.094854 | 2.125417 | 0.033552 | 0.090445 |
| CNAG_01352 | 1737.419 | -0.43921 | 0.107449 | -4.08762 | 4.36E-05 | 0.000289 |
| CNAG_01348 | 1662.762 | 0.975076 | 0.092006 | 10.59801 | 3.04E-26 | 1.80E-24 |
| CNAG_01344 | 868.5762 | 0.20522 | 0.090526 | 2.266981 | 0.023391 | 0.067635 |
| CNAG_01341 | 1265.024 | 1.078515 | 0.099517 | 10.83745 | 2.29E-27 | 1.51E-25 |
| CNAG_01340 | 3030.008 | -0.1918 | 0.075263 | -2.54835 | 0.010823 | 0.036174 |
| CNAG_01337 | 1004.983 | 0.229865 | 0.089771 | 2.56058 | 0.01045 | 0.03509 |
| CNAG_01334 | 2102.588 | -0.32585 | 0.091522 | -3.56039 | 0.00037 | 0.001989 |
| CNAG_01332 | 39096.97 | -0.19528 | 0.070814 | -2.75768 | 0.005821 | 0.021359 |
| CNAG_01323 | 10376.11 | 0.169097 | 0.077757 | 2.174693 | 0.029653 | 0.082407 |
| CNAG_01317 | 1350.234 | -0.17802 | 0.082106 | -2.16819 | 0.030144 | 0.08342 |
| CNAG_01310 | 1419.668 | -0.45601 | 0.081903 | -5.56765 | 2.58E-08 | 2.84E-07 |
| CNAG_01307 | 4177.817 | 0.431031 | 0.069895 | 6.166866 | 6.97E-10 | 9.44E-09 |
| CNAG_01306 | 565.385 | -0.31318 | 0.124697 | -2.51156 | 0.01202 | 0.039397 |
| CNAG_01304 | 1328.214 | -0.45384 | 0.082519 | -5.49985 | 3.80E-08 | 4.09E-07 |
| CNAG_01302 | 1434.1 | 0.773707 | 0.075543 | 10.2419 | 1.29E-24 | 6.97E-23 |
| CNAG_01299 | 121.5552 | -0.37865 | 0.186579 | -2.02943 | 0.042414 | 0.109567 |
| CNAG_01296 | 160.1438 | -0.39022 | 0.165432 | -2.35879 | 0.018334 | 0.055415 |
| CNAG_01293 | 1503.151 | 0.208793 | 0.099845 | 2.091173 | 0.036513 | 0.09669 |
| CNAG_01291 | 1141.067 | 0.586937 | 0.11007 | 5.332374 | 9.69E-08 | 9.82E-07 |
| CNAG_01287 | 17949.38 | 0.215766 | 0.059692 | 3.61465 | 0.000301 | 0.001651 |
| CNAG_01277 | 4437.698 | -0.21901 | 0.066504 | -3.29322 | 0.00099 | 0.004713 |
| CNAG_01275 | 1414.158 | 0.296509 | 0.075401 | 3.932433 | 8.41E-05 | 0.000531 |
| CNAG_01274 | 8243.447 | -0.26276 | 0.064774 | -4.05652 | 4.98E-05 | 0.000328 |
| CNAG_01272 | 508.6443 | 0.725073 | 0.108431 | 6.686939 | 2.28E-11 | 3.74E-10 |
| CNAG_01270 | 9149.655 | -0.13088 | 0.060747 | -2.15448 | 0.031203 | 0.085591 |
| CNAG_07429 | 2017.008 | -0.23533 | 0.073316 | -3.20984 | 0.001328 | 0.006065 |
| CNAG_01268 | 47.8784 | 1.050683 | 0.328436 | 3.199054 | 0.001379 | 0.00626 |
| CNAG_01267 | 498.0027 | 0.447135 | 0.104763 | 4.268053 | 1.97E-05 | 0.00014 |
| CNAG_01263 | 5797.257 | 0.335221 | 0.074128 | 4.522185 | 6.12E-06 | 4.76E-05 |
| CNAG_01261 | 4968.263 | 0.440789 | 0.068142 | 6.46873 | 9.88E-11 | 1.49E-09 |
| CNAG_06947 | 902.8547 | -0.30161 | 0.084454 | -3.57134 | 0.000355 | 0.001917 |

| | | | | | | |
|---|---|---|---|---|---|---|
| CNAG_06948 | 3405.662 | -0.63633 | 0.072685 | -8.7545 | 2.05E-18 | 6.27E-17 |
| CNAG_06949 | 4464.795 | 0.246143 | 0.067526 | 3.64517 | 0.000267 | 0.001493 |
| CNAG_01257 | 1014.395 | -0.25853 | 0.091301 | -2.83158 | 0.004632 | 0.017693 |
| CNAG_01255 | 1639.587 | 0.281359 | 0.08301 | 3.389458 | 0.0007 | 0.003497 |
| CNAG_01252 | 203.6246 | 0.584059 | 0.148225 | 3.940361 | 8.14E-05 | 0.000515 |
| CNAG_01251 | 2206.67 | -0.21129 | 0.100211 | -2.10849 | 0.034989 | 0.093657 |
| CNAG_01250 | 1564.179 | -0.16713 | 0.083161 | -2.00965 | 0.044468 | 0.113668 |
| CNAG_01242 | 558.304 | -0.42225 | 0.107719 | -3.91996 | 8.86E-05 | 0.000552 |
| CNAG_01236 | 4240.49 | -0.50365 | 0.065985 | -7.63273 | 2.30E-14 | 5.02E-13 |
| CNAG_01234 | 1632.931 | 0.355808 | 0.100817 | 3.52924 | 0.000417 | 0.002217 |
| CNAG_01229 | 1000.835 | -0.43101 | 0.089448 | -4.8185 | 1.45E-06 | 1.23E-05 |
| CNAG_07432 | 163.3698 | -0.48827 | 0.172273 | -2.83429 | 0.004593 | 0.017561 |
| CNAG_01224 | 53456.69 | -0.22975 | 0.078996 | -2.90841 | 0.003633 | 0.014474 |
| CNAG_01223 | 325.4751 | 0.3865 | 0.130836 | 2.954078 | 0.003136 | 0.012727 |
| CNAG_01221 | 781.4395 | 0.187313 | 0.090602 | 2.067427 | 0.038694 | 0.101502 |
| CNAG_07433 | 1203.827 | -0.28134 | 0.113765 | -2.47298 | 0.013399 | 0.043014 |
| CNAG_01217 | 2020.496 | -0.22142 | 0.068948 | -3.21139 | 0.001321 | 0.006035 |
| CNAG_01212 | 1095.748 | 0.217836 | 0.08689 | 2.507018 | 0.012175 | 0.03984 |
| CNAG_01208 | 1025.612 | -0.3878 | 0.125881 | -3.08072 | 0.002065 | 0.008844 |
| CNAG_01207 | 404.6216 | -0.459 | 0.111968 | -4.0994 | 4.14E-05 | 0.000276 |
| CNAG_07434 | 780.4192 | 0.431704 | 0.092161 | 4.684221 | 2.81E-06 | 2.31E-05 |
| CNAG_01203 | 986.491 | -0.1709 | 0.08649 | -1.97601 | 0.048154 | 0.12099 |
| CNAG_01202 | 673.2507 | -0.3114 | 0.092155 | -3.3791 | 0.000727 | 0.003613 |
| CNAG_01194 | 24.55854 | 0.793559 | 0.395968 | 2.004098 | 0.04506 | 0.114804 |
| CNAG_01189 | 2893.473 | -0.62441 | 0.211271 | -2.9555 | 0.003122 | 0.012675 |
| CNAG_01187 | 2075.639 | -0.47274 | 0.077241 | -6.12042 | 9.33E-10 | 1.25E-08 |
| CNAG_01185 | 2362.816 | -0.21096 | 0.073688 | -2.86289 | 0.004198 | 0.016299 |
| CNAG_01184 | 1239.812 | 0.266348 | 0.081197 | 3.280273 | 0.001037 | 0.004905 |
| CNAG_01183 | 1058.232 | -0.32363 | 0.084392 | -3.83481 | 0.000126 | 0.000757 |
| CNAG_01182 | 8672.821 | -0.14371 | 0.069977 | -2.05365 | 0.04001 | 0.104358 |
| CNAG_01179 | 2052.404 | -0.50917 | 0.071242 | -7.147 | 8.87E-13 | 1.68E-11 |
| CNAG_01171 | 418.1112 | -0.67232 | 0.112439 | -5.97944 | 2.24E-09 | 2.85E-08 |
| CNAG_01170 | 58945.63 | -0.19475 | 0.073799 | -2.63889 | 0.008318 | 0.029135 |
| CNAG_01168 | 10321.05 | -0.36972 | 0.067123 | -5.50802 | 3.63E-08 | 3.93E-07 |
| CNAG_01165 | 936.5873 | -0.32118 | 0.112222 | -2.86202 | 0.00421 | 0.016336 |
| CNAG_01161 | 1306.261 | -0.1822 | 0.079783 | -2.28375 | 0.022386 | 0.065216 |
| CNAG_07439 | 3329.705 | -0.55115 | 0.067952 | -8.11085 | 5.03E-16 | 1.25E-14 |
| CNAG_01158 | 5811.8 | -0.28735 | 0.061026 | -4.70869 | 2.49E-06 | 2.06E-05 |
| CNAG_01155 | 7051.981 | 0.455403 | 0.070505 | 6.459191 | 1.05E-10 | 1.59E-09 |
| CNAG_01153 | 37096.71 | -0.18728 | 0.065247 | -2.87031 | 0.004101 | 0.015993 |
| CNAG_01152 | 48269.75 | -0.26948 | 0.087707 | -3.07244 | 0.002123 | 0.009044 |
| CNAG_01150 | 10180.07 | 0.339019 | 0.062082 | 5.460856 | 4.74E-08 | 5.03E-07 |
| CNAG_01148 | 14453.08 | -0.2388 | 0.076403 | -3.1255 | 0.001775 | 0.007773 |
| CNAG_01147 | 1923.37 | 0.222894 | 0.071037 | 3.137722 | 0.001703 | 0.007516 |
| CNAG_01146 | 2984.077 | -0.5255 | 0.078777 | -6.67065 | 2.55E-11 | 4.14E-10 |
| CNAG_01143 | 853.5342 | -0.26498 | 0.090374 | -2.93204 | 0.003367 | 0.013553 |
| CNAG_01140 | 2688.374 | -0.64567 | 0.082556 | -7.82092 | 5.24E-15 | 1.20E-13 |
| CNAG_12461 | 75.51489 | -0.87176 | 0.269098 | -3.23958 | 0.001197 | 0.005544 |

| | | | | | | |
|---|---|---|---|---|---|---|
| CNAG_01139 | 55.32781 | -0.89995 | 0.278611 | -3.23015 | 0.001237 | 0.005697 |
| CNAG_01138 | 9575.89 | 0.360042 | 0.065491 | 5.497563 | 3.85E-08 | 4.14E-07 |
| CNAG_01137 | 14100.46 | 1.139126 | 0.062562 | 18.20789 | 4.47E-74 | 3.17E-71 |
| CNAG_01133 | 468.5856 | 0.226669 | 0.11096 | 2.042802 | 0.041072 | 0.106735 |
| CNAG_01132 | 1379.892 | -0.49117 | 0.083949 | -5.85079 | 4.89E-09 | 6.00E-08 |
| CNAG_01130 | 4017.855 | -0.20316 | 0.073425 | -2.76684 | 0.00566 | 0.020846 |
| CNAG_01129 | 1598.254 | -0.45719 | 0.080283 | -5.69476 | 1.24E-08 | 1.43E-07 |
| CNAG_01128 | 1587.335 | 0.175237 | 0.082969 | 2.112081 | 0.034679 | 0.092881 |
| CNAG_01123 | 1391.663 | -0.27205 | 0.079797 | -3.40927 | 0.000651 | 0.003282 |
| CNAG_01122 | 68.13906 | -0.6849 | 0.268056 | -2.55506 | 0.010617 | 0.035575 |
| CNAG_01120 | 22275.08 | 0.605678 | 0.059217 | 10.22814 | 1.48E-24 | 7.98E-23 |
| CNAG_12466 | 76.28466 | -0.55472 | 0.228924 | -2.42317 | 0.015386 | 0.048182 |
| CNAG_01118 | 346.0133 | 1.619035 | 0.12624 | 12.82505 | 1.19E-37 | 1.45E-35 |
| CNAG_01112 | 2496.325 | 0.341139 | 0.071016 | 4.803681 | 1.56E-06 | 1.32E-05 |
| CNAG_01108 | 819.6716 | -0.21547 | 0.098944 | -2.1777 | 0.029429 | 0.081934 |
| CNAG_01105 | 1401.804 | 0.18212 | 0.091525 | 1.989835 | 0.046609 | 0.11765 |
| CNAG_01104 | 1449.88 | -0.50122 | 0.076808 | -6.52565 | 6.77E-11 | 1.04E-09 |
| CNAG_01103 | 2360.435 | -0.18878 | 0.06972 | -2.70771 | 0.006775 | 0.024399 |
| CNAG_01102 | 1865.046 | 1.035824 | 0.088832 | 11.66045 | 2.03E-31 | 1.58E-29 |
| CNAG_01101 | 1323.795 | -0.35135 | 0.120424 | -2.91762 | 0.003527 | 0.01413 |
| CNAG_01098 | 1954.045 | -0.44244 | 0.078416 | -5.64218 | 1.68E-08 | 1.90E-07 |
| CNAG_01097 | 1526.62 | 0.230869 | 0.082438 | 2.800504 | 0.005102 | 0.019125 |
| CNAG_01095 | 724.9723 | 0.24145 | 0.105108 | 2.297159 | 0.02161 | 0.063471 |
| CNAG_01094 | 1828.121 | -0.18631 | 0.073637 | -2.53007 | 0.011404 | 0.037758 |
| CNAG_01091 | 6582.737 | -0.1296 | 0.060243 | -2.15135 | 0.031449 | 0.086205 |
| CNAG_01090 | 425.5922 | 1.105217 | 0.109837 | 10.06235 | 8.10E-24 | 4.08E-22 |
| CNAG_01089 | 1594.199 | 0.54837 | 0.075145 | 7.297522 | 2.93E-13 | 5.90E-12 |
| CNAG_01086 | 382.7424 | 0.667254 | 0.12308 | 5.421285 | 5.92E-08 | 6.18E-07 |
| CNAG_01084 | 429.0033 | -0.44279 | 0.13097 | -3.38087 | 0.000723 | 0.003592 |
| CNAG_01082 | 2193.215 | 0.373825 | 0.093523 | 3.997156 | 6.41E-05 | 0.000414 |
| CNAG_12474 | 14.73703 | -1.04942 | 0.519046 | -2.02182 | 0.043195 | 0.111068 |
| CNAG_01078 | 268.9968 | 0.454447 | 0.137479 | 3.30558 | 0.000948 | 0.004535 |
| CNAG_01077 | 76.56694 | 1.25304 | 0.235394 | 5.323168 | 1.02E-07 | 1.03E-06 |
| CNAG_12476 | 11.92371 | 1.213259 | 0.577112 | 2.102295 | 0.035527 | 0.094601 |
| CNAG_01076 | 97.46243 | 1.040796 | 0.205695 | 5.059894 | 4.19E-07 | 3.89E-06 |
| CNAG_01075 | 78.20411 | 0.686675 | 0.24571 | 2.794652 | 0.005196 | 0.0194 |
| CNAG_01074 | 81.84852 | 1.201897 | 0.225798 | 5.322879 | 1.02E-07 | 1.03E-06 |
| CNAG_01073 | 474.0832 | 0.224189 | 0.103953 | 2.156635 | 0.031034 | 0.085308 |
| CNAG_01061 | 540.7059 | 1.235064 | 0.115498 | 10.69339 | 1.09E-26 | 6.72E-25 |
| CNAG_01060 | 9345.746 | -0.21054 | 0.070008 | -3.00738 | 0.002635 | 0.010944 |
| CNAG_01055 | 10026.59 | 0.715111 | 0.066593 | 10.73847 | 6.71E-27 | 4.23E-25 |
| CNAG_01052 | 8512.53 | 1.277351 | 0.192214 | 6.645475 | 3.02E-11 | 4.80E-10 |
| CNAG_01050 | 971.342 | 0.23034 | 0.090954 | 2.532507 | 0.011325 | 0.037513 |
| CNAG_01047 | 313.0765 | 1.222045 | 0.144403 | 8.462724 | 2.61E-17 | 7.15E-16 |
| CNAG_01044 | 1501.386 | 0.956956 | 0.075637 | 12.65198 | 1.09E-36 | 1.23E-34 |
| CNAG_01043 | 10036.66 | 0.505203 | 0.070296 | 7.186779 | 6.63E-13 | 1.28E-11 |
| CNAG_01041 | 247.8097 | -0.32278 | 0.133841 | -2.41167 | 0.01588 | 0.049392 |
| CNAG_01037 | 528.7605 | -0.21813 | 0.106442 | -2.04927 | 0.040436 | 0.105292 |

| | | | | | | |
|---|---|---|---|---|---|---|
| CNAG_01036 | 579.6195 | 0.242048 | 0.102441 | 2.362807 | 0.018137 | 0.054925 |
| CNAG_01035 | 4452.08 | -0.16754 | 0.078017 | -2.14745 | 0.031757 | 0.086807 |
| CNAG_01032 | 340.0936 | 0.333049 | 0.139623 | 2.385348 | 0.017063 | 0.052363 |
| CNAG_01031 | 3180.227 | 0.830144 | 0.090912 | 9.131292 | 6.77E-20 | 2.45E-18 |
| CNAG_01029 | 1744.289 | 0.212082 | 0.083895 | 2.527957 | 0.011473 | 0.037954 |
| CNAG_01028 | 599.363 | 0.284779 | 0.095405 | 2.984949 | 0.002836 | 0.01168 |
| CNAG_01027 | 1042.955 | 0.441917 | 0.092534 | 4.775728 | 1.79E-06 | 1.51E-05 |
| CNAG_01026 | 777.499 | 0.611136 | 0.087238 | 7.005376 | 2.46E-12 | 4.42E-11 |
| CNAG_01022 | 690.0185 | 0.281495 | 0.098434 | 2.859739 | 0.00424 | 0.016413 |
| CNAG_01021 | 3899.804 | -0.29279 | 0.067952 | -4.30883 | 1.64E-05 | 0.000119 |
| CNAG_01019 | 26374.45 | -0.35212 | 0.063455 | -5.54918 | 2.87E-08 | 3.14E-07 |
| CNAG_01016 | 1099.661 | 0.347691 | 0.083646 | 4.156719 | 3.23E-05 | 0.000219 |
| CNAG_01011 | 485.5384 | 0.23987 | 0.11391 | 2.105782 | 0.035223 | 0.094041 |
| CNAG_01010 | 2363.477 | 0.170828 | 0.079441 | 2.150377 | 0.031525 | 0.086324 |
| CNAG_01007 | 1374.054 | 0.32507 | 0.09821 | 3.309952 | 0.000933 | 0.004476 |
| CNAG_01005 | 3621.256 | 0.256048 | 0.066794 | 3.833381 | 0.000126 | 0.000761 |
| CNAG_01004 | 1874.224 | -0.95462 | 0.076712 | -12.4442 | 1.50E-35 | 1.54E-33 |
| CNAG_00999 | 918.8093 | -0.24938 | 0.087458 | -2.85136 | 0.004353 | 0.016777 |
| CNAG_00995 | 4873.01 | 0.97974 | 0.089078 | 10.99863 | 3.88E-28 | 2.61E-26 |
| CNAG_00993 | 739.256 | -0.18387 | 0.092297 | -1.99219 | 0.046351 | 0.117251 |
| CNAG_00992 | 12496.18 | -0.25812 | 0.065872 | -3.91852 | 8.91E-05 | 0.000554 |
| CNAG_00990 | 6563.646 | 0.184832 | 0.081606 | 2.264921 | 0.023518 | 0.067899 |
| CNAG_00988 | 4157.636 | -0.23235 | 0.082439 | -2.81849 | 0.004825 | 0.018269 |
| CNAG_00985 | 340.6848 | 0.242336 | 0.116967 | 2.07183 | 0.038281 | 0.100588 |
| CNAG_00984 | 1273.931 | 1.262361 | 0.119527 | 10.56134 | 4.50E-26 | 2.60E-24 |
| CNAG_00973 | 1004.577 | 0.191052 | 0.087824 | 2.175396 | 0.0296 | 0.082354 |
| CNAG_00972 | 3669.323 | 0.202937 | 0.063917 | 3.175022 | 0.001498 | 0.00672 |
| CNAG_00967 | 896.6043 | 0.358236 | 0.093073 | 3.848963 | 0.000119 | 0.000717 |
| CNAG_07445 | 31526.75 | 0.238804 | 0.073927 | 3.230266 | 0.001237 | 0.005697 |
| CNAG_07446 | 1605.492 | 0.248661 | 0.079921 | 3.111323 | 0.001863 | 0.008107 |
| CNAG_00961 | 1397.393 | 0.960079 | 0.095299 | 10.07443 | 7.17E-24 | 3.68E-22 |
| CNAG_00940 | 2882.418 | -0.30701 | 0.069922 | -4.39074 | 1.13E-05 | 8.46E-05 |
| CNAG_00938 | 440.5637 | 0.44451 | 0.112081 | 3.965977 | 7.31E-05 | 0.000468 |
| CNAG_00932 | 240.6116 | 0.376645 | 0.156606 | 2.40504 | 0.016171 | 0.050078 |
| CNAG_00930 | 12852.53 | -0.22808 | 0.064266 | -3.54892 | 0.000387 | 0.002068 |
| CNAG_00929 | 1214.159 | 0.514738 | 0.080431 | 6.399789 | 1.56E-10 | 2.30E-09 |
| CNAG_00927 | 783.96 | -0.1796 | 0.088518 | -2.02893 | 0.042466 | 0.109617 |
| CNAG_00922 | 832.4292 | 0.203199 | 0.088741 | 2.289794 | 0.022033 | 0.064408 |
| CNAG_00921 | 369.137 | 0.352935 | 0.126847 | 2.782367 | 0.005396 | 0.020025 |
| CNAG_00920 | 589.5963 | 0.379802 | 0.104386 | 3.638424 | 0.000274 | 0.001529 |
| CNAG_00919 | 1815.226 | 1.109328 | 0.081753 | 13.56926 | 6.09E-42 | 8.65E-40 |
| CNAG_00917 | 1229.498 | 0.259054 | 0.079686 | 3.250916 | 0.00115 | 0.005353 |
| CNAG_00915 | 214.495 | 0.419831 | 0.166703 | 2.518433 | 0.011788 | 0.038815 |
| CNAG_07448 | 795.1995 | 1.521994 | 0.128625 | 11.83281 | 2.64E-32 | 2.19E-30 |
| CNAG_12498 | 209.7306 | 0.697903 | 0.152454 | 4.577794 | 4.70E-06 | 3.72E-05 |
| CNAG_12499 | 38.56846 | 1.092343 | 0.339063 | 3.221654 | 0.001275 | 0.005851 |
| CNAG_07450 | 162.0987 | 0.360924 | 0.169136 | 2.133925 | 0.032849 | 0.089012 |
| CNAG_07982 | 1639.874 | -0.22583 | 0.072931 | -3.09645 | 0.001959 | 0.008472 |

| | | | | | | |
|---|---|---|---|---|---|---|
| CNAG_00906 | 3886.928 | -0.5482 | 0.074984 | -7.31084 | 2.65E-13 | 5.38E-12 |
| CNAG_00905 | 234.0273 | -0.40807 | 0.144559 | -2.82286 | 0.00476 | 0.018093 |
| CNAG_00904 | 572.1044 | -0.47607 | 0.118137 | -4.02978 | 5.58E-05 | 0.000365 |
| CNAG_00903 | 608.3082 | -0.64773 | 0.103183 | -6.27751 | 3.44E-10 | 4.85E-09 |
| CNAG_00899 | 1026.776 | -0.73233 | 0.084651 | -8.65112 | 5.10E-18 | 1.49E-16 |
| CNAG_00897 | 3673.792 | -0.17113 | 0.073866 | -2.31671 | 0.020519 | 0.060864 |
| CNAG_12505 | 9.594742 | 1.462319 | 0.663793 | 2.202974 | 0.027597 | 0.07803 |
| CNAG_00895 | 1470.278 | 0.333367 | 0.096508 | 3.454297 | 0.000552 | 0.002846 |
| CNAG_00893 | 1113.076 | 0.372729 | 0.093919 | 3.968619 | 7.23E-05 | 0.000463 |
| CNAG_00892 | 308.9963 | 0.290372 | 0.121672 | 2.386518 | 0.017009 | 0.052217 |
| CNAG_00891 | 9321.768 | -0.1957 | 0.065429 | -2.99096 | 0.002781 | 0.011489 |
| CNAG_00886 | 33735.05 | -0.45439 | 0.062919 | -7.22192 | 5.13E-13 | 9.98E-12 |
| CNAG_00884 | 1251.636 | 0.26062 | 0.079339 | 3.284897 | 0.00102 | 0.004837 |
| CNAG_00883 | 447.4398 | -0.31341 | 0.11377 | -2.75477 | 0.005873 | 0.021529 |
| CNAG_00877 | 1051.708 | 0.197667 | 0.079809 | 2.476754 | 0.013258 | 0.04272 |
| CNAG_00875 | 801.6889 | 0.204976 | 0.093351 | 2.19575 | 0.02811 | 0.079166 |
| CNAG_00874 | 4475.332 | -0.23848 | 0.070169 | -3.39869 | 0.000677 | 0.003396 |
| CNAG_00873 | 474.4232 | 0.71811 | 0.114622 | 6.265001 | 3.73E-10 | 5.23E-09 |
| CNAG_00871 | 128.2501 | -0.81196 | 0.187214 | -4.33707 | 1.44E-05 | 0.000106 |
| CNAG_12511 | 30.86287 | 2.472362 | 0.403411 | 6.128639 | 8.86E-10 | 1.19E-08 |
| CNAG_00866 | 1554.684 | 0.824885 | 0.08482 | 9.72512 | 2.36E-22 | 1.03E-20 |
| CNAG_07456 | 132.9135 | -0.65765 | 0.180302 | -3.6475 | 0.000265 | 0.001481 |
| CNAG_07457 | 59.55037 | -0.62795 | 0.263029 | -2.38737 | 0.01697 | 0.052138 |
| CNAG_00863 | 96.9884 | -0.70741 | 0.203098 | -3.48312 | 0.000496 | 0.002591 |
| CNAG_00862 | 774.9147 | -0.24694 | 0.094166 | -2.62234 | 0.008733 | 0.030383 |
| CNAG_00861 | 249.8687 | -0.32842 | 0.133701 | -2.45639 | 0.014034 | 0.044685 |
| CNAG_02554 | 388.9578 | -0.2882 | 0.117657 | -2.44948 | 0.014306 | 0.045386 |
| CNAG_02552 | 330.9944 | -0.43768 | 0.125875 | -3.4771 | 0.000507 | 0.002641 |
| CNAG_02549 | 764.5247 | 0.444945 | 0.090099 | 4.93842 | 7.88E-07 | 6.96E-06 |
| CNAG_02548 | 1082.799 | 0.279565 | 0.118179 | 2.365618 | 0.018 | 0.054616 |
| CNAG_02546 | 3496.789 | -0.19871 | 0.076899 | -2.58406 | 0.009765 | 0.033189 |
| CNAG_02545 | 15013.96 | 0.146187 | 0.069131 | 2.11463 | 0.034462 | 0.092355 |
| CNAG_02543 | 1032.139 | 0.410202 | 0.091119 | 4.501844 | 6.74E-06 | 5.23E-05 |
| CNAG_02542 | 2618.683 | 1.48869 | 0.088325 | 16.85465 | 9.70E-64 | 4.20E-61 |
| CNAG_02541 | 3409.298 | 0.623788 | 0.067017 | 9.307889 | 1.30E-20 | 4.99E-19 |
| CNAG_02540 | 688.8213 | -0.40105 | 0.093875 | -4.27222 | 1.94E-05 | 0.000137 |
| CNAG_02539 | 1227.624 | 0.258311 | 0.083762 | 3.083881 | 0.002043 | 0.008756 |
| CNAG_02534 | 1246.734 | -0.22123 | 0.079802 | -2.77228 | 0.005567 | 0.020569 |
| CNAG_02529 | 755.4265 | 0.198711 | 0.093932 | 2.115472 | 0.03439 | 0.092246 |
| CNAG_02528 | 1567.652 | 0.60384 | 0.106171 | 5.687457 | 1.29E-08 | 1.50E-07 |
| CNAG_02527 | 3830.576 | 0.673237 | 0.068603 | 9.813553 | 9.84E-23 | 4.47E-21 |
| CNAG_02524 | 748.6166 | 0.484759 | 0.098247 | 4.934091 | 8.05E-07 | 7.09E-06 |
| CNAG_02520 | 1426.413 | -0.47371 | 0.07849 | -6.0353 | 1.59E-09 | 2.06E-08 |
| CNAG_02519 | 1640.31 | 0.33857 | 0.083235 | 4.067653 | 4.75E-05 | 0.000314 |
| CNAG_02517 | 88.7999 | 0.558771 | 0.216857 | 2.576684 | 0.009975 | 0.033773 |
| CNAG_02515 | 525.7487 | 0.706596 | 0.106012 | 6.665253 | 2.64E-11 | 4.27E-10 |
| CNAG_02512 | 1030.751 | -0.29785 | 0.087477 | -3.40488 | 0.000662 | 0.003328 |
| CNAG_02510 | 1276.392 | 0.215684 | 0.08896 | 2.42451 | 0.015329 | 0.048075 |

| | | | | | | |
|---|---|---|---|---|---|---|
| CNAG_02508 | 214.0494 | -0.60189 | 0.155231 | -3.87738 | 0.000106 | 0.000647 |
| CNAG_02507 | 4320.903 | -0.29216 | 0.078356 | -3.72865 | 0.000193 | 0.001104 |
| CNAG_02505 | 500.5902 | 0.62957 | 0.122064 | 5.157704 | 2.50E-07 | 2.40E-06 |
| CNAG_02503 | 1409.788 | -0.20811 | 0.084153 | -2.47304 | 0.013397 | 0.043014 |
| CNAG_02502 | 3183.098 | -0.66038 | 0.067124 | -9.83812 | 7.71E-23 | 3.58E-21 |
| CNAG_02500 | 7144.12 | -0.57215 | 0.065215 | -8.77332 | 1.73E-18 | 5.39E-17 |
| CNAG_02495 | 2249.649 | -0.17485 | 0.069682 | -2.50926 | 0.012099 | 0.039604 |
| CNAG_02491 | 628.596 | -0.2251 | 0.097067 | -2.31907 | 0.020391 | 0.060543 |
| CNAG_02490 | 764.3127 | -0.23252 | 0.108508 | -2.14286 | 0.032124 | 0.087542 |
| CNAG_07625 | 2918.979 | 0.207713 | 0.066101 | 3.142365 | 0.001676 | 0.007406 |
| CNAG_12527 | 38.11968 | 0.714934 | 0.318357 | 2.245699 | 0.024723 | 0.07096 |
| CNAG_02479 | 387.6302 | 0.395404 | 0.124144 | 3.185056 | 0.001447 | 0.006536 |
| CNAG_07626 | 170.6878 | -0.35685 | 0.156821 | -2.27552 | 0.022875 | 0.066487 |
| CNAG_02473 | 776.1054 | 0.576334 | 0.0947 | 6.085863 | 1.16E-09 | 1.52E-08 |
| CNAG_02469 | 1457.806 | 0.179309 | 0.085248 | 2.103373 | 0.035433 | 0.094505 |
| CNAG_02468 | 2110.48 | 0.160393 | 0.070015 | 2.29084 | 0.021973 | 0.064295 |
| CNAG_02467 | 2427.308 | 0.141501 | 0.070569 | 2.00514 | 0.044948 | 0.114595 |
| CNAG_02464 | 621.0067 | -0.27162 | 0.100308 | -2.70783 | 0.006772 | 0.024399 |
| CNAG_07627 | 1173.909 | -0.2385 | 0.102152 | -2.33475 | 0.019557 | 0.058471 |
| CNAG_02460 | 469.1498 | 0.772833 | 0.13207 | 5.851678 | 4.87E-09 | 5.98E-08 |
| CNAG_02457 | 4427.063 | -0.29457 | 0.088208 | -3.33955 | 0.000839 | 0.004078 |
| CNAG_07628 | 11866.85 | -0.14152 | 0.059913 | -2.36206 | 0.018173 | 0.054993 |
| CNAG_12536 | 10.18043 | 1.90617 | 0.68073 | 2.800183 | 0.005107 | 0.019135 |
| CNAG_02455 | 793.5454 | 0.248978 | 0.102281 | 2.434247 | 0.014923 | 0.047035 |
| CNAG_02454 | 1181.501 | -0.7168 | 0.098023 | -7.31255 | 2.62E-13 | 5.33E-12 |
| CNAG_02453 | 1140.542 | -0.26702 | 0.094743 | -2.81836 | 0.004827 | 0.018269 |
| CNAG_07629 | 1830.506 | 0.68814 | 0.072358 | 9.510201 | 1.90E-21 | 7.82E-20 |
| CNAG_07630 | 1324.121 | -0.71075 | 0.096408 | -7.37233 | 1.68E-13 | 3.47E-12 |
| CNAG_02447 | 1002.188 | 0.530851 | 0.081768 | 6.492178 | 8.46E-11 | 1.29E-09 |
| CNAG_02445 | 2719.38 | 0.400722 | 0.070077 | 5.718354 | 1.08E-08 | 1.26E-07 |
| CNAG_02444 | 521.7839 | 0.527213 | 0.115009 | 4.584114 | 4.56E-06 | 3.63E-05 |
| CNAG_02443 | 3156.824 | 0.236095 | 0.06609 | 3.572313 | 0.000354 | 0.001911 |
| CNAG_02442 | 534.0159 | -0.40486 | 0.103169 | -3.92429 | 8.70E-05 | 0.000544 |
| CNAG_02438 | 241.9203 | 0.548556 | 0.148827 | 3.685868 | 0.000228 | 0.001293 |
| CNAG_12538 | 51.72359 | 0.575237 | 0.274111 | 2.098552 | 0.035856 | 0.095405 |
| CNAG_02437 | 1265.784 | -0.56316 | 0.083497 | -6.74466 | 1.53E-11 | 2.55E-10 |
| CNAG_02435 | 4088.462 | 0.452242 | 0.075011 | 6.028979 | 1.65E-09 | 2.14E-08 |
| CNAG_02433 | 923.149 | 0.287019 | 0.084008 | 3.416561 | 0.000634 | 0.00321 |
| CNAG_02430 | 409.9636 | 0.29831 | 0.121768 | 2.449814 | 0.014293 | 0.045379 |
| CNAG_02429 | 101.4344 | -0.40988 | 0.20259 | -2.02322 | 0.04305 | 0.11077 |
| CNAG_02426 | 4298.731 | -0.20724 | 0.065931 | -3.14327 | 0.001671 | 0.00739 |
| CNAG_02417 | 233.2614 | 0.927914 | 0.178974 | 5.184625 | 2.16E-07 | 2.10E-06 |
| CNAG_02416 | 3826.16 | 0.191812 | 0.073708 | 2.602346 | 0.009259 | 0.031817 |
| CNAG_02412 | 756.4628 | -0.32822 | 0.092827 | -3.53579 | 0.000407 | 0.002166 |
| CNAG_07631 | 2467.764 | 0.549236 | 0.103921 | 5.285124 | 1.26E-07 | 1.25E-06 |
| CNAG_02409 | 592.095 | 0.415205 | 0.103236 | 4.021892 | 5.77E-05 | 0.000376 |
| CNAG_02407 | 1843.488 | 0.641388 | 0.078741 | 8.145514 | 3.78E-16 | 9.48E-15 |
| CNAG_02406 | 1388.136 | 0.297981 | 0.087164 | 3.418624 | 0.000629 | 0.003189 |

| | | | | | | |
|---|---|---|---|---|---|---|
| CNAG_02404 | 6238.892 | -0.28979 | 0.069898 | -4.14593 | 3.38E-05 | 0.000229 |
| CNAG_02403 | 3513.738 | 0.167382 | 0.073323 | 2.282787 | 0.022443 | 0.065328 |
| CNAG_02400 | 2145.726 | -0.21158 | 0.08569 | -2.46918 | 0.013542 | 0.043402 |
| CNAG_02399 | 6587.449 | 0.192491 | 0.068013 | 2.8302 | 0.004652 | 0.017744 |
| CNAG_02398 | 1305.849 | 0.634399 | 0.112137 | 5.657341 | 1.54E-08 | 1.76E-07 |
| CNAG_02395 | 1397.24 | 0.205302 | 0.078971 | 2.599728 | 0.00933 | 0.031991 |
| CNAG_02392 | 231.3547 | 0.406115 | 0.143483 | 2.830406 | 0.004649 | 0.017741 |
| CNAG_02389 | 287.502 | -0.38398 | 0.132047 | -2.90791 | 0.003639 | 0.014485 |
| CNAG_02386 | 1190.539 | 1.121094 | 0.092663 | 12.09857 | 1.07E-33 | 9.53E-32 |
| CNAG_02385 | 982.3982 | 0.490129 | 0.08927 | 5.490405 | 4.01E-08 | 4.30E-07 |
| CNAG_02382 | 3245.718 | -0.47687 | 0.070061 | -6.80651 | 1.00E-11 | 1.70E-10 |
| CNAG_07985 | 156.4244 | -0.5488 | 0.183917 | -2.98397 | 0.002845 | 0.011712 |
| CNAG_07986 | 615.3856 | -0.39604 | 0.1065 | -3.71868 | 0.0002 | 0.001144 |
| CNAG_02378 | 7860.043 | -0.45091 | 0.067006 | -6.7294 | 1.70E-11 | 2.83E-10 |
| CNAG_02376 | 1396.824 | 0.182063 | 0.084092 | 2.165044 | 0.030384 | 0.083966 |
| CNAG_02373 | 1285.016 | 0.215726 | 0.079063 | 2.728549 | 0.006361 | 0.02309 |
| CNAG_02367 | 6050.739 | -0.13208 | 0.063372 | -2.08422 | 0.03714 | 0.098085 |
| CNAG_02363 | 124.694 | 0.389948 | 0.186779 | 2.087754 | 0.03682 | 0.097306 |
| CNAG_02362 | 844.6499 | 0.613772 | 0.08892 | 6.902519 | 5.11E-12 | 8.92E-11 |
| CNAG_12554 | 129.1349 | 0.701446 | 0.175756 | 3.991032 | 6.58E-05 | 0.000424 |
| CNAG_02361 | 1499.946 | 0.265926 | 0.104308 | 2.549432 | 0.01079 | 0.036077 |
| CNAG_02360 | 1467.886 | 0.415243 | 0.076922 | 5.398243 | 6.73E-08 | 7.00E-07 |
| CNAG_02356 | 1400.787 | 0.197067 | 0.081517 | 2.417489 | 0.015628 | 0.048765 |
| CNAG_02355 | 799.9499 | 0.722103 | 0.096909 | 7.451384 | 9.24E-14 | 1.93E-12 |
| CNAG_02351 | 2411.278 | 0.481625 | 0.081333 | 5.921629 | 3.19E-09 | 3.99E-08 |
| CNAG_02349 | 337.121 | 0.443273 | 0.122817 | 3.609212 | 0.000307 | 0.001681 |
| CNAG_07987 | 587.5234 | 0.377155 | 0.099062 | 3.807259 | 0.000141 | 0.000833 |
| CNAG_02348 | 689.336 | 0.225981 | 0.09537 | 2.369527 | 0.017811 | 0.054147 |
| CNAG_02347 | 175.3842 | 0.754826 | 0.169316 | 4.45808 | 8.27E-06 | 6.33E-05 |
| CNAG_02343 | 1099.914 | -0.25421 | 0.0794 | -3.20158 | 0.001367 | 0.006227 |
| CNAG_02338 | 23141.27 | -0.1991 | 0.081629 | -2.43914 | 0.014722 | 0.046515 |
| CNAG_02337 | 1251.738 | -0.39143 | 0.09609 | -4.07353 | 4.63E-05 | 0.000307 |
| CNAG_02336 | 414.1657 | -0.38809 | 0.116708 | -3.3253 | 0.000883 | 0.004268 |
| CNAG_02335 | 12270.69 | 0.727238 | 0.075632 | 9.615479 | 6.88E-22 | 2.91E-20 |
| CNAG_02334 | 20.53513 | -0.97727 | 0.444569 | -2.19825 | 0.027931 | 0.078803 |
| CNAG_02331 | 27557.86 | -0.30203 | 0.078983 | -3.824 | 0.000131 | 0.000785 |
| CNAG_02330 | 39405.38 | -0.33263 | 0.067682 | -4.91463 | 8.90E-07 | 7.77E-06 |
| CNAG_02328 | 3281.981 | -0.69135 | 0.086609 | -7.9824 | 1.44E-15 | 3.46E-14 |
| CNAG_02325 | 962.9515 | 0.437625 | 0.084369 | 5.187047 | 2.14E-07 | 2.07E-06 |
| CNAG_02324 | 875.0951 | -0.3561 | 0.095249 | -3.73867 | 0.000185 | 0.001064 |
| CNAG_02323 | 2329.587 | -0.40796 | 0.081432 | -5.00985 | 5.45E-07 | 4.98E-06 |
| CNAG_12560 | 26.32973 | -1.11759 | 0.429498 | -2.60208 | 0.009266 | 0.031827 |
| CNAG_02322 | 890.3847 | -0.38513 | 0.102869 | -3.74387 | 0.000181 | 0.001044 |
| CNAG_02319 | 876.1289 | 0.201408 | 0.097736 | 2.060743 | 0.039328 | 0.102853 |
| CNAG_02317 | 1594.343 | -0.47438 | 0.077725 | -6.10333 | 1.04E-09 | 1.38E-08 |
| CNAG_02315 | 18264.68 | 0.283564 | 0.083096 | 3.412474 | 0.000644 | 0.00325 |
| CNAG_02314 | 241.4348 | 0.740812 | 0.139894 | 5.295526 | 1.19E-07 | 1.18E-06 |
| CNAG_02313 | 598.3791 | 0.480218 | 0.101411 | 4.735376 | 2.19E-06 | 1.82E-05 |

| | | | | | | |
|---|---|---|---|---|---|---|
| CNAG_02312 | 977.6016 | 0.517359 | 0.110769 | 4.670598 | 3.00E-06 | 2.45E-05 |
| CNAG_02310 | 1748.718 | -0.26057 | 0.076976 | -3.38507 | 0.000712 | 0.003542 |
| CNAG_02308 | 307.108 | -0.39649 | 0.122049 | -3.24859 | 0.00116 | 0.005391 |
| CNAG_02307 | 648.7002 | -0.25061 | 0.092817 | -2.70003 | 0.006933 | 0.024877 |
| CNAG_02301 | 8878.687 | -0.51914 | 0.065692 | -7.90269 | 2.73E-15 | 6.44E-14 |
| CNAG_02299 | 858.3984 | -0.34105 | 0.099259 | -3.43591 | 0.000591 | 0.003028 |
| CNAG_02298 | 2145.005 | 0.634459 | 0.100979 | 6.283081 | 3.32E-10 | 4.70E-09 |
| CNAG_02292 | 2995.775 | -0.18448 | 0.074981 | -2.46039 | 0.013879 | 0.04428 |
| CNAG_02290 | 1532.167 | -0.20235 | 0.085041 | -2.37944 | 0.017339 | 0.053022 |
| CNAG_02286 | 2032.602 | -0.36119 | 0.09267 | -3.89764 | 9.71E-05 | 0.000599 |
| CNAG_02282 | 1483.688 | 0.328436 | 0.078168 | 4.201657 | 2.65E-05 | 0.000183 |
| CNAG_02281 | 993.742 | -0.29164 | 0.087569 | -3.33044 | 0.000867 | 0.004198 |
| CNAG_07635 | 1906.737 | -0.16002 | 0.077316 | -2.06966 | 0.038485 | 0.10102 |
| CNAG_07638 | 1254.045 | 0.288804 | 0.079765 | 3.620664 | 0.000294 | 0.001619 |
| CNAG_07639 | 1757.702 | 0.459073 | 0.077566 | 5.918496 | 3.25E-09 | 4.06E-08 |
| CNAG_07641 | 151.3432 | 0.325039 | 0.164311 | 1.978193 | 0.047907 | 0.120524 |
| CNAG_07642 | 47.99989 | 0.581951 | 0.281603 | 2.066566 | 0.038775 | 0.101646 |
| CNAG_02270 | 1605.871 | -0.47015 | 0.075432 | -6.23278 | 4.58E-10 | 6.36E-09 |
| CNAG_02269 | 815.3567 | 0.205855 | 0.093784 | 2.194986 | 0.028165 | 0.079292 |
| CNAG_07164 | 851.9811 | 0.716876 | 0.112052 | 6.397718 | 1.58E-10 | 2.33E-09 |
| CNAG_02266 | 5065.643 | 0.307515 | 0.081328 | 3.781168 | 0.000156 | 0.000916 |
| CNAG_02265 | 3050.842 | -0.40542 | 0.100063 | -4.05169 | 5.08E-05 | 0.000334 |
| CNAG_02264 | 1250.742 | 0.682336 | 0.102856 | 6.633871 | 3.27E-11 | 5.18E-10 |
| CNAG_02263 | 645.2181 | 0.906544 | 0.127975 | 7.083753 | 1.40E-12 | 2.62E-11 |
| CNAG_02260 | 2144.418 | -0.3099 | 0.071539 | -4.33187 | 1.48E-05 | 0.000108 |
| CNAG_02259 | 877.2086 | -0.27893 | 0.10226 | -2.72761 | 0.00638 | 0.023146 |
| CNAG_02255 | 81.00185 | -0.45342 | 0.220704 | -2.05444 | 0.039933 | 0.104191 |
| CNAG_02254 | 45.63986 | -1.32662 | 0.306322 | -4.3308 | 1.49E-05 | 0.000109 |
| CNAG_12570 | 42.41651 | -0.92062 | 0.309526 | -2.97429 | 0.002937 | 0.01203 |
| CNAG_02240 | 2407.165 | -0.26435 | 0.092656 | -2.85307 | 0.00433 | 0.016702 |
| CNAG_02239 | 4481.58 | -0.28644 | 0.063328 | -4.52314 | 6.09E-06 | 4.75E-05 |
| CNAG_02237 | 4512.826 | -0.17896 | 0.074714 | -2.39522 | 0.016611 | 0.051115 |
| CNAG_02236 | 2216.072 | -0.34822 | 0.079973 | -4.35427 | 1.34E-05 | 9.89E-05 |
| CNAG_02235 | 789.053 | -0.2218 | 0.092571 | -2.39597 | 0.016576 | 0.05103 |
| CNAG_02234 | 51404.45 | -0.21482 | 0.075252 | -2.85464 | 0.004309 | 0.016637 |
| CNAG_02230 | 4534.038 | 0.815188 | 0.092021 | 8.858697 | 8.10E-19 | 2.59E-17 |
| CNAG_02226 | 194.7955 | 0.511968 | 0.14927 | 3.429815 | 0.000604 | 0.003083 |
| CNAG_02222 | 1023.825 | -0.18583 | 0.082838 | -2.24331 | 0.024877 | 0.071349 |
| CNAG_02220 | 1381.264 | -0.57022 | 0.085553 | -6.66504 | 2.65E-11 | 4.27E-10 |
| CNAG_02215 | 580.9516 | -0.55832 | 0.102436 | -5.45044 | 5.02E-08 | 5.32E-07 |
| CNAG_02214 | 256.1545 | -0.50576 | 0.144579 | -3.49817 | 0.000468 | 0.002467 |
| CNAG_02212 | 13613.57 | -0.16259 | 0.062368 | -2.60693 | 0.009136 | 0.031491 |
| CNAG_02211 | 482.441 | 0.403929 | 0.106817 | 3.781504 | 0.000156 | 0.000915 |
| CNAG_02209 | 9034.855 | -0.22752 | 0.067783 | -3.35655 | 0.000789 | 0.003863 |
| CNAG_02208 | 5545.647 | 0.287138 | 0.061444 | 4.673166 | 2.97E-06 | 2.42E-05 |
| CNAG_02207 | 382.9895 | -0.37845 | 0.120805 | -3.1327 | 0.001732 | 0.007628 |
| CNAG_02205 | 490.8291 | 0.336656 | 0.104019 | 3.236491 | 0.00121 | 0.005595 |
| CNAG_02199 | 331.2246 | 0.327482 | 0.119518 | 2.740022 | 0.006143 | 0.022425 |

| | | | | | | |
|---|---|---|---|---|---|---|
| CNAG_02191 | 573.5862 | -0.29754 | 0.101887 | -2.9203 | 0.003497 | 0.014024 |
| CNAG_02190 | 832.7288 | 0.848452 | 0.08616 | 9.847438 | 7.03E-23 | 3.31E-21 |
| CNAG_02189 | 4535.793 | 0.221479 | 0.082246 | 2.692898 | 0.007083 | 0.025311 |
| CNAG_02188 | 1220.254 | 0.976406 | 0.086195 | 11.32783 | 9.55E-30 | 6.72E-28 |
| CNAG_02185 | 961.5596 | -0.44042 | 0.08691 | -5.06755 | 4.03E-07 | 3.74E-06 |
| CNAG_02182 | 2522.66 | 1.296244 | 0.083332 | 15.55516 | 1.47E-54 | 3.95E-52 |
| CNAG_02179 | 1817.523 | 0.263306 | 0.074087 | 3.554 | 0.000379 | 0.002031 |
| CNAG_02176 | 1748.464 | -0.35201 | 0.099044 | -3.55409 | 0.000379 | 0.002031 |
| CNAG_02169 | 254.5684 | 0.49318 | 0.170356 | 2.894988 | 0.003792 | 0.014998 |
| CNAG_02165 | 1434.808 | 0.302959 | 0.081208 | 3.730652 | 0.000191 | 0.001096 |
| CNAG_02164 | 1154.912 | 0.613449 | 0.092787 | 6.611393 | 3.81E-11 | 5.98E-10 |
| CNAG_02163 | 394.0101 | -0.35558 | 0.119261 | -2.98156 | 0.002868 | 0.011785 |
| CNAG_02162 | 878.3634 | -0.29398 | 0.09517 | -3.08905 | 0.002008 | 0.008639 |
| CNAG_02159 | 378.379 | -0.34213 | 0.121168 | -2.82365 | 0.004748 | 0.018066 |
| CNAG_07647 | 1290.323 | 0.318247 | 0.085082 | 3.740478 | 0.000184 | 0.001057 |
| CNAG_12585 | 78.52839 | -0.65609 | 0.231734 | -2.83121 | 0.004637 | 0.017705 |
| CNAG_02154 | 862.9533 | -0.25585 | 0.085857 | -2.97995 | 0.002883 | 0.011841 |
| CNAG_02153 | 2404.365 | -0.28241 | 0.072038 | -3.92025 | 8.85E-05 | 0.000552 |
| CNAG_02144 | 52882.16 | -0.21858 | 0.063722 | -3.43027 | 0.000603 | 0.003081 |
| CNAG_02143 | 586.9729 | -0.47727 | 0.102257 | -4.66734 | 3.05E-06 | 2.48E-05 |
| CNAG_02140 | 4476.387 | 0.132688 | 0.064217 | 2.066238 | 0.038806 | 0.101682 |
| CNAG_12586 | 62.74024 | 0.698966 | 0.256938 | 2.720372 | 0.006521 | 0.023603 |
| CNAG_02134 | 3223.929 | -0.18719 | 0.072822 | -2.57051 | 0.010155 | 0.034263 |
| CNAG_02133 | 6386.399 | 0.285299 | 0.072338 | 3.943952 | 8.01E-05 | 0.000507 |
| CNAG_02131 | 1111.878 | 0.328216 | 0.084465 | 3.885804 | 0.000102 | 0.000628 |
| CNAG_02129 | 20051.59 | 1.156025 | 0.078279 | 14.76809 | 2.35E-49 | 4.59E-47 |
| CNAG_02128 | 9576.167 | -0.33658 | 0.06738 | -4.99526 | 5.88E-07 | 5.32E-06 |
| CNAG_02127 | 890.3864 | 0.229278 | 0.091935 | 2.493927 | 0.012634 | 0.041132 |
| CNAG_02124 | 844.7909 | 0.27006 | 0.092802 | 2.910057 | 0.003614 | 0.014418 |
| CNAG_02122 | 705.4632 | -0.43482 | 0.093406 | -4.65515 | 3.24E-06 | 2.62E-05 |
| CNAG_02120 | 2583.417 | -0.24213 | 0.075808 | -3.19405 | 0.001403 | 0.006358 |
| CNAG_02116 | 493.508 | 0.624455 | 0.108281 | 5.766987 | 8.07E-09 | 9.56E-08 |
| CNAG_02115 | 1827.522 | -0.34916 | 0.0727 | -4.80279 | 1.56E-06 | 1.33E-05 |
| CNAG_02111 | 1063.627 | -0.2963 | 0.081363 | -3.64173 | 0.000271 | 0.001511 |
| CNAG_02110 | 2869.773 | -0.18853 | 0.067731 | -2.78359 | 0.005376 | 0.019969 |
| CNAG_02107 | 598.8872 | 0.232506 | 0.101139 | 2.298866 | 0.021513 | 0.063233 |
| CNAG_02106 | 1730.963 | 0.490804 | 0.078072 | 6.286573 | 3.25E-10 | 4.61E-09 |
| CNAG_02105 | 260.293 | 0.315477 | 0.143879 | 2.192649 | 0.028333 | 0.079655 |
| CNAG_02104 | 548.2765 | -0.50604 | 0.105873 | -4.77971 | 1.76E-06 | 1.48E-05 |
| CNAG_12593 | 93.14421 | 0.536832 | 0.209723 | 2.559719 | 0.010476 | 0.035147 |
| CNAG_02101 | 571.9663 | -0.21498 | 0.102004 | -2.10757 | 0.035068 | 0.093691 |
| CNAG_02100 | 13098.71 | 0.181897 | 0.063206 | 2.877839 | 0.004004 | 0.015702 |
| CNAG_02096 | 322.4905 | 0.288228 | 0.124428 | 2.31642 | 0.020535 | 0.060865 |
| CNAG_02091 | 13537.5 | -0.19878 | 0.069298 | -2.86851 | 0.004124 | 0.01606 |
| CNAG_02089 | 518.5622 | 0.380031 | 0.10973 | 3.46334 | 0.000534 | 0.002764 |
| CNAG_02088 | 1260.637 | -0.15876 | 0.076278 | -2.08127 | 0.037409 | 0.098561 |
| CNAG_02087 | 146.3367 | 0.38855 | 0.170845 | 2.274286 | 0.022949 | 0.066676 |
| CNAG_02085 | 900.8593 | 0.371679 | 0.096996 | 3.831906 | 0.000127 | 0.000764 |

| | | | | | | |
|---|---|---|---|---|---|---|
| CNAG_02084 | 3980.294 | -0.44648 | 0.066832 | -6.68068 | 2.38E-11 | 3.89E-10 |
| CNAG_02083 | 181.3465 | -1.31559 | 0.158573 | -8.29644 | 1.07E-16 | 2.86E-15 |
| CNAG_02081 | 3145.666 | -0.17396 | 0.074214 | -2.34397 | 0.01908 | 0.057291 |
| CNAG_02077 | 544.2306 | -0.21067 | 0.105587 | -1.99521 | 0.04602 | 0.116557 |
| CNAG_02076 | 1558.739 | -0.21411 | 0.079265 | -2.70119 | 0.006909 | 0.024813 |
| CNAG_02074 | 2447.975 | 0.369951 | 0.071511 | 5.173374 | 2.30E-07 | 2.22E-06 |
| CNAG_02073 | 1260.055 | 0.213634 | 0.076185 | 2.804151 | 0.005045 | 0.018955 |
| CNAG_02072 | 1003.271 | 0.51387 | 0.091463 | 5.618302 | 1.93E-08 | 2.16E-07 |
| CNAG_02071 | 2370.468 | 0.16323 | 0.082637 | 1.975271 | 0.048237 | 0.121043 |
| CNAG_02070 | 6391.391 | 1.069469 | 0.092667 | 11.54095 | 8.20E-31 | 6.10E-29 |
| CNAG_02069 | 1774.233 | 0.710899 | 0.093195 | 7.628057 | 2.38E-14 | 5.20E-13 |
| CNAG_02068 | 1083.777 | -0.23458 | 0.084114 | -2.78881 | 0.00529 | 0.019725 |
| CNAG_02064 | 3383.505 | -0.22568 | 0.094782 | -2.38102 | 0.017265 | 0.052837 |
| CNAG_02063 | 1867.336 | 0.234632 | 0.089596 | 2.618776 | 0.008825 | 0.030609 |
| CNAG_02062 | 508.4946 | -0.23445 | 0.109577 | -2.13955 | 0.032391 | 0.088108 |
| CNAG_02061 | 390.4245 | 0.269202 | 0.11644 | 2.311944 | 0.020781 | 0.061522 |
| CNAG_12600 | 426.1867 | 0.742344 | 0.134852 | 5.504886 | 3.69E-08 | 4.00E-07 |
| CNAG_02058 | 46.18096 | 0.865983 | 0.297491 | 2.910953 | 0.003603 | 0.014391 |
| CNAG_02057 | 993.1108 | 0.340239 | 0.090248 | 3.770065 | 0.000163 | 0.00095 |
| CNAG_02055 | 965.9074 | 0.316575 | 0.083084 | 3.810305 | 0.000139 | 0.000823 |
| CNAG_02054 | 654.5754 | -0.26816 | 0.104472 | -2.56679 | 0.010265 | 0.034543 |
| CNAG_02050 | 609.0284 | -0.25018 | 0.110432 | -2.26546 | 0.023484 | 0.067853 |
| CNAG_02048 | 748.8377 | 0.322541 | 0.09063 | 3.558869 | 0.000372 | 0.001999 |
| CNAG_02047 | 926.577 | 0.678949 | 0.088486 | 7.672913 | 1.68E-14 | 3.72E-13 |
| CNAG_02045 | 93.70904 | -0.63655 | 0.221839 | -2.86945 | 0.004112 | 0.016021 |
| CNAG_02043 | 165.6226 | -0.92614 | 0.163644 | -5.65947 | 1.52E-08 | 1.74E-07 |
| CNAG_02041 | 2003.565 | 1.167169 | 0.100686 | 11.59221 | 4.51E-31 | 3.45E-29 |
| CNAG_02040 | 140.898 | -0.75932 | 0.196272 | -3.86873 | 0.000109 | 0.000667 |
| CNAG_02039 | 68.68623 | 0.623483 | 0.240503 | 2.59241 | 0.009531 | 0.032552 |
| CNAG_02038 | 2950.996 | 0.65589 | 0.07336 | 8.940749 | 3.87E-19 | 1.28E-17 |
| CNAG_02037 | 3112.938 | 0.162281 | 0.079737 | 2.03519 | 0.041832 | 0.108277 |
| CNAG_02035 | 16786.99 | 0.35149 | 0.067187 | 5.231535 | 1.68E-07 | 1.65E-06 |
| CNAG_02028 | 2011.45 | 0.713041 | 0.074016 | 9.633656 | 5.76E-22 | 2.46E-20 |
| CNAG_02025 | 1423.796 | -0.37794 | 0.075599 | -4.99921 | 5.76E-07 | 5.24E-06 |
| CNAG_02024 | 1600.42 | -0.29299 | 0.097198 | -3.01433 | 0.002575 | 0.01072 |
| CNAG_02023 | 139.8836 | 0.631054 | 0.169625 | 3.720296 | 0.000199 | 0.001138 |
| CNAG_02022 | 2089.792 | 0.218706 | 0.074626 | 2.930679 | 0.003382 | 0.013599 |
| CNAG_02020 | 1570.867 | 0.218599 | 0.085186 | 2.566132 | 0.010284 | 0.034593 |
| CNAG_07651 | 1009.974 | -0.49463 | 0.084421 | -5.85906 | 4.65E-09 | 5.73E-08 |
| CNAG_06524 | 180.7634 | -0.66581 | 0.204003 | -3.26374 | 0.0011 | 0.005151 |
| CNAG_06526 | 2500.607 | 0.14881 | 0.074962 | 1.985128 | 0.04713 | 0.1188 |
| CNAG_06529 | 73.14392 | -0.49765 | 0.230096 | -2.16277 | 0.030559 | 0.084387 |
| CNAG_06530 | 508.7547 | -0.34517 | 0.111157 | -3.10528 | 0.001901 | 0.008256 |
| CNAG_06532 | 640.8504 | 0.328346 | 0.108075 | 3.03812 | 0.002381 | 0.009983 |
| CNAG_06533 | 1493.405 | 0.597956 | 0.082608 | 7.238457 | 4.54E-13 | 8.92E-12 |
| CNAG_06534 | 7160.439 | -0.25024 | 0.066137 | -3.78368 | 0.000155 | 0.000908 |
| CNAG_06535 | 2392.762 | -0.64552 | 0.079108 | -8.15996 | 3.35E-16 | 8.44E-15 |
| CNAG_06536 | 306.985 | -0.42317 | 0.131592 | -3.2158 | 0.001301 | 0.005958 |

| | | | | | | |
|---|---|---|---|---|---|---|
| CNAG_06540 | 125.8116 | 0.437386 | 0.180675 | 2.42084 | 0.015485 | 0.048434 |
| CNAG_06541 | 7050.335 | 0.817709 | 0.078539 | 10.41154 | 2.20E-25 | 1.24E-23 |
| CNAG_06542 | 1577.414 | -0.15102 | 0.074412 | -2.02948 | 0.042409 | 0.109567 |
| CNAG_06545 | 4770.042 | -0.24446 | 0.062953 | -3.88323 | 0.000103 | 0.000634 |
| CNAG_06546 | 42.99951 | -0.71091 | 0.318439 | -2.23248 | 0.025583 | 0.073079 |
| CNAG_07657 | 1343.008 | -0.77642 | 0.085857 | -9.04318 | 1.52E-19 | 5.30E-18 |
| CNAG_12613 | 9.183343 | 1.536503 | 0.693037 | 2.217056 | 0.026619 | 0.075541 |
| CNAG_06550 | 2972.414 | -0.20492 | 0.096618 | -2.1209 | 0.03393 | 0.09125 |
| CNAG_06552 | 1584.69 | 0.250894 | 0.078423 | 3.199245 | 0.001378 | 0.006259 |
| CNAG_06553 | 1030.977 | 0.415336 | 0.083775 | 4.957759 | 7.13E-07 | 6.34E-06 |
| CNAG_06554 | 630.7109 | 0.471259 | 0.099584 | 4.732268 | 2.22E-06 | 1.85E-05 |
| CNAG_06555 | 1281.277 | 0.266281 | 0.076371 | 3.486678 | 0.000489 | 0.00256 |
| CNAG_06556 | 61.63709 | 1.280426 | 0.28243 | 4.533599 | 5.80E-06 | 4.53E-05 |
| CNAG_06557 | 56.89858 | 1.308075 | 0.267975 | 4.881327 | 1.05E-06 | 9.12E-06 |
| CNAG_06559 | 642.0318 | 0.553941 | 0.102866 | 5.385078 | 7.24E-08 | 7.47E-07 |
| CNAG_06560 | 1059.686 | 0.465648 | 0.090606 | 5.139245 | 2.76E-07 | 2.63E-06 |
| CNAG_06567 | 1593.101 | 0.246721 | 0.087925 | 2.806048 | 0.005015 | 0.018862 |
| CNAG_07658 | 30.5975 | 0.901755 | 0.362005 | 2.491 | 0.012738 | 0.041404 |
| CNAG_12616 | 115.7941 | 0.586579 | 0.192035 | 3.054537 | 0.002254 | 0.009529 |
| CNAG_06569 | 963.2499 | 0.402591 | 0.092512 | 4.351756 | 1.35E-05 | 1.00E-04 |
| CNAG_06570 | 459.7296 | 0.231448 | 0.105118 | 2.201803 | 0.027679 | 0.078184 |
| CNAG_07660 | 13900.43 | 0.661613 | 0.075768 | 8.732074 | 2.50E-18 | 7.62E-17 |
| CNAG_06576 | 4231.534 | 1.991892 | 0.099067 | 20.10647 | 6.48E-90 | 1.01E-86 |
| CNAG_06577 | 1210.814 | 1.862925 | 0.113585 | 16.40109 | 1.88E-60 | 6.66E-58 |
| CNAG_06578 | 817.5768 | 0.807999 | 0.088044 | 9.177267 | 4.42E-20 | 1.62E-18 |
| CNAG_07989 | 879.1577 | -0.22981 | 0.090034 | -2.55243 | 0.010698 | 0.035815 |
| CNAG_12622 | 60.7824 | 0.957376 | 0.268531 | 3.565229 | 0.000364 | 0.001957 |
| CNAG_06583 | 363.4116 | 1.109171 | 0.143626 | 7.722635 | 1.14E-14 | 2.56E-13 |
| CNAG_06584 | 764.7529 | 0.180609 | 0.087572 | 2.062415 | 0.039168 | 0.10247 |
| CNAG_12623 | 24.56038 | 0.878721 | 0.40867 | 2.150199 | 0.031539 | 0.086326 |
| CNAG_06587 | 828.9179 | -0.21741 | 0.093124 | -2.33463 | 0.019563 | 0.058471 |
| CNAG_06589 | 922.6184 | 0.330248 | 0.090533 | 3.647799 | 0.000264 | 0.001481 |
| CNAG_06591 | 4236.808 | -0.45181 | 0.112732 | -4.00778 | 6.13E-05 | 0.000398 |
| CNAG_06594 | 6487.325 | 0.261083 | 0.070293 | 3.714215 | 0.000204 | 0.001162 |
| CNAG_06595 | 279.5733 | -0.54976 | 0.180466 | -3.04636 | 0.002316 | 0.009755 |
| CNAG_06596 | 202.6582 | -0.37299 | 0.162514 | -2.2951 | 0.021727 | 0.06378 |
| CNAG_06597 | 2087.503 | -0.31054 | 0.070508 | -4.40437 | 1.06E-05 | 7.98E-05 |
| CNAG_06598 | 48.88767 | -1.00183 | 0.323983 | -3.09225 | 0.001986 | 0.008565 |
| CNAG_06599 | 878.4672 | -0.21987 | 0.094916 | -2.31649 | 0.020532 | 0.060865 |
| CNAG_12627 | 104.6776 | 0.460991 | 0.19851 | 2.322256 | 0.020219 | 0.060134 |
| CNAG_06602 | 2031.225 | 0.783212 | 0.093287 | 8.395761 | 4.63E-17 | 1.25E-15 |
| CNAG_06603 | 1881.633 | -0.16713 | 0.084055 | -1.98836 | 0.046772 | 0.118012 |
| CNAG_06605 | 62280.5 | -0.20584 | 0.077806 | -2.6455 | 0.008157 | 0.028765 |
| CNAG_06606 | 2124.936 | 0.169016 | 0.080334 | 2.103917 | 0.035386 | 0.09441 |
| CNAG_06608 | 1850.697 | 0.304525 | 0.07353 | 4.141497 | 3.45E-05 | 0.000233 |
| CNAG_12629 | 113.334 | -0.52224 | 0.193266 | -2.70219 | 0.006888 | 0.024762 |
| CNAG_06610 | 186.9565 | -0.35095 | 0.153635 | -2.2843 | 0.022354 | 0.065165 |
| CNAG_06611 | 2492.229 | -0.49448 | 0.194323 | -2.54461 | 0.01094 | 0.03647 |

| | | | | | | |
|---|---|---|---|---|---|---|
| CNAG_06612 | 1909.827 | 0.215038 | 0.077637 | 2.769769 | 0.00561 | 0.020689 |
| CNAG_12632 | 53.28428 | 0.548853 | 0.266576 | 2.058899 | 0.039504 | 0.103279 |
| CNAG_06616 | 1106.982 | 0.400016 | 0.091676 | 4.363372 | 1.28E-05 | 9.51E-05 |
| CNAG_06621 | 6197.849 | 0.61271 | 0.070698 | 8.666628 | 4.45E-18 | 1.32E-16 |
| CNAG_06622 | 179.2671 | 1.189856 | 0.163294 | 7.286606 | 3.18E-13 | 6.36E-12 |
| CNAG_06624 | 1609.716 | 0.406305 | 0.083485 | 4.866787 | 1.13E-06 | 9.79E-06 |
| CNAG_06626 | 4064.963 | -0.44313 | 0.06686 | -6.6278 | 3.41E-11 | 5.37E-10 |
| CNAG_06628 | 2023.88 | -0.2787 | 0.071297 | -3.90898 | 9.27E-05 | 0.000574 |
| CNAG_06629 | 21.76639 | -0.89333 | 0.43144 | -2.07058 | 0.038398 | 0.100828 |
| CNAG_06630 | 4852.554 | 0.293614 | 0.066231 | 4.4332 | 9.28E-06 | 7.04E-05 |
| CNAG_06632 | 1634.807 | 0.502873 | 0.078877 | 6.375445 | 1.82E-10 | 2.67E-09 |
| CNAG_06633 | 42259.19 | -0.26637 | 0.069608 | -3.82671 | 0.00013 | 0.000779 |
| CNAG_06635 | 3476.901 | -0.18039 | 0.083269 | -2.16633 | 0.030286 | 0.083723 |
| CNAG_06638 | 4216.584 | -0.54636 | 0.090661 | -6.02642 | 1.68E-09 | 2.17E-08 |
| CNAG_12642 | 41.11419 | 0.737555 | 0.305528 | 2.414033 | 0.015777 | 0.049114 |
| CNAG_06640 | 3439.61 | 0.391469 | 0.067308 | 5.816055 | 6.03E-09 | 7.26E-08 |
| CNAG_06641 | 1800.465 | -0.39042 | 0.074287 | -5.25556 | 1.48E-07 | 1.45E-06 |
| CNAG_06643 | 557.3201 | -0.2188 | 0.100075 | -2.18639 | 0.028787 | 0.080637 |
| CNAG_06644 | 3129.556 | -0.45019 | 0.079921 | -5.63292 | 1.77E-08 | 2.00E-07 |
| CNAG_06645 | 2017.142 | -0.41398 | 0.081848 | -5.05786 | 4.24E-07 | 3.92E-06 |
| CNAG_06646 | 15247.18 | 0.181557 | 0.058246 | 3.11705 | 0.001827 | 0.007968 |
| CNAG_06648 | 2272.497 | -0.13417 | 0.067998 | -1.97317 | 0.048476 | 0.121563 |
| CNAG_06656 | 1685.008 | -0.26942 | 0.087588 | -3.07596 | 0.002098 | 0.008958 |
| CNAG_06657 | 406.992 | 0.545707 | 0.115254 | 4.734803 | 2.19E-06 | 1.82E-05 |
| CNAG_06660 | 822.5904 | -0.268 | 0.093792 | -2.85737 | 0.004272 | 0.016511 |
| CNAG_06661 | 1765.996 | -0.15579 | 0.073186 | -2.12861 | 0.033286 | 0.089961 |
| CNAG_06663 | 8132.698 | 0.283107 | 0.073987 | 3.826431 | 0.00013 | 0.000779 |
| CNAG_06664 | 807.7847 | 0.25703 | 0.087794 | 2.927639 | 0.003415 | 0.013711 |
| CNAG_06668 | 2415.633 | 1.156094 | 0.087025 | 13.28469 | 2.84E-40 | 3.96E-38 |
| CNAG_06675 | 1263.426 | -0.28247 | 0.084863 | -3.32853 | 0.000873 | 0.004224 |
| CNAG_12649 | 39.81374 | -0.64415 | 0.319045 | -2.01898 | 0.043489 | 0.111651 |
| CNAG_06679 | 2912.263 | -0.19687 | 0.072696 | -2.70817 | 0.006766 | 0.024387 |
| CNAG_06680 | 854.3355 | 0.181293 | 0.086675 | 2.091635 | 0.036471 | 0.096645 |
| CNAG_07665 | 4070.482 | -0.19851 | 0.081023 | -2.45008 | 0.014282 | 0.045364 |
| CNAG_06684 | 503.1973 | -0.76713 | 0.109381 | -7.01331 | 2.33E-12 | 4.20E-11 |
| CNAG_06688 | 4819.212 | 0.334444 | 0.061141 | 5.470035 | 4.50E-08 | 4.80E-07 |
| CNAG_06689 | 2033.222 | -0.37145 | 0.069535 | -5.34195 | 9.20E-08 | 9.37E-07 |
| CNAG_06695 | 430.2418 | 0.289109 | 0.117143 | 2.467994 | 0.013587 | 0.043493 |
| CNAG_06697 | 965.8697 | -0.36849 | 0.113971 | -3.23316 | 0.001224 | 0.005647 |
| CNAG_06698 | 1662.213 | 0.297861 | 0.087711 | 3.395927 | 0.000684 | 0.003428 |
| CNAG_12655 | 106.0985 | -0.62732 | 0.198874 | -3.15435 | 0.001609 | 0.007145 |
| CNAG_05673 | 1313.232 | -0.31178 | 0.090812 | -3.43327 | 0.000596 | 0.00305 |
| CNAG_05675 | 443.1445 | -0.35796 | 0.112368 | -3.18563 | 0.001444 | 0.006527 |
| CNAG_05676 | 334.6748 | 0.272464 | 0.119598 | 2.278167 | 0.022717 | 0.066051 |
| CNAG_05678 | 780.6006 | -0.31916 | 0.08961 | -3.56162 | 0.000369 | 0.001981 |
| CNAG_07667 | 5908.232 | -0.24251 | 0.075762 | -3.201 | 0.00137 | 0.006232 |
| CNAG_05682 | 6552.293 | 0.600362 | 0.07164 | 8.38021 | 5.28E-17 | 1.42E-15 |
| CNAG_05683 | 339.4589 | 0.886508 | 0.12174 | 7.281966 | 3.29E-13 | 6.54E-12 |

| | | | | | | |
|---|---|---|---|---|---|---|
| CNAG_05684 | 2047.71 | 0.294889 | 0.070433 | 4.186814 | 2.83E-05 | 0.000194 |
| CNAG_05685 | 1035.257 | 0.183813 | 0.084676 | 2.170788 | 0.029947 | 0.082963 |
| CNAG_05686 | 810.1377 | 0.822931 | 0.087797 | 9.373158 | 7.04E-21 | 2.79E-19 |
| CNAG_05687 | 1395.599 | -0.3542 | 0.097607 | -3.62879 | 0.000285 | 0.001584 |
| CNAG_05695 | 3412.075 | -0.41094 | 0.094894 | -4.33048 | 1.49E-05 | 0.000109 |
| CNAG_05697 | 862.1686 | -0.21351 | 0.090272 | -2.36518 | 0.018021 | 0.054659 |
| CNAG_05698 | 1043.18 | 0.303105 | 0.087029 | 3.482812 | 0.000496 | 0.002592 |
| CNAG_05699 | 930.9575 | 0.321935 | 0.108546 | 2.965883 | 0.003018 | 0.0123 |
| CNAG_05707 | 688.5251 | 0.34364 | 0.101459 | 3.38699 | 0.000707 | 0.003519 |
| CNAG_05708 | 1449.01 | -0.23002 | 0.090217 | -2.5496 | 0.010785 | 0.036076 |
| CNAG_05711 | 919.9199 | 0.205225 | 0.095227 | 2.155117 | 0.031153 | 0.085514 |
| CNAG_05712 | 3642.975 | -0.19964 | 0.090857 | -2.19728 | 0.028001 | 0.078944 |
| CNAG_05714 | 1441.683 | 0.211723 | 0.076283 | 2.775488 | 0.005512 | 0.020396 |
| CNAG_05716 | 645.2976 | -0.34399 | 0.102853 | -3.34448 | 0.000824 | 0.004011 |
| CNAG_05717 | 348.7784 | -0.45674 | 0.126188 | -3.61948 | 0.000295 | 0.001626 |
| CNAG_05719 | 841.3232 | -0.2563 | 0.093807 | -2.73223 | 0.006291 | 0.022845 |
| CNAG_05720 | 1524.443 | -0.21153 | 0.076219 | -2.7753 | 0.005515 | 0.020398 |
| CNAG_05721 | 880.1921 | -0.31032 | 0.095364 | -3.25408 | 0.001138 | 0.005306 |
| CNAG_05722 | 3275.097 | -0.28838 | 0.112297 | -2.56805 | 0.010227 | 0.034447 |
| CNAG_05724 | 1113.083 | 0.362798 | 0.083868 | 4.325833 | 1.52E-05 | 0.000111 |
| CNAG_05725 | 49189.8 | -0.17046 | 0.066957 | -2.54579 | 0.010903 | 0.036363 |
| CNAG_05731 | 1154.365 | -0.17617 | 0.081532 | -2.1608 | 0.030711 | 0.084599 |
| CNAG_05736 | 105.6983 | 0.511122 | 0.201102 | 2.541608 | 0.011034 | 0.036722 |
| CNAG_05737 | 1555.348 | 0.261255 | 0.08008 | 3.262437 | 0.001105 | 0.005171 |
| CNAG_05742 | 1400.167 | -0.19125 | 0.079403 | -2.40855 | 0.016016 | 0.049737 |
| CNAG_05743 | 545.8994 | 0.262397 | 0.100331 | 2.615305 | 0.008915 | 0.030824 |
| CNAG_05745 | 4698.539 | -0.20805 | 0.079959 | -2.60192 | 0.00927 | 0.031828 |
| CNAG_05749 | 780.219 | -0.20431 | 0.096149 | -2.12492 | 0.033593 | 0.090526 |
| CNAG_05750 | 75053.74 | 0.177402 | 0.066133 | 2.682493 | 0.007308 | 0.026028 |
| CNAG_05752 | 2354.429 | -0.22894 | 0.077677 | -2.94728 | 0.003206 | 0.012976 |
| CNAG_07671 | 569.9585 | 0.367868 | 0.12204 | 3.014332 | 0.002575 | 0.01072 |
| CNAG_07672 | 1521.762 | -0.5199 | 0.08484 | -6.12804 | 8.90E-10 | 1.19E-08 |
| CNAG_05755 | 2418.759 | -0.54208 | 0.069273 | -7.82525 | 5.07E-15 | 1.17E-13 |
| CNAG_05756 | 1476.693 | 0.22628 | 0.0826 | 2.739486 | 0.006154 | 0.022446 |
| CNAG_05757 | 839.0873 | -0.96131 | 0.109702 | -8.76293 | 1.90E-18 | 5.87E-17 |
| CNAG_05759 | 12091.17 | 0.19096 | 0.061154 | 3.122631 | 0.001792 | 0.007836 |
| CNAG_05760 | 16.6468 | -1.09706 | 0.504495 | -2.17458 | 0.029662 | 0.082407 |
| CNAG_12676 | 301.5574 | -0.39107 | 0.136734 | -2.86005 | 0.004236 | 0.016405 |
| CNAG_05762 | 62409.78 | -0.22036 | 0.072903 | -3.02266 | 0.002506 | 0.010462 |
| CNAG_05763 | 344.9956 | -0.52288 | 0.118212 | -4.42322 | 9.72E-06 | 7.37E-05 |
| CNAG_05765 | 1402.834 | -0.2575 | 0.078782 | -3.2685 | 0.001081 | 0.005077 |
| CNAG_05768 | 1718.304 | 0.160578 | 0.073039 | 2.198533 | 0.027911 | 0.078777 |
| CNAG_05769 | 1034.339 | 0.173059 | 0.080954 | 2.137741 | 0.032538 | 0.088322 |
| CNAG_05770 | 3805.151 | -0.17039 | 0.065633 | -2.59611 | 0.009429 | 0.032289 |
| CNAG_05771 | 933.132 | -0.31202 | 0.122059 | -2.55632 | 0.010579 | 0.035462 |
| CNAG_05773 | 1141.857 | 0.190784 | 0.083496 | 2.284937 | 0.022317 | 0.065111 |
| CNAG_12677 | 243.4698 | -0.49929 | 0.143482 | -3.47979 | 0.000502 | 0.002616 |
| CNAG_05774 | 1021.767 | 0.503529 | 0.097135 | 5.183789 | 2.17E-07 | 2.10E-06 |

| | | | | | | |
|---|---|---|---|---|---|---|
| CNAG_05777 | 338.495 | -0.26886 | 0.129117 | -2.08228 | 0.037317 | 0.098424 |
| CNAG_05778 | 482.1304 | -0.57275 | 0.114486 | -5.00283 | 5.65E-07 | 5.15E-06 |
| CNAG_05781 | 574.492 | -0.22026 | 0.103217 | -2.13394 | 0.032848 | 0.089012 |
| CNAG_05783 | 3840.613 | 0.197257 | 0.063066 | 3.127807 | 0.001761 | 0.00773 |
| CNAG_05784 | 364.4681 | 0.406498 | 0.120635 | 3.36964 | 0.000753 | 0.003718 |
| CNAG_05789 | 1615.363 | -0.59886 | 0.079059 | -7.57485 | 3.60E-14 | 7.73E-13 |
| CNAG_05791 | 2117.21 | -0.21111 | 0.095499 | -2.21057 | 0.027065 | 0.076695 |
| CNAG_05792 | 4279.627 | -0.24557 | 0.070997 | -3.45888 | 0.000542 | 0.002805 |
| CNAG_05794 | 640.0735 | 0.311108 | 0.099622 | 3.122883 | 0.001791 | 0.007834 |
| CNAG_05799 | 6204.926 | 0.317671 | 0.063096 | 5.034755 | 4.78E-07 | 4.42E-06 |
| CNAG_05800 | 27594.15 | -0.18685 | 0.077491 | -2.41125 | 0.015898 | 0.04943 |
| CNAG_05801 | 470.8109 | -0.39402 | 0.110147 | -3.57718 | 0.000347 | 0.001885 |
| CNAG_05803 | 1454.944 | -0.16002 | 0.074102 | -2.1595 | 0.030812 | 0.084817 |
| CNAG_05808 | 852.5732 | 0.395933 | 0.092533 | 4.27881 | 1.88E-05 | 0.000135 |
| CNAG_05809 | 2798.106 | 0.431173 | 0.072478 | 5.949021 | 2.70E-09 | 3.41E-08 |
| CNAG_05811 | 730.9499 | -0.20873 | 0.094401 | -2.21111 | 0.027028 | 0.076645 |
| CNAG_05814 | 38075.78 | -0.18453 | 0.081422 | -2.26636 | 0.02343 | 0.06772 |
| CNAG_05816 | 1364.318 | -0.22115 | 0.074782 | -2.95732 | 0.003103 | 0.012614 |
| CNAG_05817 | 6055.073 | -0.25628 | 0.066805 | -3.83627 | 0.000125 | 0.000753 |
| CNAG_05820 | 368.2741 | 0.395285 | 0.118168 | 3.34512 | 0.000822 | 0.004004 |
| CNAG_05826 | 6422.358 | -0.20393 | 0.062237 | -3.27672 | 0.00105 | 0.004961 |
| CNAG_05828 | 2179.862 | -0.50232 | 0.082037 | -6.12304 | 9.18E-10 | 1.23E-08 |
| CNAG_05830 | 1374.453 | -0.18569 | 0.088653 | -2.09454 | 0.036211 | 0.096121 |
| CNAG_12684 | 106.3528 | -0.67981 | 0.201677 | -3.37078 | 0.00075 | 0.003705 |
| CNAG_05831 | 309.0704 | -0.41642 | 0.123872 | -3.36169 | 0.000775 | 0.003807 |
| CNAG_05833 | 47.40753 | -0.74629 | 0.302322 | -2.46851 | 0.013568 | 0.043448 |
| CNAG_05836 | 1725.035 | 0.491572 | 0.081716 | 6.015584 | 1.79E-09 | 2.31E-08 |
| CNAG_05839 | 10373.87 | 0.422722 | 0.074918 | 5.642463 | 1.68E-08 | 1.90E-07 |
| CNAG_05840 | 891.6908 | 0.379311 | 0.09553 | 3.970594 | 7.17E-05 | 0.00046 |
| CNAG_05846 | 1220.475 | -0.30869 | 0.086525 | -3.56767 | 0.00036 | 0.001941 |
| CNAG_05847 | 5982.212 | -0.41871 | 0.074391 | -5.62854 | 1.82E-08 | 2.04E-07 |
| CNAG_05848 | 5640.607 | -0.33048 | 0.061239 | -5.3965 | 6.80E-08 | 7.05E-07 |
| CNAG_07676 | 13395.66 | -0.3174 | 0.071244 | -4.45515 | 8.38E-06 | 6.41E-05 |
| CNAG_07677 | 380.7608 | 0.527111 | 0.114233 | 4.614345 | 3.94E-06 | 3.17E-05 |
| CNAG_05854 | 656.6178 | -0.57537 | 0.112797 | -5.10089 | 3.38E-07 | 3.19E-06 |
| CNAG_07679 | 1802.942 | -0.17642 | 0.071667 | -2.46166 | 0.013829 | 0.044159 |
| CNAG_05862 | 625.0716 | 0.214728 | 0.102128 | 2.102541 | 0.035506 | 0.094601 |
| CNAG_05864 | 1656.687 | 0.465818 | 0.079618 | 5.850624 | 4.90E-09 | 6.00E-08 |
| CNAG_05867 | 594.6275 | 0.312555 | 0.126046 | 2.479696 | 0.013149 | 0.042439 |
| CNAG_05868 | 170.4907 | 0.534733 | 0.169612 | 3.152695 | 0.001618 | 0.007181 |
| CNAG_05870 | 98.58754 | -0.46705 | 0.207584 | -2.24993 | 0.024453 | 0.070263 |
| CNAG_05871 | 971.3132 | 0.495462 | 0.092902 | 5.333148 | 9.65E-08 | 9.80E-07 |
| CNAG_05872 | 1992.376 | 1.145367 | 0.096866 | 11.82428 | 2.92E-32 | 2.40E-30 |
| CNAG_05873 | 261.2707 | 0.72672 | 0.136459 | 5.325556 | 1.01E-07 | 1.02E-06 |
| CNAG_12690 | 15.38342 | 1.032287 | 0.499935 | 2.064844 | 0.038938 | 0.101936 |
| CNAG_05874 | 1540.501 | 1.279657 | 0.105751 | 12.10069 | 1.05E-33 | 9.39E-32 |
| CNAG_05877 | 526.5045 | 0.293692 | 0.107484 | 2.732437 | 0.006287 | 0.022841 |
| CNAG_05881 | 3390.181 | 0.202243 | 0.069538 | 2.908365 | 0.003633 | 0.014474 |

| | | | | | | |
|---|---|---|---|---|---|---|
| CNAG_05884 | 6809.555 | -0.55105 | 0.064189 | -8.58471 | 9.11E-18 | 2.60E-16 |
| CNAG_05889 | 1101.081 | -0.33759 | 0.098833 | -3.41575 | 0.000636 | 0.003215 |
| CNAG_12692 | 66.57408 | -1.02494 | 0.34535 | -2.96784 | 0.002999 | 0.012241 |
| CNAG_05891 | 328.9875 | 0.535924 | 0.131011 | 4.090664 | 4.30E-05 | 0.000286 |
| CNAG_05893 | 6660.696 | 0.148069 | 0.073678 | 2.009672 | 0.044466 | 0.113668 |
| CNAG_05896 | 1343.622 | -0.18966 | 0.077328 | -2.45266 | 0.014181 | 0.045077 |
| CNAG_05899 | 2360.263 | -0.19686 | 0.08978 | -2.19271 | 0.028329 | 0.079655 |
| CNAG_05902 | 576.2938 | 0.232468 | 0.100098 | 2.322402 | 0.020211 | 0.060133 |
| CNAG_12695 | 98.66002 | -0.50924 | 0.237636 | -2.14296 | 0.032116 | 0.087542 |
| CNAG_05903 | 1108.543 | 0.19302 | 0.084675 | 2.279534 | 0.022635 | 0.065839 |
| CNAG_05905 | 1057.148 | -0.19287 | 0.080861 | -2.38516 | 0.017072 | 0.052369 |
| CNAG_05907 | 15252.22 | 0.2188 | 0.059718 | 3.663872 | 0.000248 | 0.001396 |
| CNAG_05909 | 24037.26 | 0.440342 | 0.076707 | 5.740601 | 9.43E-09 | 1.11E-07 |
| CNAG_05913 | 783.0792 | -0.34777 | 0.094318 | -3.68725 | 0.000227 | 0.001287 |
| CNAG_05914 | 335.4824 | 0.849004 | 0.138523 | 6.128978 | 8.84E-10 | 1.19E-08 |
| CNAG_05915 | 357.2785 | 0.420937 | 0.154787 | 2.719466 | 0.006539 | 0.023646 |
| CNAG_05918 | 76000.39 | 0.169313 | 0.071308 | 2.374398 | 0.017578 | 0.053668 |
| CNAG_12699 | 218.5523 | 0.532344 | 0.154975 | 3.435024 | 0.000593 | 0.003036 |
| CNAG_05929 | 461.5085 | 0.523866 | 0.120532 | 4.346267 | 1.38E-05 | 0.000102 |
| CNAG_05930 | 1270.701 | 0.253571 | 0.083841 | 3.024412 | 0.002491 | 0.010419 |
| CNAG_05931 | 2777.129 | 0.38471 | 0.066713 | 5.766688 | 8.08E-09 | 9.56E-08 |
| CNAG_05932 | 6684.503 | -0.17859 | 0.066653 | -2.67939 | 0.007376 | 0.026223 |
| CNAG_05936 | 2579.487 | -0.39777 | 0.087215 | -4.56083 | 5.10E-06 | 4.01E-05 |
| CNAG_05937 | 669.202 | 0.222971 | 0.100328 | 2.222413 | 0.026255 | 0.074644 |
| CNAG_05939 | 521.1453 | 0.980643 | 0.111037 | 8.831659 | 1.03E-18 | 3.23E-17 |
| CNAG_05940 | 3248.444 | 0.584621 | 0.088842 | 6.580472 | 4.69E-11 | 7.32E-10 |
| CNAG_07684 | 166.6848 | 0.462873 | 0.175433 | 2.638455 | 0.008328 | 0.029159 |
| CNAG_07685 | 362.9124 | 0.641813 | 0.118558 | 5.413487 | 6.18E-08 | 6.45E-07 |
| CNAG_07688 | 1316.993 | 0.348446 | 0.083462 | 4.174884 | 2.98E-05 | 0.000204 |
| CNAG_07691 | 118.7076 | 0.584254 | 0.20861 | 2.800698 | 0.005099 | 0.019123 |
| CNAG_12707 | 16.02995 | 1.212051 | 0.487957 | 2.48393 | 0.012994 | 0.042095 |
| CNAG_07692 | 892.3386 | -0.36375 | 0.086923 | -4.18473 | 2.85E-05 | 0.000196 |
| CNAG_07694 | 2040.419 | -0.20582 | 0.071367 | -2.88394 | 0.003927 | 0.015454 |
| CNAG_07695 | 8506.504 | 1.190026 | 0.065543 | 18.15633 | 1.14E-73 | 7.44E-71 |
| CNAG_07699 | 1040.26 | -0.19775 | 0.092835 | -2.13015 | 0.033159 | 0.089666 |
| CNAG_07701 | 675.5449 | 0.19257 | 0.095229 | 2.022187 | 0.043157 | 0.111008 |
| CNAG_07702 | 171.6778 | -0.75741 | 0.156688 | -4.83389 | 1.34E-06 | 1.15E-05 |
| CNAG_07703 | 279.2129 | 0.291954 | 0.134929 | 2.163751 | 0.030483 | 0.08421 |
| CNAG_05962 | 179.8431 | 0.438807 | 0.167698 | 2.616643 | 0.00888 | 0.030771 |
| CNAG_05964 | 100.0946 | -0.52641 | 0.214019 | -2.45964 | 0.013908 | 0.044355 |
| CNAG_05965 | 363.2837 | -0.37458 | 0.126039 | -2.97198 | 0.002959 | 0.012102 |
| CNAG_05968 | 409.0805 | -0.44316 | 0.110007 | -4.02844 | 5.61E-05 | 0.000366 |
| CNAG_05970 | 453.5138 | -0.28511 | 0.117253 | -2.43157 | 0.015034 | 0.047308 |
| CNAG_05972 | 92.74429 | 0.580615 | 0.224893 | 2.581738 | 0.00983 | 0.033384 |
| CNAG_12711 | 14.61462 | 1.006789 | 0.504388 | 1.99606 | 0.045927 | 0.116479 |
| CNAG_12712 | 435.7475 | 0.326127 | 0.11676 | 2.79315 | 0.00522 | 0.019472 |
| CNAG_05974 | 179.9595 | 0.318622 | 0.158838 | 2.005959 | 0.044861 | 0.114409 |
| CNAG_12714 | 110.1607 | -0.8204 | 0.239947 | -3.41908 | 0.000628 | 0.003186 |

| | | | | | | |
|---|---|---|---|---|---|---|
| CNAG_05975 | 3522.451 | -0.1716 | 0.071616 | -2.39605 | 0.016573 | 0.05103 |
| CNAG_05976 | 4591.032 | -0.27898 | 0.071959 | -3.87687 | 0.000106 | 0.000648 |
| CNAG_05977 | 3659.032 | -0.29692 | 0.113186 | -2.62328 | 0.008709 | 0.030313 |
| CNAG_10090 | 4.449642 | -2.37464 | 1.059046 | -2.24224 | 0.024946 | 0.071519 |
| CNAG_05979 | 921.1813 | 0.211762 | 0.08556 | 2.475003 | 0.013324 | 0.042877 |
| CNAG_05982 | 120.928 | 0.614128 | 0.185068 | 3.318383 | 0.000905 | 0.004356 |
| CNAG_03084 | 230.4362 | 1.352661 | 0.148667 | 9.0986 | 9.15E-20 | 3.26E-18 |
| CNAG_03085 | 13.94484 | 1.431724 | 0.556718 | 2.57172 | 0.010119 | 0.034172 |
| CNAG_12715 | 45.59801 | -0.59103 | 0.30093 | -1.96401 | 0.049529 | 0.123649 |
| CNAG_03097 | 592.9814 | 0.241098 | 0.097153 | 2.481628 | 0.013078 | 0.042262 |
| CNAG_03098 | 2152.345 | 0.344128 | 0.07863 | 4.376548 | 1.21E-05 | 9.00E-05 |
| CNAG_03101 | 315.226 | -0.29444 | 0.124936 | -2.35676 | 0.018435 | 0.055676 |
| CNAG_03103 | 1764.363 | -0.19986 | 0.094064 | -2.12477 | 0.033606 | 0.090529 |
| CNAG_03108 | 707.4713 | 0.184901 | 0.094223 | 1.962379 | 0.049718 | 0.124081 |
| CNAG_03109 | 3718.852 | -0.2071 | 0.06814 | -3.03931 | 0.002371 | 0.009949 |
| CNAG_03113 | 3608.984 | 0.779742 | 0.097481 | 7.998881 | 1.26E-15 | 3.04E-14 |
| CNAG_03114 | 2359.131 | 0.489224 | 0.098138 | 4.985063 | 6.19E-07 | 5.58E-06 |
| CNAG_03115 | 624.1746 | -0.41522 | 0.099754 | -4.16245 | 3.15E-05 | 0.000215 |
| CNAG_12720 | 88.83908 | -0.56092 | 0.226597 | -2.4754 | 0.013309 | 0.042865 |
| CNAG_03118 | 750.1654 | 0.217123 | 0.089536 | 2.424984 | 0.015309 | 0.048058 |
| CNAG_03122 | 158.8704 | -0.32657 | 0.161802 | -2.01832 | 0.043558 | 0.111781 |
| CNAG_12721 | 128.3393 | -0.56203 | 0.195539 | -2.87426 | 0.00405 | 0.01585 |
| CNAG_03124 | 3384.072 | -0.54447 | 0.072216 | -7.53952 | 4.72E-14 | 1.00E-12 |
| CNAG_03127 | 45797.26 | -0.17551 | 0.074575 | -2.35345 | 0.0186 | 0.056087 |
| CNAG_03128 | 771.9169 | 0.351668 | 0.093685 | 3.753742 | 0.000174 | 0.00101 |
| CNAG_03129 | 242.4574 | -0.49992 | 0.156251 | -3.19948 | 0.001377 | 0.006258 |
| CNAG_03136 | 2889.86 | 0.391609 | 0.068229 | 5.73964 | 9.49E-09 | 1.12E-07 |
| CNAG_12722 | 49.45307 | 1.229821 | 0.282896 | 4.347257 | 1.38E-05 | 0.000102 |
| CNAG_03139 | 4186.106 | -0.34886 | 0.065057 | -5.36236 | 8.21E-08 | 8.41E-07 |
| CNAG_03141 | 54.5919 | 0.520671 | 0.264972 | 1.965006 | 0.049414 | 0.1234 |
| CNAG_03142 | 12025.96 | 0.918456 | 0.077509 | 11.84964 | 2.16E-32 | 1.81E-30 |
| CNAG_03143 | 9624.426 | 1.511001 | 0.107075 | 14.11166 | 3.22E-45 | 5.71E-43 |
| CNAG_03144 | 1785.146 | -0.25468 | 0.079263 | -3.21308 | 0.001313 | 0.006004 |
| CNAG_03146 | 11354.1 | 0.241367 | 0.073078 | 3.302857 | 0.000957 | 0.004574 |
| CNAG_03147 | 1373.667 | 0.216369 | 0.084102 | 2.572698 | 0.010091 | 0.03412 |
| CNAG_03150 | 1025.861 | -0.26753 | 0.086682 | -3.0864 | 0.002026 | 0.008703 |
| CNAG_03152 | 1669.026 | 0.17581 | 0.076219 | 2.306662 | 0.021074 | 0.062248 |
| CNAG_03153 | 7444.347 | -0.29331 | 0.06442 | -4.55304 | 5.29E-06 | 4.16E-05 |
| CNAG_12727 | 172.5004 | 0.453246 | 0.162594 | 2.787592 | 0.00531 | 0.019762 |
| CNAG_03155 | 1089.487 | 0.346313 | 0.085717 | 4.040203 | 5.34E-05 | 0.00035 |
| CNAG_03156 | 1122.978 | 0.355188 | 0.083311 | 4.263377 | 2.01E-05 | 0.000142 |
| CNAG_03158 | 1196.803 | 0.171803 | 0.077397 | 2.219778 | 0.026434 | 0.075096 |
| CNAG_03159 | 1098.277 | -0.44051 | 0.090723 | -4.85558 | 1.20E-06 | 1.03E-05 |
| CNAG_03162 | 781.3536 | 0.57918 | 0.106109 | 5.458356 | 4.81E-08 | 5.10E-07 |
| CNAG_03167 | 1321.037 | -0.18893 | 0.091845 | -2.05703 | 0.039684 | 0.10368 |
| CNAG_03168 | 9006.1 | -0.36874 | 0.057909 | -6.36754 | 1.92E-10 | 2.81E-09 |
| CNAG_03170 | 765.1282 | -0.31628 | 0.091783 | -3.44592 | 0.000569 | 0.002928 |
| CNAG_03171 | 1970.789 | -0.2345 | 0.079974 | -2.93225 | 0.003365 | 0.013551 |

| | | | | | | |
|---|---|---|---|---|---|---|
| CNAG_03173 | 1207.547 | 0.220952 | 0.084991 | 2.599719 | 0.00933 | 0.031991 |
| CNAG_03175 | 675.3006 | -0.30926 | 0.092031 | -3.36038 | 0.000778 | 0.003818 |
| CNAG_03176 | 1053.013 | -0.92365 | 0.096258 | -9.59555 | 8.35E-22 | 3.50E-20 |
| CNAG_03178 | 1088.43 | 0.652354 | 0.096609 | 6.752548 | 1.45E-11 | 2.43E-10 |
| CNAG_03183 | 2106.818 | -0.19397 | 0.076568 | -2.53325 | 0.011301 | 0.037465 |
| CNAG_03184 | 1767.319 | -0.30873 | 0.077036 | -4.00758 | 6.13E-05 | 0.000398 |
| CNAG_03185 | 787.8214 | 0.321259 | 0.099414 | 3.231512 | 0.001231 | 0.005676 |
| CNAG_03187 | 446.5194 | 0.420075 | 0.11174 | 3.759412 | 0.00017 | 0.00099 |
| CNAG_03188 | 1900.364 | -0.4601 | 0.102606 | -4.48417 | 7.32E-06 | 5.65E-05 |
| CNAG_03191 | 1719.935 | 0.165774 | 0.07824 | 2.118787 | 0.034108 | 0.091674 |
| CNAG_03192 | 391.0663 | -0.38219 | 0.115657 | -3.30448 | 0.000952 | 0.00455 |
| CNAG_03198 | 53917.57 | -0.2515 | 0.08138 | -3.0904 | 0.001999 | 0.008604 |
| CNAG_07717 | 2857.473 | -0.23739 | 0.072684 | -3.266 | 0.001091 | 0.005116 |
| CNAG_03205 | 1669.066 | -0.67825 | 0.07643 | -8.87419 | 7.04E-19 | 2.27E-17 |
| CNAG_07718 | 4006.788 | 0.194109 | 0.067742 | 2.865418 | 0.004165 | 0.01621 |
| CNAG_03211 | 87.31884 | -0.55408 | 0.219799 | -2.52083 | 0.011708 | 0.038634 |
| CNAG_03213 | 3625.265 | 0.652554 | 0.084045 | 7.76434 | 8.21E-15 | 1.86E-13 |
| CNAG_03215 | 281.1569 | -0.51876 | 0.142208 | -3.64787 | 0.000264 | 0.001481 |
| CNAG_07719 | 4926.314 | -0.24171 | 0.069101 | -3.49789 | 0.000469 | 0.002468 |
| CNAG_07720 | 2116.664 | 0.343329 | 0.091419 | 3.755568 | 0.000173 | 0.001004 |
| CNAG_03220 | 1791.554 | 0.341963 | 0.071537 | 4.780254 | 1.75E-06 | 1.48E-05 |
| CNAG_03222 | 365.9484 | -0.31098 | 0.146299 | -2.12564 | 0.033534 | 0.090445 |
| CNAG_03225 | 20022.78 | 0.305649 | 0.078418 | 3.89768 | 9.71E-05 | 0.000599 |
| CNAG_03226 | 7919.541 | 1.076126 | 0.07382 | 14.57761 | 3.90E-48 | 7.25E-46 |
| CNAG_12740 | 111.2189 | 0.615044 | 0.18989 | 3.238946 | 0.0012 | 0.005553 |
| CNAG_03227 | 540.783 | 0.57364 | 0.120685 | 4.753203 | 2.00E-06 | 1.67E-05 |
| CNAG_03228 | 1421.499 | 0.52796 | 0.088279 | 5.980579 | 2.22E-09 | 2.84E-08 |
| CNAG_03231 | 374.3965 | -0.62882 | 0.122978 | -5.11331 | 3.17E-07 | 3.00E-06 |
| CNAG_03232 | 1180.121 | 0.314651 | 0.081146 | 3.877593 | 0.000105 | 0.000647 |
| CNAG_03235 | 2473.885 | -0.17182 | 0.070741 | -2.42883 | 0.015148 | 0.047647 |
| CNAG_03246 | 1309.378 | -0.53473 | 0.086046 | -6.21442 | 5.15E-10 | 7.09E-09 |
| CNAG_03249 | 5303.822 | -0.22797 | 0.0642 | -3.5509 | 0.000384 | 0.002053 |
| CNAG_03250 | 1045.487 | 0.494509 | 0.083564 | 5.9177 | 3.26E-09 | 4.07E-08 |
| CNAG_03258 | 1356.296 | -0.32302 | 0.109613 | -2.94689 | 0.00321 | 0.012986 |
| CNAG_03262 | 1940.9 | -0.305 | 0.076883 | -3.96708 | 7.28E-05 | 0.000466 |
| CNAG_03263 | 17074.96 | 0.77233 | 0.063372 | 12.18718 | 3.64E-34 | 3.34E-32 |
| CNAG_03264 | 1474.553 | -0.2926 | 0.074526 | -3.92621 | 8.63E-05 | 0.00054 |
| CNAG_03265 | 869.9082 | -0.29941 | 0.087308 | -3.4293 | 0.000605 | 0.003087 |
| CNAG_03268 | 3360.956 | 0.199451 | 0.082425 | 2.419776 | 0.01553 | 0.048537 |
| CNAG_03271 | 3258.937 | -0.29334 | 0.085746 | -3.42101 | 0.000624 | 0.003166 |
| CNAG_03272 | 147.3311 | 0.573266 | 0.175609 | 3.264451 | 0.001097 | 0.005141 |
| CNAG_03275 | 719.4191 | -0.34369 | 0.096382 | -3.56587 | 0.000363 | 0.001953 |
| CNAG_03281 | 23715.43 | -0.17301 | 0.082057 | -2.10841 | 0.034995 | 0.093657 |
| CNAG_03282 | 1526.421 | 0.18223 | 0.080147 | 2.273702 | 0.022984 | 0.066704 |
| CNAG_03283 | 38384.47 | -0.19422 | 0.081714 | -2.37683 | 0.017462 | 0.053358 |
| CNAG_03284 | 1166.398 | -0.51197 | 0.079178 | -6.46603 | 1.01E-10 | 1.52E-09 |
| CNAG_03285 | 1983.742 | -0.18874 | 0.070156 | -2.69027 | 0.007139 | 0.025476 |
| CNAG_03286 | 1122.141 | -0.27195 | 0.088424 | -3.07547 | 0.002102 | 0.008967 |

| | | | | | | |
|---|---|---|---|---|---|---|
| CNAG_03289 | 4736.878 | -0.23775 | 0.06319 | -3.76252 | 0.000168 | 0.000979 |
| CNAG_03291 | 1452.769 | 0.168395 | 0.08147 | 2.066952 | 0.038739 | 0.101585 |
| CNAG_03293 | 1469.12 | -0.20316 | 0.088356 | -2.29936 | 0.021485 | 0.063175 |
| CNAG_03297 | 2283.684 | -0.25635 | 0.076337 | -3.35816 | 0.000785 | 0.003846 |
| CNAG_03299 | 6114.225 | -0.1682 | 0.064286 | -2.61635 | 0.008888 | 0.030772 |
| CNAG_03300 | 2172.228 | -0.45539 | 0.074772 | -6.09039 | 1.13E-09 | 1.49E-08 |
| CNAG_03301 | 2776.169 | 0.263411 | 0.07055 | 3.733702 | 0.000189 | 0.001084 |
| CNAG_03302 | 2227.025 | -0.69484 | 0.07303 | -9.5144 | 1.83E-21 | 7.55E-20 |
| CNAG_03303 | 16885.29 | -0.19073 | 0.071067 | -2.68376 | 0.00728 | 0.025941 |
| CNAG_03310 | 811.6137 | -0.22741 | 0.091616 | -2.48223 | 0.013056 | 0.042209 |
| CNAG_03311 | 7408.077 | -0.61784 | 0.06685 | -9.24215 | 2.42E-20 | 9.02E-19 |
| CNAG_03319 | 3157.007 | -0.23132 | 0.072875 | -3.17414 | 0.001503 | 0.006732 |
| CNAG_03320 | 1525.257 | -0.20518 | 0.089126 | -2.30217 | 0.021326 | 0.062849 |
| CNAG_03321 | 2484.842 | 0.365019 | 0.080475 | 4.535825 | 5.74E-06 | 4.49E-05 |
| CNAG_03333 | 2240.014 | 0.978728 | 0.099448 | 9.841645 | 7.45E-23 | 3.48E-21 |
| CNAG_03335 | 3483.162 | -0.29308 | 0.067116 | -4.36674 | 1.26E-05 | 9.38E-05 |
| CNAG_03336 | 1948.461 | -0.5576 | 0.078073 | -7.14211 | 9.19E-13 | 1.74E-11 |
| CNAG_03337 | 1260.194 | 0.254126 | 0.080494 | 3.157065 | 0.001594 | 0.007083 |
| CNAG_03338 | 752.783 | -0.19044 | 0.096243 | -1.9788 | 0.047839 | 0.120391 |
| CNAG_03342 | 3045.033 | -0.32492 | 0.068947 | -4.71255 | 2.45E-06 | 2.03E-05 |
| CNAG_03345 | 3797.605 | 0.218648 | 0.106981 | 2.043802 | 0.040973 | 0.106526 |
| CNAG_03346 | 727.4381 | 0.457022 | 0.10199 | 4.48106 | 7.43E-06 | 5.73E-05 |
| CNAG_12757 | 25.25078 | 0.895506 | 0.412346 | 2.171736 | 0.029876 | 0.082853 |
| CNAG_03347 | 2133.654 | 0.868191 | 0.10107 | 8.590008 | 8.70E-18 | 2.50E-16 |
| CNAG_03349 | 3916.181 | -0.17342 | 0.070944 | -2.44448 | 0.014506 | 0.045925 |
| CNAG_07724 | 1782.631 | -0.3056 | 0.10199 | -2.99638 | 0.002732 | 0.011323 |
| CNAG_03352 | 903.6938 | 0.199547 | 0.094304 | 2.115995 | 0.034345 | 0.09217 |
| CNAG_03354 | 655.8682 | -0.42946 | 0.096531 | -4.44893 | 8.63E-06 | 6.58E-05 |
| CNAG_03358 | 21706.9 | 0.175629 | 0.070582 | 2.488277 | 0.012836 | 0.04167 |
| CNAG_03359 | 2609.13 | 0.525799 | 0.089242 | 5.891813 | 3.82E-09 | 4.74E-08 |
| CNAG_03365 | 2072.541 | 0.247885 | 0.073834 | 3.357343 | 0.000787 | 0.003855 |
| CNAG_12760 | 17.4431 | 1.099065 | 0.501315 | 2.192363 | 0.028353 | 0.079655 |
| CNAG_12761 | 120.7631 | -0.5945 | 0.187516 | -3.17038 | 0.001522 | 0.006816 |
| CNAG_03372 | 1764.344 | 0.363244 | 0.078911 | 4.603212 | 4.16E-06 | 3.34E-05 |
| CNAG_03375 | 9205.893 | 0.684575 | 0.077751 | 8.804675 | 1.31E-18 | 4.10E-17 |
| CNAG_03376 | 1424.108 | 0.207167 | 0.078553 | 2.637284 | 0.008357 | 0.029221 |
| CNAG_03378 | 2145.273 | -0.14873 | 0.072465 | -2.05246 | 0.040125 | 0.104588 |
| CNAG_03381 | 452.0506 | 0.507562 | 0.110072 | 4.611184 | 4.00E-06 | 3.22E-05 |
| CNAG_03383 | 1142.764 | -0.22374 | 0.086311 | -2.59224 | 0.009535 | 0.032552 |
| CNAG_03385 | 1436.328 | 0.226726 | 0.082733 | 2.740462 | 0.006135 | 0.022411 |
| CNAG_12764 | 41.81585 | -0.86595 | 0.309855 | -2.7947 | 0.005195 | 0.0194 |
| CNAG_03389 | 5022.74 | -0.19916 | 0.070593 | -2.8212 | 0.004784 | 0.018148 |
| CNAG_03390 | 1084.149 | -0.49591 | 0.08147 | -6.08707 | 1.15E-09 | 1.52E-08 |
| CNAG_03392 | 428.595 | 0.324731 | 0.118761 | 2.734324 | 0.006251 | 0.022731 |
| CNAG_03394 | 387.543 | 0.446259 | 0.113738 | 3.923578 | 8.72E-05 | 0.000545 |
| CNAG_03395 | 1768.274 | 0.329063 | 0.072279 | 4.552695 | 5.30E-06 | 4.16E-05 |
| CNAG_03396 | 2166.482 | 0.14432 | 0.071921 | 2.006637 | 0.044788 | 0.114299 |
| CNAG_03399 | 4610.012 | -0.31705 | 0.070768 | -4.48019 | 7.46E-06 | 5.75E-05 |

| | | | | | | |
|---|---|---|---|---|---|---|
| CNAG_03402 | 1335.834 | 0.177622 | 0.075544 | 2.351237 | 0.018711 | 0.056292 |
| CNAG_03403 | 760.1537 | 0.400253 | 0.093542 | 4.278846 | 1.88E-05 | 0.000135 |
| CNAG_12766 | 107.374 | 1.24672 | 0.213121 | 5.849811 | 4.92E-09 | 6.02E-08 |
| CNAG_03408 | 4329.769 | 0.885112 | 0.108674 | 8.144621 | 3.80E-16 | 9.52E-15 |
| CNAG_03409 | 1193.2 | -0.25688 | 0.126315 | -2.03367 | 0.041985 | 0.108637 |
| CNAG_03411 | 656.4367 | -0.19481 | 0.096474 | -2.01928 | 0.043458 | 0.111635 |
| CNAG_03414 | 431.6995 | 0.352495 | 0.111402 | 3.164187 | 0.001555 | 0.006939 |
| CNAG_03415 | 313.8606 | 0.268704 | 0.125397 | 2.142827 | 0.032127 | 0.087542 |
| CNAG_03423 | 1459.786 | -0.19103 | 0.091489 | -2.08806 | 0.036793 | 0.097266 |
| CNAG_03438 | 18468.68 | 0.29153 | 0.058266 | 5.003461 | 5.63E-07 | 5.14E-06 |
| CNAG_03445 | 391.0651 | 0.345614 | 0.117888 | 2.931712 | 0.003371 | 0.01356 |
| CNAG_03446 | 774.449 | 0.220154 | 0.094672 | 2.325437 | 0.020049 | 0.059717 |
| CNAG_03447 | 1242.315 | -0.20404 | 0.085952 | -2.37384 | 0.017604 | 0.053707 |
| CNAG_03448 | 1868.853 | -0.19526 | 0.076947 | -2.53753 | 0.011164 | 0.037073 |
| CNAG_03450 | 1606.537 | 0.191679 | 0.096064 | 1.995335 | 0.046006 | 0.116557 |
| CNAG_03452 | 1516.824 | 0.327662 | 0.082696 | 3.962231 | 7.43E-05 | 0.000474 |
| CNAG_03453 | 1313.888 | -0.32866 | 0.108618 | -3.02588 | 0.002479 | 0.010385 |
| CNAG_03454 | 2173.141 | -0.98384 | 0.089349 | -11.0112 | 3.37E-28 | 2.29E-26 |
| CNAG_12773 | 118.5074 | 0.723301 | 0.230084 | 3.143646 | 0.001669 | 0.007386 |
| CNAG_03457 | 8811.615 | -0.23057 | 0.06267 | -3.67911 | 0.000234 | 0.001324 |
| CNAG_03465 | 687.4842 | -0.60906 | 0.096249 | -6.32795 | 2.48E-10 | 3.59E-09 |
| CNAG_03466 | 1874.843 | -0.25234 | 0.097088 | -2.59915 | 0.009346 | 0.03203 |
| CNAG_03467 | 337.6966 | -0.31676 | 0.11976 | -2.64494 | 0.00817 | 0.028799 |
| CNAG_07735 | 2151.244 | -0.61106 | 0.079221 | -7.71333 | 1.23E-14 | 2.74E-13 |
| CNAG_07736 | 988.2488 | -0.24953 | 0.083933 | -2.97301 | 0.002949 | 0.012068 |
| CNAG_12779 | 170.5291 | -0.44846 | 0.159561 | -2.8106 | 0.004945 | 0.018652 |
| CNAG_12780 | 887.8317 | -0.45792 | 0.109045 | -4.19937 | 2.68E-05 | 0.000185 |
| CNAG_03473 | 475.3949 | 0.484501 | 0.102842 | 4.71112 | 2.46E-06 | 2.04E-05 |
| CNAG_03474 | 87.98224 | -0.89707 | 0.239599 | -3.74406 | 0.000181 | 0.001044 |
| CNAG_03475 | 1702.471 | 0.275007 | 0.079673 | 3.451699 | 0.000557 | 0.00287 |
| CNAG_03476 | 10400.98 | -0.1378 | 0.068488 | -2.01202 | 0.044218 | 0.113288 |
| CNAG_12781 | 216.1675 | 0.504716 | 0.178032 | 2.834975 | 0.004583 | 0.017532 |
| CNAG_03478 | 279.7529 | -0.35967 | 0.127479 | -2.82144 | 0.004781 | 0.018148 |
| CNAG_03479 | 2361.818 | -0.17537 | 0.084895 | -2.06566 | 0.03886 | 0.101767 |
| CNAG_03481 | 816.8103 | -0.3365 | 0.085603 | -3.93093 | 8.46E-05 | 0.000533 |
| CNAG_03482 | 14375.19 | 0.26664 | 0.0697 | 3.825541 | 0.00013 | 0.000781 |
| CNAG_03483 | 1578.742 | -0.235 | 0.085435 | -2.75062 | 0.005948 | 0.021764 |
| CNAG_03484 | 1610.125 | -0.16577 | 0.082939 | -1.99864 | 0.045647 | 0.115923 |
| CNAG_03485 | 1299.74 | -0.29095 | 0.092883 | -3.13243 | 0.001734 | 0.007631 |
| CNAG_03487 | 1061.947 | 0.272293 | 0.088276 | 3.084563 | 0.002039 | 0.008741 |
| CNAG_03489 | 897.4476 | -0.228 | 0.088872 | -2.56551 | 0.010302 | 0.03464 |
| CNAG_03492 | 2855.724 | 1.085883 | 0.107477 | 10.10337 | 5.34E-24 | 2.78E-22 |
| CNAG_03493 | 474.7629 | 0.243401 | 0.105772 | 2.301181 | 0.021381 | 0.062919 |
| CNAG_03494 | 1493.794 | -0.15659 | 0.079516 | -1.96923 | 0.048926 | 0.122416 |
| CNAG_12786 | 500.2453 | 0.577607 | 0.135369 | 4.2669 | 1.98E-05 | 0.00014 |
| CNAG_03498 | 477.0979 | -0.36773 | 0.116382 | -3.15965 | 0.00158 | 0.007028 |
| CNAG_03502 | 3100.245 | 0.400835 | 0.075355 | 5.319285 | 1.04E-07 | 1.05E-06 |
| CNAG_03503 | 846.7162 | -0.51116 | 0.121669 | -4.20123 | 2.65E-05 | 0.000183 |

| | | | | | | |
|---|---|---|---|---|---|---|
| CNAG_03507 | 13324.65 | 0.233606 | 0.09204 | 2.538092 | 0.011146 | 0.037049 |
| CNAG_03509 | 6462.621 | 0.652835 | 0.07332 | 8.903865 | 5.39E-19 | 1.75E-17 |
| CNAG_03510 | 26805.38 | -0.13397 | 0.063232 | -2.11869 | 0.034116 | 0.091674 |
| CNAG_03511 | 251.8151 | 0.640272 | 0.143964 | 4.447458 | 8.69E-06 | 6.62E-05 |
| CNAG_03513 | 1343.611 | -0.41865 | 0.089494 | -4.67792 | 2.90E-06 | 2.38E-05 |
| CNAG_07740 | 4835.611 | -0.13341 | 0.066257 | -2.01347 | 0.044065 | 0.112935 |
| CNAG_07741 | 1974.809 | 0.195119 | 0.07331 | 2.661571 | 0.007778 | 0.027515 |
| CNAG_03517 | 4996.552 | 0.640271 | 0.068221 | 9.385203 | 6.28E-21 | 2.50E-19 |
| CNAG_03518 | 86.32415 | -0.51491 | 0.218976 | -2.35146 | 0.0187 | 0.056279 |
| CNAG_03525 | 1153.455 | 0.356751 | 0.096246 | 3.706667 | 0.00021 | 0.001195 |
| CNAG_03527 | 2047.145 | 0.239945 | 0.075362 | 3.183908 | 0.001453 | 0.006558 |
| CNAG_07995 | 1587.186 | 0.160675 | 0.075357 | 2.132174 | 0.032993 | 0.089277 |
| CNAG_03534 | 308.5861 | 0.524143 | 0.124281 | 4.217398 | 2.47E-05 | 0.000172 |
| CNAG_07745 | 3835.375 | 1.415196 | 0.100665 | 14.05847 | 6.83E-45 | 1.19E-42 |
| CNAG_03543 | 1739.478 | 0.343697 | 0.077141 | 4.455443 | 8.37E-06 | 6.41E-05 |
| CNAG_03552 | 896.6971 | 0.873576 | 0.087422 | 9.992669 | 1.64E-23 | 8.16E-22 |
| CNAG_03553 | 5065.735 | 0.276191 | 0.06502 | 4.247796 | 2.16E-05 | 0.000152 |
| CNAG_03554 | 10174.61 | -0.20714 | 0.067294 | -3.07815 | 0.002083 | 0.008907 |
| CNAG_03555 | 487.8303 | 0.404396 | 0.122736 | 3.294856 | 0.000985 | 0.004689 |
| CNAG_03557 | 868.4628 | 0.242426 | 0.093382 | 2.596079 | 0.009429 | 0.032289 |
| CNAG_03559 | 516.2912 | 0.262108 | 0.102014 | 2.569343 | 0.010189 | 0.034348 |
| CNAG_03561 | 847.8344 | 0.290532 | 0.097281 | 2.986531 | 0.002822 | 0.011626 |
| CNAG_03563 | 3377.05 | 0.76039 | 0.073818 | 10.30095 | 6.98E-25 | 3.83E-23 |
| CNAG_03564 | 1578.123 | 0.886484 | 0.087701 | 10.10803 | 5.09E-24 | 2.67E-22 |
| CNAG_03565 | 3266.017 | 0.838819 | 0.072167 | 11.62323 | 3.14E-31 | 2.43E-29 |
| CNAG_03566 | 4987.727 | 0.974966 | 0.090958 | 10.71891 | 8.30E-27 | 5.18E-25 |
| CNAG_03567 | 2479.539 | 0.169069 | 0.07279 | 2.322698 | 0.020195 | 0.060109 |
| CNAG_03570 | 83.68475 | 0.531441 | 0.223173 | 2.381293 | 0.017252 | 0.052819 |
| CNAG_03572 | 2680.889 | 1.574159 | 0.081656 | 19.27783 | 8.25E-83 | 7.15E-80 |
| CNAG_03577 | 51973.66 | -0.16562 | 0.073176 | -2.26327 | 0.023619 | 0.068168 |
| CNAG_07748 | 494.8356 | -0.31141 | 0.102731 | -3.03134 | 0.002435 | 0.010204 |
| CNAG_03583 | 934.8854 | 0.199175 | 0.089052 | 2.236616 | 0.025311 | 0.072462 |
| CNAG_03592 | 3129.466 | 0.379825 | 0.069765 | 5.44431 | 5.20E-08 | 5.48E-07 |
| CNAG_06993 | 4571.13 | 0.378651 | 0.076288 | 4.963455 | 6.93E-07 | 6.20E-06 |
| CNAG_07752 | 5908.589 | 0.179042 | 0.060955 | 2.937295 | 0.003311 | 0.01336 |
| CNAG_06967 | 2694.469 | 0.134227 | 0.065875 | 2.037607 | 0.041589 | 0.107837 |
| CNAG_04102 | 364.5348 | -0.36153 | 0.11775 | -3.07036 | 0.002138 | 0.009093 |
| CNAG_04105 | 3897.213 | 1.85249 | 0.107871 | 17.17318 | 4.22E-66 | 2.06E-63 |
| CNAG_07755 | 172.836 | 0.973597 | 0.17445 | 5.580948 | 2.39E-08 | 2.64E-07 |
| CNAG_04106 | 1888.223 | 1.025835 | 0.101095 | 10.14728 | 3.41E-24 | 1.80E-22 |
| CNAG_04107 | 427.8002 | 0.564905 | 0.127966 | 4.414474 | 1.01E-05 | 7.64E-05 |
| CNAG_04108 | 1300.788 | 0.239801 | 0.084554 | 2.836078 | 0.004567 | 0.01748 |
| CNAG_04110 | 493.2954 | 0.231675 | 0.106277 | 2.179928 | 0.029263 | 0.081618 |
| CNAG_12812 | 81.53193 | 0.635349 | 0.227918 | 2.787621 | 0.00531 | 0.019762 |
| CNAG_04112 | 643.1146 | 0.658316 | 0.107745 | 6.109969 | 9.97E-10 | 1.32E-08 |
| CNAG_07756 | 1758.341 | -0.26486 | 0.080274 | -3.29945 | 0.000969 | 0.004618 |
| CNAG_04119 | 1534.979 | -0.32815 | 0.111381 | -2.94616 | 0.003217 | 0.013003 |
| CNAG_04120 | 2299.496 | 0.208022 | 0.071587 | 2.905854 | 0.003663 | 0.014553 |

| | | | | | | |
|---|---|---|---|---|---|---|
| CNAG_04121 | 2340.709 | 0.151728 | 0.072456 | 2.094075 | 0.036253 | 0.096199 |
| CNAG_04124 | 214.0491 | -0.35742 | 0.144517 | -2.47323 | 0.01339 | 0.043014 |
| CNAG_12817 | 44.64556 | 0.702596 | 0.296636 | 2.368549 | 0.017858 | 0.054269 |
| CNAG_07757 | 1129.705 | -0.21303 | 0.09562 | -2.22787 | 0.02589 | 0.073821 |
| CNAG_04136 | 1890.39 | -0.37871 | 0.073597 | -5.14577 | 2.66E-07 | 2.55E-06 |
| CNAG_12820 | 68.41902 | -0.57543 | 0.238093 | -2.41683 | 0.015656 | 0.048834 |
| CNAG_04141 | 3292.224 | -0.32613 | 0.068263 | -4.77753 | 1.77E-06 | 1.50E-05 |
| CNAG_07758 | 393.0913 | -0.30793 | 0.11429 | -2.69431 | 0.007053 | 0.025215 |
| CNAG_04147 | 1013 | -0.65077 | 0.090574 | -7.18493 | 6.72E-13 | 1.29E-11 |
| CNAG_04150 | 1751.566 | -0.24493 | 0.073742 | -3.32142 | 0.000896 | 0.004312 |
| CNAG_04160 | 28.86395 | 0.796288 | 0.37429 | 2.127464 | 0.033382 | 0.090079 |
| CNAG_04163 | 3961.601 | 1.131521 | 0.074326 | 15.22378 | 2.46E-52 | 5.82E-50 |
| CNAG_04168 | 7976.459 | -0.26973 | 0.074029 | -3.64361 | 0.000269 | 0.001501 |
| CNAG_04173 | 1603.926 | 0.336772 | 0.076293 | 4.414193 | 1.01E-05 | 7.64E-05 |
| CNAG_04175 | 2332.032 | -0.7 | 0.071785 | -9.75136 | 1.82E-22 | 8.07E-21 |
| CNAG_04177 | 1024.03 | 0.221008 | 0.085491 | 2.585165 | 0.009733 | 0.033112 |
| CNAG_04184 | 581.5363 | -0.22721 | 0.106256 | -2.13833 | 0.03249 | 0.088316 |
| CNAG_04185 | 819.394 | 0.971775 | 0.11728 | 8.285913 | 1.17E-16 | 3.09E-15 |
| CNAG_04186 | 852.9232 | 0.905827 | 0.088299 | 10.25867 | 1.08E-24 | 5.90E-23 |
| CNAG_04188 | 1025.983 | 0.220977 | 0.085729 | 2.577635 | 0.009948 | 0.033695 |
| CNAG_04189 | 17358.69 | 1.060227 | 0.063457 | 16.70772 | 1.15E-62 | 4.49E-60 |
| CNAG_04190 | 5526.788 | -0.1827 | 0.066144 | -2.76222 | 0.005741 | 0.021083 |
| CNAG_04192 | 7087.585 | -0.31515 | 0.064282 | -4.90266 | 9.45E-07 | 8.23E-06 |
| CNAG_04193 | 1078.696 | 0.193536 | 0.084089 | 2.301575 | 0.021359 | 0.062901 |
| CNAG_04195 | 857.7131 | 0.248843 | 0.091412 | 2.722218 | 0.006485 | 0.023515 |
| CNAG_04200 | 549.3563 | 0.599701 | 0.122606 | 4.891277 | 1.00E-06 | 8.69E-06 |
| CNAG_04205 | 1704.441 | -0.27574 | 0.080517 | -3.4246 | 0.000616 | 0.003134 |
| CNAG_04206 | 435.6872 | 0.54534 | 0.110558 | 4.932612 | 8.11E-07 | 7.12E-06 |
| CNAG_12830 | 57.28478 | 0.679021 | 0.273511 | 2.482614 | 0.013042 | 0.042209 |
| CNAG_04208 | 1423.488 | 0.674541 | 0.079117 | 8.525819 | 1.52E-17 | 4.21E-16 |
| CNAG_04209 | 2623.497 | -0.67281 | 0.077127 | -8.72343 | 2.70E-18 | 8.16E-17 |
| CNAG_04216 | 6.095447 | -1.93457 | 0.869879 | -2.22395 | 0.026152 | 0.074457 |
| CNAG_04219 | 3708.765 | 0.276874 | 0.079537 | 3.481068 | 0.000499 | 0.002607 |
| CNAG_04221 | 1682.419 | 0.218341 | 0.076294 | 2.861855 | 0.004212 | 0.016336 |
| CNAG_04222 | 3696.999 | -0.70336 | 0.070605 | -9.96178 | 2.24E-23 | 1.10E-21 |
| CNAG_04224 | 2540.975 | -0.1697 | 0.077535 | -2.18868 | 0.02862 | 0.080333 |
| CNAG_04230 | 314.366 | -0.53675 | 0.163818 | -3.27651 | 0.001051 | 0.004962 |
| CNAG_04236 | 3578.347 | -0.25819 | 0.072208 | -3.57559 | 0.000349 | 0.001894 |
| CNAG_04242 | 568.6742 | -0.28217 | 0.09929 | -2.84186 | 0.004485 | 0.0172 |
| CNAG_04243 | 1878.639 | -0.34161 | 0.107531 | -3.17687 | 0.001489 | 0.006689 |
| CNAG_04246 | 104.2605 | -0.48401 | 0.19858 | -2.43737 | 0.014794 | 0.046724 |
| CNAG_04248 | 727.7557 | 0.663496 | 0.093931 | 7.063655 | 1.62E-12 | 3.00E-11 |
| CNAG_04253 | 666.0195 | 0.216625 | 0.10629 | 2.038057 | 0.041544 | 0.107837 |
| CNAG_04256 | 262.9423 | 0.317239 | 0.129816 | 2.443762 | 0.014535 | 0.045998 |
| CNAG_04259 | 6415.73 | -0.3363 | 0.070635 | -4.76102 | 1.93E-06 | 1.61E-05 |
| CNAG_04263 | 504.5128 | -0.32849 | 0.114256 | -2.875 | 0.00404 | 0.01582 |
| CNAG_12837 | 10.31796 | 1.676616 | 0.644651 | 2.600813 | 0.0093 | 0.031917 |
| CNAG_04265 | 3254.494 | 0.348682 | 0.082385 | 4.232371 | 2.31E-05 | 0.000162 |

| | | | | | | |
|---|---|---|---|---|---|---|
| CNAG_04267 | 1141.577 | 0.313493 | 0.08141 | 3.850766 | 0.000118 | 0.000712 |
| CNAG_04268 | 580.8018 | 0.242908 | 0.098162 | 2.474559 | 0.01334 | 0.042913 |
| CNAG_04269 | 1284.945 | 0.302748 | 0.088434 | 3.423444 | 0.000618 | 0.003146 |
| CNAG_04270 | 562.5114 | 0.373341 | 0.10675 | 3.49734 | 0.00047 | 0.002471 |
| CNAG_04271 | 624.7144 | -0.39137 | 0.102633 | -3.81324 | 0.000137 | 0.000816 |
| CNAG_04273 | 2366.068 | 0.248423 | 0.078571 | 3.161778 | 0.001568 | 0.006985 |
| CNAG_04275 | 1191.529 | 0.835533 | 0.099783 | 8.373478 | 5.59E-17 | 1.50E-15 |
| CNAG_04276 | 2472.97 | 0.280179 | 0.072384 | 3.870721 | 0.000109 | 0.000663 |
| CNAG_04280 | 58.64069 | -0.64866 | 0.265073 | -2.44709 | 0.014402 | 0.04565 |
| CNAG_04281 | 2629.925 | 0.182316 | 0.070596 | 2.582525 | 0.009808 | 0.033323 |
| CNAG_12840 | 251.9319 | 0.313763 | 0.152939 | 2.051559 | 0.040212 | 0.104781 |
| CNAG_04287 | 3008.396 | 0.38254 | 0.077154 | 4.958159 | 7.12E-07 | 6.33E-06 |
| CNAG_04288 | 813.0117 | 0.217723 | 0.100261 | 2.171556 | 0.029889 | 0.082861 |
| CNAG_04291 | 1361.06 | 0.215709 | 0.083245 | 2.591241 | 0.009563 | 0.032632 |
| CNAG_04293 | 1225.32 | 0.236416 | 0.095414 | 2.477774 | 0.01322 | 0.04265 |
| CNAG_04297 | 618.331 | 0.219255 | 0.100975 | 2.171372 | 0.029903 | 0.082871 |
| CNAG_04298 | 3441.175 | -0.83045 | 0.092174 | -9.00961 | 2.07E-19 | 7.02E-18 |
| CNAG_04299 | 630.4653 | -0.21489 | 0.094787 | -2.26706 | 0.023387 | 0.067635 |
| CNAG_04308 | 884.4189 | -0.69944 | 0.095322 | -7.33766 | 2.17E-13 | 4.46E-12 |
| CNAG_04309 | 321.4219 | 0.502751 | 0.134077 | 3.749708 | 0.000177 | 0.001024 |
| CNAG_04311 | 579.776 | -0.30305 | 0.099077 | -3.05868 | 0.002223 | 0.009414 |
| CNAG_04313 | 273.1472 | -0.31718 | 0.136336 | -2.32644 | 0.019995 | 0.059603 |
| CNAG_04314 | 3470.192 | 0.731261 | 0.082607 | 8.852294 | 8.57E-19 | 2.73E-17 |
| CNAG_04315 | 2592.851 | 0.540211 | 0.079714 | 6.77689 | 1.23E-11 | 2.07E-10 |
| CNAG_04316 | 2129.187 | 0.212655 | 0.088405 | 2.405456 | 0.016152 | 0.050041 |
| CNAG_04322 | 1702.391 | 1.517201 | 0.096963 | 15.64727 | 3.47E-55 | 1.04E-52 |
| CNAG_04326 | 1095.346 | 0.174193 | 0.079708 | 2.185391 | 0.02886 | 0.080784 |
| CNAG_04328 | 1131.136 | -0.18836 | 0.088137 | -2.13709 | 0.032591 | 0.088404 |
| CNAG_04329 | 757.0146 | 0.233245 | 0.100415 | 2.32282 | 0.020189 | 0.060109 |
| CNAG_04334 | 2306.024 | -0.29113 | 0.083027 | -3.50649 | 0.000454 | 0.002399 |
| CNAG_04338 | 583.6387 | 0.327018 | 0.107079 | 3.053999 | 0.002258 | 0.009541 |
| CNAG_04339 | 1660.374 | 0.219067 | 0.076688 | 2.856611 | 0.004282 | 0.016543 |
| CNAG_04340 | 2114.811 | 0.157155 | 0.072779 | 2.15934 | 0.030824 | 0.08482 |
| CNAG_04341 | 1669.834 | -0.27996 | 0.109648 | -2.55331 | 0.01067 | 0.035739 |
| CNAG_04343 | 151.8524 | -0.54419 | 0.173447 | -3.13748 | 0.001704 | 0.007517 |
| CNAG_04344 | 268.6597 | -0.44038 | 0.145779 | -3.02085 | 0.002521 | 0.01052 |
| CNAG_04346 | 249.3646 | -0.3416 | 0.140168 | -2.43708 | 0.014806 | 0.046743 |
| CNAG_04347 | 4789.554 | -0.39289 | 0.064984 | -6.046 | 1.48E-09 | 1.93E-08 |
| CNAG_04348 | 2013.069 | -0.6791 | 0.072492 | -9.36796 | 7.40E-21 | 2.90E-19 |
| CNAG_12847 | 8.249275 | 1.851945 | 0.706672 | 2.620659 | 0.008776 | 0.030507 |
| CNAG_04351 | 418.6929 | 0.494468 | 0.120082 | 4.117772 | 3.83E-05 | 0.000256 |
| CNAG_04353 | 800.5143 | 0.456403 | 0.089011 | 5.127506 | 2.94E-07 | 2.79E-06 |
| CNAG_04354 | 1011.752 | 0.555614 | 0.086298 | 6.438334 | 1.21E-10 | 1.80E-09 |
| CNAG_04355 | 221.1769 | -0.54168 | 0.161443 | -3.35527 | 0.000793 | 0.003875 |
| CNAG_04356 | 1820.819 | -0.41887 | 0.0717 | -5.84206 | 5.16E-09 | 6.29E-08 |
| CNAG_04358 | 2922.877 | -0.27373 | 0.082391 | -3.3223 | 0.000893 | 0.004301 |
| CNAG_04359 | 1406.781 | 0.204756 | 0.103517 | 1.977996 | 0.047929 | 0.120541 |
| CNAG_04360 | 1557.203 | -0.30923 | 0.077396 | -3.99541 | 6.46E-05 | 0.000417 |

| | | | | | | |
|---|---|---|---|---|---|---|
| CNAG_04361 | 5873.562 | 0.340643 | 0.074532 | 4.570423 | 4.87E-06 | 3.84E-05 |
| CNAG_04364 | 3578.922 | -0.16579 | 0.067336 | -2.46212 | 0.013812 | 0.044121 |
| CNAG_04365 | 5407.368 | -0.56644 | 0.065016 | -8.7123 | 2.98E-18 | 8.94E-17 |
| CNAG_04367 | 507.5728 | -0.54298 | 0.109718 | -4.94891 | 7.46E-07 | 6.62E-06 |
| CNAG_04370 | 3497.556 | -0.45147 | 0.096619 | -4.67266 | 2.97E-06 | 2.43E-05 |
| CNAG_04371 | 926.1894 | 0.24368 | 0.093164 | 2.615604 | 0.008907 | 0.030824 |
| CNAG_04372 | 1744.171 | -0.27068 | 0.082778 | -3.26994 | 0.001076 | 0.00506 |
| CNAG_04373 | 6932.488 | 0.155662 | 0.062177 | 2.503536 | 0.012296 | 0.040177 |
| CNAG_04375 | 2840.964 | 0.187631 | 0.071163 | 2.636654 | 0.008373 | 0.029262 |
| CNAG_04376 | 2877.448 | -0.4191 | 0.073919 | -5.66968 | 1.43E-08 | 1.64E-07 |
| CNAG_04386 | 1034.295 | 1.005374 | 0.099733 | 10.08067 | 6.73E-24 | 3.48E-22 |
| CNAG_04388 | 3433.417 | 0.231325 | 0.069539 | 3.326575 | 0.000879 | 0.004251 |
| CNAG_04389 | 600.3918 | -0.25939 | 0.095997 | -2.70203 | 0.006892 | 0.024762 |
| CNAG_07765 | 276.9015 | 1.171023 | 0.160514 | 7.295443 | 2.98E-13 | 5.97E-12 |
| CNAG_04395 | 1772.96 | -0.4965 | 0.077756 | -6.38538 | 1.71E-10 | 2.52E-09 |
| CNAG_04396 | 362.1463 | 0.340287 | 0.118328 | 2.875795 | 0.00403 | 0.015789 |
| CNAG_04398 | 1172.634 | -0.18551 | 0.081625 | -2.27273 | 0.023042 | 0.066823 |
| CNAG_04401 | 2717.892 | 0.148486 | 0.075405 | 1.969179 | 0.048933 | 0.122416 |
| CNAG_04404 | 174.349 | 0.50183 | 0.161759 | 3.102336 | 0.00192 | 0.008324 |
| CNAG_04408 | 1440.646 | -0.29104 | 0.07754 | -3.75338 | 0.000174 | 0.00101 |
| CNAG_04409 | 448.8622 | 0.316738 | 0.135707 | 2.333986 | 0.019596 | 0.058549 |
| CNAG_04412 | 632.1938 | 0.25968 | 0.101932 | 2.547579 | 0.010847 | 0.036223 |
| CNAG_04414 | 973.2687 | 0.268209 | 0.095072 | 2.821104 | 0.004786 | 0.018148 |
| CNAG_04416 | 32.78958 | -0.84891 | 0.351381 | -2.41594 | 0.015695 | 0.048914 |
| CNAG_04417 | 669.3445 | 0.65466 | 0.100497 | 6.514202 | 7.31E-11 | 1.12E-09 |
| CNAG_12852 | 191.0924 | -0.36779 | 0.160625 | -2.28975 | 0.022036 | 0.064408 |
| CNAG_07769 | 2290.137 | -0.2386 | 0.117231 | -2.03528 | 0.041823 | 0.108277 |
| CNAG_04434 | 978.3457 | -0.20923 | 0.095798 | -2.18412 | 0.028954 | 0.080983 |
| CNAG_04440 | 575.3355 | -0.50124 | 0.10152 | -4.93735 | 7.92E-07 | 6.98E-06 |
| CNAG_04443 | 1364.237 | 0.319956 | 0.08143 | 3.929206 | 8.52E-05 | 0.000535 |
| CNAG_04445 | 41820.04 | -0.22353 | 0.071416 | -3.12997 | 0.001748 | 0.007678 |
| CNAG_04446 | 260.5916 | -0.48355 | 0.142414 | -3.39536 | 0.000685 | 0.003433 |
| CNAG_04448 | 65106.93 | -0.24288 | 0.07073 | -3.43388 | 0.000595 | 0.003047 |
| CNAG_04451 | 1197.258 | -0.30365 | 0.077447 | -3.92079 | 8.83E-05 | 0.000551 |
| CNAG_04452 | 1211.155 | -0.1895 | 0.090033 | -2.10483 | 0.035306 | 0.09423 |
| CNAG_04456 | 632.2115 | 0.298368 | 0.103832 | 2.87357 | 0.004059 | 0.015874 |
| CNAG_04459 | 44.42189 | 1.429047 | 0.340127 | 4.201513 | 2.65E-05 | 0.000183 |
| CNAG_12862 | 95.57403 | 0.425554 | 0.207939 | 2.046528 | 0.040704 | 0.105957 |
| CNAG_04466 | 469.6712 | 0.255146 | 0.112797 | 2.261992 | 0.023698 | 0.068293 |
| CNAG_04468 | 303.3887 | 0.481285 | 0.126236 | 3.812581 | 0.000138 | 0.000817 |
| CNAG_12863 | 16.52672 | 1.423029 | 0.494081 | 2.880153 | 0.003975 | 0.015611 |
| CNAG_04470 | 297.4702 | 0.750965 | 0.137789 | 5.450101 | 5.03E-08 | 5.32E-07 |
| CNAG_04471 | 538.7432 | 0.683563 | 0.126716 | 5.394428 | 6.87E-08 | 7.11E-07 |
| CNAG_04472 | 97.78827 | 1.165065 | 0.206193 | 5.650366 | 1.60E-08 | 1.83E-07 |
| CNAG_04473 | 1073.411 | 0.336787 | 0.081341 | 4.140431 | 3.47E-05 | 0.000234 |
| CNAG_04475 | 2040.914 | 0.284018 | 0.070295 | 4.040382 | 5.34E-05 | 0.00035 |
| CNAG_04476 | 139.2986 | 0.626923 | 0.19306 | 3.247306 | 0.001165 | 0.005412 |
| CNAG_07771 | 2221.947 | 0.364935 | 0.069843 | 5.225107 | 1.74E-07 | 1.71E-06 |

| | | | | | | |
|---|---|---|---|---|---|---|
| CNAG_04485 | 6669.458 | 0.265913 | 0.062499 | 4.254657 | 2.09E-05 | 0.000147 |
| CNAG_07775 | 962.4264 | 0.974307 | 0.113836 | 8.558837 | 1.14E-17 | 3.20E-16 |
| CNAG_04495 | 181.3039 | -0.4007 | 0.161005 | -2.48877 | 0.012819 | 0.04163 |
| CNAG_04497 | 574.4487 | -0.19566 | 0.097784 | -2.00096 | 0.045397 | 0.115589 |
| CNAG_04498 | 1453.286 | 0.195981 | 0.083465 | 2.34807 | 0.018871 | 0.056706 |
| CNAG_04499 | 3639.866 | -0.18921 | 0.074191 | -2.55028 | 0.010764 | 0.03602 |
| CNAG_04501 | 3449.385 | -0.28112 | 0.065943 | -4.26308 | 2.02E-05 | 0.000142 |
| CNAG_04503 | 695.2615 | -0.1975 | 0.098806 | -1.99885 | 0.045625 | 0.115903 |
| CNAG_04505 | 2085.026 | -0.43283 | 0.077252 | -5.60274 | 2.11E-08 | 2.35E-07 |
| CNAG_04506 | 86.47989 | 0.622139 | 0.216299 | 2.876295 | 0.004024 | 0.015772 |
| CNAG_04511 | 404.7046 | 0.458712 | 0.117065 | 3.91845 | 8.91E-05 | 0.000554 |
| CNAG_04513 | 2413.342 | -0.52222 | 0.078814 | -6.62607 | 3.45E-11 | 5.42E-10 |
| CNAG_04514 | 1516.943 | -0.22238 | 0.078036 | -2.84975 | 0.004375 | 0.016845 |
| CNAG_04515 | 2645.963 | -0.48035 | 0.081973 | -5.85984 | 4.63E-09 | 5.71E-08 |
| CNAG_12869 | 176.4357 | -0.39919 | 0.158839 | -2.51315 | 0.011966 | 0.039252 |
| CNAG_04521 | 70.9804 | -0.5043 | 0.252665 | -1.99594 | 0.045941 | 0.116479 |
| CNAG_04523 | 278.1895 | 4.959781 | 0.223265 | 22.2148 | 2.47E-109 | 4.82E-106 |
| CNAG_04524 | 1105.208 | 0.280368 | 0.096822 | 2.895721 | 0.003783 | 0.014978 |
| CNAG_04528 | 279.089 | -0.37989 | 0.170599 | -2.22678 | 0.025962 | 0.073979 |
| CNAG_04529 | 391.8999 | 0.317746 | 0.115006 | 2.762863 | 0.00573 | 0.021072 |
| CNAG_04531 | 1514.554 | 0.175641 | 0.075392 | 2.329712 | 0.019821 | 0.059153 |
| CNAG_04536 | 28.41788 | -1.62755 | 0.414679 | -3.92486 | 8.68E-05 | 0.000543 |
| CNAG_04539 | 1169.59 | 0.76801 | 0.082981 | 9.255288 | 2.14E-20 | 8.02E-19 |
| CNAG_04541 | 1429.239 | -0.18666 | 0.081016 | -2.30393 | 0.021226 | 0.062645 |
| CNAG_07782 | 691.4998 | 0.415641 | 0.090898 | 4.572637 | 4.82E-06 | 3.80E-05 |
| CNAG_12874 | 59.51824 | -1.125 | 0.263311 | -4.27249 | 1.93E-05 | 0.000137 |
| CNAG_06997 | 358.5513 | -0.78325 | 0.167738 | -4.66946 | 3.02E-06 | 2.46E-05 |
| CNAG_06999 | 580.1715 | 0.871532 | 0.101445 | 8.591206 | 8.61E-18 | 2.48E-16 |
| CNAG_12875 | 38.64572 | 0.947718 | 0.325922 | 2.907805 | 0.00364 | 0.014485 |
| CNAG_07784 | 32.0604 | -0.72027 | 0.353757 | -2.03605 | 0.041745 | 0.108124 |
| CNAG_04927 | 98.49088 | 0.558263 | 0.224903 | 2.482237 | 0.013056 | 0.042209 |
| CNAG_04926 | 182.7015 | 0.873954 | 0.159861 | 5.466954 | 4.58E-08 | 4.87E-07 |
| CNAG_04920 | 1336.674 | 1.000739 | 0.076792 | 13.03175 | 8.07E-39 | 1.05E-36 |
| CNAG_04919 | 285.9382 | -0.2681 | 0.124702 | -2.14993 | 0.031561 | 0.086331 |
| CNAG_07839 | 49184.56 | -0.17418 | 0.070398 | -2.47423 | 0.013352 | 0.042935 |
| CNAG_04913 | 1583.295 | -0.25353 | 0.075729 | -3.34789 | 0.000814 | 0.003969 |
| CNAG_04911 | 479.5338 | 0.487783 | 0.113049 | 4.314773 | 1.60E-05 | 0.000116 |
| CNAG_04908 | 921.806 | 0.632659 | 0.085871 | 7.367573 | 1.74E-13 | 3.59E-12 |
| CNAG_04906 | 2477.7 | -0.22431 | 0.070038 | -3.20267 | 0.001362 | 0.006207 |
| CNAG_07842 | 151.5463 | 0.880531 | 0.173368 | 5.078977 | 3.79E-07 | 3.54E-06 |
| CNAG_04903 | 1044.609 | 1.08429 | 0.103812 | 10.44477 | 1.55E-25 | 8.82E-24 |
| CNAG_04902 | 1624.476 | 0.214577 | 0.074149 | 2.893843 | 0.003806 | 0.01503 |
| CNAG_04899 | 1348.06 | -0.59525 | 0.07782 | -7.64915 | 2.02E-14 | 4.43E-13 |
| CNAG_04896 | 1670.207 | -0.15274 | 0.07332 | -2.08313 | 0.037239 | 0.098313 |
| CNAG_04894 | 309.5473 | 0.589139 | 0.143894 | 4.094251 | 4.24E-05 | 0.000282 |
| CNAG_04889 | 666.3531 | 0.53923 | 0.112914 | 4.775564 | 1.79E-06 | 1.51E-05 |
| CNAG_04886 | 537.5576 | 0.664967 | 0.108711 | 6.116854 | 9.54E-10 | 1.27E-08 |
| CNAG_12883 | 38.89672 | 0.69366 | 0.324448 | 2.13797 | 0.032519 | 0.088322 |

| | | | | | | |
|---|---|---|---|---|---|---|
| CNAG_04883 | 37370.24 | -0.15532 | 0.072407 | -2.14508 | 0.031946 | 0.08718 |
| CNAG_04879 | 5310.114 | 0.436238 | 0.08084 | 5.396341 | 6.80E-08 | 7.05E-07 |
| CNAG_04874 | 271.818 | -1.02941 | 0.145496 | -7.07519 | 1.49E-12 | 2.77E-11 |
| CNAG_04873 | 164.4219 | -0.54624 | 0.161663 | -3.3789 | 0.000728 | 0.003613 |
| CNAG_04872 | 1912.884 | -0.27875 | 0.077507 | -3.59644 | 0.000323 | 0.001757 |
| CNAG_04869 | 1611.108 | -0.56418 | 0.0805 | -7.00844 | 2.41E-12 | 4.33E-11 |
| CNAG_04868 | 1062.769 | -0.29169 | 0.086047 | -3.38993 | 0.000699 | 0.003493 |
| CNAG_04865 | 80.03711 | 0.724653 | 0.267098 | 2.713059 | 0.006667 | 0.02409 |
| CNAG_04864 | 3335.516 | 0.401224 | 0.078487 | 5.111975 | 3.19E-07 | 3.02E-06 |
| CNAG_04862 | 2841.019 | 0.819458 | 0.085833 | 9.547106 | 1.33E-21 | 5.57E-20 |
| CNAG_12890 | 30.76266 | 1.343195 | 0.36449 | 3.685135 | 0.000229 | 0.001295 |
| CNAG_04861 | 1803.628 | 0.544993 | 0.079778 | 6.831403 | 8.41E-12 | 1.45E-10 |
| CNAG_04859 | 970.5557 | -0.20964 | 0.088337 | -2.37321 | 0.017634 | 0.053736 |
| CNAG_04855 | 929.2033 | -0.2216 | 0.09063 | -2.44515 | 0.014479 | 0.045858 |
| CNAG_04853 | 604.5131 | -0.21941 | 0.097626 | -2.24749 | 0.024609 | 0.070658 |
| CNAG_04851 | 10397.74 | -0.36871 | 0.058448 | -6.30835 | 2.82E-10 | 4.05E-09 |
| CNAG_04849 | 494.1704 | 0.255282 | 0.122607 | 2.082122 | 0.037331 | 0.098424 |
| CNAG_04843 | 1103.258 | -0.32972 | 0.079263 | -4.15983 | 3.18E-05 | 0.000217 |
| CNAG_04837 | 595.3749 | 2.727641 | 0.254583 | 10.71414 | 8.74E-27 | 5.41E-25 |
| CNAG_04836 | 496.7626 | -0.42363 | 0.128902 | -3.28641 | 0.001015 | 0.004814 |
| CNAG_04833 | 1429.36 | 0.746369 | 0.105224 | 7.093151 | 1.31E-12 | 2.46E-11 |
| CNAG_04832 | 1170.947 | -0.27395 | 0.084025 | -3.2604 | 0.001113 | 0.005205 |
| CNAG_04830 | 927.7016 | 0.21999 | 0.093882 | 2.343254 | 0.019116 | 0.057373 |
| CNAG_12895 | 132.1852 | -0.92472 | 0.195424 | -4.73186 | 2.22E-06 | 1.85E-05 |
| CNAG_04828 | 8260.451 | -0.2046 | 0.059775 | -3.42284 | 0.00062 | 0.003151 |
| CNAG_04827 | 280.146 | 0.789368 | 0.135159 | 5.840301 | 5.21E-09 | 6.34E-08 |
| CNAG_04822 | 1231.983 | -0.32548 | 0.086249 | -3.77377 | 0.000161 | 0.000939 |
| CNAG_04820 | 1706.01 | -0.19917 | 0.081109 | -2.45559 | 0.014065 | 0.044766 |
| CNAG_04819 | 511.2956 | -0.39187 | 0.103419 | -3.78917 | 0.000151 | 0.00089 |
| CNAG_04817 | 1007.72 | -0.28924 | 0.08909 | -3.24655 | 0.001168 | 0.005421 |
| CNAG_04816 | 504.2049 | 0.547128 | 0.107469 | 5.091015 | 3.56E-07 | 3.33E-06 |
| CNAG_07845 | 150.8946 | 0.384969 | 0.188938 | 2.037538 | 0.041596 | 0.107837 |
| CNAG_12898 | 351.4718 | 0.382976 | 0.120838 | 3.169348 | 0.001528 | 0.006837 |
| CNAG_12900 | 86.92575 | 0.502919 | 0.220118 | 2.284767 | 0.022326 | 0.065111 |
| CNAG_04807 | 2294.996 | 0.526383 | 0.093788 | 5.612507 | 1.99E-08 | 2.22E-07 |
| CNAG_04805 | 1446.821 | -0.15472 | 0.075975 | -2.03647 | 0.041703 | 0.108052 |
| CNAG_12901 | 63.40217 | -0.7154 | 0.258498 | -2.76752 | 0.005649 | 0.020813 |
| CNAG_04804 | 1715.13 | -0.74996 | 0.076968 | -9.74381 | 1.96E-22 | 8.64E-21 |
| CNAG_04802 | 1575.69 | -0.51882 | 0.073825 | -7.02765 | 2.10E-12 | 3.82E-11 |
| CNAG_04799 | 44372.09 | -0.21261 | 0.074578 | -2.85084 | 0.00436 | 0.016796 |
| CNAG_04794 | 1673.157 | 0.717147 | 0.097619 | 7.346392 | 2.04E-13 | 4.19E-12 |
| CNAG_04791 | 1405.802 | -0.35768 | 0.081789 | -4.37325 | 1.22E-05 | 9.12E-05 |
| CNAG_04790 | 1010.288 | -0.21062 | 0.089962 | -2.34117 | 0.019223 | 0.057633 |
| CNAG_04789 | 1604.377 | 0.464479 | 0.092384 | 5.027677 | 4.96E-07 | 4.57E-06 |
| CNAG_04785 | 591.4588 | 0.221004 | 0.095021 | 2.325851 | 0.020027 | 0.059674 |
| CNAG_04783 | 1661.859 | -0.37653 | 0.079916 | -4.71159 | 2.46E-06 | 2.04E-05 |
| CNAG_07851 | 12957.27 | 1.262657 | 0.068475 | 18.43978 | 6.30E-76 | 4.92E-73 |
| CNAG_04777 | 769.4957 | -0.19129 | 0.093529 | -2.04528 | 0.040827 | 0.106241 |

| | | | | | | |
|---|---|---|---|---|---|---|
| CNAG_04774 | 1382.284 | -0.31871 | 0.092125 | -3.45954 | 0.000541 | 0.0028 |
| CNAG_04772 | 2386.619 | -0.24348 | 0.077305 | -3.14961 | 0.001635 | 0.007245 |
| CNAG_04770 | 2271.623 | -0.50426 | 0.074474 | -6.77096 | 1.28E-11 | 2.15E-10 |
| CNAG_04769 | 1054.581 | -0.22895 | 0.087864 | -2.60567 | 0.009169 | 0.031579 |
| CNAG_04768 | 948.2273 | -0.34692 | 0.091938 | -3.77344 | 0.000161 | 0.00094 |
| CNAG_04766 | 36.47392 | 0.701087 | 0.344103 | 2.037435 | 0.041606 | 0.107837 |
| CNAG_04762 | 47143.77 | -0.24238 | 0.083945 | -2.8873 | 0.003886 | 0.015315 |
| CNAG_04760 | 2535.004 | 0.381612 | 0.105058 | 3.63239 | 0.000281 | 0.001563 |
| CNAG_12910 | 228.4891 | 1.996554 | 0.161624 | 12.35308 | 4.69E-35 | 4.57E-33 |
| CNAG_04758 | 1745.311 | 1.833638 | 0.209159 | 8.766714 | 1.84E-18 | 5.70E-17 |
| CNAG_04757 | 1524.157 | 0.827992 | 0.079811 | 10.37443 | 3.24E-25 | 1.81E-23 |
| CNAG_04756 | 795.7556 | 0.474387 | 0.106145 | 4.469255 | 7.85E-06 | 6.03E-05 |
| CNAG_12911 | 26.20503 | 1.138112 | 0.405934 | 2.803685 | 0.005052 | 0.018974 |
| CNAG_04751 | 735.9765 | 0.349026 | 0.089367 | 3.905549 | 9.40E-05 | 0.000581 |
| CNAG_04748 | 881.1241 | 0.427502 | 0.087077 | 4.909481 | 9.13E-07 | 7.97E-06 |
| CNAG_04747 | 642.97 | 0.626376 | 0.09503 | 6.591319 | 4.36E-11 | 6.82E-10 |
| CNAG_04746 | 1775.861 | 0.672114 | 0.079687 | 8.434417 | 3.33E-17 | 9.08E-16 |
| CNAG_04744 | 1309.805 | 0.948435 | 0.096236 | 9.855338 | 6.50E-23 | 3.08E-21 |
| CNAG_04742 | 743.8265 | 0.276863 | 0.095131 | 2.910332 | 0.00361 | 0.014412 |
| CNAG_04739 | 735.702 | 0.197276 | 0.099927 | 1.974191 | 0.04836 | 0.121312 |
| CNAG_04737 | 6578.799 | 0.981463 | 0.076145 | 12.88942 | 5.16E-38 | 6.40E-36 |
| CNAG_12917 | 66.87135 | 0.580911 | 0.248553 | 2.337171 | 0.01943 | 0.058186 |
| CNAG_04736 | 487.2422 | 0.508366 | 0.118951 | 4.273733 | 1.92E-05 | 0.000137 |
| CNAG_04735 | 3048.101 | 1.066921 | 0.07197 | 14.82455 | 1.02E-49 | 2.09E-47 |
| CNAG_04734 | 1176.237 | 0.281046 | 0.079204 | 3.548362 | 0.000388 | 0.002071 |
| CNAG_04733 | 1515.459 | -0.20058 | 0.091317 | -2.19653 | 0.028054 | 0.079037 |
| CNAG_04730 | 1561.226 | 0.414708 | 0.086908 | 4.771785 | 1.83E-06 | 1.53E-05 |
| CNAG_04729 | 2480.682 | -0.41417 | 0.068346 | -6.05989 | 1.36E-09 | 1.78E-08 |
| CNAG_04727 | 442.8037 | 0.62595 | 0.125616 | 4.983056 | 6.26E-07 | 5.63E-06 |
| CNAG_04726 | 35539.77 | -0.20268 | 0.070275 | -2.88411 | 0.003925 | 0.015454 |
| CNAG_04713 | 1201.178 | -0.1585 | 0.080657 | -1.96514 | 0.049398 | 0.1234 |
| CNAG_04712 | 1034.523 | -0.32916 | 0.082528 | -3.98844 | 6.65E-05 | 0.000429 |
| CNAG_04711 | 72.9111 | -0.79709 | 0.236033 | -3.37702 | 0.000733 | 0.003631 |
| CNAG_04705 | 1641.232 | 0.229998 | 0.073483 | 3.129961 | 0.001748 | 0.007678 |
| CNAG_04704 | 32.36871 | 0.809912 | 0.342156 | 2.367088 | 0.017929 | 0.054463 |
| CNAG_12926 | 65.07278 | 0.654229 | 0.250039 | 2.616509 | 0.008883 | 0.030771 |
| CNAG_04696 | 643.0998 | 0.725799 | 0.10287 | 7.055526 | 1.72E-12 | 3.17E-11 |
| CNAG_04694 | 4420.793 | -0.56608 | 0.06493 | -8.71831 | 2.82E-18 | 8.51E-17 |
| CNAG_04690 | 982.1498 | 1.101745 | 0.088831 | 12.40277 | 2.52E-35 | 2.53E-33 |
| CNAG_04689 | 486.1518 | 0.438031 | 0.105015 | 4.171121 | 3.03E-05 | 0.000207 |
| CNAG_04688 | 405.1434 | 0.24547 | 0.109898 | 2.233621 | 0.025508 | 0.072971 |
| CNAG_04687 | 15516.44 | 0.745205 | 0.076561 | 9.733527 | 2.17E-22 | 9.51E-21 |
| CNAG_07856 | 1792.413 | 1.029614 | 0.081686 | 12.60458 | 1.99E-36 | 2.16E-34 |
| CNAG_04684 | 3516.273 | -0.18071 | 0.065657 | -2.75236 | 0.005917 | 0.021678 |
| CNAG_04680 | 1089.643 | 1.01637 | 0.093804 | 10.83506 | 2.35E-27 | 1.54E-25 |
| CNAG_04677 | 698.5664 | -0.91735 | 0.105686 | -8.67995 | 3.96E-18 | 1.17E-16 |
| CNAG_04676 | 7561.579 | 0.162997 | 0.069082 | 2.359477 | 0.018301 | 0.055335 |
| CNAG_07999 | 474.2018 | 0.287963 | 0.10896 | 2.642825 | 0.008222 | 0.028954 |

| | | | | | | |
|---|---|---|---|---|---|---|
| CNAG_04667 | 689.1706 | 0.285661 | 0.09574 | 2.983709 | 0.002848 | 0.011715 |
| CNAG_04666 | 4502.247 | -0.13835 | 0.064428 | -2.14729 | 0.03177 | 0.086812 |
| CNAG_04664 | 921.9707 | 0.214543 | 0.088214 | 2.43207 | 0.015013 | 0.047261 |
| CNAG_04659 | 15387.64 | 1.131069 | 0.082954 | 13.63481 | 2.49E-42 | 3.66E-40 |
| CNAG_12933 | 114.2058 | 0.989407 | 0.198034 | 4.996156 | 5.85E-07 | 5.31E-06 |
| CNAG_04658 | 574.7633 | 0.514468 | 0.095576 | 5.382809 | 7.33E-08 | 7.54E-07 |
| CNAG_04657 | 362.3469 | -0.28646 | 0.128829 | -2.22353 | 0.02618 | 0.074511 |
| CNAG_04654 | 1063.095 | -0.18915 | 0.083782 | -2.25759 | 0.023971 | 0.069029 |
| CNAG_04640 | 33379.6 | -0.33552 | 0.058824 | -5.70384 | 1.17E-08 | 1.36E-07 |
| CNAG_04639 | 2878.66 | -0.22414 | 0.067752 | -3.30825 | 0.000939 | 0.004497 |
| CNAG_04636 | 1743.766 | 0.249872 | 0.071326 | 3.503262 | 0.00046 | 0.002427 |
| CNAG_04635 | 2091.765 | 0.25345 | 0.070672 | 3.586269 | 0.000335 | 0.001822 |
| CNAG_04634 | 834.0277 | 0.340766 | 0.089384 | 3.812371 | 0.000138 | 0.000817 |
| CNAG_04632 | 320.965 | 1.935172 | 0.33353 | 5.802086 | 6.55E-09 | 7.84E-08 |
| CNAG_12938 | 98.00533 | 1.106445 | 0.205494 | 5.384324 | 7.27E-08 | 7.49E-07 |
| CNAG_04631 | 1110.055 | -0.44537 | 0.0887 | -5.02115 | 5.14E-07 | 4.72E-06 |
| CNAG_04630 | 1657.543 | 0.422312 | 0.096403 | 4.380697 | 1.18E-05 | 8.84E-05 |
| CNAG_04628 | 3115.489 | -0.24509 | 0.077208 | -3.17435 | 0.001502 | 0.006732 |
| CNAG_04626 | 237.5972 | 0.317017 | 0.148675 | 2.132278 | 0.032984 | 0.089277 |
| CNAG_04625 | 2885.583 | 0.669971 | 0.077459 | 8.649319 | 5.18E-18 | 1.50E-16 |
| CNAG_04621 | 5769.563 | -0.19014 | 0.076214 | -2.49484 | 0.012601 | 0.041044 |
| CNAG_04620 | 1156.64 | 0.214379 | 0.085287 | 2.513621 | 0.01195 | 0.039216 |
| CNAG_04617 | 444.1712 | 0.557625 | 0.116803 | 4.774045 | 1.81E-06 | 1.52E-05 |
| CNAG_04616 | 261.2833 | -0.3357 | 0.146764 | -2.28736 | 0.022175 | 0.064789 |
| CNAG_04615 | 357.2966 | -0.59028 | 0.118522 | -4.9804 | 6.35E-07 | 5.70E-06 |
| CNAG_04614 | 315.3463 | 0.306571 | 0.127567 | 2.403216 | 0.016252 | 0.050249 |
| CNAG_04613 | 3032.253 | -0.24741 | 0.068256 | -3.62479 | 0.000289 | 0.001599 |
| CNAG_04611 | 1076.935 | 0.600647 | 0.093198 | 6.444854 | 1.16E-10 | 1.73E-09 |
| CNAG_07858 | 8.716145 | 1.892256 | 0.708315 | 2.67149 | 0.007552 | 0.026812 |
| CNAG_04605 | 5599.391 | -0.21772 | 0.066593 | -3.26936 | 0.001078 | 0.005068 |
| CNAG_04602 | 253.8439 | 0.302634 | 0.140957 | 2.146996 | 0.031794 | 0.086815 |
| CNAG_04599 | 1079.649 | -0.19135 | 0.094434 | -2.02624 | 0.04274 | 0.110153 |
| CNAG_04589 | 468.6882 | 0.516438 | 0.129553 | 3.986306 | 6.71E-05 | 0.000432 |
| CNAG_04588 | 1739.816 | -0.30792 | 0.091933 | -3.34942 | 0.00081 | 0.00395 |
| CNAG_04587 | 3695.574 | 1.17103 | 0.0745 | 15.71863 | 1.13E-55 | 3.52E-53 |
| CNAG_04585 | 154.5088 | 0.674636 | 0.174782 | 3.85988 | 0.000113 | 0.00069 |
| CNAG_04584 | 5054.068 | -0.2007 | 0.080402 | -2.49619 | 0.012554 | 0.040939 |
| CNAG_04581 | 836.0134 | -0.23166 | 0.088443 | -2.61933 | 0.00881 | 0.030585 |
| CNAG_04577 | 4951.614 | 0.267102 | 0.079703 | 3.351196 | 0.000805 | 0.003927 |
| CNAG_07862 | 689.5208 | 0.204964 | 0.090604 | 2.2622 | 0.023685 | 0.068285 |
| CNAG_07863 | 7902.03 | -0.13535 | 0.062145 | -2.17798 | 0.029407 | 0.081904 |
| CNAG_04571 | 1138.344 | -0.40946 | 0.082867 | -4.94123 | 7.76E-07 | 6.87E-06 |
| CNAG_04570 | 5456.346 | -0.32619 | 0.080551 | -4.04948 | 5.13E-05 | 0.000337 |
| CNAG_07864 | 2116.824 | -0.44242 | 0.074538 | -5.93552 | 2.93E-09 | 3.68E-08 |
| CNAG_07866 | 359.707 | 0.353407 | 0.143969 | 2.454746 | 0.014098 | 0.044853 |
| CNAG_04565 | 1179.4 | 0.210818 | 0.083129 | 2.536024 | 0.011212 | 0.037201 |
| CNAG_04564 | 435.2407 | 0.401767 | 0.11126 | 3.611059 | 0.000305 | 0.001671 |
| CNAG_08002 | 2727.153 | -0.16836 | 0.077244 | -2.17961 | 0.029287 | 0.081655 |

| | | | | | | |
|---|---|---|---|---|---|---|
| CNAG_01464 | 25755.58 | 2.542097 | 0.091581 | 27.75783 | 1.40E-169 | 3.65E-166 |
| CNAG_12951 | 102.8894 | 0.914801 | 0.221253 | 4.134639 | 3.56E-05 | 0.00024 |
| CNAG_01470 | 5921.94 | 0.255134 | 0.068673 | 3.715212 | 0.000203 | 0.001158 |
| CNAG_01471 | 1847.962 | 0.353625 | 0.071857 | 4.92123 | 8.60E-07 | 7.53E-06 |
| CNAG_01472 | 981.0237 | 0.267522 | 0.084164 | 3.178575 | 0.00148 | 0.006661 |
| CNAG_01474 | 272.2896 | -0.34904 | 0.13014 | -2.68207 | 0.007317 | 0.026049 |
| CNAG_01475 | 7658.858 | -0.27411 | 0.062419 | -4.39144 | 1.13E-05 | 8.44E-05 |
| CNAG_01480 | 39813.78 | -0.21974 | 0.076899 | -2.85744 | 0.004271 | 0.016511 |
| CNAG_01481 | 238.477 | 0.269911 | 0.13769 | 1.96028 | 0.049963 | 0.124453 |
| CNAG_01485 | 3531.438 | 0.159148 | 0.072718 | 2.188552 | 0.028629 | 0.080333 |
| CNAG_01486 | 56246.86 | -0.21054 | 0.075552 | -2.78669 | 0.005325 | 0.019798 |
| CNAG_07587 | 80.15747 | -0.56185 | 0.233938 | -2.40171 | 0.016319 | 0.050436 |
| CNAG_01489 | 188.2388 | -0.4036 | 0.150712 | -2.67795 | 0.007408 | 0.026312 |
| CNAG_01490 | 375.9664 | -0.64958 | 0.117362 | -5.53483 | 3.12E-08 | 3.39E-07 |
| CNAG_01491 | 1637.925 | 0.266614 | 0.074851 | 3.561927 | 0.000368 | 0.00198 |
| CNAG_01493 | 468.9406 | 1.121695 | 0.125014 | 8.972525 | 2.90E-19 | 9.71E-18 |
| CNAG_12958 | 31.2037 | 1.255061 | 0.393947 | 3.185864 | 0.001443 | 0.006525 |
| CNAG_01495 | 685.4053 | -0.60288 | 0.099457 | -6.06176 | 1.35E-09 | 1.76E-08 |
| CNAG_01498 | 643.5506 | -0.2333 | 0.094934 | -2.45752 | 0.01399 | 0.044562 |
| CNAG_01501 | 351.7066 | 0.295301 | 0.122663 | 2.407416 | 0.016066 | 0.049813 |
| CNAG_01500 | 5.39868 | -1.91447 | 0.971157 | -1.97133 | 0.048687 | 0.121974 |
| CNAG_01504 | 1281.122 | 0.295539 | 0.082712 | 3.57312 | 0.000353 | 0.001906 |
| CNAG_08003 | 563.5664 | -0.29133 | 0.104069 | -2.79936 | 0.00512 | 0.019156 |
| CNAG_01508 | 1619.176 | -0.23597 | 0.07558 | -3.12212 | 0.001796 | 0.007846 |
| CNAG_01509 | 644.5968 | -0.31695 | 0.099038 | -3.20024 | 0.001373 | 0.006245 |
| CNAG_01510 | 2918.21 | 0.169571 | 0.067345 | 2.517944 | 0.011804 | 0.038853 |
| CNAG_07589 | 55.93681 | -1.24418 | 0.28622 | -4.34694 | 1.38E-05 | 0.000102 |
| CNAG_07591 | 1540.376 | 0.426191 | 0.07833 | 5.440953 | 5.30E-08 | 5.57E-07 |
| CNAG_01519 | 349.2261 | 0.258633 | 0.119056 | 2.172359 | 0.029829 | 0.082811 |
| CNAG_01520 | 1307.303 | -0.21236 | 0.076205 | -2.78675 | 0.005324 | 0.019798 |
| CNAG_01521 | 672.1847 | 0.365598 | 0.102316 | 3.573217 | 0.000353 | 0.001906 |
| CNAG_01522 | 1703.212 | 0.456551 | 0.092456 | 4.938018 | 7.89E-07 | 6.97E-06 |
| CNAG_01524 | 1621.697 | -0.17436 | 0.075144 | -2.32035 | 0.020322 | 0.060394 |
| CNAG_01526 | 2993.557 | 0.201372 | 0.071982 | 2.797516 | 0.00515 | 0.019257 |
| CNAG_12968 | 38.93687 | 0.792526 | 0.328302 | 2.414016 | 0.015778 | 0.049114 |
| CNAG_01528 | 1029.392 | -0.36473 | 0.08359 | -4.36331 | 1.28E-05 | 9.51E-05 |
| CNAG_01529 | 1506.053 | 0.554425 | 0.084367 | 6.571561 | 4.98E-11 | 7.74E-10 |
| CNAG_01530 | 2666.866 | -0.19533 | 0.08083 | -2.41656 | 0.015668 | 0.04885 |
| CNAG_01534 | 100.4065 | -0.59693 | 0.21306 | -2.80171 | 0.005083 | 0.019072 |
| CNAG_01538 | 192.5798 | -0.38752 | 0.158872 | -2.43923 | 0.014719 | 0.046515 |
| CNAG_01539 | 11338.02 | 0.34469 | 0.080877 | 4.261875 | 2.03E-05 | 0.000143 |
| CNAG_01540 | 556.3178 | -0.3035 | 0.105188 | -2.88528 | 0.003911 | 0.015406 |
| CNAG_01543 | 1399.701 | 0.290322 | 0.086083 | 3.372591 | 0.000745 | 0.003683 |
| CNAG_01545 | 316.6877 | -0.52011 | 0.128604 | -4.04427 | 5.25E-05 | 0.000344 |
| CNAG_01546 | 368.9167 | -0.32924 | 0.124885 | -2.63633 | 0.008381 | 0.029274 |
| CNAG_01547 | 2059.195 | -0.19613 | 0.072723 | -2.69699 | 0.006997 | 0.025071 |
| CNAG_01548 | 5624.871 | -0.19313 | 0.062117 | -3.10911 | 0.001877 | 0.008163 |
| CNAG_01549 | 3066.667 | -0.13648 | 0.068548 | -1.99094 | 0.046487 | 0.117472 |

| | | | | | | |
|---|---|---|---|---|---|---|
| CNAG_01550 | 1361.35 | -0.27747 | 0.076525 | -3.62581 | 0.000288 | 0.001595 |
| CNAG_01553 | 1072.925 | -0.40477 | 0.084104 | -4.81275 | 1.49E-06 | 1.27E-05 |
| CNAG_01557 | 8954.141 | 0.157364 | 0.073799 | 2.132324 | 0.03298 | 0.089277 |
| CNAG_01558 | 21547.27 | 1.310809 | 0.080132 | 16.35822 | 3.80E-60 | 1.29E-57 |
| CNAG_01559 | 1905.852 | -0.19367 | 0.076822 | -2.52101 | 0.011702 | 0.038634 |
| CNAG_01562 | 30829.86 | 0.238742 | 0.098066 | 2.434514 | 0.014912 | 0.047019 |
| CNAG_01564 | 2995.909 | -0.58615 | 0.074692 | -7.84747 | 4.24E-15 | 9.83E-14 |
| CNAG_01565 | 2762.011 | 0.173466 | 0.069757 | 2.486729 | 0.012892 | 0.04183 |
| CNAG_01567 | 181.7357 | 0.631003 | 0.153854 | 4.101314 | 4.11E-05 | 0.000274 |
| CNAG_01569 | 928.4466 | 0.38417 | 0.085876 | 4.473521 | 7.69E-06 | 5.92E-05 |
| CNAG_01571 | 944.5061 | 0.166335 | 0.084478 | 1.968967 | 0.048957 | 0.122416 |
| CNAG_01572 | 571.746 | 0.425996 | 0.110563 | 3.852965 | 0.000117 | 0.000707 |
| CNAG_01573 | 1897.377 | 0.238432 | 0.07992 | 2.9834 | 0.002851 | 0.011721 |
| CNAG_01574 | 3611.734 | 0.661125 | 0.067698 | 9.76585 | 1.58E-22 | 7.04E-21 |
| CNAG_01576 | 1632.187 | 0.175605 | 0.076632 | 2.291535 | 0.021933 | 0.064226 |
| CNAG_01577 | 39339.86 | 0.26001 | 0.068126 | 3.816604 | 0.000135 | 0.000806 |
| CNAG_01578 | 2203.538 | 0.22059 | 0.085094 | 2.592312 | 0.009533 | 0.032552 |
| CNAG_01584 | 1008.751 | 0.731425 | 0.090304 | 8.099614 | 5.51E-16 | 1.36E-14 |
| CNAG_01585 | 662.5258 | 0.847679 | 0.099663 | 8.505409 | 1.81E-17 | 4.97E-16 |
| CNAG_01586 | 16727.67 | 0.257681 | 0.078077 | 3.300336 | 0.000966 | 0.004611 |
| CNAG_01588 | 5010.773 | 0.465005 | 0.073631 | 6.315326 | 2.70E-10 | 3.88E-09 |
| CNAG_01590 | 972.7519 | 0.275106 | 0.081824 | 3.362155 | 0.000773 | 0.003804 |
| CNAG_01593 | 3054.851 | -0.26142 | 0.079517 | -3.28755 | 0.001011 | 0.004803 |
| CNAG_01594 | 4847.249 | 0.452744 | 0.074769 | 6.055277 | 1.40E-09 | 1.83E-08 |
| CNAG_01597 | 1084.675 | 0.408187 | 0.086892 | 4.697651 | 2.63E-06 | 2.17E-05 |
| CNAG_01600 | 1562.321 | -0.60853 | 0.096817 | -6.28542 | 3.27E-10 | 4.64E-09 |
| CNAG_01604 | 546.8502 | 0.681981 | 0.102424 | 6.658436 | 2.77E-11 | 4.43E-10 |
| CNAG_01605 | 350.8729 | 0.411365 | 0.134877 | 3.049929 | 0.002289 | 0.009656 |
| CNAG_01608 | 1415.545 | -0.45966 | 0.081736 | -5.62378 | 1.87E-08 | 2.09E-07 |
| CNAG_01615 | 808.8551 | 0.21434 | 0.10262 | 2.088682 | 0.036736 | 0.09716 |
| CNAG_01616 | 2769.715 | -0.18927 | 0.078009 | -2.42621 | 0.015257 | 0.047934 |
| CNAG_07594 | 1186.031 | 0.398681 | 0.078422 | 5.083791 | 3.70E-07 | 3.45E-06 |
| CNAG_01619 | 624.5736 | 0.322104 | 0.0979 | 3.290126 | 0.001001 | 0.004762 |
| CNAG_12978 | 93.47274 | 1.157261 | 0.229895 | 5.033869 | 4.81E-07 | 4.43E-06 |
| CNAG_01621 | 11391.06 | 3.284435 | 0.085464 | 38.43044 | 0 | 0 |
| CNAG_01622 | 800.4136 | -0.28607 | 0.088741 | -3.22361 | 0.001266 | 0.005814 |
| CNAG_01623 | 989.0109 | -0.42285 | 0.097764 | -4.32517 | 1.52E-05 | 0.000111 |
| CNAG_01629 | 684.7445 | -0.27944 | 0.099061 | -2.82088 | 0.004789 | 0.018152 |
| CNAG_01631 | 1543.259 | -0.14556 | 0.073681 | -1.97549 | 0.048212 | 0.121038 |
| CNAG_01632 | 582.0845 | 0.267685 | 0.097873 | 2.735026 | 0.006238 | 0.022694 |
| CNAG_01634 | 3512.732 | -0.36307 | 0.06542 | -5.54993 | 2.86E-08 | 3.13E-07 |
| CNAG_01636 | 1184.568 | -0.16443 | 0.083499 | -1.96924 | 0.048925 | 0.122416 |
| CNAG_01638 | 824.0546 | -0.42226 | 0.094148 | -4.4851 | 7.29E-06 | 5.63E-05 |
| CNAG_01639 | 7009.463 | 0.510994 | 0.07673 | 6.659644 | 2.74E-11 | 4.41E-10 |
| CNAG_01641 | 4541.884 | -0.38388 | 0.074854 | -5.12833 | 2.92E-07 | 2.79E-06 |
| CNAG_01643 | 973.5021 | -0.34989 | 0.086078 | -4.06477 | 4.81E-05 | 0.000317 |
| CNAG_01644 | 2172.336 | -0.23013 | 0.072745 | -3.16345 | 0.001559 | 0.006953 |
| CNAG_01646 | 850.3099 | -0.19103 | 0.088383 | -2.16144 | 0.030662 | 0.084523 |

| | | | | | | |
|---|---|---|---|---|---|---|
| CNAG_01653 | 142.7356 | -0.45098 | 0.199355 | -2.26218 | 0.023686 | 0.068285 |
| CNAG_01654 | 1131.069 | -0.33037 | 0.090001 | -3.67077 | 0.000242 | 0.001364 |
| CNAG_01656 | 541.6764 | 0.487224 | 0.10644 | 4.577442 | 4.71E-06 | 3.72E-05 |
| CNAG_01657 | 8907.886 | 0.474733 | 0.06697 | 7.088769 | 1.35E-12 | 2.53E-11 |
| CNAG_08004 | 125.4834 | 0.417223 | 0.19549 | 2.134248 | 0.032822 | 0.089002 |
| CNAG_01665 | 727.6429 | -0.28014 | 0.091136 | -3.07386 | 0.002113 | 0.009006 |
| CNAG_01673 | 1662.99 | -0.23454 | 0.09226 | -2.54211 | 0.011018 | 0.036684 |
| CNAG_01674 | 59.21388 | -0.53167 | 0.256699 | -2.07119 | 0.038341 | 0.100711 |
| CNAG_01675 | 880.1141 | -0.25388 | 0.091264 | -2.7818 | 0.005406 | 0.020051 |
| CNAG_01676 | 1695.894 | -0.48224 | 0.087624 | -5.50355 | 3.72E-08 | 4.02E-07 |
| CNAG_01679 | 4062.123 | -0.18814 | 0.066734 | -2.81922 | 0.004814 | 0.018237 |
| CNAG_12984 | 26.08708 | 0.838062 | 0.392029 | 2.137754 | 0.032537 | 0.088322 |
| CNAG_12985 | 1169.715 | 0.299003 | 0.091758 | 3.258585 | 0.00112 | 0.005232 |
| CNAG_01681 | 6447.551 | -0.20779 | 0.061689 | -3.36836 | 0.000756 | 0.003732 |
| CNAG_01682 | 11774.08 | 0.199351 | 0.057562 | 3.463206 | 0.000534 | 0.002764 |
| CNAG_12988 | 249.2817 | -0.43093 | 0.136362 | -3.16017 | 0.001577 | 0.00702 |
| CNAG_01683 | 10569.82 | 0.276428 | 0.0647 | 4.272427 | 1.93E-05 | 0.000137 |
| CNAG_01686 | 2477.27 | 0.559411 | 0.079507 | 7.035975 | 1.98E-12 | 3.62E-11 |
| CNAG_01687 | 1977.129 | -0.44919 | 0.077807 | -5.7731 | 7.78E-09 | 9.24E-08 |
| CNAG_01691 | 1193.256 | 0.622453 | 0.089174 | 6.980228 | 2.95E-12 | 5.22E-11 |
| CNAG_12992 | 137.0998 | 0.432918 | 0.176614 | 2.451213 | 0.014238 | 0.04524 |
| CNAG_01694 | 870.1026 | -0.3416 | 0.131239 | -2.60289 | 0.009244 | 0.031817 |
| CNAG_01696 | 4301.246 | 0.294663 | 0.071977 | 4.093853 | 4.24E-05 | 0.000282 |
| CNAG_08005 | 149.0841 | 0.574309 | 0.175018 | 3.281435 | 0.001033 | 0.004892 |
| CNAG_01708 | 1078.916 | -0.30623 | 0.100673 | -3.04187 | 0.002351 | 0.009875 |
| CNAG_01709 | 4242.506 | -0.24413 | 0.066502 | -3.67101 | 0.000242 | 0.001364 |
| CNAG_01711 | 4273.93 | 0.17461 | 0.062212 | 2.806682 | 0.005005 | 0.018834 |
| CNAG_01712 | 1115.619 | -0.35244 | 0.088544 | -3.98039 | 6.88E-05 | 0.000442 |
| CNAG_01713 | 1661.867 | 0.679268 | 0.086268 | 7.873951 | 3.44E-15 | 8.03E-14 |
| CNAG_01715 | 3709.991 | -0.55784 | 0.086629 | -6.43943 | 1.20E-10 | 1.79E-09 |
| CNAG_01721 | 3512.759 | 0.352225 | 0.079247 | 4.444636 | 8.80E-06 | 6.70E-05 |
| CNAG_01726 | 5335.235 | -0.18007 | 0.074298 | -2.42365 | 0.015366 | 0.048138 |
| CNAG_01727 | 44008.05 | -0.25773 | 0.070331 | -3.66452 | 0.000248 | 0.001393 |
| CNAG_01728 | 1207.656 | 0.378471 | 0.078943 | 4.794214 | 1.63E-06 | 1.39E-05 |
| CNAG_01729 | 1473.408 | -0.71571 | 0.093362 | -7.66594 | 1.78E-14 | 3.91E-13 |
| CNAG_01730 | 621.8661 | 0.242259 | 0.100349 | 2.414167 | 0.015771 | 0.049114 |
| CNAG_12998 | 20.62416 | 0.933476 | 0.434769 | 2.147061 | 0.031788 | 0.086815 |
| CNAG_01732 | 1675.243 | 0.23106 | 0.078251 | 2.952796 | 0.003149 | 0.012767 |
| CNAG_01735 | 1902.65 | 0.440252 | 0.077944 | 5.648279 | 1.62E-08 | 1.84E-07 |
| CNAG_01737 | 3657.343 | -0.70925 | 0.078345 | -9.05284 | 1.39E-19 | 4.92E-18 |
| CNAG_01738 | 456.36 | -0.3229 | 0.11288 | -2.86059 | 0.004229 | 0.016385 |
| CNAG_01742 | 180.1257 | 1.018947 | 0.156347 | 6.517201 | 7.16E-11 | 1.10E-09 |
| CNAG_01744 | 11131.68 | 0.298231 | 0.069577 | 4.28636 | 1.82E-05 | 0.000131 |
| CNAG_01745 | 11322.59 | 0.431255 | 0.061949 | 6.961432 | 3.37E-12 | 5.95E-11 |
| CNAG_01746 | 296.4644 | -0.38812 | 0.136214 | -2.84934 | 0.004381 | 0.01685 |
| CNAG_07599 | 1133.941 | -0.2092 | 0.088699 | -2.35857 | 0.018346 | 0.055428 |
| CNAG_01749 | 693.0057 | -0.78368 | 0.103099 | -7.60125 | 2.93E-14 | 6.34E-13 |
| CNAG_01751 | 1044.635 | 0.784926 | 0.106985 | 7.336771 | 2.19E-13 | 4.48E-12 |

| | | | | | | |
|---|---|---|---|---|---|---|
| CNAG_01752 | 7352.948 | 0.147432 | 0.059904 | 2.461153 | 0.013849 | 0.044204 |
| CNAG_01753 | 5899.009 | 0.538409 | 0.07246 | 7.430427 | 1.08E-13 | 2.25E-12 |
| CNAG_01754 | 1714.531 | -0.24577 | 0.075186 | -3.26885 | 0.00108 | 0.005073 |
| CNAG_01755 | 3037.444 | -0.4547 | 0.07177 | -6.33552 | 2.37E-10 | 3.42E-09 |
| CNAG_01756 | 738.5464 | -0.25821 | 0.092609 | -2.78814 | 0.005301 | 0.019747 |
| CNAG_08006 | 229.2531 | -0.3999 | 0.142323 | -2.80981 | 0.004957 | 0.018688 |
| CNAG_07601 | 497.2994 | -0.32337 | 0.113703 | -2.84395 | 0.004456 | 0.017104 |
| CNAG_01761 | 20527.06 | 0.826015 | 0.066959 | 12.33621 | 5.78E-35 | 5.57E-33 |
| CNAG_01764 | 813.3889 | -0.29634 | 0.099656 | -2.97365 | 0.002943 | 0.012049 |
| CNAG_01765 | 1147.517 | -0.32645 | 0.084158 | -3.87901 | 0.000105 | 0.000644 |
| CNAG_01769 | 3220.512 | 0.395722 | 0.073874 | 5.356735 | 8.47E-08 | 8.67E-07 |
| CNAG_01772 | 3395.297 | 0.152768 | 0.066351 | 2.302428 | 0.021311 | 0.06283 |
| CNAG_01777 | 1885.56 | 0.389714 | 0.081774 | 4.765726 | 1.88E-06 | 1.58E-05 |
| CNAG_01779 | 771.385 | 0.248113 | 0.092885 | 2.671193 | 0.007558 | 0.026813 |
| CNAG_01780 | 4801.815 | -0.27229 | 0.079858 | -3.40965 | 0.00065 | 0.003279 |
| CNAG_01788 | 1912.935 | -0.29128 | 0.080281 | -3.62829 | 0.000285 | 0.001586 |
| CNAG_01790 | 846.9417 | -0.26025 | 0.086056 | -3.0242 | 0.002493 | 0.01042 |
| CNAG_01791 | 1989.866 | -0.51649 | 0.076541 | -6.74782 | 1.50E-11 | 2.51E-10 |
| CNAG_01792 | 1428.294 | -0.33154 | 0.094041 | -3.52553 | 0.000423 | 0.002242 |
| CNAG_13006 | 128.4006 | -0.785 | 0.187963 | -4.17636 | 2.96E-05 | 0.000203 |
| CNAG_01793 | 652.1761 | -0.3888 | 0.096497 | -4.02916 | 5.60E-05 | 0.000365 |
| CNAG_01794 | 368.8807 | -1.86192 | 0.141017 | -13.2035 | 8.37E-40 | 1.13E-37 |
| CNAG_01795 | 385.4634 | 0.269304 | 0.117923 | 2.283725 | 0.022388 | 0.065216 |
| CNAG_01799 | 10182.6 | 0.16224 | 0.074864 | 2.167115 | 0.030226 | 0.083617 |
| CNAG_01800 | 1917.121 | 0.306008 | 0.087766 | 3.486634 | 0.000489 | 0.00256 |
| CNAG_01803 | 450.5587 | -0.29256 | 0.126372 | -2.31508 | 0.020608 | 0.061058 |
| CNAG_01807 | 860.1585 | -0.42285 | 0.118827 | -3.55852 | 0.000373 | 0.002 |
| CNAG_01808 | 2420.485 | 0.214891 | 0.070611 | 3.043294 | 0.00234 | 0.009834 |
| CNAG_01810 | 547.2964 | -0.56517 | 0.102477 | -5.51511 | 3.49E-08 | 3.78E-07 |
| CNAG_01812 | 3593.092 | -0.34271 | 0.076757 | -4.46488 | 8.01E-06 | 6.14E-05 |
| CNAG_01813 | 11597.53 | -0.15733 | 0.071937 | -2.18708 | 0.028737 | 0.080533 |
| CNAG_01818 | 1740.336 | 0.202945 | 0.078912 | 2.571793 | 0.010117 | 0.034172 |
| CNAG_01821 | 933.6063 | 0.386085 | 0.103058 | 3.746286 | 0.000179 | 0.001037 |
| CNAG_01822 | 1937.025 | 0.477328 | 0.075342 | 6.335521 | 2.37E-10 | 3.42E-09 |
| CNAG_01823 | 2922.339 | 0.191292 | 0.071085 | 2.691051 | 0.007123 | 0.025428 |
| CNAG_13008 | 157.0175 | -0.8543 | 0.172298 | -4.95828 | 7.11E-07 | 6.33E-06 |
| CNAG_07608 | 1510.373 | -0.26995 | 0.080147 | -3.36822 | 0.000757 | 0.003732 |
| CNAG_07609 | 1547.856 | -0.5874 | 0.074711 | -7.86229 | 3.77E-15 | 8.76E-14 |
| CNAG_01833 | 3874.384 | -0.26453 | 0.067325 | -3.92919 | 8.52E-05 | 0.000535 |
| CNAG_01836 | 128.4075 | -0.3816 | 0.193943 | -1.96758 | 0.049117 | 0.122776 |
| CNAG_13010 | 151.9142 | 0.377381 | 0.162735 | 2.31899 | 0.020396 | 0.060543 |
| CNAG_01839 | 2905.109 | 0.153927 | 0.070866 | 2.172072 | 0.02985 | 0.082842 |
| CNAG_01842 | 1861.129 | -0.52645 | 0.077023 | -6.83494 | 8.20E-12 | 1.41E-10 |
| CNAG_01843 | 2674.274 | 0.607146 | 0.077167 | 7.867943 | 3.61E-15 | 8.40E-14 |
| CNAG_01844 | 304.8015 | -0.34071 | 0.131686 | -2.58732 | 0.009673 | 0.032977 |
| CNAG_01846 | 6351.753 | 1.151701 | 0.073945 | 15.57516 | 1.07E-54 | 2.99E-52 |
| CNAG_01850 | 2789.441 | -0.19943 | 0.08237 | -2.42117 | 0.01547 | 0.048409 |
| CNAG_01851 | 37.67881 | 1.108748 | 0.319911 | 3.4658 | 0.000529 | 0.002744 |

| | | | | | | |
|---|---|---|---|---|---|---|
| CNAG_01860 | 759.0953 | 0.195398 | 0.090662 | 2.155238 | 0.031143 | 0.085514 |
| CNAG_13013 | 191.5037 | 0.465943 | 0.156942 | 2.968886 | 0.002989 | 0.012212 |
| CNAG_01865 | 492.2717 | -0.72559 | 0.108824 | -6.66757 | 2.60E-11 | 4.22E-10 |
| CNAG_01866 | 517.9023 | -0.37504 | 0.118111 | -3.17533 | 0.001497 | 0.006717 |
| CNAG_01867 | 2263.843 | 0.218293 | 0.070982 | 3.075328 | 0.002103 | 0.008967 |
| CNAG_01870 | 4867.491 | 0.333375 | 0.076049 | 4.383667 | 1.17E-05 | 8.73E-05 |
| CNAG_01874 | 1428.134 | 0.235389 | 0.107829 | 2.182977 | 0.029037 | 0.081105 |
| CNAG_01876 | 1426.028 | 0.473367 | 0.086995 | 5.441308 | 5.29E-08 | 5.57E-07 |
| CNAG_01877 | 9215.08 | -0.42314 | 0.075094 | -5.63488 | 1.75E-08 | 1.98E-07 |
| CNAG_01880 | 1412.552 | -0.19579 | 0.086977 | -2.25111 | 0.024379 | 0.070074 |
| CNAG_01881 | 5100.012 | 1.048402 | 0.068014 | 15.41443 | 1.31E-53 | 3.30E-51 |
| CNAG_01883 | 91.90509 | -0.91467 | 0.220553 | -4.14716 | 3.37E-05 | 0.000228 |
| CNAG_01884 | 65613.32 | -0.16259 | 0.080197 | -2.02734 | 0.042628 | 0.109937 |
| CNAG_01885 | 1353.718 | -0.39783 | 0.081532 | -4.87947 | 1.06E-06 | 9.19E-06 |
| CNAG_01890 | 54408.34 | -0.22875 | 0.058472 | -3.91223 | 9.14E-05 | 0.000567 |
| CNAG_01892 | 1851.335 | 0.371512 | 0.097253 | 3.820044 | 0.000133 | 0.000796 |
| CNAG_01893 | 1800.506 | 0.254586 | 0.079166 | 3.215859 | 0.001301 | 0.005958 |
| CNAG_01894 | 2784.456 | -0.14813 | 0.070339 | -2.10587 | 0.035216 | 0.094041 |
| CNAG_01898 | 2250.626 | -0.4757 | 0.071272 | -6.67448 | 2.48E-11 | 4.05E-10 |
| CNAG_01900 | 1123.55 | 0.361555 | 0.08439 | 4.284347 | 1.83E-05 | 0.000132 |
| CNAG_01902 | 2931.502 | -0.31572 | 0.068882 | -4.58344 | 4.57E-06 | 3.64E-05 |
| CNAG_01907 | 1914.979 | -0.30538 | 0.120363 | -2.53716 | 0.011176 | 0.037097 |
| CNAG_13021 | 375.7217 | 0.548075 | 0.134377 | 4.078654 | 4.53E-05 | 0.0003 |
| CNAG_01914 | 1552.272 | 0.330539 | 0.078153 | 4.229384 | 2.34E-05 | 0.000163 |
| CNAG_01915 | 9261.154 | 0.49094 | 0.074043 | 6.630458 | 3.35E-11 | 5.29E-10 |
| CNAG_01919 | 791.2906 | 0.242586 | 0.096901 | 2.503445 | 0.012299 | 0.040177 |
| CNAG_01920 | 13995.14 | -0.39367 | 0.067559 | -5.82705 | 5.64E-09 | 6.84E-08 |
| CNAG_01921 | 94.28011 | 0.634562 | 0.203934 | 3.111597 | 0.001861 | 0.008104 |
| CNAG_01922 | 1148.569 | 0.336943 | 0.080044 | 4.209481 | 2.56E-05 | 0.000178 |
| CNAG_01923 | 2339.581 | -0.21964 | 0.068673 | -3.19841 | 0.001382 | 0.006266 |
| CNAG_01924 | 1193 | 0.665782 | 0.081418 | 8.177359 | 2.90E-16 | 7.40E-15 |
| CNAG_01925 | 1027.715 | 1.383639 | 0.130282 | 10.62038 | 2.40E-26 | 1.43E-24 |
| CNAG_01926 | 473.4197 | 0.323604 | 0.112735 | 2.870479 | 0.004099 | 0.015993 |
| CNAG_01929 | 245.6949 | -0.58558 | 0.14062 | -4.16428 | 3.12E-05 | 0.000213 |
| CNAG_01937 | 91.63702 | -0.47434 | 0.220313 | -2.15301 | 0.031318 | 0.085877 |
| CNAG_01939 | 178.2005 | 0.521978 | 0.183413 | 2.845924 | 0.004428 | 0.017007 |
| CNAG_01941 | 1677.622 | -0.15869 | 0.079384 | -1.99903 | 0.045605 | 0.115903 |
| CNAG_01942 | 403.6559 | 0.707366 | 0.118564 | 5.966126 | 2.43E-09 | 3.09E-08 |
| CNAG_01943 | 2714.954 | -0.19751 | 0.075608 | -2.61232 | 0.008993 | 0.031038 |
| CNAG_01945 | 85.64949 | 0.437455 | 0.219691 | 1.99123 | 0.046456 | 0.117441 |
| CNAG_01946 | 1366.639 | 1.388791 | 0.079043 | 17.57016 | 4.17E-69 | 2.32E-66 |
| CNAG_01947 | 570.4618 | -0.9859 | 0.114881 | -8.58192 | 9.33E-18 | 2.66E-16 |
| CNAG_01948 | 1001.905 | -0.35268 | 0.093309 | -3.77975 | 0.000157 | 0.00092 |
| CNAG_01949 | 1236.523 | -1.23587 | 0.089574 | -13.7971 | 2.65E-43 | 4.14E-41 |
| CNAG_01952 | 446.3083 | 0.449765 | 0.116034 | 3.876161 | 0.000106 | 0.000649 |
| CNAG_01953 | 230.1651 | -0.87162 | 0.163434 | -5.33318 | 9.65E-08 | 9.80E-07 |
| CNAG_13035 | 32.48194 | -0.86781 | 0.361102 | -2.40322 | 0.016251 | 0.050249 |
| CNAG_13036 | 50.38923 | -0.88544 | 0.281158 | -3.14926 | 0.001637 | 0.00725 |

| | | | | | | |
|---|---|---|---|---|---|---|
| CNAG_01957 | 3160.701 | -0.28977 | 0.072414 | -4.00166 | 6.29E-05 | 0.000407 |
| CNAG_01958 | 2893.187 | -0.21604 | 0.066763 | -3.23593 | 0.001212 | 0.005602 |
| CNAG_01959 | 2285.706 | -0.30881 | 0.098036 | -3.14999 | 0.001633 | 0.00724 |
| CNAG_01961 | 6069.674 | 0.156788 | 0.067777 | 2.313274 | 0.020708 | 0.061329 |
| CNAG_01962 | 996.1835 | 0.27524 | 0.091056 | 3.022748 | 0.002505 | 0.010462 |
| CNAG_13037 | 50.24673 | 1.556525 | 0.318214 | 4.891449 | 1.00E-06 | 8.69E-06 |
| CNAG_01970 | 1897.529 | 0.433518 | 0.072158 | 6.007931 | 1.88E-09 | 2.41E-08 |
| CNAG_01974 | 1040.867 | 0.215211 | 0.086352 | 2.492262 | 0.012693 | 0.041285 |
| CNAG_01976 | 34288 | -0.18854 | 0.073133 | -2.57797 | 0.009938 | 0.033683 |
| CNAG_01981 | 6480.797 | -0.34368 | 0.066618 | -5.15897 | 2.48E-07 | 2.39E-06 |
| CNAG_01982 | 573.7327 | 0.366735 | 0.110335 | 3.32383 | 0.000888 | 0.004281 |
| CNAG_01984 | 52940.18 | 0.556718 | 0.060398 | 9.217553 | 3.04E-20 | 1.12E-18 |
| CNAG_01985 | 5473.857 | 0.160272 | 0.065857 | 2.433612 | 0.014949 | 0.047098 |
| CNAG_01986 | 1161.598 | 0.252836 | 0.084637 | 2.987315 | 0.002814 | 0.011603 |
| CNAG_13042 | 78.31475 | 0.813608 | 0.236198 | 3.444602 | 0.000572 | 0.00294 |
| CNAG_01987 | 1802.388 | 0.203431 | 0.083807 | 2.427365 | 0.015209 | 0.047801 |
| CNAG_01989 | 251.2613 | 0.31817 | 0.145898 | 2.180765 | 0.029201 | 0.081532 |
| CNAG_01990 | 51349.43 | -0.19812 | 0.078075 | -2.53758 | 0.011162 | 0.037073 |
| CNAG_01991 | 11756.27 | 0.268798 | 0.073189 | 3.672671 | 0.00024 | 0.001356 |
| CNAG_01993 | 153.0393 | 0.399556 | 0.166212 | 2.403898 | 0.016221 | 0.050195 |
| CNAG_01996 | 441.3278 | -0.3615 | 0.134082 | -2.69609 | 0.007016 | 0.025116 |
| CNAG_01997 | 2986.696 | -0.27083 | 0.074442 | -3.63817 | 0.000275 | 0.00153 |
| CNAG_02000 | 1958.509 | 0.308122 | 0.080639 | 3.821017 | 0.000133 | 0.000794 |
| CNAG_02004 | 2797.411 | 0.182919 | 0.068821 | 2.657905 | 0.007863 | 0.027803 |
| CNAG_02006 | 1323.295 | 0.332319 | 0.084965 | 3.911227 | 9.18E-05 | 0.000569 |
| CNAG_02008 | 2291.148 | 0.454025 | 0.101042 | 4.493429 | 7.01E-06 | 5.43E-05 |
| CNAG_02009 | 1023.195 | 0.380962 | 0.098199 | 3.879507 | 0.000105 | 0.000643 |
| CNAG_02011 | 281.6375 | -0.31181 | 0.128453 | -2.42744 | 0.015206 | 0.047801 |
| CNAG_07619 | 1039.805 | 0.419151 | 0.094511 | 4.434925 | 9.21E-06 | 6.99E-05 |
| CNAG_02016 | 12.24353 | -2.01169 | 0.63478 | -3.16911 | 0.001529 | 0.006838 |
| CNAG_07026 | 45.85985 | 1.036309 | 0.299635 | 3.458574 | 0.000543 | 0.002807 |
| CNAG_05986 | 52.6927 | 0.531706 | 0.265982 | 1.99903 | 0.045605 | 0.115903 |
| CNAG_05989 | 777.6945 | 0.254947 | 0.08912 | 2.860715 | 0.004227 | 0.016385 |
| CNAG_05990 | 396.0123 | 0.266481 | 0.119984 | 2.220968 | 0.026353 | 0.074894 |
| CNAG_05992 | 149.1987 | 0.405095 | 0.171231 | 2.365785 | 0.017992 | 0.054613 |
| CNAG_05993 | 886.0099 | 0.820564 | 0.087828 | 9.34282 | 9.38E-21 | 3.64E-19 |
| CNAG_05994 | 2147.641 | 0.943503 | 0.095737 | 9.855132 | 6.51E-23 | 3.08E-21 |
| CNAG_06000 | 2493.801 | 0.656138 | 0.069652 | 9.420263 | 4.50E-21 | 1.80E-19 |
| CNAG_06001 | 1070.688 | -0.36073 | 0.083815 | -4.30389 | 1.68E-05 | 0.000121 |
| CNAG_06005 | 1991.485 | -0.69154 | 0.074654 | -9.26332 | 1.98E-20 | 7.47E-19 |
| CNAG_06006 | 1280.428 | -0.18436 | 0.087668 | -2.10293 | 0.035472 | 0.094543 |
| CNAG_06007 | 1103.346 | -0.60894 | 0.084236 | -7.22902 | 4.87E-13 | 9.54E-12 |
| CNAG_06009 | 88.1688 | 0.723231 | 0.234025 | 3.090397 | 0.001999 | 0.008604 |
| CNAG_06011 | 85.50082 | -0.61631 | 0.24785 | -2.48662 | 0.012896 | 0.04183 |
| CNAG_06012 | 4908.301 | 0.216859 | 0.073867 | 2.935808 | 0.003327 | 0.013417 |
| CNAG_06013 | 638.8945 | 0.231728 | 0.104234 | 2.223147 | 0.026206 | 0.074557 |
| CNAG_06018 | 1336.097 | 0.182546 | 0.090812 | 2.010151 | 0.044415 | 0.113645 |
| CNAG_06022 | 511.8131 | 0.268015 | 0.115178 | 2.326952 | 0.019968 | 0.059545 |

| | | | | | | |
|---|---|---|---|---|---|---|
| CNAG_06026 | 1621.651 | -0.16087 | 0.080614 | -1.99551 | 0.045987 | 0.116557 |
| CNAG_06029 | 411.3766 | -0.42895 | 0.108888 | -3.93935 | 8.17E-05 | 0.000516 |
| CNAG_06030 | 561.3421 | 0.229972 | 0.103292 | 2.226435 | 0.025985 | 0.07401 |
| CNAG_06031 | 274.8192 | -0.38307 | 0.129816 | -2.95083 | 0.003169 | 0.012842 |
| CNAG_06032 | 582.2685 | 0.227487 | 0.105498 | 2.156312 | 0.031059 | 0.085347 |
| CNAG_06033 | 1065.05 | -0.88204 | 0.098811 | -8.92655 | 4.39E-19 | 1.44E-17 |
| CNAG_06048 | 1186.222 | 0.221256 | 0.083763 | 2.641461 | 0.008255 | 0.029019 |
| CNAG_13055 | 78.66051 | 0.802679 | 0.229554 | 3.496686 | 0.000471 | 0.002474 |
| CNAG_06050 | 187.2083 | 0.968027 | 0.173735 | 5.571874 | 2.52E-08 | 2.77E-07 |
| CNAG_06051 | 1012.147 | 0.46369 | 0.086396 | 5.367014 | 8.01E-08 | 8.21E-07 |
| CNAG_06052 | 1164.869 | 0.580289 | 0.093548 | 6.203099 | 5.54E-10 | 7.55E-09 |
| CNAG_07897 | 109.483 | 0.484705 | 0.217566 | 2.227852 | 0.02589 | 0.073821 |
| CNAG_13058 | 12.0326 | -1.29071 | 0.574992 | -2.24474 | 0.024785 | 0.07111 |
| CNAG_06061 | 5724.819 | -0.19758 | 0.064537 | -3.06144 | 0.002203 | 0.009332 |
| CNAG_06063 | 2636.711 | 1.407243 | 0.072405 | 19.43576 | 3.85E-84 | 3.75E-81 |
| CNAG_06064 | 951.1739 | 0.490513 | 0.090332 | 5.430086 | 5.63E-08 | 5.90E-07 |
| CNAG_06068 | 1958.584 | -0.30653 | 0.079469 | -3.85727 | 0.000115 | 0.000695 |
| CNAG_07902 | 1358.902 | 0.53374 | 0.086012 | 6.205438 | 5.45E-10 | 7.48E-09 |
| CNAG_06071 | 568.5196 | -0.23066 | 0.104344 | -2.21059 | 0.027064 | 0.076695 |
| CNAG_06073 | 57.15661 | -0.76528 | 0.267212 | -2.86393 | 0.004184 | 0.016262 |
| CNAG_06074 | 1214.771 | 1.048819 | 0.103313 | 10.15183 | 3.25E-24 | 1.73E-22 |
| CNAG_06075 | 11593.76 | 0.945746 | 0.078229 | 12.08942 | 1.20E-33 | 1.05E-31 |
| CNAG_06076 | 2462.182 | -0.20406 | 0.081439 | -2.50566 | 0.012222 | 0.039959 |
| CNAG_06078 | 888.082 | 0.320566 | 0.092334 | 3.471811 | 0.000517 | 0.002691 |
| CNAG_06081 | 7688.584 | -0.40803 | 0.08126 | -5.02128 | 5.13E-07 | 4.72E-06 |
| CNAG_13062 | 124.5244 | -3.29447 | 0.257742 | -12.782 | 2.07E-37 | 2.48E-35 |
| CNAG_06082 | 82.42895 | -4.42006 | 0.374413 | -11.8053 | 3.66E-32 | 2.98E-30 |
| CNAG_06085 | 766.6134 | 0.510334 | 0.097809 | 5.217655 | 1.81E-07 | 1.77E-06 |
| CNAG_06087 | 1609.556 | -0.15334 | 0.077063 | -1.98979 | 0.046614 | 0.11765 |
| CNAG_06088 | 4246.607 | 0.15404 | 0.073469 | 2.096662 | 0.036024 | 0.095785 |
| CNAG_06090 | 2778.83 | -0.35172 | 0.070405 | -4.9956 | 5.87E-07 | 5.32E-06 |
| CNAG_06095 | 90047.36 | -0.24887 | 0.068255 | -3.64624 | 0.000266 | 0.001488 |
| CNAG_06096 | 11786.51 | -0.25344 | 0.06331 | -4.0031 | 6.25E-05 | 0.000405 |
| CNAG_06097 | 504.1273 | -0.25768 | 0.118485 | -2.17482 | 0.029644 | 0.082407 |
| CNAG_06098 | 722.5889 | 0.43259 | 0.103819 | 4.166786 | 3.09E-05 | 0.000211 |
| CNAG_06100 | 874.1988 | 0.189365 | 0.095992 | 1.972722 | 0.048527 | 0.121653 |
| CNAG_06101 | 136546.9 | 0.330018 | 0.070778 | 4.662708 | 3.12E-06 | 2.54E-05 |
| CNAG_13067 | 110.8122 | -0.88568 | 0.204022 | -4.34108 | 1.42E-05 | 0.000104 |
| CNAG_06102 | 89.8804 | -0.46024 | 0.210441 | -2.18704 | 0.02874 | 0.080533 |
| CNAG_06105 | 116.8514 | -0.56878 | 0.237315 | -2.39673 | 0.016542 | 0.050985 |
| CNAG_06107 | 855.5587 | -0.23485 | 0.096255 | -2.43989 | 0.014692 | 0.046457 |
| CNAG_06109 | 13557.05 | 1.137073 | 0.09475 | 12.00071 | 3.52E-33 | 3.05E-31 |
| CNAG_07903 | 931.9005 | 0.217712 | 0.111007 | 1.961251 | 0.04985 | 0.12425 |
| CNAG_06112 | 13817.14 | 0.203873 | 0.064036 | 3.183741 | 0.001454 | 0.006558 |
| CNAG_06113 | 69847.18 | -0.24009 | 0.076386 | -3.14317 | 0.001671 | 0.00739 |
| CNAG_06114 | 2963.808 | -0.25631 | 0.069849 | -3.66948 | 0.000243 | 0.001369 |
| CNAG_06115 | 374.5759 | 0.303876 | 0.118515 | 2.564036 | 0.010346 | 0.034773 |
| CNAG_08016 | 1619.872 | 0.218175 | 0.099694 | 2.188444 | 0.028637 | 0.080333 |

| | | | | | | |
|---|---|---|---|---|---|---|
| CNAG_06119 | 173.05 | -0.36494 | 0.171054 | -2.13347 | 0.032886 | 0.089081 |
| CNAG_06121 | 1248.668 | 0.192872 | 0.096398 | 2.000786 | 0.045415 | 0.115598 |
| CNAG_06122 | 1011.282 | -0.24736 | 0.098341 | -2.51532 | 0.011893 | 0.039095 |
| CNAG_06125 | 316427.1 | -0.17056 | 0.066126 | -2.57932 | 0.009899 | 0.033575 |
| CNAG_06126 | 1360.571 | -0.55715 | 0.076083 | -7.32295 | 2.43E-13 | 4.96E-12 |
| CNAG_06127 | 3246.753 | -0.45207 | 0.072476 | -6.23746 | 4.45E-10 | 6.19E-09 |
| CNAG_06132 | 88.51604 | -0.6215 | 0.230672 | -2.69431 | 0.007053 | 0.025215 |
| CNAG_06134 | 4924.965 | -0.79272 | 0.063212 | -12.5407 | 4.47E-36 | 4.72E-34 |
| CNAG_06136 | 557.5783 | -0.31375 | 0.120563 | -2.60237 | 0.009258 | 0.031817 |
| CNAG_06138 | 2634.7 | 0.205791 | 0.079575 | 2.586116 | 0.009706 | 0.033064 |
| CNAG_06139 | 154.712 | 0.831071 | 0.167384 | 4.965051 | 6.87E-07 | 6.15E-06 |
| CNAG_06144 | 28232.05 | -0.14508 | 0.064376 | -2.25356 | 0.024224 | 0.069706 |
| CNAG_06149 | 466.1712 | -0.28567 | 0.106954 | -2.671 | 0.007563 | 0.026814 |
| CNAG_06150 | 10023.67 | -0.199 | 0.068752 | -2.89447 | 0.003798 | 0.015007 |
| CNAG_06152 | 408.8714 | 0.437907 | 0.116401 | 3.762065 | 0.000169 | 0.00098 |
| CNAG_06153 | 4285.008 | -0.32126 | 0.065803 | -4.8822 | 1.05E-06 | 9.09E-06 |
| CNAG_06154 | 708.168 | 0.292048 | 0.094347 | 3.095463 | 0.001965 | 0.008491 |
| CNAG_06156 | 918.2017 | -0.46189 | 0.128309 | -3.59984 | 0.000318 | 0.001736 |
| CNAG_06157 | 1140.34 | -0.47109 | 0.084361 | -5.58423 | 2.35E-08 | 2.60E-07 |
| CNAG_06158 | 2140.65 | 0.276757 | 0.103854 | 2.664857 | 0.007702 | 0.027272 |
| CNAG_06163 | 60.69805 | 0.848811 | 0.255374 | 3.323799 | 0.000888 | 0.004281 |
| CNAG_06169 | 964.34 | 1.248855 | 0.098736 | 12.64847 | 1.14E-36 | 1.27E-34 |
| CNAG_06173 | 740.5733 | -0.32959 | 0.101301 | -3.25355 | 0.00114 | 0.00531 |
| CNAG_06174 | 2383.159 | -0.34305 | 0.096934 | -3.53903 | 0.000402 | 0.002141 |
| CNAG_06175 | 4974.033 | -0.22975 | 0.06933 | -3.3139 | 0.00092 | 0.004418 |
| CNAG_07908 | 5721.097 | -0.15363 | 0.065643 | -2.34038 | 0.019264 | 0.057733 |
| CNAG_06186 | 559.7971 | 0.279737 | 0.102839 | 2.720154 | 0.006525 | 0.023608 |
| CNAG_07911 | 293.8715 | 0.83017 | 0.127792 | 6.496286 | 8.23E-11 | 1.25E-09 |
| CNAG_06188 | 702.2964 | 0.502166 | 0.100412 | 5.001033 | 5.70E-07 | 5.19E-06 |
| CNAG_06189 | 34.85785 | 0.6827 | 0.329179 | 2.073946 | 0.038084 | 0.100138 |
| CNAG_06190 | 1327.807 | 0.758286 | 0.083841 | 9.044347 | 1.51E-19 | 5.29E-18 |
| CNAG_06192 | 1728.255 | -0.17776 | 0.08185 | -2.17174 | 0.029875 | 0.082853 |
| CNAG_06197 | 1037.257 | -0.19015 | 0.088832 | -2.14059 | 0.032307 | 0.08794 |
| CNAG_06200 | 901.8551 | -0.25678 | 0.090956 | -2.82313 | 0.004756 | 0.018087 |
| CNAG_06201 | 1430.694 | 0.569053 | 0.097805 | 5.818242 | 5.95E-09 | 7.17E-08 |
| CNAG_06203 | 168.5899 | 0.445211 | 0.162522 | 2.739398 | 0.006155 | 0.022446 |
| CNAG_06204 | 4644.678 | 0.178333 | 0.082175 | 2.170168 | 0.029994 | 0.083064 |
| CNAG_06205 | 3783.48 | -0.285 | 0.063079 | -4.51809 | 6.24E-06 | 4.85E-05 |
| CNAG_06206 | 2559.782 | -0.49224 | 0.07073 | -6.95944 | 3.42E-12 | 6.02E-11 |
| CNAG_13083 | 76.92001 | 0.502735 | 0.236269 | 2.127808 | 0.033353 | 0.090033 |
| CNAG_06207 | 1598.042 | 1.648497 | 0.106244 | 15.51612 | 2.70E-54 | 7.02E-52 |
| CNAG_06213 | 1475.504 | 0.358886 | 0.083464 | 4.299909 | 1.71E-05 | 0.000123 |
| CNAG_06214 | 1724.896 | 0.261922 | 0.074508 | 3.515348 | 0.000439 | 0.002325 |
| CNAG_06215 | 665.4307 | -0.24864 | 0.095888 | -2.59307 | 0.009512 | 0.03253 |
| CNAG_06216 | 1006.652 | 0.450732 | 0.097985 | 4.600014 | 4.22E-06 | 3.38E-05 |
| CNAG_06218 | 1053.773 | 0.240224 | 0.087189 | 2.755215 | 0.005865 | 0.02151 |
| CNAG_06220 | 11784.82 | 1.062331 | 0.069599 | 15.26354 | 1.34E-52 | 3.26E-50 |
| CNAG_06221 | 1115.738 | -0.34228 | 0.085615 | -3.99788 | 6.39E-05 | 0.000413 |

| | | | | | | |
|---|---|---|---|---|---|---|
| CNAG_06224 | 1332.479 | -0.37776 | 0.090859 | -4.15764 | 3.22E-05 | 0.000219 |
| CNAG_06226 | 4588.918 | 0.251341 | 0.077204 | 3.255552 | 0.001132 | 0.005285 |
| CNAG_06227 | 1295.691 | 0.234617 | 0.080885 | 2.900608 | 0.003724 | 0.014776 |
| CNAG_06228 | 2205.571 | -0.52355 | 0.08316 | -6.29567 | 3.06E-10 | 4.37E-09 |
| CNAG_06229 | 1483.291 | -0.18561 | 0.076277 | -2.43337 | 0.014959 | 0.04711 |
| CNAG_13087 | 31.64338 | -1.61694 | 0.374378 | -4.31901 | 1.57E-05 | 0.000114 |
| CNAG_06231 | 50082.39 | -0.23805 | 0.073713 | -3.22936 | 0.001241 | 0.005709 |
| CNAG_06232 | 840.5581 | 0.594693 | 0.102266 | 5.815163 | 6.06E-09 | 7.28E-08 |
| CNAG_06235 | 2389.741 | -0.27421 | 0.07235 | -3.79002 | 0.000151 | 0.000887 |
| CNAG_06236 | 1608.781 | -0.19616 | 0.080877 | -2.42544 | 0.01529 | 0.048016 |
| CNAG_06237 | 1657.692 | -0.16191 | 0.080167 | -2.01963 | 0.043422 | 0.111616 |
| CNAG_06238 | 245.5819 | 1.235562 | 0.175542 | 7.038566 | 1.94E-12 | 3.56E-11 |
| CNAG_06240 | 5751.625 | -0.50866 | 0.066368 | -7.66425 | 1.80E-14 | 3.95E-13 |
| CNAG_06241 | 1661.084 | -0.45258 | 0.086657 | -5.22262 | 1.76E-07 | 1.73E-06 |
| CNAG_06242 | 3592.687 | -0.61871 | 0.073388 | -8.4307 | 3.44E-17 | 9.34E-16 |
| CNAG_06243 | 79.37952 | -0.6116 | 0.224723 | -2.72156 | 0.006497 | 0.023552 |
| CNAG_06246 | 22603.71 | -0.21664 | 0.064989 | -3.3335 | 0.000858 | 0.004154 |
| CNAG_06247 | 231.8637 | 0.401639 | 0.144181 | 2.785667 | 0.005342 | 0.019851 |
| CNAG_06248 | 8624.878 | 0.84651 | 0.070616 | 11.98751 | 4.13E-33 | 3.54E-31 |
| CNAG_06252 | 459.2874 | -0.68277 | 0.112009 | -6.09561 | 1.09E-09 | 1.44E-08 |
| CNAG_07917 | 705.5617 | -0.28963 | 0.101658 | -2.84904 | 0.004385 | 0.016858 |
| CNAG_06986 | 492.7433 | -0.38321 | 0.108487 | -3.5323 | 0.000412 | 0.002193 |
| CNAG_06259 | 112.2623 | -1.89654 | 0.207917 | -9.12163 | 7.40E-20 | 2.66E-18 |
| CNAG_06260 | 1596.2 | -0.92961 | 0.07411 | -12.5438 | 4.30E-36 | 4.60E-34 |
| CNAG_13097 | 407.7283 | 0.346563 | 0.113769 | 3.046198 | 0.002318 | 0.009755 |
| CNAG_07922 | 936.0917 | -0.26871 | 0.08841 | -3.03937 | 0.002371 | 0.009949 |
| CNAG_06267 | 1609.643 | 1.223782 | 0.115725 | 10.57488 | 3.90E-26 | 2.27E-24 |
| CNAG_13099 | 466.0541 | 0.259113 | 0.107143 | 2.418396 | 0.015589 | 0.048673 |
| CNAG_06273 | 2332.404 | -0.37719 | 0.099278 | -3.79933 | 0.000145 | 0.000858 |
| CNAG_06274 | 11105.42 | 0.486823 | 0.069019 | 7.053475 | 1.75E-12 | 3.21E-11 |
| CNAG_13104 | 94.13849 | -0.57145 | 0.244705 | -2.33525 | 0.01953 | 0.058441 |
| CNAG_06277 | 3416.323 | -0.64518 | 0.068307 | -9.44518 | 3.55E-21 | 1.43E-19 |
| CNAG_06278 | 2507.48 | -0.19063 | 0.091766 | -2.07736 | 0.037768 | 0.09944 |
| CNAG_06279 | 3500.593 | -0.16088 | 0.066661 | -2.41346 | 0.015802 | 0.04917 |
| CNAG_06280 | 1454.933 | -0.17152 | 0.083346 | -2.05786 | 0.039603 | 0.103504 |
| CNAG_06282 | 1559.427 | 0.282754 | 0.084122 | 3.361254 | 0.000776 | 0.003811 |
| CNAG_06283 | 1175.223 | -0.23482 | 0.085738 | -2.73878 | 0.006167 | 0.022467 |
| CNAG_06284 | 586.8316 | -0.48568 | 0.117796 | -4.12304 | 3.74E-05 | 0.000251 |
| CNAG_06286 | 361.2818 | 0.758656 | 0.118952 | 6.377822 | 1.80E-10 | 2.63E-09 |
| CNAG_06290 | 475.4346 | 0.666713 | 0.115503 | 5.772261 | 7.82E-09 | 9.28E-08 |
| CNAG_06291 | 242.6414 | 0.383498 | 0.161628 | 2.372724 | 0.017657 | 0.053786 |
| CNAG_06293 | 391.9879 | 0.273082 | 0.115029 | 2.374024 | 0.017595 | 0.053701 |
| CNAG_06294 | 140.8121 | 0.56513 | 0.173393 | 3.259243 | 0.001117 | 0.005223 |
| CNAG_06296 | 64.22347 | 0.728254 | 0.271759 | 2.679773 | 0.007367 | 0.026205 |
| CNAG_06297 | 709.4637 | 1.756757 | 0.241136 | 7.285329 | 3.21E-13 | 6.40E-12 |
| CNAG_06298 | 2969.625 | 1.833788 | 0.109343 | 16.77102 | 3.98E-63 | 1.63E-60 |
| CNAG_06299 | 881.2592 | -0.20856 | 0.087479 | -2.3841 | 0.017121 | 0.052459 |
| CNAG_06300 | 2092.847 | -0.2513 | 0.081468 | -3.0847 | 0.002038 | 0.008741 |

| | | | | | | |
|---|---|---|---|---|---|---|
| CNAG_06301 | 5405.068 | 0.238303 | 0.064274 | 3.707633 | 0.000209 | 0.001191 |
| CNAG_06302 | 117.3824 | 0.501688 | 0.194057 | 2.58526 | 0.009731 | 0.033112 |
| CNAG_06314 | 10198.85 | -0.2569 | 0.065002 | -3.95225 | 7.74E-05 | 0.000492 |
| CNAG_06318 | 2363.295 | -0.46592 | 0.074814 | -6.22767 | 4.73E-10 | 6.56E-09 |
| CNAG_06320 | 2769.489 | -0.15231 | 0.076769 | -1.984 | 0.047256 | 0.119039 |
| CNAG_06323 | 411.2227 | 0.273783 | 0.11549 | 2.370615 | 0.017759 | 0.054037 |
| CNAG_06329 | 196.0641 | 0.817278 | 0.163657 | 4.993846 | 5.92E-07 | 5.35E-06 |
| CNAG_06332 | 1653.148 | 0.361897 | 0.08705 | 4.157346 | 3.22E-05 | 0.000219 |
| CNAG_06336 | 811.6631 | -0.20912 | 0.086432 | -2.41943 | 0.015545 | 0.048563 |
| CNAG_13117 | 24.03939 | -1.16744 | 0.409707 | -2.84944 | 0.00438 | 0.01685 |
| CNAG_06338 | 341.6506 | -0.33853 | 0.137183 | -2.46774 | 0.013597 | 0.043505 |
| CNAG_06340 | 2205.663 | -0.406 | 0.079466 | -5.10911 | 3.24E-07 | 3.06E-06 |
| CNAG_06341 | 907.8548 | -0.71792 | 0.084113 | -8.53514 | 1.40E-17 | 3.90E-16 |
| CNAG_06345 | 1023.695 | -0.21419 | 0.092688 | -2.31082 | 0.020843 | 0.061659 |
| CNAG_13120 | 285.242 | 0.433005 | 0.134132 | 3.228203 | 0.001246 | 0.005729 |
| CNAG_06347 | 13564.62 | 0.804341 | 0.098121 | 8.197419 | 2.46E-16 | 6.30E-15 |
| CNAG_06348 | 1342.076 | -0.70335 | 0.093578 | -7.51622 | 5.64E-14 | 1.19E-12 |
| CNAG_06350 | 1575.143 | -0.49552 | 0.076412 | -6.48478 | 8.89E-11 | 1.35E-09 |
| CNAG_06351 | 2427.681 | -0.18832 | 0.072044 | -2.614 | 0.008949 | 0.030929 |
| CNAG_06353 | 3134.282 | -0.18813 | 0.065548 | -2.87014 | 0.004103 | 0.015993 |
| CNAG_06356 | 1561.39 | 0.27664 | 0.075169 | 3.680249 | 0.000233 | 0.001319 |
| CNAG_06360 | 1179.93 | 0.21203 | 0.083717 | 2.532698 | 0.011319 | 0.037508 |
| CNAG_06361 | 3442.365 | -0.14728 | 0.068269 | -2.15738 | 0.030976 | 0.085209 |
| CNAG_06363 | 1224.099 | -0.52922 | 0.081964 | -6.45672 | 1.07E-10 | 1.61E-09 |
| CNAG_06364 | 5004.111 | 0.260834 | 0.089961 | 2.899394 | 0.003739 | 0.014811 |
| CNAG_06367 | 3503.42 | -0.55759 | 0.068808 | -8.10361 | 5.34E-16 | 1.32E-14 |
| CNAG_06371 | 311.091 | 0.315129 | 0.124624 | 2.528641 | 0.011451 | 0.037896 |
| CNAG_06372 | 204.4283 | 0.704117 | 0.157674 | 4.465639 | 7.98E-06 | 6.13E-05 |
| CNAG_06374 | 1749.606 | 0.912364 | 0.111684 | 8.169187 | 3.10E-16 | 7.84E-15 |
| CNAG_06377 | 45024.72 | 0.515119 | 0.074791 | 6.887421 | 5.68E-12 | 9.87E-11 |
| CNAG_06378 | 527.0394 | -0.24976 | 0.106214 | -2.35148 | 0.018699 | 0.056279 |
| CNAG_06379 | 3283.255 | -0.63155 | 0.068932 | -9.16201 | 5.09E-20 | 1.85E-18 |
| CNAG_06381 | 8887.266 | 0.536153 | 0.066632 | 8.046421 | 8.53E-16 | 2.09E-14 |
| CNAG_06385 | 810.0598 | -0.2766 | 0.099658 | -2.77551 | 0.005512 | 0.020396 |
| CNAG_06388 | 5350.99 | 1.433442 | 0.087276 | 16.42428 | 1.28E-60 | 4.76E-58 |
| CNAG_06389 | 141.1413 | -0.4023 | 0.173412 | -2.31992 | 0.020345 | 0.060439 |
| CNAG_06392 | 1123.729 | 0.170406 | 0.079315 | 2.148465 | 0.031677 | 0.086617 |
| CNAG_13125 | 22.19235 | 1.065925 | 0.414642 | 2.570711 | 0.010149 | 0.034257 |
| CNAG_06396 | 354.7988 | 1.05265 | 0.128826 | 8.171091 | 3.06E-16 | 7.77E-15 |
| CNAG_06399 | 658.5004 | 0.317261 | 0.094522 | 3.356466 | 0.000789 | 0.003863 |
| CNAG_06400 | 84469.46 | 0.413276 | 0.060643 | 6.814864 | 9.44E-12 | 1.61E-10 |
| CNAG_13126 | 60.27702 | -0.63786 | 0.285936 | -2.23078 | 0.025696 | 0.073374 |
| CNAG_06402 | 1748.651 | -0.24659 | 0.087036 | -2.83322 | 0.004608 | 0.017611 |
| CNAG_06403 | 1624.624 | 0.187683 | 0.079186 | 2.370157 | 0.017781 | 0.054076 |
| CNAG_06404 | 5447.968 | 1.422334 | 0.084064 | 16.91972 | 3.22E-64 | 1.48E-61 |
| CNAG_06405 | 925.36 | -0.25524 | 0.092304 | -2.76525 | 0.005688 | 0.020929 |
| CNAG_06407 | 3363.323 | 0.342728 | 0.082988 | 4.129857 | 3.63E-05 | 0.000244 |
| CNAG_06409 | 574.1689 | 0.753548 | 0.118075 | 6.38195 | 1.75E-10 | 2.57E-09 |

| | | | | | | |
|---|---|---|---|---|---|---|
| CNAG_06410 | 1203.231 | 0.308531 | 0.081405 | 3.790082 | 0.000151 | 0.000887 |
| CNAG_06412 | 1005.102 | -0.71569 | 0.082006 | -8.72726 | 2.61E-18 | 7.92E-17 |
| CNAG_06420 | 3477.073 | -0.19894 | 0.070356 | -2.82767 | 0.004689 | 0.017867 |
| CNAG_06422 | 3301.558 | 0.226218 | 0.092356 | 2.449403 | 0.014309 | 0.045386 |
| CNAG_06424 | 927.1685 | -0.29124 | 0.090554 | -3.21621 | 0.001299 | 0.005956 |
| CNAG_06428 | 1077.459 | 0.387472 | 0.099905 | 3.878419 | 0.000105 | 0.000645 |
| CNAG_06432 | 4149.821 | 1.063227 | 0.086531 | 12.2872 | 1.06E-34 | 1.01E-32 |
| CNAG_07939 | 11131.59 | 0.898885 | 0.095933 | 9.369955 | 7.26E-21 | 2.86E-19 |
| CNAG_06443 | 20009.07 | -0.44287 | 0.064519 | -6.86415 | 6.69E-12 | 1.15E-10 |
| CNAG_06446 | 2915.909 | 0.782239 | 0.067683 | 11.55744 | 6.77E-31 | 5.08E-29 |
| CNAG_06447 | 35537.99 | -0.22553 | 0.080304 | -2.80848 | 0.004978 | 0.018751 |
| CNAG_06449 | 93.13515 | -0.46238 | 0.218448 | -2.11665 | 0.03429 | 0.092053 |
| CNAG_06452 | 1294.129 | 0.328427 | 0.079674 | 4.122131 | 3.75E-05 | 0.000252 |
| CNAG_06453 | 4001.456 | 1.035563 | 0.088039 | 11.76251 | 6.09E-32 | 4.80E-30 |
| CNAG_06454 | 1286.043 | -0.22218 | 0.079834 | -2.783 | 0.005386 | 0.019996 |
| CNAG_06459 | 1112.217 | 0.603435 | 0.089296 | 6.757686 | 1.40E-11 | 2.35E-10 |
| CNAG_06460 | 1757.963 | -0.16698 | 0.074744 | -2.23398 | 0.025484 | 0.07293 |
| CNAG_07942 | 1659.858 | -0.28073 | 0.088644 | -3.16692 | 0.001541 | 0.006882 |
| CNAG_06466 | 459.9842 | -0.24622 | 0.109259 | -2.25358 | 0.024222 | 0.069706 |
| CNAG_06468 | 3881.948 | -0.62655 | 0.069873 | -8.96693 | 3.05E-19 | 1.02E-17 |
| CNAG_06472 | 4788.553 | -0.53372 | 0.066817 | -7.98789 | 1.37E-15 | 3.32E-14 |
| CNAG_06474 | 5771.033 | -0.23696 | 0.071014 | -3.33682 | 0.000847 | 0.00411 |
| CNAG_06475 | 8269.977 | -0.17443 | 0.079062 | -2.2062 | 0.02737 | 0.077447 |
| CNAG_13132 | 151.7017 | -0.4423 | 0.169216 | -2.61384 | 0.008953 | 0.030929 |
| CNAG_06482 | 627.2793 | 0.453139 | 0.102644 | 4.414646 | 1.01E-05 | 7.64E-05 |
| CNAG_06484 | 121.7986 | 1.058184 | 0.189104 | 5.59578 | 2.20E-08 | 2.44E-07 |
| CNAG_06486 | 2505.615 | -0.24431 | 0.074643 | -3.27312 | 0.001064 | 0.005013 |
| CNAG_07943 | 141.5177 | -1.76491 | 0.186399 | -9.46845 | 2.84E-21 | 1.15E-19 |
| CNAG_06491 | 159.2812 | -0.6016 | 0.164929 | -3.64765 | 0.000265 | 0.001481 |
| CNAG_06493 | 1577.369 | 0.612585 | 0.108976 | 5.621271 | 1.90E-08 | 2.12E-07 |
| CNAG_06494 | 1162.252 | 0.393532 | 0.088445 | 4.449455 | 8.61E-06 | 6.57E-05 |
| CNAG_06497 | 812.6171 | 0.413793 | 0.096451 | 4.290208 | 1.79E-05 | 0.000129 |
| CNAG_06498 | 922.0555 | 0.68738 | 0.099522 | 6.906828 | 4.96E-12 | 8.67E-11 |
| CNAG_06499 | 1107.691 | 0.500012 | 0.080507 | 6.21083 | 5.27E-10 | 7.24E-09 |
| CNAG_06500 | 745.5015 | 0.311238 | 0.109503 | 2.842274 | 0.004479 | 0.017186 |
| CNAG_06505 | 1072.706 | 0.178463 | 0.082773 | 2.156062 | 0.031079 | 0.085371 |
| CNAG_06506 | 660.6532 | 0.184285 | 0.092511 | 1.992031 | 0.046368 | 0.117256 |
| CNAG_06512 | 512.5173 | -0.49446 | 0.10919 | -4.52847 | 5.94E-06 | 4.63E-05 |
| CNAG_13141 | 168.2074 | -0.55846 | 0.184604 | -3.02516 | 0.002485 | 0.010404 |
| CNAG_06517 | 3284.006 | -0.71275 | 0.092552 | -7.70112 | 1.35E-14 | 3.01E-13 |
| CNAG_13142 | 439.5184 | 0.613769 | 0.127062 | 4.830466 | 1.36E-06 | 1.17E-05 |
| CNAG_07043 | 360.4921 | 0.511198 | 0.12408 | 4.119891 | 3.79E-05 | 0.000254 |
| CNAG_05333 | 11.14682 | 2.583006 | 0.676941 | 3.815703 | 0.000136 | 0.000808 |
| CNAG_05336 | 1067.761 | 0.36054 | 0.089921 | 4.009534 | 6.08E-05 | 0.000395 |
| CNAG_05337 | 1002.436 | 0.181102 | 0.085112 | 2.127811 | 0.033353 | 0.090033 |
| CNAG_05340 | 215.6441 | -1.39521 | 0.168698 | -8.27046 | 1.33E-16 | 3.49E-15 |
| CNAG_05341 | 212.8876 | 0.423002 | 0.167801 | 2.520857 | 0.011707 | 0.038634 |
| CNAG_05343 | 1331.89 | -0.29826 | 0.096327 | -3.09631 | 0.001959 | 0.008472 |

| | | | | | | |
|---|---|---|---|---|---|---|
| CNAG_05344 | 2576.945 | 0.238342 | 0.067553 | 3.528211 | 0.000418 | 0.002224 |
| CNAG_07032 | 4062.051 | 0.212741 | 0.083001 | 2.563113 | 0.010374 | 0.03485 |
| CNAG_05349 | 5078.27 | 0.214195 | 0.069349 | 3.088652 | 0.002011 | 0.008645 |
| CNAG_13146 | 55.8919 | 0.556846 | 0.26951 | 2.066146 | 0.038815 | 0.101682 |
| CNAG_05350 | 2044.732 | 0.477868 | 0.08478 | 5.636589 | 1.73E-08 | 1.96E-07 |
| CNAG_05351 | 7594.852 | -0.19347 | 0.067385 | -2.87117 | 0.00409 | 0.015981 |
| CNAG_05352 | 1259.958 | -0.30925 | 0.078922 | -3.91844 | 8.91E-05 | 0.000554 |
| CNAG_05356 | 55.71849 | 0.523739 | 0.261273 | 2.004561 | 0.04501 | 0.114715 |
| CNAG_05357 | 17.93195 | 1.164725 | 0.481625 | 2.418322 | 0.015592 | 0.048673 |
| CNAG_13147 | 136.3623 | 0.635044 | 0.203876 | 3.114856 | 0.00184 | 0.008024 |
| CNAG_05360 | 1963.874 | -0.54287 | 0.07213 | -7.52628 | 5.22E-14 | 1.10E-12 |
| CNAG_08021 | 109.9231 | 0.389149 | 0.195254 | 1.993042 | 0.046257 | 0.117052 |
| CNAG_05363 | 2563.725 | -0.19403 | 0.076156 | -2.54786 | 0.010839 | 0.036209 |
| CNAG_05365 | 1952.526 | -0.24913 | 0.08055 | -3.09282 | 0.001983 | 0.008553 |
| CNAG_05370 | 567.1978 | -0.32861 | 0.09916 | -3.31397 | 0.00092 | 0.004418 |
| CNAG_05371 | 1883.429 | -0.46383 | 0.080222 | -5.78187 | 7.39E-09 | 8.80E-08 |
| CNAG_05375 | 25727.86 | 0.17379 | 0.059879 | 2.90235 | 0.003704 | 0.014709 |
| CNAG_05379 | 1391.16 | -0.16177 | 0.077767 | -2.08021 | 0.037507 | 0.098785 |
| CNAG_05381 | 220.8777 | 1.512279 | 0.384751 | 3.930535 | 8.48E-05 | 0.000533 |
| CNAG_07874 | 35.36483 | 0.829767 | 0.334255 | 2.482439 | 0.013049 | 0.042209 |
| CNAG_05383 | 921.7177 | 1.116757 | 0.111887 | 9.981116 | 1.84E-23 | 9.11E-22 |
| CNAG_05386 | 638.9377 | -0.24493 | 0.097563 | -2.51046 | 0.012057 | 0.039486 |
| CNAG_13158 | 42.29634 | 1.812589 | 0.340558 | 5.322408 | 1.02E-07 | 1.03E-06 |
| CNAG_05388 | 1396.068 | 0.67952 | 0.085948 | 7.906166 | 2.65E-15 | 6.28E-14 |
| CNAG_05392 | 175.9939 | 0.425172 | 0.177437 | 2.396187 | 0.016567 | 0.05103 |
| CNAG_05394 | 615.3845 | 0.21603 | 0.099938 | 2.161639 | 0.030646 | 0.084509 |
| CNAG_05401 | 1103.304 | -0.70482 | 0.277702 | -2.53806 | 0.011147 | 0.037049 |
| CNAG_05402 | 2680.022 | -0.4376 | 0.093428 | -4.68386 | 2.82E-06 | 2.31E-05 |
| CNAG_05410 | 745.9114 | 1.02892 | 0.102289 | 10.05893 | 8.39E-24 | 4.20E-22 |
| CNAG_05411 | 1266.516 | 0.475969 | 0.080376 | 5.921745 | 3.19E-09 | 3.99E-08 |
| CNAG_05413 | 707.938 | -0.20351 | 0.099771 | -2.03973 | 0.041377 | 0.107457 |
| CNAG_05415 | 1936.173 | -0.61226 | 0.074113 | -8.26118 | 1.44E-16 | 3.75E-15 |
| CNAG_13165 | 24.27277 | -0.87575 | 0.428333 | -2.04456 | 0.040898 | 0.10639 |
| CNAG_05422 | 1199.226 | 0.283288 | 0.091262 | 3.104121 | 0.001908 | 0.008283 |
| CNAG_08024 | 358.329 | -0.29779 | 0.142291 | -2.09283 | 0.036364 | 0.096427 |
| CNAG_05430 | 776.3244 | 0.593954 | 0.097062 | 6.119323 | 9.40E-10 | 1.25E-08 |
| CNAG_05431 | 1385.483 | 0.648673 | 0.080219 | 8.086325 | 6.15E-16 | 1.51E-14 |
| CNAG_05434 | 3459.741 | 0.24438 | 0.064651 | 3.779973 | 0.000157 | 0.00092 |
| CNAG_05436 | 124.1802 | -0.66485 | 0.185348 | -3.58707 | 0.000334 | 0.001817 |
| CNAG_05437 | 17511.93 | -0.27975 | 0.067445 | -4.14779 | 3.36E-05 | 0.000227 |
| CNAG_05438 | 649.5541 | -0.65222 | 0.095637 | -6.81977 | 9.12E-12 | 1.56E-10 |
| CNAG_05439 | 761.807 | 0.405476 | 0.094103 | 4.308878 | 1.64E-05 | 0.000119 |
| CNAG_05441 | 319.6465 | -0.37692 | 0.121283 | -3.10778 | 0.001885 | 0.008191 |
| CNAG_05442 | 1675.287 | -0.17835 | 0.077826 | -2.29159 | 0.021929 | 0.064226 |
| CNAG_05453 | 169.6498 | -0.51767 | 0.160954 | -3.21623 | 0.001299 | 0.005956 |
| CNAG_05457 | 4309.838 | 0.545266 | 0.073483 | 7.420284 | 1.17E-13 | 2.43E-12 |
| CNAG_05459 | 4417.873 | 0.418367 | 0.071246 | 5.872123 | 4.30E-09 | 5.31E-08 |
| CNAG_05460 | 308.406 | -0.83131 | 0.126648 | -6.56392 | 5.24E-11 | 8.12E-10 |

| | | | | | | |
|---|---|---|---|---|---|---|
| CNAG_05461 | 56.86511 | -0.97255 | 0.268767 | -3.61855 | 0.000296 | 0.001629 |
| CNAG_05462 | 10767.52 | 0.331674 | 0.068967 | 4.809162 | 1.52E-06 | 1.29E-05 |
| CNAG_05463 | 1709.763 | -0.25113 | 0.075713 | -3.31683 | 0.00091 | 0.004378 |
| CNAG_05465 | 38844.54 | -0.16644 | 0.077593 | -2.14504 | 0.031949 | 0.08718 |
| CNAG_05470 | 694.3285 | -0.21866 | 0.098374 | -2.22274 | 0.026233 | 0.074608 |
| CNAG_13180 | 21.01911 | -1.25142 | 0.446372 | -2.80353 | 0.005055 | 0.018974 |
| CNAG_07878 | 3112.887 | -0.88052 | 0.099631 | -8.83773 | 9.77E-19 | 3.07E-17 |
| CNAG_05473 | 619.7838 | -0.54174 | 0.096872 | -5.59227 | 2.24E-08 | 2.49E-07 |
| CNAG_13181 | 92.25756 | 0.452651 | 0.204422 | 2.214293 | 0.026809 | 0.07605 |
| CNAG_05475 | 2779.07 | -0.31814 | 0.068835 | -4.6218 | 3.80E-06 | 3.07E-05 |
| CNAG_05478 | 1583.486 | -0.21135 | 0.077988 | -2.70997 | 0.006729 | 0.024289 |
| CNAG_05479 | 634.3322 | -0.64963 | 0.10274 | -6.32307 | 2.56E-10 | 3.70E-09 |
| CNAG_05480 | 7277.191 | -0.21943 | 0.063311 | -3.46587 | 0.000529 | 0.002744 |
| CNAG_13183 | 17.61844 | -1.26769 | 0.491751 | -2.57791 | 0.00994 | 0.033683 |
| CNAG_05485 | 280.7604 | -0.64257 | 0.150386 | -4.2728 | 1.93E-05 | 0.000137 |
| CNAG_13185 | 108.9366 | -0.50988 | 0.198584 | -2.56755 | 0.010242 | 0.034482 |
| CNAG_05486 | 2730.489 | -0.27702 | 0.088212 | -3.14043 | 0.001687 | 0.007451 |
| CNAG_08026 | 220.4027 | -0.33305 | 0.144521 | -2.30451 | 0.021194 | 0.06258 |
| CNAG_05497 | 16844.42 | 0.300575 | 0.059008 | 5.093797 | 3.51E-07 | 3.30E-06 |
| CNAG_05504 | 2557.718 | 0.302967 | 0.069807 | 4.340044 | 1.42E-05 | 0.000105 |
| CNAG_07883 | 1118.136 | -0.1973 | 0.092134 | -2.14149 | 0.032235 | 0.087805 |
| CNAG_05509 | 2173.969 | -0.1806 | 0.079455 | -2.27296 | 0.023029 | 0.066809 |
| CNAG_05516 | 1043.768 | -0.19649 | 0.083545 | -2.3519 | 0.018678 | 0.056279 |
| CNAG_05521 | 2271.968 | 0.325383 | 0.078819 | 4.12825 | 3.66E-05 | 0.000245 |
| CNAG_05522 | 5060.169 | 0.252632 | 0.073843 | 3.421207 | 0.000623 | 0.003165 |
| CNAG_05523 | 80.12682 | -0.85703 | 0.230858 | -3.71237 | 0.000205 | 0.00117 |
| CNAG_05524 | 1242.942 | 0.219596 | 0.083258 | 2.637532 | 0.008351 | 0.029213 |
| CNAG_05525 | 46035.7 | -0.16517 | 0.079618 | -2.0745 | 0.038033 | 0.100036 |
| CNAG_05528 | 33.77831 | 0.752629 | 0.340607 | 2.209673 | 0.027128 | 0.076844 |
| CNAG_05529 | 847.5263 | -0.33718 | 0.093735 | -3.59713 | 0.000322 | 0.001753 |
| CNAG_05531 | 1471.731 | 0.320059 | 0.087192 | 3.670741 | 0.000242 | 0.001364 |
| CNAG_05532 | 165.2444 | -0.42935 | 0.164308 | -2.61307 | 0.008973 | 0.030986 |
| CNAG_05533 | 584.3482 | 0.633861 | 0.099839 | 6.348848 | 2.17E-10 | 3.16E-09 |
| CNAG_05534 | 521.6441 | -0.37395 | 0.109018 | -3.43019 | 0.000603 | 0.003081 |
| CNAG_05535 | 1758.351 | -0.48176 | 0.089189 | -5.40156 | 6.61E-08 | 6.88E-07 |
| CNAG_05536 | 255.5293 | 0.532179 | 0.142091 | 3.745347 | 0.00018 | 0.00104 |
| CNAG_05537 | 993.151 | -0.34243 | 0.084469 | -4.0539 | 5.04E-05 | 0.000331 |
| CNAG_05538 | 1634.368 | -0.15862 | 0.077191 | -2.05494 | 0.039884 | 0.1041 |
| CNAG_05540 | 8024.021 | 0.141114 | 0.064557 | 2.1859 | 0.028823 | 0.080708 |
| CNAG_13191 | 19.74802 | -1.13644 | 0.450207 | -2.52426 | 0.011594 | 0.038323 |
| CNAG_05544 | 357.4832 | 0.37602 | 0.124793 | 3.013162 | 0.002585 | 0.010755 |
| CNAG_05547 | 293.4871 | 0.919138 | 0.133816 | 6.868697 | 6.48E-12 | 1.12E-10 |
| CNAG_05548 | 162.9897 | 0.863386 | 0.164238 | 5.256915 | 1.46E-07 | 1.45E-06 |
| CNAG_05551 | 790.1423 | -0.23794 | 0.094658 | -2.51364 | 0.011949 | 0.039216 |
| CNAG_05552 | 5925.468 | -0.20374 | 0.077339 | -2.63434 | 0.00843 | 0.02941 |
| CNAG_05553 | 584.9947 | 0.341089 | 0.103353 | 3.300241 | 0.000966 | 0.004611 |
| CNAG_05555 | 54969.4 | -0.20956 | 0.074052 | -2.82992 | 0.004656 | 0.01775 |
| CNAG_05556 | 16303.97 | -0.20065 | 0.073735 | -2.72124 | 0.006504 | 0.023558 |

| | | | | | | |
|---|---|---|---|---|---|---|
| CNAG_05559 | 2263.718 | 0.251428 | 0.074166 | 3.390054 | 0.000699 | 0.003493 |
| CNAG_05566 | 515.3151 | 0.338061 | 0.104296 | 3.241369 | 0.00119 | 0.005513 |
| CNAG_05567 | 849.2166 | 0.193933 | 0.086841 | 2.233185 | 0.025537 | 0.073008 |
| CNAG_05569 | 694.6259 | 0.570822 | 0.092003 | 6.204382 | 5.49E-10 | 7.52E-09 |
| CNAG_05571 | 3167.92 | -0.60058 | 0.078788 | -7.62273 | 2.48E-14 | 5.40E-13 |
| CNAG_05573 | 1429.28 | 1.220881 | 0.085311 | 14.3109 | 1.87E-46 | 3.40E-44 |
| CNAG_05575 | 1373.762 | 0.641629 | 0.078364 | 8.187778 | 2.66E-16 | 6.81E-15 |
| CNAG_05578 | 1889.236 | 0.210962 | 0.090532 | 2.33025 | 0.019793 | 0.059091 |
| CNAG_05580 | 48.78851 | 1.443774 | 0.327518 | 4.408232 | 1.04E-05 | 7.84E-05 |
| CNAG_05584 | 1223.909 | -0.2201 | 0.085175 | -2.58407 | 0.009764 | 0.033189 |
| CNAG_05586 | 805.335 | 0.500079 | 0.088719 | 5.636636 | 1.73E-08 | 1.96E-07 |
| CNAG_05590 | 1875.431 | 0.314941 | 0.09385 | 3.355787 | 0.000791 | 0.00387 |
| CNAG_05591 | 645.3286 | 0.290146 | 0.101697 | 2.853035 | 0.00433 | 0.016702 |
| CNAG_05592 | 4025.959 | -0.81144 | 0.066565 | -12.1902 | 3.50E-34 | 3.25E-32 |
| CNAG_05595 | 1247.934 | 0.285436 | 0.081756 | 3.491309 | 0.000481 | 0.002521 |
| CNAG_05597 | 894.0709 | -0.24045 | 0.089719 | -2.68001 | 0.007362 | 0.026198 |
| CNAG_05598 | 1286.545 | 0.376294 | 0.076655 | 4.908902 | 9.16E-07 | 7.99E-06 |
| CNAG_05599 | 372.3624 | 0.829327 | 0.123977 | 6.689352 | 2.24E-11 | 3.69E-10 |
| CNAG_05600 | 1764.932 | -0.30434 | 0.077761 | -3.91384 | 9.08E-05 | 0.000564 |
| CNAG_05601 | 1121.376 | -0.26229 | 0.085231 | -3.07743 | 0.002088 | 0.008923 |
| CNAG_05603 | 1104.435 | -0.57181 | 0.088901 | -6.43206 | 1.26E-10 | 1.87E-09 |
| CNAG_05605 | 706.8289 | 0.280004 | 0.099829 | 2.804843 | 0.005034 | 0.018924 |
| CNAG_05606 | 802.56 | -0.22691 | 0.112004 | -2.02594 | 0.042771 | 0.110165 |
| CNAG_05607 | 2450.173 | 0.333019 | 0.079539 | 4.186869 | 2.83E-05 | 0.000194 |
| CNAG_05608 | 452.6187 | 0.404444 | 0.126816 | 3.189212 | 0.001427 | 0.006458 |
| CNAG_07888 | 25676.14 | 0.381057 | 0.075094 | 5.074382 | 3.89E-07 | 3.62E-06 |
| CNAG_07889 | 1618.008 | -0.41775 | 0.075949 | -5.50041 | 3.79E-08 | 4.08E-07 |
| CNAG_05614 | 1848.436 | 0.418497 | 0.083536 | 5.0098 | 5.45E-07 | 4.98E-06 |
| CNAG_05615 | 5670.37 | -0.22899 | 0.075198 | -3.0452 | 0.002325 | 0.009782 |
| CNAG_05616 | 1935.621 | 0.305044 | 0.093271 | 3.270514 | 0.001074 | 0.005053 |
| CNAG_05617 | 1258.271 | -0.29812 | 0.089294 | -3.33867 | 0.000842 | 0.004088 |
| CNAG_05618 | 1994.73 | -0.34972 | 0.074722 | -4.68026 | 2.87E-06 | 2.35E-05 |
| CNAG_05619 | 1203.288 | -0.39233 | 0.079719 | -4.92143 | 8.59E-07 | 7.53E-06 |
| CNAG_05620 | 1123.636 | 0.44498 | 0.083485 | 5.330075 | 9.82E-08 | 9.94E-07 |
| CNAG_05621 | 1187.869 | -0.41324 | 0.08208 | -5.03457 | 4.79E-07 | 4.42E-06 |
| CNAG_05622 | 772.0948 | -0.26288 | 0.120141 | -2.18809 | 0.028663 | 0.080376 |
| CNAG_05624 | 889.3339 | -0.25518 | 0.094058 | -2.713 | 0.006668 | 0.02409 |
| CNAG_05626 | 1729.89 | 1.516959 | 0.075854 | 19.99849 | 5.68E-89 | 7.38E-86 |
| CNAG_05627 | 1143.623 | -0.56662 | 0.079431 | -7.13351 | 9.78E-13 | 1.84E-11 |
| CNAG_05628 | 819.237 | 0.311831 | 0.08918 | 3.496638 | 0.000471 | 0.002474 |
| CNAG_05631 | 11536.66 | 0.187489 | 0.060316 | 3.108473 | 0.001881 | 0.008176 |
| CNAG_05632 | 257.513 | 0.583712 | 0.136458 | 4.277599 | 1.89E-05 | 0.000135 |
| CNAG_05633 | 4242.542 | 0.340525 | 0.072833 | 4.675443 | 2.93E-06 | 2.40E-05 |
| CNAG_05637 | 1043.625 | 0.298556 | 0.091265 | 3.271315 | 0.00107 | 0.005042 |
| CNAG_05639 | 1509.545 | 0.743432 | 0.084888 | 8.757849 | 1.99E-18 | 6.11E-17 |
| CNAG_05640 | 1292.576 | -0.17476 | 0.079408 | -2.20079 | 0.027751 | 0.078353 |
| CNAG_05642 | 1308.364 | -0.37359 | 0.103761 | -3.60046 | 0.000318 | 0.001734 |
| CNAG_05643 | 939.0149 | 0.481986 | 0.102463 | 4.703988 | 2.55E-06 | 2.11E-05 |

| | | | | | | |
|---|---|---|---|---|---|---|
| CNAG_05644 | 358.3596 | -0.63079 | 0.119606 | -5.27392 | 1.34E-07 | 1.32E-06 |
| CNAG_05645 | 1521.354 | 0.325999 | 0.082487 | 3.952113 | 7.75E-05 | 0.000492 |
| CNAG_05651 | 495.9795 | -0.55453 | 0.108913 | -5.09144 | 3.55E-07 | 3.33E-06 |
| CNAG_05652 | 1516.392 | 0.491594 | 0.078337 | 6.275382 | 3.49E-10 | 4.91E-09 |
| CNAG_05653 | 1365.684 | 0.93224 | 0.107073 | 8.706618 | 3.13E-18 | 9.36E-17 |
| CNAG_05654 | 471.8605 | 0.919785 | 0.118458 | 7.764624 | 8.19E-15 | 1.86E-13 |
| CNAG_05656 | 162.635 | 0.618778 | 0.162831 | 3.800127 | 0.000145 | 0.000856 |
| CNAG_05657 | 907.3743 | -1.36806 | 0.109962 | -12.4412 | 1.56E-35 | 1.58E-33 |
| CNAG_05660 | 2160.632 | -0.23905 | 0.072315 | -3.30573 | 0.000947 | 0.004535 |
| CNAG_13203 | 154.2388 | 0.545554 | 0.174526 | 3.125921 | 0.001772 | 0.007767 |
| CNAG_05662 | 4979.296 | 2.120062 | 0.247142 | 8.578314 | 9.63E-18 | 2.72E-16 |
| CNAG_05663 | 3068.972 | -0.22473 | 0.110556 | -2.03275 | 0.042077 | 0.108841 |
| CNAG_05665 | 2115.273 | 0.505974 | 0.076097 | 6.649087 | 2.95E-11 | 4.72E-10 |
| CNAG_05667 | 535.5497 | -0.58103 | 0.102202 | -5.68517 | 1.31E-08 | 1.51E-07 |
| CNAG_13206 | 289.9507 | 0.800981 | 0.14092 | 5.683929 | 1.32E-08 | 1.52E-07 |

**Table S7. Data sheets of the RNA-seq data for L-DOPA treatment.**

| | baseMean | log2FoldCh | lfcSE | stat | pvalue | padj |
|---|---|---|---|---|---|---|
| CNAG_07303 | 5.215809 | -2.41691 | 1.117363 | -2.16305 | 0.030538 | 0.063839 |
| CNAG_00002 | 777.2105 | -0.56059 | 0.209632 | -2.67417 | 0.007491 | 0.019153 |
| CNAG_12001 | 44.45174 | -1.27195 | 0.403439 | -3.15276 | 0.001617 | 0.005066 |
| CNAG_00007 | 3549.556 | -0.48982 | 0.129262 | -3.78937 | 0.000151 | 0.000628 |
| CNAG_00009 | 838.3747 | -0.35288 | 0.142968 | -2.46826 | 0.013577 | 0.031938 |
| CNAG_00011 | 1116.719 | 0.603541 | 0.197712 | 3.052629 | 0.002268 | 0.006831 |
| CNAG_00014 | 409.6928 | 0.731429 | 0.204096 | 3.583751 | 0.000339 | 0.001285 |
| CNAG_00015 | 274.6089 | 0.961797 | 0.186631 | 5.153461 | 2.56E-07 | 1.99E-06 |
| CNAG_00016 | 2990.749 | -0.31834 | 0.125818 | -2.53017 | 0.011401 | 0.027527 |
| CNAG_00017 | 492.0273 | 1.1063 | 0.259079 | 4.270125 | 1.95E-05 | 0.000101 |
| CNAG_12002 | 34.00088 | 2.14414 | 0.433779 | 4.942936 | 7.70E-07 | 5.47E-06 |
| CNAG_00018 | 1707.721 | -0.69918 | 0.149693 | -4.67075 | 3.00E-06 | 1.90E-05 |
| CNAG_00020 | 1417.702 | 0.511232 | 0.198609 | 2.574065 | 0.010051 | 0.024711 |
| CNAG_12003 | 48.53692 | 1.33116 | 0.326525 | 4.076747 | 4.57E-05 | 0.000216 |
| CNAG_00021 | 834.3204 | -0.98412 | 0.145397 | -6.76855 | 1.30E-11 | 2.24E-10 |
| CNAG_00022 | 1163.718 | -1.30674 | 0.237988 | -5.49078 | 4.00E-08 | 3.76E-07 |
| CNAG_00023 | 164.8357 | 0.762866 | 0.180711 | 4.22147 | 2.43E-05 | 0.000123 |
| CNAG_00025 | 1364.991 | 0.499258 | 0.139323 | 3.583456 | 0.000339 | 0.001285 |
| CNAG_00026 | 6056.017 | -0.62891 | 0.173395 | -3.62703 | 0.000287 | 0.001114 |
| CNAG_00030 | 238.4392 | -1.06034 | 0.207624 | -5.10702 | 3.27E-07 | 2.50E-06 |
| CNAG_00031 | 668.2287 | -0.29402 | 0.125038 | -2.35141 | 0.018702 | 0.042027 |
| CNAG_00034 | 37138.83 | -0.88395 | 0.192721 | -4.5867 | 4.50E-06 | 2.74E-05 |
| CNAG_00036 | 837.4774 | 1.7747 | 0.229261 | 7.740969 | 9.87E-15 | 2.60E-13 |
| CNAG_07307 | 51.56041 | 0.906035 | 0.365949 | 2.475853 | 0.013292 | 0.03134 |
| CNAG_00038 | 1309.458 | 1.1296 | 0.126323 | 8.942136 | 3.82E-19 | 1.63E-17 |
| CNAG_12005 | 24.19063 | 2.002427 | 0.488325 | 4.100601 | 4.12E-05 | 0.000197 |
| CNAG_00039 | 1175.732 | 0.474415 | 0.217846 | 2.177754 | 0.029424 | 0.061821 |
| CNAG_00040 | 3242.292 | -0.99312 | 0.181439 | -5.4736 | 4.41E-08 | 4.09E-07 |
| CNAG_07308 | 278.461 | 2.091331 | 0.310244 | 6.740923 | 1.57E-11 | 2.65E-10 |
| CNAG_12006 | 191.1548 | 1.25509 | 0.23486 | 5.343998 | 9.09E-08 | 7.90E-07 |
| CNAG_00045 | 635.6095 | 1.771358 | 0.15073 | 11.75184 | 6.91E-32 | 7.42E-30 |
| CNAG_00046 | 7114.898 | 0.376785 | 0.189046 | 1.99309 | 0.046252 | 0.090992 |
| CNAG_00047 | 576.7314 | 0.521276 | 0.169322 | 3.078605 | 0.00208 | 0.006336 |
| CNAG_00048 | 97.61766 | 0.680045 | 0.245766 | 2.767047 | 0.005657 | 0.015134 |
| CNAG_12007 | 3.074766 | 5.024041 | 1.715337 | 2.928894 | 0.003402 | 0.009678 |
| CNAG_00052 | 1228.553 | 1.577625 | 0.286861 | 5.499618 | 3.81E-08 | 3.62E-07 |
| CNAG_00053 | 321.0993 | -0.67494 | 0.168467 | -4.00636 | 6.17E-05 | 0.000283 |
| CNAG_00054 | 646.8075 | 2.963946 | 0.216598 | 13.68409 | 1.26E-42 | 2.39E-40 |
| CNAG_00055 | 1294.639 | -0.5708 | 0.165501 | -3.44894 | 0.000563 | 0.002012 |
| CNAG_00056 | 1227.067 | 0.783793 | 0.184364 | 4.251328 | 2.13E-05 | 0.000109 |
| CNAG_00057 | 10384.94 | 1.297369 | 0.203895 | 6.362942 | 1.98E-10 | 2.81E-09 |
| CNAG_00059 | 370.2805 | -1.07682 | 0.313899 | -3.43046 | 0.000603 | 0.002136 |
| CNAG_00061 | 25478.83 | -0.37916 | 0.129509 | -2.92768 | 0.003415 | 0.009712 |
| CNAG_00062 | 5608.088 | 0.66064 | 0.125676 | 5.256701 | 1.47E-07 | 1.21E-06 |
| CNAG_00063 | 1249.139 | -0.98638 | 0.190524 | -5.17717 | 2.25E-07 | 1.78E-06 |
| CNAG_12009 | 40.04248 | 1.049318 | 0.372366 | 2.817973 | 0.004833 | 0.013151 |

| | | | | | | |
|---|---|---|---|---|---|---|
| CNAG_12010 | 4.554955 | -2.60329 | 1.083202 | -2.40333 | 0.016246 | 0.037301 |
| CNAG_12011 | 10.06063 | -1.40166 | 0.690794 | -2.02905 | 0.042453 | 0.084544 |
| CNAG_00068 | 831.8577 | -1.39997 | 0.264058 | -5.30175 | 1.15E-07 | 9.72E-07 |
| CNAG_00070 | 1276.14 | 0.846871 | 0.145687 | 5.812936 | 6.14E-09 | 6.65E-08 |
| CNAG_00074 | 4812.398 | 0.482252 | 0.134519 | 3.585009 | 0.000337 | 0.001281 |
| CNAG_00076 | 1795.05 | 0.634464 | 0.128404 | 4.94114 | 7.77E-07 | 5.52E-06 |
| CNAG_07947 | 100.7648 | 1.144475 | 0.242348 | 4.722448 | 2.33E-06 | 1.51E-05 |
| CNAG_00077 | 5001.771 | 0.654687 | 0.129326 | 5.062288 | 4.14E-07 | 3.10E-06 |
| CNAG_00078 | 6241.714 | -0.87753 | 0.158769 | -5.52708 | 3.26E-08 | 3.16E-07 |
| CNAG_00079 | 635.3978 | -0.66939 | 0.221578 | -3.02101 | 0.002519 | 0.007492 |
| CNAG_00080 | 65.93933 | 0.786324 | 0.304342 | 2.583683 | 0.009775 | 0.024137 |
| CNAG_00081 | 3148.853 | -0.31255 | 0.132647 | -2.35625 | 0.01846 | 0.041601 |
| CNAG_00082 | 1441.44 | 0.326891 | 0.141434 | 2.311258 | 0.020819 | 0.046079 |
| CNAG_00084 | 3609.458 | -0.37701 | 0.130199 | -2.89567 | 0.003784 | 0.0106 |
| CNAG_00085 | 758.4885 | -0.4327 | 0.19308 | -2.24102 | 0.025025 | 0.05385 |
| CNAG_00086 | 1689.186 | -0.89102 | 0.133265 | -6.68613 | 2.29E-11 | 3.73E-10 |
| CNAG_00087 | 1704.526 | 0.365054 | 0.147416 | 2.476358 | 0.013273 | 0.031305 |
| CNAG_00088 | 4773.617 | 0.401168 | 0.127545 | 3.145311 | 0.001659 | 0.005184 |
| CNAG_00089 | 1337.913 | -0.38356 | 0.121135 | -3.16638 | 0.001543 | 0.004853 |
| CNAG_00090 | 1568.459 | -0.63676 | 0.17753 | -3.58677 | 0.000335 | 0.001274 |
| CNAG_00091 | 14286.99 | 3.104183 | 0.202213 | 15.35107 | 3.48E-53 | 9.67E-51 |
| CNAG_00093 | 5461.285 | 2.049661 | 0.185212 | 11.06655 | 1.82E-28 | 1.49E-26 |
| CNAG_00094 | 244.4781 | 0.70399 | 0.174012 | 4.045637 | 5.22E-05 | 0.000243 |
| CNAG_00097 | 459.3118 | 0.750498 | 0.163349 | 4.594436 | 4.34E-06 | 2.66E-05 |
| CNAG_12015 | 8.8246 | 2.139166 | 0.874208 | 2.446976 | 0.014406 | 0.033688 |
| CNAG_00098 | 1259.797 | 1.294253 | 0.172457 | 7.504799 | 6.15E-14 | 1.47E-12 |
| CNAG_00099 | 1806.048 | -0.62265 | 0.157527 | -3.95266 | 7.73E-05 | 0.000347 |
| CNAG_00101 | 1759.549 | -0.91647 | 0.230344 | -3.97871 | 6.93E-05 | 0.000315 |
| CNAG_07309 | 2292.427 | 0.392738 | 0.181844 | 2.159747 | 0.030792 | 0.064253 |
| CNAG_07310 | 1562.447 | -0.85472 | 0.228065 | -3.74769 | 0.000178 | 0.000733 |
| CNAG_00104 | 6612.922 | -0.44688 | 0.128769 | -3.47037 | 0.00052 | 0.001876 |
| CNAG_00107 | 2632.773 | 1.761914 | 0.179591 | 9.810721 | 1.01E-22 | 6.24E-21 |
| CNAG_00108 | 4585.241 | -0.40571 | 0.14858 | -2.73061 | 0.006322 | 0.016644 |
| CNAG_00109 | 1673.456 | -0.56566 | 0.214412 | -2.63822 | 0.008334 | 0.020997 |
| CNAG_00110 | 929.4177 | -1.02045 | 0.250304 | -4.07684 | 4.57E-05 | 0.000216 |
| CNAG_00111 | 3121.308 | -0.37624 | 0.178588 | -2.10673 | 0.035141 | 0.072022 |
| CNAG_07311 | 513.2772 | -0.86627 | 0.152065 | -5.69671 | 1.22E-08 | 1.26E-07 |
| CNAG_00116 | 36529.47 | -1.04897 | 0.258906 | -4.05156 | 5.09E-05 | 0.000238 |
| CNAG_00117 | 3549.399 | -0.35682 | 0.158814 | -2.24677 | 0.024655 | 0.053235 |
| CNAG_00120 | 753.3568 | -0.47347 | 0.185105 | -2.55786 | 0.010532 | 0.025746 |
| CNAG_00123 | 549.7308 | 1.201895 | 0.218959 | 5.489143 | 4.04E-08 | 3.79E-07 |
| CNAG_00124 | 987.6208 | 0.322449 | 0.156317 | 2.062792 | 0.039132 | 0.07878 |
| CNAG_00125 | 188.4452 | 1.280127 | 0.248106 | 5.159595 | 2.47E-07 | 1.93E-06 |
| CNAG_00126 | 2209.449 | -0.98138 | 0.207588 | -4.72754 | 2.27E-06 | 1.48E-05 |
| CNAG_00127 | 601.2443 | -0.70177 | 0.269544 | -2.60356 | 0.009226 | 0.022931 |
| CNAG_12018 | 293.7365 | -0.68443 | 0.208831 | -3.27743 | 0.001048 | 0.00345 |
| CNAG_12019 | 198.6296 | 1.482378 | 0.315942 | 4.691933 | 2.71E-06 | 1.73E-05 |

| | | | | | | |
|---|---|---|---|---|---|---|
| CNAG_00130 | 12416.55 | 1.064882 | 0.186699 | 5.703732 | 1.17E-08 | 1.22E-07 |
| CNAG_00132 | 189.9648 | 1.519539 | 0.247385 | 6.142399 | 8.13E-10 | 1.05E-08 |
| CNAG_00133 | 709.8663 | 0.947792 | 0.187527 | 5.054177 | 4.32E-07 | 3.23E-06 |
| CNAG_00134 | 813.6532 | 0.891723 | 0.257872 | 3.45801 | 0.000544 | 0.001954 |
| CNAG_00135 | 734.6138 | -0.54899 | 0.17597 | -3.11982 | 0.00181 | 0.005571 |
| CNAG_00136 | 7211.854 | 0.388748 | 0.146921 | 2.645965 | 0.008146 | 0.020601 |
| CNAG_00137 | 3113.142 | 1.105503 | 0.148136 | 7.462772 | 8.47E-14 | 1.97E-12 |
| CNAG_00138 | 2557.596 | -0.37254 | 0.113496 | -3.28244 | 0.001029 | 0.003402 |
| CNAG_00139 | 4942.07 | 1.207475 | 0.13957 | 8.651413 | 5.09E-18 | 1.88E-16 |
| CNAG_00141 | 1725.569 | 1.214476 | 0.1725 | 7.040427 | 1.92E-12 | 3.83E-11 |
| CNAG_12022 | 93.80807 | -1.23878 | 0.335096 | -3.69678 | 0.000218 | 0.000873 |
| CNAG_00143 | 4362.839 | -0.8286 | 0.154357 | -5.36811 | 7.96E-08 | 6.98E-07 |
| CNAG_00144 | 558.9319 | 0.847815 | 0.13114 | 6.464961 | 1.01E-10 | 1.50E-09 |
| CNAG_00145 | 1418.768 | -0.27406 | 0.139705 | -1.96169 | 0.049799 | 0.096915 |
| CNAG_00148 | 4541.728 | 0.59969 | 0.176997 | 3.388132 | 0.000704 | 0.002434 |
| CNAG_00149 | 7923.333 | -1.17129 | 0.270483 | -4.33036 | 1.49E-05 | 7.96E-05 |
| CNAG_00150 | 2958.692 | 0.33739 | 0.14616 | 2.308363 | 0.020979 | 0.046408 |
| CNAG_00151 | 2975.435 | -0.59753 | 0.128791 | -4.63953 | 3.49E-06 | 2.18E-05 |
| CNAG_00152 | 851.0682 | -0.65053 | 0.136832 | -4.75425 | 1.99E-06 | 1.30E-05 |
| CNAG_00154 | 1713.772 | 1.281942 | 0.153969 | 8.325978 | 8.36E-17 | 2.80E-15 |
| CNAG_12025 | 16.77819 | 1.80778 | 0.635173 | 2.846121 | 0.004426 | 0.012189 |
| CNAG_00156 | 1665.291 | 0.821659 | 0.171408 | 4.793582 | 1.64E-06 | 1.09E-05 |
| CNAG_00159 | 323.5826 | -0.44749 | 0.207405 | -2.15755 | 0.030963 | 0.064559 |
| CNAG_00161 | 425.5586 | -0.45306 | 0.162103 | -2.79491 | 0.005191 | 0.014045 |
| CNAG_00165 | 2767.733 | -0.64915 | 0.15371 | -4.22319 | 2.41E-05 | 0.000122 |
| CNAG_00166 | 974.1452 | -0.44888 | 0.180402 | -2.48819 | 0.012839 | 0.030491 |
| CNAG_00167 | 1037.24 | 0.522787 | 0.179942 | 2.905303 | 0.003669 | 0.010335 |
| CNAG_07314 | 649.9343 | 0.811257 | 0.156883 | 5.171093 | 2.33E-07 | 1.83E-06 |
| CNAG_07315 | 1696.62 | 0.379184 | 0.149292 | 2.539875 | 0.011089 | 0.026881 |
| CNAG_00172 | 707.6524 | -1.13431 | 0.305855 | -3.70866 | 0.000208 | 0.000839 |
| CNAG_00174 | 1233.859 | 0.537232 | 0.161076 | 3.335276 | 0.000852 | 0.002873 |
| CNAG_00175 | 875.7931 | 0.562366 | 0.208011 | 2.703539 | 0.006861 | 0.017815 |
| CNAG_00176 | 3652.366 | 0.412308 | 0.127799 | 3.226214 | 0.001254 | 0.004057 |
| CNAG_00177 | 108.1619 | 0.956787 | 0.286948 | 3.334356 | 0.000855 | 0.00288 |
| CNAG_00178 | 1589.974 | 0.874014 | 0.170862 | 5.115311 | 3.13E-07 | 2.40E-06 |
| CNAG_12030 | 27.81252 | -2.19372 | 0.487803 | -4.49715 | 6.89E-06 | 4.00E-05 |
| CNAG_00181 | 1129.181 | 0.628476 | 0.221718 | 2.834578 | 0.004589 | 0.012577 |
| CNAG_00182 | 1450.013 | 0.511795 | 0.133372 | 3.837357 | 0.000124 | 0.000528 |
| CNAG_00187 | 1865.022 | 0.28222 | 0.124136 | 2.273479 | 0.022997 | 0.050313 |
| CNAG_00189 | 881.7175 | 0.494136 | 0.142938 | 3.456984 | 0.000546 | 0.00196 |
| CNAG_00190 | 3281.729 | 1.488178 | 0.148575 | 10.01634 | 1.29E-23 | 8.21E-22 |
| CNAG_00191 | 496.8995 | -0.31002 | 0.149115 | -2.07909 | 0.037609 | 0.076411 |
| CNAG_00193 | 4120.448 | -0.87587 | 0.203722 | -4.29937 | 1.71E-05 | 9.06E-05 |
| CNAG_00194 | 2313.64 | -0.33574 | 0.124561 | -2.69534 | 0.007032 | 0.0182 |
| CNAG_07317 | 2171.182 | 0.536153 | 0.150522 | 3.56196 | 0.000368 | 0.00138 |
| CNAG_07319 | 3399.381 | 1.390531 | 0.207704 | 6.694763 | 2.16E-11 | 3.54E-10 |
| CNAG_07320 | 1140.437 | -0.28744 | 0.124476 | -2.30923 | 0.020931 | 0.046315 |

| | | | | | | |
|---|---|---|---|---|---|---|
| CNAG_07322 | 3392.788 | -0.53606 | 0.138001 | -3.88447 | 0.000103 | 0.000445 |
| CNAG_07324 | 1473.221 | 0.332143 | 0.157094 | 2.114301 | 0.03449 | 0.070833 |
| CNAG_07326 | 3061.892 | -0.64429 | 0.180575 | -3.56798 | 0.00036 | 0.001352 |
| CNAG_07334 | 647.8098 | -0.43076 | 0.199341 | -2.16094 | 0.0307 | 0.064112 |
| CNAG_07335 | 245.9256 | -0.61012 | 0.175878 | -3.46901 | 0.000522 | 0.001885 |
| CNAG_07030 | 497.9399 | -0.36398 | 0.161112 | -2.25919 | 0.023871 | 0.051911 |
| CNAG_07028 | 5156.24 | 0.743669 | 0.15496 | 4.799089 | 1.59E-06 | 1.06E-05 |
| CNAG_07338 | 951.7108 | 0.5898 | 0.152302 | 3.87258 | 0.000108 | 0.000465 |
| CNAG_07339 | 1666.75 | 0.386976 | 0.162224 | 2.385434 | 0.017059 | 0.038873 |
| CNAG_07342 | 1391.359 | 0.275253 | 0.132994 | 2.069657 | 0.038484 | 0.077772 |
| CNAG_00229 | 410.9604 | 1.247786 | 0.187084 | 6.66967 | 2.56E-11 | 4.14E-10 |
| CNAG_00231 | 577.895 | 0.520052 | 0.177711 | 2.926389 | 0.003429 | 0.009746 |
| CNAG_00232 | 25147.26 | -0.74439 | 0.252548 | -2.94752 | 0.003203 | 0.009166 |
| CNAG_00236 | 994.6797 | -0.56467 | 0.145485 | -3.8813 | 0.000104 | 0.00045 |
| CNAG_00237 | 9591.266 | -1.39178 | 0.152997 | -9.09675 | 9.31E-20 | 4.18E-18 |
| CNAG_00238 | 15158.99 | -0.6626 | 0.137551 | -4.81714 | 1.46E-06 | 9.75E-06 |
| CNAG_00239 | 628.0259 | 0.802601 | 0.166361 | 4.824451 | 1.40E-06 | 9.43E-06 |
| CNAG_00242 | 22.87801 | 0.966224 | 0.433158 | 2.23065 | 0.025704 | 0.055098 |
| CNAG_00243 | 431.6726 | -0.51517 | 0.181533 | -2.8379 | 0.004541 | 0.012469 |
| CNAG_07344 | 439.4747 | 1.27287 | 0.209603 | 6.07277 | 1.26E-09 | 1.57E-08 |
| CNAG_12039 | 113.9581 | -0.55323 | 0.224917 | -2.4597 | 0.013905 | 0.032651 |
| CNAG_00250 | 685.9511 | -1.18707 | 0.174565 | -6.80019 | 1.04E-11 | 1.83E-10 |
| CNAG_00251 | 27.47024 | 2.170126 | 0.444511 | 4.882054 | 1.05E-06 | 7.23E-06 |
| CNAG_00253 | 1181.997 | 0.691804 | 0.260619 | 2.654469 | 0.007943 | 0.020133 |
| CNAG_00254 | 1315.897 | 0.951537 | 0.241274 | 3.94381 | 8.02E-05 | 0.000359 |
| CNAG_00259 | 1832.175 | -0.54706 | 0.165686 | -3.30176 | 0.000961 | 0.003205 |
| CNAG_00260 | 2757.953 | -0.2572 | 0.117082 | -2.19672 | 0.028041 | 0.059353 |
| CNAG_00261 | 11436.57 | 0.441735 | 0.152337 | 2.899732 | 0.003735 | 0.010487 |
| CNAG_00263 | 740.8116 | 0.372474 | 0.142133 | 2.620592 | 0.008778 | 0.021934 |
| CNAG_00264 | 5324.956 | 0.339499 | 0.167947 | 2.021467 | 0.043231 | 0.085944 |
| CNAG_00265 | 1635.862 | -0.83652 | 0.145732 | -5.74014 | 9.46E-09 | 9.94E-08 |
| CNAG_00268 | 8108.835 | -0.74876 | 0.133856 | -5.59379 | 2.22E-08 | 2.21E-07 |
| CNAG_00270 | 4682.228 | -0.75682 | 0.278354 | -2.7189 | 0.00655 | 0.017121 |
| CNAG_00273 | 238.1654 | -0.70333 | 0.237539 | -2.96092 | 0.003067 | 0.00883 |
| CNAG_00279 | 5276.313 | -0.66306 | 0.213005 | -3.11288 | 0.001853 | 0.005695 |
| CNAG_00284 | 320.6185 | 0.766744 | 0.277129 | 2.76674 | 0.005662 | 0.015142 |
| CNAG_00285 | 1151.644 | 0.769233 | 0.251406 | 3.059726 | 0.002215 | 0.006696 |
| CNAG_00288 | 1486.843 | -0.71308 | 0.232913 | -3.06159 | 0.002202 | 0.006659 |
| CNAG_00292 | 4627.535 | -0.26956 | 0.10342 | -2.60645 | 0.009148 | 0.022767 |
| CNAG_00294 | 2326.582 | -0.52403 | 0.12307 | -4.25795 | 2.06E-05 | 0.000106 |
| CNAG_07347 | 10861.33 | 1.176981 | 0.236137 | 4.984323 | 6.22E-07 | 4.51E-06 |
| CNAG_07349 | 1865.677 | -0.72739 | 0.237192 | -3.06665 | 0.002165 | 0.006558 |
| CNAG_00301 | 1590.467 | 2.079908 | 0.190198 | 10.93551 | 7.80E-28 | 6.26E-26 |
| CNAG_00305 | 11356.29 | 0.262526 | 0.117006 | 2.243704 | 0.024851 | 0.053544 |
| CNAG_00308 | 1452.702 | 0.442324 | 0.19864 | 2.226763 | 0.025963 | 0.055547 |
| CNAG_00309 | 2132.22 | 0.631585 | 0.118795 | 5.316583 | 1.06E-07 | 9.04E-07 |
| CNAG_00313 | 1274.025 | 0.878312 | 0.201749 | 4.353477 | 1.34E-05 | 7.30E-05 |

| | | | | | | |
|---|---|---|---|---|---|---|
| CNAG_00314 | 728.5662 | 0.54522 | 0.131555 | 4.14442 | 3.41E-05 | 0.000167 |
| CNAG_07350 | 864.5517 | 1.251464 | 0.226683 | 5.520777 | 3.38E-08 | 3.26E-07 |
| CNAG_12047 | 26.21323 | 1.075715 | 0.436095 | 2.466701 | 0.013636 | 0.032067 |
| CNAG_00319 | 3616.281 | -0.59726 | 0.226206 | -2.64032 | 0.008283 | 0.0209 |
| CNAG_00320 | 448.795 | -0.54287 | 0.219716 | -2.47077 | 0.013482 | 0.03175 |
| CNAG_00321 | 83.07583 | 0.844062 | 0.285537 | 2.956055 | 0.003116 | 0.008958 |
| CNAG_00322 | 825.0981 | 0.289027 | 0.135744 | 2.129213 | 0.033237 | 0.068561 |
| CNAG_12048 | 114.7781 | 0.701396 | 0.26227 | 2.674323 | 0.007488 | 0.01915 |
| CNAG_00328 | 1493.481 | 0.852181 | 0.200807 | 4.243775 | 2.20E-05 | 0.000112 |
| CNAG_12049 | 7.944476 | 1.739555 | 0.80779 | 2.153474 | 0.031281 | 0.065137 |
| CNAG_00329 | 2764.446 | -1.05642 | 0.286507 | -3.68723 | 0.000227 | 0.000903 |
| CNAG_00330 | 1027.014 | -0.8926 | 0.154205 | -5.78843 | 7.10E-09 | 7.63E-08 |
| CNAG_00332 | 1854.704 | 0.424935 | 0.201023 | 2.113865 | 0.034527 | 0.070891 |
| CNAG_00334 | 62424.55 | -0.79585 | 0.114057 | -6.97769 | 3.00E-12 | 5.66E-11 |
| CNAG_00337 | 2484.078 | 1.342745 | 0.203699 | 6.591815 | 4.34E-11 | 6.74E-10 |
| CNAG_07352 | 6135.99 | -0.77036 | 0.254714 | -3.0244 | 0.002491 | 0.007417 |
| CNAG_00345 | 2395.399 | -0.30179 | 0.151864 | -1.98723 | 0.046897 | 0.092102 |
| CNAG_00347 | 1395.836 | 0.689529 | 0.323611 | 2.130732 | 0.033111 | 0.068356 |
| CNAG_00348 | 289.4319 | 0.856706 | 0.17021 | 5.03323 | 4.82E-07 | 3.57E-06 |
| CNAG_07356 | 2632.784 | -1.45821 | 0.211173 | -6.90527 | 5.01E-12 | 9.22E-11 |
| CNAG_00354 | 4947.166 | -0.32932 | 0.125734 | -2.61918 | 0.008814 | 0.022011 |
| CNAG_00357 | 441.0244 | 0.699336 | 0.245064 | 2.853686 | 0.004322 | 0.011931 |
| CNAG_12051 | 68.1 | 0.725297 | 0.267371 | 2.712706 | 0.006674 | 0.017404 |
| CNAG_00361 | 3846.398 | 0.265099 | 0.110954 | 2.389274 | 0.016882 | 0.038524 |
| CNAG_00363 | 5273.139 | 0.827581 | 0.186402 | 4.439774 | 9.01E-06 | 5.07E-05 |
| CNAG_00364 | 1312.879 | -0.5447 | 0.151184 | -3.60289 | 0.000315 | 0.001205 |
| CNAG_00365 | 2011.985 | -0.49688 | 0.127696 | -3.89112 | 9.98E-05 | 0.000435 |
| CNAG_00366 | 53.95228 | 1.073 | 0.293202 | 3.65959 | 0.000253 | 0.000993 |
| CNAG_00370 | 28965.89 | -0.90037 | 0.265248 | -3.39445 | 0.000688 | 0.00239 |
| CNAG_00374 | 1063.474 | 0.750069 | 0.200543 | 3.740193 | 0.000184 | 0.000752 |
| CNAG_00377 | 4131.759 | -0.73579 | 0.221158 | -3.32696 | 0.000878 | 0.002949 |
| CNAG_00384 | 524.8199 | 0.459938 | 0.169933 | 2.706595 | 0.006798 | 0.017675 |
| CNAG_00385 | 4026.385 | 0.517489 | 0.162268 | 3.189096 | 0.001427 | 0.004529 |
| CNAG_00386 | 5977.145 | -0.55275 | 0.147521 | -3.74689 | 0.000179 | 0.000735 |
| CNAG_00387 | 570.2841 | -0.40728 | 0.186684 | -2.18166 | 0.029135 | 0.061318 |
| CNAG_00389 | 1368.954 | -0.46817 | 0.132724 | -3.52742 | 0.00042 | 0.001549 |
| CNAG_07951 | 834.2274 | -0.61716 | 0.137359 | -4.49307 | 7.02E-06 | 4.07E-05 |
| CNAG_00392 | 334.8071 | -0.38986 | 0.196477 | -1.98424 | 0.047229 | 0.092616 |
| CNAG_00394 | 509.638 | -0.41283 | 0.209916 | -1.96664 | 0.049224 | 0.095914 |
| CNAG_00399 | 1908.943 | 1.560331 | 0.115634 | 13.49368 | 1.70E-41 | 3.15E-39 |
| CNAG_00400 | 2473.152 | 0.367127 | 0.142307 | 2.579818 | 0.009885 | 0.024379 |
| CNAG_00401 | 1139.729 | 0.646883 | 0.167891 | 3.852995 | 0.000117 | 0.0005 |
| CNAG_00403 | 844.4651 | -0.53771 | 0.200823 | -2.67755 | 0.007416 | 0.019004 |
| CNAG_00404 | 238.1282 | 0.368672 | 0.178699 | 2.063087 | 0.039104 | 0.078744 |
| CNAG_00405 | 713.3259 | 0.556939 | 0.162452 | 3.428324 | 0.000607 | 0.002146 |
| CNAG_00406 | 375.9941 | -1.1434 | 0.150807 | -7.58188 | 3.41E-14 | 8.43E-13 |
| CNAG_00407 | 8382.057 | 0.947475 | 0.218768 | 4.330957 | 1.48E-05 | 7.94E-05 |

| | | | | | | |
|---|---|---|---|---|---|---|
| CNAG_00409 | 1859.441 | 0.621118 | 0.18244 | 3.404511 | 0.000663 | 0.002318 |
| CNAG_00412 | 38.71099 | 0.705333 | 0.318516 | 2.214434 | 0.026799 | 0.05709 |
| CNAG_00413 | 3401.675 | -0.37355 | 0.137431 | -2.71807 | 0.006566 | 0.017158 |
| CNAG_00414 | 480.0561 | -0.7562 | 0.168098 | -4.49858 | 6.84E-06 | 3.98E-05 |
| CNAG_00415 | 573.3371 | -0.54172 | 0.172982 | -3.13165 | 0.001738 | 0.005387 |
| CNAG_00417 | 30819.58 | -0.57406 | 0.191532 | -2.99723 | 0.002724 | 0.008018 |
| CNAG_00418 | 17170.23 | -0.86134 | 0.10676 | -8.06801 | 7.15E-16 | 2.18E-14 |
| CNAG_12057 | 141.53 | 0.831665 | 0.227835 | 3.650292 | 0.000262 | 0.001027 |
| CNAG_00419 | 107.2509 | 1.342832 | 0.245581 | 5.467991 | 4.55E-08 | 4.21E-07 |
| CNAG_00423 | 519.2387 | -1.06393 | 0.238946 | -4.45257 | 8.48E-06 | 4.81E-05 |
| CNAG_00426 | 1410.388 | 0.431255 | 0.180449 | 2.389893 | 0.016853 | 0.038471 |
| CNAG_00428 | 736.1313 | -0.5367 | 0.204686 | -2.62208 | 0.00874 | 0.021866 |
| CNAG_00430 | 148.769 | -1.23285 | 0.217951 | -5.65654 | 1.54E-08 | 1.57E-07 |
| CNAG_07359 | 3215.254 | 1.010364 | 0.294636 | 3.429187 | 0.000605 | 0.002141 |
| CNAG_00434 | 1279.78 | -0.40817 | 0.195348 | -2.08944 | 0.036669 | 0.074767 |
| CNAG_00441 | 15276.6 | -0.34816 | 0.111589 | -3.11999 | 0.001809 | 0.00557 |
| CNAG_00442 | 1687.081 | -1.4616 | 0.158789 | -9.20466 | 3.43E-20 | 1.60E-18 |
| CNAG_00443 | 195.9744 | 0.944121 | 0.221005 | 4.271948 | 1.94E-05 | 0.000101 |
| CNAG_00444 | 374.1887 | -0.58261 | 0.257596 | -2.26171 | 0.023715 | 0.0516 |
| CNAG_00449 | 1555.141 | -0.87763 | 0.124882 | -7.02769 | 2.10E-12 | 4.14E-11 |
| CNAG_00450 | 8291.289 | -0.33364 | 0.155834 | -2.14098 | 0.032276 | 0.066957 |
| CNAG_00451 | 438.8633 | -0.67721 | 0.211576 | -3.20079 | 0.001371 | 0.004375 |
| CNAG_00456 | 10621.23 | -1.03292 | 0.117983 | -8.75485 | 2.04E-18 | 8.01E-17 |
| CNAG_00457 | 33987.9 | -0.89177 | 0.20648 | -4.31893 | 1.57E-05 | 8.36E-05 |
| CNAG_00458 | 2147.702 | -0.27054 | 0.1129 | -2.39624 | 0.016564 | 0.037891 |
| CNAG_00460 | 368.4203 | -0.94022 | 0.187458 | -5.01567 | 5.28E-07 | 3.89E-06 |
| CNAG_00461 | 5301.509 | 0.485483 | 0.155795 | 3.116162 | 0.001832 | 0.005634 |
| CNAG_00462 | 3139.424 | -1.86503 | 0.198874 | -9.37798 | 6.72E-21 | 3.38E-19 |
| CNAG_12060 | 375.9207 | 1.193931 | 0.158996 | 7.509206 | 5.95E-14 | 1.43E-12 |
| CNAG_00465 | 2815.914 | 1.508162 | 0.279552 | 5.394926 | 6.86E-08 | 6.11E-07 |
| CNAG_00466 | 866.7568 | 0.493875 | 0.196124 | 2.518185 | 0.011796 | 0.028309 |
| CNAG_00469 | 1897.183 | 0.418876 | 0.12819 | 3.267614 | 0.001085 | 0.003552 |
| CNAG_00474 | 3035.954 | 5.202829 | 0.178005 | 29.22863 | 8.39E-188 | 6.67E-184 |
| CNAG_07361 | 1902.057 | -0.79692 | 0.272947 | -2.9197 | 0.003504 | 0.009929 |
| CNAG_07362 | 8625.232 | -1.03553 | 0.200278 | -5.17047 | 2.34E-07 | 1.84E-06 |
| CNAG_07363 | 12011.51 | -0.37257 | 0.156416 | -2.38191 | 0.017223 | 0.039191 |
| CNAG_12061 | 6.308757 | 2.665066 | 0.902421 | 2.95324 | 0.003145 | 0.009024 |
| CNAG_00480 | 1943.8 | 1.508771 | 0.218651 | 6.900365 | 5.19E-12 | 9.52E-11 |
| CNAG_00485 | 2068.754 | 2.223513 | 0.198207 | 11.21812 | 3.32E-29 | 2.90E-27 |
| CNAG_00486 | 1839.694 | -0.63146 | 0.239495 | -2.63664 | 0.008373 | 0.021082 |
| CNAG_00487 | 115.5791 | 1.129111 | 0.371907 | 3.036007 | 0.002397 | 0.007167 |
| CNAG_00488 | 334.4608 | 0.478804 | 0.229522 | 2.086093 | 0.03697 | 0.075296 |
| CNAG_00492 | 64.54477 | 1.397992 | 0.295164 | 4.736325 | 2.18E-06 | 1.42E-05 |
| CNAG_00493 | 757.0207 | 0.606549 | 0.307596 | 1.971901 | 0.048621 | 0.094971 |
| CNAG_07364 | 835.2002 | 0.537956 | 0.1241 | 4.334857 | 1.46E-05 | 7.85E-05 |
| CNAG_00497 | 4198.742 | 1.585721 | 0.175049 | 9.058741 | 1.32E-19 | 5.79E-18 |
| CNAG_00498 | 739.1602 | 0.668485 | 0.163563 | 4.087024 | 4.37E-05 | 0.000208 |

| | | | | | | |
|---|---|---|---|---|---|---|
| CNAG_00499 | 854.8278 | -0.47743 | 0.210409 | -2.26907 | 0.023264 | 0.050813 |
| CNAG_00504 | 1346.898 | -0.43231 | 0.163926 | -2.6372 | 0.008359 | 0.021053 |
| CNAG_00506 | 762.0238 | -0.60657 | 0.144302 | -4.2035 | 2.63E-05 | 0.000132 |
| CNAG_00508 | 1214.985 | 0.484478 | 0.199636 | 2.426809 | 0.015232 | 0.035339 |
| CNAG_00509 | 8488.024 | -0.65097 | 0.179143 | -3.63379 | 0.000279 | 0.001088 |
| CNAG_00510 | 715.1129 | -0.63505 | 0.168557 | -3.76759 | 0.000165 | 0.000683 |
| CNAG_00511 | 105.2976 | 0.486928 | 0.223209 | 2.181491 | 0.029147 | 0.061319 |
| CNAG_00512 | 2857.96 | -0.49842 | 0.109464 | -4.5533 | 5.28E-06 | 3.16E-05 |
| CNAG_00513 | 2453.897 | -0.80181 | 0.141103 | -5.6824 | 1.33E-08 | 1.36E-07 |
| CNAG_00515 | 331.2852 | 1.459113 | 0.177151 | 8.236537 | 1.77E-16 | 5.77E-15 |
| CNAG_00516 | 229.6356 | -0.42307 | 0.189599 | -2.2314 | 0.025655 | 0.05502 |
| CNAG_07365 | 1196.379 | 0.284344 | 0.140689 | 2.021085 | 0.043271 | 0.086001 |
| CNAG_00519 | 2621.611 | -0.57179 | 0.146447 | -3.90442 | 9.45E-05 | 0.000415 |
| CNAG_07953 | 964.5317 | -0.88131 | 0.22327 | -3.9473 | 7.90E-05 | 0.000354 |
| CNAG_00521 | 2038.79 | 1.134141 | 0.159963 | 7.090044 | 1.34E-12 | 2.73E-11 |
| CNAG_00522 | 5695.671 | 1.641786 | 0.187698 | 8.746979 | 2.19E-18 | 8.49E-17 |
| CNAG_00524 | 1012.266 | 0.532561 | 0.146915 | 3.624959 | 0.000289 | 0.001121 |
| CNAG_00526 | 43.72438 | 0.999037 | 0.344234 | 2.9022 | 0.003706 | 0.010426 |
| CNAG_00527 | 3668.856 | -0.58866 | 0.126112 | -4.66774 | 3.05E-06 | 1.93E-05 |
| CNAG_00528 | 819.3239 | 0.471988 | 0.147739 | 3.194744 | 0.0014 | 0.004454 |
| CNAG_00531 | 6984.988 | 0.387301 | 0.188301 | 2.056813 | 0.039704 | 0.07983 |
| CNAG_12064 | 41.5078 | 0.812343 | 0.37944 | 2.140901 | 0.032282 | 0.066957 |
| CNAG_00532 | 1082.25 | -0.78898 | 0.161686 | -4.87969 | 1.06E-06 | 7.31E-06 |
| CNAG_00534 | 5030.711 | -0.7103 | 0.267734 | -2.65301 | 0.007978 | 0.020206 |
| CNAG_00535 | 2350.055 | -0.48966 | 0.186588 | -2.6243 | 0.008683 | 0.021744 |
| CNAG_00536 | 2352.343 | -1.24491 | 0.277256 | -4.49009 | 7.12E-06 | 4.12E-05 |
| CNAG_00537 | 1413.67 | -0.73012 | 0.275013 | -2.65488 | 0.007934 | 0.020122 |
| CNAG_00538 | 1783.586 | -0.81843 | 0.205467 | -3.98328 | 6.80E-05 | 0.000309 |
| CNAG_00543 | 3015.819 | 1.015438 | 0.209241 | 4.852957 | 1.22E-06 | 8.25E-06 |
| CNAG_00544 | 235.548 | 0.571139 | 0.213864 | 2.670574 | 0.007572 | 0.019322 |
| CNAG_00546 | 1709.191 | 1.039473 | 0.189042 | 5.498637 | 3.83E-08 | 3.63E-07 |
| CNAG_00554 | 1195.205 | 0.84635 | 0.226658 | 3.734043 | 0.000188 | 0.000769 |
| CNAG_00556 | 3027.28 | -0.35113 | 0.169803 | -2.06785 | 0.038654 | 0.078055 |
| CNAG_00557 | 1884.558 | -0.60157 | 0.154331 | -3.89788 | 9.70E-05 | 0.000424 |
| CNAG_12074 | 36.52627 | -1.13837 | 0.542557 | -2.09815 | 0.035892 | 0.073466 |
| CNAG_00559 | 2094.89 | 0.81063 | 0.19693 | 4.116341 | 3.85E-05 | 0.000186 |
| CNAG_00561 | 3168.076 | -0.31759 | 0.110409 | -2.87648 | 0.004021 | 0.011173 |
| CNAG_00562 | 1518.15 | -0.39446 | 0.127164 | -3.10195 | 0.001922 | 0.005887 |
| CNAG_00563 | 1170.534 | 0.687669 | 0.146411 | 4.696824 | 2.64E-06 | 1.70E-05 |
| CNAG_00564 | 636.9833 | 0.566827 | 0.138831 | 4.082842 | 4.45E-05 | 0.000211 |
| CNAG_00565 | 10392.76 | 0.552555 | 0.12724 | 4.342613 | 1.41E-05 | 7.61E-05 |
| CNAG_00566 | 1298.05 | 0.714141 | 0.233579 | 3.057381 | 0.002233 | 0.006741 |
| CNAG_00568 | 59.20934 | 0.653916 | 0.295952 | 2.209533 | 0.027138 | 0.057734 |
| CNAG_00569 | 2327.381 | 0.31534 | 0.132205 | 2.385231 | 0.017068 | 0.038883 |
| CNAG_00573 | 1818.162 | -1.046 | 0.197563 | -5.2945 | 1.19E-07 | 1.00E-06 |
| CNAG_00575 | 2186.642 | 2.311931 | 0.197147 | 11.72692 | 9.28E-32 | 9.45E-30 |
| CNAG_07369 | 48.68429 | 0.799991 | 0.307578 | 2.600936 | 0.009297 | 0.023082 |

| | | | | | | |
|---|---|---|---|---|---|---|
| CNAG_00581 | 14259.77 | 0.371226 | 0.171635 | 2.162878 | 0.030551 | 0.063849 |
| CNAG_00582 | 2000.016 | 0.519137 | 0.17396 | 2.984237 | 0.002843 | 0.008308 |
| CNAG_00586 | 203.2791 | 1.503659 | 0.202505 | 7.425282 | 1.13E-13 | 2.58E-12 |
| CNAG_00587 | 90.04277 | 0.672925 | 0.251147 | 2.679406 | 0.007375 | 0.018917 |
| CNAG_00588 | 80.30999 | 0.618519 | 0.302191 | 2.046783 | 0.040679 | 0.081605 |
| CNAG_00589 | 1226.122 | -0.69266 | 0.114787 | -6.03431 | 1.60E-09 | 1.97E-08 |
| CNAG_00590 | 2101.532 | -0.46798 | 0.166052 | -2.81828 | 0.004828 | 0.013143 |
| CNAG_00591 | 83.84999 | 1.0963 | 0.255278 | 4.29453 | 1.75E-05 | 9.24E-05 |
| CNAG_00592 | 1810.156 | -0.67537 | 0.272421 | -2.47914 | 0.01317 | 0.031125 |
| CNAG_00595 | 565.2274 | 1.457975 | 0.201112 | 7.24957 | 4.18E-13 | 9.10E-12 |
| CNAG_00596 | 114.5575 | 0.944597 | 0.297654 | 3.173473 | 0.001506 | 0.00475 |
| CNAG_00597 | 450.4421 | -0.42371 | 0.203065 | -2.08655 | 0.036929 | 0.07524 |
| CNAG_00598 | 148.3201 | -1.51194 | 0.251418 | -6.01364 | 1.81E-09 | 2.22E-08 |
| CNAG_00600 | 4314.558 | -0.46052 | 0.119983 | -3.83821 | 0.000124 | 0.000527 |
| CNAG_00603 | 1179.079 | -0.53867 | 0.142362 | -3.7838 | 0.000154 | 0.000641 |
| CNAG_00605 | 2361.089 | 1.75609 | 0.249968 | 7.025263 | 2.14E-12 | 4.20E-11 |
| CNAG_00607 | 1253.43 | -0.90089 | 0.186803 | -4.82267 | 1.42E-06 | 9.51E-06 |
| CNAG_00608 | 2067.548 | -0.85243 | 0.152107 | -5.60415 | 2.09E-08 | 2.08E-07 |
| CNAG_07957 | 1590.415 | -0.43196 | 0.11914 | -3.62561 | 0.000288 | 0.00112 |
| CNAG_00612 | 819.9033 | 0.611119 | 0.280823 | 2.176169 | 0.029543 | 0.062029 |
| CNAG_00613 | 960.8369 | -0.74588 | 0.216672 | -3.44244 | 0.000576 | 0.002053 |
| CNAG_07374 | 963.4661 | -0.97917 | 0.178073 | -5.49871 | 3.83E-08 | 3.63E-07 |
| CNAG_00622 | 4231.882 | -0.63595 | 0.109023 | -5.83313 | 5.44E-09 | 5.95E-08 |
| CNAG_00623 | 1382.955 | 0.448206 | 0.119876 | 3.7389 | 0.000185 | 0.000755 |
| CNAG_00625 | 2522.288 | 0.504401 | 0.217429 | 2.319845 | 0.020349 | 0.045229 |
| CNAG_00627 | 153.2503 | 1.571544 | 0.245975 | 6.389049 | 1.67E-10 | 2.39E-09 |
| CNAG_00628 | 442.1569 | -0.81361 | 0.213451 | -3.81168 | 0.000138 | 0.000582 |
| CNAG_07375 | 2682.367 | 0.537626 | 0.273979 | 1.962292 | 0.049728 | 0.096801 |
| CNAG_00637 | 602.9653 | 0.835237 | 0.236536 | 3.53112 | 0.000414 | 0.001532 |
| CNAG_00638 | 3592.542 | 1.029417 | 0.175524 | 5.864834 | 4.50E-09 | 5.01E-08 |
| CNAG_00639 | 805.6647 | -0.69988 | 0.242572 | -2.88523 | 0.003911 | 0.010894 |
| CNAG_00640 | 73231.75 | -0.96675 | 0.194834 | -4.96193 | 6.98E-07 | 5.01E-06 |
| CNAG_07958 | 151.6408 | -0.59542 | 0.273713 | -2.17534 | 0.029605 | 0.062118 |
| CNAG_00642 | 851.6809 | 1.495653 | 0.221677 | 6.746996 | 1.51E-11 | 2.55E-10 |
| CNAG_12086 | 43.15723 | -0.86826 | 0.345535 | -2.5128 | 0.011978 | 0.028684 |
| CNAG_00643 | 828.8099 | -0.53936 | 0.222746 | -2.42143 | 0.01546 | 0.035792 |
| CNAG_07959 | 1494.993 | -0.44349 | 0.179398 | -2.47208 | 0.013433 | 0.031654 |
| CNAG_00648 | 1158.281 | 0.770941 | 0.144811 | 5.323767 | 1.02E-07 | 8.72E-07 |
| CNAG_00649 | 8261.979 | -0.61816 | 0.10522 | -5.87491 | 4.23E-09 | 4.74E-08 |
| CNAG_00651 | 388.1655 | 0.432265 | 0.220526 | 1.960152 | 0.049978 | 0.097216 |
| CNAG_00652 | 930.9735 | 1.132603 | 0.123458 | 9.17403 | 4.56E-20 | 2.08E-18 |
| CNAG_00653 | 176.784 | 1.496026 | 0.305537 | 4.896387 | 9.76E-07 | 6.75E-06 |
| CNAG_00655 | 36149.24 | -1.2312 | 0.296462 | -4.15297 | 3.28E-05 | 0.000161 |
| CNAG_00656 | 45901.42 | -0.93131 | 0.158403 | -5.87939 | 4.12E-09 | 4.63E-08 |
| CNAG_00663 | 1087.836 | 0.770821 | 0.150565 | 5.119521 | 3.06E-07 | 2.35E-06 |
| CNAG_00664 | 40.81223 | 1.115947 | 0.395157 | 2.82406 | 0.004742 | 0.012939 |
| CNAG_00665 | 3908.93 | -0.3124 | 0.146491 | -2.13257 | 0.03296 | 0.068115 |

| | | | | | | |
|---|---|---|---|---|---|---|
| CNAG_00667 | 1186.672 | -0.27831 | 0.133776 | -2.08042 | 0.037487 | 0.076273 |
| CNAG_00668 | 3439.944 | 0.654524 | 0.126817 | 5.161159 | 2.45E-07 | 1.92E-06 |
| CNAG_00669 | 796.3758 | -0.79586 | 0.192464 | -4.13511 | 3.55E-05 | 0.000173 |
| CNAG_00672 | 35968.67 | -0.93162 | 0.245625 | -3.79285 | 0.000149 | 0.000621 |
| CNAG_00677 | 797.4365 | -1.02962 | 0.162935 | -6.31919 | 2.63E-10 | 3.66E-09 |
| CNAG_00679 | 205.1835 | 1.618185 | 0.215531 | 7.507893 | 6.01E-14 | 1.44E-12 |
| CNAG_00680 | 1233.165 | -1.11868 | 0.14207 | -7.87412 | 3.43E-15 | 9.60E-14 |
| CNAG_00681 | 1162.679 | -0.97032 | 0.309376 | -3.1364 | 0.00171 | 0.005311 |
| CNAG_00682 | 4184.914 | 0.765038 | 0.196197 | 3.899324 | 9.65E-05 | 0.000422 |
| CNAG_00686 | 18784.4 | 0.800413 | 0.229019 | 3.494963 | 0.000474 | 0.001723 |
| CNAG_00690 | 2087.902 | -0.48554 | 0.126443 | -3.83996 | 0.000123 | 0.000524 |
| CNAG_00692 | 478.1794 | 0.574901 | 0.223469 | 2.572623 | 0.010093 | 0.024799 |
| CNAG_00694 | 6284.553 | -0.35426 | 0.110283 | -3.21229 | 0.001317 | 0.004234 |
| CNAG_00695 | 567.9167 | 0.516141 | 0.221586 | 2.329303 | 0.019843 | 0.04424 |
| CNAG_00696 | 1888.03 | 1.259401 | 0.169367 | 7.43594 | 1.04E-13 | 2.40E-12 |
| CNAG_00697 | 4093.38 | -0.52852 | 0.105592 | -5.00532 | 5.58E-07 | 4.09E-06 |
| CNAG_00699 | 889.2494 | -0.39915 | 0.143503 | -2.78151 | 0.005411 | 0.01454 |
| CNAG_00700 | 11990.85 | -0.30103 | 0.103698 | -2.90293 | 0.003697 | 0.010406 |
| CNAG_00701 | 2948.798 | -0.53429 | 0.106782 | -5.00356 | 5.63E-07 | 4.12E-06 |
| CNAG_12093 | 256.6175 | -1.15402 | 0.287278 | -4.01708 | 5.89E-05 | 0.000271 |
| CNAG_00702 | 6628.18 | -1.47623 | 0.232532 | -6.34852 | 2.17E-10 | 3.07E-09 |
| CNAG_00703 | 36034.78 | -1.11436 | 0.284919 | -3.91116 | 9.19E-05 | 0.000405 |
| CNAG_00705 | 3845.603 | -0.35867 | 0.138586 | -2.58807 | 0.009651 | 0.023854 |
| CNAG_00706 | 1397.196 | -0.8161 | 0.163855 | -4.98066 | 6.34E-07 | 4.59E-06 |
| CNAG_00709 | 1453.971 | -0.62783 | 0.15245 | -4.11826 | 3.82E-05 | 0.000185 |
| CNAG_00711 | 770.885 | 0.993451 | 0.170346 | 5.831963 | 5.48E-09 | 5.99E-08 |
| CNAG_00714 | 706.6952 | -0.36064 | 0.168999 | -2.134 | 0.032843 | 0.067919 |
| CNAG_00715 | 1373.828 | -1.16246 | 0.236604 | -4.91309 | 8.97E-07 | 6.27E-06 |
| CNAG_00716 | 11839.18 | -1.67838 | 0.252929 | -6.63578 | 3.23E-11 | 5.14E-10 |
| CNAG_00717 | 1318.785 | 0.999995 | 0.133367 | 7.498092 | 6.48E-14 | 1.54E-12 |
| CNAG_00719 | 2205.397 | -0.42079 | 0.118725 | -3.54421 | 0.000394 | 0.001465 |
| CNAG_00720 | 2106.588 | 0.503796 | 0.123786 | 4.069898 | 4.70E-05 | 0.000222 |
| CNAG_00721 | 4042.611 | -0.80442 | 0.165505 | -4.86036 | 1.17E-06 | 7.97E-06 |
| CNAG_00722 | 1899.656 | -1.01347 | 0.194077 | -5.222 | 1.77E-07 | 1.44E-06 |
| CNAG_07381 | 1036.483 | -1.03137 | 0.24447 | -4.21878 | 2.46E-05 | 0.000124 |
| CNAG_07382 | 3684.423 | -0.65637 | 0.120743 | -5.43605 | 5.45E-08 | 4.96E-07 |
| CNAG_00727 | 1879.884 | 0.293696 | 0.128055 | 2.293518 | 0.021818 | 0.047989 |
| CNAG_00728 | 479.5886 | 1.044602 | 0.14817 | 7.050017 | 1.79E-12 | 3.59E-11 |
| CNAG_12097 | 19.67878 | -0.9273 | 0.438919 | -2.11269 | 0.034628 | 0.071062 |
| CNAG_00730 | 6302.859 | 1.74464 | 0.263937 | 6.610066 | 3.84E-11 | 6.00E-10 |
| CNAG_00732 | 2740.43 | 1.323729 | 0.199766 | 6.626381 | 3.44E-11 | 5.43E-10 |
| CNAG_12100 | 48.427 | -1.02568 | 0.337068 | -3.04294 | 0.002343 | 0.007023 |
| CNAG_00735 | 145.8135 | -0.57495 | 0.230573 | -2.49355 | 0.012647 | 0.030089 |
| CNAG_00737 | 1027.735 | 0.62525 | 0.234861 | 2.662212 | 0.007763 | 0.019739 |
| CNAG_12101 | 59.48457 | -0.75753 | 0.335571 | -2.25744 | 0.02398 | 0.052048 |
| CNAG_00742 | 1886.503 | -0.3203 | 0.107555 | -2.97799 | 0.002901 | 0.008442 |
| CNAG_00743 | 3224.951 | -1.2946 | 0.127571 | -10.1481 | 3.38E-24 | 2.26E-22 |

| | | | | | | |
|---|---|---|---|---|---|---|
| CNAG_00744 | 2590.119 | 0.712539 | 0.129604 | 5.497803 | 3.85E-08 | 3.65E-07 |
| CNAG_00745 | 2981.53 | 0.614272 | 0.233352 | 2.63239 | 0.008479 | 0.021334 |
| CNAG_00746 | 6755.083 | -0.51318 | 0.113466 | -4.52278 | 6.10E-06 | 3.60E-05 |
| CNAG_00747 | 11163.83 | -0.31737 | 0.149272 | -2.12615 | 0.033491 | 0.068996 |
| CNAG_12105 | 35.3136 | 0.6871 | 0.336741 | 2.040443 | 0.041306 | 0.082702 |
| CNAG_00750 | 1238.757 | -1.51607 | 0.260529 | -5.8192 | 5.91E-09 | 6.43E-08 |
| CNAG_12108 | 42.23534 | 1.470334 | 0.355183 | 4.139656 | 3.48E-05 | 0.00017 |
| CNAG_00751 | 18.48476 | 1.210608 | 0.593991 | 2.03809 | 0.041541 | 0.08304 |
| CNAG_12109 | 50.58834 | -2.18414 | 0.382908 | -5.70408 | 1.17E-08 | 1.22E-07 |
| CNAG_00754 | 4155.842 | -1.03913 | 0.162601 | -6.39069 | 1.65E-10 | 2.38E-09 |
| CNAG_00756 | 667.002 | -0.47278 | 0.202767 | -2.33165 | 0.019719 | 0.044026 |
| CNAG_12110 | 105.4465 | 0.927726 | 0.241896 | 3.835223 | 0.000125 | 0.000533 |
| CNAG_00758 | 556.7042 | 0.919075 | 0.155654 | 5.904615 | 3.53E-09 | 4.05E-08 |
| CNAG_00762 | 706.3651 | -0.70183 | 0.128689 | -5.45373 | 4.93E-08 | 4.53E-07 |
| CNAG_07960 | 599.0552 | 0.720213 | 0.15979 | 4.507252 | 6.57E-06 | 3.84E-05 |
| CNAG_00764 | 2565.711 | -0.88695 | 0.149174 | -5.94574 | 2.75E-09 | 3.23E-08 |
| CNAG_00765 | 629.6993 | 0.799692 | 0.162543 | 4.919868 | 8.66E-07 | 6.08E-06 |
| CNAG_00766 | 1480.123 | 1.494147 | 0.186834 | 7.997186 | 1.27E-15 | 3.80E-14 |
| CNAG_00769 | 1418.106 | 0.906879 | 0.321053 | 2.824703 | 0.004732 | 0.012918 |
| CNAG_00771 | 14687.79 | -1.02302 | 0.150084 | -6.81631 | 9.34E-12 | 1.65E-10 |
| CNAG_00774 | 7346.359 | -0.75307 | 0.224616 | -3.35269 | 0.0008 | 0.002727 |
| CNAG_00775 | 1653.57 | -0.33986 | 0.125244 | -2.71355 | 0.006657 | 0.017365 |
| CNAG_12113 | 40.39968 | 1.009595 | 0.423213 | 2.385548 | 0.017054 | 0.038872 |
| CNAG_00776 | 15106.37 | 0.593483 | 0.196883 | 3.014399 | 0.002575 | 0.007632 |
| CNAG_00778 | 1813.946 | 0.445248 | 0.125528 | 3.546993 | 0.00039 | 0.001454 |
| CNAG_00779 | 33054.15 | -0.91307 | 0.236482 | -3.86105 | 0.000113 | 0.000485 |
| CNAG_00785 | 28035.37 | -0.67499 | 0.13563 | -4.9767 | 6.47E-07 | 4.68E-06 |
| CNAG_00788 | 14516.38 | -1.31548 | 0.158918 | -8.27773 | 1.26E-16 | 4.17E-15 |
| CNAG_12114 | 202.5559 | -1.27556 | 0.250049 | -5.10122 | 3.37E-07 | 2.57E-06 |
| CNAG_00791 | 2291.568 | 0.66852 | 0.259044 | 2.580717 | 0.00986 | 0.024323 |
| CNAG_00792 | 2549.496 | 0.628568 | 0.318802 | 1.971654 | 0.048649 | 0.095003 |
| CNAG_00793 | 1959.694 | -0.77639 | 0.166514 | -4.66259 | 3.12E-06 | 1.97E-05 |
| CNAG_00794 | 2048.05 | -0.34423 | 0.164139 | -2.09721 | 0.035975 | 0.07356 |
| CNAG_12117 | 165.028 | 1.646461 | 0.246291 | 6.685023 | 2.31E-11 | 3.75E-10 |
| CNAG_00796 | 2259.353 | 1.85166 | 0.228321 | 8.109898 | 5.07E-16 | 1.58E-14 |
| CNAG_00797 | 6413.662 | -0.92175 | 0.107974 | -8.53674 | 1.38E-17 | 4.96E-16 |
| CNAG_07961 | 73.00434 | 0.820332 | 0.281481 | 2.914344 | 0.003564 | 0.010079 |
| CNAG_00798 | 166.5638 | 2.421987 | 0.389981 | 6.210519 | 5.28E-10 | 6.95E-09 |
| CNAG_00799 | 14150.75 | 0.923197 | 0.174425 | 5.292798 | 1.20E-07 | 1.01E-06 |
| CNAG_00801 | 194.2578 | 1.028147 | 0.238279 | 4.314884 | 1.60E-05 | 8.49E-05 |
| CNAG_00802 | 648.0187 | 1.136253 | 0.196788 | 5.77401 | 7.74E-09 | 8.27E-08 |
| CNAG_00803 | 1573.296 | 0.29569 | 0.134835 | 2.19298 | 0.028309 | 0.059809 |
| CNAG_00804 | 2269.403 | -0.51762 | 0.148193 | -3.49287 | 0.000478 | 0.001735 |
| CNAG_00807 | 1271.495 | 0.841293 | 0.240093 | 3.504035 | 0.000458 | 0.00167 |
| CNAG_00808 | 3899.354 | -0.70982 | 0.303966 | -2.3352 | 0.019533 | 0.043697 |
| CNAG_00810 | 1251.889 | -0.47452 | 0.133004 | -3.56771 | 0.00036 | 0.001353 |
| CNAG_00811 | 3173.971 | -1.02033 | 0.226371 | -4.50735 | 6.56E-06 | 3.84E-05 |

| | | | | | | |
|---|---|---|---|---|---|---|
| CNAG_00814 | 117.9764 | 1.157623 | 0.291911 | 3.965668 | 7.32E-05 | 0.000331 |
| CNAG_00813 | 11.22133 | 3.117156 | 0.811153 | 3.84287 | 0.000122 | 0.000519 |
| CNAG_00816 | 3125.245 | 0.53238 | 0.170245 | 3.127152 | 0.001765 | 0.005463 |
| CNAG_00819 | 12244.56 | -0.92934 | 0.321421 | -2.89135 | 0.003836 | 0.010714 |
| CNAG_00820 | 789.3654 | -1.31749 | 0.147298 | -8.94438 | 3.74E-19 | 1.61E-17 |
| CNAG_00821 | 21710.82 | -0.99752 | 0.290762 | -3.43072 | 0.000602 | 0.002134 |
| CNAG_00822 | 4017.661 | -0.62876 | 0.121061 | -5.19373 | 2.06E-07 | 1.64E-06 |
| CNAG_12119 | 55.77802 | 1.593348 | 0.30321 | 5.254931 | 1.48E-07 | 1.22E-06 |
| CNAG_00826 | 305.1559 | 2.349705 | 0.178765 | 13.14409 | 1.84E-39 | 3.05E-37 |
| CNAG_12120 | 5.511598 | 2.740832 | 1.097299 | 2.497798 | 0.012497 | 0.029775 |
| CNAG_00827 | 71.5498 | 1.266973 | 0.345197 | 3.67029 | 0.000242 | 0.000956 |
| CNAG_00830 | 884.198 | 0.879699 | 0.224857 | 3.912263 | 9.14E-05 | 0.000403 |
| CNAG_00831 | 925.1566 | 0.431376 | 0.143395 | 3.008298 | 0.002627 | 0.007772 |
| CNAG_12122 | 131.3051 | 0.56759 | 0.259511 | 2.187151 | 0.028732 | 0.060573 |
| CNAG_00833 | 79.39345 | 0.785137 | 0.26568 | 2.9552 | 0.003125 | 0.008973 |
| CNAG_00834 | 2391.731 | 2.359302 | 0.15568 | 15.15481 | 7.04E-52 | 1.80E-49 |
| CNAG_00835 | 554.9271 | 0.37775 | 0.178942 | 2.111016 | 0.034771 | 0.071337 |
| CNAG_00836 | 484.7087 | -0.665 | 0.170232 | -3.90645 | 9.37E-05 | 0.000412 |
| CNAG_00838 | 253.3647 | -0.7519 | 0.316678 | -2.37433 | 0.017581 | 0.03989 |
| CNAG_00841 | 291.1984 | -0.59088 | 0.242563 | -2.43596 | 0.014852 | 0.034564 |
| CNAG_07386 | 115.9505 | -0.90739 | 0.232454 | -3.90353 | 9.48E-05 | 0.000416 |
| CNAG_00843 | 175.4271 | 0.750513 | 0.236234 | 3.176991 | 0.001488 | 0.0047 |
| CNAG_00845 | 1573.907 | -0.9103 | 0.114892 | -7.92314 | 2.32E-15 | 6.60E-14 |
| CNAG_00846 | 681.4978 | -0.6246 | 0.137049 | -4.5575 | 5.18E-06 | 3.11E-05 |
| CNAG_00848 | 20564.91 | 4.218003 | 0.190524 | 22.13899 | 1.33E-108 | 2.65E-105 |
| CNAG_00849 | 150.0275 | 1.545251 | 0.214515 | 7.203464 | 5.87E-13 | 1.25E-11 |
| CNAG_00850 | 188.9952 | 0.556859 | 0.262098 | 2.124626 | 0.033618 | 0.06919 |
| CNAG_00851 | 179.6787 | 1.491693 | 0.417955 | 3.569025 | 0.000358 | 0.001347 |
| CNAG_00853 | 1256.51 | -0.56954 | 0.192056 | -2.96548 | 0.003022 | 0.008732 |
| CNAG_00854 | 2773.346 | -0.63336 | 0.21175 | -2.99107 | 0.00278 | 0.008154 |
| CNAG_07389 | 698.214 | 0.349427 | 0.134588 | 2.596267 | 0.009424 | 0.023372 |
| CNAG_00859 | 142.5939 | -0.94902 | 0.192072 | -4.94093 | 7.78E-07 | 5.52E-06 |
| CNAG_07459 | 221.5472 | 0.376822 | 0.181468 | 2.076522 | 0.037846 | 0.076813 |
| CNAG_12128 | 12.17184 | 1.575551 | 0.581165 | 2.71102 | 0.006708 | 0.017481 |
| CNAG_06800 | 3342.958 | 2.422315 | 0.251135 | 9.645482 | 5.14E-22 | 2.92E-20 |
| CNAG_06799 | 3695.602 | -0.69172 | 0.25751 | -2.6862 | 0.007227 | 0.018609 |
| CNAG_06798 | 1749.281 | -0.62876 | 0.139958 | -4.49246 | 7.04E-06 | 4.08E-05 |
| CNAG_06796 | 1265.656 | -0.31926 | 0.133235 | -2.39621 | 0.016566 | 0.037891 |
| CNAG_06792 | 1482.308 | 0.39574 | 0.175909 | 2.249678 | 0.024469 | 0.052936 |
| CNAG_06791 | 2571.634 | 1.44387 | 0.242137 | 5.963033 | 2.48E-09 | 2.95E-08 |
| CNAG_06790 | 3661.484 | 0.458326 | 0.154663 | 2.963385 | 0.003043 | 0.008779 |
| CNAG_07460 | 1302.189 | -0.34567 | 0.175527 | -1.96933 | 0.048915 | 0.095381 |
| CNAG_06787 | 64.40528 | 0.932621 | 0.27379 | 3.406335 | 0.000658 | 0.002306 |
| CNAG_06785 | 2379.803 | -0.71091 | 0.294074 | -2.41744 | 0.01563 | 0.036114 |
| CNAG_06782 | 2556.466 | 0.496698 | 0.17835 | 2.784958 | 0.005353 | 0.014415 |
| CNAG_12130 | 42.9361 | 0.733125 | 0.3647 | 2.010211 | 0.044409 | 0.087801 |
| CNAG_06779 | 2610.732 | -0.65222 | 0.237729 | -2.74353 | 0.006078 | 0.016089 |

| | | | | | | |
|---|---|---|---|---|---|---|
| CNAG_06778 | 2916.798 | 0.276298 | 0.115663 | 2.388819 | 0.016903 | 0.038561 |
| CNAG_06777 | 120.4397 | -1.17059 | 0.244312 | -4.79138 | 1.66E-06 | 1.10E-05 |
| CNAG_06775 | 2551.616 | -0.80441 | 0.191542 | -4.19965 | 2.67E-05 | 0.000134 |
| CNAG_06774 | 887.8696 | -1.01449 | 0.148645 | -6.8249 | 8.80E-12 | 1.56E-10 |
| CNAG_06771 | 853.3748 | 1.428093 | 0.213093 | 6.701744 | 2.06E-11 | 3.40E-10 |
| CNAG_06770 | 33719.17 | 0.340329 | 0.1418 | 2.400062 | 0.016392 | 0.03758 |
| CNAG_06767 | 12023.13 | -0.64169 | 0.293754 | -2.18444 | 0.02893 | 0.060943 |
| CNAG_06766 | 2630.628 | -0.57048 | 0.147271 | -3.87366 | 0.000107 | 0.000464 |
| CNAG_12131 | 15.04918 | -1.08116 | 0.544174 | -1.9868 | 0.046945 | 0.092173 |
| CNAG_07962 | 157.9886 | -1.34328 | 0.316961 | -4.23798 | 2.26E-05 | 0.000115 |
| CNAG_06763 | 896.0637 | -0.51381 | 0.156642 | -3.28015 | 0.001038 | 0.003425 |
| CNAG_06762 | 4569.677 | -1.43245 | 0.17625 | -8.1274 | 4.39E-16 | 1.38E-14 |
| CNAG_06760 | 906.7778 | -0.5504 | 0.119879 | -4.59127 | 4.41E-06 | 2.69E-05 |
| CNAG_12135 | 79.54064 | 1.241022 | 0.264849 | 4.68577 | 2.79E-06 | 1.78E-05 |
| CNAG_06759 | 1371.778 | 1.441176 | 0.184765 | 7.800034 | 6.19E-15 | 1.68E-13 |
| CNAG_06757 | 1039.381 | 0.479623 | 0.140635 | 3.410399 | 0.000649 | 0.002275 |
| CNAG_06756 | 812.8476 | -0.98188 | 0.254841 | -3.85289 | 0.000117 | 0.0005 |
| CNAG_06755 | 6350.936 | -0.79589 | 0.111649 | -7.12857 | 1.01E-12 | 2.10E-11 |
| CNAG_06754 | 5989.917 | -0.45398 | 0.105226 | -4.31432 | 1.60E-05 | 8.50E-05 |
| CNAG_06753 | 4739.921 | -0.47694 | 0.139706 | -3.41391 | 0.00064 | 0.00225 |
| CNAG_06752 | 559.7419 | -0.47455 | 0.187332 | -2.53318 | 0.011303 | 0.027308 |
| CNAG_06751 | 1980.53 | -0.3689 | 0.17084 | -2.15935 | 0.030823 | 0.0643 |
| CNAG_12139 | 35.07909 | -0.72802 | 0.343902 | -2.11695 | 0.034264 | 0.070428 |
| CNAG_06748 | 1965.999 | -0.42778 | 0.114728 | -3.72868 | 0.000192 | 0.000782 |
| CNAG_06747 | 22157.36 | -1.26896 | 0.197765 | -6.41651 | 1.39E-10 | 2.03E-09 |
| CNAG_06746 | 20052.37 | -0.86071 | 0.157617 | -5.46077 | 4.74E-08 | 4.38E-07 |
| CNAG_06745 | 16926.67 | -0.90702 | 0.169603 | -5.34787 | 8.90E-08 | 7.75E-07 |
| CNAG_06744 | 1131.1 | -0.62056 | 0.169213 | -3.66734 | 0.000245 | 0.000966 |
| CNAG_06742 | 2048.459 | -0.43268 | 0.125609 | -3.4447 | 0.000572 | 0.002041 |
| CNAG_06741 | 1923.738 | -0.73759 | 0.252947 | -2.916 | 0.003545 | 0.010029 |
| CNAG_06740 | 2169.923 | 0.538888 | 0.273538 | 1.970063 | 0.048831 | 0.095272 |
| CNAG_06737 | 2238.535 | 0.349369 | 0.173266 | 2.016377 | 0.043761 | 0.086822 |
| CNAG_06734 | 670.5169 | -1.12963 | 0.282877 | -3.99335 | 6.51E-05 | 0.000298 |
| CNAG_06733 | 2025.878 | 1.045084 | 0.175146 | 5.966918 | 2.42E-09 | 2.89E-08 |
| CNAG_06732 | 2793.155 | 0.704098 | 0.188728 | 3.730747 | 0.000191 | 0.000777 |
| CNAG_06729 | 645.0415 | -0.55886 | 0.17123 | -3.26379 | 0.001099 | 0.003596 |
| CNAG_06727 | 382.0397 | 1.092433 | 0.191245 | 5.712216 | 1.12E-08 | 1.16E-07 |
| CNAG_06726 | 520.3828 | 0.677607 | 0.196831 | 3.44258 | 0.000576 | 0.002053 |
| CNAG_06725 | 969.974 | -0.40852 | 0.139871 | -2.92067 | 0.003493 | 0.009905 |
| CNAG_06724 | 748.2513 | 0.668713 | 0.19419 | 3.443607 | 0.000574 | 0.002046 |
| CNAG_06723 | 3053.912 | -1.5817 | 0.199999 | -7.90855 | 2.60E-15 | 7.34E-14 |
| CNAG_06722 | 145.2193 | 1.342227 | 0.236464 | 5.676248 | 1.38E-08 | 1.41E-07 |
| CNAG_07465 | 635.6865 | -0.85724 | 0.15478 | -5.53848 | 3.05E-08 | 2.97E-07 |
| CNAG_06717 | 2075.769 | -0.56913 | 0.207966 | -2.73667 | 0.006207 | 0.016374 |
| CNAG_06716 | 2566.753 | -0.79593 | 0.22562 | -3.52773 | 0.000419 | 0.001548 |
| CNAG_06713 | 3338.57 | -0.73461 | 0.14301 | -5.13678 | 2.79E-07 | 2.16E-06 |
| CNAG_06711 | 1173.468 | -0.54238 | 0.227702 | -2.38196 | 0.017221 | 0.039191 |

| | | | | | | |
|---|---|---|---|---|---|---|
| CNAG_06710 | 2110.955 | -0.52014 | 0.245045 | -2.12264 | 0.033784 | 0.06951 |
| CNAG_06708 | 611.9189 | -0.50618 | 0.157197 | -3.22001 | 0.001282 | 0.004135 |
| CNAG_06707 | 110.815 | -0.4951 | 0.239901 | -2.06378 | 0.039038 | 0.078687 |
| CNAG_06706 | 280.3592 | 0.579372 | 0.17638 | 3.284786 | 0.001021 | 0.00338 |
| CNAG_10500 | 766813.5 | 2.191363 | 1.066599 | 2.054533 | 0.039924 | 0.080232 |
| CNAG_10501 | 16003.16 | 2.356122 | 1.069601 | 2.202804 | 0.027609 | 0.058563 |
| CNAG_10502 | 1053222 | 2.089125 | 1.056856 | 1.976735 | 0.048072 | 0.094014 |
| CNAG_10503 | 32.30304 | 1.8866 | 0.799828 | 2.358758 | 0.018336 | 0.041368 |
| CNAG_03595 | 3890.392 | 0.995966 | 0.2521 | 3.950678 | 7.79E-05 | 0.00035 |
| CNAG_12146 | 31.31839 | 1.184088 | 0.387957 | 3.052114 | 0.002272 | 0.006839 |
| CNAG_03597 | 174.5933 | 0.963831 | 0.191318 | 5.037859 | 4.71E-07 | 3.49E-06 |
| CNAG_03600 | 276.7009 | 1.318739 | 0.291087 | 4.53039 | 5.89E-06 | 3.48E-05 |
| CNAG_07964 | 1344.118 | -0.46184 | 0.197537 | -2.33798 | 0.019388 | 0.04341 |
| CNAG_03602 | 8768.433 | -1.15018 | 0.178528 | -6.4426 | 1.17E-10 | 1.73E-09 |
| CNAG_03603 | 1674.141 | -0.94273 | 0.138728 | -6.79554 | 1.08E-11 | 1.88E-10 |
| CNAG_03605 | 711.552 | -0.59243 | 0.268627 | -2.20541 | 0.027425 | 0.058237 |
| CNAG_03606 | 9286.377 | -0.39155 | 0.172431 | -2.27078 | 0.02316 | 0.050627 |
| CNAG_03607 | 378.0422 | 0.543255 | 0.187286 | 2.900673 | 0.003724 | 0.010462 |
| CNAG_03609 | 1015.614 | 0.739404 | 0.142932 | 5.173118 | 2.30E-07 | 1.81E-06 |
| CNAG_03612 | 7967.785 | -0.8168 | 0.237715 | -3.43603 | 0.00059 | 0.002099 |
| CNAG_03619 | 494.5676 | 0.630327 | 0.141867 | 4.443088 | 8.87E-06 | 5.00E-05 |
| CNAG_03621 | 15599.67 | 0.490691 | 0.193071 | 2.541497 | 0.011038 | 0.026773 |
| CNAG_03624 | 4634.178 | -0.95444 | 0.156996 | -6.07939 | 1.21E-09 | 1.51E-08 |
| CNAG_03627 | 16505.56 | 0.956087 | 0.188316 | 5.077046 | 3.83E-07 | 2.88E-06 |
| CNAG_03628 | 2530.206 | 0.510491 | 0.155065 | 3.292108 | 0.000994 | 0.003303 |
| CNAG_03629 | 7428 | -1.33232 | 0.139612 | -9.54306 | 1.39E-21 | 7.45E-20 |
| CNAG_03630 | 66.53463 | -0.60474 | 0.27872 | -2.16969 | 0.030031 | 0.062895 |
| CNAG_03631 | 1813.638 | 0.891433 | 0.198737 | 4.485486 | 7.27E-06 | 4.20E-05 |
| CNAG_03632 | 1198.482 | -0.47278 | 0.157938 | -2.99342 | 0.002759 | 0.008108 |
| CNAG_03634 | 1050.151 | -1.03716 | 0.158793 | -6.53152 | 6.51E-11 | 9.89E-10 |
| CNAG_03636 | 1254.098 | -0.79755 | 0.229453 | -3.47588 | 0.000509 | 0.001842 |
| CNAG_03637 | 2258.537 | 0.684388 | 0.168724 | 4.056253 | 4.99E-05 | 0.000234 |
| CNAG_03638 | 3157.53 | 0.54782 | 0.257801 | 2.124969 | 0.033589 | 0.069163 |
| CNAG_03639 | 417.1732 | 0.620836 | 0.276917 | 2.241958 | 0.024964 | 0.053743 |
| CNAG_07471 | 60.26821 | 1.053324 | 0.343348 | 3.067806 | 0.002156 | 0.006535 |
| CNAG_07472 | 761.1659 | -0.42611 | 0.165626 | -2.57272 | 0.01009 | 0.024799 |
| CNAG_03641 | 7056.314 | -0.59743 | 0.111804 | -5.34356 | 9.11E-08 | 7.91E-07 |
| CNAG_03642 | 259.3869 | 1.261896 | 0.399573 | 3.158112 | 0.001588 | 0.004983 |
| CNAG_03643 | 189.2327 | 0.603655 | 0.248578 | 2.428428 | 0.015164 | 0.035212 |
| CNAG_03644 | 5135.088 | -1.27648 | 0.176924 | -7.21487 | 5.40E-13 | 1.16E-11 |
| CNAG_03645 | 6874.52 | -0.40955 | 0.192445 | -2.12812 | 0.033327 | 0.068695 |
| CNAG_03646 | 1257.023 | -0.56735 | 0.125071 | -4.53621 | 5.73E-06 | 3.39E-05 |
| CNAG_03647 | 699.7721 | -0.6639 | 0.171075 | -3.88077 | 0.000104 | 0.000451 |
| CNAG_03648 | 4989.587 | 0.631354 | 0.162793 | 3.878264 | 0.000105 | 0.000455 |
| CNAG_03649 | 695.7367 | -1.42768 | 0.160105 | -8.91713 | 4.79E-19 | 1.99E-17 |
| CNAG_03651 | 1751.97 | -0.42893 | 0.158161 | -2.712 | 0.006688 | 0.017435 |
| CNAG_03654 | 1291.498 | 0.407957 | 0.164229 | 2.484072 | 0.012989 | 0.030782 |

| | | | | | | |
|---|---|---|---|---|---|---|
| CNAG_03655 | 2649.577 | -0.7322 | 0.302344 | -2.42174 | 0.015446 | 0.035773 |
| CNAG_03658 | 634.4197 | -0.49484 | 0.140381 | -3.52501 | 0.000423 | 0.001561 |
| CNAG_03661 | 618.8784 | -0.3602 | 0.176593 | -2.03972 | 0.041379 | 0.082799 |
| CNAG_03662 | 2139.38 | -0.55138 | 0.268501 | -2.05355 | 0.040019 | 0.080383 |
| CNAG_03663 | 1150.063 | -0.78663 | 0.164341 | -4.78659 | 1.70E-06 | 1.12E-05 |
| CNAG_03666 | 384.0848 | -1.1016 | 0.201792 | -5.4591 | 4.79E-08 | 4.41E-07 |
| CNAG_03667 | 4109.944 | 0.74326 | 0.231658 | 3.20843 | 0.001335 | 0.004278 |
| CNAG_03669 | 2372.469 | 1.686099 | 0.231051 | 7.297517 | 2.93E-13 | 6.45E-12 |
| CNAG_03671 | 291.5966 | -0.71306 | 0.201261 | -3.54298 | 0.000396 | 0.00147 |
| CNAG_03673 | 937.5964 | 0.32104 | 0.15663 | 2.049677 | 0.040396 | 0.081098 |
| CNAG_03675 | 3328.283 | -0.89656 | 0.121679 | -7.36825 | 1.73E-13 | 3.86E-12 |
| CNAG_03676 | 1084.534 | -0.36281 | 0.124578 | -2.91232 | 0.003588 | 0.010134 |
| CNAG_03679 | 678.3557 | 1.730977 | 0.265552 | 6.518405 | 7.11E-11 | 1.07E-09 |
| CNAG_03680 | 1166.737 | 1.022799 | 0.135957 | 7.522977 | 5.35E-14 | 1.30E-12 |
| CNAG_03683 | 1155.698 | -0.30024 | 0.120058 | -2.50075 | 0.012393 | 0.029563 |
| CNAG_03685 | 837.5046 | 1.124191 | 0.190493 | 5.901481 | 3.60E-09 | 4.11E-08 |
| CNAG_03688 | 4247.856 | 1.820354 | 0.227943 | 7.986007 | 1.39E-15 | 4.12E-14 |
| CNAG_03689 | 2771.664 | 1.28328 | 0.198299 | 6.471429 | 9.71E-11 | 1.44E-09 |
| CNAG_07473 | 5690.096 | -0.41344 | 0.111159 | -3.71933 | 0.0002 | 0.000807 |
| CNAG_03692 | 788.8265 | 1.306756 | 0.186194 | 7.01826 | 2.25E-12 | 4.41E-11 |
| CNAG_03693 | 1602.085 | -0.61788 | 0.207412 | -2.97899 | 0.002892 | 0.008424 |
| CNAG_03695 | 595.3856 | -0.55994 | 0.141921 | -3.94548 | 7.96E-05 | 0.000357 |
| CNAG_03697 | 423.8554 | 0.654175 | 0.216269 | 3.024827 | 0.002488 | 0.007412 |
| CNAG_03698 | 558.3351 | -0.30052 | 0.140062 | -2.14564 | 0.031901 | 0.066289 |
| CNAG_03702 | 832.161 | -0.47915 | 0.221455 | -2.16362 | 0.030493 | 0.063763 |
| CNAG_03705 | 4684.418 | 0.623166 | 0.205343 | 3.034749 | 0.002407 | 0.007191 |
| CNAG_03707 | 536.9054 | -0.43411 | 0.150094 | -2.89228 | 0.003825 | 0.010689 |
| CNAG_03708 | 1657.736 | -0.36722 | 0.133729 | -2.74601 | 0.006033 | 0.015978 |
| CNAG_03709 | 398.0028 | 0.630182 | 0.268583 | 2.346326 | 0.01896 | 0.042509 |
| CNAG_03712 | 322.3442 | -0.56124 | 0.172065 | -3.26179 | 0.001107 | 0.00362 |
| CNAG_03713 | 292.7512 | 1.643787 | 0.441313 | 3.72476 | 0.000196 | 0.000792 |
| CNAG_07475 | 27.43798 | 2.017515 | 0.483834 | 4.16985 | 3.05E-05 | 0.000151 |
| CNAG_03715 | 1069.591 | -1.11407 | 0.159668 | -6.9774 | 3.01E-12 | 5.66E-11 |
| CNAG_03716 | 1943.136 | -0.57726 | 0.219939 | -2.62465 | 0.008674 | 0.021742 |
| CNAG_03718 | 1405.213 | 0.620384 | 0.162646 | 3.814323 | 0.000137 | 0.000577 |
| CNAG_03719 | 1830.65 | 1.105627 | 0.330295 | 3.347391 | 0.000816 | 0.002772 |
| CNAG_03720 | 1067.888 | -0.28489 | 0.139453 | -2.04288 | 0.041065 | 0.082253 |
| CNAG_03721 | 3525.506 | 0.514569 | 0.13569 | 3.792243 | 0.000149 | 0.000622 |
| CNAG_03724 | 4890.066 | -0.66362 | 0.164844 | -4.02575 | 5.68E-05 | 0.000263 |
| CNAG_03725 | 7681.052 | -0.41246 | 0.15485 | -2.66365 | 0.00773 | 0.019667 |
| CNAG_12161 | 241.1053 | 1.396747 | 0.253606 | 5.507539 | 3.64E-08 | 3.48E-07 |
| CNAG_03727 | 2654.979 | 1.108018 | 0.271896 | 4.07516 | 4.60E-05 | 0.000217 |
| CNAG_03728 | 1154.579 | 0.909706 | 0.306784 | 2.9653 | 0.003024 | 0.008734 |
| CNAG_03729 | 499.4965 | 0.759487 | 0.159567 | 4.759664 | 1.94E-06 | 1.27E-05 |
| CNAG_03731 | 739.9512 | -0.48094 | 0.142761 | -3.36884 | 0.000755 | 0.00259 |
| CNAG_03732 | 856.3511 | 0.488374 | 0.185829 | 2.628085 | 0.008587 | 0.021558 |
| CNAG_03733 | 123.7435 | 1.2162 | 0.279743 | 4.347564 | 1.38E-05 | 7.47E-05 |

| | | | | | | |
|---|---|---|---|---|---|---|
| CNAG_03735 | 1713.6 | -0.71116 | 0.155989 | -4.55903 | 5.14E-06 | 3.09E-05 |
| CNAG_07476 | 253.6873 | 1.146517 | 0.160981 | 7.122075 | 1.06E-12 | 2.19E-11 |
| CNAG_03738 | 2139.802 | 0.562943 | 0.181159 | 3.107456 | 0.001887 | 0.005789 |
| CNAG_03739 | 34901.92 | -0.68834 | 0.151609 | -4.54023 | 5.62E-06 | 3.33E-05 |
| CNAG_03743 | 737.4167 | -0.61341 | 0.141527 | -4.33424 | 1.46E-05 | 7.86E-05 |
| CNAG_03746 | 982.2656 | 1.26251 | 0.132858 | 9.502723 | 2.04E-21 | 1.07E-19 |
| CNAG_03747 | 31935.27 | -1.00985 | 0.230711 | -4.37713 | 1.20E-05 | 6.61E-05 |
| CNAG_03748 | 4810.522 | -0.30244 | 0.146643 | -2.06244 | 0.039166 | 0.078828 |
| CNAG_07477 | 893.2449 | 0.77518 | 0.337345 | 2.297884 | 0.021568 | 0.047527 |
| CNAG_12163 | 9.046741 | 1.393889 | 0.665265 | 2.09524 | 0.03615 | 0.07388 |
| CNAG_03753 | 1101.647 | 0.503292 | 0.172211 | 2.92253 | 0.003472 | 0.00985 |
| CNAG_03754 | 7.993026 | 2.821973 | 0.877872 | 3.214561 | 0.001306 | 0.004206 |
| CNAG_07479 | 436.8169 | 1.389528 | 0.36555 | 3.801198 | 0.000144 | 0.000604 |
| CNAG_07480 | 49.01367 | 1.518689 | 0.315117 | 4.819449 | 1.44E-06 | 9.65E-06 |
| CNAG_03759 | 89.01567 | 3.996731 | 0.460552 | 8.678129 | 4.02E-18 | 1.53E-16 |
| CNAG_03762 | 3605.467 | -1.02338 | 0.144731 | -7.0709 | 1.54E-12 | 3.12E-11 |
| CNAG_03764 | 90.25136 | 1.025567 | 0.265372 | 3.864642 | 0.000111 | 0.000478 |
| CNAG_12165 | 7.460608 | 1.784258 | 0.735506 | 2.425892 | 0.015271 | 0.035418 |
| CNAG_03765 | 14656.8 | 1.090572 | 0.136561 | 7.985996 | 1.39E-15 | 4.12E-14 |
| CNAG_03766 | 1056.352 | -0.43371 | 0.169412 | -2.56008 | 0.010465 | 0.025617 |
| CNAG_12166 | 63.53157 | -1.03179 | 0.425909 | -2.42256 | 0.015412 | 0.035713 |
| CNAG_12168 | 68.34374 | -1.21299 | 0.407242 | -2.97855 | 0.002896 | 0.008433 |
| CNAG_03771 | 19092.9 | 1.359351 | 0.153196 | 8.873269 | 7.10E-19 | 2.91E-17 |
| CNAG_03772 | 17029.28 | -2.62859 | 0.245405 | -10.7113 | 9.01E-27 | 6.76E-25 |
| CNAG_03775 | 4994.291 | 0.679101 | 0.223269 | 3.041631 | 0.002353 | 0.00705 |
| CNAG_03779 | 1603.138 | -0.69438 | 0.21224 | -3.27167 | 0.001069 | 0.003509 |
| CNAG_03780 | 42384.41 | -1.29188 | 0.306797 | -4.21086 | 2.54E-05 | 0.000129 |
| CNAG_03781 | 1362.487 | -0.45531 | 0.204806 | -2.22312 | 0.026208 | 0.056011 |
| CNAG_03782 | 79.8312 | 4.202111 | 0.371887 | 11.29944 | 1.32E-29 | 1.22E-27 |
| CNAG_03783 | 75.39281 | 2.136087 | 0.288396 | 7.406779 | 1.29E-13 | 2.93E-12 |
| CNAG_03785 | 3479.562 | 0.763732 | 0.162475 | 4.700614 | 2.59E-06 | 1.67E-05 |
| CNAG_03786 | 841.9498 | -0.34827 | 0.143171 | -2.43251 | 0.014994 | 0.034858 |
| CNAG_03787 | 20601.09 | -0.69058 | 0.256488 | -2.69245 | 0.007093 | 0.018323 |
| CNAG_03789 | 2036.087 | -0.78416 | 0.183953 | -4.26282 | 2.02E-05 | 0.000104 |
| CNAG_03790 | 4322.542 | -0.33449 | 0.166375 | -2.01046 | 0.044382 | 0.087784 |
| CNAG_03791 | 2247.831 | 0.974111 | 0.267425 | 3.642561 | 0.00027 | 0.001056 |
| CNAG_03792 | 992.3894 | 0.416018 | 0.127787 | 3.255553 | 0.001132 | 0.003695 |
| CNAG_03793 | 2236.854 | -0.76297 | 0.263496 | -2.89558 | 0.003785 | 0.0106 |
| CNAG_03794 | 1910.242 | 1.658232 | 0.312224 | 5.311025 | 1.09E-07 | 9.28E-07 |
| CNAG_03806 | 1484.044 | 1.12614 | 0.138751 | 8.116239 | 4.81E-16 | 1.50E-14 |
| CNAG_03808 | 1395.833 | 1.084177 | 0.125233 | 8.657306 | 4.83E-18 | 1.80E-16 |
| CNAG_03810 | 2200.032 | -0.7666 | 0.261931 | -2.92671 | 0.003426 | 0.009739 |
| CNAG_03812 | 5484.087 | -0.82352 | 0.137813 | -5.97558 | 2.29E-09 | 2.75E-08 |
| CNAG_03813 | 2399.907 | 0.337534 | 0.16138 | 2.091549 | 0.036479 | 0.074457 |
| CNAG_03814 | 3528.402 | -0.68072 | 0.268282 | -2.53734 | 0.01117 | 0.027045 |
| CNAG_03816 | 4803.432 | 0.661098 | 0.243009 | 2.720462 | 0.006519 | 0.017062 |
| CNAG_03817 | 2014.74 | -0.79017 | 0.2339 | -3.37823 | 0.00073 | 0.002511 |

| | | | | | | |
|---|---|---|---|---|---|---|
| CNAG_03819 | 6827.004 | -1.43667 | 0.187943 | -7.64418 | 2.10E-14 | 5.33E-13 |
| CNAG_03823 | 1314.307 | 0.492094 | 0.219611 | 2.240747 | 0.025042 | 0.053868 |
| CNAG_03824 | 6880.333 | 1.441437 | 0.11979 | 12.03306 | 2.38E-33 | 2.82E-31 |
| CNAG_12173 | 68.99309 | 1.408278 | 0.293428 | 4.799404 | 1.59E-06 | 1.06E-05 |
| CNAG_03827 | 1098.971 | 0.435501 | 0.143852 | 3.027434 | 0.002466 | 0.007354 |
| CNAG_03828 | 398.3656 | 1.58126 | 0.177016 | 8.932852 | 4.15E-19 | 1.75E-17 |
| CNAG_03833 | 107.7961 | 1.001556 | 0.27585 | 3.630797 | 0.000283 | 0.001099 |
| CNAG_12175 | 216.1735 | 0.575993 | 0.223313 | 2.579303 | 0.0099 | 0.024407 |
| CNAG_03837 | 495.6285 | 0.78472 | 0.231513 | 3.389531 | 0.0007 | 0.002423 |
| CNAG_03838 | 92.67424 | 1.815616 | 0.276796 | 6.559412 | 5.40E-11 | 8.29E-10 |
| CNAG_03840 | 1899.866 | -0.45699 | 0.205018 | -2.22903 | 0.025812 | 0.055299 |
| CNAG_03841 | 3621.305 | 0.642855 | 0.296985 | 2.164603 | 0.030418 | 0.063639 |
| CNAG_03842 | 60.85488 | 1.071987 | 0.30238 | 3.545163 | 0.000392 | 0.001462 |
| CNAG_03844 | 1414.377 | 1.083268 | 0.157534 | 6.876409 | 6.14E-12 | 1.12E-10 |
| CNAG_03845 | 3966.599 | -0.53757 | 0.138051 | -3.894 | 9.86E-05 | 0.00043 |
| CNAG_03848 | 2521.138 | -1.54831 | 0.241875 | -6.40128 | 1.54E-10 | 2.23E-09 |
| CNAG_03849 | 2031.795 | 1.317981 | 0.325722 | 4.04634 | 5.20E-05 | 0.000243 |
| CNAG_03855 | 463.5988 | 0.592608 | 0.196516 | 3.015565 | 0.002565 | 0.007611 |
| CNAG_03856 | 15866.54 | 1.837633 | 0.114867 | 15.99787 | 1.32E-57 | 4.70E-55 |
| CNAG_03857 | 4896.543 | 1.24024 | 0.154753 | 8.014321 | 1.11E-15 | 3.32E-14 |
| CNAG_03858 | 3267.384 | 0.64322 | 0.154603 | 4.160455 | 3.18E-05 | 0.000157 |
| CNAG_03859 | 1388.971 | -0.77859 | 0.265201 | -2.93585 | 0.003326 | 0.009484 |
| CNAG_03861 | 9120.149 | -0.81563 | 0.175556 | -4.64596 | 3.38E-06 | 2.12E-05 |
| CNAG_03863 | 4094.821 | -0.35092 | 0.120201 | -2.91946 | 0.003506 | 0.009933 |
| CNAG_03864 | 2576.301 | 0.455671 | 0.132546 | 3.437839 | 0.000586 | 0.002086 |
| CNAG_03865 | 2358.253 | 0.62858 | 0.183406 | 3.427261 | 0.00061 | 0.002152 |
| CNAG_03868 | 1239.735 | -0.44839 | 0.154757 | -2.89737 | 0.003763 | 0.010555 |
| CNAG_03870 | 1471.175 | 0.286275 | 0.140401 | 2.038975 | 0.041453 | 0.082909 |
| CNAG_03871 | 1151.474 | -0.65307 | 0.203345 | -3.21164 | 0.00132 | 0.004242 |
| CNAG_03873 | 8124.169 | 1.317306 | 0.295812 | 4.453181 | 8.46E-06 | 4.80E-05 |
| CNAG_03875 | 1583.03 | 0.378947 | 0.135924 | 2.787935 | 0.005305 | 0.014298 |
| CNAG_03876 | 3205.819 | 0.277485 | 0.123623 | 2.244602 | 0.024794 | 0.053463 |
| CNAG_03877 | 2094.031 | 0.482011 | 0.119169 | 4.044764 | 5.24E-05 | 0.000244 |
| CNAG_03878 | 1322.652 | -0.68596 | 0.175366 | -3.91159 | 9.17E-05 | 0.000404 |
| CNAG_07482 | 1797.412 | 1.689076 | 0.251109 | 6.726465 | 1.74E-11 | 2.90E-10 |
| CNAG_03881 | 2966.649 | 2.061097 | 0.210735 | 9.780521 | 1.37E-22 | 8.16E-21 |
| CNAG_03885 | 2667.022 | -0.71193 | 0.219548 | -3.24272 | 0.001184 | 0.003852 |
| CNAG_03886 | 2022.77 | -0.27325 | 0.123425 | -2.21388 | 0.026837 | 0.057156 |
| CNAG_03890 | 1494.648 | -0.54723 | 0.120093 | -4.55668 | 5.20E-06 | 3.12E-05 |
| CNAG_03892 | 11415.44 | 0.441398 | 0.204713 | 2.156178 | 0.03107 | 0.064747 |
| CNAG_03893 | 527.2719 | -0.69777 | 0.197875 | -3.52632 | 0.000421 | 0.001555 |
| CNAG_03894 | 98.08811 | -0.97222 | 0.311114 | -3.12496 | 0.001778 | 0.005496 |
| CNAG_12183 | 36.86216 | 2.500714 | 0.534643 | 4.677352 | 2.91E-06 | 1.85E-05 |
| CNAG_03897 | 1334.117 | -0.42362 | 0.173876 | -2.43631 | 0.014838 | 0.034545 |
| CNAG_03898 | 1864.18 | -0.66013 | 0.134279 | -4.9161 | 8.83E-07 | 6.20E-06 |
| CNAG_03899 | 7793.163 | 0.913699 | 0.18652 | 4.898671 | 9.65E-07 | 6.71E-06 |
| CNAG_03901 | 718.8125 | 0.334439 | 0.151537 | 2.206972 | 0.027316 | 0.058051 |

| | | | | | | |
|---|---|---|---|---|---|---|
| CNAG_12185 | 29.19804 | 1.28475 | 0.508157 | 2.528255 | 0.011463 | 0.027635 |
| CNAG_03904 | 4459.868 | 0.292104 | 0.137278 | 2.127826 | 0.033351 | 0.068727 |
| CNAG_03906 | 537.1179 | 1.510137 | 0.170437 | 8.860364 | 7.98E-19 | 3.23E-17 |
| CNAG_03905 | 552.418 | -0.44489 | 0.226363 | -1.96537 | 0.049371 | 0.096176 |
| CNAG_03907 | 386.4624 | -0.69663 | 0.187663 | -3.71211 | 0.000206 | 0.000828 |
| CNAG_03911 | 123.8869 | 0.800524 | 0.28768 | 2.782693 | 0.005391 | 0.014501 |
| CNAG_03912 | 138.7883 | -0.42334 | 0.202486 | -2.09073 | 0.036552 | 0.074588 |
| CNAG_03913 | 596.3397 | 1.320132 | 0.18882 | 6.991476 | 2.72E-12 | 5.25E-11 |
| CNAG_03918 | 1702.366 | 0.633813 | 0.241737 | 2.621911 | 0.008744 | 0.021869 |
| CNAG_03919 | 842.8093 | -0.81405 | 0.272454 | -2.98783 | 0.00281 | 0.008226 |
| CNAG_03920 | 11070.11 | -0.43442 | 0.116267 | -3.73641 | 0.000187 | 0.000762 |
| CNAG_03921 | 532.0101 | -0.60571 | 0.219034 | -2.76536 | 0.005686 | 0.015187 |
| CNAG_03922 | 709.0913 | 3.435778 | 0.217055 | 15.82903 | 1.96E-56 | 6.24E-54 |
| CNAG_03925 | 391.4479 | -0.63954 | 0.160986 | -3.97264 | 7.11E-05 | 0.000322 |
| CNAG_03926 | 3070.962 | -0.66558 | 0.110759 | -6.00931 | 1.86E-09 | 2.28E-08 |
| CNAG_03927 | 637.4183 | -0.38112 | 0.187966 | -2.02758 | 0.042603 | 0.084822 |
| CNAG_03931 | 9311.152 | -0.45813 | 0.170594 | -2.68551 | 0.007242 | 0.018641 |
| CNAG_03935 | 4863.826 | 1.031015 | 0.115485 | 8.927671 | 4.35E-19 | 1.82E-17 |
| CNAG_03936 | 6471.408 | 0.942506 | 0.201696 | 4.672907 | 2.97E-06 | 1.88E-05 |
| CNAG_03937 | 1390.804 | 1.158204 | 0.274846 | 4.214015 | 2.51E-05 | 0.000127 |
| CNAG_03938 | 97.68533 | 1.595819 | 0.255858 | 6.237132 | 4.46E-10 | 5.92E-09 |
| CNAG_03939 | 2114.836 | -0.75812 | 0.130187 | -5.82331 | 5.77E-09 | 6.29E-08 |
| CNAG_03940 | 2626.931 | -0.75756 | 0.166749 | -4.54312 | 5.54E-06 | 3.30E-05 |
| CNAG_03941 | 6470.774 | 0.489465 | 0.234685 | 2.085623 | 0.037013 | 0.075353 |
| CNAG_03944 | 10904.01 | 0.276413 | 0.118947 | 2.323832 | 0.020134 | 0.044829 |
| CNAG_03946 | 1363.254 | 0.94345 | 0.181252 | 5.205185 | 1.94E-07 | 1.57E-06 |
| CNAG_07486 | 976.2676 | 0.728243 | 0.17486 | 4.16473 | 3.12E-05 | 0.000154 |
| CNAG_12191 | 16.18894 | 1.244076 | 0.57803 | 2.152269 | 0.031376 | 0.065283 |
| CNAG_03949 | 1070.894 | -0.9293 | 0.156536 | -5.9367 | 2.91E-09 | 3.38E-08 |
| CNAG_03951 | 1624.873 | -1.03976 | 0.193102 | -5.3845 | 7.26E-08 | 6.42E-07 |
| CNAG_03952 | 3918.141 | -0.73368 | 0.281076 | -2.61024 | 0.009048 | 0.02253 |
| CNAG_03957 | 2874.014 | -0.37891 | 0.118247 | -3.20442 | 0.001353 | 0.004326 |
| CNAG_03958 | 2466.05 | 1.346427 | 0.239752 | 5.615911 | 1.96E-08 | 1.95E-07 |
| CNAG_03959 | 5591.975 | 0.656647 | 0.193644 | 3.39101 | 0.000696 | 0.002413 |
| CNAG_03961 | 1634.495 | -0.67902 | 0.187525 | -3.62093 | 0.000294 | 0.001136 |
| CNAG_03964 | 965.2964 | -1.21273 | 0.172493 | -7.03062 | 2.06E-12 | 4.06E-11 |
| CNAG_03965 | 1645.909 | -0.63881 | 0.25325 | -2.52246 | 0.011654 | 0.028027 |
| CNAG_03966 | 973.6215 | -1.04975 | 0.234129 | -4.48364 | 7.34E-06 | 4.23E-05 |
| CNAG_03967 | 3245.457 | 0.315494 | 0.126334 | 2.4973 | 0.012514 | 0.029808 |
| CNAG_03968 | 3347.248 | -0.39701 | 0.117286 | -3.38497 | 0.000712 | 0.00246 |
| CNAG_07487 | 17131.04 | -0.53415 | 0.239171 | -2.23332 | 0.025528 | 0.054808 |
| CNAG_07488 | 3911.138 | -0.84779 | 0.209484 | -4.04703 | 5.19E-05 | 0.000242 |
| CNAG_03974 | 4014.632 | 2.56055 | 0.17877 | 14.32312 | 1.57E-46 | 3.67E-44 |
| CNAG_03976 | 1481.588 | -0.57837 | 0.127875 | -4.52293 | 6.10E-06 | 3.60E-05 |
| CNAG_03977 | 1093.747 | -1.12191 | 0.154433 | -7.26471 | 3.74E-13 | 8.18E-12 |
| CNAG_03978 | 1411.669 | 0.613354 | 0.140266 | 4.372805 | 1.23E-05 | 6.73E-05 |
| CNAG_03979 | 769.4953 | -0.43194 | 0.158474 | -2.72563 | 0.006418 | 0.016847 |

| | | | | | | |
|---|---|---|---|---|---|---|
| CNAG_03982 | 1534.046 | -0.33764 | 0.133789 | -2.52366 | 0.011614 | 0.02794 |
| CNAG_03984 | 926.8847 | -0.77827 | 0.258174 | -3.0145 | 0.002574 | 0.007632 |
| CNAG_03986 | 2503.965 | -0.73191 | 0.129976 | -5.63113 | 1.79E-08 | 1.80E-07 |
| CNAG_03987 | 5941.957 | 0.317733 | 0.13187 | 2.40944 | 0.015977 | 0.036783 |
| CNAG_03988 | 219.3306 | 0.877371 | 0.412714 | 2.125854 | 0.033515 | 0.069029 |
| CNAG_12196 | 42.77324 | 2.521474 | 0.368162 | 6.848809 | 7.45E-12 | 1.34E-10 |
| CNAG_03991 | 2108.092 | 0.825965 | 0.144814 | 5.703612 | 1.17E-08 | 1.22E-07 |
| CNAG_03992 | 860.5595 | 0.368535 | 0.176667 | 2.08604 | 0.036975 | 0.075296 |
| CNAG_03993 | 1563.657 | -1.21016 | 0.240926 | -5.02297 | 5.09E-07 | 3.75E-06 |
| CNAG_03995 | 911.5747 | 0.837327 | 0.338004 | 2.47727 | 0.013239 | 0.031246 |
| CNAG_03996 | 9445.861 | -0.97995 | 0.21101 | -4.6441 | 3.42E-06 | 2.14E-05 |
| CNAG_03997 | 57.21172 | -1.31065 | 0.397271 | -3.29914 | 0.00097 | 0.003228 |
| CNAG_03998 | 1170.607 | 0.423725 | 0.146386 | 2.894581 | 0.003797 | 0.010619 |
| CNAG_03999 | 643.1724 | 3.280671 | 0.205092 | 15.99607 | 1.36E-57 | 4.70E-55 |
| CNAG_04000 | 788.7899 | -0.53317 | 0.168302 | -3.16796 | 0.001535 | 0.004835 |
| CNAG_04003 | 667.692 | 0.83601 | 0.142224 | 5.878144 | 4.15E-09 | 4.66E-08 |
| CNAG_04004 | 79157.61 | -1.01089 | 0.21898 | -4.61638 | 3.90E-06 | 2.41E-05 |
| CNAG_04006 | 20.1764 | 1.788476 | 0.621864 | 2.875992 | 0.004028 | 0.011186 |
| CNAG_12197 | 99.18685 | -0.66247 | 0.293755 | -2.25518 | 0.024122 | 0.052313 |
| CNAG_04010 | 4326.807 | -0.24487 | 0.112323 | -2.18006 | 0.029253 | 0.06151 |
| CNAG_04011 | 21682.22 | -1.10706 | 0.314623 | -3.5187 | 0.000434 | 0.001591 |
| CNAG_04014 | 7233.727 | 0.865712 | 0.141539 | 6.11641 | 9.57E-10 | 1.22E-08 |
| CNAG_12198 | 64.69789 | 0.967346 | 0.263937 | 3.66507 | 0.000247 | 0.000975 |
| CNAG_12199 | 33.19671 | 1.39927 | 0.377605 | 3.705647 | 0.000211 | 0.000848 |
| CNAG_04017 | 2373.833 | 1.231888 | 0.186933 | 6.59001 | 4.40E-11 | 6.77E-10 |
| CNAG_07489 | 1320.746 | 1.429743 | 0.162848 | 8.779616 | 1.64E-18 | 6.48E-17 |
| CNAG_04021 | 24313.18 | -1.05771 | 0.216915 | -4.87614 | 1.08E-06 | 7.42E-06 |
| CNAG_04023 | 1790.279 | -0.77235 | 0.18913 | -4.0837 | 4.43E-05 | 0.000211 |
| CNAG_04024 | 197.6213 | -0.67807 | 0.176009 | -3.85249 | 0.000117 | 0.000501 |
| CNAG_04025 | 73.37504 | 1.293398 | 0.43212 | 2.993145 | 0.002761 | 0.008108 |
| CNAG_04030 | 697.307 | -0.7837 | 0.223465 | -3.50703 | 0.000453 | 0.001655 |
| CNAG_04032 | 8159.853 | 0.393009 | 0.128178 | 3.066121 | 0.002169 | 0.006567 |
| CNAG_04033 | 3261 | 0.366669 | 0.148015 | 2.477251 | 0.01324 | 0.031246 |
| CNAG_04036 | 243.8133 | 0.584818 | 0.169352 | 3.453265 | 0.000554 | 0.001984 |
| CNAG_04037 | 1661.991 | -0.7397 | 0.262623 | -2.8166 | 0.004854 | 0.013203 |
| CNAG_04039 | 3880.424 | 0.458696 | 0.182034 | 2.519839 | 0.011741 | 0.028188 |
| CNAG_04041 | 1026.488 | -0.31333 | 0.148555 | -2.10917 | 0.03493 | 0.071626 |
| CNAG_04043 | 570.7421 | 1.543255 | 0.215212 | 7.170871 | 7.45E-13 | 1.57E-11 |
| CNAG_04044 | 1316.292 | -0.70286 | 0.117814 | -5.96588 | 2.43E-09 | 2.90E-08 |
| CNAG_04045 | 1438.096 | -0.70568 | 0.160492 | -4.397 | 1.10E-05 | 6.06E-05 |
| CNAG_04048 | 2798.193 | -0.47942 | 0.120719 | -3.97141 | 7.14E-05 | 0.000323 |
| CNAG_04049 | 1391.898 | -0.43338 | 0.125828 | -3.44423 | 0.000573 | 0.002043 |
| CNAG_04050 | 5403.05 | -0.51192 | 0.248374 | -2.0611 | 0.039293 | 0.079044 |
| CNAG_04055 | 768.6993 | 0.555268 | 0.132413 | 4.193468 | 2.75E-05 | 0.000137 |
| CNAG_04056 | 1480.399 | -0.58227 | 0.187971 | -3.09767 | 0.00195 | 0.005963 |
| CNAG_07491 | 858.5335 | 0.939814 | 0.127154 | 7.391123 | 1.46E-13 | 3.27E-12 |
| CNAG_07492 | 126.5635 | 2.739471 | 0.411451 | 6.658073 | 2.77E-11 | 4.46E-10 |

| | | | | | | |
|---|---|---|---|---|---|---|
| CNAG_07493 | 500.5136 | 3.273028 | 0.213224 | 15.35022 | 3.53E-53 | 9.67E-51 |
| CNAG_04062 | 2071.828 | -0.50241 | 0.108434 | -4.63331 | 3.60E-06 | 2.24E-05 |
| CNAG_04064 | 6505.292 | 0.534943 | 0.206165 | 2.594728 | 0.009467 | 0.023457 |
| CNAG_04067 | 1060.961 | 1.298878 | 0.249813 | 5.199398 | 2.00E-07 | 1.61E-06 |
| CNAG_04068 | 14342.69 | -1.07501 | 0.261035 | -4.11828 | 3.82E-05 | 0.000185 |
| CNAG_04070 | 235.0909 | 1.261339 | 0.41474 | 3.041273 | 0.002356 | 0.007053 |
| CNAG_04071 | 4369.343 | 0.572949 | 0.212367 | 2.697922 | 0.006977 | 0.018071 |
| CNAG_04072 | 1693.007 | -0.83628 | 0.270849 | -3.08764 | 0.002018 | 0.006154 |
| CNAG_04073 | 1719.128 | -0.53295 | 0.177987 | -2.99432 | 0.002751 | 0.008092 |
| CNAG_12203 | 178.6501 | 0.965785 | 0.274427 | 3.519279 | 0.000433 | 0.001589 |
| CNAG_07494 | 2204.372 | -0.60874 | 0.176641 | -3.4462 | 0.000569 | 0.00203 |
| CNAG_04076 | 1481.663 | 1.470222 | 0.29482 | 4.986853 | 6.14E-07 | 4.46E-06 |
| CNAG_12204 | 17.59715 | -1.28435 | 0.467991 | -2.74439 | 0.006062 | 0.016052 |
| CNAG_04079 | 1597.165 | -0.49028 | 0.147442 | -3.32521 | 0.000884 | 0.002966 |
| CNAG_12206 | 47.07021 | -1.78492 | 0.480385 | -3.7156 | 0.000203 | 0.000819 |
| CNAG_04082 | 7185.914 | -0.70586 | 0.128634 | -5.48733 | 4.08E-08 | 3.82E-07 |
| CNAG_04083 | 1906.445 | 0.915583 | 0.236699 | 3.868126 | 0.00011 | 0.000472 |
| CNAG_12208 | 96.18157 | -1.42777 | 0.414203 | -3.44703 | 0.000567 | 0.002025 |
| CNAG_12209 | 180.6448 | 1.373764 | 0.201288 | 6.824854 | 8.80E-12 | 1.56E-10 |
| CNAG_04085 | 504.3608 | 2.577574 | 0.157995 | 16.31432 | 7.81E-60 | 3.10E-57 |
| CNAG_12210 | 582.1128 | 0.998912 | 0.190499 | 5.243666 | 1.57E-07 | 1.29E-06 |
| CNAG_04086 | 68.43611 | 1.052548 | 0.293945 | 3.580763 | 0.000343 | 0.001297 |
| CNAG_04090 | 4267.285 | 0.611787 | 0.21171 | 2.889743 | 0.003856 | 0.010761 |
| CNAG_04091 | 115.9764 | 0.886536 | 0.315424 | 2.810615 | 0.004945 | 0.013433 |
| CNAG_04092 | 63.31573 | 1.238282 | 0.275242 | 4.498882 | 6.83E-06 | 3.98E-05 |
| CNAG_04093 | 74.05258 | -0.94365 | 0.292783 | -3.22302 | 0.001268 | 0.004097 |
| CNAG_04095 | 338.455 | -0.69327 | 0.157772 | -4.39413 | 1.11E-05 | 6.14E-05 |
| CNAG_04096 | 186.6398 | -0.81975 | 0.223937 | -3.66061 | 0.000252 | 0.000989 |
| CNAG_04098 | 2432.848 | 2.688304 | 0.21962 | 12.24073 | 1.88E-34 | 2.34E-32 |
| CNAG_12212 | 496.5224 | -0.76664 | 0.185333 | -4.13656 | 3.53E-05 | 0.000172 |
| CNAG_07498 | 1123.102 | 1.23341 | 0.22854 | 5.396906 | 6.78E-08 | 6.07E-07 |
| CNAG_06973 | 112.8975 | 1.638118 | 0.27825 | 5.887225 | 3.93E-09 | 4.44E-08 |
| CNAG_07499 | 2181.588 | 0.272548 | 0.12966 | 2.10202 | 0.035552 | 0.072789 |
| CNAG_06963 | 5671.707 | -0.61885 | 0.155036 | -3.99169 | 6.56E-05 | 0.0003 |
| CNAG_06962 | 624.3542 | 1.538626 | 0.180623 | 8.518428 | 1.62E-17 | 5.74E-16 |
| CNAG_03083 | 349.7504 | 1.429386 | 0.215452 | 6.634367 | 3.26E-11 | 5.17E-10 |
| CNAG_03082 | 101.5333 | 1.042358 | 0.321739 | 3.239758 | 0.001196 | 0.003888 |
| CNAG_03081 | 2926.184 | -0.43818 | 0.105539 | -4.15186 | 3.30E-05 | 0.000162 |
| CNAG_03080 | 7403.984 | -0.56124 | 0.109074 | -5.14548 | 2.67E-07 | 2.06E-06 |
| CNAG_03078 | 3399.48 | 0.559177 | 0.142548 | 3.922733 | 8.76E-05 | 0.000389 |
| CNAG_03076 | 253.1973 | 1.078713 | 0.285632 | 3.776578 | 0.000159 | 0.000659 |
| CNAG_12217 | 652.5695 | 0.588814 | 0.229332 | 2.567523 | 0.010243 | 0.025136 |
| CNAG_03072 | 48691.77 | -0.23356 | 0.102832 | -2.27125 | 0.023132 | 0.05058 |
| CNAG_12218 | 143.8114 | 0.642828 | 0.258614 | 2.485667 | 0.012931 | 0.030662 |
| CNAG_03068 | 1199.001 | 1.413355 | 0.226582 | 6.237726 | 4.44E-10 | 5.91E-09 |
| CNAG_03067 | 406.3606 | -0.32683 | 0.157886 | -2.07004 | 0.038449 | 0.077739 |
| CNAG_07502 | 6302.66 | -0.71851 | 0.117261 | -6.12744 | 8.93E-10 | 1.14E-08 |

| | | | | | | |
|---|---|---|---|---|---|---|
| CNAG_03065 | 5639.071 | -0.72263 | 0.182698 | -3.95534 | 7.64E-05 | 0.000344 |
| CNAG_03064 | 1473.885 | -1.16134 | 0.213425 | -5.44143 | 5.29E-08 | 4.83E-07 |
| CNAG_03060 | 960.9781 | 1.339665 | 0.25745 | 5.203595 | 1.95E-07 | 1.57E-06 |
| CNAG_03058 | 24428.23 | 1.094 | 0.271527 | 4.029065 | 5.60E-05 | 0.00026 |
| CNAG_03057 | 177.6324 | -1.21496 | 0.39861 | -3.04799 | 0.002304 | 0.006926 |
| CNAG_12220 | 9.802576 | 1.323062 | 0.660114 | 2.004293 | 0.045039 | 0.088869 |
| CNAG_03055 | 1425.361 | -0.73279 | 0.231978 | -3.15889 | 0.001584 | 0.004974 |
| CNAG_03054 | 233.6019 | 1.398166 | 0.222188 | 6.292723 | 3.12E-10 | 4.30E-09 |
| CNAG_03053 | 26070.47 | -1.05087 | 0.268478 | -3.91417 | 9.07E-05 | 0.000401 |
| CNAG_03051 | 983.3837 | -0.33514 | 0.149976 | -2.23465 | 0.02544 | 0.054635 |
| CNAG_03049 | 3103.495 | -0.61181 | 0.259636 | -2.35643 | 0.018452 | 0.041593 |
| CNAG_03048 | 2097.945 | -0.63097 | 0.188744 | -3.34301 | 0.000829 | 0.002802 |
| CNAG_12222 | 49.90305 | -1.83476 | 0.643146 | -2.85279 | 0.004334 | 0.011961 |
| CNAG_03046 | 1568.243 | -2.33915 | 0.438184 | -5.33828 | 9.38E-08 | 8.11E-07 |
| CNAG_03044 | 364.4503 | -0.85156 | 0.276764 | -3.07686 | 0.002092 | 0.006364 |
| CNAG_03040 | 5387.601 | 1.708951 | 0.156711 | 10.90514 | 1.09E-27 | 8.57E-26 |
| CNAG_03039 | 1876.44 | 0.698014 | 0.166086 | 4.202741 | 2.64E-05 | 0.000133 |
| CNAG_03037 | 1122.099 | -0.95721 | 0.157698 | -6.06992 | 1.28E-09 | 1.59E-08 |
| CNAG_03036 | 1187.142 | 0.498125 | 0.150081 | 3.319041 | 0.000903 | 0.003022 |
| CNAG_03034 | 428.1564 | -0.43409 | 0.187304 | -2.31759 | 0.020472 | 0.045451 |
| CNAG_03033 | 586.7091 | -0.80763 | 0.198636 | -4.0659 | 4.78E-05 | 0.000225 |
| CNAG_03032 | 1629.896 | 0.350115 | 0.134881 | 2.595725 | 0.009439 | 0.023402 |
| CNAG_03031 | 668.8494 | 0.281958 | 0.143131 | 1.969924 | 0.048847 | 0.095272 |
| CNAG_07505 | 33.07265 | 1.311276 | 0.351663 | 3.728784 | 0.000192 | 0.000782 |
| CNAG_03027 | 3246.029 | -0.5992 | 0.112725 | -5.3156 | 1.06E-07 | 9.07E-07 |
| CNAG_03026 | 1091.544 | 0.439727 | 0.133199 | 3.301277 | 0.000962 | 0.003209 |
| CNAG_03024 | 2087.577 | 1.028507 | 0.300072 | 3.427527 | 0.000609 | 0.002151 |
| CNAG_03022 | 835.318 | -0.46511 | 0.156475 | -2.97244 | 0.002954 | 0.00858 |
| CNAG_03021 | 60.36728 | 1.503802 | 0.292661 | 5.138381 | 2.77E-07 | 2.14E-06 |
| CNAG_03018 | 1182.563 | -0.62045 | 0.218448 | -2.84027 | 0.004508 | 0.012385 |
| CNAG_03017 | 1645.709 | -0.50749 | 0.158613 | -3.19953 | 0.001377 | 0.004391 |
| CNAG_03015 | 20738.36 | -1.05859 | 0.254382 | -4.16141 | 3.16E-05 | 0.000156 |
| CNAG_03012 | 46842.82 | 1.281211 | 0.21621 | 5.925763 | 3.11E-09 | 3.61E-08 |
| CNAG_12228 | 23.42699 | 1.464692 | 0.444066 | 3.298362 | 0.000973 | 0.003235 |
| CNAG_12229 | 22.24302 | 1.605134 | 0.521584 | 3.07742 | 0.002088 | 0.006357 |
| CNAG_03011 | 887.7272 | 0.826896 | 0.187818 | 4.402646 | 1.07E-05 | 5.92E-05 |
| CNAG_03009 | 616.5646 | -0.75688 | 0.191181 | -3.95898 | 7.53E-05 | 0.000339 |
| CNAG_03008 | 731.7684 | -0.32081 | 0.147692 | -2.17214 | 0.029845 | 0.062573 |
| CNAG_03007 | 5558.311 | 2.819252 | 0.17687 | 15.93971 | 3.36E-57 | 1.11E-54 |
| CNAG_12230 | 116.0084 | 1.302608 | 0.304647 | 4.2758 | 1.90E-05 | 9.93E-05 |
| CNAG_03004 | 1729.117 | -0.3749 | 0.1782 | -2.10379 | 0.035397 | 0.072509 |
| CNAG_03002 | 1850.748 | 1.647615 | 0.353737 | 4.657741 | 3.20E-06 | 2.01E-05 |
| CNAG_03001 | 2767.415 | -0.68715 | 0.122125 | -5.62663 | 1.84E-08 | 1.84E-07 |
| CNAG_03000 | 40961.08 | -0.95177 | 0.257677 | -3.69365 | 0.000221 | 0.000883 |
| CNAG_02999 | 716.3345 | -0.51164 | 0.182489 | -2.8037 | 0.005052 | 0.013696 |
| CNAG_02996 | 716.2926 | 1.588712 | 0.167074 | 9.509031 | 1.92E-21 | 1.01E-19 |
| CNAG_02994 | 10030.43 | 0.289191 | 0.146456 | 1.974594 | 0.048314 | 0.094418 |

| | | | | | | |
|---|---|---|---|---|---|---|
| CNAG_02993 | 1611.704 | 0.433491 | 0.122171 | 3.548229 | 0.000388 | 0.001447 |
| CNAG_02990 | 5390.63 | 1.391225 | 0.213216 | 6.524953 | 6.80E-11 | 1.03E-09 |
| CNAG_02989 | 2440.342 | 0.901569 | 0.159148 | 5.664975 | 1.47E-08 | 1.50E-07 |
| CNAG_02987 | 3.438367 | 3.223536 | 1.431188 | 2.25235 | 0.0243 | 0.052627 |
| CNAG_02986 | 3251.2 | 0.786755 | 0.176313 | 4.462271 | 8.11E-06 | 4.62E-05 |
| CNAG_02984 | 61.01366 | -0.93326 | 0.323582 | -2.88416 | 0.003925 | 0.010923 |
| CNAG_02983 | 2043.821 | -0.68253 | 0.133114 | -5.1274 | 2.94E-07 | 2.26E-06 |
| CNAG_02982 | 1963.28 | -0.88615 | 0.150929 | -5.87127 | 4.32E-09 | 4.84E-08 |
| CNAG_02981 | 3952.885 | -0.30004 | 0.136019 | -2.20586 | 0.027393 | 0.058185 |
| CNAG_02980 | 722.1652 | 0.351431 | 0.152993 | 2.297043 | 0.021616 | 0.047619 |
| CNAG_02979 | 470.9472 | 0.880427 | 0.170397 | 5.166908 | 2.38E-07 | 1.87E-06 |
| CNAG_02977 | 3418.834 | 0.472933 | 0.216977 | 2.179651 | 0.029283 | 0.061557 |
| CNAG_02976 | 1534.599 | 0.316176 | 0.148554 | 2.128367 | 0.033307 | 0.06867 |
| CNAG_02973 | 992.6873 | -0.2965 | 0.142731 | -2.07736 | 0.037769 | 0.076695 |
| CNAG_02968 | 516.1453 | 0.725926 | 0.174891 | 4.150735 | 3.31E-05 | 0.000163 |
| CNAG_02967 | 2095.615 | 0.859091 | 0.185795 | 4.623874 | 3.77E-06 | 2.33E-05 |
| CNAG_02966 | 299.0179 | 1.660056 | 0.231466 | 7.171912 | 7.40E-13 | 1.57E-11 |
| CNAG_02965 | 659.9125 | 0.313553 | 0.122455 | 2.56056 | 0.01045 | 0.02559 |
| CNAG_02964 | 687.7168 | 0.543421 | 0.146757 | 3.702859 | 0.000213 | 0.000854 |
| CNAG_02963 | 628.3774 | -1.33469 | 0.191639 | -6.96461 | 3.29E-12 | 6.17E-11 |
| CNAG_02962 | 1861.789 | -0.56848 | 0.215968 | -2.63224 | 0.008482 | 0.021334 |
| CNAG_02961 | 2465.175 | -0.94426 | 0.296638 | -3.18321 | 0.001457 | 0.004607 |
| CNAG_02959 | 5528.795 | 0.586771 | 0.109736 | 5.3471 | 8.94E-08 | 7.78E-07 |
| CNAG_02958 | 3161.113 | 0.528239 | 0.163192 | 3.236919 | 0.001208 | 0.003922 |
| CNAG_07508 | 648.8403 | 0.602899 | 0.143801 | 4.192581 | 2.76E-05 | 0.000138 |
| CNAG_07510 | 4426.359 | -0.62523 | 0.120258 | -5.19911 | 2.00E-07 | 1.61E-06 |
| CNAG_02955 | 622.3308 | -0.49667 | 0.232747 | -2.13393 | 0.032848 | 0.067919 |
| CNAG_02953 | 2926.022 | 0.850823 | 0.250071 | 3.402328 | 0.000668 | 0.002331 |
| CNAG_02951 | 619.7492 | -0.60607 | 0.192938 | -3.14124 | 0.001682 | 0.00524 |
| CNAG_07511 | 4716.698 | -0.58644 | 0.141396 | -4.14751 | 3.36E-05 | 0.000165 |
| CNAG_07512 | 1080.285 | 1.715347 | 0.221566 | 7.741918 | 9.79E-15 | 2.59E-13 |
| CNAG_02944 | 539.2676 | 0.736906 | 0.253432 | 2.907705 | 0.003641 | 0.01027 |
| CNAG_02943 | 19752.92 | 1.218724 | 0.150014 | 8.124094 | 4.51E-16 | 1.42E-14 |
| CNAG_02942 | 2970.822 | 2.70153 | 0.201064 | 13.43618 | 3.71E-41 | 6.55E-39 |
| CNAG_02941 | 1727.092 | 0.681619 | 0.197027 | 3.459522 | 0.000541 | 0.001945 |
| CNAG_02940 | 1392.038 | 0.580565 | 0.248719 | 2.334216 | 0.019584 | 0.043787 |
| CNAG_02938 | 7136.613 | -0.66556 | 0.17783 | -3.74268 | 0.000182 | 0.000746 |
| CNAG_02935 | 2430.445 | -0.90926 | 0.227087 | -4.00403 | 6.23E-05 | 0.000285 |
| CNAG_02934 | 528.1121 | 1.461875 | 0.167717 | 8.716326 | 2.87E-18 | 1.10E-16 |
| CNAG_02931 | 461.4004 | -0.70109 | 0.177336 | -3.95344 | 7.70E-05 | 0.000346 |
| CNAG_02928 | 42011.42 | -0.8461 | 0.184622 | -4.58288 | 4.59E-06 | 2.78E-05 |
| CNAG_02925 | 8195.317 | 0.426778 | 0.150589 | 2.834062 | 0.004596 | 0.012584 |
| CNAG_02923 | 2637.019 | -0.68809 | 0.24181 | -2.84559 | 0.004433 | 0.012203 |
| CNAG_02922 | 2054.061 | -0.52363 | 0.172438 | -3.03664 | 0.002392 | 0.007154 |
| CNAG_02921 | 3337.009 | 0.277703 | 0.115682 | 2.400568 | 0.01637 | 0.037539 |
| CNAG_02920 | 1040.894 | -0.49414 | 0.167583 | -2.94865 | 0.003192 | 0.009139 |
| CNAG_02918 | 7935.077 | -1.04978 | 0.233671 | -4.49257 | 7.04E-06 | 4.08E-05 |

| | | | | | | |
|---|---|---|---|---|---|---|
| CNAG_02917 | 2514.823 | -0.75889 | 0.323363 | -2.34687 | 0.018932 | 0.042483 |
| CNAG_02914 | 958.0922 | -0.91014 | 0.190976 | -4.76574 | 1.88E-06 | 1.24E-05 |
| CNAG_02913 | 800.5761 | -0.36742 | 0.127842 | -2.87404 | 0.004053 | 0.011247 |
| CNAG_02910 | 857.84 | -0.44631 | 0.189264 | -2.35812 | 0.018368 | 0.041428 |
| CNAG_02907 | 893.4517 | -0.43091 | 0.139493 | -3.08914 | 0.002007 | 0.006128 |
| CNAG_02904 | 437.927 | 0.747806 | 0.174878 | 4.276155 | 1.90E-05 | 9.92E-05 |
| CNAG_02901 | 2394.629 | -0.46017 | 0.170341 | -2.70147 | 0.006903 | 0.017908 |
| CNAG_02900 | 224.1865 | 1.370075 | 0.248617 | 5.510779 | 3.57E-08 | 3.42E-07 |
| CNAG_12243 | 50.2755 | 1.931336 | 0.368142 | 5.246178 | 1.55E-07 | 1.28E-06 |
| CNAG_02899 | 46.36448 | 2.224405 | 0.420076 | 5.295242 | 1.19E-07 | 1.00E-06 |
| CNAG_02898 | 2844.45 | -0.47887 | 0.190501 | -2.51375 | 0.011945 | 0.028616 |
| CNAG_02896 | 4598.41 | -0.85324 | 0.170758 | -4.99682 | 5.83E-07 | 4.24E-06 |
| CNAG_02895 | 2976.394 | 1.080475 | 0.16688 | 6.474566 | 9.51E-11 | 1.42E-09 |
| CNAG_02893 | 337.2322 | -1.20313 | 0.173527 | -6.93341 | 4.11E-12 | 7.61E-11 |
| CNAG_02892 | 566.043 | 0.781442 | 0.14799 | 5.280364 | 1.29E-07 | 1.08E-06 |
| CNAG_02890 | 1525.567 | -0.48105 | 0.143834 | -3.3445 | 0.000824 | 0.002791 |
| CNAG_02888 | 328.351 | 0.630603 | 0.217794 | 2.89541 | 0.003787 | 0.010602 |
| CNAG_02886 | 4091.228 | 0.60445 | 0.204823 | 2.951089 | 0.003167 | 0.009074 |
| CNAG_02885 | 5783.469 | -0.56761 | 0.170246 | -3.33406 | 0.000856 | 0.002881 |
| CNAG_02883 | 1127.274 | 0.357835 | 0.182426 | 1.961536 | 0.049816 | 0.096925 |
| CNAG_12244 | 40.97456 | 1.248116 | 0.346149 | 3.605717 | 0.000311 | 0.001197 |
| CNAG_02880 | 18795.32 | -0.70256 | 0.199525 | -3.52117 | 0.00043 | 0.001581 |
| CNAG_02879 | 1880.945 | -0.6077 | 0.269045 | -2.25872 | 0.023901 | 0.051944 |
| CNAG_02878 | 1472.194 | -0.44967 | 0.133028 | -3.38022 | 0.000724 | 0.002498 |
| CNAG_02877 | 4433.044 | 2.614918 | 0.272725 | 9.588115 | 8.97E-22 | 4.92E-20 |
| CNAG_02874 | 1336.503 | 0.464528 | 0.186362 | 2.49261 | 0.012681 | 0.030159 |
| CNAG_02871 | 611.3273 | 0.295727 | 0.134366 | 2.200904 | 0.027743 | 0.058817 |
| CNAG_02870 | 1946.692 | -0.94142 | 0.170278 | -5.52872 | 3.23E-08 | 3.13E-07 |
| CNAG_02866 | 1419.547 | -0.49661 | 0.150879 | -3.29147 | 0.000997 | 0.003308 |
| CNAG_02865 | 127.3472 | 0.811173 | 0.253537 | 3.199424 | 0.001377 | 0.004391 |
| CNAG_02864 | 346.432 | 2.635024 | 0.203204 | 12.96737 | 1.87E-38 | 3.04E-36 |
| CNAG_12250 | 1216.835 | -0.90295 | 0.210489 | -4.28977 | 1.79E-05 | 9.40E-05 |
| CNAG_02862 | 409.317 | 0.806033 | 0.152325 | 5.291527 | 1.21E-07 | 1.02E-06 |
| CNAG_02860 | 9916.991 | 0.425236 | 0.144993 | 2.932812 | 0.003359 | 0.009574 |
| CNAG_02859 | 727.1346 | 0.583329 | 0.142536 | 4.092493 | 4.27E-05 | 0.000204 |
| CNAG_02858 | 19972.4 | -0.83452 | 0.15005 | -5.56159 | 2.67E-08 | 2.63E-07 |
| CNAG_02857 | 3081.084 | 0.76243 | 0.231019 | 3.300293 | 0.000966 | 0.003218 |
| CNAG_07515 | 200.7903 | -2.16919 | 0.421309 | -5.1487 | 2.62E-07 | 2.04E-06 |
| CNAG_02854 | 1837.09 | -0.65423 | 0.141357 | -4.62819 | 3.69E-06 | 2.29E-05 |
| CNAG_02853 | 5309.001 | -0.57184 | 0.117423 | -4.86991 | 1.12E-06 | 7.63E-06 |
| CNAG_02851 | 1241.125 | -0.39245 | 0.144529 | -2.71538 | 0.00662 | 0.017281 |
| CNAG_02850 | 9688.018 | -0.68942 | 0.221123 | -3.11779 | 0.001822 | 0.005605 |
| CNAG_02849 | 37.74201 | -3.58482 | 0.664649 | -5.39355 | 6.91E-08 | 6.14E-07 |
| CNAG_02848 | 333.5999 | 0.672061 | 0.182574 | 3.681031 | 0.000232 | 0.000922 |
| CNAG_02847 | 508.7487 | -0.43438 | 0.157453 | -2.75881 | 0.005801 | 0.015448 |
| CNAG_12259 | 105.873 | 0.810698 | 0.290456 | 2.791125 | 0.005253 | 0.014177 |
| CNAG_02843 | 9417.613 | 0.77698 | 0.130501 | 5.953833 | 2.62E-09 | 3.08E-08 |

| | | | | | | |
|---|---|---|---|---|---|---|
| CNAG_02842 | 2762.473 | -0.79329 | 0.202097 | -3.92527 | 8.66E-05 | 0.000385 |
| CNAG_02841 | 1689.185 | 0.540147 | 0.149871 | 3.604072 | 0.000313 | 0.001202 |
| CNAG_02840 | 1524.234 | 1.781204 | 0.19603 | 9.086384 | 1.02E-19 | 4.57E-18 |
| CNAG_02839 | 1292.676 | 4.07386 | 0.307047 | 13.26786 | 3.56E-40 | 6.01E-38 |
| CNAG_07517 | 1057.664 | -0.41396 | 0.161801 | -2.55844 | 0.010514 | 0.025714 |
| CNAG_07518 | 675.2958 | -0.43833 | 0.188141 | -2.32979 | 0.019817 | 0.044207 |
| CNAG_07519 | 3089.786 | 0.270474 | 0.126301 | 2.141509 | 0.032233 | 0.066891 |
| CNAG_12261 | 33.39499 | 1.087109 | 0.361948 | 3.003498 | 0.002669 | 0.007875 |
| CNAG_02834 | 1522.529 | 0.981315 | 0.181776 | 5.398474 | 6.72E-08 | 6.02E-07 |
| CNAG_02833 | 2360.422 | -1.05418 | 0.236997 | -4.44807 | 8.66E-06 | 4.90E-05 |
| CNAG_07520 | 150.6333 | 0.725215 | 0.212335 | 3.415427 | 0.000637 | 0.002241 |
| CNAG_02830 | 4577.475 | -0.68442 | 0.152148 | -4.49837 | 6.85E-06 | 3.98E-05 |
| CNAG_02828 | 144.5676 | 1.503434 | 0.327641 | 4.588659 | 4.46E-06 | 2.72E-05 |
| CNAG_02826 | 1069.439 | -0.84443 | 0.158845 | -5.31605 | 1.06E-07 | 9.06E-07 |
| CNAG_02825 | 5241.294 | -1.0474 | 0.103076 | -10.1614 | 2.95E-24 | 1.98E-22 |
| CNAG_07521 | 342.4462 | 0.803617 | 0.201385 | 3.990445 | 6.59E-05 | 0.000301 |
| CNAG_02824 | 1227.826 | -0.32949 | 0.140819 | -2.33981 | 0.019293 | 0.043209 |
| CNAG_12262 | 26.67334 | 1.102478 | 0.433404 | 2.543764 | 0.010967 | 0.026632 |
| CNAG_02822 | 4159.255 | 0.622311 | 0.286689 | 2.17068 | 0.029955 | 0.062769 |
| CNAG_02821 | 319.5758 | -0.93665 | 0.243079 | -3.85326 | 0.000117 | 0.0005 |
| CNAG_02819 | 861.5538 | -0.248 | 0.119224 | -2.08009 | 0.037517 | 0.076282 |
| CNAG_02818 | 2319.438 | -0.48239 | 0.157697 | -3.05897 | 0.002221 | 0.00671 |
| CNAG_02815 | 12755.86 | 2.236638 | 0.159737 | 14.00199 | 1.52E-44 | 3.17E-42 |
| CNAG_02814 | 11553.09 | 0.244998 | 0.112585 | 2.176111 | 0.029547 | 0.062029 |
| CNAG_02813 | 1433.249 | -0.69106 | 0.190788 | -3.62215 | 0.000292 | 0.001132 |
| CNAG_02812 | 4538.788 | -1.0625 | 0.179965 | -5.90389 | 3.55E-09 | 4.06E-08 |
| CNAG_02811 | 10808.63 | -1.09959 | 0.316346 | -3.47591 | 0.000509 | 0.001842 |
| CNAG_02810 | 6882.128 | -0.4664 | 0.186803 | -2.49676 | 0.012533 | 0.029844 |
| CNAG_02809 | 1596.105 | -0.50212 | 0.151138 | -3.32222 | 0.000893 | 0.002995 |
| CNAG_07522 | 2645.683 | 1.038473 | 0.165415 | 6.277982 | 3.43E-10 | 4.67E-09 |
| CNAG_02806 | 3301.093 | 0.353 | 0.128726 | 2.742266 | 0.006102 | 0.016124 |
| CNAG_02802 | 144.9544 | -1.01378 | 0.199521 | -5.08107 | 3.75E-07 | 2.83E-06 |
| CNAG_12267 | 27.31584 | 0.756386 | 0.371787 | 2.034461 | 0.041905 | 0.083579 |
| CNAG_02799 | 1076.301 | -0.59826 | 0.15751 | -3.79823 | 0.000146 | 0.000611 |
| CNAG_02798 | 1204.724 | -0.48604 | 0.216274 | -2.24733 | 0.024619 | 0.053201 |
| CNAG_02796 | 2107.179 | -0.44324 | 0.120661 | -3.67341 | 0.000239 | 0.000947 |
| CNAG_02794 | 3515.355 | -0.4333 | 0.160452 | -2.7005 | 0.006923 | 0.017953 |
| CNAG_02793 | 1854.851 | -0.38782 | 0.117524 | -3.29994 | 0.000967 | 0.003221 |
| CNAG_02791 | 512.6711 | -0.40682 | 0.173604 | -2.34338 | 0.01911 | 0.042834 |
| CNAG_12268 | 28.50297 | -3.23055 | 0.566705 | -5.70059 | 1.19E-08 | 1.24E-07 |
| CNAG_02790 | 2303.748 | -0.53802 | 0.143023 | -3.76181 | 0.000169 | 0.000697 |
| CNAG_02788 | 1134.89 | -0.89751 | 0.190271 | -4.71699 | 2.39E-06 | 1.55E-05 |
| CNAG_02785 | 1044.882 | -0.45412 | 0.228942 | -1.98356 | 0.047305 | 0.092743 |
| CNAG_02782 | 1298.479 | -0.58767 | 0.139999 | -4.19768 | 2.70E-05 | 0.000135 |
| CNAG_02780 | 2659.691 | -0.39487 | 0.111469 | -3.54243 | 0.000396 | 0.001473 |
| CNAG_02777 | 3551.266 | 0.835662 | 0.248225 | 3.36655 | 0.000761 | 0.002608 |
| CNAG_07523 | 332.6356 | -0.47187 | 0.221668 | -2.12872 | 0.033278 | 0.068628 |

| | | | | | | |
|---|---|---|---|---|---|---|
| CNAG_02775 | 1518.525 | -0.44517 | 0.18929 | -2.35177 | 0.018684 | 0.041999 |
| CNAG_02771 | 1591.653 | 0.877328 | 0.256552 | 3.419693 | 0.000627 | 0.002209 |
| CNAG_02769 | 1737.841 | -0.35265 | 0.12955 | -2.72208 | 0.006487 | 0.01699 |
| CNAG_02768 | 366.2413 | 2.273503 | 0.217399 | 10.45776 | 1.35E-25 | 9.49E-24 |
| CNAG_02767 | 594.1038 | -0.41197 | 0.153215 | -2.68887 | 0.00717 | 0.018496 |
| CNAG_02766 | 648.5144 | -0.65568 | 0.138541 | -4.73275 | 2.22E-06 | 1.44E-05 |
| CNAG_02764 | 344.7261 | 0.366219 | 0.152161 | 2.40678 | 0.016094 | 0.037014 |
| CNAG_02763 | 4481.056 | -0.60795 | 0.169591 | -3.58479 | 0.000337 | 0.001281 |
| CNAG_02762 | 11855.41 | -0.78576 | 0.264886 | -2.96642 | 0.003013 | 0.008715 |
| CNAG_12273 | 9.274607 | 1.583934 | 0.778835 | 2.033722 | 0.04198 | 0.083673 |
| CNAG_02757 | 948.4909 | 0.427719 | 0.148147 | 2.887133 | 0.003888 | 0.010839 |
| CNAG_02755 | 1281.802 | -0.48042 | 0.161705 | -2.97095 | 0.002969 | 0.008616 |
| CNAG_02754 | 48150.09 | -1.1077 | 0.327737 | -3.37985 | 0.000725 | 0.002498 |
| CNAG_02752 | 3543.645 | -0.69332 | 0.209267 | -3.31307 | 0.000923 | 0.003083 |
| CNAG_02751 | 4979.727 | 2.555496 | 0.159702 | 16.00165 | 1.24E-57 | 4.70E-55 |
| CNAG_02747 | 764.1172 | 0.258639 | 0.124954 | 2.069879 | 0.038464 | 0.077749 |
| CNAG_02741 | 912.987 | -0.34079 | 0.138727 | -2.45658 | 0.014027 | 0.032907 |
| CNAG_02740 | 427.8542 | -0.73053 | 0.160225 | -4.55942 | 5.13E-06 | 3.09E-05 |
| CNAG_02739 | 1529.54 | -0.6611 | 0.136056 | -4.85904 | 1.18E-06 | 8.02E-06 |
| CNAG_02735 | 384.2393 | 2.919442 | 0.208893 | 13.97577 | 2.19E-44 | 4.46E-42 |
| CNAG_02733 | 54.08943 | -0.72708 | 0.332048 | -2.18969 | 0.028547 | 0.060232 |
| CNAG_02725 | 5112.248 | 0.809219 | 0.145344 | 5.567628 | 2.58E-08 | 2.55E-07 |
| CNAG_02724 | 628.5831 | -1.07954 | 0.26061 | -4.14235 | 3.44E-05 | 0.000168 |
| CNAG_12276 | 2.761079 | -2.93572 | 1.440756 | -2.03763 | 0.041587 | 0.0831 |
| CNAG_12277 | 16.56335 | 1.24459 | 0.606068 | 2.053548 | 0.040019 | 0.080383 |
| CNAG_02721 | 1541.053 | -0.78352 | 0.282759 | -2.77099 | 0.005589 | 0.014987 |
| CNAG_02720 | 2162.974 | -0.67761 | 0.151267 | -4.47959 | 7.48E-06 | 4.31E-05 |
| CNAG_02719 | 332.2698 | 0.637503 | 0.202777 | 3.143862 | 0.001667 | 0.005204 |
| CNAG_07969 | 123.5481 | 2.79704 | 0.374424 | 7.470242 | 8.00E-14 | 1.89E-12 |
| CNAG_02714 | 29665.88 | -0.87941 | 0.281812 | -3.12057 | 0.001805 | 0.005564 |
| CNAG_02712 | 2058.773 | -0.49856 | 0.149341 | -3.33844 | 0.000843 | 0.002846 |
| CNAG_02709 | 659.4875 | -0.84025 | 0.239721 | -3.50513 | 0.000456 | 0.001665 |
| CNAG_02708 | 923.5684 | 0.609415 | 0.125944 | 4.838772 | 1.31E-06 | 8.81E-06 |
| CNAG_02705 | 121.1736 | -0.53623 | 0.221928 | -2.41622 | 0.015683 | 0.036225 |
| CNAG_07525 | 7.771584 | 1.87882 | 0.776399 | 2.419917 | 0.015524 | 0.035901 |
| CNAG_02704 | 2086.057 | -0.55497 | 0.210312 | -2.63877 | 0.008321 | 0.02097 |
| CNAG_02702 | 2701.431 | -0.43563 | 0.214209 | -2.03368 | 0.041984 | 0.083673 |
| CNAG_02701 | 9068.686 | 4.480936 | 0.209123 | 21.42732 | 7.43E-102 | 9.85E-99 |
| CNAG_12282 | 52.99754 | 3.067565 | 0.372864 | 8.227036 | 1.92E-16 | 6.20E-15 |
| CNAG_02700 | 2650.068 | 0.383685 | 0.192998 | 1.988026 | 0.046809 | 0.091951 |
| CNAG_02698 | 1492.459 | -0.40285 | 0.200307 | -2.01116 | 0.044308 | 0.087736 |
| CNAG_12285 | 370.7742 | 1.13832 | 0.318649 | 3.572331 | 0.000354 | 0.001336 |
| CNAG_02692 | 9327.339 | 2.858336 | 0.243902 | 11.7192 | 1.02E-31 | 1.02E-29 |
| CNAG_02691 | 1096.495 | 1.402451 | 0.28684 | 4.889308 | 1.01E-06 | 6.98E-06 |
| CNAG_02690 | 872.3762 | 0.864289 | 0.131136 | 6.59079 | 4.37E-11 | 6.77E-10 |
| CNAG_07527 | 55.88064 | 1.620577 | 0.326313 | 4.966322 | 6.82E-07 | 4.91E-06 |
| CNAG_02687 | 1943.468 | -1.0462 | 0.13811 | -7.57507 | 3.59E-14 | 8.86E-13 |

| | | | | | | |
|---|---|---|---|---|---|---|
| CNAG_02686 | 5691.914 | -0.45019 | 0.129808 | -3.46815 | 0.000524 | 0.001889 |
| CNAG_02685 | 2080.854 | 3.543326 | 0.160846 | 22.02925 | 1.51E-107 | 2.40E-104 |
| CNAG_12289 | 37.21106 | 3.870756 | 0.501398 | 7.719924 | 1.16E-14 | 3.03E-13 |
| CNAG_02684 | 2136.499 | 1.521627 | 0.312113 | 4.875241 | 1.09E-06 | 7.44E-06 |
| CNAG_02683 | 1731.62 | 0.352129 | 0.114605 | 3.072539 | 0.002122 | 0.006444 |
| CNAG_02682 | 1785.189 | 0.579136 | 0.280134 | 2.067353 | 0.038701 | 0.078129 |
| CNAG_02681 | 562.8935 | 0.816917 | 0.14899 | 5.483031 | 4.18E-08 | 3.89E-07 |
| CNAG_02679 | 749.3242 | 0.452503 | 0.144956 | 3.121662 | 0.001798 | 0.005549 |
| CNAG_02678 | 758.2865 | 0.439977 | 0.197145 | 2.231742 | 0.025632 | 0.055002 |
| CNAG_02677 | 3714.976 | -0.40358 | 0.156998 | -2.57061 | 0.010152 | 0.024928 |
| CNAG_02675 | 2912.36 | -0.60838 | 0.300852 | -2.02219 | 0.043157 | 0.085817 |
| CNAG_02674 | 949.4581 | 0.431774 | 0.172581 | 2.501858 | 0.012354 | 0.02948 |
| CNAG_02673 | 8201.156 | 1.195332 | 0.155424 | 7.690778 | 1.46E-14 | 3.75E-13 |
| CNAG_02672 | 3094.45 | -0.63072 | 0.153654 | -4.1048 | 4.05E-05 | 0.000195 |
| CNAG_02670 | 1164.37 | -0.56482 | 0.123734 | -4.56483 | 5.00E-06 | 3.01E-05 |
| CNAG_12290 | 32.13826 | 0.988374 | 0.372521 | 2.653205 | 0.007973 | 0.020202 |
| CNAG_02667 | 636.5693 | 0.637368 | 0.155875 | 4.088966 | 4.33E-05 | 0.000207 |
| CNAG_02666 | 770.0752 | 0.411991 | 0.133896 | 3.076943 | 0.002091 | 0.006364 |
| CNAG_02664 | 3768.098 | -0.70881 | 0.106574 | -6.65087 | 2.91E-11 | 4.66E-10 |
| CNAG_02663 | 1571.456 | 1.208424 | 0.138087 | 8.751152 | 2.11E-18 | 8.23E-17 |
| CNAG_12292 | 31.4581 | 1.049372 | 0.365481 | 2.871207 | 0.004089 | 0.011341 |
| CNAG_02662 | 1506.387 | 0.487317 | 0.162242 | 3.003647 | 0.002668 | 0.007874 |
| CNAG_02661 | 407.7249 | 0.487055 | 0.178806 | 2.723925 | 0.006451 | 0.016918 |
| CNAG_12293 | 180.8233 | 0.629877 | 0.179503 | 3.509007 | 0.00045 | 0.001645 |
| CNAG_12294 | 226.1881 | 0.688996 | 0.175152 | 3.933699 | 8.36E-05 | 0.000374 |
| CNAG_12295 | 79.28726 | 0.987802 | 0.378063 | 2.612796 | 0.00898 | 0.022377 |
| CNAG_12298 | 278.0393 | 0.965142 | 0.251343 | 3.839939 | 0.000123 | 0.000524 |
| CNAG_02657 | 7491.174 | -0.71535 | 0.196471 | -3.64096 | 0.000272 | 0.001061 |
| CNAG_02656 | 2852.936 | -0.44553 | 0.192216 | -2.31786 | 0.020457 | 0.045431 |
| CNAG_02655 | 3118.244 | 0.807182 | 0.154089 | 5.238404 | 1.62E-07 | 1.33E-06 |
| CNAG_07529 | 1787.853 | -0.62014 | 0.182926 | -3.39008 | 0.000699 | 0.002419 |
| CNAG_07531 | 966.7218 | -1.08115 | 0.235768 | -4.58566 | 4.53E-06 | 2.75E-05 |
| CNAG_07532 | 1019.214 | -0.47467 | 0.17799 | -2.66684 | 0.007657 | 0.0195 |
| CNAG_07533 | 138.9516 | 0.728428 | 0.350347 | 2.07916 | 0.037603 | 0.076411 |
| CNAG_07535 | 900.8105 | -0.89652 | 0.22188 | -4.04056 | 5.33E-05 | 0.000248 |
| CNAG_07536 | 708.9059 | -0.57991 | 0.158997 | -3.64731 | 0.000265 | 0.001037 |
| CNAG_07537 | 535.0589 | -0.56105 | 0.159396 | -3.51984 | 0.000432 | 0.001586 |
| CNAG_07538 | 3589.722 | 3.017498 | 0.14365 | 21.00593 | 5.79E-98 | 5.75E-95 |
| CNAG_07162 | 408.0014 | -0.63573 | 0.174183 | -3.64978 | 0.000262 | 0.001028 |
| CNAG_07540 | 672.3561 | -0.73433 | 0.242275 | -3.03099 | 0.002437 | 0.00727 |
| CNAG_07541 | 2944.711 | 1.230695 | 0.174937 | 7.035067 | 1.99E-12 | 3.96E-11 |
| CNAG_07542 | 247.8117 | 0.91386 | 0.169637 | 5.387137 | 7.16E-08 | 6.34E-07 |
| CNAG_07544 | 38.3238 | 2.096287 | 0.444319 | 4.717975 | 2.38E-06 | 1.54E-05 |
| CNAG_07545 | 791.0112 | -0.64793 | 0.234192 | -2.76665 | 0.005664 | 0.015142 |
| CNAG_07546 | 1151.195 | -0.39619 | 0.145483 | -2.72328 | 0.006464 | 0.016945 |
| CNAG_07547 | 1796.979 | -0.57821 | 0.172165 | -3.35844 | 0.000784 | 0.002675 |
| CNAG_07548 | 906.9124 | 0.50091 | 0.126517 | 3.959234 | 7.52E-05 | 0.000339 |

| | | | | | | |
|---|---|---|---|---|---|---|
| CNAG_07549 | 144.7439 | -0.99605 | 0.244071 | -4.08098 | 4.48E-05 | 0.000213 |
| CNAG_12301 | 7.134677 | -2.39053 | 0.890897 | -2.68329 | 0.00729 | 0.018735 |
| CNAG_07551 | 3282.074 | 1.381997 | 0.279963 | 4.936361 | 7.96E-07 | 5.63E-06 |
| CNAG_07552 | 1754.346 | 1.328016 | 0.326471 | 4.067789 | 4.75E-05 | 0.000223 |
| CNAG_07554 | 6254.372 | -0.88964 | 0.114386 | -7.77755 | 7.39E-15 | 1.99E-13 |
| CNAG_07555 | 503.2376 | 0.854255 | 0.177589 | 4.810296 | 1.51E-06 | 1.01E-05 |
| CNAG_07558 | 22052.64 | 0.442109 | 0.18311 | 2.414451 | 0.015759 | 0.036372 |
| CNAG_07559 | 4161.621 | 0.660292 | 0.204511 | 3.228639 | 0.001244 | 0.004024 |
| CNAG_07560 | 689.7071 | -0.84942 | 0.147933 | -5.74194 | 9.36E-09 | 9.85E-08 |
| CNAG_07561 | 52251.76 | -0.4105 | 0.137193 | -2.99214 | 0.00277 | 0.008129 |
| CNAG_07563 | 965.6417 | -0.65312 | 0.136803 | -4.77418 | 1.80E-06 | 1.19E-05 |
| CNAG_07564 | 773.3833 | 0.714848 | 0.187563 | 3.81124 | 0.000138 | 0.000582 |
| CNAG_07566 | 626.1745 | 0.320626 | 0.137025 | 2.339914 | 0.019288 | 0.043209 |
| CNAG_07567 | 1206.716 | -0.82187 | 0.18889 | -4.35107 | 1.35E-05 | 7.37E-05 |
| CNAG_07572 | 1117.166 | -1.08157 | 0.238105 | -4.54243 | 5.56E-06 | 3.30E-05 |
| CNAG_02609 | 1531.094 | -0.37795 | 0.168576 | -2.24199 | 0.024962 | 0.053743 |
| CNAG_07573 | 1401.882 | 0.939287 | 0.172109 | 5.457529 | 4.83E-08 | 4.44E-07 |
| CNAG_02606 | 771.1443 | 0.685793 | 0.187611 | 3.655391 | 0.000257 | 0.001009 |
| CNAG_02605 | 1236.565 | 0.51679 | 0.137683 | 3.753477 | 0.000174 | 0.000718 |
| CNAG_02603 | 189.7319 | -1.54576 | 0.233483 | -6.62043 | 3.58E-11 | 5.62E-10 |
| CNAG_02602 | 25.20495 | 1.365423 | 0.435925 | 3.132245 | 0.001735 | 0.005378 |
| CNAG_02600 | 265.6228 | -0.90032 | 0.163952 | -5.49136 | 3.99E-08 | 3.75E-07 |
| CNAG_02599 | 27.91824 | 0.861957 | 0.365377 | 2.359092 | 0.01832 | 0.041348 |
| CNAG_02598 | 95.35641 | 0.605187 | 0.227985 | 2.6545 | 0.007943 | 0.020133 |
| CNAG_02597 | 1757.734 | -0.59188 | 0.180756 | -3.27448 | 0.001059 | 0.003479 |
| CNAG_02593 | 1393.024 | 0.881211 | 0.156302 | 5.637862 | 1.72E-08 | 1.73E-07 |
| CNAG_02592 | 379.2939 | -0.95349 | 0.384619 | -2.47906 | 0.013173 | 0.031125 |
| CNAG_12305 | 29.77101 | 1.51929 | 0.453169 | 3.352594 | 0.000801 | 0.002727 |
| CNAG_02591 | 571.8709 | 1.919582 | 0.333514 | 5.755621 | 8.63E-09 | 9.13E-08 |
| CNAG_02590 | 454.4151 | 0.511274 | 0.15861 | 3.223458 | 0.001267 | 0.004093 |
| CNAG_02589 | 646.5203 | 0.681153 | 0.159895 | 4.260014 | 2.04E-05 | 0.000105 |
| CNAG_02588 | 264.6722 | -0.86995 | 0.250837 | -3.46819 | 0.000524 | 0.001889 |
| CNAG_02587 | 711.5187 | -0.58383 | 0.231138 | -2.52591 | 0.01154 | 0.027787 |
| CNAG_02585 | 4623.952 | 1.12089 | 0.135838 | 8.251654 | 1.56E-16 | 5.13E-15 |
| CNAG_02584 | 574.4212 | 0.83887 | 0.158708 | 5.285612 | 1.25E-07 | 1.05E-06 |
| CNAG_02580 | 1278.262 | -0.42028 | 0.17265 | -2.43428 | 0.014921 | 0.034699 |
| CNAG_02578 | 886.571 | -0.58788 | 0.181524 | -3.23859 | 0.001201 | 0.003902 |
| CNAG_02577 | 3602.693 | 1.127075 | 0.22536 | 5.001217 | 5.70E-07 | 4.16E-06 |
| CNAG_02576 | 1345.297 | 0.631881 | 0.166514 | 3.794766 | 0.000148 | 0.000618 |
| CNAG_02575 | 3968.368 | 0.347584 | 0.175937 | 1.975619 | 0.048198 | 0.094214 |
| CNAG_07574 | 124.0243 | 2.57994 | 0.288849 | 8.931806 | 4.19E-19 | 1.76E-17 |
| CNAG_12313 | 104.3997 | 1.779641 | 0.266153 | 6.686531 | 2.29E-11 | 3.73E-10 |
| CNAG_12314 | 68.29004 | 1.26507 | 0.419951 | 3.012425 | 0.002592 | 0.007676 |
| CNAG_02569 | 835.2963 | -0.362 | 0.118149 | -3.0639 | 0.002185 | 0.006613 |
| CNAG_02568 | 5323.032 | 0.579336 | 0.170876 | 3.390384 | 0.000698 | 0.002418 |
| CNAG_02566 | 1610.458 | 0.559629 | 0.253193 | 2.210282 | 0.027086 | 0.057654 |
| CNAG_02565 | 3103.754 | -1.39262 | 0.203613 | -6.83953 | 7.95E-12 | 1.42E-10 |

| | | | | | | |
|---|---|---|---|---|---|---|
| CNAG_02564 | 802.6593 | -0.86267 | 0.20714 | -4.16465 | 3.12E-05 | 0.000154 |
| CNAG_12315 | 159.6614 | 2.96547 | 0.341744 | 8.677472 | 4.05E-18 | 1.53E-16 |
| CNAG_07972 | 528.0212 | 1.470814 | 0.192827 | 7.627648 | 2.39E-14 | 6.03E-13 |
| CNAG_06879 | 1191.895 | 0.810503 | 0.1906 | 4.252365 | 2.12E-05 | 0.000109 |
| CNAG_06880 | 1940.25 | 0.622398 | 0.215254 | 2.891455 | 0.003835 | 0.010714 |
| CNAG_06881 | 2482.756 | 0.545726 | 0.190301 | 2.867703 | 0.004135 | 0.011455 |
| CNAG_06884 | 745.7562 | 1.483689 | 0.184737 | 8.031364 | 9.64E-16 | 2.92E-14 |
| CNAG_07579 | 621.5265 | -0.52861 | 0.214758 | -2.46143 | 0.013839 | 0.032504 |
| CNAG_07580 | 1275.785 | -0.3652 | 0.133372 | -2.73819 | 0.006178 | 0.016303 |
| CNAG_06890 | 2588.244 | -0.55077 | 0.217177 | -2.53605 | 0.011211 | 0.027127 |
| CNAG_06891 | 1266.943 | -0.50642 | 0.234466 | -2.15987 | 0.030783 | 0.06425 |
| CNAG_06897 | 1520.708 | -0.82744 | 0.246444 | -3.35753 | 0.000786 | 0.002682 |
| CNAG_12320 | 428.7933 | -1.70032 | 0.322482 | -5.2726 | 1.35E-07 | 1.12E-06 |
| CNAG_06899 | 9551.413 | 0.566537 | 0.119796 | 4.729171 | 2.25E-06 | 1.46E-05 |
| CNAG_06900 | 20075.56 | -0.37363 | 0.103498 | -3.61004 | 0.000306 | 0.001179 |
| CNAG_06901 | 679.7173 | 0.550008 | 0.128107 | 4.293364 | 1.76E-05 | 9.27E-05 |
| CNAG_06902 | 1074.299 | -1.01186 | 0.14858 | -6.8102 | 9.75E-12 | 1.72E-10 |
| CNAG_06905 | 2362.491 | 1.550027 | 0.17226 | 8.998164 | 2.30E-19 | 9.97E-18 |
| CNAG_06906 | 9681.462 | -0.65584 | 0.107758 | -6.08627 | 1.16E-09 | 1.45E-08 |
| CNAG_06908 | 15497.89 | -1.11233 | 0.224023 | -4.96524 | 6.86E-07 | 4.93E-06 |
| CNAG_12328 | 127.9838 | 0.833942 | 0.232816 | 3.581985 | 0.000341 | 0.001292 |
| CNAG_06913 | 661.8887 | 0.508044 | 0.187949 | 2.703093 | 0.00687 | 0.017833 |
| CNAG_06915 | 66.02042 | 0.657691 | 0.282804 | 2.325608 | 0.020039 | 0.044666 |
| CNAG_06916 | 871.0569 | -0.5023 | 0.182326 | -2.75494 | 0.00587 | 0.015606 |
| CNAG_06917 | 6961.791 | -1.12096 | 0.307733 | -3.64263 | 0.00027 | 0.001056 |
| CNAG_06918 | 102.4588 | 1.687641 | 0.248186 | 6.799901 | 1.05E-11 | 1.83E-10 |
| CNAG_06922 | 2177.399 | 0.589613 | 0.213686 | 2.75925 | 0.005793 | 0.015432 |
| CNAG_06923 | 16007.53 | -0.97013 | 0.385923 | -2.5138 | 0.011944 | 0.028616 |
| CNAG_06924 | 1157.415 | 1.014191 | 0.171731 | 5.905682 | 3.51E-09 | 4.04E-08 |
| CNAG_06925 | 1023.8 | -0.34885 | 0.1686 | -2.0691 | 0.038537 | 0.077838 |
| CNAG_06926 | 271.116 | -0.32442 | 0.157804 | -2.05588 | 0.039794 | 0.079992 |
| CNAG_06927 | 1822.756 | -0.61504 | 0.143637 | -4.28192 | 1.85E-05 | 9.70E-05 |
| CNAG_06930 | 2152.661 | -1.09566 | 0.332757 | -3.29268 | 0.000992 | 0.003298 |
| CNAG_06932 | 105.3718 | -0.67296 | 0.297985 | -2.25837 | 0.023923 | 0.051965 |
| CNAG_07583 | 730.3172 | 0.668698 | 0.195411 | 3.422012 | 0.000622 | 0.002191 |
| CNAG_06935 | 552.5127 | 2.1694 | 0.271785 | 7.982046 | 1.44E-15 | 4.22E-14 |
| CNAG_04934 | 4481.081 | 2.286163 | 0.178266 | 12.82443 | 1.20E-37 | 1.73E-35 |
| CNAG_04935 | 989.5021 | -0.43135 | 0.156908 | -2.74904 | 0.005977 | 0.015852 |
| CNAG_04936 | 2753.703 | -0.72574 | 0.126438 | -5.7399 | 9.47E-09 | 9.94E-08 |
| CNAG_04937 | 307.039 | -0.42486 | 0.183561 | -2.31455 | 0.020638 | 0.045742 |
| CNAG_04938 | 3291.144 | 2.352202 | 0.182531 | 12.88659 | 5.36E-38 | 8.34E-36 |
| CNAG_04939 | 1204.022 | -0.40644 | 0.204172 | -1.99066 | 0.046519 | 0.091449 |
| CNAG_07794 | 2677.834 | 0.885675 | 0.197287 | 4.489279 | 7.15E-06 | 4.13E-05 |
| CNAG_07796 | 818.8453 | -0.92876 | 0.172918 | -5.37108 | 7.83E-08 | 6.88E-07 |
| CNAG_04943 | 1832.192 | 1.295042 | 0.151647 | 8.539823 | 1.34E-17 | 4.86E-16 |
| CNAG_04944 | 1626.321 | 0.894732 | 0.12612 | 7.094284 | 1.30E-12 | 2.66E-11 |
| CNAG_04945 | 295.971 | 0.459818 | 0.172175 | 2.670649 | 0.00757 | 0.019322 |

| | | | | | | |
|---|---|---|---|---|---|---|
| CNAG_04946 | 257.4237 | -0.4669 | 0.199345 | -2.34216 | 0.019172 | 0.042962 |
| CNAG_12335 | 90.04055 | 3.230261 | 0.306487 | 10.53965 | 5.67E-26 | 4.10E-24 |
| CNAG_04948 | 3410.808 | -0.48662 | 0.118448 | -4.10827 | 3.99E-05 | 0.000192 |
| CNAG_04949 | 1831.938 | -1.07106 | 0.166922 | -6.41658 | 1.39E-10 | 2.03E-09 |
| CNAG_04950 | 3059.952 | -0.59332 | 0.222888 | -2.66194 | 0.007769 | 0.019748 |
| CNAG_04951 | 5103.962 | -1.52992 | 0.191936 | -7.97097 | 1.57E-15 | 4.57E-14 |
| CNAG_04952 | 2088.336 | -0.42712 | 0.161821 | -2.63948 | 0.008303 | 0.020939 |
| CNAG_04953 | 11785.78 | 0.878678 | 0.128781 | 6.823022 | 8.91E-12 | 1.58E-10 |
| CNAG_04957 | 1757.589 | -0.82925 | 0.200891 | -4.12787 | 3.66E-05 | 0.000178 |
| CNAG_04958 | 3486.944 | 0.631946 | 0.265685 | 2.378553 | 0.017381 | 0.039482 |
| CNAG_04961 | 2550.167 | -0.42387 | 0.171615 | -2.46986 | 0.013516 | 0.031813 |
| CNAG_04962 | 3671.794 | -0.61075 | 0.185662 | -3.28956 | 0.001003 | 0.003329 |
| CNAG_04963 | 4383.648 | 0.672979 | 0.1648 | 4.083613 | 4.43E-05 | 0.000211 |
| CNAG_07797 | 1110.224 | -1.84471 | 0.203223 | -9.07727 | 1.11E-19 | 4.94E-18 |
| CNAG_04968 | 415.7163 | -0.58214 | 0.18548 | -3.13855 | 0.001698 | 0.005276 |
| CNAG_04969 | 18071.11 | -0.389 | 0.123702 | -3.14463 | 0.001663 | 0.005194 |
| CNAG_04970 | 1625.676 | 0.851988 | 0.223413 | 3.813512 | 0.000137 | 0.000578 |
| CNAG_07800 | 1289.075 | 0.552596 | 0.215779 | 2.56094 | 0.010439 | 0.02557 |
| CNAG_12342 | 48.84211 | -1.03938 | 0.299363 | -3.47196 | 0.000517 | 0.001868 |
| CNAG_12343 | 8.051266 | 2.1124 | 0.746234 | 2.830746 | 0.004644 | 0.012698 |
| CNAG_04976 | 9543.384 | -1.04769 | 0.233245 | -4.4918 | 7.06E-06 | 4.09E-05 |
| CNAG_04980 | 1254.751 | -0.91552 | 0.230371 | -3.97412 | 7.06E-05 | 0.000321 |
| CNAG_04981 | 5932.724 | 1.321052 | 0.174569 | 7.567489 | 3.81E-14 | 9.36E-13 |
| CNAG_04982 | 223.0487 | -0.82245 | 0.241175 | -3.41019 | 0.000649 | 0.002275 |
| CNAG_04983 | 1362.389 | -0.81644 | 0.215079 | -3.79599 | 0.000147 | 0.000616 |
| CNAG_04984 | 3118.645 | 0.713396 | 0.164257 | 4.34317 | 1.40E-05 | 7.60E-05 |
| CNAG_04985 | 12303.31 | -0.46665 | 0.197811 | -2.35905 | 0.018322 | 0.041348 |
| CNAG_04987 | 691.7229 | 0.879076 | 0.155183 | 5.664768 | 1.47E-08 | 1.50E-07 |
| CNAG_12349 | 42.60027 | 1.695879 | 0.353768 | 4.793755 | 1.64E-06 | 1.09E-05 |
| CNAG_04990 | 3352.517 | -0.52763 | 0.125538 | -4.20293 | 2.63E-05 | 0.000133 |
| CNAG_12351 | 62.35611 | -0.90803 | 0.43865 | -2.07005 | 0.038448 | 0.077739 |
| CNAG_04992 | 733.3083 | 0.550675 | 0.174937 | 3.14784 | 0.001645 | 0.005146 |
| CNAG_04993 | 79.02306 | 1.131954 | 0.264519 | 4.279299 | 1.87E-05 | 9.80E-05 |
| CNAG_04994 | 1055.587 | -0.59618 | 0.198024 | -3.01065 | 0.002607 | 0.007715 |
| CNAG_07802 | 457.1086 | 1.193215 | 0.20029 | 5.957426 | 2.56E-09 | 3.03E-08 |
| CNAG_05001 | 785.1959 | -0.58034 | 0.259548 | -2.23598 | 0.025353 | 0.054492 |
| CNAG_05003 | 637.0037 | 0.697905 | 0.16851 | 4.141616 | 3.45E-05 | 0.000169 |
| CNAG_05004 | 1111.84 | -0.44885 | 0.196854 | -2.28014 | 0.022599 | 0.049537 |
| CNAG_05005 | 1031.464 | 1.544471 | 0.329938 | 4.681095 | 2.85E-06 | 1.82E-05 |
| CNAG_05006 | 712.9969 | -0.46634 | 0.173496 | -2.68787 | 0.007191 | 0.018546 |
| CNAG_05007 | 1309.005 | -0.86562 | 0.163556 | -5.29251 | 1.21E-07 | 1.01E-06 |
| CNAG_05009 | 658.2243 | 1.043002 | 0.208723 | 4.997054 | 5.82E-07 | 4.24E-06 |
| CNAG_12357 | 66.9697 | 1.210127 | 0.27418 | 4.413624 | 1.02E-05 | 5.66E-05 |
| CNAG_05010 | 26.16939 | 1.913308 | 0.46503 | 4.114373 | 3.88E-05 | 0.000188 |
| CNAG_05011 | 982.8566 | -0.52655 | 0.198143 | -2.65743 | 0.007874 | 0.019995 |
| CNAG_05012 | 760.4045 | 0.673356 | 0.15483 | 4.348991 | 1.37E-05 | 7.43E-05 |
| CNAG_05013 | 9673.587 | -0.50259 | 0.213663 | -2.35224 | 0.018661 | 0.041969 |

| | | | | | | |
|---|---|---|---|---|---|---|
| CNAG_05016 | 33.05743 | -1.12614 | 0.505643 | -2.22714 | 0.025938 | 0.055523 |
| CNAG_05019 | 364.9416 | -0.4955 | 0.194728 | -2.54458 | 0.010941 | 0.026586 |
| CNAG_05024 | 262.0588 | 0.93234 | 0.225047 | 4.142877 | 3.43E-05 | 0.000168 |
| CNAG_07803 | 326.6997 | -0.7842 | 0.252572 | -3.10487 | 0.001904 | 0.005838 |
| CNAG_05028 | 4465.034 | -0.45421 | 0.170365 | -2.66607 | 0.007674 | 0.019539 |
| CNAG_05030 | 960.3277 | -0.30814 | 0.123637 | -2.49231 | 0.012691 | 0.030175 |
| CNAG_05032 | 375.8386 | 2.220347 | 0.165393 | 13.42471 | 4.33E-41 | 7.48E-39 |
| CNAG_05035 | 1192.451 | -0.56384 | 0.174035 | -3.23979 | 0.001196 | 0.003888 |
| CNAG_05038 | 738.9182 | 1.418666 | 0.196799 | 7.208698 | 5.65E-13 | 1.21E-11 |
| CNAG_05039 | 1026.051 | -0.73604 | 0.210099 | -3.5033 | 0.00046 | 0.001673 |
| CNAG_05041 | 4252.486 | -1.30851 | 0.160566 | -8.14939 | 3.66E-16 | 1.16E-14 |
| CNAG_05045 | 849.9884 | -0.75528 | 0.171719 | -4.39832 | 1.09E-05 | 6.03E-05 |
| CNAG_07805 | 6320.459 | 0.732037 | 0.173047 | 4.230275 | 2.33E-05 | 0.000119 |
| CNAG_05050 | 4008.081 | -0.82499 | 0.191206 | -4.31465 | 1.60E-05 | 8.49E-05 |
| CNAG_07807 | 23712.03 | -0.95761 | 0.20301 | -4.71709 | 2.39E-06 | 1.55E-05 |
| CNAG_05053 | 630.0395 | -0.70794 | 0.259127 | -2.73203 | 0.006295 | 0.016595 |
| CNAG_12360 | 201.0564 | -1.45679 | 0.304226 | -4.78851 | 1.68E-06 | 1.11E-05 |
| CNAG_05055 | 777.7433 | -0.33738 | 0.124031 | -2.72009 | 0.006526 | 0.017076 |
| CNAG_05057 | 1027.387 | 1.076 | 0.207027 | 5.197401 | 2.02E-07 | 1.61E-06 |
| CNAG_05059 | 12922.68 | -1.03666 | 0.213836 | -4.8479 | 1.25E-06 | 8.44E-06 |
| CNAG_05060 | 1308.945 | 0.475529 | 0.204577 | 2.324452 | 0.020101 | 0.044778 |
| CNAG_05062 | 1621.172 | -0.39915 | 0.148745 | -2.68345 | 0.007287 | 0.018735 |
| CNAG_05067 | 51.65972 | 2.753318 | 0.396302 | 6.94753 | 3.72E-12 | 6.92E-11 |
| CNAG_05069 | 3809.295 | -1.14161 | 0.291622 | -3.91468 | 9.05E-05 | 0.000401 |
| CNAG_05070 | 6619.736 | -1.11725 | 0.188107 | -5.93943 | 2.86E-09 | 3.34E-08 |
| CNAG_12363 | 26.82372 | 1.767828 | 0.440973 | 4.008924 | 6.10E-05 | 0.00028 |
| CNAG_07808 | 3085.218 | -0.29547 | 0.118163 | -2.50053 | 0.012401 | 0.029573 |
| CNAG_05074 | 2044.49 | 0.813686 | 0.245825 | 3.310026 | 0.000933 | 0.003115 |
| CNAG_12365 | 29.00326 | 1.227389 | 0.381119 | 3.220486 | 0.00128 | 0.00413 |
| CNAG_05077 | 2297.277 | 1.277823 | 0.149716 | 8.534987 | 1.40E-17 | 5.02E-16 |
| CNAG_05079 | 729.1761 | -3.21489 | 0.248525 | -12.9359 | 2.82E-38 | 4.49E-36 |
| CNAG_05084 | 4215.496 | 0.419653 | 0.123976 | 3.384938 | 0.000712 | 0.00246 |
| CNAG_05085 | 1670.991 | -0.71238 | 0.203589 | -3.49909 | 0.000467 | 0.001699 |
| CNAG_05086 | 982.2872 | -0.91552 | 0.306096 | -2.99095 | 0.002781 | 0.008155 |
| CNAG_07810 | 10228.97 | -0.35353 | 0.122435 | -2.88752 | 0.003883 | 0.010834 |
| CNAG_05090 | 3397.982 | 0.39743 | 0.112065 | 3.546422 | 0.000391 | 0.001455 |
| CNAG_05091 | 1665.328 | -1.0165 | 0.223525 | -4.54758 | 5.43E-06 | 3.23E-05 |
| CNAG_05092 | 2.115566 | 3.406217 | 1.619428 | 2.103345 | 0.035436 | 0.07257 |
| CNAG_07973 | 276.3132 | 0.428826 | 0.19785 | 2.167426 | 0.030202 | 0.063205 |
| CNAG_05095 | 4252.535 | 1.805081 | 0.204856 | 8.81146 | 1.24E-18 | 4.93E-17 |
| CNAG_05097 | 2061.137 | 0.94513 | 0.203059 | 4.654458 | 3.25E-06 | 2.04E-05 |
| CNAG_05101 | 3170.88 | -0.58437 | 0.117853 | -4.9585 | 7.10E-07 | 5.09E-06 |
| CNAG_12367 | 78.91984 | -0.83024 | 0.38698 | -2.14543 | 0.031919 | 0.066307 |
| CNAG_05102 | 1187.54 | -0.3139 | 0.136472 | -2.30013 | 0.021441 | 0.047298 |
| CNAG_05103 | 824.5066 | -0.26885 | 0.123317 | -2.18017 | 0.029245 | 0.061508 |
| CNAG_05105 | 7567.1 | -0.58405 | 0.121437 | -4.80949 | 1.51E-06 | 1.01E-05 |
| CNAG_05107 | 1770.311 | -0.64981 | 0.254782 | -2.55045 | 0.010758 | 0.026182 |

| | | | | | | |
|---|---|---|---|---|---|---|
| CNAG_05109 | 3576.909 | -0.41754 | 0.115873 | -3.60346 | 0.000314 | 0.001204 |
| CNAG_05110 | 1424.496 | -0.83832 | 0.130262 | -6.43565 | 1.23E-10 | 1.81E-09 |
| CNAG_05111 | 2636.817 | -0.85497 | 0.140887 | -6.06847 | 1.29E-09 | 1.61E-08 |
| CNAG_12368 | 188.6126 | -1.40288 | 0.430432 | -3.25925 | 0.001117 | 0.003651 |
| CNAG_05114 | 357.8961 | -0.96379 | 0.181471 | -5.311 | 1.09E-07 | 9.28E-07 |
| CNAG_05115 | 940.1698 | -0.7311 | 0.166915 | -4.38009 | 1.19E-05 | 6.53E-05 |
| CNAG_05118 | 1125.551 | -0.74174 | 0.231182 | -3.20845 | 0.001335 | 0.004278 |
| CNAG_05119 | 1661.656 | 0.590085 | 0.139171 | 4.240013 | 2.24E-05 | 0.000114 |
| CNAG_05120 | 2818.373 | -0.30241 | 0.116704 | -2.59126 | 0.009563 | 0.023679 |
| CNAG_05121 | 1843.74 | -0.67436 | 0.158694 | -4.24946 | 2.14E-05 | 0.00011 |
| CNAG_05122 | 2476.478 | -0.74404 | 0.112903 | -6.5901 | 4.40E-11 | 6.77E-10 |
| CNAG_05124 | 2553.421 | -0.57553 | 0.129383 | -4.44827 | 8.66E-06 | 4.90E-05 |
| CNAG_05125 | 2182.139 | -0.94833 | 0.162381 | -5.84017 | 5.21E-09 | 5.73E-08 |
| CNAG_07814 | 1008.394 | -0.47662 | 0.13224 | -3.6042 | 0.000313 | 0.001202 |
| CNAG_05132 | 8417.017 | -1.19048 | 0.261641 | -4.55006 | 5.36E-06 | 3.20E-05 |
| CNAG_05137 | 2734.029 | 0.929386 | 0.219698 | 4.230287 | 2.33E-05 | 0.000119 |
| CNAG_05139 | 1588.431 | 0.650188 | 0.217223 | 2.993182 | 0.002761 | 0.008108 |
| CNAG_05140 | 7575.076 | -0.52031 | 0.211236 | -2.46316 | 0.013772 | 0.032357 |
| CNAG_05144 | 8507.47 | -0.41271 | 0.141518 | -2.91631 | 0.003542 | 0.010023 |
| CNAG_05145 | 1062.671 | -0.4006 | 0.177094 | -2.2621 | 0.023691 | 0.051576 |
| CNAG_05147 | 370.8285 | -1.08255 | 0.305412 | -3.54457 | 0.000393 | 0.001464 |
| CNAG_05148 | 3286.341 | -0.33735 | 0.116728 | -2.89005 | 0.003852 | 0.010754 |
| CNAG_05149 | 1004.987 | -0.42556 | 0.139641 | -3.04749 | 0.002308 | 0.006935 |
| CNAG_05150 | 2119.976 | -0.40028 | 0.122857 | -3.25815 | 0.001121 | 0.003662 |
| CNAG_05151 | 1450.83 | -0.76783 | 0.190275 | -4.03538 | 5.45E-05 | 0.000253 |
| CNAG_05152 | 2903.483 | 0.362346 | 0.157961 | 2.293902 | 0.021796 | 0.047962 |
| CNAG_05154 | 1544.633 | -1.21319 | 0.238019 | -5.09703 | 3.45E-07 | 2.62E-06 |
| CNAG_05155 | 3972.795 | 1.371637 | 0.277726 | 4.938818 | 7.86E-07 | 5.57E-06 |
| CNAG_05156 | 1411.722 | 0.547354 | 0.125295 | 4.368538 | 1.25E-05 | 6.85E-05 |
| CNAG_12377 | 247.5288 | 0.935158 | 0.317497 | 2.945408 | 0.003225 | 0.009225 |
| CNAG_05162 | 411.9671 | -0.49729 | 0.241033 | -2.06317 | 0.039097 | 0.078744 |
| CNAG_05164 | 2428.378 | -0.32012 | 0.128683 | -2.48767 | 0.012858 | 0.030526 |
| CNAG_12379 | 25.48405 | 1.043473 | 0.430949 | 2.421336 | 0.015464 | 0.035792 |
| CNAG_05167 | 6773.914 | 1.771977 | 0.228666 | 7.749208 | 9.25E-15 | 2.46E-13 |
| CNAG_05168 | 31.53438 | 1.588414 | 0.410243 | 3.871889 | 0.000108 | 0.000466 |
| CNAG_12380 | 5.481053 | -2.90852 | 1.126391 | -2.58216 | 0.009819 | 0.024237 |
| CNAG_05169 | 489.6052 | -0.60676 | 0.15503 | -3.91383 | 9.08E-05 | 0.000401 |
| CNAG_05170 | 1069.94 | 0.782551 | 0.193026 | 4.054126 | 5.03E-05 | 0.000236 |
| CNAG_05173 | 1158.755 | 0.793664 | 0.142958 | 5.551713 | 2.83E-08 | 2.76E-07 |
| CNAG_07817 | 1322.096 | -1.0051 | 0.23164 | -4.33907 | 1.43E-05 | 7.72E-05 |
| CNAG_07818 | 786.0447 | -0.83243 | 0.151757 | -5.48526 | 4.13E-08 | 3.85E-07 |
| CNAG_05178 | 291.8705 | 1.500961 | 0.191376 | 7.84298 | 4.40E-15 | 1.21E-13 |
| CNAG_05179 | 11609.53 | -1.07292 | 0.156222 | -6.86794 | 6.51E-12 | 1.18E-10 |
| CNAG_05180 | 1075.52 | 0.882212 | 0.196978 | 4.478727 | 7.51E-06 | 4.32E-05 |
| CNAG_05183 | 975.7988 | 0.479634 | 0.148356 | 3.233005 | 0.001225 | 0.003973 |
| CNAG_05184 | 48.05538 | 2.150366 | 0.420414 | 5.11487 | 3.14E-07 | 2.40E-06 |
| CNAG_05185 | 127.6718 | -0.81823 | 0.287136 | -2.84961 | 0.004377 | 0.012073 |

| | | | | | | |
|---|---|---|---|---|---|---|
| CNAG_05186 | 569.3169 | -0.60199 | 0.19005 | -3.16753 | 0.001537 | 0.004838 |
| CNAG_05189 | 924.5358 | -0.53446 | 0.136963 | -3.90218 | 9.53E-05 | 0.000418 |
| CNAG_05190 | 3034.876 | -0.59414 | 0.11451 | -5.18855 | 2.12E-07 | 1.68E-06 |
| CNAG_05192 | 151.6214 | 0.711996 | 0.278775 | 2.554019 | 0.010649 | 0.025956 |
| CNAG_05193 | 1765.219 | 0.629307 | 0.241494 | 2.605891 | 0.009164 | 0.022797 |
| CNAG_05194 | 2672.61 | -0.71762 | 0.228294 | -3.14339 | 0.00167 | 0.00521 |
| CNAG_05195 | 2946.843 | 0.61519 | 0.240428 | 2.55873 | 0.010506 | 0.025701 |
| CNAG_05196 | 1347.815 | -0.7336 | 0.182496 | -4.0198 | 5.82E-05 | 0.000269 |
| CNAG_05197 | 4504.46 | -0.96942 | 0.263724 | -3.67588 | 0.000237 | 0.000939 |
| CNAG_05198 | 1166.816 | 0.300798 | 0.135698 | 2.216674 | 0.026645 | 0.056839 |
| CNAG_05199 | 34192.58 | 0.624193 | 0.13975 | 4.466503 | 7.95E-06 | 4.53E-05 |
| CNAG_05200 | 954.8009 | -0.28704 | 0.118331 | -2.42573 | 0.015278 | 0.035424 |
| CNAG_05201 | 300.3775 | 0.866656 | 0.194573 | 4.454144 | 8.42E-06 | 4.78E-05 |
| CNAG_07819 | 35.73355 | 1.174066 | 0.464363 | 2.528339 | 0.01146 | 0.027635 |
| CNAG_05215 | 1985.354 | 0.333 | 0.107503 | 3.097591 | 0.001951 | 0.005963 |
| CNAG_05216 | 2443.422 | 0.820534 | 0.227572 | 3.605605 | 0.000311 | 0.001197 |
| CNAG_05217 | 384.3323 | -0.92852 | 0.169764 | -5.4695 | 4.51E-08 | 4.17E-07 |
| CNAG_05218 | 4145.776 | 0.268019 | 0.106364 | 2.519822 | 0.011741 | 0.028188 |
| CNAG_05220 | 177.1624 | 1.424295 | 0.226499 | 6.288296 | 3.21E-10 | 4.41E-09 |
| CNAG_05221 | 2982.812 | -1.06748 | 0.256527 | -4.16129 | 3.16E-05 | 0.000156 |
| CNAG_05224 | 189.8762 | -0.63047 | 0.174305 | -3.61705 | 0.000298 | 0.001152 |
| CNAG_05225 | 1371.202 | -0.76105 | 0.233513 | -3.25915 | 0.001117 | 0.003651 |
| CNAG_05226 | 2026.803 | -0.58564 | 0.172611 | -3.39285 | 0.000692 | 0.002399 |
| CNAG_05228 | 1486.275 | -1.2202 | 0.250612 | -4.86887 | 1.12E-06 | 7.66E-06 |
| CNAG_05229 | 1025.524 | -1.51637 | 0.19523 | -7.76713 | 8.03E-15 | 2.16E-13 |
| CNAG_05230 | 623.8783 | -0.46308 | 0.151804 | -3.05054 | 0.002284 | 0.00687 |
| CNAG_05231 | 1689.949 | -0.58186 | 0.222428 | -2.61595 | 0.008898 | 0.022206 |
| CNAG_05232 | 62055.23 | -0.91527 | 0.241357 | -3.7922 | 0.000149 | 0.000622 |
| CNAG_05234 | 2123.627 | -0.54475 | 0.20534 | -2.65293 | 0.00798 | 0.020206 |
| CNAG_05239 | 450.9411 | -0.70803 | 0.192241 | -3.68307 | 0.00023 | 0.000916 |
| CNAG_05240 | 2759.885 | -0.61824 | 0.145101 | -4.26073 | 2.04E-05 | 0.000105 |
| CNAG_05243 | 917.8231 | -0.52833 | 0.138056 | -3.82695 | 0.00013 | 0.000549 |
| CNAG_12387 | 98.91283 | -0.66843 | 0.302746 | -2.20789 | 0.027252 | 0.057942 |
| CNAG_12388 | 33.372 | -0.83892 | 0.404204 | -2.07549 | 0.037941 | 0.076965 |
| CNAG_05245 | 267.091 | -0.7868 | 0.284038 | -2.77004 | 0.005605 | 0.015017 |
| CNAG_07976 | 44.23319 | 1.144745 | 0.300246 | 3.812696 | 0.000137 | 0.00058 |
| CNAG_07822 | 649.6818 | -0.33117 | 0.144394 | -2.29348 | 0.02182 | 0.047989 |
| CNAG_12389 | 32.79039 | 0.93945 | 0.378612 | 2.481301 | 0.01309 | 0.030976 |
| CNAG_05252 | 3053.545 | 0.592845 | 0.167262 | 3.544414 | 0.000393 | 0.001464 |
| CNAG_05253 | 451.4195 | 0.395084 | 0.167848 | 2.353817 | 0.018582 | 0.041819 |
| CNAG_05256 | 1441.332 | 1.18759 | 0.149472 | 7.945253 | 1.94E-15 | 5.58E-14 |
| CNAG_05258 | 429.5345 | 1.616381 | 0.279305 | 5.787146 | 7.16E-09 | 7.68E-08 |
| CNAG_07823 | 164.9203 | 1.000216 | 0.202038 | 4.950639 | 7.40E-07 | 5.28E-06 |
| CNAG_05259 | 76.91036 | 0.605621 | 0.248466 | 2.437445 | 0.014791 | 0.034467 |
| CNAG_05260 | 5551.717 | 0.339793 | 0.169231 | 2.007872 | 0.044657 | 0.088226 |
| CNAG_05261 | 1481.234 | -0.30682 | 0.135624 | -2.26227 | 0.023681 | 0.051566 |
| CNAG_05264 | 519.7254 | 0.826868 | 0.205898 | 4.015902 | 5.92E-05 | 0.000272 |

| | | | | | | |
|---|---|---|---|---|---|---|
| CNAG_12392 | 2273.668 | 3.490258 | 0.28777 | 12.12863 | 7.45E-34 | 9.11E-32 |
| CNAG_05267 | 2470.888 | -1.26844 | 0.230975 | -5.49168 | 3.98E-08 | 3.75E-07 |
| CNAG_05268 | 51.14231 | -1.30039 | 0.414777 | -3.13514 | 0.001718 | 0.005332 |
| CNAG_05269 | 7218.402 | 0.486494 | 0.208563 | 2.332604 | 0.019669 | 0.043951 |
| CNAG_05270 | 2074.312 | -0.41793 | 0.171167 | -2.44165 | 0.01462 | 0.034109 |
| CNAG_05273 | 1731.8 | -0.80159 | 0.262298 | -3.05601 | 0.002243 | 0.006764 |
| CNAG_05274 | 839.8903 | -0.76856 | 0.207557 | -3.7029 | 0.000213 | 0.000854 |
| CNAG_05276 | 1406.115 | -0.47816 | 0.135942 | -3.51742 | 0.000436 | 0.001597 |
| CNAG_05281 | 154.7907 | -0.56981 | 0.236487 | -2.40948 | 0.015975 | 0.036783 |
| CNAG_05283 | 581.5332 | -0.96558 | 0.128635 | -7.50637 | 6.08E-14 | 1.45E-12 |
| CNAG_05284 | 555.8987 | -0.54224 | 0.16446 | -3.29712 | 0.000977 | 0.003248 |
| CNAG_05286 | 574.7402 | 0.432514 | 0.211571 | 2.044299 | 0.040924 | 0.081999 |
| CNAG_05288 | 1104.95 | 1.112602 | 0.158999 | 6.997549 | 2.60E-12 | 5.06E-11 |
| CNAG_12397 | 58.03647 | 1.535927 | 0.29932 | 5.131385 | 2.88E-07 | 2.21E-06 |
| CNAG_12399 | 16.71343 | 1.114243 | 0.527994 | 2.110331 | 0.03483 | 0.07144 |
| CNAG_05290 | 798.3017 | -0.43596 | 0.164362 | -2.65244 | 0.007991 | 0.020229 |
| CNAG_05292 | 8684.86 | 1.040839 | 0.17815 | 5.84249 | 5.14E-09 | 5.67E-08 |
| CNAG_05293 | 3249.927 | -0.99946 | 0.145886 | -6.85096 | 7.34E-12 | 1.32E-10 |
| CNAG_05294 | 1574.95 | 0.697408 | 0.149043 | 4.679251 | 2.88E-06 | 1.83E-05 |
| CNAG_05296 | 857.2664 | 0.465686 | 0.193823 | 2.402629 | 0.016278 | 0.037361 |
| CNAG_05298 | 431.166 | 0.87883 | 0.166799 | 5.268791 | 1.37E-07 | 1.14E-06 |
| CNAG_05299 | 420.4938 | 2.346117 | 0.164906 | 14.22703 | 6.23E-46 | 1.37E-43 |
| CNAG_05300 | 1073.725 | 0.963088 | 0.132765 | 7.254102 | 4.04E-13 | 8.83E-12 |
| CNAG_05301 | 4495.53 | 0.260866 | 0.123415 | 2.113728 | 0.034539 | 0.070897 |
| CNAG_05302 | 1567.99 | -0.94415 | 0.140486 | -6.7206 | 1.81E-11 | 3.00E-10 |
| CNAG_12402 | 48.21376 | 1.539331 | 0.376518 | 4.088332 | 4.34E-05 | 0.000207 |
| CNAG_05304 | 114.9343 | 0.849491 | 0.21757 | 3.904448 | 9.44E-05 | 0.000415 |
| CNAG_12403 | 16.3743 | -1.92758 | 0.71554 | -2.69388 | 0.007063 | 0.018268 |
| CNAG_05307 | 2239.893 | -0.57361 | 0.116338 | -4.93059 | 8.20E-07 | 5.78E-06 |
| CNAG_05308 | 1939.234 | 0.428756 | 0.214851 | 1.995595 | 0.045978 | 0.090543 |
| CNAG_05312 | 5912.553 | -1.88202 | 0.129911 | -14.4869 | 1.47E-47 | 3.53E-45 |
| CNAG_05313 | 1543.945 | -1.10007 | 0.282918 | -3.88828 | 0.000101 | 0.00044 |
| CNAG_05314 | 2989.467 | -0.54049 | 0.197162 | -2.74134 | 0.006119 | 0.016159 |
| CNAG_05315 | 214.6364 | 1.447727 | 0.182668 | 7.92546 | 2.27E-15 | 6.50E-14 |
| CNAG_12405 | 158.547 | 0.620861 | 0.182356 | 3.40467 | 0.000662 | 0.002318 |
| CNAG_05316 | 341.8642 | -1.07153 | 0.172727 | -6.20358 | 5.52E-10 | 7.24E-09 |
| CNAG_05317 | 484.4409 | 0.721171 | 0.175017 | 4.120574 | 3.78E-05 | 0.000183 |
| CNAG_05320 | 195.6438 | -1.18867 | 0.192809 | -6.16503 | 7.05E-10 | 9.13E-09 |
| CNAG_12407 | 247.1847 | -1.93082 | 0.305037 | -6.3298 | 2.45E-10 | 3.43E-09 |
| CNAG_05321 | 144.5405 | -0.87217 | 0.191949 | -4.54377 | 5.53E-06 | 3.29E-05 |
| CNAG_07830 | 498.3555 | -0.52379 | 0.178063 | -2.9416 | 0.003265 | 0.009326 |
| CNAG_05324 | 36.28767 | -1.53026 | 0.364575 | -4.19737 | 2.70E-05 | 0.000135 |
| CNAG_05329 | 39.28908 | -1.28613 | 0.4223 | -3.04553 | 0.002323 | 0.006975 |
| CNAG_05331 | 40.54319 | -0.79034 | 0.354398 | -2.2301 | 0.025741 | 0.05516 |
| CNAG_06876 | 835.9471 | -0.82855 | 0.195026 | -4.24842 | 2.15E-05 | 0.00011 |
| CNAG_06875 | 683.8827 | 0.712558 | 0.182031 | 3.914494 | 9.06E-05 | 0.000401 |
| CNAG_06873 | 63.28879 | 1.250246 | 0.300718 | 4.157539 | 3.22E-05 | 0.000158 |

| | | | | | | |
|---|---|---|---|---|---|---|
| CNAG_12409 | 42.02049 | -1.44668 | 0.425123 | -3.40298 | 0.000667 | 0.002326 |
| CNAG_06870 | 163.3202 | 0.699063 | 0.22404 | 3.120256 | 0.001807 | 0.005567 |
| CNAG_06868 | 4349.777 | 2.238505 | 0.196416 | 11.39674 | 4.34E-30 | 4.11E-28 |
| CNAG_06867 | 932.1456 | 1.455529 | 0.212912 | 6.836294 | 8.13E-12 | 1.45E-10 |
| CNAG_06865 | 774.8699 | -0.46687 | 0.193499 | -2.41279 | 0.015831 | 0.036525 |
| CNAG_06864 | 2448.013 | -0.52682 | 0.15359 | -3.43004 | 0.000603 | 0.002137 |
| CNAG_06863 | 660.9447 | -0.58483 | 0.172883 | -3.38278 | 0.000718 | 0.002477 |
| CNAG_07979 | 1018.758 | -0.41274 | 0.123235 | -3.3492 | 0.00081 | 0.002758 |
| CNAG_06853 | 300.1004 | 1.107217 | 0.18303 | 6.049364 | 1.45E-09 | 1.80E-08 |
| CNAG_06852 | 379.6632 | -0.49644 | 0.198373 | -2.50254 | 0.012331 | 0.02945 |
| CNAG_12414 | 121.5628 | -0.60333 | 0.292249 | -2.06444 | 0.038976 | 0.078584 |
| CNAG_07400 | 10871.93 | -0.46499 | 0.171699 | -2.70818 | 0.006765 | 0.017608 |
| CNAG_06849 | 6019.272 | -1.12343 | 0.222016 | -5.06014 | 4.19E-07 | 3.13E-06 |
| CNAG_06848 | 2.61401 | -2.86569 | 1.448265 | -1.97871 | 0.047849 | 0.093647 |
| CNAG_06847 | 16882.78 | -1.18178 | 0.286157 | -4.12984 | 3.63E-05 | 0.000177 |
| CNAG_07401 | 395.5979 | 0.634766 | 0.22101 | 2.872114 | 0.004077 | 0.011312 |
| CNAG_06846 | 652.5874 | -0.54683 | 0.2766 | -1.97696 | 0.048046 | 0.093987 |
| CNAG_07402 | 1351.531 | 0.643336 | 0.210811 | 3.051714 | 0.002275 | 0.006846 |
| CNAG_06840 | 54410.56 | -0.80762 | 0.125734 | -6.4232 | 1.33E-10 | 1.95E-09 |
| CNAG_06839 | 3645.829 | -0.89784 | 0.235281 | -3.81604 | 0.000136 | 0.000573 |
| CNAG_06835 | 678.4215 | 0.977943 | 0.18567 | 5.267098 | 1.39E-07 | 1.15E-06 |
| CNAG_06834 | 3922.616 | 0.704376 | 0.144773 | 4.865378 | 1.14E-06 | 7.78E-06 |
| CNAG_06832 | 441.4761 | -1.37551 | 0.281921 | -4.87905 | 1.07E-06 | 7.32E-06 |
| CNAG_06831 | 200.2275 | 0.915852 | 0.209442 | 4.372827 | 1.23E-05 | 6.73E-05 |
| CNAG_06830 | 4922.376 | -0.52582 | 0.123197 | -4.26809 | 1.97E-05 | 0.000102 |
| CNAG_06829 | 1884.589 | -0.84456 | 0.134497 | -6.27938 | 3.40E-10 | 4.65E-09 |
| CNAG_06827 | 36.1985 | 0.8526 | 0.359872 | 2.369175 | 0.017828 | 0.040382 |
| CNAG_06825 | 839.1502 | -0.53256 | 0.266326 | -1.99965 | 0.045538 | 0.089765 |
| CNAG_06824 | 1429.35 | -0.43516 | 0.179983 | -2.41779 | 0.015615 | 0.03609 |
| CNAG_07404 | 85.13791 | 0.715938 | 0.286359 | 2.500142 | 0.012414 | 0.029587 |
| CNAG_06818 | 764.7306 | -0.58031 | 0.190709 | -3.04291 | 0.002343 | 0.007023 |
| CNAG_12420 | 252.3418 | -1.21585 | 0.250296 | -4.85765 | 1.19E-06 | 8.07E-06 |
| CNAG_06815 | 147.2827 | -0.79842 | 0.374612 | -2.13131 | 0.033063 | 0.068301 |
| CNAG_12422 | 7.45228 | -1.84332 | 0.79112 | -2.33001 | 0.019806 | 0.044194 |
| CNAG_06813 | 3484.895 | -0.85814 | 0.158815 | -5.4034 | 6.54E-08 | 5.88E-07 |
| CNAG_12423 | 32.38781 | -1.52982 | 0.481741 | -3.17561 | 0.001495 | 0.00472 |
| CNAG_06811 | 18846.97 | -1.12709 | 0.239918 | -4.69782 | 2.63E-06 | 1.69E-05 |
| CNAG_06807 | 233.4056 | -0.7617 | 0.287866 | -2.646 | 0.008145 | 0.020601 |
| CNAG_07406 | 80.70142 | 1.513906 | 0.36598 | 4.136578 | 3.53E-05 | 0.000172 |
| CNAG_07407 | 80.94351 | 1.497833 | 0.326091 | 4.593301 | 4.36E-06 | 2.67E-05 |
| CNAG_07005 | 1054.757 | 0.504321 | 0.144002 | 3.502186 | 0.000461 | 0.00168 |
| CNAG_01455 | 12402.48 | -0.95721 | 0.256426 | -3.73288 | 0.000189 | 0.000771 |
| CNAG_12429 | 84.66806 | -0.74444 | 0.328648 | -2.26516 | 0.023503 | 0.05125 |
| CNAG_01451 | 1344.372 | -0.46956 | 0.238173 | -1.97153 | 0.048664 | 0.095008 |
| CNAG_07413 | 6028.778 | -0.93945 | 0.136293 | -6.89282 | 5.47E-12 | 1.00E-10 |
| CNAG_01446 | 12203.12 | 2.01732 | 0.224944 | 8.968086 | 3.02E-19 | 1.30E-17 |
| CNAG_01444 | 974.7183 | -0.70486 | 0.236283 | -2.98312 | 0.002853 | 0.008335 |

| | | | | | | |
|---|---|---|---|---|---|---|
| CNAG_01441 | 1522.868 | 0.789731 | 0.124987 | 6.318521 | 2.64E-10 | 3.67E-09 |
| CNAG_01439 | 2380.578 | -0.451 | 0.114878 | -3.92588 | 8.64E-05 | 0.000384 |
| CNAG_01437 | 2062.458 | -0.97457 | 0.125062 | -7.79268 | 6.56E-15 | 1.77E-13 |
| CNAG_01435 | 8951.149 | -0.58757 | 0.141298 | -4.15839 | 3.20E-05 | 0.000158 |
| CNAG_01432 | 2056.004 | -0.69128 | 0.132842 | -5.2038 | 1.95E-07 | 1.57E-06 |
| CNAG_01430 | 1042.513 | -0.53403 | 0.145395 | -3.67297 | 0.00024 | 0.000948 |
| CNAG_01428 | 30569.31 | -0.73825 | 0.170272 | -4.33572 | 1.45E-05 | 7.83E-05 |
| CNAG_01425 | 215.3935 | -0.99407 | 0.270674 | -3.67256 | 0.00024 | 0.000949 |
| CNAG_01424 | 1502.333 | 0.785798 | 0.302627 | 2.596585 | 0.009416 | 0.023358 |
| CNAG_01422 | 2873.964 | 0.828983 | 0.136111 | 6.090478 | 1.13E-09 | 1.42E-08 |
| CNAG_01421 | 5111.573 | 0.694933 | 0.129532 | 5.364966 | 8.10E-08 | 7.09E-07 |
| CNAG_07416 | 1084.558 | -0.43804 | 0.202916 | -2.15871 | 0.030872 | 0.064386 |
| CNAG_01417 | 1827.65 | 1.121181 | 0.198632 | 5.644507 | 1.66E-08 | 1.67E-07 |
| CNAG_01412 | 3119.712 | 0.460684 | 0.129266 | 3.563836 | 0.000365 | 0.001371 |
| CNAG_07418 | 952.1605 | -0.45421 | 0.196111 | -2.3161 | 0.020553 | 0.045592 |
| CNAG_01404 | 8477.965 | -0.88046 | 0.21662 | -4.06453 | 4.81E-05 | 0.000226 |
| CNAG_01402 | 6222.176 | 0.347216 | 0.106096 | 3.27266 | 0.001065 | 0.003498 |
| CNAG_01401 | 69.06397 | 1.190341 | 0.296222 | 4.018417 | 5.86E-05 | 0.00027 |
| CNAG_01400 | 7327.763 | -1.08566 | 0.156475 | -6.93824 | 3.97E-12 | 7.37E-11 |
| CNAG_01399 | 1178.941 | -0.69427 | 0.2176 | -3.19057 | 0.00142 | 0.004512 |
| CNAG_01398 | 3187.976 | -0.44612 | 0.213457 | -2.09 | 0.036618 | 0.07469 |
| CNAG_01395 | 3173.359 | -0.51579 | 0.219144 | -2.35368 | 0.018589 | 0.041819 |
| CNAG_07420 | 520.213 | -0.4139 | 0.178112 | -2.32382 | 0.020135 | 0.044829 |
| CNAG_01392 | 703.257 | -0.34423 | 0.140149 | -2.4562 | 0.014041 | 0.032919 |
| CNAG_01390 | 2985.287 | -0.92091 | 0.25387 | -3.62747 | 0.000286 | 0.001113 |
| CNAG_01389 | 633.7015 | -1.09475 | 0.156727 | -6.98505 | 2.85E-12 | 5.44E-11 |
| CNAG_01387 | 3537.926 | 0.59399 | 0.243015 | 2.444252 | 0.014515 | 0.033893 |
| CNAG_01385 | 2225.135 | 0.329895 | 0.129344 | 2.550528 | 0.010756 | 0.026182 |
| CNAG_01384 | 295.5677 | 1.040897 | 0.17003 | 6.121837 | 9.25E-10 | 1.18E-08 |
| CNAG_01381 | 1573.89 | -0.69166 | 0.22321 | -3.09868 | 0.001944 | 0.005948 |
| CNAG_01375 | 2088.011 | 0.887126 | 0.143697 | 6.173599 | 6.68E-10 | 8.68E-09 |
| CNAG_01372 | 1463.122 | -1.09384 | 0.170842 | -6.40263 | 1.53E-10 | 2.21E-09 |
| CNAG_12437 | 64.60372 | -1.16214 | 0.471323 | -2.46569 | 0.013675 | 0.032148 |
| CNAG_01369 | 128.3092 | -0.59746 | 0.235468 | -2.53732 | 0.01117 | 0.027045 |
| CNAG_01367 | 175.4867 | 1.055496 | 0.349977 | 3.015903 | 0.002562 | 0.007605 |
| CNAG_01368 | 11.34998 | 3.643044 | 0.863472 | 4.219065 | 2.45E-05 | 0.000124 |
| CNAG_01365 | 891.744 | -1.07739 | 0.162825 | -6.61683 | 3.67E-11 | 5.74E-10 |
| CNAG_01364 | 3895.677 | -0.96068 | 0.247126 | -3.88742 | 0.000101 | 0.000441 |
| CNAG_01361 | 6593.432 | -0.31969 | 0.10885 | -2.93701 | 0.003314 | 0.009455 |
| CNAG_01360 | 2531.152 | 0.733869 | 0.247301 | 2.967517 | 0.003002 | 0.00869 |
| CNAG_01359 | 3365.318 | -0.42756 | 0.170862 | -2.50235 | 0.012337 | 0.029453 |
| CNAG_01356 | 957.9734 | 0.765417 | 0.236855 | 3.23158 | 0.001231 | 0.003989 |
| CNAG_01354 | 2822.475 | -1.64968 | 0.196508 | -8.39497 | 4.66E-17 | 1.60E-15 |
| CNAG_01351 | 1707.517 | -0.60641 | 0.22572 | -2.68657 | 0.007219 | 0.018602 |
| CNAG_01350 | 1320.952 | 0.492231 | 0.163641 | 3.008003 | 0.00263 | 0.007777 |
| CNAG_01348 | 2822.84 | 1.811996 | 0.214701 | 8.439632 | 3.18E-17 | 1.10E-15 |
| CNAG_01347 | 4235.182 | 0.934156 | 0.121328 | 7.699414 | 1.37E-14 | 3.54E-13 |

| | | | | | | |
|---|---|---|---|---|---|---|
| CNAG_01345 | 851.2059 | -0.92277 | 0.231989 | -3.97765 | 6.96E-05 | 0.000316 |
| CNAG_01344 | 1376.336 | 1.554287 | 0.197605 | 7.865618 | 3.67E-15 | 1.02E-13 |
| CNAG_01341 | 2834.516 | 2.353623 | 0.242935 | 9.688289 | 3.38E-22 | 1.93E-20 |
| CNAG_01339 | 3412.018 | -0.37532 | 0.116923 | -3.20994 | 0.001328 | 0.004262 |
| CNAG_01338 | 1058.171 | 0.49014 | 0.223074 | 2.197213 | 0.028005 | 0.05931 |
| CNAG_01334 | 1901.451 | -0.72825 | 0.269267 | -2.70457 | 0.006839 | 0.017766 |
| CNAG_01333 | 727.8299 | -0.60476 | 0.295485 | -2.04668 | 0.04069 | 0.081605 |
| CNAG_01332 | 34317.43 | -0.97593 | 0.2791 | -3.49672 | 0.000471 | 0.001713 |
| CNAG_01331 | 522.8417 | 0.467455 | 0.152961 | 3.056035 | 0.002243 | 0.006764 |
| CNAG_07425 | 45.46107 | 1.071247 | 0.369545 | 2.898828 | 0.003746 | 0.010513 |
| CNAG_01324 | 569.5881 | 0.376131 | 0.149955 | 2.508291 | 0.012132 | 0.029036 |
| CNAG_01323 | 7481.49 | -1.11925 | 0.231197 | -4.84112 | 1.29E-06 | 8.72E-06 |
| CNAG_01321 | 6558.709 | -0.47246 | 0.235176 | -2.00898 | 0.04454 | 0.088016 |
| CNAG_01320 | 2406.446 | -0.62809 | 0.218494 | -2.87463 | 0.004045 | 0.011231 |
| CNAG_01318 | 3458.25 | -0.41187 | 0.148335 | -2.77663 | 0.005493 | 0.014745 |
| CNAG_01317 | 1347.215 | -0.89534 | 0.149005 | -6.00882 | 1.87E-09 | 2.28E-08 |
| CNAG_01314 | 2047.558 | -0.46676 | 0.135893 | -3.43477 | 0.000593 | 0.002106 |
| CNAG_07426 | 1776.679 | 0.666885 | 0.171421 | 3.890329 | 0.0001 | 0.000436 |
| CNAG_07427 | 14043.19 | 0.455997 | 0.105233 | 4.33322 | 1.47E-05 | 7.89E-05 |
| CNAG_01310 | 1022.595 | -0.82374 | 0.195761 | -4.2079 | 2.58E-05 | 0.00013 |
| CNAG_01308 | 1139.805 | 0.408338 | 0.178538 | 2.287125 | 0.022189 | 0.048704 |
| CNAG_01307 | 4651.193 | 0.466898 | 0.181914 | 2.566592 | 0.01027 | 0.025195 |
| CNAG_01305 | 6782.045 | -0.64145 | 0.110459 | -5.80714 | 6.35E-09 | 6.85E-08 |
| CNAG_01304 | 1204.083 | -0.644 | 0.134494 | -4.78835 | 1.68E-06 | 1.11E-05 |
| CNAG_01300 | 14240.16 | -1.13883 | 0.273606 | -4.16229 | 3.15E-05 | 0.000156 |
| CNAG_01299 | 51.31781 | 1.456659 | 0.311973 | 4.669186 | 3.02E-06 | 1.91E-05 |
| CNAG_01297 | 842.3865 | 0.428366 | 0.184642 | 2.319981 | 0.020342 | 0.045226 |
| CNAG_01287 | 11527.44 | -1.25545 | 0.129387 | -9.70304 | 2.93E-22 | 1.69E-20 |
| CNAG_01285 | 1612.784 | -0.7953 | 0.161405 | -4.92736 | 8.33E-07 | 5.87E-06 |
| CNAG_01283 | 1756.409 | -0.90877 | 0.197673 | -4.59736 | 4.28E-06 | 2.63E-05 |
| CNAG_01281 | 671.7877 | -0.43338 | 0.135052 | -3.20897 | 0.001332 | 0.004275 |
| CNAG_01279 | 1164.295 | -0.77972 | 0.168729 | -4.62113 | 3.82E-06 | 2.36E-05 |
| CNAG_07428 | 2719.083 | 0.339531 | 0.168589 | 2.013961 | 0.044014 | 0.087235 |
| CNAG_01278 | 2323.38 | 0.557466 | 0.198728 | 2.805175 | 0.005029 | 0.013638 |
| CNAG_01277 | 7966.421 | 0.496007 | 0.134682 | 3.682786 | 0.000231 | 0.000917 |
| CNAG_12450 | 30.91744 | 2.581459 | 0.525788 | 4.909699 | 9.12E-07 | 6.37E-06 |
| CNAG_01276 | 3508.749 | -0.52871 | 0.205708 | -2.57018 | 0.010165 | 0.024952 |
| CNAG_01275 | 1521.553 | 1.453022 | 0.148422 | 9.789828 | 1.25E-22 | 7.55E-21 |
| CNAG_01273 | 8410.601 | -0.84432 | 0.312663 | -2.70042 | 0.006925 | 0.017953 |
| CNAG_01272 | 811.9559 | 2.149304 | 0.172838 | 12.43535 | 1.68E-35 | 2.22E-33 |
| CNAG_01271 | 2960.455 | -0.52431 | 0.154184 | -3.40057 | 0.000672 | 0.002344 |
| CNAG_07429 | 2254.546 | -0.41498 | 0.150469 | -2.75792 | 0.005817 | 0.01548 |
| CNAG_01267 | 435.0874 | -0.86452 | 0.212251 | -4.07309 | 4.64E-05 | 0.000219 |
| CNAG_01266 | 4423.174 | 0.441842 | 0.2213 | 1.996575 | 0.045871 | 0.090355 |
| CNAG_01264 | 9531.564 | -0.77968 | 0.182525 | -4.27164 | 1.94E-05 | 0.000101 |
| CNAG_01261 | 7301.176 | 0.905493 | 0.169855 | 5.33097 | 9.77E-08 | 8.42E-07 |
| CNAG_06946 | 2091.037 | -1.35988 | 0.168392 | -8.07569 | 6.71E-16 | 2.07E-14 |

| | | | | | | |
|---|---|---|---|---|---|---|
| CNAG_06948 | 2910.131 | -0.75714 | 0.111326 | -6.80106 | 1.04E-11 | 1.83E-10 |
| CNAG_06949 | 4129.724 | -0.62643 | 0.236626 | -2.64734 | 0.008113 | 0.02053 |
| CNAG_01258 | 2800.366 | 0.479847 | 0.235842 | 2.034614 | 0.04189 | 0.08357 |
| CNAG_01256 | 245.8161 | -0.47334 | 0.177429 | -2.66779 | 0.007635 | 0.019458 |
| CNAG_01255 | 2767.316 | 1.174349 | 0.126677 | 9.270396 | 1.85E-20 | 9.10E-19 |
| CNAG_01254 | 472.5706 | 0.496581 | 0.149583 | 3.319772 | 0.000901 | 0.003015 |
| CNAG_01252 | 551.3967 | 0.547945 | 0.200666 | 2.730638 | 0.006321 | 0.016644 |
| CNAG_01251 | 3650.645 | 0.997763 | 0.336958 | 2.961086 | 0.003066 | 0.00883 |
| CNAG_01247 | 1054.803 | -0.44664 | 0.185664 | -2.40561 | 0.016145 | 0.037111 |
| CNAG_01245 | 1260.525 | -0.51131 | 0.215796 | -2.3694 | 0.017817 | 0.040381 |
| CNAG_01244 | 137.6083 | 3.64265 | 0.302709 | 12.03349 | 2.37E-33 | 2.82E-31 |
| CNAG_01242 | 2926.845 | 0.904733 | 0.265802 | 3.40378 | 0.000665 | 0.002321 |
| CNAG_01240 | 768.5472 | -0.51905 | 0.162796 | -3.18838 | 0.001431 | 0.004537 |
| CNAG_01239 | 1323.188 | -0.58113 | 0.213271 | -2.72487 | 0.006433 | 0.016875 |
| CNAG_07981 | 77.24545 | 0.911314 | 0.271404 | 3.357777 | 0.000786 | 0.002681 |
| CNAG_01236 | 3647.467 | -0.79378 | 0.128199 | -6.19181 | 5.95E-10 | 7.77E-09 |
| CNAG_01234 | 2144.726 | 0.80674 | 0.28885 | 2.792935 | 0.005223 | 0.014117 |
| CNAG_01232 | 2567.965 | 0.851511 | 0.301865 | 2.820832 | 0.00479 | 0.013061 |
| CNAG_12456 | 7.555182 | 2.474006 | 0.905804 | 2.731282 | 0.006309 | 0.016621 |
| CNAG_01230 | 5626.209 | 0.885567 | 0.162348 | 5.454755 | 4.90E-08 | 4.50E-07 |
| CNAG_01229 | 734.7625 | -0.84061 | 0.212848 | -3.94934 | 7.84E-05 | 0.000352 |
| CNAG_07432 | 150.0049 | 0.739985 | 0.303384 | 2.439101 | 0.014724 | 0.03433 |
| CNAG_01224 | 45598.54 | -1.06444 | 0.216552 | -4.91539 | 8.86E-07 | 6.21E-06 |
| CNAG_01223 | 422.3358 | 2.485368 | 0.179369 | 13.85621 | 1.17E-43 | 2.32E-41 |
| CNAG_01222 | 2774.251 | -0.6141 | 0.222629 | -2.75839 | 0.005809 | 0.015463 |
| CNAG_01221 | 1100.042 | -0.63033 | 0.160427 | -3.92905 | 8.53E-05 | 0.00038 |
| CNAG_01213 | 2081.782 | -0.49036 | 0.162393 | -3.01961 | 0.002531 | 0.007524 |
| CNAG_01212 | 1225.418 | 0.442571 | 0.186508 | 2.372932 | 0.017648 | 0.040019 |
| CNAG_12458 | 23.54695 | 0.979155 | 0.424822 | 2.304859 | 0.021174 | 0.046789 |
| CNAG_01208 | 1410.039 | 0.586501 | 0.297672 | 1.970295 | 0.048805 | 0.095259 |
| CNAG_01207 | 601.4402 | 0.414475 | 0.161144 | 2.572079 | 0.010109 | 0.02483 |
| CNAG_01204 | 13514.44 | -0.61497 | 0.261481 | -2.35186 | 0.01868 | 0.041999 |
| CNAG_01203 | 1049.632 | -0.27065 | 0.126577 | -2.13823 | 0.032498 | 0.067369 |
| CNAG_01202 | 523.0778 | -0.43116 | 0.181896 | -2.37035 | 0.017771 | 0.040288 |
| CNAG_01201 | 583.628 | -0.38942 | 0.171995 | -2.26414 | 0.023566 | 0.051358 |
| CNAG_01200 | 630.9937 | -0.55205 | 0.206876 | -2.66852 | 0.007619 | 0.019428 |
| CNAG_01197 | 2126.857 | -0.96829 | 0.154729 | -6.25799 | 3.90E-10 | 5.26E-09 |
| CNAG_01192 | 386.5002 | -0.66309 | 0.20866 | -3.17787 | 0.001484 | 0.004687 |
| CNAG_01191 | 656.0215 | 0.502152 | 0.136603 | 3.675985 | 0.000237 | 0.000939 |
| CNAG_01188 | 1077.967 | -0.44398 | 0.131992 | -3.36373 | 0.000769 | 0.00263 |
| CNAG_01187 | 1997.506 | -0.60126 | 0.147212 | -4.08435 | 4.42E-05 | 0.00021 |
| CNAG_01186 | 2053.705 | 0.411712 | 0.139293 | 2.955731 | 0.003119 | 0.008964 |
| CNAG_01184 | 1578.007 | 0.732487 | 0.16643 | 4.401166 | 1.08E-05 | 5.96E-05 |
| CNAG_01183 | 1911.839 | 0.265726 | 0.117537 | 2.260778 | 0.023773 | 0.051711 |
| CNAG_07436 | 17.79165 | 1.034969 | 0.521389 | 1.985022 | 0.047142 | 0.092492 |
| CNAG_01182 | 8492.668 | -0.56732 | 0.16443 | -3.45021 | 0.00056 | 0.002004 |
| CNAG_01181 | 37353.62 | -0.94622 | 0.217375 | -4.35296 | 1.34E-05 | 7.31E-05 |

| | | | | | | |
|---|---|---|---|---|---|---|
| CNAG_01180 | 518.5021 | -0.79308 | 0.186578 | -4.25065 | 2.13E-05 | 0.000109 |
| CNAG_01179 | 1892.955 | -0.82571 | 0.194643 | -4.24215 | 2.21E-05 | 0.000113 |
| CNAG_07437 | 730.9099 | -0.41053 | 0.176039 | -2.33205 | 0.019698 | 0.043991 |
| CNAG_07438 | 678.3816 | -0.31169 | 0.138292 | -2.25386 | 0.024205 | 0.052479 |
| CNAG_01174 | 3329.836 | 0.768185 | 0.179725 | 4.27422 | 1.92E-05 | 9.99E-05 |
| CNAG_01171 | 501.7157 | -0.52612 | 0.265614 | -1.98078 | 0.047616 | 0.093283 |
| CNAG_01170 | 52758.91 | -1.00914 | 0.271109 | -3.72227 | 0.000197 | 0.000799 |
| CNAG_01168 | 10109.32 | -0.91836 | 0.198484 | -4.62687 | 3.71E-06 | 2.30E-05 |
| CNAG_01163 | 1194.297 | 0.6272 | 0.187472 | 3.34556 | 0.000821 | 0.002785 |
| CNAG_01161 | 2508.138 | 0.438383 | 0.15129 | 2.897624 | 0.00376 | 0.01055 |
| CNAG_07439 | 3202.443 | -0.34148 | 0.122556 | -2.78634 | 0.005331 | 0.014363 |
| CNAG_01159 | 1883.669 | -0.26568 | 0.11672 | -2.27619 | 0.022835 | 0.050026 |
| CNAG_01158 | 5972.228 | -0.37412 | 0.116433 | -3.21315 | 0.001313 | 0.004225 |
| CNAG_01155 | 8682.798 | 0.567096 | 0.135112 | 4.197236 | 2.70E-05 | 0.000135 |
| CNAG_01154 | 280.3676 | 0.414988 | 0.200264 | 2.072203 | 0.038247 | 0.077429 |
| CNAG_01153 | 25745.85 | -0.55427 | 0.202109 | -2.74244 | 0.006099 | 0.016124 |
| CNAG_01152 | 42005.45 | -1.02937 | 0.233376 | -4.41077 | 1.03E-05 | 5.73E-05 |
| CNAG_01148 | 11770.59 | -0.65054 | 0.159951 | -4.06714 | 4.76E-05 | 0.000224 |
| CNAG_01146 | 2538.954 | -0.49426 | 0.23845 | -2.0728 | 0.038191 | 0.077374 |
| CNAG_01145 | 832.1088 | -1.10819 | 0.146957 | -7.54092 | 4.67E-14 | 1.14E-12 |
| CNAG_01144 | 4055.192 | 0.700363 | 0.190671 | 3.673146 | 0.00024 | 0.000948 |
| CNAG_01142 | 2118.601 | 0.882134 | 0.268678 | 3.283234 | 0.001026 | 0.003395 |
| CNAG_01141 | 1000.333 | -0.56633 | 0.242829 | -2.33221 | 0.019689 | 0.043984 |
| CNAG_01140 | 2217.491 | -0.69629 | 0.222784 | -3.12541 | 0.001776 | 0.00549 |
| CNAG_01138 | 10071.32 | -2.09868 | 0.182862 | -11.4768 | 1.73E-30 | 1.67E-28 |
| CNAG_01137 | 9140.795 | -0.39625 | 0.181632 | -2.18161 | 0.029139 | 0.061318 |
| CNAG_01136 | 2839.992 | -0.56301 | 0.135054 | -4.16879 | 3.06E-05 | 0.000152 |
| CNAG_01134 | 808.052 | -0.51306 | 0.151067 | -3.39627 | 0.000683 | 0.002377 |
| CNAG_01133 | 457.9812 | -0.50102 | 0.181794 | -2.75596 | 0.005852 | 0.015562 |
| CNAG_01132 | 1169.976 | -0.78923 | 0.13379 | -5.899 | 3.66E-09 | 4.16E-08 |
| CNAG_01131 | 430.6303 | 0.953074 | 0.165771 | 5.749342 | 8.96E-09 | 9.47E-08 |
| CNAG_12463 | 13.57317 | 1.593759 | 0.567364 | 2.809059 | 0.004969 | 0.013493 |
| CNAG_01130 | 3405.705 | -0.42792 | 0.183385 | -2.33345 | 0.019625 | 0.043864 |
| CNAG_01128 | 2322.918 | 0.422385 | 0.166988 | 2.529426 | 0.011425 | 0.02756 |
| CNAG_01127 | 1233.686 | 0.426746 | 0.133267 | 3.202182 | 0.001364 | 0.004356 |
| CNAG_01126 | 4467.578 | 0.335282 | 0.170705 | 1.964099 | 0.049519 | 0.09644 |
| CNAG_12465 | 10.51798 | 2.740964 | 0.740281 | 3.702601 | 0.000213 | 0.000855 |
| CNAG_01124 | 205.7291 | -0.93332 | 0.289899 | -3.21945 | 0.001284 | 0.004142 |
| CNAG_01123 | 1182.266 | -0.61858 | 0.14316 | -4.32093 | 1.55E-05 | 8.29E-05 |
| CNAG_01121 | 185.0343 | 1.549087 | 0.205595 | 7.53467 | 4.90E-14 | 1.19E-12 |
| CNAG_01120 | 16577.04 | -0.55756 | 0.11074 | -5.03488 | 4.78E-07 | 3.54E-06 |
| CNAG_01119 | 429.6923 | -0.70735 | 0.232579 | -3.04133 | 0.002355 | 0.007053 |
| CNAG_01118 | 7592.33 | -0.42742 | 0.195657 | -2.18453 | 0.028923 | 0.060943 |
| CNAG_12469 | 61.0564 | -2.41013 | 0.387624 | -6.21771 | 5.04E-10 | 6.65E-09 |
| CNAG_01117 | 45869.88 | -0.89295 | 0.21487 | -4.15577 | 3.24E-05 | 0.00016 |
| CNAG_01116 | 372.6243 | -0.34464 | 0.171989 | -2.00386 | 0.045085 | 0.08894 |
| CNAG_01115 | 2943.645 | 0.928371 | 0.371901 | 2.496287 | 0.01255 | 0.029875 |

| | | | | | | |
|---|---|---|---|---|---|---|
| CNAG_01113 | 1460.944 | -1.02807 | 0.229537 | -4.47889 | 7.50E-06 | 4.32E-05 |
| CNAG_01112 | 5188.532 | 1.261425 | 0.137323 | 9.185791 | 4.09E-20 | 1.88E-18 |
| CNAG_01111 | 4203.993 | -0.71481 | 0.193039 | -3.70294 | 0.000213 | 0.000854 |
| CNAG_12470 | 22.40824 | -1.18769 | 0.537499 | -2.20966 | 0.027129 | 0.05773 |
| CNAG_01108 | 1082.132 | -1.11265 | 0.182093 | -6.11035 | 9.94E-10 | 1.26E-08 |
| CNAG_01107 | 1781.246 | -0.32259 | 0.147615 | -2.18533 | 0.028865 | 0.060838 |
| CNAG_01104 | 1317.804 | -0.46537 | 0.143184 | -3.25014 | 0.001153 | 0.003759 |
| CNAG_01102 | 6345.792 | 2.508447 | 0.234456 | 10.69902 | 1.03E-26 | 7.64E-25 |
| CNAG_12472 | 17.47865 | 2.411074 | 0.566043 | 4.259524 | 2.05E-05 | 0.000105 |
| CNAG_01097 | 2439.086 | 0.765813 | 0.142941 | 5.35753 | 8.44E-08 | 7.37E-07 |
| CNAG_01095 | 1201.049 | 0.960905 | 0.13089 | 7.341291 | 2.12E-13 | 4.71E-12 |
| CNAG_01092 | 1740.827 | -0.65722 | 0.150623 | -4.36335 | 1.28E-05 | 7.00E-05 |
| CNAG_01090 | 793.6839 | 1.82608 | 0.16426 | 11.117 | 1.04E-28 | 8.77E-27 |
| CNAG_01089 | 1512.707 | -0.77238 | 0.210961 | -3.66124 | 0.000251 | 0.000987 |
| CNAG_01088 | 1250.149 | -0.37467 | 0.136618 | -2.74245 | 0.006098 | 0.016124 |
| CNAG_12473 | 17.95653 | -1.10277 | 0.493368 | -2.23519 | 0.025405 | 0.054574 |
| CNAG_01083 | 7631.149 | 0.593022 | 0.13127 | 4.517556 | 6.26E-06 | 3.68E-05 |
| CNAG_01082 | 1604.731 | 0.650913 | 0.323695 | 2.010882 | 0.044338 | 0.087748 |
| CNAG_01081 | 200.2262 | 0.921369 | 0.279344 | 3.298327 | 0.000973 | 0.003235 |
| CNAG_01080 | 195.162 | -0.90216 | 0.32783 | -2.75192 | 0.005925 | 0.01574 |
| CNAG_01077 | 40.22078 | -0.91583 | 0.341104 | -2.68491 | 0.007255 | 0.018668 |
| CNAG_01075 | 46.52481 | -1.00837 | 0.395706 | -2.54829 | 0.010825 | 0.026329 |
| CNAG_01072 | 269.0297 | -0.4604 | 0.165012 | -2.79008 | 0.00527 | 0.014218 |
| CNAG_01070 | 28.46529 | -1.09141 | 0.472962 | -2.30761 | 0.021021 | 0.046488 |
| CNAG_07443 | 54.30638 | 2.431494 | 0.3612 | 6.731717 | 1.68E-11 | 2.80E-10 |
| CNAG_01060 | 9964.917 | -0.64906 | 0.263421 | -2.46395 | 0.013742 | 0.032295 |
| CNAG_01059 | 612.2416 | 1.255645 | 0.13146 | 9.55153 | 1.28E-21 | 6.91E-20 |
| CNAG_01058 | 1323.25 | -0.3102 | 0.133539 | -2.32292 | 0.020183 | 0.044898 |
| CNAG_01057 | 2354.965 | 0.379611 | 0.169317 | 2.242015 | 0.02496 | 0.053743 |
| CNAG_01056 | 49.78544 | 1.232127 | 0.323552 | 3.808126 | 0.00014 | 0.000589 |
| CNAG_07444 | 118.17 | 0.747083 | 0.249583 | 2.993328 | 0.00276 | 0.008108 |
| CNAG_12480 | 4.273506 | -2.94385 | 1.151 | -2.55764 | 0.010538 | 0.02575 |
| CNAG_01055 | 10589.16 | -0.49137 | 0.134926 | -3.64178 | 0.000271 | 0.001058 |
| CNAG_01054 | 887.0286 | -0.52424 | 0.189453 | -2.76715 | 0.005655 | 0.015134 |
| CNAG_01052 | 13353.36 | 1.601719 | 0.277105 | 5.780198 | 7.46E-09 | 7.99E-08 |
| CNAG_01050 | 1145.59 | -0.50184 | 0.181404 | -2.76643 | 0.005667 | 0.015147 |
| CNAG_01048 | 185.3823 | 1.780798 | 0.24966 | 7.132886 | 9.83E-13 | 2.04E-11 |
| CNAG_01047 | 851.5696 | 2.736308 | 0.260722 | 10.49513 | 9.10E-26 | 6.51E-24 |
| CNAG_01045 | 2670.062 | -0.24268 | 0.120219 | -2.01867 | 0.043522 | 0.086412 |
| CNAG_01044 | 1149.048 | 0.330972 | 0.149158 | 2.218936 | 0.026491 | 0.05657 |
| CNAG_01043 | 11855.32 | 0.673947 | 0.190984 | 3.528815 | 0.000417 | 0.001543 |
| CNAG_12483 | 31.93909 | -0.75746 | 0.376734 | -2.01059 | 0.044369 | 0.087784 |
| CNAG_01041 | 269.3645 | 0.426581 | 0.180663 | 2.361201 | 0.018216 | 0.041179 |
| CNAG_01039 | 5474.518 | 0.448671 | 0.184455 | 2.432415 | 0.014999 | 0.034858 |
| CNAG_01037 | 762.9138 | 0.451239 | 0.212878 | 2.11971 | 0.03403 | 0.069999 |
| CNAG_01035 | 7603.36 | 0.7816 | 0.16483 | 4.741868 | 2.12E-06 | 1.38E-05 |
| CNAG_01034 | 71.09951 | 0.684421 | 0.259061 | 2.641926 | 0.008244 | 0.020808 |

| | | | | | | |
|---|---|---|---|---|---|---|
| CNAG_01032 | 394.7294 | 0.794994 | 0.324973 | 2.44634 | 0.014431 | 0.033737 |
| CNAG_01031 | 5077.946 | 2.196739 | 0.209864 | 10.46744 | 1.22E-25 | 8.65E-24 |
| CNAG_01027 | 1222.536 | 0.759537 | 0.125942 | 6.030867 | 1.63E-09 | 2.01E-08 |
| CNAG_01026 | 974.8293 | 0.834537 | 0.160285 | 5.206592 | 1.92E-07 | 1.55E-06 |
| CNAG_01022 | 1033.31 | 0.928076 | 0.154869 | 5.992654 | 2.06E-09 | 2.50E-08 |
| CNAG_01021 | 3293.014 | -0.60363 | 0.134182 | -4.49856 | 6.84E-06 | 3.98E-05 |
| CNAG_01020 | 1818.633 | -0.6964 | 0.233039 | -2.98837 | 0.002805 | 0.008218 |
| CNAG_01018 | 6792.381 | -0.57299 | 0.131119 | -4.36997 | 1.24E-05 | 6.81E-05 |
| CNAG_01010 | 3346.328 | 0.61928 | 0.215751 | 2.870344 | 0.0041 | 0.011364 |
| CNAG_01009 | 1044.253 | -0.55357 | 0.185469 | -2.9847 | 0.002839 | 0.008302 |
| CNAG_01008 | 845.1685 | 0.819034 | 0.133814 | 6.120684 | 9.32E-10 | 1.19E-08 |
| CNAG_01007 | 1753.303 | 1.247057 | 0.181816 | 6.858908 | 6.94E-12 | 1.26E-10 |
| CNAG_01004 | 3127.796 | 0.851083 | 0.182633 | 4.660058 | 3.16E-06 | 1.99E-05 |
| CNAG_01003 | 3317.472 | -0.27851 | 0.110174 | -2.5279 | 0.011475 | 0.027655 |
| CNAG_01001 | 181.6225 | 1.571655 | 0.233408 | 6.733498 | 1.66E-11 | 2.77E-10 |
| CNAG_01000 | 3525.955 | -0.39799 | 0.118482 | -3.35906 | 0.000782 | 0.002671 |
| CNAG_00997 | 813.3189 | -1.28632 | 0.348697 | -3.68894 | 0.000225 | 0.000897 |
| CNAG_00996 | 4335.951 | 0.213025 | 0.107774 | 1.97659 | 0.048088 | 0.094022 |
| CNAG_00995 | 8995.299 | 2.264207 | 0.176367 | 12.83804 | 1.00E-37 | 1.48E-35 |
| CNAG_00994 | 2338.781 | -0.72423 | 0.248589 | -2.91335 | 0.003576 | 0.010108 |
| CNAG_00992 | 12689.84 | -0.52827 | 0.169409 | -3.11834 | 0.001819 | 0.005597 |
| CNAG_00991 | 2246.505 | -0.36669 | 0.11933 | -3.07291 | 0.00212 | 0.006439 |
| CNAG_00990 | 6275.034 | -0.7564 | 0.26723 | -2.83052 | 0.004647 | 0.012703 |
| CNAG_00987 | 1610.519 | -0.60727 | 0.2132 | -2.84837 | 0.004394 | 0.012116 |
| CNAG_00986 | 1006.423 | -0.42531 | 0.145386 | -2.92539 | 0.00344 | 0.00977 |
| CNAG_00984 | 7784.757 | 4.129282 | 0.18335 | 22.52131 | 2.57E-112 | 6.80E-109 |
| CNAG_00982 | 1719.478 | -0.61238 | 0.172394 | -3.55222 | 0.000382 | 0.001428 |
| CNAG_00981 | 25.0946 | 1.45473 | 0.438718 | 3.315861 | 0.000914 | 0.003054 |
| CNAG_12488 | 118.4809 | -1.71338 | 0.358243 | -4.78275 | 1.73E-06 | 1.14E-05 |
| CNAG_00979 | 492.6959 | -1.62161 | 0.239982 | -6.7572 | 1.41E-11 | 2.40E-10 |
| CNAG_00978 | 2501.663 | -1.03281 | 0.272278 | -3.79322 | 0.000149 | 0.000621 |
| CNAG_00976 | 9162.302 | -0.82655 | 0.117441 | -7.03797 | 1.95E-12 | 3.88E-11 |
| CNAG_00975 | 250.1801 | -0.71555 | 0.244148 | -2.93079 | 0.003381 | 0.009626 |
| CNAG_00972 | 2095.117 | -0.7422 | 0.127705 | -5.81184 | 6.18E-09 | 6.68E-08 |
| CNAG_12489 | 322.0428 | -1.09388 | 0.256086 | -4.27155 | 1.94E-05 | 0.000101 |
| CNAG_00968 | 310.8055 | 0.536186 | 0.236516 | 2.267016 | 0.023389 | 0.051044 |
| CNAG_07445 | 32493.45 | -0.3891 | 0.126797 | -3.06868 | 0.00215 | 0.006521 |
| CNAG_00961 | 2504.188 | 1.989484 | 0.185562 | 10.72138 | 8.08E-27 | 6.11E-25 |
| CNAG_12491 | 2237.242 | 3.495378 | 0.290957 | 12.01339 | 3.02E-33 | 3.53E-31 |
| CNAG_12492 | 596.3629 | 0.542971 | 0.172894 | 3.140491 | 0.001687 | 0.005252 |
| CNAG_00938 | 527.536 | 0.514189 | 0.155497 | 3.306756 | 0.000944 | 0.00315 |
| CNAG_00935 | 10735.37 | -0.40526 | 0.196376 | -2.0637 | 0.039046 | 0.078687 |
| CNAG_00933 | 510.4658 | 0.568159 | 0.214904 | 2.643785 | 0.008198 | 0.020707 |
| CNAG_00932 | 1145.775 | 2.868044 | 0.241871 | 11.85776 | 1.96E-32 | 2.20E-30 |
| CNAG_00930 | 12863.68 | -0.81405 | 0.124756 | -6.52515 | 6.79E-11 | 1.03E-09 |
| CNAG_00929 | 1155.627 | -0.56635 | 0.203584 | -2.7819 | 0.005404 | 0.014527 |
| CNAG_00925 | 297.3377 | 0.933497 | 0.167904 | 5.559691 | 2.70E-08 | 2.66E-07 |

| | | | | | | |
|---|---|---|---|---|---|---|
| CNAG_00923 | 620.0048 | 0.950306 | 0.244457 | 3.887407 | 0.000101 | 0.000441 |
| CNAG_00921 | 852.4307 | 1.080286 | 0.208251 | 5.187417 | 2.13E-07 | 1.69E-06 |
| CNAG_00920 | 1182.618 | 0.978094 | 0.226582 | 4.316738 | 1.58E-05 | 8.43E-05 |
| CNAG_00919 | 3222.597 | -2.01026 | 0.421883 | -4.76497 | 1.89E-06 | 1.24E-05 |
| CNAG_00917 | 1274.964 | -0.36465 | 0.161018 | -2.26466 | 0.023534 | 0.051303 |
| CNAG_00915 | 312.272 | 1.890209 | 0.1659 | 11.39367 | 4.50E-30 | 4.20E-28 |
| CNAG_12499 | 128.9628 | -0.72707 | 0.318779 | -2.28081 | 0.02256 | 0.049465 |
| CNAG_12500 | 7.512711 | 2.46614 | 0.831197 | 2.966974 | 0.003007 | 0.008703 |
| CNAG_07450 | 219.9396 | 0.559016 | 0.234114 | 2.387796 | 0.01695 | 0.038646 |
| CNAG_07982 | 964.1729 | -1.3568 | 0.137277 | -9.88363 | 4.90E-23 | 3.04E-21 |
| CNAG_00908 | 37.28126 | 0.812642 | 0.360254 | 2.25575 | 0.024086 | 0.052249 |
| CNAG_00906 | 3379.571 | -0.43099 | 0.148077 | -2.91058 | 0.003608 | 0.010183 |
| CNAG_00905 | 290.9473 | 1.565361 | 0.1604 | 9.759092 | 1.69E-22 | 1.00E-20 |
| CNAG_00904 | 662.2655 | 2.627919 | 0.173703 | 15.12878 | 1.05E-51 | 2.60E-49 |
| CNAG_00903 | 651.6811 | -0.48052 | 0.146201 | -3.2867 | 0.001014 | 0.003359 |
| CNAG_00901 | 360.3833 | 0.726414 | 0.209745 | 3.463313 | 0.000534 | 0.00192 |
| CNAG_00900 | 208.3901 | -0.39627 | 0.189084 | -2.09572 | 0.036107 | 0.073812 |
| CNAG_00899 | 861.3523 | -0.75004 | 0.219105 | -3.42318 | 0.000619 | 0.002183 |
| CNAG_12503 | 22.64817 | 1.036354 | 0.479448 | 2.161559 | 0.030652 | 0.064028 |
| CNAG_00896 | 560.8143 | -1.77247 | 0.316959 | -5.59211 | 2.24E-08 | 2.23E-07 |
| CNAG_12505 | 31.39564 | -3.6998 | 0.754824 | -4.90154 | 9.51E-07 | 6.63E-06 |
| CNAG_12506 | 43.06085 | -2.63612 | 0.478791 | -5.50579 | 3.68E-08 | 3.51E-07 |
| CNAG_00895 | 20602.87 | -4.43032 | 0.832515 | -5.32161 | 1.03E-07 | 8.82E-07 |
| CNAG_00894 | 4504.303 | -0.92187 | 0.27833 | -3.31215 | 0.000926 | 0.003092 |
| CNAG_00886 | 26381.14 | -0.49376 | 0.201031 | -2.45613 | 0.014044 | 0.032919 |
| CNAG_10063 | 2.091282 | 3.392781 | 1.635475 | 2.074492 | 0.038034 | 0.077115 |
| CNAG_00883 | 840.1967 | -1.51658 | 0.292238 | -5.18952 | 2.11E-07 | 1.67E-06 |
| CNAG_12510 | 14.25785 | -1.90039 | 0.662013 | -2.87062 | 0.004097 | 0.011358 |
| CNAG_07453 | 1946.168 | -0.46689 | 0.115772 | -4.0328 | 5.51E-05 | 0.000256 |
| CNAG_00877 | 1204.232 | -0.80774 | 0.223491 | -3.61417 | 0.000301 | 0.001163 |
| CNAG_00876 | 302.8472 | 0.903026 | 0.183949 | 4.909117 | 9.15E-07 | 6.39E-06 |
| CNAG_00873 | 1006.54 | 2.098118 | 0.403669 | 5.197617 | 2.02E-07 | 1.61E-06 |
| CNAG_00870 | 48.29618 | 1.084342 | 0.374295 | 2.897027 | 0.003767 | 0.010562 |
| CNAG_12511 | 43.31428 | 2.866071 | 0.487513 | 5.878966 | 4.13E-09 | 4.64E-08 |
| CNAG_00869 | 624.9839 | -0.60321 | 0.202465 | -2.97931 | 0.002889 | 0.008421 |
| CNAG_00866 | 3115.928 | 1.848901 | 0.199893 | 9.249442 | 2.26E-20 | 1.10E-18 |
| CNAG_07456 | 301.021 | 0.76315 | 0.181316 | 4.208958 | 2.57E-05 | 0.00013 |
| CNAG_07458 | 203.6846 | 1.2102 | 0.220571 | 5.486682 | 4.10E-08 | 3.83E-07 |
| CNAG_00863 | 100.2857 | 0.470646 | 0.239236 | 1.967283 | 0.049151 | 0.095794 |
| CNAG_00862 | 592.4033 | -0.73861 | 0.248267 | -2.97506 | 0.002929 | 0.008514 |
| CNAG_00861 | 145.8064 | -0.72797 | 0.203107 | -3.58415 | 0.000338 | 0.001284 |
| CNAG_00860 | 297.2162 | -0.49336 | 0.222549 | -2.21687 | 0.026632 | 0.056826 |
| CNAG_02554 | 484.9447 | -0.43971 | 0.143545 | -3.06321 | 0.00219 | 0.006626 |
| CNAG_02553 | 137.8463 | -1.93815 | 0.292062 | -6.63608 | 3.22E-11 | 5.14E-10 |
| CNAG_02552 | 321.6267 | 0.555464 | 0.245571 | 2.261932 | 0.023702 | 0.051584 |
| CNAG_02551 | 183.2026 | -1.01492 | 0.309969 | -3.27426 | 0.001059 | 0.00348 |
| CNAG_02550 | 981.2239 | 1.628567 | 0.250165 | 6.509963 | 7.52E-11 | 1.13E-09 |

| | | | | | | |
|---|---|---|---|---|---|---|
| CNAG_02549 | 1079.368 | 0.670289 | 0.198334 | 3.379606 | 0.000726 | 0.002499 |
| CNAG_02548 | 21595.46 | -5.1498 | 1.206536 | -4.26825 | 1.97E-05 | 0.000102 |
| CNAG_02546 | 3238.391 | -0.61125 | 0.179237 | -3.41029 | 0.000649 | 0.002275 |
| CNAG_02545 | 14092.76 | -0.57459 | 0.170827 | -3.36357 | 0.000769 | 0.002631 |
| CNAG_02544 | 64.72639 | 1.298355 | 0.366389 | 3.543648 | 0.000395 | 0.001467 |
| CNAG_02543 | 1199.098 | 0.650297 | 0.145466 | 4.470428 | 7.81E-06 | 4.46E-05 |
| CNAG_02542 | 3656.266 | 0.758947 | 0.203731 | 3.725248 | 0.000195 | 0.000791 |
| CNAG_02540 | 897.2199 | -1.10932 | 0.165631 | -6.69755 | 2.12E-11 | 3.48E-10 |
| CNAG_12515 | 132.8764 | 1.760657 | 0.233197 | 7.550099 | 4.35E-14 | 1.07E-12 |
| CNAG_02537 | 10.62093 | 1.593957 | 0.70904 | 2.248049 | 0.024573 | 0.053131 |
| CNAG_02536 | 897.5579 | -0.65501 | 0.193032 | -3.39325 | 0.000691 | 0.002399 |
| CNAG_02534 | 1102.769 | -0.33429 | 0.143886 | -2.3233 | 0.020163 | 0.044865 |
| CNAG_12517 | 141.4276 | -0.87761 | 0.389031 | -2.25588 | 0.024078 | 0.052246 |
| CNAG_02531 | 338.1798 | 1.070845 | 0.157139 | 6.814632 | 9.45E-12 | 1.67E-10 |
| CNAG_02530 | 3119.827 | -0.30717 | 0.132676 | -2.31521 | 0.020601 | 0.045674 |
| CNAG_02528 | 1283.877 | 0.769717 | 0.186062 | 4.136885 | 3.52E-05 | 0.000172 |
| CNAG_02527 | 3899.208 | 1.367791 | 0.148719 | 9.197136 | 3.68E-20 | 1.70E-18 |
| CNAG_12518 | 70.89163 | 1.031904 | 0.252279 | 4.090322 | 4.31E-05 | 0.000205 |
| CNAG_02526 | 24.51124 | 1.092718 | 0.45416 | 2.406018 | 0.016127 | 0.03708 |
| CNAG_02524 | 714.992 | 0.527936 | 0.259293 | 2.036061 | 0.041744 | 0.083362 |
| CNAG_02523 | 1285.364 | 1.674869 | 0.164442 | 10.18515 | 2.31E-24 | 1.57E-22 |
| CNAG_02520 | 1442.593 | -0.56323 | 0.130034 | -4.33146 | 1.48E-05 | 7.94E-05 |
| CNAG_02519 | 2530.095 | 0.648928 | 0.23478 | 2.76398 | 0.00571 | 0.015241 |
| CNAG_02516 | 581.8195 | 0.501647 | 0.146902 | 3.414831 | 0.000638 | 0.002245 |
| CNAG_07622 | 2032.305 | 0.3081 | 0.120387 | 2.559242 | 0.01049 | 0.025671 |
| CNAG_02512 | 2222.651 | 0.914821 | 0.129538 | 7.06217 | 1.64E-12 | 3.31E-11 |
| CNAG_12522 | 46.66488 | 2.137973 | 0.351815 | 6.076989 | 1.22E-09 | 1.53E-08 |
| CNAG_02510 | 1370.621 | 0.757951 | 0.14379 | 5.27123 | 1.36E-07 | 1.13E-06 |
| CNAG_02509 | 83.07836 | 0.501109 | 0.245976 | 2.037228 | 0.041627 | 0.08315 |
| CNAG_02508 | 532.4794 | -2.17372 | 0.235405 | -9.234 | 2.61E-20 | 1.25E-18 |
| CNAG_02507 | 4399.846 | -0.52606 | 0.168528 | -3.12147 | 0.001799 | 0.005549 |
| CNAG_02506 | 1783.275 | -0.79967 | 0.223899 | -3.57156 | 0.000355 | 0.00134 |
| CNAG_02505 | 1426.216 | 1.728921 | 0.155026 | 11.15245 | 6.97E-29 | 6.02E-27 |
| CNAG_02504 | 651.2643 | -0.9085 | 0.196406 | -4.62561 | 3.73E-06 | 2.31E-05 |
| CNAG_02502 | 2594.279 | -0.68269 | 0.124288 | -5.49278 | 3.96E-08 | 3.73E-07 |
| CNAG_02500 | 19482.61 | 0.489001 | 0.208385 | 2.346626 | 0.018944 | 0.042499 |
| CNAG_02497 | 157.3386 | 1.666218 | 0.208641 | 7.986037 | 1.39E-15 | 4.12E-14 |
| CNAG_02495 | 2050.98 | -0.30065 | 0.135522 | -2.21843 | 0.026525 | 0.056629 |
| CNAG_12523 | 170.9632 | -1.64226 | 0.405014 | -4.05481 | 5.02E-05 | 0.000235 |
| CNAG_02491 | 590.1597 | -1.11185 | 0.168463 | -6.59993 | 4.11E-11 | 6.40E-10 |
| CNAG_02488 | 698.5069 | 1.919247 | 0.33834 | 5.672548 | 1.41E-08 | 1.44E-07 |
| CNAG_02486 | 4892.117 | -0.68075 | 0.250379 | -2.71888 | 0.00655 | 0.017121 |
| CNAG_02485 | 17182.01 | -0.42887 | 0.171392 | -2.50229 | 0.012339 | 0.029453 |
| CNAG_07625 | 2537.67 | -0.27327 | 0.137874 | -1.98206 | 0.047473 | 0.093026 |
| CNAG_12526 | 25.60875 | 1.01378 | 0.425768 | 2.381063 | 0.017263 | 0.03927 |
| CNAG_02482 | 1489.252 | -0.41595 | 0.184162 | -2.25863 | 0.023906 | 0.051944 |
| CNAG_12527 | 22.37138 | 1.510621 | 0.427217 | 3.535956 | 0.000406 | 0.001507 |

| | | | | | | |
|---|---|---|---|---|---|---|
| CNAG_02479 | 268.4159 | -0.85492 | 0.331614 | -2.57806 | 0.009936 | 0.024488 |
| CNAG_02475 | 673.6301 | 0.447124 | 0.125219 | 3.570728 | 0.000356 | 0.001342 |
| CNAG_07626 | 248.8011 | 0.68691 | 0.161087 | 4.264205 | 2.01E-05 | 0.000104 |
| CNAG_02473 | 2201.585 | 1.345487 | 0.164943 | 8.157308 | 3.43E-16 | 1.09E-14 |
| CNAG_12531 | 59.99577 | 1.034245 | 0.26814 | 3.857105 | 0.000115 | 0.000492 |
| CNAG_02471 | 802.2683 | -0.50763 | 0.207662 | -2.4445 | 0.014505 | 0.033881 |
| CNAG_02470 | 1353.816 | 0.464487 | 0.151382 | 3.068306 | 0.002153 | 0.006526 |
| CNAG_02469 | 1150.107 | -0.25433 | 0.121254 | -2.09753 | 0.035947 | 0.073541 |
| CNAG_02463 | 72.43378 | -1.02803 | 0.268489 | -3.82896 | 0.000129 | 0.000545 |
| CNAG_02460 | 627.668 | -1.10161 | 0.398813 | -2.76222 | 0.005741 | 0.015318 |
| CNAG_12535 | 38.14218 | -1.77764 | 0.44839 | -3.9645 | 7.35E-05 | 0.000332 |
| CNAG_02459 | 365.4704 | 0.765285 | 0.344059 | 2.224286 | 0.026129 | 0.055873 |
| CNAG_02458 | 2759.442 | 0.470073 | 0.20402 | 2.304051 | 0.02122 | 0.046876 |
| CNAG_07628 | 10821.46 | -0.46731 | 0.153987 | -3.03475 | 0.002407 | 0.007191 |
| CNAG_07629 | 2754.872 | 0.804082 | 0.180121 | 4.464132 | 8.04E-06 | 4.58E-05 |
| CNAG_02449 | 1324.71 | -0.74566 | 0.110369 | -6.75604 | 1.42E-11 | 2.41E-10 |
| CNAG_12537 | 70.30083 | 1.208229 | 0.263682 | 4.582154 | 4.60E-06 | 2.79E-05 |
| CNAG_02448 | 24.27759 | 0.895426 | 0.430657 | 2.079211 | 0.037598 | 0.076411 |
| CNAG_02444 | 1185.14 | 1.241799 | 0.176269 | 7.044891 | 1.86E-12 | 3.71E-11 |
| CNAG_02443 | 2669.771 | -0.87774 | 0.242408 | -3.62091 | 0.000294 | 0.001136 |
| CNAG_02442 | 429.6388 | -0.66268 | 0.161904 | -4.09307 | 4.26E-05 | 0.000203 |
| CNAG_02441 | 18.57564 | 2.662636 | 0.68131 | 3.90811 | 9.30E-05 | 0.000409 |
| CNAG_02440 | 5948.611 | 0.741794 | 0.110944 | 6.686198 | 2.29E-11 | 3.73E-10 |
| CNAG_02438 | 391.7374 | 1.616148 | 0.181585 | 8.900206 | 5.57E-19 | 2.31E-17 |
| CNAG_12538 | 51.52562 | 2.280461 | 0.38293 | 5.955296 | 2.60E-09 | 3.07E-08 |
| CNAG_02437 | 1063.421 | -0.74026 | 0.15013 | -4.93078 | 8.19E-07 | 5.77E-06 |
| CNAG_12540 | 49.40316 | 1.895029 | 0.316186 | 5.9934 | 2.05E-09 | 2.49E-08 |
| CNAG_02435 | 3826.964 | 0.376244 | 0.155501 | 2.419555 | 0.01554 | 0.035926 |
| CNAG_02433 | 1354.579 | 0.732102 | 0.140659 | 5.204808 | 1.94E-07 | 1.57E-06 |
| CNAG_02430 | 461.4022 | 1.045821 | 0.319373 | 3.274606 | 0.001058 | 0.003479 |
| CNAG_02427 | 901.3469 | 1.232428 | 0.159861 | 7.709354 | 1.26E-14 | 3.28E-13 |
| CNAG_02426 | 4479.382 | -0.40724 | 0.168869 | -2.4116 | 0.015883 | 0.036634 |
| CNAG_02424 | 880.8036 | -0.78329 | 0.143501 | -5.45843 | 4.80E-08 | 4.42E-07 |
| CNAG_02421 | 4073.376 | -0.74448 | 0.236359 | -3.14979 | 0.001634 | 0.005113 |
| CNAG_02418 | 12032.1 | -1.12482 | 0.128482 | -8.75466 | 2.05E-18 | 8.01E-17 |
| CNAG_02417 | 259.3887 | 1.073767 | 0.498812 | 2.152649 | 0.031346 | 0.065238 |
| CNAG_12541 | 148.549 | 0.700784 | 0.266149 | 2.633052 | 0.008462 | 0.021299 |
| CNAG_02416 | 3893.818 | -0.57556 | 0.262499 | -2.19263 | 0.028334 | 0.059847 |
| CNAG_02413 | 4610.474 | -0.38304 | 0.133838 | -2.86195 | 0.00421 | 0.011653 |
| CNAG_02412 | 617.9803 | -1.25174 | 0.210441 | -5.94817 | 2.71E-09 | 3.19E-08 |
| CNAG_12542 | 8.645919 | 1.654296 | 0.743529 | 2.224925 | 0.026086 | 0.055796 |
| CNAG_12543 | 40.42945 | -0.79203 | 0.383236 | -2.06668 | 0.038764 | 0.078178 |
| CNAG_07631 | 2645.862 | 1.572647 | 0.317924 | 4.946618 | 7.55E-07 | 5.38E-06 |
| CNAG_02409 | 884.8568 | 0.415587 | 0.164776 | 2.522137 | 0.011664 | 0.028044 |
| CNAG_02407 | 3119.648 | 0.630741 | 0.193869 | 3.253444 | 0.00114 | 0.003721 |
| CNAG_02406 | 1014.971 | 0.4251 | 0.189291 | 2.245754 | 0.02472 | 0.053347 |
| CNAG_02405 | 920.5734 | -0.36787 | 0.163687 | -2.24741 | 0.024614 | 0.053201 |

| | | | | | | |
|---|---|---|---|---|---|---|
| CNAG_02403 | 2956.793 | -0.88401 | 0.251 | -3.52197 | 0.000428 | 0.001577 |
| CNAG_02400 | 3840.155 | 0.423171 | 0.204713 | 2.067147 | 0.03872 | 0.078149 |
| CNAG_02398 | 1754.49 | 1.671938 | 0.255324 | 6.548295 | 5.82E-11 | 8.88E-10 |
| CNAG_02394 | 733.678 | -0.30674 | 0.15494 | -1.97973 | 0.047733 | 0.093467 |
| CNAG_02389 | 307.0505 | 0.756312 | 0.317967 | 2.378586 | 0.017379 | 0.039482 |
| CNAG_12549 | 57.85145 | 0.656692 | 0.262331 | 2.503297 | 0.012304 | 0.029395 |
| CNAG_02388 | 661.3557 | -0.5505 | 0.203917 | -2.69963 | 0.006942 | 0.017984 |
| CNAG_02387 | 1347.371 | 0.604277 | 0.182137 | 3.317706 | 0.000908 | 0.003035 |
| CNAG_02385 | 681.904 | -0.29786 | 0.146094 | -2.03885 | 0.041465 | 0.082909 |
| CNAG_02383 | 499.025 | -0.51068 | 0.168073 | -3.03844 | 0.002378 | 0.007117 |
| CNAG_02382 | 2973.953 | -0.68062 | 0.191169 | -3.56031 | 0.00037 | 0.001388 |
| CNAG_07986 | 679.0184 | -0.62473 | 0.137783 | -4.53415 | 5.78E-06 | 3.42E-05 |
| CNAG_02378 | 6853.396 | -0.67708 | 0.209878 | -3.22606 | 0.001255 | 0.004057 |
| CNAG_02376 | 1143.599 | -0.98 | 0.193107 | -5.07489 | 3.88E-07 | 2.91E-06 |
| CNAG_02373 | 1666.898 | 0.575831 | 0.189167 | 3.04403 | 0.002334 | 0.007007 |
| CNAG_02372 | 2900.802 | 0.286262 | 0.118262 | 2.420571 | 0.015496 | 0.035846 |
| CNAG_02371 | 376.4102 | -0.54292 | 0.153441 | -3.5383 | 0.000403 | 0.001495 |
| CNAG_07633 | 270.0906 | 0.56295 | 0.167696 | 3.356968 | 0.000788 | 0.002686 |
| CNAG_02366 | 600.4916 | -0.82282 | 0.170351 | -4.83014 | 1.36E-06 | 9.17E-06 |
| CNAG_02365 | 596.9787 | -1.13304 | 0.181511 | -6.24229 | 4.31E-10 | 5.79E-09 |
| CNAG_02362 | 1255.387 | 1.488455 | 0.126791 | 11.73944 | 8.00E-32 | 8.26E-30 |
| CNAG_02361 | 1986.333 | -0.77998 | 0.302885 | -2.57518 | 0.010019 | 0.024654 |
| CNAG_12555 | 19.91922 | -1.76742 | 0.529587 | -3.33736 | 0.000846 | 0.002855 |
| CNAG_02360 | 1937.897 | 0.510059 | 0.130307 | 3.914281 | 9.07E-05 | 0.000401 |
| CNAG_02359 | 34516.57 | -1.06406 | 0.30116 | -3.5332 | 0.000411 | 0.001521 |
| CNAG_02357 | 1092.657 | 0.703143 | 0.255243 | 2.754801 | 0.005873 | 0.015607 |
| CNAG_02355 | 1289.456 | 1.06255 | 0.166837 | 6.3688 | 1.91E-10 | 2.71E-09 |
| CNAG_02354 | 2760.404 | -0.69173 | 0.161888 | -4.2729 | 1.93E-05 | 0.0001 |
| CNAG_02351 | 2461.688 | 1.377846 | 0.132355 | 10.41022 | 2.23E-25 | 1.55E-23 |
| CNAG_02350 | 904.4119 | -0.27358 | 0.134179 | -2.03889 | 0.041461 | 0.082909 |
| CNAG_02348 | 728.7601 | 0.373972 | 0.187957 | 1.989671 | 0.046627 | 0.09164 |
| CNAG_12557 | 4.547671 | 2.184156 | 1.043246 | 2.093615 | 0.036294 | 0.074119 |
| CNAG_02343 | 1635.364 | 0.610846 | 0.123781 | 4.934889 | 8.02E-07 | 5.67E-06 |
| CNAG_02342 | 577.2899 | 0.645677 | 0.151132 | 4.272268 | 1.93E-05 | 0.000101 |
| CNAG_02338 | 16258.8 | -0.59445 | 0.234996 | -2.5296 | 0.011419 | 0.027555 |
| CNAG_02337 | 2934.459 | -2.17658 | 0.669226 | -3.25238 | 0.001144 | 0.003733 |
| CNAG_02335 | 19224.54 | 1.096374 | 0.118666 | 9.239186 | 2.48E-20 | 1.20E-18 |
| CNAG_02334 | 13.97116 | 1.126913 | 0.565847 | 1.991552 | 0.04642 | 0.091278 |
| CNAG_02333 | 493.5205 | -0.45305 | 0.222851 | -2.03298 | 0.042055 | 0.083793 |
| CNAG_02331 | 23681.16 | -1.20159 | 0.301039 | -3.99148 | 6.57E-05 | 0.0003 |
| CNAG_02330 | 27762.45 | -0.81704 | 0.200144 | -4.08228 | 4.46E-05 | 0.000212 |
| CNAG_02325 | 1235.074 | 0.506459 | 0.145027 | 3.492171 | 0.000479 | 0.001738 |
| CNAG_02323 | 5698.284 | -0.65729 | 0.226127 | -2.90673 | 0.003652 | 0.010295 |
| CNAG_12560 | 153.0386 | -1.31931 | 0.351896 | -3.74914 | 0.000177 | 0.00073 |
| CNAG_02322 | 1953.031 | -0.88273 | 0.378911 | -2.32964 | 0.019825 | 0.044213 |
| CNAG_02319 | 597.1772 | -0.81802 | 0.264619 | -3.0913 | 0.001993 | 0.006086 |
| CNAG_02317 | 1108.203 | -0.96521 | 0.140928 | -6.84892 | 7.44E-12 | 1.34E-10 |

| | | | | | | |
|---|---|---|---|---|---|---|
| CNAG_02316 | 1700.197 | -0.66115 | 0.198068 | -3.33798 | 0.000844 | 0.00285 |
| CNAG_02315 | 13695.72 | -1.13208 | 0.252882 | -4.4767 | 7.58E-06 | 4.35E-05 |
| CNAG_12562 | 41.78188 | 1.215694 | 0.321612 | 3.780005 | 0.000157 | 0.00065 |
| CNAG_02313 | 710.6406 | 1.089325 | 0.208998 | 5.212122 | 1.87E-07 | 1.52E-06 |
| CNAG_02312 | 1645.703 | 1.176353 | 0.238444 | 4.933445 | 8.08E-07 | 5.70E-06 |
| CNAG_02309 | 1576.349 | 0.443344 | 0.187125 | 2.369241 | 0.017825 | 0.040382 |
| CNAG_02305 | 491.1276 | -0.44367 | 0.208368 | -2.12927 | 0.033232 | 0.068561 |
| CNAG_02301 | 7518.334 | -1.13874 | 0.182087 | -6.25378 | 4.01E-10 | 5.39E-09 |
| CNAG_02300 | 14987.51 | -0.70894 | 0.125342 | -5.65606 | 1.55E-08 | 1.57E-07 |
| CNAG_02299 | 819.6658 | -0.32986 | 0.154317 | -2.13751 | 0.032556 | 0.067438 |
| CNAG_02298 | 3518.851 | 1.096804 | 0.253206 | 4.331666 | 1.48E-05 | 7.94E-05 |
| CNAG_02297 | 622.944 | 0.741868 | 0.198839 | 3.73099 | 0.000191 | 0.000776 |
| CNAG_02294 | 3631.611 | -0.43218 | 0.17686 | -2.4436 | 0.014542 | 0.033945 |
| CNAG_02293 | 1454.336 | -0.55672 | 0.142706 | -3.90119 | 9.57E-05 | 0.000419 |
| CNAG_02292 | 2852.996 | -0.58143 | 0.164526 | -3.53397 | 0.000409 | 0.001517 |
| CNAG_02291 | 836.5653 | -0.68782 | 0.242668 | -2.8344 | 0.004591 | 0.01258 |
| CNAG_02289 | 821.1812 | 0.78986 | 0.171511 | 4.605313 | 4.12E-06 | 2.54E-05 |
| CNAG_02285 | 19521.86 | -0.47569 | 0.238496 | -1.99453 | 0.046094 | 0.090726 |
| CNAG_02284 | 336.3327 | 0.89418 | 0.173347 | 5.158325 | 2.49E-07 | 1.94E-06 |
| CNAG_02283 | 138.092 | 1.630683 | 0.420544 | 3.877554 | 0.000106 | 0.000456 |
| CNAG_02282 | 2180.947 | 0.90342 | 0.161058 | 5.609285 | 2.03E-08 | 2.03E-07 |
| CNAG_02281 | 972.7496 | -0.41647 | 0.139134 | -2.99328 | 0.00276 | 0.008108 |
| CNAG_07635 | 1625.824 | -1.03776 | 0.176242 | -5.88826 | 3.90E-09 | 4.42E-08 |
| CNAG_07636 | 1616.638 | -0.37502 | 0.140683 | -2.66575 | 0.007682 | 0.019551 |
| CNAG_07637 | 3041.593 | -0.84073 | 0.229014 | -3.6711 | 0.000242 | 0.000953 |
| CNAG_07638 | 1468.265 | 1.385617 | 0.165783 | 8.358 | 6.38E-17 | 2.17E-15 |
| CNAG_07640 | 887.6153 | -0.72025 | 0.238787 | -3.01631 | 0.002559 | 0.007599 |
| CNAG_07641 | 133.6074 | 0.77654 | 0.202953 | 3.826206 | 0.00013 | 0.00055 |
| CNAG_02270 | 1959.924 | -0.45812 | 0.134701 | -3.40097 | 0.000671 | 0.002341 |
| CNAG_02267 | 1417.507 | -0.52022 | 0.163503 | -3.18174 | 0.001464 | 0.004629 |
| CNAG_07164 | 1187.43 | 1.487262 | 0.310965 | 4.782729 | 1.73E-06 | 1.14E-05 |
| CNAG_07163 | 1333.15 | -1.01139 | 0.20207 | -5.00512 | 5.58E-07 | 4.09E-06 |
| CNAG_02266 | 3680.719 | -1.42816 | 0.275262 | -5.18835 | 2.12E-07 | 1.68E-06 |
| CNAG_02264 | 2448.956 | 1.017122 | 0.22392 | 4.542356 | 5.56E-06 | 3.30E-05 |
| CNAG_02263 | 792.2673 | 1.831491 | 0.714111 | 2.564713 | 0.010326 | 0.025317 |
| CNAG_02262 | 3249.859 | 0.591398 | 0.289051 | 2.045998 | 0.040757 | 0.081719 |
| CNAG_02257 | 17292.18 | -0.45639 | 0.176213 | -2.58997 | 0.009598 | 0.023742 |
| CNAG_12570 | 97.68648 | 1.037631 | 0.36622 | 2.833354 | 0.004606 | 0.012608 |
| CNAG_12571 | 281.6466 | 1.180731 | 0.300681 | 3.926859 | 8.61E-05 | 0.000383 |
| CNAG_02240 | 3346.142 | 0.621537 | 0.249705 | 2.489084 | 0.012807 | 0.030423 |
| CNAG_02239 | 7299.192 | 0.654498 | 0.125975 | 5.195448 | 2.04E-07 | 1.63E-06 |
| CNAG_02235 | 544.7813 | -0.62393 | 0.178618 | -3.49308 | 0.000477 | 0.001735 |
| CNAG_02234 | 43410.99 | -1.06252 | 0.226058 | -4.70022 | 2.60E-06 | 1.67E-05 |
| CNAG_02232 | 2079.396 | -0.34295 | 0.135867 | -2.52414 | 0.011598 | 0.02791 |
| CNAG_02230 | 10315.51 | 2.434234 | 0.249616 | 9.751925 | 1.81E-22 | 1.07E-20 |
| CNAG_07646 | 189.8158 | 0.591967 | 0.195698 | 3.024899 | 0.002487 | 0.007412 |
| CNAG_02227 | 450.262 | 0.41965 | 0.1606 | 2.613021 | 0.008975 | 0.022369 |

| | | | | | | |
|---|---|---|---|---|---|---|
| CNAG_02226 | 1101.193 | 2.500566 | 0.194557 | 12.8526 | 8.32E-38 | 1.25E-35 |
| CNAG_02225 | 318.9154 | 1.790607 | 0.21777 | 8.222474 | 1.99E-16 | 6.41E-15 |
| CNAG_02223 | 3935.732 | 0.398157 | 0.166169 | 2.39609 | 0.016571 | 0.037892 |
| CNAG_02220 | 1379 | 0.339934 | 0.167076 | 2.03461 | 0.04189 | 0.08357 |
| CNAG_02219 | 810.3084 | 0.686428 | 0.123497 | 5.558248 | 2.72E-08 | 2.68E-07 |
| CNAG_02217 | 224.1793 | 1.225371 | 0.202226 | 6.059416 | 1.37E-09 | 1.70E-08 |
| CNAG_12573 | 611.94 | 3.047967 | 1.211893 | 2.515046 | 0.011902 | 0.028537 |
| CNAG_02216 | 2475.416 | 0.680595 | 0.21571 | 3.155145 | 0.001604 | 0.00503 |
| CNAG_02212 | 12175.79 | -0.65012 | 0.111752 | -5.81753 | 5.97E-09 | 6.48E-08 |
| CNAG_02211 | 662.1022 | 1.296059 | 0.214246 | 6.049404 | 1.45E-09 | 1.80E-08 |
| CNAG_02209 | 8240.496 | -0.90882 | 0.192045 | -4.73231 | 2.22E-06 | 1.44E-05 |
| CNAG_02207 | 674.6024 | 0.503331 | 0.14592 | 3.44935 | 0.000562 | 0.00201 |
| CNAG_02206 | 16.1647 | 3.43735 | 0.660119 | 5.207167 | 1.92E-07 | 1.55E-06 |
| CNAG_02205 | 534.2471 | 0.359369 | 0.176126 | 2.04041 | 0.041309 | 0.082702 |
| CNAG_02204 | 1233.913 | -0.66023 | 0.150818 | -4.37764 | 1.20E-05 | 6.60E-05 |
| CNAG_02203 | 1106.119 | -0.70304 | 0.119574 | -5.87957 | 4.11E-09 | 4.63E-08 |
| CNAG_02202 | 3602.464 | -0.83133 | 0.151253 | -5.49628 | 3.88E-08 | 3.67E-07 |
| CNAG_02198 | 458.8196 | -0.3897 | 0.136164 | -2.86201 | 0.00421 | 0.011653 |
| CNAG_02197 | 2119.243 | -0.54993 | 0.193954 | -2.83534 | 0.004578 | 0.012551 |
| CNAG_02196 | 1953.402 | -0.39548 | 0.165892 | -2.38397 | 0.017127 | 0.039006 |
| CNAG_02195 | 396.2707 | -0.99207 | 0.189481 | -5.23573 | 1.64E-07 | 1.35E-06 |
| CNAG_02194 | 974.6854 | -1.40477 | 0.177356 | -7.92065 | 2.36E-15 | 6.71E-14 |
| CNAG_02193 | 721.5058 | -1.35278 | 0.209581 | -6.45469 | 1.08E-10 | 1.61E-09 |
| CNAG_02192 | 29.38148 | 1.560358 | 0.407816 | 3.82613 | 0.00013 | 0.00055 |
| CNAG_02190 | 722.6417 | -0.51721 | 0.256827 | -2.01384 | 0.044027 | 0.087235 |
| CNAG_02189 | 3808.713 | 0.938761 | 0.203807 | 4.606119 | 4.10E-06 | 2.53E-05 |
| CNAG_12579 | 15.31994 | 1.499026 | 0.523853 | 2.86154 | 0.004216 | 0.011664 |
| CNAG_02188 | 839.7304 | 1.137473 | 0.182328 | 6.238604 | 4.41E-10 | 5.89E-09 |
| CNAG_02185 | 841.7553 | -0.36919 | 0.16497 | -2.2379 | 0.025228 | 0.054237 |
| CNAG_12580 | 12.37219 | 1.4508 | 0.592661 | 2.447943 | 0.014367 | 0.033617 |
| CNAG_02182 | 5848.313 | 2.066435 | 0.125927 | 16.40977 | 1.63E-60 | 6.81E-58 |
| CNAG_02176 | 3254.198 | 0.72452 | 0.27815 | 2.604778 | 0.009193 | 0.022864 |
| CNAG_02174 | 5247.117 | -0.34648 | 0.108352 | -3.19777 | 0.001385 | 0.004414 |
| CNAG_02173 | 801.128 | 0.597045 | 0.198697 | 3.004804 | 0.002658 | 0.007847 |
| CNAG_02170 | 170.5962 | 1.038706 | 0.254619 | 4.079446 | 4.51E-05 | 0.000214 |
| CNAG_02169 | 140.87 | 2.128318 | 0.274546 | 7.752139 | 9.04E-15 | 2.41E-13 |
| CNAG_02168 | 1697.359 | 0.870665 | 0.280523 | 3.103723 | 0.001911 | 0.005856 |
| CNAG_02167 | 1127.518 | 0.539811 | 0.252516 | 2.137734 | 0.032538 | 0.067436 |
| CNAG_02165 | 1922.333 | 0.330121 | 0.137562 | 2.399802 | 0.016404 | 0.037585 |
| CNAG_02164 | 3123.693 | 1.597887 | 0.14208 | 11.24642 | 2.41E-29 | 2.15E-27 |
| CNAG_02163 | 378.8315 | -0.86499 | 0.155751 | -5.55371 | 2.80E-08 | 2.74E-07 |
| CNAG_02159 | 470.0899 | 0.498814 | 0.229996 | 2.168791 | 0.030099 | 0.063009 |
| CNAG_07647 | 2223.3 | 1.00748 | 0.206681 | 4.874571 | 1.09E-06 | 7.46E-06 |
| CNAG_02157 | 482.8242 | -0.55206 | 0.185483 | -2.97631 | 0.002917 | 0.008482 |
| CNAG_02156 | 518.0397 | 1.381812 | 0.173092 | 7.983109 | 1.43E-15 | 4.20E-14 |
| CNAG_02154 | 852.1378 | -0.60835 | 0.245599 | -2.47701 | 0.013249 | 0.031258 |
| CNAG_02153 | 2942.735 | -0.44479 | 0.106784 | -4.1653 | 3.11E-05 | 0.000154 |

| | | | | | | |
|---|---|---|---|---|---|---|
| CNAG_02147 | 234.1744 | -1.06861 | 0.196776 | -5.43056 | 5.62E-08 | 5.10E-07 |
| CNAG_02145 | 1685.283 | -0.88335 | 0.206427 | -4.27925 | 1.88E-05 | 9.80E-05 |
| CNAG_02144 | 38037.47 | -0.70306 | 0.125428 | -5.60534 | 2.08E-08 | 2.07E-07 |
| CNAG_02143 | 619.6033 | -0.78088 | 0.163118 | -4.78719 | 1.69E-06 | 1.12E-05 |
| CNAG_02141 | 335.3481 | -0.63937 | 0.21474 | -2.97743 | 0.002907 | 0.008454 |
| CNAG_02140 | 3549.763 | -1.25784 | 0.182142 | -6.90584 | 4.99E-12 | 9.20E-11 |
| CNAG_02139 | 3431.901 | 0.495153 | 0.128128 | 3.864502 | 0.000111 | 0.000478 |
| CNAG_12586 | 77.31952 | 0.629676 | 0.258671 | 2.434275 | 0.014922 | 0.034699 |
| CNAG_02136 | 2226.857 | -0.61599 | 0.217941 | -2.82641 | 0.004707 | 0.012858 |
| CNAG_02134 | 3343.11 | -1.06952 | 0.167358 | -6.39059 | 1.65E-10 | 2.38E-09 |
| CNAG_02131 | 739.5518 | -0.65474 | 0.191468 | -3.41958 | 0.000627 | 0.002209 |
| CNAG_02130 | 4271.552 | -0.54453 | 0.180531 | -3.01625 | 0.002559 | 0.007599 |
| CNAG_02129 | 30364.43 | 1.887286 | 0.163443 | 11.54704 | 7.64E-31 | 7.50E-29 |
| CNAG_02128 | 9814.781 | -0.64884 | 0.218375 | -2.97122 | 0.002966 | 0.008611 |
| CNAG_02126 | 1069.11 | 0.532002 | 0.189101 | 2.813326 | 0.004903 | 0.013334 |
| CNAG_02125 | 66.10956 | 0.697692 | 0.268891 | 2.594705 | 0.009467 | 0.023457 |
| CNAG_02124 | 938.2335 | 0.466996 | 0.149349 | 3.126868 | 0.001767 | 0.005465 |
| CNAG_02123 | 1294.12 | -0.71186 | 0.140313 | -5.07342 | 3.91E-07 | 2.93E-06 |
| CNAG_02118 | 114.3025 | 1.730991 | 0.313543 | 5.520741 | 3.38E-08 | 3.26E-07 |
| CNAG_02117 | 992.4913 | 0.343537 | 0.157792 | 2.177145 | 0.02947 | 0.0619 |
| CNAG_02115 | 1577.534 | -0.97388 | 0.201394 | -4.83568 | 1.33E-06 | 8.94E-06 |
| CNAG_12591 | 100.447 | -1.12023 | 0.245982 | -4.55411 | 5.26E-06 | 3.15E-05 |
| CNAG_02114 | 2306.002 | 0.505791 | 0.220448 | 2.294383 | 0.021769 | 0.047915 |
| CNAG_02109 | 2028.998 | 0.571505 | 0.132679 | 4.307422 | 1.65E-05 | 8.75E-05 |
| CNAG_12593 | 281.4918 | -2.1725 | 0.362965 | -5.98543 | 2.16E-09 | 2.60E-08 |
| CNAG_02103 | 857.0484 | 1.095099 | 0.293136 | 3.735811 | 0.000187 | 0.000764 |
| CNAG_02102 | 91.24779 | 1.52971 | 0.51701 | 2.958765 | 0.003089 | 0.008886 |
| CNAG_02101 | 495.2417 | -0.49852 | 0.17907 | -2.78396 | 0.00537 | 0.014455 |
| CNAG_02100 | 13001.24 | -0.70819 | 0.118642 | -5.96909 | 2.39E-09 | 2.86E-08 |
| CNAG_02099 | 12655.11 | -0.37137 | 0.187122 | -1.98465 | 0.047183 | 0.092549 |
| CNAG_02096 | 340.6817 | -0.49185 | 0.194067 | -2.53443 | 0.011263 | 0.027227 |
| CNAG_12594 | 37.18875 | 1.38309 | 0.385457 | 3.588184 | 0.000333 | 0.001269 |
| CNAG_02094 | 3965.946 | 0.404212 | 0.117475 | 3.440829 | 0.00058 | 0.002064 |
| CNAG_02093 | 1203.056 | 1.337419 | 0.167598 | 7.979932 | 1.46E-15 | 4.26E-14 |
| CNAG_02089 | 853.5958 | 0.790814 | 0.161697 | 4.890714 | 1.00E-06 | 6.94E-06 |
| CNAG_02086 | 2108.471 | -0.75115 | 0.120979 | -6.20897 | 5.33E-10 | 7.00E-09 |
| CNAG_02084 | 3328.523 | -1.15566 | 0.131173 | -8.8102 | 1.25E-18 | 4.96E-17 |
| CNAG_02083 | 1241.225 | -1.35107 | 0.21722 | -6.21981 | 4.98E-10 | 6.57E-09 |
| CNAG_12598 | 47.7155 | -0.88209 | 0.3981 | -2.21574 | 0.026709 | 0.056929 |
| CNAG_02082 | 508.6001 | -1.14643 | 0.305162 | -3.75679 | 0.000172 | 0.00071 |
| CNAG_02079 | 12334.02 | 0.641063 | 0.160103 | 4.004058 | 6.23E-05 | 0.000285 |
| CNAG_02076 | 2099.775 | -0.54988 | 0.143191 | -3.84021 | 0.000123 | 0.000524 |
| CNAG_02075 | 904.1707 | -0.63219 | 0.142145 | -4.44748 | 8.69E-06 | 4.91E-05 |
| CNAG_02073 | 1616.291 | 1.164868 | 0.202249 | 5.759571 | 8.43E-09 | 8.95E-08 |
| CNAG_02072 | 1624.603 | 1.12569 | 0.192497 | 5.847816 | 4.98E-09 | 5.51E-08 |
| CNAG_02070 | 10052.03 | 1.855101 | 0.256894 | 7.221265 | 5.15E-13 | 1.11E-11 |
| CNAG_02069 | 2402.619 | 1.470178 | 0.243883 | 6.028212 | 1.66E-09 | 2.04E-08 |

| | | | | | | |
|---|---|---|---|---|---|---|
| CNAG_02067 | 1955.987 | -0.68248 | 0.126232 | -5.40656 | 6.42E-08 | 5.79E-07 |
| CNAG_02061 | 293.1793 | -0.56551 | 0.152979 | -3.69666 | 0.000218 | 0.000873 |
| CNAG_02060 | 1519.093 | -2.73908 | 0.192216 | -14.25 | 4.48E-46 | 1.02E-43 |
| CNAG_12600 | 1157.295 | 2.15482 | 0.188181 | 11.45077 | 2.33E-30 | 2.23E-28 |
| CNAG_02058 | 123.0927 | 1.640953 | 0.276088 | 5.943592 | 2.79E-09 | 3.26E-08 |
| CNAG_02055 | 1183.595 | 0.967533 | 0.158259 | 6.113619 | 9.74E-10 | 1.24E-08 |
| CNAG_02054 | 583.2989 | -0.46519 | 0.139085 | -3.34464 | 0.000824 | 0.002791 |
| CNAG_02051 | 1506.924 | -1.07552 | 0.159012 | -6.76375 | 1.34E-11 | 2.31E-10 |
| CNAG_02049 | 21.92868 | 1.003993 | 0.501763 | 2.000932 | 0.0454 | 0.089515 |
| CNAG_02048 | 664.7199 | -0.99806 | 0.147121 | -6.78393 | 1.17E-11 | 2.02E-10 |
| CNAG_02047 | 1248.582 | 1.561835 | 0.134066 | 11.64976 | 2.30E-31 | 2.29E-29 |
| CNAG_02045 | 156.01 | -0.58986 | 0.213071 | -2.76836 | 0.005634 | 0.015088 |
| CNAG_02043 | 220.3889 | -0.64541 | 0.212056 | -3.04359 | 0.002338 | 0.007015 |
| CNAG_02041 | 1975.727 | 0.775523 | 0.198753 | 3.901949 | 9.54E-05 | 0.000418 |
| CNAG_02040 | 322.1702 | -0.67054 | 0.29167 | -2.29896 | 0.021507 | 0.047419 |
| CNAG_02038 | 2851.61 | 0.664251 | 0.119789 | 5.545195 | 2.94E-08 | 2.86E-07 |
| CNAG_02037 | 2442.728 | -0.57288 | 0.189527 | -3.02266 | 0.002506 | 0.007454 |
| CNAG_02034 | 782.8573 | -0.80238 | 0.134025 | -5.98678 | 2.14E-09 | 2.58E-08 |
| CNAG_12605 | 116.0697 | 0.616949 | 0.228317 | 2.702159 | 0.006889 | 0.017877 |
| CNAG_12606 | 34.4284 | 0.983077 | 0.396271 | 2.480822 | 0.013108 | 0.031008 |
| CNAG_02029 | 1628.714 | 0.422914 | 0.1481 | 2.855598 | 0.004296 | 0.011868 |
| CNAG_02028 | 3740.211 | -0.7119 | 0.164666 | -4.32327 | 1.54E-05 | 8.20E-05 |
| CNAG_02026 | 361.3028 | 1.344719 | 0.203746 | 6.599967 | 4.11E-11 | 6.40E-10 |
| CNAG_02025 | 1323.112 | -0.73499 | 0.145819 | -5.04042 | 4.65E-07 | 3.45E-06 |
| CNAG_02024 | 1286.003 | -1.12631 | 0.21884 | -5.14672 | 2.65E-07 | 2.05E-06 |
| CNAG_02023 | 299.6301 | 1.184342 | 0.219092 | 5.405678 | 6.46E-08 | 5.81E-07 |
| CNAG_02020 | 1528.197 | -0.66147 | 0.206004 | -3.21096 | 0.001323 | 0.004249 |
| CNAG_06525 | 138.0058 | 0.601639 | 0.258934 | 2.323527 | 0.020151 | 0.044851 |
| CNAG_06533 | 2277.985 | 0.720263 | 0.131415 | 5.480817 | 4.23E-08 | 3.94E-07 |
| CNAG_06534 | 5648.096 | -0.80497 | 0.127873 | -6.29504 | 3.07E-10 | 4.24E-09 |
| CNAG_06535 | 2104.88 | -0.73556 | 0.136688 | -5.38131 | 7.39E-08 | 6.51E-07 |
| CNAG_06539 | 45.95181 | 0.830966 | 0.311354 | 2.668878 | 0.007611 | 0.019414 |
| CNAG_06541 | 6250.652 | 1.707607 | 0.174558 | 9.782438 | 1.34E-22 | 8.06E-21 |
| CNAG_06544 | 12488.7 | -0.7722 | 0.228847 | -3.3743 | 0.00074 | 0.002544 |
| CNAG_06545 | 3911.633 | -1.22205 | 0.113451 | -10.7716 | 4.69E-27 | 3.62E-25 |
| CNAG_06549 | 893.5401 | -0.35191 | 0.169815 | -2.07234 | 0.038233 | 0.077422 |
| CNAG_06553 | 1053.669 | 0.61535 | 0.192002 | 3.204914 | 0.001351 | 0.004322 |
| CNAG_06554 | 835.9303 | -0.74284 | 0.163211 | -4.55144 | 5.33E-06 | 3.18E-05 |
| CNAG_06555 | 1174.69 | -0.81714 | 0.125048 | -6.53458 | 6.38E-11 | 9.71E-10 |
| CNAG_06556 | 818.9171 | -0.82859 | 0.140606 | -5.89302 | 3.79E-09 | 4.30E-08 |
| CNAG_06558 | 1275.21 | -0.47237 | 0.139226 | -3.39282 | 0.000692 | 0.002399 |
| CNAG_06559 | 843.6 | 1.114792 | 0.213059 | 5.232324 | 1.67E-07 | 1.37E-06 |
| CNAG_06560 | 1065.645 | 0.709916 | 0.239188 | 2.968021 | 0.002997 | 0.008679 |
| CNAG_06563 | 5061.062 | -0.66883 | 0.190611 | -3.5089 | 0.00045 | 0.001645 |
| CNAG_06567 | 1757.241 | 1.152917 | 0.273175 | 4.220437 | 2.44E-05 | 0.000124 |
| CNAG_06568 | 1734.082 | -0.53015 | 0.235773 | -2.24856 | 0.024541 | 0.053076 |
| CNAG_07659 | 2609.657 | 0.625338 | 0.144375 | 4.331348 | 1.48E-05 | 7.94E-05 |

| | | | | | | |
|---|---|---|---|---|---|---|
| CNAG_07660 | 12015.25 | -0.67051 | 0.188647 | -3.55431 | 0.000379 | 0.001418 |
| CNAG_06573 | 1543.178 | 0.600739 | 0.203123 | 2.957515 | 0.003101 | 0.008919 |
| CNAG_06574 | 1063.601 | 1.335655 | 0.219069 | 6.09697 | 1.08E-09 | 1.37E-08 |
| CNAG_12619 | 48.48895 | 0.815496 | 0.354561 | 2.300014 | 0.021447 | 0.0473 |
| CNAG_06576 | 12160.93 | 2.675031 | 0.226822 | 11.79353 | 4.22E-32 | 4.59E-30 |
| CNAG_06577 | 2208.615 | 2.990652 | 0.22218 | 13.46048 | 2.67E-41 | 4.82E-39 |
| CNAG_06578 | 6823.399 | 0.982989 | 0.22966 | 4.280192 | 1.87E-05 | 9.77E-05 |
| CNAG_07661 | 384.9758 | 0.874437 | 0.290028 | 3.015008 | 0.00257 | 0.007622 |
| CNAG_12622 | 72.69106 | 1.794986 | 0.285891 | 6.278558 | 3.42E-10 | 4.67E-09 |
| CNAG_06583 | 265.9428 | -0.95048 | 0.180969 | -5.25216 | 1.50E-07 | 1.24E-06 |
| CNAG_06588 | 831.724 | 0.529223 | 0.221113 | 2.393448 | 0.016691 | 0.038155 |
| CNAG_06589 | 1257.602 | 0.758542 | 0.207712 | 3.651897 | 0.00026 | 0.001021 |
| CNAG_06593 | 535.9083 | 0.431694 | 0.155236 | 2.780888 | 0.005421 | 0.014562 |
| CNAG_06598 | 90.20163 | -3.15859 | 0.503254 | -6.27633 | 3.47E-10 | 4.72E-09 |
| CNAG_06599 | 792.8167 | -0.40856 | 0.172888 | -2.36317 | 0.018119 | 0.040996 |
| CNAG_06600 | 4545.657 | 0.413603 | 0.132438 | 3.123003 | 0.00179 | 0.005531 |
| CNAG_06602 | 3161.653 | 1.734175 | 0.260136 | 6.666421 | 2.62E-11 | 4.22E-10 |
| CNAG_06604 | 816.0168 | -0.54644 | 0.192538 | -2.83809 | 0.004539 | 0.012466 |
| CNAG_06605 | 49064.45 | -0.89769 | 0.250977 | -3.57677 | 0.000348 | 0.001316 |
| CNAG_06606 | 2626.282 | 0.582829 | 0.199251 | 2.925104 | 0.003443 | 0.009775 |
| CNAG_06608 | 1496.219 | -0.4993 | 0.15234 | -3.27753 | 0.001047 | 0.00345 |
| CNAG_12629 | 209.9164 | 1.275261 | 0.280421 | 4.547674 | 5.42E-06 | 3.23E-05 |
| CNAG_06609 | 37.87022 | 1.463612 | 0.396972 | 3.686938 | 0.000227 | 0.000903 |
| CNAG_06610 | 235.6492 | 0.590453 | 0.217108 | 2.719626 | 0.006536 | 0.017094 |
| CNAG_06614 | 811.4306 | 0.741527 | 0.183122 | 4.049362 | 5.14E-05 | 0.00024 |
| CNAG_12632 | 80.80964 | 0.94997 | 0.297568 | 3.192444 | 0.001411 | 0.004487 |
| CNAG_12633 | 31.6983 | 0.821325 | 0.364524 | 2.253145 | 0.02425 | 0.052547 |
| CNAG_06616 | 1340.753 | 1.135175 | 0.190516 | 5.958419 | 2.55E-09 | 3.02E-08 |
| CNAG_07664 | 2125.752 | -0.42966 | 0.114183 | -3.76288 | 0.000168 | 0.000694 |
| CNAG_06621 | 5299.953 | -0.75897 | 0.194696 | -3.89824 | 9.69E-05 | 0.000424 |
| CNAG_06623 | 903.034 | 0.720448 | 0.211569 | 3.405266 | 0.000661 | 0.002314 |
| CNAG_06625 | 4335.564 | 0.64537 | 0.231165 | 2.791816 | 0.005241 | 0.014156 |
| CNAG_06626 | 3986.662 | -0.52071 | 0.155529 | -3.348 | 0.000814 | 0.002769 |
| CNAG_06627 | 2628.882 | -0.40628 | 0.109654 | -3.70509 | 0.000211 | 0.00085 |
| CNAG_12636 | 21.96318 | 1.443999 | 0.447046 | 3.230093 | 0.001238 | 0.004005 |
| CNAG_06629 | 31.28149 | 1.298195 | 0.388715 | 3.339708 | 0.000839 | 0.002835 |
| CNAG_06630 | 4019.899 | -0.75112 | 0.158109 | -4.75067 | 2.03E-06 | 1.32E-05 |
| CNAG_06631 | 2254.419 | -0.31943 | 0.15182 | -2.10402 | 0.035377 | 0.072487 |
| CNAG_06632 | 2278.841 | 0.573575 | 0.122035 | 4.70007 | 2.60E-06 | 1.67E-05 |
| CNAG_06633 | 37055.71 | -0.91589 | 0.255578 | -3.58358 | 0.000339 | 0.001285 |
| CNAG_06634 | 917.6936 | -0.24668 | 0.115034 | -2.14438 | 0.032002 | 0.066446 |
| CNAG_06635 | 3687.13 | -0.45754 | 0.206448 | -2.21627 | 0.026673 | 0.056883 |
| CNAG_06636 | 219.9576 | 1.117521 | 0.183271 | 6.097641 | 1.08E-09 | 1.36E-08 |
| CNAG_06638 | 10232.22 | -0.99084 | 0.130593 | -7.58725 | 3.27E-14 | 8.11E-13 |
| CNAG_06641 | 1826.298 | -0.97267 | 0.20117 | -4.83506 | 1.33E-06 | 8.96E-06 |
| CNAG_06643 | 548.5152 | -0.64514 | 0.143731 | -4.48856 | 7.17E-06 | 4.14E-05 |
| CNAG_06644 | 3536.07 | -0.83989 | 0.195817 | -4.28916 | 1.79E-05 | 9.42E-05 |

| | | | | | | |
|---|---|---|---|---|---|---|
| CNAG_06645 | 2241.223 | -0.96616 | 0.175548 | -5.50369 | 3.72E-08 | 3.55E-07 |
| CNAG_06648 | 2286.01 | -0.35217 | 0.136002 | -2.58948 | 0.009612 | 0.023764 |
| CNAG_06650 | 209.1591 | 0.764112 | 0.177819 | 4.29714 | 1.73E-05 | 9.14E-05 |
| CNAG_06652 | 206.044 | 0.679326 | 0.219349 | 3.097011 | 0.001955 | 0.005972 |
| CNAG_06654 | 549.8244 | -0.5877 | 0.205685 | -2.85727 | 0.004273 | 0.011814 |
| CNAG_06655 | 525.1391 | 0.795347 | 0.171242 | 4.644573 | 3.41E-06 | 2.13E-05 |
| CNAG_06656 | 1613.589 | -0.92023 | 0.221174 | -4.16067 | 3.17E-05 | 0.000157 |
| CNAG_06657 | 653.0325 | 0.44412 | 0.214451 | 2.070965 | 0.038362 | 0.077615 |
| CNAG_06658 | 503.8034 | 1.297803 | 0.160117 | 8.105365 | 5.26E-16 | 1.63E-14 |
| CNAG_06659 | 794.0542 | 0.662032 | 0.127867 | 5.177501 | 2.25E-07 | 1.78E-06 |
| CNAG_06661 | 1712.887 | -0.57354 | 0.142863 | -4.01461 | 5.95E-05 | 0.000274 |
| CNAG_12644 | 24.68528 | 1.232236 | 0.485793 | 2.536548 | 0.011195 | 0.027096 |
| CNAG_06663 | 6462.542 | -1.39128 | 0.315524 | -4.40945 | 1.04E-05 | 5.75E-05 |
| CNAG_06664 | 1153.357 | 0.539464 | 0.144082 | 3.744147 | 0.000181 | 0.000742 |
| CNAG_06666 | 7406.11 | 1.051079 | 0.150597 | 6.979431 | 2.96E-12 | 5.63E-11 |
| CNAG_06668 | 5591.367 | 1.863303 | 0.174523 | 10.67657 | 1.31E-26 | 9.64E-25 |
| CNAG_06669 | 1859.369 | -0.7123 | 0.246056 | -2.89488 | 0.003793 | 0.010612 |
| CNAG_06671 | 2649.705 | 0.457074 | 0.182584 | 2.503357 | 0.012302 | 0.029395 |
| CNAG_12647 | 150.0606 | 0.874639 | 0.193926 | 4.510172 | 6.48E-06 | 3.80E-05 |
| CNAG_06672 | 22.12899 | 1.380918 | 0.546772 | 2.525583 | 0.011551 | 0.027804 |
| CNAG_06673 | 3097.601 | 0.558278 | 0.126087 | 4.427709 | 9.52E-06 | 5.35E-05 |
| CNAG_06674 | 669.1995 | -0.908 | 0.203043 | -4.47197 | 7.75E-06 | 4.43E-05 |
| CNAG_06679 | 3161.354 | -0.73326 | 0.138215 | -5.3052 | 1.13E-07 | 9.54E-07 |
| CNAG_07177 | 2840.555 | -1.2644 | 0.294167 | -4.29825 | 1.72E-05 | 9.10E-05 |
| CNAG_06684 | 467.4622 | -0.91706 | 0.182924 | -5.01335 | 5.35E-07 | 3.93E-06 |
| CNAG_06685 | 2315.012 | -0.3629 | 0.117041 | -3.1006 | 0.001931 | 0.005911 |
| CNAG_06690 | 2691.246 | -0.71598 | 0.173603 | -4.12425 | 3.72E-05 | 0.000181 |
| CNAG_06692 | 806.4776 | -0.52069 | 0.203741 | -2.55566 | 0.010599 | 0.025866 |
| CNAG_06693 | 689.4349 | 1.16279 | 0.156682 | 7.421348 | 1.16E-13 | 2.65E-12 |
| CNAG_06694 | 640.5801 | 0.620171 | 0.193456 | 3.20575 | 0.001347 | 0.004311 |
| CNAG_06695 | 486.9776 | 1.187783 | 0.17021 | 6.978325 | 2.99E-12 | 5.66E-11 |
| CNAG_07990 | 995.296 | -1.61292 | 0.156755 | -10.2894 | 7.87E-25 | 5.43E-23 |
| CNAG_12653 | 14.37475 | -1.45651 | 0.52211 | -2.78966 | 0.005276 | 0.014227 |
| CNAG_06698 | 792.8623 | 0.993662 | 0.130697 | 7.602782 | 2.90E-14 | 7.24E-13 |
| CNAG_06699 | 88531.13 | -0.61364 | 0.125812 | -4.87747 | 1.07E-06 | 7.37E-06 |
| CNAG_05671 | 896.2535 | 0.689252 | 0.130446 | 5.28382 | 1.27E-07 | 1.06E-06 |
| CNAG_05672 | 2264.693 | -0.44684 | 0.160575 | -2.78275 | 0.00539 | 0.014501 |
| CNAG_05673 | 1422.148 | 0.320183 | 0.159204 | 2.011151 | 0.044309 | 0.087736 |
| CNAG_07666 | 21.37283 | 1.226775 | 0.443843 | 2.763987 | 0.00571 | 0.015241 |
| CNAG_05676 | 1061.783 | 2.833818 | 0.153334 | 18.48139 | 2.92E-76 | 1.36E-73 |
| CNAG_07668 | 1097.274 | 0.593368 | 0.146648 | 4.04619 | 5.21E-05 | 0.000243 |
| CNAG_05682 | 6975.13 | 0.912235 | 0.141348 | 6.453842 | 1.09E-10 | 1.61E-09 |
| CNAG_05683 | 829.4698 | 3.093695 | 0.321528 | 9.621847 | 6.47E-22 | 3.64E-20 |
| CNAG_05684 | 2305.541 | 0.76066 | 0.16607 | 4.580356 | 4.64E-06 | 2.81E-05 |
| CNAG_12663 | 227.7349 | 1.596839 | 0.333036 | 4.794796 | 1.63E-06 | 1.08E-05 |
| CNAG_05686 | 1161.365 | 1.228228 | 0.195009 | 6.298318 | 3.01E-10 | 4.16E-09 |
| CNAG_05689 | 1397.393 | -0.24997 | 0.120443 | -2.0754 | 0.03795 | 0.076965 |

| | | | | | | |
|---|---|---|---|---|---|---|
| CNAG_05694 | 4523.308 | -0.48749 | 0.122691 | -3.97333 | 7.09E-05 | 0.000321 |
| CNAG_12666 | 45.40922 | 0.839539 | 0.332929 | 2.521672 | 0.01168 | 0.028073 |
| CNAG_05697 | 574.0339 | 0.619471 | 0.170213 | 3.639382 | 0.000273 | 0.001067 |
| CNAG_05699 | 726.5972 | -0.93414 | 0.227174 | -4.112 | 3.92E-05 | 0.000189 |
| CNAG_05700 | 2037.641 | -0.31649 | 0.135425 | -2.33698 | 0.01944 | 0.043513 |
| CNAG_12667 | 129.5731 | 0.763325 | 0.37503 | 2.035367 | 0.041814 | 0.083481 |
| CNAG_05702 | 1207.022 | -0.29547 | 0.131517 | -2.24661 | 0.024665 | 0.053242 |
| CNAG_05705 | 428.3794 | -0.72621 | 0.184681 | -3.93224 | 8.42E-05 | 0.000375 |
| CNAG_05707 | 978.2395 | 0.705181 | 0.170151 | 4.144451 | 3.41E-05 | 0.000167 |
| CNAG_05711 | 711.4776 | -0.56565 | 0.221305 | -2.55597 | 0.010589 | 0.025858 |
| CNAG_05718 | 577.7874 | 1.379094 | 0.161117 | 8.559566 | 1.13E-17 | 4.13E-16 |
| CNAG_05719 | 1190.015 | -0.87392 | 0.145573 | -6.00329 | 1.93E-09 | 2.35E-08 |
| CNAG_05723 | 709.8298 | 1.508511 | 0.154864 | 9.740904 | 2.02E-22 | 1.17E-20 |
| CNAG_05724 | 996.8716 | 0.789852 | 0.177502 | 4.449822 | 8.59E-06 | 4.87E-05 |
| CNAG_05725 | 49620.6 | -0.82967 | 0.141401 | -5.86749 | 4.42E-09 | 4.94E-08 |
| CNAG_05726 | 1613.877 | -0.64483 | 0.173649 | -3.71344 | 0.000204 | 0.000825 |
| CNAG_12671 | 30.61619 | 1.467149 | 0.389592 | 3.76586 | 0.000166 | 0.000687 |
| CNAG_05732 | 2714.287 | 0.793934 | 0.255064 | 3.112689 | 0.001854 | 0.005697 |
| CNAG_05734 | 2274.185 | 0.36934 | 0.173361 | 2.130467 | 0.033133 | 0.068383 |
| CNAG_05735 | 223.4558 | 1.26619 | 0.202438 | 6.254703 | 3.98E-10 | 5.36E-09 |
| CNAG_05736 | 67.64624 | 1.019388 | 0.29672 | 3.435521 | 0.000591 | 0.002101 |
| CNAG_05737 | 1784.313 | 0.575939 | 0.155895 | 3.694406 | 0.00022 | 0.00088 |
| CNAG_05738 | 108.4428 | 1.85424 | 0.378501 | 4.898901 | 9.64E-07 | 6.71E-06 |
| CNAG_05739 | 115.6057 | 1.530752 | 0.276034 | 5.545522 | 2.93E-08 | 2.86E-07 |
| CNAG_12673 | 66.88003 | 1.034155 | 0.308656 | 3.350508 | 0.000807 | 0.002746 |
| CNAG_05742 | 2550.557 | -1.29775 | 0.170482 | -7.61224 | 2.69E-14 | 6.77E-13 |
| CNAG_05745 | 3890.283 | -0.50092 | 0.197301 | -2.53887 | 0.011121 | 0.02695 |
| CNAG_05746 | 124.5189 | 0.525657 | 0.251515 | 2.089963 | 0.036621 | 0.07469 |
| CNAG_05747 | 996.7486 | 0.312099 | 0.119997 | 2.600883 | 0.009298 | 0.023082 |
| CNAG_05748 | 1175.795 | -0.47112 | 0.21953 | -2.14605 | 0.031869 | 0.066238 |
| CNAG_05749 | 684.0578 | -0.62797 | 0.215976 | -2.90757 | 0.003642 | 0.010271 |
| CNAG_05750 | 71367.37 | -0.45615 | 0.14269 | -3.1968 | 0.00139 | 0.004426 |
| CNAG_05751 | 1110.443 | -0.45907 | 0.152036 | -3.0195 | 0.002532 | 0.007524 |
| CNAG_05752 | 1793.976 | -1.24042 | 0.182757 | -6.78728 | 1.14E-11 | 1.98E-10 |
| CNAG_05753 | 10381.83 | -0.57151 | 0.110066 | -5.1924 | 2.08E-07 | 1.65E-06 |
| CNAG_07671 | 582.2324 | 0.512644 | 0.225678 | 2.271567 | 0.023113 | 0.050551 |
| CNAG_07672 | 1142.64 | -0.64871 | 0.167783 | -3.86637 | 0.00011 | 0.000475 |
| CNAG_05755 | 1925.592 | -1.22071 | 0.21922 | -5.56845 | 2.57E-08 | 2.55E-07 |
| CNAG_05756 | 1403.154 | -0.65303 | 0.157346 | -4.15028 | 3.32E-05 | 0.000163 |
| CNAG_07673 | 21.33842 | 1.344967 | 0.530589 | 2.534856 | 0.011249 | 0.027203 |
| CNAG_05757 | 518.5253 | -0.78016 | 0.228475 | -3.41465 | 0.000639 | 0.002245 |
| CNAG_05759 | 11603.57 | -0.31532 | 0.138909 | -2.26996 | 0.02321 | 0.050722 |
| CNAG_12676 | 197.1311 | 1.054742 | 0.243727 | 4.327553 | 1.51E-05 | 8.05E-05 |
| CNAG_05762 | 44869.22 | -0.96207 | 0.276061 | -3.485 | 0.000492 | 0.001785 |
| CNAG_05763 | 247.2849 | -0.71018 | 0.166867 | -4.25595 | 2.08E-05 | 0.000107 |
| CNAG_05764 | 2738.582 | -1.04641 | 0.227749 | -4.59458 | 4.34E-06 | 2.66E-05 |
| CNAG_05768 | 2381.426 | 0.396641 | 0.12536 | 3.16401 | 0.001556 | 0.004891 |

| | | | | | | |
|---|---|---|---|---|---|---|
| CNAG_05770 | 6812.55 | 0.608717 | 0.181931 | 3.345869 | 0.00082 | 0.002783 |
| CNAG_05771 | 1756.242 | 0.988803 | 0.293755 | 3.366085 | 0.000762 | 0.002609 |
| CNAG_05772 | 1024.101 | 0.302545 | 0.133382 | 2.268252 | 0.023314 | 0.050907 |
| CNAG_05775 | 1023.196 | 0.477029 | 0.168837 | 2.825376 | 0.004723 | 0.012895 |
| CNAG_05777 | 366.6272 | -0.80624 | 0.223936 | -3.60034 | 0.000318 | 0.001215 |
| CNAG_05778 | 573.8333 | 0.632098 | 0.152443 | 4.146454 | 3.38E-05 | 0.000166 |
| CNAG_05782 | 3080.889 | -0.47253 | 0.141633 | -3.33631 | 0.000849 | 0.002863 |
| CNAG_05784 | 480.3418 | 0.340863 | 0.155031 | 2.198677 | 0.027901 | 0.059104 |
| CNAG_12679 | 29.63195 | -1.04188 | 0.400575 | -2.60097 | 0.009296 | 0.023082 |
| CNAG_05786 | 408.9752 | 0.337818 | 0.170637 | 1.979744 | 0.047732 | 0.093467 |
| CNAG_05787 | 2180.14 | -0.4049 | 0.182108 | -2.22344 | 0.026186 | 0.05598 |
| CNAG_05789 | 1680.004 | -0.76997 | 0.168413 | -4.57192 | 4.83E-06 | 2.92E-05 |
| CNAG_05792 | 7339.643 | 0.706654 | 0.196541 | 3.595458 | 0.000324 | 0.001237 |
| CNAG_07675 | 1068.365 | 0.566502 | 0.210656 | 2.689229 | 0.007162 | 0.018482 |
| CNAG_05798 | 988.2102 | -0.40085 | 0.158135 | -2.53485 | 0.011249 | 0.027203 |
| CNAG_05799 | 6904.03 | 0.576675 | 0.184694 | 3.122331 | 0.001794 | 0.005539 |
| CNAG_05800 | 22237.37 | -0.99061 | 0.240608 | -4.1171 | 3.84E-05 | 0.000186 |
| CNAG_05801 | 332.5277 | -1.10036 | 0.163395 | -6.73439 | 1.65E-11 | 2.76E-10 |
| CNAG_05803 | 643.5944 | -0.51298 | 0.147442 | -3.47921 | 0.000503 | 0.001823 |
| CNAG_05805 | 917.2634 | -0.55627 | 0.184622 | -3.01304 | 0.002586 | 0.007663 |
| CNAG_05807 | 508.105 | 0.734728 | 0.147911 | 4.967367 | 6.79E-07 | 4.89E-06 |
| CNAG_05810 | 468.2518 | -0.87237 | 0.187415 | -4.65475 | 3.24E-06 | 2.04E-05 |
| CNAG_05811 | 780.2214 | -0.75793 | 0.230807 | -3.28381 | 0.001024 | 0.003391 |
| CNAG_05813 | 702.9794 | -1.23314 | 0.247486 | -4.98269 | 6.27E-07 | 4.54E-06 |
| CNAG_05814 | 29874.78 | -0.9916 | 0.269806 | -3.67525 | 0.000238 | 0.000941 |
| CNAG_05815 | 205.4164 | 0.860898 | 0.217996 | 3.949149 | 7.84E-05 | 0.000352 |
| CNAG_05816 | 1208.274 | -0.53204 | 0.123949 | -4.2924 | 1.77E-05 | 9.30E-05 |
| CNAG_05818 | 1532.758 | 0.80636 | 0.234691 | 3.435841 | 0.000591 | 0.002099 |
| CNAG_05819 | 766.1738 | -0.59128 | 0.270004 | -2.18991 | 0.028531 | 0.060214 |
| CNAG_05820 | 704.8476 | -0.77479 | 0.303418 | -2.55355 | 0.010663 | 0.025983 |
| CNAG_05823 | 352.8805 | -0.58374 | 0.163508 | -3.57007 | 0.000357 | 0.001344 |
| CNAG_05824 | 791.2992 | 0.61726 | 0.134428 | 4.59176 | 4.40E-06 | 2.69E-05 |
| CNAG_05825 | 3277.826 | -0.66435 | 0.166415 | -3.99216 | 6.55E-05 | 0.000299 |
| CNAG_05827 | 4510.908 | -0.71601 | 0.212379 | -3.37136 | 0.000748 | 0.00257 |
| CNAG_05833 | 78.3389 | 0.658734 | 0.284273 | 2.317256 | 0.02049 | 0.045478 |
| CNAG_05836 | 1288.602 | 0.64645 | 0.130571 | 4.950956 | 7.38E-07 | 5.28E-06 |
| CNAG_05837 | 5.521661 | 2.800809 | 0.992255 | 2.82267 | 0.004763 | 0.012991 |
| CNAG_05839 | 8361.748 | -1.12102 | 0.241319 | -4.64539 | 3.39E-06 | 2.13E-05 |
| CNAG_05840 | 783.1102 | -0.32574 | 0.135141 | -2.41035 | 0.015937 | 0.036735 |
| CNAG_05847 | 8089.914 | -0.4488 | 0.151534 | -2.9617 | 0.003059 | 0.008818 |
| CNAG_05848 | 5485.102 | -0.59038 | 0.126268 | -4.67563 | 2.93E-06 | 1.86E-05 |
| CNAG_07676 | 12657.43 | -0.56323 | 0.129356 | -4.35413 | 1.34E-05 | 7.28E-05 |
| CNAG_07677 | 529.5817 | 1.189864 | 0.159346 | 7.467154 | 8.19E-14 | 1.92E-12 |
| CNAG_05853 | 443.8658 | -1.59704 | 0.184213 | -8.66956 | 4.34E-18 | 1.63E-16 |
| CNAG_05854 | 553.2839 | -0.58773 | 0.195487 | -3.00651 | 0.002643 | 0.007809 |
| CNAG_07678 | 131.2789 | 0.883691 | 0.231815 | 3.812058 | 0.000138 | 0.000581 |
| CNAG_07679 | 1685.851 | -0.88812 | 0.170516 | -5.20842 | 1.90E-07 | 1.54E-06 |

| | | | | | | |
|---|---|---|---|---|---|---|
| CNAG_05858 | 836.1699 | -0.72571 | 0.191135 | -3.79683 | 0.000147 | 0.000614 |
| CNAG_05859 | 751.6144 | -0.43151 | 0.192183 | -2.2453 | 0.024749 | 0.053395 |
| CNAG_05860 | 897.5911 | 1.016498 | 0.19682 | 5.164606 | 2.41E-07 | 1.89E-06 |
| CNAG_05861 | 179.3079 | 1.508648 | 0.288728 | 5.22516 | 1.74E-07 | 1.42E-06 |
| CNAG_05862 | 820.3352 | 1.136888 | 0.162209 | 7.008765 | 2.40E-12 | 4.69E-11 |
| CNAG_05863 | 767.6634 | -1.51959 | 0.179971 | -8.44354 | 3.08E-17 | 1.07E-15 |
| CNAG_05864 | 4189.695 | 1.915438 | 0.153813 | 12.45306 | 1.35E-35 | 1.81E-33 |
| CNAG_05866 | 471.2296 | 0.889058 | 0.252185 | 3.525422 | 0.000423 | 0.00156 |
| CNAG_05867 | 336.8493 | -1.8867 | 0.274465 | -6.87411 | 6.24E-12 | 1.14E-10 |
| CNAG_05870 | 546.9335 | -0.97659 | 0.308152 | -3.16918 | 0.001529 | 0.004818 |
| CNAG_05871 | 1394.684 | 0.912337 | 0.298125 | 3.060248 | 0.002212 | 0.006687 |
| CNAG_05872 | 4322.131 | 2.002612 | 0.252981 | 7.916049 | 2.45E-15 | 6.93E-14 |
| CNAG_05873 | 197.8644 | 1.374271 | 0.271012 | 5.070891 | 3.96E-07 | 2.97E-06 |
| CNAG_05874 | 708.0415 | -0.37048 | 0.179746 | -2.0611 | 0.039293 | 0.079044 |
| CNAG_05875 | 1217.385 | -1.1973 | 0.21029 | -5.69357 | 1.24E-08 | 1.28E-07 |
| CNAG_05876 | 293.8776 | -1.22022 | 0.174677 | -6.98556 | 2.84E-12 | 5.43E-11 |
| CNAG_05881 | 3465.473 | -0.37568 | 0.184729 | -2.03369 | 0.041982 | 0.083673 |
| CNAG_07680 | 728.8998 | -0.70509 | 0.271752 | -2.59459 | 0.00947 | 0.023458 |
| CNAG_07681 | 1149.835 | -0.4782 | 0.14869 | -3.21606 | 0.0013 | 0.004188 |
| CNAG_05884 | 5745.553 | -0.42657 | 0.18584 | -2.29539 | 0.021711 | 0.047801 |
| CNAG_05886 | 2645.593 | -0.40348 | 0.197778 | -2.04004 | 0.041346 | 0.082755 |
| CNAG_05887 | 1861.739 | -0.42454 | 0.127569 | -3.3279 | 0.000875 | 0.00294 |
| CNAG_05888 | 3156.855 | -0.57475 | 0.292893 | -1.96233 | 0.049724 | 0.096801 |
| CNAG_05889 | 1647.405 | 0.377942 | 0.12086 | 3.127092 | 0.001765 | 0.005463 |
| CNAG_05891 | 518.9264 | 0.798258 | 0.173759 | 4.594057 | 4.35E-06 | 2.66E-05 |
| CNAG_05892 | 3305.484 | -0.80977 | 0.208518 | -3.88343 | 0.000103 | 0.000447 |
| CNAG_05894 | 2973.847 | 0.849778 | 0.235857 | 3.602929 | 0.000315 | 0.001205 |
| CNAG_05896 | 1214.002 | -0.75022 | 0.148972 | -5.03598 | 4.75E-07 | 3.52E-06 |
| CNAG_05899 | 3777.029 | -1.27359 | 0.225855 | -5.63896 | 1.71E-08 | 1.72E-07 |
| CNAG_05900 | 8672.836 | -0.62063 | 0.102147 | -6.07585 | 1.23E-09 | 1.54E-08 |
| CNAG_05901 | 507.3845 | -0.51971 | 0.243134 | -2.13756 | 0.032553 | 0.067438 |
| CNAG_12695 | 66.51062 | -1.33985 | 0.374097 | -3.58156 | 0.000342 | 0.001294 |
| CNAG_05904 | 25813.45 | -1.07073 | 0.303401 | -3.52909 | 0.000417 | 0.001543 |
| CNAG_05905 | 1239.052 | -0.2829 | 0.142486 | -1.98543 | 0.047096 | 0.092425 |
| CNAG_05907 | 20057.24 | -0.757 | 0.143247 | -5.28461 | 1.26E-07 | 1.05E-06 |
| CNAG_05909 | 16315.97 | -1.47587 | 0.22785 | -6.47736 | 9.33E-11 | 1.40E-09 |
| CNAG_05910 | 154.2908 | 0.457663 | 0.202003 | 2.265627 | 0.023474 | 0.051215 |
| CNAG_05911 | 84.0884 | -0.97455 | 0.468508 | -2.08011 | 0.037516 | 0.076282 |
| CNAG_12698 | 186.6623 | -1.63071 | 0.278022 | -5.86539 | 4.48E-09 | 5.00E-08 |
| CNAG_05912 | 879.8823 | 0.614772 | 0.204829 | 3.001396 | 0.002687 | 0.007922 |
| CNAG_05913 | 3922.837 | -0.79648 | 0.147586 | -5.39668 | 6.79E-08 | 6.07E-07 |
| CNAG_05914 | 500.67 | -1.81918 | 0.18357 | -9.91001 | 3.77E-23 | 2.36E-21 |
| CNAG_05915 | 662.662 | 1.956508 | 0.289378 | 6.761077 | 1.37E-11 | 2.34E-10 |
| CNAG_05917 | 1385.218 | -0.45389 | 0.151992 | -2.98624 | 0.002824 | 0.008263 |
| CNAG_05919 | 1076.041 | -0.65868 | 0.183029 | -3.59877 | 0.00032 | 0.001222 |
| CNAG_05920 | 543.2724 | -0.44754 | 0.142221 | -3.14681 | 0.001651 | 0.005161 |
| CNAG_05921 | 1165.947 | 0.663886 | 0.141997 | 4.675343 | 2.93E-06 | 1.86E-05 |

| | | | | | | |
|---|---|---|---|---|---|---|
| CNAG_12699 | 157.8095 | 1.442113 | 0.227758 | 6.331765 | 2.42E-10 | 3.40E-09 |
| CNAG_12700 | 25.56107 | 1.411432 | 0.469284 | 3.00763 | 0.002633 | 0.007783 |
| CNAG_05924 | 84.62874 | 1.190869 | 0.249182 | 4.779107 | 1.76E-06 | 1.16E-05 |
| CNAG_05926 | 2051.95 | -0.54632 | 0.148411 | -3.68111 | 0.000232 | 0.000922 |
| CNAG_05927 | 1388.556 | -0.33707 | 0.114508 | -2.94362 | 0.003244 | 0.009272 |
| CNAG_05930 | 1651.716 | 0.41121 | 0.122085 | 3.368226 | 0.000757 | 0.002594 |
| CNAG_05935 | 3426.141 | -0.85002 | 0.289261 | -2.93859 | 0.003297 | 0.00941 |
| CNAG_05937 | 1007.262 | 0.483434 | 0.204902 | 2.359339 | 0.018308 | 0.041339 |
| CNAG_05938 | 50.3443 | 0.861619 | 0.331746 | 2.597224 | 0.009398 | 0.023322 |
| CNAG_05939 | 1280.762 | 3.534858 | 0.191091 | 18.49827 | 2.13E-76 | 1.06E-73 |
| CNAG_12702 | 35.72721 | 1.508245 | 0.359325 | 4.197446 | 2.70E-05 | 0.000135 |
| CNAG_05940 | 4212.066 | 1.66149 | 0.214 | 7.763958 | 8.23E-15 | 2.20E-13 |
| CNAG_12703 | 138.8485 | -0.68096 | 0.288618 | -2.35938 | 0.018305 | 0.041339 |
| CNAG_05941 | 178.7865 | -0.64876 | 0.237896 | -2.72707 | 0.00639 | 0.016791 |
| CNAG_12704 | 619.4443 | 0.570927 | 0.273423 | 2.088074 | 0.036791 | 0.074998 |
| CNAG_07683 | 328.2446 | 1.151944 | 0.412935 | 2.789651 | 0.005276 | 0.014227 |
| CNAG_07684 | 228.6168 | 0.474767 | 0.229057 | 2.0727 | 0.0382 | 0.077374 |
| CNAG_07687 | 92.05715 | 0.934988 | 0.22779 | 4.104601 | 4.05E-05 | 0.000195 |
| CNAG_07688 | 1814.93 | 0.575288 | 0.15819 | 3.636697 | 0.000276 | 0.001077 |
| CNAG_07689 | 3681.849 | -0.51135 | 0.114849 | -4.45239 | 8.49E-06 | 4.81E-05 |
| CNAG_07690 | 312.3993 | 1.025034 | 0.412898 | 2.482533 | 0.013045 | 0.030887 |
| CNAG_12707 | 15.69795 | 2.483605 | 0.562785 | 4.413061 | 1.02E-05 | 5.68E-05 |
| CNAG_07188 | 222.9442 | 0.660622 | 0.195784 | 3.374232 | 0.00074 | 0.002544 |
| CNAG_07693 | 12.81441 | 1.461383 | 0.639618 | 2.284777 | 0.022326 | 0.048992 |
| CNAG_07701 | 573.3986 | -0.48943 | 0.176384 | -2.7748 | 0.005524 | 0.014823 |
| CNAG_07702 | 187.639 | -0.91709 | 0.199425 | -4.59864 | 4.25E-06 | 2.62E-05 |
| CNAG_05963 | 1246.938 | 0.395515 | 0.185263 | 2.134877 | 0.032771 | 0.067795 |
| CNAG_05965 | 680.4448 | 0.59292 | 0.15259 | 3.885708 | 0.000102 | 0.000444 |
| CNAG_05968 | 293.7265 | -0.79369 | 0.197232 | -4.02416 | 5.72E-05 | 0.000264 |
| CNAG_05972 | 46.99 | -1.62195 | 0.367798 | -4.40991 | 1.03E-05 | 5.75E-05 |
| CNAG_05973 | 244.3749 | 0.456523 | 0.218728 | 2.087173 | 0.036873 | 0.075145 |
| CNAG_05976 | 4151.618 | -0.63666 | 0.14397 | -4.42216 | 9.77E-06 | 5.47E-05 |
| CNAG_05977 | 6505.447 | 1.100081 | 0.318192 | 3.457284 | 0.000546 | 0.001958 |
| CNAG_05978 | 7853.24 | -0.4633 | 0.10487 | -4.4179 | 9.97E-06 | 5.58E-05 |
| CNAG_05979 | 946.052 | -0.79063 | 0.162036 | -4.87935 | 1.06E-06 | 7.32E-06 |
| CNAG_05980 | 3739.746 | -0.79182 | 0.205809 | -3.84736 | 0.000119 | 0.00051 |
| CNAG_05981 | 447.1017 | -1.04977 | 0.201901 | -5.19945 | 2.00E-07 | 1.61E-06 |
| CNAG_05982 | 127.3196 | 0.830734 | 0.24478 | 3.393805 | 0.000689 | 0.002395 |
| CNAG_06944 | 87.07363 | -0.96067 | 0.273846 | -3.50806 | 0.000451 | 0.001649 |
| CNAG_03084 | 194.9222 | 1.232774 | 0.228628 | 5.392056 | 6.97E-08 | 6.18E-07 |
| CNAG_03086 | 986.6133 | -0.51267 | 0.153152 | -3.34745 | 0.000816 | 0.002772 |
| CNAG_03091 | 9.678931 | 1.491658 | 0.747642 | 1.995149 | 0.046027 | 0.090616 |
| CNAG_07708 | 91.28168 | 1.265009 | 0.286401 | 4.416908 | 1.00E-05 | 5.59E-05 |
| CNAG_03097 | 461.6247 | -0.34729 | 0.137433 | -2.527 | 0.011504 | 0.027718 |
| CNAG_03098 | 2024.649 | -0.66249 | 0.219066 | -3.02414 | 0.002493 | 0.00742 |
| CNAG_03100 | 1611.869 | 0.329997 | 0.132577 | 2.489094 | 0.012807 | 0.030423 |
| CNAG_03106 | 1626.576 | -1.13585 | 0.177764 | -6.38967 | 1.66E-10 | 2.39E-09 |

| | | | | | | |
|---|---|---|---|---|---|---|
| CNAG_03107 | 16.54807 | 1.93974 | 0.523749 | 3.703568 | 0.000213 | 0.000854 |
| CNAG_12716 | 42.75924 | 0.855477 | 0.319103 | 2.680886 | 0.007343 | 0.018845 |
| CNAG_03108 | 956.3275 | 0.771746 | 0.143056 | 5.394712 | 6.86E-08 | 6.11E-07 |
| CNAG_03109 | 5951.02 | 0.596946 | 0.110909 | 5.382293 | 7.35E-08 | 6.49E-07 |
| CNAG_03110 | 5006.545 | 1.34442 | 0.171267 | 7.849841 | 4.17E-15 | 1.15E-13 |
| CNAG_03113 | 9172.344 | 1.380596 | 0.219988 | 6.275774 | 3.48E-10 | 4.73E-09 |
| CNAG_03114 | 4130.271 | 1.660831 | 0.211104 | 7.867355 | 3.62E-15 | 1.01E-13 |
| CNAG_03115 | 883.0503 | 0.504531 | 0.129882 | 3.88453 | 0.000103 | 0.000445 |
| CNAG_12720 | 73.50889 | 0.765119 | 0.314101 | 2.435898 | 0.014855 | 0.034564 |
| CNAG_03116 | 2015.574 | -0.42261 | 0.125535 | -3.36648 | 0.000761 | 0.002608 |
| CNAG_03122 | 318.9179 | -1.12432 | 0.312017 | -3.60339 | 0.000314 | 0.001204 |
| CNAG_03125 | 420.212 | -0.82926 | 0.304004 | -2.7278 | 0.006376 | 0.01677 |
| CNAG_03127 | 42700.78 | -1.2065 | 0.308441 | -3.91162 | 9.17E-05 | 0.000404 |
| CNAG_03134 | 308.1252 | -0.41525 | 0.188476 | -2.20319 | 0.027581 | 0.058529 |
| CNAG_03135 | 1163.408 | 0.568546 | 0.226348 | 2.51182 | 0.012011 | 0.028756 |
| CNAG_03136 | 2622.319 | -0.4516 | 0.172372 | -2.61991 | 0.008795 | 0.02197 |
| CNAG_12722 | 42.528 | 0.796191 | 0.322393 | 2.469624 | 0.013526 | 0.031825 |
| CNAG_03138 | 1889.003 | 0.702133 | 0.20625 | 3.404286 | 0.000663 | 0.002319 |
| CNAG_03139 | 6653.178 | -0.24349 | 0.105263 | -2.31315 | 0.020715 | 0.045884 |
| CNAG_03140 | 13083.5 | -0.24181 | 0.115971 | -2.08509 | 0.037061 | 0.075432 |
| CNAG_03141 | 195.1698 | -0.53811 | 0.266613 | -2.01831 | 0.043559 | 0.086466 |
| CNAG_03142 | 18126 | 1.722636 | 0.230645 | 7.468762 | 8.10E-14 | 1.90E-12 |
| CNAG_03143 | 16821.56 | 2.206082 | 0.29854 | 7.389562 | 1.47E-13 | 3.30E-12 |
| CNAG_03145 | 44.9838 | 0.831118 | 0.292516 | 2.841272 | 0.004493 | 0.01235 |
| CNAG_03148 | 2031.341 | -0.84556 | 0.326459 | -2.5901 | 0.009595 | 0.023742 |
| CNAG_03150 | 967.2869 | -0.49803 | 0.143156 | -3.47896 | 0.000503 | 0.001824 |
| CNAG_03154 | 626.2224 | 0.775359 | 0.214631 | 3.612513 | 0.000303 | 0.00117 |
| CNAG_03159 | 945.1001 | -0.49758 | 0.205004 | -2.4272 | 0.015216 | 0.035322 |
| CNAG_03160 | 473.466 | 0.934022 | 0.35009 | 2.667944 | 0.007632 | 0.019455 |
| CNAG_03162 | 1476.38 | 0.932629 | 0.232968 | 4.003253 | 6.25E-05 | 0.000286 |
| CNAG_03163 | 601.7208 | 0.683728 | 0.166243 | 4.112828 | 3.91E-05 | 0.000189 |
| CNAG_03165 | 2427.872 | -0.80428 | 0.179184 | -4.48858 | 7.17E-06 | 4.14E-05 |
| CNAG_03168 | 9085.733 | -1.30395 | 0.157779 | -8.26438 | 1.40E-16 | 4.65E-15 |
| CNAG_12730 | 15.94837 | 1.306289 | 0.554717 | 2.354875 | 0.018529 | 0.041732 |
| CNAG_03170 | 611.1578 | -0.4091 | 0.156388 | -2.61596 | 0.008898 | 0.022206 |
| CNAG_03171 | 2270.114 | 0.354436 | 0.128414 | 2.760097 | 0.005778 | 0.015403 |
| CNAG_03172 | 500.719 | 0.429305 | 0.155547 | 2.759971 | 0.005781 | 0.015403 |
| CNAG_03173 | 2003.1 | 0.659773 | 0.180066 | 3.664062 | 0.000248 | 0.000978 |
| CNAG_03176 | 6628.778 | 2.114982 | 0.138234 | 15.30002 | 7.64E-53 | 2.02E-50 |
| CNAG_12732 | 73.3114 | 0.840403 | 0.289204 | 2.905918 | 0.003662 | 0.010318 |
| CNAG_12733 | 156.7534 | 0.377571 | 0.187907 | 2.009349 | 0.0445 | 0.08796 |
| CNAG_03178 | 1587.853 | 0.648737 | 0.29446 | 2.203139 | 0.027585 | 0.058529 |
| CNAG_12734 | 311.2131 | 0.962388 | 0.214888 | 4.478556 | 7.51E-06 | 4.32E-05 |
| CNAG_03181 | 37.58068 | 1.257834 | 0.358212 | 3.51142 | 0.000446 | 0.001632 |
| CNAG_03183 | 1468.786 | -0.95334 | 0.215543 | -4.42297 | 9.74E-06 | 5.46E-05 |
| CNAG_03184 | 1480.246 | -1.39642 | 0.187977 | -7.42869 | 1.10E-13 | 2.52E-12 |
| CNAG_03185 | 1167.406 | 0.745977 | 0.229609 | 3.248903 | 0.001159 | 0.003773 |

| | | | | | | |
|---|---|---|---|---|---|---|
| CNAG_03186 | 2497.081 | -0.53104 | 0.22784 | -2.33075 | 0.019767 | 0.04412 |
| CNAG_03187 | 497.6416 | 1.26592 | 0.308754 | 4.100094 | 4.13E-05 | 0.000198 |
| CNAG_03189 | 1496.506 | -0.40133 | 0.167877 | -2.39062 | 0.01682 | 0.038406 |
| CNAG_03193 | 43.31226 | 2.39601 | 0.453972 | 5.277881 | 1.31E-07 | 1.09E-06 |
| CNAG_03195 | 222.8374 | 1.285436 | 0.166126 | 7.737721 | 1.01E-14 | 2.65E-13 |
| CNAG_03196 | 2510.271 | -1.00366 | 0.210351 | -4.77136 | 1.83E-06 | 1.20E-05 |
| CNAG_03197 | 172.349 | 0.506107 | 0.230588 | 2.194849 | 0.028174 | 0.059557 |
| CNAG_03198 | 46401 | -1.07352 | 0.247175 | -4.34317 | 1.40E-05 | 7.60E-05 |
| CNAG_07717 | 4518.096 | 0.459414 | 0.185433 | 2.477521 | 0.01323 | 0.031241 |
| CNAG_03202 | 2188.555 | 0.348692 | 0.136493 | 2.554642 | 0.01063 | 0.025917 |
| CNAG_03204 | 491.303 | 0.985813 | 0.181737 | 5.424389 | 5.82E-08 | 5.27E-07 |
| CNAG_03205 | 1478.291 | -1.00239 | 0.160788 | -6.23421 | 4.54E-10 | 6.02E-09 |
| CNAG_03206 | 670.7081 | -0.82691 | 0.18691 | -4.42413 | 9.68E-06 | 5.43E-05 |
| CNAG_03211 | 48.76455 | 0.692569 | 0.333985 | 2.073649 | 0.038112 | 0.077235 |
| CNAG_03212 | 680.1161 | -1.46457 | 0.244993 | -5.978 | 2.26E-09 | 2.71E-08 |
| CNAG_03213 | 4390.977 | 0.936178 | 0.207511 | 4.511456 | 6.44E-06 | 3.78E-05 |
| CNAG_03215 | 352.0165 | 1.108621 | 0.165341 | 6.705051 | 2.01E-11 | 3.33E-10 |
| CNAG_07719 | 8083.101 | 0.630485 | 0.118349 | 5.327358 | 9.97E-08 | 8.57E-07 |
| CNAG_03221 | 9105.899 | -1.01772 | 0.262984 | -3.86989 | 0.000109 | 0.000469 |
| CNAG_03226 | 3516.585 | -1.69422 | 0.263592 | -6.42743 | 1.30E-10 | 1.90E-09 |
| CNAG_12740 | 54.05898 | -1.47756 | 0.409138 | -3.6114 | 0.000305 | 0.001174 |
| CNAG_03230 | 160.9057 | 2.396291 | 0.215687 | 11.11004 | 1.12E-28 | 9.28E-27 |
| CNAG_03232 | 1884.784 | 1.346777 | 0.1646 | 8.182121 | 2.79E-16 | 8.90E-15 |
| CNAG_03233 | 146.6599 | 0.669894 | 0.238723 | 2.806161 | 0.005014 | 0.013606 |
| CNAG_03234 | 96.22928 | 1.355778 | 0.252439 | 5.370721 | 7.84E-08 | 6.89E-07 |
| CNAG_03238 | 155.1646 | 0.553139 | 0.253455 | 2.182392 | 0.029081 | 0.061228 |
| CNAG_03239 | 597.5709 | -1.11764 | 0.176651 | -6.32682 | 2.50E-10 | 3.49E-09 |
| CNAG_03240 | 3185.605 | 0.53258 | 0.184463 | 2.887186 | 0.003887 | 0.010839 |
| CNAG_03242 | 335.2912 | 0.405952 | 0.162082 | 2.504612 | 0.012259 | 0.029313 |
| CNAG_03246 | 1294.117 | -0.35095 | 0.127118 | -2.76082 | 0.005766 | 0.015374 |
| CNAG_03247 | 3325.49 | 0.492898 | 0.224399 | 2.196531 | 0.028054 | 0.059365 |
| CNAG_03248 | 3014.735 | 0.310761 | 0.11469 | 2.709582 | 0.006737 | 0.017545 |
| CNAG_03250 | 1003.96 | 0.935963 | 0.173674 | 5.389186 | 7.08E-08 | 6.28E-07 |
| CNAG_03259 | 111.1711 | 1.167877 | 0.237602 | 4.915259 | 8.87E-07 | 6.21E-06 |
| CNAG_03260 | 1340.438 | -0.69761 | 0.15033 | -4.64054 | 3.48E-06 | 2.17E-05 |
| CNAG_03261 | 3317.734 | -0.46292 | 0.156359 | -2.96064 | 0.00307 | 0.008835 |
| CNAG_03264 | 1635.896 | -0.47601 | 0.13489 | -3.5289 | 0.000417 | 0.001543 |
| CNAG_03266 | 7108.337 | -0.3429 | 0.149828 | -2.28866 | 0.022099 | 0.048535 |
| CNAG_03267 | 4121.853 | -0.64344 | 0.164412 | -3.91361 | 9.09E-05 | 0.000402 |
| CNAG_03268 | 5014.449 | 0.650187 | 0.217295 | 2.992191 | 0.00277 | 0.008129 |
| CNAG_03270 | 4170.582 | -0.70933 | 0.119529 | -5.93443 | 2.95E-09 | 3.43E-08 |
| CNAG_03275 | 748.3796 | -0.36344 | 0.148575 | -2.44617 | 0.014438 | 0.033743 |
| CNAG_03276 | 328.6787 | -0.56748 | 0.250516 | -2.26524 | 0.023498 | 0.05125 |
| CNAG_03277 | 221.7017 | -0.58879 | 0.247829 | -2.37578 | 0.017512 | 0.039746 |
| CNAG_12750 | 60.05136 | 0.724836 | 0.307932 | 2.353881 | 0.018579 | 0.041819 |
| CNAG_03279 | 418.8433 | -0.39825 | 0.16259 | -2.44944 | 0.014308 | 0.033488 |
| CNAG_03280 | 834.2109 | -0.6985 | 0.142548 | -4.90012 | 9.58E-07 | 6.67E-06 |

| | | | | | | |
|---|---|---|---|---|---|---|
| CNAG_03282 | 1877.523 | 0.477581 | 0.176472 | 2.706275 | 0.006804 | 0.017686 |
| CNAG_03283 | 33321.77 | -1.07029 | 0.295215 | -3.62545 | 0.000288 | 0.00112 |
| CNAG_03284 | 970.171 | -0.95449 | 0.128884 | -7.40574 | 1.30E-13 | 2.94E-12 |
| CNAG_03285 | 1893.317 | -0.54955 | 0.127977 | -4.29413 | 1.75E-05 | 9.25E-05 |
| CNAG_03290 | 1494.604 | 0.409568 | 0.156161 | 2.622735 | 0.008723 | 0.02183 |
| CNAG_03291 | 1323.558 | -0.43621 | 0.154739 | -2.81897 | 0.004818 | 0.013119 |
| CNAG_12751 | 32.30105 | 1.07117 | 0.428418 | 2.500292 | 0.012409 | 0.029584 |
| CNAG_07722 | 4.98819 | 2.500707 | 1.194505 | 2.093509 | 0.036304 | 0.074119 |
| CNAG_03292 | 17.394 | 1.707265 | 0.487084 | 3.50507 | 0.000456 | 0.001665 |
| CNAG_03293 | 1379.121 | -0.88134 | 0.216636 | -4.06828 | 4.74E-05 | 0.000223 |
| CNAG_03294 | 287.1748 | 0.536385 | 0.161097 | 3.329581 | 0.00087 | 0.002926 |
| CNAG_03296 | 680.6673 | -0.86526 | 0.148559 | -5.82437 | 5.73E-09 | 6.26E-08 |
| CNAG_03298 | 4225.239 | -0.8225 | 0.261945 | -3.13996 | 0.00169 | 0.005257 |
| CNAG_03300 | 1687.363 | -0.67809 | 0.127044 | -5.33749 | 9.42E-08 | 8.14E-07 |
| CNAG_03302 | 1800.577 | -0.75726 | 0.129556 | -5.84502 | 5.07E-09 | 5.59E-08 |
| CNAG_03303 | 13591.11 | -0.83202 | 0.187366 | -4.44062 | 8.97E-06 | 5.06E-05 |
| CNAG_12753 | 155.757 | 0.829738 | 0.230418 | 3.601017 | 0.000317 | 0.001213 |
| CNAG_03307 | 29.96698 | 1.730749 | 0.47851 | 3.616956 | 0.000298 | 0.001152 |
| CNAG_03308 | 1087.438 | 0.602788 | 0.212492 | 2.836756 | 0.004557 | 0.012505 |
| CNAG_03310 | 711.3058 | -0.62413 | 0.147921 | -4.21935 | 2.45E-05 | 0.000124 |
| CNAG_03311 | 6246.677 | -1.22135 | 0.157658 | -7.74687 | 9.42E-15 | 2.49E-13 |
| CNAG_03314 | 1089.046 | 0.44878 | 0.193914 | 2.314324 | 0.02065 | 0.045757 |
| CNAG_03316 | 2437.492 | 0.739548 | 0.207486 | 3.564332 | 0.000365 | 0.001369 |
| CNAG_03318 | 204.1021 | 0.502257 | 0.18311 | 2.742934 | 0.006089 | 0.016112 |
| CNAG_03319 | 3489.558 | -0.35895 | 0.162496 | -2.20897 | 0.027177 | 0.057802 |
| CNAG_03321 | 2644.047 | 0.260022 | 0.122963 | 2.114634 | 0.034461 | 0.070793 |
| CNAG_03322 | 8583.657 | -1.03479 | 0.234388 | -4.41486 | 1.01E-05 | 5.64E-05 |
| CNAG_03326 | 2478.631 | 1.493089 | 0.119059 | 12.54072 | 4.47E-36 | 6.23E-34 |
| CNAG_03327 | 47.98833 | 0.734747 | 0.309472 | 2.374199 | 0.017587 | 0.039894 |
| CNAG_03328 | 2303.059 | 0.740424 | 0.126658 | 5.845846 | 5.04E-09 | 5.57E-08 |
| CNAG_03330 | 414.7591 | 0.438693 | 0.221191 | 1.983326 | 0.047331 | 0.092771 |
| CNAG_03331 | 2633.287 | -0.37597 | 0.108334 | -3.4705 | 0.000519 | 0.001876 |
| CNAG_03333 | 2336.615 | 1.400019 | 0.17786 | 7.87145 | 3.51E-15 | 9.77E-14 |
| CNAG_03335 | 2638.873 | -0.56613 | 0.134903 | -4.19657 | 2.71E-05 | 0.000135 |
| CNAG_03336 | 2988.563 | 0.502879 | 0.186524 | 2.696052 | 0.007017 | 0.018167 |
| CNAG_03337 | 1357.564 | 0.687643 | 0.157727 | 4.359704 | 1.30E-05 | 7.11E-05 |
| CNAG_03338 | 795.2477 | -0.58664 | 0.18793 | -3.12156 | 0.001799 | 0.005549 |
| CNAG_03340 | 62.92256 | 0.667747 | 0.263952 | 2.529805 | 0.011413 | 0.027547 |
| CNAG_03341 | 1970.636 | -0.52535 | 0.192613 | -2.72749 | 0.006382 | 0.01678 |
| CNAG_03342 | 2463.4 | -0.66767 | 0.125958 | -5.30077 | 1.15E-07 | 9.76E-07 |
| CNAG_03343 | 1032.082 | -0.50579 | 0.148935 | -3.39607 | 0.000684 | 0.002377 |
| CNAG_03345 | 6340.71 | 1.197844 | 0.371726 | 3.222382 | 0.001271 | 0.004105 |
| CNAG_07723 | 41.77314 | 1.60818 | 0.465981 | 3.451173 | 0.000558 | 0.001998 |
| CNAG_03347 | 4478.361 | 0.916507 | 0.12368 | 7.410287 | 1.26E-13 | 2.86E-12 |
| CNAG_03349 | 3958.776 | -0.3399 | 0.1554 | -2.18726 | 0.028723 | 0.060572 |
| CNAG_03353 | 2203.111 | -0.72258 | 0.197685 | -3.65521 | 0.000257 | 0.001009 |
| CNAG_03355 | 5624.413 | 0.537707 | 0.191305 | 2.81073 | 0.004943 | 0.013432 |

| | | | | | | |
|---|---|---|---|---|---|---|
| CNAG_03357 | 1126.707 | -0.52189 | 0.135992 | -3.83763 | 0.000124 | 0.000528 |
| CNAG_03359 | 2186.011 | -0.90018 | 0.302072 | -2.98001 | 0.002882 | 0.008408 |
| CNAG_03360 | 1801.494 | -0.45779 | 0.229735 | -1.99267 | 0.046297 | 0.091059 |
| CNAG_03363 | 829.6426 | -0.39662 | 0.120653 | -3.28727 | 0.001012 | 0.003354 |
| CNAG_03364 | 750.9439 | -0.45363 | 0.139985 | -3.24056 | 0.001193 | 0.00388 |
| CNAG_03365 | 3015.714 | 0.270526 | 0.121243 | 2.231268 | 0.025663 | 0.055025 |
| CNAG_03366 | 431.7754 | 1.167151 | 0.167051 | 6.986802 | 2.81E-12 | 5.40E-11 |
| CNAG_12761 | 153.8575 | -0.66142 | 0.320538 | -2.06347 | 0.039068 | 0.078711 |
| CNAG_12762 | 47.30185 | 1.045067 | 0.305739 | 3.418173 | 0.00063 | 0.002219 |
| CNAG_03372 | 2788.886 | 0.777917 | 0.215907 | 3.603014 | 0.000315 | 0.001205 |
| CNAG_03374 | 127.8355 | 0.736601 | 0.250488 | 2.940664 | 0.003275 | 0.009351 |
| CNAG_03376 | 1530.637 | 0.517641 | 0.240437 | 2.152921 | 0.031325 | 0.06521 |
| CNAG_03377 | 755.3989 | -0.63137 | 0.21388 | -2.95197 | 0.003158 | 0.009057 |
| CNAG_03378 | 2280.813 | -0.48867 | 0.162613 | -3.00513 | 0.002655 | 0.007842 |
| CNAG_03381 | 536.7999 | 0.599228 | 0.183304 | 3.269039 | 0.001079 | 0.003539 |
| CNAG_03382 | 3061.649 | 0.885213 | 0.210739 | 4.200526 | 2.66E-05 | 0.000134 |
| CNAG_03384 | 40.61589 | 1.123566 | 0.337563 | 3.328463 | 0.000873 | 0.002937 |
| CNAG_03385 | 1974.469 | 1.823443 | 0.221115 | 8.246592 | 1.63E-16 | 5.33E-15 |
| CNAG_03386 | 3627.81 | -0.74577 | 0.200811 | -3.7138 | 0.000204 | 0.000824 |
| CNAG_03387 | 373.1175 | -0.4122 | 0.171821 | -2.399 | 0.01644 | 0.037646 |
| CNAG_03389 | 7905.537 | -0.59143 | 0.142535 | -4.14932 | 3.33E-05 | 0.000164 |
| CNAG_03390 | 1128.894 | -0.286 | 0.123966 | -2.30704 | 0.021052 | 0.046545 |
| CNAG_03391 | 818.1959 | -0.48163 | 0.138874 | -3.46811 | 0.000524 | 0.001889 |
| CNAG_03392 | 715.327 | 0.915702 | 0.202673 | 4.518119 | 6.24E-06 | 3.67E-05 |
| CNAG_03394 | 407.5347 | -0.65208 | 0.218894 | -2.97897 | 0.002892 | 0.008424 |
| CNAG_03395 | 1517.391 | -0.83008 | 0.224961 | -3.68989 | 0.000224 | 0.000894 |
| CNAG_03397 | 4552.307 | 0.645432 | 0.126397 | 5.106376 | 3.28E-07 | 2.50E-06 |
| CNAG_03398 | 61.96981 | -2.19039 | 0.441406 | -4.96231 | 6.97E-07 | 5.00E-06 |
| CNAG_03400 | 1137.878 | 1.555945 | 0.162705 | 9.56296 | 1.14E-21 | 6.23E-20 |
| CNAG_03401 | 1434.525 | -0.33945 | 0.145264 | -2.33677 | 0.019451 | 0.043526 |
| CNAG_03403 | 832.8278 | -0.4805 | 0.162777 | -2.95188 | 0.003158 | 0.009057 |
| CNAG_03404 | 354.4324 | 0.665562 | 0.215664 | 3.086103 | 0.002028 | 0.006184 |
| CNAG_03405 | 2972.283 | 0.824776 | 0.107899 | 7.643989 | 2.11E-14 | 5.33E-13 |
| CNAG_03406 | 574.6689 | -0.57509 | 0.175868 | -3.27001 | 0.001075 | 0.003528 |
| CNAG_12767 | 97.78523 | 1.057574 | 0.41347 | 2.557805 | 0.010534 | 0.025746 |
| CNAG_03408 | 8196.069 | 2.248837 | 0.261348 | 8.604768 | 7.65E-18 | 2.81E-16 |
| CNAG_03412 | 1379.517 | 0.680646 | 0.151126 | 4.503842 | 6.67E-06 | 3.90E-05 |
| CNAG_03413 | 1270.619 | -0.27543 | 0.139211 | -1.9785 | 0.047872 | 0.09367 |
| CNAG_03416 | 1765.824 | -0.40817 | 0.167311 | -2.43959 | 0.014704 | 0.034293 |
| CNAG_03417 | 655.7264 | 0.401035 | 0.140891 | 2.846416 | 0.004421 | 0.012182 |
| CNAG_03420 | 1851.292 | 0.582348 | 0.18349 | 3.173727 | 0.001505 | 0.004747 |
| CNAG_03424 | 149.1596 | -0.9608 | 0.427636 | -2.24678 | 0.024654 | 0.053235 |
| CNAG_03426 | 1352.439 | -1.67728 | 0.178989 | -9.37086 | 7.19E-21 | 3.60E-19 |
| CNAG_07732 | 743.0702 | 0.473711 | 0.15947 | 2.970537 | 0.002973 | 0.008624 |
| CNAG_03432 | 836.4701 | 0.551455 | 0.124673 | 4.423199 | 9.73E-06 | 5.45E-05 |
| CNAG_03433 | 2971.971 | -0.52129 | 0.206878 | -2.5198 | 0.011742 | 0.028188 |
| CNAG_03435 | 6578.224 | -0.61321 | 0.205676 | -2.98142 | 0.002869 | 0.008375 |

| | | | | | | |
|---|---|---|---|---|---|---|
| CNAG_03436 | 3480.687 | -0.45541 | 0.136173 | -3.34437 | 0.000825 | 0.002791 |
| CNAG_12769 | 73.32976 | 0.899393 | 0.348333 | 2.581993 | 0.009823 | 0.024241 |
| CNAG_03437 | 155.9288 | 0.78383 | 0.270237 | 2.900527 | 0.003725 | 0.010464 |
| CNAG_12770 | 1069.224 | -0.91882 | 0.198991 | -4.6174 | 3.89E-06 | 2.40E-05 |
| CNAG_03439 | 8045.292 | -0.35666 | 0.165436 | -2.15589 | 0.031092 | 0.064777 |
| CNAG_03441 | 1291.924 | -0.68961 | 0.130148 | -5.29866 | 1.17E-07 | 9.85E-07 |
| CNAG_03442 | 583.0863 | -0.61398 | 0.157354 | -3.90193 | 9.54E-05 | 0.000418 |
| CNAG_03445 | 720.6091 | 1.379318 | 0.158211 | 8.718206 | 2.83E-18 | 1.08E-16 |
| CNAG_03447 | 1062.443 | -0.77133 | 0.287657 | -2.68143 | 0.007331 | 0.018827 |
| CNAG_03448 | 1964.906 | -0.43219 | 0.164922 | -2.62058 | 0.008778 | 0.021934 |
| CNAG_03452 | 1220.699 | -0.41596 | 0.124288 | -3.34677 | 0.000818 | 0.002777 |
| CNAG_03453 | 954.0932 | -1.1471 | 0.249658 | -4.59467 | 4.33E-06 | 2.66E-05 |
| CNAG_12772 | 23.34025 | 1.353314 | 0.486408 | 2.782259 | 0.005398 | 0.014516 |
| CNAG_03454 | 2303.418 | 0.59105 | 0.221035 | 2.674015 | 0.007495 | 0.019156 |
| CNAG_12773 | 135.504 | 1.765257 | 0.279611 | 6.31327 | 2.73E-10 | 3.79E-09 |
| CNAG_03455 | 330.0205 | 1.14507 | 0.225428 | 5.079537 | 3.78E-07 | 2.86E-06 |
| CNAG_03456 | 1848.68 | -0.91765 | 0.208637 | -4.39833 | 1.09E-05 | 6.03E-05 |
| CNAG_03457 | 7994.009 | -0.65972 | 0.130741 | -5.04604 | 4.51E-07 | 3.36E-06 |
| CNAG_03458 | 1891.118 | -0.38881 | 0.145342 | -2.67514 | 0.00747 | 0.019115 |
| CNAG_03463 | 3419.655 | 0.313994 | 0.117379 | 2.675045 | 0.007472 | 0.019115 |
| CNAG_12776 | 234.7642 | -0.67783 | 0.281444 | -2.40839 | 0.016023 | 0.036872 |
| CNAG_07736 | 447.9161 | -1.57114 | 0.181105 | -8.67527 | 4.13E-18 | 1.55E-16 |
| CNAG_12780 | 936.1566 | -0.59896 | 0.21396 | -2.79942 | 0.005119 | 0.01386 |
| CNAG_03473 | 358.8328 | -0.59397 | 0.199133 | -2.98279 | 0.002856 | 0.008341 |
| CNAG_03475 | 2277.265 | 0.72279 | 0.176033 | 4.105987 | 4.03E-05 | 0.000194 |
| CNAG_03476 | 8094.926 | -0.63483 | 0.114647 | -5.53722 | 3.07E-08 | 2.99E-07 |
| CNAG_03477 | 183.3122 | -0.90841 | 0.277076 | -3.27855 | 0.001043 | 0.003442 |
| CNAG_12781 | 139.6914 | -1.46978 | 0.328639 | -4.47231 | 7.74E-06 | 4.43E-05 |
| CNAG_12782 | 45.94104 | 0.810058 | 0.30222 | 2.680356 | 0.007354 | 0.018869 |
| CNAG_03480 | 715.042 | 1.407007 | 0.138116 | 10.18713 | 2.26E-24 | 1.55E-22 |
| CNAG_03481 | 559.0382 | -1.25338 | 0.176379 | -7.10622 | 1.19E-12 | 2.45E-11 |
| CNAG_03483 | 1807.146 | -0.90756 | 0.194253 | -4.67206 | 2.98E-06 | 1.89E-05 |
| CNAG_03484 | 1109.723 | -1.21442 | 0.272269 | -4.46036 | 8.18E-06 | 4.65E-05 |
| CNAG_03485 | 2065.263 | 0.594415 | 0.117497 | 5.05898 | 4.22E-07 | 3.15E-06 |
| CNAG_03492 | 8366.198 | 3.513742 | 0.186522 | 18.8382 | 3.67E-79 | 2.08E-76 |
| CNAG_03493 | 710.559 | 0.385213 | 0.132647 | 2.904036 | 0.003684 | 0.010373 |
| CNAG_03494 | 1859.015 | -0.33817 | 0.128543 | -2.63076 | 0.008519 | 0.02141 |
| CNAG_12785 | 19.49327 | 2.079844 | 0.540322 | 3.849266 | 0.000118 | 0.000507 |
| CNAG_12786 | 584.8094 | 1.995378 | 0.285257 | 6.995012 | 2.65E-12 | 5.14E-11 |
| CNAG_03498 | 3537.951 | 1.221782 | 0.290926 | 4.199633 | 2.67E-05 | 0.000134 |
| CNAG_03499 | 414.4909 | 0.986516 | 0.148868 | 6.62678 | 3.43E-11 | 5.43E-10 |
| CNAG_12788 | 54.37729 | 1.695886 | 0.309183 | 5.485059 | 4.13E-08 | 3.85E-07 |
| CNAG_03500 | 356.0445 | -0.40914 | 0.16083 | -2.54394 | 0.010961 | 0.026627 |
| CNAG_12789 | 50.52344 | 1.345752 | 0.354803 | 3.792961 | 0.000149 | 0.000621 |
| CNAG_03502 | 3126.712 | 0.652816 | 0.14815 | 4.406462 | 1.05E-05 | 5.83E-05 |
| CNAG_03503 | 1232.537 | 0.837828 | 0.231554 | 3.61828 | 0.000297 | 0.001147 |
| CNAG_03504 | 235.9506 | 1.077616 | 0.171499 | 6.283495 | 3.31E-10 | 4.54E-09 |

| | | | | | | |
|---|---|---|---|---|---|---|
| CNAG_03507 | 11897.22 | -0.97373 | 0.227166 | -4.28644 | 1.82E-05 | 9.52E-05 |
| CNAG_03509 | 6840.05 | -0.61874 | 0.220876 | -2.80128 | 0.00509 | 0.01379 |
| CNAG_03510 | 20067.33 | -0.6658 | 0.22013 | -3.02459 | 0.00249 | 0.007415 |
| CNAG_03512 | 1466.19 | -0.57048 | 0.191377 | -2.98094 | 0.002874 | 0.008386 |
| CNAG_03513 | 1292.831 | -1.25861 | 0.237491 | -5.29959 | 1.16E-07 | 9.81E-07 |
| CNAG_07740 | 4909.034 | -0.2737 | 0.130672 | -2.09457 | 0.036209 | 0.073983 |
| CNAG_07741 | 2287.757 | 0.263738 | 0.130612 | 2.019251 | 0.043461 | 0.086314 |
| CNAG_03517 | 6071.724 | 0.789251 | 0.151432 | 5.211907 | 1.87E-07 | 1.52E-06 |
| CNAG_12792 | 20.94023 | -2.00167 | 0.636694 | -3.14384 | 0.001667 | 0.005204 |
| CNAG_03519 | 30.89492 | 0.838084 | 0.355636 | 2.356579 | 0.018444 | 0.041588 |
| CNAG_03521 | 124.7561 | -1.13629 | 0.298072 | -3.81214 | 0.000138 | 0.000581 |
| CNAG_03524 | 1841.778 | -0.96381 | 0.234804 | -4.10472 | 4.05E-05 | 0.000195 |
| CNAG_03525 | 1345.852 | 1.370477 | 0.172095 | 7.963483 | 1.67E-15 | 4.83E-14 |
| CNAG_03526 | 197.396 | 0.953192 | 0.242649 | 3.928276 | 8.56E-05 | 0.000381 |
| CNAG_07995 | 2349.143 | 0.801077 | 0.120727 | 6.635419 | 3.24E-11 | 5.14E-10 |
| CNAG_03534 | 593.1612 | 0.664315 | 0.222448 | 2.98639 | 0.002823 | 0.008262 |
| CNAG_07745 | 10582.03 | 0.680746 | 0.207674 | 3.277964 | 0.001046 | 0.003447 |
| CNAG_07746 | 19935.57 | -1.87158 | 0.226604 | -8.25923 | 1.47E-16 | 4.83E-15 |
| CNAG_12794 | 409.5044 | -0.83391 | 0.305592 | -2.72884 | 0.006356 | 0.016723 |
| CNAG_03538 | 322.7807 | -1.45338 | 0.389396 | -3.7324 | 0.00019 | 0.000772 |
| CNAG_03539 | 738.4286 | -1.00851 | 0.228647 | -4.41078 | 1.03E-05 | 5.73E-05 |
| CNAG_03540 | 1187.125 | 0.561682 | 0.129187 | 4.347829 | 1.37E-05 | 7.46E-05 |
| CNAG_03542 | 1605.488 | -0.37421 | 0.141499 | -2.6446 | 0.008179 | 0.020677 |
| CNAG_12795 | 172.4731 | 0.434493 | 0.176804 | 2.457489 | 0.013991 | 0.032833 |
| CNAG_03543 | 1854.749 | 0.520872 | 0.185458 | 2.808569 | 0.004976 | 0.013509 |
| CNAG_03544 | 5098.413 | -0.90498 | 0.167606 | -5.39943 | 6.69E-08 | 6.00E-07 |
| CNAG_03548 | 36.72053 | 1.032344 | 0.372967 | 2.767924 | 0.005641 | 0.015103 |
| CNAG_03550 | 294.6315 | -0.64718 | 0.149234 | -4.33669 | 1.45E-05 | 7.80E-05 |
| CNAG_03552 | 627.5558 | -0.58559 | 0.237017 | -2.47068 | 0.013486 | 0.03175 |
| CNAG_03553 | 6016.494 | 0.344715 | 0.144878 | 2.379342 | 0.017344 | 0.039425 |
| CNAG_03557 | 1195.447 | 0.501505 | 0.181326 | 2.765766 | 0.005679 | 0.015173 |
| CNAG_03560 | 244.7795 | 0.330974 | 0.165579 | 1.998889 | 0.04562 | 0.089883 |
| CNAG_03562 | 425.166 | -0.8499 | 0.219548 | -3.87112 | 0.000108 | 0.000467 |
| CNAG_03563 | 5482.062 | 1.818982 | 0.18143 | 10.02579 | 1.17E-23 | 7.52E-22 |
| CNAG_03564 | 2016.574 | 1.598746 | 0.215462 | 7.420089 | 1.17E-13 | 2.66E-12 |
| CNAG_03565 | 5681.452 | 2.371709 | 0.234706 | 10.10503 | 5.25E-24 | 3.48E-22 |
| CNAG_03566 | 11820 | 2.075669 | 0.234203 | 8.862674 | 7.81E-19 | 3.18E-17 |
| CNAG_12800 | 131.6413 | 0.516543 | 0.240339 | 2.149226 | 0.031617 | 0.065731 |
| CNAG_03569 | 23.21226 | 1.572984 | 0.595779 | 2.640214 | 0.008285 | 0.0209 |
| CNAG_03570 | 64.20211 | 1.404117 | 0.297257 | 4.723581 | 2.32E-06 | 1.50E-05 |
| CNAG_12801 | 42.61394 | 2.167359 | 0.369827 | 5.860468 | 4.62E-09 | 5.12E-08 |
| CNAG_03572 | 6252.863 | 2.67118 | 0.190028 | 14.05677 | 7.00E-45 | 1.50E-42 |
| CNAG_03573 | 750.7462 | 0.478272 | 0.182223 | 2.624643 | 0.008674 | 0.021742 |
| CNAG_12804 | 21.46813 | 2.636907 | 0.589468 | 4.473364 | 7.70E-06 | 4.41E-05 |
| CNAG_03575 | 804.5943 | 0.581226 | 0.160891 | 3.612538 | 0.000303 | 0.00117 |
| CNAG_03577 | 41013.46 | -0.93654 | 0.183156 | -5.11334 | 3.17E-07 | 2.42E-06 |
| CNAG_03578 | 785.6439 | -0.92984 | 0.15777 | -5.89361 | 3.78E-09 | 4.29E-08 |

| | | | | | | |
|---|---|---|---|---|---|---|
| CNAG_07749 | 951.0989 | -0.45228 | 0.160414 | -2.81947 | 0.00481 | 0.013108 |
| CNAG_03584 | 2300.639 | -0.36483 | 0.128601 | -2.83689 | 0.004556 | 0.012504 |
| CNAG_03587 | 647.9181 | -0.65548 | 0.250828 | -2.61326 | 0.008968 | 0.02236 |
| CNAG_03588 | 8573.218 | -0.80582 | 0.124122 | -6.49215 | 8.46E-11 | 1.27E-09 |
| CNAG_03589 | 1374.964 | -0.54755 | 0.160591 | -3.40961 | 0.000651 | 0.002279 |
| CNAG_03590 | 2805.792 | -0.50381 | 0.141215 | -3.56766 | 0.00036 | 0.001353 |
| CNAG_03592 | 3809.196 | -0.79859 | 0.128366 | -6.22123 | 4.93E-10 | 6.52E-09 |
| CNAG_12806 | 22.25701 | 2.720156 | 0.54382 | 5.001944 | 5.68E-07 | 4.15E-06 |
| CNAG_12807 | 41.79633 | 1.347596 | 0.390325 | 3.452497 | 0.000555 | 0.001989 |
| CNAG_06977 | 75.66666 | -0.69354 | 0.302559 | -2.29226 | 0.021891 | 0.048104 |
| CNAG_06993 | 3820.488 | 0.498323 | 0.123405 | 4.038106 | 5.39E-05 | 0.00025 |
| CNAG_06994 | 1170.964 | 0.568114 | 0.130193 | 4.363623 | 1.28E-05 | 7.00E-05 |
| CNAG_06966 | 62.89035 | 1.388564 | 0.288842 | 4.807341 | 1.53E-06 | 1.02E-05 |
| CNAG_06968 | 2180.263 | -0.6289 | 0.262774 | -2.3933 | 0.016698 | 0.038159 |
| CNAG_04101 | 2290.372 | -0.88139 | 0.158292 | -5.56813 | 2.57E-08 | 2.55E-07 |
| CNAG_12810 | 22.6395 | 6.032422 | 1.084286 | 5.563498 | 2.64E-08 | 2.61E-07 |
| CNAG_04103 | 452.5193 | -1.59826 | 0.163121 | -9.79799 | 1.15E-22 | 7.02E-21 |
| CNAG_04104 | 1001.829 | -0.59287 | 0.148494 | -3.99254 | 6.54E-05 | 0.000299 |
| CNAG_04105 | 7130.739 | 2.418903 | 0.276806 | 8.738632 | 2.36E-18 | 9.10E-17 |
| CNAG_07755 | 161.0579 | 1.448299 | 0.2428 | 5.964992 | 2.45E-09 | 2.91E-08 |
| CNAG_04106 | 4118.491 | 3.035452 | 0.247146 | 12.28204 | 1.13E-34 | 1.45E-32 |
| CNAG_04107 | 506.5762 | 1.2234 | 0.224618 | 5.446572 | 5.13E-08 | 4.70E-07 |
| CNAG_04110 | 851.5525 | 0.670481 | 0.174564 | 3.840899 | 0.000123 | 0.000523 |
| CNAG_04111 | 2410.893 | -0.30167 | 0.121431 | -2.48428 | 0.012981 | 0.030772 |
| CNAG_12812 | 88.08804 | 1.500562 | 0.27392 | 5.478111 | 4.30E-08 | 4.00E-07 |
| CNAG_04112 | 815.9357 | 0.953621 | 0.161192 | 5.916074 | 3.30E-09 | 3.81E-08 |
| CNAG_04114 | 27651.76 | -1.11877 | 0.283575 | -3.94524 | 7.97E-05 | 0.000357 |
| CNAG_07756 | 1369.656 | -1.20999 | 0.162262 | -7.457 | 8.85E-14 | 2.05E-12 |
| CNAG_04115 | 928.2815 | -0.64483 | 0.153111 | -4.21151 | 2.54E-05 | 0.000128 |
| CNAG_04117 | 803.5507 | 0.905755 | 0.158147 | 5.727283 | 1.02E-08 | 1.07E-07 |
| CNAG_12814 | 24.38939 | 1.272439 | 0.400242 | 3.17917 | 0.001477 | 0.004668 |
| CNAG_04120 | 2406.598 | -0.33613 | 0.124826 | -2.69277 | 0.007086 | 0.018317 |
| CNAG_04122 | 336.6849 | -0.34089 | 0.153843 | -2.21583 | 0.026703 | 0.056929 |
| CNAG_12816 | 94.28141 | 0.851252 | 0.248251 | 3.428991 | 0.000606 | 0.002141 |
| CNAG_04129 | 113.2616 | 0.696495 | 0.217225 | 3.206325 | 0.001344 | 0.004304 |
| CNAG_04130 | 2072.306 | -0.68684 | 0.208108 | -3.3004 | 0.000965 | 0.003218 |
| CNAG_04132 | 2824.204 | -0.62893 | 0.150708 | -4.17317 | 3.00E-05 | 0.000149 |
| CNAG_07757 | 1573.024 | 0.666467 | 0.262617 | 2.537794 | 0.011155 | 0.027025 |
| CNAG_04136 | 1782.163 | -0.81859 | 0.129227 | -6.33447 | 2.38E-10 | 3.35E-09 |
| CNAG_12820 | 64.16834 | -1.5564 | 0.286809 | -5.42662 | 5.74E-08 | 5.21E-07 |
| CNAG_04141 | 3675.421 | -0.57224 | 0.127204 | -4.49863 | 6.84E-06 | 3.98E-05 |
| CNAG_04142 | 1378.345 | -0.72593 | 0.198876 | -3.65018 | 0.000262 | 0.001027 |
| CNAG_07759 | 22.71913 | -3.30293 | 0.655429 | -5.03934 | 4.67E-07 | 3.47E-06 |
| CNAG_04146 | 2139.206 | -0.5847 | 0.168734 | -3.46523 | 0.00053 | 0.001907 |
| CNAG_04147 | 1009.633 | -0.53433 | 0.136429 | -3.91653 | 8.98E-05 | 0.000399 |
| CNAG_04148 | 1427.623 | -0.70677 | 0.225213 | -3.13822 | 0.0017 | 0.00528 |
| CNAG_04149 | 2016.607 | -0.5898 | 0.114824 | -5.13658 | 2.80E-07 | 2.16E-06 |

| | | | | | | |
|---|---|---|---|---|---|---|
| CNAG_12822 | 19.02869 | 1.320499 | 0.519193 | 2.543368 | 0.010979 | 0.026654 |
| CNAG_04156 | 2729.947 | -0.85295 | 0.212195 | -4.01963 | 5.83E-05 | 0.000269 |
| CNAG_04157 | 785.542 | -0.68941 | 0.158836 | -4.34042 | 1.42E-05 | 7.68E-05 |
| CNAG_04159 | 1647.857 | 0.728104 | 0.322359 | 2.258671 | 0.023904 | 0.051944 |
| CNAG_04163 | 7519.364 | 2.141726 | 0.251636 | 8.511199 | 1.72E-17 | 6.08E-16 |
| CNAG_04167 | 229.4743 | 0.882497 | 0.16974 | 5.1991 | 2.00E-07 | 1.61E-06 |
| CNAG_04168 | 4803.182 | -1.3609 | 0.195327 | -6.96729 | 3.23E-12 | 6.07E-11 |
| CNAG_04175 | 2011.076 | -1.19563 | 0.17719 | -6.74772 | 1.50E-11 | 2.54E-10 |
| CNAG_04177 | 1380.155 | 0.327963 | 0.122077 | 2.686535 | 0.00722 | 0.018602 |
| CNAG_04178 | 493.1591 | -0.93322 | 0.171148 | -5.45272 | 4.96E-08 | 4.55E-07 |
| CNAG_04179 | 8429.776 | -0.69311 | 0.122365 | -5.66427 | 1.48E-08 | 1.50E-07 |
| CNAG_04180 | 2238.544 | -0.60666 | 0.16998 | -3.56902 | 0.000358 | 0.001347 |
| CNAG_04181 | 1209.874 | -0.90553 | 0.210911 | -4.29341 | 1.76E-05 | 9.27E-05 |
| CNAG_12827 | 1275.048 | -0.9495 | 0.299751 | -3.16763 | 0.001537 | 0.004838 |
| CNAG_04182 | 1817.888 | -0.70017 | 0.212949 | -3.28798 | 0.001009 | 0.003347 |
| CNAG_04183 | 19507.13 | -0.50443 | 0.165203 | -3.05342 | 0.002262 | 0.006818 |
| CNAG_07997 | 74.0655 | 0.932458 | 0.309606 | 3.011759 | 0.002597 | 0.00769 |
| CNAG_04185 | 1700.335 | 0.59424 | 0.228155 | 2.604547 | 0.0092 | 0.022872 |
| CNAG_04186 | 1569.486 | -0.88755 | 0.206962 | -4.28847 | 1.80E-05 | 9.44E-05 |
| CNAG_04187 | 2068.828 | 0.886482 | 0.133823 | 6.624267 | 3.49E-11 | 5.50E-10 |
| CNAG_12829 | 66.00899 | 1.798567 | 0.381135 | 4.718981 | 2.37E-06 | 1.53E-05 |
| CNAG_04188 | 1701.805 | 0.681196 | 0.216967 | 3.139629 | 0.001692 | 0.005261 |
| CNAG_04189 | 7187.697 | -1.46683 | 0.155352 | -9.44198 | 3.66E-21 | 1.88E-19 |
| CNAG_04190 | 5041.428 | -0.75882 | 0.133516 | -5.68334 | 1.32E-08 | 1.36E-07 |
| CNAG_04192 | 5918.783 | -0.61805 | 0.124551 | -4.96219 | 6.97E-07 | 5.00E-06 |
| CNAG_04195 | 2370.614 | 2.044103 | 0.165571 | 12.3458 | 5.13E-35 | 6.69E-33 |
| CNAG_04197 | 2022.645 | 0.645661 | 0.230345 | 2.803018 | 0.005063 | 0.01372 |
| CNAG_04199 | 326.8415 | 0.818004 | 0.260612 | 3.138777 | 0.001697 | 0.005274 |
| CNAG_04200 | 878.344 | 1.233418 | 0.321398 | 3.837669 | 0.000124 | 0.000528 |
| CNAG_04201 | 90.11558 | 0.610962 | 0.242923 | 2.515041 | 0.011902 | 0.028537 |
| CNAG_04203 | 1835.164 | 0.40515 | 0.188882 | 2.144995 | 0.031953 | 0.066362 |
| CNAG_04204 | 1218.2 | -0.935 | 0.218754 | -4.27421 | 1.92E-05 | 9.99E-05 |
| CNAG_04205 | 1563.633 | -0.57466 | 0.130201 | -4.41362 | 1.02E-05 | 5.66E-05 |
| CNAG_04206 | 1557.928 | 1.184503 | 0.199514 | 5.936942 | 2.90E-09 | 3.38E-08 |
| CNAG_12830 | 65.74125 | 1.310281 | 0.338268 | 3.873499 | 0.000107 | 0.000464 |
| CNAG_04208 | 1938.175 | 0.513381 | 0.124941 | 4.108981 | 3.97E-05 | 0.000191 |
| CNAG_04210 | 2237.148 | -0.41044 | 0.168351 | -2.43797 | 0.01477 | 0.034427 |
| CNAG_04212 | 2078.43 | -0.51165 | 0.229271 | -2.23163 | 0.025639 | 0.055002 |
| CNAG_04215 | 15206.26 | -0.93301 | 0.162675 | -5.73542 | 9.73E-09 | 1.02E-07 |
| CNAG_04219 | 3894.344 | -0.41894 | 0.182727 | -2.2927 | 0.021865 | 0.048061 |
| CNAG_04220 | 1562.149 | 0.408667 | 0.133147 | 3.069281 | 0.002146 | 0.00651 |
| CNAG_04222 | 3031.586 | -0.94866 | 0.143251 | -6.62235 | 3.54E-11 | 5.56E-10 |
| CNAG_04224 | 2288.838 | -0.64028 | 0.238828 | -2.68092 | 0.007342 | 0.018845 |
| CNAG_04225 | 145.1467 | 0.916036 | 0.202931 | 4.514024 | 6.36E-06 | 3.74E-05 |
| CNAG_04227 | 1126.67 | -0.92446 | 0.255087 | -3.62408 | 0.00029 | 0.001124 |
| CNAG_04228 | 460.1911 | 1.127675 | 0.157906 | 7.141432 | 9.24E-13 | 1.94E-11 |
| CNAG_04231 | 349.5866 | -0.71753 | 0.20469 | -3.50544 | 0.000456 | 0.001664 |

| | | | | | | |
|---|---|---|---|---|---|---|
| CNAG_04235 | 1553.226 | -0.59405 | 0.188516 | -3.15119 | 0.001626 | 0.005091 |
| CNAG_04236 | 2767.568 | -0.91383 | 0.13516 | -6.76106 | 1.37E-11 | 2.34E-10 |
| CNAG_04239 | 326.0677 | -0.81032 | 0.163946 | -4.94262 | 7.71E-07 | 5.48E-06 |
| CNAG_04242 | 502.6002 | -0.7782 | 0.208742 | -3.72804 | 0.000193 | 0.000784 |
| CNAG_12834 | 74.55906 | 1.246645 | 0.273199 | 4.563139 | 5.04E-06 | 3.03E-05 |
| CNAG_04244 | 1231.29 | 0.619617 | 0.225413 | 2.748801 | 0.005981 | 0.015859 |
| CNAG_04245 | 7021.923 | -0.9591 | 0.112151 | -8.55191 | 1.21E-17 | 4.39E-16 |
| CNAG_04248 | 1192.516 | 0.862232 | 0.171929 | 5.01506 | 5.30E-07 | 3.90E-06 |
| CNAG_07763 | 894.5105 | 0.992244 | 0.148082 | 6.700618 | 2.08E-11 | 3.41E-10 |
| CNAG_04253 | 669.0648 | 0.438009 | 0.176088 | 2.487449 | 0.012866 | 0.030536 |
| CNAG_04255 | 337.8184 | -0.50346 | 0.210446 | -2.39235 | 0.016741 | 0.038247 |
| CNAG_04256 | 529.8846 | 1.478979 | 0.183185 | 8.073683 | 6.82E-16 | 2.09E-14 |
| CNAG_04257 | 4242.46 | -0.53483 | 0.139503 | -3.83379 | 0.000126 | 0.000536 |
| CNAG_04259 | 5143.838 | -0.50215 | 0.18686 | -2.68732 | 0.007203 | 0.01857 |
| CNAG_04260 | 2046.563 | -0.41074 | 0.156405 | -2.62612 | 0.008636 | 0.021662 |
| CNAG_04261 | 276.948 | -0.76721 | 0.225833 | -3.39722 | 0.000681 | 0.002369 |
| CNAG_04262 | 326.1466 | 0.469873 | 0.175732 | 2.673804 | 0.0075 | 0.019161 |
| CNAG_04263 | 533.9701 | -1.27946 | 0.199604 | -6.41 | 1.46E-10 | 2.11E-09 |
| CNAG_04266 | 1045.185 | -0.6186 | 0.133308 | -4.64035 | 3.48E-06 | 2.17E-05 |
| CNAG_04268 | 845.9737 | 0.88338 | 0.19761 | 4.470311 | 7.81E-06 | 4.46E-05 |
| CNAG_04269 | 1867.26 | 0.470735 | 0.136779 | 3.441563 | 0.000578 | 0.002059 |
| CNAG_04270 | 824.0834 | 0.685367 | 0.149088 | 4.597055 | 4.29E-06 | 2.63E-05 |
| CNAG_04276 | 3368.559 | -0.72582 | 0.10599 | -6.84801 | 7.49E-12 | 1.34E-10 |
| CNAG_04278 | 1305.875 | -0.41192 | 0.18755 | -2.19634 | 0.028067 | 0.059378 |
| CNAG_04280 | 130.3603 | 0.566753 | 0.267822 | 2.116154 | 0.034332 | 0.070546 |
| CNAG_04283 | 2729.47 | 0.814737 | 0.130596 | 6.238618 | 4.41E-10 | 5.89E-09 |
| CNAG_12840 | 235.142 | 1.704955 | 0.214979 | 7.930812 | 2.18E-15 | 6.25E-14 |
| CNAG_04284 | 4466.405 | -0.27723 | 0.116667 | -2.37622 | 0.017491 | 0.03971 |
| CNAG_04288 | 1379.689 | 1.068911 | 0.288643 | 3.703226 | 0.000213 | 0.000854 |
| CNAG_04289 | 3703.571 | -0.61982 | 0.14544 | -4.26166 | 2.03E-05 | 0.000105 |
| CNAG_04290 | 3681.986 | -0.61226 | 0.295845 | -2.06951 | 0.038498 | 0.077779 |
| CNAG_04292 | 2752.343 | -0.58497 | 0.214592 | -2.72595 | 0.006412 | 0.016837 |
| CNAG_04293 | 1649.173 | 0.789414 | 0.289914 | 2.72293 | 0.006471 | 0.016952 |
| CNAG_04294 | 401.5228 | 0.51679 | 0.161111 | 3.207668 | 0.001338 | 0.004286 |
| CNAG_04295 | 771.1678 | -0.46464 | 0.138319 | -3.3592 | 0.000782 | 0.00267 |
| CNAG_04297 | 481.5277 | -0.80336 | 0.195829 | -4.10233 | 4.09E-05 | 0.000196 |
| CNAG_04307 | 450.5155 | -0.40281 | 0.178968 | -2.25077 | 0.0244 | 0.052815 |
| CNAG_04309 | 315.8449 | 0.416757 | 0.191578 | 2.175392 | 0.029601 | 0.062118 |
| CNAG_04310 | 1371.064 | -0.92515 | 0.122835 | -7.53164 | 5.01E-14 | 1.22E-12 |
| CNAG_04313 | 524.5698 | 1.694864 | 0.176581 | 9.598212 | 8.13E-22 | 4.55E-20 |
| CNAG_04314 | 4321.687 | 1.473956 | 0.146563 | 10.0568 | 8.57E-24 | 5.63E-22 |
| CNAG_04315 | 3142.176 | 0.69 | 0.148959 | 4.63215 | 3.62E-06 | 2.25E-05 |
| CNAG_04317 | 83.14973 | 0.596892 | 0.246548 | 2.420993 | 0.015478 | 0.035815 |
| CNAG_04320 | 3157.98 | 0.658592 | 0.180831 | 3.642036 | 0.00027 | 0.001057 |
| CNAG_04321 | 2534.753 | 0.58931 | 0.174348 | 3.380077 | 0.000725 | 0.002498 |
| CNAG_04322 | 4593.431 | 2.411687 | 0.221143 | 10.90555 | 1.08E-27 | 8.57E-26 |
| CNAG_04323 | 3202.761 | 0.474797 | 0.20833 | 2.279068 | 0.022663 | 0.049664 |

| | | | | | | |
|---|---|---|---|---|---|---|
| CNAG_04325 | 3095.169 | 3.478768 | 0.146233 | 23.78924 | 4.32E-125 | 1.71E-121 |
| CNAG_04326 | 1483.119 | 0.766839 | 0.144512 | 5.306407 | 1.12E-07 | 9.49E-07 |
| CNAG_12843 | 24.03689 | 1.271629 | 0.48456 | 2.624295 | 0.008683 | 0.021744 |
| CNAG_04331 | 44.74372 | 1.548874 | 0.368981 | 4.19771 | 2.70E-05 | 0.000135 |
| CNAG_04332 | 907.713 | -0.4111 | 0.153631 | -2.67592 | 0.007452 | 0.019084 |
| CNAG_04333 | 73.36433 | 1.164522 | 0.322754 | 3.608079 | 0.000308 | 0.001186 |
| CNAG_04334 | 2030.927 | -0.63836 | 0.129925 | -4.91334 | 8.95E-07 | 6.27E-06 |
| CNAG_04338 | 917.4034 | 1.165952 | 0.195654 | 5.959244 | 2.53E-09 | 3.01E-08 |
| CNAG_04339 | 2162.929 | 0.520372 | 0.170166 | 3.05802 | 0.002228 | 0.006729 |
| CNAG_04340 | 2011.129 | -0.36547 | 0.137589 | -2.65625 | 0.007902 | 0.020047 |
| CNAG_04343 | 251.4318 | 0.624419 | 0.238778 | 2.615058 | 0.008921 | 0.02225 |
| CNAG_04345 | 817.3422 | -0.44936 | 0.181159 | -2.48049 | 0.01312 | 0.031027 |
| CNAG_04347 | 5367.18 | -1.01086 | 0.133176 | -7.59035 | 3.19E-14 | 7.95E-13 |
| CNAG_04348 | 1919.825 | -0.91646 | 0.183019 | -5.00746 | 5.52E-07 | 4.05E-06 |
| CNAG_12847 | 6.407591 | 2.428589 | 0.86124 | 2.819875 | 0.004804 | 0.013096 |
| CNAG_04350 | 845.3031 | 0.690584 | 0.138784 | 4.975968 | 6.49E-07 | 4.69E-06 |
| CNAG_04351 | 556.6744 | 0.994588 | 0.141405 | 7.033609 | 2.01E-12 | 3.99E-11 |
| CNAG_04353 | 1162.174 | 0.867409 | 0.158472 | 5.473582 | 4.41E-08 | 4.09E-07 |
| CNAG_04354 | 1355.85 | 0.501194 | 0.147722 | 3.392826 | 0.000692 | 0.002399 |
| CNAG_04356 | 1952.985 | -0.32229 | 0.126073 | -2.55635 | 0.010578 | 0.025838 |
| CNAG_04361 | 6388.428 | 0.7448 | 0.183812 | 4.051961 | 5.08E-05 | 0.000238 |
| CNAG_04362 | 4037.028 | -0.29395 | 0.12255 | -2.39865 | 0.016456 | 0.037671 |
| CNAG_04363 | 69.05449 | 1.576803 | 0.300212 | 5.25229 | 1.50E-07 | 1.24E-06 |
| CNAG_04364 | 6487.994 | 0.351248 | 0.105253 | 3.337191 | 0.000846 | 0.002855 |
| CNAG_04365 | 4974.926 | -0.63985 | 0.116326 | -5.5005 | 3.79E-08 | 3.61E-07 |
| CNAG_04367 | 366.3704 | -0.74158 | 0.150188 | -4.93767 | 7.91E-07 | 5.60E-06 |
| CNAG_04368 | 1336.038 | -0.37559 | 0.131334 | -2.85982 | 0.004239 | 0.011723 |
| CNAG_04371 | 1029.603 | 0.687212 | 0.18344 | 3.746246 | 0.00018 | 0.000737 |
| CNAG_04372 | 1908.259 | -0.78045 | 0.278194 | -2.80542 | 0.005025 | 0.013633 |
| CNAG_12849 | 10.2376 | 2.470761 | 0.733379 | 3.36901 | 0.000754 | 0.002589 |
| CNAG_04375 | 2308.956 | -0.56945 | 0.176274 | -3.23051 | 0.001236 | 0.004003 |
| CNAG_04376 | 2740.168 | -0.89834 | 0.174468 | -5.14906 | 2.62E-07 | 2.04E-06 |
| CNAG_04378 | 3367.027 | -0.586 | 0.230284 | -2.54467 | 0.010938 | 0.026586 |
| CNAG_04380 | 1157.844 | 0.591378 | 0.164708 | 3.590462 | 0.00033 | 0.00126 |
| CNAG_04383 | 1328.643 | -0.46367 | 0.196343 | -2.36151 | 0.018201 | 0.041156 |
| CNAG_04386 | 1412.366 | 1.944664 | 0.208905 | 9.308861 | 1.29E-20 | 6.38E-19 |
| CNAG_04388 | 4301.498 | 0.912477 | 0.170379 | 5.355569 | 8.53E-08 | 7.45E-07 |
| CNAG_07766 | 1010.517 | 1.224383 | 0.216375 | 5.658621 | 1.53E-08 | 1.55E-07 |
| CNAG_07765 | 858.9353 | 3.822831 | 0.228971 | 16.69569 | 1.41E-62 | 6.22E-60 |
| CNAG_04392 | 538.9923 | 0.850214 | 0.169023 | 5.030174 | 4.90E-07 | 3.62E-06 |
| CNAG_04394 | 41.13837 | 1.311003 | 0.409338 | 3.202736 | 0.001361 | 0.004349 |
| CNAG_04395 | 1592.628 | -1.16024 | 0.200392 | -5.78984 | 7.05E-09 | 7.58E-08 |
| CNAG_04397 | 629.6212 | -0.44685 | 0.170118 | -2.62673 | 0.008621 | 0.02163 |
| CNAG_04399 | 1107.207 | -0.4429 | 0.171011 | -2.58991 | 0.0096 | 0.023742 |
| CNAG_04400 | 1258.815 | -1.12374 | 0.243965 | -4.60614 | 4.10E-06 | 2.53E-05 |
| CNAG_04402 | 1749.823 | -0.91714 | 0.16857 | -5.44067 | 5.31E-08 | 4.84E-07 |
| CNAG_04404 | 94.79903 | 0.598062 | 0.282514 | 2.11693 | 0.034266 | 0.070428 |

| | | | | | | |
|---|---|---|---|---|---|---|
| CNAG_04405 | 1289.449 | 0.389692 | 0.131263 | 2.968781 | 0.00299 | 0.008664 |
| CNAG_04408 | 1758.719 | -0.46947 | 0.110022 | -4.26706 | 1.98E-05 | 0.000102 |
| CNAG_04412 | 862.2095 | 1.11865 | 0.225036 | 4.970978 | 6.66E-07 | 4.81E-06 |
| CNAG_04413 | 36.72698 | 3.022681 | 0.44955 | 6.723791 | 1.77E-11 | 2.94E-10 |
| CNAG_04414 | 1845.424 | 0.886403 | 0.119303 | 7.429832 | 1.09E-13 | 2.50E-12 |
| CNAG_04415 | 198.236 | 1.238673 | 0.249903 | 4.956612 | 7.17E-07 | 5.13E-06 |
| CNAG_04416 | 519.3414 | 4.496399 | 0.223893 | 20.08279 | 1.04E-89 | 9.21E-87 |
| CNAG_07767 | 54.53371 | 1.540597 | 0.337576 | 4.563701 | 5.03E-06 | 3.03E-05 |
| CNAG_04417 | 491.1069 | 0.629406 | 0.134332 | 4.685442 | 2.79E-06 | 1.78E-05 |
| CNAG_12852 | 173.1047 | 0.754919 | 0.274559 | 2.749565 | 0.005967 | 0.015837 |
| CNAG_07769 | 3808.89 | 0.969518 | 0.328881 | 2.94793 | 0.003199 | 0.009157 |
| CNAG_04433 | 1606.335 | 0.536722 | 0.228302 | 2.350931 | 0.018727 | 0.04207 |
| CNAG_04434 | 888.1219 | -1.27716 | 0.182604 | -6.99417 | 2.67E-12 | 5.16E-11 |
| CNAG_04436 | 998.0949 | -0.59039 | 0.177722 | -3.32201 | 0.000894 | 0.002996 |
| CNAG_04437 | 805.8678 | -0.66791 | 0.20952 | -3.1878 | 0.001434 | 0.004542 |
| CNAG_04438 | 1018.546 | -0.49817 | 0.130949 | -3.8043 | 0.000142 | 0.000598 |
| CNAG_04440 | 647.5042 | -0.50811 | 0.207237 | -2.45183 | 0.014213 | 0.033305 |
| CNAG_04441 | 18171.47 | -0.72587 | 0.154704 | -4.69202 | 2.71E-06 | 1.73E-05 |
| CNAG_04443 | 2396.583 | 1.289846 | 0.161625 | 7.980495 | 1.46E-15 | 4.26E-14 |
| CNAG_04445 | 34970.77 | -0.74505 | 0.196298 | -3.79548 | 0.000147 | 0.000617 |
| CNAG_04446 | 157.5928 | 0.669466 | 0.232421 | 2.880408 | 0.003972 | 0.011042 |
| CNAG_04447 | 730.1736 | -0.57513 | 0.156992 | -3.66345 | 0.000249 | 0.00098 |
| CNAG_04448 | 53001.49 | -0.91345 | 0.210896 | -4.33131 | 1.48E-05 | 7.94E-05 |
| CNAG_04451 | 1165.073 | -0.52559 | 0.128817 | -4.08013 | 4.50E-05 | 0.000213 |
| CNAG_04452 | 1048.456 | -0.81472 | 0.227141 | -3.58685 | 0.000335 | 0.001274 |
| CNAG_04453 | 844.8054 | -0.34484 | 0.134972 | -2.55491 | 0.010622 | 0.025905 |
| CNAG_04454 | 13.35447 | 1.46579 | 0.534502 | 2.742349 | 0.0061 | 0.016124 |
| CNAG_04455 | 309.9925 | 0.542509 | 0.183575 | 2.955244 | 0.003124 | 0.008973 |
| CNAG_04456 | 812.9594 | 0.334405 | 0.168957 | 1.979234 | 0.04779 | 0.093554 |
| CNAG_04457 | 60.58762 | 1.692044 | 0.306876 | 5.513778 | 3.51E-08 | 3.37E-07 |
| CNAG_04459 | 54.84426 | 2.732116 | 0.405142 | 6.743608 | 1.55E-11 | 2.61E-10 |
| CNAG_04460 | 2892.787 | -0.4533 | 0.108936 | -4.16117 | 3.17E-05 | 0.000156 |
| CNAG_04462 | 374.7069 | 0.369521 | 0.166678 | 2.216979 | 0.026625 | 0.056825 |
| CNAG_04465 | 2954.858 | -0.59607 | 0.203418 | -2.93026 | 0.003387 | 0.009639 |
| CNAG_04466 | 463.5234 | 0.981358 | 0.200764 | 4.888108 | 1.02E-06 | 7.02E-06 |
| CNAG_04469 | 365.2721 | -0.6715 | 0.24052 | -2.79186 | 0.005241 | 0.014156 |
| CNAG_04470 | 1013.564 | -1.16756 | 0.237207 | -4.92212 | 8.56E-07 | 6.02E-06 |
| CNAG_04471 | 689.5265 | -0.50296 | 0.177012 | -2.84141 | 0.004491 | 0.012349 |
| CNAG_04473 | 1148.426 | -0.63126 | 0.15703 | -4.01999 | 5.82E-05 | 0.000269 |
| CNAG_04474 | 214.3697 | 0.844546 | 0.172473 | 4.896685 | 9.75E-07 | 6.75E-06 |
| CNAG_04476 | 101.1503 | 1.044857 | 0.293618 | 3.55856 | 0.000373 | 0.001396 |
| CNAG_07771 | 2497.865 | 0.273504 | 0.124995 | 2.188115 | 0.028661 | 0.060457 |
| CNAG_04478 | 3449.722 | 0.333258 | 0.151693 | 2.196922 | 0.028026 | 0.059338 |
| CNAG_04479 | 471.6854 | -1.21838 | 0.175367 | -6.94758 | 3.72E-12 | 6.92E-11 |
| CNAG_07772 | 144.7322 | 0.781972 | 0.192756 | 4.056794 | 4.98E-05 | 0.000234 |
| CNAG_04485 | 7240.922 | 0.327474 | 0.156094 | 2.09793 | 0.035911 | 0.073487 |
| CNAG_04486 | 61.02837 | 0.686836 | 0.303535 | 2.262789 | 0.023649 | 0.051511 |

| | | | | | | |
|---|---|---|---|---|---|---|
| CNAG_07775 | 1606.814 | 1.712925 | 0.285157 | 6.00696 | 1.89E-09 | 2.30E-08 |
| CNAG_07777 | 238.357 | -0.59528 | 0.181425 | -3.28116 | 0.001034 | 0.003414 |
| CNAG_04493 | 747.459 | -0.50873 | 0.146544 | -3.47151 | 0.000518 | 0.00187 |
| CNAG_04494 | 1842.541 | -0.46988 | 0.21731 | -2.16225 | 0.030599 | 0.063934 |
| CNAG_04496 | 793.9003 | -1.26891 | 0.21651 | -5.86078 | 4.61E-09 | 5.12E-08 |
| CNAG_04497 | 665.1585 | -0.52085 | 0.149144 | -3.49227 | 0.000479 | 0.001738 |
| CNAG_04501 | 3818.458 | -0.49984 | 0.134988 | -3.70288 | 0.000213 | 0.000854 |
| CNAG_04503 | 742.5382 | -0.45719 | 0.219764 | -2.08035 | 0.037493 | 0.076273 |
| CNAG_04505 | 2311.657 | 0.601186 | 0.241315 | 2.491295 | 0.012728 | 0.030253 |
| CNAG_12867 | 36.69069 | 2.36022 | 0.402643 | 5.86182 | 4.58E-09 | 5.10E-08 |
| CNAG_04506 | 68.12763 | 1.392824 | 0.279279 | 4.987216 | 6.13E-07 | 4.45E-06 |
| CNAG_04507 | 728.038 | 1.118541 | 0.226033 | 4.948571 | 7.48E-07 | 5.33E-06 |
| CNAG_04512 | 451.0593 | -0.38883 | 0.197624 | -1.9675 | 0.049125 | 0.095768 |
| CNAG_04513 | 2402.751 | -0.35561 | 0.176838 | -2.01092 | 0.044333 | 0.087748 |
| CNAG_04515 | 1727.997 | -0.55613 | 0.146752 | -3.78957 | 0.000151 | 0.000627 |
| CNAG_12869 | 155.5881 | -1.91896 | 0.321853 | -5.96223 | 2.49E-09 | 2.96E-08 |
| CNAG_04518 | 663.8676 | -0.53301 | 0.154336 | -3.45357 | 0.000553 | 0.001983 |
| CNAG_04519 | 334.3014 | -0.52879 | 0.200893 | -2.63217 | 0.008484 | 0.021334 |
| CNAG_04520 | 427.4157 | -0.31019 | 0.137654 | -2.25342 | 0.024232 | 0.052523 |
| CNAG_04521 | 375.7199 | 1.37311 | 0.177477 | 7.736847 | 1.02E-14 | 2.66E-13 |
| CNAG_04522 | 1293.56 | -0.66766 | 0.186798 | -3.57421 | 0.000351 | 0.001328 |
| CNAG_04523 | 108.5481 | 0.747968 | 0.285015 | 2.624308 | 0.008683 | 0.021744 |
| CNAG_04524 | 1392.208 | 0.803397 | 0.214612 | 3.743494 | 0.000181 | 0.000744 |
| CNAG_04528 | 194.7188 | -1.10077 | 0.260503 | -4.22558 | 2.38E-05 | 0.000121 |
| CNAG_04529 | 427.8862 | -1.02063 | 0.252412 | -4.0435 | 5.27E-05 | 0.000245 |
| CNAG_07778 | 7406.647 | -0.56113 | 0.105623 | -5.31259 | 1.08E-07 | 9.21E-07 |
| CNAG_04534 | 201.998 | -1.09929 | 0.222777 | -4.93451 | 8.04E-07 | 5.68E-06 |
| CNAG_04535 | 78.15657 | -0.76105 | 0.267584 | -2.84415 | 0.004453 | 0.012252 |
| CNAG_04536 | 36.6286 | -1.15119 | 0.365099 | -3.1531 | 0.001615 | 0.005062 |
| CNAG_04539 | 1079.957 | 0.357022 | 0.173422 | 2.05869 | 0.039524 | 0.079488 |
| CNAG_04541 | 1232.917 | -0.39857 | 0.155957 | -2.55565 | 0.010599 | 0.025866 |
| CNAG_04546 | 62.03755 | 1.520756 | 0.298584 | 5.093227 | 3.52E-07 | 2.67E-06 |
| CNAG_12873 | 118.7335 | 1.41054 | 0.221231 | 6.375862 | 1.82E-10 | 2.59E-09 |
| CNAG_07780 | 941.9218 | 0.864166 | 0.143657 | 6.015489 | 1.79E-09 | 2.20E-08 |
| CNAG_07781 | 1097.806 | 0.971869 | 0.276232 | 3.518312 | 0.000434 | 0.001593 |
| CNAG_06998 | 13337.54 | -0.37455 | 0.169739 | -2.20662 | 0.02734 | 0.058087 |
| CNAG_06999 | 1286.754 | 1.664299 | 0.176353 | 9.437293 | 3.83E-21 | 1.95E-19 |
| CNAG_12876 | 476.986 | -0.61206 | 0.211617 | -2.8923 | 0.003824 | 0.010689 |
| CNAG_07784 | 735.872 | 1.398085 | 0.221746 | 6.304909 | 2.88E-10 | 3.99E-09 |
| CNAG_07002 | 144.1876 | 0.707921 | 0.288867 | 2.450686 | 0.014258 | 0.033392 |
| CNAG_04931 | 2.241939 | 3.505295 | 1.584259 | 2.212577 | 0.026927 | 0.057331 |
| CNAG_04926 | 102.8055 | 0.607875 | 0.244549 | 2.485695 | 0.01293 | 0.030662 |
| CNAG_04925 | 1410.134 | -0.40851 | 0.168145 | -2.42953 | 0.015118 | 0.035126 |
| CNAG_04922 | 257.2537 | 0.882087 | 0.165103 | 5.342634 | 9.16E-08 | 7.94E-07 |
| CNAG_04921 | 47.5342 | 1.480119 | 0.56591 | 2.615466 | 0.008911 | 0.02223 |
| CNAG_04920 | 2373.494 | 2.132758 | 0.198571 | 10.74053 | 6.57E-27 | 5.02E-25 |
| CNAG_07839 | 43385.38 | -1.00518 | 0.268778 | -3.73981 | 0.000184 | 0.000753 |

| | | | | | | |
|---|---|---|---|---|---|---|
| CNAG_04914 | 6049.696 | -0.66808 | 0.302597 | -2.20782 | 0.027257 | 0.057942 |
| CNAG_04911 | 543.5153 | 0.617895 | 0.183016 | 3.376174 | 0.000735 | 0.002528 |
| CNAG_07840 | 28.82336 | 1.478802 | 0.535591 | 2.761066 | 0.005761 | 0.015367 |
| CNAG_04908 | 736.1696 | 0.421523 | 0.152174 | 2.770007 | 0.005606 | 0.015017 |
| CNAG_04907 | 89.91186 | 0.789338 | 0.266487 | 2.96201 | 0.003056 | 0.008812 |
| CNAG_04906 | 4401.702 | 0.321462 | 0.142713 | 2.252509 | 0.02429 | 0.05262 |
| CNAG_04905 | 41.09815 | -1.82023 | 0.385169 | -4.72579 | 2.29E-06 | 1.49E-05 |
| CNAG_04904 | 13509.41 | 0.263919 | 0.109729 | 2.405183 | 0.016164 | 0.037143 |
| CNAG_04903 | 3289.99 | 4.016241 | 0.212593 | 18.89167 | 1.34E-79 | 8.16E-77 |
| CNAG_04902 | 2776.11 | 0.496636 | 0.173896 | 2.855937 | 0.004291 | 0.01186 |
| CNAG_04901 | 360.0345 | -1.02283 | 0.173221 | -5.90477 | 3.53E-09 | 4.05E-08 |
| CNAG_04900 | 1629.873 | -0.71636 | 0.132497 | -5.40664 | 6.42E-08 | 5.79E-07 |
| CNAG_04899 | 1524.979 | -0.7241 | 0.162868 | -4.44591 | 8.75E-06 | 4.94E-05 |
| CNAG_04898 | 145.4048 | 1.061039 | 0.207636 | 5.110103 | 3.22E-07 | 2.46E-06 |
| CNAG_04896 | 1257.197 | -0.40533 | 0.120415 | -3.3661 | 0.000762 | 0.002609 |
| CNAG_04894 | 182.0871 | 1.489238 | 0.207483 | 7.177651 | 7.09E-13 | 1.51E-11 |
| CNAG_04891 | 86.3423 | 2.161455 | 0.302923 | 7.13533 | 9.66E-13 | 2.01E-11 |
| CNAG_04889 | 1047.584 | 1.031278 | 0.185064 | 5.57255 | 2.51E-08 | 2.49E-07 |
| CNAG_04887 | 159.4706 | 0.683424 | 0.292751 | 2.334491 | 0.01957 | 0.043767 |
| CNAG_04886 | 1319.138 | 1.424848 | 0.153038 | 9.31039 | 1.27E-20 | 6.33E-19 |
| CNAG_12883 | 56.6491 | 1.86334 | 0.343987 | 5.416895 | 6.06E-08 | 5.48E-07 |
| CNAG_04884 | 28130.58 | -1.1492 | 0.310874 | -3.69668 | 0.000218 | 0.000873 |
| CNAG_04883 | 27845.7 | -0.7357 | 0.236452 | -3.1114 | 0.001862 | 0.005719 |
| CNAG_12884 | 28.7377 | 0.995551 | 0.463179 | 2.149388 | 0.031604 | 0.065722 |
| CNAG_04882 | 1359.947 | -0.32079 | 0.139442 | -2.30052 | 0.021419 | 0.047263 |
| CNAG_04880 | 429.7505 | 0.605965 | 0.240984 | 2.514548 | 0.011919 | 0.028568 |
| CNAG_04879 | 6961.109 | 0.923634 | 0.255848 | 3.610091 | 0.000306 | 0.001179 |
| CNAG_12885 | 122.0404 | -2.6812 | 0.903044 | -2.96907 | 0.002987 | 0.008659 |
| CNAG_04877 | 1424.001 | -5.98492 | 1.321154 | -4.53007 | 5.90E-06 | 3.48E-05 |
| CNAG_04876 | 2144.667 | -0.42155 | 0.187589 | -2.24721 | 0.024627 | 0.053204 |
| CNAG_04875 | 992.1489 | -0.66694 | 0.187378 | -3.55936 | 0.000372 | 0.001393 |
| CNAG_04874 | 863.8176 | -1.11336 | 0.202522 | -5.49748 | 3.85E-08 | 3.65E-07 |
| CNAG_04873 | 203.0658 | -0.57667 | 0.180816 | -3.18927 | 0.001426 | 0.004528 |
| CNAG_04872 | 1777.077 | -0.80255 | 0.139849 | -5.73871 | 9.54E-09 | 1.00E-07 |
| CNAG_04871 | 436.7681 | -0.4653 | 0.178448 | -2.60746 | 0.009122 | 0.022707 |
| CNAG_12886 | 86.59126 | -0.7778 | 0.24838 | -3.13149 | 0.001739 | 0.005388 |
| CNAG_04869 | 2021.02 | -1.61064 | 0.167833 | -9.59672 | 8.25E-22 | 4.59E-20 |
| CNAG_04868 | 913.7871 | -0.33405 | 0.134685 | -2.48023 | 0.01313 | 0.031041 |
| CNAG_04867 | 701.1543 | 1.008302 | 0.217913 | 4.627083 | 3.71E-06 | 2.30E-05 |
| CNAG_04865 | 76.59406 | 0.621422 | 0.264692 | 2.347712 | 0.018889 | 0.042399 |
| CNAG_04864 | 3599.543 | -0.6048 | 0.155889 | -3.87966 | 0.000105 | 0.000453 |
| CNAG_12889 | 95.56136 | -1.51286 | 0.33477 | -4.51909 | 6.21E-06 | 3.66E-05 |
| CNAG_04862 | 5522.086 | -1.10038 | 0.324784 | -3.38802 | 0.000704 | 0.002434 |
| CNAG_12890 | 15.03917 | -1.30059 | 0.53379 | -2.43653 | 0.014829 | 0.034544 |
| CNAG_04861 | 1575.512 | 1.369571 | 0.138103 | 9.917001 | 3.51E-23 | 2.21E-21 |
| CNAG_04860 | 2485.265 | 0.923273 | 0.145897 | 6.328234 | 2.48E-10 | 3.46E-09 |
| CNAG_04859 | 930.3518 | -0.85463 | 0.160329 | -5.33045 | 9.80E-08 | 8.43E-07 |

| | | | | | | |
|---|---|---|---|---|---|---|
| CNAG_04857 | 3326.293 | 0.58734 | 0.184088 | 3.19053 | 0.00142 | 0.004512 |
| CNAG_04853 | 1042.25 | 0.518293 | 0.21084 | 2.458229 | 0.013962 | 0.032776 |
| CNAG_04851 | 26576.07 | 1.173573 | 0.12412 | 9.455129 | 3.23E-21 | 1.66E-19 |
| CNAG_12892 | 197.1012 | 1.11471 | 0.238151 | 4.68068 | 2.86E-06 | 1.82E-05 |
| CNAG_04849 | 705.9836 | 1.011578 | 0.283407 | 3.569346 | 0.000358 | 0.001347 |
| CNAG_04845 | 1389.625 | -0.37771 | 0.172407 | -2.19078 | 0.028468 | 0.060097 |
| CNAG_04843 | 981.2917 | -0.60725 | 0.160296 | -3.78832 | 0.000152 | 0.00063 |
| CNAG_04840 | 6444.59 | -0.57153 | 0.279203 | -2.047 | 0.040659 | 0.081584 |
| CNAG_04839 | 848.3073 | -0.96597 | 0.20048 | -4.81828 | 1.45E-06 | 9.70E-06 |
| CNAG_04837 | 173.0774 | -1.38375 | 0.19741 | -7.00952 | 2.39E-12 | 4.68E-11 |
| CNAG_04835 | 155.1204 | 0.537067 | 0.221301 | 2.426864 | 0.01523 | 0.035339 |
| CNAG_04833 | 1772.232 | 1.260234 | 0.174777 | 7.210524 | 5.57E-13 | 1.20E-11 |
| CNAG_04831 | 259.9463 | 1.05037 | 0.18197 | 5.772231 | 7.82E-09 | 8.33E-08 |
| CNAG_04830 | 761.4839 | -0.66146 | 0.153411 | -4.31166 | 1.62E-05 | 8.59E-05 |
| CNAG_04828 | 9849.897 | 0.427795 | 0.206546 | 2.071185 | 0.038342 | 0.077601 |
| CNAG_04824 | 644.7034 | -0.97066 | 0.280429 | -3.46136 | 0.000537 | 0.001933 |
| CNAG_04817 | 1251.888 | -0.69646 | 0.223929 | -3.11019 | 0.00187 | 0.005738 |
| CNAG_04816 | 800.9765 | 1.146789 | 0.190846 | 6.008962 | 1.87E-09 | 2.28E-08 |
| CNAG_07845 | 89.51658 | -1.11435 | 0.29932 | -3.72293 | 0.000197 | 0.000797 |
| CNAG_04815 | 1826.904 | 0.55146 | 0.203573 | 2.708899 | 0.006751 | 0.017576 |
| CNAG_12898 | 561.3834 | -0.99761 | 0.276692 | -3.60549 | 0.000312 | 0.001197 |
| CNAG_04809 | 1405.428 | 0.354045 | 0.131493 | 2.692501 | 0.007092 | 0.018323 |
| CNAG_12900 | 198.6281 | -0.82281 | 0.323673 | -2.54209 | 0.011019 | 0.026735 |
| CNAG_04805 | 1488.592 | -0.45113 | 0.121293 | -3.71939 | 0.0002 | 0.000807 |
| CNAG_04803 | 7488.824 | 0.54711 | 0.24562 | 2.227464 | 0.025916 | 0.055492 |
| CNAG_04802 | 1218.362 | -0.54436 | 0.132971 | -4.0938 | 4.24E-05 | 0.000203 |
| CNAG_04800 | 6828.782 | -0.51388 | 0.098867 | -5.1977 | 2.02E-07 | 1.61E-06 |
| CNAG_04799 | 41178.63 | -1.22683 | 0.28887 | -4.24701 | 2.17E-05 | 0.000111 |
| CNAG_04798 | 3932.397 | -0.61175 | 0.163608 | -3.73915 | 0.000185 | 0.000755 |
| CNAG_04797 | 2591.185 | -0.43026 | 0.107841 | -3.9898 | 6.61E-05 | 0.000301 |
| CNAG_04794 | 2474.587 | 2.075434 | 0.245324 | 8.459957 | 2.67E-17 | 9.36E-16 |
| CNAG_04793 | 35.80674 | -1.43152 | 0.469802 | -3.04706 | 0.002311 | 0.006942 |
| CNAG_04792 | 2434.457 | 0.660484 | 0.217857 | 3.031735 | 0.002432 | 0.007255 |
| CNAG_04791 | 1272.794 | -0.61799 | 0.112937 | -5.47201 | 4.45E-08 | 4.12E-07 |
| CNAG_04789 | 1775.409 | 0.971872 | 0.183539 | 5.295176 | 1.19E-07 | 1.00E-06 |
| CNAG_04788 | 1910.073 | -0.76084 | 0.237129 | -3.20854 | 0.001334 | 0.004278 |
| CNAG_04787 | 401.0619 | 0.776941 | 0.166546 | 4.665034 | 3.09E-06 | 1.95E-05 |
| CNAG_12903 | 74.12723 | 0.829651 | 0.27263 | 3.043139 | 0.002341 | 0.007023 |
| CNAG_04785 | 626.5591 | -0.66495 | 0.234808 | -2.83188 | 0.004628 | 0.012658 |
| CNAG_04783 | 1505.076 | -0.4951 | 0.124288 | -3.9835 | 6.79E-05 | 0.000309 |
| CNAG_04777 | 931.6976 | -0.45553 | 0.136114 | -3.34666 | 0.000818 | 0.002777 |
| CNAG_04773 | 63.59954 | 0.564584 | 0.280363 | 2.013762 | 0.044035 | 0.087235 |
| CNAG_12906 | 66.11613 | 1.450088 | 0.305901 | 4.740381 | 2.13E-06 | 1.39E-05 |
| CNAG_04770 | 2267.278 | -0.92586 | 0.182713 | -5.0673 | 4.03E-07 | 3.02E-06 |
| CNAG_04769 | 1193.374 | -0.30764 | 0.130174 | -2.36329 | 0.018114 | 0.040994 |
| CNAG_04768 | 1220.495 | -0.68257 | 0.165159 | -4.13278 | 3.58E-05 | 0.000174 |
| CNAG_04763 | 2848.652 | 0.621002 | 0.306919 | 2.023343 | 0.043038 | 0.085602 |

| | | | | | | |
|---|---|---|---|---|---|---|
| CNAG_04762 | 39867.25 | -0.98879 | 0.239338 | -4.13138 | 3.61E-05 | 0.000175 |
| CNAG_04761 | 415.5378 | -0.40257 | 0.173637 | -2.31846 | 0.020424 | 0.045371 |
| CNAG_04760 | 33099.45 | -0.64947 | 0.211092 | -3.07674 | 0.002093 | 0.006364 |
| CNAG_12910 | 559.6893 | -1.40452 | 0.259163 | -5.41946 | 5.98E-08 | 5.40E-07 |
| CNAG_04758 | 19874.37 | -0.90575 | 0.209918 | -4.31478 | 1.60E-05 | 8.49E-05 |
| CNAG_04753 | 555.9771 | -0.62766 | 0.141731 | -4.42852 | 9.49E-06 | 5.33E-05 |
| CNAG_04751 | 1375.934 | 1.230806 | 0.224182 | 5.490208 | 4.01E-08 | 3.77E-07 |
| CNAG_07853 | 51.3055 | 2.116019 | 0.356187 | 5.940747 | 2.84E-09 | 3.32E-08 |
| CNAG_12916 | 67.50749 | 0.795703 | 0.29569 | 2.691009 | 0.007124 | 0.01839 |
| CNAG_04750 | 333.1582 | 1.52685 | 0.261346 | 5.842264 | 5.15E-09 | 5.67E-08 |
| CNAG_04747 | 969.2673 | 1.200467 | 0.157043 | 7.644205 | 2.10E-14 | 5.33E-13 |
| CNAG_04746 | 2062.11 | 0.377328 | 0.171367 | 2.201874 | 0.027674 | 0.058687 |
| CNAG_04744 | 3302.21 | 2.69929 | 0.245163 | 11.01021 | 3.41E-28 | 2.77E-26 |
| CNAG_04741 | 2522.639 | 0.503863 | 0.20122 | 2.504037 | 0.012279 | 0.029352 |
| CNAG_04737 | 11278.11 | 1.218588 | 0.291679 | 4.177833 | 2.94E-05 | 0.000146 |
| CNAG_04736 | 699.6732 | -0.93403 | 0.2306 | -4.05045 | 5.11E-05 | 0.000239 |
| CNAG_04732 | 1215.114 | -0.36083 | 0.160764 | -2.24448 | 0.024802 | 0.053466 |
| CNAG_04730 | 1679.757 | 0.614014 | 0.1215 | 5.0536 | 4.34E-07 | 3.23E-06 |
| CNAG_12918 | 30.43895 | -1.13547 | 0.450568 | -2.52008 | 0.011733 | 0.028188 |
| CNAG_04729 | 2268.456 | -0.86347 | 0.146279 | -5.90287 | 3.57E-09 | 4.08E-08 |
| CNAG_04728 | 1335.764 | -0.5046 | 0.211747 | -2.38304 | 0.01717 | 0.039093 |
| CNAG_04727 | 299.0197 | 1.015026 | 0.202296 | 5.017539 | 5.23E-07 | 3.85E-06 |
| CNAG_04726 | 29706.31 | -1.00864 | 0.174013 | -5.79635 | 6.78E-09 | 7.30E-08 |
| CNAG_04725 | 1158.212 | -0.3985 | 0.173361 | -2.29864 | 0.021525 | 0.047446 |
| CNAG_04724 | 3052.692 | 1.008351 | 0.209893 | 4.804106 | 1.55E-06 | 1.04E-05 |
| CNAG_04721 | 1108.847 | -0.36101 | 0.142457 | -2.53416 | 0.011272 | 0.02724 |
| CNAG_12921 | 134.4506 | 0.57018 | 0.225651 | 2.526825 | 0.01151 | 0.027723 |
| CNAG_04717 | 28.25016 | 0.821932 | 0.357986 | 2.295988 | 0.021677 | 0.047739 |
| CNAG_04716 | 4205.379 | -0.40802 | 0.130183 | -3.13421 | 0.001723 | 0.005344 |
| CNAG_04714 | 798.1972 | 0.671298 | 0.130056 | 5.161622 | 2.45E-07 | 1.92E-06 |
| CNAG_04712 | 984.5449 | -0.3457 | 0.145293 | -2.3793 | 0.017346 | 0.039425 |
| CNAG_04711 | 33.13132 | 1.084142 | 0.394238 | 2.749971 | 0.00596 | 0.015828 |
| CNAG_04710 | 2215.097 | -0.25386 | 0.113897 | -2.22886 | 0.025823 | 0.055307 |
| CNAG_04709 | 3099.776 | -0.66071 | 0.113237 | -5.83474 | 5.39E-09 | 5.90E-08 |
| CNAG_04707 | 284.4198 | 0.621541 | 0.189352 | 3.282464 | 0.001029 | 0.003402 |
| CNAG_04705 | 1844.286 | 0.539623 | 0.158779 | 3.398582 | 0.000677 | 0.00236 |
| CNAG_04704 | 43.32203 | -1.04239 | 0.350032 | -2.97798 | 0.002902 | 0.008442 |
| CNAG_04700 | 237.357 | -0.45493 | 0.224553 | -2.02595 | 0.04277 | 0.085112 |
| CNAG_04697 | 1083.274 | -1.05134 | 0.210974 | -4.9833 | 6.25E-07 | 4.53E-06 |
| CNAG_04695 | 50.63395 | 1.49797 | 0.349577 | 4.285097 | 1.83E-05 | 9.57E-05 |
| CNAG_04694 | 4021.925 | -0.65949 | 0.129633 | -5.08736 | 3.63E-07 | 2.75E-06 |
| CNAG_04692 | 1272.793 | -0.86641 | 0.147213 | -5.88542 | 3.97E-09 | 4.48E-08 |
| CNAG_04691 | 68.97791 | 1.177253 | 0.428474 | 2.747545 | 0.006004 | 0.015914 |
| CNAG_04690 | 794.3843 | -0.4378 | 0.133633 | -3.27613 | 0.001052 | 0.003463 |
| CNAG_04689 | 827.1002 | -0.34123 | 0.170227 | -2.00454 | 0.045012 | 0.08884 |
| CNAG_04688 | 649.9956 | 0.401621 | 0.184803 | 2.173236 | 0.029763 | 0.062416 |
| CNAG_04687 | 16840.14 | 0.921633 | 0.139846 | 6.590349 | 4.39E-11 | 6.77E-10 |

| | | | | | | |
|---|---|---|---|---|---|---|
| CNAG_04686 | 525.5554 | -0.82215 | 0.281833 | -2.91716 | 0.003532 | 0.009999 |
| CNAG_07856 | 840.4385 | 0.953624 | 0.175832 | 5.423511 | 5.84E-08 | 5.29E-07 |
| CNAG_04683 | 1334.815 | -0.95405 | 0.223519 | -4.26831 | 1.97E-05 | 0.000102 |
| CNAG_04681 | 229.7749 | 1.517405 | 0.473514 | 3.204562 | 0.001353 | 0.004325 |
| CNAG_04680 | 1164.302 | 1.014468 | 0.208358 | 4.868876 | 1.12E-06 | 7.66E-06 |
| CNAG_04679 | 1880.099 | -0.70236 | 0.26083 | -2.69277 | 0.007086 | 0.018317 |
| CNAG_04675 | 1069.071 | 1.133092 | 0.304317 | 3.723395 | 0.000197 | 0.000796 |
| CNAG_12929 | 267.8376 | 2.387415 | 0.214239 | 11.1437 | 7.69E-29 | 6.57E-27 |
| CNAG_04673 | 162.3214 | 0.966864 | 0.185248 | 5.219302 | 1.80E-07 | 1.46E-06 |
| CNAG_12930 | 32.34344 | 1.164544 | 0.435612 | 2.673354 | 0.00751 | 0.019175 |
| CNAG_04671 | 1418.333 | -0.52698 | 0.142425 | -3.70008 | 0.000216 | 0.000863 |
| CNAG_04669 | 947.5293 | -0.33685 | 0.14102 | -2.3887 | 0.016908 | 0.038562 |
| CNAG_04668 | 953.6871 | 0.427581 | 0.180961 | 2.362832 | 0.018136 | 0.041021 |
| CNAG_04666 | 7206.183 | 0.751054 | 0.136141 | 5.516759 | 3.45E-08 | 3.32E-07 |
| CNAG_04664 | 890.6925 | -0.44749 | 0.150675 | -2.96988 | 0.002979 | 0.00864 |
| CNAG_04663 | 526.1937 | 1.043223 | 0.178068 | 5.858582 | 4.67E-09 | 5.17E-08 |
| CNAG_08000 | 64.14481 | 1.608939 | 0.278586 | 5.775375 | 7.68E-09 | 8.21E-08 |
| CNAG_12933 | 30.20256 | -0.93286 | 0.395291 | -2.35994 | 0.018278 | 0.041295 |
| CNAG_04658 | 565.7853 | 0.826265 | 0.27237 | 3.033614 | 0.002416 | 0.007213 |
| CNAG_04657 | 2043.343 | 2.007048 | 0.240739 | 8.337025 | 7.62E-17 | 2.57E-15 |
| CNAG_04656 | 1379.29 | 0.335514 | 0.13339 | 2.515289 | 0.011893 | 0.028534 |
| CNAG_04655 | 1163.806 | -1.2694 | 0.1867 | -6.79914 | 1.05E-11 | 1.84E-10 |
| CNAG_04653 | 281.7125 | -0.68758 | 0.195833 | -3.51103 | 0.000446 | 0.001634 |
| CNAG_04652 | 2520.48 | 0.661963 | 0.232454 | 2.847715 | 0.004403 | 0.012137 |
| CNAG_04651 | 4251.675 | 0.981136 | 0.159646 | 6.145692 | 7.96E-10 | 1.03E-08 |
| CNAG_04649 | 512.3606 | 0.577671 | 0.143424 | 4.027729 | 5.63E-05 | 0.000261 |
| CNAG_04646 | 562.9894 | -0.72818 | 0.231673 | -3.14314 | 0.001671 | 0.005212 |
| CNAG_07857 | 3978.559 | -0.52594 | 0.124554 | -4.22258 | 2.42E-05 | 0.000123 |
| CNAG_04638 | 297.7611 | 1.555568 | 0.229348 | 6.782574 | 1.18E-11 | 2.04E-10 |
| CNAG_04636 | 1950.657 | 0.387564 | 0.143545 | 2.699956 | 0.006935 | 0.017973 |
| CNAG_04635 | 4102.406 | 1.672148 | 0.21861 | 7.648995 | 2.03E-14 | 5.18E-13 |
| CNAG_04634 | 1160.639 | 0.960776 | 0.156004 | 6.158663 | 7.34E-10 | 9.49E-09 |
| CNAG_12937 | 26.7104 | 1.130024 | 0.47875 | 2.360366 | 0.018257 | 0.04126 |
| CNAG_04632 | 3988.155 | -0.68153 | 0.336679 | -2.02427 | 0.042942 | 0.085432 |
| CNAG_04630 | 2940.085 | 1.348506 | 0.258484 | 5.216972 | 1.82E-07 | 1.48E-06 |
| CNAG_04628 | 3006.52 | -0.45575 | 0.213841 | -2.13127 | 0.033067 | 0.068301 |
| CNAG_04626 | 181.4537 | 1.141911 | 0.239733 | 4.763266 | 1.90E-06 | 1.25E-05 |
| CNAG_04625 | 5731.083 | 0.915592 | 0.158756 | 5.767293 | 8.06E-09 | 8.56E-08 |
| CNAG_04622 | 411.3502 | 1.429033 | 0.250946 | 5.694575 | 1.24E-08 | 1.27E-07 |
| CNAG_04618 | 763.8307 | -1.13675 | 0.208573 | -5.45015 | 5.03E-08 | 4.61E-07 |
| CNAG_04617 | 221.3967 | -0.57542 | 0.196834 | -2.92337 | 0.003463 | 0.009827 |
| CNAG_04615 | 266.5174 | -0.78846 | 0.199206 | -3.95799 | 7.56E-05 | 0.00034 |
| CNAG_04614 | 251.7462 | 0.570986 | 0.168065 | 3.397406 | 0.00068 | 0.002369 |
| CNAG_04613 | 3011.917 | -0.38484 | 0.195049 | -1.97306 | 0.048489 | 0.094737 |
| CNAG_04612 | 2212.719 | -0.99444 | 0.216805 | -4.58677 | 4.50E-06 | 2.74E-05 |
| CNAG_04611 | 2109.229 | 0.631719 | 0.235831 | 2.678688 | 0.007391 | 0.018945 |
| CNAG_07858 | 19.91401 | 1.040118 | 0.460582 | 2.258268 | 0.023929 | 0.051965 |

| | | | | | | |
|---|---|---|---|---|---|---|
| CNAG_04610 | 21.50288 | 2.005314 | 0.457803 | 4.380301 | 1.19E-05 | 6.53E-05 |
| CNAG_04606 | 517.4225 | 1.859704 | 0.195268 | 9.523859 | 1.67E-21 | 8.90E-20 |
| CNAG_04605 | 5101.638 | -0.80411 | 0.185698 | -4.3302 | 1.49E-05 | 7.96E-05 |
| CNAG_04604 | 3990.749 | -0.7704 | 0.17115 | -4.50131 | 6.75E-06 | 3.94E-05 |
| CNAG_04603 | 434.0193 | -0.4545 | 0.1854 | -2.45142 | 0.014229 | 0.033333 |
| CNAG_04601 | 19395.27 | -0.74379 | 0.121731 | -6.11008 | 9.96E-10 | 1.26E-08 |
| CNAG_04600 | 1059.925 | 0.604237 | 0.234664 | 2.5749 | 0.010027 | 0.024667 |
| CNAG_04598 | 275.0662 | 0.934499 | 0.160112 | 5.836539 | 5.33E-09 | 5.85E-08 |
| CNAG_04589 | 939.2177 | 1.940622 | 0.163341 | 11.88078 | 1.49E-32 | 1.69E-30 |
| CNAG_04588 | 1532.838 | -0.5618 | 0.215047 | -2.61244 | 0.00899 | 0.022393 |
| CNAG_04587 | 5083.775 | 1.577124 | 0.222926 | 7.074668 | 1.50E-12 | 3.04E-11 |
| CNAG_04585 | 243.9033 | 1.696977 | 0.238001 | 7.130119 | 1.00E-12 | 2.08E-11 |
| CNAG_04584 | 5187.429 | -0.74985 | 0.211972 | -3.53751 | 0.000404 | 0.001498 |
| CNAG_04582 | 254.7639 | 0.70133 | 0.171159 | 4.097538 | 4.18E-05 | 0.0002 |
| CNAG_04576 | 310.6479 | 1.035681 | 0.246642 | 4.199118 | 2.68E-05 | 0.000135 |
| CNAG_12946 | 69.79461 | 1.117678 | 0.305898 | 3.653759 | 0.000258 | 0.001014 |
| CNAG_07862 | 650.7369 | -0.84039 | 0.14843 | -5.66184 | 1.50E-08 | 1.52E-07 |
| CNAG_07863 | 12337.62 | 0.733073 | 0.201774 | 3.633138 | 0.00028 | 0.001091 |
| CNAG_04571 | 890.6342 | -1.1452 | 0.152504 | -7.50932 | 5.94E-14 | 1.43E-12 |
| CNAG_07865 | 2833.672 | 0.636232 | 0.144411 | 4.405707 | 1.05E-05 | 5.85E-05 |
| CNAG_04567 | 564.1076 | -0.35454 | 0.153952 | -2.30291 | 0.021284 | 0.046992 |
| CNAG_04566 | 9218.959 | 0.402842 | 0.13422 | 3.00136 | 0.002688 | 0.007922 |
| CNAG_07866 | 426.2796 | -0.5883 | 0.184069 | -3.19606 | 0.001393 | 0.004435 |
| CNAG_04564 | 434.112 | 0.499439 | 0.17607 | 2.836593 | 0.00456 | 0.012507 |
| CNAG_04561 | 512.4147 | 0.471382 | 0.218591 | 2.156453 | 0.031048 | 0.064719 |
| CNAG_04553 | 61.51869 | 2.01119 | 0.306743 | 6.556596 | 5.50E-11 | 8.41E-10 |
| CNAG_04552 | 13.66016 | 1.36528 | 0.568663 | 2.400861 | 0.016357 | 0.03752 |
| CNAG_07868 | 291.3529 | -0.71336 | 0.208914 | -3.41461 | 0.000639 | 0.002245 |
| CNAG_08001 | 74.55993 | -0.77265 | 0.26237 | -2.94487 | 0.003231 | 0.009238 |
| CNAG_01464 | 4946.695 | -1.20855 | 0.204593 | -5.9071 | 3.48E-09 | 4.01E-08 |
| CNAG_12950 | 10.50614 | -3.60016 | 0.825779 | -4.35972 | 1.30E-05 | 7.11E-05 |
| CNAG_01465 | 347.5617 | 0.875752 | 0.364976 | 2.399476 | 0.016419 | 0.037608 |
| CNAG_01466 | 773.9926 | 0.958524 | 0.186467 | 5.140438 | 2.74E-07 | 2.12E-06 |
| CNAG_01467 | 521.6224 | -0.28667 | 0.133993 | -2.13943 | 0.032401 | 0.067186 |
| CNAG_01468 | 142.7591 | -0.76044 | 0.370801 | -2.05082 | 0.040285 | 0.080895 |
| CNAG_12951 | 87.52462 | 1.046848 | 0.260338 | 4.021105 | 5.79E-05 | 0.000268 |
| CNAG_01470 | 3853.7 | -1.56367 | 0.203234 | -7.69391 | 1.43E-14 | 3.68E-13 |
| CNAG_01471 | 1215.944 | -0.87407 | 0.163948 | -5.33141 | 9.75E-08 | 8.41E-07 |
| CNAG_01473 | 181.159 | 1.100167 | 0.212276 | 5.182708 | 2.19E-07 | 1.73E-06 |
| CNAG_12954 | 116.67 | -0.93883 | 0.239793 | -3.91517 | 9.03E-05 | 0.0004 |
| CNAG_01475 | 10269.54 | -0.3445 | 0.128421 | -2.68256 | 0.007306 | 0.01877 |
| CNAG_01476 | 2691.637 | -0.46515 | 0.123919 | -3.75362 | 0.000174 | 0.000718 |
| CNAG_01480 | 32206.08 | -0.84953 | 0.239943 | -3.54055 | 0.000399 | 0.001483 |
| CNAG_01481 | 270.8942 | 1.1674 | 0.203292 | 5.742472 | 9.33E-09 | 9.83E-08 |
| CNAG_01486 | 45072.38 | -0.95148 | 0.254868 | -3.73324 | 0.000189 | 0.000771 |
| CNAG_01487 | 26.08257 | 1.058653 | 0.465525 | 2.274108 | 0.022959 | 0.050258 |
| CNAG_01489 | 150.505 | -1.24095 | 0.296519 | -4.18505 | 2.85E-05 | 0.000142 |

| | | | | | | |
|---|---|---|---|---|---|---|
| CNAG_01490 | 472.6067 | 0.410589 | 0.170412 | 2.409385 | 0.015979 | 0.036783 |
| CNAG_01493 | 1058.859 | 1.237131 | 0.232621 | 5.318224 | 1.05E-07 | 8.97E-07 |
| CNAG_12957 | 16.83516 | -1.04886 | 0.48145 | -2.17854 | 0.029366 | 0.061714 |
| CNAG_01495 | 1087.304 | 1.126939 | 0.181765 | 6.199963 | 5.65E-10 | 7.39E-09 |
| CNAG_01498 | 461.0215 | -0.74071 | 0.195355 | -3.79161 | 0.00015 | 0.000623 |
| CNAG_08003 | 538.8346 | -0.58502 | 0.212783 | -2.74936 | 0.005971 | 0.015842 |
| CNAG_01506 | 98.72248 | 1.333009 | 0.22586 | 5.901928 | 3.59E-09 | 4.10E-08 |
| CNAG_01507 | 567.5718 | -0.33325 | 0.164486 | -2.02598 | 0.042767 | 0.085112 |
| CNAG_01508 | 2111.373 | -0.62915 | 0.139825 | -4.49956 | 6.81E-06 | 3.97E-05 |
| CNAG_01509 | 544.6401 | -0.68329 | 0.214393 | -3.18709 | 0.001437 | 0.004551 |
| CNAG_01511 | 38.13081 | 1.913081 | 0.371672 | 5.147225 | 2.64E-07 | 2.05E-06 |
| CNAG_01512 | 471.8925 | 1.524159 | 0.200353 | 7.607383 | 2.80E-14 | 7.01E-13 |
| CNAG_07589 | 89.20854 | -0.72951 | 0.27099 | -2.69202 | 0.007102 | 0.01834 |
| CNAG_07591 | 2266.289 | 1.119483 | 0.160316 | 6.982993 | 2.89E-12 | 5.51E-11 |
| CNAG_01518 | 1526.5 | -0.46834 | 0.171732 | -2.72717 | 0.006388 | 0.016791 |
| CNAG_01519 | 595.5433 | 0.868876 | 0.156206 | 5.562359 | 2.66E-08 | 2.62E-07 |
| CNAG_01520 | 1300.224 | -0.30392 | 0.149156 | -2.03758 | 0.041592 | 0.0831 |
| CNAG_01522 | 1461.073 | -0.48513 | 0.149184 | -3.25191 | 0.001146 | 0.003738 |
| CNAG_01523 | 5297.082 | -0.32988 | 0.167454 | -1.96998 | 0.048841 | 0.095272 |
| CNAG_01524 | 2645.229 | 0.860279 | 0.167632 | 5.131937 | 2.87E-07 | 2.21E-06 |
| CNAG_01525 | 311.831 | 0.430054 | 0.177058 | 2.42889 | 0.015145 | 0.035178 |
| CNAG_12967 | 44.56283 | 1.264563 | 0.317385 | 3.984321 | 6.77E-05 | 0.000308 |
| CNAG_12968 | 77.04696 | 1.972437 | 0.273675 | 7.207218 | 5.71E-13 | 1.22E-11 |
| CNAG_01528 | 975.7016 | -0.7543 | 0.126667 | -5.95502 | 2.60E-09 | 3.07E-08 |
| CNAG_01529 | 1099.234 | -0.46385 | 0.170681 | -2.71766 | 0.006575 | 0.017173 |
| CNAG_01531 | 928.6669 | -0.66397 | 0.181286 | -3.66257 | 0.00025 | 0.000983 |
| CNAG_01534 | 174.4485 | 1.147795 | 0.256279 | 4.478697 | 7.51E-06 | 4.32E-05 |
| CNAG_01538 | 200.5905 | 0.971876 | 0.28297 | 3.43455 | 0.000594 | 0.002106 |
| CNAG_01539 | 25635.48 | -0.92866 | 0.168328 | -5.51698 | 3.45E-08 | 3.32E-07 |
| CNAG_01542 | 912.0421 | -0.91175 | 0.199168 | -4.57779 | 4.70E-06 | 2.84E-05 |
| CNAG_01544 | 5738.517 | -0.95376 | 0.216561 | -4.40413 | 1.06E-05 | 5.89E-05 |
| CNAG_01545 | 588.1984 | -0.43593 | 0.171546 | -2.54118 | 0.011048 | 0.026789 |
| CNAG_01549 | 2882.854 | -0.54599 | 0.121271 | -4.50221 | 6.72E-06 | 3.93E-05 |
| CNAG_01550 | 1271.396 | -0.63782 | 0.163538 | -3.90016 | 9.61E-05 | 0.000421 |
| CNAG_01551 | 1225.814 | -0.96371 | 0.205194 | -4.69659 | 2.65E-06 | 1.70E-05 |
| CNAG_01552 | 1371.376 | -0.62327 | 0.269007 | -2.31693 | 0.020507 | 0.045504 |
| CNAG_01555 | 86.27468 | 0.570557 | 0.268548 | 2.124603 | 0.03362 | 0.06919 |
| CNAG_01556 | 1290.581 | 0.654427 | 0.166408 | 3.932676 | 8.40E-05 | 0.000375 |
| CNAG_01558 | 26931.68 | 1.419912 | 0.149809 | 9.478168 | 2.59E-21 | 1.34E-19 |
| CNAG_01559 | 1583.15 | -0.74316 | 0.198296 | -3.74774 | 0.000178 | 0.000733 |
| CNAG_01560 | 99.67169 | 1.054133 | 0.265849 | 3.96516 | 7.33E-05 | 0.000332 |
| CNAG_01561 | 2667.63 | 0.599335 | 0.248231 | 2.414427 | 0.01576 | 0.036372 |
| CNAG_01562 | 26070.5 | 0.812391 | 0.137327 | 5.915742 | 3.30E-09 | 3.82E-08 |
| CNAG_01564 | 2884.236 | -0.8468 | 0.159033 | -5.32467 | 1.01E-07 | 8.69E-07 |
| CNAG_01566 | 796.5241 | -0.97133 | 0.253471 | -3.83213 | 0.000127 | 0.000539 |
| CNAG_01569 | 964.2754 | -0.39234 | 0.172451 | -2.27506 | 0.022902 | 0.05016 |
| CNAG_01577 | 61396.07 | -0.53854 | 0.161529 | -3.33405 | 0.000856 | 0.002881 |

| | | | | | | |
|---|---|---|---|---|---|---|
| CNAG_01580 | 2661.964 | 0.250747 | 0.120811 | 2.075524 | 0.037938 | 0.076965 |
| CNAG_01583 | 2865.318 | -0.37133 | 0.184952 | -2.0077 | 0.044676 | 0.088241 |
| CNAG_01584 | 1381.585 | 1.267793 | 0.214592 | 5.907914 | 3.46E-09 | 4.00E-08 |
| CNAG_01585 | 1384.189 | 2.578735 | 0.231998 | 11.11533 | 1.06E-28 | 8.84E-27 |
| CNAG_01586 | 16102.7 | -0.69916 | 0.235868 | -2.96418 | 0.003035 | 0.008763 |
| CNAG_01587 | 2217.384 | -0.54869 | 0.163978 | -3.34612 | 0.00082 | 0.002782 |
| CNAG_12974 | 58.37785 | 2.43877 | 0.442467 | 5.511757 | 3.55E-08 | 3.41E-07 |
| CNAG_01588 | 6183.824 | 0.884502 | 0.197517 | 4.478104 | 7.53E-06 | 4.32E-05 |
| CNAG_01590 | 801.3 | -0.67565 | 0.165491 | -4.0827 | 4.45E-05 | 0.000211 |
| CNAG_01592 | 192.9267 | -0.68533 | 0.20486 | -3.34536 | 0.000822 | 0.002786 |
| CNAG_01593 | 2798.235 | -0.86504 | 0.24113 | -3.58747 | 0.000334 | 0.001272 |
| CNAG_01594 | 7670.545 | -0.58256 | 0.163129 | -3.57114 | 0.000355 | 0.001341 |
| CNAG_01595 | 654.2025 | -0.9789 | 0.21123 | -4.63426 | 3.58E-06 | 2.23E-05 |
| CNAG_01596 | 1207.046 | -0.9603 | 0.234646 | -4.09255 | 4.27E-05 | 0.000204 |
| CNAG_01598 | 4279.535 | -1.11925 | 0.280855 | -3.98515 | 6.74E-05 | 0.000307 |
| CNAG_01599 | 854.0656 | -1.12463 | 0.383527 | -2.93234 | 0.003364 | 0.009585 |
| CNAG_01600 | 1531.399 | -0.77509 | 0.239788 | -3.23238 | 0.001228 | 0.00398 |
| CNAG_01601 | 327.0946 | 0.531089 | 0.17143 | 3.097997 | 0.001948 | 0.005959 |
| CNAG_01603 | 6919.337 | -0.50726 | 0.221243 | -2.29275 | 0.021862 | 0.048061 |
| CNAG_01605 | 638.5882 | 1.14895 | 0.189821 | 6.052815 | 1.42E-09 | 1.76E-08 |
| CNAG_01607 | 1355.565 | -0.61921 | 0.206378 | -3.00039 | 0.002696 | 0.007944 |
| CNAG_01608 | 1355.248 | -0.65975 | 0.116379 | -5.66897 | 1.44E-08 | 1.47E-07 |
| CNAG_01609 | 2620.534 | -0.99123 | 0.126163 | -7.85679 | 3.94E-15 | 1.09E-13 |
| CNAG_01610 | 1717.436 | 0.804667 | 0.31236 | 2.576093 | 0.009992 | 0.024612 |
| CNAG_01611 | 732.3153 | 1.017557 | 0.180238 | 5.645635 | 1.65E-08 | 1.66E-07 |
| CNAG_01613 | 2233.516 | 0.690656 | 0.255065 | 2.707767 | 0.006774 | 0.017618 |
| CNAG_01614 | 1626.214 | -0.2596 | 0.116819 | -2.22226 | 0.026265 | 0.056119 |
| CNAG_01615 | 1104.75 | 0.541965 | 0.195544 | 2.771579 | 0.005579 | 0.014965 |
| CNAG_07594 | 894.3915 | 0.359392 | 0.161848 | 2.220551 | 0.026381 | 0.056351 |
| CNAG_01619 | 868.4353 | 0.454585 | 0.139268 | 3.2641 | 0.001098 | 0.003594 |
| CNAG_01620 | 839.5024 | -0.99857 | 0.19591 | -5.09708 | 3.45E-07 | 2.62E-06 |
| CNAG_12978 | 162.4187 | 1.208983 | 0.354505 | 3.410338 | 0.000649 | 0.002275 |
| CNAG_01621 | 7336.807 | 1.780353 | 0.246453 | 7.223892 | 5.05E-13 | 1.09E-11 |
| CNAG_01625 | 563.4569 | -0.78731 | 0.162133 | -4.85592 | 1.20E-06 | 8.13E-06 |
| CNAG_01626 | 2693.83 | -0.51489 | 0.125557 | -4.10083 | 4.12E-05 | 0.000197 |
| CNAG_01628 | 21678.15 | -0.95414 | 0.232552 | -4.10292 | 4.08E-05 | 0.000196 |
| CNAG_01629 | 675.7061 | -1.20717 | 0.259365 | -4.65434 | 3.25E-06 | 2.04E-05 |
| CNAG_01631 | 1527.536 | -0.34975 | 0.116966 | -2.99018 | 0.002788 | 0.008172 |
| CNAG_01632 | 851.6002 | 0.439262 | 0.142711 | 3.077991 | 0.002084 | 0.006347 |
| CNAG_01634 | 3288.995 | -0.35294 | 0.110705 | -3.18807 | 0.001432 | 0.00454 |
| CNAG_01638 | 1069.536 | -0.35915 | 0.149438 | -2.40332 | 0.016247 | 0.037301 |
| CNAG_01640 | 3996.508 | 0.914862 | 0.263856 | 3.467273 | 0.000526 | 0.001894 |
| CNAG_01644 | 1818.637 | -1.045 | 0.157017 | -6.65531 | 2.83E-11 | 4.54E-10 |
| CNAG_01645 | 97.83634 | 1.273249 | 0.236687 | 5.379464 | 7.47E-08 | 6.57E-07 |
| CNAG_01648 | 19735.94 | -1.31832 | 0.230806 | -5.71182 | 1.12E-08 | 1.16E-07 |
| CNAG_01650 | 1203.659 | -0.63234 | 0.1455 | -4.34595 | 1.39E-05 | 7.52E-05 |
| CNAG_01653 | 3161.175 | 1.635958 | 0.359031 | 4.556592 | 5.20E-06 | 3.12E-05 |

| | | | | | | |
|---|---|---|---|---|---|---|
| CNAG_01655 | 5403.541 | 0.697862 | 0.245452 | 2.843176 | 0.004467 | 0.012285 |
| CNAG_01660 | 1168.888 | -0.27276 | 0.130284 | -2.09359 | 0.036296 | 0.074119 |
| CNAG_12982 | 6.124772 | 2.394698 | 0.964276 | 2.483416 | 0.013013 | 0.03082 |
| CNAG_01665 | 532.6417 | -0.71028 | 0.187239 | -3.79343 | 0.000149 | 0.00062 |
| CNAG_01666 | 1655.507 | -0.82463 | 0.283371 | -2.91007 | 0.003613 | 0.010196 |
| CNAG_01667 | 938.2104 | -0.50227 | 0.168292 | -2.98453 | 0.00284 | 0.008303 |
| CNAG_01668 | 18.47569 | 1.166468 | 0.513949 | 2.269617 | 0.023231 | 0.050754 |
| CNAG_01669 | 1001.512 | -0.32943 | 0.152198 | -2.16447 | 0.030428 | 0.063644 |
| CNAG_01671 | 150.1181 | 0.620281 | 0.183268 | 3.384556 | 0.000713 | 0.002462 |
| CNAG_01674 | 66.85076 | -0.96065 | 0.257908 | -3.72479 | 0.000195 | 0.000792 |
| CNAG_01679 | 3228.969 | -0.44077 | 0.183568 | -2.40112 | 0.016345 | 0.037504 |
| CNAG_01680 | 80.32814 | 0.710074 | 0.24124 | 2.943431 | 0.003246 | 0.009274 |
| CNAG_12985 | 1380.675 | -0.67367 | 0.231355 | -2.91185 | 0.003593 | 0.010146 |
| CNAG_01681 | 9303.489 | -0.96155 | 0.15586 | -6.16934 | 6.86E-10 | 8.90E-09 |
| CNAG_07596 | 191.8114 | 2.114417 | 0.231033 | 9.152 | 5.59E-20 | 2.54E-18 |
| CNAG_01683 | 5584.181 | 0.317564 | 0.135172 | 2.349334 | 0.018807 | 0.042227 |
| CNAG_01686 | 4591.883 | 1.014989 | 0.184969 | 5.487359 | 4.08E-08 | 3.82E-07 |
| CNAG_01687 | 1638.789 | -0.49074 | 0.197703 | -2.48223 | 0.013056 | 0.030904 |
| CNAG_01688 | 5215.015 | -0.36223 | 0.177121 | -2.04508 | 0.040847 | 0.08188 |
| CNAG_12991 | 97.32666 | 0.62253 | 0.228003 | 2.730359 | 0.006327 | 0.016651 |
| CNAG_01690 | 856.2562 | 0.876936 | 0.153859 | 5.699598 | 1.20E-08 | 1.24E-07 |
| CNAG_01691 | 1682.381 | 1.33037 | 0.18753 | 7.094167 | 1.30E-12 | 2.66E-11 |
| CNAG_12992 | 90.84264 | 1.999035 | 0.286476 | 6.978012 | 2.99E-12 | 5.66E-11 |
| CNAG_01692 | 2027.528 | 0.288371 | 0.129216 | 2.231696 | 0.025635 | 0.055002 |
| CNAG_01693 | 1069.247 | -0.61127 | 0.150149 | -4.07107 | 4.68E-05 | 0.000221 |
| CNAG_12993 | 253.8494 | 4.10606 | 1.406564 | 2.919213 | 0.003509 | 0.009937 |
| CNAG_01695 | 163.9948 | 1.785078 | 0.197031 | 9.059902 | 1.31E-19 | 5.76E-18 |
| CNAG_01699 | 728.6099 | 0.463486 | 0.189766 | 2.442406 | 0.01459 | 0.034047 |
| CNAG_01702 | 440.5116 | 0.391982 | 0.158334 | 2.47567 | 0.013299 | 0.031347 |
| CNAG_01703 | 280.1995 | 0.565661 | 0.159483 | 3.546831 | 0.00039 | 0.001454 |
| CNAG_01704 | 5098.194 | 0.499408 | 0.176215 | 2.834087 | 0.004596 | 0.012584 |
| CNAG_01713 | 1812.328 | 0.792241 | 0.24823 | 3.191567 | 0.001415 | 0.004499 |
| CNAG_01714 | 26.83944 | 1.017415 | 0.382663 | 2.658772 | 0.007843 | 0.019922 |
| CNAG_01715 | 3411.854 | -0.60995 | 0.205993 | -2.96102 | 0.003066 | 0.00883 |
| CNAG_01716 | 1327.673 | -0.76756 | 0.127338 | -6.02771 | 1.66E-09 | 2.04E-08 |
| CNAG_01719 | 130.4212 | 0.70054 | 0.221458 | 3.163315 | 0.00156 | 0.004901 |
| CNAG_01721 | 1859.677 | -1.6264 | 0.189259 | -8.5935 | 8.44E-18 | 3.09E-16 |
| CNAG_01722 | 7190.533 | 0.727328 | 0.275129 | 2.643584 | 0.008203 | 0.020713 |
| CNAG_01723 | 1405.706 | 0.42312 | 0.152707 | 2.770797 | 0.005592 | 0.014991 |
| CNAG_12996 | 10.51626 | 1.397013 | 0.620283 | 2.25222 | 0.024308 | 0.052631 |
| CNAG_01727 | 78863.58 | 0.660382 | 0.14117 | 4.677928 | 2.90E-06 | 1.85E-05 |
| CNAG_12998 | 21.77811 | -1.44942 | 0.584931 | -2.47793 | 0.013215 | 0.031214 |
| CNAG_01732 | 2673.698 | 0.588404 | 0.149989 | 3.922968 | 8.75E-05 | 0.000388 |
| CNAG_01733 | 6998.083 | -0.51785 | 0.223025 | -2.32192 | 0.020237 | 0.045006 |
| CNAG_01735 | 3066.854 | 2.13781 | 0.189363 | 11.2895 | 1.48E-29 | 1.35E-27 |
| CNAG_01737 | 3761.762 | -0.86482 | 0.138566 | -6.24125 | 4.34E-10 | 5.82E-09 |
| CNAG_01739 | 1025.3 | 0.91073 | 0.200929 | 4.532602 | 5.83E-06 | 3.44E-05 |

| | | | | | | |
|---|---|---|---|---|---|---|
| CNAG_01742 | 333.4892 | 1.485815 | 0.174803 | 8.499939 | 1.90E-17 | 6.67E-16 |
| CNAG_01743 | 832.315 | 1.753221 | 0.18996 | 9.22942 | 2.72E-20 | 1.29E-18 |
| CNAG_01744 | 9733.519 | 0.390838 | 0.128035 | 3.052596 | 0.002269 | 0.006831 |
| CNAG_01745 | 10666.05 | 0.646276 | 0.118892 | 5.435841 | 5.45E-08 | 4.96E-07 |
| CNAG_01746 | 494.7675 | 1.100219 | 0.166221 | 6.619034 | 3.62E-11 | 5.67E-10 |
| CNAG_07600 | 727.086 | 1.067186 | 0.247765 | 4.307259 | 1.65E-05 | 8.75E-05 |
| CNAG_01750 | 14499.68 | 2.373868 | 0.173414 | 13.68904 | 1.18E-42 | 2.29E-40 |
| CNAG_01751 | 2789.388 | 2.211216 | 0.255517 | 8.653888 | 4.98E-18 | 1.85E-16 |
| CNAG_01752 | 6609.959 | -0.80258 | 0.173 | -4.63918 | 3.50E-06 | 2.18E-05 |
| CNAG_01753 | 6922.828 | 0.972594 | 0.15144 | 6.422292 | 1.34E-10 | 1.96E-09 |
| CNAG_01755 | 2840.915 | -0.61002 | 0.12456 | -4.89736 | 9.71E-07 | 6.74E-06 |
| CNAG_01758 | 107.285 | 0.480047 | 0.212091 | 2.263398 | 0.023611 | 0.051444 |
| CNAG_13000 | 12.45636 | 1.733108 | 0.598628 | 2.895133 | 0.00379 | 0.010608 |
| CNAG_08008 | 685.3608 | 0.792925 | 0.275413 | 2.879038 | 0.003989 | 0.011086 |
| CNAG_01764 | 1179.533 | 0.424805 | 0.177007 | 2.399932 | 0.016398 | 0.037583 |
| CNAG_01765 | 1275.854 | 0.371633 | 0.178779 | 2.078725 | 0.037643 | 0.076459 |
| CNAG_07604 | 357.6251 | 0.647836 | 0.168487 | 3.845023 | 0.000121 | 0.000515 |
| CNAG_01771 | 1620.398 | 0.505034 | 0.224408 | 2.250515 | 0.024416 | 0.052835 |
| CNAG_01772 | 2525.562 | -0.92568 | 0.124076 | -7.46059 | 8.61E-14 | 2.00E-12 |
| CNAG_01773 | 719.9459 | 0.283204 | 0.133754 | 2.117357 | 0.03423 | 0.07039 |
| CNAG_13002 | 11.52858 | 1.767611 | 0.612514 | 2.885832 | 0.003904 | 0.010876 |
| CNAG_01776 | 217.7817 | 0.495744 | 0.18783 | 2.639331 | 0.008307 | 0.020941 |
| CNAG_01779 | 660.5379 | 0.328817 | 0.1397 | 2.353743 | 0.018585 | 0.041819 |
| CNAG_13004 | 31.03489 | 0.77596 | 0.384096 | 2.020225 | 0.04336 | 0.086135 |
| CNAG_01789 | 2806.358 | -0.67168 | 0.204039 | -3.2919 | 0.000995 | 0.003304 |
| CNAG_01791 | 2103.062 | -0.41484 | 0.131833 | -3.14671 | 0.001651 | 0.005161 |
| CNAG_01796 | 408.2626 | 0.843763 | 0.149223 | 5.654364 | 1.56E-08 | 1.58E-07 |
| CNAG_01797 | 1659.395 | -0.81055 | 0.186606 | -4.34365 | 1.40E-05 | 7.59E-05 |
| CNAG_01799 | 7086.034 | -0.74724 | 0.231329 | -3.2302 | 0.001237 | 0.004005 |
| CNAG_01800 | 1780.139 | -1.63441 | 0.316308 | -5.16715 | 2.38E-07 | 1.87E-06 |
| CNAG_01803 | 559.9347 | 0.406334 | 0.169567 | 2.3963 | 0.016562 | 0.037891 |
| CNAG_13007 | 173.4295 | 0.811536 | 0.26435 | 3.069928 | 0.002141 | 0.006499 |
| CNAG_01806 | 27.58516 | 1.175958 | 0.420068 | 2.799451 | 0.005119 | 0.01386 |
| CNAG_01808 | 2015.949 | -0.54532 | 0.123454 | -4.41716 | 1.00E-05 | 5.59E-05 |
| CNAG_01809 | 1359.588 | -0.57154 | 0.170918 | -3.34393 | 0.000826 | 0.002794 |
| CNAG_01810 | 549.7051 | -0.82989 | 0.194061 | -4.27643 | 1.90E-05 | 9.92E-05 |
| CNAG_01812 | 3096.691 | -0.83624 | 0.192904 | -4.33503 | 1.46E-05 | 7.85E-05 |
| CNAG_01813 | 15920.01 | -0.59307 | 0.205895 | -2.88044 | 0.003971 | 0.011042 |
| CNAG_01816 | 148.0092 | -0.39232 | 0.193055 | -2.03218 | 0.042135 | 0.083933 |
| CNAG_01818 | 1599.59 | -0.45267 | 0.131432 | -3.44414 | 0.000573 | 0.002043 |
| CNAG_01820 | 24220.26 | -0.67195 | 0.138703 | -4.84452 | 1.27E-06 | 8.58E-06 |
| CNAG_01821 | 1403.276 | 1.70679 | 0.20802 | 8.204938 | 2.31E-16 | 7.39E-15 |
| CNAG_01822 | 1487.571 | -0.30794 | 0.130223 | -2.36471 | 0.018044 | 0.040848 |
| CNAG_01823 | 3675.666 | 0.457989 | 0.178564 | 2.564854 | 0.010322 | 0.025314 |
| CNAG_13008 | 154.172 | -0.68029 | 0.287455 | -2.36661 | 0.017952 | 0.040651 |
| CNAG_01824 | 4581.56 | 0.563584 | 0.210667 | 2.675232 | 0.007468 | 0.019115 |
| CNAG_01828 | 1009.336 | -0.59187 | 0.153086 | -3.86627 | 0.000111 | 0.000475 |

| | | | | | | |
|---|---|---|---|---|---|---|
| CNAG_07607 | 1308.38 | 0.454883 | 0.133669 | 3.403053 | 0.000666 | 0.002326 |
| CNAG_07609 | 1602.109 | -0.77116 | 0.13878 | -5.5567 | 2.75E-08 | 2.70E-07 |
| CNAG_01832 | 711.4152 | 0.338037 | 0.142019 | 2.380232 | 0.017302 | 0.039347 |
| CNAG_01836 | 41.2134 | -1.22568 | 0.348172 | -3.52031 | 0.000431 | 0.001584 |
| CNAG_13010 | 267.1419 | 1.448356 | 0.228177 | 6.34751 | 2.19E-10 | 3.09E-09 |
| CNAG_01840 | 23260.23 | -0.53023 | 0.112723 | -4.70383 | 2.55E-06 | 1.65E-05 |
| CNAG_01841 | 75.28866 | 0.873464 | 0.298016 | 2.930926 | 0.00338 | 0.009625 |
| CNAG_01842 | 1559.691 | -0.87187 | 0.226611 | -3.84742 | 0.000119 | 0.00051 |
| CNAG_01844 | 523.4775 | 0.386498 | 0.157913 | 2.447547 | 0.014383 | 0.033644 |
| CNAG_01847 | 284.4879 | 2.006729 | 0.25015 | 8.022097 | 1.04E-15 | 3.14E-14 |
| CNAG_01848 | 3872.629 | 0.809486 | 0.278935 | 2.902059 | 0.003707 | 0.010427 |
| CNAG_01854 | 13944.62 | 0.519616 | 0.141136 | 3.681654 | 0.000232 | 0.000921 |
| CNAG_01856 | 334.7915 | 1.648435 | 0.260198 | 6.335304 | 2.37E-10 | 3.34E-09 |
| CNAG_01858 | 64.53579 | 0.585369 | 0.260855 | 2.244043 | 0.02483 | 0.053511 |
| CNAG_01861 | 2184.606 | 0.978123 | 0.231406 | 4.226865 | 2.37E-05 | 0.000121 |
| CNAG_07610 | 12.14255 | -1.29375 | 0.592469 | -2.18365 | 0.028988 | 0.061049 |
| CNAG_01863 | 4289.887 | -0.55871 | 0.20454 | -2.73156 | 0.006304 | 0.016613 |
| CNAG_13013 | 134.3118 | -1.34215 | 0.313132 | -4.28619 | 1.82E-05 | 9.53E-05 |
| CNAG_01864 | 1262 | -0.78374 | 0.149033 | -5.25886 | 1.45E-07 | 1.20E-06 |
| CNAG_01865 | 1086.812 | -1.36307 | 0.152548 | -8.93536 | 4.06E-19 | 1.72E-17 |
| CNAG_01866 | 934.1104 | 1.053207 | 0.269009 | 3.915142 | 9.04E-05 | 0.0004 |
| CNAG_01867 | 2676.19 | 0.352997 | 0.157227 | 2.245148 | 0.024759 | 0.053402 |
| CNAG_01871 | 1191.987 | -0.48104 | 0.172991 | -2.78071 | 0.005424 | 0.014565 |
| CNAG_01873 | 6.458666 | 3.007629 | 1.018949 | 2.951696 | 0.00316 | 0.009059 |
| CNAG_01874 | 2789.952 | 0.833658 | 0.219473 | 3.798453 | 0.000146 | 0.000611 |
| CNAG_01876 | 1465.299 | 0.571032 | 0.177757 | 3.212427 | 0.001316 | 0.004234 |
| CNAG_01877 | 8343.073 | -0.75724 | 0.19069 | -3.97106 | 7.16E-05 | 0.000324 |
| CNAG_01878 | 1464.657 | 0.451236 | 0.131542 | 3.430345 | 0.000603 | 0.002136 |
| CNAG_01880 | 1202.518 | -0.29179 | 0.144725 | -2.01615 | 0.043784 | 0.086846 |
| CNAG_01882 | 583.0035 | -0.90794 | 0.194273 | -4.6735 | 2.96E-06 | 1.88E-05 |
| CNAG_07611 | 28.76741 | 1.488991 | 0.401575 | 3.707881 | 0.000209 | 0.000841 |
| CNAG_01883 | 121.0179 | 2.136228 | 0.530532 | 4.026579 | 5.66E-05 | 0.000262 |
| CNAG_01884 | 55254.05 | -0.90679 | 0.158377 | -5.72556 | 1.03E-08 | 1.08E-07 |
| CNAG_01885 | 1125.998 | -0.77588 | 0.166886 | -4.64919 | 3.33E-06 | 2.09E-05 |
| CNAG_13015 | 97.00212 | -1.18064 | 0.272221 | -4.33705 | 1.44E-05 | 7.79E-05 |
| CNAG_01889 | 4133.842 | 3.595972 | 0.189135 | 19.01269 | 1.34E-80 | 9.67E-78 |
| CNAG_01890 | 47371.17 | -0.37225 | 0.179248 | -2.07673 | 0.037827 | 0.076794 |
| CNAG_01891 | 749.5437 | 0.433661 | 0.142923 | 3.034226 | 0.002412 | 0.007201 |
| CNAG_01892 | 2442.323 | 0.740204 | 0.269977 | 2.741725 | 0.006112 | 0.016145 |
| CNAG_01893 | 2625.111 | 0.902206 | 0.141945 | 6.356044 | 2.07E-10 | 2.93E-09 |
| CNAG_01894 | 2376.308 | -0.43186 | 0.125939 | -3.42917 | 0.000605 | 0.002141 |
| CNAG_01896 | 16099.69 | 0.935159 | 0.135355 | 6.908958 | 4.88E-12 | 9.02E-11 |
| CNAG_01897 | 5467.897 | -1.01294 | 0.128428 | -7.8872 | 3.09E-15 | 8.68E-14 |
| CNAG_01898 | 1631.079 | -0.71232 | 0.132011 | -5.39596 | 6.82E-08 | 6.08E-07 |
| CNAG_01902 | 2341.736 | -0.7022 | 0.174446 | -4.02533 | 5.69E-05 | 0.000263 |
| CNAG_01903 | 406.0368 | -1.3776 | 0.205187 | -6.71388 | 1.90E-11 | 3.14E-10 |
| CNAG_01904 | 891.5203 | -0.40044 | 0.200803 | -1.99422 | 0.046128 | 0.090771 |

| | | | | | | |
|---|---|---|---|---|---|---|
| CNAG_01906 | 576.2489 | -0.33033 | 0.150168 | -2.19974 | 0.027825 | 0.058976 |
| CNAG_01907 | 1199.082 | -1.08865 | 0.325523 | -3.34432 | 0.000825 | 0.002791 |
| CNAG_01908 | 363.9971 | -0.68862 | 0.192854 | -3.57069 | 0.000356 | 0.001342 |
| CNAG_13021 | 489.8642 | 0.796157 | 0.164223 | 4.848034 | 1.25E-06 | 8.44E-06 |
| CNAG_01912 | 6198.959 | 0.543924 | 0.182559 | 2.979439 | 0.002888 | 0.008421 |
| CNAG_01913 | 284.9271 | 0.7321 | 0.171389 | 4.271558 | 1.94E-05 | 0.000101 |
| CNAG_01914 | 1445.626 | -0.37798 | 0.157223 | -2.4041 | 0.016212 | 0.037242 |
| CNAG_01915 | 10591.02 | 0.424675 | 0.141621 | 2.998672 | 0.002712 | 0.007983 |
| CNAG_13022 | 29.45576 | -0.76089 | 0.368145 | -2.06681 | 0.038752 | 0.078173 |
| CNAG_01917 | 293.7818 | -1.03109 | 0.182902 | -5.63743 | 1.73E-08 | 1.73E-07 |
| CNAG_01919 | 1271.192 | 1.138027 | 0.175447 | 6.486449 | 8.79E-11 | 1.32E-09 |
| CNAG_01920 | 22290.49 | 0.496355 | 0.130828 | 3.793951 | 0.000148 | 0.00062 |
| CNAG_01922 | 987.4754 | 0.574619 | 0.249662 | 2.301581 | 0.021359 | 0.047144 |
| CNAG_01924 | 1425.983 | 0.596423 | 0.193516 | 3.082041 | 0.002056 | 0.006266 |
| CNAG_01925 | 486.6026 | 0.977735 | 0.246882 | 3.960335 | 7.48E-05 | 0.000338 |
| CNAG_01934 | 68.42027 | 0.821358 | 0.25059 | 3.277701 | 0.001047 | 0.003449 |
| CNAG_08011 | 220.0807 | 0.684038 | 0.194131 | 3.523589 | 0.000426 | 0.001569 |
| CNAG_01942 | 634.3924 | 1.057753 | 0.293136 | 3.608403 | 0.000308 | 0.001186 |
| CNAG_01943 | 2575.731 | -0.50625 | 0.168782 | -2.99943 | 0.002705 | 0.007966 |
| CNAG_01947 | 472.3497 | -1.36318 | 0.298049 | -4.57368 | 4.79E-06 | 2.89E-05 |
| CNAG_01949 | 1769.866 | -1.67437 | 0.322076 | -5.19868 | 2.01E-07 | 1.61E-06 |
| CNAG_01951 | 25965.07 | -1.08772 | 0.243033 | -4.47559 | 7.62E-06 | 4.37E-05 |
| CNAG_13034 | 6.381678 | 1.864393 | 0.791568 | 2.355317 | 0.018507 | 0.041694 |
| CNAG_01953 | 544.7344 | 1.058467 | 0.149714 | 7.069937 | 1.55E-12 | 3.13E-11 |
| CNAG_13035 | 69.23935 | 1.400165 | 0.372961 | 3.75419 | 0.000174 | 0.000717 |
| CNAG_01954 | 978.3304 | 5.19529 | 0.270402 | 19.21323 | 2.87E-82 | 2.28E-79 |
| CNAG_01957 | 5500.312 | 0.615605 | 0.177901 | 3.460377 | 0.000539 | 0.001939 |
| CNAG_01959 | 1655.048 | -1.13362 | 0.248887 | -4.55476 | 5.24E-06 | 3.14E-05 |
| CNAG_01960 | 180.1647 | 1.501248 | 0.231894 | 6.473853 | 9.55E-11 | 1.42E-09 |
| CNAG_01963 | 368.1061 | -0.50313 | 0.182541 | -2.75627 | 0.005847 | 0.015553 |
| CNAG_01965 | 76.10516 | -1.33219 | 0.256245 | -5.19889 | 2.00E-07 | 1.61E-06 |
| CNAG_01969 | 215.1733 | 0.483695 | 0.224993 | 2.149819 | 0.03157 | 0.065668 |
| CNAG_01970 | 2508.646 | 0.879952 | 0.159282 | 5.524507 | 3.30E-08 | 3.19E-07 |
| CNAG_01972 | 1949.693 | 0.80987 | 0.215859 | 3.75185 | 0.000176 | 0.000723 |
| CNAG_01974 | 847.4947 | -0.83513 | 0.205031 | -4.07319 | 4.64E-05 | 0.000219 |
| CNAG_01976 | 30345.25 | -1.14359 | 0.26988 | -4.2374 | 2.26E-05 | 0.000115 |
| CNAG_01981 | 8259.497 | -0.4354 | 0.125865 | -3.4593 | 0.000542 | 0.001945 |
| CNAG_01982 | 617.1063 | 0.624306 | 0.166214 | 3.756041 | 0.000173 | 0.000712 |
| CNAG_01983 | 1721.746 | -0.86168 | 0.226778 | -3.79968 | 0.000145 | 0.000608 |
| CNAG_01985 | 4977.067 | -0.2949 | 0.12064 | -2.44449 | 0.014506 | 0.033881 |
| CNAG_01986 | 1524.111 | 1.121601 | 0.190117 | 5.899529 | 3.65E-09 | 4.15E-08 |
| CNAG_13042 | 44.64373 | 1.583411 | 0.400436 | 3.954213 | 7.68E-05 | 0.000345 |
| CNAG_01987 | 2215.157 | 0.560214 | 0.157446 | 3.558139 | 0.000373 | 0.001398 |
| CNAG_01988 | 3597.959 | 1.138726 | 0.277015 | 4.110696 | 3.94E-05 | 0.00019 |
| CNAG_01989 | 212.3376 | 1.868547 | 0.202812 | 9.21322 | 3.16E-20 | 1.50E-18 |
| CNAG_01990 | 41431.64 | -1.01443 | 0.184233 | -5.5062 | 3.67E-08 | 3.51E-07 |
| CNAG_01991 | 8598.857 | -0.86493 | 0.178941 | -4.83358 | 1.34E-06 | 9.02E-06 |

| | | | | | | |
|---|---|---|---|---|---|---|
| CNAG_01992 | 101.0048 | 1.580479 | 0.26688 | 5.92206 | 3.18E-09 | 3.68E-08 |
| CNAG_01993 | 404.8912 | 0.659634 | 0.205607 | 3.208223 | 0.001336 | 0.004279 |
| CNAG_01994 | 40.91381 | 1.170926 | 0.352536 | 3.321437 | 0.000896 | 0.003001 |
| CNAG_01997 | 3031.046 | -1.10182 | 0.177964 | -6.19126 | 5.97E-10 | 7.79E-09 |
| CNAG_01999 | 559.4627 | -0.3944 | 0.1771 | -2.22698 | 0.025948 | 0.055531 |
| CNAG_02000 | 2552.424 | 0.997136 | 0.194624 | 5.123395 | 3.00E-07 | 2.30E-06 |
| CNAG_02001 | 2074.059 | 0.823407 | 0.18238 | 4.514797 | 6.34E-06 | 3.72E-05 |
| CNAG_02002 | 2828.588 | -0.82664 | 0.20397 | -4.05274 | 5.06E-05 | 0.000237 |
| CNAG_08013 | 874.0688 | -0.27752 | 0.137613 | -2.01666 | 0.043731 | 0.086784 |
| CNAG_02004 | 3595.788 | 0.771645 | 0.123041 | 6.27144 | 3.58E-10 | 4.84E-09 |
| CNAG_02005 | 405.6325 | -4.33272 | 0.940399 | -4.60732 | 4.08E-06 | 2.52E-05 |
| CNAG_02006 | 1505.006 | 0.90215 | 0.184222 | 4.897071 | 9.73E-07 | 6.74E-06 |
| CNAG_02007 | 7367.185 | -0.49326 | 0.1138 | -4.33442 | 1.46E-05 | 7.86E-05 |
| CNAG_02008 | 3226.368 | 1.334063 | 0.20883 | 6.388272 | 1.68E-10 | 2.40E-09 |
| CNAG_02009 | 1497.572 | 0.601754 | 0.175367 | 3.431394 | 0.0006 | 0.00213 |
| CNAG_02010 | 580.1039 | -0.48205 | 0.16477 | -2.92558 | 0.003438 | 0.009767 |
| CNAG_02011 | 731.9068 | -0.48033 | 0.181653 | -2.64424 | 0.008188 | 0.020686 |
| CNAG_07619 | 1285.899 | 1.023764 | 0.17102 | 5.986218 | 2.15E-09 | 2.59E-08 |
| CNAG_13043 | 110.2366 | -0.49403 | 0.227324 | -2.17324 | 0.029762 | 0.062416 |
| CNAG_07026 | 41.50189 | 0.86733 | 0.340631 | 2.546242 | 0.010889 | 0.026476 |
| CNAG_05990 | 461.0397 | 0.486391 | 0.163874 | 2.968077 | 0.002997 | 0.008679 |
| CNAG_05993 | 4174.95 | -1.32505 | 0.417333 | -3.17504 | 0.001498 | 0.004728 |
| CNAG_05994 | 4863.652 | 0.993318 | 0.26394 | 3.763418 | 0.000168 | 0.000693 |
| CNAG_05995 | 1704.016 | -0.92006 | 0.131605 | -6.99108 | 2.73E-12 | 5.25E-11 |
| CNAG_05997 | 4106.404 | -1.01959 | 0.198097 | -5.14696 | 2.65E-07 | 2.05E-06 |
| CNAG_05999 | 1033.788 | 0.47088 | 0.171251 | 2.749656 | 0.005966 | 0.015837 |
| CNAG_06001 | 1113.154 | -0.75149 | 0.198087 | -3.79377 | 0.000148 | 0.00062 |
| CNAG_06002 | 548.4 | -0.60919 | 0.173042 | -3.52048 | 0.000431 | 0.001584 |
| CNAG_06004 | 507.9144 | -0.27691 | 0.129959 | -2.13073 | 0.033111 | 0.068356 |
| CNAG_06005 | 1625.32 | -0.70666 | 0.17113 | -4.12938 | 3.64E-05 | 0.000177 |
| CNAG_06006 | 1348.049 | -0.38044 | 0.146885 | -2.59005 | 0.009596 | 0.023742 |
| CNAG_06007 | 926.6096 | -1.01907 | 0.221655 | -4.59756 | 4.27E-06 | 2.63E-05 |
| CNAG_06008 | 2822.206 | -1.20324 | 0.149062 | -8.07205 | 6.91E-16 | 2.11E-14 |
| CNAG_06009 | 974.153 | 1.278125 | 0.198814 | 6.428749 | 1.29E-10 | 1.89E-09 |
| CNAG_13046 | 177.324 | 3.914676 | 0.407975 | 9.595378 | 8.36E-22 | 4.61E-20 |
| CNAG_06012 | 5773.027 | -0.59112 | 0.193537 | -3.05431 | 0.002256 | 0.0068 |
| CNAG_06018 | 884.4276 | 0.406414 | 0.201172 | 2.020226 | 0.04336 | 0.086135 |
| CNAG_06021 | 5606.902 | 1.3166 | 0.168713 | 7.803803 | 6.01E-15 | 1.63E-13 |
| CNAG_06026 | 2268.959 | -0.45756 | 0.143418 | -3.19043 | 0.001421 | 0.004512 |
| CNAG_06027 | 118.6324 | 2.962348 | 0.263495 | 11.24251 | 2.52E-29 | 2.23E-27 |
| CNAG_06028 | 483.4542 | -0.52988 | 0.147753 | -3.58627 | 0.000335 | 0.001276 |
| CNAG_06029 | 389.0015 | -0.61168 | 0.154287 | -3.9646 | 7.35E-05 | 0.000332 |
| CNAG_06030 | 590.1055 | 0.301238 | 0.131919 | 2.283507 | 0.022401 | 0.049143 |
| CNAG_06033 | 1032.29 | -0.59213 | 0.211569 | -2.79876 | 0.00513 | 0.013883 |
| CNAG_06034 | 1999.188 | 0.351722 | 0.174952 | 2.010396 | 0.044389 | 0.087784 |
| CNAG_06035 | 712.4227 | 1.040001 | 0.160609 | 6.475346 | 9.46E-11 | 1.42E-09 |
| CNAG_06036 | 4069.298 | 0.548749 | 0.175749 | 3.122354 | 0.001794 | 0.005539 |

| | | | | | | |
|---|---|---|---|---|---|---|
| CNAG_13055 | 50.33539 | 1.163234 | 0.316404 | 3.676421 | 0.000237 | 0.000938 |
| CNAG_06052 | 1028.865 | 0.446799 | 0.164994 | 2.707972 | 0.00677 | 0.017613 |
| CNAG_06055 | 205.428 | 0.666557 | 0.197139 | 3.381152 | 0.000722 | 0.00249 |
| CNAG_06060 | 3593.177 | 1.16544 | 0.256894 | 4.536651 | 5.72E-06 | 3.39E-05 |
| CNAG_06061 | 4954.102 | -0.44078 | 0.125262 | -3.51889 | 0.000433 | 0.00159 |
| CNAG_06063 | 1343.425 | -0.64854 | 0.133776 | -4.84798 | 1.25E-06 | 8.44E-06 |
| CNAG_06064 | 1237.816 | 0.971046 | 0.252277 | 3.849123 | 0.000119 | 0.000507 |
| CNAG_06065 | 1831.974 | 0.568031 | 0.235866 | 2.408274 | 0.016028 | 0.036873 |
| CNAG_06066 | 59.35672 | 1.741251 | 0.482246 | 3.61071 | 0.000305 | 0.001177 |
| CNAG_07901 | 2755.316 | 0.912177 | 0.273571 | 3.334332 | 0.000855 | 0.00288 |
| CNAG_07902 | 9801.368 | -0.57205 | 0.147846 | -3.86921 | 0.000109 | 0.000471 |
| CNAG_06072 | 1525.955 | -0.59269 | 0.182516 | -3.24731 | 0.001165 | 0.003792 |
| CNAG_13060 | 20.03477 | 1.046552 | 0.452451 | 2.313073 | 0.020719 | 0.045884 |
| CNAG_13061 | 232.5635 | -0.62171 | 0.269635 | -2.30573 | 0.021126 | 0.046694 |
| CNAG_06074 | 1968.784 | 1.837625 | 0.280224 | 6.557712 | 5.46E-11 | 8.37E-10 |
| CNAG_06075 | 14178.65 | 0.857111 | 0.198809 | 4.31122 | 1.62E-05 | 8.61E-05 |
| CNAG_06079 | 4814.985 | -0.3569 | 0.131064 | -2.72313 | 0.006467 | 0.016947 |
| CNAG_06080 | 4216.965 | 0.38471 | 0.129425 | 2.972454 | 0.002954 | 0.00858 |
| CNAG_06082 | 10.37644 | -1.97194 | 0.72471 | -2.72101 | 0.006508 | 0.017039 |
| CNAG_13063 | 1211.315 | 0.441412 | 0.173596 | 2.542746 | 0.010999 | 0.026693 |
| CNAG_06085 | 1017.177 | 0.897253 | 0.177624 | 5.051421 | 4.39E-07 | 3.27E-06 |
| CNAG_06086 | 837.6739 | -0.54445 | 0.163601 | -3.32792 | 0.000875 | 0.00294 |
| CNAG_06087 | 1206.537 | -1.44384 | 0.212909 | -6.78146 | 1.19E-11 | 2.05E-10 |
| CNAG_06095 | 70698.83 | -0.90795 | 0.242139 | -3.74969 | 0.000177 | 0.000729 |
| CNAG_06096 | 11341.02 | -1.36592 | 0.135958 | -10.0466 | 9.51E-24 | 6.19E-22 |
| CNAG_06097 | 464.0876 | -0.48253 | 0.194104 | -2.48592 | 0.012922 | 0.030659 |
| CNAG_06100 | 819.9915 | 0.689177 | 0.155203 | 4.44049 | 8.98E-06 | 5.06E-05 |
| CNAG_06101 | 119193.9 | -0.77819 | 0.153217 | -5.07901 | 3.79E-07 | 2.86E-06 |
| CNAG_13067 | 155.2584 | 0.481375 | 0.221772 | 2.170585 | 0.029963 | 0.062769 |
| CNAG_06102 | 172.2447 | 1.173725 | 0.20384 | 5.758066 | 8.51E-09 | 9.01E-08 |
| CNAG_06103 | 4783.081 | 0.780022 | 0.198807 | 3.923513 | 8.73E-05 | 0.000388 |
| CNAG_06104 | 665.1195 | -1.89673 | 0.226816 | -8.3624 | 6.15E-17 | 2.10E-15 |
| CNAG_13069 | 20.53869 | -2.89087 | 0.611122 | -4.73044 | 2.24E-06 | 1.46E-05 |
| CNAG_06105 | 105.2853 | 1.708945 | 0.372514 | 4.587594 | 4.48E-06 | 2.73E-05 |
| CNAG_06106 | 7914.293 | 0.495945 | 0.187548 | 2.644369 | 0.008184 | 0.020685 |
| CNAG_06109 | 21560.74 | 1.90181 | 0.271521 | 7.00428 | 2.48E-12 | 4.83E-11 |
| CNAG_06112 | 11978.07 | -0.53993 | 0.123501 | -4.37188 | 1.23E-05 | 6.75E-05 |
| CNAG_06113 | 61073.26 | -0.80816 | 0.223583 | -3.61459 | 0.000301 | 0.001162 |
| CNAG_06118 | 1228.658 | -0.89058 | 0.235472 | -3.7821 | 0.000156 | 0.000645 |
| CNAG_06119 | 205.5649 | 1.782989 | 0.193081 | 9.234401 | 2.60E-20 | 1.25E-18 |
| CNAG_06121 | 3424.663 | 1.691985 | 0.225965 | 7.48783 | 7.00E-14 | 1.66E-12 |
| CNAG_06122 | 1175.721 | -0.89458 | 0.182831 | -4.89292 | 9.94E-07 | 6.86E-06 |
| CNAG_06123 | 8264.354 | -0.75287 | 0.172788 | -4.35717 | 1.32E-05 | 7.19E-05 |
| CNAG_06125 | 303615.7 | -0.44973 | 0.115528 | -3.89286 | 9.91E-05 | 0.000432 |
| CNAG_06126 | 1014.206 | -1.32129 | 0.201035 | -6.57247 | 4.95E-11 | 7.61E-10 |
| CNAG_06127 | 3099.008 | -0.4338 | 0.15213 | -2.85153 | 0.004351 | 0.012004 |
| CNAG_06134 | 10528.66 | 0.363824 | 0.141279 | 2.57521 | 0.010018 | 0.024654 |

| | | | | | | |
|---|---|---|---|---|---|---|
| CNAG_07905 | 67.29487 | 1.06393 | 0.381111 | 2.791655 | 0.005244 | 0.014158 |
| CNAG_06136 | 732.1771 | 1.055573 | 0.215534 | 4.897474 | 9.71E-07 | 6.74E-06 |
| CNAG_06138 | 1776.579 | -1.24752 | 0.240079 | -5.19629 | 2.03E-07 | 1.62E-06 |
| CNAG_06140 | 6148.501 | -0.67284 | 0.300244 | -2.24098 | 0.025027 | 0.05385 |
| CNAG_06141 | 946.6629 | -0.65122 | 0.223024 | -2.91996 | 0.003501 | 0.009924 |
| CNAG_06144 | 21051.57 | -0.61666 | 0.194692 | -3.16736 | 0.001538 | 0.004839 |
| CNAG_06145 | 915.3392 | 0.348083 | 0.154177 | 2.257688 | 0.023965 | 0.052029 |
| CNAG_06147 | 2455.513 | -0.98659 | 0.317125 | -3.11104 | 0.001864 | 0.005724 |
| CNAG_06149 | 518.4282 | -0.53977 | 0.159704 | -3.37982 | 0.000725 | 0.002498 |
| CNAG_06152 | 612.7142 | 1.130706 | 0.185968 | 6.080107 | 1.20E-09 | 1.51E-08 |
| CNAG_06153 | 7399.439 | 0.736556 | 0.133324 | 5.524536 | 3.30E-08 | 3.19E-07 |
| CNAG_06154 | 889.3086 | 0.493577 | 0.18024 | 2.738437 | 0.006173 | 0.016296 |
| CNAG_06157 | 1171.873 | -0.29795 | 0.145465 | -2.04825 | 0.040536 | 0.081358 |
| CNAG_06159 | 132.1555 | 1.006549 | 0.227981 | 4.415063 | 1.01E-05 | 5.64E-05 |
| CNAG_06161 | 1459.184 | 0.821581 | 0.252866 | 3.249079 | 0.001158 | 0.003772 |
| CNAG_06162 | 626.9209 | -0.50777 | 0.255701 | -1.98581 | 0.047055 | 0.092366 |
| CNAG_06164 | 376.5253 | 0.352364 | 0.175765 | 2.004748 | 0.04499 | 0.088818 |
| CNAG_06166 | 1547.86 | 0.672257 | 0.268199 | 2.506558 | 0.012191 | 0.02917 |
| CNAG_08019 | 444.289 | 0.668652 | 0.2447 | 2.732536 | 0.006285 | 0.016575 |
| CNAG_06167 | 4801.227 | 0.500575 | 0.134285 | 3.7277 | 0.000193 | 0.000784 |
| CNAG_06168 | 4016.548 | -0.80062 | 0.137687 | -5.81475 | 6.07E-09 | 6.58E-08 |
| CNAG_06169 | 3498.42 | 3.461289 | 0.18711 | 18.4987 | 2.12E-76 | 1.06E-73 |
| CNAG_06170 | 3057.937 | 0.346565 | 0.162228 | 2.136285 | 0.032656 | 0.06761 |
| CNAG_06175 | 7789.522 | 0.777484 | 0.250457 | 3.104261 | 0.001908 | 0.005848 |
| CNAG_07908 | 2804.403 | -1.89791 | 0.147455 | -12.8712 | 6.54E-38 | 1.00E-35 |
| CNAG_07909 | 88.58309 | -1.34707 | 0.250223 | -5.38346 | 7.31E-08 | 6.45E-07 |
| CNAG_06180 | 2365.493 | -1.35638 | 0.270036 | -5.02295 | 5.09E-07 | 3.75E-06 |
| CNAG_13075 | 35.35055 | 0.848194 | 0.360985 | 2.349665 | 0.01879 | 0.042201 |
| CNAG_07910 | 5.882076 | 2.31634 | 0.950724 | 2.436395 | 0.014834 | 0.034545 |
| CNAG_06183 | 934.3235 | -0.44384 | 0.124988 | -3.55107 | 0.000384 | 0.001433 |
| CNAG_06184 | 585.2704 | -0.48484 | 0.237172 | -2.04427 | 0.040927 | 0.081999 |
| CNAG_06186 | 1106.734 | -0.52671 | 0.125657 | -4.1916 | 2.77E-05 | 0.000138 |
| CNAG_07911 | 212.5932 | -1.42409 | 0.302941 | -4.70089 | 2.59E-06 | 1.67E-05 |
| CNAG_06188 | 877.4999 | -0.89488 | 0.226961 | -3.94288 | 8.05E-05 | 0.00036 |
| CNAG_06190 | 1334.095 | 0.919909 | 0.181215 | 5.076347 | 3.85E-07 | 2.89E-06 |
| CNAG_06191 | 586.5711 | -0.60358 | 0.179464 | -3.36324 | 0.00077 | 0.002633 |
| CNAG_06193 | 1159.174 | 0.735561 | 0.361503 | 2.034729 | 0.041878 | 0.08357 |
| CNAG_13077 | 20.19787 | 2.701082 | 0.543792 | 4.967122 | 6.80E-07 | 4.90E-06 |
| CNAG_13078 | 6.798204 | 2.884407 | 1.07677 | 2.678758 | 0.00739 | 0.018945 |
| CNAG_06200 | 1659.278 | 0.776619 | 0.188857 | 4.112201 | 3.92E-05 | 0.000189 |
| CNAG_13080 | 140.3594 | 0.785872 | 0.316429 | 2.483568 | 0.013007 | 0.030816 |
| CNAG_06201 | 2472.475 | 1.441931 | 0.159304 | 9.051466 | 1.41E-19 | 6.16E-18 |
| CNAG_06202 | 90.30429 | 0.556949 | 0.260065 | 2.141572 | 0.032228 | 0.066891 |
| CNAG_06203 | 256.5733 | -0.68115 | 0.253815 | -2.68366 | 0.007282 | 0.018732 |
| CNAG_13082 | 700.1597 | -1.73031 | 0.353364 | -4.89669 | 9.75E-07 | 6.75E-06 |
| CNAG_06204 | 5071.004 | -2.05251 | 0.173423 | -11.8353 | 2.56E-32 | 2.83E-30 |
| CNAG_06205 | 4761.677 | -0.3529 | 0.120193 | -2.93613 | 0.003323 | 0.009479 |

| | | | | | | |
|---|---|---|---|---|---|---|
| CNAG_06206 | 2249.13 | -0.79787 | 0.138334 | -5.76768 | 8.04E-09 | 8.55E-08 |
| CNAG_06207 | 1791.376 | -1.35555 | 0.220985 | -6.13413 | 8.56E-10 | 1.10E-08 |
| CNAG_06208 | 37432.18 | 0.48886 | 0.106469 | 4.591577 | 4.40E-06 | 2.69E-05 |
| CNAG_06213 | 1709.843 | 1.325273 | 0.249643 | 5.308672 | 1.10E-07 | 9.38E-07 |
| CNAG_06214 | 2045.407 | 0.460413 | 0.173294 | 2.656837 | 0.007888 | 0.020024 |
| CNAG_13084 | 46.06722 | 1.388079 | 0.349317 | 3.973695 | 7.08E-05 | 0.000321 |
| CNAG_06216 | 1728.867 | 0.987316 | 0.182976 | 5.395868 | 6.82E-08 | 6.08E-07 |
| CNAG_06217 | 200.6591 | 0.469319 | 0.172235 | 2.724872 | 0.006433 | 0.016875 |
| CNAG_06220 | 12864.59 | 2.105051 | 0.136374 | 15.43592 | 9.39E-54 | 2.76E-51 |
| CNAG_06221 | 914.1366 | -0.79711 | 0.152358 | -5.23184 | 1.68E-07 | 1.37E-06 |
| CNAG_06222 | 35383.36 | -1.06794 | 0.283588 | -3.76582 | 0.000166 | 0.000687 |
| CNAG_06226 | 3702.119 | -2.01692 | 0.3153 | -6.39681 | 1.59E-10 | 2.29E-09 |
| CNAG_06227 | 1732.178 | 0.966422 | 0.184572 | 5.23601 | 1.64E-07 | 1.35E-06 |
| CNAG_06228 | 1920.62 | -0.49667 | 0.145015 | -3.42493 | 0.000615 | 0.00217 |
| CNAG_13087 | 26.0759 | -2.12825 | 0.487208 | -4.36827 | 1.25E-05 | 6.85E-05 |
| CNAG_06231 | 37350.46 | -0.85743 | 0.213499 | -4.01609 | 5.92E-05 | 0.000272 |
| CNAG_06232 | 870.0846 | 0.433535 | 0.186481 | 2.324823 | 0.020081 | 0.044747 |
| CNAG_13089 | 229.8415 | 1.041675 | 0.171103 | 6.088011 | 1.14E-09 | 1.44E-08 |
| CNAG_06235 | 2154.112 | -0.42341 | 0.133619 | -3.16882 | 0.001531 | 0.004822 |
| CNAG_06237 | 2380.129 | 0.450442 | 0.16523 | 2.72615 | 0.006408 | 0.016832 |
| CNAG_06238 | 646.6783 | 3.131871 | 0.266533 | 11.75039 | 7.03E-32 | 7.45E-30 |
| CNAG_06239 | 240.7518 | 0.431963 | 0.179218 | 2.410273 | 0.015941 | 0.036735 |
| CNAG_06240 | 14961.17 | 0.904018 | 0.133196 | 6.787123 | 1.14E-11 | 1.98E-10 |
| CNAG_06244 | 1618.165 | 0.583383 | 0.152404 | 3.827874 | 0.000129 | 0.000547 |
| CNAG_06247 | 165.1945 | 0.401317 | 0.179171 | 2.239856 | 0.0251 | 0.053977 |
| CNAG_06249 | 413.2646 | 0.37827 | 0.169186 | 2.235819 | 0.025364 | 0.0545 |
| CNAG_06252 | 442.9206 | -0.61124 | 0.175753 | -3.47782 | 0.000506 | 0.00183 |
| CNAG_06259 | 319.8805 | -0.76724 | 0.201759 | -3.80278 | 0.000143 | 0.000601 |
| CNAG_06260 | 1718.149 | -0.66465 | 0.12309 | -5.39971 | 6.67E-08 | 5.99E-07 |
| CNAG_07921 | 1005.524 | -0.31947 | 0.140477 | -2.27418 | 0.022955 | 0.050258 |
| CNAG_13097 | 435.6443 | -0.59664 | 0.201148 | -2.96617 | 0.003015 | 0.008719 |
| CNAG_06263 | 23.39551 | 1.035494 | 0.419021 | 2.471223 | 0.013465 | 0.031721 |
| CNAG_06265 | 541.8821 | 0.390913 | 0.139963 | 2.792976 | 0.005223 | 0.014117 |
| CNAG_06267 | 3292.705 | 3.365344 | 0.177185 | 18.99334 | 1.94E-80 | 1.28E-77 |
| CNAG_13099 | 788.1813 | 1.418284 | 0.203671 | 6.963599 | 3.32E-12 | 6.20E-11 |
| CNAG_07923 | 20.77869 | 1.451521 | 0.469942 | 3.08872 | 0.00201 | 0.006134 |
| CNAG_13100 | 91.06707 | -0.69781 | 0.239542 | -2.91308 | 0.003579 | 0.010113 |
| CNAG_07924 | 1434.057 | -0.42164 | 0.212004 | -1.98883 | 0.04672 | 0.0918 |
| CNAG_06272 | 612.3673 | 0.568778 | 0.192517 | 2.954427 | 0.003132 | 0.008992 |
| CNAG_06273 | 2168.118 | -0.58699 | 0.291287 | -2.01517 | 0.043887 | 0.087029 |
| CNAG_06274 | 8273.175 | -0.85132 | 0.238435 | -3.57043 | 0.000356 | 0.001343 |
| CNAG_06277 | 3054.206 | -0.36876 | 0.167931 | -2.19592 | 0.028098 | 0.059426 |
| CNAG_06279 | 2921.811 | -0.46623 | 0.161815 | -2.88128 | 0.003961 | 0.011019 |
| CNAG_06283 | 1128.035 | -0.59031 | 0.160692 | -3.67357 | 0.000239 | 0.000947 |
| CNAG_06285 | 533.7644 | 1.296339 | 0.20314 | 6.381512 | 1.75E-10 | 2.50E-09 |
| CNAG_06286 | 349.499 | 1.158729 | 0.157894 | 7.338638 | 2.16E-13 | 4.79E-12 |
| CNAG_06288 | 2157.494 | -0.73245 | 0.187197 | -3.91271 | 9.13E-05 | 0.000403 |

| | | | | | | |
|---|---|---|---|---|---|---|
| CNAG_06289 | 2277.747 | 0.484451 | 0.133483 | 3.629314 | 0.000284 | 0.001105 |
| CNAG_13106 | 60.40101 | 1.533679 | 0.281956 | 5.439419 | 5.35E-08 | 4.87E-07 |
| CNAG_06290 | 411.6428 | 1.695978 | 0.21664 | 7.828571 | 4.93E-15 | 1.35E-13 |
| CNAG_06291 | 148.4142 | 1.381843 | 0.195752 | 7.059161 | 1.68E-12 | 3.37E-11 |
| CNAG_06292 | 57.38041 | 0.82824 | 0.308655 | 2.683383 | 0.007288 | 0.018735 |
| CNAG_06293 | 415.6124 | -0.59243 | 0.147877 | -4.00624 | 6.17E-05 | 0.000283 |
| CNAG_06296 | 63.86876 | 2.278038 | 0.317954 | 7.164689 | 7.80E-13 | 1.64E-11 |
| CNAG_06297 | 932.2337 | -1.92132 | 0.264218 | -7.27172 | 3.55E-13 | 7.79E-12 |
| CNAG_06298 | 3401.578 | -2.08395 | 0.323475 | -6.44237 | 1.18E-10 | 1.73E-09 |
| CNAG_13108 | 57.32374 | 1.317113 | 0.433538 | 3.038056 | 0.002381 | 0.007124 |
| CNAG_07928 | 68.68854 | -1.57782 | 0.35239 | -4.47749 | 7.55E-06 | 4.33E-05 |
| CNAG_06301 | 6322.571 | 0.443652 | 0.135735 | 3.268512 | 0.001081 | 0.003543 |
| CNAG_06302 | 137.8962 | 1.284116 | 0.244095 | 5.260733 | 1.43E-07 | 1.19E-06 |
| CNAG_06308 | 420.3452 | -0.54735 | 0.169154 | -3.23583 | 0.001213 | 0.003935 |
| CNAG_06309 | 142.0079 | 0.924794 | 0.239301 | 3.864559 | 0.000111 | 0.000478 |
| CNAG_06310 | 789.182 | 0.87386 | 0.130976 | 6.671922 | 2.52E-11 | 4.09E-10 |
| CNAG_06311 | 1112.997 | 0.356964 | 0.178523 | 1.999534 | 0.045551 | 0.089768 |
| CNAG_06314 | 8698.309 | -0.59132 | 0.119453 | -4.95021 | 7.41E-07 | 5.29E-06 |
| CNAG_06315 | 2795.986 | -0.68052 | 0.191594 | -3.55191 | 0.000382 | 0.001429 |
| CNAG_06316 | 3328.23 | -0.55621 | 0.177848 | -3.12746 | 0.001763 | 0.00546 |
| CNAG_06318 | 2126.87 | -0.86375 | 0.227016 | -3.80478 | 0.000142 | 0.000597 |
| CNAG_06320 | 3235.558 | 0.532709 | 0.142233 | 3.745322 | 0.00018 | 0.000739 |
| CNAG_06321 | 106.8702 | 1.19764 | 0.329706 | 3.632444 | 0.000281 | 0.001093 |
| CNAG_06322 | 601.2305 | -0.94998 | 0.152268 | -6.23884 | 4.41E-10 | 5.89E-09 |
| CNAG_06324 | 542.2937 | -1.24923 | 0.248597 | -5.02512 | 5.03E-07 | 3.72E-06 |
| CNAG_06325 | 2251.212 | -0.63112 | 0.132015 | -4.78066 | 1.75E-06 | 1.15E-05 |
| CNAG_06326 | 1033.987 | 0.641688 | 0.154323 | 4.158078 | 3.21E-05 | 0.000158 |
| CNAG_06330 | 295.86 | -0.53105 | 0.179171 | -2.96395 | 0.003037 | 0.008766 |
| CNAG_06332 | 2038.55 | 0.315509 | 0.11802 | 2.673354 | 0.00751 | 0.019175 |
| CNAG_06334 | 2301.863 | 1.347586 | 0.214537 | 6.281366 | 3.36E-10 | 4.60E-09 |
| CNAG_06335 | 1238.854 | -0.79591 | 0.211146 | -3.76949 | 0.000164 | 0.000678 |
| CNAG_13118 | 42.36409 | 2.039625 | 0.363064 | 5.617819 | 1.93E-08 | 1.94E-07 |
| CNAG_06339 | 637.4768 | 0.928483 | 0.220091 | 4.218625 | 2.46E-05 | 0.000124 |
| CNAG_06340 | 2042.357 | -0.80534 | 0.225198 | -3.57615 | 0.000349 | 0.001319 |
| CNAG_06341 | 825.5964 | -1.1517 | 0.264872 | -4.34812 | 1.37E-05 | 7.46E-05 |
| CNAG_06345 | 897.3282 | -0.62078 | 0.184374 | -3.36696 | 0.00076 | 0.002605 |
| CNAG_06346 | 6430.218 | -2.01435 | 0.330282 | -6.09887 | 1.07E-09 | 1.35E-08 |
| CNAG_06347 | 12077.16 | 3.038419 | 0.195229 | 15.56338 | 1.29E-54 | 3.95E-52 |
| CNAG_06352 | 1625.174 | -0.8465 | 0.236066 | -3.58586 | 0.000336 | 0.001277 |
| CNAG_06353 | 2554.499 | -0.55379 | 0.121932 | -4.54183 | 5.58E-06 | 3.31E-05 |
| CNAG_06355 | 606.064 | 0.583958 | 0.266376 | 2.192229 | 0.028363 | 0.059892 |
| CNAG_06357 | 547.6756 | -0.72249 | 0.280348 | -2.57711 | 0.009963 | 0.024548 |
| CNAG_06358 | 380.7226 | -0.68776 | 0.258889 | -2.6566 | 0.007893 | 0.020032 |
| CNAG_06359 | 483.8197 | 0.404945 | 0.188033 | 2.15359 | 0.031272 | 0.065135 |
| CNAG_06361 | 6105.973 | 0.762812 | 0.114658 | 6.652931 | 2.87E-11 | 4.60E-10 |
| CNAG_06362 | 1346.303 | 0.672 | 0.255441 | 2.630749 | 0.00852 | 0.02141 |
| CNAG_06363 | 1021.016 | -1.04835 | 0.167147 | -6.272 | 3.56E-10 | 4.83E-09 |

| | | | | | | |
|---|---|---|---|---|---|---|
| CNAG_06365 | 1019.452 | 0.839769 | 0.145452 | 5.773511 | 7.76E-09 | 8.28E-08 |
| CNAG_06367 | 3024.776 | -0.73384 | 0.130135 | -5.63905 | 1.71E-08 | 1.72E-07 |
| CNAG_06368 | 640.3864 | -0.77468 | 0.173437 | -4.46665 | 7.95E-06 | 4.53E-05 |
| CNAG_06369 | 754.0376 | -0.43042 | 0.145876 | -2.95056 | 0.003172 | 0.009086 |
| CNAG_06370 | 3457.632 | -1.19297 | 0.164717 | -7.24257 | 4.40E-13 | 9.56E-12 |
| CNAG_06372 | 354.9425 | 0.982051 | 0.264499 | 3.712869 | 0.000205 | 0.000826 |
| CNAG_06374 | 6527.991 | -1.28057 | 0.155636 | -8.22797 | 1.90E-16 | 6.18E-15 |
| CNAG_06377 | 38960.21 | -0.40878 | 0.194896 | -2.09742 | 0.035956 | 0.073542 |
| CNAG_06378 | 501.3514 | -0.48618 | 0.174566 | -2.78509 | 0.005351 | 0.014414 |
| CNAG_06379 | 2866.844 | -0.44546 | 0.175921 | -2.53217 | 0.011336 | 0.027379 |
| CNAG_06381 | 8933.18 | 0.834342 | 0.136146 | 6.128289 | 8.88E-10 | 1.14E-08 |
| CNAG_06382 | 500.731 | 0.481396 | 0.152411 | 3.158534 | 0.001586 | 0.004978 |
| CNAG_06385 | 623.5707 | -0.77878 | 0.166137 | -4.6876 | 2.76E-06 | 1.77E-05 |
| CNAG_06386 | 1053.831 | -0.63848 | 0.184874 | -3.45362 | 0.000553 | 0.001983 |
| CNAG_06388 | 4302.2 | -1.54726 | 0.27969 | -5.53207 | 3.16E-08 | 3.07E-07 |
| CNAG_13123 | 23.26738 | -1.96325 | 0.534751 | -3.67133 | 0.000241 | 0.000953 |
| CNAG_06389 | 1362.178 | -0.74832 | 0.264451 | -2.82971 | 0.004659 | 0.012731 |
| CNAG_13124 | 94.62026 | 1.110744 | 0.264588 | 4.198018 | 2.69E-05 | 0.000135 |
| CNAG_06391 | 94.05762 | -0.72245 | 0.268947 | -2.68622 | 0.007227 | 0.018609 |
| CNAG_06392 | 1019.187 | -0.83708 | 0.199061 | -4.20516 | 2.61E-05 | 0.000132 |
| CNAG_06396 | 1048.257 | 3.173816 | 0.266843 | 11.89393 | 1.27E-32 | 1.47E-30 |
| CNAG_07936 | 2102.048 | 0.818465 | 0.263792 | 3.10269 | 0.001918 | 0.005874 |
| CNAG_06399 | 897.9589 | 1.138161 | 0.32306 | 3.523063 | 0.000427 | 0.001571 |
| CNAG_06400 | 56227.19 | -0.52802 | 0.205023 | -2.57541 | 0.010012 | 0.024653 |
| CNAG_13126 | 19.4759 | -1.4196 | 0.526736 | -2.6951 | 0.007037 | 0.018207 |
| CNAG_13128 | 106.3671 | 3.387127 | 1.352238 | 2.50483 | 0.012251 | 0.029304 |
| CNAG_06404 | 1999.299 | -1.18866 | 0.22808 | -5.21159 | 1.87E-07 | 1.52E-06 |
| CNAG_06406 | 874.7334 | -0.65306 | 0.164746 | -3.96405 | 7.37E-05 | 0.000332 |
| CNAG_06407 | 2842.926 | -0.85069 | 0.255902 | -3.3243 | 0.000886 | 0.002974 |
| CNAG_06411 | 195.1836 | 0.730979 | 0.19089 | 3.829322 | 0.000128 | 0.000545 |
| CNAG_06412 | 981.6674 | -0.54322 | 0.165447 | -3.28335 | 0.001026 | 0.003395 |
| CNAG_06413 | 181.7171 | 0.408808 | 0.176771 | 2.312647 | 0.020742 | 0.045923 |
| CNAG_06414 | 476.2503 | -0.60841 | 0.224833 | -2.70606 | 0.006809 | 0.017692 |
| CNAG_06415 | 1528.64 | 0.285204 | 0.124951 | 2.282524 | 0.022458 | 0.049256 |
| CNAG_06421 | 3831.404 | -1.09497 | 0.255159 | -4.29132 | 1.78E-05 | 9.34E-05 |
| CNAG_06425 | 711.4704 | -0.66686 | 0.166158 | -4.01339 | 5.99E-05 | 0.000275 |
| CNAG_06426 | 985.6855 | -0.80201 | 0.262348 | -3.05706 | 0.002235 | 0.006745 |
| CNAG_06427 | 880.1445 | 1.376919 | 0.241496 | 5.701624 | 1.19E-08 | 1.23E-07 |
| CNAG_06428 | 1757.216 | 1.301173 | 0.138211 | 9.414419 | 4.76E-21 | 2.41E-19 |
| CNAG_07937 | 2802.608 | -0.32997 | 0.128188 | -2.57414 | 0.010049 | 0.024711 |
| CNAG_07938 | 1366.399 | 1.159973 | 0.314127 | 3.692695 | 0.000222 | 0.000885 |
| CNAG_06431 | 812.7307 | 0.400616 | 0.150482 | 2.662214 | 0.007763 | 0.019739 |
| CNAG_06432 | 7099.155 | 1.219491 | 0.125194 | 9.740839 | 2.02E-22 | 1.17E-20 |
| CNAG_07939 | 13828.06 | 1.858162 | 0.338151 | 5.495066 | 3.91E-08 | 3.69E-07 |
| CNAG_13131 | 59.94245 | 0.637594 | 0.264047 | 2.414697 | 0.015748 | 0.036366 |
| CNAG_06439 | 2942.163 | -0.41347 | 0.18849 | -2.19362 | 0.028263 | 0.059727 |
| CNAG_07940 | 386.5895 | 0.807379 | 0.235411 | 3.429662 | 0.000604 | 0.002139 |

| | | | | | | |
|---|---|---|---|---|---|---|
| CNAG_06443 | 56821.23 | 1.160644 | 0.107421 | 10.80463 | 3.27E-27 | 2.55E-25 |
| CNAG_06446 | 2315.881 | -0.32149 | 0.110798 | -2.90164 | 0.003712 | 0.010438 |
| CNAG_06447 | 29575.62 | -0.91816 | 0.219904 | -4.17526 | 2.98E-05 | 0.000148 |
| CNAG_06449 | 133.3932 | 0.774026 | 0.204063 | 3.793076 | 0.000149 | 0.000621 |
| CNAG_06452 | 1855.223 | 0.852943 | 0.136154 | 6.264543 | 3.74E-10 | 5.05E-09 |
| CNAG_06453 | 7541.479 | 2.730502 | 0.129343 | 21.11051 | 6.37E-99 | 7.23E-96 |
| CNAG_06456 | 2472.182 | -0.86845 | 0.121664 | -7.13811 | 9.46E-13 | 1.98E-11 |
| CNAG_07941 | 2535.349 | -0.38748 | 0.186825 | -2.07403 | 0.038076 | 0.077182 |
| CNAG_06459 | 2226.76 | 1.19563 | 0.155421 | 7.692856 | 1.44E-14 | 3.70E-13 |
| CNAG_06460 | 3299.548 | 0.946699 | 0.129437 | 7.313998 | 2.59E-13 | 5.74E-12 |
| CNAG_06464 | 1173.49 | -0.5 | 0.193363 | -2.58581 | 0.009715 | 0.024004 |
| CNAG_06466 | 361.8598 | -0.75581 | 0.198522 | -3.80716 | 0.000141 | 0.000591 |
| CNAG_06468 | 3674.305 | -0.82347 | 0.164783 | -4.99731 | 5.81E-07 | 4.24E-06 |
| CNAG_06471 | 1242.061 | -0.76278 | 0.209756 | -3.63653 | 0.000276 | 0.001077 |
| CNAG_06472 | 3990.732 | -0.63595 | 0.139719 | -4.55166 | 5.32E-06 | 3.18E-05 |
| CNAG_06473 | 1551.469 | 0.489417 | 0.110366 | 4.434512 | 9.23E-06 | 5.19E-05 |
| CNAG_06474 | 6189.762 | -0.48215 | 0.146917 | -3.28178 | 0.001032 | 0.003408 |
| CNAG_13133 | 37.47929 | 0.970884 | 0.362653 | 2.677169 | 0.007425 | 0.019019 |
| CNAG_06482 | 586.764 | 0.573898 | 0.143326 | 4.004139 | 6.22E-05 | 0.000285 |
| CNAG_06485 | 294.5617 | -0.64365 | 0.183384 | -3.50985 | 0.000448 | 0.00164 |
| CNAG_06487 | 1065.241 | -1.04448 | 0.227664 | -4.5878 | 4.48E-06 | 2.73E-05 |
| CNAG_07943 | 1329.717 | 2.145612 | 0.213901 | 10.03086 | 1.12E-23 | 7.21E-22 |
| CNAG_06490 | 2314.595 | 0.455175 | 0.178314 | 2.552663 | 0.01069 | 0.026041 |
| CNAG_06491 | 153.1315 | -0.69598 | 0.283343 | -2.45633 | 0.014037 | 0.032919 |
| CNAG_06492 | 539.8058 | -0.81979 | 0.18169 | -4.51203 | 6.42E-06 | 3.77E-05 |
| CNAG_06493 | 1790.371 | 1.254108 | 0.311529 | 4.025649 | 5.68E-05 | 0.000263 |
| CNAG_06494 | 1830.774 | 1.405078 | 0.167806 | 8.373223 | 5.61E-17 | 1.92E-15 |
| CNAG_06497 | 930.6993 | 0.835848 | 0.132021 | 6.331189 | 2.43E-10 | 3.41E-09 |
| CNAG_06498 | 1432.193 | 2.17816 | 0.185407 | 11.74798 | 7.23E-32 | 7.56E-30 |
| CNAG_06499 | 999.7608 | 0.292129 | 0.13451 | 2.171811 | 0.02987 | 0.062608 |
| CNAG_06500 | 719.1125 | -0.89882 | 0.167981 | -5.35072 | 8.76E-08 | 7.64E-07 |
| CNAG_06503 | 65.74907 | 0.758158 | 0.327381 | 2.31583 | 0.020568 | 0.045613 |
| CNAG_06506 | 582.4857 | -0.28605 | 0.129756 | -2.20454 | 0.027486 | 0.05835 |
| CNAG_06510 | 780.8659 | -0.31709 | 0.148683 | -2.13265 | 0.032953 | 0.068115 |
| CNAG_06511 | 1996.767 | 1.153711 | 0.22686 | 5.085555 | 3.67E-07 | 2.77E-06 |
| CNAG_06512 | 879.3628 | 0.956637 | 0.177665 | 5.384505 | 7.26E-08 | 6.42E-07 |
| CNAG_06515 | 379.1845 | 0.620689 | 0.189492 | 3.275542 | 0.001055 | 0.003468 |
| CNAG_13141 | 217.0418 | -1.33392 | 0.266881 | -4.99818 | 5.79E-07 | 4.22E-06 |
| CNAG_06517 | 4082.058 | -1.95726 | 0.212586 | -9.20689 | 3.36E-20 | 1.58E-18 |
| CNAG_06520 | 482.4641 | -1.07502 | 0.234513 | -4.58405 | 4.56E-06 | 2.77E-05 |
| CNAG_06521 | 170.6628 | 0.599952 | 0.207118 | 2.896659 | 0.003772 | 0.010571 |
| CNAG_07043 | 391.3022 | 0.603479 | 0.153672 | 3.927062 | 8.60E-05 | 0.000383 |
| CNAG_05336 | 1234.458 | 0.400282 | 0.125079 | 3.200225 | 0.001373 | 0.004382 |
| CNAG_05337 | 1176.836 | 0.342231 | 0.155862 | 2.195727 | 0.028111 | 0.059439 |
| CNAG_05339 | 2355.017 | 0.70564 | 0.189213 | 3.729335 | 0.000192 | 0.000781 |
| CNAG_05340 | 460.7888 | -0.7862 | 0.185109 | -4.24723 | 2.16E-05 | 0.000111 |
| CNAG_05341 | 285.3404 | 1.7288 | 0.194595 | 8.884094 | 6.44E-19 | 2.65E-17 |

| | | | | | | |
|---|---|---|---|---|---|---|
| CNAG_13144 | 21.71922 | 1.453459 | 0.558763 | 2.601209 | 0.00929 | 0.023082 |
| CNAG_05345 | 8.196203 | 1.955611 | 0.744452 | 2.626915 | 0.008616 | 0.021625 |
| CNAG_07869 | 18.85935 | -3.60089 | 0.673683 | -5.34508 | 9.04E-08 | 7.86E-07 |
| CNAG_05348 | 5560.847 | -0.52439 | 0.192962 | -2.71755 | 0.006577 | 0.017173 |
| CNAG_05349 | 5341.014 | 0.539085 | 0.159963 | 3.370062 | 0.000752 | 0.002581 |
| CNAG_13146 | 48.12884 | 0.987861 | 0.310202 | 3.184576 | 0.00145 | 0.004587 |
| CNAG_05351 | 14355.84 | 0.408462 | 0.203563 | 2.006559 | 0.044797 | 0.088458 |
| CNAG_05352 | 2695.088 | 1.110914 | 0.13381 | 8.302159 | 1.02E-16 | 3.41E-15 |
| CNAG_05354 | 1012.783 | -0.79962 | 0.168086 | -4.7572 | 1.96E-06 | 1.28E-05 |
| CNAG_05355 | 5677.648 | 0.307023 | 0.123087 | 2.494358 | 0.012619 | 0.030029 |
| CNAG_13147 | 29.38503 | -1.17994 | 0.422328 | -2.79388 | 0.005208 | 0.014085 |
| CNAG_05358 | 64.57686 | -1.89891 | 0.43404 | -4.37496 | 1.21E-05 | 6.67E-05 |
| CNAG_05359 | 985.9689 | -0.41759 | 0.127908 | -3.26476 | 0.001096 | 0.003587 |
| CNAG_05360 | 1777.286 | -0.73964 | 0.127321 | -5.80926 | 6.28E-09 | 6.77E-08 |
| CNAG_05365 | 2018.331 | -0.53226 | 0.187051 | -2.84554 | 0.004434 | 0.012203 |
| CNAG_05366 | 6785.339 | -0.65673 | 0.129229 | -5.08194 | 3.74E-07 | 2.82E-06 |
| CNAG_05367 | 2255.964 | -1.18798 | 0.275188 | -4.31696 | 1.58E-05 | 8.43E-05 |
| CNAG_07873 | 1501.848 | -0.73823 | 0.197259 | -3.74243 | 0.000182 | 0.000746 |
| CNAG_05369 | 476.8578 | -0.34596 | 0.147445 | -2.3464 | 0.018956 | 0.042509 |
| CNAG_05370 | 782.886 | 0.567144 | 0.12199 | 4.649105 | 3.33E-06 | 2.09E-05 |
| CNAG_05371 | 1575.282 | -0.71182 | 0.14877 | -4.7847 | 1.71E-06 | 1.13E-05 |
| CNAG_05372 | 3332.508 | -0.37412 | 0.121717 | -3.07372 | 0.002114 | 0.006424 |
| CNAG_05377 | 52.6942 | 1.484371 | 0.372296 | 3.987072 | 6.69E-05 | 0.000305 |
| CNAG_05383 | 1829.157 | 2.825741 | 0.226704 | 12.46444 | 1.17E-35 | 1.60E-33 |
| CNAG_13157 | 21.93735 | 1.627507 | 0.455399 | 3.573808 | 0.000352 | 0.001329 |
| CNAG_05387 | 7806.013 | -0.78381 | 0.136408 | -5.74606 | 9.13E-09 | 9.64E-08 |
| CNAG_13158 | 9.839205 | 2.278478 | 0.708272 | 3.216953 | 0.001296 | 0.004176 |
| CNAG_05388 | 2323.508 | 0.759412 | 0.165958 | 4.575929 | 4.74E-06 | 2.86E-05 |
| CNAG_05390 | 718.6779 | 0.383561 | 0.136394 | 2.812151 | 0.004921 | 0.013378 |
| CNAG_05391 | 1535.427 | -1.25282 | 0.141906 | -8.82854 | 1.06E-18 | 4.26E-17 |
| CNAG_13160 | 15.0444 | -1.94619 | 0.741738 | -2.62383 | 0.008695 | 0.021767 |
| CNAG_05392 | 1151.504 | -2.90686 | 0.315878 | -9.20248 | 3.50E-20 | 1.63E-18 |
| CNAG_05393 | 1028.198 | -0.37608 | 0.136892 | -2.74726 | 0.00601 | 0.015923 |
| CNAG_05395 | 1206.728 | 0.737694 | 0.181139 | 4.072526 | 4.65E-05 | 0.000219 |
| CNAG_05398 | 4469.195 | -1.1441 | 0.192429 | -5.94558 | 2.75E-09 | 3.23E-08 |
| CNAG_05402 | 2417.497 | -1.20139 | 0.225 | -5.33949 | 9.32E-08 | 8.07E-07 |
| CNAG_05406 | 305.1098 | -0.75317 | 0.169043 | -4.45548 | 8.37E-06 | 4.76E-05 |
| CNAG_05408 | 4268.553 | -0.29063 | 0.119983 | -2.42225 | 0.015425 | 0.035733 |
| CNAG_05410 | 1473.846 | 1.659787 | 0.233519 | 7.107722 | 1.18E-12 | 2.43E-11 |
| CNAG_05411 | 2035.898 | 0.651267 | 0.123031 | 5.293524 | 1.20E-07 | 1.01E-06 |
| CNAG_05412 | 50.85807 | 3.338141 | 0.44713 | 7.465711 | 8.29E-14 | 1.94E-12 |
| CNAG_05414 | 260.1327 | -0.37299 | 0.156005 | -2.39092 | 0.016806 | 0.038385 |
| CNAG_05415 | 1412.379 | -1.42596 | 0.211097 | -6.755 | 1.43E-11 | 2.43E-10 |
| CNAG_05416 | 4239.082 | -0.45033 | 0.140864 | -3.19687 | 0.001389 | 0.004426 |
| CNAG_05419 | 2480.536 | 1.071717 | 0.209885 | 5.106199 | 3.29E-07 | 2.50E-06 |
| CNAG_05420 | 401.7238 | -0.72895 | 0.242854 | -3.00161 | 0.002686 | 0.007921 |
| CNAG_13168 | 114.9943 | 0.667412 | 0.250212 | 2.667382 | 0.007644 | 0.019475 |

| | | | | | | |
|---|---|---|---|---|---|---|
| CNAG_05423 | 372.4674 | -0.51336 | 0.144653 | -3.54891 | 0.000387 | 0.001444 |
| CNAG_05425 | 14780.77 | -1.19959 | 0.160555 | -7.4715 | 7.93E-14 | 1.87E-12 |
| CNAG_05426 | 128.6912 | 1.153377 | 0.281234 | 4.101129 | 4.11E-05 | 0.000197 |
| CNAG_13172 | 76.14692 | -0.8265 | 0.314183 | -2.63063 | 0.008523 | 0.02141 |
| CNAG_05430 | 838.4851 | 0.726947 | 0.187209 | 3.883079 | 0.000103 | 0.000447 |
| CNAG_05431 | 1306.178 | -0.50484 | 0.15409 | -3.27627 | 0.001052 | 0.003462 |
| CNAG_05434 | 2424.538 | -0.8864 | 0.193376 | -4.58384 | 4.57E-06 | 2.77E-05 |
| CNAG_05435 | 875.4889 | -0.59459 | 0.152569 | -3.89721 | 9.73E-05 | 0.000425 |
| CNAG_05436 | 121.2127 | -0.57433 | 0.247691 | -2.31875 | 0.020409 | 0.045349 |
| CNAG_05437 | 14317.64 | -0.58714 | 0.168778 | -3.47878 | 0.000504 | 0.001824 |
| CNAG_05438 | 597.1646 | -1.41719 | 0.236837 | -5.98381 | 2.18E-09 | 2.62E-08 |
| CNAG_05439 | 749.8201 | 0.566315 | 0.12925 | 4.381556 | 1.18E-05 | 6.50E-05 |
| CNAG_05440 | 1261.769 | -0.56587 | 0.195059 | -2.90101 | 0.00372 | 0.010455 |
| CNAG_05441 | 493.9045 | 0.730794 | 0.138612 | 5.272232 | 1.35E-07 | 1.12E-06 |
| CNAG_05442 | 1919.28 | -0.33986 | 0.159036 | -2.13702 | 0.032596 | 0.067503 |
| CNAG_05443 | 160.2872 | 0.53179 | 0.230902 | 2.303097 | 0.021273 | 0.046981 |
| CNAG_05446 | 2860.153 | -0.30323 | 0.115284 | -2.63028 | 0.008531 | 0.021426 |
| CNAG_07876 | 90.68279 | 0.908639 | 0.278003 | 3.268453 | 0.001081 | 0.003543 |
| CNAG_05449 | 10619.71 | 0.85196 | 0.23067 | 3.693418 | 0.000221 | 0.000883 |
| CNAG_05450 | 336.9453 | 0.630339 | 0.199618 | 3.157731 | 0.00159 | 0.004988 |
| CNAG_05454 | 58.16226 | 1.637859 | 0.548155 | 2.987946 | 0.002809 | 0.008226 |
| CNAG_05455 | 8739.483 | -0.85008 | 0.332683 | -2.55522 | 0.010612 | 0.02589 |
| CNAG_05456 | 745.882 | -0.29129 | 0.127316 | -2.28791 | 0.022143 | 0.048618 |
| CNAG_05457 | 5402.298 | 1.245138 | 0.207297 | 6.006551 | 1.90E-09 | 2.30E-08 |
| CNAG_05458 | 241.5273 | 2.113513 | 0.22222 | 9.510893 | 1.89E-21 | 1.00E-19 |
| CNAG_05460 | 466.3211 | 0.432763 | 0.15101 | 2.865795 | 0.00416 | 0.011521 |
| CNAG_05463 | 2138.325 | 0.366908 | 0.177522 | 2.066837 | 0.03875 | 0.078173 |
| CNAG_05465 | 30504.12 | -0.85463 | 0.172303 | -4.96003 | 7.05E-07 | 5.05E-06 |
| CNAG_08025 | 1826.996 | -0.51079 | 0.140905 | -3.62503 | 0.000289 | 0.001121 |
| CNAG_05468 | 1088.738 | -0.39832 | 0.141283 | -2.81933 | 0.004812 | 0.013109 |
| CNAG_05469 | 2444.482 | 0.35664 | 0.113566 | 3.140381 | 0.001687 | 0.005252 |
| CNAG_05473 | 469.8033 | -0.64312 | 0.165023 | -3.89717 | 9.73E-05 | 0.000425 |
| CNAG_13181 | 99.30572 | 0.621687 | 0.256491 | 2.423822 | 0.015358 | 0.0356 |
| CNAG_05475 | 4689.589 | 0.721184 | 0.178105 | 4.049207 | 5.14E-05 | 0.00024 |
| CNAG_05476 | 344.2489 | -0.72573 | 0.280886 | -2.58373 | 0.009774 | 0.024137 |
| CNAG_05477 | 1964.382 | -0.34154 | 0.143692 | -2.3769 | 0.017459 | 0.039648 |
| CNAG_05478 | 1949.054 | -0.35756 | 0.157708 | -2.26723 | 0.023376 | 0.05103 |
| CNAG_05482 | 437.1089 | 0.423493 | 0.146786 | 2.885106 | 0.003913 | 0.010894 |
| CNAG_08026 | 218.4035 | -0.69486 | 0.214657 | -3.23707 | 0.001208 | 0.003922 |
| CNAG_05496 | 6737.925 | -0.47808 | 0.140455 | -3.4038 | 0.000665 | 0.002321 |
| CNAG_05497 | 11447.4 | -1.43375 | 0.156897 | -9.13814 | 6.35E-20 | 2.87E-18 |
| CNAG_05499 | 3377.748 | -0.88185 | 0.164542 | -5.35939 | 8.35E-08 | 7.31E-07 |
| CNAG_05500 | 519.6682 | 0.603422 | 0.163597 | 3.688462 | 0.000226 | 0.000899 |
| CNAG_05505 | 744.9845 | 0.396284 | 0.137291 | 2.886455 | 0.003896 | 0.010859 |
| CNAG_07884 | 4060.901 | -0.84215 | 0.151634 | -5.55382 | 2.79E-08 | 2.74E-07 |
| CNAG_05507 | 2089.043 | -0.5068 | 0.197775 | -2.5625 | 0.010392 | 0.025463 |
| CNAG_05509 | 1970.154 | -1.45514 | 0.212238 | -6.85618 | 7.07E-12 | 1.28E-10 |

| | | | | | | |
|---|---|---|---|---|---|---|
| CNAG_05512 | 1658.04 | -0.62618 | 0.119697 | -5.23139 | 1.68E-07 | 1.37E-06 |
| CNAG_05515 | 3416.793 | 0.692848 | 0.136467 | 5.077033 | 3.83E-07 | 2.88E-06 |
| CNAG_05516 | 927.9199 | -0.672 | 0.158997 | -4.22649 | 2.37E-05 | 0.000121 |
| CNAG_05521 | 3706.543 | 0.867959 | 0.165852 | 5.233334 | 1.66E-07 | 1.36E-06 |
| CNAG_13188 | 142.1512 | 1.731098 | 0.202999 | 8.527626 | 1.49E-17 | 5.32E-16 |
| CNAG_05525 | 40304.53 | -1.09579 | 0.280453 | -3.90721 | 9.34E-05 | 0.000411 |
| CNAG_05528 | 18.51503 | 1.331854 | 0.470082 | 2.83324 | 0.004608 | 0.012608 |
| CNAG_05531 | 1941.923 | 0.728363 | 0.268745 | 2.71024 | 0.006723 | 0.017516 |
| CNAG_05534 | 1079.795 | 0.562667 | 0.175007 | 3.215115 | 0.001304 | 0.0042 |
| CNAG_05535 | 1554.425 | -0.50938 | 0.234873 | -2.16876 | 0.030101 | 0.063009 |
| CNAG_05537 | 963.9415 | -1.09571 | 0.164222 | -6.67213 | 2.52E-11 | 4.09E-10 |
| CNAG_05543 | 125.647 | 1.069929 | 0.235523 | 4.54278 | 5.55E-06 | 3.30E-05 |
| CNAG_05544 | 1477.376 | 2.895072 | 0.235966 | 12.26904 | 1.33E-34 | 1.68E-32 |
| CNAG_05546 | 989.0788 | -0.47309 | 0.219027 | -2.15997 | 0.030775 | 0.06425 |
| CNAG_05547 | 1003.038 | 3.325793 | 0.261485 | 12.71889 | 4.64E-37 | 6.59E-35 |
| CNAG_05548 | 390.3206 | 1.091659 | 0.224335 | 4.866193 | 1.14E-06 | 7.75E-06 |
| CNAG_13192 | 31.5853 | 1.573643 | 0.445794 | 3.529977 | 0.000416 | 0.001538 |
| CNAG_05553 | 586.0468 | -0.49881 | 0.15533 | -3.21131 | 0.001321 | 0.004245 |
| CNAG_05554 | 2458.293 | -0.96187 | 0.301664 | -3.18854 | 0.00143 | 0.004536 |
| CNAG_05555 | 43697.33 | -0.95473 | 0.184867 | -5.16443 | 2.41E-07 | 1.89E-06 |
| CNAG_05556 | 14241.7 | -0.98946 | 0.282359 | -3.50425 | 0.000458 | 0.001669 |
| CNAG_08027 | 805.1203 | 0.361971 | 0.122052 | 2.965724 | 0.00302 | 0.008729 |
| CNAG_05563 | 602.0247 | -0.80455 | 0.167251 | -4.81041 | 1.51E-06 | 1.01E-05 |
| CNAG_05565 | 228.9488 | -2.11382 | 0.296098 | -7.13894 | 9.41E-13 | 1.97E-11 |
| CNAG_05572 | 1028.649 | 0.405396 | 0.184332 | 2.19927 | 0.027859 | 0.059031 |
| CNAG_05573 | 1110.477 | 0.763613 | 0.212626 | 3.591346 | 0.000329 | 0.001256 |
| CNAG_05576 | 697.9617 | -1.28764 | 0.17608 | -7.31284 | 2.62E-13 | 5.77E-12 |
| CNAG_05578 | 1660.659 | -0.78208 | 0.24548 | -3.18591 | 0.001443 | 0.004568 |
| CNAG_05581 | 4458.413 | 0.849074 | 0.270241 | 3.141907 | 0.001679 | 0.005232 |
| CNAG_05583 | 870.7321 | -0.55562 | 0.149444 | -3.71794 | 0.000201 | 0.000811 |
| CNAG_13195 | 15.29305 | -1.48905 | 0.568213 | -2.62059 | 0.008778 | 0.021934 |
| CNAG_05584 | 969.3965 | -0.25854 | 0.129157 | -2.00178 | 0.045309 | 0.089358 |
| CNAG_05586 | 573.658 | 0.306531 | 0.143562 | 2.135183 | 0.032746 | 0.067764 |
| CNAG_07887 | 2231.705 | 0.776978 | 0.252663 | 3.075148 | 0.002104 | 0.006396 |
| CNAG_05590 | 3438.415 | 0.499968 | 0.195881 | 2.552406 | 0.010698 | 0.026052 |
| CNAG_05592 | 12731.55 | 1.292441 | 0.152932 | 8.451063 | 2.89E-17 | 1.01E-15 |
| CNAG_08028 | 742.1318 | -0.55734 | 0.133392 | -4.17821 | 2.94E-05 | 0.000146 |
| CNAG_05595 | 1253.771 | 0.84394 | 0.163921 | 5.148472 | 2.63E-07 | 2.04E-06 |
| CNAG_05596 | 3583.947 | -0.66314 | 0.179026 | -3.70419 | 0.000212 | 0.000852 |
| CNAG_05598 | 1708.42 | 0.787082 | 0.152835 | 5.149891 | 2.61E-07 | 2.03E-06 |
| CNAG_05599 | 575.9312 | 0.561899 | 0.169216 | 3.320599 | 0.000898 | 0.003008 |
| CNAG_05600 | 1887.788 | -0.45613 | 0.145509 | -3.13469 | 0.00172 | 0.005338 |
| CNAG_05602 | 716.9023 | -0.36951 | 0.166777 | -2.21559 | 0.02672 | 0.056936 |
| CNAG_05603 | 865.2206 | -0.8822 | 0.143829 | -6.13366 | 8.59E-10 | 1.10E-08 |
| CNAG_05606 | 699.5317 | -0.70113 | 0.258224 | -2.71518 | 0.006624 | 0.017285 |
| CNAG_05607 | 3311.086 | 1.047489 | 0.204112 | 5.131936 | 2.87E-07 | 2.21E-06 |
| CNAG_05608 | 807.4946 | 1.644594 | 0.197268 | 8.336836 | 7.63E-17 | 2.57E-15 |

| | | | | | | |
|---|---|---|---|---|---|---|
| CNAG_05609 | 1423.344 | -0.46356 | 0.230092 | -2.01467 | 0.04394 | 0.087112 |
| CNAG_05610 | 506.6208 | -0.48104 | 0.228112 | -2.10879 | 0.034963 | 0.071676 |
| CNAG_07889 | 1414.737 | -0.67104 | 0.160229 | -4.18799 | 2.81E-05 | 0.00014 |
| CNAG_05613 | 3409.191 | 0.535114 | 0.156846 | 3.41171 | 0.000646 | 0.002267 |
| CNAG_05614 | 1770.891 | -0.46526 | 0.193103 | -2.40938 | 0.01598 | 0.036783 |
| CNAG_05616 | 2910.782 | 1.254246 | 0.257611 | 4.868766 | 1.12E-06 | 7.66E-06 |
| CNAG_05618 | 2642.67 | -0.49837 | 0.12225 | -4.07665 | 4.57E-05 | 0.000216 |
| CNAG_05619 | 1200.143 | -0.25698 | 0.12409 | -2.0709 | 0.038368 | 0.077615 |
| CNAG_05621 | 1194.684 | -0.50567 | 0.140874 | -3.58951 | 0.000331 | 0.001263 |
| CNAG_05623 | 5429.848 | -0.826 | 0.168279 | -4.90853 | 9.18E-07 | 6.40E-06 |
| CNAG_05624 | 967.305 | -1.02392 | 0.160417 | -6.38289 | 1.74E-10 | 2.48E-09 |
| CNAG_08029 | 617.3068 | -0.42228 | 0.147957 | -2.85407 | 0.004316 | 0.011921 |
| CNAG_05627 | 978.1637 | -1.00693 | 0.161693 | -6.22742 | 4.74E-10 | 6.28E-09 |
| CNAG_05630 | 1318.4 | -0.47269 | 0.139382 | -3.39131 | 0.000696 | 0.002411 |
| CNAG_05631 | 8593.795 | -1.10523 | 0.125015 | -8.84082 | 9.50E-19 | 3.83E-17 |
| CNAG_05632 | 365.2155 | -0.49669 | 0.151459 | -3.27938 | 0.00104 | 0.003433 |
| CNAG_13197 | 149.4965 | 0.5822 | 0.22706 | 2.564081 | 0.010345 | 0.025355 |
| CNAG_05633 | 2976.404 | -0.92936 | 0.209469 | -4.43673 | 9.13E-06 | 5.14E-05 |
| CNAG_05635 | 516.9609 | -0.82675 | 0.32426 | -2.54966 | 0.010783 | 0.026234 |
| CNAG_05638 | 6296.089 | 0.724908 | 0.201922 | 3.590041 | 0.000331 | 0.001261 |
| CNAG_05639 | 1732.984 | 0.53986 | 0.153545 | 3.515968 | 0.000438 | 0.001605 |
| CNAG_05640 | 2019.904 | -0.5167 | 0.163871 | -3.15307 | 0.001616 | 0.005062 |
| CNAG_05641 | 41.41962 | 2.965132 | 0.479479 | 6.184065 | 6.25E-10 | 8.14E-09 |
| CNAG_13200 | 40.93158 | 1.760762 | 0.380462 | 4.627954 | 3.69E-06 | 2.29E-05 |
| CNAG_05647 | 942.9512 | -0.64101 | 0.149239 | -4.29521 | 1.75E-05 | 9.21E-05 |
| CNAG_05649 | 440.28 | 0.484941 | 0.197955 | 2.449758 | 0.014295 | 0.033468 |
| CNAG_05652 | 2589.538 | 1.081696 | 0.146228 | 7.397343 | 1.39E-13 | 3.13E-12 |
| CNAG_05653 | 13908.59 | -1.34262 | 0.167447 | -8.01817 | 1.07E-15 | 3.23E-14 |
| CNAG_05654 | 345.4224 | 1.955974 | 0.241228 | 8.108411 | 5.13E-16 | 1.59E-14 |
| CNAG_05655 | 776.1542 | -0.79613 | 0.268676 | -2.96314 | 0.003045 | 0.008783 |
| CNAG_05657 | 1537.307 | -0.72231 | 0.217526 | -3.32057 | 0.000898 | 0.003008 |
| CNAG_05658 | 114.9674 | -2.88894 | 0.271267 | -10.6498 | 1.75E-26 | 1.27E-24 |
| CNAG_05660 | 2354.96 | -0.48436 | 0.137509 | -3.52235 | 0.000428 | 0.001575 |
| CNAG_13202 | 15.79204 | 1.862923 | 0.772894 | 2.410323 | 0.015938 | 0.036735 |
| CNAG_13203 | 214.6857 | 1.192433 | 0.302236 | 3.945372 | 7.97E-05 | 0.000357 |
| CNAG_05662 | 3319.274 | -2.74454 | 0.243948 | -11.2505 | 2.30E-29 | 2.08E-27 |
| CNAG_05664 | 3707.795 | -0.57878 | 0.140434 | -4.12138 | 3.77E-05 | 0.000183 |
| CNAG_13204 | 317.5297 | 0.767405 | 0.244262 | 3.141723 | 0.00168 | 0.005234 |
| CNAG_05665 | 2832.545 | 0.748514 | 0.128603 | 5.820336 | 5.87E-09 | 6.39E-08 |
| CNAG_13206 | 395.6514 | 1.351681 | 0.200623 | 6.737403 | 1.61E-11 | 2.71E-10 |
| CNAG_07892 | 217.6739 | -0.9129 | 0.343188 | -2.66005 | 0.007813 | 0.019853 |
| CNAG_11003 | 3.85239 | 4.316338 | 2.021551 | 2.135162 | 0.032748 | 0.067764 |
| CNAG_11005 | 5.992957 | 4.010114 | 1.749958 | 2.291549 | 0.021932 | 0.048181 |
| CNAG_11016 | 5.231253 | 3.906606 | 1.718029 | 2.273888 | 0.022973 | 0.050273 |

**Table S8. Data sheets of the RNA-seq data for the WT vs. *cir1Δ* mutant.**

| | baseMean | log2FoldCh | lfcSE | stat | pvalue | padj |
|---|---|---|---|---|---|---|
| CNAG_04548 | 29.49797 | 1.849894 | 0.413161 | 4.477413 | 7.56E-06 | 3.42E-05 |
| CNAG_07304 | 568.791 | -0.72459 | 0.119374 | -6.06994 | 1.28E-09 | 9.29E-09 |
| CNAG_07305 | 59.90067 | -0.70012 | 0.310414 | -2.25544 | 0.024106 | 0.049907 |
| CNAG_00002 | 708.4145 | -0.77868 | 0.09605 | -8.10701 | 5.19E-16 | 6.05E-15 |
| CNAG_00003 | 491.0802 | -0.64586 | 0.10644 | -6.06786 | 1.30E-09 | 9.40E-09 |
| CNAG_00004 | 832.1575 | -0.25856 | 0.094413 | -2.73863 | 0.00617 | 0.015028 |
| CNAG_12001 | 99.62981 | 2.13244 | 0.273079 | 7.808872 | 5.77E-15 | 6.32E-14 |
| CNAG_00006 | 3855.291 | -0.27477 | 0.067321 | -4.08146 | 4.48E-05 | 0.000175 |
| CNAG_00007 | 4144.377 | -0.29968 | 0.061216 | -4.89547 | 9.81E-07 | 5.04E-06 |
| CNAG_00009 | 891.8573 | -0.34468 | 0.091836 | -3.75323 | 0.000175 | 0.000608 |
| CNAG_00010 | 4925.036 | 1.257745 | 0.067484 | 18.6376 | 1.59E-77 | 8.51E-76 |
| CNAG_00011 | 1064.399 | 1.458651 | 0.09982 | 14.61282 | 2.33E-48 | 7.63E-47 |
| CNAG_00012 | 1785.724 | -0.22546 | 0.079456 | -2.83754 | 0.004546 | 0.011476 |
| CNAG_00013 | 397.1122 | -0.28382 | 0.115604 | -2.45515 | 0.014082 | 0.03124 |
| CNAG_00015 | 232.4405 | 0.315939 | 0.148986 | 2.1206 | 0.033956 | 0.067069 |
| CNAG_00016 | 2912.48 | -0.47658 | 0.0676 | -7.05004 | 1.79E-12 | 1.68E-11 |
| CNAG_00018 | 2384.28 | 0.415973 | 0.088753 | 4.686838 | 2.77E-06 | 1.34E-05 |
| CNAG_00021 | 1051.99 | -0.30558 | 0.097012 | -3.14992 | 0.001633 | 0.00458 |
| CNAG_00022 | 2120.207 | 1.200887 | 0.099259 | 12.09849 | 1.08E-33 | 2.44E-32 |
| CNAG_00023 | 207.3176 | 0.563018 | 0.16586 | 3.394536 | 0.000687 | 0.002108 |
| CNAG_00024 | 13749.08 | -0.13196 | 0.062593 | -2.10824 | 0.03501 | 0.068868 |
| CNAG_00025 | 1299.259 | 0.324855 | 0.078509 | 4.137783 | 3.51E-05 | 0.00014 |
| CNAG_00026 | 5765.296 | -0.22484 | 0.057657 | -3.89966 | 9.63E-05 | 0.000355 |
| CNAG_00027 | 589.8108 | -0.30737 | 0.106373 | -2.88959 | 0.003857 | 0.009916 |
| CNAG_00028 | 570.5793 | -0.66634 | 0.107252 | -6.21284 | 5.20E-10 | 3.91E-09 |
| CNAG_00029 | 198.7606 | 0.448392 | 0.164949 | 2.718365 | 0.006561 | 0.015869 |
| CNAG_00030 | 163.5758 | -0.59647 | 0.181523 | -3.28594 | 0.001016 | 0.002986 |
| CNAG_07306 | 14.5706 | 1.343968 | 0.626473 | 2.145292 | 0.03193 | 0.063698 |
| CNAG_00034 | 47674.08 | -0.22085 | 0.067831 | -3.25582 | 0.001131 | 0.003285 |
| CNAG_00038 | 845.3812 | -0.22695 | 0.091667 | -2.47584 | 0.013292 | 0.029731 |
| CNAG_00039 | 1490.853 | 0.225677 | 0.083242 | 2.7111 | 0.006706 | 0.016189 |
| CNAG_07308 | 149.473 | 1.395493 | 0.240913 | 5.792511 | 6.93E-09 | 4.66E-08 |
| CNAG_00043 | 1028.574 | -0.49895 | 0.083373 | -5.98453 | 2.17E-09 | 1.54E-08 |
| CNAG_00048 | 57.43961 | -1.01436 | 0.296107 | -3.42567 | 0.000613 | 0.001903 |
| CNAG_00050 | 391.0857 | -0.23247 | 0.115573 | -2.01143 | 0.04428 | 0.084605 |
| CNAG_00052 | 277.7986 | -1.21882 | 0.183493 | -6.64228 | 3.09E-11 | 2.61E-10 |
| CNAG_00053 | 704.9547 | 0.901339 | 0.108483 | 8.3086 | 9.68E-17 | 1.17E-15 |
| CNAG_00054 | 236.5174 | 2.089649 | 0.197578 | 10.57633 | 3.84E-26 | 6.85E-25 |
| CNAG_00055 | 980.8145 | -0.67618 | 0.09508 | -7.11172 | 1.15E-12 | 1.09E-11 |
| CNAG_00057 | 3856.094 | -0.71537 | 0.237657 | -3.01009 | 0.002612 | 0.006992 |
| CNAG_00062 | 4281.298 | 0.287662 | 0.060325 | 4.768535 | 1.86E-06 | 9.19E-06 |
| CNAG_00063 | 1333.084 | -0.45584 | 0.078897 | -5.77766 | 7.57E-09 | 5.05E-08 |
| CNAG_12011 | 13.68901 | -1.6806 | 0.672175 | -2.50025 | 0.012411 | 0.027966 |
| CNAG_00068 | 874.0638 | -1.60241 | 0.112267 | -14.2732 | 3.22E-46 | 1.02E-44 |
| CNAG_12013 | 22.37455 | -1.14781 | 0.454163 | -2.52731 | 0.011494 | 0.026169 |
| CNAG_00070 | 1011.075 | -0.19192 | 0.092241 | -2.08066 | 0.037465 | 0.073166 |

| | | | | | | |
|---|---|---|---|---|---|---|
| CNAG_00072 | 2291.302 | -0.16792 | 0.07304 | -2.29896 | 0.021507 | 0.045112 |
| CNAG_00073 | 3611.763 | -0.26481 | 0.082931 | -3.19318 | 0.001407 | 0.003999 |
| CNAG_00074 | 3891.007 | -0.58606 | 0.078982 | -7.42016 | 1.17E-13 | 1.19E-12 |
| CNAG_00075 | 1471.069 | -0.8536 | 0.078917 | -10.8164 | 2.88E-27 | 5.38E-26 |
| CNAG_00076 | 1594.614 | 0.216994 | 0.084124 | 2.579457 | 0.009896 | 0.022889 |
| CNAG_00077 | 3281.64 | 0.302706 | 0.083127 | 3.64151 | 0.000271 | 0.000905 |
| CNAG_00078 | 2493.755 | 0.399832 | 0.065043 | 6.14724 | 7.88E-10 | 5.83E-09 |
| CNAG_00079 | 441.5475 | -0.50879 | 0.14605 | -3.48364 | 0.000495 | 0.001559 |
| CNAG_00081 | 2944.419 | -0.2418 | 0.070197 | -3.44464 | 0.000572 | 0.001784 |
| CNAG_00082 | 1153.761 | 0.179384 | 0.081027 | 2.213879 | 0.026837 | 0.05489 |
| CNAG_00083 | 1479.53 | -0.51998 | 0.07628 | -6.81675 | 9.31E-12 | 8.29E-11 |
| CNAG_00088 | 3316.172 | -0.27078 | 0.064833 | -4.17656 | 2.96E-05 | 0.000119 |
| CNAG_00090 | 1465.254 | -0.29482 | 0.077082 | -3.82475 | 0.000131 | 0.000468 |
| CNAG_00092 | 3343.798 | -0.48381 | 0.080975 | -5.97476 | 2.30E-09 | 1.63E-08 |
| CNAG_00094 | 393.8972 | 0.842823 | 0.171632 | 4.910652 | 9.08E-07 | 4.68E-06 |
| CNAG_00096 | 545.6639 | -0.27298 | 0.111937 | -2.4387 | 0.01474 | 0.032477 |
| CNAG_00099 | 2019.318 | 0.181536 | 0.072552 | 2.502163 | 0.012344 | 0.027854 |
| CNAG_00100 | 7410.898 | -0.16154 | 0.0733 | -2.20381 | 0.027538 | 0.056098 |
| CNAG_00101 | 1453.87 | -0.36379 | 0.095495 | -3.80948 | 0.000139 | 0.000495 |
| CNAG_07309 | 2930.754 | 0.901329 | 0.080636 | 11.17773 | 5.24E-29 | 1.04E-27 |
| CNAG_07310 | 3196.16 | 1.175353 | 0.069464 | 16.92037 | 3.18E-64 | 1.46E-62 |
| CNAG_00104 | 9504.794 | 0.448211 | 0.063514 | 7.056886 | 1.70E-12 | 1.60E-11 |
| CNAG_00106 | 3010.925 | -0.42885 | 0.07538 | -5.68921 | 1.28E-08 | 8.27E-08 |
| CNAG_00107 | 1641.667 | -0.49952 | 0.130431 | -3.8298 | 0.000128 | 0.000459 |
| CNAG_00108 | 4450.198 | -0.21703 | 0.06636 | -3.27056 | 0.001073 | 0.003129 |
| CNAG_00109 | 1728.724 | -0.29498 | 0.088245 | -3.34272 | 0.00083 | 0.002493 |
| CNAG_07312 | 738.4456 | -0.24131 | 0.092161 | -2.61837 | 0.008835 | 0.020731 |
| CNAG_00115 | 4979.398 | -0.16437 | 0.076921 | -2.13689 | 0.032607 | 0.064815 |
| CNAG_00117 | 3807.06 | -0.58444 | 0.079504 | -7.35118 | 1.96E-13 | 1.98E-12 |
| CNAG_00118 | 347.8509 | -0.36328 | 0.139292 | -2.60805 | 0.009106 | 0.02133 |
| CNAG_00121 | 5292.164 | -0.24393 | 0.057187 | -4.26556 | 1.99E-05 | 8.27E-05 |
| CNAG_00123 | 457.9927 | 0.459547 | 0.158018 | 2.908199 | 0.003635 | 0.009407 |
| CNAG_00126 | 3120.373 | 0.583712 | 0.104429 | 5.589581 | 2.28E-08 | 1.43E-07 |
| CNAG_00127 | 608.9325 | -0.4026 | 0.105065 | -3.83189 | 0.000127 | 0.000456 |
| CNAG_12019 | 170.7909 | 0.569405 | 0.178094 | 3.197226 | 0.001388 | 0.003953 |
| CNAG_00129 | 976.8929 | 0.502262 | 0.093223 | 5.387728 | 7.14E-08 | 4.27E-07 |
| CNAG_00131 | 175.9531 | -0.58332 | 0.170915 | -3.41292 | 0.000643 | 0.00198 |
| CNAG_00135 | 746.4342 | -0.47778 | 0.100058 | -4.77498 | 1.80E-06 | 8.90E-06 |
| CNAG_00137 | 1826.05 | 0.260384 | 0.077258 | 3.370332 | 0.000751 | 0.002285 |
| CNAG_00139 | 2298.778 | -0.50922 | 0.094288 | -5.40064 | 6.64E-08 | 3.99E-07 |
| CNAG_00140 | 4641.241 | -0.20984 | 0.079151 | -2.65113 | 0.008022 | 0.018977 |
| CNAG_00141 | 386.0639 | -0.58995 | 0.121832 | -4.84233 | 1.28E-06 | 6.48E-06 |
| CNAG_00144 | 371.0994 | 0.43494 | 0.121224 | 3.587912 | 0.000333 | 0.001096 |
| CNAG_00146 | 915.9672 | -0.37004 | 0.111283 | -3.32524 | 0.000883 | 0.002634 |
| CNAG_00150 | 3427.811 | 0.477085 | 0.08793 | 5.425736 | 5.77E-08 | 3.49E-07 |
| CNAG_00152 | 911.5428 | -0.20704 | 0.087702 | -2.36076 | 0.018238 | 0.039112 |
| CNAG_00156 | 1859.066 | 0.917971 | 0.085528 | 10.73303 | 7.12E-27 | 1.31E-25 |

| | | | | | | |
|---|---|---|---|---|---|---|
| CNAG_00158 | 260.0033 | 0.542024 | 0.145231 | 3.732155 | 0.00019 | 0.000657 |
| CNAG_00159 | 463.9156 | 0.423706 | 0.137822 | 3.074304 | 0.00211 | 0.005774 |
| CNAG_00161 | 767.7702 | 1.929976 | 0.113446 | 17.01227 | 6.66E-65 | 3.11E-63 |
| CNAG_00162 | 17637.63 | 2.126017 | 0.075813 | 28.04282 | 4.89E-173 | 7.94E-171 |
| CNAG_00163 | 1667.74 | 0.693215 | 0.082102 | 8.443372 | 3.08E-17 | 3.81E-16 |
| CNAG_12026 | 45.75919 | 1.344526 | 0.320136 | 4.199854 | 2.67E-05 | 0.000108 |
| CNAG_00164 | 603.8415 | -3.60357 | 0.131206 | -27.4649 | 4.61E-166 | 6.81E-164 |
| CNAG_00165 | 3036.546 | 0.185414 | 0.068705 | 2.698709 | 0.006961 | 0.016714 |
| CNAG_12028 | 10.83929 | 2.25756 | 0.721982 | 3.126892 | 0.001767 | 0.004921 |
| CNAG_07314 | 954.9889 | 1.669225 | 0.100694 | 16.57726 | 1.02E-61 | 4.42E-60 |
| CNAG_00171 | 620.691 | -0.35717 | 0.099668 | -3.58363 | 0.000339 | 0.001111 |
| CNAG_00172 | 1089.929 | -0.40547 | 0.099957 | -4.05644 | 4.98E-05 | 0.000193 |
| CNAG_00176 | 2787.186 | 0.501832 | 0.067567 | 7.427149 | 1.11E-13 | 1.13E-12 |
| CNAG_00177 | 100.7193 | 0.556307 | 0.21587 | 2.577043 | 0.009965 | 0.02303 |
| CNAG_00180 | 859.325 | -0.41258 | 0.095984 | -4.29843 | 1.72E-05 | 7.23E-05 |
| CNAG_00181 | 581.0464 | -0.45898 | 0.107208 | -4.28123 | 1.86E-05 | 7.75E-05 |
| CNAG_00185 | 1314.42 | -0.20278 | 0.075441 | -2.68788 | 0.007191 | 0.017225 |
| CNAG_00188 | 640.2396 | -0.62494 | 0.099962 | -6.25179 | 4.06E-10 | 3.09E-09 |
| CNAG_00189 | 754.5839 | -0.54902 | 0.109968 | -4.99257 | 5.96E-07 | 3.15E-06 |
| CNAG_00190 | 1258.689 | -0.28476 | 0.083217 | -3.42191 | 0.000622 | 0.001928 |
| CNAG_00191 | 845.1393 | 0.473869 | 0.093883 | 5.047436 | 4.48E-07 | 2.41E-06 |
| CNAG_00192 | 1957.658 | -1.85439 | 0.096759 | -19.165 | 7.25E-82 | 4.09E-80 |
| CNAG_00193 | 4466.256 | -0.30939 | 0.062902 | -4.91853 | 8.72E-07 | 4.51E-06 |
| CNAG_07316 | 497.2665 | -1.68548 | 0.118481 | -14.2257 | 6.34E-46 | 1.99E-44 |
| CNAG_07317 | 1386.882 | -0.5073 | 0.105336 | -4.81596 | 1.46E-06 | 7.33E-06 |
| CNAG_07318 | 2644.015 | -0.27764 | 0.074403 | -3.73162 | 0.00019 | 0.000658 |
| CNAG_07319 | 1603.902 | -0.7575 | 0.136615 | -5.5448 | 2.94E-08 | 1.83E-07 |
| CNAG_07320 | 1138.428 | -0.21628 | 0.080975 | -2.67099 | 0.007563 | 0.018026 |
| CNAG_07321 | 934.0709 | 0.501585 | 0.085362 | 5.87595 | 4.20E-09 | 2.88E-08 |
| CNAG_07322 | 3589.606 | -0.15028 | 0.066293 | -2.26698 | 0.023392 | 0.048601 |
| CNAG_07323 | 2429.867 | -0.21969 | 0.068459 | -3.2091 | 0.001331 | 0.003804 |
| CNAG_07324 | 1083.838 | 0.402805 | 0.091673 | 4.393918 | 1.11E-05 | 4.85E-05 |
| CNAG_07326 | 4631.039 | 0.601767 | 0.090055 | 6.682238 | 2.35E-11 | 2.00E-10 |
| CNAG_07327 | 846.603 | 1.05406 | 0.115307 | 9.141304 | 6.17E-20 | 8.71E-19 |
| CNAG_07331 | 174.1312 | -0.34249 | 0.172054 | -1.9906 | 0.046525 | 0.088169 |
| CNAG_07334 | 964.8688 | 0.752081 | 0.085535 | 8.792702 | 1.46E-18 | 1.91E-17 |
| CNAG_07335 | 244.8595 | 0.807044 | 0.144605 | 5.581008 | 2.39E-08 | 1.50E-07 |
| CNAG_07336 | 167.5221 | -0.48613 | 0.176039 | -2.76147 | 0.005754 | 0.01416 |
| CNAG_07029 | 678.3519 | -0.28125 | 0.118427 | -2.37491 | 0.017553 | 0.037824 |
| CNAG_07028 | 3203.825 | 0.167583 | 0.070742 | 2.368921 | 0.01784 | 0.038375 |
| CNAG_07027 | 1082.56 | 0.200654 | 0.089115 | 2.25163 | 0.024346 | 0.050335 |
| CNAG_00231 | 395.7034 | -0.42439 | 0.140146 | -3.02822 | 0.00246 | 0.00663 |
| CNAG_00232 | 32233.65 | -0.32832 | 0.066346 | -4.94863 | 7.47E-07 | 3.90E-06 |
| CNAG_00234 | 697.0856 | -0.30803 | 0.094752 | -3.2509 | 0.00115 | 0.003336 |
| CNAG_00235 | 4031.02 | 0.488376 | 0.077109 | 6.333606 | 2.39E-10 | 1.87E-09 |
| CNAG_00236 | 778.1946 | 0.515922 | 0.092567 | 5.573477 | 2.50E-08 | 1.57E-07 |
| CNAG_00237 | 25159.71 | 1.407787 | 0.066907 | 21.04088 | 2.77E-98 | 1.99E-96 |

| | | | | | | |
|---|---|---|---|---|---|---|
| CNAG_00238 | 18429.5 | -0.25063 | 0.057932 | -4.32622 | 1.52E-05 | 6.43E-05 |
| CNAG_00240 | 1033.702 | -0.26806 | 0.09428 | -2.84325 | 0.004466 | 0.01129 |
| CNAG_07344 | 446.3303 | 0.333974 | 0.123487 | 2.70452 | 0.00684 | 0.016454 |
| CNAG_00249 | 774.9446 | 0.316285 | 0.09823 | 3.219841 | 0.001283 | 0.003676 |
| CNAG_00250 | 1469.311 | -0.73473 | 0.104731 | -7.01536 | 2.29E-12 | 2.13E-11 |
| CNAG_00253 | 823.5459 | -0.38743 | 0.100934 | -3.83845 | 0.000124 | 0.000445 |
| CNAG_00254 | 895.4873 | -0.33391 | 0.109607 | -3.04648 | 0.002315 | 0.006277 |
| CNAG_00257 | 3880.453 | 0.25097 | 0.068416 | 3.668293 | 0.000244 | 0.000825 |
| CNAG_00260 | 3079.778 | -0.28 | 0.064216 | -4.36027 | 1.30E-05 | 5.58E-05 |
| CNAG_00261 | 15646.08 | 1.492057 | 0.079227 | 18.83262 | 4.08E-79 | 2.21E-77 |
| CNAG_00264 | 1559.264 | -0.48914 | 0.125607 | -3.89421 | 9.85E-05 | 0.000362 |
| CNAG_00266 | 848.7154 | -0.45518 | 0.09315 | -4.88647 | 1.03E-06 | 5.26E-06 |
| CNAG_00269 | 72.12152 | 2.219789 | 0.423411 | 5.242631 | 1.58E-07 | 9.04E-07 |
| CNAG_12041 | 33.55763 | 2.438777 | 0.470824 | 5.17981 | 2.22E-07 | 1.24E-06 |
| CNAG_00270 | 3458.129 | -0.68702 | 0.078 | -8.80796 | 1.27E-18 | 1.67E-17 |
| CNAG_00274 | 539.8007 | 0.393719 | 0.106122 | 3.710072 | 0.000207 | 0.000711 |
| CNAG_00275 | 1074.694 | 2.512146 | 0.129613 | 19.38189 | 1.10E-83 | 6.42E-82 |
| CNAG_00276 | 462.9181 | 1.211699 | 0.118214 | 10.25006 | 1.18E-24 | 2.05E-23 |
| CNAG_00278 | 1169.177 | -0.19177 | 0.089195 | -2.15 | 0.031555 | 0.063094 |
| CNAG_00279 | 3909.761 | -0.37746 | 0.085966 | -4.39079 | 1.13E-05 | 4.90E-05 |
| CNAG_07345 | 284.3202 | -0.46274 | 0.159192 | -2.90682 | 0.003651 | 0.009445 |
| CNAG_07114 | 1715.285 | -0.39697 | 0.081285 | -4.88366 | 1.04E-06 | 5.33E-06 |
| CNAG_07346 | 9104.222 | -0.12184 | 0.059503 | -2.04758 | 0.040601 | 0.078479 |
| CNAG_00283 | 369.3849 | -0.84881 | 0.123038 | -6.8987 | 5.25E-12 | 4.73E-11 |
| CNAG_00287 | 5233.665 | 0.143974 | 0.068429 | 2.103991 | 0.035379 | 0.069476 |
| CNAG_00288 | 1282.363 | 0.325391 | 0.082531 | 3.942641 | 8.06E-05 | 0.000301 |
| CNAG_00289 | 554.155 | -0.52428 | 0.106714 | -4.91297 | 8.97E-07 | 4.63E-06 |
| CNAG_00290 | 1183.636 | -0.30316 | 0.081154 | -3.73559 | 0.000187 | 0.000649 |
| CNAG_12042 | 26.50789 | 1.429164 | 0.43932 | 3.25313 | 0.001141 | 0.003312 |
| CNAG_00291 | 1554.841 | 0.491664 | 0.077543 | 6.340523 | 2.29E-10 | 1.79E-09 |
| CNAG_00293 | 3062.979 | -0.33564 | 0.072021 | -4.66025 | 3.16E-06 | 1.51E-05 |
| CNAG_00294 | 2670.59 | 0.168382 | 0.084409 | 1.994835 | 0.046061 | 0.087432 |
| CNAG_07348 | 507.4209 | -0.5679 | 0.112368 | -5.05391 | 4.33E-07 | 2.34E-06 |
| CNAG_07349 | 1711.653 | -0.50288 | 0.071924 | -6.99179 | 2.71E-12 | 2.50E-11 |
| CNAG_00299 | 1662.095 | 0.258891 | 0.077149 | 3.355744 | 0.000792 | 0.002389 |
| CNAG_00301 | 1502.092 | 1.670724 | 0.106327 | 15.71312 | 1.23E-55 | 4.67E-54 |
| CNAG_00302 | 244.0802 | -1.21142 | 0.173097 | -6.99852 | 2.59E-12 | 2.39E-11 |
| CNAG_00304 | 567.529 | -0.91174 | 0.114145 | -7.98755 | 1.38E-15 | 1.57E-14 |
| CNAG_12043 | 32.45517 | -0.76572 | 0.370654 | -2.06586 | 0.038842 | 0.07551 |
| CNAG_00306 | 7319.368 | -1.69562 | 0.103512 | -16.381 | 2.62E-60 | 1.11E-58 |
| CNAG_00307 | 1607.181 | 0.372285 | 0.092821 | 4.010798 | 6.05E-05 | 0.000231 |
| CNAG_00308 | 1005.474 | 0.384492 | 0.09046 | 4.250426 | 2.13E-05 | 8.80E-05 |
| CNAG_00309 | 1643.543 | 0.487359 | 0.080595 | 6.047029 | 1.48E-09 | 1.06E-08 |
| CNAG_00312 | 573.1096 | -0.28815 | 0.104424 | -2.7594 | 0.005791 | 0.014224 |
| CNAG_12046 | 543.8169 | 1.883244 | 0.190384 | 9.891795 | 4.52E-23 | 7.43E-22 |
| CNAG_07350 | 586.941 | -0.60177 | 0.117618 | -5.11627 | 3.12E-07 | 1.71E-06 |
| CNAG_00317 | 967.3674 | -0.26886 | 0.10399 | -2.58544 | 0.009725 | 0.02254 |

| | | | | | | |
|---|---|---|---|---|---|---|
| CNAG_00318 | 1677.626 | -0.37539 | 0.075747 | -4.95591 | 7.20E-07 | 3.77E-06 |
| CNAG_00319 | 3166.92 | -0.25125 | 0.082791 | -3.03473 | 0.002408 | 0.006508 |
| CNAG_07351 | 2129.983 | 0.516039 | 0.080709 | 6.393842 | 1.62E-10 | 1.28E-09 |
| CNAG_00329 | 3787.338 | 0.352853 | 0.076697 | 4.600608 | 4.21E-06 | 1.98E-05 |
| CNAG_00331 | 267.9383 | 1.936404 | 0.149836 | 12.92348 | 3.32E-38 | 8.59E-37 |
| CNAG_00332 | 1342.748 | 0.306041 | 0.077587 | 3.944506 | 8.00E-05 | 0.000299 |
| CNAG_00333 | 1339.826 | 0.219197 | 0.084934 | 2.580795 | 0.009857 | 0.022813 |
| CNAG_00335 | 1606.665 | -0.2146 | 0.072793 | -2.94805 | 0.003198 | 0.008387 |
| CNAG_00336 | 681.5817 | -0.34635 | 0.099088 | -3.49534 | 0.000473 | 0.0015 |
| CNAG_00337 | 1287.521 | -0.35218 | 0.084645 | -4.16068 | 3.17E-05 | 0.000127 |
| CNAG_07352 | 5102.126 | -0.44976 | 0.0778 | -5.78101 | 7.43E-09 | 4.96E-08 |
| CNAG_07354 | 1491.087 | -0.45076 | 0.084282 | -5.34827 | 8.88E-08 | 5.24E-07 |
| CNAG_00343 | 1315.124 | -0.48105 | 0.084487 | -5.69379 | 1.24E-08 | 8.07E-08 |
| CNAG_00345 | 2773.018 | -0.323 | 0.06995 | -4.61752 | 3.88E-06 | 1.84E-05 |
| CNAG_00347 | 1422.419 | 0.198119 | 0.075932 | 2.609169 | 0.009076 | 0.021272 |
| CNAG_00349 | 1258.754 | 4.475888 | 0.114404 | 39.12353 | 0 | 0 |
| CNAG_07356 | 6043.167 | 1.20662 | 0.059146 | 20.40072 | 1.65E-92 | 1.07E-90 |
| CNAG_00352 | 996.6743 | 0.367653 | 0.095149 | 3.863986 | 0.000112 | 0.000405 |
| CNAG_00353 | 516.0946 | -0.40369 | 0.110154 | -3.6648 | 0.000248 | 0.000836 |
| CNAG_00354 | 3572.122 | -0.44521 | 0.077608 | -5.73662 | 9.66E-09 | 6.36E-08 |
| CNAG_00355 | 2726.163 | -0.27004 | 0.087658 | -3.08058 | 0.002066 | 0.005662 |
| CNAG_00358 | 1138.8 | -0.18785 | 0.081711 | -2.29898 | 0.021506 | 0.045112 |
| CNAG_00359 | 551.6016 | -0.55444 | 0.115855 | -4.78558 | 1.70E-06 | 8.50E-06 |
| CNAG_00360 | 322.9154 | 1.175396 | 0.162052 | 7.25322 | 4.07E-13 | 4.04E-12 |
| CNAG_00361 | 2392.303 | -0.1788 | 0.087742 | -2.03783 | 0.041567 | 0.079984 |
| CNAG_00362 | 1927.222 | -0.29123 | 0.082092 | -3.54759 | 0.000389 | 0.001257 |
| CNAG_00363 | 3897.218 | -0.37258 | 0.064704 | -5.75829 | 8.50E-09 | 5.64E-08 |
| CNAG_00364 | 1421.158 | -0.24834 | 0.08641 | -2.87396 | 0.004054 | 0.010361 |
| CNAG_00365 | 1708.41 | -0.40573 | 0.077874 | -5.21004 | 1.89E-07 | 1.07E-06 |
| CNAG_00368 | 1072.182 | -0.17686 | 0.084683 | -2.08849 | 0.036753 | 0.071927 |
| CNAG_00370 | 36343.35 | -0.18611 | 0.069479 | -2.67865 | 0.007392 | 0.017666 |
| CNAG_00371 | 248.8602 | -0.35499 | 0.141276 | -2.51271 | 0.011981 | 0.027148 |
| CNAG_00372 | 2794.784 | -0.42938 | 0.080269 | -5.34925 | 8.83E-08 | 5.22E-07 |
| CNAG_00373 | 625.326 | -0.45444 | 0.100594 | -4.51759 | 6.25E-06 | 2.87E-05 |
| CNAG_00374 | 671.0683 | 0.323293 | 0.12207 | 2.648423 | 0.008087 | 0.019119 |
| CNAG_00375 | 2208.617 | -0.2834 | 0.072648 | -3.90104 | 9.58E-05 | 0.000353 |
| CNAG_12053 | 8.611709 | -2.16937 | 0.802799 | -2.70226 | 0.006887 | 0.016557 |
| CNAG_00378 | 201.713 | -1.48459 | 0.161083 | -9.2163 | 3.08E-20 | 4.42E-19 |
| CNAG_07950 | 87.31114 | 0.94674 | 0.283343 | 3.341318 | 0.000834 | 0.002503 |
| CNAG_00383 | 2640.508 | -0.26905 | 0.072536 | -3.70924 | 0.000208 | 0.000713 |
| CNAG_00387 | 491.3496 | -0.265 | 0.112693 | -2.3515 | 0.018698 | 0.039984 |
| CNAG_00388 | 1067.757 | -0.17334 | 0.086608 | -2.00146 | 0.045343 | 0.086311 |
| CNAG_00389 | 1497.867 | 0.236196 | 0.073516 | 3.212833 | 0.001314 | 0.003759 |
| CNAG_07951 | 1109.648 | -0.42398 | 0.102633 | -4.13102 | 3.61E-05 | 0.000143 |
| CNAG_00392 | 336.678 | -0.54234 | 0.124787 | -4.34613 | 1.39E-05 | 5.92E-05 |
| CNAG_00393 | 6109.625 | -0.60659 | 0.115138 | -5.26838 | 1.38E-07 | 7.91E-07 |
| CNAG_00395 | 1544.313 | 0.764838 | 0.076314 | 10.02224 | 1.22E-23 | 2.02E-22 |

| | | | | | | |
|---|---|---|---|---|---|---|
| CNAG_00396 | 848.9417 | -0.29071 | 0.086437 | -3.36319 | 0.00077 | 0.002334 |
| CNAG_00398 | 1016.897 | 0.661659 | 0.092193 | 7.176861 | 7.13E-13 | 6.89E-12 |
| CNAG_00399 | 1442.959 | -0.5089 | 0.104401 | -4.87446 | 1.09E-06 | 5.56E-06 |
| CNAG_00400 | 3723.099 | 0.751581 | 0.077446 | 9.704548 | 2.88E-22 | 4.54E-21 |
| CNAG_00402 | 6098.712 | 0.232356 | 0.070442 | 3.298523 | 0.000972 | 0.00287 |
| CNAG_00406 | 499.022 | -0.43474 | 0.146292 | -2.97174 | 0.002961 | 0.007822 |
| CNAG_00407 | 11033.21 | 1.027028 | 0.084877 | 12.10025 | 1.05E-33 | 2.39E-32 |
| CNAG_00408 | 878.8752 | -0.31274 | 0.093633 | -3.34012 | 0.000837 | 0.002513 |
| CNAG_00409 | 1792.342 | 0.72892 | 0.082149 | 8.873158 | 7.11E-19 | 9.46E-18 |
| CNAG_00410 | 4149.686 | -0.23704 | 0.062162 | -3.81335 | 0.000137 | 0.000487 |
| CNAG_00414 | 429.6272 | 0.650301 | 0.11253 | 5.778895 | 7.52E-09 | 5.02E-08 |
| CNAG_00415 | 562.8747 | -0.47074 | 0.108085 | -4.35523 | 1.33E-05 | 5.69E-05 |
| CNAG_00418 | 21632.93 | -0.23418 | 0.05717 | -4.0961 | 4.20E-05 | 0.000165 |
| CNAG_00420 | 437.4556 | -0.37314 | 0.113218 | -3.29579 | 0.000981 | 0.002896 |
| CNAG_00421 | 616.1406 | -0.46605 | 0.104026 | -4.48009 | 7.46E-06 | 3.37E-05 |
| CNAG_00423 | 479.8652 | -0.5263 | 0.112611 | -4.67364 | 2.96E-06 | 1.42E-05 |
| CNAG_00424 | 1846.882 | -0.50427 | 0.088678 | -5.68648 | 1.30E-08 | 8.39E-08 |
| CNAG_07359 | 2392.586 | 0.170681 | 0.073887 | 2.31002 | 0.020887 | 0.043921 |
| CNAG_00433 | 453.401 | 0.77062 | 0.139319 | 5.531354 | 3.18E-08 | 1.97E-07 |
| CNAG_00434 | 1226.007 | -0.42863 | 0.079938 | -5.36197 | 8.23E-08 | 4.89E-07 |
| CNAG_00437 | 261.1519 | -0.61739 | 0.163976 | -3.76511 | 0.000166 | 0.000583 |
| CNAG_00440 | 416.6591 | -0.41592 | 0.13086 | -3.17837 | 0.001481 | 0.004184 |
| CNAG_00441 | 18084.14 | 0.256071 | 0.055918 | 4.57937 | 4.66E-06 | 2.18E-05 |
| CNAG_00442 | 1901.416 | -0.19537 | 0.072413 | -2.69801 | 0.006976 | 0.016745 |
| CNAG_00443 | 408.0111 | 0.724308 | 0.143463 | 5.048739 | 4.45E-07 | 2.40E-06 |
| CNAG_00444 | 1029.412 | 2.294119 | 0.10376 | 22.10994 | 2.54E-108 | 2.04E-106 |
| CNAG_00445 | 2858.051 | -0.30361 | 0.086072 | -3.52735 | 0.00042 | 0.001348 |
| CNAG_00447 | 8001.673 | -0.23629 | 0.0566 | -4.17484 | 2.98E-05 | 0.00012 |
| CNAG_00448 | 3090.96 | -0.33547 | 0.064473 | -5.20326 | 1.96E-07 | 1.10E-06 |
| CNAG_00449 | 2400.232 | 0.314718 | 0.078332 | 4.017764 | 5.88E-05 | 0.000225 |
| CNAG_00450 | 5800.685 | 0.224862 | 0.06744 | 3.334244 | 0.000855 | 0.002563 |
| CNAG_00451 | 1609.288 | 2.278371 | 0.083577 | 27.26075 | 1.24E-163 | 1.77E-161 |
| CNAG_00452 | 68.0323 | 1.320508 | 0.2634 | 5.013317 | 5.35E-07 | 2.85E-06 |
| CNAG_00453 | 2295.846 | -0.19629 | 0.074305 | -2.64172 | 0.008249 | 0.019478 |
| CNAG_00455 | 2597.829 | 0.145135 | 0.065941 | 2.200998 | 0.027736 | 0.056445 |
| CNAG_00456 | 4601.064 | 0.569186 | 0.082193 | 6.924978 | 4.36E-12 | 3.94E-11 |
| CNAG_00457 | 26848.99 | 0.438094 | 0.062057 | 7.059516 | 1.67E-12 | 1.57E-11 |
| CNAG_00460 | 315.9912 | -1.09893 | 0.161666 | -6.7975 | 1.06E-11 | 9.38E-11 |
| CNAG_00462 | 12045.24 | 1.596675 | 0.066334 | 24.07022 | 5.13E-128 | 4.90E-126 |
| CNAG_00464 | 5468.769 | -0.34139 | 0.066018 | -5.17119 | 2.33E-07 | 1.30E-06 |
| CNAG_00465 | 1284.544 | -0.62488 | 0.151443 | -4.12617 | 3.69E-05 | 0.000146 |
| CNAG_00469 | 1174.848 | 0.19307 | 0.084894 | 2.274258 | 0.022951 | 0.047782 |
| CNAG_00473 | 956.9382 | -0.26876 | 0.097792 | -2.74825 | 0.005992 | 0.014645 |
| CNAG_00474 | 159.3296 | -0.49295 | 0.2173 | -2.26852 | 0.023298 | 0.048433 |
| CNAG_00475 | 1994.49 | 0.551431 | 0.071612 | 7.700286 | 1.36E-14 | 1.46E-13 |
| CNAG_00476 | 46.60672 | 0.937106 | 0.361154 | 2.594755 | 0.009466 | 0.022042 |
| CNAG_07361 | 2611.263 | 1.297101 | 0.070487 | 18.40205 | 1.26E-75 | 6.68E-74 |

| | | | | | | |
|---|---|---|---|---|---|---|
| CNAG_07362 | 12772.62 | 0.184521 | 0.056824 | 3.247217 | 0.001165 | 0.003375 |
| CNAG_07363 | 6291.932 | -0.51746 | 0.062879 | -8.2294 | 1.88E-16 | 2.23E-15 |
| CNAG_00484 | 395.2795 | -0.27588 | 0.126903 | -2.17395 | 0.029709 | 0.059875 |
| CNAG_00486 | 1546.511 | -0.37544 | 0.088015 | -4.26567 | 1.99E-05 | 8.27E-05 |
| CNAG_00490 | 464.4588 | -0.50892 | 0.115411 | -4.40967 | 1.04E-05 | 4.54E-05 |
| CNAG_00493 | 842.9338 | 0.232622 | 0.111259 | 2.09082 | 0.036544 | 0.071591 |
| CNAG_00496 | 828.2752 | -0.20329 | 0.092567 | -2.19619 | 0.028079 | 0.057042 |
| CNAG_00498 | 805.003 | 0.713388 | 0.120729 | 5.909022 | 3.44E-09 | 2.40E-08 |
| CNAG_00500 | 2123.466 | -0.20137 | 0.082087 | -2.45315 | 0.014161 | 0.031406 |
| CNAG_00501 | 178.7423 | 0.492549 | 0.161315 | 3.053346 | 0.002263 | 0.006154 |
| CNAG_00503 | 1502.394 | 1.104758 | 0.085466 | 12.92634 | 3.20E-38 | 8.30E-37 |
| CNAG_00504 | 1311.587 | 0.3003 | 0.102729 | 2.923239 | 0.003464 | 0.009016 |
| CNAG_00505 | 537.3144 | 0.602227 | 0.111005 | 5.425238 | 5.79E-08 | 3.50E-07 |
| CNAG_00506 | 767.2382 | -0.29783 | 0.093455 | -3.18686 | 0.001438 | 0.004075 |
| CNAG_00509 | 9555.016 | -0.29933 | 0.064222 | -4.66085 | 3.15E-06 | 1.51E-05 |
| CNAG_00512 | 3263.203 | -0.21856 | 0.066811 | -3.27137 | 0.00107 | 0.003124 |
| CNAG_00514 | 1375.026 | -0.18315 | 0.089087 | -2.0558 | 0.039802 | 0.077082 |
| CNAG_07365 | 947.9411 | 0.615277 | 0.09453 | 6.508774 | 7.58E-11 | 6.18E-10 |
| CNAG_00519 | 2632.464 | -0.7969 | 0.073957 | -10.7752 | 4.51E-27 | 8.35E-26 |
| CNAG_07953 | 951.222 | -0.51224 | 0.115333 | -4.4414 | 8.94E-06 | 3.97E-05 |
| CNAG_00520 | 6741.521 | 0.528282 | 0.092337 | 5.721258 | 1.06E-08 | 6.93E-08 |
| CNAG_00521 | 1025.576 | -0.28315 | 0.097443 | -2.90584 | 0.003663 | 0.009468 |
| CNAG_00522 | 2223.189 | -0.55694 | 0.090165 | -6.17692 | 6.54E-10 | 4.88E-09 |
| CNAG_00523 | 878.9278 | -0.50101 | 0.087286 | -5.73988 | 9.47E-09 | 6.26E-08 |
| CNAG_00524 | 671.0548 | -0.401 | 0.101398 | -3.9547 | 7.66E-05 | 0.000288 |
| CNAG_00528 | 556.6911 | -0.38888 | 0.106137 | -3.66399 | 0.000248 | 0.000838 |
| CNAG_00529 | 1150.863 | -0.41287 | 0.10063 | -4.10281 | 4.08E-05 | 0.000161 |
| CNAG_00531 | 1786.825 | -1.58196 | 0.115757 | -13.6662 | 1.62E-42 | 4.69E-41 |
| CNAG_12064 | 52.39721 | 1.674691 | 0.332478 | 5.037001 | 4.73E-07 | 2.54E-06 |
| CNAG_00535 | 2618.95 | -0.42755 | 0.07738 | -5.52535 | 3.29E-08 | 2.04E-07 |
| CNAG_00536 | 3617.03 | 0.728365 | 0.090945 | 8.008858 | 1.16E-15 | 1.33E-14 |
| CNAG_00539 | 707.5348 | 1.355498 | 0.126597 | 10.70718 | 9.42E-27 | 1.72E-25 |
| CNAG_12067 | 100.9061 | 0.795711 | 0.249852 | 3.184723 | 0.001449 | 0.004101 |
| CNAG_00540 | 480.5287 | 5.720876 | 0.231848 | 24.67516 | 1.98E-134 | 2.06E-132 |
| CNAG_12068 | 19.73429 | 2.709537 | 0.569049 | 4.761515 | 1.92E-06 | 9.48E-06 |
| CNAG_00541 | 407.734 | 0.582777 | 0.124584 | 4.677798 | 2.90E-06 | 1.40E-05 |
| CNAG_00542 | 392.8636 | 2.0925 | 0.141438 | 14.79447 | 1.59E-49 | 5.34E-48 |
| CNAG_00543 | 1649.37 | 0.674828 | 0.084029 | 8.030874 | 9.68E-16 | 1.12E-14 |
| CNAG_00545 | 1412.688 | 0.320925 | 0.077038 | 4.165791 | 3.10E-05 | 0.000124 |
| CNAG_00546 | 1796.824 | 0.491266 | 0.094713 | 5.186902 | 2.14E-07 | 1.20E-06 |
| CNAG_00547 | 3036.527 | -0.29194 | 0.069608 | -4.19404 | 2.74E-05 | 0.000111 |
| CNAG_07955 | 254.2929 | -1.30832 | 0.145917 | -8.96621 | 3.07E-19 | 4.19E-18 |
| CNAG_00550 | 650.1725 | 0.853608 | 0.102296 | 8.344461 | 7.15E-17 | 8.69E-16 |
| CNAG_00551 | 5701.426 | -0.23915 | 0.059042 | -4.05044 | 5.11E-05 | 0.000197 |
| CNAG_12069 | 88.87125 | 0.591586 | 0.241429 | 2.450354 | 0.014272 | 0.031625 |
| CNAG_07367 | 807.0699 | 5.620102 | 0.170011 | 33.05725 | 1.22E-239 | 3.43E-237 |
| CNAG_12070 | 79.01648 | 3.958264 | 0.344518 | 11.48929 | 1.49E-30 | 3.12E-29 |

| | | | | | | |
|---|---|---|---|---|---|---|
| CNAG_12071 | 11.83123 | 5.212269 | 1.178658 | 4.422206 | 9.77E-06 | 4.31E-05 |
| CNAG_07368 | 1984.932 | -0.15734 | 0.069318 | -2.2699 | 0.023214 | 0.048281 |
| CNAG_00556 | 2829.099 | -0.30046 | 0.081279 | -3.69667 | 0.000218 | 0.000745 |
| CNAG_00557 | 1654.748 | -0.35514 | 0.09407 | -3.77526 | 0.00016 | 0.000563 |
| CNAG_00558 | 2017.596 | -0.19497 | 0.070958 | -2.74772 | 0.006001 | 0.014661 |
| CNAG_00560 | 4060.827 | -0.20211 | 0.068345 | -2.95725 | 0.003104 | 0.008167 |
| CNAG_00564 | 1095.405 | 0.798824 | 0.088536 | 9.022594 | 1.84E-19 | 2.53E-18 |
| CNAG_00565 | 6926.407 | 0.151823 | 0.062786 | 2.418095 | 0.015602 | 0.034117 |
| CNAG_00566 | 1041.003 | -0.56823 | 0.085552 | -6.64192 | 3.10E-11 | 2.61E-10 |
| CNAG_00567 | 1710.448 | -0.28577 | 0.071724 | -3.98432 | 6.77E-05 | 0.000256 |
| CNAG_00570 | 3442.716 | -0.36667 | 0.066428 | -5.51983 | 3.39E-08 | 2.10E-07 |
| CNAG_00572 | 1606.34 | 0.288404 | 0.086221 | 3.344949 | 0.000823 | 0.002474 |
| CNAG_00573 | 3521.879 | 1.053018 | 0.079502 | 13.24524 | 4.81E-40 | 1.32E-38 |
| CNAG_00575 | 2056.568 | 1.083216 | 0.374782 | 2.890257 | 0.003849 | 0.009898 |
| CNAG_00576 | 1407.426 | 0.231697 | 0.077573 | 2.98682 | 0.002819 | 0.0075 |
| CNAG_00577 | 703.8161 | -0.26515 | 0.093812 | -2.82644 | 0.004707 | 0.011841 |
| CNAG_00580 | 4271.003 | 0.197032 | 0.063958 | 3.080649 | 0.002065 | 0.005662 |
| CNAG_00581 | 5809.919 | -0.35874 | 0.079738 | -4.49897 | 6.83E-06 | 3.11E-05 |
| CNAG_00582 | 2176.39 | 0.166453 | 0.084815 | 1.96255 | 0.049699 | 0.093183 |
| CNAG_00583 | 4517.357 | 0.396479 | 0.06711 | 5.907867 | 3.47E-09 | 2.41E-08 |
| CNAG_00586 | 431.465 | 2.36855 | 0.177639 | 13.33349 | 1.48E-40 | 4.09E-39 |
| CNAG_00587 | 525.1943 | 3.861883 | 0.174071 | 22.18566 | 4.72E-109 | 3.88E-107 |
| CNAG_00588 | 1120.537 | 3.16049 | 0.283442 | 11.15038 | 7.13E-29 | 1.41E-27 |
| CNAG_12078 | 142.6597 | 1.209842 | 0.232599 | 5.201417 | 1.98E-07 | 1.11E-06 |
| CNAG_00589 | 1153.75 | -0.20436 | 0.090358 | -2.26164 | 0.02372 | 0.049196 |
| CNAG_12079 | 288.1586 | 0.382701 | 0.177704 | 2.153584 | 0.031273 | 0.062607 |
| CNAG_00592 | 768.0805 | 0.958529 | 0.097574 | 9.823603 | 8.91E-23 | 1.43E-21 |
| CNAG_00593 | 883.986 | 0.690598 | 0.092561 | 7.460992 | 8.59E-14 | 8.85E-13 |
| CNAG_00594 | 3393.397 | 0.185863 | 0.067376 | 2.75858 | 0.005805 | 0.014251 |
| CNAG_00595 | 512.6608 | 0.650603 | 0.157446 | 4.132226 | 3.59E-05 | 0.000143 |
| CNAG_00596 | 240.8583 | 1.206579 | 0.161571 | 7.467802 | 8.15E-14 | 8.43E-13 |
| CNAG_12080 | 15.98176 | 3.714935 | 0.736149 | 5.046447 | 4.50E-07 | 2.42E-06 |
| CNAG_00597 | 538.8334 | 1.047606 | 0.160391 | 6.531569 | 6.51E-11 | 5.34E-10 |
| CNAG_12081 | 51.065 | 1.154155 | 0.347133 | 3.32482 | 0.000885 | 0.002637 |
| CNAG_00600 | 2970.695 | -0.19585 | 0.076012 | -2.57661 | 0.009977 | 0.023052 |
| CNAG_00602 | 5220.697 | -0.25783 | 0.071891 | -3.58633 | 0.000335 | 0.001101 |
| CNAG_00604 | 536.7545 | 0.284359 | 0.118232 | 2.405094 | 0.016168 | 0.035204 |
| CNAG_07370 | 131.6267 | 0.530891 | 0.245586 | 2.161733 | 0.030639 | 0.061444 |
| CNAG_00613 | 1227.129 | -0.35111 | 0.078865 | -4.45207 | 8.50E-06 | 3.80E-05 |
| CNAG_07373 | 17586.65 | 0.236169 | 0.065501 | 3.605596 | 0.000311 | 0.001029 |
| CNAG_07374 | 1110.991 | -0.19984 | 0.087435 | -2.28556 | 0.02228 | 0.046565 |
| CNAG_00624 | 1469.859 | -0.42631 | 0.0744 | -5.72993 | 1.00E-08 | 6.61E-08 |
| CNAG_00626 | 9998.39 | 0.449575 | 0.054748 | 8.211748 | 2.18E-16 | 2.58E-15 |
| CNAG_00627 | 206.6885 | 1.261289 | 0.227886 | 5.534742 | 3.12E-08 | 1.94E-07 |
| CNAG_00635 | 3594.882 | 0.210438 | 0.067277 | 3.127945 | 0.00176 | 0.004906 |
| CNAG_00638 | 1895.893 | -0.31832 | 0.119929 | -2.65423 | 0.007949 | 0.018836 |
| CNAG_00639 | 796.327 | 0.545799 | 0.104802 | 5.207884 | 1.91E-07 | 1.08E-06 |

| | | | | | | |
|---|---|---|---|---|---|---|
| CNAG_00640 | 92132.66 | -0.24455 | 0.059991 | -4.07648 | 4.57E-05 | 0.000178 |
| CNAG_00641 | 4573.781 | -0.24219 | 0.071744 | -3.37578 | 0.000736 | 0.002246 |
| CNAG_00642 | 795.1184 | 0.449268 | 0.088475 | 5.077921 | 3.82E-07 | 2.08E-06 |
| CNAG_07959 | 1758.173 | -0.18232 | 0.076173 | -2.39355 | 0.016686 | 0.036225 |
| CNAG_00648 | 881.5939 | -0.33318 | 0.131995 | -2.52421 | 0.011596 | 0.026357 |
| CNAG_00649 | 8376.147 | -0.12711 | 0.063953 | -1.98755 | 0.046861 | 0.088723 |
| CNAG_00651 | 318.8435 | 0.728593 | 0.13658 | 5.334556 | 9.58E-08 | 5.63E-07 |
| CNAG_00652 | 751.2915 | 0.798641 | 0.10178 | 7.846744 | 4.27E-15 | 4.74E-14 |
| CNAG_00653 | 229.3201 | 1.560594 | 0.187013 | 8.344823 | 7.13E-17 | 8.68E-16 |
| CNAG_00654 | 3918.735 | 3.354689 | 0.097075 | 34.55788 | 1.09E-261 | 3.84E-259 |
| CNAG_00656 | 62188.68 | -0.20626 | 0.062302 | -3.31068 | 0.000931 | 0.002761 |
| CNAG_00658 | 493.9736 | -0.32211 | 0.106818 | -3.0155 | 0.002566 | 0.006885 |
| CNAG_00659 | 374.3972 | -0.42464 | 0.123868 | -3.42819 | 0.000608 | 0.001887 |
| CNAG_00660 | 700.7039 | -0.23387 | 0.102973 | -2.27116 | 0.023137 | 0.048134 |
| CNAG_00662 | 2817.699 | 1.058867 | 0.077383 | 13.6834 | 1.28E-42 | 3.76E-41 |
| CNAG_00663 | 1938.013 | 1.778469 | 0.082247 | 21.62352 | 1.08E-103 | 8.35E-102 |
| CNAG_00664 | 109.1452 | 2.127225 | 0.237373 | 8.961513 | 3.20E-19 | 4.36E-18 |
| CNAG_00665 | 3985.046 | -0.15746 | 0.065743 | -2.3951 | 0.016616 | 0.036101 |
| CNAG_00668 | 3422.753 | 0.913773 | 0.066953 | 13.64788 | 2.08E-42 | 5.99E-41 |
| CNAG_00669 | 915.8334 | -0.24692 | 0.09009 | -2.74086 | 0.006128 | 0.014935 |
| CNAG_00671 | 1775.205 | 0.141441 | 0.07042 | 2.008535 | 0.044586 | 0.08509 |
| CNAG_00672 | 49158.09 | -0.26517 | 0.072524 | -3.65635 | 0.000256 | 0.000859 |
| CNAG_00673 | 2287.699 | -0.31734 | 0.070746 | -4.48564 | 7.27E-06 | 3.29E-05 |
| CNAG_00674 | 1407.117 | -0.28918 | 0.116555 | -2.48106 | 0.013099 | 0.029371 |
| CNAG_00676 | 804.8599 | -0.19203 | 0.087538 | -2.19369 | 0.028258 | 0.057363 |
| CNAG_00678 | 5737.769 | 0.842179 | 0.062926 | 13.38362 | 7.54E-41 | 2.11E-39 |
| CNAG_00680 | 1489.114 | -0.47759 | 0.085066 | -5.61443 | 1.97E-08 | 1.25E-07 |
| CNAG_00681 | 1634.956 | -0.33319 | 0.073794 | -4.51511 | 6.33E-06 | 2.90E-05 |
| CNAG_12088 | 13.40043 | -1.14244 | 0.570544 | -2.00237 | 0.045245 | 0.086164 |
| CNAG_12090 | 2.944063 | 3.077477 | 1.47645 | 2.084377 | 0.037126 | 0.072556 |
| CNAG_00683 | 1353.908 | -0.41942 | 0.105963 | -3.95823 | 7.55E-05 | 0.000284 |
| CNAG_00684 | 3457.075 | -0.43169 | 0.065284 | -6.61251 | 3.78E-11 | 3.17E-10 |
| CNAG_00686 | 11858.96 | -0.24993 | 0.071683 | -3.48656 | 0.000489 | 0.001546 |
| CNAG_00691 | 850.8162 | 0.329131 | 0.089124 | 3.692951 | 0.000222 | 0.000755 |
| CNAG_00692 | 466.6789 | 0.418344 | 0.114928 | 3.640055 | 0.000273 | 0.000909 |
| CNAG_00694 | 6509.986 | -0.1437 | 0.060249 | -2.38512 | 0.017074 | 0.036898 |
| CNAG_00696 | 706.6754 | 0.213979 | 0.095036 | 2.251558 | 0.02435 | 0.050335 |
| CNAG_00698 | 1719.67 | -0.41541 | 0.078767 | -5.27385 | 1.34E-07 | 7.69E-07 |
| CNAG_00699 | 230.3554 | 0.863581 | 0.165773 | 5.209411 | 1.89E-07 | 1.07E-06 |
| CNAG_12092 | 237.1898 | 0.523011 | 0.171495 | 3.049717 | 0.002291 | 0.006216 |
| CNAG_00700 | 14045.61 | 0.189211 | 0.054317 | 3.483477 | 0.000495 | 0.001559 |
| CNAG_00701 | 2493.789 | -0.18524 | 0.077478 | -2.39084 | 0.01681 | 0.036435 |
| CNAG_12093 | 255.4261 | 0.332856 | 0.140648 | 2.366597 | 0.017952 | 0.038603 |
| CNAG_00702 | 3850.365 | -0.23764 | 0.080702 | -2.94472 | 0.003232 | 0.008459 |
| CNAG_00703 | 43403.11 | -0.15963 | 0.070562 | -2.26227 | 0.02368 | 0.049138 |
| CNAG_00705 | 4113.844 | -0.1472 | 0.065403 | -2.25075 | 0.024402 | 0.050429 |
| CNAG_00707 | 2436.795 | -0.4046 | 0.070509 | -5.73822 | 9.57E-09 | 6.31E-08 |

| | | | | | | |
|---|---|---|---|---|---|---|
| CNAG_00708 | 2209.699 | -0.29493 | 0.078744 | -3.74544 | 0.00018 | 0.000626 |
| CNAG_00711 | 370.8835 | -0.36197 | 0.121905 | -2.96926 | 0.002985 | 0.007878 |
| CNAG_00713 | 1860.908 | -0.22706 | 0.07413 | -3.06294 | 0.002192 | 0.005972 |
| CNAG_00715 | 2601.093 | 1.105202 | 0.094148 | 11.73901 | 8.04E-32 | 1.72E-30 |
| CNAG_00716 | 34602.19 | 1.10056 | 0.091643 | 12.00925 | 3.18E-33 | 7.08E-32 |
| CNAG_00720 | 1654.148 | 0.252958 | 0.079132 | 3.196679 | 0.00139 | 0.003958 |
| CNAG_00721 | 2910.399 | -0.40444 | 0.070876 | -5.7063 | 1.15E-08 | 7.52E-08 |
| CNAG_07381 | 1651.419 | -0.26477 | 0.082924 | -3.1929 | 0.001409 | 0.004002 |
| CNAG_12098 | 12.1312 | 1.580763 | 0.659217 | 2.39794 | 0.016488 | 0.035851 |
| CNAG_12099 | 21.31527 | -1.15523 | 0.460985 | -2.50599 | 0.012211 | 0.027585 |
| CNAG_00733 | 1940.858 | -0.23626 | 0.073641 | -3.20822 | 0.001336 | 0.003813 |
| CNAG_00735 | 479.7023 | 3.659307 | 0.150768 | 24.27116 | 3.95E-130 | 3.92E-128 |
| CNAG_12101 | 81.52035 | 0.506475 | 0.247644 | 2.045168 | 0.040838 | 0.078825 |
| CNAG_00738 | 86.02155 | -1.15382 | 0.254912 | -4.52635 | 6.00E-06 | 2.76E-05 |
| CNAG_00739 | 1955.817 | -0.26103 | 0.07635 | -3.41888 | 0.000629 | 0.001947 |
| CNAG_00741 | 12499.69 | -0.33332 | 0.060077 | -5.54825 | 2.89E-08 | 1.80E-07 |
| CNAG_00744 | 1715.956 | 0.54808 | 0.082436 | 6.648527 | 2.96E-11 | 2.50E-10 |
| CNAG_00745 | 2684.223 | 0.501162 | 0.095031 | 5.273672 | 1.34E-07 | 7.69E-07 |
| CNAG_00747 | 8330.659 | -0.30418 | 0.061699 | -4.93008 | 8.22E-07 | 4.27E-06 |
| CNAG_00748 | 387.7121 | -0.43733 | 0.124605 | -3.5097 | 0.000449 | 0.001429 |
| CNAG_12105 | 24.60691 | -1.10065 | 0.431517 | -2.55066 | 0.010752 | 0.02468 |
| CNAG_00750 | 1096.594 | -0.58825 | 0.114441 | -5.14026 | 2.74E-07 | 1.52E-06 |
| CNAG_00752 | 470.0058 | -0.26455 | 0.112002 | -2.36203 | 0.018175 | 0.038989 |
| CNAG_00753 | 1238.57 | 0.26173 | 0.080684 | 3.243881 | 0.001179 | 0.003409 |
| CNAG_00754 | 9760.054 | 0.970532 | 0.06123 | 15.8506 | 1.39E-56 | 5.44E-55 |
| CNAG_00755 | 778.4898 | 0.490548 | 0.111319 | 4.406673 | 1.05E-05 | 4.60E-05 |
| CNAG_00756 | 488.346 | -0.42021 | 0.124409 | -3.3777 | 0.000731 | 0.002231 |
| CNAG_00757 | 1436.674 | -0.19593 | 0.087588 | -2.2369 | 0.025293 | 0.052006 |
| CNAG_00758 | 1072.687 | 1.183083 | 0.097892 | 12.08554 | 1.26E-33 | 2.84E-32 |
| CNAG_00760 | 1940.311 | -0.26651 | 0.070439 | -3.78355 | 0.000155 | 0.000545 |
| CNAG_00761 | 1617.719 | -0.26561 | 0.080696 | -3.29149 | 0.000997 | 0.002934 |
| CNAG_00765 | 922.0556 | 1.018724 | 0.107352 | 9.489566 | 2.32E-21 | 3.52E-20 |
| CNAG_00766 | 501.8959 | -0.52226 | 0.117373 | -4.44962 | 8.60E-06 | 3.84E-05 |
| CNAG_12111 | 22.94239 | -1.2386 | 0.454708 | -2.72395 | 0.006451 | 0.015637 |
| CNAG_00769 | 1501.102 | 0.21029 | 0.098039 | 2.144957 | 0.031956 | 0.063724 |
| CNAG_00770 | 1562.674 | 0.317034 | 0.074244 | 4.270164 | 1.95E-05 | 8.11E-05 |
| CNAG_00774 | 6657.061 | -0.35278 | 0.057227 | -6.16448 | 7.07E-10 | 5.26E-09 |
| CNAG_12113 | 121.2239 | 2.115956 | 0.220389 | 9.600999 | 7.92E-22 | 1.22E-20 |
| CNAG_00776 | 20997.53 | 0.909812 | 0.056327 | 16.15235 | 1.09E-58 | 4.49E-57 |
| CNAG_00779 | 42416.85 | -0.17125 | 0.070637 | -2.42431 | 0.015337 | 0.033629 |
| CNAG_00781 | 1163.475 | -0.61892 | 0.113492 | -5.4534 | 4.94E-08 | 3.00E-07 |
| CNAG_00782 | 902.4532 | 0.614726 | 0.095728 | 6.421561 | 1.35E-10 | 1.08E-09 |
| CNAG_00783 | 1017.238 | -0.22656 | 0.089611 | -2.52829 | 0.011462 | 0.026111 |
| CNAG_00787 | 659.3878 | -0.37082 | 0.100204 | -3.70071 | 0.000215 | 0.000735 |
| CNAG_12114 | 242.0139 | -0.85845 | 0.176004 | -4.87745 | 1.07E-06 | 5.48E-06 |
| CNAG_00790 | 724.4073 | -0.40107 | 0.091791 | -4.36938 | 1.25E-05 | 5.37E-05 |
| CNAG_00791 | 1950.014 | 0.751065 | 0.092938 | 8.081332 | 6.41E-16 | 7.43E-15 |

| | | | | | | |
|---|---|---|---|---|---|---|
| CNAG_00793 | 2025.447 | -0.23228 | 0.073311 | -3.16837 | 0.001533 | 0.004322 |
| CNAG_12117 | 147.4699 | 0.432548 | 0.187951 | 2.301383 | 0.02137 | 0.044848 |
| CNAG_00796 | 2891.956 | 1.671698 | 0.086372 | 19.35457 | 1.87E-83 | 1.08E-81 |
| CNAG_00797 | 9102.64 | 0.324898 | 0.064642 | 5.026137 | 5.00E-07 | 2.68E-06 |
| CNAG_07961 | 142.8089 | 1.177914 | 0.194208 | 6.065211 | 1.32E-09 | 9.53E-09 |
| CNAG_00798 | 81.59156 | 0.860242 | 0.287016 | 2.997188 | 0.002725 | 0.007271 |
| CNAG_00799 | 11953.73 | 0.258372 | 0.07551 | 3.421704 | 0.000622 | 0.001929 |
| CNAG_00800 | 1484.556 | 3.253029 | 0.13934 | 23.34606 | 1.51E-120 | 1.34E-118 |
| CNAG_00803 | 1308.066 | 0.296045 | 0.084146 | 3.518242 | 0.000434 | 0.00139 |
| CNAG_00805 | 480.1864 | -0.3838 | 0.130422 | -2.94278 | 0.003253 | 0.008507 |
| CNAG_00806 | 7138.843 | -0.21088 | 0.104017 | -2.02741 | 0.04262 | 0.08178 |
| CNAG_00807 | 746.4357 | -0.29438 | 0.106823 | -2.75572 | 0.005856 | 0.014351 |
| CNAG_00810 | 1275.618 | -0.47413 | 0.081283 | -5.83303 | 5.44E-09 | 3.71E-08 |
| CNAG_00811 | 2443.384 | -0.17798 | 0.076327 | -2.33176 | 0.019713 | 0.041857 |
| CNAG_00814 | 120.8409 | 1.00115 | 0.201447 | 4.9698 | 6.70E-07 | 3.52E-06 |
| CNAG_00815 | 2837.788 | 1.557054 | 0.075641 | 20.58485 | 3.75E-94 | 2.52E-92 |
| CNAG_00816 | 2174.345 | -0.22185 | 0.092127 | -2.40812 | 0.016035 | 0.034941 |
| CNAG_00818 | 789.0289 | 0.454782 | 0.122899 | 3.700443 | 0.000215 | 0.000735 |
| CNAG_00820 | 908.397 | -0.98866 | 0.092879 | -10.6446 | 1.85E-26 | 3.31E-25 |
| CNAG_00821 | 26496.93 | -0.19886 | 0.074941 | -2.65363 | 0.007963 | 0.018864 |
| CNAG_00822 | 5849.895 | 0.1776 | 0.061101 | 2.906647 | 0.003653 | 0.009448 |
| CNAG_00823 | 856.5344 | 0.291421 | 0.093967 | 3.101302 | 0.001927 | 0.005334 |
| CNAG_00828 | 314.0889 | 0.387563 | 0.143607 | 2.698776 | 0.006959 | 0.016714 |
| CNAG_00829 | 2666.345 | 0.137806 | 0.069851 | 1.972867 | 0.048511 | 0.091335 |
| CNAG_00830 | 714.3475 | 0.415494 | 0.099034 | 4.195464 | 2.72E-05 | 0.00011 |
| CNAG_00831 | 1086.809 | 0.194898 | 0.082499 | 2.362445 | 0.018155 | 0.038966 |
| CNAG_00832 | 353.3065 | -0.2776 | 0.129825 | -2.1383 | 0.032493 | 0.064635 |
| CNAG_00834 | 823.6927 | 0.392122 | 0.156072 | 2.512448 | 0.01199 | 0.027161 |
| CNAG_00836 | 987.3982 | 1.251948 | 0.095976 | 13.04439 | 6.84E-39 | 1.81E-37 |
| CNAG_00837 | 874.6321 | 0.552041 | 0.098099 | 5.627367 | 1.83E-08 | 1.17E-07 |
| CNAG_00838 | 965.5996 | 2.952668 | 0.118535 | 24.90957 | 5.86E-137 | 6.35E-135 |
| CNAG_00839 | 289.4732 | 1.445234 | 0.149558 | 9.663397 | 4.31E-22 | 6.70E-21 |
| CNAG_00840 | 405.6491 | 3.643071 | 0.153544 | 23.72651 | 1.92E-124 | 1.72E-122 |
| CNAG_00841 | 516.7955 | 0.420917 | 0.134466 | 3.130273 | 0.001746 | 0.004871 |
| CNAG_07386 | 335.0616 | 1.660035 | 0.133986 | 12.38963 | 2.97E-35 | 7.07E-34 |
| CNAG_12123 | 12.22564 | 3.59251 | 0.800139 | 4.489857 | 7.13E-06 | 3.24E-05 |
| CNAG_00843 | 917.2593 | 2.797725 | 0.123264 | 22.69698 | 4.80E-114 | 4.06E-112 |
| CNAG_00844 | 459.5852 | 5.976354 | 0.244845 | 24.40874 | 1.38E-131 | 1.39E-129 |
| CNAG_00845 | 2754.43 | 0.361189 | 0.076469 | 4.72332 | 2.32E-06 | 1.13E-05 |
| CNAG_00847 | 651.0425 | -0.23164 | 0.097405 | -2.37815 | 0.0174 | 0.037513 |
| CNAG_00848 | 3073.122 | 1.091748 | 0.295015 | 3.700659 | 0.000215 | 0.000735 |
| CNAG_00849 | 162.1783 | 1.190115 | 0.204699 | 5.813968 | 6.10E-09 | 4.14E-08 |
| CNAG_00851 | 109.0244 | -0.54518 | 0.232403 | -2.34582 | 0.018986 | 0.040512 |
| CNAG_00852 | 1957.968 | -0.19241 | 0.069652 | -2.76244 | 0.005737 | 0.014126 |
| CNAG_00854 | 2975.653 | -0.86957 | 0.096531 | -9.00822 | 2.09E-19 | 2.88E-18 |
| CNAG_07387 | 1107.006 | 2.805631 | 0.12965 | 21.63999 | 7.55E-104 | 5.90E-102 |
| CNAG_12126 | 23.94457 | 3.196999 | 0.534048 | 5.986355 | 2.15E-09 | 1.52E-08 |

| | | | | | | |
|---|---|---|---|---|---|---|
| CNAG_07388 | 11.62635 | 5.183479 | 1.166816 | 4.442412 | 8.90E-06 | 3.96E-05 |
| CNAG_07389 | 879.4098 | 0.688906 | 0.097775 | 7.045801 | 1.84E-12 | 1.73E-11 |
| CNAG_07391 | 129.265 | 1.223187 | 0.200349 | 6.105278 | 1.03E-09 | 7.50E-09 |
| CNAG_07392 | 558.0798 | 0.996276 | 0.111606 | 8.926704 | 4.39E-19 | 5.95E-18 |
| CNAG_00858 | 467.0977 | 0.444341 | 0.130464 | 3.405848 | 0.00066 | 0.002026 |
| CNAG_00859 | 198.2663 | 0.405048 | 0.165677 | 2.444798 | 0.014493 | 0.032037 |
| CNAG_07393 | 136.6967 | -0.96661 | 0.212889 | -4.54045 | 5.61E-06 | 2.60E-05 |
| CNAG_07459 | 197.54 | -0.38135 | 0.153704 | -2.48107 | 0.013099 | 0.029371 |
| CNAG_06801 | 872.6173 | -0.27905 | 0.10263 | -2.71896 | 0.006549 | 0.015851 |
| CNAG_06799 | 3577.029 | -0.33855 | 0.065527 | -5.16659 | 2.38E-07 | 1.33E-06 |
| CNAG_06796 | 1242.735 | -0.20047 | 0.082458 | -2.43115 | 0.015051 | 0.033064 |
| CNAG_06795 | 3204.851 | 0.542488 | 0.077429 | 7.006237 | 2.45E-12 | 2.26E-11 |
| CNAG_06794 | 1587.141 | -0.43191 | 0.090686 | -4.76274 | 1.91E-06 | 9.42E-06 |
| CNAG_06793 | 1357.297 | 0.259722 | 0.087415 | 2.971134 | 0.002967 | 0.007832 |
| CNAG_06784 | 153.6575 | 0.626513 | 0.184862 | 3.38909 | 0.000701 | 0.002146 |
| CNAG_06782 | 1955.887 | 0.4502 | 0.086525 | 5.203092 | 1.96E-07 | 1.10E-06 |
| CNAG_06781 | 902.4883 | -0.38094 | 0.10033 | -3.79689 | 0.000147 | 0.000519 |
| CNAG_06779 | 2728.669 | -0.39406 | 0.076243 | -5.16845 | 2.36E-07 | 1.32E-06 |
| CNAG_06777 | 371.2748 | 0.955134 | 0.132733 | 7.195904 | 6.20E-13 | 6.02E-12 |
| CNAG_06776 | 173.5202 | 0.436998 | 0.177622 | 2.460273 | 0.013883 | 0.030873 |
| CNAG_06775 | 2961.941 | -0.12791 | 0.064027 | -1.99768 | 0.045751 | 0.086946 |
| CNAG_06774 | 1391.57 | -0.19033 | 0.087652 | -2.17143 | 0.029899 | 0.060197 |
| CNAG_06772 | 1115.676 | -0.44393 | 0.105492 | -4.20816 | 2.57E-05 | 0.000104 |
| CNAG_06770 | 27216.81 | -0.24353 | 0.054034 | -4.50698 | 6.58E-06 | 3.01E-05 |
| CNAG_06767 | 11377.96 | -0.39032 | 0.068847 | -5.66936 | 1.43E-08 | 9.24E-08 |
| CNAG_06766 | 3559.146 | 0.222028 | 0.072896 | 3.045817 | 0.00232 | 0.006289 |
| CNAG_07962 | 93.10753 | -1.82962 | 0.233586 | -7.83277 | 4.77E-15 | 5.28E-14 |
| CNAG_06765 | 1641.414 | -0.34406 | 0.0872 | -3.94566 | 7.96E-05 | 0.000298 |
| CNAG_06764 | 1795.193 | -1.5719 | 0.088919 | -17.6778 | 6.22E-70 | 3.08E-68 |
| CNAG_06763 | 1052.403 | -0.36391 | 0.088881 | -4.09432 | 4.23E-05 | 0.000166 |
| CNAG_12134 | 225.3926 | 0.395066 | 0.162212 | 2.435487 | 0.014872 | 0.032749 |
| CNAG_06762 | 1982.759 | -1.60539 | 0.072607 | -22.1107 | 2.49E-108 | 2.03E-106 |
| CNAG_06761 | 655.872 | 2.078238 | 0.113174 | 18.36318 | 2.59E-75 | 1.36E-73 |
| CNAG_06760 | 964.465 | -1.07951 | 0.095117 | -11.3494 | 7.47E-30 | 1.54E-28 |
| CNAG_06758 | 1666.669 | 1.470499 | 0.238849 | 6.156598 | 7.43E-10 | 5.52E-09 |
| CNAG_12136 | 20.14768 | 2.566369 | 0.584697 | 4.389227 | 1.14E-05 | 4.94E-05 |
| CNAG_06755 | 6664.28 | -0.22206 | 0.05942 | -3.7371 | 0.000186 | 0.000645 |
| CNAG_06753 | 5491.506 | -0.19555 | 0.063299 | -3.08925 | 0.002007 | 0.005527 |
| CNAG_06752 | 1098.275 | -0.36924 | 0.087173 | -4.23566 | 2.28E-05 | 9.31E-05 |
| CNAG_06751 | 3478.083 | 0.704919 | 0.074727 | 9.433301 | 3.97E-21 | 5.99E-20 |
| CNAG_06750 | 697.0571 | -0.79742 | 0.09846 | -8.09886 | 5.55E-16 | 6.46E-15 |
| CNAG_06749 | 956.7191 | 0.350504 | 0.110356 | 3.176116 | 0.001493 | 0.004212 |
| CNAG_06747 | 27192.95 | -1.12491 | 0.093629 | -12.0146 | 2.98E-33 | 6.65E-32 |
| CNAG_06746 | 27756.83 | -1.0379 | 0.095253 | -10.8963 | 1.20E-27 | 2.28E-26 |
| CNAG_06745 | 24402.53 | -1.22666 | 0.099647 | -12.31 | 8.00E-35 | 1.87E-33 |
| CNAG_06743 | 541.78 | 3.225349 | 0.135743 | 23.76064 | 8.53E-125 | 7.79E-123 |
| CNAG_06742 | 3302.986 | 0.1996 | 0.068324 | 2.921387 | 0.003485 | 0.009064 |

| | | | | | | |
|---|---|---|---|---|---|---|
| CNAG_06741 | 3331.393 | 0.185594 | 0.06226 | 2.98096 | 0.002873 | 0.007618 |
| CNAG_06738 | 1916.903 | -0.25881 | 0.082414 | -3.14034 | 0.001688 | 0.004723 |
| CNAG_06737 | 1418.929 | -0.25706 | 0.093735 | -2.74241 | 0.006099 | 0.014878 |
| CNAG_07463 | 1511.178 | 0.227818 | 0.077006 | 2.958451 | 0.003092 | 0.00814 |
| CNAG_06730 | 4700.192 | -0.3619 | 0.073714 | -4.90958 | 9.13E-07 | 4.70E-06 |
| CNAG_06729 | 858.9593 | 0.695432 | 0.107455 | 6.471849 | 9.68E-11 | 7.84E-10 |
| CNAG_06728 | 2540.669 | 0.250019 | 0.070427 | 3.550031 | 0.000385 | 0.001247 |
| CNAG_06727 | 334.5539 | 1.190843 | 0.150591 | 7.907821 | 2.62E-15 | 2.94E-14 |
| CNAG_06723 | 8391.441 | 1.283334 | 0.063135 | 20.32681 | 7.45E-92 | 4.77E-90 |
| CNAG_06722 | 93.49608 | 1.289657 | 0.227525 | 5.668205 | 1.44E-08 | 9.29E-08 |
| CNAG_07464 | 3169.43 | -0.15565 | 0.063207 | -2.46259 | 0.013794 | 0.030717 |
| CNAG_07465 | 850.4193 | -0.19188 | 0.091882 | -2.08831 | 0.03677 | 0.071929 |
| CNAG_06717 | 2383.106 | 0.404548 | 0.075816 | 5.335951 | 9.50E-08 | 5.60E-07 |
| CNAG_06713 | 3425.277 | -0.24598 | 0.074502 | -3.3017 | 0.000961 | 0.002841 |
| CNAG_06712 | 1025.771 | -0.24305 | 0.092187 | -2.63652 | 0.008376 | 0.019739 |
| CNAG_06711 | 694.071 | -0.43576 | 0.115945 | -3.75838 | 0.000171 | 0.000598 |
| CNAG_06710 | 1719.24 | -0.41854 | 0.076068 | -5.50215 | 3.75E-08 | 2.31E-07 |
| CNAG_06703 | 966.7072 | 0.40325 | 0.087289 | 4.619714 | 3.84E-06 | 1.82E-05 |
| CNAG_07466 | 885.6219 | -1.01962 | 0.4393 | -2.32102 | 0.020286 | 0.042905 |
| CNAG_03595 | 3212.028 | 0.364795 | 0.080161 | 4.550784 | 5.34E-06 | 2.48E-05 |
| CNAG_03596 | 7943.144 | 0.194646 | 0.058785 | 3.311147 | 0.000929 | 0.002757 |
| CNAG_03598 | 1938.487 | -0.23201 | 0.06878 | -3.37321 | 0.000743 | 0.002265 |
| CNAG_07964 | 1873.408 | 0.417851 | 0.08221 | 5.082737 | 3.72E-07 | 2.03E-06 |
| CNAG_03602 | 12506.82 | 0.128906 | 0.053936 | 2.389952 | 0.016851 | 0.036513 |
| CNAG_03603 | 2397.253 | -0.19508 | 0.070821 | -2.75452 | 0.005878 | 0.014394 |
| CNAG_03604 | 1161.112 | 0.323634 | 0.083496 | 3.876025 | 0.000106 | 0.000387 |
| CNAG_03606 | 11359.81 | 0.28333 | 0.056491 | 5.015505 | 5.29E-07 | 2.83E-06 |
| CNAG_03607 | 249.1935 | -0.31707 | 0.141945 | -2.23378 | 0.025497 | 0.0524 |
| CNAG_03608 | 153.9924 | -0.36499 | 0.182138 | -2.0039 | 0.045081 | 0.085893 |
| CNAG_03609 | 566.5416 | -0.51103 | 0.106998 | -4.77605 | 1.79E-06 | 8.87E-06 |
| CNAG_12147 | 58.52828 | -0.76825 | 0.297258 | -2.58446 | 0.009753 | 0.022591 |
| CNAG_03612 | 7176.922 | -0.27954 | 0.067217 | -4.15873 | 3.20E-05 | 0.000128 |
| CNAG_03618 | 356.3774 | -0.66752 | 0.131457 | -5.07787 | 3.82E-07 | 2.08E-06 |
| CNAG_03620 | 1580.937 | 0.161642 | 0.074968 | 2.156143 | 0.031072 | 0.062252 |
| CNAG_03621 | 15393.83 | 0.257472 | 0.072832 | 3.535164 | 0.000408 | 0.001311 |
| CNAG_03623 | 1873.74 | 1.426811 | 0.079089 | 18.04057 | 9.36E-73 | 4.75E-71 |
| CNAG_03625 | 705.9968 | -0.24306 | 0.109974 | -2.21015 | 0.027094 | 0.055352 |
| CNAG_03629 | 15899.7 | 0.852129 | 0.056176 | 15.16897 | 5.68E-52 | 2.00E-50 |
| CNAG_03630 | 240.9745 | 1.050665 | 0.182505 | 5.756926 | 8.57E-09 | 5.67E-08 |
| CNAG_03631 | 961.7198 | -0.35106 | 0.096271 | -3.64659 | 0.000266 | 0.000889 |
| CNAG_03633 | 610.183 | -0.29891 | 0.105146 | -2.84277 | 0.004472 | 0.011303 |
| CNAG_03636 | 1385.833 | -0.21145 | 0.088407 | -2.39173 | 0.016769 | 0.036375 |
| CNAG_03638 | 1793.506 | -0.2601 | 0.075835 | -3.42974 | 0.000604 | 0.001877 |
| CNAG_03639 | 254.9577 | 0.377027 | 0.15636 | 2.411274 | 0.015897 | 0.034687 |
| CNAG_03642 | 323.1557 | 0.348715 | 0.13153 | 2.651228 | 0.00802 | 0.018977 |
| CNAG_03645 | 11574.81 | 0.230159 | 0.059421 | 3.87337 | 0.000107 | 0.000391 |
| CNAG_03646 | 1295.609 | -0.17495 | 0.082419 | -2.12271 | 0.033778 | 0.066816 |

| | | | | | | |
|---|---|---|---|---|---|---|
| CNAG_03648 | 3454.955 | 0.209726 | 0.068617 | 3.056469 | 0.00224 | 0.006092 |
| CNAG_03649 | 1266.062 | 0.538624 | 0.089124 | 6.043566 | 1.51E-09 | 1.08E-08 |
| CNAG_03652 | 681.6938 | 0.319858 | 0.111665 | 2.864443 | 0.004177 | 0.010654 |
| CNAG_03654 | 1260.551 | 0.393508 | 0.079723 | 4.935913 | 7.98E-07 | 4.15E-06 |
| CNAG_03655 | 3554.727 | -0.36524 | 0.084218 | -4.33687 | 1.45E-05 | 6.16E-05 |
| CNAG_03660 | 1194.115 | -0.38286 | 0.088481 | -4.32702 | 1.51E-05 | 6.41E-05 |
| CNAG_03663 | 4153.187 | 2.468461 | 0.087695 | 28.14832 | 2.51E-174 | 4.26E-172 |
| CNAG_03664 | 3701.265 | 1.155786 | 0.066966 | 17.25934 | 9.52E-67 | 4.58E-65 |
| CNAG_03665 | 2480.444 | 0.203102 | 0.068351 | 2.971442 | 0.002964 | 0.007827 |
| CNAG_03666 | 952.6017 | 2.217248 | 0.109325 | 20.28122 | 1.88E-91 | 1.20E-89 |
| CNAG_03667 | 2620.478 | 0.366538 | 0.113417 | 3.231779 | 0.00123 | 0.003539 |
| CNAG_03670 | 1123.251 | -0.28661 | 0.109938 | -2.60699 | 0.009134 | 0.021384 |
| CNAG_03671 | 370.3873 | 0.381344 | 0.123194 | 3.095483 | 0.001965 | 0.005429 |
| CNAG_03672 | 866.2158 | 0.656904 | 0.096831 | 6.784041 | 1.17E-11 | 1.02E-10 |
| CNAG_03674 | 4869.026 | 0.315686 | 0.068147 | 4.632421 | 3.61E-06 | 1.72E-05 |
| CNAG_03675 | 5650.592 | 0.260979 | 0.067094 | 3.889736 | 0.0001 | 0.000368 |
| CNAG_03676 | 833.3323 | -0.23125 | 0.087549 | -2.64137 | 0.008257 | 0.019493 |
| CNAG_03677 | 3721.719 | -0.63213 | 0.077003 | -8.20909 | 2.23E-16 | 2.63E-15 |
| CNAG_03679 | 294.183 | -0.30488 | 0.133855 | -2.27768 | 0.022746 | 0.047417 |
| CNAG_03680 | 716.478 | 0.259784 | 0.099906 | 2.600292 | 0.009314 | 0.021755 |
| CNAG_03681 | 3137.241 | -0.14421 | 0.067216 | -2.14551 | 0.031912 | 0.063698 |
| CNAG_03682 | 13108.78 | -0.52559 | 0.066444 | -7.91018 | 2.57E-15 | 2.89E-14 |
| CNAG_03683 | 814.7479 | 0.678645 | 0.124648 | 5.444497 | 5.20E-08 | 3.15E-07 |
| CNAG_03684 | 4777.389 | 0.172277 | 0.061814 | 2.787009 | 0.00532 | 0.013219 |
| CNAG_03685 | 420.4969 | -0.57291 | 0.130701 | -4.38336 | 1.17E-05 | 5.06E-05 |
| CNAG_03688 | 1748.529 | -0.29229 | 0.112725 | -2.59296 | 0.009515 | 0.022109 |
| CNAG_03694 | 1424.645 | 0.242022 | 0.082253 | 2.942395 | 0.003257 | 0.008512 |
| CNAG_03695 | 721.7648 | 0.974998 | 0.097463 | 10.00382 | 1.47E-23 | 2.42E-22 |
| CNAG_03696 | 278.54 | 1.821443 | 0.144698 | 12.58792 | 2.46E-36 | 6.08E-35 |
| CNAG_03697 | 320.5701 | 0.296914 | 0.141509 | 2.098207 | 0.035887 | 0.070388 |
| CNAG_03699 | 3992.71 | 0.1725 | 0.062339 | 2.767123 | 0.005655 | 0.013942 |
| CNAG_03700 | 1155.751 | -0.2106 | 0.081753 | -2.57607 | 0.009993 | 0.023081 |
| CNAG_03701 | 6506.444 | -0.17495 | 0.058512 | -2.98992 | 0.002791 | 0.007427 |
| CNAG_03705 | 1928.788 | -1.28127 | 0.085036 | -15.0675 | 2.65E-51 | 9.13E-50 |
| CNAG_03706 | 8087.18 | -0.26773 | 0.055993 | -4.78158 | 1.74E-06 | 8.65E-06 |
| CNAG_03707 | 977.7683 | 0.471108 | 0.084014 | 5.607486 | 2.05E-08 | 1.30E-07 |
| CNAG_03708 | 1723.487 | -0.26365 | 0.075514 | -3.49137 | 0.000481 | 0.00152 |
| CNAG_03710 | 2034.168 | 0.19899 | 0.072064 | 2.761298 | 0.005757 | 0.014161 |
| CNAG_12158 | 3.518367 | 2.724256 | 1.345643 | 2.024501 | 0.042919 | 0.082274 |
| CNAG_03712 | 378.6808 | -0.37607 | 0.131012 | -2.87047 | 0.004099 | 0.010473 |
| CNAG_03713 | 217.9844 | -1.319 | 0.639149 | -2.06368 | 0.039048 | 0.075839 |
| CNAG_03715 | 1514.538 | -0.22616 | 0.110959 | -2.03822 | 0.041528 | 0.079928 |
| CNAG_03716 | 3324.743 | -0.68474 | 0.079042 | -8.66302 | 4.59E-18 | 5.89E-17 |
| CNAG_03718 | 1432.943 | 0.261847 | 0.094167 | 2.780672 | 0.005425 | 0.013455 |
| CNAG_03719 | 1471.948 | -0.25048 | 0.109517 | -2.28713 | 0.022188 | 0.046422 |
| CNAG_03721 | 2693.97 | 0.184436 | 0.065344 | 2.82255 | 0.004764 | 0.011959 |
| CNAG_03725 | 6077.278 | -0.43639 | 0.062337 | -7.00039 | 2.55E-12 | 2.36E-11 |

| | | | | | | |
|---|---|---|---|---|---|---|
| CNAG_03726 | 3096.019 | -0.61052 | 0.079714 | -7.65891 | 1.88E-14 | 2.01E-13 |
| CNAG_12161 | 180.603 | 0.760389 | 0.177427 | 4.285638 | 1.82E-05 | 7.61E-05 |
| CNAG_03727 | 1735.33 | 0.232787 | 0.092314 | 2.52168 | 0.01168 | 0.02654 |
| CNAG_03728 | 787.6637 | -0.63439 | 0.128994 | -4.91796 | 8.74E-07 | 4.52E-06 |
| CNAG_03731 | 890.9305 | -0.3354 | 0.092419 | -3.62906 | 0.000284 | 0.000948 |
| CNAG_03733 | 296.0824 | 2.203602 | 0.162263 | 13.58043 | 5.23E-42 | 1.48E-40 |
| CNAG_03734 | 284.3304 | 0.766544 | 0.134084 | 5.71688 | 1.08E-08 | 7.09E-08 |
| CNAG_03735 | 1223.394 | -0.35057 | 0.101009 | -3.47063 | 0.000519 | 0.00163 |
| CNAG_07476 | 281.9448 | 1.867598 | 0.148897 | 12.54291 | 4.35E-36 | 1.06E-34 |
| CNAG_03742 | 4463.661 | 0.22873 | 0.062538 | 3.65743 | 0.000255 | 0.000857 |
| CNAG_03743 | 602.131 | 0.44849 | 0.10532 | 4.25834 | 2.06E-05 | 8.52E-05 |
| CNAG_03746 | 921.3316 | 4.69875 | 0.133483 | 35.20122 | 1.92E-271 | 7.78E-269 |
| CNAG_07477 | 968.3183 | 0.670911 | 0.11011 | 6.09311 | 1.11E-09 | 8.07E-09 |
| CNAG_03752 | 327.2693 | -0.361 | 0.154703 | -2.33354 | 0.01962 | 0.041713 |
| CNAG_03755 | 963.0318 | 1.407556 | 0.112738 | 12.48521 | 8.99E-36 | 2.17E-34 |
| CNAG_07479 | 300.4344 | 0.522985 | 0.135376 | 3.863219 | 0.000112 | 0.000405 |
| CNAG_07480 | 38.27334 | 0.664935 | 0.336486 | 1.976119 | 0.048141 | 0.090787 |
| CNAG_03760 | 1785.252 | -0.15283 | 0.077336 | -1.97614 | 0.048139 | 0.090787 |
| CNAG_03761 | 3414.181 | -0.15359 | 0.065535 | -2.34357 | 0.0191 | 0.040704 |
| CNAG_03763 | 3739.615 | 0.161739 | 0.078331 | 2.064811 | 0.038941 | 0.075666 |
| CNAG_03764 | 129.8356 | 1.830447 | 0.22045 | 8.303223 | 1.01E-16 | 1.22E-15 |
| CNAG_12165 | 7.449947 | 1.782489 | 0.809142 | 2.202938 | 0.027599 | 0.056194 |
| CNAG_03765 | 6633.157 | -0.26239 | 0.080998 | -3.23952 | 0.001197 | 0.003453 |
| CNAG_03766 | 1326.312 | 0.320227 | 0.078747 | 4.066534 | 4.77E-05 | 0.000185 |
| CNAG_12166 | 82.35345 | 0.46482 | 0.235653 | 1.972477 | 0.048555 | 0.091395 |
| CNAG_12168 | 51.7309 | -0.88837 | 0.326547 | -2.72048 | 0.006519 | 0.015788 |
| CNAG_03767 | 2468.69 | 0.19707 | 0.070368 | 2.800558 | 0.005101 | 0.012731 |
| CNAG_03768 | 1302.302 | 0.641693 | 0.103043 | 6.227443 | 4.74E-10 | 3.58E-09 |
| CNAG_03769 | 9327.863 | -0.3049 | 0.064768 | -4.70749 | 2.51E-06 | 1.22E-05 |
| CNAG_03771 | 10287.94 | -0.82704 | 0.262463 | -3.15109 | 0.001627 | 0.004565 |
| CNAG_03772 | 7199.003 | -2.73168 | 0.0967 | -28.249 | 1.47E-175 | 2.53E-173 |
| CNAG_07965 | 15860.78 | -0.66434 | 0.067169 | -9.89053 | 4.58E-23 | 7.50E-22 |
| CNAG_03775 | 3966.318 | -0.21583 | 0.060643 | -3.55901 | 0.000372 | 0.001211 |
| CNAG_03776 | 755.4853 | -0.50567 | 0.09115 | -5.54762 | 2.90E-08 | 1.81E-07 |
| CNAG_03777 | 354.319 | -0.61018 | 0.133462 | -4.57193 | 4.83E-06 | 2.25E-05 |
| CNAG_07481 | 582.3554 | -0.36515 | 0.107275 | -3.40382 | 0.000665 | 0.00204 |
| CNAG_03781 | 1583.362 | -0.30099 | 0.071912 | -4.18561 | 2.84E-05 | 0.000115 |
| CNAG_12169 | 5.79013 | -2.88235 | 1.072404 | -2.68774 | 0.007194 | 0.017227 |
| CNAG_03782 | 51.72812 | 2.318165 | 0.39321 | 5.895488 | 3.74E-09 | 2.59E-08 |
| CNAG_03784 | 18.95683 | -1.69999 | 0.547909 | -3.10268 | 0.001918 | 0.005315 |
| CNAG_03788 | 625.9915 | -0.7389 | 0.105913 | -6.97652 | 3.03E-12 | 2.77E-11 |
| CNAG_03790 | 4026.609 | -0.23826 | 0.071743 | -3.32101 | 0.000897 | 0.002665 |
| CNAG_03794 | 716.1781 | -0.57914 | 0.135512 | -4.27374 | 1.92E-05 | 8.00E-05 |
| CNAG_03796 | 1443.723 | -0.51319 | 0.075987 | -6.75361 | 1.44E-11 | 1.25E-10 |
| CNAG_03805 | 2345.949 | -0.14792 | 0.069487 | -2.12878 | 0.033272 | 0.065977 |
| CNAG_03806 | 830.7638 | -0.24349 | 0.092699 | -2.62663 | 0.008624 | 0.020281 |
| CNAG_03807 | 109.9753 | -0.5624 | 0.226972 | -2.47785 | 0.013218 | 0.029596 |

| | | | | | | |
|---|---|---|---|---|---|---|
| CNAG_03808 | 566.4195 | 0.848836 | 0.106442 | 7.974671 | 1.53E-15 | 1.74E-14 |
| CNAG_03809 | 2699.794 | -0.50757 | 0.083841 | -6.05397 | 1.41E-09 | 1.02E-08 |
| CNAG_03810 | 2090.181 | -0.16608 | 0.081711 | -2.03254 | 0.042099 | 0.080894 |
| CNAG_03811 | 1422.197 | -0.17862 | 0.079676 | -2.24186 | 0.02497 | 0.051499 |
| CNAG_03812 | 4034.925 | -0.30834 | 0.065272 | -4.72385 | 2.31E-06 | 1.13E-05 |
| CNAG_03814 | 3050.201 | -0.26425 | 0.082275 | -3.21176 | 0.001319 | 0.003772 |
| CNAG_03815 | 1515.012 | -0.48201 | 0.081469 | -5.91652 | 3.29E-09 | 2.29E-08 |
| CNAG_03817 | 3189.042 | 0.281243 | 0.072404 | 3.884385 | 0.000103 | 0.000375 |
| CNAG_03818 | 1542.884 | -0.18613 | 0.080178 | -2.32145 | 0.020263 | 0.042878 |
| CNAG_03819 | 10238.52 | -0.39392 | 0.065348 | -6.02806 | 1.66E-09 | 1.19E-08 |
| CNAG_03822 | 1362.834 | 0.210746 | 0.084124 | 2.505193 | 0.012238 | 0.027632 |
| CNAG_12174 | 34.55009 | 0.884281 | 0.375127 | 2.357284 | 0.018409 | 0.039418 |
| CNAG_03826 | 1165.982 | -0.39242 | 0.081574 | -4.81064 | 1.50E-06 | 7.52E-06 |
| CNAG_03828 | 173.7103 | -0.47345 | 0.186797 | -2.5346 | 0.011258 | 0.025739 |
| CNAG_03829 | 1118.698 | -0.51938 | 0.082038 | -6.331 | 2.44E-10 | 1.90E-09 |
| CNAG_03830 | 26.24843 | -0.88817 | 0.440514 | -2.01622 | 0.043777 | 0.083702 |
| CNAG_03834 | 601.2459 | -0.2548 | 0.109253 | -2.3322 | 0.01969 | 0.041841 |
| CNAG_03835 | 399.7989 | -0.44345 | 0.148547 | -2.98523 | 0.002834 | 0.007537 |
| CNAG_03836 | 1206.835 | -0.19457 | 0.088908 | -2.18847 | 0.028636 | 0.058013 |
| CNAG_03837 | 232.181 | -0.51903 | 0.160917 | -3.22542 | 0.001258 | 0.003609 |
| CNAG_03839 | 2222.829 | -0.21504 | 0.069525 | -3.09295 | 0.001982 | 0.00547 |
| CNAG_03841 | 3334.929 | 0.501141 | 0.079845 | 6.276407 | 3.46E-10 | 2.65E-09 |
| CNAG_03844 | 665.0681 | 0.416929 | 0.10785 | 3.86582 | 0.000111 | 0.000402 |
| CNAG_03847 | 1737.15 | -0.49089 | 0.084904 | -5.78164 | 7.40E-09 | 4.95E-08 |
| CNAG_03848 | 316.1861 | 0.423483 | 0.150419 | 2.81536 | 0.004872 | 0.012211 |
| CNAG_03853 | 4651.283 | -0.28221 | 0.073294 | -3.85035 | 0.000118 | 0.000426 |
| CNAG_03854 | 636.0906 | -0.37439 | 0.101537 | -3.68717 | 0.000227 | 0.00077 |
| CNAG_03855 | 386.4081 | -0.39562 | 0.127252 | -3.10898 | 0.001877 | 0.005213 |
| CNAG_03856 | 7952.723 | 0.215328 | 0.059077 | 3.644884 | 0.000268 | 0.000895 |
| CNAG_03857 | 3400.031 | 0.801598 | 0.070852 | 11.3137 | 1.12E-29 | 2.28E-28 |
| CNAG_03859 | 3074.191 | 0.173417 | 0.071851 | 2.413548 | 0.015798 | 0.03449 |
| CNAG_03861 | 12784.04 | 0.354587 | 0.073123 | 4.84921 | 1.24E-06 | 6.27E-06 |
| CNAG_03864 | 1794.302 | -0.4157 | 0.085919 | -4.83827 | 1.31E-06 | 6.60E-06 |
| CNAG_03865 | 1553.642 | -0.48436 | 0.118751 | -4.07879 | 4.53E-05 | 0.000176 |
| CNAG_03866 | 561.5421 | 0.515903 | 0.118443 | 4.35571 | 1.33E-05 | 5.68E-05 |
| CNAG_03870 | 1175.583 | 0.18516 | 0.078982 | 2.344328 | 0.019061 | 0.040643 |
| CNAG_03871 | 1043.903 | -0.2065 | 0.09042 | -2.28382 | 0.022382 | 0.046742 |
| CNAG_03872 | 1231.915 | -0.34631 | 0.097699 | -3.54465 | 0.000393 | 0.001269 |
| CNAG_03873 | 2820.896 | -0.79702 | 0.218067 | -3.65494 | 0.000257 | 0.000863 |
| CNAG_03874 | 2782.805 | 2.534447 | 0.08185 | 30.96471 | 1.61E-210 | 3.54E-208 |
| CNAG_03875 | 1496.167 | -0.52527 | 0.081982 | -6.40708 | 1.48E-10 | 1.18E-09 |
| CNAG_03876 | 2692.559 | -0.51923 | 0.085744 | -6.05556 | 1.40E-09 | 1.01E-08 |
| CNAG_03877 | 1551.092 | -0.25858 | 0.093413 | -2.76811 | 0.005638 | 0.013907 |
| CNAG_03878 | 1724.711 | -0.17106 | 0.079827 | -2.14284 | 0.032126 | 0.064031 |
| CNAG_03882 | 782.9503 | -0.303 | 0.090678 | -3.34155 | 0.000833 | 0.002501 |
| CNAG_03884 | 2432.595 | -0.50248 | 0.069695 | -7.2097 | 5.61E-13 | 5.48E-12 |
| CNAG_03885 | 2156.129 | -0.33095 | 0.081513 | -4.06009 | 4.91E-05 | 0.00019 |

| | | | | | | |
|---|---|---|---|---|---|---|
| CNAG_03887 | 670.7872 | -0.3688 | 0.100141 | -3.68285 | 0.000231 | 0.000782 |
| CNAG_03888 | 3082.239 | -0.15164 | 0.064283 | -2.35897 | 0.018326 | 0.03928 |
| CNAG_03890 | 1809.013 | -0.34031 | 0.074075 | -4.59417 | 4.34E-06 | 2.04E-05 |
| CNAG_03891 | 26894.58 | 1.053919 | 0.066397 | 15.87309 | 9.73E-57 | 3.82E-55 |
| CNAG_03892 | 12655.85 | 0.802669 | 0.067359 | 11.91623 | 9.74E-33 | 2.14E-31 |
| CNAG_03894 | 45.57833 | -2.31914 | 0.3828 | -6.05837 | 1.38E-09 | 9.93E-09 |
| CNAG_12179 | 57.90976 | -0.8335 | 0.292083 | -2.85363 | 0.004322 | 0.010993 |
| CNAG_12181 | 120.7518 | 0.972672 | 0.20059 | 4.849055 | 1.24E-06 | 6.27E-06 |
| CNAG_03896 | 299.2441 | -0.33236 | 0.132703 | -2.50455 | 0.012261 | 0.027674 |
| CNAG_12182 | 40.74616 | -0.90007 | 0.338182 | -2.66151 | 0.007779 | 0.018488 |
| CNAG_03898 | 2271.565 | 0.605401 | 0.084592 | 7.156709 | 8.26E-13 | 7.95E-12 |
| CNAG_03899 | 5537.731 | 0.705595 | 0.072789 | 9.693731 | 3.21E-22 | 5.02E-21 |
| CNAG_03900 | 1167.038 | -0.27922 | 0.093637 | -2.98195 | 0.002864 | 0.007598 |
| CNAG_03901 | 736.7676 | 0.253693 | 0.097734 | 2.59576 | 0.009438 | 0.021994 |
| CNAG_03902 | 3438.86 | 0.222186 | 0.067283 | 3.302281 | 0.000959 | 0.002839 |
| CNAG_03903 | 1189.753 | 0.392375 | 0.085469 | 4.590849 | 4.41E-06 | 2.07E-05 |
| CNAG_12185 | 47.88524 | -1.29115 | 0.330687 | -3.90446 | 9.44E-05 | 0.000348 |
| CNAG_03904 | 3357.156 | 0.153146 | 0.067785 | 2.259303 | 0.023865 | 0.049457 |
| CNAG_03906 | 427.4322 | 0.368436 | 0.126901 | 2.903343 | 0.003692 | 0.00953 |
| CNAG_03905 | 548.8287 | -0.46312 | 0.129989 | -3.56276 | 0.000367 | 0.001195 |
| CNAG_03908 | 1043.805 | 0.192402 | 0.088961 | 2.162765 | 0.030559 | 0.06133 |
| CNAG_03909 | 669.4801 | 1.169867 | 0.143199 | 8.169534 | 3.10E-16 | 3.64E-15 |
| CNAG_03910 | 1862.151 | 0.814941 | 0.320315 | 2.544188 | 0.010953 | 0.025092 |
| CNAG_03911 | 273.6764 | 1.023427 | 0.143647 | 7.124617 | 1.04E-12 | 9.95E-12 |
| CNAG_03912 | 1362.29 | 2.618863 | 0.090733 | 28.86337 | 3.44E-183 | 6.08E-181 |
| CNAG_03914 | 855.9375 | 0.280139 | 0.093041 | 3.010916 | 0.002605 | 0.006978 |
| CNAG_03915 | 572.7982 | 0.390488 | 0.099844 | 3.910958 | 9.19E-05 | 0.00034 |
| CNAG_03916 | 11586.75 | -0.18419 | 0.069242 | -2.6601 | 0.007812 | 0.01856 |
| CNAG_03919 | 636.5416 | -0.63007 | 0.116387 | -5.41357 | 6.18E-08 | 3.73E-07 |
| CNAG_03920 | 7782.429 | -0.14447 | 0.056207 | -2.57032 | 0.01016 | 0.023421 |
| CNAG_12188 | 49.8418 | 0.687238 | 0.315098 | 2.181028 | 0.029181 | 0.058928 |
| CNAG_03922 | 940.0425 | 3.041622 | 0.15532 | 19.58297 | 2.16E-85 | 1.29E-83 |
| CNAG_03924 | 383.3117 | 0.415471 | 0.124801 | 3.329081 | 0.000871 | 0.002605 |
| CNAG_03927 | 1189.769 | 0.968445 | 0.09571 | 10.11852 | 4.57E-24 | 7.73E-23 |
| CNAG_03930 | 697.9234 | -0.20935 | 0.098226 | -2.13133 | 0.033062 | 0.065608 |
| CNAG_03931 | 10352.27 | 0.114169 | 0.056042 | 2.037204 | 0.04163 | 0.080087 |
| CNAG_03933 | 433.7794 | -0.22836 | 0.115325 | -1.98011 | 0.047691 | 0.090065 |
| CNAG_03937 | 749.518 | -1.01462 | 0.104726 | -9.68825 | 3.38E-22 | 5.29E-21 |
| CNAG_03939 | 4862.001 | 1.893163 | 0.078179 | 24.21571 | 1.52E-129 | 1.49E-127 |
| CNAG_03940 | 4407.726 | 0.275596 | 0.062957 | 4.377551 | 1.20E-05 | 5.19E-05 |
| CNAG_07485 | 714.4179 | -0.24484 | 0.092928 | -2.63475 | 0.00842 | 0.019837 |
| CNAG_03946 | 931.2078 | -0.2725 | 0.111138 | -2.4519 | 0.01421 | 0.031497 |
| CNAG_03949 | 1082.808 | -0.77074 | 0.082858 | -9.30193 | 1.38E-20 | 2.01E-19 |
| CNAG_03950 | 5367.558 | -0.15671 | 0.066447 | -2.35844 | 0.018352 | 0.039316 |
| CNAG_03955 | 1106.278 | -0.20178 | 0.093844 | -2.15019 | 0.03154 | 0.063094 |
| CNAG_03956 | 804.3783 | 0.254929 | 0.08976 | 2.840129 | 0.00451 | 0.011394 |
| CNAG_03957 | 1813.46 | -0.60459 | 0.0821 | -7.36407 | 1.78E-13 | 1.80E-12 |

| | | | | | | |
|---|---|---|---|---|---|---|
| CNAG_03960 | 1315.393 | 0.559352 | 0.07741 | 7.225816 | 4.98E-13 | 4.90E-12 |
| CNAG_03961 | 2089.726 | -0.17988 | 0.069056 | -2.60478 | 0.009193 | 0.021507 |
| CNAG_03963 | 475.8948 | -0.35783 | 0.117576 | -3.04338 | 0.002339 | 0.006338 |
| CNAG_03965 | 1845.176 | 0.165865 | 0.079763 | 2.079466 | 0.037575 | 0.073362 |
| CNAG_03966 | 1989.782 | 0.671846 | 0.077647 | 8.65252 | 5.04E-18 | 6.44E-17 |
| CNAG_03967 | 2393.284 | -0.27075 | 0.070845 | -3.82171 | 0.000133 | 0.000473 |
| CNAG_03968 | 3614.566 | -0.41799 | 0.072647 | -5.7538 | 8.73E-09 | 5.77E-08 |
| CNAG_07487 | 15823.83 | -0.20952 | 0.086337 | -2.42683 | 0.015231 | 0.033432 |
| CNAG_07488 | 3164.919 | -0.31959 | 0.08278 | -3.86069 | 0.000113 | 0.000409 |
| CNAG_03971 | 18.91067 | -1.07176 | 0.510554 | -2.09922 | 0.035798 | 0.07023 |
| CNAG_03973 | 1997.724 | 1.005432 | 0.075282 | 13.35548 | 1.10E-40 | 3.06E-39 |
| CNAG_03974 | 1372.652 | 0.693879 | 0.120552 | 5.755835 | 8.62E-09 | 5.71E-08 |
| CNAG_03976 | 1111.845 | -0.60427 | 0.125684 | -4.80782 | 1.53E-06 | 7.62E-06 |
| CNAG_03977 | 1470.77 | -0.2422 | 0.092986 | -2.60472 | 0.009195 | 0.021507 |
| CNAG_03981 | 1443.7 | 0.268033 | 0.081419 | 3.292024 | 0.000995 | 0.002929 |
| CNAG_03982 | 1822.952 | 0.194723 | 0.072815 | 2.674206 | 0.007491 | 0.01787 |
| CNAG_03983 | 6410.341 | -0.62815 | 0.056946 | -11.0307 | 2.72E-28 | 5.25E-27 |
| CNAG_03984 | 1586.935 | 1.160977 | 0.113052 | 10.26937 | 9.68E-25 | 1.68E-23 |
| CNAG_03985 | 6645.362 | 2.145939 | 0.073664 | 29.13136 | 1.44E-186 | 2.66E-184 |
| CNAG_03986 | 3468.734 | -0.17151 | 0.080245 | -2.13731 | 0.032572 | 0.064778 |
| CNAG_03987 | 4011.175 | -0.38931 | 0.079674 | -4.88626 | 1.03E-06 | 5.26E-06 |
| CNAG_03988 | 142.1106 | -0.88263 | 0.202001 | -4.36943 | 1.25E-05 | 5.37E-05 |
| CNAG_03989 | 794.9008 | -0.41507 | 0.093564 | -4.43626 | 9.15E-06 | 4.06E-05 |
| CNAG_03990 | 43.57508 | -2.17886 | 0.375442 | -5.80345 | 6.50E-09 | 4.38E-08 |
| CNAG_03991 | 890.222 | -0.5392 | 0.091657 | -5.88285 | 4.03E-09 | 2.77E-08 |
| CNAG_03992 | 616.9222 | 0.252234 | 0.098245 | 2.56741 | 0.010246 | 0.023586 |
| CNAG_03994 | 748.6258 | -0.34887 | 0.09908 | -3.52111 | 0.00043 | 0.001376 |
| CNAG_03995 | 529.9081 | -0.41081 | 0.134103 | -3.06337 | 0.002189 | 0.005965 |
| CNAG_03996 | 6807.393 | 0.242093 | 0.067009 | 3.612842 | 0.000303 | 0.001002 |
| CNAG_03997 | 120.7226 | 0.760606 | 0.219865 | 3.459429 | 0.000541 | 0.001695 |
| CNAG_03998 | 1273.808 | 0.403728 | 0.079432 | 5.082676 | 3.72E-07 | 2.03E-06 |
| CNAG_03999 | 337.7861 | 1.882006 | 0.162927 | 11.55122 | 7.28E-31 | 1.53E-29 |
| CNAG_04000 | 907.0582 | -0.23608 | 0.0887 | -2.66154 | 0.007778 | 0.018488 |
| CNAG_04002 | 1412.974 | -0.53353 | 0.090292 | -5.9089 | 3.44E-09 | 2.40E-08 |
| CNAG_04004 | 99206.93 | -0.33413 | 0.057001 | -5.8618 | 4.58E-09 | 3.13E-08 |
| CNAG_04005 | 3204.746 | -0.21361 | 0.069226 | -3.08568 | 0.002031 | 0.005582 |
| CNAG_12197 | 120.5523 | -0.62368 | 0.206287 | -3.02336 | 0.0025 | 0.006724 |
| CNAG_04011 | 25099.18 | -0.38139 | 0.074469 | -5.12155 | 3.03E-07 | 1.67E-06 |
| CNAG_04013 | 1470.232 | 0.204 | 0.094427 | 2.160409 | 0.030741 | 0.061634 |
| CNAG_04014 | 4936.87 | 0.189055 | 0.064779 | 2.918487 | 0.003517 | 0.009134 |
| CNAG_12198 | 50.82786 | -1.2021 | 0.308976 | -3.89059 | 0.0001 | 0.000367 |
| CNAG_04015 | 52.43847 | -1.10821 | 0.300537 | -3.68745 | 0.000227 | 0.000769 |
| CNAG_04017 | 1196.78 | 0.821158 | 0.085547 | 9.598906 | 8.08E-22 | 1.24E-20 |
| CNAG_07489 | 1175.621 | 0.457763 | 0.120967 | 3.78419 | 0.000154 | 0.000544 |
| CNAG_04021 | 30411.85 | -0.21032 | 0.073704 | -2.8536 | 0.004323 | 0.010993 |
| CNAG_04022 | 2896.522 | -0.57045 | 0.071559 | -7.97173 | 1.56E-15 | 1.78E-14 |
| CNAG_04023 | 1660.112 | -0.82501 | 0.121457 | -6.79255 | 1.10E-11 | 9.69E-11 |

| | | | | | | |
|---|---|---|---|---|---|---|
| CNAG_04024 | 268.5063 | -0.34017 | 0.139806 | -2.43314 | 0.014968 | 0.032935 |
| CNAG_04025 | 102.8059 | 1.194736 | 0.297875 | 4.010865 | 6.05E-05 | 0.000231 |
| CNAG_04026 | 875.118 | 0.274448 | 0.098389 | 2.789412 | 0.00528 | 0.013129 |
| CNAG_04027 | 473.6225 | 1.391235 | 0.13053 | 10.65836 | 1.59E-26 | 2.87E-25 |
| CNAG_04028 | 13487.94 | -0.14333 | 0.066912 | -2.14211 | 0.032185 | 0.064113 |
| CNAG_04029 | 514.457 | 1.235129 | 0.110245 | 11.2035 | 3.92E-29 | 7.83E-28 |
| CNAG_04031 | 625.2437 | -0.26119 | 0.105271 | -2.48112 | 0.013097 | 0.029371 |
| CNAG_04032 | 6355.326 | 0.144692 | 0.061238 | 2.362784 | 0.018138 | 0.03894 |
| CNAG_04033 | 2193.001 | -0.21259 | 0.072317 | -2.93975 | 0.003285 | 0.008582 |
| CNAG_04035 | 1101.1 | 0.270467 | 0.086854 | 3.114039 | 0.001845 | 0.00513 |
| CNAG_04036 | 257.5827 | 1.02071 | 0.15937 | 6.404674 | 1.51E-10 | 1.19E-09 |
| CNAG_04037 | 2108.83 | 0.201378 | 0.089725 | 2.244398 | 0.024807 | 0.051188 |
| CNAG_12200 | 114.4661 | 1.506982 | 0.234569 | 6.424461 | 1.32E-10 | 1.06E-09 |
| CNAG_04039 | 3161.673 | 0.803606 | 0.074836 | 10.73817 | 6.74E-27 | 1.24E-25 |
| CNAG_04040 | 2032.135 | -0.32612 | 0.071427 | -4.5658 | 4.98E-06 | 2.31E-05 |
| CNAG_04044 | 1867.174 | -0.21156 | 0.081533 | -2.59474 | 0.009466 | 0.022042 |
| CNAG_04046 | 787.806 | -0.45408 | 0.112368 | -4.04103 | 5.32E-05 | 0.000205 |
| CNAG_04048 | 2861.161 | -0.25614 | 0.076202 | -3.36138 | 0.000776 | 0.002347 |
| CNAG_04050 | 5555.769 | -0.24393 | 0.078759 | -3.09717 | 0.001954 | 0.005401 |
| CNAG_04052 | 1544.934 | 0.273404 | 0.077596 | 3.523456 | 0.000426 | 0.001365 |
| CNAG_04053 | 988.9508 | 0.204919 | 0.089998 | 2.276927 | 0.022791 | 0.047498 |
| CNAG_04055 | 744.4831 | 0.399052 | 0.096827 | 4.121279 | 3.77E-05 | 0.000149 |
| CNAG_04056 | 2603.306 | 1.218776 | 0.072788 | 16.74408 | 6.26E-63 | 2.81E-61 |
| CNAG_07491 | 1077.513 | 1.279498 | 0.113045 | 11.3185 | 1.06E-29 | 2.16E-28 |
| CNAG_07492 | 39.75301 | 1.260264 | 0.352681 | 3.573382 | 0.000352 | 0.001151 |
| CNAG_07493 | 163.2604 | 1.399439 | 0.186241 | 7.514147 | 5.73E-14 | 5.98E-13 |
| CNAG_04062 | 3167.545 | 0.212394 | 0.063066 | 3.367812 | 0.000758 | 0.002301 |
| CNAG_04064 | 7286.707 | 0.489169 | 0.078853 | 6.203533 | 5.52E-10 | 4.14E-09 |
| CNAG_04065 | 181.067 | -1.16993 | 0.19531 | -5.99013 | 2.10E-09 | 1.49E-08 |
| CNAG_04067 | 373.1085 | -0.45722 | 0.136217 | -3.35659 | 0.000789 | 0.002383 |
| CNAG_04068 | 17653.83 | -0.15425 | 0.075268 | -2.04935 | 0.040428 | 0.07822 |
| CNAG_12201 | 122.8126 | -0.87331 | 0.203249 | -4.29674 | 1.73E-05 | 7.28E-05 |
| CNAG_04069 | 3026.105 | -0.16917 | 0.06986 | -2.4215 | 0.015457 | 0.033845 |
| CNAG_04070 | 130.1682 | -0.64315 | 0.201226 | -3.19614 | 0.001393 | 0.003964 |
| CNAG_04071 | 2707.554 | 0.179212 | 0.070904 | 2.527517 | 0.011487 | 0.026161 |
| CNAG_04073 | 2869.957 | 0.398556 | 0.072605 | 5.489362 | 4.03E-08 | 2.47E-07 |
| CNAG_04076 | 848.5114 | 0.373691 | 0.087434 | 4.273971 | 1.92E-05 | 7.99E-05 |
| CNAG_12205 | 28.81701 | 1.64877 | 0.458397 | 3.596815 | 0.000322 | 0.001061 |
| CNAG_04078 | 1963.956 | 0.25949 | 0.072788 | 3.565015 | 0.000364 | 0.001185 |
| CNAG_04079 | 1829.294 | 0.233135 | 0.079141 | 2.945795 | 0.003221 | 0.008432 |
| CNAG_12209 | 150.7929 | 0.422599 | 0.20553 | 2.05614 | 0.039769 | 0.077036 |
| CNAG_12210 | 256.8104 | 0.600274 | 0.137048 | 4.380019 | 1.19E-05 | 5.13E-05 |
| CNAG_04086 | 176.8476 | 0.628561 | 0.171668 | 3.6615 | 0.000251 | 0.000845 |
| CNAG_04087 | 221.8126 | 0.310953 | 0.149734 | 2.076699 | 0.037829 | 0.073789 |
| CNAG_04089 | 2587.229 | -0.23952 | 0.082478 | -2.90403 | 0.003684 | 0.009515 |
| CNAG_04090 | 2492.377 | -0.53077 | 0.083886 | -6.32721 | 2.50E-10 | 1.95E-09 |
| CNAG_04091 | 204.9462 | 1.159249 | 0.179525 | 6.457326 | 1.07E-10 | 8.59E-10 |

| | | | | | | |
|---|---|---|---|---|---|---|
| CNAG_04092 | 93.73175 | -0.61645 | 0.218032 | -2.82733 | 0.004694 | 0.011815 |
| CNAG_04093 | 71.92766 | -0.90903 | 0.274693 | -3.30924 | 0.000936 | 0.002773 |
| CNAG_04097 | 162.434 | 0.909424 | 0.17496 | 5.197905 | 2.02E-07 | 1.13E-06 |
| CNAG_04098 | 452.9101 | -0.48534 | 0.117884 | -4.11711 | 3.84E-05 | 0.000152 |
| CNAG_04099 | 427.9522 | 0.258182 | 0.111265 | 2.32043 | 0.020318 | 0.04295 |
| CNAG_07497 | 59.43192 | -1.35253 | 0.305909 | -4.42135 | 9.81E-06 | 4.33E-05 |
| CNAG_12212 | 594.6968 | 0.458687 | 0.112041 | 4.093915 | 4.24E-05 | 0.000166 |
| CNAG_06984 | 1253.675 | 0.319977 | 0.08433 | 3.794336 | 0.000148 | 0.000524 |
| CNAG_07498 | 876.1435 | 0.7091 | 0.089106 | 7.957947 | 1.75E-15 | 1.98E-14 |
| CNAG_06973 | 126.3821 | 0.623627 | 0.211694 | 2.945887 | 0.00322 | 0.008432 |
| CNAG_06963 | 8683.637 | -0.33374 | 0.084614 | -3.94432 | 8.00E-05 | 0.000299 |
| CNAG_06962 | 428.1289 | -0.57627 | 0.118919 | -4.84591 | 1.26E-06 | 6.37E-06 |
| CNAG_03081 | 3104.815 | 0.198868 | 0.061566 | 3.230134 | 0.001237 | 0.003557 |
| CNAG_03079 | 703.9364 | -0.3117 | 0.098007 | -3.1804 | 0.001471 | 0.004158 |
| CNAG_03078 | 1530.993 | -0.22284 | 0.078353 | -2.8441 | 0.004454 | 0.011274 |
| CNAG_03076 | 212.5413 | 0.717108 | 0.15354 | 4.670492 | 3.00E-06 | 1.44E-05 |
| CNAG_12217 | 829.1505 | 1.230059 | 0.120186 | 10.23466 | 1.39E-24 | 2.38E-23 |
| CNAG_03072 | 44265.27 | 0.41312 | 0.053327 | 7.746978 | 9.41E-15 | 1.02E-13 |
| CNAG_03071 | 779.4531 | 0.50613 | 0.100947 | 5.013824 | 5.34E-07 | 2.85E-06 |
| CNAG_03070 | 1207.86 | -0.38432 | 0.082732 | -4.64537 | 3.39E-06 | 1.62E-05 |
| CNAG_03069 | 831.035 | -0.42582 | 0.098224 | -4.33523 | 1.46E-05 | 6.20E-05 |
| CNAG_03068 | 400.1139 | -1.67928 | 0.141223 | -11.891 | 1.32E-32 | 2.88E-31 |
| CNAG_07502 | 9716.78 | 0.331171 | 0.06006 | 5.513986 | 3.51E-08 | 2.17E-07 |
| CNAG_03065 | 5346.353 | -0.39289 | 0.07485 | -5.24901 | 1.53E-07 | 8.77E-07 |
| CNAG_03064 | 2425.134 | 0.197644 | 0.086901 | 2.274359 | 0.022944 | 0.047782 |
| CNAG_03063 | 3856.242 | 0.420295 | 0.066843 | 6.287815 | 3.22E-10 | 2.48E-09 |
| CNAG_07967 | 1070.897 | -0.29775 | 0.101161 | -2.94331 | 0.003247 | 0.008495 |
| CNAG_03059 | 902.5381 | 0.443535 | 0.090642 | 4.893267 | 9.92E-07 | 5.09E-06 |
| CNAG_03058 | 10784.55 | -0.94757 | 0.210454 | -4.50251 | 6.72E-06 | 3.06E-05 |
| CNAG_03057 | 293.2537 | -0.70713 | 0.169906 | -4.16188 | 3.16E-05 | 0.000126 |
| CNAG_12220 | 7.122724 | 1.845759 | 0.818242 | 2.255761 | 0.024086 | 0.049877 |
| CNAG_03056 | 63.30021 | -1.72398 | 0.296668 | -5.81114 | 6.20E-09 | 4.21E-08 |
| CNAG_03051 | 1071.808 | 0.624848 | 0.08626 | 7.243775 | 4.36E-13 | 4.31E-12 |
| CNAG_03049 | 2717.636 | -0.27058 | 0.068171 | -3.96914 | 7.21E-05 | 0.000272 |
| CNAG_03047 | 815.1413 | 0.907579 | 0.092773 | 9.782791 | 1.33E-22 | 2.12E-21 |
| CNAG_12222 | 65.39244 | 0.536919 | 0.269318 | 1.993623 | 0.046193 | 0.087642 |
| CNAG_03040 | 2114.273 | -0.60734 | 0.096622 | -6.28572 | 3.26E-10 | 2.50E-09 |
| CNAG_03038 | 3202.722 | 0.206263 | 0.08814 | 2.340169 | 0.019275 | 0.041034 |
| CNAG_12223 | 101.9334 | 0.529867 | 0.260496 | 2.034072 | 0.041944 | 0.080635 |
| CNAG_03033 | 769.1853 | 0.216822 | 0.09481 | 2.286907 | 0.022201 | 0.046436 |
| CNAG_03031 | 636.4608 | -0.52304 | 0.09848 | -5.31111 | 1.09E-07 | 6.35E-07 |
| CNAG_03027 | 3485.684 | -0.13748 | 0.068729 | -2.00035 | 0.045463 | 0.086498 |
| CNAG_03025 | 1699.73 | -0.23681 | 0.076801 | -3.08345 | 0.002046 | 0.005613 |
| CNAG_03024 | 1424.822 | 0.212868 | 0.081926 | 2.598278 | 0.009369 | 0.021845 |
| CNAG_03023 | 1810.117 | 0.263119 | 0.08102 | 3.247562 | 0.001164 | 0.003372 |
| CNAG_03022 | 674.5312 | -0.5152 | 0.103587 | -4.97356 | 6.57E-07 | 3.46E-06 |
| CNAG_03020 | 760.0473 | -0.33469 | 0.09563 | -3.49986 | 0.000465 | 0.001477 |

| | | | | | | |
|---|---|---|---|---|---|---|
| CNAG_03019 | 1613.886 | 0.296231 | 0.084508 | 3.505347 | 0.000456 | 0.001451 |
| CNAG_03016 | 1083.369 | -0.19751 | 0.081673 | -2.41834 | 0.015591 | 0.034103 |
| CNAG_03013 | 1136.279 | 0.834093 | 0.08543 | 9.763474 | 1.62E-22 | 2.55E-21 |
| CNAG_03012 | 57063.79 | 0.198323 | 0.097231 | 2.039712 | 0.041379 | 0.07968 |
| CNAG_12229 | 56.68434 | 1.345707 | 0.309547 | 4.347341 | 1.38E-05 | 5.89E-05 |
| CNAG_03011 | 608.7692 | 3.097129 | 0.11894 | 26.03951 | 1.77E-149 | 2.28E-147 |
| CNAG_03009 | 675.4265 | -0.37923 | 0.125143 | -3.03034 | 0.002443 | 0.006593 |
| CNAG_03007 | 12481.93 | 2.797286 | 0.086586 | 32.30632 | 5.70E-229 | 1.49E-226 |
| CNAG_12230 | 85.24824 | 0.467178 | 0.227478 | 2.053731 | 0.040002 | 0.07745 |
| CNAG_03004 | 1195.077 | -0.83292 | 0.094832 | -8.78312 | 1.59E-18 | 2.07E-17 |
| CNAG_03000 | 56225.19 | -0.27348 | 0.079815 | -3.42639 | 0.000612 | 0.001898 |
| CNAG_02999 | 777.1163 | -0.25096 | 0.102551 | -2.44717 | 0.014398 | 0.031853 |
| CNAG_02998 | 1830.42 | -0.40834 | 0.070874 | -5.76154 | 8.34E-09 | 5.54E-08 |
| CNAG_02995 | 3710.444 | -0.40922 | 0.081879 | -4.9979 | 5.80E-07 | 3.07E-06 |
| CNAG_02994 | 9596.967 | -0.96398 | 0.085172 | -11.318 | 1.07E-29 | 2.17E-28 |
| CNAG_02993 | 1074.547 | -0.24552 | 0.092587 | -2.65171 | 0.008009 | 0.018956 |
| CNAG_02992 | 583.6595 | 0.463527 | 0.10977 | 4.222713 | 2.41E-05 | 9.82E-05 |
| CNAG_02991 | 7961.987 | -0.20497 | 0.066269 | -3.09295 | 0.001982 | 0.00547 |
| CNAG_02990 | 2419.551 | 0.654934 | 0.102759 | 6.37349 | 1.85E-10 | 1.46E-09 |
| CNAG_02986 | 1133.015 | -0.25653 | 0.12441 | -2.06201 | 0.039207 | 0.076092 |
| CNAG_02984 | 62.08059 | -2.10612 | 0.311021 | -6.77165 | 1.27E-11 | 1.11E-10 |
| CNAG_02982 | 3053.373 | 0.192366 | 0.073324 | 2.623486 | 0.008704 | 0.020446 |
| CNAG_02978 | 397.4024 | 1.807336 | 0.137395 | 13.15436 | 1.61E-39 | 4.32E-38 |
| CNAG_02977 | 2846.645 | 0.187668 | 0.06765 | 2.774078 | 0.005536 | 0.013689 |
| CNAG_02974 | 26253.5 | -0.36967 | 0.072537 | -5.09624 | 3.46E-07 | 1.89E-06 |
| CNAG_02973 | 1004.69 | -0.27283 | 0.082498 | -3.30705 | 0.000943 | 0.002793 |
| CNAG_02969 | 1130.82 | 1.578982 | 0.123852 | 12.7489 | 3.16E-37 | 7.98E-36 |
| CNAG_02968 | 743.2274 | 1.80435 | 0.121658 | 14.83138 | 9.18E-50 | 3.11E-48 |
| CNAG_02966 | 260.3101 | 1.160928 | 0.210389 | 5.518003 | 3.43E-08 | 2.12E-07 |
| CNAG_02964 | 458.073 | 0.241491 | 0.117273 | 2.059225 | 0.039473 | 0.076553 |
| CNAG_02963 | 742.3496 | -0.23716 | 0.109167 | -2.17247 | 0.02982 | 0.060069 |
| CNAG_02960 | 416.8997 | -0.78285 | 0.138274 | -5.66157 | 1.50E-08 | 9.65E-08 |
| CNAG_02959 | 27170.45 | 5.385603 | 0.093582 | 57.54935 | 0 | 0 |
| CNAG_02958 | 10857.39 | 4.862178 | 0.088041 | 55.22649 | 0 | 0 |
| CNAG_07508 | 866.8292 | 0.561479 | 0.101002 | 5.559097 | 2.71E-08 | 1.70E-07 |
| CNAG_07509 | 2243.583 | 0.180614 | 0.073887 | 2.444485 | 0.014506 | 0.032039 |
| CNAG_07510 | 4104.222 | -0.31294 | 0.069937 | -4.47459 | 7.66E-06 | 3.46E-05 |
| CNAG_02955 | 491.4327 | -0.48282 | 0.108879 | -4.43448 | 9.23E-06 | 4.09E-05 |
| CNAG_02951 | 812.0128 | -0.30011 | 0.098343 | -3.05166 | 0.002276 | 0.006182 |
| CNAG_02950 | 2254.446 | 0.398645 | 0.072511 | 5.497733 | 3.85E-08 | 2.37E-07 |
| CNAG_02948 | 10134.68 | -0.25233 | 0.056048 | -4.50197 | 6.73E-06 | 3.07E-05 |
| CNAG_07511 | 6366.415 | 0.167888 | 0.072404 | 2.318777 | 0.020407 | 0.043075 |
| CNAG_07512 | 613.9858 | 0.862814 | 0.103986 | 8.297413 | 1.06E-16 | 1.28E-15 |
| CNAG_02944 | 9978.511 | 4.921192 | 0.070282 | 70.02076 | 0 | 0 |
| CNAG_02943 | 14274.28 | -0.23427 | 0.072901 | -3.21358 | 0.001311 | 0.003752 |
| CNAG_02941 | 1246.918 | -0.44728 | 0.07912 | -5.65312 | 1.58E-08 | 1.01E-07 |
| CNAG_02940 | 1171.947 | 0.2761 | 0.085184 | 3.24121 | 0.00119 | 0.003437 |

| | | | | | | |
|---|---|---|---|---|---|---|
| CNAG_02939 | 1293.446 | -0.51592 | 0.08667 | -5.95275 | 2.64E-09 | 1.85E-08 |
| CNAG_02938 | 9654.985 | 0.35374 | 0.078299 | 4.517813 | 6.25E-06 | 2.87E-05 |
| CNAG_02936 | 5318.053 | -0.20749 | 0.061143 | -3.39357 | 0.00069 | 0.002113 |
| CNAG_02935 | 2825.85 | 1.860464 | 0.06868 | 27.08885 | 1.33E-161 | 1.87E-159 |
| CNAG_02934 | 157.7101 | 1.33767 | 0.177214 | 7.54832 | 4.41E-14 | 4.62E-13 |
| CNAG_02933 | 700.4543 | -0.71046 | 0.141703 | -5.01375 | 5.34E-07 | 2.85E-06 |
| CNAG_02928 | 58207.12 | -0.1448 | 0.071485 | -2.02564 | 0.042802 | 0.082069 |
| CNAG_02925 | 4782.434 | -0.7956 | 0.089075 | -8.93174 | 4.19E-19 | 5.69E-18 |
| CNAG_02923 | 2794.414 | -0.20479 | 0.078693 | -2.60233 | 0.009259 | 0.021639 |
| CNAG_02922 | 1859.653 | -0.30403 | 0.073079 | -4.16028 | 3.18E-05 | 0.000127 |
| CNAG_02921 | 3636.334 | 0.434502 | 0.068973 | 6.299579 | 2.98E-10 | 2.31E-09 |
| CNAG_02919 | 544.7185 | 0.341441 | 0.109275 | 3.124603 | 0.00178 | 0.004958 |
| CNAG_12238 | 165.8138 | 1.122372 | 0.198622 | 5.650806 | 1.60E-08 | 1.02E-07 |
| CNAG_02918 | 10417.61 | -0.73571 | 0.094133 | -7.81557 | 5.47E-15 | 6.03E-14 |
| CNAG_02910 | 678.6401 | -0.34143 | 0.146522 | -2.3302 | 0.019795 | 0.042021 |
| CNAG_07513 | 1729.324 | 0.335157 | 0.082109 | 4.081868 | 4.47E-05 | 0.000174 |
| CNAG_02904 | 307.6311 | -0.32029 | 0.132479 | -2.41767 | 0.01562 | 0.034148 |
| CNAG_02903 | 1563.707 | 0.217299 | 0.085939 | 2.528524 | 0.011454 | 0.026101 |
| CNAG_02902 | 430.449 | -0.26122 | 0.113782 | -2.29579 | 0.021688 | 0.045479 |
| CNAG_02901 | 2586.876 | -0.35254 | 0.094151 | -3.74445 | 0.000181 | 0.000628 |
| CNAG_02900 | 283.6691 | 1.701873 | 0.165188 | 10.30261 | 6.86E-25 | 1.19E-23 |
| CNAG_12243 | 62.34547 | 0.757637 | 0.316654 | 2.392636 | 0.016728 | 0.036305 |
| CNAG_02898 | 2852.961 | -0.38151 | 0.077387 | -4.92997 | 8.22E-07 | 4.27E-06 |
| CNAG_02897 | 1021.888 | -0.27847 | 0.097906 | -2.84431 | 0.004451 | 0.01127 |
| CNAG_02896 | 4106.511 | -1.25793 | 0.082763 | -15.1991 | 3.58E-52 | 1.27E-50 |
| CNAG_02893 | 270.8289 | 0.717938 | 0.150346 | 4.775239 | 1.79E-06 | 8.90E-06 |
| CNAG_02888 | 430.3248 | 1.236371 | 0.119191 | 10.37305 | 3.29E-25 | 5.77E-24 |
| CNAG_02885 | 4446.753 | -0.35522 | 0.066254 | -5.36153 | 8.25E-08 | 4.89E-07 |
| CNAG_12244 | 61.44315 | -0.69612 | 0.287015 | -2.42536 | 0.015293 | 0.033541 |
| CNAG_02880 | 21901.02 | -0.16842 | 0.0542 | -3.10744 | 0.001887 | 0.005235 |
| CNAG_02879 | 1566.105 | -0.32296 | 0.075318 | -4.28795 | 1.80E-05 | 7.54E-05 |
| CNAG_02878 | 1552.261 | 0.194155 | 0.076827 | 2.527183 | 0.011498 | 0.026171 |
| CNAG_02875 | 1414.369 | -0.21423 | 0.085279 | -2.51213 | 0.012 | 0.02717 |
| CNAG_02871 | 458.9689 | -0.42239 | 0.118709 | -3.55819 | 0.000373 | 0.001214 |
| CNAG_02870 | 2571.796 | -0.63197 | 0.087495 | -7.22289 | 5.09E-13 | 5.00E-12 |
| CNAG_02869 | 227.149 | -0.82948 | 0.153179 | -5.41509 | 6.13E-08 | 3.70E-07 |
| CNAG_12248 | 126.089 | -0.9639 | 0.223313 | -4.31636 | 1.59E-05 | 6.70E-05 |
| CNAG_02868 | 1129.952 | -0.45305 | 0.101405 | -4.46769 | 7.91E-06 | 3.55E-05 |
| CNAG_02867 | 2185.057 | 0.574823 | 0.078439 | 7.328262 | 2.33E-13 | 2.33E-12 |
| CNAG_02866 | 1114.918 | -0.5676 | 0.093575 | -6.06576 | 1.31E-09 | 9.51E-09 |
| CNAG_12249 | 41.66207 | 1.546807 | 0.35403 | 4.369145 | 1.25E-05 | 5.37E-05 |
| CNAG_02865 | 227.5908 | 1.102678 | 0.179118 | 6.156156 | 7.45E-10 | 5.52E-09 |
| CNAG_02864 | 708.8817 | 1.212536 | 0.136215 | 8.90164 | 5.50E-19 | 7.42E-18 |
| CNAG_12250 | 1636.238 | 0.708615 | 0.084702 | 8.365934 | 5.96E-17 | 7.29E-16 |
| CNAG_12252 | 29.39639 | 1.25731 | 0.468779 | 2.682098 | 0.007316 | 0.017495 |
| CNAG_02860 | 9163.356 | -0.14413 | 0.068322 | -2.10963 | 0.03489 | 0.068666 |
| CNAG_02859 | 578.9676 | -0.39692 | 0.117986 | -3.36413 | 0.000768 | 0.002328 |

| | | | | | | |
|---|---|---|---|---|---|---|
| CNAG_02858 | 22206.69 | -0.25408 | 0.055049 | -4.61545 | 3.92E-06 | 1.85E-05 |
| CNAG_02856 | 3695.082 | 0.167519 | 0.062068 | 2.698938 | 0.006956 | 0.016713 |
| CNAG_02854 | 1516.054 | -0.52505 | 0.07603 | -6.90587 | 4.99E-12 | 4.50E-11 |
| CNAG_02852 | 1081.212 | 1.174118 | 0.100468 | 11.68648 | 1.49E-31 | 3.17E-30 |
| CNAG_02850 | 12791.95 | -0.52601 | 0.067678 | -7.77217 | 7.72E-15 | 8.39E-14 |
| CNAG_02848 | 382.2539 | 0.419414 | 0.147474 | 2.843992 | 0.004455 | 0.011274 |
| CNAG_02847 | 621.825 | -0.20773 | 0.105387 | -1.97108 | 0.048715 | 0.091571 |
| CNAG_02846 | 410.3426 | 0.384145 | 0.137008 | 2.803807 | 0.00505 | 0.012619 |
| CNAG_12259 | 67.19973 | 1.012101 | 0.349403 | 2.896658 | 0.003772 | 0.00972 |
| CNAG_02840 | 735.2634 | 0.390327 | 0.115908 | 3.367549 | 0.000758 | 0.002303 |
| CNAG_02839 | 1388.165 | 5.30373 | 0.131708 | 40.26898 | 0 | 0 |
| CNAG_07519 | 2867.946 | 2.437587 | 0.073116 | 33.33865 | 1.06E-243 | 3.20E-241 |
| CNAG_02834 | 611.3254 | -0.25644 | 0.099903 | -2.56689 | 0.010262 | 0.023614 |
| CNAG_02833 | 2884.369 | 0.443286 | 0.088606 | 5.002883 | 5.65E-07 | 3.00E-06 |
| CNAG_07520 | 216.2484 | 0.758763 | 0.160052 | 4.740743 | 2.13E-06 | 1.05E-05 |
| CNAG_02830 | 5172.701 | -0.94767 | 0.085052 | -11.1422 | 7.81E-29 | 1.54E-27 |
| CNAG_02828 | 148.5009 | 0.502919 | 0.192911 | 2.606995 | 0.009134 | 0.021384 |
| CNAG_02827 | 1190.376 | 0.22531 | 0.10149 | 2.220015 | 0.026418 | 0.054114 |
| CNAG_02826 | 914.243 | 0.584532 | 0.094121 | 6.210403 | 5.28E-10 | 3.97E-09 |
| CNAG_02825 | 5664.456 | -0.14083 | 0.064525 | -2.18263 | 0.029063 | 0.058748 |
| CNAG_07521 | 211.4469 | -0.627 | 0.152835 | -4.10244 | 4.09E-05 | 0.000161 |
| CNAG_12262 | 9.069974 | -1.89385 | 0.783671 | -2.41663 | 0.015665 | 0.034217 |
| CNAG_02822 | 4769.818 | -0.5493 | 0.065941 | -8.33017 | 8.07E-17 | 9.79E-16 |
| CNAG_02821 | 394.2103 | -0.38297 | 0.126056 | -3.03809 | 0.002381 | 0.00644 |
| CNAG_02819 | 815.8178 | 0.41059 | 0.098154 | 4.183119 | 2.88E-05 | 0.000116 |
| CNAG_02818 | 2108.433 | 0.814328 | 0.075789 | 10.74473 | 6.27E-27 | 1.16E-25 |
| CNAG_02817 | 3964.763 | -0.19868 | 0.063782 | -3.11502 | 0.001839 | 0.005115 |
| CNAG_02815 | 4220.686 | 0.783371 | 0.287198 | 2.72763 | 0.006379 | 0.015487 |
| CNAG_02814 | 7153.239 | -0.6262 | 0.079389 | -7.88771 | 3.08E-15 | 3.44E-14 |
| CNAG_02812 | 3751.418 | -0.24494 | 0.064432 | -3.80148 | 0.000144 | 0.00051 |
| CNAG_07522 | 1664.705 | -0.46794 | 0.106831 | -4.38015 | 1.19E-05 | 5.13E-05 |
| CNAG_02801 | 7856.096 | -0.21305 | 0.068944 | -3.09021 | 0.002 | 0.005515 |
| CNAG_02794 | 3112.826 | -0.38526 | 0.097202 | -3.96343 | 7.39E-05 | 0.000278 |
| CNAG_02790 | 2267.268 | 0.620686 | 0.068687 | 9.036462 | 1.62E-19 | 2.25E-18 |
| CNAG_02788 | 1383.201 | 0.954211 | 0.087567 | 10.8969 | 1.19E-27 | 2.27E-26 |
| CNAG_02787 | 1067.872 | -0.56106 | 0.084416 | -6.64645 | 3.00E-11 | 2.54E-10 |
| CNAG_02786 | 1589.147 | -0.19891 | 0.075345 | -2.63999 | 0.008291 | 0.019559 |
| CNAG_02783 | 1013.958 | 0.274556 | 0.087497 | 3.137901 | 0.001702 | 0.004758 |
| CNAG_02782 | 1339.408 | -0.35445 | 0.078847 | -4.49533 | 6.95E-06 | 3.16E-05 |
| CNAG_02781 | 69.68701 | 0.810764 | 0.258429 | 3.137283 | 0.001705 | 0.004766 |
| CNAG_02780 | 2408.792 | -0.31259 | 0.070727 | -4.41962 | 9.89E-06 | 4.36E-05 |
| CNAG_02779 | 2464.811 | -0.17936 | 0.068207 | -2.62965 | 0.008547 | 0.020108 |
| CNAG_02777 | 9556.577 | 2.050762 | 0.103911 | 19.73575 | 1.06E-86 | 6.55E-85 |
| CNAG_12271 | 120.161 | 3.805009 | 0.279688 | 13.60448 | 3.77E-42 | 1.07E-40 |
| CNAG_07523 | 454.6127 | 0.246108 | 0.114627 | 2.147032 | 0.031791 | 0.063488 |
| CNAG_02776 | 359.6756 | 0.366785 | 0.136951 | 2.678214 | 0.007402 | 0.017684 |
| CNAG_02775 | 4863.762 | 1.704045 | 0.080324 | 21.21475 | 6.98E-100 | 5.11E-98 |

| | | | | | | |
|---|---|---|---|---|---|---|
| CNAG_02773 | 3847.16 | 0.178322 | 0.061685 | 2.890847 | 0.003842 | 0.009882 |
| CNAG_02771 | 1342.11 | 0.480089 | 0.086376 | 5.558146 | 2.73E-08 | 1.70E-07 |
| CNAG_02770 | 804.912 | -0.34088 | 0.097871 | -3.48298 | 0.000496 | 0.001562 |
| CNAG_02769 | 1655.951 | -0.23251 | 0.075254 | -3.08964 | 0.002004 | 0.005523 |
| CNAG_02768 | 89.65675 | -0.68108 | 0.243856 | -2.79295 | 0.005223 | 0.013006 |
| CNAG_02763 | 4325.378 | -0.2027 | 0.061542 | -3.29361 | 0.000989 | 0.002916 |
| CNAG_02762 | 11724.94 | -0.24308 | 0.081507 | -2.98234 | 0.002861 | 0.007591 |
| CNAG_12272 | 25.83888 | 1.171141 | 0.431757 | 2.7125 | 0.006678 | 0.01613 |
| CNAG_02759 | 220.8623 | 0.414368 | 0.147563 | 2.808088 | 0.004984 | 0.012464 |
| CNAG_02758 | 641.1155 | 1.754309 | 0.108475 | 16.17245 | 7.89E-59 | 3.26E-57 |
| CNAG_02753 | 3805.318 | -0.5624 | 0.064767 | -8.68343 | 3.84E-18 | 4.95E-17 |
| CNAG_02752 | 3490.059 | 1.045955 | 0.078813 | 13.27133 | 3.40E-40 | 9.32E-39 |
| CNAG_02750 | 354.8119 | 0.347336 | 0.135857 | 2.556634 | 0.010569 | 0.024288 |
| CNAG_02745 | 2825.472 | 0.140686 | 0.065438 | 2.149909 | 0.031562 | 0.063094 |
| CNAG_02744 | 911.9062 | 0.368674 | 0.091565 | 4.02638 | 5.66E-05 | 0.000217 |
| CNAG_02742 | 1419.123 | -0.23993 | 0.083809 | -2.86278 | 0.004199 | 0.0107 |
| CNAG_02739 | 1426.898 | -0.51551 | 0.085681 | -6.01664 | 1.78E-09 | 1.27E-08 |
| CNAG_02738 | 2325.382 | 0.165257 | 0.073868 | 2.237195 | 0.025274 | 0.05198 |
| CNAG_02736 | 6937.413 | -0.1255 | 0.05952 | -2.10847 | 0.034991 | 0.068846 |
| CNAG_02735 | 169.4656 | 1.160996 | 0.23253 | 4.992888 | 5.95E-07 | 3.15E-06 |
| CNAG_02734 | 19.07818 | 2.040913 | 0.581477 | 3.509878 | 0.000448 | 0.001428 |
| CNAG_02733 | 83.25345 | 0.583315 | 0.252957 | 2.305984 | 0.021112 | 0.044328 |
| CNAG_02730 | 917.0801 | -0.34466 | 0.103625 | -3.32604 | 0.000881 | 0.002627 |
| CNAG_02726 | 4607.304 | -0.16582 | 0.074068 | -2.23881 | 0.025168 | 0.051803 |
| CNAG_02725 | 3767.889 | 0.132007 | 0.06169 | 2.139835 | 0.032368 | 0.064451 |
| CNAG_12274 | 149.4084 | 0.787107 | 0.200125 | 3.933086 | 8.39E-05 | 0.000312 |
| CNAG_02723 | 1393.03 | -0.35068 | 0.078713 | -4.45513 | 8.38E-06 | 3.75E-05 |
| CNAG_12277 | 48.16274 | 2.381169 | 0.350259 | 6.798317 | 1.06E-11 | 9.34E-11 |
| CNAG_02721 | 2228.211 | -0.23893 | 0.077008 | -3.10265 | 0.001918 | 0.005315 |
| CNAG_02718 | 1366.645 | -0.29002 | 0.086008 | -3.37201 | 0.000746 | 0.002273 |
| CNAG_02717 | 1276.726 | -0.35686 | 0.084476 | -4.22441 | 2.40E-05 | 9.76E-05 |
| CNAG_02716 | 827.7525 | 0.22444 | 0.091905 | 2.442082 | 0.014603 | 0.032236 |
| CNAG_02715 | 1019.052 | 0.226243 | 0.090477 | 2.50056 | 0.0124 | 0.027957 |
| CNAG_02714 | 34971.21 | -0.15397 | 0.070675 | -2.17862 | 0.02936 | 0.059259 |
| CNAG_02713 | 748.4753 | -0.40544 | 0.113338 | -3.57727 | 0.000347 | 0.001136 |
| CNAG_02712 | 2205.642 | -0.7847 | 0.082798 | -9.47728 | 2.61E-21 | 3.96E-20 |
| CNAG_02711 | 46.1168 | -1.25396 | 0.317104 | -3.95442 | 7.67E-05 | 0.000288 |
| CNAG_02710 | 8732.624 | -0.17604 | 0.061607 | -2.85739 | 0.004271 | 0.010877 |
| CNAG_02708 | 525.4763 | -0.48217 | 0.125024 | -3.85661 | 0.000115 | 0.000416 |
| CNAG_02705 | 152.7764 | -1.60172 | 0.225954 | -7.08871 | 1.35E-12 | 1.28E-11 |
| CNAG_02704 | 1752.818 | -0.3083 | 0.092502 | -3.33293 | 0.000859 | 0.002575 |
| CNAG_02703 | 2637.826 | 0.360315 | 0.081672 | 4.411714 | 1.03E-05 | 4.50E-05 |
| CNAG_02702 | 2049.399 | -0.33586 | 0.08561 | -3.92311 | 8.74E-05 | 0.000325 |
| CNAG_02701 | 374.5981 | 0.608484 | 0.208586 | 2.917188 | 0.003532 | 0.00916 |
| CNAG_12282 | 10.99249 | 1.711717 | 0.678865 | 2.521441 | 0.011688 | 0.02655 |
| CNAG_12285 | 285.8993 | 1.302626 | 0.146594 | 8.88592 | 6.34E-19 | 8.50E-18 |
| CNAG_02697 | 1623.376 | 1.60856 | 0.087328 | 18.41971 | 9.13E-76 | 4.85E-74 |

| | | | | | | |
|---|---|---|---|---|---|---|
| CNAG_02695 | 1704.055 | -0.18593 | 0.075892 | -2.44999 | 0.014286 | 0.031648 |
| CNAG_02694 | 60.00179 | 0.550812 | 0.275707 | 1.997818 | 0.045736 | 0.086938 |
| CNAG_02692 | 6111.295 | 1.427691 | 0.094182 | 15.1589 | 6.62E-52 | 2.32E-50 |
| CNAG_02690 | 1142.358 | 0.694509 | 0.107509 | 6.459998 | 1.05E-10 | 8.46E-10 |
| CNAG_02689 | 453.6595 | -0.31144 | 0.110367 | -2.82188 | 0.004774 | 0.011977 |
| CNAG_02687 | 3026.148 | 0.468014 | 0.066398 | 7.048587 | 1.81E-12 | 1.69E-11 |
| CNAG_02686 | 3772.054 | -0.26635 | 0.066817 | -3.9863 | 6.71E-05 | 0.000254 |
| CNAG_02684 | 1300.4 | 0.594786 | 0.088601 | 6.713082 | 1.91E-11 | 1.63E-10 |
| CNAG_02683 | 1731.156 | -0.68079 | 0.090241 | -7.54412 | 4.55E-14 | 4.77E-13 |
| CNAG_02675 | 3944.987 | -0.36407 | 0.072611 | -5.01401 | 5.33E-07 | 2.85E-06 |
| CNAG_02674 | 549.876 | 0.839302 | 0.138214 | 6.072481 | 1.26E-09 | 9.16E-09 |
| CNAG_02673 | 4521.968 | -0.33711 | 0.105529 | -3.19444 | 0.001401 | 0.003986 |
| CNAG_02672 | 4515.097 | 0.448045 | 0.068691 | 6.522585 | 6.91E-11 | 5.66E-10 |
| CNAG_02671 | 7126.599 | -0.18252 | 0.067377 | -2.70894 | 0.00675 | 0.01628 |
| CNAG_02670 | 1107.84 | -0.28279 | 0.088625 | -3.19092 | 0.001418 | 0.004028 |
| CNAG_02666 | 734.3896 | 0.372013 | 0.092801 | 4.008707 | 6.11E-05 | 0.000233 |
| CNAG_02665 | 202.095 | 1.15127 | 0.183836 | 6.26249 | 3.79E-10 | 2.89E-09 |
| CNAG_02664 | 3681.5 | -0.27488 | 0.071002 | -3.87151 | 0.000108 | 0.000393 |
| CNAG_02663 | 1188.403 | 0.579051 | 0.096801 | 5.981881 | 2.21E-09 | 1.56E-08 |
| CNAG_12292 | 67.754 | 1.847354 | 0.307487 | 6.007904 | 1.88E-09 | 1.34E-08 |
| CNAG_12295 | 79.49224 | 1.318919 | 0.277405 | 4.754484 | 1.99E-06 | 9.81E-06 |
| CNAG_02658 | 1845.084 | -0.15916 | 0.080589 | -1.97496 | 0.048273 | 0.090959 |
| CNAG_02657 | 8262.559 | -0.22984 | 0.063887 | -3.59764 | 0.000321 | 0.001058 |
| CNAG_02656 | 3018.164 | -0.17377 | 0.073847 | -2.35311 | 0.018617 | 0.039821 |
| CNAG_02655 | 1649.271 | -0.50211 | 0.081408 | -6.16783 | 6.92E-10 | 5.16E-09 |
| CNAG_02654 | 1604.475 | 0.418671 | 0.086549 | 4.837394 | 1.32E-06 | 6.62E-06 |
| CNAG_07529 | 1659.513 | -0.17519 | 0.082112 | -2.13361 | 0.032875 | 0.0653 |
| CNAG_07530 | 1943.043 | -0.19626 | 0.095364 | -2.05798 | 0.039592 | 0.076748 |
| CNAG_07531 | 728.1733 | 0.26156 | 0.094586 | 2.765327 | 0.005687 | 0.01401 |
| CNAG_07532 | 968.5041 | -0.25684 | 0.11754 | -2.18514 | 0.028879 | 0.058434 |
| CNAG_07533 | 120.6888 | 0.430129 | 0.218132 | 1.971872 | 0.048624 | 0.091464 |
| CNAG_07538 | 1575.935 | 1.905783 | 0.122539 | 15.55246 | 1.53E-54 | 5.63E-53 |
| CNAG_07539 | 2348.846 | 0.249471 | 0.065283 | 3.821355 | 0.000133 | 0.000473 |
| CNAG_07540 | 334.5358 | -0.37538 | 0.151983 | -2.46988 | 0.013516 | 0.03018 |
| CNAG_07541 | 1900.062 | 0.91284 | 0.075825 | 12.03884 | 2.22E-33 | 4.97E-32 |
| CNAG_07542 | 221.95 | 0.890232 | 0.16172 | 5.504789 | 3.70E-08 | 2.28E-07 |
| CNAG_12299 | 21.2287 | 3.975148 | 0.689994 | 5.76113 | 8.36E-09 | 5.55E-08 |
| CNAG_07544 | 120.402 | 1.145788 | 0.21947 | 5.220705 | 1.78E-07 | 1.01E-06 |
| CNAG_07545 | 1081.98 | 0.194521 | 0.085524 | 2.274453 | 0.022939 | 0.047782 |
| CNAG_07547 | 1108.479 | 0.522035 | 0.110531 | 4.722969 | 2.32E-06 | 1.13E-05 |
| CNAG_07548 | 1165.382 | 0.991488 | 0.083923 | 11.81427 | 3.29E-32 | 7.10E-31 |
| CNAG_07549 | 423.9964 | 0.838444 | 0.133487 | 6.281101 | 3.36E-10 | 2.58E-09 |
| CNAG_07550 | 641.4152 | 0.722107 | 0.099008 | 7.293382 | 3.02E-13 | 3.01E-12 |
| CNAG_07552 | 1489.991 | 0.704496 | 0.081967 | 8.594876 | 8.34E-18 | 1.05E-16 |
| CNAG_07553 | 1048.33 | 0.392637 | 0.085303 | 4.602876 | 4.17E-06 | 1.96E-05 |
| CNAG_07554 | 4446.986 | -0.1684 | 0.066162 | -2.5453 | 0.010918 | 0.025027 |
| CNAG_07108 | 2424.731 | -0.17339 | 0.077441 | -2.23894 | 0.02516 | 0.051798 |

| | | | | | | |
|---|---|---|---|---|---|---|
| CNAG_07555 | 396.8036 | 0.933767 | 0.134105 | 6.962965 | 3.33E-12 | 3.03E-11 |
| CNAG_07556 | 900.8896 | -0.35854 | 0.086666 | -4.13703 | 3.52E-05 | 0.00014 |
| CNAG_07557 | 816.4633 | -0.91075 | 0.102497 | -8.88568 | 6.35E-19 | 8.51E-18 |
| CNAG_07558 | 17881.99 | 0.503108 | 0.069448 | 7.24443 | 4.34E-13 | 4.29E-12 |
| CNAG_07561 | 47826.76 | -0.13516 | 0.060218 | -2.24446 | 0.024803 | 0.051188 |
| CNAG_07565 | 541.9002 | -0.44758 | 0.102693 | -4.35845 | 1.31E-05 | 5.62E-05 |
| CNAG_07566 | 487.6013 | 0.281268 | 0.110464 | 2.546244 | 0.010889 | 0.024966 |
| CNAG_07571 | 1798.754 | -0.23146 | 0.087123 | -2.65669 | 0.007891 | 0.018705 |
| CNAG_07572 | 1624.498 | 0.402372 | 0.086288 | 4.663115 | 3.11E-06 | 1.49E-05 |
| CNAG_02609 | 1607.053 | 0.174366 | 0.076743 | 2.272067 | 0.023082 | 0.048032 |
| CNAG_07573 | 792.9257 | -0.37308 | 0.089153 | -4.1847 | 2.86E-05 | 0.000115 |
| CNAG_02605 | 844.2059 | 0.233637 | 0.094882 | 2.462392 | 0.013801 | 0.030725 |
| CNAG_02604 | 90.67147 | 0.761912 | 0.228231 | 3.338346 | 0.000843 | 0.002528 |
| CNAG_12302 | 200.5664 | 2.372431 | 0.18526 | 12.80597 | 1.52E-37 | 3.86E-36 |
| CNAG_02603 | 215.959 | 2.196834 | 0.173346 | 12.67309 | 8.34E-37 | 2.09E-35 |
| CNAG_02602 | 193.1596 | 2.005705 | 0.174322 | 11.50573 | 1.23E-30 | 2.59E-29 |
| CNAG_02600 | 417.9334 | 0.31704 | 0.115074 | 2.755086 | 0.005868 | 0.014374 |
| CNAG_02599 | 38.19121 | -0.78195 | 0.354284 | -2.20714 | 0.027304 | 0.055705 |
| CNAG_02598 | 173.2087 | 0.587566 | 0.198694 | 2.957138 | 0.003105 | 0.008168 |
| CNAG_12303 | 29.92101 | 0.897252 | 0.426391 | 2.104295 | 0.035353 | 0.069458 |
| CNAG_02595 | 412.5918 | -0.33249 | 0.142503 | -2.33323 | 0.019636 | 0.041737 |
| CNAG_02594 | 353.0005 | -0.33072 | 0.131262 | -2.51952 | 0.011751 | 0.02668 |
| CNAG_02593 | 719.084 | -0.27541 | 0.114823 | -2.39852 | 0.016461 | 0.035804 |
| CNAG_02591 | 143.2009 | -0.45852 | 0.230058 | -1.99308 | 0.046252 | 0.087713 |
| CNAG_02590 | 449.1127 | 1.300269 | 0.11469 | 11.33723 | 8.58E-30 | 1.76E-28 |
| CNAG_02589 | 596.6411 | 0.382002 | 0.101675 | 3.757084 | 0.000172 | 0.000601 |
| CNAG_02587 | 2085.141 | 1.497356 | 0.071469 | 20.95123 | 1.83E-97 | 1.28E-95 |
| CNAG_02586 | 137.8576 | 2.040228 | 0.208811 | 9.770678 | 1.50E-22 | 2.38E-21 |
| CNAG_02585 | 3062.216 | 0.540184 | 0.079289 | 6.812889 | 9.57E-12 | 8.51E-11 |
| CNAG_12306 | 33.71509 | -1.15278 | 0.399871 | -2.88287 | 0.003941 | 0.010095 |
| CNAG_02581 | 3553.66 | 1.455969 | 0.06795 | 21.4271 | 7.47E-102 | 5.57E-100 |
| CNAG_02580 | 3772.633 | 1.578499 | 0.062006 | 25.45723 | 5.87E-143 | 7.12E-141 |
| CNAG_02579 | 1574.616 | 0.463304 | 0.103324 | 4.483984 | 7.33E-06 | 3.32E-05 |
| CNAG_02577 | 2787.046 | -0.4297 | 0.085435 | -5.02956 | 4.92E-07 | 2.64E-06 |
| CNAG_02575 | 2703.577 | -0.23901 | 0.066342 | -3.60267 | 0.000315 | 0.00104 |
| CNAG_02574 | 1261.597 | 0.165112 | 0.079481 | 2.077393 | 0.037765 | 0.073717 |
| CNAG_12311 | 42.746 | 0.695745 | 0.336255 | 2.0691 | 0.038537 | 0.075024 |
| CNAG_07574 | 99.36881 | 0.634227 | 0.229811 | 2.759771 | 0.005784 | 0.014212 |
| CNAG_12313 | 153.319 | 0.562711 | 0.242789 | 2.317691 | 0.020466 | 0.043174 |
| CNAG_12314 | 99.60187 | 2.827129 | 0.286713 | 9.860469 | 6.18E-23 | 9.96E-22 |
| CNAG_02569 | 749.5491 | -0.2655 | 0.095792 | -2.77167 | 0.005577 | 0.013779 |
| CNAG_02568 | 3326.95 | -0.28999 | 0.090977 | -3.18748 | 0.001435 | 0.004069 |
| CNAG_02567 | 618.3446 | -0.32114 | 0.120514 | -2.66474 | 0.007705 | 0.018333 |
| CNAG_02566 | 2084.2 | 0.18245 | 0.088242 | 2.067625 | 0.038675 | 0.075254 |
| CNAG_02565 | 7937.487 | 1.172001 | 0.060067 | 19.51158 | 8.75E-85 | 5.16E-83 |
| CNAG_02564 | 1712.727 | 1.00186 | 0.078196 | 12.81221 | 1.40E-37 | 3.57E-36 |
| CNAG_02563 | 1262.083 | 0.178936 | 0.079449 | 2.252204 | 0.024309 | 0.050276 |

| | | | | | | |
|---|---|---|---|---|---|---|
| CNAG_07972 | 202.3628 | 0.544642 | 0.159415 | 3.416505 | 0.000634 | 0.001957 |
| CNAG_06879 | 632.7217 | -0.35626 | 0.097791 | -3.64311 | 0.000269 | 0.0009 |
| CNAG_06880 | 907.5488 | -0.2053 | 0.090281 | -2.27404 | 0.022964 | 0.047797 |
| CNAG_06881 | 1599.383 | -0.19336 | 0.082306 | -2.34921 | 0.018813 | 0.040203 |
| CNAG_07576 | 561.1288 | 0.397733 | 0.115315 | 3.44911 | 0.000562 | 0.001756 |
| CNAG_07579 | 596.3554 | -0.26402 | 0.107274 | -2.46117 | 0.013849 | 0.030805 |
| CNAG_07580 | 1288.691 | -0.25154 | 0.099545 | -2.5269 | 0.011507 | 0.026185 |
| CNAG_06887 | 464.6232 | 0.44928 | 0.111769 | 4.019708 | 5.83E-05 | 0.000223 |
| CNAG_12318 | 34.25002 | 0.911617 | 0.368388 | 2.474613 | 0.013338 | 0.029815 |
| CNAG_06891 | 1054.973 | -0.52252 | 0.099694 | -5.24119 | 1.60E-07 | 9.10E-07 |
| CNAG_06892 | 1228.268 | -0.27523 | 0.091321 | -3.01388 | 0.002579 | 0.006917 |
| CNAG_06893 | 1121.43 | -0.54936 | 0.089529 | -6.13619 | 8.45E-10 | 6.21E-09 |
| CNAG_06896 | 1807.901 | -0.19391 | 0.076762 | -2.52609 | 0.011534 | 0.026231 |
| CNAG_06897 | 1049.748 | -0.69607 | 0.084123 | -8.2744 | 1.29E-16 | 1.54E-15 |
| CNAG_12319 | 35.97954 | 1.530299 | 0.360516 | 4.244747 | 2.19E-05 | 8.99E-05 |
| CNAG_12322 | 17.75523 | 2.067403 | 0.562419 | 3.675913 | 0.000237 | 0.000803 |
| CNAG_12323 | 97.24229 | 1.395041 | 0.247272 | 5.641717 | 1.68E-08 | 1.08E-07 |
| CNAG_06898 | 2498.388 | 0.473068 | 0.069533 | 6.803514 | 1.02E-11 | 9.04E-11 |
| CNAG_06899 | 6865.633 | 0.271353 | 0.07228 | 3.754207 | 0.000174 | 0.000607 |
| CNAG_06900 | 24646.99 | 0.326523 | 0.067573 | 4.832138 | 1.35E-06 | 6.78E-06 |
| CNAG_06901 | 651.9261 | 1.346073 | 0.113676 | 11.84129 | 2.39E-32 | 5.17E-31 |
| CNAG_06902 | 1108.302 | -0.44501 | 0.086733 | -5.13078 | 2.89E-07 | 1.60E-06 |
| CNAG_06903 | 913.9355 | -0.27892 | 0.101084 | -2.75926 | 0.005793 | 0.014226 |
| CNAG_06905 | 1346.312 | -0.82716 | 0.089084 | -9.28508 | 1.62E-20 | 2.35E-19 |
| CNAG_06906 | 11201.92 | -0.30235 | 0.067624 | -4.47106 | 7.78E-06 | 3.50E-05 |
| CNAG_06908 | 12870.16 | 0.277174 | 0.062771 | 4.415634 | 1.01E-05 | 4.43E-05 |
| CNAG_06909 | 639.7245 | 0.266439 | 0.099598 | 2.675154 | 0.007469 | 0.01783 |
| CNAG_12328 | 157.1227 | 0.77452 | 0.182503 | 4.243868 | 2.20E-05 | 9.02E-05 |
| CNAG_06910 | 503.3857 | 2.528741 | 0.122742 | 20.60206 | 2.63E-94 | 1.78E-92 |
| CNAG_06911 | 8.749487 | 3.702141 | 0.974172 | 3.800295 | 0.000145 | 0.000512 |
| CNAG_06912 | 358.764 | 0.769757 | 0.137957 | 5.579695 | 2.41E-08 | 1.51E-07 |
| CNAG_06913 | 678.2259 | 1.716682 | 0.112697 | 15.23277 | 2.14E-52 | 7.67E-51 |
| CNAG_06914 | 1579.045 | -0.31531 | 0.073398 | -4.29593 | 1.74E-05 | 7.30E-05 |
| CNAG_06916 | 1462.174 | 0.448777 | 0.080948 | 5.544052 | 2.96E-08 | 1.84E-07 |
| CNAG_12331 | 74.34948 | 0.952478 | 0.261655 | 3.640205 | 0.000272 | 0.000909 |
| CNAG_06920 | 4145.38 | -0.23002 | 0.090852 | -2.53179 | 0.011348 | 0.025903 |
| CNAG_06922 | 1377.784 | -0.96084 | 0.270933 | -3.54642 | 0.000391 | 0.001262 |
| CNAG_06923 | 14457.53 | 2.099632 | 0.075563 | 27.78663 | 6.29E-170 | 9.47E-168 |
| CNAG_06924 | 1106.951 | -0.30592 | 0.128484 | -2.38101 | 0.017265 | 0.037268 |
| CNAG_06925 | 923.2477 | -0.40717 | 0.09177 | -4.43688 | 9.13E-06 | 4.05E-05 |
| CNAG_06929 | 1640.382 | -0.66822 | 0.075085 | -8.8995 | 5.61E-19 | 7.55E-18 |
| CNAG_06930 | 1942.762 | -0.16123 | 0.073099 | -2.2056 | 0.027412 | 0.055869 |
| CNAG_06931 | 149.0415 | -1.15425 | 0.182557 | -6.32264 | 2.57E-10 | 2.00E-09 |
| CNAG_06932 | 212.2935 | 0.871521 | 0.17595 | 4.95324 | 7.30E-07 | 3.82E-06 |
| CNAG_07583 | 751.9705 | 0.366884 | 0.101472 | 3.615617 | 0.0003 | 0.000992 |
| CNAG_06934 | 349.5647 | -0.36148 | 0.135361 | -2.67051 | 0.007574 | 0.018047 |
| CNAG_12333 | 97.0693 | 0.570861 | 0.236044 | 2.418451 | 0.015587 | 0.034102 |

| | | | | | | |
|---|---|---|---|---|---|---|
| CNAG_06935 | 358.2044 | 1.541798 | 0.195488 | 7.886922 | 3.10E-15 | 3.45E-14 |
| CNAG_07793 | 339.885 | 0.412586 | 0.14511 | 2.843255 | 0.004466 | 0.01129 |
| CNAG_04932 | 489.9836 | -0.39425 | 0.107721 | -3.65991 | 0.000252 | 0.00085 |
| CNAG_04933 | 135.9218 | -0.46926 | 0.20322 | -2.30911 | 0.020937 | 0.043997 |
| CNAG_04936 | 2845.58 | -0.31063 | 0.090385 | -3.43679 | 0.000589 | 0.001832 |
| CNAG_04937 | 382.3149 | -0.30447 | 0.124731 | -2.441 | 0.014647 | 0.032315 |
| CNAG_07796 | 1534.995 | 0.963685 | 0.090387 | 10.6618 | 1.54E-26 | 2.77E-25 |
| CNAG_04943 | 1513.661 | 0.574534 | 0.084458 | 6.80262 | 1.03E-11 | 9.09E-11 |
| CNAG_04945 | 248.8406 | -0.46418 | 0.168687 | -2.75175 | 0.005928 | 0.014508 |
| CNAG_04948 | 3475.145 | -0.2436 | 0.06375 | -3.82119 | 0.000133 | 0.000473 |
| CNAG_04949 | 1178.045 | -0.6393 | 0.107655 | -5.93843 | 2.88E-09 | 2.01E-08 |
| CNAG_04950 | 1880.662 | -0.44589 | 0.085313 | -5.22655 | 1.73E-07 | 9.82E-07 |
| CNAG_04951 | 7956.066 | 1.072241 | 0.062626 | 17.12134 | 1.03E-65 | 4.86E-64 |
| CNAG_04952 | 2272.483 | 0.171424 | 0.069151 | 2.478989 | 0.013176 | 0.029518 |
| CNAG_12336 | 110.9508 | -0.40836 | 0.207039 | -1.97239 | 0.048565 | 0.091395 |
| CNAG_04953 | 8751.313 | -0.51448 | 0.067557 | -7.6155 | 2.63E-14 | 2.79E-13 |
| CNAG_04954 | 2758.315 | 0.204269 | 0.087213 | 2.342202 | 0.01917 | 0.040843 |
| CNAG_04955 | 745.0789 | -0.37713 | 0.128639 | -2.9317 | 0.003371 | 0.008791 |
| CNAG_04959 | 992.3606 | -0.23633 | 0.100638 | -2.3483 | 0.018859 | 0.040286 |
| CNAG_04960 | 528.0498 | 2.619765 | 0.169112 | 15.49127 | 3.97E-54 | 1.45E-52 |
| CNAG_04962 | 4259.113 | -0.18985 | 0.07428 | -2.55589 | 0.010592 | 0.024333 |
| CNAG_12339 | 147.228 | 0.425416 | 0.206885 | 2.056295 | 0.039754 | 0.077025 |
| CNAG_04963 | 5423.409 | 0.555034 | 0.082344 | 6.740435 | 1.58E-11 | 1.37E-10 |
| CNAG_07797 | 3647.622 | 1.9583 | 0.06424 | 30.48413 | 4.23E-204 | 8.82E-202 |
| CNAG_07798 | 424.7204 | 3.909511 | 0.156448 | 24.98921 | 8.01E-138 | 8.80E-136 |
| CNAG_04967 | 393.7943 | 0.586788 | 0.11819 | 4.964793 | 6.88E-07 | 3.61E-06 |
| CNAG_04968 | 732.7209 | 0.968993 | 0.094613 | 10.2416 | 1.29E-24 | 2.22E-23 |
| CNAG_04969 | 12984.26 | -0.22961 | 0.060808 | -3.77595 | 0.000159 | 0.000562 |
| CNAG_07800 | 879.907 | -0.2005 | 0.091771 | -2.18477 | 0.028906 | 0.058473 |
| CNAG_04973 | 2499.694 | 0.293119 | 0.074437 | 3.937787 | 8.22E-05 | 0.000307 |
| CNAG_12342 | 41.54759 | 1.837355 | 0.352004 | 5.219696 | 1.79E-07 | 1.02E-06 |
| CNAG_12345 | 81.89419 | -0.57431 | 0.244912 | -2.34497 | 0.019029 | 0.040584 |
| CNAG_12346 | 266.5849 | 1.016817 | 0.195087 | 5.212121 | 1.87E-07 | 1.06E-06 |
| CNAG_04981 | 5124.736 | 1.360711 | 0.216356 | 6.289235 | 3.19E-10 | 2.46E-09 |
| CNAG_04982 | 280.4606 | 1.48055 | 0.14788 | 10.01184 | 1.35E-23 | 2.24E-22 |
| CNAG_04983 | 1359.899 | 0.452238 | 0.075167 | 6.016442 | 1.78E-09 | 1.27E-08 |
| CNAG_04985 | 15040.49 | -0.32052 | 0.081623 | -3.92681 | 8.61E-05 | 0.00032 |
| CNAG_12348 | 64.20114 | 1.070315 | 0.295001 | 3.628178 | 0.000285 | 0.00095 |
| CNAG_12349 | 48.59229 | 2.647918 | 0.348027 | 7.608366 | 2.78E-14 | 2.94E-13 |
| CNAG_04988 | 201.5274 | 4.506704 | 0.248518 | 18.13431 | 1.71E-73 | 8.84E-72 |
| CNAG_12350 | 213.6776 | 1.783522 | 0.195966 | 9.101175 | 8.94E-20 | 1.25E-18 |
| CNAG_04989 | 725.0531 | -0.41136 | 0.102397 | -4.01734 | 5.89E-05 | 0.000225 |
| CNAG_04991 | 2065.11 | -0.28318 | 0.084529 | -3.35008 | 0.000808 | 0.002433 |
| CNAG_12354 | 24.22724 | 1.45634 | 0.434496 | 3.351795 | 0.000803 | 0.002421 |
| CNAG_04993 | 47.31311 | -1.00094 | 0.324114 | -3.08825 | 0.002013 | 0.005542 |
| CNAG_04994 | 1019.132 | -0.35779 | 0.107489 | -3.32866 | 0.000873 | 0.002607 |
| CNAG_07801 | 2105.491 | -0.47128 | 0.069705 | -6.76102 | 1.37E-11 | 1.19E-10 |

| | | | | | | |
|---|---|---|---|---|---|---|
| CNAG_07802 | 888.0694 | 4.756849 | 0.323808 | 14.69032 | 7.44E-49 | 2.45E-47 |
| CNAG_04997 | 1289.079 | 0.391217 | 0.08451 | 4.629243 | 3.67E-06 | 1.74E-05 |
| CNAG_04999 | 278.7704 | -0.44261 | 0.138802 | -3.18876 | 0.001429 | 0.004054 |
| CNAG_05002 | 19.90533 | 1.034946 | 0.493398 | 2.097587 | 0.035942 | 0.070479 |
| CNAG_05003 | 955.2489 | 1.249398 | 0.090779 | 13.76309 | 4.25E-43 | 1.28E-41 |
| CNAG_05004 | 1405.703 | 0.371134 | 0.076786 | 4.833329 | 1.34E-06 | 6.74E-06 |
| CNAG_05005 | 550.0823 | 0.426923 | 0.111946 | 3.813643 | 0.000137 | 0.000487 |
| CNAG_05011 | 803.5507 | -0.20727 | 0.093244 | -2.22285 | 0.026226 | 0.053774 |
| CNAG_05012 | 671.0477 | 0.341175 | 0.096738 | 3.526785 | 0.000421 | 0.00135 |
| CNAG_05013 | 9137.499 | -0.30488 | 0.072458 | -4.20762 | 2.58E-05 | 0.000105 |
| CNAG_05015 | 335.0409 | 1.893688 | 0.131301 | 14.42249 | 3.74E-47 | 1.20E-45 |
| CNAG_05016 | 32.448 | 1.771368 | 0.403398 | 4.39112 | 1.13E-05 | 4.90E-05 |
| CNAG_05017 | 719.6035 | 1.248052 | 0.09502 | 13.13465 | 2.08E-39 | 5.59E-38 |
| CNAG_05018 | 146.9048 | 1.124596 | 0.206462 | 5.446981 | 5.12E-08 | 3.11E-07 |
| CNAG_05020 | 326.5855 | 0.908408 | 0.12959 | 7.009844 | 2.39E-12 | 2.21E-11 |
| CNAG_05023 | 907.7706 | -0.66486 | 0.088613 | -7.50298 | 6.24E-14 | 6.49E-13 |
| CNAG_05024 | 256.4629 | 0.525251 | 0.143631 | 3.656941 | 0.000255 | 0.000857 |
| CNAG_05028 | 4161.375 | -0.40561 | 0.065159 | -6.22498 | 4.82E-10 | 3.63E-09 |
| CNAG_05030 | 934.0634 | -0.23955 | 0.092715 | -2.58369 | 0.009775 | 0.022635 |
| CNAG_05031 | 2219.891 | 1.283286 | 0.105416 | 12.17354 | 4.30E-34 | 9.90E-33 |
| CNAG_05032 | 949.1459 | 3.74759 | 0.287351 | 13.04187 | 7.07E-39 | 1.87E-37 |
| CNAG_05033 | 2213.652 | -0.1753 | 0.075462 | -2.32295 | 0.020182 | 0.042718 |
| CNAG_05036 | 1827.725 | 0.38678 | 0.074252 | 5.209032 | 1.90E-07 | 1.07E-06 |
| CNAG_05038 | 406.5965 | 0.339821 | 0.123014 | 2.762448 | 0.005737 | 0.014126 |
| CNAG_05039 | 802.35 | -0.31566 | 0.104967 | -3.00725 | 0.002636 | 0.007053 |
| CNAG_05041 | 10220.64 | 0.982959 | 0.066609 | 14.75718 | 2.77E-49 | 9.21E-48 |
| CNAG_05043 | 1263.938 | -0.40706 | 0.092246 | -4.41275 | 1.02E-05 | 4.48E-05 |
| CNAG_07805 | 4517.346 | 0.231309 | 0.071282 | 3.244976 | 0.001175 | 0.0034 |
| CNAG_07807 | 25738.57 | -1.10582 | 0.084811 | -13.0386 | 7.38E-39 | 1.94E-37 |
| CNAG_05053 | 662.6556 | -0.35686 | 0.108355 | -3.29343 | 0.00099 | 0.002916 |
| CNAG_05057 | 1311.571 | 0.441489 | 0.096174 | 4.590506 | 4.42E-06 | 2.07E-05 |
| CNAG_05059 | 12107.97 | -0.43815 | 0.079908 | -5.48317 | 4.18E-08 | 2.55E-07 |
| CNAG_05060 | 870.4215 | -0.49555 | 0.100222 | -4.94453 | 7.63E-07 | 3.98E-06 |
| CNAG_05061 | 65.32604 | -1.36737 | 0.308108 | -4.43795 | 9.08E-06 | 4.03E-05 |
| CNAG_12361 | 55.31546 | -0.96071 | 0.292071 | -3.28931 | 0.001004 | 0.002953 |
| CNAG_05067 | 51.39233 | 0.901125 | 0.324796 | 2.774434 | 0.00553 | 0.013681 |
| CNAG_05068 | 7225.774 | -0.27472 | 0.071315 | -3.85223 | 0.000117 | 0.000423 |
| CNAG_05069 | 5579.525 | 0.835188 | 0.096159 | 8.685443 | 3.77E-18 | 4.88E-17 |
| CNAG_12362 | 20.45336 | 2.324042 | 0.512804 | 4.532031 | 5.84E-06 | 2.69E-05 |
| CNAG_05070 | 14629.43 | 0.68106 | 0.058527 | 11.63669 | 2.68E-31 | 5.68E-30 |
| CNAG_05072 | 11.54336 | -1.76605 | 0.680793 | -2.5941 | 0.009484 | 0.022056 |
| CNAG_07808 | 3010.839 | 0.705937 | 0.084193 | 8.384742 | 5.08E-17 | 6.23E-16 |
| CNAG_05077 | 2015.802 | 0.169181 | 0.085403 | 1.980964 | 0.047595 | 0.089945 |
| CNAG_05079 | 73.28773 | -2.07735 | 0.322044 | -6.45052 | 1.11E-10 | 8.96E-10 |
| CNAG_05082 | 1383.624 | -0.17808 | 0.076448 | -2.32948 | 0.019834 | 0.042069 |
| CNAG_05083 | 2436.412 | -0.35526 | 0.072617 | -4.89228 | 9.97E-07 | 5.11E-06 |
| CNAG_05085 | 2140.562 | 0.786827 | 0.070753 | 11.12082 | 9.94E-29 | 1.94E-27 |

| | | | | | | |
|---|---|---|---|---|---|---|
| CNAG_05086 | 1152.817 | 0.87687 | 0.098234 | 8.926363 | 4.40E-19 | 5.95E-18 |
| CNAG_07810 | 9738.756 | -0.32224 | 0.079066 | -4.07559 | 4.59E-05 | 0.000178 |
| CNAG_05090 | 2380.403 | 0.442873 | 0.086706 | 5.107756 | 3.26E-07 | 1.79E-06 |
| CNAG_05091 | 1808.589 | -0.30853 | 0.073113 | -4.2199 | 2.44E-05 | 9.92E-05 |
| CNAG_05095 | 1569.343 | -0.40495 | 0.127725 | -3.17048 | 0.001522 | 0.004292 |
| CNAG_05096 | 1292.447 | -0.19151 | 0.08477 | -2.2592 | 0.023871 | 0.049457 |
| CNAG_05097 | 1177.917 | -0.40455 | 0.103374 | -3.91349 | 9.10E-05 | 0.000337 |
| CNAG_05098 | 1216.253 | 0.190289 | 0.085742 | 2.219317 | 0.026465 | 0.054184 |
| CNAG_05105 | 9325.032 | -0.20987 | 0.06399 | -3.27967 | 0.001039 | 0.003045 |
| CNAG_05109 | 4779.12 | 0.527276 | 0.063888 | 8.253106 | 1.54E-16 | 1.84E-15 |
| CNAG_05110 | 1680.742 | -0.2185 | 0.111369 | -1.96198 | 0.049764 | 0.093264 |
| CNAG_05111 | 3080.163 | -0.321 | 0.081008 | -3.96263 | 7.41E-05 | 0.000279 |
| CNAG_12368 | 143.3247 | 1.552887 | 0.211247 | 7.351061 | 1.97E-13 | 1.98E-12 |
| CNAG_05112 | 2301.202 | 1.006007 | 0.076467 | 13.15608 | 1.57E-39 | 4.25E-38 |
| CNAG_05113 | 1117.493 | 0.312703 | 0.079913 | 3.913029 | 9.11E-05 | 0.000337 |
| CNAG_05114 | 256.7402 | 2.369697 | 0.159936 | 14.81657 | 1.14E-49 | 3.86E-48 |
| CNAG_05115 | 1606.862 | 1.110322 | 0.074678 | 14.86803 | 5.32E-50 | 1.82E-48 |
| CNAG_12369 | 27.75856 | 2.055094 | 0.478831 | 4.291901 | 1.77E-05 | 7.43E-05 |
| CNAG_05117 | 4567.603 | 0.253958 | 0.067696 | 3.751465 | 0.000176 | 0.000612 |
| CNAG_05118 | 1355.787 | 0.388879 | 0.089159 | 4.361645 | 1.29E-05 | 5.55E-05 |
| CNAG_05119 | 575.8568 | 1.274026 | 0.136506 | 9.333129 | 1.03E-20 | 1.51E-19 |
| CNAG_05120 | 2100.934 | -0.4123 | 0.077203 | -5.34055 | 9.27E-08 | 5.47E-07 |
| CNAG_05121 | 2730.212 | 0.291094 | 0.082601 | 3.524088 | 0.000425 | 0.001363 |
| CNAG_05123 | 504.3681 | -0.49683 | 0.113761 | -4.3673 | 1.26E-05 | 5.41E-05 |
| CNAG_05124 | 2296.328 | -0.28153 | 0.080055 | -3.51664 | 0.000437 | 0.001397 |
| CNAG_05125 | 2716.751 | -0.47431 | 0.092792 | -5.11158 | 3.19E-07 | 1.75E-06 |
| CNAG_07814 | 988.4651 | -0.43556 | 0.091771 | -4.74614 | 2.07E-06 | 1.02E-05 |
| CNAG_05130 | 1544.575 | -1.23167 | 0.098751 | -12.4726 | 1.05E-35 | 2.53E-34 |
| CNAG_05132 | 11617.38 | 0.462987 | 0.072879 | 6.352804 | 2.11E-10 | 1.66E-09 |
| CNAG_05131 | 376.9968 | -0.41223 | 0.117419 | -3.51071 | 0.000447 | 0.001425 |
| CNAG_05134 | 1169.327 | -0.30972 | 0.083713 | -3.69985 | 0.000216 | 0.000736 |
| CNAG_05136 | 6004.101 | -0.13809 | 0.069273 | -1.99348 | 0.046209 | 0.087651 |
| CNAG_05137 | 1755.779 | -0.2048 | 0.102592 | -1.99626 | 0.045905 | 0.087198 |
| CNAG_05138 | 3125.271 | 1.110811 | 0.080524 | 13.79471 | 2.74E-43 | 8.32E-42 |
| CNAG_12370 | 18.05912 | 2.004128 | 0.669131 | 2.995123 | 0.002743 | 0.007318 |
| CNAG_05140 | 10255.64 | 0.12173 | 0.057355 | 2.122406 | 0.033804 | 0.066834 |
| CNAG_05141 | 1712.432 | 0.579137 | 0.075357 | 7.685225 | 1.53E-14 | 1.64E-13 |
| CNAG_05144 | 11230.34 | -0.44646 | 0.079205 | -5.63678 | 1.73E-08 | 1.11E-07 |
| CNAG_05147 | 219.1772 | 0.821716 | 0.193004 | 4.257499 | 2.07E-05 | 8.55E-05 |
| CNAG_05149 | 1119.569 | 0.236133 | 0.086694 | 2.723748 | 0.006455 | 0.015642 |
| CNAG_05150 | 2973.953 | 0.356125 | 0.063554 | 5.6035 | 2.10E-08 | 1.33E-07 |
| CNAG_05153 | 1216.98 | 0.181734 | 0.092317 | 1.968587 | 0.049001 | 0.092066 |
| CNAG_05154 | 7007.322 | 1.851586 | 0.062938 | 29.41943 | 3.10E-190 | 5.86E-188 |
| CNAG_12373 | 6.681207 | 1.78673 | 0.894214 | 1.998102 | 0.045706 | 0.0869 |
| CNAG_05157 | 143.3155 | -0.50479 | 0.182127 | -2.77164 | 0.005577 | 0.013779 |
| CNAG_12377 | 149.6482 | 0.567683 | 0.254114 | 2.23397 | 0.025485 | 0.052388 |
| CNAG_05159 | 122.584 | -0.53072 | 0.244078 | -2.17441 | 0.029675 | 0.05982 |

| | | | | | | |
|---|---|---|---|---|---|---|
| CNAG_05161 | 137.2744 | -1.79375 | 0.213288 | -8.41 | 4.10E-17 | 5.05E-16 |
| CNAG_05163 | 601.4705 | 0.327541 | 0.107829 | 3.037595 | 0.002385 | 0.006448 |
| CNAG_05164 | 2271.49 | -0.15869 | 0.071227 | -2.22794 | 0.025885 | 0.053143 |
| CNAG_05166 | 4634.123 | -0.39392 | 0.069023 | -5.707 | 1.15E-08 | 7.49E-08 |
| CNAG_05167 | 2328.707 | -0.52594 | 0.117605 | -4.47209 | 7.75E-06 | 3.49E-05 |
| CNAG_12380 | 17.99518 | 1.428376 | 0.520617 | 2.743622 | 0.006077 | 0.014827 |
| CNAG_05169 | 3110.038 | 2.702743 | 0.109397 | 24.70572 | 9.28E-135 | 9.80E-133 |
| CNAG_12381 | 69.96935 | 2.084069 | 0.34994 | 5.955495 | 2.59E-09 | 1.82E-08 |
| CNAG_05170 | 1033.9 | 0.596352 | 0.089463 | 6.66589 | 2.63E-11 | 2.23E-10 |
| CNAG_05171 | 1422.345 | -0.4661 | 0.089883 | -5.18568 | 2.15E-07 | 1.20E-06 |
| CNAG_05177 | 1456.422 | 0.227718 | 0.086574 | 2.630328 | 0.00853 | 0.020074 |
| CNAG_05179 | 17994.69 | 0.464056 | 0.060751 | 7.638688 | 2.19E-14 | 2.34E-13 |
| CNAG_07974 | 626.1113 | 1.11659 | 0.104539 | 10.68106 | 1.25E-26 | 2.26E-25 |
| CNAG_05180 | 510.1348 | 0.501982 | 0.1104 | 4.546924 | 5.44E-06 | 2.52E-05 |
| CNAG_12383 | 16.96595 | 1.114616 | 0.527643 | 2.112444 | 0.034648 | 0.068239 |
| CNAG_05181 | 811.1847 | 0.349849 | 0.097495 | 3.588381 | 0.000333 | 0.001094 |
| CNAG_05182 | 811.9484 | -0.37707 | 0.098427 | -3.83097 | 0.000128 | 0.000457 |
| CNAG_05185 | 156.3645 | -0.58834 | 0.211815 | -2.7776 | 0.005476 | 0.013566 |
| CNAG_05186 | 499.7637 | -0.35836 | 0.127335 | -2.81429 | 0.004889 | 0.012241 |
| CNAG_05187 | 1602.931 | 0.361433 | 0.073595 | 4.911086 | 9.06E-07 | 4.67E-06 |
| CNAG_05189 | 1387.132 | 0.288566 | 0.083663 | 3.449132 | 0.000562 | 0.001756 |
| CNAG_05190 | 2652.407 | 0.322733 | 0.069991 | 4.611058 | 4.01E-06 | 1.89E-05 |
| CNAG_05192 | 118.4266 | 0.477778 | 0.192558 | 2.481219 | 0.013093 | 0.029371 |
| CNAG_05194 | 4024.229 | 1.301325 | 0.075339 | 17.27284 | 7.53E-67 | 3.64E-65 |
| CNAG_05196 | 1398.305 | -0.47529 | 0.102119 | -4.65425 | 3.25E-06 | 1.55E-05 |
| CNAG_05197 | 5626.599 | 0.403848 | 0.070613 | 5.719196 | 1.07E-08 | 7.00E-08 |
| CNAG_05198 | 1090.519 | 0.455592 | 0.087778 | 5.190248 | 2.10E-07 | 1.18E-06 |
| CNAG_05199 | 28688.81 | 0.568871 | 0.060934 | 9.335857 | 1.00E-20 | 1.48E-19 |
| CNAG_05200 | 809.4788 | 0.197031 | 0.087657 | 2.247751 | 0.024592 | 0.050797 |
| CNAG_05212 | 59.10329 | -0.62568 | 0.270824 | -2.31028 | 0.020873 | 0.043906 |
| CNAG_05217 | 415.4146 | -0.65667 | 0.127491 | -5.15069 | 2.60E-07 | 1.44E-06 |
| CNAG_05218 | 2932.12 | -0.22938 | 0.072224 | -3.17598 | 0.001493 | 0.004213 |
| CNAG_05219 | 1735.686 | -0.40949 | 0.07927 | -5.16582 | 2.39E-07 | 1.33E-06 |
| CNAG_05221 | 2675.099 | -1.18651 | 0.075933 | -15.6257 | 4.86E-55 | 1.83E-53 |
| CNAG_05224 | 174.721 | -0.5736 | 0.166136 | -3.4526 | 0.000555 | 0.001737 |
| CNAG_05226 | 1536.912 | -0.2271 | 0.078464 | -2.89428 | 0.0038 | 0.009784 |
| CNAG_05229 | 570.6193 | -1.53384 | 0.1166 | -13.1547 | 1.60E-39 | 4.32E-38 |
| CNAG_05230 | 683.7068 | -0.30471 | 0.123929 | -2.45878 | 0.013941 | 0.03096 |
| CNAG_05231 | 1700.637 | -0.50361 | 0.086937 | -5.79278 | 6.92E-09 | 4.65E-08 |
| CNAG_05232 | 83884.47 | -0.34219 | 0.069452 | -4.92708 | 8.35E-07 | 4.32E-06 |
| CNAG_05234 | 3295.855 | 0.20499 | 0.095432 | 2.148012 | 0.031713 | 0.063367 |
| CNAG_05235 | 22620.91 | -0.19384 | 0.067958 | -2.85237 | 0.004339 | 0.011026 |
| CNAG_05236 | 1697.382 | -0.52071 | 0.076757 | -6.78391 | 1.17E-11 | 1.02E-10 |
| CNAG_05241 | 1581.134 | -0.1829 | 0.076462 | -2.39207 | 0.016754 | 0.036351 |
| CNAG_05242 | 4061.156 | -0.12499 | 0.063657 | -1.96343 | 0.049596 | 0.093034 |
| CNAG_07975 | 334.1651 | 1.440868 | 0.176762 | 8.151478 | 3.60E-16 | 4.21E-15 |
| CNAG_05245 | 334.1354 | -0.31375 | 0.158461 | -1.98 | 0.047703 | 0.090065 |

| | | | | | | |
|---|---|---|---|---|---|---|
| CNAG_05248 | 3066.916 | -0.17159 | 0.065968 | -2.6011 | 0.009292 | 0.02171 |
| CNAG_07822 | 653.6228 | -0.42077 | 0.105599 | -3.98463 | 6.76E-05 | 0.000256 |
| CNAG_12389 | 171.9461 | 2.348205 | 0.238384 | 9.850507 | 6.82E-23 | 1.10E-21 |
| CNAG_05251 | 796.9937 | 1.629703 | 0.101504 | 16.05561 | 5.22E-58 | 2.09E-56 |
| CNAG_05252 | 2431.576 | 0.97652 | 0.066849 | 14.60791 | 2.50E-48 | 8.16E-47 |
| CNAG_05253 | 1080.852 | 1.693833 | 0.105026 | 16.12769 | 1.63E-58 | 6.66E-57 |
| CNAG_05256 | 2128.427 | 2.228433 | 0.092132 | 24.18741 | 3.02E-129 | 2.92E-127 |
| CNAG_05258 | 369.8197 | 3.57619 | 0.167193 | 21.38959 | 1.67E-101 | 1.23E-99 |
| CNAG_05259 | 99.38353 | 0.736059 | 0.249523 | 2.949857 | 0.003179 | 0.008346 |
| CNAG_05262 | 3014.407 | -0.33388 | 0.066045 | -5.0553 | 4.30E-07 | 2.32E-06 |
| CNAG_05264 | 366.7278 | 0.842401 | 0.390429 | 2.157631 | 0.030957 | 0.062035 |
| CNAG_12392 | 95.38313 | 1.593755 | 0.311633 | 5.11421 | 3.15E-07 | 1.73E-06 |
| CNAG_05266 | 135.2581 | 0.487034 | 0.187548 | 2.596848 | 0.009408 | 0.02193 |
| CNAG_05267 | 3150.78 | 0.447121 | 0.073293 | 6.100419 | 1.06E-09 | 7.73E-09 |
| CNAG_05268 | 65.76442 | -0.85486 | 0.282433 | -3.02678 | 0.002472 | 0.006657 |
| CNAG_05270 | 2973.033 | 0.166399 | 0.067399 | 2.468862 | 0.013554 | 0.030242 |
| CNAG_05273 | 2121.856 | 0.28265 | 0.087418 | 3.233301 | 0.001224 | 0.003524 |
| CNAG_07824 | 447.3292 | 0.611747 | 0.127884 | 4.783608 | 1.72E-06 | 8.57E-06 |
| CNAG_05276 | 1420.416 | -0.18051 | 0.085853 | -2.10253 | 0.035507 | 0.069693 |
| CNAG_05277 | 3466.7 | 0.898932 | 0.068906 | 13.0457 | 6.72E-39 | 1.79E-37 |
| CNAG_05278 | 61.7579 | 7.624976 | 1.052415 | 7.24522 | 4.32E-13 | 4.28E-12 |
| CNAG_12395 | 21.96557 | 3.173725 | 0.591152 | 5.368714 | 7.93E-08 | 4.73E-07 |
| CNAG_05279 | 2631.03 | 6.126184 | 0.249191 | 24.58428 | 1.86E-133 | 1.89E-131 |
| CNAG_05280 | 1114.549 | 0.284893 | 0.080093 | 3.557051 | 0.000375 | 0.001218 |
| CNAG_05282 | 1497.062 | -0.31736 | 0.08947 | -3.54716 | 0.000389 | 0.001258 |
| CNAG_05283 | 1215.124 | 1.110832 | 0.083616 | 13.28498 | 2.83E-40 | 7.80E-39 |
| CNAG_12396 | 12.93101 | 1.858902 | 0.685384 | 2.712206 | 0.006684 | 0.016139 |
| CNAG_05288 | 804.1079 | 0.195837 | 0.095713 | 2.046091 | 0.040747 | 0.078669 |
| CNAG_12400 | 38.51375 | 0.898263 | 0.376109 | 2.388308 | 0.016926 | 0.036648 |
| CNAG_05290 | 719.1727 | 0.679484 | 0.126197 | 5.384296 | 7.27E-08 | 4.35E-07 |
| CNAG_12401 | 13.92106 | 1.532528 | 0.605308 | 2.531817 | 0.011347 | 0.025903 |
| CNAG_05291 | 1973.892 | 0.302427 | 0.080337 | 3.764455 | 0.000167 | 0.000585 |
| CNAG_05292 | 4126.643 | -0.48625 | 0.077514 | -6.27304 | 3.54E-10 | 2.71E-09 |
| CNAG_05293 | 2957.086 | -0.82293 | 0.094977 | -8.66453 | 4.53E-18 | 5.82E-17 |
| CNAG_05297 | 402.3901 | -0.31514 | 0.129897 | -2.42609 | 0.015263 | 0.033483 |
| CNAG_05298 | 436.5836 | 0.472016 | 0.117258 | 4.025431 | 5.69E-05 | 0.000218 |
| CNAG_05299 | 251.3216 | 2.529455 | 0.249328 | 10.1451 | 3.48E-24 | 5.91E-23 |
| CNAG_05300 | 624.4155 | 2.068554 | 0.116803 | 17.7097 | 3.53E-70 | 1.76E-68 |
| CNAG_05301 | 3102.69 | -0.1702 | 0.070222 | -2.42379 | 0.01536 | 0.033669 |
| CNAG_05302 | 1648.588 | 0.641128 | 0.081689 | 7.848433 | 4.21E-15 | 4.68E-14 |
| CNAG_05303 | 151.0915 | 0.691016 | 0.184073 | 3.754033 | 0.000174 | 0.000607 |
| CNAG_12402 | 32.41819 | 1.876859 | 0.488442 | 3.842544 | 0.000122 | 0.000439 |
| CNAG_05306 | 758.6465 | -0.29178 | 0.104849 | -2.78284 | 0.005389 | 0.013382 |
| CNAG_05307 | 1945.364 | -0.17331 | 0.078894 | -2.19681 | 0.028034 | 0.056979 |
| CNAG_05309 | 2693.974 | -0.61949 | 0.09393 | -6.59524 | 4.25E-11 | 3.54E-10 |
| CNAG_05310 | 62.6933 | -1.20365 | 0.301628 | -3.99051 | 6.59E-05 | 0.00025 |
| CNAG_05311 | 7830.278 | -0.20912 | 0.070558 | -2.9638 | 0.003039 | 0.008007 |

| | | | | | | |
|---|---|---|---|---|---|---|
| CNAG_05312 | 3299.63 | -0.53008 | 0.090541 | -5.85462 | 4.78E-09 | 3.27E-08 |
| CNAG_05313 | 1060.437 | -0.2633 | 0.095757 | -2.74963 | 0.005966 | 0.014593 |
| CNAG_05315 | 291.9634 | 0.895501 | 0.169058 | 5.296994 | 1.18E-07 | 6.83E-07 |
| CNAG_12405 | 204.7944 | 0.561534 | 0.170389 | 3.295591 | 0.000982 | 0.002897 |
| CNAG_05316 | 1392.86 | 1.760264 | 0.092722 | 18.98439 | 2.30E-80 | 1.28E-78 |
| CNAG_05317 | 744.2843 | 0.323216 | 0.098656 | 3.276176 | 0.001052 | 0.003076 |
| CNAG_05318 | 870.0095 | 0.608821 | 0.096632 | 6.300397 | 2.97E-10 | 2.30E-09 |
| CNAG_05320 | 563.85 | 0.87678 | 0.125773 | 6.971118 | 3.14E-12 | 2.87E-11 |
| CNAG_12407 | 474.6261 | 1.103692 | 0.154485 | 7.144325 | 9.04E-13 | 8.66E-12 |
| CNAG_05321 | 307.5637 | 1.021512 | 0.129011 | 7.918006 | 2.41E-15 | 2.72E-14 |
| CNAG_07830 | 1862.322 | 2.128592 | 0.0757 | 28.11882 | 5.77E-174 | 9.57E-172 |
| CNAG_05324 | 645.1815 | 4.513806 | 0.151582 | 29.77789 | 7.55E-195 | 1.46E-192 |
| CNAG_05327 | 2061.017 | 0.325903 | 0.074185 | 4.393086 | 1.12E-05 | 4.86E-05 |
| CNAG_07977 | 1475.823 | 1.211218 | 0.076701 | 15.79136 | 3.57E-56 | 1.37E-54 |
| CNAG_05329 | 839.1827 | 5.311932 | 0.175415 | 30.28216 | 1.97E-201 | 4.00E-199 |
| CNAG_05330 | 119.6345 | 4.965588 | 0.348919 | 14.23133 | 5.86E-46 | 1.84E-44 |
| CNAG_05331 | 52.04084 | 0.960884 | 0.320577 | 2.997361 | 0.002723 | 0.007269 |
| CNAG_06876 | 779.2441 | 1.368371 | 0.100185 | 13.65848 | 1.80E-42 | 5.20E-41 |
| CNAG_06875 | 609.493 | 0.333635 | 0.097782 | 3.412033 | 0.000645 | 0.001986 |
| CNAG_06874 | 550.5545 | -0.38057 | 0.122542 | -3.10562 | 0.001899 | 0.005266 |
| CNAG_06873 | 110.8584 | 1.491463 | 0.230971 | 6.457364 | 1.07E-10 | 8.59E-10 |
| CNAG_12409 | 23.17785 | -1.00846 | 0.442666 | -2.27816 | 0.022717 | 0.047369 |
| CNAG_06871 | 1359.419 | -0.28467 | 0.097255 | -2.92705 | 0.003422 | 0.008912 |
| CNAG_12411 | 19.48531 | 0.966965 | 0.484718 | 1.994901 | 0.046054 | 0.087432 |
| CNAG_06867 | 727.372 | 0.397621 | 0.130497 | 3.046965 | 0.002312 | 0.006269 |
| CNAG_06864 | 2532.572 | -0.21576 | 0.089076 | -2.42219 | 0.015427 | 0.033789 |
| CNAG_06863 | 338.4678 | -0.38995 | 0.133204 | -2.92746 | 0.003417 | 0.008904 |
| CNAG_12412 | 348.271 | 0.315261 | 0.147741 | 2.133879 | 0.032853 | 0.065272 |
| CNAG_06853 | 326.0104 | 1.118203 | 0.145059 | 7.708609 | 1.27E-14 | 1.37E-13 |
| CNAG_07400 | 10186.01 | -0.21976 | 0.069463 | -3.16376 | 0.001557 | 0.004386 |
| CNAG_06849 | 4344.826 | -0.59848 | 0.06547 | -9.14126 | 6.17E-20 | 8.71E-19 |
| CNAG_07401 | 177.0753 | -0.42519 | 0.194878 | -2.18182 | 0.029123 | 0.058839 |
| CNAG_06846 | 382.6651 | -0.33515 | 0.139111 | -2.40923 | 0.015986 | 0.034845 |
| CNAG_06845 | 750.5663 | 0.189469 | 0.092849 | 2.040624 | 0.041288 | 0.079543 |
| CNAG_07403 | 911.792 | -0.23172 | 0.093389 | -2.48129 | 0.013091 | 0.029371 |
| CNAG_06839 | 3383.295 | -0.27282 | 0.065563 | -4.16125 | 3.17E-05 | 0.000127 |
| CNAG_06837 | 1278.167 | -0.51672 | 0.09185 | -5.62566 | 1.85E-08 | 1.18E-07 |
| CNAG_06836 | 204.3801 | 1.704907 | 0.192043 | 8.877736 | 6.82E-19 | 9.11E-18 |
| CNAG_12417 | 37.2923 | 1.75197 | 0.372295 | 4.705861 | 2.53E-06 | 1.23E-05 |
| CNAG_06835 | 794.9398 | 1.234079 | 0.135537 | 9.105138 | 8.62E-20 | 1.21E-18 |
| CNAG_06834 | 3261.638 | 0.643057 | 0.078762 | 8.164613 | 3.22E-16 | 3.79E-15 |
| CNAG_06833 | 27.9277 | 1.403582 | 0.456817 | 3.072523 | 0.002123 | 0.005801 |
| CNAG_06832 | 920.3238 | -0.95112 | 0.096367 | -9.86972 | 5.63E-23 | 9.12E-22 |
| CNAG_06829 | 2553.808 | 0.308911 | 0.068702 | 4.496424 | 6.91E-06 | 3.14E-05 |
| CNAG_06828 | 2575.363 | 1.427275 | 0.075482 | 18.90892 | 9.63E-80 | 5.33E-78 |
| CNAG_06827 | 49.42285 | 1.041851 | 0.3452 | 3.018111 | 0.002544 | 0.006832 |
| CNAG_06821 | 2895.425 | 1.551905 | 0.064507 | 24.05797 | 6.89E-128 | 6.51E-126 |

| | | | | | | |
|---|---|---|---|---|---|---|
| CNAG_06820 | 1439.494 | 0.509245 | 0.074755 | 6.812211 | 9.61E-12 | 8.54E-11 |
| CNAG_06819 | 2807.721 | 0.345364 | 0.079771 | 4.329446 | 1.49E-05 | 6.36E-05 |
| CNAG_07404 | 115.385 | 0.801751 | 0.247383 | 3.240925 | 0.001191 | 0.003439 |
| CNAG_07405 | 450.8453 | 2.085257 | 0.124001 | 16.81646 | 1.85E-63 | 8.40E-62 |
| CNAG_06818 | 1035.31 | 2.050663 | 0.090748 | 22.59727 | 4.61E-113 | 3.86E-111 |
| CNAG_06817 | 1193.794 | 3.065309 | 0.098589 | 31.09195 | 3.09E-212 | 7.18E-210 |
| CNAG_06816 | 340.0522 | 1.070711 | 0.124717 | 8.585135 | 9.07E-18 | 1.14E-16 |
| CNAG_12420 | 252.9743 | 2.547229 | 0.181866 | 14.00606 | 1.43E-44 | 4.42E-43 |
| CNAG_06815 | 125.4043 | 1.334185 | 0.225921 | 5.905546 | 3.51E-09 | 2.44E-08 |
| CNAG_06814 | 478.9307 | 1.496282 | 0.115648 | 12.93826 | 2.74E-38 | 7.13E-37 |
| CNAG_12421 | 190.4192 | -1.26329 | 0.1755 | -7.19823 | 6.10E-13 | 5.93E-12 |
| CNAG_06809 | 1241.476 | -0.38262 | 0.077268 | -4.95181 | 7.35E-07 | 3.84E-06 |
| CNAG_06808 | 605.9115 | 1.228807 | 0.121677 | 10.0989 | 5.59E-24 | 9.38E-23 |
| CNAG_06805 | 33.0125 | 1.917873 | 0.401044 | 4.782198 | 1.73E-06 | 8.62E-06 |
| CNAG_06804 | 1380.488 | 0.352979 | 0.083232 | 4.240922 | 2.23E-05 | 9.13E-05 |
| CNAG_07406 | 54.93966 | 0.968245 | 0.393585 | 2.460067 | 0.013891 | 0.030883 |
| CNAG_07407 | 71.61637 | 0.79104 | 0.299317 | 2.642813 | 0.008222 | 0.019421 |
| CNAG_06971 | 4050.666 | 0.159789 | 0.062603 | 2.55239 | 0.010699 | 0.024572 |
| CNAG_07408 | 2062.916 | -0.25742 | 0.069533 | -3.7021 | 0.000214 | 0.000731 |
| CNAG_07409 | 3660.77 | 0.271444 | 0.068006 | 3.991477 | 6.57E-05 | 0.000249 |
| CNAG_07005 | 576.1913 | -0.43309 | 0.116119 | -3.72967 | 0.000192 | 0.000663 |
| CNAG_07004 | 18224.74 | -0.33796 | 0.063394 | -5.33116 | 9.76E-08 | 5.73E-07 |
| CNAG_07410 | 4413.381 | 0.172776 | 0.060649 | 2.848803 | 0.004388 | 0.011133 |
| CNAG_07015 | 1398.029 | -0.25379 | 0.083981 | -3.02199 | 0.002511 | 0.006752 |
| CNAG_01457 | 1598.423 | -0.39639 | 0.073737 | -5.37577 | 7.63E-08 | 4.55E-07 |
| CNAG_01456 | 29.17747 | -1.83892 | 0.453971 | -4.05075 | 5.11E-05 | 0.000197 |
| CNAG_12428 | 57.94383 | -1.08559 | 0.318963 | -3.4035 | 0.000665 | 0.002042 |
| CNAG_01454 | 3979.85 | -0.13834 | 0.064345 | -2.14995 | 0.031559 | 0.063094 |
| CNAG_01451 | 1274.038 | 0.419753 | 0.127815 | 3.284057 | 0.001023 | 0.003002 |
| CNAG_07414 | 731.8338 | -0.37022 | 0.092076 | -4.02075 | 5.80E-05 | 0.000222 |
| CNAG_01446 | 3272.351 | -0.81524 | 0.124593 | -6.54327 | 6.02E-11 | 4.95E-10 |
| CNAG_01445 | 381.0788 | -0.47668 | 0.146904 | -3.24484 | 0.001175 | 0.0034 |
| CNAG_01444 | 921.4469 | -0.43515 | 0.10566 | -4.11836 | 3.82E-05 | 0.000151 |
| CNAG_01443 | 826.2441 | 1.197111 | 0.104811 | 11.4216 | 3.26E-30 | 6.76E-29 |
| CNAG_01442 | 2785.478 | 0.169416 | 0.078872 | 2.147986 | 0.031715 | 0.063367 |
| CNAG_01441 | 1642.068 | 0.518177 | 0.097635 | 5.307309 | 1.11E-07 | 6.48E-07 |
| CNAG_01440 | 367.6182 | -0.4922 | 0.121569 | -4.04867 | 5.15E-05 | 0.000199 |
| CNAG_01439 | 2406.278 | -0.2884 | 0.07476 | -3.85773 | 0.000114 | 0.000414 |
| CNAG_07415 | 2189.223 | -0.38215 | 0.079042 | -4.8348 | 1.33E-06 | 6.70E-06 |
| CNAG_01431 | 447.9901 | -0.27954 | 0.117409 | -2.38089 | 0.017271 | 0.037268 |
| CNAG_01430 | 1081.559 | -0.27635 | 0.106715 | -2.5896 | 0.009609 | 0.022289 |
| CNAG_12433 | 27.66627 | 0.954041 | 0.447586 | 2.131524 | 0.033046 | 0.065592 |
| CNAG_01426 | 2020.998 | -0.43327 | 0.078936 | -5.48888 | 4.05E-08 | 2.48E-07 |
| CNAG_01425 | 211.7247 | -0.49022 | 0.160176 | -3.06051 | 0.00221 | 0.006017 |
| CNAG_01424 | 909.5627 | 0.526314 | 0.109995 | 4.784903 | 1.71E-06 | 8.52E-06 |
| CNAG_01423 | 3020.289 | -0.15825 | 0.066677 | -2.37342 | 0.017624 | 0.037967 |
| CNAG_01422 | 1566.352 | -0.27827 | 0.088003 | -3.16202 | 0.001567 | 0.004411 |

| | | | | | | |
|---|---|---|---|---|---|---|
| CNAG_01420 | 874.3747 | 0.903063 | 0.089198 | 10.12426 | 4.31E-24 | 7.30E-23 |
| CNAG_01419 | 1333.771 | 0.377807 | 0.092692 | 4.075959 | 4.58E-05 | 0.000178 |
| CNAG_07416 | 1080.944 | -0.29903 | 0.103088 | -2.9007 | 0.003723 | 0.009604 |
| CNAG_01418 | 1011.259 | -0.43137 | 0.122413 | -3.52389 | 0.000425 | 0.001364 |
| CNAG_01416 | 1617.684 | -0.21997 | 0.077936 | -2.82239 | 0.004767 | 0.011962 |
| CNAG_07418 | 819.1113 | -0.36503 | 0.099912 | -3.65355 | 0.000259 | 0.000867 |
| CNAG_01408 | 1382.9 | -0.37368 | 0.086311 | -4.32941 | 1.50E-05 | 6.36E-05 |
| CNAG_01405 | 1726.749 | -0.49785 | 0.075989 | -6.55166 | 5.69E-11 | 4.70E-10 |
| CNAG_01402 | 3784.006 | -0.17377 | 0.075923 | -2.28872 | 0.022095 | 0.046275 |
| CNAG_01400 | 13099.25 | 0.750417 | 0.057338 | 13.08753 | 3.88E-39 | 1.04E-37 |
| CNAG_01399 | 1025.703 | 0.23153 | 0.097738 | 2.368878 | 0.017842 | 0.038375 |
| CNAG_01395 | 2654.35 | -0.29907 | 0.067322 | -4.44244 | 8.89E-06 | 3.96E-05 |
| CNAG_07419 | 610.634 | -0.55483 | 0.111215 | -4.9888 | 6.08E-07 | 3.21E-06 |
| CNAG_01392 | 666.3458 | -0.28657 | 0.109113 | -2.62636 | 0.00863 | 0.020292 |
| CNAG_01391 | 1423.612 | -0.52469 | 0.084375 | -6.21847 | 5.02E-10 | 3.79E-09 |
| CNAG_01389 | 821.4383 | -0.28803 | 0.096545 | -2.98338 | 0.002851 | 0.00757 |
| CNAG_01388 | 1141.011 | -0.25767 | 0.091784 | -2.8074 | 0.004994 | 0.012487 |
| CNAG_01385 | 2055.247 | 0.222417 | 0.078767 | 2.82375 | 0.004747 | 0.011918 |
| CNAG_07421 | 74.44494 | -0.70718 | 0.272675 | -2.59347 | 0.009501 | 0.022083 |
| CNAG_01382 | 494.9979 | -0.52339 | 0.106364 | -4.92078 | 8.62E-07 | 4.46E-06 |
| CNAG_01381 | 1436.649 | -0.28948 | 0.08469 | -3.41806 | 0.000631 | 0.001948 |
| CNAG_01380 | 750.7387 | -0.38477 | 0.102202 | -3.76479 | 0.000167 | 0.000584 |
| CNAG_01379 | 1225.438 | 0.22708 | 0.089143 | 2.547369 | 0.010854 | 0.0249 |
| CNAG_01378 | 2213.198 | -0.23949 | 0.068199 | -3.5117 | 0.000445 | 0.00142 |
| CNAG_01377 | 2717.208 | -0.18283 | 0.067933 | -2.69128 | 0.007118 | 0.017066 |
| CNAG_01375 | 1400.126 | 0.245073 | 0.094319 | 2.598351 | 0.009367 | 0.021845 |
| CNAG_01372 | 1690.558 | -0.77771 | 0.091854 | -8.46686 | 2.52E-17 | 3.13E-16 |
| CNAG_01371 | 1163.959 | -0.33067 | 0.082914 | -3.98812 | 6.66E-05 | 0.000252 |
| CNAG_01370 | 76.79685 | -0.83865 | 0.252804 | -3.31738 | 0.000909 | 0.002697 |
| CNAG_12438 | 50.38505 | -0.83896 | 0.326114 | -2.5726 | 0.010094 | 0.023285 |
| CNAG_01367 | 178.3791 | 0.992982 | 0.18146 | 5.47219 | 4.45E-08 | 2.71E-07 |
| CNAG_01365 | 866.7227 | -0.74102 | 0.097226 | -7.6216 | 2.51E-14 | 2.67E-13 |
| CNAG_01364 | 3479.36 | -0.39721 | 0.074576 | -5.32622 | 1.00E-07 | 5.87E-07 |
| CNAG_01363 | 2093.251 | 0.209091 | 0.074841 | 2.793803 | 0.005209 | 0.01298 |
| CNAG_01362 | 3084.091 | -0.28146 | 0.068374 | -4.11643 | 3.85E-05 | 0.000152 |
| CNAG_01360 | 2232.745 | 0.341957 | 0.084497 | 4.04697 | 5.19E-05 | 0.0002 |
| CNAG_01359 | 3117.308 | -0.28212 | 0.079918 | -3.53012 | 0.000415 | 0.001335 |
| CNAG_01352 | 1892.656 | -0.31462 | 0.089872 | -3.50077 | 0.000464 | 0.001473 |
| CNAG_01350 | 761.8542 | -0.29976 | 0.096682 | -3.10044 | 0.001932 | 0.005348 |
| CNAG_01348 | 1021.243 | -0.44395 | 0.132066 | -3.36154 | 0.000775 | 0.002346 |
| CNAG_01347 | 1962.954 | -0.24271 | 0.072084 | -3.36705 | 0.00076 | 0.002305 |
| CNAG_01346 | 930.2224 | -0.31072 | 0.085322 | -3.64173 | 0.000271 | 0.000905 |
| CNAG_12440 | 49.37505 | 0.594102 | 0.302739 | 1.962424 | 0.049713 | 0.093189 |
| CNAG_01345 | 873.322 | -0.33194 | 0.09276 | -3.57842 | 0.000346 | 0.001132 |
| CNAG_01344 | 1102.533 | 0.68156 | 0.114803 | 5.936774 | 2.91E-09 | 2.03E-08 |
| CNAG_01343 | 717.3022 | -0.19376 | 0.092907 | -2.08554 | 0.03702 | 0.072401 |
| CNAG_01342 | 857.3186 | -0.38255 | 0.087477 | -4.37314 | 1.22E-05 | 5.28E-05 |

| | | | | | | |
|---|---|---|---|---|---|---|
| CNAG_01340 | 3179.947 | -0.1909 | 0.07114 | -2.68345 | 0.007287 | 0.017435 |
| CNAG_01338 | 647.4369 | -0.48448 | 0.107178 | -4.52034 | 6.17E-06 | 2.84E-05 |
| CNAG_01336 | 120.0598 | -0.74351 | 0.198226 | -3.75084 | 0.000176 | 0.000613 |
| CNAG_01335 | 887.6743 | 0.262348 | 0.089909 | 2.917933 | 0.003524 | 0.009144 |
| CNAG_01333 | 1106.078 | -0.18507 | 0.080228 | -2.30674 | 0.021069 | 0.044262 |
| CNAG_01332 | 40468.67 | -0.23806 | 0.077465 | -3.07309 | 0.002119 | 0.005792 |
| CNAG_01331 | 429.3237 | 1.275925 | 0.122159 | 10.44477 | 1.55E-25 | 2.73E-24 |
| CNAG_12443 | 254.4594 | -0.60832 | 0.162863 | -3.73517 | 0.000188 | 0.00065 |
| CNAG_01324 | 434.7166 | 0.364342 | 0.120431 | 3.025315 | 0.002484 | 0.006685 |
| CNAG_01323 | 13348.85 | 0.68163 | 0.075403 | 9.03982 | 1.57E-19 | 2.18E-18 |
| CNAG_01321 | 9189.215 | 0.290437 | 0.074232 | 3.912561 | 9.13E-05 | 0.000338 |
| CNAG_01319 | 1050.777 | -0.31326 | 0.085339 | -3.67081 | 0.000242 | 0.000818 |
| CNAG_01317 | 1280.698 | -0.51078 | 0.09856 | -5.1824 | 2.19E-07 | 1.23E-06 |
| CNAG_01316 | 1736.422 | 0.322372 | 0.072781 | 4.429331 | 9.45E-06 | 4.18E-05 |
| CNAG_01314 | 1237.865 | -0.56482 | 0.086024 | -6.5658 | 5.18E-11 | 4.29E-10 |
| CNAG_01313 | 630.5879 | -0.30224 | 0.105304 | -2.87015 | 0.004103 | 0.01048 |
| CNAG_07427 | 11436.94 | -0.19588 | 0.074228 | -2.63895 | 0.008316 | 0.01961 |
| CNAG_01310 | 1844.005 | 0.189684 | 0.076856 | 2.468052 | 0.013585 | 0.030302 |
| CNAG_01307 | 2994.308 | -0.72766 | 0.073912 | -9.84494 | 7.21E-23 | 1.16E-21 |
| CNAG_01306 | 557.6999 | -0.52008 | 0.107784 | -4.82517 | 1.40E-06 | 7.01E-06 |
| CNAG_01305 | 6851.738 | -0.20959 | 0.05918 | -3.5416 | 0.000398 | 0.001281 |
| CNAG_01302 | 1256.672 | 0.33551 | 0.096269 | 3.485119 | 0.000492 | 0.001552 |
| CNAG_01301 | 698.3895 | -0.41592 | 0.096159 | -4.32536 | 1.52E-05 | 6.46E-05 |
| CNAG_01299 | 175.7139 | 0.528005 | 0.214475 | 2.461851 | 0.013822 | 0.030763 |
| CNAG_01295 | 1034.21 | -0.71355 | 0.12137 | -5.87914 | 4.12E-09 | 2.83E-08 |
| CNAG_01293 | 1737.143 | 0.459354 | 0.080273 | 5.72239 | 1.05E-08 | 6.88E-08 |
| CNAG_01289 | 299.9433 | -0.34672 | 0.149567 | -2.31812 | 0.020443 | 0.043135 |
| CNAG_01288 | 1831.802 | 0.80116 | 0.071153 | 11.25968 | 2.08E-29 | 4.19E-28 |
| CNAG_01287 | 23692.78 | 0.781389 | 0.054807 | 14.25699 | 4.06E-46 | 1.28E-44 |
| CNAG_01286 | 272.7706 | -0.29343 | 0.137822 | -2.12903 | 0.033252 | 0.065951 |
| CNAG_01285 | 1950.389 | -0.1912 | 0.078348 | -2.44045 | 0.014669 | 0.032355 |
| CNAG_01284 | 2655.682 | -0.14177 | 0.068574 | -2.06745 | 0.038692 | 0.075254 |
| CNAG_01283 | 2484.651 | -0.24541 | 0.079901 | -3.07142 | 0.00213 | 0.005817 |
| CNAG_01282 | 1876.684 | -0.18943 | 0.07462 | -2.53867 | 0.011127 | 0.02547 |
| CNAG_01281 | 576.469 | -0.37843 | 0.107674 | -3.51459 | 0.00044 | 0.001407 |
| CNAG_01279 | 1564.592 | -0.44801 | 0.077333 | -5.79329 | 6.90E-09 | 4.64E-08 |
| CNAG_07428 | 2530.841 | 0.416415 | 0.067931 | 6.129953 | 8.79E-10 | 6.45E-09 |
| CNAG_01278 | 2492.259 | 0.531718 | 0.082536 | 6.442281 | 1.18E-10 | 9.43E-10 |
| CNAG_01276 | 3893.991 | 0.446782 | 0.065079 | 6.865168 | 6.64E-12 | 5.94E-11 |
| CNAG_01274 | 9016.198 | -0.13409 | 0.063442 | -2.11357 | 0.034552 | 0.068081 |
| CNAG_01273 | 9680.944 | -0.35977 | 0.065067 | -5.52926 | 3.22E-08 | 2.00E-07 |
| CNAG_01272 | 1446.564 | 2.629865 | 0.329001 | 7.993486 | 1.31E-15 | 1.50E-14 |
| CNAG_01271 | 2763.873 | -0.23543 | 0.080738 | -2.91603 | 0.003545 | 0.009188 |
| CNAG_01270 | 9444.939 | -0.18063 | 0.059024 | -3.06027 | 0.002211 | 0.006019 |
| CNAG_01268 | 44.34688 | 0.772267 | 0.341297 | 2.262744 | 0.023651 | 0.049091 |
| CNAG_01263 | 6158.607 | 0.36737 | 0.060804 | 6.041896 | 1.52E-09 | 1.09E-08 |
| CNAG_01262 | 727.4044 | -0.26572 | 0.125351 | -2.11978 | 0.034024 | 0.067153 |

| | | | | | | |
|---|---|---|---|---|---|---|
| CNAG_07430 | 1544.572 | 0.302793 | 0.086794 | 3.488629 | 0.000486 | 0.001534 |
| CNAG_06946 | 4045.942 | 0.733073 | 0.067648 | 10.83651 | 2.31E-27 | 4.35E-26 |
| CNAG_06949 | 3565.539 | -0.59263 | 0.083645 | -7.08507 | 1.39E-12 | 1.31E-11 |
| CNAG_07431 | 1497.128 | 0.331562 | 0.100419 | 3.30177 | 0.000961 | 0.002841 |
| CNAG_01257 | 1540.966 | 0.728807 | 0.139977 | 5.20661 | 1.92E-07 | 1.08E-06 |
| CNAG_01254 | 373.6897 | -0.27793 | 0.128461 | -2.16357 | 0.030497 | 0.061236 |
| CNAG_01253 | 3931.575 | -0.23031 | 0.067426 | -3.41579 | 0.000636 | 0.001961 |
| CNAG_01252 | 139.2392 | -0.66864 | 0.184218 | -3.62962 | 0.000284 | 0.000946 |
| CNAG_01251 | 2823.317 | 0.34842 | 0.065375 | 5.329537 | 9.85E-08 | 5.77E-07 |
| CNAG_01249 | 1176.658 | 0.358341 | 0.081923 | 4.374123 | 1.22E-05 | 5.26E-05 |
| CNAG_01245 | 1393.129 | -0.44635 | 0.090538 | -4.93 | 8.22E-07 | 4.27E-06 |
| CNAG_01244 | 96.9469 | 1.191573 | 0.266504 | 4.471131 | 7.78E-06 | 3.50E-05 |
| CNAG_01243 | 2336.725 | 0.390031 | 0.067113 | 5.811553 | 6.19E-09 | 4.20E-08 |
| CNAG_01242 | 3166.214 | 3.077547 | 0.07549 | 40.76755 | 0 | 0 |
| CNAG_01241 | 1130.394 | -0.29828 | 0.088586 | -3.36718 | 0.000759 | 0.002305 |
| CNAG_12453 | 240.5769 | 0.394423 | 0.158625 | 2.486511 | 0.0129 | 0.029013 |
| CNAG_12454 | 11.03374 | 1.452886 | 0.677575 | 2.144243 | 0.032013 | 0.063823 |
| CNAG_07981 | 108.0254 | 1.957868 | 0.235437 | 8.315881 | 9.11E-17 | 1.10E-15 |
| CNAG_01237 | 442.8714 | 0.316192 | 0.113853 | 2.777189 | 0.005483 | 0.013579 |
| CNAG_01235 | 7291.199 | -0.19791 | 0.066888 | -2.95888 | 0.003088 | 0.008132 |
| CNAG_01233 | 801.3455 | 0.229337 | 0.091125 | 2.516717 | 0.011845 | 0.026871 |
| CNAG_01232 | 1946.516 | 0.391454 | 0.076792 | 5.097624 | 3.44E-07 | 1.88E-06 |
| CNAG_01231 | 813.0892 | 0.660369 | 0.098321 | 6.716452 | 1.86E-11 | 1.60E-10 |
| CNAG_12456 | 58.09923 | 2.39526 | 0.326467 | 7.336902 | 2.19E-13 | 2.19E-12 |
| CNAG_01230 | 17281.4 | 1.842432 | 0.073187 | 25.17437 | 7.65E-140 | 8.88E-138 |
| CNAG_01229 | 3830.6 | 2.421271 | 0.112974 | 21.43204 | 6.72E-102 | 5.06E-100 |
| CNAG_01228 | 800.7058 | -0.68464 | 0.103317 | -6.62663 | 3.43E-11 | 2.89E-10 |
| CNAG_07432 | 252.5063 | 0.608779 | 0.156138 | 3.898973 | 9.66E-05 | 0.000355 |
| CNAG_01224 | 57444.81 | -0.15635 | 0.070883 | -2.20567 | 0.027407 | 0.055869 |
| CNAG_01223 | 542.4388 | 1.413264 | 0.160364 | 8.812855 | 1.22E-18 | 1.60E-17 |
| CNAG_01222 | 2110.153 | -0.52005 | 0.089703 | -5.79749 | 6.73E-09 | 4.53E-08 |
| CNAG_01217 | 2098.67 | -0.25401 | 0.070841 | -3.58556 | 0.000336 | 0.001103 |
| CNAG_01214 | 356.304 | 0.578273 | 0.148532 | 3.893246 | 9.89E-05 | 0.000363 |
| CNAG_01213 | 2554.216 | 0.169124 | 0.0661 | 2.558595 | 0.01051 | 0.024165 |
| CNAG_01212 | 1197.908 | 0.325497 | 0.087493 | 3.72028 | 0.000199 | 0.000686 |
| CNAG_01211 | 3635.135 | -0.22246 | 0.06279 | -3.54285 | 0.000396 | 0.001276 |
| CNAG_01208 | 1303.139 | 0.184794 | 0.086712 | 2.131128 | 0.033079 | 0.065624 |
| CNAG_07434 | 621.2933 | -0.35499 | 0.108064 | -3.28495 | 0.00102 | 0.002994 |
| CNAG_01204 | 12968.52 | -0.21226 | 0.087021 | -2.43924 | 0.014718 | 0.032437 |
| CNAG_01203 | 1229.082 | 0.313562 | 0.080868 | 3.877442 | 0.000106 | 0.000385 |
| CNAG_01200 | 622.8057 | -0.21435 | 0.108258 | -1.98002 | 0.047701 | 0.090065 |
| CNAG_01197 | 2586.471 | 0.159905 | 0.078197 | 2.044902 | 0.040865 | 0.078839 |
| CNAG_01196 | 1461.846 | -0.37252 | 0.075227 | -4.95196 | 7.35E-07 | 3.84E-06 |
| CNAG_01192 | 346.0905 | -0.31058 | 0.125513 | -2.47453 | 0.013341 | 0.029815 |
| CNAG_01188 | 958.8408 | -0.52404 | 0.096794 | -5.41398 | 6.16E-08 | 3.72E-07 |
| CNAG_01187 | 2712.875 | 0.193291 | 0.083151 | 2.324563 | 0.020095 | 0.042557 |
| CNAG_01185 | 2425.607 | -0.28046 | 0.0789 | -3.55457 | 0.000379 | 0.001227 |

| | | | | | | |
|---|---|---|---|---|---|---|
| CNAG_01182 | 8740.112 | -0.26949 | 0.057352 | -4.69885 | 2.62E-06 | 1.27E-05 |
| CNAG_01180 | 559.4952 | 0.397759 | 0.115521 | 3.443176 | 0.000575 | 0.001793 |
| CNAG_01178 | 868.9376 | -0.29942 | 0.107718 | -2.77966 | 0.005442 | 0.013493 |
| CNAG_07437 | 645.7539 | -0.42838 | 0.107125 | -3.99889 | 6.36E-05 | 0.000242 |
| CNAG_01175 | 1170.028 | -0.55385 | 0.082344 | -6.72606 | 1.74E-11 | 1.50E-10 |
| CNAG_01174 | 3683.143 | 0.522114 | 0.09428 | 5.537896 | 3.06E-08 | 1.90E-07 |
| CNAG_01173 | 1295.94 | -0.24734 | 0.095144 | -2.59968 | 0.009331 | 0.021788 |
| CNAG_01172 | 1795.519 | -0.45121 | 0.070932 | -6.36113 | 2.00E-10 | 1.58E-09 |
| CNAG_01170 | 60488.08 | -0.2647 | 0.068564 | -3.86061 | 0.000113 | 0.000409 |
| CNAG_01168 | 11464.79 | -0.18784 | 0.062196 | -3.0201 | 0.002527 | 0.006792 |
| CNAG_01167 | 2296.103 | 0.200298 | 0.066776 | 2.999562 | 0.002704 | 0.007222 |
| CNAG_01166 | 759.7753 | -0.3548 | 0.095369 | -3.72033 | 0.000199 | 0.000686 |
| CNAG_01165 | 900.7655 | -0.62257 | 0.121594 | -5.12011 | 3.05E-07 | 1.68E-06 |
| CNAG_01163 | 1071.395 | 0.485134 | 0.096451 | 5.029857 | 4.91E-07 | 2.63E-06 |
| CNAG_01162 | 905.4934 | -0.24634 | 0.08853 | -2.78257 | 0.005393 | 0.013389 |
| CNAG_07439 | 4338.571 | 0.123685 | 0.062166 | 1.989593 | 0.046636 | 0.088358 |
| CNAG_01159 | 1933.835 | -0.32445 | 0.097776 | -3.31833 | 0.000906 | 0.002689 |
| CNAG_01158 | 6399.104 | -0.13744 | 0.064267 | -2.13862 | 0.032466 | 0.064607 |
| CNAG_01156 | 3125.706 | -0.30074 | 0.063927 | -4.7045 | 2.54E-06 | 1.23E-05 |
| CNAG_01154 | 299.2748 | 0.451031 | 0.137135 | 3.288961 | 0.001006 | 0.002956 |
| CNAG_01152 | 52378.57 | -0.16518 | 0.066238 | -2.4938 | 0.012638 | 0.028463 |
| CNAG_01151 | 527.9899 | -0.67316 | 0.114203 | -5.8944 | 3.76E-09 | 2.60E-08 |
| CNAG_01150 | 10043.38 | 0.175738 | 0.055157 | 3.18613 | 0.001442 | 0.004084 |
| CNAG_01149 | 2102.857 | 0.282117 | 0.080503 | 3.504438 | 0.000458 | 0.001454 |
| CNAG_01147 | 1672.685 | -0.32912 | 0.083079 | -3.96147 | 7.45E-05 | 0.00028 |
| CNAG_01146 | 3970.788 | 0.20079 | 0.06144 | 3.268069 | 0.001083 | 0.003155 |
| CNAG_01145 | 1176.094 | -0.44703 | 0.115974 | -3.85458 | 0.000116 | 0.000419 |
| CNAG_01144 | 3335.934 | 0.304761 | 0.070864 | 4.300653 | 1.70E-05 | 7.16E-05 |
| CNAG_01143 | 1067.549 | 0.243857 | 0.083568 | 2.918085 | 0.003522 | 0.009143 |
| CNAG_01140 | 3822.884 | 0.289018 | 0.067638 | 4.272992 | 1.93E-05 | 8.01E-05 |
| CNAG_12461 | 245.3238 | 1.914247 | 0.18483 | 10.35679 | 3.90E-25 | 6.83E-24 |
| CNAG_01138 | 67505.67 | 3.841985 | 0.055919 | 68.70658 | 0 | 0 |
| CNAG_01137 | 17753.72 | 1.50746 | 0.072946 | 20.66541 | 7.10E-95 | 4.89E-93 |
| CNAG_01136 | 3448.257 | -0.28536 | 0.113598 | -2.51204 | 0.012004 | 0.02717 |
| CNAG_01133 | 501.4316 | 0.275825 | 0.107408 | 2.568004 | 0.010229 | 0.023552 |
| CNAG_01132 | 1811.254 | 0.189828 | 0.086077 | 2.205321 | 0.027432 | 0.055895 |
| CNAG_01131 | 400.1403 | 0.480029 | 0.140433 | 3.418216 | 0.00063 | 0.001948 |
| CNAG_01129 | 1711.851 | -0.38779 | 0.082349 | -4.70905 | 2.49E-06 | 1.21E-05 |
| CNAG_01128 | 1390.035 | -0.36525 | 0.082376 | -4.43395 | 9.25E-06 | 4.10E-05 |
| CNAG_01127 | 701.9703 | -0.36283 | 0.097999 | -3.70242 | 0.000214 | 0.000731 |
| CNAG_01126 | 3548.529 | 0.201344 | 0.068178 | 2.953224 | 0.003145 | 0.008264 |
| CNAG_01123 | 1510.977 | -0.16653 | 0.075155 | -2.21578 | 0.026706 | 0.05465 |
| CNAG_01121 | 92.76694 | 0.580384 | 0.248006 | 2.340198 | 0.019274 | 0.041034 |
| CNAG_01120 | 17100.03 | -0.24343 | 0.058911 | -4.13214 | 3.59E-05 | 0.000143 |
| CNAG_12466 | 117.6591 | 0.554607 | 0.214051 | 2.591012 | 0.009569 | 0.022217 |
| CNAG_01119 | 672.7811 | 0.342356 | 0.101066 | 3.38744 | 0.000705 | 0.002157 |
| CNAG_01118 | 464.7329 | 2.074244 | 0.122767 | 16.89576 | 4.83E-64 | 2.21E-62 |

| | | | | | | |
|---|---|---|---|---|---|---|
| CNAG_12469 | 12.52967 | 2.135194 | 0.646831 | 3.301008 | 0.000963 | 0.002847 |
| CNAG_01117 | 52523.4 | 0.132492 | 0.055494 | 2.387501 | 0.016963 | 0.036719 |
| CNAG_01116 | 291.2001 | -0.53271 | 0.133458 | -3.99155 | 6.56E-05 | 0.000249 |
| CNAG_01114 | 347.1643 | -0.41592 | 0.151725 | -2.74128 | 0.00612 | 0.014925 |
| CNAG_01113 | 1554.553 | -0.28664 | 0.092882 | -3.08604 | 0.002028 | 0.005577 |
| CNAG_01111 | 4578.916 | -0.1917 | 0.065286 | -2.9363 | 0.003322 | 0.008667 |
| CNAG_12470 | 38.0379 | 1.206936 | 0.418422 | 2.884495 | 0.00392 | 0.010055 |
| CNAG_12471 | 32.47593 | 2.182544 | 0.418137 | 5.219682 | 1.79E-07 | 1.02E-06 |
| CNAG_01108 | 4363.617 | 3.07931 | 0.08568 | 35.93947 | 7.39E-283 | 3.34E-280 |
| CNAG_01107 | 1767.089 | -0.17253 | 0.076989 | -2.24099 | 0.025027 | 0.051576 |
| CNAG_01106 | 3434.355 | -0.32113 | 0.07542 | -4.25789 | 2.06E-05 | 8.54E-05 |
| CNAG_01103 | 2361.144 | -0.34044 | 0.068172 | -4.99382 | 5.92E-07 | 3.13E-06 |
| CNAG_01097 | 1353.131 | -0.25897 | 0.092052 | -2.81325 | 0.004904 | 0.012274 |
| CNAG_01096 | 480.2473 | -0.36763 | 0.11039 | -3.33029 | 0.000868 | 0.002596 |
| CNAG_01094 | 2169.378 | 0.169866 | 0.067227 | 2.52673 | 0.011513 | 0.02619 |
| CNAG_01092 | 1922.535 | -0.29269 | 0.075693 | -3.86673 | 0.00011 | 0.0004 |
| CNAG_01091 | 6658.76 | -0.24248 | 0.059439 | -4.07945 | 4.51E-05 | 0.000176 |
| CNAG_01090 | 380.798 | 0.755625 | 0.148827 | 5.077201 | 3.83E-07 | 2.08E-06 |
| CNAG_01089 | 1506.425 | 0.283463 | 0.075243 | 3.76731 | 0.000165 | 0.000579 |
| CNAG_01082 | 2601.044 | 0.673825 | 0.081554 | 8.262324 | 1.43E-16 | 1.71E-15 |
| CNAG_01081 | 382.9221 | 2.031832 | 0.170674 | 11.90476 | 1.12E-32 | 2.45E-31 |
| CNAG_01080 | 233.8248 | 0.962316 | 0.152879 | 6.294643 | 3.08E-10 | 2.38E-09 |
| CNAG_01079 | 81.43918 | 1.062399 | 0.239713 | 4.431967 | 9.34E-06 | 4.13E-05 |
| CNAG_01078 | 526.7559 | 1.77733 | 0.146041 | 12.17007 | 4.49E-34 | 1.03E-32 |
| CNAG_01077 | 118.1479 | 1.994959 | 0.213509 | 9.343697 | 9.30E-21 | 1.37E-19 |
| CNAG_12476 | 17.06924 | 1.813566 | 0.561631 | 3.229107 | 0.001242 | 0.003565 |
| CNAG_01076 | 557.0657 | 3.970526 | 0.149205 | 26.61117 | 5.04E-156 | 6.94E-154 |
| CNAG_01075 | 1403.917 | 5.468708 | 0.141305 | 38.70146 | 0 | 0 |
| CNAG_01074 | 92.60523 | 1.358058 | 0.237084 | 5.728172 | 1.02E-08 | 6.67E-08 |
| CNAG_01073 | 594.9086 | 0.671135 | 0.109682 | 6.118931 | 9.42E-10 | 6.89E-09 |
| CNAG_01072 | 428.8517 | 0.333808 | 0.132469 | 2.519892 | 0.011739 | 0.02666 |
| CNAG_01070 | 33.59571 | 0.936632 | 0.372594 | 2.513814 | 0.011943 | 0.027071 |
| CNAG_01069 | 630.6072 | 0.291517 | 0.104131 | 2.799508 | 0.005118 | 0.012768 |
| CNAG_07442 | 3824.921 | 0.186556 | 0.069399 | 2.688176 | 0.007184 | 0.017215 |
| CNAG_07443 | 66.65091 | 1.266167 | 0.285114 | 4.440911 | 8.96E-06 | 3.98E-05 |
| CNAG_01065 | 345.1568 | 0.425005 | 0.126043 | 3.371907 | 0.000746 | 0.002273 |
| CNAG_01064 | 244.4291 | -0.64701 | 0.155008 | -4.17404 | 2.99E-05 | 0.00012 |
| CNAG_01063 | 3130.487 | -0.16724 | 0.074196 | -2.25403 | 0.024195 | 0.050077 |
| CNAG_01062 | 1996.646 | 0.490165 | 0.068907 | 7.113444 | 1.13E-12 | 1.08E-11 |
| CNAG_01061 | 617.1712 | 1.407611 | 0.121106 | 11.62299 | 3.15E-31 | 6.65E-30 |
| CNAG_01060 | 8694.024 | -0.61577 | 0.068176 | -9.03213 | 1.68E-19 | 2.34E-18 |
| CNAG_01059 | 314.2859 | -0.30564 | 0.129677 | -2.35691 | 0.018428 | 0.039447 |
| CNAG_01058 | 1339.235 | -0.23646 | 0.09809 | -2.41066 | 0.015924 | 0.034727 |
| CNAG_01057 | 1906.061 | -0.33109 | 0.070548 | -4.69316 | 2.69E-06 | 1.30E-05 |
| CNAG_07444 | 253.7611 | 1.075165 | 0.20559 | 5.229661 | 1.70E-07 | 9.66E-07 |
| CNAG_01055 | 12414.69 | 1.082171 | 0.065157 | 16.6087 | 6.03E-62 | 2.65E-60 |
| CNAG_01054 | 799.4585 | -0.23333 | 0.093831 | -2.4867 | 0.012893 | 0.029006 |

| | | | | | | |
|---|---|---|---|---|---|---|
| CNAG_01051 | 2088.873 | 0.18884 | 0.073197 | 2.579899 | 0.009883 | 0.022866 |
| CNAG_01049 | 1581.828 | 0.304936 | 0.096887 | 3.147337 | 0.001648 | 0.004618 |
| CNAG_01048 | 150.5836 | 0.990971 | 0.221621 | 4.471469 | 7.77E-06 | 3.50E-05 |
| CNAG_01047 | 170.6498 | -0.4442 | 0.217393 | -2.0433 | 0.041023 | 0.079069 |
| CNAG_01046 | 1991.05 | -0.43631 | 0.078488 | -5.55901 | 2.71E-08 | 1.70E-07 |
| CNAG_01044 | 1543.289 | 0.911595 | 0.103098 | 8.84205 | 9.40E-19 | 1.24E-17 |
| CNAG_01043 | 6539.098 | -0.99375 | 0.078696 | -12.6277 | 1.48E-36 | 3.70E-35 |
| CNAG_01042 | 643.7699 | -1.31387 | 0.106156 | -12.3768 | 3.49E-35 | 8.25E-34 |
| CNAG_01041 | 220.7664 | -0.92296 | 0.154706 | -5.9659 | 2.43E-09 | 1.72E-08 |
| CNAG_01040 | 134.9821 | 1.493442 | 0.213882 | 6.98255 | 2.90E-12 | 2.66E-11 |
| CNAG_12485 | 34.81236 | 0.902876 | 0.387532 | 2.329808 | 0.019816 | 0.042043 |
| CNAG_01038 | 773.6461 | -0.23682 | 0.097583 | -2.42686 | 0.01523 | 0.033432 |
| CNAG_01036 | 487.659 | -0.41735 | 0.11329 | -3.68389 | 0.00023 | 0.00078 |
| CNAG_01034 | 112.8403 | 0.44861 | 0.213035 | 2.105811 | 0.035221 | 0.069232 |
| CNAG_01032 | 376.6367 | 0.471888 | 0.133158 | 3.543808 | 0.000394 | 0.001273 |
| CNAG_01031 | 1937.739 | -0.70575 | 0.304369 | -2.31875 | 0.020409 | 0.043075 |
| CNAG_01029 | 1483.98 | -0.41512 | 0.104679 | -3.96566 | 7.32E-05 | 0.000276 |
| CNAG_01028 | 463.1188 | -0.66166 | 0.11562 | -5.72278 | 1.05E-08 | 6.87E-08 |
| CNAG_01025 | 1006.796 | -0.64088 | 0.10317 | -6.21187 | 5.24E-10 | 3.93E-09 |
| CNAG_01023 | 1229.798 | -0.37417 | 0.0914 | -4.09379 | 4.24E-05 | 0.000166 |
| CNAG_01021 | 3915.744 | -0.4384 | 0.069202 | -6.33513 | 2.37E-10 | 1.86E-09 |
| CNAG_01020 | 1468.999 | -0.33015 | 0.080759 | -4.08805 | 4.35E-05 | 0.00017 |
| CNAG_01019 | 22425.22 | -1.16815 | 0.079411 | -14.7102 | 5.54E-49 | 1.83E-47 |
| CNAG_01018 | 6977.898 | -0.12935 | 0.065565 | -1.97287 | 0.048511 | 0.091335 |
| CNAG_01015 | 786.4554 | -0.22912 | 0.102912 | -2.2264 | 0.025987 | 0.053313 |
| CNAG_01014 | 1719.407 | 0.385414 | 0.117637 | 3.276303 | 0.001052 | 0.003076 |
| CNAG_01013 | 939.7223 | 0.182314 | 0.09072 | 2.009634 | 0.04447 | 0.084922 |
| CNAG_01010 | 1908.597 | -0.65357 | 0.06915 | -9.45144 | 3.34E-21 | 5.05E-20 |
| CNAG_01009 | 989.5335 | -0.36198 | 0.09509 | -3.80672 | 0.000141 | 0.0005 |
| CNAG_01008 | 593.4608 | -0.39853 | 0.136631 | -2.91686 | 0.003536 | 0.009167 |
| CNAG_01006 | 382.3441 | -0.2666 | 0.129598 | -2.0571 | 0.039677 | 0.076894 |
| CNAG_01004 | 2283.246 | -0.39546 | 0.084421 | -4.68444 | 2.81E-06 | 1.35E-05 |
| CNAG_01002 | 548.6005 | -0.55858 | 0.112797 | -4.95213 | 7.34E-07 | 3.83E-06 |
| CNAG_01001 | 365.3954 | 0.285095 | 0.134458 | 2.120325 | 0.033979 | 0.067082 |
| CNAG_00999 | 1693.989 | 1.15991 | 0.071704 | 16.17631 | 7.41E-59 | 3.07E-57 |
| CNAG_00997 | 1234.693 | 0.760202 | 0.107092 | 7.098578 | 1.26E-12 | 1.19E-11 |
| CNAG_00994 | 1900.965 | -0.39645 | 0.083663 | -4.73869 | 2.15E-06 | 1.06E-05 |
| CNAG_00993 | 699.4328 | -0.52629 | 0.099456 | -5.29169 | 1.21E-07 | 7.03E-07 |
| CNAG_00992 | 15998.33 | 0.310512 | 0.058325 | 5.323819 | 1.02E-07 | 5.94E-07 |
| CNAG_00989 | 900.4566 | -0.44 | 0.101115 | -4.35145 | 1.35E-05 | 5.79E-05 |
| CNAG_00988 | 5173.49 | 0.257751 | 0.073955 | 3.485258 | 0.000492 | 0.001552 |
| CNAG_00987 | 1362.664 | -0.56943 | 0.079027 | -7.20545 | 5.79E-13 | 5.63E-12 |
| CNAG_00986 | 1046.844 | -0.23944 | 0.086882 | -2.75589 | 0.005853 | 0.014351 |
| CNAG_00982 | 1829.457 | -0.34194 | 0.090044 | -3.79744 | 0.000146 | 0.000518 |
| CNAG_12487 | 28.27434 | -0.91108 | 0.424702 | -2.14522 | 0.031935 | 0.063698 |
| CNAG_12488 | 113.1367 | -1.02769 | 0.207338 | -4.95658 | 7.17E-07 | 3.76E-06 |
| CNAG_00979 | 346.7778 | 0.490353 | 0.193793 | 2.530292 | 0.011397 | 0.025991 |

| | | | | | | |
|---|---|---|---|---|---|---|
| CNAG_00978 | 5868.876 | 1.045342 | 0.059979 | 17.42853 | 5.01E-68 | 2.44E-66 |
| CNAG_00977 | 1969.13 | 0.352553 | 0.076464 | 4.610682 | 4.01E-06 | 1.89E-05 |
| CNAG_00976 | 7551.328 | -0.33929 | 0.061209 | -5.54319 | 2.97E-08 | 1.85E-07 |
| CNAG_00972 | 5461.195 | 1.036695 | 0.075969 | 13.64629 | 2.12E-42 | 6.10E-41 |
| CNAG_00971 | 356.079 | -0.48115 | 0.131648 | -3.65479 | 0.000257 | 0.000863 |
| CNAG_00970 | 3732.659 | -0.13032 | 0.061704 | -2.11204 | 0.034683 | 0.068274 |
| CNAG_00969 | 189.8661 | -0.83391 | 0.158952 | -5.24632 | 1.55E-07 | 8.88E-07 |
| CNAG_00966 | 2276.502 | 0.257238 | 0.078595 | 3.272954 | 0.001064 | 0.003109 |
| CNAG_00964 | 375.3805 | 0.76054 | 0.133365 | 5.702675 | 1.18E-08 | 7.68E-08 |
| CNAG_07446 | 1454.971 | -0.16835 | 0.081058 | -2.07692 | 0.037809 | 0.073766 |
| CNAG_00961 | 852.8509 | -0.48591 | 0.163676 | -2.96874 | 0.00299 | 0.007889 |
| CNAG_00960 | 3647.439 | -0.14921 | 0.063355 | -2.3552 | 0.018513 | 0.039619 |
| CNAG_12491 | 51.63093 | 0.905416 | 0.451315 | 2.006173 | 0.044838 | 0.08551 |
| CNAG_12492 | 382.3648 | -0.30476 | 0.138394 | -2.2021 | 0.027658 | 0.0563 |
| CNAG_00941 | 113.2531 | -0.74375 | 0.207346 | -3.58701 | 0.000334 | 0.001099 |
| CNAG_00940 | 2989.848 | -0.3442 | 0.068676 | -5.01191 | 5.39E-07 | 2.87E-06 |
| CNAG_00936 | 1584.873 | 1.109779 | 0.30659 | 3.619748 | 0.000295 | 0.000978 |
| CNAG_00935 | 10297.12 | -0.59675 | 0.083566 | -7.14105 | 9.26E-13 | 8.86E-12 |
| CNAG_00934 | 107.0924 | 1.22233 | 0.24625 | 4.963776 | 6.91E-07 | 3.63E-06 |
| CNAG_00930 | 13857.36 | -0.14461 | 0.058485 | -2.4726 | 0.013413 | 0.029968 |
| CNAG_00928 | 780.6188 | 0.330392 | 0.097923 | 3.374006 | 0.000741 | 0.002259 |
| CNAG_00925 | 308.6275 | 0.6155 | 0.134863 | 4.563892 | 5.02E-06 | 2.33E-05 |
| CNAG_00923 | 487.9718 | 0.576877 | 0.114618 | 5.033048 | 4.83E-07 | 2.59E-06 |
| CNAG_00916 | 1950.661 | -0.58631 | 0.071966 | -8.14705 | 3.73E-16 | 4.36E-15 |
| CNAG_00915 | 266.2126 | 0.823741 | 0.195008 | 4.224143 | 2.40E-05 | 9.76E-05 |
| CNAG_00914 | 6039.559 | 0.410361 | 0.071429 | 5.745047 | 9.19E-09 | 6.07E-08 |
| CNAG_07448 | 565.9349 | 0.699716 | 0.13331 | 5.248788 | 1.53E-07 | 8.78E-07 |
| CNAG_12499 | 151.8747 | 3.415096 | 0.243235 | 14.04029 | 8.83E-45 | 2.75E-43 |
| CNAG_07449 | 195.367 | 0.719312 | 0.161452 | 4.455257 | 8.38E-06 | 3.75E-05 |
| CNAG_07450 | 172.0367 | 0.385859 | 0.16902 | 2.282915 | 0.022435 | 0.04683 |
| CNAG_07982 | 2389.557 | 0.656605 | 0.068093 | 9.642707 | 5.28E-22 | 8.19E-21 |
| CNAG_00909 | 936.7265 | 0.994861 | 0.101425 | 9.808833 | 1.03E-22 | 1.65E-21 |
| CNAG_00908 | 29.81876 | 0.88906 | 0.446994 | 1.988975 | 0.046704 | 0.088466 |
| CNAG_00907 | 130.9669 | 0.815992 | 0.220138 | 3.706731 | 0.00021 | 0.000719 |
| CNAG_00906 | 5178.014 | 0.18692 | 0.069417 | 2.69273 | 0.007087 | 0.016997 |
| CNAG_00905 | 511.9902 | 1.411981 | 0.186131 | 7.585948 | 3.30E-14 | 3.49E-13 |
| CNAG_00904 | 1033.311 | 0.969631 | 0.122094 | 7.941671 | 1.99E-15 | 2.25E-14 |
| CNAG_00903 | 662.8758 | -0.5129 | 0.120989 | -4.23928 | 2.24E-05 | 9.18E-05 |
| CNAG_00902 | 757.3756 | -0.55523 | 0.096189 | -5.77227 | 7.82E-09 | 5.21E-08 |
| CNAG_00897 | 2419.339 | -2.43663 | 0.095368 | -25.5497 | 5.53E-144 | 6.92E-142 |
| CNAG_00896 | 340.9561 | 0.302034 | 0.135883 | 2.222758 | 0.026232 | 0.053774 |
| CNAG_12505 | 112.2884 | 5.366901 | 0.427168 | 12.56392 | 3.33E-36 | 8.21E-35 |
| CNAG_12506 | 54.10843 | 0.827621 | 0.298583 | 2.771832 | 0.005574 | 0.013779 |
| CNAG_00895 | 1038.194 | -0.9404 | 0.096103 | -9.78538 | 1.30E-22 | 2.07E-21 |
| CNAG_00893 | 1093.848 | 0.199908 | 0.096896 | 2.063118 | 0.039101 | 0.075919 |
| CNAG_00891 | 13370.87 | 0.642402 | 0.073098 | 8.788284 | 1.52E-18 | 1.98E-17 |
| CNAG_00890 | 1750.491 | 0.175599 | 0.073345 | 2.394151 | 0.016659 | 0.036185 |

| | | | | | | |
|---|---|---|---|---|---|---|
| CNAG_00886 | 37300.14 | -0.2816 | 0.05819 | -4.83939 | 1.30E-06 | 6.56E-06 |
| CNAG_00884 | 1382.399 | 0.394122 | 0.074485 | 5.291271 | 1.21E-07 | 7.04E-07 |
| CNAG_00883 | 609.6072 | 0.427221 | 0.110403 | 3.869638 | 0.000109 | 0.000396 |
| CNAG_07453 | 2365.365 | -0.27058 | 0.075087 | -3.60354 | 0.000314 | 0.001037 |
| CNAG_07454 | 13.07649 | 2.615423 | 0.66743 | 3.918648 | 8.90E-05 | 0.00033 |
| CNAG_00879 | 2856.496 | 0.439308 | 0.09 | 4.881224 | 1.05E-06 | 5.39E-06 |
| CNAG_00877 | 1222.624 | 0.462497 | 0.088661 | 5.216482 | 1.82E-07 | 1.03E-06 |
| CNAG_00875 | 681.8511 | -0.42559 | 0.108688 | -3.91572 | 9.01E-05 | 0.000334 |
| CNAG_00873 | 424.6345 | 0.332826 | 0.122493 | 2.717112 | 0.006585 | 0.015921 |
| CNAG_00872 | 2044.24 | 0.186476 | 0.070506 | 2.644815 | 0.008174 | 0.019318 |
| CNAG_07455 | 17.59246 | 3.346372 | 0.66007 | 5.069723 | 3.98E-07 | 2.16E-06 |
| CNAG_00871 | 550.8597 | 2.443552 | 0.124852 | 19.57151 | 2.71E-85 | 1.61E-83 |
| CNAG_00870 | 75.60761 | 0.740276 | 0.319046 | 2.320283 | 0.020326 | 0.042952 |
| CNAG_12511 | 26.71083 | 2.138439 | 0.458038 | 4.66869 | 3.03E-06 | 1.45E-05 |
| CNAG_00869 | 1976.259 | 1.761523 | 0.103284 | 17.05517 | 3.20E-65 | 1.50E-63 |
| CNAG_12512 | 18.31125 | 3.144855 | 0.603027 | 5.215115 | 1.84E-07 | 1.04E-06 |
| CNAG_00867 | 83.20172 | 2.637454 | 0.290351 | 9.083686 | 1.05E-19 | 1.47E-18 |
| CNAG_07456 | 224.0475 | 0.698115 | 0.179848 | 3.881688 | 0.000104 | 0.000379 |
| CNAG_00864 | 20.88274 | -1.59441 | 0.498451 | -3.19874 | 0.00138 | 0.003934 |
| CNAG_07457 | 55.18671 | -1.13267 | 0.306465 | -3.69593 | 0.000219 | 0.000747 |
| CNAG_00863 | 101.0154 | -0.73541 | 0.215807 | -3.4077 | 0.000655 | 0.002013 |
| CNAG_00862 | 706.6025 | -0.73462 | 0.099645 | -7.37242 | 1.68E-13 | 1.69E-12 |
| CNAG_00860 | 394.9065 | 0.465827 | 0.124478 | 3.742257 | 0.000182 | 0.000633 |
| CNAG_02558 | 119.4073 | -0.7009 | 0.220059 | -3.18507 | 0.001447 | 0.004097 |
| CNAG_02555 | 1101.056 | -0.32987 | 0.082855 | -3.98127 | 6.85E-05 | 0.000259 |
| CNAG_02549 | 735.6237 | 0.220433 | 0.106447 | 2.070828 | 0.038375 | 0.074745 |
| CNAG_02548 | 3698.579 | 2.63527 | 0.220695 | 11.94079 | 7.25E-33 | 1.61E-31 |
| CNAG_10064 | 5.95209 | 3.544977 | 1.155003 | 3.069235 | 0.002146 | 0.005857 |
| CNAG_02546 | 5165.663 | 0.708721 | 0.065769 | 10.77597 | 4.47E-27 | 8.30E-26 |
| CNAG_02545 | 13569.65 | -0.29626 | 0.065262 | -4.53951 | 5.64E-06 | 2.60E-05 |
| CNAG_02543 | 1034.75 | 0.291426 | 0.08529 | 3.416879 | 0.000633 | 0.001956 |
| CNAG_02542 | 1966.361 | 0.785048 | 0.105458 | 7.44418 | 9.75E-14 | 1.00E-12 |
| CNAG_02541 | 3208.828 | 0.355128 | 0.078322 | 4.534234 | 5.78E-06 | 2.67E-05 |
| CNAG_02540 | 1021.753 | 0.569843 | 0.086315 | 6.601897 | 4.06E-11 | 3.40E-10 |
| CNAG_02539 | 1702.555 | 0.928912 | 0.083706 | 11.0973 | 1.29E-28 | 2.52E-27 |
| CNAG_12515 | 112.7204 | 0.98231 | 0.214285 | 4.584123 | 4.56E-06 | 2.13E-05 |
| CNAG_02536 | 798.4722 | -0.23256 | 0.106341 | -2.1869 | 0.02875 | 0.058202 |
| CNAG_02532 | 501.4713 | 1.23408 | 0.12058 | 10.23456 | 1.39E-24 | 2.38E-23 |
| CNAG_12516 | 107.2199 | 0.461685 | 0.221494 | 2.084416 | 0.037122 | 0.072556 |
| CNAG_02531 | 257.3539 | 0.366293 | 0.144669 | 2.531933 | 0.011344 | 0.025903 |
| CNAG_02529 | 608.1006 | -0.62925 | 0.118124 | -5.32702 | 9.98E-08 | 5.85E-07 |
| CNAG_02528 | 1125.701 | -0.46569 | 0.099244 | -4.69237 | 2.70E-06 | 1.30E-05 |
| CNAG_02527 | 5354.113 | 1.296765 | 0.092661 | 13.99477 | 1.68E-44 | 5.15E-43 |
| CNAG_12518 | 123.7797 | 0.599721 | 0.293975 | 2.04004 | 0.041346 | 0.079636 |
| CNAG_02526 | 415.6408 | 4.412258 | 0.216974 | 20.33541 | 6.25E-92 | 4.03E-90 |
| CNAG_02525 | 132.1359 | 1.201995 | 0.205447 | 5.850637 | 4.90E-09 | 3.34E-08 |
| CNAG_02523 | 957.9047 | 0.502109 | 0.091093 | 5.512073 | 3.55E-08 | 2.19E-07 |

| | | | | | | |
|---|---|---|---|---|---|---|
| CNAG_02517 | 94.32553 | 0.58883 | 0.223544 | 2.634067 | 0.008437 | 0.019871 |
| CNAG_02516 | 642.0123 | 0.634374 | 0.105108 | 6.035457 | 1.59E-09 | 1.13E-08 |
| CNAG_02515 | 519.6383 | 0.565228 | 0.126639 | 4.463305 | 8.07E-06 | 3.62E-05 |
| CNAG_07622 | 1244.495 | -0.22434 | 0.101964 | -2.20015 | 0.027797 | 0.056549 |
| CNAG_02512 | 1322.255 | 0.284673 | 0.077044 | 3.694958 | 0.00022 | 0.000749 |
| CNAG_02511 | 584.0263 | 0.205959 | 0.104287 | 1.974922 | 0.048277 | 0.090959 |
| CNAG_12522 | 132.9902 | 1.375213 | 0.308914 | 4.451768 | 8.52E-06 | 3.80E-05 |
| CNAG_02510 | 1600.064 | 0.661073 | 0.102638 | 6.44084 | 1.19E-10 | 9.51E-10 |
| CNAG_02508 | 305.4746 | 0.326281 | 0.145276 | 2.245945 | 0.024707 | 0.051009 |
| CNAG_02507 | 4682.186 | -0.1914 | 0.079736 | -2.40041 | 0.016377 | 0.035638 |
| CNAG_02503 | 1495.369 | -0.17527 | 0.076356 | -2.29549 | 0.021705 | 0.045504 |
| CNAG_02502 | 4326.722 | 0.156879 | 0.061265 | 2.560667 | 0.010447 | 0.024035 |
| CNAG_02501 | 1077.942 | -0.31164 | 0.102135 | -3.05127 | 0.002279 | 0.006188 |
| CNAG_02500 | 11808.25 | 0.708851 | 0.069547 | 10.19236 | 2.14E-24 | 3.65E-23 |
| CNAG_02497 | 160.8685 | 0.671769 | 0.208936 | 3.215183 | 0.001304 | 0.003734 |
| CNAG_02496 | 548.1451 | -0.2932 | 0.116083 | -2.52583 | 0.011543 | 0.026243 |
| CNAG_02495 | 2251.785 | -0.32416 | 0.076261 | -4.25074 | 2.13E-05 | 8.79E-05 |
| CNAG_12525 | 22.45864 | 0.99119 | 0.473005 | 2.095516 | 0.036125 | 0.070821 |
| CNAG_02492 | 877.1868 | -0.25327 | 0.099423 | -2.54742 | 0.010852 | 0.0249 |
| CNAG_02490 | 969.4046 | 0.3093 | 0.089096 | 3.471527 | 0.000518 | 0.001625 |
| CNAG_02488 | 532.0962 | 1.524336 | 0.129254 | 11.7933 | 4.23E-32 | 9.09E-31 |
| CNAG_02487 | 987.6137 | -0.57497 | 0.11747 | -4.89464 | 9.85E-07 | 5.05E-06 |
| CNAG_02486 | 4658.327 | -0.18237 | 0.060498 | -3.01454 | 0.002574 | 0.006904 |
| CNAG_02485 | 16804.44 | -0.34171 | 0.071475 | -4.78082 | 1.75E-06 | 8.67E-06 |
| CNAG_02481 | 2312.506 | 0.159888 | 0.073093 | 2.187467 | 0.028708 | 0.058132 |
| CNAG_02479 | 508.4982 | 0.923975 | 0.115105 | 8.027212 | 9.97E-16 | 1.15E-14 |
| CNAG_12528 | 26.5949 | 0.866241 | 0.413599 | 2.094396 | 0.036225 | 0.070999 |
| CNAG_02478 | 1083.409 | 1.486347 | 0.092238 | 16.11434 | 2.02E-58 | 8.18E-57 |
| CNAG_02476 | 444.2318 | 1.102332 | 0.119161 | 9.250752 | 2.23E-20 | 3.23E-19 |
| CNAG_02475 | 881.9709 | 0.509023 | 0.09868 | 5.158304 | 2.49E-07 | 1.38E-06 |
| CNAG_07626 | 391.3638 | 1.532161 | 0.124522 | 12.3043 | 8.59E-35 | 2.00E-33 |
| CNAG_02469 | 1682.086 | 0.428107 | 0.095793 | 4.469095 | 7.86E-06 | 3.53E-05 |
| CNAG_02466 | 421.6873 | 0.487297 | 0.110944 | 4.392257 | 1.12E-05 | 4.88E-05 |
| CNAG_02464 | 835.6742 | 0.429167 | 0.100305 | 4.278598 | 1.88E-05 | 7.83E-05 |
| CNAG_02463 | 186.411 | 1.018076 | 0.182889 | 5.566634 | 2.60E-08 | 1.63E-07 |
| CNAG_07627 | 1229.753 | -0.24315 | 0.087495 | -2.77896 | 0.005453 | 0.013518 |
| CNAG_02458 | 2456.966 | 0.352773 | 0.074658 | 4.725184 | 2.30E-06 | 1.12E-05 |
| CNAG_02457 | 5511.929 | 0.207105 | 0.060629 | 3.415939 | 0.000636 | 0.001961 |
| CNAG_07628 | 12246.84 | -0.19236 | 0.068621 | -2.80326 | 0.005059 | 0.012636 |
| CNAG_12536 | 14.36655 | 2.434539 | 0.700211 | 3.476868 | 0.000507 | 0.001595 |
| CNAG_02455 | 1004.306 | 0.714144 | 0.114014 | 6.263623 | 3.76E-10 | 2.87E-09 |
| CNAG_02454 | 1801.946 | 0.419962 | 0.074795 | 5.614869 | 1.97E-08 | 1.25E-07 |
| CNAG_02452 | 152.1631 | -0.43487 | 0.181882 | -2.39095 | 0.016805 | 0.036435 |
| CNAG_02445 | 2202.135 | -0.33912 | 0.081129 | -4.17998 | 2.92E-05 | 0.000117 |
| CNAG_02444 | 364.1082 | -0.6856 | 0.124773 | -5.49476 | 3.91E-08 | 2.40E-07 |
| CNAG_02443 | 3683.578 | 0.507226 | 0.102699 | 4.938965 | 7.85E-07 | 4.09E-06 |
| CNAG_02440 | 3642.231 | 0.620873 | 0.066376 | 9.353875 | 8.45E-21 | 1.25E-19 |

| | | | | | | |
|---|---|---|---|---|---|---|
| CNAG_02438 | 162.1094 | -0.80998 | 0.194912 | -4.15563 | 3.24E-05 | 0.00013 |
| CNAG_02437 | 1869.034 | 0.443735 | 0.075384 | 5.886303 | 3.95E-09 | 2.72E-08 |
| CNAG_02435 | 3283.171 | -0.29929 | 0.080482 | -3.71873 | 0.0002 | 0.000689 |
| CNAG_02434 | 1685.226 | 0.602218 | 0.092294 | 6.524982 | 6.80E-11 | 5.58E-10 |
| CNAG_02432 | 964.5717 | 0.380716 | 0.092434 | 4.118781 | 3.81E-05 | 0.000151 |
| CNAG_02431 | 808.8768 | -0.69365 | 0.092769 | -7.47719 | 7.59E-14 | 7.87E-13 |
| CNAG_02429 | 81.94989 | -1.52014 | 0.28246 | -5.38178 | 7.38E-08 | 4.41E-07 |
| CNAG_02427 | 460.786 | -0.31292 | 0.122977 | -2.54456 | 0.010942 | 0.025073 |
| CNAG_02426 | 4379.529 | -0.30042 | 0.068587 | -4.3801 | 1.19E-05 | 5.13E-05 |
| CNAG_02425 | 284.862 | -0.58066 | 0.140945 | -4.11976 | 3.79E-05 | 0.00015 |
| CNAG_02422 | 1161.089 | -0.51729 | 0.080757 | -6.40542 | 1.50E-10 | 1.19E-09 |
| CNAG_02421 | 4555.462 | -0.15458 | 0.065483 | -2.36065 | 0.018243 | 0.039113 |
| CNAG_02420 | 70.05482 | 0.74521 | 0.268605 | 2.774372 | 0.005531 | 0.013681 |
| CNAG_02418 | 10505.16 | -0.14909 | 0.060699 | -2.45629 | 0.014038 | 0.031166 |
| CNAG_02417 | 141.3121 | -0.56246 | 0.215933 | -2.60477 | 0.009194 | 0.021507 |
| CNAG_12541 | 182.7254 | -0.50857 | 0.172524 | -2.94783 | 0.0032 | 0.00839 |
| CNAG_02415 | 1782.168 | 0.562026 | 0.077821 | 7.222024 | 5.12E-13 | 5.03E-12 |
| CNAG_02413 | 3567.284 | -0.64411 | 0.069987 | -9.20333 | 3.47E-20 | 4.95E-19 |
| CNAG_07984 | 17.24001 | -1.50098 | 0.545791 | -2.75011 | 0.005958 | 0.014576 |
| CNAG_12543 | 41.01777 | -1.20673 | 0.400445 | -3.01348 | 0.002583 | 0.006924 |
| CNAG_02409 | 658.5635 | 0.560079 | 0.102042 | 5.488728 | 4.05E-08 | 2.48E-07 |
| CNAG_02407 | 1730.354 | 0.367961 | 0.085123 | 4.3227 | 1.54E-05 | 6.53E-05 |
| CNAG_02405 | 662.5594 | 0.787129 | 0.116953 | 6.730296 | 1.69E-11 | 1.46E-10 |
| CNAG_02404 | 8525.972 | 0.451476 | 0.080529 | 5.60636 | 2.07E-08 | 1.31E-07 |
| CNAG_02403 | 3824.781 | 0.265439 | 0.062758 | 4.229594 | 2.34E-05 | 9.55E-05 |
| CNAG_02402 | 959.7893 | -0.38284 | 0.102911 | -3.72012 | 0.000199 | 0.000686 |
| CNAG_02400 | 2547.021 | 0.147982 | 0.06682 | 2.214641 | 0.026785 | 0.054796 |
| CNAG_02397 | 355.3747 | 0.347311 | 0.129414 | 2.683725 | 0.007281 | 0.017426 |
| CNAG_02387 | 1850.033 | 0.954118 | 0.084587 | 11.27977 | 1.65E-29 | 3.34E-28 |
| CNAG_02386 | 1013.712 | 0.655448 | 0.094241 | 6.955051 | 3.52E-12 | 3.20E-11 |
| CNAG_02385 | 1023.462 | 0.472036 | 0.090123 | 5.237668 | 1.63E-07 | 9.26E-07 |
| CNAG_02383 | 572.7471 | -0.29879 | 0.128949 | -2.31712 | 0.020497 | 0.043228 |
| CNAG_07985 | 226.8192 | 0.411336 | 0.150593 | 2.731436 | 0.006306 | 0.015327 |
| CNAG_07986 | 792.4544 | 0.212938 | 0.090706 | 2.347561 | 0.018897 | 0.040345 |
| CNAG_02378 | 8614.322 | -0.30677 | 0.066297 | -4.62712 | 3.71E-06 | 1.76E-05 |
| CNAG_02377 | 1655.562 | -0.3441 | 0.080941 | -4.25128 | 2.13E-05 | 8.77E-05 |
| CNAG_02376 | 1302.189 | -0.15936 | 0.078397 | -2.03269 | 0.042084 | 0.080883 |
| CNAG_02373 | 1161.431 | -0.21395 | 0.08739 | -2.44824 | 0.014356 | 0.031785 |
| CNAG_02372 | 1983.983 | -0.407 | 0.06904 | -5.89517 | 3.74E-09 | 2.59E-08 |
| CNAG_12551 | 27.67231 | 1.674737 | 0.431141 | 3.884427 | 0.000103 | 0.000375 |
| CNAG_02368 | 1320.19 | -0.21736 | 0.081963 | -2.65198 | 0.008002 | 0.018952 |
| CNAG_02367 | 6304.017 | -0.15401 | 0.057251 | -2.69006 | 0.007144 | 0.017124 |
| CNAG_02366 | 1246.524 | 0.380043 | 0.085535 | 4.443113 | 8.87E-06 | 3.95E-05 |
| CNAG_02365 | 828.7584 | 2.72287 | 0.108759 | 25.03586 | 2.49E-138 | 2.77E-136 |
| CNAG_02362 | 650.5263 | -0.22201 | 0.111089 | -1.99846 | 0.045667 | 0.086847 |
| CNAG_12554 | 123.4374 | 0.478001 | 0.191069 | 2.501721 | 0.012359 | 0.027881 |
| CNAG_02361 | 1601.361 | 0.312584 | 0.080638 | 3.876383 | 0.000106 | 0.000386 |

| | | | | | | |
|---|---|---|---|---|---|---|
| CNAG_02358 | 370.7217 | -0.58474 | 0.12797 | -4.56936 | 4.89E-06 | 2.28E-05 |
| CNAG_02357 | 968.6082 | 0.228814 | 0.088998 | 2.570993 | 0.010141 | 0.023383 |
| CNAG_02356 | 1293.741 | -0.17134 | 0.087025 | -1.96883 | 0.048972 | 0.092034 |
| CNAG_02353 | 372.259 | 0.731939 | 0.129998 | 5.630383 | 1.80E-08 | 1.15E-07 |
| CNAG_07634 | 2237.043 | 0.479492 | 0.088077 | 5.444032 | 5.21E-08 | 3.15E-07 |
| CNAG_02351 | 2679.591 | 0.621234 | 0.088281 | 7.037038 | 1.96E-12 | 1.84E-11 |
| CNAG_07987 | 472.011 | -0.39674 | 0.114936 | -3.45187 | 0.000557 | 0.00174 |
| CNAG_02345 | 580.7666 | -0.29807 | 0.108765 | -2.7405 | 0.006135 | 0.014946 |
| CNAG_12557 | 56.34422 | 5.155652 | 0.589345 | 8.748108 | 2.17E-18 | 2.82E-17 |
| CNAG_02343 | 2393.029 | 1.492089 | 0.06793 | 21.96512 | 6.21E-107 | 4.95E-105 |
| CNAG_02342 | 533.6535 | 0.483851 | 0.117207 | 4.128161 | 3.66E-05 | 0.000145 |
| CNAG_02340 | 626.369 | -0.24761 | 0.101177 | -2.44725 | 0.014395 | 0.031853 |
| CNAG_02338 | 24243.49 | -0.20451 | 0.073203 | -2.79367 | 0.005211 | 0.012981 |
| CNAG_02336 | 638.276 | 0.668878 | 0.102575 | 6.520853 | 6.99E-11 | 5.72E-10 |
| CNAG_02335 | 8196.066 | -0.53412 | 0.09769 | -5.46751 | 4.56E-08 | 2.78E-07 |
| CNAG_12558 | 173.3537 | -0.72939 | 0.219014 | -3.33034 | 0.000867 | 0.002596 |
| CNAG_02333 | 425.4437 | -0.38195 | 0.124268 | -3.07362 | 0.002115 | 0.005784 |
| CNAG_02332 | 1119.977 | -0.46516 | 0.085168 | -5.46169 | 4.72E-08 | 2.87E-07 |
| CNAG_02331 | 28831.43 | -0.31182 | 0.069046 | -4.51619 | 6.30E-06 | 2.88E-05 |
| CNAG_02330 | 43423.93 | -0.17785 | 0.07497 | -2.37226 | 0.01768 | 0.038066 |
| CNAG_02328 | 4667.413 | 0.256799 | 0.063198 | 4.06343 | 4.84E-05 | 0.000188 |
| CNAG_02327 | 521.2414 | -0.23992 | 0.103354 | -2.32135 | 0.020268 | 0.042878 |
| CNAG_02326 | 4083.647 | -0.24182 | 0.064682 | -3.73864 | 0.000185 | 0.000642 |
| CNAG_02325 | 793.6832 | -0.2366 | 0.090129 | -2.6251 | 0.008662 | 0.020361 |
| CNAG_02324 | 952.0468 | -0.23897 | 0.100149 | -2.38609 | 0.017028 | 0.03683 |
| CNAG_02323 | 2369.601 | -0.51399 | 0.074146 | -6.93212 | 4.15E-12 | 3.76E-11 |
| CNAG_12560 | 22.48223 | -2.37188 | 0.552286 | -4.29466 | 1.75E-05 | 7.34E-05 |
| CNAG_02322 | 948.6456 | -0.33303 | 0.08831 | -3.77116 | 0.000162 | 0.000571 |
| CNAG_02318 | 890.6368 | 0.461544 | 0.098727 | 4.674957 | 2.94E-06 | 1.41E-05 |
| CNAG_02317 | 2509.293 | 0.659798 | 0.069396 | 9.507778 | 1.95E-21 | 2.98E-20 |
| CNAG_02316 | 1744.714 | -0.30349 | 0.1079 | -2.81267 | 0.004913 | 0.012291 |
| CNAG_02315 | 21494.39 | 0.571614 | 0.0722 | 7.917103 | 2.43E-15 | 2.74E-14 |
| CNAG_02314 | 149.981 | -0.78094 | 0.178208 | -4.38221 | 1.17E-05 | 5.09E-05 |
| CNAG_02312 | 919.2693 | 0.238664 | 0.094104 | 2.536158 | 0.011208 | 0.025639 |
| CNAG_02310 | 1823.443 | -0.28025 | 0.078412 | -3.57411 | 0.000351 | 0.001148 |
| CNAG_02305 | 857.3013 | 1.220867 | 0.111097 | 10.98921 | 4.31E-28 | 8.30E-27 |
| CNAG_02304 | 181.9371 | -0.57536 | 0.182455 | -3.15346 | 0.001613 | 0.004533 |
| CNAG_02302 | 296.1312 | 0.693478 | 0.130056 | 5.332147 | 9.71E-08 | 5.70E-07 |
| CNAG_02301 | 10414.37 | -0.15578 | 0.071874 | -2.16742 | 0.030203 | 0.06075 |
| CNAG_02300 | 11759.61 | -0.56209 | 0.062067 | -9.05628 | 1.35E-19 | 1.88E-18 |
| CNAG_02299 | 1073.853 | 0.182267 | 0.08507 | 2.142566 | 0.032148 | 0.06406 |
| CNAG_02294 | 4675.407 | -0.20967 | 0.062386 | -3.36078 | 0.000777 | 0.002351 |
| CNAG_02292 | 2671.25 | -0.74258 | 0.079221 | -9.37347 | 7.02E-21 | 1.04E-19 |
| CNAG_02289 | 468.4094 | -0.23124 | 0.112949 | -2.04728 | 0.040631 | 0.078499 |
| CNAG_02287 | 746.8986 | -0.33453 | 0.103392 | -3.23551 | 0.001214 | 0.003499 |
| CNAG_02286 | 2282.794 | -0.14553 | 0.069167 | -2.10401 | 0.035378 | 0.069476 |
| CNAG_02285 | 27123.64 | 0.240754 | 0.103764 | 2.320212 | 0.020329 | 0.042952 |

| | | | | | | |
|---|---|---|---|---|---|---|
| CNAG_02284 | 235.8217 | -0.562 | 0.14671 | -3.83071 | 0.000128 | 0.000457 |
| CNAG_02282 | 1253.563 | -0.29164 | 0.091069 | -3.20247 | 0.001363 | 0.003886 |
| CNAG_02281 | 1037.512 | -0.30769 | 0.091896 | -3.34821 | 0.000813 | 0.002446 |
| CNAG_02280 | 1138.219 | -0.24838 | 0.110842 | -2.24087 | 0.025034 | 0.051579 |
| CNAG_07635 | 1778.458 | -0.54617 | 0.070358 | -7.76279 | 8.31E-15 | 9.03E-14 |
| CNAG_07638 | 1663.493 | 0.855541 | 0.072743 | 11.7612 | 6.19E-32 | 1.32E-30 |
| CNAG_07640 | 662.9757 | -0.27756 | 0.109727 | -2.52957 | 0.01142 | 0.02603 |
| CNAG_02270 | 1572.568 | -0.72004 | 0.081812 | -8.80114 | 1.35E-18 | 1.77E-17 |
| CNAG_02269 | 686.0902 | -0.45992 | 0.11236 | -4.09326 | 4.25E-05 | 0.000167 |
| CNAG_02268 | 469.1995 | -0.25168 | 0.118856 | -2.1175 | 0.034217 | 0.067488 |
| CNAG_12567 | 34.6486 | -1.49065 | 0.377466 | -3.94909 | 7.84E-05 | 0.000294 |
| CNAG_02267 | 1649.019 | -0.18284 | 0.075072 | -2.43554 | 0.014869 | 0.032749 |
| CNAG_07164 | 610.9024 | -0.30955 | 0.106036 | -2.9193 | 0.003508 | 0.009119 |
| CNAG_07163 | 2074.541 | 0.841797 | 0.075082 | 11.21172 | 3.57E-29 | 7.15E-28 |
| CNAG_02266 | 6788.835 | 0.893063 | 0.075184 | 11.87835 | 1.53E-32 | 3.34E-31 |
| CNAG_02265 | 3889.434 | 0.180001 | 0.063912 | 2.816399 | 0.004857 | 0.012176 |
| CNAG_02264 | 2280.555 | 1.818539 | 0.105457 | 17.24433 | 1.23E-66 | 5.90E-65 |
| CNAG_02263 | 405.4538 | -0.46496 | 0.133001 | -3.49594 | 0.000472 | 0.001498 |
| CNAG_02262 | 3134.535 | 0.470026 | 0.073748 | 6.373388 | 1.85E-10 | 1.46E-09 |
| CNAG_02260 | 2258.387 | -0.29791 | 0.070274 | -4.23934 | 2.24E-05 | 9.18E-05 |
| CNAG_02259 | 1132.402 | 0.317484 | 0.093161 | 3.407919 | 0.000655 | 0.002012 |
| CNAG_02258 | 1476.611 | -0.46334 | 0.07337 | -6.31519 | 2.70E-10 | 2.10E-09 |
| CNAG_02257 | 19369.72 | -0.12909 | 0.059594 | -2.16616 | 0.030299 | 0.060913 |
| CNAG_12570 | 74.14115 | 0.626969 | 0.257747 | 2.432501 | 0.014995 | 0.032967 |
| CNAG_12571 | 62.69955 | 1.617559 | 0.316061 | 5.117873 | 3.09E-07 | 1.70E-06 |
| CNAG_02240 | 2579.249 | -0.19746 | 0.067201 | -2.93839 | 0.003299 | 0.008617 |
| CNAG_02237 | 4580.069 | -0.28309 | 0.079013 | -3.58284 | 0.00034 | 0.001114 |
| CNAG_02236 | 2231.945 | -0.48534 | 0.08084 | -6.00379 | 1.93E-09 | 1.37E-08 |
| CNAG_02234 | 54647.86 | -0.17455 | 0.063114 | -2.76557 | 0.005682 | 0.014004 |
| CNAG_02232 | 2064.647 | -0.32226 | 0.0727 | -4.43278 | 9.30E-06 | 4.12E-05 |
| CNAG_07646 | 192.9672 | -0.63822 | 0.177311 | -3.59942 | 0.000319 | 0.001052 |
| CNAG_02226 | 290.7281 | 1.295351 | 0.192703 | 6.72202 | 1.79E-11 | 1.54E-10 |
| CNAG_02220 | 2138.508 | 0.555004 | 0.088675 | 6.258879 | 3.88E-10 | 2.95E-09 |
| CNAG_02219 | 401.4697 | -0.32794 | 0.130585 | -2.5113 | 0.012029 | 0.027211 |
| CNAG_02217 | 248.0087 | 0.881595 | 0.179727 | 4.905192 | 9.33E-07 | 4.80E-06 |
| CNAG_02215 | 904.7927 | 0.577362 | 0.09027 | 6.395963 | 1.60E-10 | 1.26E-09 |
| CNAG_02214 | 471.1354 | 0.99057 | 0.126748 | 7.815288 | 5.48E-15 | 6.03E-14 |
| CNAG_02213 | 2320.999 | -0.25379 | 0.074219 | -3.41955 | 0.000627 | 0.001943 |
| CNAG_02212 | 14265.89 | -0.16704 | 0.067426 | -2.47739 | 0.013235 | 0.029626 |
| CNAG_02210 | 889.3993 | -0.19471 | 0.090668 | -2.14745 | 0.031758 | 0.063437 |
| CNAG_02209 | 9774.168 | -0.13385 | 0.066818 | -2.00322 | 0.045153 | 0.086011 |
| CNAG_02208 | 6309.214 | 0.493114 | 0.064058 | 7.697913 | 1.38E-14 | 1.49E-13 |
| CNAG_02205 | 378.5779 | -0.59425 | 0.13694 | -4.3395 | 1.43E-05 | 6.10E-05 |
| CNAG_02204 | 1858.213 | 0.36369 | 0.074193 | 4.901955 | 9.49E-07 | 4.88E-06 |
| CNAG_02203 | 1351.173 | 0.264511 | 0.081921 | 3.228872 | 0.001243 | 0.003567 |
| CNAG_02202 | 4318.931 | -0.25438 | 0.067199 | -3.7854 | 0.000153 | 0.000542 |
| CNAG_02201 | 402.8127 | 0.495608 | 0.165727 | 2.990511 | 0.002785 | 0.007415 |

| | | | | | | |
|---|---|---|---|---|---|---|
| CNAG_02198 | 530.0916 | -0.37607 | 0.103843 | -3.62157 | 0.000293 | 0.000971 |
| CNAG_02197 | 1975.271 | -0.23779 | 0.07844 | -3.03145 | 0.002434 | 0.006572 |
| CNAG_02193 | 719.9626 | -0.48156 | 0.109591 | -4.39417 | 1.11E-05 | 4.84E-05 |
| CNAG_02192 | 43.79218 | 0.660569 | 0.332463 | 1.986895 | 0.046934 | 0.08884 |
| CNAG_02190 | 897.7692 | 0.90834 | 0.099621 | 9.117917 | 7.66E-20 | 1.08E-18 |
| CNAG_02189 | 4139.646 | -0.17899 | 0.08831 | -2.0268 | 0.042683 | 0.081862 |
| CNAG_02187 | 785.0951 | -0.21535 | 0.092421 | -2.33005 | 0.019803 | 0.042026 |
| CNAG_02183 | 977.0796 | 0.18991 | 0.08772 | 2.16496 | 0.030391 | 0.061067 |
| CNAG_02182 | 4996.153 | 2.465408 | 0.106585 | 23.13095 | 2.26E-118 | 1.93E-116 |
| CNAG_02181 | 1008.951 | -0.49593 | 0.085462 | -5.80295 | 6.52E-09 | 4.39E-08 |
| CNAG_02180 | 2752.141 | 0.146875 | 0.073652 | 1.994175 | 0.046133 | 0.087548 |
| CNAG_02179 | 1535.918 | -0.37298 | 0.079795 | -4.6742 | 2.95E-06 | 1.42E-05 |
| CNAG_02177 | 6125.277 | -0.40906 | 0.062196 | -6.57694 | 4.80E-11 | 3.98E-10 |
| CNAG_02176 | 1828.888 | -0.36238 | 0.089855 | -4.03293 | 5.51E-05 | 0.000212 |
| CNAG_02175 | 2207.645 | -0.20454 | 0.079114 | -2.58533 | 0.009729 | 0.022541 |
| CNAG_02174 | 4590.067 | -0.36617 | 0.064377 | -5.68789 | 1.29E-08 | 8.33E-08 |
| CNAG_02171 | 908.9165 | -0.2255 | 0.087159 | -2.58717 | 0.009677 | 0.02244 |
| CNAG_02168 | 1718.291 | 0.295349 | 0.09928 | 2.974919 | 0.002931 | 0.007754 |
| CNAG_02166 | 7743.14 | -0.23434 | 0.057834 | -4.05193 | 5.08E-05 | 0.000196 |
| CNAG_02165 | 1102.655 | -0.65376 | 0.106287 | -6.1509 | 7.70E-10 | 5.70E-09 |
| CNAG_02164 | 1125.922 | 0.433311 | 0.113753 | 3.809238 | 0.000139 | 0.000495 |
| CNAG_02163 | 394.2006 | -0.52015 | 0.144214 | -3.60679 | 0.00031 | 0.001025 |
| CNAG_02161 | 2479.829 | -0.15743 | 0.066748 | -2.35864 | 0.018342 | 0.039305 |
| CNAG_02156 | 556.8918 | 1.293404 | 0.12506 | 10.34223 | 4.54E-25 | 7.93E-24 |
| CNAG_02154 | 825.0226 | -0.56992 | 0.101212 | -5.63093 | 1.79E-08 | 1.14E-07 |
| CNAG_02153 | 2344.987 | -0.52899 | 0.087122 | -6.07182 | 1.26E-09 | 9.19E-09 |
| CNAG_02152 | 936.1077 | 0.933541 | 0.09445 | 9.884001 | 4.88E-23 | 7.99E-22 |
| CNAG_02151 | 1436.805 | -0.37281 | 0.099195 | -3.75838 | 0.000171 | 0.000598 |
| CNAG_02147 | 842.367 | 2.089229 | 0.099306 | 21.03834 | 2.92E-98 | 2.09E-96 |
| CNAG_02143 | 778.6342 | 0.230126 | 0.111073 | 2.071847 | 0.03828 | 0.074595 |
| CNAG_02141 | 516.5441 | 0.263177 | 0.110994 | 2.37108 | 0.017736 | 0.038168 |
| CNAG_02140 | 6646.148 | 0.975164 | 0.069665 | 13.99783 | 1.61E-44 | 4.95E-43 |
| CNAG_02139 | 2745.232 | -0.67433 | 0.073193 | -9.21312 | 3.17E-20 | 4.54E-19 |
| CNAG_02138 | 371.2661 | 0.556219 | 0.14646 | 3.797742 | 0.000146 | 0.000517 |
| CNAG_12586 | 65.64317 | 0.698802 | 0.291925 | 2.393775 | 0.016676 | 0.036212 |
| CNAG_02134 | 3098.066 | -0.47462 | 0.082181 | -5.77533 | 7.68E-09 | 5.12E-08 |
| CNAG_02133 | 5403.462 | -0.34233 | 0.058173 | -5.88466 | 3.99E-09 | 2.75E-08 |
| CNAG_02131 | 1984.337 | 1.504181 | 0.075234 | 19.99334 | 6.29E-89 | 3.97E-87 |
| CNAG_02130 | 5279.926 | -0.33588 | 0.078792 | -4.26292 | 2.02E-05 | 8.36E-05 |
| CNAG_02128 | 10016 | -0.34751 | 0.060787 | -5.71692 | 1.08E-08 | 7.09E-08 |
| CNAG_02127 | 712.0854 | -0.61068 | 0.092147 | -6.62731 | 3.42E-11 | 2.88E-10 |
| CNAG_02124 | 899.9151 | 0.310934 | 0.09781 | 3.178973 | 0.001478 | 0.004177 |
| CNAG_02123 | 1388.113 | 0.775077 | 0.090864 | 8.530079 | 1.46E-17 | 1.83E-16 |
| CNAG_02121 | 1027.681 | -0.25391 | 0.091968 | -2.76086 | 0.005765 | 0.014174 |
| CNAG_02120 | 4972.992 | 1.254182 | 0.063126 | 19.86789 | 7.72E-88 | 4.83E-86 |
| CNAG_02119 | 814.6334 | -0.37655 | 0.089182 | -4.22224 | 2.42E-05 | 9.84E-05 |
| CNAG_02117 | 916.1622 | -0.44967 | 0.095987 | -4.68463 | 2.80E-06 | 1.35E-05 |

| | | | | | | |
|---|---|---|---|---|---|---|
| CNAG_02115 | 1652.054 | -0.89429 | 0.07736 | -11.5601 | 6.57E-31 | 1.38E-29 |
| CNAG_02114 | 2752.276 | 0.533714 | 0.078612 | 6.789206 | 1.13E-11 | 9.91E-11 |
| CNAG_02113 | 1739.393 | -0.2686 | 0.081983 | -3.27627 | 0.001052 | 0.003076 |
| CNAG_02111 | 1151.189 | -0.19846 | 0.081242 | -2.44278 | 0.014575 | 0.032182 |
| CNAG_02110 | 3014.102 | -0.18564 | 0.071884 | -2.58246 | 0.00981 | 0.02271 |
| CNAG_02109 | 1838.329 | 0.208678 | 0.088903 | 2.347264 | 0.018912 | 0.040367 |
| CNAG_02107 | 937.1705 | 1.165204 | 0.105041 | 11.09281 | 1.36E-28 | 2.64E-27 |
| CNAG_02105 | 185.9056 | -0.92715 | 0.185412 | -5.0005 | 5.72E-07 | 3.04E-06 |
| CNAG_02104 | 2317.213 | 2.553399 | 0.076327 | 33.45342 | 2.30E-245 | 7.46E-243 |
| CNAG_12593 | 153.7627 | 1.514446 | 0.210071 | 7.209201 | 5.63E-13 | 5.50E-12 |
| CNAG_02103 | 836.1567 | 0.568935 | 0.101878 | 5.584495 | 2.34E-08 | 1.47E-07 |
| CNAG_02100 | 13571.52 | 0.147563 | 0.066154 | 2.230602 | 0.025708 | 0.052805 |
| CNAG_02099 | 13953.42 | 0.149251 | 0.065243 | 2.287614 | 0.02216 | 0.046374 |
| CNAG_02096 | 270.5046 | -0.36766 | 0.135704 | -2.70929 | 0.006743 | 0.016268 |
| CNAG_02094 | 2176.524 | -0.35423 | 0.081321 | -4.35592 | 1.33E-05 | 5.68E-05 |
| CNAG_02093 | 975.0277 | 0.601651 | 0.095329 | 6.311302 | 2.77E-10 | 2.15E-09 |
| CNAG_02091 | 14558.67 | -0.12398 | 0.060545 | -2.04779 | 0.04058 | 0.078457 |
| CNAG_02089 | 547.5381 | 0.397727 | 0.138067 | 2.880692 | 0.003968 | 0.010152 |
| CNAG_02088 | 1101.882 | -0.78899 | 0.089201 | -8.84501 | 9.15E-19 | 1.21E-17 |
| CNAG_02084 | 4169.406 | -0.45284 | 0.067501 | -6.70867 | 1.96E-11 | 1.68E-10 |
| CNAG_02083 | 1843.461 | 3.650457 | 0.08736 | 41.7862 | 0 | 0 |
| CNAG_12597 | 100.5393 | 1.030305 | 0.23634 | 4.359421 | 1.30E-05 | 5.60E-05 |
| CNAG_12598 | 94.77128 | 1.531732 | 0.233576 | 6.557746 | 5.46E-11 | 4.52E-10 |
| CNAG_02079 | 6590.19 | -1.18816 | 0.096032 | -12.3725 | 3.68E-35 | 8.67E-34 |
| CNAG_02077 | 547.2243 | -0.34897 | 0.103557 | -3.36984 | 0.000752 | 0.002288 |
| CNAG_02076 | 1487.989 | -0.52803 | 0.085429 | -6.18093 | 6.37E-10 | 4.76E-09 |
| CNAG_02071 | 2644.508 | 0.327048 | 0.086652 | 3.774275 | 0.00016 | 0.000565 |
| CNAG_02067 | 2846.841 | 0.613433 | 0.07179 | 8.544846 | 1.29E-17 | 1.61E-16 |
| CNAG_02066 | 121.3151 | 0.430799 | 0.213538 | 2.017432 | 0.04365 | 0.0835 |
| CNAG_02064 | 3610.97 | -0.17246 | 0.061982 | -2.78243 | 0.005395 | 0.01339 |
| CNAG_02062 | 966.3409 | 1.234059 | 0.090281 | 13.66908 | 1.55E-42 | 4.53E-41 |
| CNAG_02061 | 432.5537 | 0.409217 | 0.120045 | 3.408856 | 0.000652 | 0.002007 |
| CNAG_02060 | 1330.485 | -0.83891 | 0.115797 | -7.2446 | 4.34E-13 | 4.29E-12 |
| CNAG_12600 | 394.5593 | 0.442632 | 0.155168 | 2.852596 | 0.004336 | 0.011025 |
| CNAG_02057 | 741.6325 | -0.70689 | 0.099967 | -7.07128 | 1.54E-12 | 1.45E-11 |
| CNAG_02053 | 38.43995 | -1.20251 | 0.381376 | -3.15309 | 0.001616 | 0.004536 |
| CNAG_12603 | 20.53524 | -1.0066 | 0.484654 | -2.07694 | 0.037807 | 0.073766 |
| CNAG_02049 | 69.07862 | 2.346075 | 0.29702 | 7.898718 | 2.82E-15 | 3.16E-14 |
| CNAG_02048 | 864.6425 | 0.562478 | 0.090792 | 6.195221 | 5.82E-10 | 4.36E-09 |
| CNAG_02044 | 159.0894 | 0.823781 | 0.170404 | 4.834285 | 1.34E-06 | 6.71E-06 |
| CNAG_02043 | 659.3559 | 2.261997 | 0.107691 | 21.0045 | 5.97E-98 | 4.22E-96 |
| CNAG_12604 | 4.785441 | 3.925355 | 1.355199 | 2.896517 | 0.003773 | 0.009721 |
| CNAG_07649 | 6307.124 | 0.609632 | 0.060355 | 10.10077 | 5.48E-24 | 9.22E-23 |
| CNAG_02041 | 2063.891 | 1.127422 | 0.265422 | 4.247658 | 2.16E-05 | 8.89E-05 |
| CNAG_02040 | 290.5437 | 1.082761 | 0.161106 | 6.720786 | 1.81E-11 | 1.55E-10 |
| CNAG_02039 | 108.5635 | 1.502606 | 0.2277 | 6.599053 | 4.14E-11 | 3.46E-10 |
| CNAG_02038 | 2830.853 | 0.437361 | 0.075268 | 5.810737 | 6.22E-09 | 4.21E-08 |

| | | | | | | |
|---|---|---|---|---|---|---|
| CNAG_02037 | 2821.645 | -0.26909 | 0.078003 | -3.44976 | 0.000561 | 0.001753 |
| CNAG_02035 | 12806.45 | -0.61174 | 0.059928 | -10.2078 | 1.83E-24 | 3.13E-23 |
| CNAG_02030 | 10123.13 | 0.911363 | 0.09035 | 10.08701 | 6.31E-24 | 1.06E-22 |
| CNAG_02029 | 1103.648 | -0.32472 | 0.09597 | -3.38351 | 0.000716 | 0.002188 |
| CNAG_02027 | 1428.811 | 0.75035 | 0.077348 | 9.700927 | 2.99E-22 | 4.70E-21 |
| CNAG_02026 | 196.0705 | 0.560761 | 0.166419 | 3.369571 | 0.000753 | 0.002289 |
| CNAG_02024 | 1640.495 | -0.36927 | 0.089604 | -4.1212 | 3.77E-05 | 0.000149 |
| CNAG_02023 | 93.04264 | -0.6967 | 0.226032 | -3.0823 | 0.002054 | 0.005633 |
| CNAG_02021 | 1010.775 | -0.3598 | 0.085639 | -4.20135 | 2.65E-05 | 0.000107 |
| CNAG_02020 | 1340.402 | -0.3991 | 0.090693 | -4.40059 | 1.08E-05 | 4.71E-05 |
| CNAG_07651 | 1061.089 | -0.49046 | 0.084232 | -5.8227 | 5.79E-09 | 3.94E-08 |
| CNAG_07652 | 14.92404 | -1.09606 | 0.552682 | -1.98317 | 0.047349 | 0.089542 |
| CNAG_06524 | 11903.13 | 6.660527 | 0.265914 | 25.04769 | 1.85E-138 | 2.09E-136 |
| CNAG_06525 | 194.1101 | 0.381853 | 0.175714 | 2.173151 | 0.029769 | 0.059981 |
| CNAG_06526 | 2173.671 | -0.42079 | 0.079539 | -5.29034 | 1.22E-07 | 7.07E-07 |
| CNAG_06527 | 34.10265 | -1.22777 | 0.379315 | -3.23681 | 0.001209 | 0.003485 |
| CNAG_12608 | 35.48674 | -0.95 | 0.375344 | -2.53101 | 0.011374 | 0.025946 |
| CNAG_06529 | 130.994 | 0.936306 | 0.203989 | 4.589991 | 4.43E-06 | 2.07E-05 |
| CNAG_06531 | 1204.65 | 1.183443 | 0.10637 | 11.12567 | 9.41E-29 | 1.85E-27 |
| CNAG_06534 | 8628.562 | 0.155802 | 0.065308 | 2.385651 | 0.017049 | 0.036855 |
| CNAG_06536 | 559.4933 | 1.020711 | 0.110659 | 9.223941 | 2.86E-20 | 4.13E-19 |
| CNAG_06539 | 165.1746 | 1.695483 | 0.217882 | 7.781657 | 7.16E-15 | 7.80E-14 |
| CNAG_06540 | 171.9774 | 1.050047 | 0.18074 | 5.809714 | 6.26E-09 | 4.24E-08 |
| CNAG_06541 | 5801.022 | 0.222732 | 0.111303 | 2.001136 | 0.045378 | 0.086357 |
| CNAG_06544 | 12398.42 | -0.73008 | 0.061917 | -11.7912 | 4.34E-32 | 9.30E-31 |
| CNAG_06545 | 4565.542 | -0.55237 | 0.076865 | -7.18626 | 6.66E-13 | 6.44E-12 |
| CNAG_06549 | 898.1072 | -0.44638 | 0.090848 | -4.91344 | 8.95E-07 | 4.62E-06 |
| CNAG_06550 | 3559.551 | 0.178539 | 0.064267 | 2.778092 | 0.005468 | 0.01355 |
| CNAG_06551 | 372.5635 | 1.891276 | 0.175304 | 10.78858 | 3.90E-27 | 7.25E-26 |
| CNAG_06554 | 1400.016 | 2.017842 | 0.081997 | 24.60859 | 1.02E-133 | 1.05E-131 |
| CNAG_06555 | 1935.554 | 1.118308 | 0.073364 | 15.24322 | 1.83E-52 | 6.57E-51 |
| CNAG_06556 | 382.7093 | 4.282597 | 0.178615 | 23.97673 | 4.86E-127 | 4.54E-125 |
| CNAG_06557 | 437.5445 | 4.624025 | 0.181178 | 25.52202 | 1.12E-143 | 1.38E-141 |
| CNAG_06559 | 774.1712 | 0.87585 | 0.126961 | 6.898553 | 5.25E-12 | 4.73E-11 |
| CNAG_06563 | 5253.998 | -0.31001 | 0.061644 | -5.02911 | 4.93E-07 | 2.64E-06 |
| CNAG_06565 | 900.7694 | -0.38519 | 0.098055 | -3.92828 | 8.56E-05 | 0.000318 |
| CNAG_06566 | 1388.743 | -0.42173 | 0.078846 | -5.34874 | 8.86E-08 | 5.23E-07 |
| CNAG_06567 | 1219.505 | -0.74622 | 0.09063 | -8.23377 | 1.81E-16 | 2.16E-15 |
| CNAG_06568 | 1593.284 | -1.71583 | 0.083282 | -20.6026 | 2.60E-94 | 1.78E-92 |
| CNAG_12616 | 189.2387 | 1.532499 | 0.173552 | 8.830222 | 1.04E-18 | 1.38E-17 |
| CNAG_06569 | 931.257 | 0.188646 | 0.088723 | 2.126239 | 0.033483 | 0.066363 |
| CNAG_06570 | 506.2656 | 0.356896 | 0.108917 | 3.276775 | 0.00105 | 0.003073 |
| CNAG_07660 | 10627.15 | -0.18291 | 0.067874 | -2.69488 | 0.007041 | 0.016893 |
| CNAG_06573 | 1249.401 | 0.288432 | 0.088191 | 3.270548 | 0.001073 | 0.003129 |
| CNAG_06576 | 3567.04 | 1.583163 | 0.250754 | 6.3136 | 2.73E-10 | 2.12E-09 |
| CNAG_06577 | 1001.566 | 1.408274 | 0.368995 | 3.816516 | 0.000135 | 0.000482 |
| CNAG_06578 | 1014.675 | 1.171521 | 0.116937 | 10.01837 | 1.27E-23 | 2.10E-22 |

| | | | | | | |
|---|---|---|---|---|---|---|
| CNAG_12620 | 95.53988 | 0.514437 | 0.248419 | 2.070846 | 0.038373 | 0.074745 |
| CNAG_07989 | 920.3455 | -0.23649 | 0.094222 | -2.50989 | 0.012077 | 0.027313 |
| CNAG_07661 | 304.8418 | 0.591475 | 0.129979 | 4.550526 | 5.35E-06 | 2.48E-05 |
| CNAG_12622 | 90.45288 | 1.662988 | 0.251998 | 6.599202 | 4.13E-11 | 3.46E-10 |
| CNAG_06583 | 366.4325 | 1.026905 | 0.153122 | 6.706448 | 1.99E-11 | 1.70E-10 |
| CNAG_06584 | 846.645 | 0.322785 | 0.08601 | 3.752863 | 0.000175 | 0.000609 |
| CNAG_06588 | 1155.928 | 0.733754 | 0.085032 | 8.629132 | 6.18E-18 | 7.88E-17 |
| CNAG_06589 | 797.146 | -0.22268 | 0.110278 | -2.01923 | 0.043464 | 0.083182 |
| CNAG_06590 | 282.5501 | -0.75178 | 0.147585 | -5.09386 | 3.51E-07 | 1.92E-06 |
| CNAG_06591 | 4582.476 | -0.35052 | 0.078556 | -4.46196 | 8.12E-06 | 3.64E-05 |
| CNAG_06592 | 2160.723 | 0.156082 | 0.077458 | 2.01504 | 0.0439 | 0.083919 |
| CNAG_06594 | 5776.146 | -0.20892 | 0.065551 | -3.1872 | 0.001437 | 0.004071 |
| CNAG_06597 | 2189.753 | -0.31114 | 0.067639 | -4.6 | 4.22E-06 | 1.98E-05 |
| CNAG_06598 | 52.61524 | -0.89678 | 0.353143 | -2.53943 | 0.011103 | 0.025422 |
| CNAG_12627 | 131.1066 | 0.877389 | 0.194968 | 4.500165 | 6.79E-06 | 3.10E-05 |
| CNAG_06602 | 1355.351 | -0.45323 | 0.115812 | -3.91347 | 9.10E-05 | 0.000337 |
| CNAG_06603 | 1901.355 | -0.28429 | 0.074458 | -3.81807 | 0.000135 | 0.000479 |
| CNAG_06605 | 65669.78 | -0.19109 | 0.061851 | -3.08954 | 0.002005 | 0.005523 |
| CNAG_06606 | 2383.366 | 0.346181 | 0.082783 | 4.181778 | 2.89E-05 | 0.000116 |
| CNAG_06607 | 2387.877 | 0.402517 | 0.071677 | 5.615692 | 1.96E-08 | 1.24E-07 |
| CNAG_06608 | 1876.622 | 0.213911 | 0.071919 | 2.974323 | 0.002936 | 0.007767 |
| CNAG_06610 | 259.7129 | 0.446899 | 0.138257 | 3.232369 | 0.001228 | 0.003534 |
| CNAG_12630 | 81.16586 | -0.48734 | 0.238381 | -2.04438 | 0.040916 | 0.078919 |
| CNAG_06611 | 3455.8 | 0.33311 | 0.062586 | 5.322453 | 1.02E-07 | 5.98E-07 |
| CNAG_06612 | 1698.227 | -0.26861 | 0.082867 | -3.24149 | 0.001189 | 0.003435 |
| CNAG_06614 | 995.0128 | 0.790473 | 0.12016 | 6.578509 | 4.75E-11 | 3.95E-10 |
| CNAG_06616 | 878.1151 | -0.40782 | 0.101395 | -4.02212 | 5.77E-05 | 0.000221 |
| CNAG_06617 | 1548.454 | 0.547188 | 0.0854 | 6.407331 | 1.48E-10 | 1.18E-09 |
| CNAG_12635 | 1.404214 | 4.011354 | 2.018199 | 1.987591 | 0.046857 | 0.088723 |
| CNAG_06621 | 7997.407 | 1.077191 | 0.064778 | 16.62908 | 4.29E-62 | 1.90E-60 |
| CNAG_06622 | 276.6771 | 1.936454 | 0.183913 | 10.52921 | 6.34E-26 | 1.13E-24 |
| CNAG_06623 | 10833.01 | 1.745868 | 0.108289 | 16.12234 | 1.78E-58 | 7.22E-57 |
| CNAG_06624 | 1299.27 | -0.34426 | 0.089903 | -3.82924 | 0.000129 | 0.00046 |
| CNAG_06625 | 2520.035 | -0.15779 | 0.072554 | -2.17475 | 0.029649 | 0.059783 |
| CNAG_06628 | 2487.6 | 0.185375 | 0.08618 | 2.15101 | 0.031475 | 0.062982 |
| CNAG_06631 | 1893.247 | -0.18352 | 0.087876 | -2.08842 | 0.03676 | 0.071927 |
| CNAG_06633 | 45493.07 | -0.18644 | 0.068883 | -2.70656 | 0.006798 | 0.016363 |
| CNAG_06635 | 3605.617 | -0.21643 | 0.070717 | -3.06049 | 0.00221 | 0.006017 |
| CNAG_06636 | 281.4348 | 0.44404 | 0.149822 | 2.963776 | 0.003039 | 0.008007 |
| CNAG_06638 | 8329.498 | 1.118952 | 0.081647 | 13.70478 | 9.51E-43 | 2.81E-41 |
| CNAG_12642 | 73.83616 | 1.834861 | 0.26137 | 7.020161 | 2.22E-12 | 2.07E-11 |
| CNAG_06643 | 567.8747 | -0.31164 | 0.110263 | -2.82631 | 0.004709 | 0.011842 |
| CNAG_06645 | 2701.82 | 0.303559 | 0.080736 | 3.75989 | 0.00017 | 0.000595 |
| CNAG_06646 | 13198.47 | -0.39511 | 0.05647 | -6.99669 | 2.62E-12 | 2.41E-11 |
| CNAG_06647 | 1563.318 | 0.530744 | 0.080216 | 6.616455 | 3.68E-11 | 3.09E-10 |
| CNAG_06649 | 247.9662 | 1.020099 | 0.142251 | 7.171113 | 7.44E-13 | 7.17E-12 |
| CNAG_12643 | 2.53375 | 4.871188 | 1.779941 | 2.736713 | 0.006206 | 0.015111 |

| | | | | | | |
|---|---|---|---|---|---|---|
| CNAG_06650 | 2892.261 | 4.894956 | 0.106018 | 46.17105 | 0 | 0 |
| CNAG_06651 | 285.0091 | 5.594552 | 0.273071 | 20.48751 | 2.78E-93 | 1.82E-91 |
| CNAG_06652 | 224.7739 | 1.039835 | 0.163154 | 6.373321 | 1.85E-10 | 1.46E-09 |
| CNAG_06656 | 2281.66 | 0.444066 | 0.073207 | 6.065867 | 1.31E-09 | 9.51E-09 |
| CNAG_06657 | 303.1378 | -0.42535 | 0.135002 | -3.15071 | 0.001629 | 0.00457 |
| CNAG_06658 | 714.0265 | 1.749418 | 0.120285 | 14.54396 | 6.38E-48 | 2.07E-46 |
| CNAG_06660 | 822.0004 | -0.42787 | 0.103774 | -4.12309 | 3.74E-05 | 0.000148 |
| CNAG_06661 | 1741.925 | -0.35149 | 0.088157 | -3.98706 | 6.69E-05 | 0.000253 |
| CNAG_06663 | 9808.138 | 0.630087 | 0.062517 | 10.07859 | 6.87E-24 | 1.15E-22 |
| CNAG_06664 | 629.6517 | -0.66503 | 0.097638 | -6.81118 | 9.68E-12 | 8.59E-11 |
| CNAG_06666 | 2858.628 | 1.009984 | 0.079466 | 12.7096 | 5.23E-37 | 1.32E-35 |
| CNAG_06669 | 1524.097 | -0.34668 | 0.091706 | -3.78034 | 0.000157 | 0.000552 |
| CNAG_06670 | 1856.235 | -0.31037 | 0.07556 | -4.10768 | 4.00E-05 | 0.000158 |
| CNAG_06671 | 2069.844 | 0.196748 | 0.075708 | 2.598774 | 0.009356 | 0.021839 |
| CNAG_12646 | 10.9569 | 1.45831 | 0.660955 | 2.20637 | 0.027358 | 0.055787 |
| CNAG_12647 | 201.8711 | 0.832118 | 0.164022 | 5.073202 | 3.91E-07 | 2.12E-06 |
| CNAG_06674 | 856.2055 | -0.25551 | 0.088437 | -2.88916 | 0.003863 | 0.009925 |
| CNAG_06676 | 452.5856 | -0.35296 | 0.111813 | -3.15667 | 0.001596 | 0.004488 |
| CNAG_06678 | 1037.394 | -0.54259 | 0.103386 | -5.24824 | 1.54E-07 | 8.80E-07 |
| CNAG_06679 | 2983.574 | -0.27255 | 0.066726 | -4.0846 | 4.42E-05 | 0.000172 |
| CNAG_06680 | 751.8678 | -0.34253 | 0.103072 | -3.32321 | 0.00089 | 0.002648 |
| CNAG_06681 | 769.904 | -0.40196 | 0.121471 | -3.30914 | 0.000936 | 0.002773 |
| CNAG_07665 | 5035.624 | 0.270308 | 0.090371 | 2.991102 | 0.00278 | 0.007408 |
| CNAG_07177 | 4747.938 | 0.590991 | 0.097914 | 6.035845 | 1.58E-09 | 1.13E-08 |
| CNAG_06690 | 2194.134 | -0.46581 | 0.068742 | -6.77631 | 1.23E-11 | 1.08E-10 |
| CNAG_06692 | 857.251 | 0.380393 | 0.088188 | 4.313421 | 1.61E-05 | 6.78E-05 |
| CNAG_06693 | 544.0034 | 0.581685 | 0.110276 | 5.274786 | 1.33E-07 | 7.67E-07 |
| CNAG_06694 | 253.2691 | -0.9857 | 0.141912 | -6.94582 | 3.76E-12 | 3.42E-11 |
| CNAG_06696 | 4931.863 | -0.35481 | 0.061103 | -5.80682 | 6.37E-09 | 4.30E-08 |
| CNAG_06697 | 1037.256 | -0.29127 | 0.090646 | -3.21325 | 0.001312 | 0.003755 |
| CNAG_07990 | 1308.991 | -0.50961 | 0.081061 | -6.28671 | 3.24E-10 | 2.49E-09 |
| CNAG_06698 | 1332.955 | -0.50847 | 0.094695 | -5.36954 | 7.89E-08 | 4.71E-07 |
| CNAG_06699 | 107055.7 | -0.19361 | 0.054938 | -3.52421 | 0.000425 | 0.001363 |
| CNAG_05671 | 634.5978 | -0.2087 | 0.102756 | -2.03104 | 0.042251 | 0.081167 |
| CNAG_05673 | 1387.815 | -0.28741 | 0.077782 | -3.69509 | 0.00022 | 0.000749 |
| CNAG_05674 | 953.5084 | -0.48126 | 0.085842 | -5.60631 | 2.07E-08 | 1.31E-07 |
| CNAG_05676 | 376.8417 | 0.452265 | 0.139215 | 3.248674 | 0.001159 | 0.00336 |
| CNAG_05678 | 1086.117 | 0.474574 | 0.082428 | 5.757422 | 8.54E-09 | 5.66E-08 |
| CNAG_05679 | 2130.506 | -0.31815 | 0.069962 | -4.54744 | 5.43E-06 | 2.51E-05 |
| CNAG_05683 | 201.367 | -0.70774 | 0.248515 | -2.84789 | 0.004401 | 0.011157 |
| CNAG_05685 | 1186.61 | 0.415377 | 0.088009 | 4.719699 | 2.36E-06 | 1.15E-05 |
| CNAG_12663 | 111.422 | -0.8765 | 0.314099 | -2.79051 | 0.005262 | 0.013088 |
| CNAG_05686 | 558.6661 | -0.28601 | 0.115179 | -2.48322 | 0.01302 | 0.029241 |
| CNAG_05688 | 1066.846 | -0.44291 | 0.08064 | -5.49247 | 3.96E-08 | 2.43E-07 |
| CNAG_05689 | 1469.287 | -0.39981 | 0.088611 | -4.51196 | 6.42E-06 | 2.94E-05 |
| CNAG_05690 | 3600.809 | -0.24467 | 0.060771 | -4.02607 | 5.67E-05 | 0.000217 |
| CNAG_05692 | 1185.242 | -0.45946 | 0.081346 | -5.64819 | 1.62E-08 | 1.04E-07 |

| | | | | | | |
|---|---|---|---|---|---|---|
| CNAG_05694 | 4494.972 | -0.1908 | 0.071414 | -2.67168 | 0.007547 | 0.018 |
| CNAG_05695 | 4425.786 | 0.222124 | 0.064345 | 3.452099 | 0.000556 | 0.00174 |
| CNAG_05697 | 1147.595 | 0.446853 | 0.086002 | 5.19582 | 2.04E-07 | 1.14E-06 |
| CNAG_05698 | 876.847 | -0.33959 | 0.09342 | -3.63509 | 0.000278 | 0.000927 |
| CNAG_05699 | 1004.797 | 0.395428 | 0.095801 | 4.127607 | 3.67E-05 | 0.000145 |
| CNAG_05700 | 1869.737 | -0.20627 | 0.06981 | -2.95476 | 0.003129 | 0.008228 |
| CNAG_12667 | 198.0589 | 0.801316 | 0.227067 | 3.52899 | 0.000417 | 0.00134 |
| CNAG_05702 | 1258.888 | -0.21674 | 0.082982 | -2.61187 | 0.009005 | 0.021117 |
| CNAG_05703 | 2187.043 | -0.18797 | 0.072178 | -2.60431 | 0.009206 | 0.021527 |
| CNAG_05705 | 486.2795 | -0.32543 | 0.124746 | -2.6087 | 0.009089 | 0.021295 |
| CNAG_05707 | 736.017 | 0.391049 | 0.112835 | 3.465662 | 0.000529 | 0.001657 |
| CNAG_05708 | 1422.371 | -0.44722 | 0.089503 | -4.99672 | 5.83E-07 | 3.09E-06 |
| CNAG_12668 | 95.77436 | -1.13111 | 0.228195 | -4.95677 | 7.17E-07 | 3.76E-06 |
| CNAG_05711 | 769.964 | -0.48295 | 0.109697 | -4.40259 | 1.07E-05 | 4.68E-05 |
| CNAG_05714 | 1266.001 | -0.31126 | 0.085106 | -3.65729 | 0.000255 | 0.000857 |
| CNAG_05715 | 893.9332 | -0.40047 | 0.104941 | -3.81619 | 0.000136 | 0.000482 |
| CNAG_05717 | 358.4681 | -0.53145 | 0.135422 | -3.9244 | 8.69E-05 | 0.000323 |
| CNAG_05719 | 900.2185 | -0.19596 | 0.094987 | -2.06305 | 0.039108 | 0.075919 |
| CNAG_05720 | 1462.305 | -0.50723 | 0.07934 | -6.39311 | 1.63E-10 | 1.28E-09 |
| CNAG_05721 | 1116.409 | 0.244735 | 0.081503 | 3.002782 | 0.002675 | 0.007148 |
| CNAG_05722 | 4170.526 | 0.272164 | 0.061867 | 4.399186 | 1.09E-05 | 4.74E-05 |
| CNAG_05724 | 1546.485 | 1.019672 | 0.082057 | 12.42631 | 1.88E-35 | 4.48E-34 |
| CNAG_05725 | 51171.06 | -0.19715 | 0.056256 | -3.50454 | 0.000457 | 0.001454 |
| CNAG_12670 | 67.92234 | 0.639563 | 0.257282 | 2.485848 | 0.012924 | 0.029051 |
| CNAG_05727 | 33.58321 | 2.346283 | 0.408969 | 5.737068 | 9.63E-09 | 6.35E-08 |
| CNAG_05731 | 1677.538 | 0.68805 | 0.076514 | 8.992524 | 2.42E-19 | 3.32E-18 |
| CNAG_05732 | 1713.358 | -1.07287 | 0.112951 | -9.49857 | 2.13E-21 | 3.24E-20 |
| CNAG_05737 | 1300.338 | -0.41287 | 0.091394 | -4.51748 | 6.26E-06 | 2.87E-05 |
| CNAG_05738 | 130.3272 | 1.587178 | 0.266107 | 5.964439 | 2.45E-09 | 1.73E-08 |
| CNAG_05740 | 535.3075 | -0.30864 | 0.126258 | -2.44456 | 0.014503 | 0.032039 |
| CNAG_05742 | 1343.683 | -0.48152 | 0.084722 | -5.68357 | 1.32E-08 | 8.52E-08 |
| CNAG_05743 | 478.3062 | -0.25576 | 0.106554 | -2.40032 | 0.016381 | 0.035638 |
| CNAG_05744 | 660.2837 | -0.32266 | 0.099901 | -3.22978 | 0.001239 | 0.003561 |
| CNAG_05747 | 809.4054 | -0.21355 | 0.09568 | -2.23194 | 0.025619 | 0.052637 |
| CNAG_05749 | 786.0784 | -0.33285 | 0.108366 | -3.07156 | 0.002129 | 0.005816 |
| CNAG_05750 | 70535.33 | -0.13882 | 0.069135 | -2.00798 | 0.044645 | 0.085182 |
| CNAG_05751 | 1386.388 | -0.23309 | 0.11082 | -2.10332 | 0.035438 | 0.069575 |
| CNAG_05752 | 2369.488 | -0.36354 | 0.086176 | -4.21852 | 2.46E-05 | 9.98E-05 |
| CNAG_05753 | 9648.222 | -0.12539 | 0.060836 | -2.06113 | 0.03929 | 0.076236 |
| CNAG_07672 | 2175.701 | 0.391972 | 0.078834 | 4.972124 | 6.62E-07 | 3.49E-06 |
| CNAG_05755 | 3201.566 | 0.173287 | 0.070561 | 2.455839 | 0.014056 | 0.031197 |
| CNAG_05756 | 1564.075 | 0.249691 | 0.100404 | 2.486864 | 0.012887 | 0.029 |
| CNAG_05757 | 942.5591 | -0.6881 | 0.099316 | -6.9284 | 4.26E-12 | 3.85E-11 |
| CNAG_05758 | 1002.13 | -0.24862 | 0.090017 | -2.76187 | 0.005747 | 0.014147 |
| CNAG_05759 | 10944.13 | -0.23881 | 0.065932 | -3.622 | 0.000292 | 0.000971 |
| CNAG_12675 | 94.8963 | 1.149169 | 0.226515 | 5.073265 | 3.91E-07 | 2.12E-06 |
| CNAG_05760 | 32.28191 | 0.776522 | 0.376984 | 2.059828 | 0.039415 | 0.076459 |

| | | | | | | |
|---|---|---|---|---|---|---|
| CNAG_05761 | 1178.286 | 0.194691 | 0.088697 | 2.195015 | 0.028162 | 0.057198 |
| CNAG_05762 | 65203.54 | -0.23426 | 0.075353 | -3.10888 | 0.001878 | 0.005213 |
| CNAG_05763 | 516.9623 | 0.507744 | 0.107752 | 4.712142 | 2.45E-06 | 1.19E-05 |
| CNAG_05764 | 2973.673 | -0.36787 | 0.080322 | -4.57988 | 4.65E-06 | 2.17E-05 |
| CNAG_05768 | 1607.972 | -0.17048 | 0.080317 | -2.12259 | 0.033789 | 0.066821 |
| CNAG_05769 | 938.0488 | -0.25329 | 0.085738 | -2.95422 | 0.003135 | 0.00824 |
| CNAG_05771 | 1167.274 | 0.205051 | 0.091015 | 2.25294 | 0.024263 | 0.050193 |
| CNAG_05774 | 813.9959 | -0.26069 | 0.091539 | -2.84791 | 0.004401 | 0.011157 |
| CNAG_05776 | 1137.759 | -0.23943 | 0.087516 | -2.73579 | 0.006223 | 0.015149 |
| CNAG_05777 | 351.8169 | -0.29998 | 0.125952 | -2.38169 | 0.017233 | 0.037224 |
| CNAG_05778 | 715.7151 | 0.44942 | 0.097681 | 4.600873 | 4.21E-06 | 1.98E-05 |
| CNAG_05782 | 3058.462 | -0.48672 | 0.071513 | -6.80601 | 1.00E-11 | 8.89E-11 |
| CNAG_05783 | 4364.034 | 0.405535 | 0.064662 | 6.271599 | 3.57E-10 | 2.73E-09 |
| CNAG_05784 | 277.2093 | -0.54569 | 0.158545 | -3.44184 | 0.000578 | 0.001801 |
| CNAG_12679 | 54.1046 | 1.430673 | 0.297479 | 4.809327 | 1.51E-06 | 7.57E-06 |
| CNAG_05786 | 364.8285 | -0.43383 | 0.123183 | -3.52185 | 0.000429 | 0.001372 |
| CNAG_05788 | 1693.099 | -0.22375 | 0.091131 | -2.4553 | 0.014077 | 0.031235 |
| CNAG_05789 | 2268.328 | 0.289106 | 0.078345 | 3.690172 | 0.000224 | 0.000762 |
| CNAG_05790 | 1562.683 | 0.662679 | 0.079601 | 8.324956 | 8.44E-17 | 1.02E-15 |
| CNAG_05793 | 888.9825 | -0.50974 | 0.08624 | -5.91073 | 3.41E-09 | 2.37E-08 |
| CNAG_07675 | 1082.649 | 0.24148 | 0.0963 | 2.507587 | 0.012156 | 0.027483 |
| CNAG_05797 | 4457.83 | -0.15132 | 0.064007 | -2.36415 | 0.018072 | 0.038838 |
| CNAG_05798 | 1203.501 | -0.3355 | 0.100814 | -3.32793 | 0.000875 | 0.002613 |
| CNAG_05799 | 6608.682 | 0.356063 | 0.063036 | 5.648551 | 1.62E-08 | 1.04E-07 |
| CNAG_05802 | 276.3479 | 0.980823 | 0.138275 | 7.093284 | 1.31E-12 | 1.24E-11 |
| CNAG_05803 | 2434.174 | 1.015321 | 0.068432 | 14.837 | 8.44E-50 | 2.87E-48 |
| CNAG_05804 | 2993.716 | 0.129836 | 0.064243 | 2.02102 | 0.043278 | 0.082884 |
| CNAG_05807 | 440.0747 | 0.621075 | 0.122408 | 5.073824 | 3.90E-07 | 2.12E-06 |
| CNAG_05809 | 2911.465 | 0.410097 | 0.069006 | 5.942908 | 2.80E-09 | 1.96E-08 |
| CNAG_05810 | 532.2629 | -0.2809 | 0.106405 | -2.63992 | 0.008293 | 0.019559 |
| CNAG_05811 | 768.768 | -0.20236 | 0.092264 | -2.19324 | 0.02829 | 0.057414 |
| CNAG_05812 | 1062.79 | -0.34823 | 0.085001 | -4.09676 | 4.19E-05 | 0.000165 |
| CNAG_05814 | 39991.86 | -0.18197 | 0.069732 | -2.60958 | 0.009065 | 0.021253 |
| CNAG_05815 | 306.5484 | 0.70594 | 0.168437 | 4.191122 | 2.78E-05 | 0.000112 |
| CNAG_05817 | 8240.169 | 0.467146 | 0.065205 | 7.164303 | 7.82E-13 | 7.53E-12 |
| CNAG_05818 | 2394.86 | 1.566144 | 0.07544 | 20.76009 | 9.94E-96 | 6.90E-94 |
| CNAG_05819 | 1050.591 | -0.27069 | 0.095804 | -2.8255 | 0.004721 | 0.011861 |
| CNAG_05820 | 424.1291 | 0.624382 | 0.119348 | 5.23161 | 1.68E-07 | 9.56E-07 |
| CNAG_12683 | 349.0375 | 0.54754 | 0.135089 | 4.053191 | 5.05E-05 | 0.000196 |
| CNAG_05821 | 32.22518 | 0.781347 | 0.398088 | 1.962749 | 0.049675 | 0.093161 |
| CNAG_05822 | 1487.059 | -0.31597 | 0.080292 | -3.93524 | 8.31E-05 | 0.00031 |
| CNAG_05825 | 4056.606 | -0.18334 | 0.063283 | -2.89717 | 0.003765 | 0.009707 |
| CNAG_05826 | 6645.308 | -0.24801 | 0.066939 | -3.70503 | 0.000211 | 0.000724 |
| CNAG_05828 | 2510.708 | -0.1965 | 0.095741 | -2.05243 | 0.040128 | 0.077675 |
| CNAG_05829 | 2166.484 | -0.5151 | 0.074905 | -6.87668 | 6.13E-12 | 5.49E-11 |
| CNAG_05831 | 301.102 | -0.68217 | 0.139785 | -4.88015 | 1.06E-06 | 5.41E-06 |
| CNAG_05835 | 1560.033 | -0.58866 | 0.079166 | -7.43584 | 1.04E-13 | 1.06E-12 |

| | | | | | | |
|---|---|---|---|---|---|---|
| CNAG_05836 | 1626.726 | 0.218491 | 0.093909 | 2.326626 | 0.019985 | 0.042368 |
| CNAG_05839 | 11065.22 | 0.463559 | 0.070376 | 6.586915 | 4.49E-11 | 3.74E-10 |
| CNAG_05841 | 699.2273 | 0.417923 | 0.11448 | 3.650629 | 0.000262 | 0.000876 |
| CNAG_05843 | 936.2309 | 0.581679 | 0.086807 | 6.700852 | 2.07E-11 | 1.77E-10 |
| CNAG_05845 | 2958.892 | -0.24944 | 0.070356 | -3.54539 | 0.000392 | 0.001266 |
| CNAG_05847 | 41109.52 | 3.385007 | 0.063693 | 53.14585 | 0 | 0 |
| CNAG_05848 | 5642.634 | -0.49111 | 0.080152 | -6.12725 | 8.94E-10 | 6.55E-09 |
| CNAG_07676 | 17621.73 | 0.332906 | 0.056382 | 5.904472 | 3.54E-09 | 2.46E-08 |
| CNAG_07677 | 406.2587 | 0.567079 | 0.133754 | 4.239708 | 2.24E-05 | 9.17E-05 |
| CNAG_05851 | 1420.216 | 0.53519 | 0.079514 | 6.730798 | 1.69E-11 | 1.46E-10 |
| CNAG_05852 | 1487.308 | 0.581924 | 0.076457 | 7.611124 | 2.72E-14 | 2.88E-13 |
| CNAG_05853 | 787.7304 | -0.1941 | 0.09884 | -1.96375 | 0.049559 | 0.093001 |
| CNAG_05855 | 1174.82 | -0.33368 | 0.104005 | -3.20833 | 0.001335 | 0.003813 |
| CNAG_07678 | 150.6252 | -0.50598 | 0.205115 | -2.46682 | 0.013632 | 0.030398 |
| CNAG_07679 | 1760.93 | -0.40743 | 0.091671 | -4.4445 | 8.81E-06 | 3.93E-05 |
| CNAG_05859 | 504.4987 | -0.52313 | 0.12304 | -4.25174 | 2.12E-05 | 8.76E-05 |
| CNAG_05860 | 432.6775 | -0.39381 | 0.126555 | -3.1118 | 0.00186 | 0.005167 |
| CNAG_05862 | 549.7671 | -0.29937 | 0.119072 | -2.5142 | 0.01193 | 0.027053 |
| CNAG_05863 | 861.1287 | -1.01301 | 0.113846 | -8.89811 | 5.68E-19 | 7.63E-18 |
| CNAG_05864 | 2273.278 | 1.081369 | 0.119744 | 9.030692 | 1.71E-19 | 2.36E-18 |
| CNAG_05866 | 483.6915 | 0.784715 | 0.113888 | 6.890225 | 5.57E-12 | 5.00E-11 |
| CNAG_05867 | 420.3886 | -0.96607 | 0.149871 | -6.44601 | 1.15E-10 | 9.22E-10 |
| CNAG_05868 | 194.6641 | 0.731428 | 0.157154 | 4.654205 | 3.25E-06 | 1.55E-05 |
| CNAG_05870 | 91.03128 | -0.94269 | 0.260283 | -3.62177 | 0.000293 | 0.000971 |
| CNAG_05871 | 717.9209 | -0.52096 | 0.106747 | -4.88032 | 1.06E-06 | 5.41E-06 |
| CNAG_05872 | 2610.036 | 1.59104 | 0.269503 | 5.903614 | 3.56E-09 | 2.46E-08 |
| CNAG_05874 | 1122.582 | 0.463833 | 0.127859 | 3.627682 | 0.000286 | 0.000951 |
| CNAG_05875 | 3754.714 | 1.758624 | 0.070777 | 24.84756 | 2.75E-136 | 2.94E-134 |
| CNAG_05876 | 573.2032 | 0.985799 | 0.11561 | 8.526928 | 1.50E-17 | 1.87E-16 |
| CNAG_05878 | 5295.487 | -0.12985 | 0.063424 | -2.04735 | 0.040624 | 0.078499 |
| CNAG_05880 | 724.1231 | 0.572031 | 0.103034 | 5.55187 | 2.83E-08 | 1.77E-07 |
| CNAG_05881 | 4570.707 | 0.8177 | 0.059778 | 13.67894 | 1.36E-42 | 3.98E-41 |
| CNAG_05884 | 10099.01 | 0.462766 | 0.057027 | 8.114824 | 4.86E-16 | 5.68E-15 |
| CNAG_05886 | 2291.617 | -0.31067 | 0.08076 | -3.8469 | 0.00012 | 0.000432 |
| CNAG_05887 | 1939.651 | -0.4194 | 0.077786 | -5.39167 | 6.98E-08 | 4.18E-07 |
| CNAG_05888 | 1813.212 | -0.44861 | 0.077242 | -5.80791 | 6.33E-09 | 4.28E-08 |
| CNAG_05889 | 1080.509 | -0.56791 | 0.107017 | -5.30673 | 1.12E-07 | 6.49E-07 |
| CNAG_05892 | 2617.552 | -0.40041 | 0.075624 | -5.29479 | 1.19E-07 | 6.91E-07 |
| CNAG_05893 | 6325.749 | -0.13974 | 0.067103 | -2.08252 | 0.037295 | 0.072851 |
| CNAG_05894 | 2588.161 | 0.372524 | 0.068365 | 5.449042 | 5.06E-08 | 3.08E-07 |
| CNAG_05895 | 3190.159 | -0.15547 | 0.063037 | -2.4663 | 0.013652 | 0.030433 |
| CNAG_05898 | 545.1958 | -0.56843 | 0.13416 | -4.23697 | 2.27E-05 | 9.27E-05 |
| CNAG_05900 | 8936.938 | -0.12363 | 0.055979 | -2.20857 | 0.027205 | 0.055516 |
| CNAG_05901 | 606.1403 | -0.32343 | 0.105941 | -3.05291 | 0.002266 | 0.006161 |
| CNAG_05902 | 499.4046 | -0.33335 | 0.131464 | -2.53567 | 0.011223 | 0.025668 |
| CNAG_12695 | 102.3212 | -0.56383 | 0.256277 | -2.20008 | 0.027801 | 0.056549 |
| CNAG_05903 | 954.8289 | -0.39609 | 0.089652 | -4.41812 | 9.96E-06 | 4.39E-05 |

| | | | | | | |
|---|---|---|---|---|---|---|
| CNAG_05904 | 32431.53 | -0.17078 | 0.078978 | -2.16234 | 0.030592 | 0.061381 |
| CNAG_05905 | 1289.939 | 0.237952 | 0.086468 | 2.751905 | 0.005925 | 0.014505 |
| CNAG_05907 | 16721.5 | 0.334193 | 0.063734 | 5.243593 | 1.57E-07 | 9.00E-07 |
| CNAG_05908 | 168.0279 | 0.486155 | 0.168474 | 2.885639 | 0.003906 | 0.010022 |
| CNAG_12696 | 33.23852 | 1.121845 | 0.373287 | 3.005316 | 0.002653 | 0.007093 |
| CNAG_05909 | 25433.97 | 0.460848 | 0.066888 | 6.889796 | 5.59E-12 | 5.01E-11 |
| CNAG_05911 | 150.9264 | -0.64991 | 0.176195 | -3.68857 | 0.000226 | 0.000767 |
| CNAG_12698 | 256.4066 | -0.56998 | 0.150258 | -3.79331 | 0.000149 | 0.000525 |
| CNAG_05912 | 932.0002 | 0.200283 | 0.085537 | 2.34147 | 0.019208 | 0.040913 |
| CNAG_05913 | 1909.699 | 1.656487 | 0.086603 | 19.12729 | 1.50E-81 | 8.39E-80 |
| CNAG_05914 | 379.2217 | 1.01494 | 0.144635 | 7.017261 | 2.26E-12 | 2.11E-11 |
| CNAG_05915 | 388.4839 | 0.511617 | 0.140402 | 3.643946 | 0.000268 | 0.000898 |
| CNAG_05921 | 761.2049 | -0.27488 | 0.090655 | -3.03216 | 0.002428 | 0.006561 |
| CNAG_05926 | 1967.517 | -0.31569 | 0.073134 | -4.31668 | 1.58E-05 | 6.69E-05 |
| CNAG_05927 | 1394.8 | -0.41942 | 0.074605 | -5.62183 | 1.89E-08 | 1.20E-07 |
| CNAG_05929 | 360.0313 | -0.2944 | 0.131436 | -2.23987 | 0.025099 | 0.0517 |
| CNAG_05930 | 1069.307 | -0.399 | 0.086399 | -4.61815 | 3.87E-06 | 1.83E-05 |
| CNAG_05931 | 2284.725 | -0.30765 | 0.07704 | -3.99336 | 6.51E-05 | 0.000247 |
| CNAG_05932 | 7829.644 | 0.140639 | 0.064622 | 2.176341 | 0.02953 | 0.059558 |
| CNAG_05937 | 736.6723 | 0.349357 | 0.102208 | 3.418103 | 0.000631 | 0.001948 |
| CNAG_12702 | 51.87046 | 1.730859 | 0.34303 | 5.0458 | 4.52E-07 | 2.43E-06 |
| CNAG_07685 | 353.3726 | 0.46253 | 0.141109 | 3.277811 | 0.001046 | 0.003063 |
| CNAG_07686 | 1957.716 | 0.160172 | 0.072503 | 2.209186 | 0.027162 | 0.055442 |
| CNAG_12705 | 7.413872 | 3.859265 | 1.118053 | 3.451772 | 0.000557 | 0.00174 |
| CNAG_07687 | 139.2701 | 1.861156 | 0.247829 | 7.50984 | 5.92E-14 | 6.17E-13 |
| CNAG_07689 | 3361.548 | -0.29043 | 0.067201 | -4.32179 | 1.55E-05 | 6.55E-05 |
| CNAG_07690 | 430.3046 | -0.30917 | 0.126408 | -2.44585 | 0.014451 | 0.031956 |
| CNAG_07691 | 152.7546 | 1.042312 | 0.190672 | 5.466518 | 4.59E-08 | 2.80E-07 |
| CNAG_12707 | 27.59879 | 2.15741 | 0.456919 | 4.721642 | 2.34E-06 | 1.14E-05 |
| CNAG_07188 | 326.0514 | 1.089742 | 0.155475 | 7.009103 | 2.40E-12 | 2.22E-11 |
| CNAG_07992 | 41.59829 | 1.039566 | 0.338185 | 3.073956 | 0.002112 | 0.005779 |
| CNAG_07693 | 15.92936 | 1.254194 | 0.59141 | 2.120686 | 0.033948 | 0.067069 |
| CNAG_07694 | 2117.929 | -0.23942 | 0.07124 | -3.36071 | 0.000777 | 0.002351 |
| CNAG_07695 | 7572.613 | 0.835097 | 0.070615 | 11.82609 | 2.86E-32 | 6.19E-31 |
| CNAG_07703 | 231.556 | -0.39573 | 0.169136 | -2.33971 | 0.019299 | 0.041063 |
| CNAG_05964 | 106.0095 | -0.49109 | 0.221375 | -2.21835 | 0.026531 | 0.054305 |
| CNAG_05965 | 368.4091 | -0.49222 | 0.140163 | -3.51176 | 0.000445 | 0.00142 |
| CNAG_05967 | 412.5318 | -0.81197 | 0.147639 | -5.49972 | 3.80E-08 | 2.34E-07 |
| CNAG_05969 | 471.2135 | -0.311 | 0.157021 | -1.98063 | 0.047632 | 0.089995 |
| CNAG_05970 | 476.0293 | -0.28221 | 0.117802 | -2.39564 | 0.016592 | 0.036058 |
| CNAG_12712 | 445.7508 | 0.259605 | 0.108743 | 2.387326 | 0.016971 | 0.036726 |
| CNAG_12713 | 75.23501 | 2.714783 | 0.291088 | 9.32633 | 1.10E-20 | 1.61E-19 |
| CNAG_05975 | 4129.714 | 0.149901 | 0.06381 | 2.349165 | 0.018816 | 0.040203 |
| CNAG_05976 | 5689.809 | 0.20728 | 0.063799 | 3.248973 | 0.001158 | 0.003357 |
| CNAG_05977 | 4659.428 | 0.265495 | 0.067771 | 3.9175 | 8.95E-05 | 0.000332 |
| CNAG_05979 | 962.4853 | 0.199406 | 0.100754 | 1.979138 | 0.0478 | 0.090228 |
| CNAG_05980 | 4357.644 | -0.32648 | 0.074171 | -4.40177 | 1.07E-05 | 4.70E-05 |

| | | | | | | |
|---|---|---|---|---|---|---|
| CNAG_05981 | 471.021 | -0.56383 | 0.118252 | -4.76805 | 1.86E-06 | 9.20E-06 |
| CNAG_07706 | 529.0163 | 0.271525 | 0.111016 | 2.445809 | 0.014453 | 0.031956 |
| CNAG_03084 | 472.2164 | 2.572927 | 0.351177 | 7.326582 | 2.36E-13 | 2.36E-12 |
| CNAG_03085 | 68.45281 | 4.038878 | 0.417125 | 9.682657 | 3.57E-22 | 5.56E-21 |
| CNAG_03086 | 1120.436 | 0.287917 | 0.104766 | 2.748184 | 0.005993 | 0.014645 |
| CNAG_03090 | 273.176 | 0.834369 | 0.161034 | 5.181319 | 2.20E-07 | 1.23E-06 |
| CNAG_03092 | 951.1183 | -0.25601 | 0.093716 | -2.73171 | 0.006301 | 0.015319 |
| CNAG_12715 | 39.7449 | -1.39589 | 0.345071 | -4.04523 | 5.23E-05 | 0.000201 |
| CNAG_03097 | 507.9971 | -0.35325 | 0.123337 | -2.86412 | 0.004182 | 0.010662 |
| CNAG_03101 | 439.2993 | 0.497169 | 0.115631 | 4.299631 | 1.71E-05 | 7.19E-05 |
| CNAG_03106 | 2055.659 | -0.26008 | 0.082477 | -3.15335 | 0.001614 | 0.004533 |
| CNAG_03108 | 798.463 | 0.377504 | 0.101683 | 3.712567 | 0.000205 | 0.000705 |
| CNAG_03110 | 2130.182 | 0.846424 | 0.090012 | 9.403442 | 5.28E-21 | 7.90E-20 |
| CNAG_03115 | 1546.139 | 1.646773 | 0.088208 | 18.66926 | 8.81E-78 | 4.74E-76 |
| CNAG_07711 | 49.88375 | 1.820482 | 0.371057 | 4.906204 | 9.29E-07 | 4.78E-06 |
| CNAG_12718 | 77.17537 | 1.667077 | 0.264758 | 6.296615 | 3.04E-10 | 2.35E-09 |
| CNAG_12719 | 4.623654 | 3.757957 | 1.369686 | 2.743663 | 0.006076 | 0.014827 |
| CNAG_12720 | 135.0057 | 0.518639 | 0.218304 | 2.375764 | 0.017513 | 0.037747 |
| CNAG_03116 | 2208.586 | -0.32456 | 0.077347 | -4.19616 | 2.71E-05 | 0.00011 |
| CNAG_03120 | 9257.024 | 0.331448 | 0.072013 | 4.602613 | 4.17E-06 | 1.96E-05 |
| CNAG_03121 | 1516.529 | -0.21916 | 0.076434 | -2.86733 | 0.004139 | 0.010567 |
| CNAG_03123 | 743.3144 | -0.502 | 0.093444 | -5.37223 | 7.78E-08 | 4.64E-07 |
| CNAG_03125 | 603.8943 | -0.40399 | 0.097366 | -4.14914 | 3.34E-05 | 0.000133 |
| CNAG_03127 | 48643.23 | -0.13852 | 0.068282 | -2.02869 | 0.04249 | 0.081549 |
| CNAG_03128 | 2926.277 | 2.854045 | 0.089548 | 31.87153 | 6.62E-223 | 1.63E-220 |
| CNAG_03130 | 834.8777 | -0.43975 | 0.102698 | -4.28197 | 1.85E-05 | 7.73E-05 |
| CNAG_03132 | 47.4629 | 1.640332 | 0.348313 | 4.709357 | 2.48E-06 | 1.21E-05 |
| CNAG_03133 | 802.6362 | 2.121197 | 0.129317 | 16.40304 | 1.82E-60 | 7.78E-59 |
| CNAG_03135 | 1103.441 | 0.229359 | 0.099009 | 2.316541 | 0.020529 | 0.043261 |
| CNAG_03136 | 2394.845 | -0.27629 | 0.076824 | -3.59644 | 0.000323 | 0.001062 |
| CNAG_12722 | 47.65008 | 1.055901 | 0.314407 | 3.35839 | 0.000784 | 0.002369 |
| CNAG_03139 | 4207.567 | -0.49446 | 0.07654 | -6.4602 | 1.05E-10 | 8.46E-10 |
| CNAG_12723 | 20.80785 | 2.707543 | 0.536043 | 5.05098 | 4.40E-07 | 2.37E-06 |
| CNAG_03142 | 8060.698 | -0.24056 | 0.121986 | -1.972 | 0.048609 | 0.091458 |
| CNAG_03146 | 12868.05 | 0.440451 | 0.065629 | 6.711259 | 1.93E-11 | 1.65E-10 |
| CNAG_03148 | 2694.374 | -0.35267 | 0.077888 | -4.52794 | 5.96E-06 | 2.74E-05 |
| CNAG_03150 | 1515.206 | 0.657046 | 0.079708 | 8.243186 | 1.68E-16 | 2.00E-15 |
| CNAG_03151 | 887.8501 | 0.291385 | 0.094714 | 3.076483 | 0.002095 | 0.005734 |
| CNAG_03152 | 1755.276 | 0.182311 | 0.076573 | 2.380859 | 0.017272 | 0.037268 |
| CNAG_03153 | 8137.128 | -0.1656 | 0.062098 | -2.66681 | 0.007657 | 0.018231 |
| CNAG_03154 | 1198.846 | 1.940473 | 0.098907 | 19.61911 | 1.06E-85 | 6.39E-84 |
| CNAG_12726 | 39.04536 | 0.822126 | 0.378704 | 2.170894 | 0.029939 | 0.060249 |
| CNAG_03156 | 1724.607 | 1.224475 | 0.089159 | 13.73363 | 6.39E-43 | 1.89E-41 |
| CNAG_03158 | 1110.154 | -0.18461 | 0.079858 | -2.31169 | 0.020795 | 0.043754 |
| CNAG_03159 | 1218.617 | -0.25672 | 0.097831 | -2.62413 | 0.008687 | 0.020413 |
| CNAG_03161 | 354.4363 | 0.462404 | 0.125556 | 3.682863 | 0.000231 | 0.000782 |
| CNAG_03163 | 503.978 | 0.320386 | 0.105774 | 3.028963 | 0.002454 | 0.006615 |

| | | | | | | |
|---|---|---|---|---|---|---|
| CNAG_03166 | 210.3726 | -0.45651 | 0.193605 | -2.35795 | 0.018376 | 0.039358 |
| CNAG_03167 | 1345.407 | -0.28204 | 0.091174 | -3.0934 | 0.001979 | 0.005465 |
| CNAG_03168 | 12291.89 | 0.387368 | 0.063184 | 6.130798 | 8.74E-10 | 6.42E-09 |
| CNAG_12730 | 37.07372 | 1.356664 | 0.382081 | 3.550726 | 0.000384 | 0.001244 |
| CNAG_03170 | 1179.129 | 0.719691 | 0.081107 | 8.873363 | 7.10E-19 | 9.46E-18 |
| CNAG_03172 | 552.3662 | 0.334237 | 0.118723 | 2.815267 | 0.004874 | 0.012211 |
| CNAG_12731 | 51.12334 | 2.013837 | 0.378058 | 5.32679 | 1.00E-07 | 5.85E-07 |
| CNAG_03173 | 1570.343 | 0.75236 | 0.107829 | 6.977309 | 3.01E-12 | 2.75E-11 |
| CNAG_03176 | 2228.251 | 1.057545 | 0.085607 | 12.35343 | 4.67E-35 | 1.10E-33 |
| CNAG_12733 | 337.9669 | 0.559522 | 0.175387 | 3.190209 | 0.001422 | 0.004036 |
| CNAG_03178 | 798.8059 | -0.32343 | 0.10008 | -3.23172 | 0.00123 | 0.003539 |
| CNAG_03182 | 1677.029 | -0.19647 | 0.07434 | -2.64287 | 0.008221 | 0.019421 |
| CNAG_03183 | 2724.746 | 0.389278 | 0.077626 | 5.014778 | 5.31E-07 | 2.84E-06 |
| CNAG_03185 | 980.1492 | 0.736274 | 0.098655 | 7.4631 | 8.45E-14 | 8.72E-13 |
| CNAG_03186 | 2560.752 | -0.52987 | 0.075421 | -7.02541 | 2.13E-12 | 1.99E-11 |
| CNAG_03187 | 452.3346 | 0.333234 | 0.117912 | 2.826128 | 0.004711 | 0.011845 |
| CNAG_03190 | 3683.652 | -0.16609 | 0.068921 | -2.40991 | 0.015956 | 0.034789 |
| CNAG_03191 | 1529.28 | -0.32625 | 0.074125 | -4.4014 | 1.08E-05 | 4.70E-05 |
| CNAG_03192 | 406.758 | -0.41129 | 0.118541 | -3.46963 | 0.000521 | 0.001636 |
| CNAG_03193 | 34.83883 | -0.72958 | 0.356564 | -2.04614 | 0.040743 | 0.078669 |
| CNAG_03194 | 121.3339 | -0.99057 | 0.245831 | -4.02948 | 5.59E-05 | 0.000215 |
| CNAG_03196 | 2432.824 | -0.37869 | 0.080211 | -4.72122 | 2.34E-06 | 1.14E-05 |
| CNAG_12735 | 14.5142 | 1.272471 | 0.609116 | 2.089044 | 0.036704 | 0.071852 |
| CNAG_03199 | 133.6702 | 0.736633 | 0.244699 | 3.010358 | 0.002609 | 0.006988 |
| CNAG_03204 | 265.9572 | 0.776903 | 0.155543 | 4.994795 | 5.89E-07 | 3.12E-06 |
| CNAG_03205 | 2275.43 | 0.15229 | 0.072248 | 2.10788 | 0.035041 | 0.068913 |
| CNAG_03206 | 857.2381 | 0.821438 | 0.087276 | 9.411954 | 4.87E-21 | 7.30E-20 |
| CNAG_03210 | 992.1379 | -0.31637 | 0.100077 | -3.1613 | 0.001571 | 0.004419 |
| CNAG_03212 | 829.9802 | -0.3518 | 0.115654 | -3.04181 | 0.002352 | 0.006369 |
| CNAG_03216 | 1061.717 | 0.556778 | 0.09896 | 5.626275 | 1.84E-08 | 1.17E-07 |
| CNAG_07720 | 1793.486 | -0.26609 | 0.07853 | -3.38843 | 0.000703 | 0.00215 |
| CNAG_03222 | 522.9654 | 0.54727 | 0.112723 | 4.854977 | 1.20E-06 | 6.10E-06 |
| CNAG_03223 | 5370.501 | 0.76422 | 0.082635 | 9.248142 | 2.28E-20 | 3.30E-19 |
| CNAG_12739 | 57.45058 | 1.636411 | 0.312083 | 5.243509 | 1.58E-07 | 9.00E-07 |
| CNAG_03224 | 40.78848 | 1.05222 | 0.370944 | 2.836599 | 0.00456 | 0.011506 |
| CNAG_03225 | 14410.01 | -0.90662 | 0.072837 | -12.4471 | 1.45E-35 | 3.47E-34 |
| CNAG_03226 | 10612.54 | 1.570201 | 0.060347 | 26.01961 | 2.97E-149 | 3.77E-147 |
| CNAG_12740 | 185.5085 | 1.594283 | 0.17406 | 9.159408 | 5.22E-20 | 7.39E-19 |
| CNAG_03227 | 500.0812 | 0.250996 | 0.112559 | 2.229904 | 0.025754 | 0.052887 |
| CNAG_03229 | 26.80175 | -3.7031 | 0.566338 | -6.53867 | 6.21E-11 | 5.10E-10 |
| CNAG_12741 | 7.414266 | -4.34142 | 1.245717 | -3.48508 | 0.000492 | 0.001552 |
| CNAG_03230 | 55.70652 | 1.121004 | 0.302029 | 3.711572 | 0.000206 | 0.000708 |
| CNAG_03231 | 438.3286 | -0.25817 | 0.129693 | -1.99061 | 0.046524 | 0.088169 |
| CNAG_03232 | 1015.163 | -0.25346 | 0.114683 | -2.21011 | 0.027097 | 0.055352 |
| CNAG_03234 | 160.1105 | 1.461958 | 0.222224 | 6.57875 | 4.74E-11 | 3.94E-10 |
| CNAG_03235 | 2507.679 | -0.28041 | 0.073068 | -3.83775 | 0.000124 | 0.000446 |
| CNAG_03237 | 694.1036 | 0.236872 | 0.11214 | 2.112287 | 0.034662 | 0.068249 |

| | | | | | | |
|---|---|---|---|---|---|---|
| CNAG_03238 | 181.8372 | 2.722762 | 0.188857 | 14.41708 | 4.04E-47 | 1.29E-45 |
| CNAG_03239 | 629.1135 | -0.35245 | 0.104607 | -3.36932 | 0.000754 | 0.00229 |
| CNAG_03240 | 2710.473 | 0.181445 | 0.066457 | 2.730266 | 0.006328 | 0.015373 |
| CNAG_03245 | 9298.725 | -0.17648 | 0.064524 | -2.73503 | 0.006238 | 0.015179 |
| CNAG_03246 | 1751.861 | 0.208758 | 0.095268 | 2.191276 | 0.028432 | 0.057673 |
| CNAG_03247 | 2369.973 | 0.248574 | 0.07112 | 3.495133 | 0.000474 | 0.001501 |
| CNAG_03248 | 1615.98 | -0.41958 | 0.078215 | -5.36441 | 8.12E-08 | 4.83E-07 |
| CNAG_03249 | 5288.497 | -0.39334 | 0.063187 | -6.22504 | 4.81E-10 | 3.63E-09 |
| CNAG_03251 | 15235.79 | -0.13864 | 0.068275 | -2.03062 | 0.042293 | 0.081229 |
| CNAG_12745 | 15.21748 | -1.93327 | 0.58066 | -3.32944 | 0.00087 | 0.002602 |
| CNAG_12746 | 6.182892 | -2.28205 | 0.935287 | -2.43995 | 0.014689 | 0.032389 |
| CNAG_12748 | 65.0347 | -1.22443 | 0.284082 | -4.31013 | 1.63E-05 | 6.87E-05 |
| CNAG_03259 | 131.8268 | 1.153353 | 0.224919 | 5.127864 | 2.93E-07 | 1.62E-06 |
| CNAG_03261 | 3535.215 | -0.24791 | 0.066025 | -3.75481 | 0.000173 | 0.000606 |
| CNAG_03262 | 2053.874 | -0.27903 | 0.071594 | -3.89744 | 9.72E-05 | 0.000357 |
| CNAG_03263 | 14582.29 | 0.267896 | 0.062117 | 4.312786 | 1.61E-05 | 6.79E-05 |
| CNAG_03266 | 5218.36 | -0.25382 | 0.068445 | -3.70839 | 0.000209 | 0.000715 |
| CNAG_03268 | 2649.046 | -0.70411 | 0.104727 | -6.72332 | 1.78E-11 | 1.53E-10 |
| CNAG_03269 | 1187.77 | -0.25104 | 0.086463 | -2.90347 | 0.00369 | 0.009529 |
| CNAG_03271 | 3464.009 | -0.25168 | 0.064667 | -3.89199 | 9.94E-05 | 0.000365 |
| CNAG_03273 | 2191.356 | -0.19374 | 0.079973 | -2.42255 | 0.015412 | 0.033765 |
| CNAG_03275 | 914.4967 | 0.222039 | 0.104147 | 2.131976 | 0.033009 | 0.06555 |
| CNAG_03279 | 532.3013 | 0.83815 | 0.117194 | 7.151789 | 8.57E-13 | 8.23E-12 |
| CNAG_03280 | 978.4387 | 0.319849 | 0.094941 | 3.368941 | 0.000755 | 0.002293 |
| CNAG_03281 | 24387.14 | -0.23612 | 0.059653 | -3.95816 | 7.55E-05 | 0.000284 |
| CNAG_03282 | 1328.539 | -0.37596 | 0.084208 | -4.46469 | 8.02E-06 | 3.60E-05 |
| CNAG_03283 | 38806.68 | -0.31202 | 0.067495 | -4.62292 | 3.78E-06 | 1.79E-05 |
| CNAG_03284 | 1599.634 | 0.290964 | 0.078085 | 3.726263 | 0.000194 | 0.000672 |
| CNAG_03288 | 1348.69 | -0.60253 | 0.079018 | -7.62526 | 2.44E-14 | 2.60E-13 |
| CNAG_03289 | 5837.225 | 0.226987 | 0.059148 | 3.837598 | 0.000124 | 0.000446 |
| CNAG_03290 | 1207.076 | -0.20506 | 0.079835 | -2.56849 | 0.010214 | 0.023525 |
| CNAG_03291 | 1564.102 | 0.23745 | 0.079211 | 2.997683 | 0.00272 | 0.007264 |
| CNAG_03296 | 607.0495 | -0.47238 | 0.11834 | -3.9917 | 6.56E-05 | 0.000249 |
| CNAG_03297 | 2302.667 | -0.38581 | 0.08356 | -4.61721 | 3.89E-06 | 1.84E-05 |
| CNAG_03298 | 4476.919 | -0.24297 | 0.108581 | -2.23767 | 0.025242 | 0.051942 |
| CNAG_03300 | 2506.499 | -0.1491 | 0.075349 | -1.97873 | 0.047847 | 0.090295 |
| CNAG_03303 | 17906.41 | -0.15856 | 0.073553 | -2.15576 | 0.031102 | 0.062297 |
| CNAG_12753 | 234.5905 | -0.37103 | 0.155315 | -2.38892 | 0.016898 | 0.036597 |
| CNAG_03306 | 775.9346 | -0.65394 | 0.104804 | -6.23963 | 4.39E-10 | 3.33E-09 |
| CNAG_03308 | 1399.8 | 0.381929 | 0.086833 | 4.398415 | 1.09E-05 | 4.75E-05 |
| CNAG_03309 | 1765.774 | -0.69305 | 0.095193 | -7.28045 | 3.33E-13 | 3.31E-12 |
| CNAG_03314 | 1216.798 | 0.4491 | 0.100879 | 4.451864 | 8.51E-06 | 3.80E-05 |
| CNAG_03316 | 2156.581 | 0.276205 | 0.067685 | 4.08076 | 4.49E-05 | 0.000175 |
| CNAG_03317 | 2354.664 | -0.32835 | 0.069444 | -4.72828 | 2.26E-06 | 1.11E-05 |
| CNAG_03318 | 198.9116 | 1.890252 | 0.166109 | 11.37958 | 5.29E-30 | 1.09E-28 |
| CNAG_03319 | 3174.858 | -0.36849 | 0.098029 | -3.75902 | 0.000171 | 0.000597 |
| CNAG_03321 | 2156.737 | -0.16467 | 0.081925 | -2.00995 | 0.044437 | 0.084884 |

| | | | | | | |
|---|---|---|---|---|---|---|
| CNAG_03323 | 3840.949 | -0.32007 | 0.089885 | -3.56088 | 0.00037 | 0.001203 |
| CNAG_03328 | 1355.372 | -0.48173 | 0.107877 | -4.46555 | 7.99E-06 | 3.59E-05 |
| CNAG_03329 | 823.6213 | -0.25089 | 0.099892 | -2.5116 | 0.012019 | 0.027196 |
| CNAG_03331 | 2216.382 | -0.47924 | 0.073112 | -6.55484 | 5.57E-11 | 4.61E-10 |
| CNAG_03335 | 4562.354 | 0.341912 | 0.058114 | 5.883445 | 4.02E-09 | 2.77E-08 |
| CNAG_03337 | 1333.135 | 0.275728 | 0.089385 | 3.084712 | 0.002037 | 0.005597 |
| CNAG_03342 | 3742.511 | 0.145545 | 0.064936 | 2.241374 | 0.025002 | 0.051551 |
| CNAG_03343 | 1060.202 | -0.19055 | 0.09318 | -2.04495 | 0.04086 | 0.078839 |
| CNAG_03345 | 4098.498 | 0.293502 | 0.077019 | 3.810769 | 0.000139 | 0.000492 |
| CNAG_07723 | 59.76205 | -1.03806 | 0.294589 | -3.52374 | 0.000425 | 0.001364 |
| CNAG_03346 | 412.6671 | -1.81934 | 0.125492 | -14.4977 | 1.25E-47 | 4.03E-46 |
| CNAG_03347 | 2122.64 | 0.747257 | 0.106533 | 7.014306 | 2.31E-12 | 2.15E-11 |
| CNAG_03349 | 4086.508 | -0.19056 | 0.068165 | -2.7955 | 0.005182 | 0.012916 |
| CNAG_07724 | 1825.89 | -0.38544 | 0.098779 | -3.90206 | 9.54E-05 | 0.000352 |
| CNAG_03352 | 1020.417 | 0.390991 | 0.093086 | 4.200323 | 2.67E-05 | 0.000108 |
| CNAG_03353 | 2411.298 | -0.19492 | 0.073338 | -2.65786 | 0.007864 | 0.018651 |
| CNAG_03354 | 856.1046 | 0.224065 | 0.10084 | 2.221993 | 0.026284 | 0.053867 |
| CNAG_03355 | 5350.264 | 0.534478 | 0.071904 | 7.433227 | 1.06E-13 | 1.08E-12 |
| CNAG_03356 | 695.4869 | -0.37048 | 0.096185 | -3.85179 | 0.000117 | 0.000423 |
| CNAG_03357 | 1210.892 | -0.18577 | 0.088746 | -2.09325 | 0.036327 | 0.071182 |
| CNAG_03358 | 27827.09 | 0.679391 | 0.053206 | 12.76904 | 2.44E-37 | 6.18E-36 |
| CNAG_03359 | 2966.502 | 0.715945 | 0.095822 | 7.471621 | 7.92E-14 | 8.20E-13 |
| CNAG_03361 | 581.8787 | -0.21229 | 0.098077 | -2.16455 | 0.030422 | 0.061114 |
| CNAG_03363 | 1859.21 | 1.311373 | 0.085772 | 15.28901 | 9.05E-53 | 3.27E-51 |
| CNAG_03365 | 1742.146 | -0.41089 | 0.077689 | -5.28892 | 1.23E-07 | 7.12E-07 |
| CNAG_03366 | 463.8954 | 0.577908 | 0.115951 | 4.984073 | 6.23E-07 | 3.28E-06 |
| CNAG_12760 | 18.15565 | 1.077987 | 0.490766 | 2.19654 | 0.028053 | 0.057004 |
| CNAG_03367 | 1364.645 | -0.27657 | 0.075562 | -3.66012 | 0.000252 | 0.00085 |
| CNAG_12761 | 119.961 | -0.80329 | 0.2002 | -4.01245 | 6.01E-05 | 0.00023 |
| CNAG_03370 | 4315.321 | -0.46948 | 0.071731 | -6.54499 | 5.95E-11 | 4.90E-10 |
| CNAG_03372 | 1501.917 | -0.22585 | 0.098922 | -2.28313 | 0.022423 | 0.046815 |
| CNAG_03373 | 1686.272 | -0.25615 | 0.085653 | -2.99049 | 0.002785 | 0.007415 |
| CNAG_03375 | 5962.825 | -0.7152 | 0.065493 | -10.9202 | 9.23E-28 | 1.77E-26 |
| CNAG_03376 | 1272.1 | -0.26219 | 0.082938 | -3.16134 | 0.00157 | 0.004419 |
| CNAG_03378 | 2569.557 | 0.227516 | 0.069797 | 3.259665 | 0.001115 | 0.003244 |
| CNAG_03380 | 738.4693 | -0.22411 | 0.096616 | -2.31962 | 0.020362 | 0.042998 |
| CNAG_03381 | 675.8875 | 1.294686 | 0.115872 | 11.17339 | 5.50E-29 | 1.09E-27 |
| CNAG_03382 | 1455.667 | -0.40591 | 0.083398 | -4.86718 | 1.13E-06 | 5.75E-06 |
| CNAG_03386 | 2319.894 | 0.313796 | 0.069898 | 4.489325 | 7.14E-06 | 3.24E-05 |
| CNAG_03389 | 5275.637 | -0.19621 | 0.071079 | -2.76045 | 0.005772 | 0.014187 |
| CNAG_03390 | 1263.519 | -0.15418 | 0.076856 | -2.00606 | 0.04485 | 0.085513 |
| CNAG_03391 | 719.4727 | -0.30501 | 0.124003 | -2.45973 | 0.013904 | 0.030903 |
| CNAG_03392 | 455.9731 | 0.361378 | 0.124366 | 2.905768 | 0.003664 | 0.009468 |
| CNAG_03395 | 4538.935 | 2.176032 | 0.078235 | 27.81388 | 2.95E-170 | 4.52E-168 |
| CNAG_03397 | 2848.771 | -0.14639 | 0.062948 | -2.32551 | 0.020045 | 0.042467 |
| CNAG_03400 | 315.9039 | 1.010238 | 0.140696 | 7.180293 | 6.96E-13 | 6.72E-12 |
| CNAG_03401 | 1873.952 | 0.550971 | 0.074281 | 7.417349 | 1.19E-13 | 1.21E-12 |

| | | | | | | |
|---|---|---|---|---|---|---|
| CNAG_03405 | 2961.436 | 0.547496 | 0.072116 | 7.591891 | 3.15E-14 | 3.34E-13 |
| CNAG_03406 | 648.5251 | 0.268457 | 0.098863 | 2.715434 | 0.006619 | 0.015997 |
| CNAG_03407 | 32.41668 | -0.87253 | 0.371589 | -2.3481 | 0.018869 | 0.040297 |
| CNAG_03413 | 1093.767 | -0.71801 | 0.083904 | -8.55754 | 1.15E-17 | 1.45E-16 |
| CNAG_03416 | 1543.501 | -0.2277 | 0.082219 | -2.76947 | 0.005615 | 0.013859 |
| CNAG_03419 | 1587.026 | -0.28987 | 0.073693 | -3.93341 | 8.37E-05 | 0.000312 |
| CNAG_03420 | 1612.89 | 1.038702 | 0.082991 | 12.51589 | 6.11E-36 | 1.48E-34 |
| CNAG_03421 | 154.1653 | 0.626173 | 0.191852 | 3.263837 | 0.001099 | 0.003199 |
| CNAG_03425 | 1434.832 | 0.231369 | 0.102123 | 2.265579 | 0.023477 | 0.048766 |
| CNAG_03426 | 1870.482 | -0.93526 | 0.087546 | -10.6831 | 1.22E-26 | 2.22E-25 |
| CNAG_03427 | 536.1417 | 0.221985 | 0.102235 | 2.171323 | 0.029907 | 0.060199 |
| CNAG_03431 | 773.2234 | -0.44564 | 0.097648 | -4.56373 | 5.03E-06 | 2.33E-05 |
| CNAG_03432 | 1173.805 | 0.928347 | 0.119611 | 7.761356 | 8.40E-15 | 9.12E-14 |
| CNAG_03434 | 1386.722 | -0.41245 | 0.088089 | -4.68219 | 2.84E-06 | 1.37E-05 |
| CNAG_03435 | 9118.811 | 0.169206 | 0.070056 | 2.415309 | 0.015722 | 0.034333 |
| CNAG_03436 | 2575.009 | 0.447285 | 0.079976 | 5.592728 | 2.24E-08 | 1.41E-07 |
| CNAG_12770 | 865.544 | -0.81705 | 0.124664 | -6.55407 | 5.60E-11 | 4.62E-10 |
| CNAG_07733 | 25.82767 | 1.726485 | 0.490378 | 3.520724 | 0.00043 | 0.001377 |
| CNAG_03439 | 9211.138 | -0.17998 | 0.061597 | -2.92196 | 0.003478 | 0.00905 |
| CNAG_03440 | 416.8385 | -0.33483 | 0.116178 | -2.88205 | 0.003951 | 0.010115 |
| CNAG_03441 | 1493.923 | -0.18803 | 0.07679 | -2.44869 | 0.014338 | 0.031754 |
| CNAG_03446 | 700.1372 | -0.20856 | 0.10089 | -2.06722 | 0.038713 | 0.075278 |
| CNAG_03449 | 424.0807 | 0.321188 | 0.12023 | 2.671449 | 0.007552 | 0.018007 |
| CNAG_03450 | 2694.892 | 1.278572 | 0.077074 | 16.58892 | 8.38E-62 | 3.66E-60 |
| CNAG_03452 | 1668.031 | 0.448084 | 0.088637 | 5.055261 | 4.30E-07 | 2.32E-06 |
| CNAG_03454 | 2250.633 | -1.0415 | 0.106476 | -9.78157 | 1.35E-22 | 2.14E-21 |
| CNAG_03455 | 468.5504 | 0.743696 | 0.108362 | 6.863053 | 6.74E-12 | 6.03E-11 |
| CNAG_03456 | 1801.692 | -0.36476 | 0.080151 | -4.5509 | 5.34E-06 | 2.48E-05 |
| CNAG_03458 | 1790.094 | -0.62452 | 0.072612 | -8.60081 | 7.92E-18 | 1.00E-16 |
| CNAG_03459 | 6044.185 | -0.14608 | 0.066029 | -2.21238 | 0.02694 | 0.055073 |
| CNAG_03461 | 773.5077 | 2.087963 | 0.139771 | 14.93846 | 1.85E-50 | 6.35E-49 |
| CNAG_03463 | 2863.365 | 0.292807 | 0.069236 | 4.22914 | 2.35E-05 | 9.57E-05 |
| CNAG_03464 | 758.1376 | 3.402955 | 0.112538 | 30.23823 | 7.45E-201 | 1.48E-198 |
| CNAG_12774 | 18.79915 | 2.833285 | 0.587649 | 4.82139 | 1.43E-06 | 7.14E-06 |
| CNAG_12776 | 254.2729 | -0.29896 | 0.147348 | -2.02892 | 0.042467 | 0.081523 |
| CNAG_12777 | 22.17856 | 2.240813 | 0.498757 | 4.492799 | 7.03E-06 | 3.20E-05 |
| CNAG_07734 | 7.739304 | 3.986292 | 1.074392 | 3.710277 | 0.000207 | 0.000711 |
| CNAG_03465 | 2525.417 | 2.261959 | 0.081305 | 27.82055 | 2.45E-170 | 3.83E-168 |
| CNAG_03467 | 358.588 | -0.27715 | 0.122241 | -2.26727 | 0.023374 | 0.048576 |
| CNAG_07736 | 755.2722 | -1.55346 | 0.112328 | -13.8297 | 1.69E-43 | 5.14E-42 |
| CNAG_03473 | 481.8728 | 0.397535 | 0.107419 | 3.700773 | 0.000215 | 0.000735 |
| CNAG_03475 | 2464.046 | 1.03465 | 0.07135 | 14.50101 | 1.19E-47 | 3.85E-46 |
| CNAG_03476 | 10824.54 | -0.16286 | 0.063307 | -2.57254 | 0.010096 | 0.023285 |
| CNAG_12781 | 163.517 | -0.43153 | 0.178288 | -2.42042 | 0.015503 | 0.033936 |
| CNAG_03479 | 2366.2 | -0.32126 | 0.082173 | -3.90958 | 9.25E-05 | 0.000341 |
| CNAG_03480 | 781.144 | 1.223538 | 0.113652 | 10.76563 | 5.00E-27 | 9.24E-26 |
| CNAG_03481 | 1188.746 | 0.570882 | 0.080296 | 7.109703 | 1.16E-12 | 1.10E-11 |

| | | | | | | |
|---|---|---|---|---|---|---|
| CNAG_03482 | 48147.37 | 2.592493 | 0.054149 | 47.87699 | 0 | 0 |
| CNAG_03483 | 1664.727 | -0.22077 | 0.077083 | -2.86401 | 0.004183 | 0.010662 |
| CNAG_03484 | 1399.44 | -0.82013 | 0.110419 | -7.42744 | 1.11E-13 | 1.13E-12 |
| CNAG_03485 | 2140.941 | 0.889583 | 0.094895 | 9.374414 | 6.96E-21 | 1.04E-19 |
| CNAG_03486 | 4927.493 | 0.225494 | 0.076497 | 2.947752 | 0.003201 | 0.00839 |
| CNAG_03489 | 878.2861 | -0.45747 | 0.095474 | -4.79161 | 1.65E-06 | 8.25E-06 |
| CNAG_03492 | 2580.609 | 0.75579 | 0.311317 | 2.427723 | 0.015194 | 0.033368 |
| CNAG_03493 | 528.4586 | 0.397373 | 0.114566 | 3.468504 | 0.000523 | 0.001641 |
| CNAG_03494 | 1521.813 | -0.2488 | 0.08888 | -2.79925 | 0.005122 | 0.012775 |
| CNAG_12786 | 327.8924 | -0.84369 | 0.188066 | -4.48611 | 7.25E-06 | 3.29E-05 |
| CNAG_03498 | 7663.09 | 4.709352 | 0.083825 | 56.18106 | 0 | 0 |
| CNAG_12787 | 85.92498 | 0.86896 | 0.249666 | 3.480485 | 0.000501 | 0.001576 |
| CNAG_03502 | 3300.719 | 0.438271 | 0.103742 | 4.22463 | 2.39E-05 | 9.75E-05 |
| CNAG_03503 | 931.7551 | -0.34651 | 0.118366 | -2.92742 | 0.003418 | 0.008904 |
| CNAG_03508 | 1055.206 | -0.40591 | 0.094656 | -4.28828 | 1.80E-05 | 7.54E-05 |
| CNAG_03509 | 4873.037 | -0.23783 | 0.067348 | -3.53128 | 0.000414 | 0.00133 |
| CNAG_03510 | 26558.93 | -0.31375 | 0.072304 | -4.33925 | 1.43E-05 | 6.10E-05 |
| CNAG_03511 | 316.895 | 1.047814 | 0.141014 | 7.430576 | 1.08E-13 | 1.10E-12 |
| CNAG_03512 | 1729.447 | 0.284768 | 0.079322 | 3.590018 | 0.000331 | 0.001088 |
| CNAG_12791 | 2.748698 | 3.001983 | 1.504295 | 1.995608 | 0.045977 | 0.087313 |
| CNAG_07743 | 73.25834 | 0.742384 | 0.253828 | 2.924748 | 0.003447 | 0.008975 |
| CNAG_12792 | 13.5 | -1.62452 | 0.615964 | -2.63736 | 0.008355 | 0.019696 |
| CNAG_03520 | 250.0637 | -0.44499 | 0.152609 | -2.91588 | 0.003547 | 0.00919 |
| CNAG_03522 | 256.6275 | -0.38409 | 0.154625 | -2.48398 | 0.012992 | 0.029188 |
| CNAG_03524 | 3095.827 | -0.6162 | 0.064326 | -9.57937 | 9.76E-22 | 1.50E-20 |
| CNAG_03526 | 265.1309 | 0.438979 | 0.156041 | 2.813227 | 0.004905 | 0.012274 |
| CNAG_03533 | 1123.615 | -0.54478 | 0.089843 | -6.06367 | 1.33E-09 | 9.61E-09 |
| CNAG_03534 | 328.2239 | 0.55876 | 0.131697 | 4.242779 | 2.21E-05 | 9.06E-05 |
| CNAG_07744 | 2196.712 | 0.143679 | 0.070209 | 2.046455 | 0.040712 | 0.078637 |
| CNAG_07745 | 4140.008 | 1.470661 | 0.094275 | 15.59974 | 7.31E-55 | 2.72E-53 |
| CNAG_07746 | 10376.49 | 0.126943 | 0.058721 | 2.161799 | 0.030634 | 0.061444 |
| CNAG_03539 | 1305.68 | 0.505807 | 0.078141 | 6.472995 | 9.61E-11 | 7.79E-10 |
| CNAG_03541 | 1703.818 | 0.152187 | 0.073717 | 2.064459 | 0.038974 | 0.075713 |
| CNAG_03542 | 1885.163 | 0.42797 | 0.080743 | 5.300384 | 1.16E-07 | 6.71E-07 |
| CNAG_12795 | 332.0213 | 0.766334 | 0.142907 | 5.362459 | 8.21E-08 | 4.88E-07 |
| CNAG_03543 | 3433.371 | 1.70896 | 0.094621 | 18.06103 | 6.46E-73 | 3.30E-71 |
| CNAG_03544 | 7895.165 | 0.563882 | 0.070397 | 8.010031 | 1.15E-15 | 1.32E-14 |
| CNAG_12796 | 26.98867 | 1.505479 | 0.46884 | 3.211069 | 0.001322 | 0.003779 |
| CNAG_03550 | 339.9965 | -0.66996 | 0.122092 | -5.48734 | 4.08E-08 | 2.50E-07 |
| CNAG_03552 | 1304.831 | 1.549863 | 0.095423 | 16.24197 | 2.55E-59 | 1.07E-57 |
| CNAG_12799 | 20.08189 | 2.047167 | 0.506966 | 4.038074 | 5.39E-05 | 0.000207 |
| CNAG_03553 | 5237.363 | 0.236812 | 0.068338 | 3.465298 | 0.00053 | 0.001659 |
| CNAG_03555 | 482.949 | 0.253543 | 0.126855 | 1.998686 | 0.045642 | 0.08682 |
| CNAG_03556 | 308.7895 | -0.48272 | 0.147675 | -3.26876 | 0.00108 | 0.003148 |
| CNAG_03557 | 898.1946 | 0.202693 | 0.087616 | 2.313411 | 0.0207 | 0.043599 |
| CNAG_03560 | 409.6252 | 1.338128 | 0.132311 | 10.11349 | 4.81E-24 | 8.12E-23 |
| CNAG_03562 | 592.2024 | 0.36385 | 0.097792 | 3.720664 | 0.000199 | 0.000685 |

| | | | | | | |
|---|---|---|---|---|---|---|
| CNAG_03565 | 3590.595 | 0.943045 | 0.099284 | 9.49842 | 2.13E-21 | 3.24E-20 |
| CNAG_03566 | 2614.121 | -1.05468 | 0.244188 | -4.31915 | 1.57E-05 | 6.62E-05 |
| CNAG_03567 | 2733.325 | 0.300895 | 0.072268 | 4.163607 | 3.13E-05 | 0.000125 |
| CNAG_03570 | 163.4919 | 1.83487 | 0.275903 | 6.650412 | 2.92E-11 | 2.48E-10 |
| CNAG_12801 | 76.49322 | 2.401483 | 0.418616 | 5.736716 | 9.65E-09 | 6.36E-08 |
| CNAG_03573 | 677.1761 | 0.766967 | 0.100062 | 7.664949 | 1.79E-14 | 1.92E-13 |
| CNAG_03575 | 562.4521 | -0.35021 | 0.107061 | -3.27111 | 0.001071 | 0.003125 |
| CNAG_03577 | 54884.43 | -0.14643 | 0.059142 | -2.47586 | 0.013291 | 0.029731 |
| CNAG_03576 | 55.57131 | -0.72064 | 0.29818 | -2.4168 | 0.015657 | 0.034212 |
| CNAG_03578 | 1510.129 | 0.686462 | 0.079623 | 8.621399 | 6.61E-18 | 8.39E-17 |
| CNAG_07749 | 750.9711 | -0.43258 | 0.091225 | -4.74191 | 2.12E-06 | 1.04E-05 |
| CNAG_03583 | 826.8051 | -0.30439 | 0.098204 | -3.09958 | 0.001938 | 0.00536 |
| CNAG_03584 | 2364.431 | -0.39946 | 0.080948 | -4.93473 | 8.03E-07 | 4.17E-06 |
| CNAG_07996 | 1016.919 | -0.40099 | 0.082911 | -4.83642 | 1.32E-06 | 6.65E-06 |
| CNAG_03586 | 513.6189 | -0.35198 | 0.105792 | -3.32712 | 0.000877 | 0.002619 |
| CNAG_03587 | 667.4555 | -0.27866 | 0.108224 | -2.57488 | 0.010027 | 0.023148 |
| CNAG_03589 | 1769.42 | 0.364623 | 0.087023 | 4.189956 | 2.79E-05 | 0.000112 |
| CNAG_03590 | 2565.712 | -0.30527 | 0.086019 | -3.54887 | 0.000387 | 0.001252 |
| CNAG_03591 | 4956.451 | -0.22495 | 0.065961 | -3.4104 | 0.000649 | 0.001997 |
| CNAG_07751 | 2455.128 | 2.49988 | 0.075991 | 32.89688 | 2.44E-237 | 6.60E-235 |
| CNAG_12806 | 59.76047 | 4.831634 | 0.479281 | 10.081 | 6.70E-24 | 1.12E-22 |
| CNAG_12807 | 126.0081 | 5.293662 | 1.124631 | 4.707022 | 2.51E-06 | 1.22E-05 |
| CNAG_06977 | 141.5437 | 0.430764 | 0.20206 | 2.131861 | 0.033018 | 0.065553 |
| CNAG_06978 | 293.1094 | 0.494608 | 0.157675 | 3.136879 | 0.001708 | 0.004769 |
| CNAG_06993 | 3787.32 | -0.29224 | 0.092969 | -3.14339 | 0.00167 | 0.004676 |
| CNAG_06994 | 982.8045 | -0.3044 | 0.091339 | -3.33261 | 0.00086 | 0.002577 |
| CNAG_06966 | 38.04036 | -1.84571 | 0.402529 | -4.58528 | 4.53E-06 | 2.12E-05 |
| CNAG_06967 | 2326.867 | -0.46149 | 0.066552 | -6.93427 | 4.08E-12 | 3.70E-11 |
| CNAG_06968 | 1458.4 | -0.82254 | 0.105042 | -7.83062 | 4.85E-15 | 5.36E-14 |
| CNAG_06969 | 996.9388 | -0.59983 | 0.092518 | -6.48341 | 8.97E-11 | 7.29E-10 |
| CNAG_06942 | 540.7463 | -0.58551 | 0.103852 | -5.63787 | 1.72E-08 | 1.10E-07 |
| CNAG_06943 | 222.7318 | 0.412769 | 0.145441 | 2.83805 | 0.004539 | 0.011461 |
| CNAG_04100 | 118.7084 | -0.75129 | 0.195836 | -3.83631 | 0.000125 | 0.000449 |
| CNAG_04101 | 1814.781 | -0.89599 | 0.074144 | -12.0845 | 1.27E-33 | 2.87E-32 |
| CNAG_04102 | 355.3392 | -0.6172 | 0.125693 | -4.91036 | 9.09E-07 | 4.68E-06 |
| CNAG_04103 | 423.4972 | 2.334165 | 0.169438 | 13.77595 | 3.56E-43 | 1.07E-41 |
| CNAG_04104 | 1075.31 | 0.357138 | 0.095059 | 3.757025 | 0.000172 | 0.000601 |
| CNAG_04108 | 1133.192 | -0.30219 | 0.101667 | -2.97233 | 0.002955 | 0.007812 |
| CNAG_04110 | 441.0806 | -0.22997 | 0.117114 | -1.96368 | 0.049567 | 0.093001 |
| CNAG_04111 | 2338.893 | -0.20135 | 0.069747 | -2.88688 | 0.003891 | 0.009989 |
| CNAG_12812 | 95.75616 | 0.890112 | 0.293856 | 3.029072 | 0.002453 | 0.006615 |
| CNAG_04112 | 439.5254 | -0.56166 | 0.142599 | -3.93877 | 8.19E-05 | 0.000306 |
| CNAG_04113 | 3742.86 | -0.20319 | 0.07831 | -2.5947 | 0.009467 | 0.022042 |
| CNAG_04115 | 1297.729 | 0.355449 | 0.087435 | 4.065299 | 4.80E-05 | 0.000186 |
| CNAG_04122 | 224.1128 | -0.77591 | 0.163578 | -4.7434 | 2.10E-06 | 1.03E-05 |
| CNAG_12817 | 25.84617 | -1.15188 | 0.41767 | -2.75788 | 0.005818 | 0.014278 |
| CNAG_04131 | 3118.811 | 0.139247 | 0.068288 | 2.039123 | 0.041438 | 0.079774 |

| | | | | | | |
|---|---|---|---|---|---|---|
| CNAG_04132 | 2602.374 | -0.41777 | 0.070947 | -5.88841 | 3.90E-09 | 2.69E-08 |
| CNAG_07757 | 1401.705 | 0.266031 | 0.083418 | 3.189122 | 0.001427 | 0.00405 |
| CNAG_04137 | 250.9376 | -0.55847 | 0.143809 | -3.88343 | 0.000103 | 0.000376 |
| CNAG_04139 | 123.1084 | -0.65067 | 0.215143 | -3.02436 | 0.002492 | 0.006704 |
| CNAG_12820 | 65.65765 | -0.92004 | 0.269164 | -3.41814 | 0.00063 | 0.001948 |
| CNAG_04141 | 2908.962 | -0.96125 | 0.07028 | -13.6775 | 1.38E-42 | 4.05E-41 |
| CNAG_04142 | 1869.106 | 0.476033 | 0.117196 | 4.061862 | 4.87E-05 | 0.000189 |
| CNAG_04143 | 29.80756 | 1.354155 | 0.391983 | 3.454625 | 0.000551 | 0.001725 |
| CNAG_12821 | 524.5282 | 0.489239 | 0.139113 | 3.516836 | 0.000437 | 0.001396 |
| CNAG_07758 | 505.8195 | 0.285608 | 0.10725 | 2.663019 | 0.007744 | 0.018416 |
| CNAG_04148 | 1833.809 | 0.193378 | 0.07458 | 2.592879 | 0.009518 | 0.022109 |
| CNAG_04150 | 1690.796 | -0.52216 | 0.082487 | -6.33021 | 2.45E-10 | 1.91E-09 |
| CNAG_04151 | 2558.041 | -0.13804 | 0.070258 | -1.96471 | 0.049448 | 0.092843 |
| CNAG_04152 | 837.9981 | -0.37823 | 0.087317 | -4.33176 | 1.48E-05 | 6.29E-05 |
| CNAG_04155 | 2619.717 | -0.37984 | 0.090236 | -4.20938 | 2.56E-05 | 0.000104 |
| CNAG_04156 | 2308.718 | -0.38869 | 0.075435 | -5.1526 | 2.57E-07 | 1.43E-06 |
| CNAG_04157 | 905.4701 | -0.37027 | 0.110257 | -3.35824 | 0.000784 | 0.002369 |
| CNAG_04158 | 1418.989 | -0.20521 | 0.081041 | -2.53211 | 0.011338 | 0.0259 |
| CNAG_04159 | 1366.832 | -0.2172 | 0.078813 | -2.7559 | 0.005853 | 0.014351 |
| CNAG_04161 | 1005.554 | -0.28167 | 0.082892 | -3.39803 | 0.000679 | 0.002083 |
| CNAG_04163 | 1998.03 | -0.90442 | 0.251219 | -3.60011 | 0.000318 | 0.00105 |
| CNAG_04164 | 1076.06 | -0.18332 | 0.08285 | -2.21262 | 0.026924 | 0.055053 |
| CNAG_04165 | 437.3919 | -0.30769 | 0.138178 | -2.22675 | 0.025964 | 0.053292 |
| CNAG_04166 | 1836.174 | -0.2316 | 0.082637 | -2.80258 | 0.00507 | 0.012659 |
| CNAG_04167 | 170.4251 | 0.671214 | 0.172392 | 3.893541 | 9.88E-05 | 0.000363 |
| CNAG_04168 | 6472.061 | -1.27106 | 0.084299 | -15.078 | 2.26E-51 | 7.82E-50 |
| CNAG_04169 | 545.6578 | 0.639088 | 0.109492 | 5.836826 | 5.32E-09 | 3.62E-08 |
| CNAG_04170 | 682.6 | 0.306357 | 0.106157 | 2.885892 | 0.003903 | 0.010017 |
| CNAG_04171 | 3133.066 | 0.308297 | 0.079344 | 3.885569 | 0.000102 | 0.000373 |
| CNAG_04172 | 765.9888 | -0.31064 | 0.117144 | -2.65176 | 0.008007 | 0.018956 |
| CNAG_04175 | 2807.24 | -0.2286 | 0.09338 | -2.44811 | 0.014361 | 0.031788 |
| CNAG_04177 | 928.5381 | -0.19899 | 0.088932 | -2.23757 | 0.025249 | 0.051942 |
| CNAG_04178 | 413.0029 | -0.76698 | 0.144462 | -5.30921 | 1.10E-07 | 6.42E-07 |
| CNAG_04180 | 2559.599 | -0.19653 | 0.069462 | -2.82936 | 0.004664 | 0.011752 |
| CNAG_12827 | 1144.077 | -0.29415 | 0.11225 | -2.6205 | 0.00878 | 0.020614 |
| CNAG_04184 | 908.7566 | 0.816952 | 0.088053 | 9.277965 | 1.73E-20 | 2.51E-19 |
| CNAG_07997 | 114.1534 | 0.719581 | 0.24397 | 2.949465 | 0.003183 | 0.008352 |
| CNAG_04185 | 843.0872 | 0.928566 | 0.155733 | 5.96256 | 2.48E-09 | 1.74E-08 |
| CNAG_04186 | 939.4431 | 1.013128 | 0.109825 | 9.224922 | 2.84E-20 | 4.10E-19 |
| CNAG_04187 | 1574.288 | 0.530502 | 0.075934 | 6.986339 | 2.82E-12 | 2.59E-11 |
| CNAG_04189 | 20311.82 | 1.286983 | 0.057847 | 22.24813 | 1.18E-109 | 9.75E-108 |
| CNAG_04190 | 5780.054 | -0.19292 | 0.064512 | -2.99052 | 0.002785 | 0.007415 |
| CNAG_04191 | 1647.322 | -0.2981 | 0.082324 | -3.62109 | 0.000293 | 0.000973 |
| CNAG_04192 | 7765.069 | -0.17881 | 0.056783 | -3.14902 | 0.001638 | 0.004593 |
| CNAG_04193 | 1150.005 | 0.235504 | 0.081515 | 2.889084 | 0.003864 | 0.009925 |
| CNAG_04194 | 4551.731 | -0.16359 | 0.06466 | -2.52995 | 0.011408 | 0.026009 |
| CNAG_04195 | 2100.156 | 2.038069 | 0.080449 | 25.3337 | 1.36E-141 | 1.62E-139 |

| | | | | | | |
|---|---|---|---|---|---|---|
| CNAG_04196 | 3956.058 | -0.34751 | 0.07291 | -4.76636 | 1.88E-06 | 9.27E-06 |
| CNAG_04198 | 1363.661 | 0.276281 | 0.0848 | 3.258024 | 0.001122 | 0.003261 |
| CNAG_04200 | 886.3866 | 1.522187 | 0.10995 | 13.84437 | 1.38E-43 | 4.20E-42 |
| CNAG_04201 | 177.3294 | 1.928538 | 0.267147 | 7.219014 | 5.24E-13 | 5.13E-12 |
| CNAG_04202 | 1969.384 | 1.568638 | 0.087146 | 18.00009 | 1.95E-72 | 9.76E-71 |
| CNAG_04204 | 1321.856 | -0.30337 | 0.09855 | -3.0783 | 0.002082 | 0.005703 |
| CNAG_04205 | 1773.113 | -0.30468 | 0.089095 | -3.41969 | 0.000627 | 0.001943 |
| CNAG_04206 | 635.7365 | 1.274941 | 0.108691 | 11.72991 | 8.96E-32 | 1.91E-30 |
| CNAG_12830 | 67.41798 | 0.935317 | 0.296337 | 3.156258 | 0.001598 | 0.004493 |
| CNAG_04208 | 1005.587 | -0.41827 | 0.089619 | -4.66718 | 3.05E-06 | 1.46E-05 |
| CNAG_04209 | 4740.993 | 0.850339 | 0.098377 | 8.643697 | 5.44E-18 | 6.94E-17 |
| CNAG_04210 | 4982.188 | 2.554371 | 0.080569 | 31.7043 | 1.36E-220 | 3.24E-218 |
| CNAG_04215 | 13216.55 | 0.298071 | 0.059167 | 5.037793 | 4.71E-07 | 2.53E-06 |
| CNAG_12832 | 129.7409 | 0.914195 | 0.234492 | 3.898626 | 9.67E-05 | 0.000356 |
| CNAG_04217 | 2120.597 | 1.115081 | 0.314747 | 3.542788 | 0.000396 | 0.001276 |
| CNAG_04218 | 154.676 | -0.68627 | 0.19001 | -3.61177 | 0.000304 | 0.001006 |
| CNAG_04219 | 4750.55 | 0.765423 | 0.068802 | 11.12507 | 9.47E-29 | 1.86E-27 |
| CNAG_04221 | 1914.536 | 0.429603 | 0.069939 | 6.142558 | 8.12E-10 | 5.99E-09 |
| CNAG_04222 | 5056.236 | 0.142885 | 0.072301 | 1.976242 | 0.048127 | 0.090787 |
| CNAG_04224 | 2299.048 | -0.66885 | 0.082957 | -8.06263 | 7.47E-16 | 8.63E-15 |
| CNAG_04225 | 249.7219 | 0.5438 | 0.203085 | 2.6777 | 0.007413 | 0.017706 |
| CNAG_04227 | 1424.233 | 0.197497 | 0.080183 | 2.46307 | 0.013775 | 0.030686 |
| CNAG_04228 | 244.1602 | 0.713495 | 0.174322 | 4.092977 | 4.26E-05 | 0.000167 |
| CNAG_04231 | 315.0142 | -0.37308 | 0.133831 | -2.78771 | 0.005308 | 0.013194 |
| CNAG_04234 | 799.8823 | -0.19415 | 0.096848 | -2.00473 | 0.044992 | 0.085763 |
| CNAG_04236 | 3819.328 | -0.20452 | 0.067572 | -3.02676 | 0.002472 | 0.006657 |
| CNAG_04237 | 706.9689 | -0.48962 | 0.119058 | -4.1125 | 3.91E-05 | 0.000155 |
| CNAG_04239 | 356.1775 | -0.46627 | 0.153393 | -3.03967 | 0.002368 | 0.00641 |
| CNAG_04240 | 290.4571 | -0.34435 | 0.14929 | -2.30661 | 0.021077 | 0.044266 |
| CNAG_04241 | 2502.326 | 2.683379 | 0.077486 | 34.63029 | 8.85E-263 | 3.42E-260 |
| CNAG_04242 | 1468.773 | 1.801944 | 0.087771 | 20.52996 | 1.16E-93 | 7.68E-92 |
| CNAG_12835 | 20.56384 | 1.199186 | 0.486872 | 2.463043 | 0.013776 | 0.030686 |
| CNAG_04243 | 2023.651 | -0.2572 | 0.08694 | -2.95837 | 0.003093 | 0.00814 |
| CNAG_04244 | 1058.983 | 0.526694 | 0.099354 | 5.301205 | 1.15E-07 | 6.69E-07 |
| CNAG_04245 | 8885.769 | 0.292654 | 0.061401 | 4.766232 | 1.88E-06 | 9.27E-06 |
| CNAG_04246 | 159.6716 | 0.587798 | 0.190548 | 3.08478 | 0.002037 | 0.005597 |
| CNAG_04248 | 527.4485 | -0.35116 | 0.108235 | -3.24446 | 0.001177 | 0.003404 |
| CNAG_07763 | 438.9119 | -0.61091 | 0.113942 | -5.36158 | 8.25E-08 | 4.89E-07 |
| CNAG_04253 | 548.888 | -0.51672 | 0.123744 | -4.17572 | 2.97E-05 | 0.000119 |
| CNAG_04256 | 287.6402 | 0.424661 | 0.149038 | 2.849354 | 0.004381 | 0.01112 |
| CNAG_04257 | 6691.423 | 0.908518 | 0.059833 | 15.1843 | 4.49E-52 | 1.59E-50 |
| CNAG_04259 | 6759.154 | -0.32409 | 0.084814 | -3.82124 | 0.000133 | 0.000473 |
| CNAG_04262 | 355.3273 | 0.45497 | 0.130238 | 3.493377 | 0.000477 | 0.00151 |
| CNAG_04263 | 520.9823 | -0.38341 | 0.134759 | -2.84512 | 0.004439 | 0.011245 |
| CNAG_04264 | 853.2895 | -0.17398 | 0.087797 | -1.98157 | 0.047527 | 0.089837 |
| CNAG_04265 | 2683.983 | -0.34668 | 0.085106 | -4.07354 | 4.63E-05 | 0.00018 |
| CNAG_04267 | 1445.415 | 0.770674 | 0.077913 | 9.891466 | 4.53E-23 | 7.44E-22 |

| | | | | | | |
|---|---|---|---|---|---|---|
| CNAG_12838 | 24.69194 | 2.22658 | 0.457932 | 4.862246 | 1.16E-06 | 5.89E-06 |
| CNAG_04269 | 1122.592 | -0.21827 | 0.100812 | -2.16513 | 0.030378 | 0.061056 |
| CNAG_04271 | 978.387 | 0.703585 | 0.092104 | 7.639033 | 2.19E-14 | 2.34E-13 |
| CNAG_04272 | 2688.908 | 0.599222 | 0.071697 | 8.357683 | 6.40E-17 | 7.79E-16 |
| CNAG_04273 | 1974.393 | -0.43477 | 0.06839 | -6.35722 | 2.05E-10 | 1.61E-09 |
| CNAG_04275 | 800.786 | -0.35143 | 0.142299 | -2.46962 | 0.013526 | 0.030186 |
| CNAG_04276 | 2514.372 | 0.196657 | 0.068809 | 2.858027 | 0.004263 | 0.010858 |
| CNAG_04277 | 1235.368 | 0.661379 | 0.090251 | 7.328243 | 2.33E-13 | 2.33E-12 |
| CNAG_04278 | 1474.891 | 0.377131 | 0.076313 | 4.941917 | 7.74E-07 | 4.03E-06 |
| CNAG_04281 | 2450.358 | -0.1604 | 0.075735 | -2.11786 | 0.034187 | 0.067445 |
| CNAG_04284 | 4343.728 | -0.66153 | 0.083769 | -7.89706 | 2.86E-15 | 3.19E-14 |
| CNAG_04285 | 709.8897 | -0.37669 | 0.126298 | -2.98258 | 0.002858 | 0.007587 |
| CNAG_04286 | 2512.025 | -0.281 | 0.08016 | -3.5055 | 0.000456 | 0.00145 |
| CNAG_04288 | 710.0328 | -0.32293 | 0.112595 | -2.86811 | 0.004129 | 0.010545 |
| CNAG_04289 | 4082.679 | -0.29491 | 0.083976 | -3.51181 | 0.000445 | 0.00142 |
| CNAG_04294 | 347.7864 | 0.565691 | 0.122393 | 4.621928 | 3.80E-06 | 1.80E-05 |
| CNAG_04297 | 752.9171 | 0.595135 | 0.10781 | 5.520222 | 3.39E-08 | 2.10E-07 |
| CNAG_04299 | 634.3847 | -0.34879 | 0.103248 | -3.37822 | 0.00073 | 0.002228 |
| CNAG_04302 | 158.7116 | -0.47456 | 0.194969 | -2.43402 | 0.014932 | 0.032865 |
| CNAG_04303 | 4372.933 | -0.15491 | 0.067711 | -2.28788 | 0.022144 | 0.046365 |
| CNAG_04304 | 7655.161 | -0.18002 | 0.062417 | -2.88419 | 0.003924 | 0.010062 |
| CNAG_04305 | 525.59 | -0.2676 | 0.116902 | -2.28911 | 0.022073 | 0.046239 |
| CNAG_04307 | 318.0846 | 3.635913 | 0.169571 | 21.44178 | 5.45E-102 | 4.14E-100 |
| CNAG_04309 | 353.923 | 0.616953 | 0.13831 | 4.460654 | 8.17E-06 | 3.66E-05 |
| CNAG_04310 | 1030.921 | -0.17349 | 0.085847 | -2.02086 | 0.043294 | 0.082896 |
| CNAG_04312 | 2523.024 | 0.484981 | 0.074456 | 6.513643 | 7.33E-11 | 5.99E-10 |
| CNAG_04313 | 457.2462 | 0.908217 | 0.125339 | 7.246064 | 4.29E-13 | 4.26E-12 |
| CNAG_04315 | 2656.986 | 0.481986 | 0.086031 | 5.60244 | 2.11E-08 | 1.33E-07 |
| CNAG_04319 | 256.0929 | -0.35296 | 0.142141 | -2.48314 | 0.013023 | 0.029241 |
| CNAG_04320 | 2657.543 | 0.570161 | 0.063414 | 8.99108 | 2.45E-19 | 3.36E-18 |
| CNAG_04321 | 1986.855 | 0.384393 | 0.083916 | 4.580682 | 4.63E-06 | 2.16E-05 |
| CNAG_04322 | 1234.27 | 0.736481 | 0.325359 | 2.263593 | 0.023599 | 0.048995 |
| CNAG_04325 | 7426.228 | 3.795506 | 0.11419 | 33.23848 | 3.00E-242 | 8.70E-240 |
| CNAG_04327 | 7223.642 | 0.172847 | 0.060641 | 2.850317 | 0.004368 | 0.01109 |
| CNAG_04330 | 52.33667 | 1.020008 | 0.331531 | 3.076661 | 0.002093 | 0.005733 |
| CNAG_12843 | 44.08418 | 0.829396 | 0.344001 | 2.411028 | 0.015908 | 0.034701 |
| CNAG_04333 | 138.4455 | 0.935351 | 0.249192 | 3.753541 | 0.000174 | 0.000608 |
| CNAG_04334 | 2479.42 | -0.21427 | 0.090674 | -2.36312 | 0.018122 | 0.038915 |
| CNAG_04335 | 1740.468 | -0.23647 | 0.076077 | -3.10827 | 0.001882 | 0.005222 |
| CNAG_04336 | 343.5665 | -0.41053 | 0.149339 | -2.74899 | 0.005978 | 0.014618 |
| CNAG_04338 | 466.1821 | -0.48068 | 0.123598 | -3.88905 | 0.000101 | 0.000368 |
| CNAG_04340 | 1943.17 | -0.23142 | 0.075331 | -3.07212 | 0.002125 | 0.005807 |
| CNAG_04341 | 1807.322 | -0.18147 | 0.07671 | -2.36564 | 0.017999 | 0.038693 |
| CNAG_12845 | 23.60756 | 3.38583 | 0.563035 | 6.013531 | 1.82E-09 | 1.29E-08 |
| CNAG_04344 | 423.1715 | 0.685727 | 0.147522 | 4.648308 | 3.35E-06 | 1.59E-05 |
| CNAG_04345 | 759.7339 | 0.297079 | 0.14496 | 2.049383 | 0.040425 | 0.07822 |
| CNAG_04347 | 5348.149 | -0.19321 | 0.060113 | -3.21412 | 0.001308 | 0.003746 |

| | | | | | | |
|---|---|---|---|---|---|---|
| CNAG_04348 | 3142.573 | 0.502338 | 0.07204 | 6.973059 | 3.10E-12 | 2.83E-11 |
| CNAG_04349 | 1552.564 | 0.388105 | 0.083783 | 4.632256 | 3.62E-06 | 1.72E-05 |
| CNAG_12847 | 9.522774 | 2.036823 | 0.810752 | 2.512265 | 0.011996 | 0.027167 |
| CNAG_04350 | 1212.217 | 1.267266 | 0.09034 | 14.02768 | 1.06E-44 | 3.27E-43 |
| CNAG_04351 | 421.2563 | 0.389005 | 0.120463 | 3.229235 | 0.001241 | 0.003565 |
| CNAG_04358 | 3070.463 | -0.27063 | 0.074682 | -3.62376 | 0.00029 | 0.000964 |
| CNAG_04359 | 1237.44 | -0.31263 | 0.094671 | -3.30223 | 0.000959 | 0.002839 |
| CNAG_04360 | 1641.37 | -0.29446 | 0.076623 | -3.843 | 0.000122 | 0.000438 |
| CNAG_04361 | 4322.155 | -0.76218 | 0.087825 | -8.67841 | 4.01E-18 | 5.17E-17 |
| CNAG_04362 | 3615.74 | -0.25901 | 0.063245 | -4.09529 | 4.22E-05 | 0.000165 |
| CNAG_04364 | 4436.238 | 0.303629 | 0.077221 | 3.931974 | 8.43E-05 | 0.000314 |
| CNAG_04365 | 7113.379 | 0.13798 | 0.062991 | 2.19048 | 0.028489 | 0.057761 |
| CNAG_04368 | 1290.765 | -0.31482 | 0.086776 | -3.62798 | 0.000286 | 0.00095 |
| CNAG_04369 | 289.041 | -0.54094 | 0.134241 | -4.02962 | 5.59E-05 | 0.000215 |
| CNAG_04372 | 1882.496 | -0.18327 | 0.075292 | -2.43419 | 0.014925 | 0.032858 |
| CNAG_04373 | 8005.141 | 0.407245 | 0.060918 | 6.685128 | 2.31E-11 | 1.96E-10 |
| CNAG_04375 | 2652.021 | -0.14816 | 0.068371 | -2.16707 | 0.03023 | 0.060789 |
| CNAG_04377 | 2005.513 | -0.22802 | 0.096141 | -2.37169 | 0.017707 | 0.038115 |
| CNAG_04380 | 1348.627 | 0.306475 | 0.084441 | 3.629458 | 0.000284 | 0.000946 |
| CNAG_04382 | 1545.029 | 0.172736 | 0.077075 | 2.241136 | 0.025017 | 0.05157 |
| CNAG_04383 | 1399.147 | -0.21514 | 0.086896 | -2.47585 | 0.013292 | 0.029731 |
| CNAG_04387 | 719.5862 | -0.73444 | 0.098388 | -7.46467 | 8.35E-14 | 8.62E-13 |
| CNAG_07766 | 697.1233 | 0.380223 | 0.099237 | 3.831462 | 0.000127 | 0.000456 |
| CNAG_07765 | 135.9897 | -0.94572 | 0.302321 | -3.12819 | 0.001759 | 0.004904 |
| CNAG_04393 | 2277.012 | 0.230832 | 0.07221 | 3.19666 | 0.00139 | 0.003958 |
| CNAG_04396 | 301.28 | -0.33203 | 0.13415 | -2.47506 | 0.013321 | 0.029787 |
| CNAG_04397 | 463.9313 | -0.30507 | 0.111781 | -2.7292 | 0.006349 | 0.015418 |
| CNAG_04398 | 1141.478 | -0.42737 | 0.087765 | -4.86943 | 1.12E-06 | 5.70E-06 |
| CNAG_04399 | 963.0344 | -0.30041 | 0.095796 | -3.13595 | 0.001713 | 0.004781 |
| CNAG_04401 | 2261.805 | -0.57424 | 0.076754 | -7.48157 | 7.34E-14 | 7.62E-13 |
| CNAG_04402 | 1836.793 | -0.25314 | 0.088789 | -2.851 | 0.004358 | 0.01107 |
| CNAG_04405 | 936.4631 | -0.33805 | 0.096536 | -3.5018 | 0.000462 | 0.001468 |
| CNAG_04409 | 345.8262 | -0.62907 | 0.15956 | -3.94255 | 8.06E-05 | 0.000301 |
| CNAG_04411 | 565.8979 | 0.245613 | 0.117819 | 2.084665 | 0.0371 | 0.072539 |
| CNAG_04415 | 253.6549 | -0.35361 | 0.146616 | -2.41183 | 0.015873 | 0.034643 |
| CNAG_04416 | 135.1849 | 2.365735 | 0.217715 | 10.86618 | 1.67E-27 | 3.16E-26 |
| CNAG_04417 | 976.4353 | 1.368636 | 0.134151 | 10.20222 | 1.94E-24 | 3.31E-23 |
| CNAG_12853 | 219.0377 | -1.03217 | 0.159241 | -6.48181 | 9.06E-11 | 7.36E-10 |
| CNAG_07769 | 2984.121 | 0.372769 | 0.065489 | 5.692106 | 1.25E-08 | 8.14E-08 |
| CNAG_04432 | 700.5972 | -0.36376 | 0.092363 | -3.9384 | 8.20E-05 | 0.000306 |
| CNAG_04434 | 916.0327 | -0.59328 | 0.115875 | -5.12001 | 3.06E-07 | 1.68E-06 |
| CNAG_07770 | 1899.458 | 0.279581 | 0.078995 | 3.539226 | 0.000401 | 0.001292 |
| CNAG_04436 | 925.7561 | 0.210584 | 0.086615 | 2.431268 | 0.015046 | 0.033062 |
| CNAG_04437 | 635.2994 | -0.83121 | 0.099957 | -8.31566 | 9.12E-17 | 1.10E-15 |
| CNAG_04438 | 823.8402 | -0.20432 | 0.103478 | -1.97457 | 0.048316 | 0.091012 |
| CNAG_04439 | 3606.757 | -0.20434 | 0.062799 | -3.25383 | 0.001139 | 0.003306 |
| CNAG_04440 | 603.2056 | -0.50654 | 0.122157 | -4.14666 | 3.37E-05 | 0.000135 |

| | | | | | | |
|---|---|---|---|---|---|---|
| CNAG_04443 | 1023.959 | -0.71821 | 0.086635 | -8.29004 | 1.13E-16 | 1.36E-15 |
| CNAG_04444 | 4837.277 | -0.35001 | 0.065351 | -5.3558 | 8.52E-08 | 5.05E-07 |
| CNAG_04445 | 44810.06 | -0.15916 | 0.076132 | -2.09061 | 0.036563 | 0.071593 |
| CNAG_04447 | 849.4895 | 0.302276 | 0.090868 | 3.326536 | 0.000879 | 0.002624 |
| CNAG_04449 | 1296.243 | 0.319054 | 0.094017 | 3.393564 | 0.00069 | 0.002113 |
| CNAG_04450 | 3470.287 | 0.164616 | 0.063822 | 2.579297 | 0.0099 | 0.022893 |
| CNAG_04455 | 364.1478 | 0.384048 | 0.123897 | 3.099734 | 0.001937 | 0.005359 |
| CNAG_04457 | 28.09225 | -0.87207 | 0.40394 | -2.1589 | 0.030858 | 0.061852 |
| CNAG_04458 | 772.2769 | -0.40102 | 0.090961 | -4.40869 | 1.04E-05 | 4.56E-05 |
| CNAG_04459 | 69.4329 | 2.163756 | 0.421816 | 5.129619 | 2.90E-07 | 1.60E-06 |
| CNAG_12856 | 166.7236 | 0.774609 | 0.23622 | 3.279177 | 0.001041 | 0.003049 |
| CNAG_04460 | 2608.456 | -0.46032 | 0.068654 | -6.70496 | 2.01E-11 | 1.72E-10 |
| CNAG_12860 | 16.55816 | 1.39202 | 0.537201 | 2.591246 | 0.009563 | 0.022208 |
| CNAG_04465 | 3035.888 | -0.27465 | 0.070046 | -3.92096 | 8.82E-05 | 0.000327 |
| CNAG_04466 | 492.6355 | 0.253092 | 0.112131 | 2.257098 | 0.024002 | 0.049717 |
| CNAG_04467 | 768.1813 | 0.34874 | 0.101605 | 3.432318 | 0.000598 | 0.001861 |
| CNAG_12863 | 145.9734 | 4.916215 | 0.310424 | 15.83709 | 1.73E-56 | 6.71E-55 |
| CNAG_04469 | 646.1785 | 5.005497 | 0.161303 | 31.03171 | 2.01E-211 | 4.55E-209 |
| CNAG_04470 | 1356.706 | 3.410557 | 0.100188 | 34.04156 | 5.41E-254 | 1.83E-251 |
| CNAG_04471 | 749.9792 | 1.295941 | 0.114192 | 11.34884 | 7.52E-30 | 1.54E-28 |
| CNAG_04472 | 226.0888 | 2.622711 | 0.198949 | 13.18285 | 1.10E-39 | 2.99E-38 |
| CNAG_12864 | 21.59614 | 2.818814 | 0.583922 | 4.827382 | 1.38E-06 | 6.94E-06 |
| CNAG_04474 | 251.3096 | 0.954612 | 0.162332 | 5.880632 | 4.09E-09 | 2.81E-08 |
| CNAG_04475 | 2389.194 | 0.559705 | 0.07846 | 7.133602 | 9.78E-13 | 9.33E-12 |
| CNAG_04476 | 169.9747 | 0.972572 | 0.249037 | 3.905331 | 9.41E-05 | 0.000347 |
| CNAG_07771 | 1827.681 | -0.33352 | 0.072652 | -4.59058 | 4.42E-06 | 2.07E-05 |
| CNAG_04478 | 2931.099 | 0.265981 | 0.084748 | 3.138491 | 0.001698 | 0.00475 |
| CNAG_04479 | 556.4628 | -0.55447 | 0.129049 | -4.29659 | 1.73E-05 | 7.28E-05 |
| CNAG_07774 | 487.5299 | 0.526371 | 0.110127 | 4.779688 | 1.76E-06 | 8.72E-06 |
| CNAG_04485 | 5924.183 | -0.21 | 0.079791 | -2.6319 | 0.008491 | 0.019987 |
| CNAG_04488 | 2465.59 | -0.27511 | 0.076351 | -3.60322 | 0.000314 | 0.001038 |
| CNAG_07775 | 564.2893 | -0.60429 | 0.183173 | -3.299 | 0.00097 | 0.002866 |
| CNAG_07777 | 271.6706 | -0.35886 | 0.138336 | -2.59412 | 0.009483 | 0.022056 |
| CNAG_04494 | 1650.21 | -0.34159 | 0.095717 | -3.56876 | 0.000359 | 0.00117 |
| CNAG_04495 | 184.9344 | -0.49302 | 0.167155 | -2.94946 | 0.003183 | 0.008352 |
| CNAG_04496 | 1046.504 | -0.42923 | 0.099194 | -4.32716 | 1.51E-05 | 6.41E-05 |
| CNAG_04497 | 585.4135 | -0.29 | 0.106664 | -2.71878 | 0.006552 | 0.015855 |
| CNAG_04501 | 3623.471 | -0.2779 | 0.068553 | -4.05384 | 5.04E-05 | 0.000195 |
| CNAG_04503 | 698.4389 | -0.33767 | 0.096573 | -3.49651 | 0.000471 | 0.001495 |
| CNAG_04504 | 2738.914 | 1.113768 | 0.07627 | 14.60302 | 2.69E-48 | 8.74E-47 |
| CNAG_12866 | 76.81096 | 0.908053 | 0.254476 | 3.56832 | 0.000359 | 0.001171 |
| CNAG_04505 | 2358.23 | -0.18973 | 0.082938 | -2.28762 | 0.022159 | 0.046374 |
| CNAG_12867 | 21.30836 | 1.267126 | 0.489213 | 2.590134 | 0.009594 | 0.022261 |
| CNAG_04510 | 2299.193 | -0.21947 | 0.073764 | -2.97535 | 0.002927 | 0.007748 |
| CNAG_04511 | 307.4679 | -0.47423 | 0.141562 | -3.34997 | 0.000808 | 0.002433 |
| CNAG_04512 | 461.5559 | -0.28524 | 0.113324 | -2.51704 | 0.011835 | 0.026854 |
| CNAG_04514 | 1847.871 | 0.209991 | 0.071114 | 2.952874 | 0.003148 | 0.008271 |

| | | | | | | |
|---|---|---|---|---|---|---|
| CNAG_04515 | 2186.919 | -1.5057 | 0.098987 | -15.211 | 2.99E-52 | 1.07E-50 |
| CNAG_12869 | 139.1626 | -1.64183 | 0.230812 | -7.11327 | 1.13E-12 | 1.08E-11 |
| CNAG_04518 | 578.4792 | -0.27859 | 0.114573 | -2.43155 | 0.015034 | 0.033045 |
| CNAG_04519 | 168.9949 | 0.553988 | 0.17945 | 3.087149 | 0.002021 | 0.005559 |
| CNAG_12871 | 381.9505 | 1.962366 | 0.144113 | 13.61682 | 3.18E-42 | 9.07E-41 |
| CNAG_04521 | 2224.449 | 5.652485 | 0.117147 | 48.25123 | 0 | 0 |
| CNAG_04522 | 1243.057 | 0.81986 | 0.094615 | 8.665251 | 4.51E-18 | 5.79E-17 |
| CNAG_04523 | 30.45696 | 1.208954 | 0.391565 | 3.087492 | 0.002019 | 0.005554 |
| CNAG_04524 | 1694.854 | 1.162084 | 0.090082 | 12.9003 | 4.48E-38 | 1.15E-36 |
| CNAG_04527 | 665.4656 | -0.26372 | 0.100965 | -2.61195 | 0.009003 | 0.021117 |
| CNAG_04528 | 291.0624 | -0.39786 | 0.154461 | -2.57582 | 0.01 | 0.023091 |
| CNAG_04529 | 312.7125 | -0.50169 | 0.146759 | -3.4185 | 0.00063 | 0.001948 |
| CNAG_04530 | 2065.873 | -0.27333 | 0.080187 | -3.40862 | 0.000653 | 0.002008 |
| CNAG_04531 | 1226.633 | -0.6371 | 0.091302 | -6.97801 | 2.99E-12 | 2.74E-11 |
| CNAG_07778 | 8817.827 | -0.29339 | 0.068381 | -4.29052 | 1.78E-05 | 7.47E-05 |
| CNAG_07779 | 71.30721 | 3.894425 | 0.356776 | 10.9156 | 9.71E-28 | 1.86E-26 |
| CNAG_04535 | 102.7336 | 0.861518 | 0.21187 | 4.066249 | 4.78E-05 | 0.000186 |
| CNAG_04536 | 75.89429 | 1.255344 | 0.270819 | 4.635363 | 3.56E-06 | 1.69E-05 |
| CNAG_04537 | 490.1641 | -0.35278 | 0.145854 | -2.41869 | 0.015576 | 0.034088 |
| CNAG_04539 | 1137.841 | 0.590134 | 0.103625 | 5.694913 | 1.23E-08 | 8.02E-08 |
| CNAG_04541 | 1504.555 | -0.17638 | 0.084986 | -2.0754 | 0.03795 | 0.074005 |
| CNAG_04544 | 470.4639 | 0.255747 | 0.121019 | 2.113281 | 0.034577 | 0.068114 |
| CNAG_04545 | 702.5206 | -0.25307 | 0.092795 | -2.72715 | 0.006388 | 0.015504 |
| CNAG_04546 | 57.21206 | 1.050033 | 0.290227 | 3.61797 | 0.000297 | 0.000984 |
| CNAG_12873 | 118.0525 | 0.724858 | 0.215302 | 3.3667 | 0.000761 | 0.002307 |
| CNAG_07780 | 647.6429 | 0.340508 | 0.099328 | 3.428109 | 0.000608 | 0.001887 |
| CNAG_07781 | 1218.093 | 0.832405 | 0.09042 | 9.20596 | 3.39E-20 | 4.84E-19 |
| CNAG_07020 | 1709.533 | 0.195951 | 0.077825 | 2.517833 | 0.011808 | 0.026801 |
| CNAG_07782 | 987.6819 | 1.122647 | 0.092275 | 12.16637 | 4.69E-34 | 1.07E-32 |
| CNAG_06997 | 565.0639 | 0.461283 | 0.125061 | 3.68847 | 0.000226 | 0.000767 |
| CNAG_06999 | 341.4196 | -0.76415 | 0.16994 | -4.49659 | 6.91E-06 | 3.14E-05 |
| CNAG_12876 | 573.6393 | -0.24973 | 0.107323 | -2.32688 | 0.019972 | 0.042351 |
| CNAG_07784 | 488.4333 | 4.482754 | 0.16974 | 26.40947 | 1.07E-153 | 1.45E-151 |
| CNAG_06953 | 79.0404 | -1.04772 | 0.246726 | -4.24649 | 2.17E-05 | 8.93E-05 |
| CNAG_04927 | 117.1542 | 0.844996 | 0.205823 | 4.105458 | 4.04E-05 | 0.000159 |
| CNAG_04926 | 179.5091 | 0.725712 | 0.164114 | 4.421996 | 9.78E-06 | 4.32E-05 |
| CNAG_04925 | 1540.547 | -0.37198 | 0.072974 | -5.09748 | 3.44E-07 | 1.88E-06 |
| CNAG_04921 | 36.32467 | 0.840088 | 0.351375 | 2.390858 | 0.016809 | 0.036435 |
| CNAG_04920 | 1979.958 | 1.694896 | 0.123208 | 13.75633 | 4.67E-43 | 1.40E-41 |
| CNAG_04919 | 475.3407 | 0.926412 | 0.118542 | 7.815036 | 5.49E-15 | 6.03E-14 |
| CNAG_07838 | 827.7808 | -0.27632 | 0.088096 | -3.13656 | 0.001709 | 0.004773 |
| CNAG_07839 | 50250.59 | -0.25772 | 0.072266 | -3.56621 | 0.000362 | 0.00118 |
| CNAG_04914 | 4730.539 | -0.60735 | 0.068391 | -8.88052 | 6.65E-19 | 8.90E-18 |
| CNAG_04913 | 1635.881 | -0.3026 | 0.082843 | -3.65265 | 0.00026 | 0.00087 |
| CNAG_04911 | 576.0887 | 0.807788 | 0.108276 | 7.460474 | 8.62E-14 | 8.87E-13 |
| CNAG_04909 | 1642.072 | 2.263187 | 0.077803 | 29.08867 | 4.99E-186 | 9.02E-184 |
| CNAG_12880 | 36.28588 | 0.983441 | 0.381567 | 2.577374 | 0.009955 | 0.023014 |

| | | | | | | |
|---|---|---|---|---|---|---|
| CNAG_04908 | 926.5003 | 0.530334 | 0.124481 | 4.260377 | 2.04E-05 | 8.45E-05 |
| CNAG_04906 | 2993.129 | 0.184861 | 0.064123 | 2.882917 | 0.00394 | 0.010095 |
| CNAG_04905 | 87.22985 | 0.908046 | 0.234109 | 3.878735 | 0.000105 | 0.000383 |
| CNAG_07842 | 158.839 | 0.875406 | 0.174025 | 5.030332 | 4.90E-07 | 2.63E-06 |
| CNAG_04903 | 1158.993 | 1.198809 | 0.341246 | 3.513037 | 0.000443 | 0.001415 |
| CNAG_04901 | 560.8244 | 1.491422 | 0.108469 | 13.74973 | 5.11E-43 | 1.53E-41 |
| CNAG_04899 | 1575.295 | -0.23343 | 0.081462 | -2.86553 | 0.004163 | 0.010624 |
| CNAG_04898 | 185.2022 | 0.482628 | 0.169548 | 2.846565 | 0.004419 | 0.011197 |
| CNAG_04894 | 347.0558 | 0.749208 | 0.19514 | 3.839328 | 0.000123 | 0.000444 |
| CNAG_04892 | 193.0196 | -0.67935 | 0.170803 | -3.97737 | 6.97E-05 | 0.000263 |
| CNAG_04891 | 94.38786 | 1.747955 | 0.243005 | 7.193089 | 6.33E-13 | 6.14E-12 |
| CNAG_04890 | 110.9697 | 3.555587 | 0.287057 | 12.38634 | 3.10E-35 | 7.34E-34 |
| CNAG_04889 | 1020.384 | 1.367997 | 0.109427 | 12.50149 | 7.33E-36 | 1.77E-34 |
| CNAG_04887 | 239.4791 | 1.828331 | 0.163019 | 11.21548 | 3.42E-29 | 6.87E-28 |
| CNAG_04886 | 477.991 | 0.254259 | 0.117539 | 2.163181 | 0.030527 | 0.061281 |
| CNAG_12883 | 56.57471 | 1.402576 | 0.36102 | 3.885031 | 0.000102 | 0.000374 |
| CNAG_07844 | 174.9827 | 4.314451 | 0.250669 | 17.21176 | 2.17E-66 | 1.03E-64 |
| CNAG_04883 | 37856.6 | -0.26537 | 0.078163 | -3.39515 | 0.000686 | 0.002104 |
| CNAG_04882 | 1238.459 | -0.37905 | 0.079014 | -4.79722 | 1.61E-06 | 8.03E-06 |
| CNAG_04880 | 472.3937 | 0.550636 | 0.123329 | 4.464761 | 8.02E-06 | 3.60E-05 |
| CNAG_04879 | 3675.307 | -0.85572 | 0.078893 | -10.8466 | 2.07E-27 | 3.90E-26 |
| CNAG_12885 | 70.86427 | 0.767714 | 0.2606 | 2.945944 | 0.00322 | 0.008432 |
| CNAG_04877 | 34.47079 | 1.243258 | 0.455897 | 2.727056 | 0.00639 | 0.015504 |
| CNAG_04876 | 1933.753 | 0.21863 | 0.070482 | 3.101932 | 0.001923 | 0.005324 |
| CNAG_04874 | 549.8908 | 0.904636 | 0.126511 | 7.150676 | 8.64E-13 | 8.29E-12 |
| CNAG_04873 | 289.5207 | 0.868168 | 0.134454 | 6.456983 | 1.07E-10 | 8.60E-10 |
| CNAG_04872 | 2675.406 | 0.51763 | 0.06847 | 7.559923 | 4.03E-14 | 4.24E-13 |
| CNAG_04871 | 438.5878 | 0.280833 | 0.112089 | 2.505443 | 0.01223 | 0.02762 |
| CNAG_12886 | 171.8027 | 0.922662 | 0.180878 | 5.101032 | 3.38E-07 | 1.85E-06 |
| CNAG_04870 | 56.17476 | 0.587118 | 0.292907 | 2.004455 | 0.045021 | 0.085799 |
| CNAG_04869 | 3483.26 | 1.294807 | 0.079199 | 16.3487 | 4.44E-60 | 1.88E-58 |
| CNAG_04868 | 1478.635 | 0.493533 | 0.104201 | 4.736356 | 2.18E-06 | 1.07E-05 |
| CNAG_04866 | 901.4675 | 0.386639 | 0.103663 | 3.729781 | 0.000192 | 0.000663 |
| CNAG_04865 | 101.3745 | 1.149945 | 0.323755 | 3.551893 | 0.000382 | 0.001239 |
| CNAG_12888 | 46.92136 | 2.254059 | 0.340983 | 6.61047 | 3.83E-11 | 3.21E-10 |
| CNAG_04864 | 2188.266 | -1.14947 | 0.087939 | -13.0712 | 4.81E-39 | 1.28E-37 |
| CNAG_04863 | 1107.163 | 1.625056 | 0.096701 | 16.80489 | 2.25E-63 | 1.01E-61 |
| CNAG_12889 | 103.3623 | 4.739899 | 0.368383 | 12.86677 | 6.92E-38 | 1.77E-36 |
| CNAG_04862 | 10918.01 | 3.190223 | 0.07032 | 45.36703 | 0 | 0 |
| CNAG_12890 | 48.21804 | 2.104236 | 0.362379 | 5.806731 | 6.37E-09 | 4.30E-08 |
| CNAG_04861 | 2039.636 | 0.722291 | 0.084073 | 8.59127 | 8.60E-18 | 1.09E-16 |
| CNAG_04860 | 2096.062 | 0.229304 | 0.085705 | 2.675499 | 0.007462 | 0.017817 |
| CNAG_04859 | 885.9186 | -0.68803 | 0.105985 | -6.4918 | 8.48E-11 | 6.91E-10 |
| CNAG_04857 | 3919.748 | 0.38279 | 0.09455 | 4.048554 | 5.15E-05 | 0.000199 |
| CNAG_04853 | 579.8694 | -0.51732 | 0.106259 | -4.86844 | 1.12E-06 | 5.72E-06 |
| CNAG_04851 | 13322.32 | 0.222387 | 0.055511 | 4.006153 | 6.17E-05 | 0.000235 |
| CNAG_12892 | 141.7236 | -1.17008 | 0.21919 | -5.3382 | 9.39E-08 | 5.53E-07 |

| | | | | | | |
|---|---|---|---|---|---|---|
| CNAG_04849 | 403.3195 | -0.5012 | 0.1247 | -4.01924 | 5.84E-05 | 0.000223 |
| CNAG_04848 | 2954.941 | -0.45986 | 0.065599 | -7.01011 | 2.38E-12 | 2.21E-11 |
| CNAG_04847 | 980.2396 | 0.18501 | 0.084722 | 2.183715 | 0.028983 | 0.058601 |
| CNAG_04846 | 2519.608 | 1.217549 | 0.105994 | 11.48701 | 1.53E-30 | 3.20E-29 |
| CNAG_04845 | 1128.78 | -0.42084 | 0.08674 | -4.85177 | 1.22E-06 | 6.20E-06 |
| CNAG_04842 | 1351.386 | 0.187446 | 0.079358 | 2.362022 | 0.018176 | 0.038989 |
| CNAG_04840 | 5801.675 | -0.42221 | 0.062496 | -6.75579 | 1.42E-11 | 1.23E-10 |
| CNAG_04838 | 2777.035 | -0.17883 | 0.069595 | -2.56956 | 0.010183 | 0.023466 |
| CNAG_04837 | 133.324 | -0.66905 | 0.209491 | -3.1937 | 0.001405 | 0.003993 |
| CNAG_04836 | 676.624 | 0.341658 | 0.099557 | 3.431787 | 0.0006 | 0.001864 |
| CNAG_04833 | 1010.677 | -0.31285 | 0.134534 | -2.32546 | 0.020048 | 0.042467 |
| CNAG_04829 | 416.6248 | -0.39454 | 0.138743 | -2.84365 | 0.00446 | 0.011282 |
| CNAG_12895 | 220.458 | 0.514793 | 0.154651 | 3.328736 | 0.000872 | 0.002607 |
| CNAG_04828 | 7826.192 | -0.54314 | 0.064745 | -8.38893 | 4.91E-17 | 6.02E-16 |
| CNAG_04827 | 257.8583 | 0.479145 | 0.165913 | 2.887936 | 0.003878 | 0.009958 |
| CNAG_04824 | 554.745 | -0.56047 | 0.103584 | -5.41084 | 6.27E-08 | 3.78E-07 |
| CNAG_04823 | 1883.063 | -0.55183 | 0.080025 | -6.89572 | 5.36E-12 | 4.82E-11 |
| CNAG_04822 | 1543.099 | 0.197977 | 0.076193 | 2.598379 | 0.009367 | 0.021845 |
| CNAG_04820 | 1790.474 | -0.19831 | 0.083355 | -2.37906 | 0.017357 | 0.037441 |
| CNAG_04819 | 1233.434 | 1.610855 | 0.091272 | 17.64904 | 1.04E-69 | 5.07E-68 |
| CNAG_04818 | 565.761 | 3.616379 | 0.137294 | 26.34036 | 6.62E-153 | 8.82E-151 |
| CNAG_07845 | 118.7506 | -0.45791 | 0.212551 | -2.15437 | 0.031212 | 0.0625 |
| CNAG_04813 | 1250.813 | -0.352 | 0.089899 | -3.91547 | 9.02E-05 | 0.000334 |
| CNAG_04810 | 432.3292 | -0.3073 | 0.119638 | -2.56859 | 0.010211 | 0.023525 |
| CNAG_04808 | 3944.613 | -0.13168 | 0.064303 | -2.04782 | 0.040578 | 0.078457 |
| CNAG_04807 | 1488.21 | -0.97434 | 0.115804 | -8.41373 | 3.97E-17 | 4.90E-16 |
| CNAG_04806 | 5165.772 | -0.20037 | 0.064142 | -3.12393 | 0.001785 | 0.004967 |
| CNAG_04805 | 1424.632 | -0.3564 | 0.078937 | -4.51495 | 6.33E-06 | 2.90E-05 |
| CNAG_04804 | 1709.57 | -0.95862 | 0.077038 | -12.4435 | 1.52E-35 | 3.62E-34 |
| CNAG_04802 | 1826.075 | -0.19244 | 0.075236 | -2.55786 | 0.010532 | 0.024209 |
| CNAG_04800 | 6425.96 | -0.36325 | 0.062808 | -5.7835 | 7.32E-09 | 4.90E-08 |
| CNAG_04799 | 46517.5 | -0.21547 | 0.070237 | -3.06779 | 0.002156 | 0.005882 |
| CNAG_04798 | 2701.914 | -0.39408 | 0.084348 | -4.6721 | 2.98E-06 | 1.43E-05 |
| CNAG_04796 | 4036.942 | 0.615802 | 0.066201 | 9.30201 | 1.38E-20 | 2.01E-19 |
| CNAG_04795 | 671.7788 | 0.493396 | 0.122458 | 4.029091 | 5.60E-05 | 0.000215 |
| CNAG_04792 | 2030.593 | 0.384647 | 0.072876 | 5.278088 | 1.31E-07 | 7.54E-07 |
| CNAG_04791 | 1548.782 | -0.20133 | 0.075768 | -2.65721 | 0.007879 | 0.018682 |
| CNAG_04790 | 1045.995 | -0.25323 | 0.093563 | -2.70655 | 0.006799 | 0.016363 |
| CNAG_04788 | 2388.88 | -0.37976 | 0.084324 | -4.50355 | 6.68E-06 | 3.05E-05 |
| CNAG_04787 | 199.0635 | -0.54652 | 0.175034 | -3.12235 | 0.001794 | 0.004992 |
| CNAG_04785 | 674.9248 | 0.437202 | 0.113155 | 3.863754 | 0.000112 | 0.000405 |
| CNAG_04784 | 103.3832 | 2.764237 | 0.279968 | 9.873415 | 5.43E-23 | 8.81E-22 |
| CNAG_04783 | 1851.051 | -0.18489 | 0.076791 | -2.40772 | 0.016052 | 0.03497 |
| CNAG_04782 | 1775.389 | -0.3079 | 0.072727 | -4.23367 | 2.30E-05 | 9.39E-05 |
| CNAG_07848 | 1026.853 | -0.34477 | 0.097403 | -3.5396 | 0.000401 | 0.00129 |
| CNAG_07851 | 7246.306 | -0.30089 | 0.067224 | -4.47587 | 7.61E-06 | 3.44E-05 |
| CNAG_04777 | 1050.699 | 0.527891 | 0.089857 | 5.874775 | 4.23E-09 | 2.90E-08 |

| | | | | | | |
|---|---|---|---|---|---|---|
| CNAG_04776 | 4722.717 | -0.30964 | 0.070092 | -4.41753 | 9.98E-06 | 4.40E-05 |
| CNAG_04775 | 492.5194 | -0.37442 | 0.111043 | -3.37189 | 0.000747 | 0.002273 |
| CNAG_04774 | 2362.801 | 0.952925 | 0.073409 | 12.98104 | 1.57E-38 | 4.11E-37 |
| CNAG_12905 | 56.86448 | 1.956782 | 0.338185 | 5.786134 | 7.20E-09 | 4.83E-08 |
| CNAG_04773 | 181.3694 | 2.04332 | 0.211027 | 9.682763 | 3.57E-22 | 5.56E-21 |
| CNAG_12906 | 255.9963 | 2.766887 | 0.182737 | 15.14139 | 8.64E-52 | 3.01E-50 |
| CNAG_04772 | 2566.76 | -0.16606 | 0.076274 | -2.17715 | 0.02947 | 0.059451 |
| CNAG_04770 | 2991.439 | 0.189052 | 0.069835 | 2.707131 | 0.006787 | 0.016349 |
| CNAG_04768 | 1316.009 | 0.448293 | 0.085494 | 5.243567 | 1.58E-07 | 9.00E-07 |
| CNAG_04767 | 947.1594 | -0.39965 | 0.095936 | -4.16587 | 3.10E-05 | 0.000124 |
| CNAG_04765 | 1257.22 | -0.49021 | 0.093275 | -5.25557 | 1.48E-07 | 8.47E-07 |
| CNAG_04764 | 576.8756 | -0.30326 | 0.10078 | -3.00916 | 0.00262 | 0.007011 |
| CNAG_04763 | 2786.684 | 0.172175 | 0.080982 | 2.12609 | 0.033496 | 0.066371 |
| CNAG_04761 | 601.7613 | -0.2218 | 0.109908 | -2.018 | 0.043591 | 0.083405 |
| CNAG_04758 | 913.4176 | 0.351986 | 0.108109 | 3.255847 | 0.001131 | 0.003285 |
| CNAG_04756 | 509.7001 | -1.12236 | 0.125321 | -8.95592 | 3.37E-19 | 4.58E-18 |
| CNAG_12912 | 56.70851 | 0.66464 | 0.28736 | 2.312914 | 0.020727 | 0.043631 |
| CNAG_04755 | 2944.124 | 0.325909 | 0.082074 | 3.97092 | 7.16E-05 | 0.00027 |
| CNAG_12914 | 127.9346 | 1.639486 | 0.209219 | 7.836225 | 4.64E-15 | 5.14E-14 |
| CNAG_04753 | 2118.192 | 2.04778 | 0.073487 | 27.86573 | 6.95E-171 | 1.11E-168 |
| CNAG_04752 | 1476.989 | 0.521769 | 0.083246 | 6.267812 | 3.66E-10 | 2.80E-09 |
| CNAG_04751 | 601.2702 | -0.37739 | 0.098201 | -3.84307 | 0.000122 | 0.000438 |
| CNAG_04750 | 135.2371 | -0.86471 | 0.204637 | -4.22561 | 2.38E-05 | 9.71E-05 |
| CNAG_04749 | 1987.857 | 0.456595 | 0.073248 | 6.233569 | 4.56E-10 | 3.45E-09 |
| CNAG_04747 | 729.4129 | 0.807479 | 0.10856 | 7.438091 | 1.02E-13 | 1.05E-12 |
| CNAG_04746 | 1307.347 | -0.28814 | 0.080565 | -3.57656 | 0.000348 | 0.001138 |
| CNAG_04743 | 4385.723 | 0.268898 | 0.061956 | 4.340135 | 1.42E-05 | 6.08E-05 |
| CNAG_04742 | 879.1484 | 0.575823 | 0.089357 | 6.444086 | 1.16E-10 | 9.33E-10 |
| CNAG_04738 | 3694.606 | -0.27595 | 0.074746 | -3.69181 | 0.000223 | 0.000758 |
| CNAG_04737 | 5028.361 | 0.223069 | 0.104999 | 2.12449 | 0.033629 | 0.06657 |
| CNAG_04736 | 596.0919 | 0.868159 | 0.117139 | 7.411354 | 1.25E-13 | 1.27E-12 |
| CNAG_04735 | 3766.564 | 1.403166 | 0.071438 | 19.64172 | 6.80E-86 | 4.13E-84 |
| CNAG_04734 | 1243.365 | 0.300769 | 0.079449 | 3.785704 | 0.000153 | 0.000541 |
| CNAG_04733 | 1604.896 | -0.17147 | 0.074807 | -2.29212 | 0.021899 | 0.045887 |
| CNAG_04732 | 1171.847 | -0.29053 | 0.081218 | -3.57714 | 0.000347 | 0.001136 |
| CNAG_04731 | 499.8423 | -0.21392 | 0.106502 | -2.00858 | 0.044582 | 0.08509 |
| CNAG_04730 | 1646.574 | 0.428003 | 0.09067 | 4.720441 | 2.35E-06 | 1.15E-05 |
| CNAG_04729 | 3308.143 | 0.292199 | 0.071632 | 4.079154 | 4.52E-05 | 0.000176 |
| CNAG_04724 | 1839.187 | 0.323874 | 0.071364 | 4.538348 | 5.67E-06 | 2.62E-05 |
| CNAG_12920 | 39.72841 | 0.984232 | 0.342175 | 2.876403 | 0.004022 | 0.010288 |
| CNAG_04719 | 789.1179 | -0.29206 | 0.095753 | -3.05013 | 0.002287 | 0.00621 |
| CNAG_04718 | 238.1775 | -0.66368 | 0.169229 | -3.92182 | 8.79E-05 | 0.000326 |
| CNAG_04716 | 4239.217 | -0.17326 | 0.062296 | -2.78121 | 0.005416 | 0.013437 |
| CNAG_04715 | 2617.862 | 0.271512 | 0.090103 | 3.013341 | 0.002584 | 0.006924 |
| CNAG_04714 | 891.9219 | 0.998688 | 0.092167 | 10.83565 | 2.33E-27 | 4.38E-26 |
| CNAG_04713 | 1445.907 | 0.232957 | 0.079299 | 2.93771 | 0.003306 | 0.008633 |
| CNAG_04712 | 1525.596 | 0.608104 | 0.076223 | 7.977923 | 1.49E-15 | 1.70E-14 |

| | | | | | | |
|---|---|---|---|---|---|---|
| CNAG_04711 | 74.07886 | -0.9227 | 0.316504 | -2.9153 | 0.003553 | 0.009204 |
| CNAG_04710 | 2107.418 | -0.2382 | 0.071675 | -3.32332 | 0.00089 | 0.002648 |
| CNAG_04709 | 3241.691 | -0.39364 | 0.084217 | -4.67409 | 2.95E-06 | 1.42E-05 |
| CNAG_04708 | 484.9134 | -0.34262 | 0.117944 | -2.90491 | 0.003674 | 0.009491 |
| CNAG_04707 | 284.0546 | 1.102592 | 0.145627 | 7.571343 | 3.69E-14 | 3.89E-13 |
| CNAG_07855 | 4744.044 | 0.369005 | 0.061307 | 6.018983 | 1.76E-09 | 1.25E-08 |
| CNAG_04705 | 2260.096 | 0.888332 | 0.067138 | 13.23141 | 5.78E-40 | 1.58E-38 |
| CNAG_04704 | 78.19078 | 2.416737 | 0.269611 | 8.963778 | 3.14E-19 | 4.28E-18 |
| CNAG_12926 | 82.24474 | 1.07396 | 0.246072 | 4.36442 | 1.27E-05 | 5.48E-05 |
| CNAG_04703 | 2331.947 | 0.295742 | 0.083505 | 3.541621 | 0.000398 | 0.001281 |
| CNAG_04698 | 4125.193 | -0.18765 | 0.059269 | -3.16605 | 0.001545 | 0.004355 |
| CNAG_04697 | 1245.027 | -0.47365 | 0.09464 | -5.00477 | 5.59E-07 | 2.98E-06 |
| CNAG_04696 | 727.1423 | 0.897399 | 0.112708 | 7.962132 | 1.69E-15 | 1.92E-14 |
| CNAG_04695 | 95.68835 | 0.638658 | 0.275997 | 2.314003 | 0.020668 | 0.043542 |
| CNAG_04694 | 6165.313 | 0.294821 | 0.061887 | 4.763861 | 1.90E-06 | 9.38E-06 |
| CNAG_04691 | 124.2103 | 1.756828 | 0.217666 | 8.071217 | 6.96E-16 | 8.06E-15 |
| CNAG_04690 | 1235.546 | 1.470936 | 0.086866 | 16.93343 | 2.55E-64 | 1.18E-62 |
| CNAG_04689 | 708.8216 | 1.183143 | 0.10926 | 10.82871 | 2.52E-27 | 4.71E-26 |
| CNAG_04688 | 1068.144 | 2.166457 | 0.086247 | 25.11925 | 3.06E-139 | 3.51E-137 |
| CNAG_07856 | 1860.005 | 1.006916 | 0.097749 | 10.30106 | 6.97E-25 | 1.21E-23 |
| CNAG_04684 | 3683.634 | -0.18567 | 0.070322 | -2.64032 | 0.008283 | 0.019548 |
| CNAG_04683 | 1450.513 | -0.19345 | 0.087534 | -2.21002 | 0.027104 | 0.055352 |
| CNAG_04681 | 188.8275 | 0.671706 | 0.169778 | 3.956368 | 7.61E-05 | 0.000286 |
| CNAG_04677 | 2061.993 | 1.723547 | 0.258405 | 6.669957 | 2.56E-11 | 2.17E-10 |
| CNAG_04673 | 127.859 | -0.46863 | 0.190543 | -2.45946 | 0.013915 | 0.03091 |
| CNAG_07999 | 502.5953 | 0.312853 | 0.114473 | 2.732988 | 0.006276 | 0.015264 |
| CNAG_04671 | 1275.437 | -0.29771 | 0.085126 | -3.49723 | 0.00047 | 0.001491 |
| CNAG_04669 | 1308.156 | 0.313547 | 0.076278 | 4.110598 | 3.95E-05 | 0.000156 |
| CNAG_04668 | 620.2378 | -0.3511 | 0.107227 | -3.27436 | 0.001059 | 0.003095 |
| CNAG_04667 | 575.358 | -0.38618 | 0.103247 | -3.74031 | 0.000184 | 0.000638 |
| CNAG_04666 | 5251.775 | 0.165907 | 0.056793 | 2.921287 | 0.003486 | 0.009064 |
| CNAG_04665 | 452.1273 | -0.49102 | 0.111453 | -4.40564 | 1.05E-05 | 4.62E-05 |
| CNAG_04662 | 2019.223 | 0.158745 | 0.071836 | 2.20983 | 0.027117 | 0.055365 |
| CNAG_12933 | 60.77652 | -0.94272 | 0.291339 | -3.2358 | 0.001213 | 0.003496 |
| CNAG_04658 | 765.4076 | 1.058954 | 0.109217 | 9.695887 | 3.14E-22 | 4.93E-21 |
| CNAG_04657 | 480.4941 | 0.375817 | 0.125919 | 2.984604 | 0.002839 | 0.007547 |
| CNAG_04655 | 1663.514 | 0.745254 | 0.081213 | 9.176582 | 4.45E-20 | 6.33E-19 |
| CNAG_04654 | 1077.794 | -0.29842 | 0.090904 | -3.28284 | 0.001028 | 0.003014 |
| CNAG_04652 | 1233.637 | 0.213878 | 0.085556 | 2.499848 | 0.012425 | 0.02799 |
| CNAG_04651 | 2181.828 | -0.21493 | 0.076971 | -2.79239 | 0.005232 | 0.013021 |
| CNAG_04650 | 1466.895 | -0.19226 | 0.079244 | -2.42615 | 0.01526 | 0.033483 |
| CNAG_04647 | 1238.291 | -0.20488 | 0.086679 | -2.36369 | 0.018094 | 0.038876 |
| CNAG_04646 | 675.4883 | -0.2655 | 0.099692 | -2.66317 | 0.007741 | 0.018413 |
| CNAG_07857 | 4651.79 | -0.206 | 0.077479 | -2.65875 | 0.007843 | 0.018608 |
| CNAG_04642 | 2109.348 | 0.375775 | 0.070751 | 5.311243 | 1.09E-07 | 6.35E-07 |
| CNAG_04641 | 1476.931 | -0.3018 | 0.086501 | -3.48901 | 0.000485 | 0.001533 |
| CNAG_04638 | 367.2114 | 0.557991 | 0.163309 | 3.416789 | 0.000634 | 0.001956 |

| | | | | | | |
|---|---|---|---|---|---|---|
| CNAG_04637 | 6622.257 | -0.14479 | 0.064019 | -2.26163 | 0.02372 | 0.049196 |
| CNAG_12934 | 31.05135 | 1.130077 | 0.411838 | 2.743984 | 0.00607 | 0.01482 |
| CNAG_04635 | 2683.759 | 0.749505 | 0.086596 | 8.655163 | 4.92E-18 | 6.30E-17 |
| CNAG_04634 | 1049.953 | 0.782912 | 0.106735 | 7.335135 | 2.21E-13 | 2.22E-12 |
| CNAG_04633 | 1257.48 | 0.463213 | 0.080044 | 5.786972 | 7.17E-09 | 4.81E-08 |
| CNAG_12938 | 47.0696 | -1.1608 | 0.326532 | -3.55494 | 0.000378 | 0.001226 |
| CNAG_04630 | 1621.816 | 0.241455 | 0.100704 | 2.39766 | 0.0165 | 0.035869 |
| CNAG_04628 | 3350.216 | -0.16908 | 0.062689 | -2.69714 | 0.006994 | 0.016783 |
| CNAG_04626 | 269.7551 | 0.519277 | 0.191656 | 2.709413 | 0.00674 | 0.016266 |
| CNAG_04623 | 145.6624 | 0.898845 | 0.267224 | 3.363634 | 0.000769 | 0.002331 |
| CNAG_04621 | 5495.95 | -0.50707 | 0.072762 | -6.96889 | 3.19E-12 | 2.91E-11 |
| CNAG_04617 | 905.7667 | 1.927932 | 0.102277 | 18.85005 | 2.94E-79 | 1.61E-77 |
| CNAG_04615 | 591.7843 | 0.701973 | 0.102472 | 6.850383 | 7.37E-12 | 6.57E-11 |
| CNAG_04614 | 250.8909 | -0.52417 | 0.142896 | -3.66819 | 0.000244 | 0.000825 |
| CNAG_04613 | 3292.759 | -0.14155 | 0.068772 | -2.05822 | 0.039569 | 0.076722 |
| CNAG_04612 | 4015.624 | 0.418975 | 0.081133 | 5.164025 | 2.42E-07 | 1.34E-06 |
| CNAG_04609 | 9337.656 | 0.124772 | 0.061932 | 2.014682 | 0.043938 | 0.083971 |
| CNAG_04606 | 259.4606 | 1.066474 | 0.171593 | 6.215123 | 5.13E-10 | 3.86E-09 |
| CNAG_04605 | 5191.082 | -0.63748 | 0.071011 | -8.9772 | 2.78E-19 | 3.80E-18 |
| CNAG_04603 | 561.9257 | 0.455842 | 0.124578 | 3.659086 | 0.000253 | 0.000852 |
| CNAG_04601 | 21636.08 | -0.21991 | 0.060055 | -3.66184 | 0.00025 | 0.000845 |
| CNAG_04600 | 1050.518 | 0.479907 | 0.084801 | 5.659225 | 1.52E-08 | 9.77E-08 |
| CNAG_04599 | 1812.189 | 1.001598 | 0.077546 | 12.91615 | 3.65E-38 | 9.42E-37 |
| CNAG_04597 | 550.6653 | 0.621573 | 0.113068 | 5.497339 | 3.86E-08 | 2.37E-07 |
| CNAG_12943 | 14.98676 | 1.895333 | 0.58447 | 3.24282 | 0.001184 | 0.003421 |
| CNAG_04596 | 758.6796 | 0.279953 | 0.098258 | 2.849172 | 0.004383 | 0.011123 |
| CNAG_04595 | 393.0421 | 0.28232 | 0.118702 | 2.378394 | 0.017388 | 0.037498 |
| CNAG_04594 | 544.9614 | 1.050001 | 0.109098 | 9.624419 | 6.31E-22 | 9.76E-21 |
| CNAG_04590 | 196.4843 | -0.69359 | 0.162367 | -4.27177 | 1.94E-05 | 8.05E-05 |
| CNAG_04589 | 458.4424 | 0.339373 | 0.17183 | 1.975054 | 0.048262 | 0.090959 |
| CNAG_04584 | 5023.115 | -0.37536 | 0.08561 | -4.38459 | 1.16E-05 | 5.04E-05 |
| CNAG_04583 | 3915.492 | -0.15094 | 0.071182 | -2.12044 | 0.033969 | 0.06708 |
| CNAG_04581 | 1043.722 | 0.266535 | 0.087968 | 3.029909 | 0.002446 | 0.006599 |
| CNAG_04580 | 6822.232 | 0.539548 | 0.060968 | 8.84964 | 8.78E-19 | 1.16E-17 |
| CNAG_07861 | 1038.618 | -0.26548 | 0.114443 | -2.31979 | 0.020352 | 0.042989 |
| CNAG_04576 | 470.3276 | 0.658918 | 0.137879 | 4.778966 | 1.76E-06 | 8.74E-06 |
| CNAG_12946 | 101.079 | 0.847513 | 0.278962 | 3.038101 | 0.002381 | 0.00644 |
| CNAG_04574 | 421.3525 | 0.303036 | 0.130579 | 2.320711 | 0.020302 | 0.042929 |
| CNAG_07862 | 3529.849 | 3.24875 | 0.101401 | 32.0385 | 3.18E-225 | 8.06E-223 |
| CNAG_07863 | 9204.621 | 0.164756 | 0.061792 | 2.666294 | 0.007669 | 0.018254 |
| CNAG_07865 | 2910.96 | 0.638582 | 0.06793 | 9.400564 | 5.43E-21 | 8.11E-20 |
| CNAG_04566 | 6738.779 | 0.498377 | 0.058991 | 8.448338 | 2.95E-17 | 3.66E-16 |
| CNAG_04563 | 681.139 | -0.37478 | 0.102049 | -3.6725 | 0.00024 | 0.000813 |
| CNAG_04561 | 415.2263 | -0.37961 | 0.135514 | -2.80129 | 0.00509 | 0.012706 |
| CNAG_04558 | 168.6141 | -0.48007 | 0.198081 | -2.4236 | 0.015367 | 0.033677 |
| CNAG_04557 | 214.2613 | -0.65441 | 0.164819 | -3.9705 | 7.17E-05 | 0.000271 |
| CNAG_04553 | 22.25841 | 2.198796 | 0.496576 | 4.427918 | 9.51E-06 | 4.20E-05 |

| | | | | | | |
|---|---|---|---|---|---|---|
| CNAG_07868 | 313.003 | 0.313 | 0.142876 | 2.190714 | 0.028473 | 0.057741 |
| CNAG_01461 | 1376.557 | 0.305404 | 0.08516 | 3.586228 | 0.000335 | 0.001101 |
| CNAG_08002 | 3324.865 | 0.257863 | 0.07764 | 3.321243 | 0.000896 | 0.002663 |
| CNAG_01463 | 689.0847 | 1.226532 | 0.110004 | 11.14989 | 7.17E-29 | 1.41E-27 |
| CNAG_12948 | 60.59855 | 1.066155 | 0.277743 | 3.838636 | 0.000124 | 0.000445 |
| CNAG_12949 | 33.59957 | 1.552233 | 0.379464 | 4.090592 | 4.30E-05 | 0.000168 |
| CNAG_01464 | 24947.22 | 2.40627 | 0.111411 | 21.59807 | 1.87E-103 | 1.44E-101 |
| CNAG_12950 | 46.04185 | 1.588659 | 0.340773 | 4.661932 | 3.13E-06 | 1.50E-05 |
| CNAG_01466 | 1205.658 | 1.258303 | 0.099472 | 12.64984 | 1.12E-36 | 2.80E-35 |
| CNAG_12951 | 105.0709 | 0.858918 | 0.24154 | 3.556014 | 0.000377 | 0.001222 |
| CNAG_01470 | 8810.195 | 1.076782 | 0.067061 | 16.05675 | 5.13E-58 | 2.06E-56 |
| CNAG_01471 | 2889.329 | 1.259458 | 0.066853 | 18.83915 | 3.61E-79 | 1.97E-77 |
| CNAG_01472 | 1168.69 | 0.586746 | 0.081189 | 7.226899 | 4.94E-13 | 4.87E-12 |
| CNAG_01473 | 125.1268 | -1.39423 | 0.207695 | -6.71287 | 1.91E-11 | 1.63E-10 |
| CNAG_12954 | 134.3387 | -0.40986 | 0.207863 | -1.97175 | 0.048638 | 0.091469 |
| CNAG_01476 | 2297.719 | -0.47753 | 0.094121 | -5.07354 | 3.90E-07 | 2.12E-06 |
| CNAG_01477 | 2027.065 | -0.3524 | 0.080666 | -4.36864 | 1.25E-05 | 5.38E-05 |
| CNAG_01478 | 526.8113 | -0.57606 | 0.109365 | -5.26735 | 1.38E-07 | 7.95E-07 |
| CNAG_01479 | 478.8856 | -0.48638 | 0.110156 | -4.41532 | 1.01E-05 | 4.44E-05 |
| CNAG_01480 | 41098.03 | -0.27179 | 0.068454 | -3.97032 | 7.18E-05 | 0.000271 |
| CNAG_01483 | 1019.242 | -0.37575 | 0.087711 | -4.28396 | 1.84E-05 | 7.67E-05 |
| CNAG_01486 | 58753.56 | -0.22501 | 0.073343 | -3.06788 | 0.002156 | 0.005882 |
| CNAG_01489 | 163.2601 | -1.14674 | 0.196217 | -5.84425 | 5.09E-09 | 3.47E-08 |
| CNAG_01490 | 974.2733 | 1.606308 | 0.08912 | 18.02419 | 1.26E-72 | 6.35E-71 |
| CNAG_01491 | 1418.504 | -0.29094 | 0.0889 | -3.27272 | 0.001065 | 0.003111 |
| CNAG_01492 | 8256.777 | -0.47459 | 0.059285 | -8.00518 | 1.19E-15 | 1.37E-14 |
| CNAG_01493 | 262.0623 | -0.5317 | 0.170893 | -3.11132 | 0.001863 | 0.005174 |
| CNAG_12960 | 40.52686 | -1.62232 | 0.34909 | -4.64729 | 3.36E-06 | 1.60E-05 |
| CNAG_01494 | 89.34482 | -1.52058 | 0.254935 | -5.96457 | 2.45E-09 | 1.73E-08 |
| CNAG_01495 | 949.4206 | 0.249432 | 0.110688 | 2.253456 | 0.02423 | 0.050139 |
| CNAG_01497 | 242.2539 | 0.607731 | 0.141238 | 4.302873 | 1.69E-05 | 7.10E-05 |
| CNAG_01498 | 805.089 | 0.271448 | 0.090935 | 2.985096 | 0.002835 | 0.007537 |
| CNAG_01499 | 518.1244 | 1.041728 | 0.103803 | 10.03558 | 1.06E-23 | 1.77E-22 |
| CNAG_12961 | 63.64945 | 1.117699 | 0.350592 | 3.18803 | 0.001432 | 0.004062 |
| CNAG_12962 | 82.6556 | 2.60468 | 0.282783 | 9.210893 | 3.23E-20 | 4.63E-19 |
| CNAG_01504 | 1330.407 | 0.267762 | 0.089134 | 3.004033 | 0.002664 | 0.007121 |
| CNAG_01506 | 153.4773 | 1.626237 | 0.235438 | 6.907281 | 4.94E-12 | 4.46E-11 |
| CNAG_01510 | 3037.481 | 0.14779 | 0.065251 | 2.264946 | 0.023516 | 0.048834 |
| CNAG_12963 | 197.832 | 0.501321 | 0.172703 | 2.902797 | 0.003698 | 0.009543 |
| CNAG_01512 | 1267.667 | 2.138551 | 0.089649 | 23.8546 | 9.07E-126 | 8.38E-124 |
| CNAG_01513 | 608.1731 | -0.82321 | 0.104226 | -7.89831 | 2.83E-15 | 3.17E-14 |
| CNAG_07590 | 2199.896 | -0.36899 | 0.080652 | -4.57507 | 4.76E-06 | 2.22E-05 |
| CNAG_07591 | 1254.248 | -0.28875 | 0.083045 | -3.47702 | 0.000507 | 0.001595 |
| CNAG_01519 | 385.5613 | 0.38999 | 0.118415 | 3.293426 | 0.00099 | 0.002916 |
| CNAG_01520 | 1384.16 | -0.18404 | 0.074207 | -2.48014 | 0.013133 | 0.029439 |
| CNAG_01522 | 1641.649 | 0.236411 | 0.087314 | 2.707603 | 0.006777 | 0.016336 |
| CNAG_01523 | 3989.639 | -0.42349 | 0.062392 | -6.78757 | 1.14E-11 | 1.00E-10 |

| | | | | | | |
|---|---|---|---|---|---|---|
| CNAG_01524 | 1657.691 | -0.25648 | 0.075093 | -3.41549 | 0.000637 | 0.001962 |
| CNAG_01525 | 288.6828 | 0.7402 | 0.15021 | 4.927772 | 8.32E-07 | 4.31E-06 |
| CNAG_12967 | 212.7659 | 2.26393 | 0.180268 | 12.5587 | 3.56E-36 | 8.75E-35 |
| CNAG_01526 | 4339.263 | 0.978428 | 0.063971 | 15.29485 | 8.28E-53 | 3.00E-51 |
| CNAG_12968 | 105.7123 | 2.605074 | 0.244708 | 10.64566 | 1.83E-26 | 3.29E-25 |
| CNAG_01527 | 54.83171 | 3.064558 | 0.374606 | 8.180747 | 2.82E-16 | 3.33E-15 |
| CNAG_01528 | 1676.61 | 0.813218 | 0.07537 | 10.78962 | 3.85E-27 | 7.18E-26 |
| CNAG_01529 | 1498.115 | 0.421954 | 0.089243 | 4.728143 | 2.27E-06 | 1.11E-05 |
| CNAG_01531 | 1097.263 | -0.48729 | 0.082688 | -5.89304 | 3.79E-09 | 2.62E-08 |
| CNAG_01532 | 1899.261 | -0.2116 | 0.074875 | -2.82598 | 0.004714 | 0.011847 |
| CNAG_01533 | 1807.66 | 1.184298 | 0.074606 | 15.87396 | 9.60E-57 | 3.79E-55 |
| CNAG_01535 | 4783.081 | 0.218647 | 0.058337 | 3.748006 | 0.000178 | 0.00062 |
| CNAG_12969 | 34.47326 | 1.944206 | 0.419607 | 4.633397 | 3.60E-06 | 1.71E-05 |
| CNAG_01539 | 12073.55 | 0.381822 | 0.082222 | 4.643815 | 3.42E-06 | 1.63E-05 |
| CNAG_12970 | 56.32869 | 0.995131 | 0.286485 | 3.473585 | 0.000514 | 0.001614 |
| CNAG_01540 | 1088.248 | 1.248253 | 0.155188 | 8.04349 | 8.73E-16 | 1.01E-14 |
| CNAG_01541 | 356.5454 | 1.247839 | 0.159994 | 7.799262 | 6.23E-15 | 6.81E-14 |
| CNAG_01542 | 292.1181 | 3.475725 | 0.176933 | 19.64431 | 6.47E-86 | 3.95E-84 |
| CNAG_01544 | 5537.488 | 0.26776 | 0.063192 | 4.237261 | 2.26E-05 | 9.26E-05 |
| CNAG_01545 | 878.7536 | 1.799638 | 0.093674 | 19.21172 | 2.95E-82 | 1.68E-80 |
| CNAG_01548 | 5614.319 | -0.35258 | 0.060141 | -5.86259 | 4.56E-09 | 3.12E-08 |
| CNAG_01549 | 3194.767 | -0.15862 | 0.069562 | -2.2803 | 0.02259 | 0.047128 |
| CNAG_01550 | 1473.259 | -0.18119 | 0.08237 | -2.19973 | 0.027826 | 0.056586 |
| CNAG_01551 | 574.8408 | -0.46076 | 0.128727 | -3.57941 | 0.000344 | 0.001128 |
| CNAG_01552 | 743.696 | -0.28 | 0.10527 | -2.65984 | 0.007818 | 0.018569 |
| CNAG_01554 | 1353.316 | -0.27083 | 0.091349 | -2.96477 | 0.003029 | 0.007988 |
| CNAG_01558 | 10926.01 | -0.55204 | 0.108619 | -5.08237 | 3.73E-07 | 2.03E-06 |
| CNAG_01560 | 76.6077 | -0.63483 | 0.282605 | -2.24636 | 0.024681 | 0.050967 |
| CNAG_01561 | 1869.352 | -0.30235 | 0.090247 | -3.35022 | 0.000807 | 0.002433 |
| CNAG_01562 | 19430.9 | -1.69192 | 0.0822 | -20.583 | 3.90E-94 | 2.60E-92 |
| CNAG_01563 | 1579.579 | -0.26382 | 0.088601 | -2.97766 | 0.002905 | 0.007695 |
| CNAG_01565 | 2401.765 | -0.39105 | 0.085111 | -4.59453 | 4.34E-06 | 2.04E-05 |
| CNAG_01566 | 1006.604 | -0.29912 | 0.091931 | -3.25375 | 0.001139 | 0.003306 |
| CNAG_01567 | 210.4955 | 0.855731 | 0.156917 | 5.453407 | 4.94E-08 | 3.00E-07 |
| CNAG_01568 | 6099.012 | -0.13795 | 0.070223 | -1.96441 | 0.049482 | 0.092885 |
| CNAG_01572 | 684.6005 | 0.745701 | 0.114481 | 6.513766 | 7.33E-11 | 5.99E-10 |
| CNAG_01574 | 4257.227 | 0.925974 | 0.086721 | 10.6776 | 1.30E-26 | 2.35E-25 |
| CNAG_01575 | 5299.924 | 0.134021 | 0.065439 | 2.04804 | 0.040556 | 0.078448 |
| CNAG_01576 | 1723.864 | 0.192725 | 0.082484 | 2.33653 | 0.019464 | 0.041392 |
| CNAG_01577 | 33814.57 | -0.32209 | 0.060027 | -5.36583 | 8.06E-08 | 4.79E-07 |
| CNAG_01583 | 2504.068 | -0.16111 | 0.069648 | -2.31319 | 0.020712 | 0.043614 |
| CNAG_01584 | 673.3143 | -0.52874 | 0.138357 | -3.82154 | 0.000133 | 0.000473 |
| CNAG_01586 | 14952.77 | -0.20053 | 0.087723 | -2.28598 | 0.022255 | 0.046538 |
| CNAG_01587 | 2048.136 | -0.21102 | 0.08286 | -2.54677 | 0.010873 | 0.024936 |
| CNAG_01592 | 184.6095 | 0.466178 | 0.164894 | 2.827129 | 0.004697 | 0.011819 |
| CNAG_01593 | 3671.467 | 0.138407 | 0.068534 | 2.019535 | 0.043432 | 0.08314 |
| CNAG_01594 | 5685.922 | 0.721405 | 0.080867 | 8.920836 | 4.63E-19 | 6.25E-18 |

| | | | | | | |
|---|---|---|---|---|---|---|
| CNAG_01596 | 1793.527 | 0.459464 | 0.08178 | 5.6183 | 1.93E-08 | 1.22E-07 |
| CNAG_01598 | 4256.041 | -0.27364 | 0.060394 | -4.53092 | 5.87E-06 | 2.70E-05 |
| CNAG_01600 | 1839.798 | -0.22207 | 0.094672 | -2.34573 | 0.01899 | 0.040512 |
| CNAG_01601 | 463.6564 | 0.836511 | 0.111035 | 7.533777 | 4.93E-14 | 5.15E-13 |
| CNAG_01602 | 1170.942 | 0.349538 | 0.09021 | 3.874694 | 0.000107 | 0.000389 |
| CNAG_01603 | 4136.171 | 0.480547 | 0.070807 | 6.786708 | 1.15E-11 | 1.00E-10 |
| CNAG_01604 | 907.9513 | 1.641777 | 0.096552 | 17.00409 | 7.66E-65 | 3.56E-63 |
| CNAG_01605 | 542.7246 | 1.283427 | 0.144915 | 8.856392 | 8.26E-19 | 1.10E-17 |
| CNAG_01609 | 2448.384 | -0.34005 | 0.067907 | -5.00765 | 5.51E-07 | 2.93E-06 |
| CNAG_01610 | 1005.981 | -0.3936 | 0.091212 | -4.31525 | 1.59E-05 | 6.73E-05 |
| CNAG_12977 | 56.54712 | 0.971854 | 0.292528 | 3.32226 | 0.000893 | 0.002656 |
| CNAG_01613 | 1914.763 | 0.196603 | 0.079877 | 2.461319 | 0.013843 | 0.0308 |
| CNAG_01614 | 1556.004 | -0.35452 | 0.08283 | -4.28011 | 1.87E-05 | 7.79E-05 |
| CNAG_01615 | 626.4529 | -0.75215 | 0.110819 | -6.78724 | 1.14E-11 | 1.00E-10 |
| CNAG_01616 | 3300.052 | 0.178521 | 0.062502 | 2.856266 | 0.004287 | 0.010912 |
| CNAG_07593 | 1550.832 | 0.34153 | 0.076507 | 4.464034 | 8.04E-06 | 3.61E-05 |
| CNAG_01619 | 682.8328 | 0.426307 | 0.094918 | 4.491339 | 7.08E-06 | 3.22E-05 |
| CNAG_12978 | 89.51796 | 0.95793 | 0.257369 | 3.722003 | 0.000198 | 0.000682 |
| CNAG_01621 | 2845.052 | 0.637304 | 0.107021 | 5.954916 | 2.60E-09 | 1.82E-08 |
| CNAG_01622 | 781.0034 | -0.53171 | 0.092864 | -5.7257 | 1.03E-08 | 6.76E-08 |
| CNAG_01626 | 2270.683 | -0.43489 | 0.069986 | -6.21405 | 5.16E-10 | 3.89E-09 |
| CNAG_01627 | 1041.752 | 0.189836 | 0.086986 | 2.182386 | 0.029081 | 0.05877 |
| CNAG_01631 | 1537.35 | -0.30854 | 0.07357 | -4.1939 | 2.74E-05 | 0.000111 |
| CNAG_01633 | 672.9171 | -0.25283 | 0.102377 | -2.46963 | 0.013525 | 0.030186 |
| CNAG_01634 | 4382.875 | 0.15542 | 0.067085 | 2.316762 | 0.020517 | 0.043258 |
| CNAG_01636 | 1143.986 | -0.4319 | 0.08655 | -4.99016 | 6.03E-07 | 3.19E-06 |
| CNAG_01637 | 3100.924 | -0.14912 | 0.070443 | -2.11694 | 0.034265 | 0.067565 |
| CNAG_01639 | 7860.516 | 0.670802 | 0.061101 | 10.97866 | 4.84E-28 | 9.30E-27 |
| CNAG_01641 | 5827.808 | 0.214536 | 0.061867 | 3.467713 | 0.000525 | 0.001645 |
| CNAG_01643 | 1254.507 | 0.251656 | 0.090631 | 2.776701 | 0.005491 | 0.013595 |
| CNAG_01644 | 1938.18 | -0.79862 | 0.071023 | -11.2445 | 2.47E-29 | 4.96E-28 |
| CNAG_01645 | 138.2921 | 0.872557 | 0.258752 | 3.372172 | 0.000746 | 0.002273 |
| CNAG_01648 | 23364.05 | -0.67409 | 0.08938 | -7.54187 | 4.63E-14 | 4.85E-13 |
| CNAG_01650 | 1619.552 | -0.33577 | 0.105119 | -3.19418 | 0.001402 | 0.003988 |
| CNAG_01651 | 202.7324 | -0.34428 | 0.171565 | -2.00669 | 0.044783 | 0.085425 |
| CNAG_01652 | 1375.289 | 0.215648 | 0.086548 | 2.491655 | 0.012715 | 0.02862 |
| CNAG_01653 | 4598.715 | 5.715659 | 0.112314 | 50.89017 | 0 | 0 |
| CNAG_12980 | 7.150087 | 3.78792 | 1.08427 | 3.493521 | 0.000477 | 0.001509 |
| CNAG_01654 | 1252.45 | -0.15913 | 0.078893 | -2.01707 | 0.043688 | 0.083553 |
| CNAG_12981 | 23.98624 | 1.001018 | 0.482929 | 2.072806 | 0.03819 | 0.074439 |
| CNAG_01655 | 4844.696 | 0.218883 | 0.057978 | 3.775253 | 0.00016 | 0.000563 |
| CNAG_01656 | 531.8065 | 0.318019 | 0.119968 | 2.650857 | 0.008029 | 0.018987 |
| CNAG_01657 | 6646.726 | -0.51629 | 0.063787 | -8.09395 | 5.78E-16 | 6.71E-15 |
| CNAG_08004 | 133.7896 | 0.453826 | 0.209691 | 2.164261 | 0.030444 | 0.061144 |
| CNAG_01665 | 719.8431 | -0.4759 | 0.098035 | -4.8544 | 1.21E-06 | 6.12E-06 |
| CNAG_01668 | 83.38234 | 2.890876 | 0.293048 | 9.864846 | 5.91E-23 | 9.55E-22 |
| CNAG_01669 | 853.5653 | -0.25013 | 0.102011 | -2.45199 | 0.014207 | 0.031497 |

| | | | | | | |
|---|---|---|---|---|---|---|
| CNAG_01670 | 305.2806 | -0.42014 | 0.15055 | -2.79069 | 0.00526 | 0.013085 |
| CNAG_01673 | 1737.929 | -0.24697 | 0.084616 | -2.9187 | 0.003515 | 0.009131 |
| CNAG_01674 | 135.0829 | 1.416179 | 0.196797 | 7.196133 | 6.19E-13 | 6.02E-12 |
| CNAG_01675 | 892.9991 | -0.36346 | 0.08554 | -4.24898 | 2.15E-05 | 8.85E-05 |
| CNAG_01676 | 1916.713 | -0.23701 | 0.078181 | -3.03162 | 0.002432 | 0.006571 |
| CNAG_01677 | 1664.872 | -0.25322 | 0.079057 | -3.20299 | 0.00136 | 0.00388 |
| CNAG_01678 | 2611.137 | -0.34663 | 0.071138 | -4.87262 | 1.10E-06 | 5.61E-06 |
| CNAG_01679 | 4209.135 | -0.22739 | 0.071076 | -3.19928 | 0.001378 | 0.003928 |
| CNAG_12984 | 37.67941 | 1.507274 | 0.393089 | 3.834438 | 0.000126 | 0.000452 |
| CNAG_12985 | 903.1014 | -0.6411 | 0.099749 | -6.42708 | 1.30E-10 | 1.04E-09 |
| CNAG_12987 | 174.0332 | 0.721531 | 0.228729 | 3.154519 | 0.001608 | 0.004518 |
| CNAG_01682 | 10676.52 | -0.22357 | 0.056548 | -3.95361 | 7.70E-05 | 0.000288 |
| CNAG_07596 | 129.8324 | -0.61808 | 0.294091 | -2.10165 | 0.035584 | 0.069827 |
| CNAG_12989 | 54.4519 | 1.026096 | 0.317762 | 3.229137 | 0.001242 | 0.003565 |
| CNAG_01683 | 8836.042 | -0.39295 | 0.088546 | -4.43782 | 9.09E-06 | 4.03E-05 |
| CNAG_01686 | 1682.173 | -0.73475 | 0.114495 | -6.41732 | 1.39E-10 | 1.11E-09 |
| CNAG_12991 | 98.29096 | 0.988705 | 0.245458 | 4.027998 | 5.63E-05 | 0.000216 |
| CNAG_01689 | 61.47184 | 1.073352 | 0.275009 | 3.902973 | 9.50E-05 | 0.00035 |
| CNAG_01690 | 732.5371 | 0.480825 | 0.12549 | 3.831566 | 0.000127 | 0.000456 |
| CNAG_01691 | 1071.48 | 0.230483 | 0.092772 | 2.484395 | 0.012977 | 0.029162 |
| CNAG_01692 | 1509.529 | -0.42052 | 0.080276 | -5.23837 | 1.62E-07 | 9.23E-07 |
| CNAG_01695 | 178.2578 | 0.868498 | 0.194138 | 4.473616 | 7.69E-06 | 3.47E-05 |
| CNAG_01696 | 4271.873 | 0.147149 | 0.073674 | 1.997294 | 0.045793 | 0.087005 |
| CNAG_01698 | 575.0264 | 0.269561 | 0.112148 | 2.403611 | 0.016234 | 0.035337 |
| CNAG_01699 | 917.0707 | 0.53031 | 0.087209 | 6.080891 | 1.20E-09 | 8.70E-09 |
| CNAG_07598 | 5238.9 | 0.273053 | 0.060077 | 4.545019 | 5.49E-06 | 2.54E-05 |
| CNAG_12994 | 304.4258 | 0.613866 | 0.162969 | 3.766773 | 0.000165 | 0.00058 |
| CNAG_01701 | 1334.867 | 0.495819 | 0.11267 | 4.400626 | 1.08E-05 | 4.71E-05 |
| CNAG_01702 | 613.8367 | 0.622193 | 0.098176 | 6.33753 | 2.33E-10 | 1.83E-09 |
| CNAG_01704 | 7899.673 | 0.987604 | 0.076643 | 12.88582 | 5.41E-38 | 1.39E-36 |
| CNAG_08005 | 161.9636 | 0.65394 | 0.183442 | 3.564833 | 0.000364 | 0.001186 |
| CNAG_01707 | 1467.605 | -0.22823 | 0.093477 | -2.44154 | 0.014625 | 0.032275 |
| CNAG_01709 | 5519.012 | 0.363363 | 0.063593 | 5.713917 | 1.10E-08 | 7.21E-08 |
| CNAG_01711 | 3793.125 | -0.32132 | 0.061957 | -5.18616 | 2.15E-07 | 1.20E-06 |
| CNAG_01712 | 1178.347 | -0.32988 | 0.0825 | -3.99852 | 6.37E-05 | 0.000242 |
| CNAG_01714 | 71.10475 | 1.580522 | 0.339507 | 4.655351 | 3.23E-06 | 1.54E-05 |
| CNAG_01715 | 5198.375 | 0.3136 | 0.066773 | 4.696494 | 2.65E-06 | 1.28E-05 |
| CNAG_01716 | 1253.266 | -0.21381 | 0.105038 | -2.03551 | 0.0418 | 0.080395 |
| CNAG_01717 | 799.2558 | -0.32894 | 0.095659 | -3.43872 | 0.000584 | 0.001819 |
| CNAG_01719 | 216.5917 | 0.866299 | 0.170337 | 5.0858 | 3.66E-07 | 2.00E-06 |
| CNAG_01721 | 5368.869 | 1.2109 | 0.081862 | 14.79195 | 1.65E-49 | 5.52E-48 |
| CNAG_01722 | 3888.825 | -0.44049 | 0.069956 | -6.29669 | 3.04E-10 | 2.35E-09 |
| CNAG_01727 | 46865.22 | -0.21154 | 0.077938 | -2.71418 | 0.006644 | 0.016053 |
| CNAG_01728 | 1034.499 | -0.18899 | 0.087027 | -2.17158 | 0.029888 | 0.06019 |
| CNAG_01729 | 2163.749 | 0.324275 | 0.072816 | 4.45334 | 8.45E-06 | 3.78E-05 |
| CNAG_01732 | 1722.953 | 0.177347 | 0.082928 | 2.13857 | 0.03247 | 0.064607 |
| CNAG_01733 | 6152.503 | -0.4459 | 0.072623 | -6.13997 | 8.25E-10 | 6.07E-09 |

| | | | | | | |
|---|---|---|---|---|---|---|
| CNAG_01735 | 1185.054 | -1.32795 | 0.107729 | -12.3267 | 6.51E-35 | 1.52E-33 |
| CNAG_01736 | 424.4583 | -0.64424 | 0.119243 | -5.40275 | 6.56E-08 | 3.94E-07 |
| CNAG_01737 | 3588.793 | -0.98039 | 0.088476 | -11.0808 | 1.56E-28 | 3.01E-27 |
| CNAG_01741 | 408.2973 | -0.3335 | 0.134152 | -2.48599 | 0.012919 | 0.029048 |
| CNAG_01742 | 71.82826 | -2.70509 | 0.309668 | -8.73544 | 2.43E-18 | 3.15E-17 |
| CNAG_01743 | 309.4366 | -0.27237 | 0.128234 | -2.12404 | 0.033667 | 0.066629 |
| CNAG_01744 | 8366.651 | -0.74415 | 0.059045 | -12.603 | 2.03E-36 | 5.05E-35 |
| CNAG_01745 | 7925.409 | -0.81936 | 0.07861 | -10.4231 | 1.95E-25 | 3.42E-24 |
| CNAG_07600 | 602.9265 | 0.444179 | 0.104401 | 4.254537 | 2.09E-05 | 8.66E-05 |
| CNAG_01750 | 1572.564 | -0.52569 | 0.138319 | -3.8006 | 0.000144 | 0.000512 |
| CNAG_01751 | 699.2442 | -0.43716 | 0.166093 | -2.63203 | 0.008488 | 0.019985 |
| CNAG_01753 | 3988.523 | -0.78571 | 0.08605 | -9.13088 | 6.79E-20 | 9.55E-19 |
| CNAG_01754 | 2200.266 | 0.327471 | 0.067356 | 4.861809 | 1.16E-06 | 5.90E-06 |
| CNAG_01755 | 5101.757 | 0.823059 | 0.067757 | 12.14717 | 5.94E-34 | 1.36E-32 |
| CNAG_08006 | 243.1226 | -0.36328 | 0.152244 | -2.38618 | 0.017025 | 0.03683 |
| CNAG_01758 | 113.8314 | -0.60563 | 0.20299 | -2.98354 | 0.002849 | 0.007568 |
| CNAG_01759 | 23.32815 | -1.71668 | 0.505748 | -3.39434 | 0.000688 | 0.002108 |
| CNAG_07601 | 476.9639 | -0.63311 | 0.118323 | -5.35069 | 8.76E-08 | 5.18E-07 |
| CNAG_01760 | 331.2346 | 0.360463 | 0.145384 | 2.479382 | 0.013161 | 0.029493 |
| CNAG_01761 | 12938.34 | -0.58808 | 0.065206 | -9.01876 | 1.90E-19 | 2.62E-18 |
| CNAG_01764 | 1003.135 | 0.181351 | 0.086159 | 2.104854 | 0.035304 | 0.069379 |
| CNAG_07604 | 425.2685 | 0.538403 | 0.136278 | 3.95077 | 7.79E-05 | 0.000292 |
| CNAG_01768 | 1589.034 | 0.46817 | 0.100593 | 4.654119 | 3.25E-06 | 1.55E-05 |
| CNAG_01773 | 553.691 | -0.38653 | 0.108699 | -3.55599 | 0.000377 | 0.001222 |
| CNAG_13002 | 10.22632 | 1.428006 | 0.673682 | 2.119704 | 0.034031 | 0.067153 |
| CNAG_01776 | 210.2642 | 0.664073 | 0.17325 | 3.833036 | 0.000127 | 0.000454 |
| CNAG_01777 | 1491.263 | -0.43144 | 0.109777 | -3.93013 | 8.49E-05 | 0.000316 |
| CNAG_01778 | 1573.026 | -0.19981 | 0.081022 | -2.46615 | 0.013657 | 0.030438 |
| CNAG_01780 | 5157.107 | -0.19855 | 0.074355 | -2.67031 | 0.007578 | 0.018052 |
| CNAG_01789 | 2955.037 | -0.19095 | 0.074624 | -2.55887 | 0.010501 | 0.024153 |
| CNAG_01790 | 887.3911 | -0.26427 | 0.094135 | -2.8073 | 0.004996 | 0.012487 |
| CNAG_01792 | 1778.478 | 0.176669 | 0.080189 | 2.203161 | 0.027583 | 0.056176 |
| CNAG_13006 | 148.1429 | -0.43681 | 0.187328 | -2.33181 | 0.019711 | 0.041857 |
| CNAG_01793 | 875.4238 | 0.329598 | 0.091211 | 3.613569 | 0.000302 | 0.001 |
| CNAG_01794 | 1648.059 | 2.146031 | 0.253511 | 8.465243 | 2.56E-17 | 3.17E-16 |
| CNAG_01795 | 311.5357 | -0.51386 | 0.135346 | -3.79661 | 0.000147 | 0.000519 |
| CNAG_01796 | 401.0963 | 1.360657 | 0.157787 | 8.623405 | 6.50E-18 | 8.25E-17 |
| CNAG_01797 | 1956.758 | -0.2351 | 0.076226 | -3.08425 | 0.002041 | 0.005602 |
| CNAG_01799 | 11767.21 | 0.414891 | 0.073045 | 5.679941 | 1.35E-08 | 8.70E-08 |
| CNAG_01800 | 2297.946 | 0.635342 | 0.111531 | 5.69654 | 1.22E-08 | 7.95E-08 |
| CNAG_01802 | 2235.179 | 1.428805 | 0.0743 | 19.23027 | 2.07E-82 | 1.18E-80 |
| CNAG_01803 | 598.5234 | 0.375238 | 0.127735 | 2.937622 | 0.003307 | 0.008633 |
| CNAG_13007 | 138.9394 | -0.46862 | 0.201726 | -2.32304 | 0.020177 | 0.042718 |
| CNAG_01809 | 1317.502 | -0.30367 | 0.100212 | -3.0303 | 0.002443 | 0.006593 |
| CNAG_01813 | 12079.99 | -0.18008 | 0.053289 | -3.37928 | 0.000727 | 0.00222 |
| CNAG_01816 | 197.1749 | 0.412194 | 0.167593 | 2.459498 | 0.013913 | 0.03091 |
| CNAG_01824 | 3502.081 | -0.15331 | 0.071417 | -2.14664 | 0.031822 | 0.063534 |

| | | | | | | |
|---|---|---|---|---|---|---|
| CNAG_01828 | 1181.246 | -0.29515 | 0.080366 | -3.67263 | 0.00024 | 0.000813 |
| CNAG_01832 | 590.2706 | -0.35989 | 0.109653 | -3.28205 | 0.001031 | 0.003022 |
| CNAG_01834 | 1847.638 | -0.22534 | 0.073489 | -3.06638 | 0.002167 | 0.005908 |
| CNAG_01835 | 2243.464 | -0.2597 | 0.068826 | -3.77332 | 0.000161 | 0.000567 |
| CNAG_01836 | 265.7054 | 1.31533 | 0.158303 | 8.308921 | 9.66E-17 | 1.16E-15 |
| CNAG_01837 | 2838.49 | 0.403177 | 0.068289 | 5.904015 | 3.55E-09 | 2.46E-08 |
| CNAG_01839 | 2406.459 | -0.58345 | 0.079497 | -7.33924 | 2.15E-13 | 2.16E-12 |
| CNAG_01841 | 223.9684 | -0.54736 | 0.208703 | -2.62268 | 0.008724 | 0.020488 |
| CNAG_01843 | 2542.432 | 0.362402 | 0.082424 | 4.396779 | 1.10E-05 | 4.79E-05 |
| CNAG_01845 | 5825.171 | 0.348485 | 0.065588 | 5.313268 | 1.08E-07 | 6.29E-07 |
| CNAG_01846 | 15998.86 | 2.751478 | 0.104733 | 26.2714 | 4.07E-152 | 5.34E-150 |
| CNAG_01847 | 286.6384 | -0.49452 | 0.175329 | -2.82053 | 0.004794 | 0.012024 |
| CNAG_01848 | 2772.102 | 0.198753 | 0.066662 | 2.981506 | 0.002868 | 0.007606 |
| CNAG_01852 | 2696.529 | -0.31517 | 0.084929 | -3.71095 | 0.000206 | 0.000709 |
| CNAG_01853 | 1228.237 | -0.25172 | 0.089942 | -2.79864 | 0.005132 | 0.012795 |
| CNAG_01854 | 23628.15 | 1.238998 | 0.074159 | 16.7074 | 1.16E-62 | 5.17E-61 |
| CNAG_01855 | 1191.397 | 0.751194 | 0.092781 | 8.096388 | 5.66E-16 | 6.58E-15 |
| CNAG_01856 | 285.7656 | 0.771902 | 0.143829 | 5.366808 | 8.01E-08 | 4.77E-07 |
| CNAG_01858 | 35.12308 | -1.74619 | 0.408628 | -4.27331 | 1.93E-05 | 8.01E-05 |
| CNAG_01861 | 1214.456 | 0.400776 | 0.097047 | 4.129687 | 3.63E-05 | 0.000144 |
| CNAG_01863 | 4493.617 | -0.28124 | 0.0817 | -3.44236 | 0.000577 | 0.001798 |
| CNAG_01864 | 1286.559 | -0.22993 | 0.079395 | -2.89601 | 0.003779 | 0.009733 |
| CNAG_01865 | 543.3564 | -0.53712 | 0.113284 | -4.74134 | 2.12E-06 | 1.04E-05 |
| CNAG_01868 | 1163.654 | -0.33455 | 0.08183 | -4.08839 | 4.34E-05 | 0.00017 |
| CNAG_01870 | 4177.938 | -0.23732 | 0.068355 | -3.47191 | 0.000517 | 0.001624 |
| CNAG_01871 | 917.5772 | -0.60201 | 0.099415 | -6.05557 | 1.40E-09 | 1.01E-08 |
| CNAG_01872 | 202.0434 | -0.43154 | 0.159522 | -2.70519 | 0.006827 | 0.016426 |
| CNAG_01876 | 1341.534 | 0.191366 | 0.086227 | 2.219337 | 0.026464 | 0.054184 |
| CNAG_01879 | 891.4736 | -0.48108 | 0.087953 | -5.46972 | 4.51E-08 | 2.75E-07 |
| CNAG_01880 | 1421.555 | -0.32957 | 0.096415 | -3.41826 | 0.00063 | 0.001948 |
| CNAG_01881 | 4172.309 | 0.477224 | 0.063003 | 7.574655 | 3.60E-14 | 3.80E-13 |
| CNAG_01882 | 505.909 | -0.79235 | 0.150005 | -5.28212 | 1.28E-07 | 7.38E-07 |
| CNAG_07611 | 24.65236 | 1.225239 | 0.513554 | 2.385805 | 0.017042 | 0.036849 |
| CNAG_07612 | 78.18468 | -1.58011 | 0.252796 | -6.25055 | 4.09E-10 | 3.11E-09 |
| CNAG_13015 | 51.00597 | -2.46097 | 0.34141 | -7.20824 | 5.67E-13 | 5.53E-12 |
| CNAG_01889 | 525.0027 | -0.65005 | 0.125863 | -5.16471 | 2.41E-07 | 1.34E-06 |
| CNAG_01891 | 686.3532 | 0.271657 | 0.098382 | 2.761252 | 0.005758 | 0.014161 |
| CNAG_01892 | 1516.158 | -0.34158 | 0.103976 | -3.28516 | 0.001019 | 0.002993 |
| CNAG_01893 | 1435.606 | -0.58651 | 0.082182 | -7.13677 | 9.55E-13 | 9.13E-12 |
| CNAG_01897 | 4476.417 | -0.39616 | 0.064235 | -6.16735 | 6.94E-10 | 5.17E-09 |
| CNAG_01898 | 2982.163 | 0.22751 | 0.07377 | 3.084048 | 0.002042 | 0.005604 |
| CNAG_01900 | 933.323 | -0.3075 | 0.089566 | -3.43322 | 0.000596 | 0.001855 |
| CNAG_13017 | 139.236 | 0.674592 | 0.218205 | 3.091558 | 0.001991 | 0.005492 |
| CNAG_01901 | 10.63588 | 1.565967 | 0.749014 | 2.090705 | 0.036555 | 0.071593 |
| CNAG_01902 | 2555.075 | -1.00509 | 0.070696 | -14.2171 | 7.18E-46 | 2.24E-44 |
| CNAG_01904 | 1335.877 | 0.879144 | 0.077598 | 11.32953 | 9.37E-30 | 1.91E-28 |
| CNAG_01905 | 2345.201 | 0.26769 | 0.070123 | 3.817421 | 0.000135 | 0.00048 |

| | | | | | | |
|---|---|---|---|---|---|---|
| CNAG_01906 | 500.7163 | -0.37253 | 0.106922 | -3.48412 | 0.000494 | 0.001557 |
| CNAG_13020 | 58.37859 | 1.942038 | 0.30549 | 6.357135 | 2.06E-10 | 1.61E-09 |
| CNAG_01908 | 2458.667 | 2.669371 | 0.074433 | 35.86269 | 1.17E-281 | 4.99E-279 |
| CNAG_01909 | 465.5863 | -0.28999 | 0.114521 | -2.53222 | 0.011334 | 0.0259 |
| CNAG_01910 | 1176.004 | -0.19055 | 0.083785 | -2.27427 | 0.02295 | 0.047782 |
| CNAG_01912 | 4896.121 | -0.18496 | 0.07763 | -2.38252 | 0.017195 | 0.03715 |
| CNAG_01913 | 262.207 | 0.784953 | 0.156873 | 5.003762 | 5.62E-07 | 2.99E-06 |
| CNAG_01914 | 1274.712 | -0.38559 | 0.09194 | -4.19396 | 2.74E-05 | 0.000111 |
| CNAG_01915 | 8669.28 | 0.19609 | 0.060475 | 3.242484 | 0.001185 | 0.003424 |
| CNAG_01916 | 2262.735 | 0.500304 | 0.080793 | 6.192434 | 5.92E-10 | 4.43E-09 |
| CNAG_13022 | 57.86226 | 0.628216 | 0.287988 | 2.181396 | 0.029154 | 0.058888 |
| CNAG_01919 | 691.5372 | -0.29125 | 0.102711 | -2.83567 | 0.004573 | 0.011536 |
| CNAG_01921 | 126.286 | 1.176132 | 0.192878 | 6.09781 | 1.08E-09 | 7.85E-09 |
| CNAG_13023 | 22.80761 | 1.698066 | 0.475521 | 3.570963 | 0.000356 | 0.001161 |
| CNAG_01924 | 766.6617 | -0.77417 | 0.099483 | -7.78188 | 7.15E-15 | 7.80E-14 |
| CNAG_01925 | 895.5735 | 1.000138 | 0.147703 | 6.771256 | 1.28E-11 | 1.11E-10 |
| CNAG_01926 | 396.4383 | -0.32935 | 0.120152 | -2.7411 | 0.006123 | 0.014928 |
| CNAG_01928 | 1237.556 | -0.3454 | 0.082081 | -4.20809 | 2.58E-05 | 0.000104 |
| CNAG_01929 | 356.4759 | 0.385753 | 0.129667 | 2.974945 | 0.00293 | 0.007754 |
| CNAG_01930 | 1989.422 | 0.441793 | 0.077675 | 5.687731 | 1.29E-08 | 8.33E-08 |
| CNAG_01936 | 92.20081 | 1.08337 | 0.310347 | 3.490836 | 0.000482 | 0.001523 |
| CNAG_01938 | 3047.72 | -0.16272 | 0.073748 | -2.2064 | 0.027356 | 0.055787 |
| CNAG_01942 | 383.3093 | 0.465814 | 0.125448 | 3.713204 | 0.000205 | 0.000704 |
| CNAG_13033 | 101.4379 | 2.029565 | 0.236753 | 8.572493 | 1.01E-17 | 1.27E-16 |
| CNAG_01944 | 21.33703 | 1.662007 | 0.508729 | 3.26698 | 0.001087 | 0.003164 |
| CNAG_01945 | 108.3131 | 0.884363 | 0.256992 | 3.441207 | 0.000579 | 0.001804 |
| CNAG_01946 | 1006.426 | 0.623742 | 0.121121 | 5.149757 | 2.61E-07 | 1.45E-06 |
| CNAG_01947 | 1911.37 | 1.926479 | 0.294211 | 6.547957 | 5.83E-11 | 4.81E-10 |
| CNAG_01948 | 1470.265 | 0.57997 | 0.076498 | 7.581545 | 3.41E-14 | 3.60E-13 |
| CNAG_01949 | 6496.545 | 2.615744 | 0.249348 | 10.49035 | 9.57E-26 | 1.70E-24 |
| CNAG_01951 | 30819.16 | -0.16828 | 0.078562 | -2.14204 | 0.032191 | 0.064113 |
| CNAG_13034 | 9.492597 | 2.109664 | 0.745262 | 2.830766 | 0.004644 | 0.011711 |
| CNAG_01952 | 1255.404 | 2.418577 | 0.088607 | 27.29554 | 4.79E-164 | 6.96E-162 |
| CNAG_01953 | 591.5363 | 1.47608 | 0.137614 | 10.72624 | 7.66E-27 | 1.40E-25 |
| CNAG_13035 | 55.9242 | 0.611677 | 0.289233 | 2.114826 | 0.034445 | 0.067887 |
| CNAG_01954 | 63.37195 | 1.49256 | 0.28891 | 5.166169 | 2.39E-07 | 1.33E-06 |
| CNAG_13036 | 86.06338 | 0.589837 | 0.232775 | 2.533938 | 0.011279 | 0.02578 |
| CNAG_01958 | 2965.044 | -0.29114 | 0.07328 | -3.97294 | 7.10E-05 | 0.000268 |
| CNAG_01959 | 2274.248 | -0.48614 | 0.071523 | -6.797 | 1.07E-11 | 9.41E-11 |
| CNAG_01960 | 200.1807 | 0.927886 | 0.201372 | 4.60783 | 4.07E-06 | 1.92E-05 |
| CNAG_01961 | 5575.974 | -0.23193 | 0.062352 | -3.71968 | 0.000199 | 0.000687 |
| CNAG_01962 | 779.5885 | -0.62658 | 0.092112 | -6.80239 | 1.03E-11 | 9.09E-11 |
| CNAG_01963 | 230.2542 | 1.400707 | 0.171364 | 8.173891 | 2.99E-16 | 3.52E-15 |
| CNAG_01964 | 823.0087 | 1.427093 | 0.131233 | 10.87449 | 1.53E-27 | 2.89E-26 |
| CNAG_01965 | 712.9114 | 3.776662 | 0.123628 | 30.5486 | 5.90E-205 | 1.26E-202 |
| CNAG_07613 | 119.1671 | 0.791825 | 0.221288 | 3.578262 | 0.000346 | 0.001132 |
| CNAG_01968 | 27.29019 | 1.252348 | 0.442629 | 2.82934 | 0.004664 | 0.011752 |

| | | | | | | |
|---|---|---|---|---|---|---|
| CNAG_01969 | 288.5904 | 0.315845 | 0.135785 | 2.326068 | 0.020015 | 0.04242 |
| CNAG_01972 | 1662.282 | 0.182883 | 0.090417 | 2.02267 | 0.043107 | 0.082597 |
| CNAG_01973 | 2606.961 | -0.25759 | 0.072965 | -3.53031 | 0.000415 | 0.001335 |
| CNAG_01975 | 1376.224 | 0.292511 | 0.08335 | 3.509419 | 0.000449 | 0.00143 |
| CNAG_01976 | 36463.94 | -0.14777 | 0.073041 | -2.02311 | 0.043062 | 0.082529 |
| CNAG_01978 | 298.1564 | -0.39556 | 0.146136 | -2.70677 | 0.006794 | 0.016362 |
| CNAG_01981 | 6712.118 | -0.38677 | 0.058731 | -6.58543 | 4.54E-11 | 3.78E-10 |
| CNAG_01982 | 590.2342 | 0.316366 | 0.114025 | 2.77454 | 0.005528 | 0.013681 |
| CNAG_13040 | 104.4732 | 0.614048 | 0.305563 | 2.009563 | 0.044478 | 0.084922 |
| CNAG_01983 | 2146.188 | 0.232218 | 0.085183 | 2.726127 | 0.006408 | 0.015543 |
| CNAG_01984 | 41288.61 | -0.2575 | 0.052897 | -4.86789 | 1.13E-06 | 5.73E-06 |
| CNAG_01986 | 1370.437 | 0.549155 | 0.138761 | 3.957573 | 7.57E-05 | 0.000284 |
| CNAG_13042 | 97.06914 | 1.173745 | 0.249535 | 4.70372 | 2.55E-06 | 1.24E-05 |
| CNAG_01987 | 1945.859 | 0.279401 | 0.07462 | 3.744334 | 0.000181 | 0.000628 |
| CNAG_01990 | 53628.57 | -0.21297 | 0.064098 | -3.32251 | 0.000892 | 0.002654 |
| CNAG_01991 | 13334.45 | 0.467756 | 0.064838 | 7.214241 | 5.42E-13 | 5.31E-12 |
| CNAG_01992 | 124.5093 | 0.60408 | 0.207295 | 2.914115 | 0.003567 | 0.009236 |
| CNAG_01993 | 250.2752 | 1.384313 | 0.147024 | 9.415557 | 4.71E-21 | 7.08E-20 |
| CNAG_01994 | 79.20204 | 0.848565 | 0.31072 | 2.730965 | 0.006315 | 0.015345 |
| CNAG_01995 | 211.6614 | 4.571167 | 0.251791 | 18.15463 | 1.18E-73 | 6.15E-72 |
| CNAG_01997 | 3030.574 | -0.3811 | 0.087784 | -4.34131 | 1.42E-05 | 6.05E-05 |
| CNAG_08012 | 2522.001 | -0.36475 | 0.088265 | -4.13239 | 3.59E-05 | 0.000143 |
| CNAG_01999 | 587.0693 | -0.43362 | 0.131545 | -3.29632 | 0.00098 | 0.002891 |
| CNAG_02000 | 1532.042 | -0.58237 | 0.093263 | -6.2444 | 4.25E-10 | 3.23E-09 |
| CNAG_02001 | 1213.861 | 0.327511 | 0.081883 | 3.999746 | 6.34E-05 | 0.000241 |
| CNAG_08013 | 771.5507 | 0.577659 | 0.098232 | 5.880571 | 4.09E-09 | 2.81E-08 |
| CNAG_02004 | 2923.978 | 0.17213 | 0.072826 | 2.363565 | 0.0181 | 0.038879 |
| CNAG_02005 | 28.89942 | 0.879785 | 0.413641 | 2.126929 | 0.033426 | 0.066265 |
| CNAG_02006 | 1150.643 | -0.19878 | 0.089313 | -2.22566 | 0.026037 | 0.053401 |
| CNAG_02007 | 7954.175 | -0.21758 | 0.065455 | -3.32404 | 0.000887 | 0.002643 |
| CNAG_02009 | 1181.017 | 0.61645 | 0.097821 | 6.301846 | 2.94E-10 | 2.28E-09 |
| CNAG_02011 | 436.7141 | 0.737595 | 0.114254 | 6.455752 | 1.08E-10 | 8.67E-10 |
| CNAG_02013 | 377.7059 | -0.31418 | 0.12679 | -2.47798 | 0.013213 | 0.029593 |
| CNAG_13043 | 158.7199 | -0.7279 | 0.188416 | -3.86326 | 0.000112 | 0.000405 |
| CNAG_02016 | 39.74492 | 1.545181 | 0.372789 | 4.144923 | 3.40E-05 | 0.000136 |
| CNAG_06956 | 286.1157 | -0.75683 | 0.146823 | -5.15473 | 2.54E-07 | 1.41E-06 |
| CNAG_13045 | 129.7858 | -0.86677 | 0.196633 | -4.40807 | 1.04E-05 | 4.57E-05 |
| CNAG_05986 | 32.70191 | -1.16556 | 0.383307 | -3.0408 | 0.00236 | 0.006388 |
| CNAG_05989 | 689.5866 | -0.2277 | 0.108077 | -2.10682 | 0.035133 | 0.069077 |
| CNAG_05990 | 311.9276 | -0.60866 | 0.156517 | -3.88878 | 0.000101 | 0.000369 |
| CNAG_05991 | 133.3364 | -0.58012 | 0.194439 | -2.98354 | 0.002849 | 0.007568 |
| CNAG_05992 | 117.9794 | -0.4065 | 0.205735 | -1.97583 | 0.048174 | 0.090827 |
| CNAG_05993 | 609.2117 | -0.29545 | 0.129258 | -2.28573 | 0.02227 | 0.046556 |
| CNAG_05995 | 1646.063 | 0.265477 | 0.080752 | 3.287553 | 0.001011 | 0.00297 |
| CNAG_08014 | 115.8849 | 2.176807 | 0.233437 | 9.325015 | 1.11E-20 | 1.63E-19 |
| CNAG_05996 | 1208.327 | 1.454345 | 0.08796 | 16.53424 | 2.08E-61 | 8.99E-60 |
| CNAG_05997 | 5903.273 | 0.406413 | 0.068149 | 5.963592 | 2.47E-09 | 1.73E-08 |

| | | | | | | |
|---|---|---|---|---|---|---|
| CNAG_05998 | 1330.26 | 0.330297 | 0.082014 | 4.027321 | 5.64E-05 | 0.000217 |
| CNAG_05999 | 793.7848 | -0.33679 | 0.118588 | -2.84002 | 0.004511 | 0.011394 |
| CNAG_06000 | 1593.41 | -0.81578 | 0.097535 | -8.36398 | 6.06E-17 | 7.40E-16 |
| CNAG_06001 | 1061.194 | -0.55827 | 0.10364 | -5.38668 | 7.18E-08 | 4.30E-07 |
| CNAG_06003 | 96.05841 | 1.073406 | 0.239272 | 4.486128 | 7.25E-06 | 3.29E-05 |
| CNAG_06006 | 1337.591 | -0.19985 | 0.101377 | -1.97137 | 0.048681 | 0.091529 |
| CNAG_06008 | 1089.697 | 1.689201 | 0.087381 | 19.3314 | 2.92E-83 | 1.69E-81 |
| CNAG_06009 | 318.5889 | 3.02462 | 0.191414 | 15.80148 | 3.04E-56 | 1.18E-54 |
| CNAG_06013 | 550.9414 | -0.34588 | 0.119565 | -2.89283 | 0.003818 | 0.009823 |
| CNAG_06016 | 1712.809 | 0.32708 | 0.086208 | 3.794069 | 0.000148 | 0.000524 |
| CNAG_06017 | 179.0337 | 1.433507 | 0.2129 | 6.733229 | 1.66E-11 | 1.44E-10 |
| CNAG_06019 | 1310.989 | -0.31379 | 0.104875 | -2.99204 | 0.002771 | 0.007387 |
| CNAG_06020 | 138.6405 | 0.508835 | 0.202929 | 2.507448 | 0.012161 | 0.027487 |
| CNAG_13051 | 41.70825 | 1.152891 | 0.34835 | 3.309578 | 0.000934 | 0.002771 |
| CNAG_06021 | 2483.568 | 0.325283 | 0.093775 | 3.468755 | 0.000523 | 0.00164 |
| CNAG_06023 | 372.2496 | -0.46857 | 0.118126 | -3.96671 | 7.29E-05 | 0.000275 |
| CNAG_06026 | 1510.393 | -0.55345 | 0.085353 | -6.48425 | 8.92E-11 | 7.26E-10 |
| CNAG_06027 | 22.49336 | 1.528335 | 0.530436 | 2.881279 | 0.003961 | 0.010136 |
| CNAG_06028 | 704.3165 | 0.561989 | 0.100926 | 5.568338 | 2.57E-08 | 1.61E-07 |
| CNAG_06029 | 575.1484 | 0.403116 | 0.103639 | 3.889624 | 0.0001 | 0.000368 |
| CNAG_06030 | 605.1458 | 0.301057 | 0.105542 | 2.852482 | 0.004338 | 0.011025 |
| CNAG_06031 | 561.4359 | 1.289433 | 0.166866 | 7.727373 | 1.10E-14 | 1.19E-13 |
| CNAG_13053 | 30.75384 | 0.856914 | 0.422144 | 2.029907 | 0.042366 | 0.081349 |
| CNAG_06034 | 1971.194 | 0.302798 | 0.095144 | 3.182518 | 0.00146 | 0.00413 |
| CNAG_06035 | 621.4994 | 0.45642 | 0.111385 | 4.097679 | 4.17E-05 | 0.000164 |
| CNAG_06048 | 1024.304 | -0.35404 | 0.088657 | -3.99335 | 6.51E-05 | 0.000247 |
| CNAG_06049 | 1460.385 | -0.37932 | 0.075874 | -4.99936 | 5.75E-07 | 3.05E-06 |
| CNAG_13055 | 120.7904 | 1.592523 | 0.258688 | 6.156164 | 7.45E-10 | 5.52E-09 |
| CNAG_06050 | 242.161 | 1.400902 | 0.158714 | 8.826596 | 1.08E-18 | 1.42E-17 |
| CNAG_06051 | 1416.182 | 1.119044 | 0.117372 | 9.534187 | 1.51E-21 | 2.32E-20 |
| CNAG_06052 | 1601.523 | 1.18166 | 0.09909 | 11.92513 | 8.75E-33 | 1.93E-31 |
| CNAG_13056 | 36.67863 | 3.621844 | 0.488247 | 7.41806 | 1.19E-13 | 1.21E-12 |
| CNAG_13057 | 419.0959 | 1.852964 | 0.135818 | 13.64304 | 2.22E-42 | 6.36E-41 |
| CNAG_07897 | 245.1798 | 2.044705 | 0.205963 | 9.927539 | 3.16E-23 | 5.21E-22 |
| CNAG_06055 | 797.6791 | 1.554841 | 0.140197 | 11.09039 | 1.40E-28 | 2.71E-27 |
| CNAG_06056 | 574.2178 | 1.371578 | 0.112027 | 12.24325 | 1.83E-34 | 4.24E-33 |
| CNAG_07898 | 61.39751 | 0.907028 | 0.281703 | 3.219807 | 0.001283 | 0.003676 |
| CNAG_06060 | 2277.261 | 0.450991 | 0.06852 | 6.58191 | 4.64E-11 | 3.86E-10 |
| CNAG_13059 | 28.3067 | 0.953243 | 0.407362 | 2.340042 | 0.019282 | 0.041037 |
| CNAG_06061 | 6057.817 | -0.17186 | 0.064262 | -2.6744 | 0.007486 | 0.017865 |
| CNAG_06063 | 3616.425 | 1.916314 | 0.075788 | 25.28524 | 4.64E-141 | 5.47E-139 |
| CNAG_06064 | 760.7448 | -0.26508 | 0.098757 | -2.68417 | 0.007271 | 0.017408 |
| CNAG_06065 | 1262.188 | -0.44786 | 0.09474 | -4.72728 | 2.28E-06 | 1.11E-05 |
| CNAG_06066 | 53.947 | 1.084753 | 0.323953 | 3.348491 | 0.000813 | 0.002445 |
| CNAG_06071 | 588.7678 | -0.27209 | 0.098723 | -2.75613 | 0.005849 | 0.01435 |
| CNAG_13060 | 67.92098 | 3.620986 | 0.345695 | 10.47451 | 1.13E-25 | 2.00E-24 |
| CNAG_13061 | 242.6266 | 0.297439 | 0.146207 | 2.034371 | 0.041914 | 0.080596 |

| | | | | | | |
|---|---|---|---|---|---|---|
| CNAG_06074 | 667.979 | -0.71923 | 0.142361 | -5.05214 | 4.37E-07 | 2.36E-06 |
| CNAG_06075 | 6904.386 | -0.59812 | 0.111767 | -5.35147 | 8.72E-08 | 5.16E-07 |
| CNAG_06076 | 2588.443 | -0.19895 | 0.064415 | -3.08862 | 0.002011 | 0.005537 |
| CNAG_06077 | 2655.354 | -0.1602 | 0.070875 | -2.26028 | 0.023804 | 0.049356 |
| CNAG_06081 | 10366.04 | 0.326016 | 0.061819 | 5.273712 | 1.34E-07 | 7.69E-07 |
| CNAG_13063 | 622.956 | -0.54888 | 0.107201 | -5.12007 | 3.05E-07 | 1.68E-06 |
| CNAG_06085 | 1299.183 | 1.545004 | 0.096948 | 15.93637 | 3.54E-57 | 1.40E-55 |
| CNAG_06087 | 1564.722 | -0.3967 | 0.084444 | -4.69782 | 2.63E-06 | 1.27E-05 |
| CNAG_06088 | 3460.521 | -0.64215 | 0.06924 | -9.27419 | 1.79E-20 | 2.60E-19 |
| CNAG_13065 | 147.311 | 0.689533 | 0.198155 | 3.479765 | 0.000502 | 0.001579 |
| CNAG_06094 | 181.1933 | -0.66954 | 0.161576 | -4.14382 | 3.42E-05 | 0.000136 |
| CNAG_06095 | 96910.64 | -0.17046 | 0.068971 | -2.47141 | 0.013458 | 0.03006 |
| CNAG_06096 | 19198.5 | 0.891302 | 0.065677 | 13.57101 | 5.95E-42 | 1.68E-40 |
| CNAG_06098 | 550.6801 | -0.50113 | 0.157746 | -3.17679 | 0.001489 | 0.004204 |
| CNAG_06100 | 944.2578 | 0.26789 | 0.099255 | 2.699005 | 0.006955 | 0.016713 |
| CNAG_06103 | 3545.761 | 0.193404 | 0.070975 | 2.724971 | 0.006431 | 0.015593 |
| CNAG_06104 | 578.2713 | -1.68605 | 0.129917 | -12.978 | 1.63E-38 | 4.26E-37 |
| CNAG_06105 | 124.0833 | -0.5208 | 0.205719 | -2.53159 | 0.011355 | 0.02591 |
| CNAG_06106 | 4552.307 | -0.65812 | 0.082537 | -7.97358 | 1.54E-15 | 1.75E-14 |
| CNAG_06107 | 829.337 | -0.49575 | 0.105225 | -4.71132 | 2.46E-06 | 1.20E-05 |
| CNAG_06108 | 527.6809 | -0.28681 | 0.107867 | -2.65897 | 0.007838 | 0.018601 |
| CNAG_06113 | 75164.22 | -0.16212 | 0.078411 | -2.06752 | 0.038685 | 0.075254 |
| CNAG_06115 | 492.0308 | 0.848253 | 0.145606 | 5.825674 | 5.69E-09 | 3.87E-08 |
| CNAG_07904 | 1571.281 | -0.16973 | 0.084795 | -2.00168 | 0.045319 | 0.086285 |
| CNAG_08016 | 1690.669 | 0.202593 | 0.103064 | 1.965711 | 0.049332 | 0.092667 |
| CNAG_06118 | 1508.229 | -0.30628 | 0.083472 | -3.66928 | 0.000243 | 0.000822 |
| CNAG_06120 | 1248.313 | -0.32016 | 0.087532 | -3.65766 | 0.000255 | 0.000856 |
| CNAG_06122 | 1735.724 | 1.010294 | 0.083476 | 12.10283 | 1.02E-33 | 2.32E-32 |
| CNAG_06125 | 335725.8 | -0.13705 | 0.061986 | -2.21095 | 0.027039 | 0.055262 |
| CNAG_06127 | 4200.349 | 0.182284 | 0.081086 | 2.24802 | 0.024575 | 0.050774 |
| CNAG_06128 | 368.371 | -0.43578 | 0.138907 | -3.13717 | 0.001706 | 0.004766 |
| CNAG_06129 | 840.768 | -0.30394 | 0.091505 | -3.32162 | 0.000895 | 0.002661 |
| CNAG_06134 | 7555.058 | 0.384672 | 0.062628 | 6.142199 | 8.14E-10 | 6.00E-09 |
| CNAG_06136 | 857.0709 | 0.716895 | 0.132636 | 5.404989 | 6.48E-08 | 3.90E-07 |
| CNAG_06138 | 3742.196 | 0.936832 | 0.068164 | 13.74383 | 5.55E-43 | 1.65E-41 |
| CNAG_06140 | 4832.84 | -0.42677 | 0.098625 | -4.32724 | 1.51E-05 | 6.41E-05 |
| CNAG_06142 | 1008.835 | 0.292933 | 0.087214 | 3.358783 | 0.000783 | 0.002366 |
| CNAG_06144 | 29151.08 | -0.19456 | 0.077815 | -2.50035 | 0.012407 | 0.027966 |
| CNAG_06145 | 976.6917 | 0.504107 | 0.084807 | 5.944182 | 2.78E-09 | 1.95E-08 |
| CNAG_08018 | 1138.376 | -0.32246 | 0.103092 | -3.12787 | 0.001761 | 0.004906 |
| CNAG_06147 | 2795.445 | 0.420297 | 0.118113 | 3.558435 | 0.000373 | 0.001213 |
| CNAG_06148 | 915.1966 | -0.2154 | 0.096744 | -2.22649 | 0.025981 | 0.053313 |
| CNAG_06151 | 3050.127 | -0.1853 | 0.068447 | -2.70724 | 0.006785 | 0.016349 |
| CNAG_06157 | 1199.916 | -0.46231 | 0.080013 | -5.77791 | 7.56E-09 | 5.05E-08 |
| CNAG_06158 | 2255.801 | 0.288699 | 0.074495 | 3.875419 | 0.000106 | 0.000388 |
| CNAG_06162 | 1210.178 | 0.4851 | 0.089849 | 5.399037 | 6.70E-08 | 4.02E-07 |
| CNAG_06164 | 279.5409 | -0.33329 | 0.137011 | -2.43258 | 0.014992 | 0.032967 |

| | | | | | | |
|---|---|---|---|---|---|---|
| CNAG_06167 | 3750.047 | 0.202484 | 0.065486 | 3.092036 | 0.001988 | 0.005485 |
| CNAG_06168 | 8342.771 | 0.67608 | 0.06055 | 11.16566 | 6.00E-29 | 1.19E-27 |
| CNAG_06169 | 699.7949 | 0.417947 | 0.148001 | 2.823953 | 0.004744 | 0.011915 |
| CNAG_06170 | 2297.452 | -0.40925 | 0.074049 | -5.52676 | 3.26E-08 | 2.02E-07 |
| CNAG_06173 | 742.1221 | -0.48392 | 0.110559 | -4.377 | 1.20E-05 | 5.20E-05 |
| CNAG_06175 | 5945.228 | 0.151396 | 0.06554 | 2.309958 | 0.02089 | 0.043921 |
| CNAG_07908 | 9364.887 | 0.972432 | 0.061766 | 15.74389 | 7.56E-56 | 2.89E-54 |
| CNAG_07909 | 226.7368 | 1.642422 | 0.17373 | 9.453875 | 3.27E-21 | 4.94E-20 |
| CNAG_06180 | 4559.364 | 1.064392 | 0.070396 | 15.12006 | 1.19E-51 | 4.15E-50 |
| CNAG_06185 | 956.5084 | 0.261933 | 0.088356 | 2.964507 | 0.003032 | 0.007993 |
| CNAG_06186 | 728.82 | 0.804746 | 0.117282 | 6.861646 | 6.81E-12 | 6.08E-11 |
| CNAG_13076 | 13.54578 | 4.823436 | 0.995011 | 4.847619 | 1.25E-06 | 6.32E-06 |
| CNAG_07911 | 1420.995 | 3.564105 | 0.103134 | 34.55804 | 1.08E-261 | 3.84E-259 |
| CNAG_06188 | 1137.533 | 1.44968 | 0.0894 | 16.21573 | 3.90E-59 | 1.63E-57 |
| CNAG_06191 | 863.9461 | 0.894418 | 0.090507 | 9.882334 | 4.97E-23 | 8.11E-22 |
| CNAG_06193 | 1175.562 | 0.324524 | 0.097253 | 3.336889 | 0.000847 | 0.00254 |
| CNAG_06194 | 60.05841 | 2.520912 | 0.333222 | 7.565253 | 3.87E-14 | 4.07E-13 |
| CNAG_06197 | 1265.012 | 0.240693 | 0.102278 | 2.353317 | 0.018607 | 0.039809 |
| CNAG_06200 | 877.4704 | -0.50661 | 0.090065 | -5.62495 | 1.86E-08 | 1.18E-07 |
| CNAG_06201 | 1035.546 | -0.48669 | 0.115313 | -4.22058 | 2.44E-05 | 9.90E-05 |
| CNAG_13081 | 16.88897 | 1.862264 | 0.565813 | 3.291306 | 0.000997 | 0.002935 |
| CNAG_06203 | 529.379 | 2.603123 | 0.131862 | 19.74131 | 9.53E-87 | 5.91E-85 |
| CNAG_13082 | 1143.947 | 0.750989 | 0.090652 | 8.284277 | 1.19E-16 | 1.42E-15 |
| CNAG_06204 | 6314.81 | 0.817562 | 0.079304 | 10.30921 | 6.40E-25 | 1.12E-23 |
| CNAG_06206 | 3388.933 | 0.212776 | 0.078276 | 2.718286 | 0.006562 | 0.015869 |
| CNAG_06207 | 1011.767 | 0.577332 | 0.135437 | 4.262732 | 2.02E-05 | 8.37E-05 |
| CNAG_06209 | 96.07593 | 0.701206 | 0.243208 | 2.883155 | 0.003937 | 0.010092 |
| CNAG_07912 | 102.1258 | 0.52765 | 0.233454 | 2.260189 | 0.02381 | 0.049356 |
| CNAG_07914 | 4610.116 | -0.33693 | 0.065704 | -5.128 | 2.93E-07 | 1.62E-06 |
| CNAG_06214 | 1331.993 | -0.69368 | 0.088761 | -7.81516 | 5.49E-15 | 6.03E-14 |
| CNAG_13084 | 54.5278 | 0.823867 | 0.340893 | 2.416792 | 0.015658 | 0.034212 |
| CNAG_06216 | 714.2896 | -0.73861 | 0.115523 | -6.39363 | 1.62E-10 | 1.28E-09 |
| CNAG_06218 | 912.1412 | -0.32425 | 0.099237 | -3.2674 | 0.001085 | 0.003161 |
| CNAG_06219 | 1652.598 | -0.63384 | 0.07432 | -8.52857 | 1.48E-17 | 1.84E-16 |
| CNAG_06220 | 5880.974 | -1.09341 | 0.237786 | -4.59829 | 4.26E-06 | 2.00E-05 |
| CNAG_13085 | 204.7453 | 0.370549 | 0.179168 | 2.068161 | 0.038625 | 0.075178 |
| CNAG_06221 | 1490.518 | 0.353308 | 0.08189 | 4.314443 | 1.60E-05 | 6.75E-05 |
| CNAG_06222 | 40352.15 | -0.22759 | 0.083236 | -2.73422 | 0.006253 | 0.015212 |
| CNAG_06223 | 1287.324 | -0.36252 | 0.079304 | -4.57122 | 4.85E-06 | 2.26E-05 |
| CNAG_06224 | 1273.854 | -0.70778 | 0.091452 | -7.73937 | 9.99E-15 | 1.08E-13 |
| CNAG_06225 | 117.6344 | 1.666085 | 0.291801 | 5.709664 | 1.13E-08 | 7.38E-08 |
| CNAG_06226 | 6759.996 | 1.052833 | 0.068709 | 15.32317 | 5.35E-53 | 1.95E-51 |
| CNAG_06227 | 1493.97 | 0.477649 | 0.073582 | 6.491343 | 8.51E-11 | 6.93E-10 |
| CNAG_06229 | 1433.508 | -0.45409 | 0.083906 | -5.41191 | 6.24E-08 | 3.76E-07 |
| CNAG_13087 | 39.74563 | -0.77674 | 0.35392 | -2.19467 | 0.028187 | 0.057233 |
| CNAG_06231 | 52349.31 | -0.25058 | 0.068141 | -3.6774 | 0.000236 | 0.000798 |
| CNAG_06232 | 630.749 | -0.33194 | 0.098703 | -3.36297 | 0.000771 | 0.002335 |

| | | | | | | |
|---|---|---|---|---|---|---|
| CNAG_13089 | 251.0394 | -0.37796 | 0.190396 | -1.98514 | 0.047129 | 0.089147 |
| CNAG_06235 | 2461.189 | -0.33366 | 0.065392 | -5.10244 | 3.35E-07 | 1.84E-06 |
| CNAG_06236 | 1597.785 | -0.37297 | 0.083137 | -4.48629 | 7.25E-06 | 3.29E-05 |
| CNAG_06237 | 2329.597 | 0.61961 | 0.070181 | 8.828768 | 1.06E-18 | 1.39E-17 |
| CNAG_06238 | 347.165 | 1.815958 | 0.183838 | 9.878025 | 5.18E-23 | 8.44E-22 |
| CNAG_06239 | 486.1242 | 1.991559 | 0.124585 | 15.9855 | 1.61E-57 | 6.43E-56 |
| CNAG_06240 | 8882.27 | 0.59066 | 0.064634 | 9.138538 | 6.33E-20 | 8.92E-19 |
| CNAG_06241 | 5295.096 | 2.089887 | 0.089923 | 23.24093 | 1.76E-119 | 1.52E-117 |
| CNAG_06242 | 10034.01 | 1.763058 | 0.083113 | 21.21286 | 7.27E-100 | 5.27E-98 |
| CNAG_13091 | 163.4241 | 1.28055 | 0.207752 | 6.163842 | 7.10E-10 | 5.28E-09 |
| CNAG_06246 | 22335.11 | -0.41135 | 0.06396 | -6.43134 | 1.26E-10 | 1.01E-09 |
| CNAG_06248 | 7399.99 | 0.365301 | 0.06006 | 6.082271 | 1.18E-09 | 8.63E-09 |
| CNAG_06249 | 266.6735 | -0.38017 | 0.149471 | -2.54342 | 0.010977 | 0.025141 |
| CNAG_06251 | 413.9747 | 1.03906 | 0.136379 | 7.618905 | 2.56E-14 | 2.72E-13 |
| CNAG_06252 | 1041.909 | 1.326685 | 0.105836 | 12.53526 | 4.79E-36 | 1.17E-34 |
| CNAG_07917 | 1077.024 | 0.718706 | 0.092749 | 7.748932 | 9.27E-15 | 1.00E-13 |
| CNAG_07008 | 105.091 | 0.479389 | 0.211324 | 2.268501 | 0.023299 | 0.048433 |
| CNAG_07011 | 1308.056 | -0.52008 | 0.081101 | -6.41275 | 1.43E-10 | 1.14E-09 |
| CNAG_06985 | 783.3759 | -0.29372 | 0.090293 | -3.25292 | 0.001142 | 0.003313 |
| CNAG_06986 | 536.5764 | -0.26137 | 0.113186 | -2.3092 | 0.020933 | 0.043997 |
| CNAG_13094 | 98.35944 | -0.76575 | 0.236349 | -3.23991 | 0.001196 | 0.00345 |
| CNAG_06259 | 497.613 | 2.121455 | 0.129204 | 16.4194 | 1.39E-60 | 5.97E-59 |
| CNAG_06260 | 1622.098 | -1.06868 | 0.089609 | -11.9261 | 8.65E-33 | 1.91E-31 |
| CNAG_07921 | 1130.885 | -0.27129 | 0.108792 | -2.49371 | 0.012642 | 0.028463 |
| CNAG_07922 | 1158.185 | 0.212093 | 0.082418 | 2.573369 | 0.010071 | 0.023243 |
| CNAG_06265 | 313.4909 | -0.54958 | 0.128161 | -4.28819 | 1.80E-05 | 7.54E-05 |
| CNAG_13099 | 504.2077 | 0.339164 | 0.115112 | 2.946367 | 0.003215 | 0.008425 |
| CNAG_13100 | 104.0605 | 0.967556 | 0.220699 | 4.38406 | 1.16E-05 | 5.05E-05 |
| CNAG_07924 | 2240.013 | 0.543821 | 0.095782 | 5.677689 | 1.37E-08 | 8.81E-08 |
| CNAG_06272 | 414.8203 | 0.386822 | 0.12341 | 3.134432 | 0.001722 | 0.004804 |
| CNAG_06274 | 11719.11 | 0.50048 | 0.098466 | 5.082783 | 3.72E-07 | 2.03E-06 |
| CNAG_06276 | 1341.008 | 0.244322 | 0.088258 | 2.768283 | 0.005635 | 0.013905 |
| CNAG_13104 | 137.3187 | 0.400409 | 0.183225 | 2.185343 | 0.028864 | 0.058417 |
| CNAG_06277 | 4805.237 | 0.260363 | 0.061524 | 4.231894 | 2.32E-05 | 9.46E-05 |
| CNAG_06278 | 2264.581 | -0.70064 | 0.07133 | -9.82254 | 9.00E-23 | 1.44E-21 |
| CNAG_07927 | 275.1368 | -0.99311 | 0.150304 | -6.60733 | 3.91E-11 | 3.28E-10 |
| CNAG_06282 | 1391.969 | -0.17421 | 0.084346 | -2.06542 | 0.038884 | 0.075573 |
| CNAG_06283 | 1173.627 | -0.39456 | 0.087046 | -4.53273 | 5.82E-06 | 2.68E-05 |
| CNAG_06285 | 283.9851 | -0.39309 | 0.141898 | -2.7702 | 0.005602 | 0.013832 |
| CNAG_06287 | 2701.728 | -0.21755 | 0.07807 | -2.78658 | 0.005327 | 0.013232 |
| CNAG_06288 | 2280.038 | -0.34792 | 0.071449 | -4.86945 | 1.12E-06 | 5.70E-06 |
| CNAG_06290 | 520.0453 | 0.763066 | 0.175897 | 4.338148 | 1.44E-05 | 6.13E-05 |
| CNAG_06291 | 462.3321 | 1.668814 | 0.171552 | 9.727741 | 2.30E-22 | 3.62E-21 |
| CNAG_06293 | 308.205 | -0.61219 | 0.162471 | -3.76798 | 0.000165 | 0.000578 |
| CNAG_06294 | 169.4888 | 0.884217 | 0.177635 | 4.977719 | 6.43E-07 | 3.39E-06 |
| CNAG_06297 | 528.3282 | 1.075917 | 0.170456 | 6.311977 | 2.75E-10 | 2.14E-09 |
| CNAG_06298 | 2896.7 | 1.696721 | 0.108696 | 15.6098 | 6.24E-55 | 2.34E-53 |

| | | | | | | |
|---|---|---|---|---|---|---|
| CNAG_06299 | 1131.778 | 0.359395 | 0.08458 | 4.249191 | 2.15E-05 | 8.84E-05 |
| CNAG_13108 | 37.07939 | 3.903559 | 0.497704 | 7.843141 | 4.39E-15 | 4.87E-14 |
| CNAG_06300 | 2562.578 | 0.198764 | 0.076731 | 2.590384 | 0.009587 | 0.022251 |
| CNAG_06302 | 130.6913 | 0.642227 | 0.214628 | 2.992284 | 0.002769 | 0.007384 |
| CNAG_13110 | 499.3985 | -0.43706 | 0.108981 | -4.01041 | 6.06E-05 | 0.000231 |
| CNAG_06309 | 266.4615 | 0.726636 | 0.220066 | 3.301894 | 0.00096 | 0.002841 |
| CNAG_06310 | 768.9585 | 0.579779 | 0.094345 | 6.145277 | 7.98E-10 | 5.90E-09 |
| CNAG_06311 | 1551.019 | 0.783696 | 0.083548 | 9.380198 | 6.58E-21 | 9.82E-20 |
| CNAG_06313 | 9990.69 | -0.63781 | 0.067048 | -9.51272 | 1.86E-21 | 2.84E-20 |
| CNAG_06314 | 10552.4 | -0.30207 | 0.062388 | -4.84188 | 1.29E-06 | 6.49E-06 |
| CNAG_06315 | 2746.821 | -0.32645 | 0.080551 | -4.05268 | 5.06E-05 | 0.000196 |
| CNAG_06316 | 3258.513 | 0.252416 | 0.0652 | 3.871404 | 0.000108 | 0.000393 |
| CNAG_06320 | 2514.648 | -0.63531 | 0.0908 | -6.99688 | 2.62E-12 | 2.41E-11 |
| CNAG_06322 | 583.5923 | -0.65075 | 0.105965 | -6.14119 | 8.19E-10 | 6.03E-09 |
| CNAG_13113 | 27.33144 | 0.977446 | 0.450385 | 2.170244 | 0.029988 | 0.060333 |
| CNAG_06323 | 446.9104 | 0.368275 | 0.130214 | 2.828241 | 0.00468 | 0.011787 |
| CNAG_06324 | 623.0548 | 0.501308 | 0.126768 | 3.954524 | 7.67E-05 | 0.000288 |
| CNAG_06325 | 2462.014 | -0.3428 | 0.077922 | -4.39921 | 1.09E-05 | 4.74E-05 |
| CNAG_06327 | 2259.374 | 0.513931 | 0.085976 | 5.977576 | 2.26E-09 | 1.60E-08 |
| CNAG_06329 | 334.934 | 1.806592 | 0.148268 | 12.18463 | 3.75E-34 | 8.67E-33 |
| CNAG_06330 | 452.9037 | 1.002877 | 0.121954 | 8.223411 | 1.98E-16 | 2.34E-15 |
| CNAG_06331 | 425.2422 | 1.409816 | 0.133084 | 10.59347 | 3.20E-26 | 5.72E-25 |
| CNAG_06336 | 1375.636 | 1.009596 | 0.098514 | 10.2483 | 1.20E-24 | 2.08E-23 |
| CNAG_13119 | 5.120899 | 3.352802 | 1.177326 | 2.84781 | 0.004402 | 0.011157 |
| CNAG_06339 | 979.4262 | 0.541918 | 0.109369 | 4.954938 | 7.24E-07 | 3.78E-06 |
| CNAG_06340 | 2790.547 | 0.157381 | 0.070267 | 2.239773 | 0.025106 | 0.0517 |
| CNAG_06343 | 743.7277 | -0.59004 | 0.095168 | -6.20003 | 5.65E-10 | 4.23E-09 |
| CNAG_06345 | 1046.091 | -0.29816 | 0.10011 | -2.97835 | 0.002898 | 0.00768 |
| CNAG_06346 | 2931.502 | -2.57746 | 0.110513 | -23.3228 | 2.61E-120 | 2.28E-118 |
| CNAG_13120 | 217.6473 | -0.4933 | 0.178212 | -2.76803 | 0.00564 | 0.013907 |
| CNAG_06347 | 6774.593 | -1.70235 | 0.104142 | -16.3464 | 4.62E-60 | 1.94E-58 |
| CNAG_06348 | 2493.18 | 0.894631 | 0.083543 | 10.70866 | 9.27E-27 | 1.69E-25 |
| CNAG_06352 | 1952.128 | -0.26694 | 0.088523 | -3.01546 | 0.002566 | 0.006885 |
| CNAG_06354 | 500.3047 | 0.348926 | 0.118581 | 2.942527 | 0.003255 | 0.008511 |
| CNAG_06355 | 270.5631 | -0.56647 | 0.156106 | -3.62872 | 0.000285 | 0.000948 |
| CNAG_06357 | 546.7886 | 0.278497 | 0.107703 | 2.585777 | 0.009716 | 0.022525 |
| CNAG_06358 | 412.6546 | 0.400756 | 0.127434 | 3.144817 | 0.001662 | 0.004655 |
| CNAG_06360 | 1040.911 | -0.29692 | 0.097068 | -3.05893 | 0.002221 | 0.006044 |
| CNAG_06361 | 4017.749 | 0.159689 | 0.068474 | 2.332099 | 0.019696 | 0.041841 |
| CNAG_06365 | 462.6091 | -0.37851 | 0.117389 | -3.22443 | 0.001262 | 0.00362 |
| CNAG_06368 | 598.3816 | -0.55191 | 0.111201 | -4.96323 | 6.93E-07 | 3.64E-06 |
| CNAG_06369 | 649.3642 | -0.30932 | 0.115299 | -2.68278 | 0.007301 | 0.017464 |
| CNAG_06370 | 2652.519 | -0.64194 | 0.074864 | -8.57482 | 9.92E-18 | 1.25E-16 |
| CNAG_06371 | 347.7795 | 0.47815 | 0.13178 | 3.62841 | 0.000285 | 0.000949 |
| CNAG_06372 | 306.4218 | 1.461071 | 0.159491 | 9.160825 | 5.15E-20 | 7.31E-19 |
| CNAG_06374 | 4844.457 | 2.723878 | 0.125019 | 21.78774 | 3.03E-105 | 2.39E-103 |
| CNAG_06375 | 205.9661 | -1.91934 | 0.168377 | -11.3991 | 4.22E-30 | 8.74E-29 |

| | | | | | | |
|---|---|---|---|---|---|---|
| CNAG_06377 | 36996.68 | -0.14868 | 0.060824 | -2.4445 | 0.014505 | 0.032039 |
| CNAG_06380 | 1283.374 | -0.29316 | 0.080891 | -3.62416 | 0.00029 | 0.000963 |
| CNAG_13122 | 114.8944 | -0.53216 | 0.262526 | -2.02707 | 0.042655 | 0.081827 |
| CNAG_06383 | 328.1615 | 0.419895 | 0.124114 | 3.38315 | 0.000717 | 0.00219 |
| CNAG_06384 | 1232.776 | 0.253804 | 0.07891 | 3.216391 | 0.001298 | 0.003719 |
| CNAG_06385 | 783.0877 | -0.55124 | 0.101281 | -5.44269 | 5.25E-08 | 3.18E-07 |
| CNAG_06386 | 1360.394 | -0.27166 | 0.081582 | -3.32996 | 0.000869 | 0.002598 |
| CNAG_06390 | 197.7906 | 0.495755 | 0.156032 | 3.177266 | 0.001487 | 0.004199 |
| CNAG_06392 | 942.3536 | -0.5187 | 0.086557 | -5.99263 | 2.06E-09 | 1.47E-08 |
| CNAG_06393 | 619.6121 | -0.39589 | 0.106384 | -3.72131 | 0.000198 | 0.000684 |
| CNAG_06396 | 386.4617 | 1.130254 | 0.143391 | 7.882301 | 3.21E-15 | 3.58E-14 |
| CNAG_07936 | 1947.441 | 0.942426 | 0.074811 | 12.59746 | 2.18E-36 | 5.40E-35 |
| CNAG_06399 | 536.6224 | -0.4277 | 0.115282 | -3.71006 | 0.000207 | 0.000711 |
| CNAG_06400 | 83351.39 | 0.254143 | 0.062007 | 4.098622 | 4.16E-05 | 0.000164 |
| CNAG_13126 | 55.22741 | -1.19703 | 0.321486 | -3.72344 | 0.000197 | 0.000679 |
| CNAG_06401 | 693.5521 | -0.30016 | 0.098346 | -3.05207 | 0.002273 | 0.006176 |
| CNAG_06402 | 2104.44 | 0.154411 | 0.076362 | 2.022097 | 0.043166 | 0.08269 |
| CNAG_06404 | 13693.85 | 2.96606 | 0.088749 | 33.42069 | 6.86E-245 | 2.15E-242 |
| CNAG_06405 | 1358.961 | 0.653041 | 0.078045 | 8.367489 | 5.89E-17 | 7.20E-16 |
| CNAG_06406 | 869.3644 | -0.18996 | 0.089779 | -2.11583 | 0.034359 | 0.067735 |
| CNAG_06407 | 3803.798 | 0.530166 | 0.093615 | 5.663251 | 1.49E-08 | 9.56E-08 |
| CNAG_06408 | 286.9907 | 0.457636 | 0.151614 | 3.018421 | 0.002541 | 0.006827 |
| CNAG_06409 | 408.345 | -0.28766 | 0.140775 | -2.0434 | 0.041012 | 0.079068 |
| CNAG_06410 | 964.3909 | -0.49483 | 0.094926 | -5.21282 | 1.86E-07 | 1.05E-06 |
| CNAG_06412 | 1460.213 | 0.295385 | 0.080279 | 3.679491 | 0.000234 | 0.000792 |
| CNAG_06413 | 139.3752 | -0.97412 | 0.198095 | -4.91744 | 8.77E-07 | 4.53E-06 |
| CNAG_06414 | 675.0753 | 0.529846 | 0.100466 | 5.273907 | 1.34E-07 | 7.69E-07 |
| CNAG_06415 | 1235.296 | 0.617758 | 0.100579 | 6.142019 | 8.15E-10 | 6.00E-09 |
| CNAG_06416 | 1637.345 | -0.33266 | 0.086926 | -3.82701 | 0.00013 | 0.000464 |
| CNAG_06417 | 413.5796 | -0.38518 | 0.131967 | -2.91876 | 0.003514 | 0.009131 |
| CNAG_06419 | 365.4729 | -0.462 | 0.125225 | -3.68939 | 0.000225 | 0.000764 |
| CNAG_06420 | 3447.366 | -0.38047 | 0.06931 | -5.48944 | 4.03E-08 | 2.47E-07 |
| CNAG_06421 | 3500.238 | -0.36298 | 0.096297 | -3.76936 | 0.000164 | 0.000575 |
| CNAG_06423 | 524.0156 | -0.50053 | 0.109359 | -4.57692 | 4.72E-06 | 2.20E-05 |
| CNAG_06424 | 997.0277 | -0.21405 | 0.088959 | -2.40613 | 0.016122 | 0.035113 |
| CNAG_07937 | 1956.56 | -0.24564 | 0.070498 | -3.48437 | 0.000493 | 0.001556 |
| CNAG_07938 | 1180.727 | 0.508002 | 0.090545 | 5.610514 | 2.02E-08 | 1.28E-07 |
| CNAG_06431 | 489.4846 | -0.58259 | 0.110332 | -5.28031 | 1.29E-07 | 7.45E-07 |
| CNAG_06432 | 2323.579 | -0.6242 | 0.1146 | -5.44678 | 5.13E-08 | 3.11E-07 |
| CNAG_07939 | 6074.986 | -1.02914 | 0.323527 | -3.18099 | 0.001468 | 0.004151 |
| CNAG_13129 | 173.1888 | -1.20276 | 0.208458 | -5.76977 | 7.94E-09 | 5.28E-08 |
| CNAG_06437 | 1621.943 | -0.43715 | 0.101297 | -4.31549 | 1.59E-05 | 6.72E-05 |
| CNAG_13131 | 37.91817 | -2.07831 | 0.389572 | -5.33486 | 9.56E-08 | 5.63E-07 |
| CNAG_06438 | 9.217949 | -2.69543 | 0.819005 | -3.2911 | 0.000998 | 0.002936 |
| CNAG_06439 | 1943.37 | -0.30506 | 0.074442 | -4.09799 | 4.17E-05 | 0.000164 |
| CNAG_06440 | 929.5038 | -0.63638 | 0.085002 | -7.48665 | 7.07E-14 | 7.34E-13 |
| CNAG_07940 | 212.011 | -0.6067 | 0.158348 | -3.83144 | 0.000127 | 0.000456 |

| | | | | | | |
|---|---|---|---|---|---|---|
| CNAG_06443 | 30492.8 | 0.604726 | 0.071925 | 8.4077 | 4.18E-17 | 5.14E-16 |
| CNAG_06444 | 1027.411 | -0.2933 | 0.087437 | -3.35439 | 0.000795 | 0.0024 |
| CNAG_06446 | 2453.337 | 0.239762 | 0.065518 | 3.65949 | 0.000253 | 0.000851 |
| CNAG_06450 | 2339.413 | -0.19041 | 0.065255 | -2.91786 | 0.003524 | 0.009144 |
| CNAG_06452 | 1119.728 | -0.21728 | 0.093791 | -2.31666 | 0.020522 | 0.043259 |
| CNAG_06454 | 1592.05 | 0.252212 | 0.075873 | 3.324151 | 0.000887 | 0.002642 |
| CNAG_07941 | 4426.618 | 0.668801 | 0.063591 | 10.51714 | 7.20E-26 | 1.28E-24 |
| CNAG_06459 | 850.5833 | -0.2595 | 0.106357 | -2.43988 | 0.014692 | 0.032389 |
| CNAG_06460 | 2673.243 | 0.798789 | 0.091071 | 8.771092 | 1.77E-18 | 2.30E-17 |
| CNAG_06464 | 984.8213 | -0.23495 | 0.09345 | -2.51414 | 0.011932 | 0.027053 |
| CNAG_06465 | 1808.912 | -0.43012 | 0.089119 | -4.82637 | 1.39E-06 | 6.97E-06 |
| CNAG_06466 | 439.3082 | -0.56078 | 0.118296 | -4.74048 | 2.13E-06 | 1.05E-05 |
| CNAG_06468 | 5194.15 | 0.138931 | 0.063491 | 2.188217 | 0.028654 | 0.058036 |
| CNAG_06469 | 3147.177 | -0.25146 | 0.070108 | -3.58677 | 0.000335 | 0.001099 |
| CNAG_06472 | 6559.502 | 0.271922 | 0.066145 | 4.111002 | 3.94E-05 | 0.000155 |
| CNAG_06474 | 5957.511 | -0.28969 | 0.074896 | -3.86792 | 0.00011 | 0.000399 |
| CNAG_13134 | 193.5674 | -0.71452 | 0.166967 | -4.2794 | 1.87E-05 | 7.81E-05 |
| CNAG_06482 | 514.8719 | -0.22775 | 0.107191 | -2.12475 | 0.033607 | 0.066543 |
| CNAG_06483 | 1310.504 | 0.760094 | 0.081233 | 9.356977 | 8.21E-21 | 1.22E-19 |
| CNAG_06484 | 99.51664 | 0.482815 | 0.21742 | 2.220657 | 0.026374 | 0.054038 |
| CNAG_06485 | 299.9853 | 1.067345 | 0.148075 | 7.20815 | 5.67E-13 | 5.53E-12 |
| CNAG_06487 | 1604.724 | 0.213406 | 0.074043 | 2.88218 | 0.003949 | 0.010113 |
| CNAG_07943 | 639.3607 | 2.191708 | 0.140217 | 15.63079 | 4.49E-55 | 1.70E-53 |
| CNAG_06489 | 5676.122 | -0.23618 | 0.102988 | -2.29329 | 0.021831 | 0.045756 |
| CNAG_06493 | 1000.479 | -0.91741 | 0.401657 | -2.28406 | 0.022368 | 0.046725 |
| CNAG_06497 | 651.1786 | -0.35554 | 0.125625 | -2.83014 | 0.004653 | 0.01173 |
| CNAG_06499 | 1032.991 | 0.194832 | 0.090542 | 2.151844 | 0.03141 | 0.062866 |
| CNAG_06500 | 809.2307 | 0.396824 | 0.11305 | 3.510161 | 0.000448 | 0.001427 |
| CNAG_06501 | 1678.38 | 0.33362 | 0.09109 | 3.662526 | 0.00025 | 0.000843 |
| CNAG_06503 | 144.7733 | 2.906641 | 0.217985 | 13.33416 | 1.46E-40 | 4.06E-39 |
| CNAG_06505 | 938.9552 | -0.36145 | 0.085326 | -4.23609 | 2.27E-05 | 9.30E-05 |
| CNAG_06506 | 591.7325 | -0.27989 | 0.105249 | -2.65933 | 0.00783 | 0.018592 |
| CNAG_06508 | 5537.129 | 0.310233 | 0.078795 | 3.937197 | 8.24E-05 | 0.000307 |
| CNAG_06510 | 852.5143 | -0.28957 | 0.098872 | -2.92874 | 0.003403 | 0.008872 |
| CNAG_06512 | 1636.087 | 2.070139 | 0.087247 | 23.72733 | 1.88E-124 | 1.70E-122 |
| CNAG_06515 | 281.6783 | 0.285864 | 0.133761 | 2.137128 | 0.032588 | 0.064793 |
| CNAG_06516 | 389.0127 | 1.622666 | 0.141532 | 11.46499 | 1.98E-30 | 4.11E-29 |
| CNAG_06517 | 3773.514 | -0.3888 | 0.088491 | -4.39366 | 1.11E-05 | 4.85E-05 |
| CNAG_06518 | 14.21652 | -1.63396 | 0.604301 | -2.70389 | 0.006853 | 0.01648 |
| CNAG_06519 | 447.9681 | -0.34629 | 0.119468 | -2.89861 | 0.003748 | 0.009665 |
| CNAG_06520 | 489.9676 | -0.49224 | 0.10786 | -4.56373 | 5.03E-06 | 2.33E-05 |
| CNAG_05333 | 7.392897 | 1.796938 | 0.823977 | 2.180811 | 0.029197 | 0.058946 |
| CNAG_05334 | 39.09676 | -0.86111 | 0.368264 | -2.33829 | 0.019372 | 0.041209 |
| CNAG_05336 | 1149.731 | 0.426093 | 0.115467 | 3.69019 | 0.000224 | 0.000762 |
| CNAG_05337 | 1128.001 | 0.365005 | 0.087651 | 4.164291 | 3.12E-05 | 0.000125 |
| CNAG_05339 | 1526.021 | -0.22635 | 0.078217 | -2.89382 | 0.003806 | 0.009795 |
| CNAG_05340 | 434.9493 | 0.726496 | 0.171315 | 4.240709 | 2.23E-05 | 9.14E-05 |

| | | | | | | |
|---|---|---|---|---|---|---|
| CNAG_05341 | 418.5035 | 1.758742 | 0.161275 | 10.90526 | 1.09E-27 | 2.08E-26 |
| CNAG_05343 | 1408.173 | -0.27566 | 0.108032 | -2.55163 | 0.010722 | 0.024618 |
| CNAG_05344 | 2795.028 | 0.325526 | 0.06552 | 4.968315 | 6.75E-07 | 3.55E-06 |
| CNAG_13145 | 31.44399 | 0.892927 | 0.407816 | 2.189532 | 0.028558 | 0.057886 |
| CNAG_05346 | 138.4807 | -0.44265 | 0.194257 | -2.27868 | 0.022686 | 0.047316 |
| CNAG_07032 | 3311.287 | -0.56318 | 0.070164 | -8.02669 | 1.00E-15 | 1.15E-14 |
| CNAG_05348 | 5097.948 | -0.3839 | 0.062235 | -6.16856 | 6.89E-10 | 5.14E-09 |
| CNAG_05351 | 9219.481 | 0.224393 | 0.074366 | 3.017398 | 0.00255 | 0.006846 |
| CNAG_05352 | 1618.148 | 0.277652 | 0.073304 | 3.787657 | 0.000152 | 0.000537 |
| CNAG_05354 | 1083.89 | -0.28498 | 0.090557 | -3.14702 | 0.001649 | 0.004621 |
| CNAG_05356 | 178.5587 | 2.692567 | 0.20133 | 13.37392 | 8.59E-41 | 2.40E-39 |
| CNAG_05357 | 47.59441 | 2.860715 | 0.400359 | 7.145369 | 8.98E-13 | 8.60E-12 |
| CNAG_13147 | 410.1513 | 2.654722 | 0.168263 | 15.77726 | 4.46E-56 | 1.71E-54 |
| CNAG_13148 | 58.25823 | 5.722221 | 0.620385 | 9.223654 | 2.87E-20 | 4.13E-19 |
| CNAG_07870 | 11.23604 | 7.015354 | 1.31581 | 5.331586 | 9.74E-08 | 5.72E-07 |
| CNAG_05358 | 208.6 | 3.555495 | 0.213659 | 16.64096 | 3.52E-62 | 1.56E-60 |
| CNAG_05359 | 1097.254 | -0.18067 | 0.090698 | -1.99202 | 0.046369 | 0.087913 |
| CNAG_08021 | 133.945 | 0.755641 | 0.186702 | 4.047314 | 5.18E-05 | 0.0002 |
| CNAG_08022 | 2447.666 | 0.258058 | 0.071739 | 3.597174 | 0.000322 | 0.00106 |
| CNAG_05364 | 385.3338 | 0.250311 | 0.126242 | 1.982783 | 0.047392 | 0.089602 |
| CNAG_05365 | 2773.571 | 0.578135 | 0.064035 | 9.028372 | 1.74E-19 | 2.41E-18 |
| CNAG_07873 | 1565.861 | -0.21141 | 0.078837 | -2.68159 | 0.007327 | 0.017517 |
| CNAG_05371 | 2507.333 | 0.249586 | 0.07444 | 3.35286 | 0.0008 | 0.002412 |
| CNAG_05373 | 648.8763 | -0.24618 | 0.10971 | -2.24393 | 0.024837 | 0.051237 |
| CNAG_05374 | 1535.251 | 0.463392 | 0.076708 | 6.040989 | 1.53E-09 | 1.10E-08 |
| CNAG_05375 | 24284.28 | -0.12965 | 0.059117 | -2.19309 | 0.028301 | 0.057422 |
| CNAG_05377 | 139.8929 | 2.768565 | 0.233364 | 11.86372 | 1.83E-32 | 3.97E-31 |
| CNAG_05378 | 462.8577 | 0.390106 | 0.123238 | 3.165478 | 0.001548 | 0.004362 |
| CNAG_05379 | 2638.46 | 1.277535 | 0.070527 | 18.11405 | 2.47E-73 | 1.27E-71 |
| CNAG_05380 | 1371.999 | 0.418383 | 0.08129 | 5.146782 | 2.65E-07 | 1.47E-06 |
| CNAG_05381 | 294.0478 | 1.960582 | 0.483928 | 4.051393 | 5.09E-05 | 0.000197 |
| CNAG_13155 | 87.76734 | 1.73097 | 0.245987 | 7.036842 | 1.97E-12 | 1.84E-11 |
| CNAG_13156 | 158.357 | 1.798877 | 0.182141 | 9.876298 | 5.27E-23 | 8.57E-22 |
| CNAG_05383 | 725.2154 | 0.462342 | 0.141284 | 3.272432 | 0.001066 | 0.003113 |
| CNAG_05384 | 2069.077 | -0.26614 | 0.071099 | -3.74326 | 0.000182 | 0.000631 |
| CNAG_05387 | 8274.853 | -1.49092 | 0.118865 | -12.5431 | 4.34E-36 | 1.06E-34 |
| CNAG_05391 | 1752.761 | -0.28231 | 0.092638 | -3.04741 | 0.002308 | 0.006262 |
| CNAG_05392 | 180.3928 | 0.364724 | 0.185614 | 1.964957 | 0.049419 | 0.092809 |
| CNAG_05393 | 1007.089 | -0.17374 | 0.083384 | -2.08366 | 0.037192 | 0.072666 |
| CNAG_05394 | 657.3767 | 0.266104 | 0.099691 | 2.669276 | 0.007602 | 0.018103 |
| CNAG_07875 | 741.0708 | 0.470929 | 0.126647 | 3.718441 | 0.0002 | 0.00069 |
| CNAG_05396 | 320.2088 | -0.63738 | 0.147515 | -4.32077 | 1.55E-05 | 6.58E-05 |
| CNAG_05397 | 203.441 | 1.256531 | 0.161326 | 7.788765 | 6.77E-15 | 7.39E-14 |
| CNAG_05398 | 3823.886 | 0.344647 | 0.068756 | 5.0126 | 5.37E-07 | 2.86E-06 |
| CNAG_05401 | 1654.963 | 0.385792 | 0.074345 | 5.189206 | 2.11E-07 | 1.18E-06 |
| CNAG_05402 | 2982.656 | -0.24615 | 0.064073 | -3.84169 | 0.000122 | 0.00044 |
| CNAG_05403 | 483.17 | 0.337755 | 0.107583 | 3.139484 | 0.001692 | 0.004735 |

| | | | | | | |
|---|---|---|---|---|---|---|
| CNAG_05404 | 1437.738 | 0.469794 | 0.076745 | 6.121477 | 9.27E-10 | 6.79E-09 |
| CNAG_05410 | 568.6152 | 0.275594 | 0.11574 | 2.381151 | 0.017259 | 0.037268 |
| CNAG_05411 | 1531.784 | 0.811785 | 0.077508 | 10.47363 | 1.14E-25 | 2.02E-24 |
| CNAG_05412 | 49.80278 | 2.624956 | 0.35428 | 7.409273 | 1.27E-13 | 1.29E-12 |
| CNAG_05413 | 697.7319 | -0.40425 | 0.09432 | -4.28594 | 1.82E-05 | 7.61E-05 |
| CNAG_05415 | 1862.954 | -0.95377 | 0.101332 | -9.41234 | 4.85E-21 | 7.29E-20 |
| CNAG_05416 | 3619.854 | -0.21599 | 0.077346 | -2.79252 | 0.00523 | 0.013019 |
| CNAG_05419 | 1756.011 | -0.24817 | 0.083477 | -2.97288 | 0.00295 | 0.007801 |
| CNAG_05420 | 578.219 | -0.36838 | 0.119439 | -3.08424 | 0.002041 | 0.005602 |
| CNAG_13168 | 218.5939 | 0.353241 | 0.15335 | 2.303492 | 0.021251 | 0.04461 |
| CNAG_05423 | 751.8424 | 1.252165 | 0.103958 | 12.04488 | 2.06E-33 | 4.63E-32 |
| CNAG_05424 | 15238.29 | 0.918624 | 0.068209 | 13.46773 | 2.42E-41 | 6.81E-40 |
| CNAG_05425 | 13525.19 | -0.24521 | 0.054121 | -4.53071 | 5.88E-06 | 2.71E-05 |
| CNAG_05426 | 157.1185 | 0.444575 | 0.20955 | 2.121572 | 0.033874 | 0.066957 |
| CNAG_08024 | 359.6774 | -0.44048 | 0.122545 | -3.59443 | 0.000325 | 0.00107 |
| CNAG_05428 | 1822.333 | -0.27952 | 0.081972 | -3.40995 | 0.00065 | 0.002 |
| CNAG_05431 | 980.7471 | -0.45015 | 0.10059 | -4.4751 | 7.64E-06 | 3.45E-05 |
| CNAG_05434 | 4727.412 | 0.883453 | 0.072218 | 12.23321 | 2.07E-34 | 4.78E-33 |
| CNAG_05437 | 18290.44 | -0.29503 | 0.081996 | -3.5981 | 0.000321 | 0.001057 |
| CNAG_05438 | 747.7716 | -0.32985 | 0.097245 | -3.39198 | 0.000694 | 0.002124 |
| CNAG_05440 | 979.93 | -0.41845 | 0.09656 | -4.33362 | 1.47E-05 | 6.25E-05 |
| CNAG_05441 | 446.8728 | 0.441889 | 0.141676 | 3.119018 | 0.001815 | 0.005047 |
| CNAG_13178 | 80.53969 | -0.70486 | 0.246033 | -2.8649 | 0.004171 | 0.010642 |
| CNAG_05444 | 1658.575 | 1.572879 | 0.089059 | 17.66111 | 8.36E-70 | 4.12E-68 |
| CNAG_05448 | 18.30143 | 1.514748 | 0.535586 | 2.828204 | 0.004681 | 0.011787 |
| CNAG_07876 | 162.754 | 0.626942 | 0.168371 | 3.723582 | 0.000196 | 0.000678 |
| CNAG_05449 | 9037.467 | 0.658089 | 0.104626 | 6.289936 | 3.18E-10 | 2.45E-09 |
| CNAG_05450 | 350.4994 | 0.322299 | 0.135856 | 2.372363 | 0.017675 | 0.038066 |
| CNAG_05455 | 9900.422 | 0.278006 | 0.069362 | 4.008047 | 6.12E-05 | 0.000233 |
| CNAG_05457 | 3881.833 | 0.152284 | 0.062593 | 2.432935 | 0.014977 | 0.032945 |
| CNAG_05458 | 267.743 | 1.665772 | 0.228056 | 7.304223 | 2.79E-13 | 2.78E-12 |
| CNAG_05459 | 3604.832 | -0.29115 | 0.068545 | -4.24762 | 2.16E-05 | 8.89E-05 |
| CNAG_05460 | 465.5538 | 0.317645 | 0.119456 | 2.65909 | 0.007835 | 0.018599 |
| CNAG_05461 | 151.1317 | 1.501946 | 0.223199 | 6.729187 | 1.71E-11 | 1.47E-10 |
| CNAG_05462 | 9386.991 | -0.19069 | 0.064587 | -2.9525 | 0.003152 | 0.008278 |
| CNAG_05463 | 1729.119 | -0.37078 | 0.083717 | -4.42896 | 9.47E-06 | 4.18E-05 |
| CNAG_05464 | 688.8991 | -0.46182 | 0.108752 | -4.24652 | 2.17E-05 | 8.93E-05 |
| CNAG_05468 | 1084.763 | -0.29069 | 0.081706 | -3.55773 | 0.000374 | 0.001215 |
| CNAG_05470 | 837.3481 | 0.186505 | 0.087753 | 2.125342 | 0.033558 | 0.066464 |
| CNAG_07878 | 4465.691 | 0.150629 | 0.06917 | 2.177654 | 0.029432 | 0.05939 |
| CNAG_05472 | 61.47351 | 0.780361 | 0.30082 | 2.594115 | 0.009483 | 0.022056 |
| CNAG_07879 | 268.513 | -0.50195 | 0.140641 | -3.56899 | 0.000358 | 0.001169 |
| CNAG_13182 | 10.19985 | 2.465593 | 0.736377 | 3.348274 | 0.000813 | 0.002446 |
| CNAG_05479 | 3602.749 | 2.974927 | 0.074789 | 39.77749 | 0 | 0 |
| CNAG_05480 | 7815.294 | -0.14832 | 0.069496 | -2.13417 | 0.032829 | 0.065241 |
| CNAG_13183 | 18.91325 | -1.13916 | 0.536634 | -2.12279 | 0.033771 | 0.066816 |
| CNAG_07881 | 28.48793 | -1.1824 | 0.455012 | -2.5986 | 0.00936 | 0.021844 |

| | | | | | | |
|---|---|---|---|---|---|---|
| CNAG_05482 | 282.2259 | -0.52215 | 0.134152 | -3.89224 | 9.93E-05 | 0.000364 |
| CNAG_05483 | 1104.585 | -0.36196 | 0.08452 | -4.2826 | 1.85E-05 | 7.71E-05 |
| CNAG_05484 | 843.5024 | -0.26953 | 0.092521 | -2.91318 | 0.003578 | 0.009261 |
| CNAG_08026 | 191.7243 | -1.04351 | 0.174907 | -5.96605 | 2.43E-09 | 1.72E-08 |
| CNAG_05496 | 6015.336 | -0.2098 | 0.070491 | -2.97621 | 0.002918 | 0.007729 |
| CNAG_05497 | 22059.24 | 0.836101 | 0.053742 | 15.55772 | 1.41E-54 | 5.21E-53 |
| CNAG_05499 | 3806.739 | -0.53986 | 0.06319 | -8.54345 | 1.30E-17 | 1.63E-16 |
| CNAG_05501 | 257.265 | 1.404728 | 0.179575 | 7.822495 | 5.18E-15 | 5.71E-14 |
| CNAG_05502 | 1835.562 | 0.849001 | 0.070796 | 11.9922 | 3.90E-33 | 8.67E-32 |
| CNAG_05503 | 427.9764 | -0.56121 | 0.12423 | -4.51749 | 6.26E-06 | 2.87E-05 |
| CNAG_05505 | 531.6081 | -0.25494 | 0.118831 | -2.1454 | 0.031921 | 0.063698 |
| CNAG_07883 | 1366.226 | 0.239647 | 0.083906 | 2.856143 | 0.004288 | 0.010913 |
| CNAG_07884 | 5997.638 | 0.151863 | 0.074332 | 2.04304 | 0.041049 | 0.0791 |
| CNAG_05507 | 2239.2 | -0.2412 | 0.092981 | -2.59411 | 0.009484 | 0.022056 |
| CNAG_05508 | 450.7679 | -0.26254 | 0.115065 | -2.28166 | 0.022509 | 0.046972 |
| CNAG_05509 | 3139.787 | 0.670108 | 0.084481 | 7.932049 | 2.16E-15 | 2.43E-14 |
| CNAG_05511 | 4097.337 | 0.261935 | 0.071623 | 3.657127 | 0.000255 | 0.000857 |
| CNAG_05512 | 1635.127 | -0.2064 | 0.079579 | -2.59362 | 0.009497 | 0.02208 |
| CNAG_05516 | 1050.137 | -0.33128 | 0.099137 | -3.34164 | 0.000833 | 0.002501 |
| CNAG_05517 | 1675.14 | -0.30387 | 0.085639 | -3.54823 | 0.000388 | 0.001254 |
| CNAG_05518 | 1176.659 | -0.20705 | 0.099841 | -2.07378 | 0.0381 | 0.07428 |
| CNAG_05519 | 600.5453 | 0.31216 | 0.100836 | 3.095719 | 0.001963 | 0.005426 |
| CNAG_05520 | 605.9062 | 0.310193 | 0.10252 | 3.025693 | 0.002481 | 0.006679 |
| CNAG_05521 | 2566.391 | 0.511451 | 0.094658 | 5.403152 | 6.55E-08 | 3.94E-07 |
| CNAG_13188 | 140.6003 | 0.779488 | 0.190354 | 4.094929 | 4.22E-05 | 0.000166 |
| CNAG_05523 | 210.6718 | 1.535548 | 0.176146 | 8.717468 | 2.84E-18 | 3.68E-17 |
| CNAG_13189 | 48.45548 | -0.98499 | 0.327617 | -3.00651 | 0.002643 | 0.007068 |
| CNAG_05524 | 1113.449 | -0.23811 | 0.085922 | -2.77123 | 0.005585 | 0.013792 |
| CNAG_05525 | 47762.56 | -0.20035 | 0.069676 | -2.87539 | 0.004035 | 0.010317 |
| CNAG_05526 | 417.127 | -0.27735 | 0.129657 | -2.13912 | 0.032426 | 0.064551 |
| CNAG_05527 | 1822.543 | 0.346949 | 0.077014 | 4.505014 | 6.64E-06 | 3.03E-05 |
| CNAG_05529 | 930.6531 | -0.19277 | 0.087641 | -2.19953 | 0.02784 | 0.056599 |
| CNAG_05537 | 1254.265 | 0.203882 | 0.096141 | 2.120657 | 0.033951 | 0.067069 |
| CNAG_05539 | 1089.8 | 0.376348 | 0.087035 | 4.324081 | 1.53E-05 | 6.49E-05 |
| CNAG_05540 | 9583.24 | 0.478747 | 0.056126 | 8.529871 | 1.47E-17 | 1.83E-16 |
| CNAG_05543 | 153.4894 | 0.931466 | 0.187401 | 4.970454 | 6.68E-07 | 3.51E-06 |
| CNAG_05546 | 611.6221 | -0.30986 | 0.105531 | -2.93618 | 0.003323 | 0.008667 |
| CNAG_05547 | 249.7992 | 0.426939 | 0.151693 | 2.814498 | 0.004885 | 0.012237 |
| CNAG_05553 | 491.0408 | -0.29993 | 0.115251 | -2.60245 | 0.009256 | 0.021637 |
| CNAG_05554 | 3633.556 | 0.669305 | 0.073029 | 9.164905 | 4.96E-20 | 7.05E-19 |
| CNAG_05558 | 1017.541 | 0.360166 | 0.08532 | 4.221344 | 2.43E-05 | 9.87E-05 |
| CNAG_05563 | 638.2739 | -0.24823 | 0.097843 | -2.53701 | 0.01118 | 0.025584 |
| CNAG_05564 | 618.6709 | -0.28962 | 0.110526 | -2.62034 | 0.008784 | 0.020617 |
| CNAG_05566 | 437.835 | -0.2634 | 0.115271 | -2.28503 | 0.022311 | 0.046618 |
| CNAG_05567 | 723.304 | -0.4347 | 0.092535 | -4.69773 | 2.63E-06 | 1.27E-05 |
| CNAG_05568 | 936.9637 | -0.30043 | 0.085796 | -3.50165 | 0.000462 | 0.001469 |
| CNAG_05570 | 794.6046 | -0.21971 | 0.106113 | -2.07055 | 0.038401 | 0.074777 |

| | | | | | | |
|---|---|---|---|---|---|---|
| CNAG_05571 | 4359.338 | 0.234721 | 0.068251 | 3.439077 | 0.000584 | 0.001818 |
| CNAG_05573 | 1041.657 | 0.393055 | 0.119829 | 3.280142 | 0.001038 | 0.003041 |
| CNAG_05575 | 925.0902 | -0.63836 | 0.088298 | -7.22968 | 4.84E-13 | 4.78E-12 |
| CNAG_05577 | 751.1243 | -0.35032 | 0.089823 | -3.90008 | 9.62E-05 | 0.000354 |
| CNAG_05578 | 2321.189 | 0.608725 | 0.091546 | 6.64938 | 2.94E-11 | 2.49E-10 |
| CNAG_05580 | 38.17557 | 0.842001 | 0.424086 | 1.985449 | 0.047095 | 0.089123 |
| CNAG_05581 | 5758.291 | 0.688787 | 0.071631 | 9.615834 | 6.86E-22 | 1.06E-20 |
| CNAG_05582 | 102.7275 | 0.756322 | 0.233934 | 3.233061 | 0.001225 | 0.003526 |
| CNAG_05584 | 1298.366 | -0.18683 | 0.087907 | -2.12533 | 0.033559 | 0.066464 |
| CNAG_07887 | 1874.824 | 0.592185 | 0.098782 | 5.994865 | 2.04E-09 | 1.45E-08 |
| CNAG_05590 | 1923.235 | 0.256213 | 0.102205 | 2.506856 | 0.012181 | 0.027525 |
| CNAG_05591 | 531.1361 | -0.4286 | 0.106878 | -4.0102 | 6.07E-05 | 0.000231 |
| CNAG_05592 | 6611.226 | 0.542867 | 0.066572 | 8.154621 | 3.50E-16 | 4.11E-15 |
| CNAG_08028 | 409.2906 | -1.01034 | 0.142145 | -7.10779 | 1.18E-12 | 1.12E-11 |
| CNAG_05595 | 2316.093 | 1.548779 | 0.104984 | 14.75251 | 2.96E-49 | 9.83E-48 |
| CNAG_05596 | 3681.064 | -0.3663 | 0.082031 | -4.46535 | 7.99E-06 | 3.59E-05 |
| CNAG_05598 | 1502.297 | 0.639549 | 0.085936 | 7.442142 | 9.91E-14 | 1.02E-12 |
| CNAG_05602 | 1359.724 | 0.821678 | 0.088264 | 9.309284 | 1.29E-20 | 1.88E-19 |
| CNAG_05605 | 763.5273 | 0.356694 | 0.095247 | 3.744945 | 0.00018 | 0.000627 |
| CNAG_05606 | 844.3683 | -0.21719 | 0.093905 | -2.31285 | 0.020731 | 0.043631 |
| CNAG_05607 | 1702.556 | -1.00814 | 0.136149 | -7.40463 | 1.32E-13 | 1.33E-12 |
| CNAG_07888 | 27487.9 | 0.43159 | 0.063141 | 6.83532 | 8.18E-12 | 7.29E-11 |
| CNAG_05609 | 1873.824 | 0.17016 | 0.072405 | 2.350118 | 0.018767 | 0.040121 |
| CNAG_07889 | 2065.437 | 0.172054 | 0.078594 | 2.189157 | 0.028585 | 0.057926 |
| CNAG_05612 | 858.0849 | -0.39181 | 0.088778 | -4.41332 | 1.02E-05 | 4.47E-05 |
| CNAG_05613 | 2079.617 | 0.194157 | 0.071366 | 2.720567 | 0.006517 | 0.015788 |
| CNAG_05615 | 6092.74 | -0.15634 | 0.076498 | -2.04372 | 0.040981 | 0.079026 |
| CNAG_05616 | 2088.405 | 0.37811 | 0.086701 | 4.361063 | 1.29E-05 | 5.56E-05 |
| CNAG_05617 | 1777.871 | 0.528026 | 0.084429 | 6.254089 | 4.00E-10 | 3.04E-09 |
| CNAG_05618 | 2504.828 | 0.184102 | 0.067016 | 2.747162 | 0.006011 | 0.014682 |
| CNAG_05621 | 1299.285 | -0.27786 | 0.0867 | -3.20488 | 0.001351 | 0.003856 |
| CNAG_05623 | 5529.689 | -0.19553 | 0.072183 | -2.70878 | 0.006753 | 0.016283 |
| CNAG_05626 | 1033.431 | 0.261348 | 0.084245 | 3.102249 | 0.001921 | 0.005321 |
| CNAG_05627 | 1339.486 | -0.20176 | 0.101628 | -1.98522 | 0.047119 | 0.089147 |
| CNAG_05628 | 870.6824 | 0.344148 | 0.091364 | 3.766795 | 0.000165 | 0.00058 |
| CNAG_05629 | 2227.985 | 0.227797 | 0.069822 | 3.262526 | 0.001104 | 0.003212 |
| CNAG_05631 | 15954.73 | 0.862894 | 0.055393 | 15.57765 | 1.03E-54 | 3.83E-53 |
| CNAG_13197 | 208.1841 | 0.614373 | 0.168216 | 3.652284 | 0.00026 | 0.000871 |
| CNAG_05633 | 4860.968 | 0.559851 | 0.096793 | 5.784008 | 7.29E-09 | 4.89E-08 |
| CNAG_05634 | 1010.847 | -0.2994 | 0.08935 | -3.35085 | 0.000806 | 0.002428 |
| CNAG_05637 | 1077.492 | 0.255953 | 0.094103 | 2.719926 | 0.00653 | 0.01581 |
| CNAG_05638 | 3691.335 | 0.177853 | 0.06454 | 2.755698 | 0.005857 | 0.014351 |
| CNAG_05639 | 977.4448 | -0.61942 | 0.095704 | -6.47222 | 9.66E-11 | 7.83E-10 |
| CNAG_05640 | 1256.495 | -0.41975 | 0.086733 | -4.8396 | 1.30E-06 | 6.56E-06 |
| CNAG_05642 | 1394.509 | -0.32203 | 0.09059 | -3.55476 | 0.000378 | 0.001227 |
| CNAG_05643 | 706.3935 | -0.47755 | 0.144713 | -3.29996 | 0.000967 | 0.002857 |
| CNAG_05644 | 395.998 | -0.44857 | 0.122862 | -3.65106 | 0.000261 | 0.000875 |

| | | | | | | |
|---|---|---|---|---|---|---|
| CNAG_05645 | 1322.215 | -0.20563 | 0.08607 | -2.38912 | 0.016889 | 0.036587 |
| CNAG_05650 | 8365.346 | -0.17871 | 0.06013 | -2.97205 | 0.002958 | 0.007817 |
| CNAG_05652 | 1729.767 | 0.692662 | 0.102283 | 6.771996 | 1.27E-11 | 1.11E-10 |
| CNAG_05653 | 1247.006 | 0.614528 | 0.101662 | 6.044824 | 1.50E-09 | 1.07E-08 |
| CNAG_05654 | 1058.943 | 2.372662 | 0.275033 | 8.626834 | 6.31E-18 | 8.02E-17 |
| CNAG_05656 | 170.8465 | 0.623933 | 0.174342 | 3.57878 | 0.000345 | 0.00113 |
| CNAG_05657 | 1000.2 | -1.13033 | 0.110885 | -10.1937 | 2.12E-24 | 3.60E-23 |
| CNAG_05658 | 65.61507 | -1.8822 | 0.302228 | -6.22777 | 4.73E-10 | 3.58E-09 |
| CNAG_05661 | 993.7504 | 0.190643 | 0.081341 | 2.343751 | 0.019091 | 0.040695 |
| CNAG_05662 | 1193.332 | -2.1719 | 0.478425 | -4.53968 | 5.63E-06 | 2.60E-05 |
| CNAG_05664 | 3653.358 | -0.29066 | 0.061137 | -4.75435 | 1.99E-06 | 9.81E-06 |
| CNAG_13206 | 256.8247 | 0.396081 | 0.181354 | 2.184022 | 0.028961 | 0.05857 |
| CNAG_05668 | 116.2856 | -0.79247 | 0.210325 | -3.76785 | 0.000165 | 0.000578 |
| CNAG_11000 | 3.380565 | -4.11158 | 1.643758 | -2.50133 | 0.012373 | 0.027904 |
